# Perturbation-based propagation of bacterial composition uncertainty improves robustness and reproducibility of microbiome differential abundance analysis

**DOI:** 10.64898/2026.10.23.760456

**Authors:** Madison E. Andrews, Tessa Vincennie, Milica Koprivica, Mackenzie Browning, Steven Bracco, Michael Mullan, Aaron T. Wright, Fiona Crawford, Benoit Mouzon

## Abstract

Differential abundance (DA) analysis remains the analytical bottleneck of microbiome science; despite dozens of available methods, results remain poorly reproducible across tools, datasets, and study designs. Three intertwined challenges drive this failure: compositional data structure, stochastic count uncertainty, and complex experimental design; yet no existing method addresses all three simultaneously. Here, we introduce PARALLAX (Perturbation-based Assessment of Robustness Across Latent Logical Axes), a unified DA framework that resolves this gap by propagating compositional uncertainty through per-replicate linear mixed modeling across a structured scale perturbation grid, then summarizes inference through two stability-based metrics: P_post_ (effect consistency across Monte Carlo replicates) and R_scale_ (robustness to scale perturbation). PARALLAX retains uncertainty through the complete inferential pipeline while natively supporting repeated measures and multifactorial designs. Under repeated subsampling, PARALLAX identified fully stable DA taxa in 14.4% of cases, versus 1.7% and 0% for comparators. We applied PARALLAX to two independent longitudinal shotgun metagenomic datasets from mechanistically distinct mouse models: Gulf War toxicant exposure and repetitive head trauma. In both, PARALLAX resolved robust, sex-specific, and time-dependent genus-level shifts while accommodating repeated measures and multifactorial design, tasks that challenged both comparator frameworks. PARALLAX is released as an open-source, AI-compatible R workflow, lowering the barrier to rigorous microbiome DA analysis across complex study designs.

**Importance:** Microbiome studies link gut microbes to immune, metabolic, and neurological health, but identifying microbes that truly change with disease or treatment remains difficult. Differential abundance results often vary across computational tools because microbiome data are noisy, sparse, and compositional, especially in longitudinal or multifactorial studies. This instability limits reproducibility and creates difficulty in prioritizing microbes for biological interpretation. PARALLAX addresses this gap by focusing on consistency rather than relying on a single model fit. It analyzes data under multiple plausible representations that capture uncertainty in counts and composition, then identifies bacteria with consistent effects using stability and robustness metrics. Using ground-truth simulations and two mechanistically distinct mouse models, Gulf War toxicant exposure and traumatic brain injury, PARALLAX recovers microbial signals that remain reliable across complex study designs. This open-source R workflow provides a more reproducible path from microbiome sequencing to biological insight.

## Introduction

High-throughput sequencing technologies such as 16S rRNA amplicon profiling and shotgun metagenomics have enabled large-scale characterization of microbiome bacterial communities, driving the development of robust statistical methods for differential abundance (DA) analysis(1–9). DA analysis remains one of the least reproducible components of the microbiome analysis pipeline. Across 38 datasets and 14 methods, agreement among methods was often no better than chance, and effect direction and significance varied with analytical choice(10).

Three interconnected analytical challenges contribute to this instability. First, sequencing data is compositional; sequencing returns relative rather than absolute abundances. Therefore, a change in any one taxon mathematically affects all others(11). Second, taxa counts are stochastic realizations of an underlying community, with biological and technical variability that can inflate false positives or obscure stable signals(12). Third, many microbiome studies employ longitudinal, paired, and multifactorial designs that require modeling beyond simple two-group comparisons(12,13).

Existing methods address these challenges only in part. ALDEx2 models compositional uncertainty with Dirichlet-multinomial Monte Carlo (MC) sampling and centered log-ratio (CLR) transformation, yet aggregates MC replicates prior to modeling, discarding replicate-level variability at the inference stage. Additionally, ALDEx2 yields limited support for repeated measures or complex covariate structures(11). ANCOM-BC2 supports covariate adjustment, repeated measures, bias correction, and scale normalization, but inference is conducted on a single model fit, which does not account for the stochastic nature of the data(12). MaAsLin2, LinDA, and corncob add modeling flexibility but likewise do not propagate compositional uncertainty through model fitting(14–16).

PARALLAX is motivated by the principle that reliable DA signals should remain consistent when data are perturbed within plausible bounds of compositional and sampling uncertainty. This stability-based view of inference has precedent in sensitivity analysis and ensemble learning but has not been systematically integrated into microbiome DA workflows.

Here, we introduce PARALLAX (Perturbation-based Assessment of Robustness Across Latent Logical Axes), a DA framework that addresses these limitations through three coordinated features. First, PARALLAX retains each MC replicate as an independent realization and fits a full linear model to each replicate with limma. allowing repeated measures and multi-covariate designs while propagating compositional uncertainty through the inference stage. Second, PARALLAX applies a structured scale perturbation grid in CLR space to assess sensitivity to scale variation. Third, PARALLAX replaces single-fit p-value summaries with stability metrics; P_post_ represents the proportion of MC replicates supporting an effect beyond a defined threshold, and R_scale_ defines the proportion of scale perturbations under which that effect persists. We benchmark PARALLAX against ALDEx2 and ANCOM-BC2 and apply it to longitudinal shotgun metagenomic datasets from mouse models of Gulf War (GW) toxic exposure and traumatic brain injury (TBI).

## Results

### Application

PARALLAX identified 44 unique robust DA genera in sex-collapsed post-hoc contrasts and 71 unique robust genera in sex-stratified in the GW dataset.

In the TBI dataset, PARALLAX identified 52 unique robust DA genera in sex-collapsed post-hoc contrasts and 70 unique robust genera in sex-stratified contrasts.

For visualization, the 15 genera with the largest absolute log₂ fold-change were selected from each result, yielding 30 representative taxa within each dataset. The heatmap (**Fig. 1A, 2A**) shows robust effects across Exposure/Injury, Time, and Sex contrasts, with several genera exhibiting DA across multiple contrasts. Multiple genera yielded decreased abundance in male GW samples at initialization. In female TBI mice, multiple genera exhibited elevated abundance at the conclusion of the experiment. Exposure/Injury x Time variation was observed in the sex-collapsed interaction, suggesting that many interaction effects were not sex-dependent (**Fig. 1C, 2C**). P_post_ and R_scale_ were both high among q_post_ ≤ 0.05 taxa (**Fig. 1B, 2B**), with most taxa reaching R_scale_ = 1.

**Figure 1.**
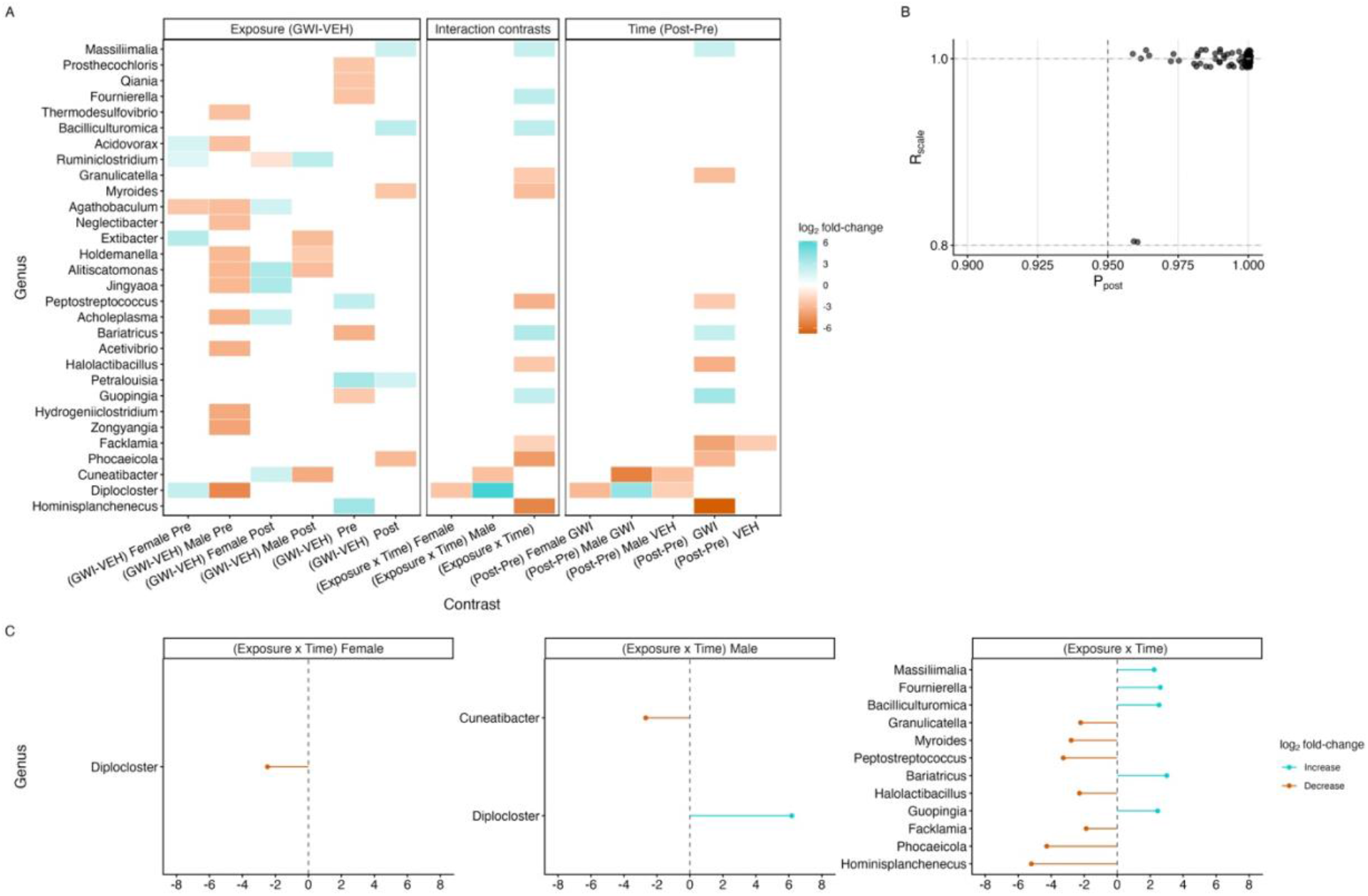
(A) Log₂ fold-change estimates for 30 representative genera across Exposure, Time, and Interaction contrasts. Colors indicate direction and magnitude of DA. (B) Relationship between posterior consistency (P_post_) and scale robustness (R_scale_) for retained taxa (q_post_≤0.05). (C) Forest-style plots of log₂ fold-change estimates for interaction contrasts for quantitative comparison of effect sizes within each condition.

**Figure 2.**
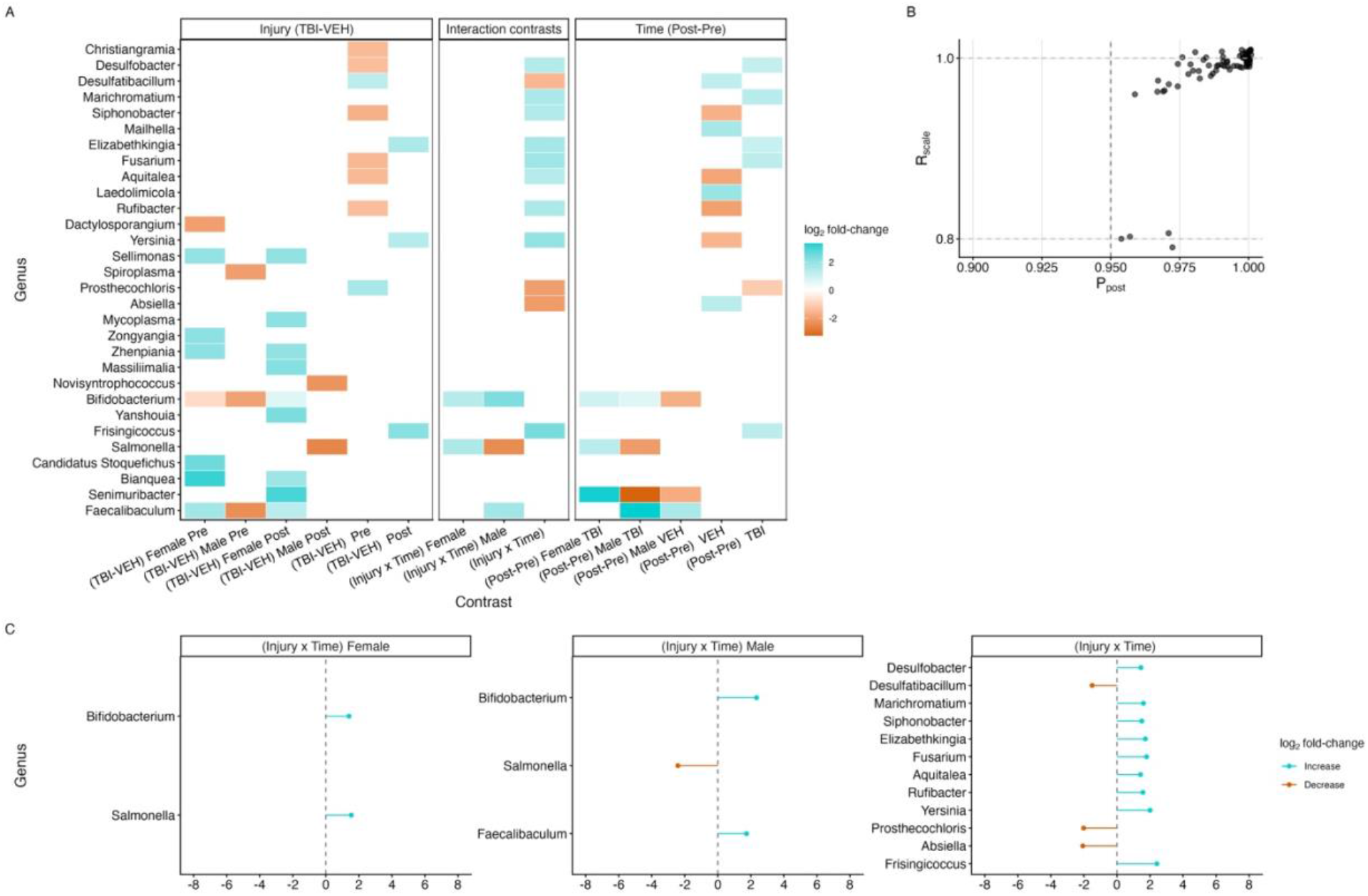
(A) Log₂ fold-change estimates for 30 representative genera across Injury, Time, and Interaction contrasts. Colors indicate direction and magnitude of DA. (B) Relationship between posterior consistency (P_post_) and scale robustness (R_scale_) for retained taxa (q_post_≤0.05). (C) Forest-style plots of log₂ fold-change estimates for interaction contrasts for quantitative comparison of effect sizes within each condition.

**Figure 3.**
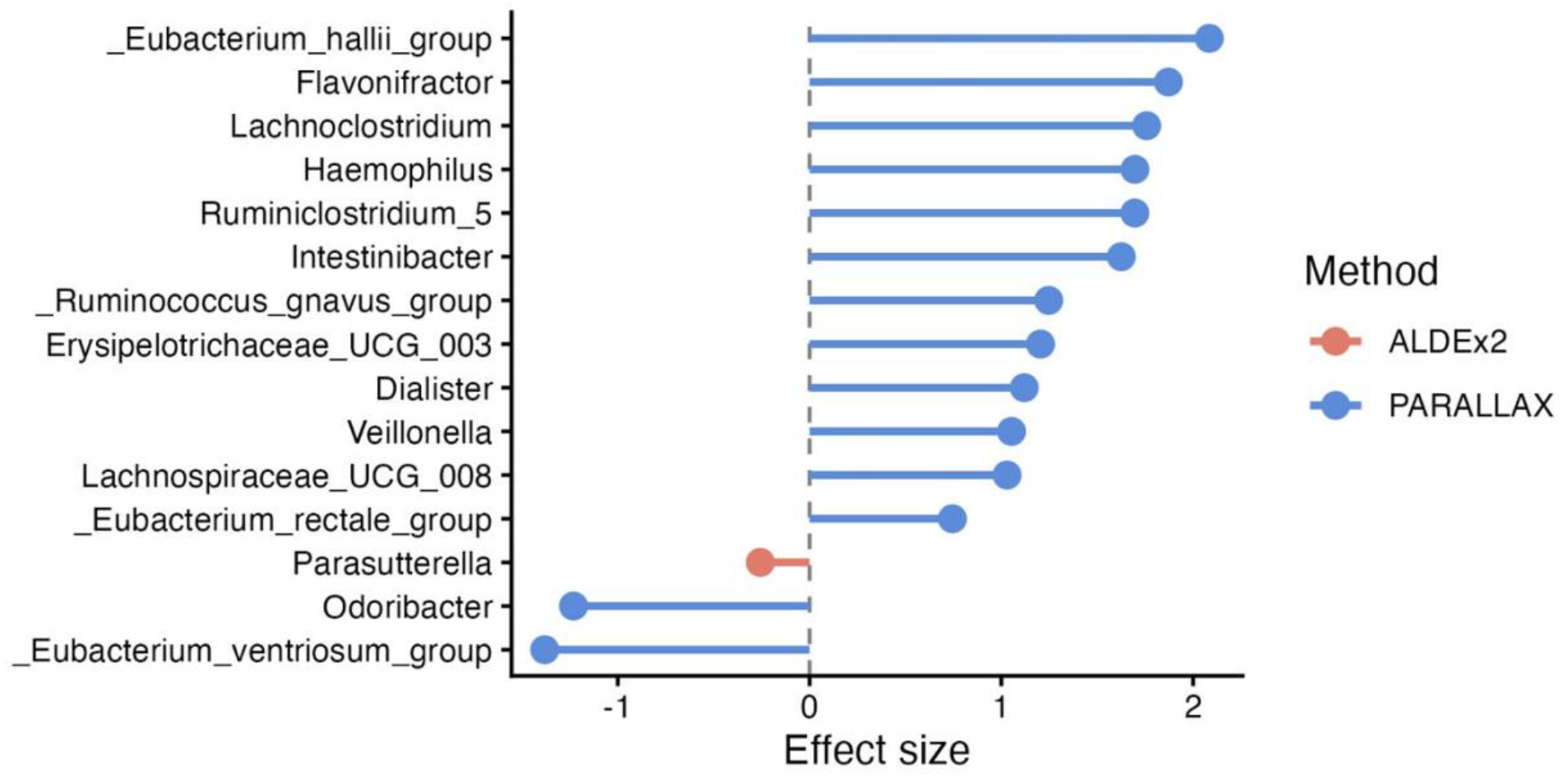
Genera exhibiting significant changes in abundance in IBD patients identified by the three DA methods with log_2_ fold-change effect size directions. Relevant hits were filtered by the Disease main effect. ANCOM-BC2 returned no relevant hits.

On the HMP2 dataset derived from the The Inflammatory Bowel Disease Multiomics Database, PARALLAX identified 14 genera significantly associated with IBD status, whereas ALDEx2 identified only a single genus (*Parasutterella*) and ANCOM-BC2 identified no significant taxa.

The taxa detected by PARALLAX were broadly consistent with established features of IBD-associated dysbiosis. Notably, the *Ruminococcus gnavus* group exhibited a positive association with disease, corroborating numerous reports that have identified *R. gnavus* as one of the most reproducibly enriched taxa in both Crohn’s disease and ulcerative colitis(17,18). PARALLAX also identified positive disease associations for *Haemophilus*(19) and *Veillonella*(17), genera belonging to bacterial communities that are frequently enriched during intestinal inflammation and are characteristic of the expansion of facultative anaerobic and Proteobacteria-associated taxa observed in IBD. Conversely, several taxa with known roles in short-chain fatty acid production and maintenance of gut homeostasis, including members of the *Eubacterium* (*E. hallii*, *E. rectale*, and *E. ventriosum* groups)(17), Lachnospiraceae (*Lachnospiraceae UCG-008*, *Lachnoclostridium*)(20), *Odoribacter*(21), and *Ruminiclostridium*(17,18), were significantly altered, consistent with the well-documented depletion of butyrate-producing Firmicutes in IBD. Although ALDEx2 identified only *Parasutterella*, this genus has also been implicated in IBD-associated alterations in microbial metabolism and bile acid homeostasis(22). Collectively, these findings indicate that PARALLAX identifies several well-established components of the IBD microbial signature while delivering a substantially larger set of disease-associated taxa than the comparator differential abundance methods.

### Ground Truth

Ground-truth simulations evaluated detection accuracy for known DA taxa across 100 independently generated simulation replicates, incorporating taxon-specific negative-binomial overdispersion. Precision and recall for the exposure main effect is shown in Fig. 4. Additional results for the other main effects and interaction effect are provided in Supplementary Figures S2–S4.

**Figure 4.**
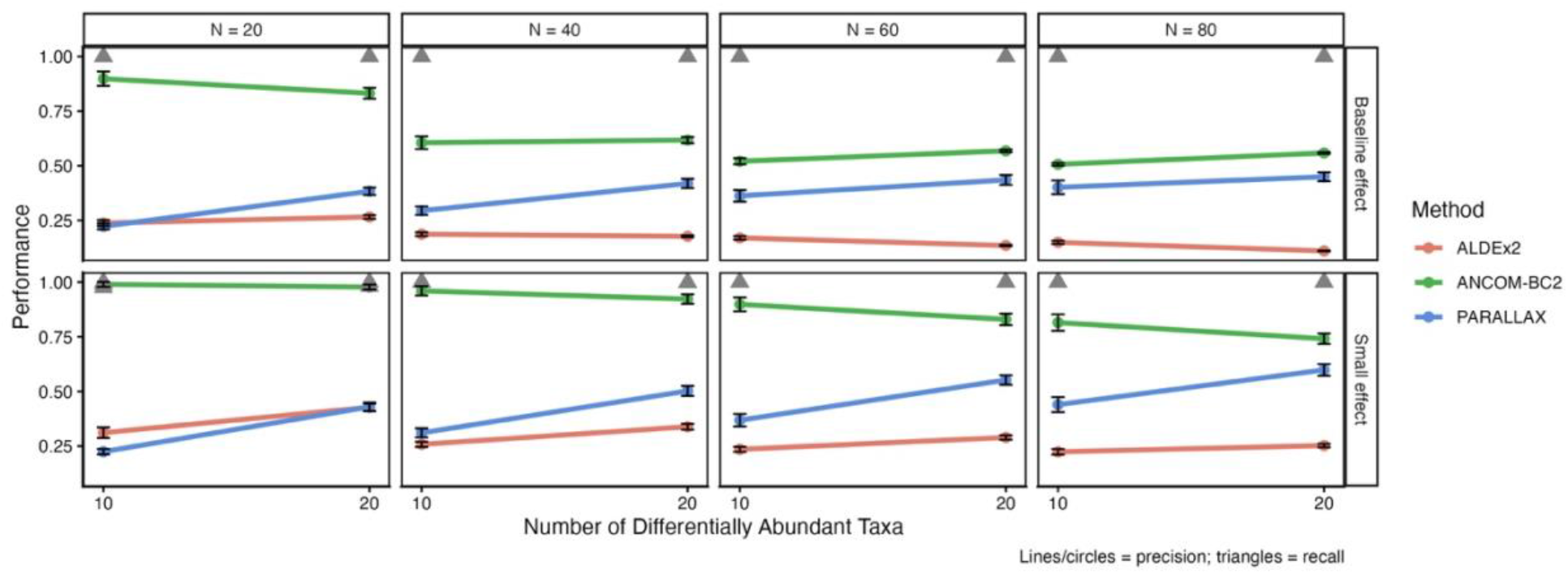
Precision and recall are shown for PARALLAX, ANCOM-BC2, and ALDEx2 across sample sizes (N = 20, 40, 60, 80), numbers of true differentially abundant taxa (DA = 10, 20), and effect size regimes (baseline and reduced) for the exposure main effect. Error bars show 95% CI across replicates.

For the exposure main effect, ANCOM-BC2 achieved the highest precision under nearly all conditions tested, ranging from 0.90 (N = 20, 10 DA taxa, baseline effect) to 0.64 (N = 80, 20 DA taxa, baseline effect), and from 0.99 to 0.79 across the same range under the reduced-effect condition. PARALLAX showed intermediate precision (0.35–0.45 under baseline effect; 0.38–0.63 under reduced effect) that, unlike either comparator, increased with sample size in both effect-size regimes. ALDEx2 showed the lowest precision overall, declining from 0.29 to 0.10 as N increased from 20 to 80 under baseline effect, and from 0.43 to 0.25 under reduced effect. Recall was maintained across all conditions tested for all three methods at or near 1.0, indicating that differences between methods for this effect were driven entirely by false-positive behavior rather than sensitivity.

For the exposure × time interaction, only PARALLAX’s model specification includes an interaction term; ALDEx2 and ANCOM-BC2 were not configured to test this effect in the present benchmark and are therefore not compared directly (**Fig. S4**). PARALLAX’s precision for the interaction effect was lower than for the exposure main effect, ranging from 0.20 to 0.52 under baseline effect and 0.13 to 0.53 under reduced effect, in both cases increasing with sample size. Recall was near-complete under baseline effect (0.95–1.00) but dropped substantially at small sample sizes under the reduced-effect condition (0.60 at N = 20, rising to 0.93 at N = 80.

R_scale_ separated true DA taxa from non-DA taxa in both baseline and reduced-effect datasets (**Fig. S5A, S5B**). True DA taxa exhibited high stability (R_scale_ > 0.95), whereas non-DA taxa showed lower, broader R_scale_ distributions. R_scale_ also increased with estimated effect size (**Fig. S5C, S5D**). True DA taxa clustered at larger log_2_ fold-changes and high R_scale_, while non-DA taxa exhibited smaller effects and lower stability.

### Comparison of method quantitative metrics

PARALLAX effect sizes agreed strongly with ALDEx2 (Pearson r = 0.84) and moderately with ANCOM-BC2 (r = 0.72), demonstrating broad agreement in direction and relative magnitude (**Fig. 5A, 5B**). Despite this general alignment, the distribution of effect sizes differed substantially across methods. PARALLAX produced a wider range of effect sizes (approximately −5 to 10) compared to ALDEx2 (approximately −1 to 3) and ANCOM-BC2 (approximately −2 to 6). For ALDEx2, the correspondence with PARALLAX is primarily aligned along the diagonal; however, PARALLAX extends these effects, separating and amplifying signals that are consistently supported across perturbations. In contrast, the relationship between PARALLAX and ANCOM-BC2 shows greater dispersion, particularly in the vertical direction (i.e., PARALLAX values vary more for a given ANCOM-BC2 estimate).

**Figure 5.**
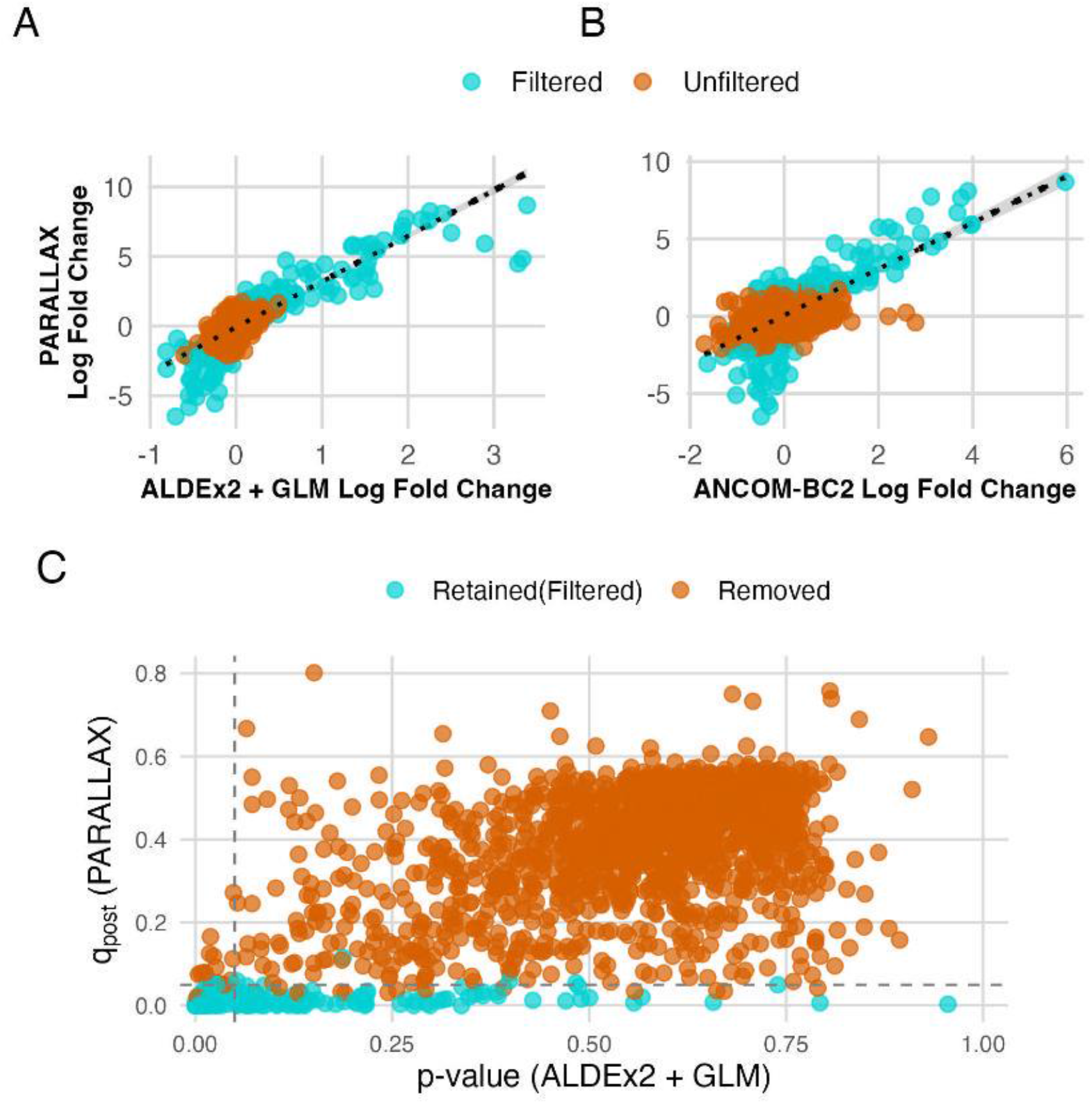
(A) Comparison of PARALLAX and ALDEx2 effect sizes shows strong agreement with an expanded dynamic range for PARALLAX. (B) Comparison with ANCOM-BC2 shows moderate agreement and greater dispersion, indicating reduced alignment in effect magnitude. (C) PARALLAX posterior lack-of-consistency metric (q_post_) versus ALDEx2 p-values. The horizontal dashed line indicates the nominal ALDEx2 p = 0.05 threshold, while the vertical dashed line indicates the PARALLAX decision criterion q_post_ = 0.05 (equivalent to P_post_ ≥ 0.95). Blue points represent taxa satisfying the PARALLAX robustness criteria (q_post_ ≤ 0.05 and R_scale_ ≥ 0.8).

Figure **5C** compares the PARALLAX consistency metric with ALDEx2 p-values. Blue points indicate taxa passing PARALLAX filters (q_post_ ≤ 0.05 and R_scale_ ≥ 0.8); orange points did not meet both criteria. PARALLAX and ALDEx2 agreed for taxa in the lower left region, but PARALLAX retained additional perturbation-consistent taxa that did not meet ALDEx2 p-value criteria.

### Subsampling stability

PARALLAX produced more stable results under subsampling (**Fig. 6**). Fully stable taxa were absent for ALDEx2 (0%) and rare for ANCOM-BC2 (1.7%) but more common for PARALLAX (14.4%). The detection-frequency distributions were also more informative for PARALLAX, spanning low to high stability rather than concentrating near zero.

**Figure 6.**
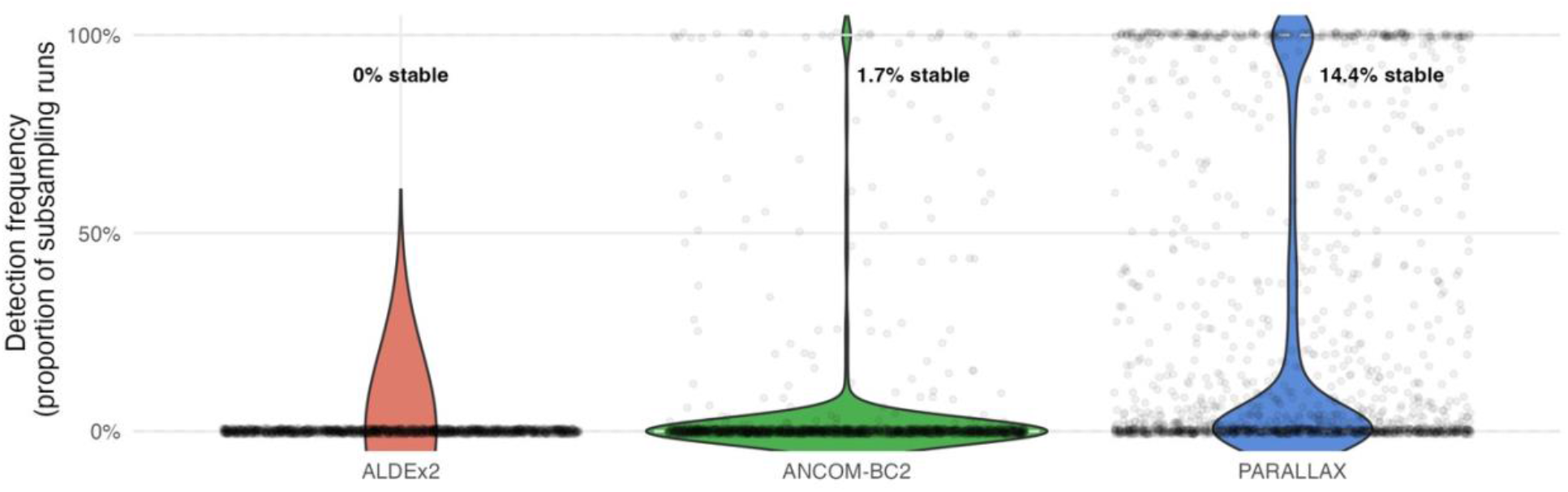
Subsampling stability comparison across methods. Violin plots of detection frequency across 50 subsampling runs for ALDEx2, ANCOM-BC2, and PARALLAX. PARALLAX identifies a larger proportion of fully stable taxa (14.4%) and demonstrates a broader distribution of detection frequencies, indicating improved reproducibility and discrimination between stable and unstable signals compared to existing methods.

### Empirical stability and R_scale_

Empirical stability increased with R_scale_, indicating that taxa whose effects were preserved across a greater proportion of scale perturbation conditions were more reproducibly detected under subsampling (**Fig. 7**). The majority of taxa (*N* = 1445) exhibited R_scale_ = 0, reflecting sensitivity to scale perturbation. Intermediate R_scale_ levels (0.2–0.8) contained relatively few taxa (*N* = 13-19), resulting in greater variability in mean stability estimates at these levels. In contrast, taxa with R_scale_ = 1.0 (*N* = 189) showed the highest proportion of fully stable signals, with many taxa consistently detected across all subsampling iterations.

**Figure 7.**
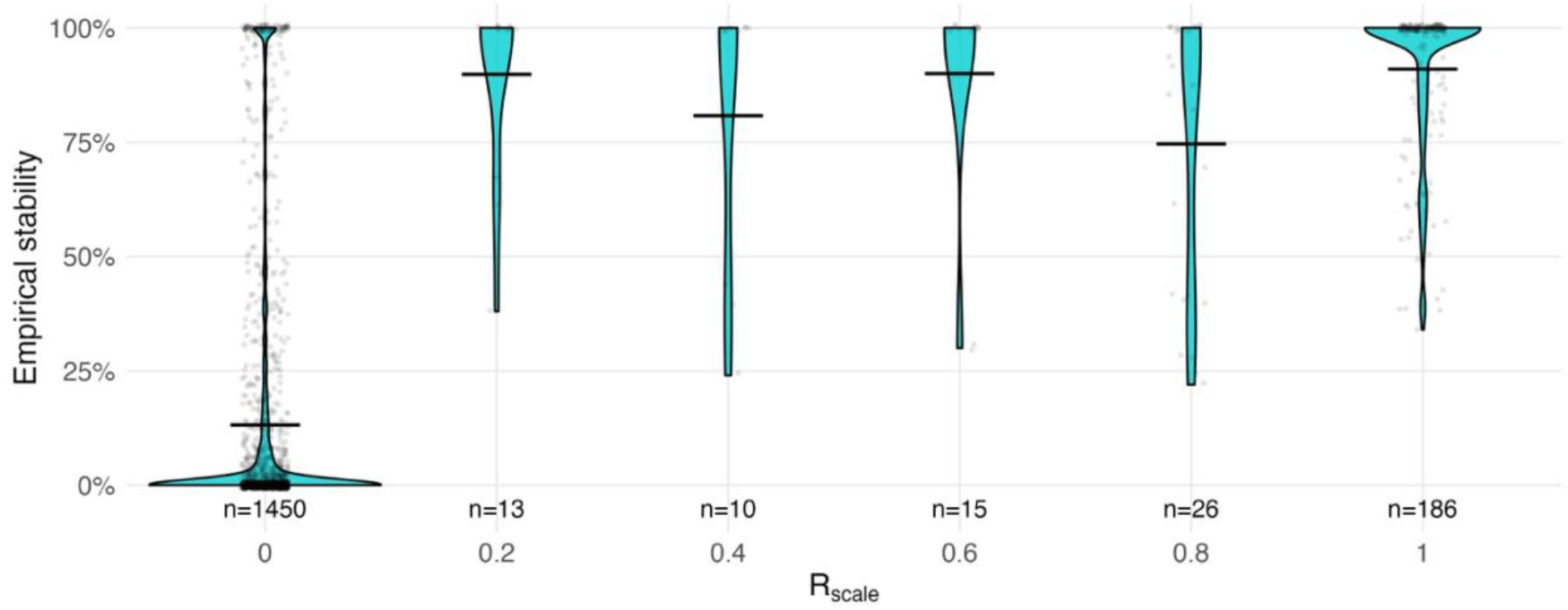
Relationship between scale robustness and empirical stability. Empirical stability (detection frequency across subsampling) increases with R_scale_, with higher R_scale_ values enriched for highly stable taxa. Persistence of variability at R_scale_ = 1 indicates that scale robustness captures a complementary dimension of stability distinct from subsampling stability.

## Discussion

PARALLAX was developed as a stability-based DA framework and evaluated its performance relative to two established methods, ALDEx2 and ANCOM-BC2, selected for compositional uncertainty modeling and scale-aware regression, respectively.

### Ground-truth validation

Ground-truth simulations showed that, for the exposure main effect, ANCOM-BC2 achieved the highest precision among the three methods across nearly all conditions tested, while PARALLAX showed intermediate precision that increased with sample size, a pattern which was not observed for either comparator, both of which showed flat or declining precision as N increased. This distinguishes PARALLAX’s failure mode from those of the other two methods: rather than accumulating false positives as more data become available, PARALLAX’s false-positive rate declines with additional samples, consistent with its consistency- and stability-based detection criteria becoming better calibrated as Monte Carlo and perturbation estimates stabilize.

ALDEx2’s effect-size-dependent precision is consistent with a known limitation of standard CLR-based normalization, where a true, large-magnitude shift in a subset of taxa can measurably displace the compositional reference and cause unaffected taxa to appear differentially abundant. ANCOM-BC2’s bias-correction step appears to mitigate this instability substantially, achieving the highest precision of the three methods under both effect-size regimes for this effect.

It is important to note that the detection criteria used in PARALLAX are not based on classical hypothesis testing. The q_post_ metric represents a probability of effect consistency across MC replicates rather than a calibrated p-value and does not provide formal control of the false discovery rate. In the present simulations, this yielded precision that, while lower than ANCOM-BC2’s for the exposure main effect, was accompanied by maintained recall and, distinctively, by precision that improved rather than degraded with sample size. These results should be interpreted as empirical evidence of PARALLAX’s behavior under the conditions tested rather than a guarantee of superior detection accuracy in all settings; formal characterization of error rates remains an important area for future work.

A further distinction between methods emerged for the exposure × time interaction. PARALLAX’s modeling framework explicitly incorporates interaction terms, allowing direct evaluation of interaction-specific detection accuracy; ALDEx2 and ANCOM-BC2, as benchmarked here, do not model this term and consequently cannot be evaluated on the same basis. This should be understood as a difference in modeling capability rather than a demonstrated performance advantage, since no comparable output exists for the other two methods under this benchmark configuration. Within PARALLAX alone, precision and recall for the interaction effect were both lower and more sample-size-dependent than for the corresponding main effect, particularly under the reduced-effect condition at small N, indicating that detecting interaction terms is a harder inferential problem than detecting main effects even within the same framework.

The simulations used in this study were designed to provide controlled evaluation of method performance while preserving key features of microbiome data, including compositional structure and variability in sequencing depth. By specifying known ground-truth taxa and applying defined effect sizes across covariates, this framework enables direct comparison of true and false detections. However, these simulations represent a limited set of conditions and do not capture the full diversity of real-world microbiome datasets. Factors such as more complex sparsity patterns, zero inflation, taxon co-occurrence structure, and more heterogeneous sequencing depth distributions may influence method performance in practice.

These findings support PARALLAX as a conservative workflow for prioritizing DA signals that persist across compositional and scale perturbations rather than marginal effects sensitive to modeling assumptions.

Beyond binary detection accuracy, simulation results further demonstrate that PARALLAX leverages stability-based inference to distinguish true biological signal from noise. The stability metric R_scale_ exhibited clear separation between ground-truth DA and non-DA taxa (**Fig. S5A, S5B**), with DA taxa showing high stability and non-DA taxa displaying substantially lower values. Furthermore, R_scale_ showed a positive association with effect size (**Fig. S5C, S5D**), with true DA taxa characterized by both larger estimated log₂ fold changes and higher stability. However, this relationship was not strictly linear, suggesting that R_scale_ captures robustness of signal beyond magnitude alone. Together, these findings demonstrate that stability under perturbation provides an additional axis of inference that complements traditional significance- and effect size-based approaches.

### Benchmarking comparison

Benchmarking showed that PARALLAX preserves the broad signal structure recovered by established methods while extending the dynamic range of effect size estimates (**Fig. 5A, 5B**).

PARALLAX also produced more stable results under repeated subsampling (**Fig. 6**). This improvement in stability likely reflects the ability of PARALLAX to jointly model compositional uncertainty and scale perturbation while retaining replicate-level variability throughout inference. In contrast, ALDEx2 collapses uncertainty prior to model fitting, and ANCOM-BC2 relies on point estimates, which may limit the ability of those methods to identify reproducible signals.

One potential explanation for the greater number of disease-associated taxa identified by PARALLAX in the HMP2 dataset is its ability to accommodate the hierarchical structure of the HMP2 data through participant-specific random effects (**Fig. 3**). Because multiple biopsies were obtained from individual participants, observations are not independent. By modeling participant-level variation directly, PARALLAX may provide increased power to detect disease-associated taxa while accounting for within-subject correlation.

PARALLAX identified additional consistent signals that were missed by ALDEx2 p-value thresholds (**Fig. 5C**), highlighting the difference between stability across plausible data realizations and significance in a single test.

A central contribution of PARALLAX is the introduction of R_scale_, a metric designed to isolate robustness to scale perturbation. While posterior consistency (P_post_) evaluates stability across both compositional and scale uncertainty, R_scale_ specifically measures invariance across scale perturbations. Empirical analysis demonstrated a strong, though not deterministic, relationship between R_scale_ and subsampling-based stability (**Fig. 7**). Taxa with higher R_scale_ values were enriched for highly stable signals, but variability remained at R_scale_ = 1, indicating that scale robustness represents only one dimension of overall stability.

### Biological interpretation

Application of PARALLAX to the GW toxic exposure dataset identified robust and structured shifts in gut microbial composition across exposure, time, and their interaction. GW-exposed animals showed consistent decreases relative to vehicle controls in multiple genera with established roles in intestinal homeostasis and host immune modulation, including *Ruminiclostridium, Agathobaculum, Acetivibrio, Hydrogeniiclostridium*, and *Diplocloster*. Reduction of these taxa is consistent with prior reports of butyrate depletion and increased intestinal permeability in GW preclinical models, supporting a mechanism whereby toxicant-induced microbial depletion could promotes systemic inflammation via barrier disruption and TLR4 activation(1,23). Additional decreases in *Peptostreptococcus, Bariatricus, Halolactibacillus, Guopingia*, and *Hominisplanchenecus*, and increases in *Fournierella, Granulicatella, and Bacilliculturomica*, were identified in Exposure × Time interaction contrasts, suggesting that some microbial shifts intensify or emerge longitudinally rather than being fully established at the pre-treatment timepoint. Notably, *Diplocloster* showed divergent sex-dependent trajectories decreasing in females but increasing in males in the Exposure × Time interaction; while *Cuneatibacter* increased specifically in GW-exposed males, highlighting sex-specific microbial responses that would not be detectable without the stratified contrast framework enabled by PARALLAX. Collectively, these findings reveal a multifactorial pattern of GW toxicant-associated gut dysbiosis involving fermentative and immunomodulatory genera across exposure and temporal dimensions and demonstrate the capacity of PARALLAX to resolve overlapping biological signals in complex longitudinal study designs.

Application of PARALLAX to the TBI dataset revealed distinct and reproducible shifts in gut microbial composition following repetitive head injury (RHI), with patterns that differed in character from those observed in the GW toxic exposure model. Among the most consistent findings, *Bifidobacterium* exhibited divergent sex-dependent trajectories in the Injury × Time interaction, decreasing in females but increasing in males mirroring the pattern of sex-stratified microbiome responses observed in the GW dataset and reinforcing the importance of sex-stratified analyses in preclinical neurotrauma models. *Faecalibaculum*, a genus associated with short-chain fatty acid production and intestinal immune regulation, increased in TBI-exposed males over time, while *Salmonella*-classified reads decreased across both sexes in interaction contrasts. In the sex-collapsed Injury × Time analysis, robust increases were observed in *Marichromatium*, *Siphonobacter*, *Elizabethkingia*, *Fusarium*, *Aquitalea*, and *Rufibacter*, while *Desulfobacter*, *Desulfatibacillum*, *Prosthecochloris*, *Absiella*, and *Frisingicoccus* were consistently decreased. Several of these taxa have been reported in the context of opportunistic infection and immune dysregulation, raising the possibility that TBI-associated gut dysbiosis involves shifts toward taxa with pathobiont characteristics alongside loss of commensal fermenters. PARALLAX has demonstrated the ability to resolve structured, multifactorial microbiome responses across two biologically distinct models of neurological injury, identifying both shared and model-specific patterns of gut dysbiosis that would be difficult to detect with conventional single-fit DA methods.

### Limitations

Despite these advantages, several limitations should be considered. First, PARALLAX introduces increased computational complexity due to the generation and propagation of MC replicates across perturbation conditions. Second, the framework relies on assumptions inherent to the underlying MC sampling strategy (i.e. Dirichlet-based modeling), which may not fully capture all aspects of true microbiome distributions. Third, PARALLAX does not directly measure reproducibility across independent datasets, instead inferring stability through perturbation-based methods. Additionally, unlike ANCOM-BC2, PARALLAX does not explicitly estimate absolute abundances, and therefore its results remain relative. Furthermore, while PARALLAX was evaluated on two representative datasets, broader validation across diverse microbiome datasets will be necessary to fully characterize its performance. Finally, PARALLAX evaluates the robustness of detected features but does not independently assess taxonomic assignment accuracy. Consequently, biological interpretation remains dependent on the quality of the underlying taxonomic annotation.

## Conclusion

PARALLAX provides a stability-based DA framework that propagates compositional and scale-related uncertainty through modeling, enabling inference to prioritize effects that remain consistent across plausible data realizations. Across benchmarking analyses, PARALLAX showed comparatively stable and sample-size-robust precision, improving rather than degrading as N increased, along with maintained recall, improved subsampling reproducibility, and additional complementary scale-robustness insight relative to ALDEx2 and ANCOM-BC2. For the exposure main effect specifically, ANCOM-BC2 achieved the highest raw precision of the three methods under the conditions tested. These findings support stability-based inference as a useful complement to traditional significance-based approaches for reproducible microbiome DA analysis.

## Materials and Methods

### GW Toxic Exposure Animal Model

Analysis was conducted on a longitudinal C57BL/6J mouse model of GW toxic exposure(24). Sixty-four mice were assigned by sex and exposure status, yielding four groups: male vehicle, male GW, female vehicle, and female GW. Details of dosing and housing are provided in the Supplementary Methods. Fecal pellets were collected from 40 mice (10 per sex-by-exposure group) at 8 and 12 months after the final toxic exposure and submitted to Transnetyx for shotgun metagenomic sequencing.

### TBI Animal Model

Analysis was conducted on a longitudinal C57BL/6J mouse model of repeated head injury (RHI) developed and validated by The Roskamp Institute(25–27). Male and female mice (aged 3–4 months) received five repeated closed-head injuries at 48-hour intervals under isoflurane anesthesia using an electromagnetic impactor with parameters validated to remain below the threshold for skull fracture or gross hemorrhage(26). Sham animals underwent identical anesthesia and handling without impact. This paradigm produces progressive and long-lasting neuropathological and neurobehavioral deficits including axonal injury, neuroinflammation, and cognitive impairment(25–27). As part of a longitudinal aging study, fecal pellets were collected at 20 months post-injury (n=5 per group per sex) and 24 months post-injury (n=9–10 per group per sex) and submitted to Transnetyx for shotgun metagenomic sequencing.

### PARALLAX Statistical Workflow

The workflow was developed in R v. 4.5.2 (2025-10-31) using RStudio v. 2026.5.0.218 on macOS v. 15.7.7. Data wrangling used tidyverse v. 2.0.0. The workflow is summarized in Fig. 8.

**Figure 8.**
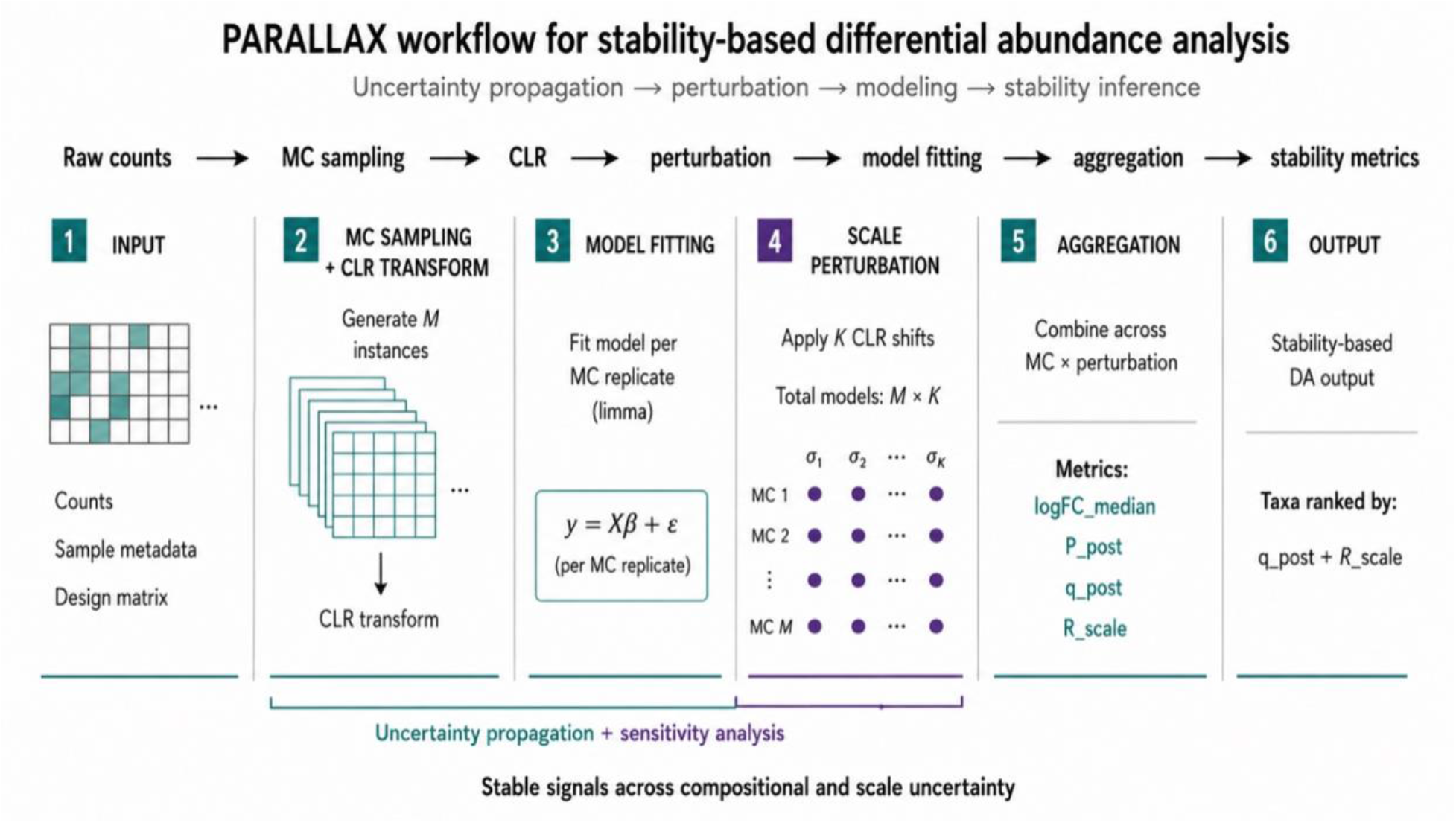
PARALLAX workflow for stability-based differential abundance analysis. Raw count data are expanded into Monte Carlo compositional replicates, transformed using CLR, and evaluated across a structured scale perturbation grid. Per-replicate linear models are fit and aggregated to compute stability-based metrics (P_post_, q_post_, R_scale_), enabling prioritization of taxa with consistent effects across compositional and scale uncertainty.

Raw count data were transformed with the CLR transformation implemented in ALDEx2 v. 1.42.0. ALDEx2’s Dirichlet–multinomial framework generated *M* MC replicates of the dataset (standard *M* = 128), each representing a plausible realization of the underlying microbial composition.

Unlike the standard ALDEx2 workflow, PARALLAX retained each MC replicate for downstream modeling, rather than limiting the propagation of compositional uncertainty to the transformation stage. A structured perturbation grid with K = 5 levels spanning [0,1] was then applied to evaluate sensitivity to scale variation. The grid provides a tractable approximation of continuous scale perturbation and can be adjusted by the user. Moderate variation in the grid size or spacing did not qualitatively alter results (**Fig. S1**). For each MC replicate, sample-wise perturbations were applied as additive CLR-space shifts drawn from a normal distribution *N*(0, *σ*^2^) producing *M x K* MC replicates that span compositional and scale uncertainty. Each replicate was fit independently to a common model design with limma v. 3.66.0. CLR-transformed MC replicates allowed limma to accommodate repeated measures, covariates, and interaction terms; within-subject correlation was estimated via duplicateCorrelation.

Following model fitting, effect estimates were aggregated across all *M* x *K* realizations. Instead of relying on p-values derived from a single model fit, summary statistics were defined based on the distribution of effect estimates across the full perturbation space. For each feature and model coefficient, the median log_2_ fold-change was computed to provide a robust estimate of effect size. In addition, a probability-based metric derived from MC replicates, P_post_, was defined as the proportion of simulations in which the inferred effect exceeded a predefined threshold *δ* in either direction. The complementary metric, q_post_ = 1 – P_post_, represents the proportion of Monte Carlo and scale-perturbation realizations that failed to support the effect-size criterion. Thus, q_post_ functions as a posterior consistency measure rather than a p-value, significance level, or calibrated estimate of false discovery rate (FDR). Consequently, inference in PARALLAX reflects stability-based filtering rather than hypothesis testing in the classical frequentist sense.

The effect-size threshold was set to log(1.2), corresponding to an ∼20% change in relative abundance, and can user-adjustable. Smaller thresholds increase sensitivity; larger thresholds prioritize stronger effects. R_scale_ was calculated by grouping MC replicates by perturbation level and estimating the proportion of levels in which the effect remained consistent. R_scale_ therefore distinguishes global consistency from scale-dependent effects. Post-hoc contrasts used the same MC and scale-perturbation framework. Sex-collapsed and sex-stratified contrasts were evaluated to support group-specific interpretation while preserving stability-based inference.

### Choice of Comparator Methods for Benchmarking

PARALLAX was benchmarked against ALDEx2 v. 1.42.0 and ANCOM-BC2 v. 2.12.1. ALDEx2 was selected because it also employs centered log-ratio (CLR) transformation and MC sampling. However, it collapses MC replicates before modeling and has limited native support for repeated measures or fully specified multifactor models. ANCOM-BC2 was selected because it supports covariates, repeated measures, bias correction, and scale normalization. However, it conducts point estimates and binary detection outcomes rather than replicate-level uncertainty.

### Evaluation on Simulated Data with Ground Truth

Simulated microbiome count data were generated from taxa abundances in the GW toxic exposure dataset to preserve realistic compositional structure. A subset of 120 taxa was sampled, baseline log-abundances were initialized from a normal distribution, and metadata were generated for 40 subjects with longitudinal paired samples. Exposure and Sex were subject-level covariates; Time varied within subject.

Performance was evaluated across key dimensions including sample size, effect magnitude, and number of true differential taxa, representing primary drivers of statistical detectability. Differential abundance was introduced in log space by applying additive effect sizes to a predefined subset of 20 taxa corresponding to Exposure, Time, Sex, and Exposure × Time interaction effects. Effect sizes were drawn from uniform distributions to produce moderate (log_2_ fold-change ∼1.0–1.5) or reduced (scaled to ∼0.3–0.7) signal strengths. For each sample, perturbed log-abundances were exponentiated and normalized to generate compositional proportions. Sequencing counts were then simulated via Poisson sampling using these proportions and sample-specific sequencing depths, including a subset of high-load samples to introduce variability in library size. Ground-truth labels for differential taxa were retained for benchmarking. Additional simulations varied sample size (N = 20, 40, 60, 80) and numbers of true DA taxa (N = 10, 20), while preserving compositional structure and effect distributions.

Detection accuracy was assessed by comparing detected DA taxa to ground-truth for each covariate. Precision and recall were calculated as TP/(TP+FP) and TP/(TP+FN). R_scale_ performance was evaluated by comparing value of the metric for true DA and non-DA taxa and by relating R_scale_ to estimated log_2_ fold-change.

### Benchmarking Against Established Methods

Benchmarking was performed following the recommendation outlined by Bokulich et al. in regard to comparator method selection and optimization, selection of test datasets, and selection of performance metrics(28). Four complementary comparisons were conducted. First, concordance in effect direction was evaluated between methods by fitting comparable models to the GWI dataset and isolating the main effect of time. This effect was selected due to its sensitivity to model specification, particularly given ALDEx2’s limitations in handling repeated measures. The correlation of estimated effects between PARALLAX and each comparison method was assessed with the Pearson correlation coefficient. Next, unfiltered consistency metric results from PARALLAX were compared with raw ALDEx2 p-values. Adjusted p-values were not used as the small sample size caused convergence to a common insignificant value.

Similar to previous microbiome benchmarking studies that introduced stochastic variation through repeated random sampling of metagenomic sequence data, pipeline robustness was evaluated by repeatedly subsampling the available samples and comparing the consistency of DA results across iterations(29). The dataset was randomly subsampled 50 times, with each iteration retaining 70% of subjects (without replacement) while preserving the paired structure. The number of iterations was set to 50 to ensure stable averaged performance metrics while maintaining computational efficiency. Repeated resampling methods rely on aggregating results across iterations, but the marginal gain in precision diminishes as the number of iterations increases, making moderate iteration counts a practical and widely accepted compromise in benchmarking studies. For this study, stable performance was evaluated at 30 and 50 iterations, with minimal change between the two scenarios regarding performance of the three methods. For each subsample, all three methods were independently applied, and taxa identified as significant or robust (according to method-specific criteria) were recorded. Detection frequency across iterations was then computed for each taxon, providing an empirical measure of stability.

For PARALLAX, taxa were considered perturbation-consistent and scale-robust if they satisfied posterior stability thresholds (q_post_ ≤ 0.05 and R_scale_ ≥ 0.8). The threshold q_post_ ≤ 0.05 was chosen to require posterior support for the effect in at least 95% of perturbation realizations (P_post_ ≥ 0.95). Although not calibrated to control FDR, empirical results demonstrate that this threshold combination effectively suppresses false positives under the simulated conditions. The R_scale_ threshold was empirically determined using ground truth simulations. Across both baseline and shrunken-effect size datasets, very few taxa (fewer than 10 out of 720) achieved R_scale_ ≥ 0.8 without also exhibiting strong directional consistency (P_post_ ≥ 0.95), indicating that R_scale_ ≥ 0.8 corresponds to highly stable and reproducible signals. This threshold was therefore considered appropriately conservative for defining scale-robust DA.

For ALDEx2, taxa were considered significant at nominal *p* ≤ 0.05, and ANCOM-BC2 taxa were considered significant using the method’s native diff_robust = TRUE indicator. Finally, subsampling stability estimates were compared with PARALLAX Rscale values to assess whether robustness to scale perturbation captured a complementary dimension of uncertainty.All three methods were then applied to the evaluation of the publicly available HMP2 16S dataset from The Inflammatory Bowel Disease Multiomics Database (30). Data was evaluated over sex, diagnosis (simplified into IBD/nonIBD), and biopsy location metadata variables. Participants lacking a biopsy from both the small (ileum) and large intestine (rectum) were excluded from this analysis. A random effect was included for the Participant ID over both biopsy samples. Results comparison was done for the diagnosis variable for verification against current IBD literature.

### Application of PARALLAX to the GW Toxic Exposure and TBI Datasets

The PARALLAX DA framework was applied to a GW toxic exposure and a TBI gut microbiome dataset generated via shotgun metagenomic sequencing (Transnetyx). For this analysis, taxa were evaluated at the genus level, which remains a commonly used and reliably inferred taxonomic resolution in microbiome studies(31,32). This stated, PARALLAX is not restricted to a specific taxonomic level and can be applied at any resolution supported by the data.

After CLR-transformation and generation of the *M x K* realizations, a repeated-measures model incorporated Sex, Exposure status, Time, and one two-way interaction [ ∼ (Sex + Exposure + Time + (Exposure * Time)]. Each experimental group contained *N* = 10 subjects. Taxa were classified as perturbation-consistent and scale-robust if they satisfied q_post_ ≤ 0.05 and R_scale_ ≥ 0.8. Taxa with 0.05 < q_post_ ≤ 0.1 or 0.6 ≤ R_scale_ < 0.8 were considered exploratory and not included in primary analyses.

Post-hoc contrasts were applied to robust taxa. Contrasts were either stratified by sex or evaluated after collapsing across sex depending on observed sex effects; taxa exhibiting isolated, main effects of Sex were excluded from downstream contrast analysis.

### Computational considerations

The PARALLAX framework requires fitting models across multiple Monte Carlo (MC) instances and scale perturbations. In the present implementation, this corresponds to *M x K* = 128 x 5 = 640 model fits per analysis. Computational performance was evaluated on a dataset containing 80 samples and ∼1700 genera. On a standard MacBook Pro (M4, 16 GB), the full analysis required approximately 18 seconds of wall-clock time. Memory usage was approximately 1 GB. Computation scaled approximately linearly with the number of taxa, samples, MC replicates, and perturbation levels. Importantly, model fitting is independent across MC replicates and perturbation conditions, enabling straightforward parallelization to substantially reduce runtime for larger datasets.

## Declarations

### Ethics approval and consent to participate

All animal procedures were approved by The Roskamp Institute’s IACUC: VA TBI project [145], GWI project [149].

### Consent for publication

Not applicable.

## Availability of Data and Materials

### Data

The two mouse shotgun metagenomics datasets and metadata used and analyzed during the current study are available in the additional files as additional files 5-8. Files 5 and 6 pertain to the R149 GWI study, and files 7 and 8 pertain to the R145 TBI study.

The human 16S IBD dataset used and analyzed during the current study is available in the IBDMD repository, https://ibdmdb.org/results.

### Code

The source code is publicly available on GitHub at: https://github.com/TheRoskampInstitute/PARALLAX under the MIT license. The version used in this study (v1.0.0) has been archived in Zenodo and is available at DOI: 10.5281/zenodo.22851027

## Competing interests

The authors declare that they have no competing interests.

## Funding

This work was funded by the Department of Defense [CDMRP Log Number: TX220288].

## Authors’ Contributions

MA, Conceptualization, Methodology, Formal analysis, Writing-original draft, Writing-review and editing; TV, Conceptualization, Data generation, Writing-review; MK, Data generation, Exploratory analysis, Writing-review; MB, Conceptualization, Data generation; SB, Data generation; MM, Funding acquisition; AW, Conceptualization, Writing-review; FC, Funding acquisition; BM, Conceptualization, Data generation, Funding acquisition, Writing-review and editing

## Supporting information

GWI R149 metadata

Simulation source code

AI-parsable walkthrough

GWI R149 data

TBI R145 data

TBI R145 metadata

Simulation results

## Acknowledgements

The authors would like to acknowledge Gogce Crynen, Ph.D. for her assistance in reviewing the benchmarking method rigor for this work.

## Authors’ information

Fiona Crawford PhD is a Senior Research Career Scientist with the James A. Haley Veteran’s Hospital in Tampa Florida.

## Notes

https://github.com/TheRoskampInstitute/PARALLAX

### Competing Interest Statement

The authors have declared no competing interest.

