## Supplementary material for "Perturbation-based propagation of bacterial composition uncertainty improves robustness and reproducibility of microbiome differential abundance analysis": Simulation source code

### Additional file 4

Script created by Madison Andrews 8.18.2026.

Portions of this script were developed with assistance from Claude (Claude Sonnet 5, Anthropic), accessed via claude.ai, August 2026.

Compiled 9.3.2026 by Madison Andrews. Optimized for MacOS Sequoia 15.7.7, 16 GB RAM.

Microbiome data simulation and comparison between PARALLAX, ANCOM-BC2, & ALDEx2. Script simulates data based on a real dataset for N = 20-80, DA taxa = 10,20, and effect size of reduced/baseline. Simulates known ground truth and calculates precision/recall for each method. Has the capability of simulating poisson, negative binomial, and zero-inflated negative binomial distributions. Published results from this script utilizing the Roskamp Disease shotgun metagenomics dataset as the real data are included in the PARALLAX publication as [Additional_file_3.XLSX].

##### NOTE:

*Runtime for 100 replicates with these specifications was ~6 hours wall clock time with parallelization (recommend 6 cores). In comparison, 2 cores took ~8 hours. Script separates generation of baseline/reduced effect size data for this reason. See script parameters in number of replicates, variety of N, and number of Monte Carlo instances in ALDEx2 for computational load reduction.*

#========================================== # DO BEFORE RUNNING EITHER/BOTH SIMULATION(S)

Sys.setenv(OMP_NUM_THREADS = "1") Sys.setenv(OPENBLAS_NUM_THREADS = "1") Sys.setenv(MKL_NUM_THREADS = "1") Sys.setenv(VECLIB_MAXIMUM_THREADS = "1")

### Import packages

library(tidyverse) library(ALDEx2) library(ANCOMBC) library(limma) library(matrixStats) library(scales)

if (requireNamespace("RhpcBLASctl", quietly = TRUE)) { RhpcBLASctl::blas_set_num_threads(1) RhpcBLASctl::omp_set_num_threads(1) }

#### Set for the local user and desired file directory (macOS syntax)

setwd(“ “)

#### MODEL BASELINE DATA FIRST

#============================================================
### 0. Load real data # # NOTE: only rownames(counts) (the taxon names) are used, to sample a # realistic pool of taxa for the simulation. meta_sim2 is loaded for # completeness but is not consumed by the simulation itself. #============================================================

#### use real counts/metadata. Set up for N = 40 and “Genus” taxonomic classification

counts_raw <- read_csv(“counts.csv”) %>% dplyr::select(-1) %>% # drop leading numeric row-index column as.matrix()

counts <- counts_raw rownames(counts) <- paste0(“Genus_”, seq_len(nrow(counts_raw)))

meta_sim2 <- read_csv(“meta.csv”) %>% dplyr::select(-1)

#============================================================
### 1. Simulation functions # (build_master_cohort / subsample_cohort / small_effect) # # FEATURES: # # (a) Count generation supports “poisson”, “nb” # (negative binomial, default), or “zinb” (zero-inflated negative # binomial), via the `distribution` argument. # # (b) NB overdispersion is TAXON-SPECIFIC: each taxon gets its own NB # “size” parameter (size = 1 / dispersion), drawn once per master # cohort (Stream 1, seeded, independent of N) from a log-normal # distribution controlled by `nb_size_mean` / `nb_size_logsd`. # Smaller size => more overdispersion (variance = mu + mu^2/size). # # (c) If distribution == “zinb”, each taxon also gets a structural # zero-inflation probability, drawn from a Beta distribution # controlled by `zero_infl_prob` (mean) and `zero_infl_conc` # (concentration -> higher = less taxon-to-taxon variability in # the zero-inflation rate). # # (d) build_master_cohort() returns `size_taxon` and `pi_zero` in # the output list, so you can inspect/reuse the dispersion and # zero-inflation parameters that were actually drawn for a given # master cohort.
#============================================================

#### —- helper: single NB / ZINB draw for one sample's taxon vector ————–

simulate_counts_1sample <- function(mu, distribution, size_taxon, pi_zero) {

n_taxa <- length(mu)

counts <- switch( distribution, poisson = rpois(n_taxa, lambda = mu), nb = rnbinom(n_taxa, mu = mu, size = size_taxon), zinb = rnbinom(n_taxa, mu = mu, size = size_taxon), stop("distribution must be one of 'poisson', 'nb', 'zinb' ") )

if (distribution == "zinb") { structural_zero <- rbinom(n_taxa, 1, pi_zero) counts[structural_zero == 1] <- 0 }

counts }

build_master_cohort <- function( counts, n_subjects_max = 80, # change if desired for higher N simulation n_taxa = 120, n_DA = 20, prop_high_load = 0.2, seed = 123, distribution = c("nb", "poisson", "zinb"), nb_size_mean = 5, # mean NB size (dispersion) across taxa; lower = more overdispersed nb_size_logsd = 0.4, # log-scale SD -> taxon-to-taxon variability in dispersion nb_size_floor = 0.1, # hard floor so no taxon gets an absurdly tiny size zero_infl_prob = 0.1, # mean structural zero-inflation probability (zinb only) zero_infl_conc = 5, # Beta concentration for zero_infl_prob (zinb only) effect_scale = 1 ) {

distribution <- match.arg(distribution)

strata <- expand.grid(Disease = c("Disease", "Control"), Sex = c("M", "F"), stringsAsFactors = FALSE) n_strata <- nrow(strata) # 4

stopifnot( "n_subjects_max must be divisible by the number of Disease x Sex strata (4)" = (n_subjects_max %% n_strata) == 0 ) n_per_stratum <- n_subjects_max / n_strata

### —- Stream 1 (seed): taxon-level ground truth — independent of N —- set.seed(seed)

taxa_all <- rownames(counts) taxa <- sample(taxa_all, n_taxa)

prop_disease <- 0.35 prop_time <- 0.25 prop_sex <- 0.20 prop_interaction <- 0.20

n_DA_disease <- round(n_DA * prop_disease) n_DA_time <- round(n_DA * prop_time) n_DA_sex <- round(n_DA * prop_sex) n_used <- n_DA_disease + n_DA_time + n_DA_sex n_DA_interaction <- n_DA - n_used

idx <- sample(seq_len(n_taxa))

start <- 1 DA_disease <- idx[start:(start + n_DA_disease - 1)] start <- start + n_DA_disease

DA_time <- idx[start:(start + n_DA_time - 1)] start <- start + n_DA_time

DA_sex <- idx[start:(start + n_DA_sex - 1)] start <- start + n_DA_sex

DA_interaction <- idx[start:(start + n_DA_interaction - 1)]

effect_disease <- runif(length(DA_disease), 1.0, 1.5) effect_time <- runif(length(DA_time), 1.0, 1.5) effect_sex <- runif(length(DA_sex), 1.0, 1.5) effect_interaction <- runif(length(DA_interaction), 1.0, 1.5)

base <- rnorm(n_taxa, mean = 0, sd = 1)

### —- taxon-specific overdispersion (and zero-inflation, if requested) —- # Drawn in the same seeded stream as the rest of the taxon-level ground # truth, so they stay tied to the taxon identities/effects rather than to N. size_taxon <- exp(rnorm(n_taxa, mean = log(nb_size_mean), sd = nb_size_logsd)) size_taxon <- pmax(size_taxon, nb_size_floor)

if (distribution == “zinb”) { a <- zero_infl_prob * zero_infl_conc b <- (1 - zero_infl_prob) * zero_infl_conc pi_zero <- rbeta(n_taxa, a, b) } else { pi_zero <- rep(0, n_taxa) }

### —- Stream 2 (seed + 1000): balanced subject allocation + nested rank —- set.seed(seed + 1000)

subject_table <- do.call(rbind, lapply(seq_len(n_strata), function(s) { data.frame( Disease = strata$Disease[s], Sex = strata$Sex[s], nested_rank = sample(seq_len(n_per_stratum)), stringsAsFactors = FALSE ) })) subject_table <- subject_table[order(subject_table$Disease, subject_{t}able$Sex, subject_table$nested_{r}ank), ]subject_{t}able$ID <- paste0(“ID_”, seq_len(n_subjects_max)) rownames(subject_table) <- NULL

sim_meta <- expand.grid(ID = subject_table$ID, Time = c("Pre", "Post"), stringsAsFactors = FALSE)sim_{m}eta$Disease <- subject_table$Disease[match(sim_{m}eta$ID, subject_table$ID)]sim_{m}eta$Sex <- subject_table$Sex[match(sim_{m}eta$ID, subject_table$ID)]sim_{m}eta$SampleID <- paste0(“Sample”, seq_len(nrow(sim_meta)))

### —- Stream 3 (seed + 2000): count generation —- set.seed(seed + 2000)

counts_mat <- matrix(0, nrow = n_taxa, ncol = nrow(sim_meta)) rownames(counts_mat) <- taxa

depth <- 20000 n_high_load <- round(prop_high_load * nrow(sim_meta)) high_load_idx <- sample(seq_len(nrow(sim_meta)), n_high_load)

for (i in seq_len(nrow(sim_meta))) {

```
row <- sim_meta[i, ]
log_theta <- base

if (row$Disease == "Disease") {
  log_theta[DA_disease] <- log_theta[DA_disease] + effect_disease
} else {
  log_theta[DA_disease] <- log_theta[DA_disease] - effect_disease
}
```

```
if (row$Time == "Post") {
  log_theta[DA_time] <- log_theta[DA_time] + effect_time
} else {
  log_theta[DA_time] <- log_theta[DA_time] - effect_time
}

if (row$Sex == "F") {
  log_theta[DA_sex] <- log_theta[DA_sex] + effect_sex
} else {
  log_theta[DA_sex] <- log_theta[DA_sex] - effect_sex
}

if (row$Disease == "Disease" && row$Time == "Post") {
  log_theta[DA_interaction] <- log_theta[DA_interaction] + effect_interaction
}

theta <- exp(log_theta)
theta <- theta / sum(theta)

depth_i <- depth
if (i %in% high_load_idx) {
  depth_i <- depth_i * runif(1, 2, 4)
}

mu_i <- theta * depth_i

counts_mat[, i] <- simulate_counts_1sample(
  mu           = mu_i,
  distribution = distribution,
  size_taxon   = size_taxon,
  pi_zero      = pi_zero
)
```

}

colnames(counts_mat) <- sim_meta$SampleID

truth <- data.frame( Taxon = taxa, DA_Disease = seq_len(n_taxa) %in% DA_disease, DA_Time = seq_len(n_taxa) %in% DA_time, DA_Sex = seq_len(n_taxa) %in% DA_sex, DA_Interaction = seq_len(n_taxa) %in% DA_interaction ) truth$is.DA <- rowSums(truth[, 2:5]) > 0 list( counts = counts_mat, metadata = sim_meta, truth = truth, subject_table = subject_table, n_strata = n_strata, n_subjects_max = n_subjects_max, distribution = distribution, size_taxon = setNames(size_taxon, taxa), pi_zero = setNames(pi_zero, taxa) ) }

subsample_cohort <- function(master, n_subjects) {

stopifnot( "n_subjects must be divisible by the number of Disease x Sex strata (4)" = (n_subjects %% master$n_strata$) == 0, "$n_subjectscannotexceedthemastercohortsize$" = $n_subjects <= master$n_subjects_max )

per_stratum <- n_subjects / master$n_stratakeep_ids < -master$subject_table$ID[master$subject_table$nested_rank <= per_stratum]

meta_sub <- master$metadata[master$metadata$IDmeta_sub < -meta_sub[order(meta_sub$ID, meta_sub$Time), ]

counts_sub <- master$counts[, meta_sub$SampleID, drop = FALSE]

list( counts = counts_sub, metadata = meta_sub, truth = master$truth ) }

small_effect <- function(sim_counts, scale_factor = 0.5) { log_counts <- log(sim_counts + 1) log_counts_small <- log_counts * scale_factor noise <- matrix(rnorm(length(log_counts_small), mean = 0, sd = 0.05), nrow = nrow(log_counts_small)) log_counts_small <- log_counts_small + noise counts_small <- exp(log_counts_small) - 1 counts_small[counts_small < 0] <- 0 counts_small <- round(counts_small) counts_small }

#==================================================
### 2. PARALLAX pipeline (unchanged) #==========================================

runmypipeline <- function(counts, meta) {

x <- aldex.clr(counts, conds = rep("A", ncol(counts)), mc.samples = 128, denom = "all") mc_list <- getMonteCarloInstances(x) n_samples <- ncol(counts) n_features <- nrow(counts) M <- ncol(mc_list[[1]])

names(mc_list) <- colnames(counts) M <- ncol(mc_list[[1]])

clr_mats <- lapply(seq_len(M), function(mc) { mat <- sapply(mc_list, function(sample_mat) { vec <- sample_mat[, mc] vec_aligned <- vec[rownames(counts)] return(vec_aligned) }) mat <- as.matrix(mat) rownames(mat) <- rownames(counts) colnames(mat) <- colnames(counts) mat })

clr_array <- simplify2array(clr_mats) clr_mean <- apply(clr_array, c(1, 2), mean) clr_mean_t <- t(clr_mean)

meta <- meta %>% mutate( Sex = factor(Sex, levels = c("M", "F")), Disease = factor(Disease, levels = c("Control", "Disease")), Time = factor(Time, levels = c("Pre", "Post")), MouseID = factor(ID) ) if (!all(colnames(counts) %in% rownames(meta))) { stop("Metadata rows do not match count columns") }

meta <- meta[match(colnames(counts), rownames(meta)), , drop = FALSE] rownames(meta) <- colnames(counts)

if (any(is.na(meta))) { stop("NA values introduced in metadata alignment") }

mf <- model.frame(~ (Sex + Disease + Time)^2, data = meta, na.action = na.fail) design <- model.matrix(~ (Sex + Disease + Time)^2, data = mf) rownames(design) <- colnames(counts) colnames(design) <- make.names(colnames(design))

sigma_grid <- c(0, 0.25, 0.5, 0.75, 1)

expr_ref <- clr_mats[[1]] corfit <- duplicateCorrelation(expr_ref, design, block = meta$ID)consensus_{c}or < -corfit$consensus

fit_list_all <- vector("list", length(sigma_grid) * M) idx <- 1

for (sigma in sigma_grid) { message("Running sigma =", sigma) for (mc in seq_len(M)) { expr <- clr_mats[[mc]] stopifnot(ncol(expr) == nrow(design)) stopifnot(all(colnames(expr) == rownames(design)))

```
  log_s <- rnorm(ncol(expr), mean = 0, sd = sigma)
  expr <- sweep(expr, 2, log_s, FUN = "+")

  fit <- lmFit(expr, design, block = meta$ID, correlation = consensus_cor)
  fit <- eBayes(fit)

  fit_list_all[[idx]] <- fit
  idx <- idx + 1
}
```

}

coef_names <- colnames(fit_list_all[[1]]$coefficients) n_coef <- length(coef_names) n_features <- nrow(counts)

result_table <- data.frame( Coefficient = rep(coef_names, each = n_features), Taxon = rep(rownames(counts), times = n_coef), logFC_median = NA, P_post = NA, q_post = NA, R_scale = NA )

n_sigma <- length(sigma_grid) rep_per_sigma <- M sigma_labels <- rep(seq_len(n_sigma), each = rep_per_sigma)

row_idx <- 1

for (coef_idx in seq_len(n_coef)) {

```
theta_array <- do.call(cbind, lapply(fit_list_all, function(fit) {
  vec <- fit$coefficients[, coef_idx]
  vec[is.infinite(vec)] <- NA
  vec
}))
vals <- as.vector(theta_array)

delta <- log(1.2)

P_pos <- rowMeans(theta_array > delta, na.rm = TRUE)
P_neg <- rowMeans(theta_array < -delta, na.rm = TRUE)

P_post <- pmax(P_pos, P_neg)
q_post <- 1 - P_post

logFC_median <- apply(theta_array, 1, median, na.rm = TRUE)

R_scale_vec <- apply(theta_array, 1, function(theta_vec) {
  theta_by_sigma <- split(theta_vec, sigma_labels)
  sig_sigma <- sapply(theta_by_sigma, function(vec) {
    max(mean(vec > delta), mean(vec < -delta))
  })
  mean(sig_sigma)
})

result_table$logFC_median[row_idx:(row_idx + n_features - 1)] <- logFC_median
result_table$P_post[row_idx:(row_idx + n_features - 1)] <- P_post
result_table$q_post[row_idx:(row_idx + n_features - 1)] <- q_post
result_table$R_scale[row_idx:(row_idx + n_features - 1)] <- R_scale_vec

row_idx <- row_idx + n_features
```

} result_table }

#==========================================================================
### 3. Detection accuracy + sim_detection # # `truth` is passed explicitly (not a global), since with the replicate loop # below there are many distinct ground-truth tables in flight at once.
#==========================================================================

detection_accuracy <- function(truth, detected_disease, detected_sex, detected_time, detected_interaction) {

truth_vec_d <- truth |> filter(DA_Disease == TRUE) |> pull(Taxon) d_intersect <- detected_disease[detected_disease %in% truth_vec_d] disease <- tibble( var = "disease",

TP = length(d_intersect), FP = length(detected_disease) - length(d_intersect), FN = length(truth_vec_d) - length(d_intersect), precision = TP / (TP + FP), recall = TP / (TP + FN) )

truth_vec_s <- truth |> filter(DA_Sex == TRUE) |> pull(Taxon) s_intersect <- detected_sex[detected_sex %in% truth_vec_s] sex <- tibble( var = "sex", TP = length(s_intersect), FP = length(detected_sex) - length(s_intersect), FN = length(truth_vec_s) - length(s_intersect), precision = TP / (TP + FP), recall = TP / (TP + FN) )

truth_vec_t <- truth |> filter(DA_Time == TRUE) |> pull(Taxon) t_intersect <- detected_time[detected_time %in% truth_vec_t] time <- tibble( var = "time", TP = length(t_intersect), FP = length(detected_time) - length(t_intersect), FN = length(truth_vec_t) - length(t_intersect), precision = TP / (TP + FP), recall = TP / (TP + FN) )

truth_vec_i <- truth |> filter(DA_Interaction == TRUE) |> pull(Taxon) i_intersect <- detected_interaction[detected_interaction %in% truth_vec_i] interaction <- tibble( var = "interaction", TP = length(i_intersect), FP = length(detected_interaction) - length(i_intersect), FN = length(truth_vec_i) - length(i_intersect), precision = TP / (TP + FP), recall = TP / (TP + FN) )

bind_rows(disease, sex, time, interaction) }

sim_detection <- function(sim_counts, sim_meta, truth) {

sim_meta <- sim_meta |> mutate( Sex = factor(Sex), Disease = factor(Disease), Time = factor(Time), MouseID = factor(ID) ) rownames(sim_meta) <- sim_meta$SampleID

#——————————————— ALDEx2 model_matrix <- model.matrix(~ Sex + Disease + Time, data = sim_meta)

x <- aldex.clr(sim_counts, conds = model_matrix, mc.samples = 128)

glm_test <- aldex.glm(x) glm_effect <- aldex.glm.effect(x)

aldex_res <- cbind(glm_test, glm_effect)

res_noID <- aldex_res |> dplyr::select(contains("pval"), -contains("Intercept"))

colnames(res_noID) <- str_remove_all(colnames(res_noID), "M|Disease|Pre")

#——————————————— ANCOM-BC2 counts_mat <- as.matrix(sim_counts)

ancom_res <- ancombc2( data = counts_mat, meta_data = sim_meta, fix_formula = "Sex + Time + Disease", rand_formula = NULL, p_adj_method = "BH", prv_cut = 0.1, lib_cut = 0, group = NULL, struc_zero = FALSE, neg_lb = TRUE )

res_ancom <- ancom_res$res

ancom_effects <- res_ancom |> dplyr::select(taxon, contains("diff_robust"), -contains("Intercept")) colnames(ancom_effects) <- str_remove_all(colnames(ancom_effects), “M|Disease|Pre”)

#——————————— PARALLAX result_table <- runmypipeline(sim_counts, sim_meta)

parallax_res <- result_table |> dplyr::filter(Coefficient != “X.Intercept.”) |> mutate( Coefficient = str_remove_all(Coefficient, “F|Disease|Post”), is.DA = ifelse((q_post <= 0.05 & R_scale >= 0.8), “TRUE”, “FALSE”) )

parallax_res_f <- parallax_res |> dplyr::filter(is.DA == “TRUE”)

p_disease <- parallax_res_f |> dplyr::filter(Coefficient == “Disease”) |> pull(Taxon) p_sex <- parallax_res_f |> dplyr::filter(Coefficient == “Sex”) |> pull(Taxon) p_time <- parallax_res_f |> dplyr::filter(Coefficient == “Time”) |> pull(Taxon) p_interaction <- parallax_res_f |> dplyr::filter(Coefficient == “Disease.Time”) |> pull(Taxon)

res_noID <- rownames_to_column(res_noID, var = “Taxon”) al_disease <- res_noID |> dplyr::filter(`Disease:pval` <= 0.05) |> pull(Taxon) al_sex <- res_noID |> dplyr::filter(`Sex:pval` <= 0.05) |> pull(Taxon) al_time <- res_noID |> dplyr::filter(`Time:pval` <= 0.05) |> pull(Taxon) al_interaction <- NULL

an_disease <- ancom_effects |> dplyr::filter(diff_robust_Disease == TRUE) |> pull(taxon) an_sex <- ancom_effects |> dplyr::filter(diff_robust_Sex == TRUE) |> pull(taxon) an_time <- ancom_effects |> dplyr::filter(diff_robust_Time == TRUE) |> pull(taxon) an_interaction <- NULL

detection_acc_p <- detection_accuracy(truth, p_disease, p_sex, p_time, p_interaction) detection_acc_al <- detection_accuracy(truth, al_disease, al_sex, al_time, al_interaction) detection_acc_an <- detection_accuracy(truth, an_disease, an_sex, an_time, an_interaction)

list(parallax = detection_acc_p, ancom = detection_acc_an, aldex2 = detection_acc_al) }

#==========================================================
### 4. Replicated benchmark: many independent master cohorts x N x DA x effect # # FEATURES: # # * Everything is wrapped in an outer `for (rep_id in 1:n_reps)` loop. # Each replicate draws a FRESH pair of master cohorts (DA=10, DA=20) with # their own seeds, so N = 20/40/60/80 are nested nested subsets # *within* a replicate, but the taxon identities, effect sizes, and # (now) NB dispersion / zero-inflation parameters vary *across* # replicates. That’s what lets you summarize sensitivity to the specific # random realization, and gives you a real empirical false-positive rate # under the null cells (e.g. wherever a factor’s true DA set is empty) # rather than a single point estimate. # # * Results are written incrementally (appended to disk after every # replicate x N x effect x method combo) rather than only at the very # end. With n_reps = 100 and PARALLAX alone fitting sigma_grid(5) x # mc.samples(128) = 640 lmFit models per run, a full pass here is a # multi-hour-to-multi-day job on a single machine – you do not want to # lose 99 replicates of progress to a crash

on the 100th. # # * distribution is fixed to “nb” for both master cohorts below. Switch to # “zinb” (and tune zero_infl_prob / zero_infl_conc) if you want to also # stress-test the zero-inflated case reviewers mentioned; keep # “poisson” only if you specifically want a Poisson comparison arm. # # PRACTICAL NOTE ON RUNTIME: before committing to n_reps = 100, run with a # small n_reps (e.g. 2-3) first and time it, so you know what a full 100-rep # pass will cost. Consider also reducing mc.samples in aldex.clr()/ # runmypipeline() for the exploratory pass, or parallelizing the rep_id loop # across cores (e.g. with parallel::mclapply or the furrr package) once # you've confirmed the pipeline behaves as expected. #==========================================================================

library(parallel) library(BiocParallel)

N_values <- c(20, 40, 60, 80) n_reps <- 100

base_seed_DA20 <- 123 base_seed_DA10 <- 456

—- parallel setup ————————————————————————

I'm using macOS (per the setwd() path above), so I use fork-based

parallelism via mclapply(). Forking is memory-cheap up front: child

processes share the parent's memory copy-on-write, so `counts` and the

loaded packages aren't duplicated just by forking.

The RAM cost instead comes from what EACH replicate allocates while it

runs: ALDEx2's CLR Monte Carlo arrays, 640 limma fit objects inside

runmypipeline(), and ANCOM-BC2's internals. With 16 GB total (and macOS,

the browser, etc. already using some of that), running many replicates

at once compounds those per-replicate peaks.

Two safety choices below address this directly:

1. n_cores defaults to a conservative value, not detectCores()-1.

Start here; only raise it if you watch Activity Monitor during a

short run and see comfortable headroom.

2. mc.preschedule = FALSE means each replicate gets its OWN forked

process that exits (and returns its memory to the OS) as soon as run_replicate <- function(rep_id) {

### Force ancombc2 (and anything else BiocParallel-aware) to stay serial # *inside* this worker. Without this, ancombc2 could try to spin up its # own internal parallel workers on top of the mclapply fork, multiplying # memory/CPU use well beyond n_cores. BiocParallel::register(BiocParallel::SerialParam())

seed_DA20 <- base_seed_DA20 + rep_id * 97 seed_DA10 <- base_seed_DA10 + rep_id * 131

master_DA20 <- build_master_cohort( counts, n_subjects_max = 80, n_DA = 20, seed = seed_DA20, distribution = "nb", nb_size_mean = 5, nb_size_logsd = 0.4 ) master_DA10 <- build_master_cohort( counts, n_subjects_max = 80, n_DA = 10, seed = seed_DA10, distribution = "nb", nb_size_mean = 5, nb_size_logsd = 0.4 )

DA_masters <- list(`10` = master_DA10, `20` = master_DA20)

rep_results <- list()

for (da_label in names(DA_masters)) {

```
master <- DA_masters[[da_label]]

for (n in N_values) {

  cohort <- subsample_cohort(master, n)


    res <- sim_detection(cohort$counts, cohort$metadata, cohort$truth)

    for (method in names(res)) {
      acc <- res[[method]]
      acc$N <- n
      acc$DA_number <- as.integer(da_label)
      acc$Effect_size <- "Baseline"
      acc$source <- method
      acc$replicate <- rep_id
      rep_results[[length(rep_results) + 1]] <- acc
    }
}
```

}

out <- bind_rows(rep_results) |> mutate(precision = replace(precision, is.nan(precision), 0))

### Belt-and-suspenders: with mc.preschedule = FALSE this process is about # to exit anyway (returning all its memory to the OS), but explicitly # dropping the big objects and forcing a full GC first costs nothing. rm(master_DA20, master_DA10, DA_masters, rep_results) gc(full = TRUE)

out }

out_file <- “results_base.csv” # EDIT FILENAME if (file.exists(out_file)) file.remove(out_file)

#### Process replicates in batches of size n_cores, and checkpoint to disk

#### after each batch. This bounds how many replicates' worth of results sit

#### unsaved in memory at once, and means a crash mid-run only costs you the

#### current (small) batch, not the whole job.

rep_batches <- split(seq_len(n_reps), ceiling(seq_len(n_reps) / n_cores))

all_results <- list()

for (batch in rep_batches) {

message(sprintf(“### Replicates %d-%d of %d ###”, min(batch), max(batch), n_reps))

if (.Platform$OS.type == “windows”) { # No forking on Windows; falls back to sequential. A PSOCK cluster # (parallel::makeCluster + parLapply) is possible instead, but each # worker there gets its own full copy of `counts` and loaded packages # via serialization – meaningfully more RAM-hungry than forking, so # it isn't the default recommendation on a 16 GB machine. batch_out <- lapply(batch, run_replicate) } else { batch_out <- parallel::mclapply( batch, run_replicate, mc.cores = n_cores, mc.preschedule = FALSE ) }

### A worker that throws a normal R error returns a “try-error” object. # A worker that CRASHES (e.g. a segfault from fork + threaded BLAS) # instead returns NULL, with no catchable error at all – checking only # for “try-error” misses this and lets bind_rows() silently drop it. # Both cases are treated as failures here, and the real message (when # there is one) is printed so you're not debugging blind. ok <- vapply(batch_out, function(x) !is.null(x) && !inherits(x, “try-error”), logical(1)) if (!all(ok)) { for (i in which(!ok)) { rid <- batch[i]

res <- batch_out[[i]] if (is.null(res)) { message(sprintf("Replicate %d: worker returned NULL (likely crashed/segfaulted, not a normal R error)", rid)) } else { message(sprintf("Replicate %d failed: %s", rid, conditionMessage(attr(res, "condition")))) } } warning(sprintf("Replicate(s) %s failed in this batch", paste(batch[!ok], collapse = ","))) }

batch_df <- bind_rows(batch_out[ok]) write_csv(batch_df, out_file, append = file.exists(out_file)) all_results[[length(all_results) + 1]] <- batch_df }

res_cleaned <- bind_rows(all_results)

### res_cleaned now has one row per (replicate x DA_number x N x Effect_size x

### source x var), so false-positive behavior for a given cell can be

### summarized e.g. as:

### res_cleaned |>

### filter(source == "parallax", var == "disease") |>

### group_by(N, DA_number, Effect_size) |>

### summarise(mean_FP = mean(FP), mean_precision = mean(precision), .groups = "drop")

#=================================================================
### Run the following for REDUCED effect simulations #==================================

#=================================================================
### 0. Re/load real data # # NOTE: only rownames(counts) (the taxon names) are used, to sample a # realistic pool of taxa for the simulation. meta_sim2 is loaded for # completeness but is not consumed by the simulation itself. #==================================
### Change filenames accordingly. Again, set for N = 40 counts_raw <- read_csv("counts.csv") %>% dplyr::select(-1) %>% # drop leading numeric row-index column as.matrix()

counts <- counts_raw rownames(counts) <- paste0("Genus_", seq_len(nrow(counts_raw)))

meta_sim2 <- read_csv("meta.csv") %>% dplyr::select(-1)

#========================================================================
### 1. Corrected simulation functions # (build_master_cohort / subsample_cohort / small_effect) # # FEATURES: # # (a) Count generation supports "poisson" (old behavior), "nb" # (negative binomial, default), or "zinb" (zero-inflated negative # binomial), via the `distribution` argument. # # (b) NB overdispersion is TAXON-SPECIFIC: each taxon gets its own NB # "size" parameter (size = 1 / dispersion), drawn once per master # cohort (Stream 1, seeded, independent of N) from a log-normal # distribution controlled by `nb_size_mean` / `nb_size_logsd`. # Smaller size => more overdispersion (variance = mu + mu^2/size). # # (c) If distribution == "zinb", each taxon also gets a structural # zero-inflation probability, drawn from a Beta distribution # controlled by `zero_infl_prob` (mean) and `zero_infl_conc` # (concentration -> higher = less taxon-to-taxon variability in # the zero-inflation rate). # # (d) build_master_cohort() returns `size_taxon` and `pi_zero` in # the output list, so you can inspect/reuse the dispersion and # zero-inflation parameters that were actually drawn for a given # master cohort.
#========================================================================

#### —- helper: single NB / ZINB draw for one sample's taxon vector ————–

simulate_counts_1sample <- function(mu, distribution, size_taxon, pi_zero) {

n_taxa <- length(mu)

counts <- switch( distribution, poisson = rpois(n_taxa, lambda = mu), nb = rnbinom(n_taxa, mu = mu, size = size_taxon), zinb = rnbinom(n_taxa, mu = mu, size = size_taxon), stop("distribution must be one of 'poisson', 'nb', 'zinb' ") )

if (distribution == "zinb") { structural_zero <- rbinom(n_taxa, 1, pi_zero) counts[structural_zero == 1] <- 0 }

counts }

build_master_cohort <- function( counts, n_subjects_max = 80, n_taxa = 120, n_DA = 20, prop_high_load = 0.2, seed = 123, distribution = c("nb", "poisson", "zinb"), nb_size_mean = 5, # mean NB size (dispersion) across taxa; lower = more overdispersed nb_size_logsd = 0.4, # log-scale SD -> taxon-to-taxon variability in dispersion nb_size_floor = 0.1, # hard floor so no taxon gets an absurdly tiny size zero_infl_prob = 0.1, # mean structural zero-inflation probability (zinb only) zero_infl_conc = 5, # Beta concentration for zero_infl_prob (zinb only) effect_scale = 1 ) {

distribution <- match.arg(distribution)

strata <- expand.grid(Disease = c(“DIsease”, “Control”), Sex = c(“M”, “F”), stringsAsFactors = FALSE) n_strata <- nrow(strata) # 4

stopifnot( “n_subjects_max must be divisible by the number of Disease x Sex strata (4)” = (n_subjects_max %% n_strata) == 0 ) n_per_stratum <- n_subjects_max / n_strata

### —- Stream 1 (seed): taxon-level ground truth — independent of N —- set.seed(seed)

taxa_all <- rownames(counts) taxa <- sample(taxa_all, n_taxa)

prop_disease <- 0.35 prop_time <- 0.25 prop_sex <- 0.20 prop_interaction <- 0.20

n_DA_disease <- round(n_DA * prop_disease) n_DA_time <- round(n_DA * prop_time) n_DA_sex <- round(n_DA * prop_sex) n_used <- n_DA_disease + n_DA_time + n_DA_sex n_DA_interaction <- n_DA - n_used

idx <- sample(seq_len(n_taxa))

start <- 1 DA_disease <- idx[start:(start + n_DA_disease - 1)] start <- start + n_DA_disease

DA_time <- idx[start:(start + n_DA_time - 1)] start <- start + n_DA_time

DA_sex <- idx[start:(start + n_DA_sex - 1)] start <- start + n_DA_sex

DA_interaction <- idx[start:(start + n_DA_interaction - 1)]

effect_disease <- runif(length(DA_disease), 1.0, 1.5) effect_time <- runif(length(DA_time), 1.0, 1.5) effect_sex <- runif(length(DA_sex), 1.0, 1.5) effect_interaction <- runif(length(DA_interaction), 1.0, 1.5)

base <- rnorm(n_taxa, mean = 0, sd = 1)

### —- taxon-specific overdispersion (and zero-inflation, if requested) —- # Drawn in the same seeded stream as the rest of the taxon-level ground # truth, so they stay tied to the taxon identities/effects rather than to N. size_taxon <- exp(rnorm(n_taxa, mean = log(nb_size_mean), sd = nb_size_logsd)) size_taxon <- pmax(size_taxon, nb_size_floor)

if (distribution == “zinb”) { a <- zero_infl_prob * zero_infl_conc b <- (1 - zero_infl_prob) * zero_infl_conc pi_zero <- rbeta(n_taxa, a, b) } else { pi_zero <- rep(0, n_taxa) }

### —- Stream 2 (seed + 1000): balanced subject allocation + nested rank —- set.seed(seed + 1000)

subject_table <- do.call(rbind, lapply(seq_len(n_strata), function(s) { data.frame( Disease = strata$Disease[s], Sex = strata$Sex[s], nested_rank = sample(seq_len(n_per_stratum)), stringsAsFactors = FALSE ) })) subject_table <- subject_table[order(subject_table$Disease, subject_t able$Sex, subject_table$nested_r ank), ]subject_t able$ID <- paste0(“ID_”, seq_len(n_subjects_max)) rownames(subject_table) <- NULL sim_meta <- expand.grid(ID = subject_table$ID, Time = c("Pre", "Post"), stringsAsFactors = FALSE) sim_m eta$Disease <- subject_table$Disease[match(sim_m eta$ID, subject_table$ID)] sim_m eta$Sex <- subject_table$Sex[match(sim_m eta$ID, subject_table$ID)] sim_m eta$SampleID <- paste0("Sample", seq_len(nrow(sim_meta)))

### —- Stream 3 (seed + 2000): count generation —- set.seed(seed + 2000)

counts_mat <- matrix(0, nrow = n_taxa, ncol = nrow(sim_meta)) rownames(counts_mat) <- taxa

depth <- 20000 n_high_load <- round(prop_high_load * nrow(sim_meta)) high_load_idx <- sample(seq_len(nrow(sim_meta)), n_high_load)

for (i in seq_len(nrow(sim_meta))) {

```
row <- sim_meta[i, ]
log_theta <- base

if (row$Disease == "Disease") {
  log_theta[DA_disease] <- log_theta[DA_disease] + effect_scale * effect_disease
} else {
  log_theta[DA_disease] <- log_theta[DA_disease] - effect_scale * effect_disease
}

if (row$Time == "Post") {
  log_theta[DA_time] <- log_theta[DA_time] + effect_scale * effect_time
} else {
  log_theta[DA_time] <- log_theta[DA_time] - effect_scale * effect_time
}

if (row$Sex == "F") {
  log_theta[DA_sex] <- log_theta[DA_sex] + effect_scale * effect_sex
} else {
  log_theta[DA_sex] <- log_theta[DA_sex] - effect_scale * effect_sex
}

if (row$Disease == "Disease" && row$Time == "Post") {
  log_theta[DA_interaction] <- log_theta[DA_interaction] + effect_scale * effect_interaction
}

theta <- exp(log_theta)
theta <- theta / sum(theta)

depth_i <- depth
```

```
if (i %in% high_load_idx) {
  depth_i <- depth_i * runif(1, 2, 4)
}

mu_i <- theta * depth_i

counts_mat[, i] <- simulate_counts_1sample(
  mu           = mu_i,
  distribution = distribution,
  size_taxon   = size_taxon,
  pi_zero      = pi_zero
)
```

}

colnames(counts_mat) <- sim_meta$SampleID

truth <- data.frame( Taxon = taxa, DA_Disease = seq_len(n_taxa) %in% DA_disease, DA_Time = seq_len(n_taxa) %in% DA_time, DA_Sex = seq_len(n_taxa) %in% DA_sex, DA_Interaction = seq_len(n_taxa) %in% DA_interaction ) truth$is.DA <- rowSums(truth[, 2:5]) > 0

list( counts = counts_mat, metadata = sim_meta, truth = truth, subject_table = subject_table, n_strata = n_strata, n_subjects_max = n_subjects_max, distribution = distribution, size_taxon = setNames(size_taxon, taxa), pi_zero = setNames(pi_zero, taxa) ) }

subsample_cohort <- function(master, n_subjects) {

stopifnot( “n_subjects must be divisible by the number of Disease x Sex strata (4)” = (n_subjects %% master$n_strata) == 0, "n_subjects cannot exceed the master cohort size" = n_subjects <= master$n_subjects_max )

per_stratum <- n_subjects / master$n_strata keep_ids <- master$subject_table$ID[master$subject_table$nested_rank <= per_stratum]

meta_sub <- master$metadata[master$metadata$ID meta_sub <- meta_sub[order(meta_sub$ID, meta_sub$Time), ]

counts_sub <- master$counts[, meta_sub$SampleID, drop = FALSE]

list( counts = counts_sub, metadata = meta_sub, truth = master$truth ) }

#==========================================================
### 2. PARALLAX pipeline (unchanged) #==========================================

runmypipeline <- function(counts, meta) {

x <- aldex.clr(counts, conds = rep("A", ncol(counts)), mc.samples = 128, denom = "all") mc__list <- getMonteCarloInstances(x) n__samples <- ncol(counts) n__features <- nrow(counts) M <- ncol(mc__list[[1]])

names(mc__list) <- colnames(counts) M <- ncol(mc__list[[1]])

clr__mats <- lapply(seq__len(M), function(mc) { mat <- sapply(mc__list, function(sample__mat) { vec <- sample__mat[, mc] vec__aligned <- vec[rownames(counts)] return(vec__aligned) }) mat <- as.matrix(mat) rownames(mat) <- rownames(counts) colnames(mat) <- colnames(counts) mat })

clr__array <- simplify2array(clr__mats) clr__mean <- apply(clr__array, c(1, 2), mean) clr__mean__t <- t(clr__mean)

meta <- meta %>% mutate( Sex = factor(Sex, levels = c("M", "F")), Disease = factor(Disease, levels = c("Control", "Disease")), Time = factor(Time, levels = c("Pre", "Post")), MouseID = factor(ID) )

if (!all(colnames(counts) %in% rownames(meta))) { stop("Metadata rows do not match count columns") }

meta <- meta[match(colnames(counts), rownames(meta)), , drop = FALSE] rownames(meta) <- colnames(counts)

if (any(is.na(meta))) { stop("NA values introduced in metadata alignment") }

mf <- model.frame(~ (Sex + Disease + Time)^2, data = meta, na.action = na.fail) design <- model.matrix(~ (Sex + Disease + Time)^2, data = mf) rownames(design) <- colnames(counts) colnames(design) <- make.names(colnames(design))

sigma__grid <- c(0, 0.25, 0.5, 0.75, 1)

expr__ref <- clr__mats[[1]] corfit <- duplicateCorrelation(expr__ref, design, block = meta$ID)consensus_cor < -corfit$consensus

fit__list__all <- vector("list", length(sigma__grid) * M) idx <- 1

for (sigma in sigma__grid) { message("Running sigma =", sigma) for (mc in seq__len(M)) { expr <- clr__mats[[mc]] stopifnot(ncol(expr) == nrow(design)) stopifnot(all(colnames(expr) == rownames(design)))

```
  log_s <- rnorm(ncol(expr), mean = 0, sd = sigma)
  expr <- sweep(expr, 2, log_s, FUN = "+")

  fit <- lmFit(expr, design, block = meta$ID, correlation = consensus_cor)
  fit <- eBayes(fit)

  fit_list_all[[idx]] <- fit
```

```
  idx <- idx + 1
}
```

}

coef_names <- colnames(fit_list_all[[1]]$coefficients) n_coef <- length(coef_names) n_features <- nrow(counts)

result_table <- data.frame( Coefficient = rep(coef_names, each = n_features), Taxon = rep(rownames(counts), times = n_coef), logFC_median = NA, P_post = NA, q_post = NA, R_scale = NA )

n_sigma <- length(sigma_grid) rep_per_sigma <- M sigma_labels <- rep(seq_len(n_sigma), each = rep_per_sigma)

row_idx <- 1

for (coef_idx in seq_len(n_coef)) {

```
theta_array <- do.call(cbind, lapply(fit_list_all, function(fit) {
  vec <- fit$coefficients[, coef_idx]
  vec[is.infinite(vec)] <- NA
  vec
}))
vals <- as.vector(theta_array)

delta <- log(1.2)

P_pos <- rowMeans(theta_array > delta, na.rm = TRUE)
P_neg <- rowMeans(theta_array < -delta, na.rm = TRUE)

P_post <- pmax(P_pos, P_neg)
q_post <- 1 - P_post

logFC_median <- apply(theta_array, 1, median, na.rm = TRUE)

R_scale_vec <- apply(theta_array, 1, function(theta_vec) {
  theta_by_sigma <- split(theta_vec, sigma_labels)
  sig_sigma <- sapply(theta_by_sigma, function(vec) {
    max(mean(vec > delta), mean(vec < -delta))
  })
  mean(sig_sigma)
})
```

```
result_table$logFC_median[row_idx:(row_idx + n_features - 1)] <- logFC_median
result_table$P_post[row_idx:(row_idx + n_features - 1)] <- P_post
result_table$q_post[row_idx:(row_idx + n_features - 1)] <- q_post
result_table$R_scale[row_idx:(row_idx + n_features - 1)] <- R_scale_vec

row_idx <- row_idx + n_features
```

} result_table }

#==================================================================================
### 3. Detection accuracy + sim_detection # # `truth` is passed explicitly (not a global), since with the replicate loop # below there are many distinct ground-truth tables in flight at once.
#==================================================================================

detection_accuracy <- function(truth, detected_disease, detected_sex, detected_time, detected_interaction) {

truth_vec_d <- truth |> filter(DA_Disease == TRUE) |> pull(Taxon) d_intersect <- detected_disease[detected_disease %in% truth_vec_d] disease <- tibble( var = "disease", TP = length(d_intersect), FP = length(detected_disease) - length(d_intersect), FN = length(truth_vec_d) - length(d_intersect), precision = TP / (TP + FP), recall = TP / (TP + FN) )

truth_vec_s <- truth |> filter(DA_Sex == TRUE) |> pull(Taxon) s_intersect <- detected_sex[detected_sex %in% truth_vec_s] sex <- tibble( var = "sex", TP = length(s_intersect), FP = length(detected_sex) - length(s_intersect), FN = length(truth_vec_s) - length(s_intersect), precision = TP / (TP + FP), recall = TP / (TP + FN) )

truth_vec_t <- truth |> filter(DA_Time == TRUE) |> pull(Taxon) t_intersect <- detected_time[detected_time %in% truth_vec_t] time <- tibble( var = "time", TP = length(t_intersect), FP = length(detected_time) - length(t_intersect), FN = length(truth_vec_t) - length(t_intersect), precision = TP / (TP + FP), recall = TP / (TP + FN) )

truth_vec_i <- truth |> filter(DA_Interaction == TRUE) |> pull(Taxon) i_intersect <- detected_interaction[detected_interaction %in% truth_vec_i] interaction <- tibble( var = "interaction", TP = length(i_intersect), FP = length(detected_interaction) - length(i_intersect), FN = length(truth_vec_i) - length(i_intersect), precision = TP / (TP + FP), recall = TP / (TP + FN) )

bind_rows(disease, sex, time, interaction) }

sim_detection <- function(sim_counts, sim_meta, truth) {

sim_meta <- sim_meta |> mutate( Sex = factor(Sex), Disease = factor(Disease), Time = factor(Time), MouseID = factor(ID) ) rownames(sim_meta) <- sim_meta$SampleID

#———————————— ALDEx2 model_matrix <- model.matrix(~ Sex + Disease + Time, data = sim_meta)

x <- aldex.clr(sim_counts, conds = model_matrix, mc.samples = 128)

glm_test <- aldex.glm(x) glm_effect <- aldex.glm.effect(x)

aldex_res <- cbind(glm_test, glm_effect)

res_noID <- aldex_res |> dplyr::select(contains("pval"), -contains("Intercept"))

colnames(res_noID) <- str_remove_all(colnames(res_noID), "M|Disease|Pre")

#———————————— ANCOM-BC2 counts_mat <- as.matrix(sim_counts)

ancom_res <- ancombc2( data = counts_mat, meta_data = sim_meta, fix_formula = "Sex + Time + Disease", rand_formula = NULL, p_adj_method = "BH", prv_cut = 0.1, lib_cut = 0, group = NULL, struc_zero = FALSE, neg_lb = TRUE )

res_ancom <- ancom_res$res

ancom_effects <- res_ancom |> dplyr::select(taxon, contains("diff_robust"), -contains("Intercept"))

colnames(ancom_effects) <- str_remove_all(colnames(ancom_effects), "M|Disease|Pre")

#———————————— PARALLAX result_table <- runmypipeline(sim_counts, sim_meta)

parallax_res <- result_table |> dplyr::filter(Coefficient != "X.Intercept.") |> mutate( Coefficient = str_remove_all(Coefficient, "F|Disease|Post"), is.DA = ifelse((q_post <= 0.05 & R_scale >= 0.8), "TRUE", "FALSE") )

parallax_res_f <- parallax_res |> dplyr::filter(is.DA == "TRUE")

p_disease <- parallax_res_f |> dplyr::filter(Coefficient == "Disease") |> pull(Taxon) p_sex <- parallax_res_f |> dplyr::filter(Coefficient == "Sex") |> pull(Taxon) p_time <- parallax_res_f |> dplyr::filter(Coefficient == "Time") |> pull(Taxon) p_interaction <- parallax_res_f |> dplyr::filter(Coefficient == "Disease.Time") |> pull(Taxon)

res_noID <- rownames_to_column(res_noID, var = "Taxon") al_disease <- res_noID |> dplyr::filter(`Disease:pval` <= 0.05) |> pull(Taxon) al_sex <- res_noID |> dplyr::filter(`Sex:pval` <= 0.05) |> pull(Taxon) al_time <- res_noID |> dplyr::filter(`Time:pval` <= 0.05) |> pull(Taxon) al_interaction <- NULL

an_disease <- ancom_effects |> dplyr::filter(diff_robust_Disease == TRUE) |> pull(taxon) an_sex <- ancom_effects |> dplyr::filter(diff_robust_Sex == TRUE) |> pull(taxon) an_time <- ancom_effects |> dplyr::filter(diff_robust_Time == TRUE) |> pull(taxon) an_interaction <- NULL

detection_acc_p <- detection_accuracy(truth, p_disease, p_sex, p_time, p_interaction) detection_acc_al <- detection_accuracy(truth, al_disease, al_sex, al_time, al_interaction) detection_acc_an <- detection_accuracy(truth, an_disease, an_sex, an_time, an_interaction)

list(parallax = detection_acc_p, ancom = detection_acc_an, aldex2 = detection_acc_al) }

#==================================================================== # 4. Replicated benchmark: many independent master cohorts x N x DA x effect # # FEATURES: # # * Everything is wrapped in an outer `for (rep_id in 1:n_reps)` loop. # Each replicate draws a FRESH pair of master cohorts (DA=10, DA=20) with # their own seeds, so N = 20/40/60/80 are nested nested subsets # *within* a replicate, and the taxon identities, effect sizes, and # (now) NB dispersion / zero-inflation parameters vary *across* # replicates. That's what lets you summarize sensitivity to the specific # random realization, and gives you a real empirical false-positive rate # under the null cells (e.g. wherever a factor's true DA set is empty) # rather than a single point estimate. # # * Results are written incrementally (appended to disk after every # replicate x N x effect x method combo) rather than only at the very # end. With n_reps = 100 and PARALLAX alone fitting sigma_grid(5) x # mc.samples(128) = 640 lmFit models per run, a full pass here is a # multi-hour-to-multi-day job on a single machine – you do not want to # lose 99 replicates of progress to a crash on the 100th. # # * distribution is fixed to "nb" for both master cohorts below. Switch to # "zinb" (and tune zero_infl_prob / zero_infl_conc) if you want to also # stress-test the zero-inflated case reviewers mentioned; keep # "poisson" only if you specifically want a Poisson comparison arm. # # PRACTICAL NOTE ON RUNTIME: before committing to n_reps = 100, run with a # small n_reps (e.g. 2-3) first and time it, so you know what a full 100-rep # pass will cost. Consider also reducing mc.samples in aldex.clr()/ # runmypipeline() for the exploratory pass, or parallelizing the rep_id loop # across cores (e.g. with parallel::mclapply or the furrr package) once # you've confirmed the pipeline behaves as expected. #====================================================================

library(parallel) library(BiocParallel)

N_values <- c(20, 40, 60, 80) n_reps <- 100

base_seed_DA20 <- 123 base_seed_DA10 <- 456

—- parallel setup ————————————————————————————

I'm on macOS (per the setwd() path above), so I use fork-based

parallelism via mclapply(). Forking is memory-cheap up front: child

processes share the parent's memory copy-on-write, so `counts` and the

loaded packages aren't duplicated just by forking.

The RAM cost instead comes from what EACH replicate allocates while it

runs: ALDEx2's CLR Monte Carlo arrays, 640 limma fit objects inside

runmypipeline(), and ANCOM-BC2's internals. With 16 GB total (and macOS,

the browser, etc. already using some of that), running many replicates

at once compounds those per-replicate peaks.

Two safety choices below address this directly:

1. n_cores defaults to a conservative value, not detectCores()-1.

Start here; only raise it if you watch Activity Monitor during a

short run and see comfortable headroom.

2. mc.preschedule = FALSE means each replicate gets its OWN forked

process that exits (and returns its memory to the OS) as soon as run_replicate <- function(rep_id) {

### Force ancombc2 (and anything else BiocParallel-aware) to stay serial # *inside* this worker. Without this, ancombc2 could try to spin up its # own internal parallel workers on top of the mclapply fork, multiplying # memory/CPU use well beyond n_cores. BiocParallel::register(BiocParallel::SerialParam())

seed_DA20 <- base_seed_DA20 + rep_id * 97 seed_DA10 <- base_seed_DA10 + rep_id * 131

master_DA20 <- build_master_cohort( counts, n_subjects_max = 80, n_DA = 20, seed = seed_DA20, distribution = "nb", nb_size_mean = 5, nb_size_logsd = 0.4, effect_scale = 0.5 ) master_DA10 <- build_master_cohort( counts, n_subjects_max = 80, n_DA = 10, seed = seed_DA10, distribution = "nb", nb_size_mean = 5, nb_size_logsd = 0.4, effect_scale = 0.5 )

DA_masters <- list(`10` = master_DA10, `20` = master_DA20)

rep_results <- list()

for (da_label in names(DA_masters)) {

```
master <- DA_masters[[da_label]]

for (n in N_values) {

  cohort <- subsample_cohort(master, n)

  res <- sim_detection(cohort$counts, cohort$metadata, cohort$truth)

  for (method in names(res)) {
    acc <- res[[method]]
    acc$N <- n
    acc$DA_number <- as.integer(da_label)
    acc$Effect_size <- "Small"
    acc$source <- method
    acc$replicate <- rep_id
    rep_results[[length(rep_results) + 1]] <- acc
  }
}
```

}

out <- bind_rows(rep_results) |> mutate(precision = replace(precision, is.nan(precision), 0))

### Belt-and-suspenders: with mc.preschedule = FALSE this process is about # to exit anyway (returning all its memory to the OS), but explicitly # dropping the big objects and forcing a full GC first costs nothing. rm(master_DA20, master_DA10, DA_masters, rep_results) gc(full = TRUE)

out }

out_file <- "results_small.csv" # EDIT FILENAME if (file.exists(out_file)) file.remove(out_file)

### Process replicates in batches of size n_cores, and checkpoint to disk

### after each batch. This bounds how many replicates' worth of results sit

### unsaved in memory at once, and means a crash mid-run only costs you the

### current (small) batch, not the whole job.

rep_batches <- split(seq_len(n_reps), ceiling(seq_len(n_reps) / n_cores))

all_results <- list()

for (batch in rep_batches) {

message(sprintf("### Replicates %d-%d of %d ###", min(batch), max(batch), n_reps))

if (.Platform$OS.type == "windows") { # No forking on Windows; falls back to sequential. A PSOCK cluster # (parallel::makeCluster + parLapply) is possible instead, but each # worker there gets its own full copy of `counts` and loaded packages # via serialization – meaningfully more RAM-hungry than forking, so # it isn't the default recommendation on a 16 GB machine. batch_out <- lapply(batch, run_replicate) } else { batch_out <- parallel::mclapply( batch, run_replicate, mc.cores = n_cores, mc.preschedule = FALSE ) }

### A worker that throws a normal R error returns a "try-error" object. # A worker that CRASHES (e.g. a segfault from fork + threaded BLAS) # instead returns NULL, with no catchable error at all – checking only # for "try-error" misses this and lets bind_rows() silently drop it. # Both cases are treated as failures here, and the real message (when # there is one) is printed so you're not debugging blind. ok <- vapply(batch_out, function(x) !is.null(x) && !inherits(x, "try-error"), logical(1)) if (!all(ok)) { for (i in which(!ok)) { rid <- batch[i]

res <- batch_out[[i]] if (is.null(res)) { message(sprintf("Replicate %d: worker returned NULL (likely crashed/segfaulted, not a normal R error)", rid)) } else { message(sprintf("Replicate %d failed: %s", rid, conditionMessage(attr(res, "condition")))) } } warning(sprintf("Replicate(s) %s failed in this batch", paste(batch[!ok], collapse = ","))) }

batch_df <- bind_rows(batch_out[ok]) write_csv(batch_df, out_file, append = file.exists(out_file)) all_results[[length(all_results) + 1]] <- batch_df }

res_cleaned <- bind_rows(all_results)

#### import the filtered baseline sims generated in the original V3 sim

base <- read.csv("results_base.csv")

#### combine the baseline sims with the small effect sims

all_res <- bind_rows(res_cleaned, base)

write.csv(all_res, "results.csv")

#========================================== # VISUALIZATION #========================================== all_res_averaged <- all_res |> group_by( N, DA_number, Effect_size, source, var ) |> summarize( mean_precision = mean(precision), ci_precision = 1.96 * sd(precision) / sqrt(n()),

```
mean_recall = mean(recall),
ci_recall = 1.96 * sd(recall) / sqrt(n()),

.groups = "drop"
```

)

all_res_plot <- all_res_averaged %>% mutate( N = str_c("N =", N), N = factor(N, levels = str_c("N =", N_values)), source = recode(source, aldex2 = "ALDEx2", ancom = "ANCOM-BC2", parallax = "PARALLAX"), Effect_size = str_c(Effect_size, " effect") )

covariate_plot <- function(df, var_name) { ggplot(df %>% filter(var == var_name), aes(x = DA_number, group = source)) + geom_line(aes(y = mean_precision, color = source), linewidth = 1.2) + geom_point(aes(y = mean_precision, color = source), size = 2) + geom_point(aes(y = mean_recall, color = "black"), shape = 17, size = 3) + geom_errorbar( aes( ymin = mean_precision - ci_precision, ymax = mean_precision + ci_precision ), color

= “black”, width = 0.5 ) + facet_grid(Effect_size ~ N) + scale_x_continuous(breaks = c(10, 20)) + scale_color_manual(values = c( “ALDEx2” = “#E07A6A”, “ANCOM-BC2” = “#4CAF50”, “PARALLAX” = “#5B8BD9” )) + guides(color = guide_legend(title = “Method”)) + labs(x = “Number of Differentially Abundant Taxa”, y = “Performance”, caption = “Lines/circles = precision; triangles = recall”) + theme_classic() + theme(panel.grid.minor = element_blank(), panel.border = element_rect(color = “black”, fill = NA, linewidth = 0.8)) }

disease_plot <- covariate_plot(all_res_plot, “disease”) time_plot <- covariate_plot(all_res_plot, “time”) sex_plot <- covariate_plot(all_res_plot, “sex”)

interaction_plot <- ggplot(all_res_plot %>% filter(var == “interaction”, source == “PARALLAX”), aes(x = DA_number)) + geom_line(aes(y = mean_precision, color = source), linewidth = 1.2) + geom_point(aes(y = mean_precision, color = source), size = 2) + geom_point(aes(y = mean_recall, color = “black”), shape = 17, size = 3) + geom_errorbar( aes( ymin = mean_precision - ci_precision, ymax = mean_precision + ci_precision ), color = “black”, width = 0.5 ) + facet_grid(Effect_size ~ N) + scale_x_continuous(breaks = c(10, 20)) + scale_color_manual(values = c( “ALDEx2” = “#E07A6A”, “ANCOM-BC2” = “#4CAF50”, “PARALLAX” = “#5B8BD9” )) + labs(x = “Number of Differentially Abundant Taxa”, y = “Performance”, caption = “Lines/circles = precision; triangles = recall”) + theme_classic() + theme(panel.grid.minor = element_blank(), panel.border = element_rect(color = “black”, fill = NA, linewidth = 0.8), legend.position = “null”)

ggsave(“disease.jpg”, disease_plot, height = 4, width = 11) ggsave(“time.jpg”, time_plot, height = 4, width = 11) ggsave(“sex.jpg”, sex_plot, height = 4, width = 11) ggsave(“interaction.jpg”, interaction_plot, height = 4, width = 11)

message(“Done. Results in results.csv; figures written as .jpg files.”)
