## Supplementary material for "Perturbation-based propagation of bacterial composition uncertainty improves robustness and reproducibility of microbiome differential abundance analysis": AI-parsable walkthrough

Supplementary File S1

PARALLAX AI-Executable Workflow Specification

This document is a fully executable specification of the PARALLAX microbiome analysis pipeline.
It is intentionally written to support both human users and AI systems.

#### AI EXECUTION NOTES

- All code is provided in sequential order and should be executed as written
- Inputs, transformations, and outputs are explicitly defined inline
- User-modifiable parameters are marked as "ACTION ITEM"
- The workflow assumes compositional count data with repeated measures

#### REQUIRED INPUTS

1. Taxonomic count data (CSV)

The count input file must contain the following columns: - Sample.Name - Tax.Name - Reads.with.Children - Rank

2. Metadata file (CSV)

Required metadata columns (case-sensitive): - Name: Sample identifier (must match count data Sample.Name) - Mouse_ID: repeated measures identifier - Pre.or.Post.Tx: timepoint (e.g., "01. Pre-Tx", "02. Post-Tx") - Sex: biological sex - Injury: disease condition - Treatment: treatment group (optional unless used in model)

All columns MUST exist and be spelled exactly as above.

No spaces vs dots matter (R auto-renaming behavior)

The metadata must contain a column named exactly "Mouse_ID". The design_matrix() function will create "MouseID" internally.

ASSUMPTION: Each subject must have exactly 2 samples (e.g., Pre and Post). Samples must be balanced across timepoints.

If not, this workflow will fail or produce incorrect pairing.

Data must be balanced across all factor combinations so that sorting by (Time, Sex, Injury, Treatment) yields equal-sized Pre and Post partitions.

No missing values are allowed in metadata used for modeling.

#### OUTPUTS

- PARALLAX_res_unfiltered.csv
- PARALLAX_res_filtered.csv
- metadata.csv

#### WORKFLOW SUMMARY

1. Load data
2. Generate taxon-specific matrices
3. Perform CLR transformation + Monte Carlo sampling
4. Construct design matrix
5. Fit mixed models
6. Aggregate posterior statistics
7. Filter significant taxa

The following sections contain the full executable implementation.

#### Execution Instructions

Run all code sequentially in R.

Each section produces intermediate files required for subsequent steps.

All outputs are automatically saved as .CSV files.

This analysis is an example for a Sex x Disease x Time design. It is modifiable by the user at highlighted sections marked "ACTION ITEM"

##### Load required packages

```
library(tidyverse); library(vegan); library(lme4); library(lmerTest);
library(emmeans); library(matrixStats); library(limma); library(ALDEx2)
```

##### Set working directory

**ACTION ITEM (USER-DEFINED PARAMETER – REQUIRED):**

You need 2 folders. One for the scripts (parent), and one for the generated output (child).

*For your script, this will be set up as* `setwd("/insert/file/directory")`

*Use the* `getwd()` *function to determine your current directory for help with formatting*

For example, the parent file containing the scripts is "PARALLAX" and the child file for the output is "PARALLAX Saved Files".

#### Separating data by taxon

**ACTION ITEM:** If you would like, you can customize the saved CSV names in each of the 3 following functions.

##### Generating the species data table

Used in alpha diversity and the species data DA analysis.

```
generate_species_data <- function(data, metadata){
  # Filter to species rank and select necessary columns
  species_data <- data %>%
  dplyr::mutate(
    Rank = tolower(Rank)
  ) %>%
  dplyr::filter(Rank == "species") %>%
  dplyr::select(Sample_Name = Sample.Name,
              Tax_Name = Tax.Name,
              Reads_with_Children = Reads.with.Children) %>%
    dplyr::mutate(Sample_Name = tolower(Sample_Name))

  wide_data <- species_data %>%
  tidyr::pivot_wider(
    names_from = Tax_Name,                # Taxa names become columns
    values_from = Reads_with_Children,    # Values = read counts
    values_fill = 0                       # Fill missing with 0
  )

# Add metadata

# Convert sample names in metadata to lowercase so they can match
  metadata$Name <- tolower(metadata$Name)

# Should read true before reordering.
  checkpoint <- setequal(wide_data$Sample_Name, metadata$Name)

  if(checkpoint == TRUE) {
    # Reorder metadata to match species matrix based on Sample Name
    metadata_ordered <- metadata[match(metadata$Name,wide_data$Sample_Name), ]
    normal_species <- wide_data %>% dplyr::select(-Sample_Name)
    full_results_md <- cbind(metadata_ordered, normal_species)
```

```r
  # Change sample column name
  colnames(full_results_md)[colnames(full_results_md) == "Name"] <- "Sample"
  full_results_md$Sample <- toupper(full_results_md$Sample)

  # Export combined data (can do it, but it is not necessary)
  # write.csv(full_results_md, "R149CombinedRawSpeciesDataAndmeta.csv")
  } else { stop("Sample names do not match")
  }

  model_data <- full_results_md |>
    dplyr::mutate(
      Time = Pre.or.Post.Tx,
      Mouse_ID = Mouse.ID
    ) |>
    relocate(Time, Mouse_ID, .before = Injury)

  rows <- model_data |>
    nrow()

  Subject <- c(1:(rows/2), 1:(rows/2))

# Rearrange data by cohorts
model_ordered <- model_data |>
  arrange(Time, Sex, Injury, Treatment)

# Split data into pre and post time point tables
pretx_df <- model_ordered |>
  slice(1:(rows/2))
posttx_df <- model_ordered |>
  slice(((rows/2)+1):n())

# Reorder posttx_df to match the Mouse ID order in  pre-tx_df
posttx_ordered <- posttx_df[match(pretx_df$Mouse_ID, posttx_df$Mouse_ID), ]

# Combine the ordered posttx_df rows with the pretx_df samples
model_ordered[((rows/2)+1):rows, ] <- posttx_ordered

# Should read TRUE if the pre- and post-tx samples are in the same order
checkpoint2 <- all(model_ordered$Mouse_ID[1:(rows/2)] == model_ordered$Mouse_ID[((rows/2)+1)

if (checkpoint2 == TRUE) {
# Place identical Mouse ID samples together
```

```
model_ordered$Subject <- Subject
model_interleave <- model_ordered[order(model_ordered$Subject, model_ordered$Time), ]

# Format table for input into ALDEx2
sideways_data <- model_interleave |>
  dplyr::select(Sample, where(is.numeric))
aldex2_species_data <- sideways_data |>
  t() |>
  as_tibble(rownames = "Species")

colnames(aldex2_species_data)[1:ncol(aldex2_species_data)]<- aldex2_species_data[1,]
aldex2_species_data<-aldex2_species_data[-1,]

# Export raw counts table to csv for ALDEx2 analysis
write.csv(aldex2_species_data,"readcountsWithchildren_species_data_ordered.csv")

# Make and export the metadata file for limma analysis
metadata_limma <- model_interleave |>
  dplyr::select(Sample, Mouse_ID, Sex, Injury, Time)

write.csv(metadata_limma,"metadata.csv")
} else {stop("PRE and POST-TX samples are not in the same order")}

list(species_data = aldex2_species_data, metadata = metadata_limma)
}
```

##### Generating the genus data table

Used in the genus data DA analysis.

```
generate_genus_data <- function(data, metadata){
  # Filter to genus rank and select necessary columns
  genus_data <- data %>%
  dplyr::mutate(
    Rank = tolower(Rank)
  ) %>%
  dplyr::filter(Rank == "genus") %>%
  dplyr::select(Sample_Name = Sample.Name,
                Tax_Name = Tax.Name,
                Reads_with_Children = Reads.with.Children) %>%
    dplyr::mutate(Sample_Name = tolower(Sample_Name))
```

```
  wide_data <- genus_data %>%
  tidyr::pivot_wider(
    names_from = Tax_Name,                # Taxa names become columns
    values_from = Reads_with_Children,    # Values = read counts
    values_fill = 0                       # Fill missing with 0
  )

# Add metadata

# Convert sample names in metadata to lowercase so they can match
  metadata$Name <- tolower(metadata$Name)

# Should read true before reordering.
  checkpoint <- setequal(wide_data$Sample_Name, metadata$Name)

  if(checkpoint == TRUE) {
    # Reorder metadata to match genus matrix based on Sample Name
    metadata_ordered <- metadata[match(metadata$Name,wide_data$Sample_Name), ]
    normal_genus <- wide_data %>% dplyr::select(-Sample_Name)
    full_results_md <- cbind(metadata_ordered, normal_genus)

  # Change sample column name
  colnames(full_results_md)[colnames(full_results_md) == "Name"] <- "Sample"
  full_results_md$Sample <- toupper(full_results_md$Sample)


  } else { stop("Sample names do not match")
  }

  model_data <- full_results_md |>
    dplyr::mutate(
      Time = Pre.or.Post.Tx,
      Mouse_ID = Mouse.ID
    ) |>
    relocate(Time, Mouse_ID, .before = Injury)

  rows <- model_data |>
    nrow()

  Subject <- c(1:(rows/2), 1:(rows/2))

# Rearrange data by cohorts
```

```
model_ordered <- model_data |>
  arrange(Time, Sex, Injury, Treatment)

# Split data into pre and post time point tables
pretx_df <- model_ordered |>
  slice(1:(rows/2))
posttx_df <- model_ordered |>
  slice(((rows/2)+1):n())

# Reorder posttx_df to match the Mouse ID order in  pre-tx_df
posttx_ordered <- posttx_df[match(pretx_df$Mouse_ID, posttx_df$Mouse_ID), ]

# Combine the ordered posttx_df rows with the pretx_df samples
model_ordered[((rows/2)+1):rows, ] <- posttx_ordered

# Should read TRUE if the pre- and post-tx samples are in the same order
checkpoint2 <- all(model_ordered$Mouse_ID[1:(rows/2)] == model_ordered$Mouse_ID[((rows/2)+1)

if (checkpoint2 == TRUE) {
# Place identical Mouse ID samples together
model_ordered$Subject <- Subject
model_interleave <- model_ordered[order(model_ordered$Subject, model_ordered$Time), ]

# Format table for input into ALDEx2
sideways_data <- model_interleave |>
  dplyr::select(Sample, where(is.numeric))
aldex2_genus_data <- sideways_data |>
  t() |>
  as_tibble(rownames = "Genus")

colnames(aldex2_genus_data)[1:ncol(aldex2_genus_data)]<- aldex2_genus_data[1,]
aldex2_genus_data<-aldex2_genus_data[-1,]

# Export raw counts table to csv for ALDEx2 analysis
write.csv(aldex2_genus_data,"readcountsWithchildren_genus_data_ordered.csv")

# Make and export the metadata file for limma analysis
metadata_limma <- model_interleave |>
  dplyr::select(Sample, Mouse_ID, Sex, Injury, Treatment, Time)

write.csv(metadata_limma,"metadata.csv")
```

```
} else {stop("PRE and POST-TX samples are not in the same order")}

list(genus_data = aldex2_genus_data, metadata = metadata_limma)
}
```

##### Generating the phylum data table

Used in the phylum data DA analysis.

```
generate_phylum_data <- function(data, metadata){
  # Filter to phylum rank and select necessary columns
  phylum_data <- data %>%
  dplyr::mutate(
    Rank = tolower(Rank)
  ) %>%
  dplyr::filter(Rank == "phylum") %>%
  dplyr::select(Sample_Name = Sample.Name,
                Tax_Name = Tax.Name,
                Reads_with_Children = Reads.with.Children) %>%
    dplyr::mutate(Sample_Name = tolower(Sample_Name))

  wide_data <- phylum_data %>%
  tidyr::pivot_wider(
    names_from = Tax_Name,                # Taxa names become columns
    values_from = Reads_with_Children,    # Values = read counts
    values_fill = 0                       # Fill missing with 0
  )

# Add metadata

# Convert sample names in metadata to lowercase so they can match
  metadata$Name <- tolower(metadata$Name)

# Should read true before reordering.
  checkpoint <- setequal(wide_data$Sample_Name, metadata$Name)

  if(checkpoint == TRUE) {
    # Reorder metadata to match phylum matrix based on Sample Name
    metadata_ordered <- metadata[match(metadata$Name,wide_data$Sample_Name), ]
    normal_phylum <- wide_data %>% dplyr::select(-Sample_Name)
    full_results_md <- cbind(metadata_ordered, normal_phylum)
```

```
  # Change sample column name
  colnames(full_results_md)[colnames(full_results_md) == "Name"] <- "Sample"
  full_results_md$Sample <- toupper(full_results_md$Sample)


  } else { stop("Sample names do not match")
  }

   model_data <- full_results_md |>
    dplyr::mutate(
      Time = Pre.or.Post.Tx,
      Mouse_ID = Mouse.ID
    ) |>
    relocate(Time, Mouse_ID, .before = Injury)

  rows <- model_data |>
    nrow()

  Subject <- c(1:(rows/2), 1:(rows/2))

# Rearrange data by cohorts
model_ordered <- model_data |>
  arrange(Time, Sex, Injury, Treatment)

# Split data into pre and post time point tables
pretx_df <- model_ordered |>
  slice(1:(rows/2))
posttx_df <- model_ordered |>
  slice(((rows/2)+1):n())

# Reorder posttx_df to match the Mouse ID order in  pre-tx_df
posttx_ordered <- posttx_df[match(pretx_df$Mouse_ID, posttx_df$Mouse_ID), ]

# Combine the ordered posttx_df rows with the pretx_df samples
model_ordered[((rows/2)+1):rows, ] <- posttx_ordered

# Should read TRUE if the pre- and post-tx samples are in the same order
checkpoint2 <- all(model_ordered$Mouse_ID[1:(rows/2)] == model_ordered$Mouse_ID[((rows/2)+1)

if (checkpoint2 == TRUE) {
# Place identical Mouse ID samples together
model_ordered$Subject <- Subject
```

```
model_interleave <- model_ordered[order(model_ordered$Subject, model_ordered$Time), ]

# Format table for input into ALDEx2
sideways_data <- model_interleave |>
  dplyr::select(Sample, where(is.numeric))
aldex2_phylum_data <- sideways_data |>
  t() |>
  as_tibble(rownames = "Genus")

colnames(aldex2_phylum_data)[1:ncol(aldex2_phylum_data)]<- aldex2_phylum_data[1,]
aldex2_phylum_data<-aldex2_phylum_data[-1,]

# Export raw counts table to csv for ALDEx2 analysis
write.csv(aldex2_phylum_data,"readcountsWithchildren_phylum_data_ordered.csv")

# Make and export the metadata file for limma analysis
metadata_limma <- model_interleave |>
  dplyr::select(Sample, Mouse_ID, Sex, Injury, Treatment, Time)

write.csv(metadata_limma,"metadata.csv")

} else {stop("PRE and POST-TX samples are not in the same order")}

list(phylum_data = aldex2_phylum_data, metadata = metadata_limma)
}
```

### Functions for the PARALLAX workflow

#### CLR transformation and MC replicate generation with ALDEx2

##### About:

*aldex.clr takes raw count data and generates Monte Carlo (MC) instances and applies a centered-log ratio (CLR) transformation. ALDEx2 models uncertainty by treating data as a probability distribution since it is noisy and compositional. Simulating and analyzing multiple probable versions of the data with MC replicates allows for proper compositionality considerations.*

**Data needs to enter with rows = genera; columns = SampleID; cells = counts**

- conds assigns all samples to the same condition
- 128 monte carlo replicates generated per sample
- CLR denominator is the geometric mean of all features (`denom=all`)

*There should be no need to customize this function*

```
# Output is a list: clr_mats (matrix of MC replicates) and M (# cols in 1st MC replicate)

clr_MC <- function(counts) {
    x <- aldex.clr(counts, conds = rep("A", ncol(counts)),
                   mc.samples = 128, denom = "all")
  # Extracts MC CLR transformed data from 'x'. Each element is an MC replicate
  mc_list <- getMonteCarloInstances(x)
  # Counts the number of samples.
  n_samples <- ncol(counts)
  # Counts the number of features (taxa in this case)
  n_features <- nrow(counts)

  # We want one matrix per MC replicate.
  # MC matrices will be features(r) × samples(c).
  # This way each matrix element in the list is a full simulated dataset and
  # statistics can be run per each of the 128 datasets.

  # Build MC replicate matrices safely WITH alignment

  names(mc_list) <- colnames(counts)
  # Counts the number of columns in the first MC replicate CLR matrix.
  # Confirming dimensions.
  M <- ncol(mc_list[[1]])
```

```r
  # clr_mats is the output of the entire function
  clr_mats <- lapply(seq_len(M), function(mc) {

    mat <- sapply(mc_list, function(sample_mat) {

      vec <- sample_mat[, mc]

      # enforce correct ordering + completeness
      vec_aligned <- vec[rownames(counts)]

      return(vec_aligned)
    })

    # ensure matrix structure
    mat <- as.matrix(mat)

    rownames(mat) <- rownames(counts)
    colnames(mat) <- colnames(counts)

    mat
  })
  list(clr_mats = clr_mats, M = M)
}
```

##### Creating the model design matrix

###### Creates reproducible design matrix for each MC replicate

**ACTION ITEMS (USER-DEFINED PARAMETERS – REQUIRED):**

**ACTION ITEM 1:** Change the factor levels for sex, disease, and time according to the data you are working with: `levels = c(x,y)`. The levels need to match the text input in your metadata EXACTLY.

Look for this block to find the levels to change:

```r
meta <- meta %>%
    mutate(
      Sex = factor(Sex, levels = c("Male", "Female")),
      Injury = factor(Injury, levels = c("SHAM", "GWI")),
      Time = factor(Time, levels = c("01. Pre-Tx", "02. Post-Tx")),
      Treatment = factor(Treatment, levels = c("VEHICLE", "ST266")),
      MouseID = factor(ID)
```

```
)
```

- Check your metadata manually with `View(meta)` in the Console before proceeding to ensure match between level names in this function and in the metadata

**ACTION ITEM 2:** Change the model frame `mf` and model matrix `design` to match your experimental parameters. Currently in this example it is set up for 3 main effects and interactions `(Sex + Disease + Time)^2`.

These 2 lines are what you are changing:

```
mf <- model.frame(~ (Sex + Injury + Time)^2, data = meta, na.action = na.fail)

design <- model.matrix(~ (Sex + Injury + Time)^2, data = mf)
```

- Main effects are indicated by `+` and manually iterated interactions by `*`
- Main effects must match the factor names you create in the beginning of the function EXACTLY (ex. Sex, Disease, Time, MouseID)
- Full 2-way/3-way/etc. models are indicated by surrounding the main effect iteration by `()^n` where `n` is the interaction level

```
# Output is design matrix and altered metadata

design_matrix <- function(counts, meta) {
   #==========================
  # Converts metadata columns into defined factor levels
  # CHANGE THE FACTOR LEVELS ACCORDING TO DATASET IMPORTED!!!
  meta <- meta %>%
    dplyr::mutate(
      Sex = factor(Sex, levels = c("Male", "Female")),
      Injury = factor(Injury, levels = c("SHAM", "GWI")),
      Time = factor(Time, levels = c("01. Pre-Tx", "02. Post-Tx")),
      Treatment = factor(Treatment, levels = c("VEHICLE", "ST266")),
      Cohort = factor(paste(Sex, Injury, Time, sep="_")),
      MouseID = factor(Mouse_ID)
    )
  rownames(meta) <- meta$Sample

  meta <- meta |>
    dplyr::select(-Sample)

  # check alignment BEFORE forcing
  if (!all(colnames(counts) %in% rownames(meta))) {
```

```
    stop("Metadata rows do not match count columns")
  }

  # reorder safely
  meta <- meta[match(colnames(counts), rownames(meta)), , drop = FALSE]
  rownames(meta) <- colnames(counts)

  if (any(is.na(meta))) {
    stop("NA values introduced in metadata alignment")
  }

  # Build model frame explicitly.
  # na.action = na.pass prevents silent row dropping, keeps rows with
  # potentially missing values

  # CHANGE THIS ACCORDING TO YOUR EXPERIMENT
  mf <- model.frame(~ (Sex + Injury + Time)^2, data = meta, na.action = na.fail)

  # Build design matrix
  # Each row is a sample, columns are the main effects or interactions

  # CHANGE THIS ACCORDING TO YOUR EXPERIMENT
  design <- model.matrix(~ (Sex + Injury + Time)^2, data = mf)

  # Now assign names safely
  rownames(design) <- colnames(counts)  # must match # rows or model will break

  # make.names is used to eliminate prohibited characters
  colnames(design) <- make.names(colnames(design))

  list(design = design, meta = meta)
}
```

##### Modeling and summary statistics

Linear mixed model defined in the previous function is applied to each MC replicate.

**ACTION ITEM (USER-DEFINED PARAMETER – REQUIRED):**

**ACTION ITEM:** Determine whether to keep or change default delta value.

*delta has been predefined in the inputs list as log(1.2) for this function. It is CUSTOMIZABLE. Change this with the knowledge that it will change the conservative nature of identifying DA taxa and may generate more false positives/false negatives depending on the data.*

```
# Output: result table with summary statistics

model_stats <- function(counts, clr_mats, M, meta, design,
                  sigma_grid, delta = log(1.2),
                  block){
  # Fitting the limma linear mixed model per MC replicate

  # Estimate correlation ONCE
  expr_ref <- clr_mats[[1]]

   if (is.null(block) == FALSE) {
    # duplicateCorrelation accounts for repeated measures
    corfit <- duplicateCorrelation(expr_ref, design, block = block)
    consensus_cor <- corfit$consensus
  } else {
    consensus_cor = NULL
    message("No repeated measures")
  }

  fit_list_all <- vector("list", length(sigma_grid) * M)
  idx <- 1

  for (sigma in sigma_grid) {

    message("Running sigma = ", sigma)

    for (mc in seq_len(M)) {

      expr <- clr_mats[[mc]]

      # Sanity check:
      # Need same number of samples in design matrix as expr data
      stopifnot(ncol(expr) == nrow(design))
      # Makes sure sample order is identical
      stopifnot(all(colnames(expr) == rownames(design)))

      # Apply scale perturbation
      log_s <- rnorm(ncol(expr), mean = 0, sd = sigma)
      expr <- sweep(expr, 2, log_s, FUN = "+")
```

```
    # limma
    fit <- lmFit(expr, design,
                 block = block,
                 correlation = consensus_cor)
    fit <- eBayes(fit)

    fit_list_all[[idx]] <- fit
    idx <- idx + 1
  }
}

# Aggregate across MC replicates
# Combines results from MC replicates into a single summary

# Gets the coefficient names that reside in column names.
# These are for each factor/interaction
coef_names <- colnames(fit_list_all[[1]]$coefficients)
# Counts number of coefficient names
n_coef <- length(coef_names)
n_features <- nrow(counts)

# parameters
# estimate noise scale

result_table <- data.frame(
  Coefficient = rep(coef_names, each = n_features),
  Taxon = rep(rownames(counts), times = n_coef),
  logFC_median = NA,
  P_post = NA,
  q_post = NA,
  R_scale = NA
)

# needed for R_scale
n_sigma <- length(sigma_grid)
rep_per_sigma <- M
sigma_labels <- rep(seq_len(n_sigma), each = rep_per_sigma)

row_idx <- 1

for (coef_idx in seq_len(n_coef)) {
```

```r
  theta_array <- do.call(cbind, lapply(fit_list_all, function(fit) {
    vec <- fit$coefficients[, coef_idx]


    # preserve structure
    vec[is.infinite(vec)] <- NA

    vec
  }))
  vals <- as.vector(theta_array)


  # vectorized statistics
  # What percent of instances have the effect exceeding the delta threshold
  # in either direction
  P_pos <- rowMeans(theta_array > delta, na.rm = TRUE)
  P_neg <- rowMeans(theta_array < -delta, na.rm = TRUE)

  P_post <- pmax(P_pos, P_neg)
  q_post <- 1 - P_post

  logFC_median <- apply(theta_array, 1, median, na.rm = TRUE)

  # scale robustness (R_scale)
  R_scale_vec <- apply(theta_array, 1, function(theta_vec) {
    theta_by_sigma <- split(theta_vec, sigma_labels)

    sig_sigma <- sapply(theta_by_sigma, function(vec) {
      max(mean(vec > delta), mean(vec < -delta))
    })

    mean(sig_sigma)
  })

  # assign block correctly
  result_table$logFC_median[row_idx:(row_idx+n_features-1)] <- logFC_median
  result_table$P_post[row_idx:(row_idx+n_features-1)] <- P_post
  result_table$q_post[row_idx:(row_idx+n_features-1)] <- q_post
  result_table$R_scale[row_idx:(row_idx+n_features-1)] <- R_scale_vec

  row_idx <- row_idx + n_features
}
```

```
  result_table

}
```

##### PARALLAX running function

There is no need to edit this function.

```
# Output is summary statistics table

# EDIT functions according to instructions before running this one
# load functions before running this one

parallax.main <- function(counts, meta, sigma_grid, block) {

  # CLR transform and generate MC replicates
  clrMC <- clr_MC(counts)

  # Unpack outputs from list
  clr_mats <- clrMC$clr_mats
  M <- clrMC$M

  # Outline model design (main effects/interactions)
  design <- design_matrix(counts, meta)

  # Unpack list outputs
  meta <- design$meta
  design <- design$design

  results <- model_stats(counts, clr_mats, M, meta,
                         design, sigma_grid,
                         delta = log(1.2), block)
  list(results = results, meta = meta)

}
```

##### Function for splitting data by time

Done in the case you would like to analyze your data without the confound of Time if necessary.

No edits needed. Ensure that your *Time* column is named *Time* or change the following code to match.

```r
split_timepoints <- function(counts, meta) {

  time_vec <- levels(meta$Time)

  time_split <- list()
  split_meta <- list()

  for (i in seq_along(time_vec)) {

    split_meta[[i]] <- meta |>
      dplyr::filter(Time == time_vec[i])

    time_samples <- as.character(split_meta[[i]]$Sample)

    tmp <- counts[, time_samples, drop = FALSE]

    # Remove Subject row if present
    tmp <- tmp[rownames(tmp) != "Subject", , drop = FALSE]

    # Keep taxa present in at least 5 samples
    tmp <- tmp[rowSums(tmp > 0) >= 5, , drop = FALSE]

    time_split[[i]] <- tmp

  }

  names(time_split) <- time_vec
  names(split_meta) <- time_vec

  list(
    split_counts = time_split,
    split_meta = split_meta
  )
}
```

### Using the PARALLAX workflow

#### Load data

This data is obtained by downloading the numerical data and metadata for a set of samples from OneCodex. See separate tutorial/email OneCodex for help with this.

**ACTION ITEM (USER-DEFINED PARAMETER – REQUIRED):**

*Fill in the " " inside the* `read.csv()` *function with the your data and metadata CSV files*

```
# data <- read.csv("data.csv")
# metadata <- read.csv("metadata.csv")
```

#### Generate separated data by taxon

These functions save .CSV files; there is no need to define outputs.

```
generate_genus_data(data, metadata)
generate_species_data(data, metadata)
generate_phylum_data(data, metadata)
```

##### Loading and filtering the desired data for use

**ACTION ITEMS (USER-DEFINED PARAMETERS – REQUIRED):**

**ACTION ITEM 1:** Replace the name of the .CSV file to load a taxon's data.

Here are the default filenames generated. If the user changed those, use the updated names.

**Genus:** "readcountsWithchildren_genus_data_ordered.csv"

**Species:** "readcountsWithchildren_species_data_ordered.csv"

**Phylum:** "readcountsWithchildren_phylum_data_ordered.csv"

**Metadata:** "metadata.csv"

**ACTION ITEM 2:**

- User needs to ensure the column "Sample" in the **COUNTS** contains the taxa names
- User needs to ensure the column "Sample" in the **METADATA** contains SampleIDs
- User needs to **MANUALLY CHECK** that the rownames of COUNTS end up containing taxa names if the sanity check stops the run
  - This can be done with the function `View(counts)` in the Console (letter case matters)
  - counts should have only the SampleIDs as column names

```
# Import raw counts file. Set rowNames to the first column.
# This example uses the genus data

counts <- read.csv("readcountsWithchildren_genus_data_ordered.csv",
                   row.names = 1, check.names = FALSE)

# Import metadata file. Set rowNames to first column.

meta <- read.csv("metadata.csv", row.names = 1, stringsAsFactors = TRUE)

# Manually rename the rows of the counts df to taxa names. counts$Sample
# contains these for this dataset.
# You may have to alter 'Sample' to the appropriate column name in the
# imported dataset.
rownames(counts) <- counts$Sample
counts <- counts |>
  dplyr::select(-Sample)
```

```r
## Sanity check:
# Ensures sampleID order and identifiers are the same in both files.
# In this metadata file meta$Sample contains SampleID.

stopifnot(all(colnames(counts) == meta$Sample))

# Filter low-abundance taxa
# This step removes entire rows (not all zero-inflation goes away)
# This step is optional. Helps with zero-inflation, but if all counts are low,
# you should avoid.

counts <- counts |>
  dplyr::mutate(across(everything(), as.numeric))
counts <- counts[rownames(counts) != "Subject", ]
counts<- counts |> as.matrix()
counts <- counts[rowSums(counts > 0) >= 5, ]
```

##### Splitting the data

Use this capability if the data needs to be split to conduct analysis without the effect of time. Each time point receives its own dataset. Run `str(split_counts)` and `str(split_meta)` in the console to determine how many list elements to unpack for **BOTH** lists and the names of those list elements. The example below is for two time points.

**NOTE:** *In the case that analyses split by time are intended to be run, ensure block = NULL in the later step and that Time does not appear in the model spec in design_matrix()*

```r
split_data <- split_timepoints(counts, meta)

# Unpack results
split_counts <- split_data$split_counts
split_meta <- split_data$split_meta

# Check how many elements must be unpacked for BOTH lists
posttx_counts <- split_counts$`02. Post-Tx`
pretx_counts <- split_counts$`01. Pre-Tx`

pretx_meta <- split_meta$`01. Pre-Tx`
posttx_meta <- split_meta$`02. Post-Tx`
```

##### Define scale sensitivity grid

Here it is defined with $K = 5$ levels over [0,1]. 0 introducing the least scale variation, 1 introducing the most.

Changing $K$ moderately does not have an impact on data distribution ($R_{scale}$) further down the line. Customize as desired for more/less detail. See manuscript for evidence.

**ACTION ITEM (USER-DEFINED PARAMETER – REQUIRED):**

**ACTION ITEM:** User needs to decide what value of K they will use for the sigma grid.

Instructions for changing the sigma grid: All values of sigma are spread evenly on the interval [0,1].

```
# K = 5
sigma_grid <- c(0, 0.25, 0.5, 0.75, 1)
```

##### Adjust block for repeated measures variable

**ACTION ITEM (USER-DEFINED PARAMETER – REQUIRED):**

**ACTION ITEM:** User needs to define `block`

Notes for user:

- Blocking variable should be something like Mouse ID
- Check column names in metadata
- **NOTE:** *If not dealing with repeated measures, set `block = NULL`*

```
block = meta$MouseID # Change 'MouseID' if necessary
```

#### Run PARALLAX

In the case that analyses split by time are run, ensure that `counts` and `meta` reflect the appropriately split inputs generated by `split_data()`.

```
# Run the workflow
parallax_res <- parallax.main(counts, meta, sigma_grid, block)

# Unpack outputs
parallax_res_raw <- parallax_res$results
meta <- parallax_res$meta
```

##### Save and filter/polish the results

**ACTION ITEM (USER-DEFINED PARAMETER – REQUIRED):**

**ACTION ITEM:** Change these removals to the SECOND factor level you define for each variable in the design_matrix() function.

```
Coefficient = str_remove_all(Coefficient, "Female|GWI|02..Post.Tx")
```

```
# Save raw results. Change filename as desired.
write.csv(parallax_res_raw, "PARALLAX_res_unfiltered.csv")

unfiltered_parallax_res <- parallax_res_raw |>
  dplyr::filter(Coefficient != "X.Intercept.") |> #remove intercept rows
  #(not biologically meaningful)
  dplyr::mutate(
    Coefficient = str_remove_all(Coefficient, "Female|GWI|02..Post.Tx")
  )

# filter according to parameters defined in manuscript
# You can change q_post to 0.1 if looking for exploratory results
# Examine the data distribution for R_scale before altering that parameter
filtered_parallax_res <- unfiltered_parallax_res |> # Filter significant results
  filter(q_post <= 0.05 & R_scale >= 0.8 & abs(logFC_median) >= 0.2)

# Export filtered result table to a csv. Change filename as desired.
write.csv(filtered_parallax_res, "PARALLAX_res_filtered.csv", row.names = FALSE)

# Export final form of metadata file
write.csv(meta, "metadata_final.csv", row.names = FALSE)
```

##### Interpretation of Outputs

Each row represents a taxon–coefficient pair.

- logFC_median: Median effect size
- P_post: Posterior probability effect exceeds threshold
- q_post: Statistical significance (1 − P_post)
- R_scale: Robustness to compositional scaling

Default significance thresholds: - q_post <= 0.05
- R_scale >= 0.8
- |logFC_median| >= 0.2

These values can be adjusted depending on analysis goals.
