## Supplementary methods and figures for "Perturbation-based propagation of bacterial composition uncertainty improves robustness and reproducibility of microbiome differential abundance analysis"

### Supplementary Information.

#### Supplementary Scripts.

An AI-parseable workflow of the PARALLAX pipeline is included as Additional File 2. A traditional walk-through of both the basic method and the post-hoc analysis with embedded code for an example experimental design are included in the GitHub repository cited.

#### Supplementary Methods.

*Animal Model of GW toxic exposure*

Male and Female, C57BL/6 J mice (aged 4-5 months, 20–26 g, Jackson Laboratories, Bar Harbor ME) were housed under standard laboratory conditions (23 ± 1 °C, 50 ± 5% humidity, and 12-h light/dark cycle) with free access to food and water throughout the study. All procedures were carried out under Institutional Animal Care and Use Committee approval and in accordance with the National Institute of Health Guide for the Care and Use of Laboratory Animals. Mice were exposed to GW toxicants through co-administration of pyridostigmine bromide (PB; 0.7 mg/kg; Thermo Fisher Scientific, Waltham, MA) and permethrin (PER; 200 mg/kg; 98.3% mixture consisting of 27.2% cis and 71.1% trans isomers; Sigma-Aldrich, St. Louis, MO). These compounds were delivered in a single 50 µL intraperitoneal (i.p.) injection using dimethyl sulfoxide (DMSO; Sigma-Aldrich, St. Louis, MO) as the vehicle. Control animals received DMSO alone. Injections were administered once daily for 10 consecutive days, as previously described(23,32).

Veterans were typically instructed to self-administer PB at a dose of 90 mg/day (approximately 1.2 mg/kg), taken as one 30 mg tablet every 8 hours. In the present study, mice received a dose of 0.7 mg/kg (i.p.). Pharmacokinetic data indicate that the area under the curve (AUC) for orally administered PB is approximately 39 ng/mL·h in rats (5.82 mg/kg) and 231 ng/mL·h in humans (0.85 mg/kg). To account for interspecies differences in metabolism, a correction factor of 12 was applied, resulting in an estimated equivalent intraperitoneal dose of approximately 0.9 mg/mL in mice(33).

*Animal Model of Repetitive Head Injury*

The TBI dataset was generated using a well-characterized murine model of repetitive mild traumatic brain injury (r-mTBI) developed and validated by our laboratory(25–27). Male and female C57BL/6J mice (aged 3–4 months, Jackson Laboratories, Bar Harbor, ME) were housed under standard laboratory conditions with free access to food and water. All procedures were conducted under Institutional Animal Care and Use Committee approval and in accordance with the National Institutes of Health Guide for the Care and Use of Laboratory Animals.

Animals received five repeated closed-head injuries administered at 48-hour intervals using an electromagnetic controlled impact device (Impact One™ Stereotaxic Impactor, Stoelting, Wood Dale, IL) under isoflurane anesthesia (1.5 L/min oxygen, 3% isoflurane). Injuries were delivered at a strike velocity of 5 m/s, depth of 1.0 mm, and dwell time of 200 ms, with the impact centered over the sagittal suture midway between bregma and lambda. These parameters were selected based on pilot experiments establishing the threshold below which skull fracture, hemorrhage, and gross edema do not occur(26), placing the model at the mild end of the preclinical TBI severity spectrum. Sham animals underwent identical anesthesia and handling procedures without receiving an impact, controlling for the effects of repeated anesthetic exposure.

This r-mTBI paradigm produces a reproducible and progressive neuropathological phenotype including persistent axonal injury, tau hyperphosphorylation, neuroinflammation, white matter degeneration, and long-lasting behavioral deficits across cognitive, motor, and anxiety-related domains that evolve over months to years following injury(25–27).

*Transnetyx Microbiome Analysis*

Transnetyx Microbiome kits containing barcoded sample collection tubes were provided by Transnetyx (Cordova, TN, USA). Mouse fecal samples were placed in individual tubes containing DNA stabilization buffer to ensure reproducibility, stability, and traceability, and shipped for DNA extraction, library preparation, and sequencing by Transnetyx (Cordova, TN USA). DNA extraction was optimized and fully automated using a robust process for reproducible extraction of inhibitor-free, high molecular weight genomic DNA that captures the true microbial diversity of stool samples. After DNA extraction and quality control (QC), genomic DNA was converted into sequencing libraries using a method optimized for minimal bias. Unique dual indexed (UDI) adapters were used to ensure that reads and/or organisms are not mis-assigned. After QC, the libraries were sequenced using the shotgun sequencing method (a depth of 2 million 2x150 bp read pairs), which enables species and strain level taxonomic resolution. Sequencing data were uploaded automatically onto One Codex [1] analysis software and analyzed against the One Codex database consisting of >148K whole microbial reference genomes. The classification results were filtered through several statistical post-processing steps designed to eliminate false positive results caused by contamination or sequencing artifacts.

The One Codex Database consists of ~148K complete microbial genomes, including 71K distinct bacterial genomes, 72K viral genomes, and thousands of archaeal and eukaryotic genomes. Human and mouse genomes are included to screen out host reads, and the complete list of organisms is found here [https://app.onecodex.com/references]. The database is assembled from both of public and private sources, with a combination of automated and manual curation steps to remove low quality or mislabeled records.

DNA extraction is performed using the Qiagen DNeasy 96 PowerSoil Pro QIAcube HT extraction kit and protocol. Library preparation is performed using the KAPA HyperPlus library preparation protocol. Sequencing is performed using the Illumina NextSeq 2000 instrument and protocol. Raw data (in the form of FASTQ files) is analyzed using the One Codex database as above.

Comparing a microbial sample against the One Codex database consists of three sequential steps: K-mer based classification. Every individual sequence (NGS read or contig) is compared against the One Codex database by exact alignment using k-mers where k=31 (see Ames et al., 2014 [2] and Wood et al., 2014 [3] for details on k-mer based classification). Artifact filtering. Based on the relative frequency of unique k-mers in the sample, sequencing artifacts are filtered out of the sample. This filtering should run automatically on most WGS data and does not eliminate low abundance or low confidence hits, only probable sequencing or reference genome artifacts. Species-level abundance estimation. The relative abundance of each microbial species is estimated based on the depth and coverage of sequencing across every available reference genome(34–36).

**Supplementary Figures.**

*Methods: PARALLAX Statistical Workflow*

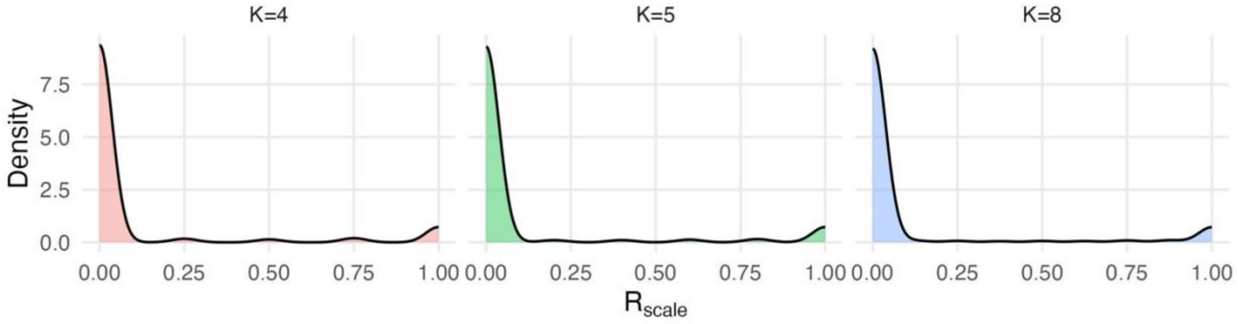


Figure S1. Comparison of $R_{scale}$ distribution across different scale perturbation grids marked by the size of *K*. No significant variation in distribution is observed based on the moderate size variation of the scale perturbation grid.

*Results: Ground Truth*

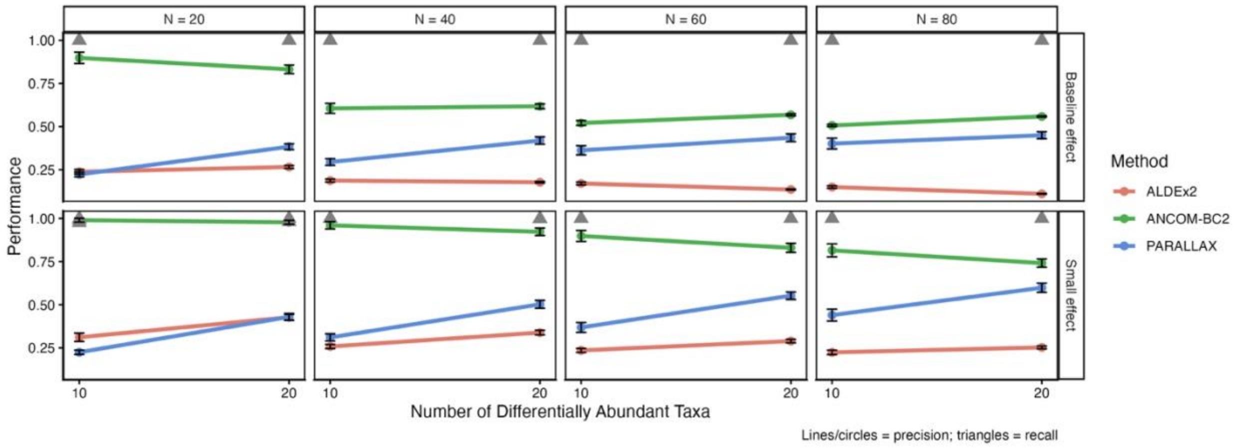


Figure S2. Precision and recall are shown for PARALLAX, ANCOM-BC2, and ALDEx2 across sample sizes (N = 20, 40, 60, 80), numbers of true differentially abundant taxa (DA = 10, 20), and effect size regimes (baseline and reduced) for the time main effect. Error bars show 95% CI across replicates.

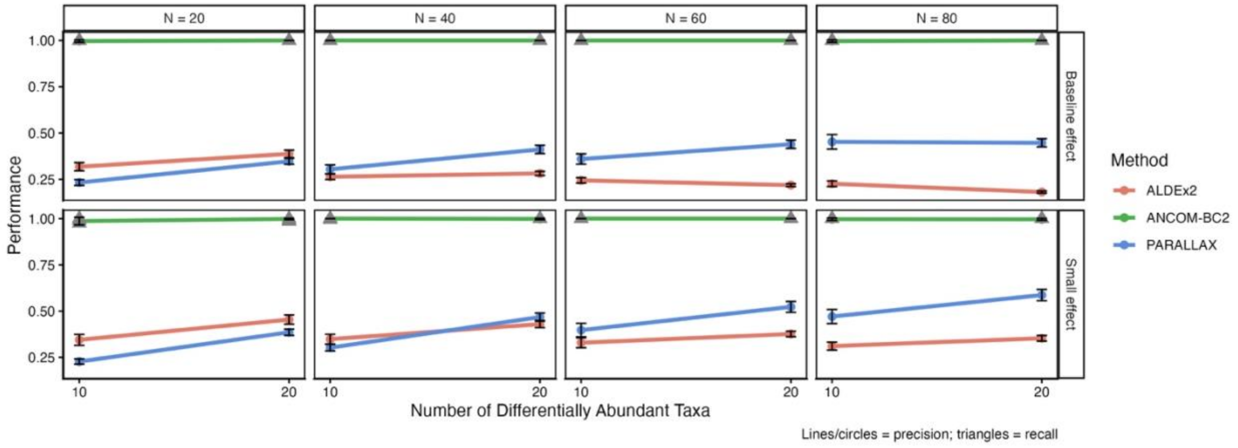

Figure S3. Precision and recall are shown for PARALLAX, ANCOM-BC2, and ALDEx2 across sample sizes (N = 20, 40, 60, 80), numbers of true differentially abundant taxa (DA = 10, 20), and effect size regimes (baseline and reduced) for the sex main effect. Error bars show 95% CI across replicates.

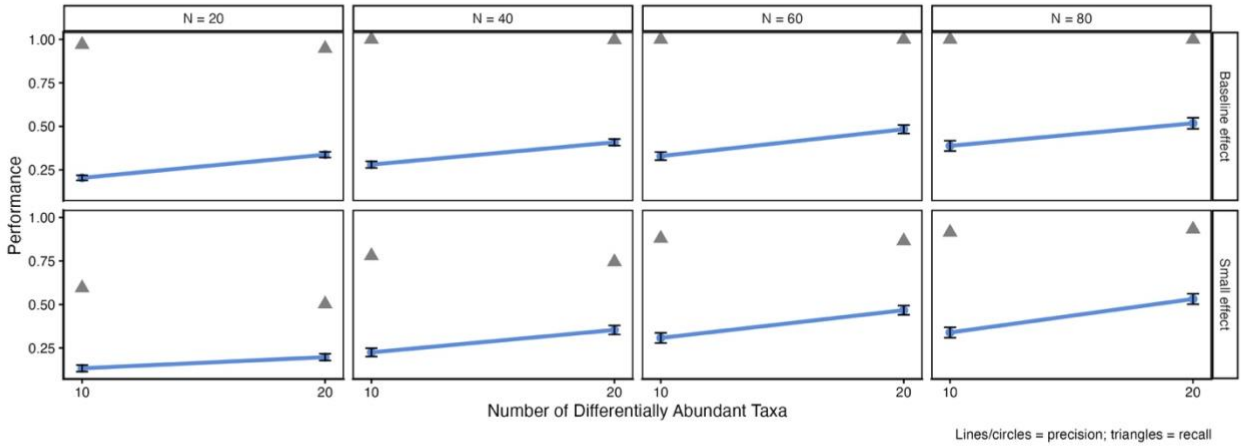


Figure S4. Precision and recall are shown for PARALLAX across sample sizes (N = 20, 40, 60, 80), numbers of true differentially abundant taxa (DA = 10, 20), and effect size regimes (baseline and reduced) for the exposure x time interaction effect. Error bars show 95% CI across replicates.

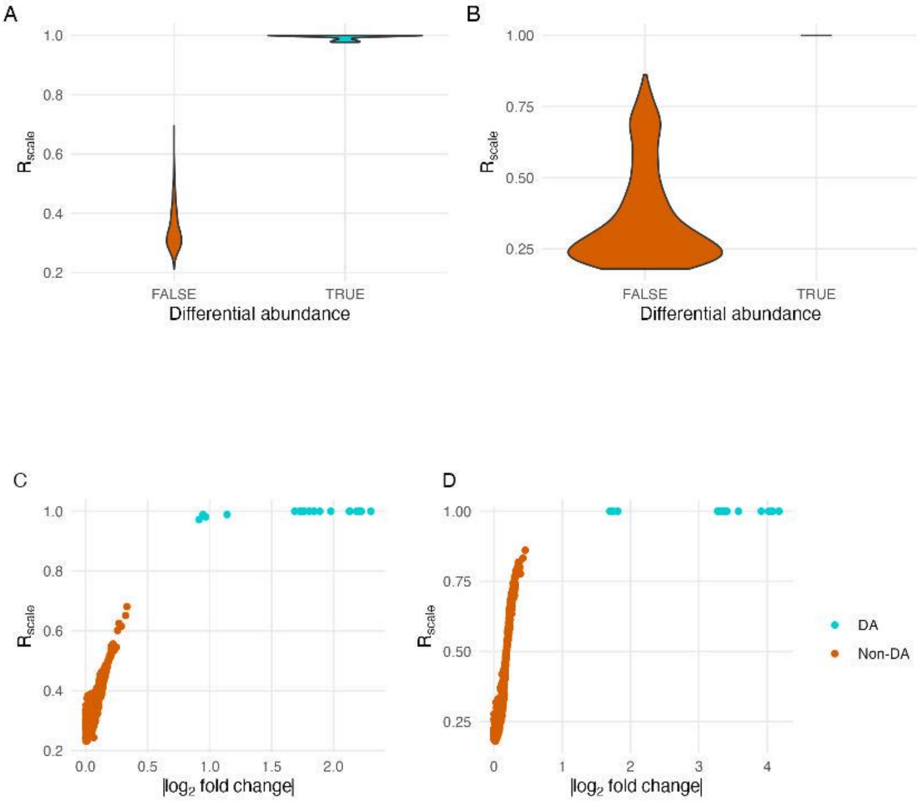

Figure S5. (A) Relationship between $R_{scale}$ and DA status for a reduced-effect simulated dataset. (B) Relationship between $R_{scale}$ and DA status for a baseline simulated dataset. $R_{scale}$ demonstrates clear separation of true DA from non-DA taxa. (C) Relationship between $R_{scale}$ and $log_2$ fold-change for a reduced-effect simulated dataset. (D) Relationship between $R_{scale}$ and $log_2$ fold-change for a baseline simulated dataset. Higher $R_{scale}$ values are associated with true DA taxa and correspond to larger effect sizes.
