## Supplementary material for "Perturbation-based propagation of bacterial composition uncertainty improves robustness and reproducibility of microbiome differential abundance analysis": Simulation results

Ground Truth Simulation Results

| | var | TP | FP | FN | precision | recall | N | DA_numbe | Effect_size | source | replicate |
|---|---|---|---|---|---|---|---|---|---|---|---|
| 1 | disease | 4 | 7 | 0 | 0.363636364 | 1 | # | 10 | Baseline | paralle | 1 |
| 2 | sex | 2 | 5 | 0 | 0.285714286 | 1 | # | 10 | Baseline | paralle | 1 |
| 3 | time | 2 | 11 | 0 | 0.153846154 | 1 | # | 10 | Baseline | paralle | 1 |
| 4 | interaction | 2 | 3 | 0 | 0.4 | 1 | # | 10 | Baseline | paralle | 1 |
| 5 | disease | 4 | 0 | 0 | 1 | 1 | # | 10 | Baseline | ancom | 1 |
| 6 | sex | 2 | 0 | 0 | 1 | 1 | # | 10 | Baseline | ancom | 1 |
| 7 | time | 2 | 0 | 0 | 1 | 1 | # | 10 | Baseline | ancom | 1 |
| 8 | interaction | 0 | 0 | 2 | 0 | 0 | # | 10 | Baseline | ancom | 1 |
| 9 | disease | 4 | 10 | 0 | 0.285714286 | 1 | # | 10 | Baseline | aldex2 | 1 |
| # | sex | 2 | 6 | 0 | 0.25 | 1 | # | 10 | Baseline | aldex2 | 1 |
| # | time | 2 | 8 | 0 | 0.2 | 1 | # | 10 | Baseline | aldex2 | 1 |
| # | interaction | 0 | 0 | 2 | 0 | 0 | # | 10 | Baseline | aldex2 | 1 |
| # | disease | 4 | 0 | 0 | 1 | 1 | # | 10 | Small | paralle | 1 |
| # | sex | 2 | 0 | 0 | 1 | 1 | # | 10 | Small | paralle | 1 |
| # | time | 2 | 0 | 0 | 1 | 1 | # | 10 | Small | paralle | 1 |
| # | interaction | 2 | 0 | 0 | 1 | 1 | # | 10 | Small | paralle | 1 |
| # | disease | 4 | 0 | 0 | 1 | 1 | # | 10 | Small | ancom | 1 |
| # | sex | 2 | 0 | 0 | 1 | 1 | # | 10 | Small | ancom | 1 |
| # | time | 2 | 0 | 0 | 1 | 1 | # | 10 | Small | ancom | 1 |
| # | interaction | 0 | 0 | 2 | 0 | 0 | # | 10 | Small | ancom | 1 |
| # | disease | 4 | 1 | 0 | 0.8 | 1 | # | 10 | Small | aldex2 | 1 |
| # | sex | 2 | 0 | 0 | 1 | 1 | # | 10 | Small | aldex2 | 1 |
| # | time | 2 | 2 | 0 | 0.5 | 1 | # | 10 | Small | aldex2 | 1 |
| # | interaction | 0 | 0 | 2 | 0 | 0 | # | 10 | Small | aldex2 | 1 |
| # | disease | 4 | 3 | 0 | 0.571428571 | 1 | # | 10 | Baseline | paralle | 1 |
| # | sex | 2 | 4 | 0 | 0.333333333 | 1 | # | 10 | Baseline | paralle | 1 |
| # | time | 2 | 8 | 0 | 0.2 | 1 | # | 10 | Baseline | paralle | 1 |
| # | interaction | 2 | 2 | 0 | 0.5 | 1 | # | 10 | Baseline | paralle | 1 |
| # | disease | 4 | 1 | 0 | 0.8 | 1 | # | 10 | Baseline | ancom | 1 |
| # | sex | 2 | 0 | 0 | 1 | 1 | # | 10 | Baseline | ancom | 1 |
| # | time | 2 | 0 | 0 | 1 | 1 | # | 10 | Baseline | ancom | 1 |
| # | interaction | 0 | 0 | 2 | 0 | 0 | # | 10 | Baseline | ancom | 1 |
| # | disease | 4 | 17 | 0 | 0.19047619 | 1 | # | 10 | Baseline | aldex2 | 1 |
| # | sex | 2 | 5 | 0 | 0.285714286 | 1 | # | 10 | Baseline | aldex2 | 1 |
| # | time | 2 | 11 | 0 | 0.153846154 | 1 | # | 10 | Baseline | aldex2 | 1 |
| # | interaction | 0 | 0 | 2 | 0 | 0 | # | 10 | Baseline | aldex2 | 1 |
| # | disease | 4 | 0 | 0 | 1 | 1 | # | 10 | Small | paralle | 1 |
| # | sex | 2 | 0 | 0 | 1 | 1 | # | 10 | Small | paralle | 1 |
| # | time | 2 | 0 | 0 | 1 | 1 | # | 10 | Small | paralle | 1 |
| # | interaction | 2 | 0 | 0 | 1 | 1 | # | 10 | Small | paralle | 1 |
| # | disease | 4 | 1 | 0 | 0.8 | 1 | # | 10 | Small | ancom | 1 |
| # | sex | 2 | 0 | 0 | 1 | 1 | # | 10 | Small | ancom | 1 |
| # | time | 2 | 0 | 0 | 1 | 1 | # | 10 | Small | ancom | 1 |
| # | interaction | 0 | 0 | 2 | 0 | 0 | # | 10 | Small | ancom | 1 |
| # | disease | 4 | 1 | 0 | 0.8 | 1 | # | 10 | Small | aldex2 | 1 |
| # | sex | 2 | 0 | 0 | 1 | 1 | # | 10 | Small | aldex2 | 1 |
| # | time | 2 | 1 | 0 | 0.666666667 | 1 | # | 10 | Small | aldex2 | 1 |
| # | interaction | 0 | 0 | 2 | 0 | 0 | # | 10 | Small | aldex2 | 1 |
| # | disease | 4 | 3 | 0 | 0.571428571 | 1 | # | 10 | Baseline | paralle | 1 |
| # | sex | 2 | 2 | 0 | 0.5 | 1 | # | 10 | Baseline | paralle | 1 |
| # | time | 2 | 5 | 0 | 0.285714286 | 1 | # | 10 | Baseline | paralle | 1 |
| # | interaction | 2 | 2 | 0 | 0.5 | 1 | # | 10 | Baseline | paralle | 1 |
| # | disease | 4 | 1 | 0 | 0.8 | 1 | # | 10 | Baseline | ancom | 1 |
| # | sex | 2 | 0 | 0 | 1 | 1 | # | 10 | Baseline | ancom | 1 |
| # | time | 2 | 2 | 0 | 0.5 | 1 | # | 10 | Baseline | ancom | 1 |
| # | interaction | 0 | 0 | 2 | 0 | 0 | # | 10 | Baseline | ancom | 1 |
| # | disease | 4 | 18 | 0 | 0.181818182 | 1 | # | 10 | Baseline | aldex2 | 1 |
| # | sex | 2 | 6 | 0 | 0.25 | 1 | # | 10 | Baseline | aldex2 | 1 |
| # | time | 2 | 10 | 0 | 0.166666667 | 1 | # | 10 | Baseline | aldex2 | 1 |
| # | interaction | 0 | 0 | 2 | 0 | 0 | # | 10 | Baseline | aldex2 | 1 |
| # | disease | 4 | 0 | 0 | 1 | 1 | # | 10 | Small | paralle | 1 |
| # | sex | 2 | 0 | 0 | 1 | 1 | # | 10 | Small | paralle | 1 |
| # | time | 2 | 0 | 0 | 1 | 1 | # | 10 | Small | paralle | 1 |
| # | interaction | 2 | 0 | 0 | 1 | 1 | # | 10 | Small | paralle | 1 |
| # | disease | 4 | 1 | 0 | 0.8 | 1 | # | 10 | Small | ancom | 1 |
| # | sex | 2 | 0 | 0 | 1 | 1 | # | 10 | Small | ancom | 1 |
| # | time | 2 | 2 | 0 | 0.5 | 1 | # | 10 | Small | ancom | 1 |
| # | interaction | 0 | 0 | 2 | 0 | 0 | # | 10 | Small | ancom | 1 |
| # | disease | 4 | 1 | 0 | 0.8 | 1 | # | 10 | Small | aldex2 | 1 |
| # | sex | 2 | 0 | 0 | 1 | 1 | # | 10 | Small | aldex2 | 1 |
| # | time | 2 | 3 | 0 | 0.4 | 1 | # | 10 | Small | aldex2 | 1 |
| # | interaction | 0 | 0 | 2 | 0 | 0 | # | 10 | Small | aldex2 | 1 |
| # | disease | 4 | 4 | 0 | 0.5 | 1 | # | 10 | Baseline | paralle | 1 |
| # | sex | 2 | 3 | 0 | 0.4 | 1 | # | 10 | Baseline | paralle | 1 |
| # | time | 2 | 7 | 0 | 0.222222222 | 1 | # | 10 | Baseline | paralle | 1 |
| # | interaction | 2 | 4 | 0 | 0.333333333 | 1 | # | 10 | Baseline | paralle | 1 |
| # | disease | 4 | 1 | 0 | 0.8 | 1 | # | 10 | Baseline | ancom | 1 |
| # | sex | 2 | 0 | 0 | 1 | 1 | # | 10 | Baseline | ancom | 1 |
| # | time | 2 | 2 | 0 | 0.5 | 1 | # | 10 | Baseline | ancom | 1 |
| # | interaction | 0 | 0 | 2 | 0 | 0 | # | 10 | Baseline | ancom | 1 |
| # | disease | 4 | 18 | 0 | 0.181818182 | 1 | # | 10 | Baseline | aldex2 | 1 |
| # | sex | 2 | 6 | 0 | 0.25 | 1 | # | 10 | Baseline | aldex2 | 1 |
| # | time | 2 | 17 | 0 | 0.105263158 | 1 | # | 10 | Baseline | aldex2 | 1 |
| # | interaction | 0 | 0 | 2 | 0 | 0 | # | 10 | Baseline | aldex2 | 1 |
| # | disease | 4 | 0 | 0 | 1 | 1 | # | 10 | Small | paralle | 1 |
| # | sex | 2 | 0 | 0 | 1 | 1 | # | 10 | Small | paralle | 1 |
| # | time | 2 | 0 | 0 | 1 | 1 | # | 10 | Small | paralle | 1 |
| # | interaction | 2 | 0 | 0 | 1 | 1 | # | 10 | Small | paralle | 1 |
| # | disease | 4 | 1 | 0 | 0.8 | 1 | # | 10 | Small | ancom | 1 |
| # | sex | 2 | 0 | 0 | 1 | 1 | # | 10 | Small | ancom | 1 |
| # | time | 2 | 2 | 0 | 0.5 | 1 | # | 10 | Small | ancom | 1 |
| # | interaction | 0 | 0 | 2 | 0 | 0 | # | 10 | Small | ancom | 1 |
| # | disease | 4 | 1 | 0 | 0.8 | 1 | # | 10 | Small | aldex2 | 1 |
| # | sex | 2 | 0 | 0 | 1 | 1 | # | 10 | Small | aldex2 | 1 |
| # | time | 2 | 3 | 0 | 0.4 | 1 | # | 10 | Small | aldex2 | 1 |
| # | interaction | 0 | 0 | 2 | 0 | 0 | # | 10 | Small | aldex2 | 1 |
| # | disease | 7 | 6 | 0 | 0.538461538 | 1 | # | 20 | Baseline | paralle | 1 |
| # | sex | 4 | 6 | 0 | 0.4 | 1 | # | 20 | Baseline | paralle | 1 |
| # | time | 5 | 8 | 0 | 0.384615385 | 1 | # | 20 | Baseline | paralle | 1 |
| # | interaction | 4 | 4 | 0 | 0.5 | 1 | # | 20 | Baseline | paralle | 1 |
| # | disease | 7 | 1 | 0 | 0.875 | 1 | # | 20 | Baseline | ancom | 1 |
| # | sex | 4 | 0 | 0 | 1 | 1 | # | 20 | Baseline | ancom | 1 |
| # | time | 5 | 1 | 0 | 0.833333333 | 1 | # | 20 | Baseline | ancom | 1 |
| # | interaction | 0 | 0 | 4 | 0 | 0 | # | 20 | Baseline | ancom | 1 |
| # | disease | 7 | 22 | 0 | 0.24137931 | 1 | # | 20 | Baseline | aldex2 | 1 |
| # | sex | 4 | 6 | 0 | 0.4 | 1 | # | 20 | Baseline | aldex2 | 1 |
| # | time | 5 | 14 | 0 | 0.263157895 | 1 | # | 20 | Baseline | aldex2 | 1 |
| # | interaction | 0 | 0 | 4 | 0 | 0 | # | 20 | Baseline | aldex2 | 1 |
| # | disease | 7 | 0 | 0 | 1 | 1 | # | 20 | Small | paralle | 1 |
| # | sex | 4 | 0 | 0 | 1 | 1 | # | 20 | Small | paralle | 1 |
| # | time | 5 | 0 | 0 | 1 | 1 | # | 20 | Small | paralle | 1 |
| # | interaction | 3 | 0 | 1 | 1 | 0.75 | # | 20 | Small | paralle | 1 |
| # | disease | 7 | 1 | 0 | 0.875 | 1 | # | 20 | Small | ancom | 1 |
| # | sex | 4 | 0 | 0 | 1 | 1 | # | 20 | Small | ancom | 1 |
| # | time | 5 | 1 | 0 | 0.833333333 | 1 | # | 20 | Small | ancom | 1 |
| # | interaction | 0 | 0 | 4 | 0 | 0 | # | 20 | Small | ancom | 1 |
| # | disease | 7 | 2 | 0 | 0.777777778 | 1 | # | 20 | Small | aldex2 | 1 |
| # | sex | 4 | 0 | 0 | 1 | 1 | # | 20 | Small | aldex2 | 1 |
| # | time | 5 | 2 | 0 | 0.714285714 | 1 | # | 20 | Small | aldex2 | 1 |
| # | interaction | 0 | 0 | 4 | 0 | 0 | # | 20 | Small | aldex2 | 1 |
| # | disease | 7 | 10 | 0 | 0.411764706 | 1 | # | 20 | Baseline | paralle | 1 |
| # | sex | 4 | 4 | 0 | 0.5 | 1 | # | 20 | Baseline | paralle | 1 |
| # | time | 5 | 4 | 0 | 0.555555556 | 1 | # | 20 | Baseline | paralle | 1 |
| # | interaction | 4 | 3 | 0 | 0.571428571 | 1 | # | 20 | Baseline | paralle | 1 |
| # | disease | 7 | 4 | 0 | 0.636363636 | 1 | # | 20 | Baseline | ancom | 1 |
| # | sex | 4 | 0 | 0 | 1 | 1 | # | 20 | Baseline | ancom | 1 |
| # | time | 5 | 3 | 0 | 0.625 | 1 | # | 20 | Baseline | ancom | 1 |
| # | interaction | 0 | 0 | 4 | 0 | 0 | # | 20 | Baseline | ancom | 1 |
| # | disease | 7 | 39 | 0 | 0.152173913 | 1 | # | 20 | Baseline | aldex2 | 1 |
| # | sex | 4 | 9 | 0 | 0.307692308 | 1 | # | 20 | Baseline | aldex2 | 1 |
| # | time | 5 | 27 | 0 | 0.15625 | 1 | # | 20 | Baseline | aldex2 | 1 |
| # | interaction | 0 | 0 | 4 | 0 | 0 | # | 20 | Baseline | aldex2 | 1 |
| # | disease | 7 | 0 | 0 | 1 | 1 | # | 20 | Small | paralle | 1 |
| # | sex | 4 | 0 | 0 | 1 | 1 | # | 20 | Small | paralle | 1 |
| # | time | 5 | 0 | 0 | 1 | 1 | # | 20 | Small | paralle | 1 |
| # | interaction | 4 | 0 | 0 | 1 | 1 | # | 20 | Small | paralle | 1 |
| # | disease | 7 | 3 | 0 | 0.7 | 1 | # | 20 | Small | ancom | 1 |
| # | sex | 4 | 0 | 0 | 1 | 1 | # | 20 | Small | ancom | 1 |
| # | time | 5 | 3 | 0 | 0.625 | 1 | # | 20 | Small | ancom | 1 |
| # | interaction | 0 | 0 | 4 | 0 | 0 | # | 20 | Small | ancom | 1 |
| # | disease | 7 | 7 | 0 | 0.5 | 1 | # | 20 | Small | aldex2 | 1 |
| # | sex | 4 | 0 | 0 | 1 | 1 | # | 20 | Small | aldex2 | 1 |
| # | time | 5 | 3 | 0 | 0.625 | 1 | # | 20 | Small | aldex2 | 1 |
| # | interaction | 0 | 0 | 4 | 0 | 0 | # | 20 | Small | aldex2 | 1 |
| # | disease | 7 | 6 | 0 | 0.538461538 | 1 | # | 20 | Baseline | paralle | 1 |
| # | sex | 4 | 4 | 0 | 0.5 | 1 | # | 20 | Baseline | paralle | 1 |
| # | time | 5 | 9 | 0 | 0.357142857 | 1 | # | 20 | Baseline | paralle | 1 |
| # | interaction | 4 | 2 | 0 | 0.666666667 | 1 | # | 20 | Baseline | paralle | 1 |
| # | disease | 7 | 4 | 0 | 0.636363636 | 1 | # | 20 | Baseline | ancom | 1 |
| # | sex | 4 | 0 | 0 | 1 | 1 | # | 20 | Baseline | ancom | 1 |
| # | time | 5 | 5 | 0 | 0.5 | 1 | # | 20 | Baseline | ancom | 1 |
| # | interaction | 0 | 0 | 4 | 0 | 0 | # | 20 | Baseline | ancom | 1 |
| # | disease | 7 | 52 | 0 | 0.118644068 | 1 | # | 20 | Baseline | aldex2 | 1 |
| # | sex | 4 | 15 | 0 | 0.210526316 | 1 | # | 20 | Baseline | aldex2 | 1 |
| # | time | 5 | 36 | 0 | 0.12195122 | 1 | # | 20 | Baseline | aldex2 | 1 |
| # | interaction | 0 | 0 | 4 | 0 | 0 | # | 20 | Baseline | aldex2 | 1 |
| # | disease | 7 | 0 | 0 | 1 | 1 | # | 20 | Small | paralle | 1 |
| # | sex | 4 | 0 | 0 | 1 | 1 | # | 20 | Small | paralle | 1 |
| # | time | 5 | 0 | 0 | 1 | 1 | # | 20 | Small | paralle | 1 |
| # | interaction | 4 | 0 | 0 | 1 | 1 | # | 20 | Small | paralle | 1 |
| # | disease | 7 | 4 | 0 | 0.636363636 | 1 | # | 20 | Small | ancom | 1 |
| # | sex | 4 | 0 | 0 | 1 | 1 | # | 20 | Small | ancom | 1 |
| # | time | 5 | 5 | 0 | 0.5 | 1 | # | 20 | Small | ancom | 1 |
| # | interaction | 0 | 0 | 4 | 0 | 0 | # | 20 | Small | ancom | 1 |
| # | disease | 7 | 8 | 0 | 0.466666667 | 1 | # | 20 | Small | aldex2 | 1 |
| # | sex | 4 | 0 | 0 | 1 | 1 | # | 20 | Small | aldex2 | 1 |
| # | time | 5 | 5 | 0 | 0.5 | 1 | # | 20 | Small | aldex2 | 1 |
| # | interaction | 0 | 0 | 4 | 0 | 0 | # | 20 | Small | aldex2 | 1 |
| # | disease | 7 | 5 | 0 | 0.583333333 | 1 | # | 20 | Baseline | paralle | 1 |
| # | sex | 4 | 5 | 0 | 0.444444444 | 1 | # | 20 | Baseline | paralle | 1 |
| # | time | 5 | 6 | 0 | 0.454545455 | 1 | # | 20 | Baseline | paralle | 1 |
| # | interaction | 4 | 3 | 0 | 0.571428571 | 1 | # | 20 | Baseline | paralle | 1 |
| # | disease | 7 | 4 | 0 | 0.636363636 | 1 | # | 20 | Baseline | ancom | 1 |
| # | sex | 4 | 0 | 0 | 1 | 1 | # | 20 | Baseline | ancom | 1 |
| # | time | 5 | 4 | 0 | 0.555555556 | 1 | # | 20 | Baseline | ancom | 1 |
| # | interaction | 0 | 0 | 4 | 0 | 0 | # | 20 | Baseline | ancom | 1 |
| # | disease | 7 | 58 | 0 | 0.107692308 | 1 | # | 20 | Baseline | aldex2 | 1 |
| # | sex | 4 | 21 | 0 | 0.16 | 1 | # | 20 | Baseline | aldex2 | 1 |
| # | time | 5 | 69 | 0 | 0.092592593 | 1 | # | 20 | Baseline | aldex2 | 1 |
| # | interaction | 0 | 0 | 4 | 0 | 0 | # | 20 | Baseline | aldex2 | 1 |
| # | disease | 7 | 0 | 0 | 1 | 1 | # | 20 | Small | paralle | 1 |
| # | sex | 4 | 0 | 0 | 1 | 1 | # | 20 | Small | paralle | 1 |
| # | time | 5 | 0 | 0 | 1 | 1 | # | 20 | Small | paralle | 1 |
| # | interaction | 4 | 0 | 0 | 1 | 1 | # | 20 | Small | paralle | 1 |
| # | disease | 7 | 4 | 0 | 0.636363636 | 1 | # | 20 | Small | ancom | 1 |
| # | sex | 4 | 0 | 0 | 1 | 1 | # | 20 | Small | ancom | 1 |
| # | time | 5 | 5 | 0 | 0.5 | 1 | # | 20 | Small | ancom | 1 |
| # | interaction | 0 | 0 | 4 | 0 | 0 | # | 20 | Small | ancom | 1 |
| # | disease | 7 | 7 | 0 | 0.5 | 1 | # | 20 | Small | aldex2 | 1 |
| # | sex | 4 | 1 | 0 | 0.8 | 1 | # | 20 | Small | aldex2 | 1 |
| # | time | 5 | 6 | 0 | 0.454545455 | 1 | # | 20 | Small | aldex2 | 1 |
| # | interaction | 0 | 0 | 4 | 0 | 0 | # | 20 | Small | aldex2 | 1 |
| # | disease | 4 | 7 | 0 | 0.363636364 | 1 | # | 10 | Baseline | paralle | 2 |
| # | sex | 2 | 11 | 0 | 0.153846154 | 1 | # | 10 | Baseline | paralle | 2 |
| # | time | 2 | 9 | 0 | 0.181818182 | 1 | # | 10 | Baseline | paralle | 2 |
| # | interaction | 1 | 8 | 1 | 0.111111111 | 0.5 | # | 10 | Baseline | paralle | 2 |
| # | disease | 4 | 0 | 0 | 1 | 1 | # | 10 | Baseline | ancom | 2 |
| # | sex | 2 | 0 | 0 | 1 | 1 | # | 10 | Baseline | ancom | 2 |
| # | time | 2 | 1 | 0 | 0.666666667 | 1 | # | 10 | Baseline | ancom | 2 |
| # | interaction | 0 | 0 | 2 | 0 | 0 | # | 10 | Baseline | ancom | 2 |
| # | disease | 4 | 9 | 0 | 0.307692308 | 1 | # | 10 | Baseline | aldex2 | 2 |
| # | sex | 2 | 3 | 0 | 0.4 | 1 | # | 10 | Baseline | aldex2 | 2 |
| # | time | 2 | 8 | 0 | 0.2 | 1 | # | 10 | Baseline | aldex2 | 2 |
| # | interaction | 0 | 0 | 2 | 0 | 0 | # | 10 | Baseline | aldex2 | 2 |
| # | disease | 4 | 0 | 0 | 1 | 1 | # | 10 | Small | paralle | 2 |
| # | sex | 2 | 0 | 0 | 1 | 1 | # | 10 | Small | paralle | 2 |
| # | time | 2 | 0 | 0 | 1 | 1 | # | 10 | Small | paralle | 2 |
| # | interaction | 1 | 1 | 1 | 0.5 | 0.5 | # | 10 | Small | paralle | 2 |
| # | disease | 4 | 0 | 0 | 1 | 1 | # | 10 | Small | ancom | 2 |
| # | sex | 2 | 0 | 0 | 1 | 1 | # | 10 | Small | ancom | 2 |
| # | time | 2 | 0 | 0 | 1 | 1 | # | 10 | Small | ancom | 2 |
| # | interaction | 0 | 0 | 2 | 0 | 0 | # | 10 | Small | ancom | 2 |
| # | disease | 4 | 0 | 0 | 1 | 1 | # | 10 | Small | aldex2 | 2 |
| # | sex | 2 | 0 | 0 | 1 | 1 | # | 10 | Small | aldex2 | 2 |
| # | time | 2 | 1 | 0 | 0.666666667 | 1 | # | 10 | Small | aldex2 | 2 |
| # | interaction | 0 | 0 | 2 | 0 | 0 | # | 10 | Small | aldex2 | 2 |
| # | disease | 4 | 5 | 0 | 0.444444444 | 1 | # | 10 | Baseline | paralle | 2 |
| # | sex | 2 | 5 | 0 | 0.285714286 | 1 | # | 10 | Baseline | paralle | 2 |
| # | time | 2 | 1 | 0 | 0.666666667 | 1 | # | 10 | Baseline | paralle | 2 |
| # | interaction | 2 | 6 | 0 | 0.25 | 1 | # | 10 | Baseline | paralle | 2 |
| # | disease | 4 | 2 | 0 | 0.666666667 | 1 | # | 10 | Baseline | ancom | 2 |
| # | sex | 2 | 0 | 0 | 1 | 1 | # | 10 | Baseline | ancom | 2 |
| # | time | 2 | 1 | 0 | 0.666666667 | 1 | # | 10 | Baseline | ancom | 2 |
| # | interaction | 0 | 0 | 2 | 0 | 0 | # | 10 | Baseline | ancom | 2 |
| # | disease | 4 | 9 | 0 | 0.307692308 | 1 | # | 10 | Baseline | aldex2 | 2 |
| # | sex | 2 | 5 | 0 | 0.285714286 | 1 | # | 10 | Baseline | aldex2 | 2 |
| # | time | 2 | 9 | 0 | 0.181818182 | 1 | # | 10 | Baseline | aldex2 | 2 |
| # | interaction | 0 | 0 | 2 | 0 | 0 | # | 10 | Baseline | aldex2 | 2 |
| # | disease | 4 | 0 | 0 | 1 | 1 | # | 10 | Small | paralle | 2 |
| # | sex | 2 | 0 | 0 | 1 | 1 | # | 10 | Small | paralle | 2 |
| # | time | 2 | 0 | 0 | 1 | 1 | # | 10 | Small | paralle | 2 |
| # | interaction | 1 | 0 | 1 | 1 | 0.5 | # | 10 | Small | paralle | 2 |
| # | disease | 4 | 1 | 0 | 0.8 | 1 | # | 10 | Small | ancom | 2 |
| # | sex | 2 | 0 | 0 | 1 | 1 | # | 10 | Small | ancom | 2 |
| # | time | 2 | 1 | 0 | 0.666666667 | 1 | # | 10 | Small | ancom | 2 |
| # | interaction | 0 | 0 | 2 | 0 | 0 | # | 10 | Small | ancom | 2 |
| # | disease | 4 | 1 | 0 | 0.8 | 1 | # | 10 | Small | aldex2 | 2 |
| # | sex | 2 | 0 | 0 | 1 | 1 | # | 10 | Small | aldex2 | 2 |
| # | time | 2 | 1 | 0 | 0.666666667 | 1 | # | 10 | Small | aldex2 | 2 |
| # | interaction | 0 | 0 | 2 | 0 | 0 | # | 10 | Small | aldex2 | 2 |
| # | disease | 4 | 7 | 0 | 0.363636364 | 1 | # | 10 | Baseline | paralle | 2 |
| # | sex | 2 | 6 | 0 | 0.25 | 1 | # | 10 | Baseline | paralle | 2 |
| # | time | 2 | 2 | 0 | 0.5 | 1 | # | 10 | Baseline | paralle | 2 |
| # | interaction | 2 | 4 | 0 | 0.333333333 | 1 | # | 10 | Baseline | paralle | 2 |
| # | disease | 4 | 2 | 0 | 0.666666667 | 1 | # | 10 | Baseline | ancom | 2 |
| # | sex | 2 | 0 | 0 | 1 | 1 | # | 10 | Baseline | ancom | 2 |
| # | time | 2 | 2 | 0 | 0.5 | 1 | # | 10 | Baseline | ancom | 2 |
| # | interaction | 0 | 0 | 2 | 0 | 0 | # | 10 | Baseline | ancom | 2 |
| # | disease | 4 | 17 | 0 | 0.19047619 | 1 | # | 10 | Baseline | aldex2 | 2 |
| # | sex | 2 | 4 | 0 | 0.333333333 | 1 | # | 10 | Baseline | aldex2 | 2 |
| # | time | 2 | 9 | 0 | 0.181818182 | 1 | # | 10 | Baseline | aldex2 | 2 |
| # | interaction | 0 | 0 | 2 | 0 | 0 | # | 10 | Baseline | aldex2 | 2 |
| # | disease | 4 | 0 | 0 | 1 | 1 | # | 10 | Small | paralle | 2 |
| # | sex | 2 | 1 | 0 | 0.666666667 | 1 | # | 10 | Small | paralle | 2 |
| # | time | 2 | 0 | 0 | 1 | 1 | # | 10 | Small | paralle | 2 |
| # | interaction | 1 | 0 | 1 | 1 | 0.5 | # | 10 | Small | paralle | 2 |
| # | disease | 4 | 2 | 0 | 0.666666667 | 1 | # | 10 | Small | ancom | 2 |
| # | sex | 2 | 0 | 0 | 1 | 1 | # | 10 | Small | ancom | 2 |
| # | time | 2 | 1 | 0 | 0.666666667 | 1 | # | 10 | Small | ancom | 2 |
| # | interaction | 0 | 0 | 2 | 0 | 0 | # | 10 | Small | ancom | 2 |
| # | disease | 4 | 1 | 0 | 0.8 | 1 | # | 10 | Small | aldex2 | 2 |
| # | sex | 2 | 0 | 0 | 1 | 1 | # | 10 | Small | aldex2 | 2 |
| # | time | 2 | 1 | 0 | 0.666666667 | 1 | # | 10 | Small | aldex2 | 2 |
| # | interaction | 0 | 0 | 2 | 0 | 0 | # | 10 | Small | aldex2 | 2 |
| # | disease | 4 | 5 | 0 | 0.444444444 | 1 | # | 10 | Baseline | paralle | 2 |
| # | sex | 2 | 2 | 0 | 0.5 | 1 | # | 10 | Baseline | paralle | 2 |
| # | time | 2 | 4 | 0 | 0.333333333 | 1 | # | 10 | Baseline | paralle | 2 |
| # | interaction | 2 | 3 | 0 | 0.4 | 1 | # | 10 | Baseline | paralle | 2 |
| # | disease | 4 | 2 | 0 | 0.666666667 | 1 | # | 10 | Baseline | ancom | 2 |
| # | sex | 2 | 0 | 0 | 1 | 1 | # | 10 | Baseline | ancom | 2 |
| # | time | 2 | 2 | 0 | 0.5 | 1 | # | 10 | Baseline | ancom | 2 |
| # | interaction | 0 | 0 | 2 | 0 | 0 | # | 10 | Baseline | ancom | 2 |
| # | disease | 4 | 20 | 0 | 0.166666667 | 1 | # | 10 | Baseline | aldex2 | 2 |
| # | sex | 2 | 5 | 0 | 0.285714286 | 1 | # | 10 | Baseline | aldex2 | 2 |
| # | time | 2 | 7 | 0 | 0.222222222 | 1 | # | 10 | Baseline | aldex2 | 2 |
| # | interaction | 0 | 0 | 2 | 0 | 0 | # | 10 | Baseline | aldex2 | 2 |
| # | disease | 4 | 0 | 0 | 1 | 1 | # | 10 | Small | paralle | 2 |
| # | sex | 2 | 1 | 0 | 0.666666667 | 1 | # | 10 | Small | paralle | 2 |
| # | time | 2 | 0 | 0 | 1 | 1 | # | 10 | Small | paralle | 2 |
| # | interaction | 2 | 0 | 0 | 1 | 1 | # | 10 | Small | paralle | 2 |
| # | disease | 4 | 2 | 0 | 0.666666667 | 1 | # | 10 | Small | ancom | 2 |
| # | sex | 2 | 0 | 0 | 1 | 1 | # | 10 | Small | ancom | 2 |
| # | time | 2 | 2 | 0 | 0.5 | 1 | # | 10 | Small | ancom | 2 |
| # | interaction | 0 | 0 | 2 | 0 | 0 | # | 10 | Small | ancom | 2 |
| # | disease | 4 | 1 | 0 | 0.8 | 1 | # | 10 | Small | aldex2 | 2 |
| # | sex | 2 | 0 | 0 | 1 | 1 | # | 10 | Small | aldex2 | 2 |
| # | time | 2 | 2 | 0 | 0.5 | 1 | # | 10 | Small | aldex2 | 2 |
| # | interaction | 0 | 0 | 2 | 0 | 0 | # | 10 | Small | aldex2 | 2 |
| # | disease | 7 | 10 | 0 | 0.411764706 | 1 | # | 20 | Baseline | paralle | 2 |
| # | sex | 4 | 10 | 0 | 0.285714286 | 1 | # | 20 | Baseline | paralle | 2 |
| # | time | 5 | 5 | 0 | 0.5 | 1 | # | 20 | Baseline | paralle | 2 |
| # | interaction | 4 | 3 | 0 | 0.571428571 | 1 | # | 20 | Baseline | paralle | 2 |
| # | disease | 7 | 1 | 0 | 0.875 | 1 | # | 20 | Baseline | ancom | 2 |
| # | sex | 4 | 0 | 0 | 1 | 1 | # | 20 | Baseline | ancom | 2 |
| # | time | 5 | 1 | 0 | 0.833333333 | 1 | # | 20 | Baseline | ancom | 2 |
| # | interaction | 0 | 0 | 4 | 0 | 0 | # | 20 | Baseline | ancom | 2 |
| # | disease | 7 | 21 | 0 | 0.25 | 1 | # | 20 | Baseline | aldex2 | 2 |
| # | sex | 4 | 3 | 0 | 0.571428571 | 1 | # | 20 | Baseline | aldex2 | 2 |
| # | time | 5 | 11 | 0 | 0.3125 | 1 | # | 20 | Baseline | aldex2 | 2 |
| # | interaction | 0 | 0 | 4 | 0 | 0 | # | 20 | Baseline | aldex2 | 2 |
| # | disease | 7 | 1 | 0 | 0.875 | 1 | # | 20 | Small | paralle | 2 |
| # | sex | 4 | 0 | 0 | 1 | 1 | # | 20 | Small | paralle | 2 |
| # | time | 5 | 0 | 0 | 1 | 1 | # | 20 | Small | paralle | 2 |
| # | interaction | 1 | 0 | 3 | 1 | 0.25 | # | 20 | Small | paralle | 2 |
| # | disease | 7 | 1 | 0 | 0.875 | 1 | # | 20 | Small | ancom | 2 |
| # | sex | 4 | 0 | 0 | 1 | 1 | # | 20 | Small | ancom | 2 |
| # | time | 5 | 1 | 0 | 0.833333333 | 1 | # | 20 | Small | ancom | 2 |
| # | interaction | 0 | 0 | 4 | 0 | 0 | # | 20 | Small | ancom | 2 |
| # | disease | 7 | 1 | 0 | 0.875 | 1 | # | 20 | Small | aldex2 | 2 |
| # | sex | 4 | 0 | 0 | 1 | 1 | # | 20 | Small | aldex2 | 2 |
| # | time | 5 | 1 | 0 | 0.833333333 | 1 | # | 20 | Small | aldex2 | 2 |
| # | interaction | 0 | 0 | 4 | 0 | 0 | # | 20 | Small | aldex2 | 2 |
| # | disease | 7 | 10 | 0 | 0.411764706 | 1 | # | 20 | Baseline | paralle | 2 |
| # | sex | 4 | 8 | 0 | 0.333333333 | 1 | # | 20 | Baseline | paralle | 2 |
| # | time | 5 | 9 | 0 | 0.357142857 | 1 | # | 20 | Baseline | paralle | 2 |
| # | interaction | 4 | 3 | 0 | 0.571428571 | 1 | # | 20 | Baseline | paralle | 2 |
| # | disease | 7 | 3 | 0 | 0.7 | 1 | # | 20 | Baseline | ancom | 2 |
| # | sex | 4 | 0 | 0 | 1 | 1 | # | 20 | Baseline | ancom | 2 |
| # | time | 5 | 5 | 0 | 0.5 | 1 | # | 20 | Baseline | ancom | 2 |
| # | interaction | 0 | 0 | 4 | 0 | 0 | # | 20 | Baseline | ancom | 2 |
| # | disease | 7 | 32 | 0 | 0.179487179 | 1 | # | 20 | Baseline | aldex2 | 2 |
| # | sex | 4 | 9 | 0 | 0.307692308 | 1 | # | 20 | Baseline | aldex2 | 2 |
| # | time | 5 | 25 | 0 | 0.166666667 | 1 | # | 20 | Baseline | aldex2 | 2 |
| # | interaction | 0 | 0 | 4 | 0 | 0 | # | 20 | Baseline | aldex2 | 2 |
| # | disease | 7 | 0 | 0 | 1 | 1 | # | 20 | Small | paralle | 2 |
| # | sex | 4 | 0 | 0 | 1 | 1 | # | 20 | Small | paralle | 2 |
| # | time | 5 | 0 | 0 | 1 | 1 | # | 20 | Small | paralle | 2 |
| # | interaction | 3 | 0 | 1 | 1 | 0.75 | # | 20 | Small | paralle | 2 |
| # | disease | 7 | 3 | 0 | 0.7 | 1 | # | 20 | Small | ancom | 2 |
| # | sex | 4 | 0 | 0 | 1 | 1 | # | 20 | Small | ancom | 2 |
| # | time | 5 | 4 | 0 | 0.555555556 | 1 | # | 20 | Small | ancom | 2 |
| # | interaction | 0 | 0 | 4 | 0 | 0 | # | 20 | Small | ancom | 2 |
| # | disease | 7 | 2 | 0 | 0.777777778 | 1 | # | 20 | Small | aldex2 | 2 |
| # | sex | 4 | 0 | 0 | 1 | 1 | # | 20 | Small | aldex2 | 2 |

| | | | | | | | | | | | |
|---|---|---|---|---|---|---|---|---|---|---|---|
| # | time | 5 | 3 | 0 | 0.625 | 1 | # | 20 | Small | sides2 | 2 |
| # | interaction | 0 | 0 | 4 | 0 | 0 | # | 20 | Small | sides2 | 2 |
| # | disease | 7 | 13 | 0 | 0.35 | 1 | # | 20 | Baseline | paralle | 2 |
| # | sex | 4 | 5 | 0 | 0.444444444 | 1 | # | 20 | Baseline | paralle | 2 |
| # | time | 5 | 6 | 0 | 0.454545455 | 1 | # | 20 | Baseline | paralle | 2 |
| # | interaction | 4 | 5 | 0 | 0.444444444 | 1 | # | 20 | Baseline | paralle | 2 |
| # | disease | 7 | 4 | 0 | 0.636363636 | 1 | # | 20 | Baseline | ancom | 2 |
| # | sex | 4 | 0 | 0 | 1 | 1 | # | 20 | Baseline | ancom | 2 |
| # | time | 5 | 4 | 0 | 0.555555556 | 1 | # | 20 | Baseline | ancom | 2 |
| # | interaction | 0 | 0 | 4 | 0 | 0 | # | 20 | Baseline | ancom | 2 |
| # | disease | 7 | 54 | 0 | 0.114754098 | 1 | # | 20 | Baseline | sides2 | 2 |
| # | sex | 4 | 15 | 0 | 0.210526316 | 1 | # | 20 | Baseline | sides2 | 2 |
| # | time | 5 | 37 | 0 | 0.119047619 | 1 | # | 20 | Baseline | sides2 | 2 |
| # | interaction | 0 | 0 | 4 | 0 | 0 | # | 20 | Baseline | sides2 | 2 |
| # | disease | 7 | 0 | 0 | 1 | 1 | # | 20 | Small | paralle | 2 |
| # | sex | 4 | 0 | 0 | 1 | 1 | # | 20 | Small | paralle | 2 |
| # | time | 5 | 0 | 0 | 1 | 1 | # | 20 | Small | paralle | 2 |
| # | interaction | 4 | 0 | 0 | 1 | 1 | # | 20 | Small | paralle | 2 |
| # | disease | 7 | 4 | 0 | 0.636363636 | 1 | # | 20 | Small | ancom | 2 |
| # | sex | 4 | 0 | 0 | 1 | 1 | # | 20 | Small | ancom | 2 |
| # | time | 5 | 4 | 0 | 0.555555556 | 1 | # | 20 | Small | ancom | 2 |
| # | interaction | 0 | 0 | 4 | 0 | 0 | # | 20 | Small | ancom | 2 |
| # | disease | 7 | 4 | 0 | 0.636363636 | 1 | # | 20 | Small | sides2 | 2 |
| # | sex | 4 | 0 | 0 | 1 | 1 | # | 20 | Small | sides2 | 2 |
| # | time | 5 | 4 | 0 | 0.555555556 | 1 | # | 20 | Small | sides2 | 2 |
| # | interaction | 0 | 0 | 4 | 0 | 0 | # | 20 | Small | sides2 | 2 |
| # | disease | 7 | 11 | 0 | 0.388888889 | 1 | # | 20 | Baseline | paralle | 2 |
| # | sex | 4 | 5 | 0 | 0.444444444 | 1 | # | 20 | Baseline | paralle | 2 |
| # | time | 5 | 4 | 0 | 0.555555556 | 1 | # | 20 | Baseline | paralle | 2 |
| # | interaction | 4 | 2 | 0 | 0.666666667 | 1 | # | 20 | Baseline | paralle | 2 |
| # | disease | 7 | 4 | 0 | 0.636363636 | 1 | # | 20 | Baseline | ancom | 2 |
| # | sex | 4 | 0 | 0 | 1 | 1 | # | 20 | Baseline | ancom | 2 |
| # | time | 5 | 4 | 0 | 0.555555556 | 1 | # | 20 | Baseline | ancom | 2 |
| # | interaction | 0 | 0 | 4 | 0 | 0 | # | 20 | Baseline | ancom | 2 |
| # | disease | 7 | 59 | 0 | 0.106060606 | 1 | # | 20 | Baseline | sides2 | 2 |
| # | sex | 4 | 16 | 0 | 0.2 | 1 | # | 20 | Baseline | sides2 | 2 |
| # | time | 5 | 61 | 0 | 0.108695652 | 1 | # | 20 | Baseline | sides2 | 2 |
| # | interaction | 0 | 0 | 4 | 0 | 0 | # | 20 | Baseline | sides2 | 2 |
| # | disease | 7 | 0 | 0 | 1 | 1 | # | 20 | Small | paralle | 2 |
| # | sex | 4 | 0 | 0 | 1 | 1 | # | 20 | Small | paralle | 2 |
| # | time | 5 | 0 | 0 | 1 | 1 | # | 20 | Small | paralle | 2 |
| # | interaction | 4 | 0 | 0 | 1 | 1 | # | 20 | Small | paralle | 2 |
| # | disease | 7 | 4 | 0 | 0.636363636 | 1 | # | 20 | Small | ancom | 2 |
| # | sex | 4 | 0 | 0 | 1 | 1 | # | 20 | Small | ancom | 2 |
| # | time | 5 | 4 | 0 | 0.555555556 | 1 | # | 20 | Small | ancom | 2 |
| # | interaction | 0 | 0 | 4 | 0 | 0 | # | 20 | Small | ancom | 2 |
| # | disease | 7 | 6 | 0 | 0.538461538 | 1 | # | 20 | Small | sides2 | 2 |
| # | sex | 4 | 0 | 0 | 1 | 1 | # | 20 | Small | sides2 | 2 |
| # | time | 5 | 4 | 0 | 0.555555556 | 1 | # | 20 | Small | sides2 | 2 |
| # | interaction | 0 | 0 | 4 | 0 | 0 | # | 20 | Small | sides2 | 2 |
| # | disease | 4 | 7 | 0 | 0.363636364 | 1 | # | 10 | Baseline | paralle | 3 |
| # | sex | 2 | 5 | 0 | 0.285714286 | 1 | # | 10 | Baseline | paralle | 3 |
| # | time | 2 | 4 | 0 | 0.333333333 | 1 | # | 10 | Baseline | paralle | 3 |
| # | interaction | 2 | 7 | 0 | 0.222222222 | 1 | # | 10 | Baseline | paralle | 3 |
| # | disease | 4 | 0 | 0 | 1 | 1 | # | 10 | Baseline | ancom | 3 |
| # | sex | 2 | 0 | 0 | 1 | 1 | # | 10 | Baseline | ancom | 3 |
| # | time | 2 | 0 | 0 | 1 | 1 | # | 10 | Baseline | ancom | 3 |
| # | interaction | 0 | 0 | 2 | 0 | 0 | # | 10 | Baseline | ancom | 3 |
| # | disease | 4 | 7 | 0 | 0.363636364 | 1 | # | 10 | Baseline | sides2 | 3 |
| # | sex | 2 | 5 | 0 | 0.285714286 | 1 | # | 10 | Baseline | sides2 | 3 |
| # | time | 2 | 6 | 0 | 0.25 | 1 | # | 10 | Baseline | sides2 | 3 |
| # | interaction | 0 | 0 | 2 | 0 | 0 | # | 10 | Baseline | sides2 | 3 |
| # | disease | 4 | 0 | 0 | 1 | 1 | # | 10 | Small | paralle | 3 |
| # | sex | 2 | 1 | 0 | 0.666666667 | 1 | # | 10 | Small | paralle | 3 |
| # | time | 2 | 0 | 0 | 1 | 1 | # | 10 | Small | paralle | 3 |
| # | interaction | 2 | 0 | 0 | 1 | 1 | # | 10 | Small | paralle | 3 |
| # | disease | 4 | 0 | 0 | 1 | 1 | # | 10 | Small | ancom | 3 |
| # | sex | 2 | 0 | 0 | 1 | 1 | # | 10 | Small | ancom | 3 |
| # | time | 2 | 0 | 0 | 1 | 1 | # | 10 | Small | ancom | 3 |
| # | interaction | 0 | 0 | 2 | 0 | 0 | # | 10 | Small | ancom | 3 |
| # | disease | 4 | 1 | 0 | 0.8 | 1 | # | 10 | Small | sides2 | 3 |
| # | sex | 2 | 0 | 0 | 1 | 1 | # | 10 | Small | sides2 | 3 |
| # | time | 2 | 1 | 0 | 0.666666667 | 1 | # | 10 | Small | sides2 | 3 |
| # | interaction | 0 | 0 | 2 | 0 | 0 | # | 10 | Small | sides2 | 3 |
| # | disease | 4 | 4 | 0 | 0.5 | 1 | # | 10 | Baseline | paralle | 3 |
| # | sex | 2 | 4 | 0 | 0.333333333 | 1 | # | 10 | Baseline | paralle | 3 |
| # | time | 2 | 3 | 0 | 0.4 | 1 | # | 10 | Baseline | paralle | 3 |
| # | interaction | 2 | 4 | 0 | 0.333333333 | 1 | # | 10 | Baseline | paralle | 3 |
| # | disease | 4 | 2 | 0 | 0.666666667 | 1 | # | 10 | Baseline | ancom | 3 |
| # | sex | 2 | 0 | 0 | 1 | 1 | # | 10 | Baseline | ancom | 3 |
| # | time | 2 | 0 | 0 | 1 | 1 | # | 10 | Baseline | ancom | 3 |
| # | interaction | 0 | 0 | 2 | 0 | 0 | # | 10 | Baseline | ancom | 3 |
| # | disease | 4 | 15 | 0 | 0.210526316 | 1 | # | 10 | Baseline | sides2 | 3 |
| # | sex | 2 | 5 | 0 | 0.285714286 | 1 | # | 10 | Baseline | sides2 | 3 |
| # | time | 2 | 11 | 0 | 0.153846154 | 1 | # | 10 | Baseline | sides2 | 3 |
| # | interaction | 0 | 0 | 2 | 0 | 0 | # | 10 | Baseline | sides2 | 3 |
| # | disease | 4 | 0 | 0 | 1 | 1 | # | 10 | Small | paralle | 3 |
| # | sex | 2 | 1 | 0 | 0.666666667 | 1 | # | 10 | Small | paralle | 3 |
| # | time | 2 | 0 | 0 | 1 | 1 | # | 10 | Small | paralle | 3 |
| # | interaction | 2 | 0 | 0 | 1 | 1 | # | 10 | Small | paralle | 3 |
| # | disease | 4 | 2 | 0 | 0.666666667 | 1 | # | 10 | Small | ancom | 3 |
| # | sex | 2 | 0 | 0 | 1 | 1 | # | 10 | Small | ancom | 3 |
| # | time | 2 | 0 | 0 | 1 | 1 | # | 10 | Small | ancom | 3 |
| # | interaction | 0 | 0 | 2 | 0 | 0 | # | 10 | Small | ancom | 3 |
| # | disease | 4 | 2 | 0 | 0.666666667 | 1 | # | 10 | Small | sides2 | 3 |
| # | sex | 2 | 0 | 0 | 1 | 1 | # | 10 | Small | sides2 | 3 |
| # | time | 2 | 2 | 0 | 0.5 | 1 | # | 10 | Small | sides2 | 3 |
| # | interaction | 0 | 0 | 2 | 0 | 0 | # | 10 | Small | sides2 | 3 |
| # | disease | 4 | 1 | 0 | 0.8 | 1 | # | 10 | Baseline | paralle | 3 |
| # | sex | 2 | 4 | 0 | 0.333333333 | 1 | # | 10 | Baseline | paralle | 3 |
| # | time | 2 | 3 | 0 | 0.4 | 1 | # | 10 | Baseline | paralle | 3 |
| # | interaction | 2 | 3 | 0 | 0.4 | 1 | # | 10 | Baseline | paralle | 3 |
| # | disease | 4 | 2 | 0 | 0.666666667 | 1 | # | 10 | Baseline | ancom | 3 |
| # | sex | 2 | 0 | 0 | 1 | 1 | # | 10 | Baseline | ancom | 3 |
| # | time | 2 | 2 | 0 | 0.5 | 1 | # | 10 | Baseline | ancom | 3 |
| # | interaction | 0 | 0 | 2 | 0 | 0 | # | 10 | Baseline | ancom | 3 |
| # | disease | 4 | 18 | 0 | 0.181818182 | 1 | # | 10 | Baseline | sides2 | 3 |
| # | sex | 2 | 4 | 0 | 0.333333333 | 1 | # | 10 | Baseline | sides2 | 3 |
| # | time | 2 | 14 | 0 | 0.125 | 1 | # | 10 | Baseline | sides2 | 3 |
| # | interaction | 0 | 0 | 2 | 0 | 0 | # | 10 | Baseline | sides2 | 3 |
| # | disease | 4 | 0 | 0 | 1 | 1 | # | 10 | Small | paralle | 3 |
| # | sex | 2 | 0 | 0 | 1 | 1 | # | 10 | Small | paralle | 3 |
| # | time | 2 | 0 | 0 | 1 | 1 | # | 10 | Small | paralle | 3 |
| # | interaction | 2 | 0 | 0 | 1 | 1 | # | 10 | Small | paralle | 3 |
| # | disease | 4 | 2 | 0 | 0.666666667 | 1 | # | 10 | Small | ancom | 3 |
| # | sex | 2 | 0 | 0 | 1 | 1 | # | 10 | Small | ancom | 3 |
| # | time | 2 | 2 | 0 | 0.5 | 1 | # | 10 | Small | ancom | 3 |
| # | interaction | 0 | 0 | 2 | 0 | 0 | # | 10 | Small | ancom | 3 |
| # | disease | 4 | 2 | 0 | 0.666666667 | 1 | # | 10 | Small | sides2 | 3 |
| # | sex | 2 | 0 | 0 | 1 | 1 | # | 10 | Small | sides2 | 3 |
| # | time | 2 | 3 | 0 | 0.4 | 1 | # | 10 | Small | sides2 | 3 |
| # | interaction | 0 | 0 | 2 | 0 | 0 | # | 10 | Small | sides2 | 3 |
| # | disease | 4 | 2 | 0 | 0.666666667 | 1 | # | 10 | Baseline | paralle | 3 |
| # | sex | 2 | 1 | 0 | 0.666666667 | 1 | # | 10 | Baseline | paralle | 3 |
| # | time | 2 | 6 | 0 | 0.25 | 1 | # | 10 | Baseline | paralle | 3 |
| # | interaction | 2 | 2 | 0 | 0.5 | 1 | # | 10 | Baseline | paralle | 3 |
| # | disease | 4 | 2 | 0 | 0.666666667 | 1 | # | 10 | Baseline | ancom | 3 |
| # | sex | 2 | 0 | 0 | 1 | 1 | # | 10 | Baseline | ancom | 3 |
| # | time | 2 | 2 | 0 | 0.5 | 1 | # | 10 | Baseline | ancom | 3 |
| # | interaction | 0 | 0 | 2 | 0 | 0 | # | 10 | Baseline | ancom | 3 |
| # | disease | 4 | 25 | 0 | 0.137931034 | 1 | # | 10 | Baseline | sides2 | 3 |
| # | sex | 2 | 1 | 0 | 0.666666667 | 1 | # | 10 | Baseline | sides2 | 3 |
| # | time | 2 | 14 | 0 | 0.125 | 1 | # | 10 | Baseline | sides2 | 3 |
| # | interaction | 0 | 0 | 2 | 0 | 0 | # | 10 | Baseline | sides2 | 3 |
| # | disease | 4 | 0 | 0 | 1 | 1 | # | 10 | Small | paralle | 3 |
| # | sex | 2 | 0 | 0 | 1 | 1 | # | 10 | Small | paralle | 3 |
| # | time | 2 | 0 | 0 | 1 | 1 | # | 10 | Small | paralle | 3 |
| # | interaction | 2 | 0 | 0 | 1 | 1 | # | 10 | Small | paralle | 3 |
| # | disease | 4 | 2 | 0 | 0.666666667 | 1 | # | 10 | Small | ancom | 3 |
| # | sex | 2 | 0 | 0 | 1 | 1 | # | 10 | Small | ancom | 3 |
| # | time | 2 | 2 | 0 | 0.5 | 1 | # | 10 | Small | ancom | 3 |
| # | interaction | 0 | 0 | 2 | 0 | 0 | # | 10 | Small | ancom | 3 |
| # | disease | 4 | 2 | 0 | 0.666666667 | 1 | # | 10 | Small | sides2 | 3 |
| # | sex | 2 | 0 | 0 | 1 | 1 | # | 10 | Small | sides2 | 3 |
| # | time | 2 | 3 | 0 | 0.4 | 1 | # | 10 | Small | sides2 | 3 |
| # | interaction | 0 | 0 | 2 | 0 | 0 | # | 10 | Small | sides2 | 3 |
| # | disease | 7 | 12 | 0 | 0.368421053 | 1 | # | 20 | Baseline | paralle | 3 |
| # | sex | 4 | 9 | 0 | 0.307692308 | 1 | # | 20 | Baseline | paralle | 3 |
| # | time | 5 | 11 | 0 | 0.3125 | 1 | # | 20 | Baseline | paralle | 3 |
| # | interaction | 4 | 9 | 0 | 0.307692308 | 1 | # | 20 | Baseline | paralle | 3 |
| # | disease | 7 | 2 | 0 | 0.777777778 | 1 | # | 20 | Baseline | ancom | 3 |
| # | sex | 4 | 0 | 0 | 1 | 1 | # | 20 | Baseline | ancom | 3 |
| # | time | 5 | 1 | 0 | 0.833333333 | 1 | # | 20 | Baseline | ancom | 3 |
| # | interaction | 0 | 0 | 4 | 0 | 0 | # | 20 | Baseline | ancom | 3 |
| # | disease | 7 | 16 | 0 | 0.304347826 | 1 | # | 20 | Baseline | sides2 | 3 |
| # | sex | 4 | 5 | 0 | 0.444444444 | 1 | # | 20 | Baseline | sides2 | 3 |
| # | time | 5 | 16 | 0 | 0.238095238 | 1 | # | 20 | Baseline | sides2 | 3 |
| # | interaction | 0 | 0 | 4 | 0 | 0 | # | 20 | Baseline | sides2 | 3 |
| # | disease | 7 | 0 | 0 | 1 | 1 | # | 20 | Small | paralle | 3 |
| # | sex | 4 | 0 | 0 | 1 | 1 | # | 20 | Small | paralle | 3 |
| # | time | 5 | 0 | 0 | 1 | 1 | # | 20 | Small | paralle | 3 |
| # | interaction | 3 | 1 | 1 | 0.75 | 0.75 | # | 20 | Small | paralle | 3 |
| # | disease | 7 | 2 | 0 | 0.777777778 | 1 | # | 20 | Small | ancom | 3 |
| # | sex | 4 | 0 | 0 | 1 | 1 | # | 20 | Small | ancom | 3 |
| # | time | 5 | 1 | 0 | 0.833333333 | 1 | # | 20 | Small | ancom | 3 |
| # | interaction | 0 | 0 | 4 | 0 | 0 | # | 20 | Small | ancom | 3 |
| # | disease | 7 | 1 | 0 | 0.875 | 1 | # | 20 | Small | sides2 | 3 |
| # | sex | 4 | 0 | 0 | 1 | 1 | # | 20 | Small | sides2 | 3 |
| # | time | 5 | 1 | 0 | 0.833333333 | 1 | # | 20 | Small | sides2 | 3 |
| # | interaction | 0 | 0 | 4 | 0 | 0 | # | 20 | Small | sides2 | 3 |
| # | disease | 7 | 6 | 0 | 0.538461538 | 1 | # | 20 | Baseline | paralle | 3 |
| # | sex | 4 | 8 | 0 | 0.333333333 | 1 | # | 20 | Baseline | paralle | 3 |
| # | time | 5 | 10 | 0 | 0.333333333 | 1 | # | 20 | Baseline | paralle | 3 |
| # | interaction | 4 | 3 | 0 | 0.571428571 | 1 | # | 20 | Baseline | paralle | 3 |
| # | disease | 7 | 4 | 0 | 0.636363636 | 1 | # | 20 | Baseline | ancom | 3 |
| # | sex | 4 | 0 | 0 | 1 | 1 | # | 20 | Baseline | ancom | 3 |
| # | time | 5 | 3 | 0 | 0.625 | 1 | # | 20 | Baseline | ancom | 3 |
| # | interaction | 0 | 0 | 4 | 0 | 0 | # | 20 | Baseline | ancom | 3 |
| # | disease | 7 | 38 | 0 | 0.155555556 | 1 | # | 20 | Baseline | sides2 | 3 |
| # | sex | 4 | 12 | 0 | 0.25 | 1 | # | 20 | Baseline | sides2 | 3 |
| # | time | 5 | 20 | 0 | 0.2 | 1 | # | 20 | Baseline | sides2 | 3 |
| # | interaction | 0 | 0 | 4 | 0 | 0 | # | 20 | Baseline | sides2 | 3 |
| # | disease | 7 | 0 | 0 | 1 | 1 | # | 20 | Small | paralle | 3 |
| # | sex | 4 | 0 | 0 | 1 | 1 | # | 20 | Small | paralle | 3 |
| # | time | 5 | 0 | 0 | 1 | 1 | # | 20 | Small | paralle | 3 |
| # | interaction | 4 | 0 | 0 | 1 | 1 | # | 20 | Small | paralle | 3 |
| # | disease | 7 | 4 | 0 | 0.636363636 | 1 | # | 20 | Small | ancom | 3 |
| # | sex | 4 | 0 | 0 | 1 | 1 | # | 20 | Small | ancom | 3 |
| # | time | 5 | 2 | 0 | 0.714285714 | 1 | # | 20 | Small | ancom | 3 |
| # | interaction | 0 | 0 | 4 | 0 | 0 | # | 20 | Small | ancom | 3 |
| # | disease | 7 | 3 | 0 | 0.7 | 1 | # | 20 | Small | sides2 | 3 |
| # | sex | 4 | 0 | 0 | 1 | 1 | # | 20 | Small | sides2 | 3 |
| # | time | 5 | 2 | 0 | 0.714285714 | 1 | # | 20 | Small | sides2 | 3 |
| # | interaction | 0 | 0 | 4 | 0 | 0 | # | 20 | Small | sides2 | 3 |
| # | disease | 7 | 12 | 0 | 0.368421053 | 1 | # | 20 | Baseline | paralle | 3 |
| # | sex | 4 | 7 | 0 | 0.363636364 | 1 | # | 20 | Baseline | paralle | 3 |
| # | time | 5 | 8 | 0 | 0.384615385 | 1 | # | 20 | Baseline | paralle | 3 |
| # | interaction | 4 | 1 | 0 | 0.8 | 1 | # | 20 | Baseline | paralle | 3 |
| # | disease | 7 | 4 | 0 | 0.636363636 | 1 | # | 20 | Baseline | ancom | 3 |
| # | sex | 4 | 0 | 0 | 1 | 1 | # | 20 | Baseline | ancom | 3 |
| # | time | 5 | 4 | 0 | 0.555555556 | 1 | # | 20 | Baseline | ancom | 3 |
| # | interaction | 0 | 0 | 4 | 0 | 0 | # | 20 | Baseline | ancom | 3 |
| # | disease | 7 | 52 | 0 | 0.118644068 | 1 | # | 20 | Baseline | sides2 | 3 |
| # | sex | 4 | 16 | 0 | 0.2 | 1 | # | 20 | Baseline | sides2 | 3 |
| # | time | 5 | 38 | 0 | 0.11627907 | 1 | # | 20 | Baseline | sides2 | 3 |
| # | interaction | 0 | 0 | 4 | 0 | 0 | # | 20 | Baseline | sides2 | 3 |
| # | disease | 7 | 0 | 0 | 1 | 1 | # | 20 | Small | paralle | 3 |
| # | sex | 4 | 0 | 0 | 1 | 1 | # | 20 | Small | paralle | 3 |
| # | time | 5 | 0 | 0 | 1 | 1 | # | 20 | Small | paralle | 3 |
| # | interaction | 4 | 0 | 0 | 1 | 1 | # | 20 | Small | paralle | 3 |
| # | disease | 7 | 4 | 0 | 0.636363636 | 1 | # | 20 | Small | ancom | 3 |
| # | sex | 4 | 0 | 0 | 1 | 1 | # | 20 | Small | ancom | 3 |
| # | time | 5 | 4 | 0 | 0.555555556 | 1 | # | 20 | Small | ancom | 3 |
| # | interaction | 0 | 0 | 4 | 0 | 0 | # | 20 | Small | ancom | 3 |
| # | disease | 7 | 3 | 0 | 0.7 | 1 | # | 20 | Small | sides2 | 3 |
| # | sex | 4 | 0 | 0 | 1 | 1 | # | 20 | Small | sides2 | 3 |
| # | time | 5 | 2 | 0 | 0.714285714 | 1 | # | 20 | Small | sides2 | 3 |
| # | interaction | 0 | 0 | 4 | 0 | 0 | # | 20 | Small | sides2 | 3 |
| # | disease | 7 | 10 | 0 | 0.411764706 | 1 | # | 20 | Baseline | paralle | 3 |
| # | sex | 4 | 9 | 0 | 0.307692308 | 1 | # | 20 | Baseline | paralle | 3 |
| # | time | 5 | 8 | 0 | 0.384615385 | 1 | # | 20 | Baseline | paralle | 3 |
| # | interaction | 4 | 1 | 0 | 0.8 | 1 | # | 20 | Baseline | paralle | 3 |
| # | disease | 7 | 4 | 0 | 0.636363636 | 1 | # | 20 | Baseline | ancom | 3 |
| # | sex | 4 | 0 | 0 | 1 | 1 | # | 20 | Baseline | ancom | 3 |
| # | time | 5 | 4 | 0 | 0.555555556 | 1 | # | 20 | Baseline | ancom | 3 |
| # | interaction | 0 | 0 | 4 | 0 | 0 | # | 20 | Baseline | ancom | 3 |
| # | disease | 7 | 53 | 0 | 0.1 | 1 | # | 20 | Baseline | sides2 | 3 |
| # | sex | 4 | 18 | 0 | 0.181818182 | 1 | # | 20 | Baseline | sides2 | 3 |
| # | time | 5 | 63 | 0 | 0.104166667 | 1 | # | 20 | Baseline | sides2 | 3 |
| # | interaction | 0 | 0 | 4 | 0 | 0 | # | 20 | Baseline | sides2 | 3 |
| # | disease | 7 | 0 | 0 | 1 | 1 | # | 20 | Small | paralle | 3 |
| # | sex | 4 | 0 | 0 | 1 | 1 | # | 20 | Small | paralle | 3 |
| # | time | 5 | 0 | 0 | 1 | 1 | # | 20 | Small | paralle | 3 |
| # | interaction | 4 | 0 | 0 | 1 | 1 | # | 20 | Small | paralle | 3 |
| # | disease | 7 | 4 | 0 | 0.636363636 | 1 | # | 20 | Small | ancom | 3 |
| # | sex | 4 | 0 | 0 | 1 | 1 | # | 20 | Small | ancom | 3 |
| # | time | 5 | 4 | 0 | 0.555555556 | 1 | # | 20 | Small | ancom | 3 |
| # | interaction | 0 | 0 | 4 | 0 | 0 | # | 20 | Small | ancom | 3 |
| # | disease | 7 | 7 | 0 | 0.5 | 1 | # | 20 | Small | sides2 | 3 |
| # | sex | 4 | 0 | 0 | 1 | 1 | # | 20 | Small | sides2 | 3 |
| # | time | 5 | 4 | 0 | 0.555555556 | 1 | # | 20 | Small | sides2 | 3 |
| # | interaction | 0 | 0 | 4 | 0 | 0 | # | 20 | Small | sides2 | 3 |
| # | disease | 4 | 10 | 0 | 0.285714286 | 1 | # | 10 | Baseline | paralle | 4 |
| # | sex | 2 | 8 | 0 | 0.2 | 1 | # | 10 | Baseline | paralle | 4 |
| # | time | 2 | 10 | 0 | 0.166666667 | 1 | # | 10 | Baseline | paralle | 4 |
| # | interaction | 2 | 11 | 0 | 0.153846154 | 1 | # | 10 | Baseline | paralle | 4 |
| # | disease | 4 | 1 | 0 | 0.8 | 1 | # | 10 | Baseline | ancom | 4 |
| # | sex | 2 | 0 | 0 | 1 | 1 | # | 10 | Baseline | ancom | 4 |
| # | time | 2 | 1 | 0 | 0.666666667 | 1 | # | 10 | Baseline | ancom | 4 |
| # | interaction | 0 | 0 | 2 | 0 | 0 | # | 10 | Baseline | ancom | 4 |
| # | disease | 4 | 11 | 0 | 0.266666667 | 1 | # | 10 | Baseline | sides2 | 4 |
| # | sex | 2 | 6 | 0 | 0.25 | 1 | # | 10 | Baseline | sides2 | 4 |
| # | time | 2 | 6 | 0 | 0.25 | 1 | # | 10 | Baseline | sides2 | 4 |
| # | interaction | 0 | 0 | 2 | 0 | 0 | # | 10 | Baseline | sides2 | 4 |
| # | disease | 4 | 1 | 0 | 0.8 | 1 | # | 10 | Small | paralle | 4 |
| # | sex | 2 | 0 | 0 | 1 | 1 | # | 10 | Small | paralle | 4 |
| # | time | 2 | 1 | 0 | 0.666666667 | 1 | # | 10 | Small | paralle | 4 |
| # | interaction | 1 | 0 | 1 | 1 | 0.5 | # | 10 | Small | paralle | 4 |
| # | disease | 4 | 1 | 0 | 0.8 | 1 | # | 10 | Small | ancom | 4 |
| # | sex | 2 | 0 | 0 | 1 | 1 | # | 10 | Small | ancom | 4 |
| # | time | 2 | 1 | 0 | 0.666666667 | 1 | # | 10 | Small | ancom | 4 |
| # | interaction | 0 | 0 | 2 | 0 | 0 | # | 10 | Small | ancom | 4 |
| # | disease | 4 | 2 | 0 | 0.666666667 | 1 | # | 10 | Small | sides2 | 4 |
| # | sex | 2 | 1 | 0 | 0.666666667 | 1 | # | 10 | Small | sides2 | 4 |
| # | time | 2 | 2 | 0 | 0.5 | 1 | # | 10 | Small | sides2 | 4 |
| # | interaction | 0 | 0 | 2 | 0 | 0 | # | 10 | Small | sides2 | 4 |
| # | disease | 4 | 11 | 0 | 0.266666667 | 1 | # | 10 | Baseline | paralle | 4 |
| # | sex | 2 | 7 | 0 | 0.222222222 | 1 | # | 10 | Baseline | paralle | 4 |
| # | time | 2 | 8 | 0 | 0.2 | 1 | # | 10 | Baseline | paralle | 4 |
| # | interaction | 2 | 9 | 0 | 0.181818182 | 1 | # | 10 | Baseline | paralle | 4 |
| # | disease | 4 | 2 | 0 | 0.666666667 | 1 | # | 10 | Baseline | ancom | 4 |
| # | sex | 2 | 0 | 0 | 1 | 1 | # | 10 | Baseline | ancom | 4 |
| # | time | 2 | 2 | 0 | 0.5 | 1 | # | 10 | Baseline | ancom | 4 |
| # | interaction | 0 | 0 | 2 | 0 | 0 | # | 10 | Baseline | ancom | 4 |
| # | disease | 4 | 15 | 0 | 0.210526316 | 1 | # | 10 | Baseline | sides2 | 4 |
| # | sex | 2 | 5 | 0 | 0.285714286 | 1 | # | 10 | Baseline | sides2 | 4 |
| # | time | 2 | 7 | 0 | 0.222222222 | 1 | # | 10 | Baseline | sides2 | 4 |
| # | interaction | 0 | 0 | 2 | 0 | 0 | # | 10 | Baseline | sides2 | 4 |
| # | disease | 4 | 0 | 0 | 1 | 1 | # | 10 | Small | paralle | 4 |
| # | sex | 2 | 0 | 0 | 1 | 1 | # | 10 | Small | paralle | 4 |
| # | time | 2 | 0 | 0 | 1 | 1 | # | 10 | Small | paralle | 4 |
| # | interaction | 1 | 0 | 1 | 1 | 0.5 | # | 10 | Small | paralle | 4 |
| # | disease | 4 | 1 | 0 | 0.8 | 1 | # | 10 | Small | ancom | 4 |
| # | sex | 2 | 0 | 0 | 1 | 1 | # | 10 | Small | ancom | 4 |
| # | time | 2 | 2 | 0 | 0.5 | 1 | # | 10 | Small | ancom | 4 |
| # | interaction | 0 | 0 | 2 | 0 | 0 | # | 10 | Small | ancom | 4 |
| # | disease | 4 | 1 | 0 | 0.8 | 1 | # | 10 | Small | sides2 | 4 |
| # | sex | 2 | 0 | 0 | 1 | 1 | # | 10 | Small | sides2 | 4 |
| # | time | 2 | 2 | 0 | 0.5 | 1 | # | 10 | Small | sides2 | 4 |
| # | interaction | 0 | 0 | 2 | 0 | 0 | # | 10 | Small | sides2 | 4 |
| # | disease | 4 | 9 | 0 | 0.307692308 | 1 | # | 10 | Baseline | paralle | 4 |
| # | sex | 2 | 3 | 0 | 0.4 | 1 | # | 10 | Baseline | paralle | 4 |
| # | time | 2 | 5 | 0 | 0.285714286 | 1 | # | 10 | Baseline | paralle | 4 |
| # | interaction | 2 | 10 | 0 | 0.166666667 | 1 | # | 10 | Baseline | paralle | 4 |
| # | disease | 4 | 2 | 0 | 0.666666667 | 1 | # | 10 | Baseline | ancom | 4 |
| # | sex | 2 | 0 | 0 | 1 | 1 | # | 10 | Baseline | ancom | 4 |
| # | time | 2 | 2 | 0 | 0.5 | 1 | # | 10 | Baseline | ancom | 4 |
| # | interaction | 0 | 0 | 2 | 0 | 0 | # | 10 | Baseline | ancom | 4 |
| # | disease | 4 | 18 | 0 | 0.181818182 | 1 | # | 10 | Baseline | sides2 | 4 |
| # | sex | 2 | 8 | 0 | 0.2 | 1 | # | 10 | Baseline | sides2 | 4 |
| # | time | 2 | 11 | 0 | 0.153846154 | 1 | # | 10 | Baseline | sides2 | 4 |
| # | interaction | 0 | 0 | 2 | 0 | 0 | # | 10 | Baseline | sides2 | 4 |
| # | disease | 4 | 0 | 0 | 1 | 1 | # | 10 | Small | paralle | 4 |
| # | sex | 2 | 0 | 0 | 1 | 1 | # | 10 | Small | paralle | 4 |
| # | time | 2 | 0 | 0 | 1 | 1 | # | 10 | Small | paralle | 4 |
| # | interaction | 2 | 0 | 0 | 1 | 1 | # | 10 | Small | paralle | 4 |
| # | disease | 4 | 2 | 0 | 0.666666667 | 1 | # | 10 | Small | ancom | 4 |
| # | sex | 2 | 0 | 0 | 1 | 1 | # | 10 | Small | ancom | 4 |
| # | time | 2 | 2 | 0 | 0.5 | 1 | # | 10 | Small | ancom | 4 |
| # | interaction | 0 | 0 | 2 | 0 | 0 | # | 10 | Small | ancom | 4 |
| # | disease | 4 | 3 | 0 | 0.571428571 | 1 | # | 10 | Small | sides2 | 4 |
| # | sex | 2 | 0 | 0 | 1 | 1 | # | 10 | Small | sides2 | 4 |
| # | time | 2 | 2 | 0 | 0.5 | 1 | # | 10 | Small | sides2 | 4 |
| # | interaction | 0 | 0 | 2 | 0 | 0 | # | 10 | Small | sides2 | 4 |
| # | disease | 4 | 4 | 0 | 0.5 | 1 | # | 10 | Baseline | paralle | 4 |
| # | sex | 2 | 1 | 0 | 0.666666667 | 1 | # | 10 | Baseline | paralle | 4 |
| # | time | 2 | 6 | 0 | 0.25 | 1 | # | 10 | Baseline | paralle | 4 |
| # | interaction | 2 | 9 | 0 | 0.181818182 | 1 | # | 10 | Baseline | paralle | 4 |
| # | disease | 4 | 2 | 0 | 0.666666667 | 1 | # | 10 | Baseline | ancom | 4 |
| # | sex | 2 | 0 | 0 | 1 | 1 | # | 10 | Baseline | ancom | 4 |
| # | time | 2 | 2 | 0 | 0.5 | 1 | # | 10 | Baseline | ancom | 4 |
| # | interaction | 0 | 0 | 2 | 0 | 0 | # | 10 | Baseline | ancom | 4 |
| # | disease | 4 | 19 | 0 | 0.173913043 | 1 | # | 10 | Baseline | sides2 | 4 |
| # | sex | 2 | 6 | 0 | 0.25 | 1 | # | 10 | Baseline | sides2 | 4 |
| # | time | 2 | 9 | 0 | 0.181818182 | 1 | # | 10 | Baseline | sides2 | 4 |
| # | interaction | 0 | 0 | 2 | 0 | 0 | # | 10 | Baseline | sides2 | 4 |
| # | disease | 4 | 0 | 0 | 1 | 1 | # | 10 | Small | paralle | 4 |
| # | sex | 2 | 0 | 0 | 1 | 1 | # | 10 | Small | paralle | 4 |
| # | time | 2 | 0 | 0 | 1 | 1 | # | 10 | Small | paralle | 4 |
| # | interaction | 2 | 0 | 0 | 1 | 1 | # | 10 | Small | paralle | 4 |
| # | disease | 4 | 2 | 0 | 0.666666667 | 1 | # | 10 | Small | ancom | 4 |
| # | sex | 2 | 0 | 0 | 1 | 1 | # | 10 | Small | ancom | 4 |
| # | time | 2 | 2 | 0 | 0.5 | 1 | # | 10 | Small | ancom | 4 |
| # | interaction | 0 | 0 | 2 | 0 | 0 | # | 10 | Small | ancom | 4 |
| # | disease | 4 | 2 | 0 | 0.666666667 | 1 | # | 10 | Small | sides2 | 4 |
| # | sex | 2 | 0 | 0 | 1 | 1 | # | 10 | Small | sides2 | 4 |
| # | time | 2 | 2 | 0 | 0.5 | 1 | # | 10 | Small | sides2 | 4 |

| | | | | | | | | | | | |
|---|---|---|---|---|---|---|---|---|---|---|---|
| # | interaction | 0 | 0 | 2 | 0 | 0 | # | 10 | Small | aldex2 | 4 |
| # | disease | 7 | 8 | 0 | 0.466666667 | 1 | # | 20 | Baseline | paralle | 4 |
| # | sex | 4 | 11 | 0 | 0.266666667 | 1 | # | 20 | Baseline | paralle | 4 |
| # | time | 5 | 14 | 0 | 0.263157895 | 1 | # | 20 | Baseline | paralle | 4 |
| # | interaction | 3 | 6 | 1 | 0.333333333 | 0.75 | # | 20 | Baseline | paralle | 4 |
| # | disease | 7 | 2 | 0 | 0.777777778 | 1 | # | 20 | Baseline | ancom | 4 |
| # | sex | 4 | 0 | 0 | 1 | 1 | # | 20 | Baseline | ancom | 4 |
| # | time | 5 | 4 | 0 | 0.555555556 | 1 | # | 20 | Baseline | ancom | 4 |
| # | interaction | 0 | 0 | 4 | 0 | 0 | # | 20 | Baseline | ancom | 4 |
| # | disease | 7 | 22 | 0 | 0.24137931 | 1 | # | 20 | Baseline | aldex2 | 4 |
| # | sex | 4 | 7 | 0 | 0.363636364 | 1 | # | 20 | Baseline | aldex2 | 4 |
| # | time | 5 | 16 | 0 | 0.238095238 | 1 | # | 20 | Baseline | aldex2 | 4 |
| # | interaction | 0 | 0 | 4 | 0 | 0 | # | 20 | Baseline | aldex2 | 4 |
| # | disease | 7 | 0 | 0 | 1 | 1 | # | 20 | Small | paralle | 4 |
| # | sex | 4 | 0 | 0 | 1 | 1 | # | 20 | Small | paralle | 4 |
| # | time | 5 | 0 | 0 | 1 | 1 | # | 20 | Small | paralle | 4 |
| # | interaction | 1 | 0 | 3 | 1 | 0.25 | # | 20 | Small | paralle | 4 |
| # | disease | 7 | 1 | 0 | 0.875 | 1 | # | 20 | Small | ancom | 4 |
| # | sex | 4 | 0 | 0 | 1 | 1 | # | 20 | Small | ancom | 4 |
| # | time | 5 | 4 | 0 | 0.555555556 | 1 | # | 20 | Small | ancom | 4 |
| # | interaction | 0 | 0 | 4 | 0 | 0 | # | 20 | Small | ancom | 4 |
| # | disease | 7 | 1 | 0 | 0.875 | 1 | # | 20 | Small | aldex2 | 4 |
| # | sex | 4 | 0 | 0 | 1 | 1 | # | 20 | Small | aldex2 | 4 |
| # | time | 5 | 5 | 0 | 0.5 | 1 | # | 20 | Small | aldex2 | 4 |
| # | interaction | 0 | 0 | 4 | 0 | 0 | # | 20 | Small | aldex2 | 4 |
| # | disease | 7 | 13 | 0 | 0.35 | 1 | # | 20 | Baseline | paralle | 4 |
| # | sex | 4 | 3 | 0 | 0.571428571 | 1 | # | 20 | Baseline | paralle | 4 |
| # | time | 5 | 13 | 0 | 0.277777778 | 1 | # | 20 | Baseline | paralle | 4 |
| # | interaction | 4 | 8 | 0 | 0.333333333 | 1 | # | 20 | Baseline | paralle | 4 |
| # | disease | 7 | 4 | 0 | 0.636363636 | 1 | # | 20 | Baseline | ancom | 4 |
| # | sex | 4 | 0 | 0 | 1 | 1 | # | 20 | Baseline | ancom | 4 |
| # | time | 5 | 4 | 0 | 0.555555556 | 1 | # | 20 | Baseline | ancom | 4 |
| # | interaction | 0 | 0 | 4 | 0 | 0 | # | 20 | Baseline | ancom | 4 |
| # | disease | 7 | 33 | 0 | 0.175 | 1 | # | 20 | Baseline | aldex2 | 4 |
| # | sex | 4 | 6 | 0 | 0.4 | 1 | # | 20 | Baseline | aldex2 | 4 |
| # | time | 5 | 24 | 0 | 0.172413793 | 1 | # | 20 | Baseline | aldex2 | 4 |
| # | interaction | 0 | 0 | 4 | 0 | 0 | # | 20 | Baseline | aldex2 | 4 |
| # | disease | 7 | 0 | 0 | 1 | 1 | # | 20 | Small | paralle | 4 |
| # | sex | 4 | 0 | 0 | 1 | 1 | # | 20 | Small | paralle | 4 |
| # | time | 5 | 0 | 0 | 1 | 1 | # | 20 | Small | paralle | 4 |
| # | interaction | 3 | 0 | 1 | 1 | 0.75 | # | 20 | Small | paralle | 4 |
| # | disease | 7 | 4 | 0 | 0.636363636 | 1 | # | 20 | Small | ancom | 4 |
| # | sex | 4 | 0 | 0 | 1 | 1 | # | 20 | Small | ancom | 4 |
| # | time | 5 | 4 | 0 | 0.555555556 | 1 | # | 20 | Small | ancom | 4 |
| # | interaction | 0 | 0 | 4 | 0 | 0 | # | 20 | Small | ancom | 4 |
| # | disease | 7 | 5 | 0 | 0.583333333 | 1 | # | 20 | Small | aldex2 | 4 |
| # | sex | 4 | 0 | 0 | 1 | 1 | # | 20 | Small | aldex2 | 4 |
| # | time | 5 | 4 | 0 | 0.555555556 | 1 | # | 20 | Small | aldex2 | 4 |
| # | interaction | 0 | 0 | 4 | 0 | 0 | # | 20 | Small | aldex2 | 4 |
| # | disease | 7 | 10 | 0 | 0.411764706 | 1 | # | 20 | Baseline | paralle | 4 |
| # | sex | 4 | 3 | 0 | 0.571428571 | 1 | # | 20 | Baseline | paralle | 4 |
| # | time | 5 | 8 | 0 | 0.384615385 | 1 | # | 20 | Baseline | paralle | 4 |
| # | interaction | 4 | 5 | 0 | 0.444444444 | 1 | # | 20 | Baseline | paralle | 4 |
| # | disease | 7 | 4 | 0 | 0.636363636 | 1 | # | 20 | Baseline | ancom | 4 |
| # | sex | 4 | 0 | 0 | 1 | 1 | # | 20 | Baseline | ancom | 4 |
| # | time | 5 | 4 | 0 | 0.555555556 | 1 | # | 20 | Baseline | ancom | 4 |
| # | interaction | 0 | 0 | 4 | 0 | 0 | # | 20 | Baseline | ancom | 4 |
| # | disease | 7 | 51 | 0 | 0.120689655 | 1 | # | 20 | Baseline | aldex2 | 4 |
| # | sex | 4 | 12 | 0 | 0.25 | 1 | # | 20 | Baseline | aldex2 | 4 |
| # | time | 5 | 36 | 0 | 0.12195122 | 1 | # | 20 | Baseline | aldex2 | 4 |
| # | interaction | 0 | 0 | 4 | 0 | 0 | # | 20 | Baseline | aldex2 | 4 |
| # | disease | 7 | 0 | 0 | 1 | 1 | # | 20 | Small | paralle | 4 |
| # | sex | 4 | 0 | 0 | 1 | 1 | # | 20 | Small | paralle | 4 |
| # | time | 5 | 0 | 0 | 1 | 1 | # | 20 | Small | paralle | 4 |
| # | interaction | 3 | 0 | 1 | 1 | 0.75 | # | 20 | Small | paralle | 4 |
| # | disease | 7 | 4 | 0 | 0.636363636 | 1 | # | 20 | Small | ancom | 4 |
| # | sex | 4 | 0 | 0 | 1 | 1 | # | 20 | Small | ancom | 4 |
| # | time | 5 | 4 | 0 | 0.555555556 | 1 | # | 20 | Small | ancom | 4 |
| # | interaction | 0 | 0 | 4 | 0 | 0 | # | 20 | Small | ancom | 4 |
| # | disease | 7 | 8 | 0 | 0.466666667 | 1 | # | 20 | Small | aldex2 | 4 |
| # | sex | 4 | 0 | 0 | 1 | 1 | # | 20 | Small | aldex2 | 4 |
| # | time | 5 | 4 | 0 | 0.555555556 | 1 | # | 20 | Small | aldex2 | 4 |
| # | interaction | 0 | 0 | 4 | 0 | 0 | # | 20 | Small | aldex2 | 4 |
| # | disease | 7 | 15 | 0 | 0.318181818 | 1 | # | 20 | Baseline | paralle | 4 |
| # | sex | 4 | 3 | 0 | 0.571428571 | 1 | # | 20 | Baseline | paralle | 4 |
| # | time | 5 | 7 | 0 | 0.416666667 | 1 | # | 20 | Baseline | paralle | 4 |
| # | interaction | 4 | 3 | 0 | 0.571428571 | 1 | # | 20 | Baseline | paralle | 4 |
| # | disease | 7 | 4 | 0 | 0.636363636 | 1 | # | 20 | Baseline | ancom | 4 |
| # | sex | 4 | 0 | 0 | 1 | 1 | # | 20 | Baseline | ancom | 4 |
| # | time | 5 | 4 | 0 | 0.555555556 | 1 | # | 20 | Baseline | ancom | 4 |
| # | interaction | 0 | 0 | 4 | 0 | 0 | # | 20 | Baseline | ancom | 4 |
| # | disease | 7 | 57 | 0 | 0.109375 | 1 | # | 20 | Baseline | aldex2 | 4 |
| # | sex | 4 | 14 | 0 | 0.222222222 | 1 | # | 20 | Baseline | aldex2 | 4 |
| # | time | 5 | 63 | 0 | 0.104166667 | 1 | # | 20 | Baseline | aldex2 | 4 |
| # | interaction | 0 | 0 | 4 | 0 | 0 | # | 20 | Baseline | aldex2 | 4 |
| # | disease | 7 | 0 | 0 | 1 | 1 | # | 20 | Small | paralle | 4 |
| # | sex | 4 | 1 | 0 | 0.8 | 1 | # | 20 | Small | paralle | 4 |
| # | time | 5 | 0 | 0 | 1 | 1 | # | 20 | Small | paralle | 4 |
| # | interaction | 3 | 0 | 1 | 1 | 0.75 | # | 20 | Small | paralle | 4 |
| # | disease | 7 | 4 | 0 | 0.636363636 | 1 | # | 20 | Small | ancom | 4 |
| # | sex | 4 | 0 | 0 | 1 | 1 | # | 20 | Small | ancom | 4 |
| # | time | 5 | 4 | 0 | 0.555555556 | 1 | # | 20 | Small | ancom | 4 |
| # | interaction | 0 | 0 | 4 | 0 | 0 | # | 20 | Small | ancom | 4 |
| # | disease | 7 | 12 | 0 | 0.368421053 | 1 | # | 20 | Small | aldex2 | 4 |
| # | sex | 4 | 0 | 0 | 1 | 1 | # | 20 | Small | aldex2 | 4 |
| # | time | 5 | 5 | 0 | 0.5 | 1 | # | 20 | Small | aldex2 | 4 |
| # | interaction | 0 | 0 | 4 | 0 | 0 | # | 20 | Small | aldex2 | 4 |
| # | disease | 4 | 10 | 0 | 0.285714286 | 1 | # | 10 | Baseline | paralle | 5 |
| # | sex | 2 | 5 | 0 | 0.285714286 | 1 | # | 10 | Baseline | paralle | 5 |
| # | time | 2 | 7 | 0 | 0.222222222 | 1 | # | 10 | Baseline | paralle | 5 |
| # | interaction | 2 | 6 | 0 | 0.25 | 1 | # | 10 | Baseline | paralle | 5 |
| # | disease | 4 | 1 | 0 | 0.8 | 1 | # | 10 | Baseline | ancom | 5 |
| # | sex | 2 | 0 | 0 | 1 | 1 | # | 10 | Baseline | ancom | 5 |
| # | time | 2 | 0 | 0 | 1 | 1 | # | 10 | Baseline | ancom | 5 |
| # | interaction | 0 | 0 | 2 | 0 | 0 | # | 10 | Baseline | ancom | 5 |
| # | disease | 4 | 9 | 0 | 0.307692308 | 1 | # | 10 | Baseline | aldex2 | 5 |
| # | sex | 2 | 7 | 0 | 0.222222222 | 1 | # | 10 | Baseline | aldex2 | 5 |
| # | time | 2 | 4 | 0 | 0.333333333 | 1 | # | 10 | Baseline | aldex2 | 5 |
| # | interaction | 0 | 0 | 2 | 0 | 0 | # | 10 | Baseline | aldex2 | 5 |
| # | disease | 4 | 0 | 0 | 1 | 1 | # | 10 | Small | paralle | 5 |
| # | sex | 2 | 0 | 0 | 1 | 1 | # | 10 | Small | paralle | 5 |
| # | time | 2 | 0 | 0 | 1 | 1 | # | 10 | Small | paralle | 5 |
| # | interaction | 2 | 0 | 0 | 1 | 1 | # | 10 | Small | paralle | 5 |
| # | disease | 4 | 0 | 0 | 1 | 1 | # | 10 | Small | ancom | 5 |
| # | sex | 2 | 0 | 0 | 1 | 1 | # | 10 | Small | ancom | 5 |
| # | time | 2 | 0 | 0 | 1 | 1 | # | 10 | Small | ancom | 5 |
| # | interaction | 0 | 0 | 2 | 0 | 0 | # | 10 | Small | ancom | 5 |
| # | disease | 4 | 0 | 0 | 1 | 1 | # | 10 | Small | aldex2 | 5 |
| # | sex | 2 | 0 | 0 | 1 | 1 | # | 10 | Small | aldex2 | 5 |
| # | time | 2 | 1 | 0 | 0.666666667 | 1 | # | 10 | Small | aldex2 | 5 |
| # | interaction | 0 | 0 | 2 | 0 | 0 | # | 10 | Small | aldex2 | 5 |
| # | disease | 4 | 6 | 0 | 0.4 | 1 | # | 10 | Baseline | paralle | 5 |
| # | sex | 2 | 1 | 0 | 0.666666667 | 1 | # | 10 | Baseline | paralle | 5 |
| # | time | 2 | 9 | 0 | 0.181818182 | 1 | # | 10 | Baseline | paralle | 5 |
| # | interaction | 2 | 5 | 0 | 0.285714286 | 1 | # | 10 | Baseline | paralle | 5 |
| # | disease | 4 | 1 | 0 | 0.8 | 1 | # | 10 | Baseline | ancom | 5 |
| # | sex | 2 | 0 | 0 | 1 | 1 | # | 10 | Baseline | ancom | 5 |
| # | time | 2 | 1 | 0 | 0.666666667 | 1 | # | 10 | Baseline | ancom | 5 |
| # | interaction | 0 | 0 | 2 | 0 | 0 | # | 10 | Baseline | ancom | 5 |
| # | disease | 4 | 13 | 0 | 0.235294118 | 1 | # | 10 | Baseline | aldex2 | 5 |
| # | sex | 2 | 3 | 0 | 0.4 | 1 | # | 10 | Baseline | aldex2 | 5 |
| # | time | 2 | 5 | 0 | 0.285714286 | 1 | # | 10 | Baseline | aldex2 | 5 |
| # | interaction | 0 | 0 | 2 | 0 | 0 | # | 10 | Baseline | aldex2 | 5 |
| # | disease | 4 | 0 | 0 | 1 | 1 | # | 10 | Small | paralle | 5 |
| # | sex | 2 | 0 | 0 | 1 | 1 | # | 10 | Small | paralle | 5 |
| # | time | 2 | 0 | 0 | 1 | 1 | # | 10 | Small | paralle | 5 |
| # | interaction | 2 | 0 | 0 | 1 | 1 | # | 10 | Small | paralle | 5 |
| # | disease | 4 | 1 | 0 | 0.8 | 1 | # | 10 | Small | ancom | 5 |
| # | sex | 2 | 0 | 0 | 1 | 1 | # | 10 | Small | ancom | 5 |
| # | time | 2 | 1 | 0 | 0.666666667 | 1 | # | 10 | Small | ancom | 5 |
| # | interaction | 0 | 0 | 2 | 0 | 0 | # | 10 | Small | ancom | 5 |
| # | disease | 4 | 1 | 0 | 0.8 | 1 | # | 10 | Small | aldex2 | 5 |
| # | sex | 2 | 0 | 0 | 1 | 1 | # | 10 | Small | aldex2 | 5 |
| # | time | 2 | 1 | 0 | 0.666666667 | 1 | # | 10 | Small | aldex2 | 5 |
| # | interaction | 0 | 0 | 2 | 0 | 0 | # | 10 | Small | aldex2 | 5 |
| # | disease | 4 | 6 | 0 | 0.4 | 1 | # | 10 | Baseline | paralle | 5 |
| # | sex | 2 | 1 | 0 | 0.666666667 | 1 | # | 10 | Baseline | paralle | 5 |
| # | time | 2 | 3 | 0 | 0.4 | 1 | # | 10 | Baseline | paralle | 5 |
| # | interaction | 2 | 4 | 0 | 0.333333333 | 1 | # | 10 | Baseline | paralle | 5 |
| # | disease | 4 | 2 | 0 | 0.666666667 | 1 | # | 10 | Baseline | ancom | 5 |
| # | sex | 2 | 0 | 0 | 1 | 1 | # | 10 | Baseline | ancom | 5 |
| # | time | 2 | 1 | 0 | 0.666666667 | 1 | # | 10 | Baseline | ancom | 5 |
| # | interaction | 0 | 0 | 2 | 0 | 0 | # | 10 | Baseline | ancom | 5 |
| # | disease | 4 | 21 | 0 | 0.16 | 1 | # | 10 | Baseline | aldex2 | 5 |
| # | sex | 2 | 4 | 0 | 0.333333333 | 1 | # | 10 | Baseline | aldex2 | 5 |
| # | time | 2 | 10 | 0 | 0.166666667 | 1 | # | 10 | Baseline | aldex2 | 5 |
| # | interaction | 0 | 0 | 2 | 0 | 0 | # | 10 | Baseline | aldex2 | 5 |
| # | disease | 4 | 0 | 0 | 1 | 1 | # | 10 | Small | paralle | 5 |
| # | sex | 2 | 0 | 0 | 1 | 1 | # | 10 | Small | paralle | 5 |
| # | time | 2 | 0 | 0 | 1 | 1 | # | 10 | Small | paralle | 5 |
| # | interaction | 2 | 1 | 0 | 0.666666667 | 1 | # | 10 | Small | paralle | 5 |
| # | disease | 4 | 2 | 0 | 0.666666667 | 1 | # | 10 | Small | ancom | 5 |
| # | sex | 2 | 0 | 0 | 1 | 1 | # | 10 | Small | ancom | 5 |
| # | time | 2 | 1 | 0 | 0.666666667 | 1 | # | 10 | Small | ancom | 5 |
| # | interaction | 0 | 0 | 2 | 0 | 0 | # | 10 | Small | ancom | 5 |
| # | disease | 4 | 2 | 0 | 0.666666667 | 1 | # | 10 | Small | aldex2 | 5 |
| # | sex | 2 | 0 | 0 | 1 | 1 | # | 10 | Small | aldex2 | 5 |
| # | time | 2 | 1 | 0 | 0.666666667 | 1 | # | 10 | Small | aldex2 | 5 |
| # | interaction | 0 | 0 | 2 | 0 | 0 | # | 10 | Small | aldex2 | 5 |
| # | disease | 4 | 7 | 0 | 0.363636364 | 1 | # | 10 | Baseline | paralle | 5 |
| # | sex | 2 | 1 | 0 | 0.666666667 | 1 | # | 10 | Baseline | paralle | 5 |
| # | time | 2 | 5 | 0 | 0.285714286 | 1 | # | 10 | Baseline | paralle | 5 |
| # | interaction | 2 | 6 | 0 | 0.25 | 1 | # | 10 | Baseline | paralle | 5 |
| # | disease | 4 | 2 | 0 | 0.666666667 | 1 | # | 10 | Baseline | ancom | 5 |
| # | sex | 2 | 0 | 0 | 1 | 1 | # | 10 | Baseline | ancom | 5 |
| # | time | 2 | 1 | 0 | 0.666666667 | 1 | # | 10 | Baseline | ancom | 5 |
| # | interaction | 0 | 0 | 2 | 0 | 0 | # | 10 | Baseline | ancom | 5 |
| # | disease | 4 | 24 | 0 | 0.142857143 | 1 | # | 10 | Baseline | aldex2 | 5 |
| # | sex | 2 | 8 | 0 | 0.2 | 1 | # | 10 | Baseline | aldex2 | 5 |
| # | time | 2 | 11 | 0 | 0.153846154 | 1 | # | 10 | Baseline | aldex2 | 5 |
| # | interaction | 0 | 0 | 2 | 0 | 0 | # | 10 | Baseline | aldex2 | 5 |
| # | disease | 4 | 0 | 0 | 1 | 1 | # | 10 | Small | paralle | 5 |
| # | sex | 2 | 0 | 0 | 1 | 1 | # | 10 | Small | paralle | 5 |
| # | time | 2 | 0 | 0 | 1 | 1 | # | 10 | Small | paralle | 5 |
| # | interaction | 2 | 0 | 0 | 1 | 1 | # | 10 | Small | paralle | 5 |
| # | disease | 4 | 2 | 0 | 0.666666667 | 1 | # | 10 | Small | ancom | 5 |
| # | sex | 2 | 0 | 0 | 1 | 1 | # | 10 | Small | ancom | 5 |
| # | time | 2 | 1 | 0 | 0.666666667 | 1 | # | 10 | Small | ancom | 5 |
| # | interaction | 0 | 0 | 2 | 0 | 0 | # | 10 | Small | ancom | 5 |
| # | disease | 4 | 3 | 0 | 0.571428571 | 1 | # | 10 | Small | aldex2 | 5 |
| # | sex | 2 | 0 | 0 | 1 | 1 | # | 10 | Small | aldex2 | 5 |
| # | time | 2 | 1 | 0 | 0.666666667 | 1 | # | 10 | Small | aldex2 | 5 |
| # | interaction | 0 | 0 | 2 | 0 | 0 | # | 10 | Small | aldex2 | 5 |
| # | disease | 7 | 12 | 0 | 0.368421053 | 1 | # | 20 | Baseline | paralle | 5 |
| # | sex | 4 | 8 | 0 | 0.333333333 | 1 | # | 20 | Baseline | paralle | 5 |
| # | time | 5 | 9 | 0 | 0.357142857 | 1 | # | 20 | Baseline | paralle | 5 |
| # | interaction | 4 | 6 | 0 | 0.4 | 1 | # | 20 | Baseline | paralle | 5 |
| # | disease | 7 | 0 | 0 | 1 | 1 | # | 20 | Baseline | ancom | 5 |
| # | sex | 4 | 0 | 0 | 1 | 1 | # | 20 | Baseline | ancom | 5 |
| # | time | 5 | 0 | 0 | 1 | 1 | # | 20 | Baseline | ancom | 5 |
| # | interaction | 0 | 0 | 4 | 0 | 0 | # | 20 | Baseline | ancom | 5 |
| # | disease | 7 | 21 | 0 | 0.25 | 1 | # | 20 | Baseline | aldex2 | 5 |
| # | sex | 4 | 8 | 0 | 0.333333333 | 1 | # | 20 | Baseline | aldex2 | 5 |
| # | time | 5 | 10 | 0 | 0.333333333 | 1 | # | 20 | Baseline | aldex2 | 5 |
| # | interaction | 0 | 0 | 4 | 0 | 0 | # | 20 | Baseline | aldex2 | 5 |
| # | disease | 7 | 1 | 0 | 0.875 | 1 | # | 20 | Small | paralle | 5 |
| # | sex | 4 | 0 | 0 | 1 | 1 | # | 20 | Small | paralle | 5 |
| # | time | 5 | 0 | 0 | 1 | 1 | # | 20 | Small | paralle | 5 |
| # | interaction | 1 | 0 | 3 | 1 | 0.25 | # | 20 | Small | paralle | 5 |
| # | disease | 7 | 0 | 0 | 1 | 1 | # | 20 | Small | ancom | 5 |
| # | sex | 4 | 0 | 0 | 1 | 1 | # | 20 | Small | ancom | 5 |
| # | time | 5 | 1 | 0 | 0.833333333 | 1 | # | 20 | Small | ancom | 5 |
| # | interaction | 0 | 0 | 4 | 0 | 0 | # | 20 | Small | ancom | 5 |
| # | disease | 7 | 1 | 0 | 0.875 | 1 | # | 20 | Small | aldex2 | 5 |
| # | sex | 4 | 0 | 0 | 1 | 1 | # | 20 | Small | aldex2 | 5 |
| # | time | 5 | 0 | 0 | 1 | 1 | # | 20 | Small | aldex2 | 5 |
| # | interaction | 0 | 0 | 4 | 0 | 0 | # | 20 | Small | aldex2 | 5 |
| # | disease | 7 | 10 | 0 | 0.411764706 | 1 | # | 20 | Baseline | paralle | 5 |
| # | sex | 4 | 13 | 0 | 0.235294118 | 1 | # | 20 | Baseline | paralle | 5 |
| # | time | 5 | 3 | 0 | 0.625 | 1 | # | 20 | Baseline | paralle | 5 |
| # | interaction | 4 | 4 | 0 | 0.5 | 1 | # | 20 | Baseline | paralle | 5 |
| # | disease | 7 | 1 | 0 | 0.875 | 1 | # | 20 | Baseline | ancom | 5 |
| # | sex | 4 | 0 | 0 | 1 | 1 | # | 20 | Baseline | ancom | 5 |
| # | time | 5 | 2 | 0 | 0.714285714 | 1 | # | 20 | Baseline | ancom | 5 |
| # | interaction | 0 | 0 | 4 | 0 | 0 | # | 20 | Baseline | ancom | 5 |
| # | disease | 7 | 31 | 0 | 0.184210526 | 1 | # | 20 | Baseline | aldex2 | 5 |
| # | sex | 4 | 10 | 0 | 0.285714286 | 1 | # | 20 | Baseline | aldex2 | 5 |
| # | time | 5 | 20 | 0 | 0.2 | 1 | # | 20 | Baseline | aldex2 | 5 |
| # | interaction | 0 | 0 | 4 | 0 | 0 | # | 20 | Baseline | aldex2 | 5 |
| # | disease | 7 | 0 | 0 | 1 | 1 | # | 20 | Small | paralle | 5 |
| # | sex | 4 | 0 | 0 | 1 | 1 | # | 20 | Small | paralle | 5 |
| # | time | 5 | 0 | 0 | 1 | 1 | # | 20 | Small | paralle | 5 |
| # | interaction | 2 | 0 | 2 | 1 | 0.5 | # | 20 | Small | paralle | 5 |
| # | disease | 7 | 0 | 0 | 1 | 1 | # | 20 | Small | ancom | 5 |
| # | sex | 4 | 0 | 0 | 1 | 1 | # | 20 | Small | ancom | 5 |
| # | time | 5 | 1 | 0 | 0.833333333 | 1 | # | 20 | Small | ancom | 5 |
| # | interaction | 0 | 0 | 4 | 0 | 0 | # | 20 | Small | ancom | 5 |
| # | disease | 7 | 3 | 0 | 0.7 | 1 | # | 20 | Small | aldex2 | 5 |
| # | sex | 4 | 0 | 0 | 1 | 1 | # | 20 | Small | aldex2 | 5 |
| # | time | 5 | 2 | 0 | 0.714285714 | 1 | # | 20 | Small | aldex2 | 5 |
| # | interaction | 0 | 0 | 4 | 0 | 0 | # | 20 | Small | aldex2 | 5 |
| # | disease | 7 | 11 | 0 | 0.388888889 | 1 | # | 20 | Baseline | paralle | 5 |
| # | sex | 4 | 5 | 0 | 0.444444444 | 1 | # | 20 | Baseline | paralle | 5 |
| # | time | 5 | 5 | 0 | 0.5 | 1 | # | 20 | Baseline | paralle | 5 |
| # | interaction | 4 | 6 | 0 | 0.4 | 1 | # | 20 | Baseline | paralle | 5 |
| # | disease | 7 | 4 | 0 | 0.636363636 | 1 | # | 20 | Baseline | ancom | 5 |
| # | sex | 4 | 0 | 0 | 1 | 1 | # | 20 | Baseline | ancom | 5 |
| # | time | 5 | 4 | 0 | 0.555555556 | 1 | # | 20 | Baseline | ancom | 5 |
| # | interaction | 0 | 0 | 4 | 0 | 0 | # | 20 | Baseline | ancom | 5 |
| # | disease | 7 | 52 | 0 | 0.118644068 | 1 | # | 20 | Baseline | aldex2 | 5 |
| # | sex | 4 | 16 | 0 | 0.2 | 1 | # | 20 | Baseline | aldex2 | 5 |
| # | time | 5 | 29 | 0 | 0.147058824 | 1 | # | 20 | Baseline | aldex2 | 5 |
| # | interaction | 0 | 0 | 4 | 0 | 0 | # | 20 | Baseline | aldex2 | 5 |
| # | disease | 7 | 1 | 0 | 0.875 | 1 | # | 20 | Small | paralle | 5 |
| # | sex | 4 | 0 | 0 | 1 | 1 | # | 20 | Small | paralle | 5 |
| # | time | 5 | 0 | 0 | 1 | 1 | # | 20 | Small | paralle | 5 |
| # | interaction | 3 | 0 | 1 | 1 | 0.75 | # | 20 | Small | paralle | 5 |
| # | disease | 7 | 4 | 0 | 0.636363636 | 1 | # | 20 | Small | ancom | 5 |
| # | sex | 4 | 0 | 0 | 1 | 1 | # | 20 | Small | ancom | 5 |
| # | time | 5 | 2 | 0 | 0.714285714 | 1 | # | 20 | Small | ancom | 5 |
| # | interaction | 0 | 0 | 4 | 0 | 0 | # | 20 | Small | ancom | 5 |
| # | disease | 7 | 6 | 0 | 0.538461538 | 1 | # | 20 | Small | aldex2 | 5 |
| # | sex | 4 | 0 | 0 | 1 | 1 | # | 20 | Small | aldex2 | 5 |
| # | time | 5 | 1 | 0 | 0.833333333 | 1 | # | 20 | Small | aldex2 | 5 |
| # | interaction | 0 | 0 | 4 | 0 | 0 | # | 20 | Small | aldex2 | 5 |
| # | disease | 7 | 11 | 0 | 0.388888889 | 1 | # | 20 | Baseline | paralle | 5 |
| # | sex | 4 | 5 | 0 | 0.444444444 | 1 | # | 20 | Baseline | paralle | 5 |
| # | time | 5 | 3 | 0 | 0.625 | 1 | # | 20 | Baseline | paralle | 5 |
| # | interaction | 4 | 6 | 0 | 0.4 | 1 | # | 20 | Baseline | paralle | 5 |
| # | disease | 7 | 4 | 0 | 0.636363636 | 1 | # | 20 | Baseline | ancom | 5 |
| # | sex | 4 | 0 | 0 | 1 | 1 | # | 20 | Baseline | ancom | 5 |
| # | time | 5 | 4 | 0 | 0.555555556 | 1 | # | 20 | Baseline | ancom | 5 |
| # | interaction | 0 | 0 | 4 | 0 | 0 | # | 20 | Baseline | ancom | 5 |
| # | disease | 7 | 82 | 0 | 0.101449275 | 1 | # | 20 | Baseline | aldex2 | 5 |
| # | sex | 4 | 24 | 0 | 0.142857143 | 1 | # | 20 | Baseline | aldex2 | 5 |
| # | time | 5 | 60 | 0 | 0.111111111 | 1 | # | 20 | Baseline | aldex2 | 5 |
| # | interaction | 0 | 0 | 4 | 0 | 0 | # | 20 | Baseline | aldex2 | 5 |
| # | disease | 7 | 0 | 0 | 1 | 1 | # | 20 | Small | paralle | 5 |
| # | sex | 4 | 0 | 0 | 1 | 1 | # | 20 | Small | paralle | 5 |
| # | time | 5 | 0 | 0 | 1 | 1 | # | 20 | Small | paralle | 5 |
| # | interaction | 4 | 0 | 0 | 1 | 1 | # | 20 | Small | paralle | 5 |
| # | disease | 7 | 4 | 0 | 0.636363636 | 1 | # | 20 | Small | ancom | 5 |
| # | sex | 4 | 0 | 0 | 1 | 1 | # | 20 | Small | ancom | 5 |
| # | time | 5 | 4 | 0 | 0.555555556 | 1 | # | 20 | Small | ancom | 5 |
| # | interaction | 0 | 0 | 4 | 0 | 0 | # | 20 | Small | ancom | 5 |
| # | disease | 7 | 12 | 0 | 0.368421053 | 1 | # | 20 | Small | aldex2 | 5 |
| # | sex | 4 | 1 | 0 | 0.8 | 1 | # | 20 | Small | aldex2 | 5 |
| # | time | 5 | 4 | 0 | 0.555555556 | 1 | # | 20 | Small | aldex2 | 5 |
| # | interaction | 0 | 0 | 4 | 0 | 0 | # | 20 | Small | aldex2 | 5 |
| # | disease | 4 | 8 | 0 | 0.333333333 | 1 | # | 10 | Baseline | paralle | 6 |
| # | sex | 2 | 6 | 0 | 0.25 | 1 | # | 10 | Baseline | paralle | 6 |
| # | time | 2 | 7 | 0 | 0.222222222 | 1 | # | 10 | Baseline | paralle | 6 |
| # | interaction | 2 | 12 | 0 | 0.142857143 | 1 | # | 10 | Baseline | paralle | 6 |
| # | disease | 4 | 1 | 0 | 0.8 | 1 | # | 10 | Baseline | ancom | 6 |
| # | sex | 2 | 0 | 0 | 1 | 1 | # | 10 | Baseline | ancom | 6 |
| # | time | 2 | 2 | 0 | 0.5 | 1 | # | 10 | Baseline | ancom | 6 |
| # | interaction | 0 | 0 | 2 | 0 | 0 | # | 10 | Baseline | ancom | 6 |
| # | disease | 4 | 16 | 0 | 0.2 | 1 | # | 10 | Baseline | aldex2 | 6 |
| # | sex | 2 | 4 | 0 | 0.333333333 | 1 | # | 10 | Baseline | aldex2 | 6 |
| # | time | 2 | 8 | 0 | 0.2 | 1 | # | 10 | Baseline | aldex2 | 6 |
| # | interaction | 0 | 0 | 2 | 0 | 0 | # | 10 | Baseline | aldex2 | 6 |
| # | disease | 4 | 0 | 0 | 1 | 1 | # | 10 | Small | paralle | 6 |
| # | sex | 2 | 0 | 0 | 1 | 1 | # | 10 | Small | paralle | 6 |
| # | time | 2 | 0 | 0 | 1 | 1 | # | 10 | Small | paralle | 6 |
| # | interaction | 0 | 0 | 2 | 0 | 0 | # | 10 | Small | paralle | 6 |
| # | disease | 4 | 1 | 0 | 0.8 | 1 | # | 10 | Small | ancom | 6 |
| # | sex | 2 | 0 | 0 | 1 | 1 | # | 10 | Small | ancom | 6 |
| # | time | 2 | 1 | 0 | 0.666666667 | 1 | # | 10 | Small | ancom | 6 |
| # | interaction | 0 | 0 | 2 | 0 | 0 | # | 10 | Small | ancom | 6 |
| # | disease | 4 | 1 | 0 | 0.8 | 1 | # | 10 | Small | aldex2 | 6 |
| # | sex | 2 | 0 | 0 | 1 | 1 | # | 10 | Small | aldex2 | 6 |
| # | time | 2 | 2 | 0 | 0.5 | 1 | # | 10 | Small | aldex2 | 6 |
| # | interaction | 0 | 0 | 2 | 0 | 0 | # | 10 | Small | aldex2 | 6 |
| # | disease | 4 | 8 | 0 | 0.333333333 | 1 | # | 10 | Baseline | paralle | 6 |
| # | sex | 2 | 3 | 0 | 0.4 | 1 | # | 10 | Baseline | paralle | 6 |
| # | time | 2 | 2 | 0 | 0.5 | 1 | # | 10 | Baseline | paralle | 6 |
| # | interaction | 2 | 9 | 0 | 0.181818182 | 1 | # | 10 | Baseline | paralle | 6 |
| # | disease | 4 | 2 | 0 | 0.666666667 | 1 | # | 10 | Baseline | ancom | 6 |
| # | sex | 2 | 0 | 0 | 1 | 1 | # | 10 | Baseline | ancom | 6 |
| # | time | 2 | 2 | 0 | 0.5 | 1 | # | 10 | Baseline | ancom | 6 |
| # | interaction | 0 | 0 | 2 | 0 | 0 | # | 10 | Baseline | ancom | 6 |
| # | disease | 4 | 16 | 0 | 0.2 | 1 | # | 10 | Baseline | aldex2 | 6 |
| # | sex | 2 | 5 | 0 | 0.285714286 | 1 | # | 10 | Baseline | aldex2 | 6 |
| # | time | 2 | 12 | 0 | 0.142857143 | 1 | # | 10 | Baseline | aldex2 | 6 |
| # | interaction | 0 | 0 | 2 | 0 | 0 | # | 10 | Baseline | aldex2 | 6 |
| # | disease | 4 | 0 | 0 | 1 | 1 | # | 10 | Small | paralle | 6 |
| # | sex | 2 | 0 | 0 | 1 | 1 | # | 10 | Small | paralle | 6 |
| # | time | 2 | 0 | 0 | 1 | 1 | # | 10 | Small | paralle | 6 |
| # | interaction | 2 | 0 | 0 | 1 | 1 | # | 10 | Small | paralle | 6 |
| # | disease | 4 | 2 | 0 | 0.666666667 | 1 | # | 10 | Small | ancom | 6 |
| # | sex | 2 | 0 | 0 | 1 | 1 | # | 10 | Small | ancom | 6 |
| # | time | 2 | 2 | 0 | 0.5 | 1 | # | 10 | Small | ancom | 6 |
| # | interaction | 0 | 0 | 2 | 0 | 0 | # | 10 | Small | ancom | 6 |
| # | disease | 4 | 5 | 0 | 0.444444444 | 1 | # | 10 | Small | aldex2 | 6 |
| # | sex | 2 | 0 | 0 | 1 | 1 | # | 10 | Small | aldex2 | 6 |
| # | time | 2 | 2 | 0 | 0.5 | 1 | # | 10 | Small | aldex2 | 6 |
| # | interaction | 0 | 0 | 2 | 0 | 0 | # | 10 | Small | aldex2 | 6 |

| | | | | | | | | | | | |
|---|---|---|---|---|---|---|---|---|---|---|---|
| # | disease | 4 | 7 | 0 | 0.363636364 | 1 | # | 10 | Baseline | paralle | 6 |
| # | sex | 2 | 5 | 0 | 0.285714286 | 1 | # | 10 | Baseline | paralle | 6 |
| # | time | 2 | 2 | 0 | 0.5 | 1 | # | 10 | Baseline | paralle | 6 |
| # | interaction | 2 | 5 | 0 | 0.285714286 | 1 | # | 10 | Baseline | paralle | 6 |
| # | disease | 4 | 2 | 0 | 0.666666667 | 1 | # | 10 | Baseline | ancom | 6 |
| # | sex | 2 | 0 | 0 | 1 | 1 | # | 10 | Baseline | ancom | 6 |
| # | time | 2 | 2 | 0 | 0.5 | 1 | # | 10 | Baseline | ancom | 6 |
| # | interaction | 0 | 0 | 2 | 0 | 0 | # | 10 | Baseline | ancom | 6 |
| # | disease | 4 | 15 | 0 | 0.210526316 | 1 | # | 10 | Baseline | aldex2 | 6 |
| # | sex | 2 | 3 | 0 | 0.4 | 1 | # | 10 | Baseline | aldex2 | 6 |
| # | time | 2 | 13 | 0 | 0.133333333 | 1 | # | 10 | Baseline | aldex2 | 6 |
| # | interaction | 0 | 0 | 2 | 0 | 0 | # | 10 | Baseline | aldex2 | 6 |
| # | disease | 4 | 0 | 0 | 1 | 1 | # | 10 | Small | paralle | 6 |
| # | sex | 2 | 0 | 0 | 1 | 1 | # | 10 | Small | paralle | 6 |
| # | time | 2 | 0 | 0 | 1 | 1 | # | 10 | Small | paralle | 6 |
| # | interaction | 2 | 0 | 0 | 1 | 1 | # | 10 | Small | paralle | 6 |
| # | disease | 4 | 2 | 0 | 0.666666667 | 1 | # | 10 | Small | ancom | 6 |
| # | sex | 2 | 0 | 0 | 1 | 1 | # | 10 | Small | ancom | 6 |
| # | time | 2 | 2 | 0 | 0.5 | 1 | # | 10 | Small | ancom | 6 |
| # | interaction | 0 | 0 | 2 | 0 | 0 | # | 10 | Small | ancom | 6 |
| # | disease | 4 | 3 | 0 | 0.571428571 | 1 | # | 10 | Small | aldex2 | 6 |
| # | sex | 2 | 0 | 0 | 1 | 1 | # | 10 | Small | aldex2 | 6 |
| # | time | 2 | 2 | 0 | 0.5 | 1 | # | 10 | Small | aldex2 | 6 |
| # | interaction | 0 | 0 | 2 | 0 | 0 | # | 10 | Small | aldex2 | 6 |
| # | disease | 4 | 6 | 0 | 0.4 | 1 | # | 10 | Baseline | paralle | 6 |
| # | sex | 2 | 2 | 0 | 0.5 | 1 | # | 10 | Baseline | paralle | 6 |
| # | time | 2 | 1 | 0 | 0.666666667 | 1 | # | 10 | Baseline | paralle | 6 |
| # | interaction | 2 | 3 | 0 | 0.4 | 1 | # | 10 | Baseline | paralle | 6 |
| # | disease | 4 | 2 | 0 | 0.666666667 | 1 | # | 10 | Baseline | ancom | 6 |
| # | sex | 2 | 0 | 0 | 1 | 1 | # | 10 | Baseline | ancom | 6 |
| # | time | 2 | 2 | 0 | 0.5 | 1 | # | 10 | Baseline | ancom | 6 |
| # | interaction | 0 | 0 | 2 | 0 | 0 | # | 10 | Baseline | ancom | 6 |
| # | disease | 4 | 25 | 0 | 0.137931034 | 1 | # | 10 | Baseline | aldex2 | 6 |
| # | sex | 2 | 6 | 0 | 0.25 | 1 | # | 10 | Baseline | aldex2 | 6 |
| # | time | 2 | 16 | 0 | 0.111111111 | 1 | # | 10 | Baseline | aldex2 | 6 |
| # | interaction | 0 | 0 | 2 | 0 | 0 | # | 10 | Baseline | aldex2 | 6 |
| # | disease | 4 | 0 | 0 | 1 | 1 | # | 10 | Small | paralle | 6 |
| # | sex | 2 | 0 | 0 | 1 | 1 | # | 10 | Small | paralle | 6 |
| # | time | 2 | 0 | 0 | 1 | 1 | # | 10 | Small | paralle | 6 |
| # | interaction | 2 | 0 | 0 | 1 | 1 | # | 10 | Small | paralle | 6 |
| # | disease | 4 | 2 | 0 | 0.666666667 | 1 | # | 10 | Small | ancom | 6 |
| # | sex | 2 | 0 | 0 | 1 | 1 | # | 10 | Small | ancom | 6 |
| # | time | 2 | 2 | 0 | 0.5 | 1 | # | 10 | Small | ancom | 6 |
| # | interaction | 0 | 0 | 2 | 0 | 0 | # | 10 | Small | ancom | 6 |
| # | disease | 4 | 4 | 0 | 0.5 | 1 | # | 10 | Small | aldex2 | 6 |
| # | sex | 2 | 0 | 0 | 1 | 1 | # | 10 | Small | aldex2 | 6 |
| # | time | 2 | 2 | 0 | 0.5 | 1 | # | 10 | Small | aldex2 | 6 |
| # | interaction | 0 | 0 | 2 | 0 | 0 | # | 10 | Small | aldex2 | 6 |
| # | disease | 7 | 8 | 0 | 0.466666667 | 1 | # | 20 | Baseline | paralle | 6 |
| # | sex | 4 | 6 | 0 | 0.4 | 1 | # | 20 | Baseline | paralle | 6 |
| # | time | 5 | 10 | 0 | 0.333333333 | 1 | # | 20 | Baseline | paralle | 6 |
| # | interaction | 4 | 8 | 0 | 0.333333333 | 1 | # | 20 | Baseline | paralle | 6 |
| # | disease | 7 | 1 | 0 | 0.875 | 1 | # | 20 | Baseline | ancom | 6 |
| # | sex | 4 | 0 | 0 | 1 | 1 | # | 20 | Baseline | ancom | 6 |
| # | time | 5 | 1 | 0 | 0.833333333 | 1 | # | 20 | Baseline | ancom | 6 |
| # | interaction | 0 | 0 | 4 | 0 | 0 | # | 20 | Baseline | ancom | 6 |
| # | disease | 7 | 21 | 0 | 0.25 | 1 | # | 20 | Baseline | aldex2 | 6 |
| # | sex | 4 | 6 | 0 | 0.4 | 1 | # | 20 | Baseline | aldex2 | 6 |
| # | time | 5 | 16 | 0 | 0.238095238 | 1 | # | 20 | Baseline | aldex2 | 6 |
| # | interaction | 0 | 0 | 4 | 0 | 0 | # | 20 | Baseline | aldex2 | 6 |
| # | disease | 7 | 0 | 0 | 1 | 1 | # | 20 | Small | paralle | 6 |
| # | sex | 4 | 0 | 0 | 1 | 1 | # | 20 | Small | paralle | 6 |
| # | time | 5 | 0 | 0 | 1 | 1 | # | 20 | Small | paralle | 6 |
| # | interaction | 3 | 0 | 1 | 1 | 0.75 | # | 20 | Small | paralle | 6 |
| # | disease | 7 | 1 | 0 | 0.875 | 1 | # | 20 | Small | ancom | 6 |
| # | sex | 4 | 0 | 0 | 1 | 1 | # | 20 | Small | ancom | 6 |
| # | time | 5 | 1 | 0 | 0.833333333 | 1 | # | 20 | Small | ancom | 6 |
| # | interaction | 0 | 0 | 4 | 0 | 0 | # | 20 | Small | ancom | 6 |
| # | disease | 7 | 1 | 0 | 0.875 | 1 | # | 20 | Small | aldex2 | 6 |
| # | sex | 4 | 1 | 0 | 0.8 | 1 | # | 20 | Small | aldex2 | 6 |
| # | time | 5 | 1 | 0 | 0.833333333 | 1 | # | 20 | Small | aldex2 | 6 |
| # | interaction | 0 | 0 | 4 | 0 | 0 | # | 20 | Small | aldex2 | 6 |
| # | disease | 7 | 10 | 0 | 0.411764706 | 1 | # | 20 | Baseline | paralle | 6 |
| # | sex | 4 | 4 | 0 | 0.5 | 1 | # | 20 | Baseline | paralle | 6 |
| # | time | 5 | 5 | 0 | 0.5 | 1 | # | 20 | Baseline | paralle | 6 |
| # | interaction | 4 | 8 | 0 | 0.333333333 | 1 | # | 20 | Baseline | paralle | 6 |
| # | disease | 7 | 4 | 0 | 0.636363636 | 1 | # | 20 | Baseline | ancom | 6 |
| # | sex | 4 | 0 | 0 | 1 | 1 | # | 20 | Baseline | ancom | 6 |
| # | time | 5 | 4 | 0 | 0.555555556 | 1 | # | 20 | Baseline | ancom | 6 |
| # | interaction | 0 | 0 | 4 | 0 | 0 | # | 20 | Baseline | ancom | 6 |
| # | disease | 7 | 37 | 0 | 0.159090909 | 1 | # | 20 | Baseline | aldex2 | 6 |
| # | sex | 4 | 10 | 0 | 0.285714286 | 1 | # | 20 | Baseline | aldex2 | 6 |
| # | time | 5 | 30 | 0 | 0.142857143 | 1 | # | 20 | Baseline | aldex2 | 6 |
| # | interaction | 0 | 0 | 4 | 0 | 0 | # | 20 | Baseline | aldex2 | 6 |
| # | disease | 7 | 0 | 0 | 1 | 1 | # | 20 | Small | paralle | 6 |
| # | sex | 4 | 1 | 0 | 0.8 | 1 | # | 20 | Small | paralle | 6 |
| # | time | 5 | 0 | 0 | 1 | 1 | # | 20 | Small | paralle | 6 |
| # | interaction | 3 | 0 | 1 | 1 | 0.75 | # | 20 | Small | paralle | 6 |
| # | disease | 7 | 4 | 0 | 0.636363636 | 1 | # | 20 | Small | ancom | 6 |
| # | sex | 4 | 0 | 0 | 1 | 1 | # | 20 | Small | ancom | 6 |
| # | time | 5 | 3 | 0 | 0.625 | 1 | # | 20 | Small | ancom | 6 |
| # | interaction | 0 | 0 | 4 | 0 | 0 | # | 20 | Small | ancom | 6 |
| # | disease | 7 | 3 | 0 | 0.7 | 1 | # | 20 | Small | aldex2 | 6 |
| # | sex | 4 | 0 | 0 | 1 | 1 | # | 20 | Small | aldex2 | 6 |
| # | time | 5 | 2 | 0 | 0.714285714 | 1 | # | 20 | Small | aldex2 | 6 |
| # | interaction | 0 | 0 | 4 | 0 | 0 | # | 20 | Small | aldex2 | 6 |
| # | disease | 7 | 10 | 0 | 0.411764706 | 1 | # | 20 | Baseline | paralle | 6 |
| # | sex | 4 | 8 | 0 | 0.333333333 | 1 | # | 20 | Baseline | paralle | 6 |
| # | time | 5 | 7 | 0 | 0.416666667 | 1 | # | 20 | Baseline | paralle | 6 |
| # | interaction | 4 | 3 | 0 | 0.571428571 | 1 | # | 20 | Baseline | paralle | 6 |
| # | disease | 7 | 4 | 0 | 0.636363636 | 1 | # | 20 | Baseline | ancom | 6 |
| # | sex | 4 | 0 | 0 | 1 | 1 | # | 20 | Baseline | ancom | 6 |
| # | time | 5 | 4 | 0 | 0.555555556 | 1 | # | 20 | Baseline | ancom | 6 |
| # | interaction | 0 | 0 | 4 | 0 | 0 | # | 20 | Baseline | ancom | 6 |
| # | disease | 7 | 68 | 0 | 0.127272727 | 1 | # | 20 | Baseline | aldex2 | 6 |
| # | sex | 4 | 15 | 0 | 0.210526316 | 1 | # | 20 | Baseline | aldex2 | 6 |
| # | time | 5 | 38 | 0 | 0.11627907 | 1 | # | 20 | Baseline | aldex2 | 6 |
| # | interaction | 0 | 0 | 4 | 0 | 0 | # | 20 | Baseline | aldex2 | 6 |
| # | disease | 7 | 0 | 0 | 1 | 1 | # | 20 | Small | paralle | 6 |
| # | sex | 4 | 0 | 0 | 1 | 1 | # | 20 | Small | paralle | 6 |
| # | time | 5 | 0 | 0 | 1 | 1 | # | 20 | Small | paralle | 6 |
| # | interaction | 3 | 0 | 1 | 1 | 0.75 | # | 20 | Small | paralle | 6 |
| # | disease | 7 | 4 | 0 | 0.636363636 | 1 | # | 20 | Small | ancom | 6 |
| # | sex | 4 | 0 | 0 | 1 | 1 | # | 20 | Small | ancom | 6 |
| # | time | 5 | 4 | 0 | 0.555555556 | 1 | # | 20 | Small | ancom | 6 |
| # | interaction | 0 | 0 | 4 | 0 | 0 | # | 20 | Small | ancom | 6 |
| # | disease | 7 | 6 | 0 | 0.538461538 | 1 | # | 20 | Small | aldex2 | 6 |
| # | sex | 4 | 1 | 0 | 0.8 | 1 | # | 20 | Small | aldex2 | 6 |
| # | time | 5 | 5 | 0 | 0.5 | 1 | # | 20 | Small | aldex2 | 6 |
| # | interaction | 0 | 0 | 4 | 0 | 0 | # | 20 | Small | aldex2 | 6 |
| # | disease | 7 | 11 | 0 | 0.388888889 | 1 | # | 20 | Baseline | paralle | 6 |
| # | sex | 4 | 6 | 0 | 0.4 | 1 | # | 20 | Baseline | paralle | 6 |
| # | time | 5 | 6 | 0 | 0.454545455 | 1 | # | 20 | Baseline | paralle | 6 |
| # | interaction | 4 | 4 | 0 | 0.5 | 1 | # | 20 | Baseline | paralle | 6 |
| # | disease | 7 | 4 | 0 | 0.636363636 | 1 | # | 20 | Baseline | ancom | 6 |
| # | sex | 4 | 0 | 0 | 1 | 1 | # | 20 | Baseline | ancom | 6 |
| # | time | 5 | 4 | 0 | 0.555555556 | 1 | # | 20 | Baseline | ancom | 6 |
| # | interaction | 0 | 0 | 4 | 0 | 0 | # | 20 | Baseline | ancom | 6 |
| # | disease | 7 | 58 | 0 | 0.107692308 | 1 | # | 20 | Baseline | aldex2 | 6 |
| # | sex | 4 | 19 | 0 | 0.173913043 | 1 | # | 20 | Baseline | aldex2 | 6 |
| # | time | 5 | 67 | 0 | 0.096153846 | 1 | # | 20 | Baseline | aldex2 | 6 |
| # | interaction | 0 | 0 | 4 | 0 | 0 | # | 20 | Baseline | aldex2 | 6 |
| # | disease | 7 | 0 | 0 | 1 | 1 | # | 20 | Small | paralle | 6 |
| # | sex | 4 | 0 | 0 | 1 | 1 | # | 20 | Small | paralle | 6 |
| # | time | 5 | 0 | 0 | 1 | 1 | # | 20 | Small | paralle | 6 |
| # | interaction | 4 | 0 | 0 | 1 | 1 | # | 20 | Small | paralle | 6 |
| # | disease | 7 | 4 | 0 | 0.636363636 | 1 | # | 20 | Small | ancom | 6 |
| # | sex | 4 | 0 | 0 | 1 | 1 | # | 20 | Small | ancom | 6 |
| # | time | 5 | 4 | 0 | 0.555555556 | 1 | # | 20 | Small | ancom | 6 |
| # | interaction | 0 | 0 | 4 | 0 | 0 | # | 20 | Small | ancom | 6 |
| # | disease | 7 | 9 | 0 | 0.4375 | 1 | # | 20 | Small | aldex2 | 6 |
| # | sex | 4 | 1 | 0 | 0.8 | 1 | # | 20 | Small | aldex2 | 6 |
| # | time | 5 | 6 | 0 | 0.454545455 | 1 | # | 20 | Small | aldex2 | 6 |
| # | interaction | 0 | 0 | 4 | 0 | 0 | # | 20 | Small | aldex2 | 6 |
| # | disease | 4 | 6 | 0 | 0.4 | 1 | # | 10 | Baseline | paralle | 7 |
| # | sex | 2 | 7 | 0 | 0.222222222 | 1 | # | 10 | Baseline | paralle | 7 |
| # | time | 2 | 12 | 0 | 0.142857143 | 1 | # | 10 | Baseline | paralle | 7 |
| # | interaction | 2 | 6 | 0 | 0.25 | 1 | # | 10 | Baseline | paralle | 7 |
| # | disease | 4 | 1 | 0 | 0.8 | 1 | # | 10 | Baseline | ancom | 7 |
| # | sex | 2 | 0 | 0 | 1 | 1 | # | 10 | Baseline | ancom | 7 |
| # | time | 2 | 0 | 0 | 1 | 1 | # | 10 | Baseline | ancom | 7 |
| # | interaction | 0 | 0 | 2 | 0 | 0 | # | 10 | Baseline | ancom | 7 |
| # | disease | 4 | 11 | 0 | 0.266666667 | 1 | # | 10 | Baseline | aldex2 | 7 |
| # | sex | 2 | 7 | 0 | 0.222222222 | 1 | # | 10 | Baseline | aldex2 | 7 |
| # | time | 2 | 6 | 0 | 0.25 | 1 | # | 10 | Baseline | aldex2 | 7 |
| # | interaction | 0 | 0 | 2 | 0 | 0 | # | 10 | Baseline | aldex2 | 7 |
| # | disease | 4 | 0 | 0 | 1 | 1 | # | 10 | Small | paralle | 7 |
| # | sex | 2 | 0 | 0 | 1 | 1 | # | 10 | Small | paralle | 7 |
| # | time | 2 | 0 | 0 | 1 | 1 | # | 10 | Small | paralle | 7 |
| # | interaction | 1 | 0 | 1 | 1 | 0.5 | # | 10 | Small | paralle | 7 |
| # | disease | 4 | 1 | 0 | 0.8 | 1 | # | 10 | Small | ancom | 7 |
| # | sex | 2 | 0 | 0 | 1 | 1 | # | 10 | Small | ancom | 7 |
| # | time | 2 | 0 | 0 | 1 | 1 | # | 10 | Small | ancom | 7 |
| # | interaction | 0 | 0 | 2 | 0 | 0 | # | 10 | Small | ancom | 7 |
| # | disease | 4 | 1 | 0 | 0.8 | 1 | # | 10 | Small | aldex2 | 7 |
| # | sex | 2 | 0 | 0 | 1 | 1 | # | 10 | Small | aldex2 | 7 |
| # | time | 2 | 0 | 0 | 1 | 1 | # | 10 | Small | aldex2 | 7 |
| # | interaction | 0 | 0 | 2 | 0 | 0 | # | 10 | Small | aldex2 | 7 |
| # | disease | 4 | 6 | 0 | 0.4 | 1 | # | 10 | Baseline | paralle | 7 |
| # | sex | 2 | 4 | 0 | 0.333333333 | 1 | # | 10 | Baseline | paralle | 7 |
| # | time | 2 | 3 | 0 | 0.4 | 1 | # | 10 | Baseline | paralle | 7 |
| # | interaction | 2 | 5 | 0 | 0.285714286 | 1 | # | 10 | Baseline | paralle | 7 |
| # | disease | 4 | 1 | 0 | 0.8 | 1 | # | 10 | Baseline | ancom | 7 |
| # | sex | 2 | 0 | 0 | 1 | 1 | # | 10 | Baseline | ancom | 7 |
| # | time | 2 | 2 | 0 | 0.5 | 1 | # | 10 | Baseline | ancom | 7 |
| # | interaction | 0 | 0 | 2 | 0 | 0 | # | 10 | Baseline | ancom | 7 |
| # | disease | 4 | 20 | 0 | 0.166666667 | 1 | # | 10 | Baseline | aldex2 | 7 |
| # | sex | 2 | 8 | 0 | 0.2 | 1 | # | 10 | Baseline | aldex2 | 7 |
| # | time | 2 | 9 | 0 | 0.181818182 | 1 | # | 10 | Baseline | aldex2 | 7 |
| # | interaction | 0 | 0 | 2 | 0 | 0 | # | 10 | Baseline | aldex2 | 7 |
| # | disease | 4 | 0 | 0 | 1 | 1 | # | 10 | Small | paralle | 7 |
| # | sex | 2 | 0 | 0 | 1 | 1 | # | 10 | Small | paralle | 7 |
| # | time | 2 | 0 | 0 | 1 | 1 | # | 10 | Small | paralle | 7 |
| # | interaction | 2 | 0 | 0 | 1 | 1 | # | 10 | Small | paralle | 7 |
| # | disease | 4 | 1 | 0 | 0.8 | 1 | # | 10 | Small | ancom | 7 |
| # | sex | 2 | 0 | 0 | 1 | 1 | # | 10 | Small | ancom | 7 |
| # | time | 2 | 2 | 0 | 0.5 | 1 | # | 10 | Small | ancom | 7 |
| # | interaction | 0 | 0 | 2 | 0 | 0 | # | 10 | Small | ancom | 7 |
| # | disease | 4 | 2 | 0 | 0.666666667 | 1 | # | 10 | Small | aldex2 | 7 |
| # | sex | 2 | 0 | 0 | 1 | 1 | # | 10 | Small | aldex2 | 7 |
| # | time | 2 | 2 | 0 | 0.5 | 1 | # | 10 | Small | aldex2 | 7 |
| # | interaction | 0 | 0 | 2 | 0 | 0 | # | 10 | Small | aldex2 | 7 |
| # | disease | 4 | 8 | 0 | 0.333333333 | 1 | # | 10 | Baseline | paralle | 7 |
| # | sex | 2 | 1 | 0 | 0.666666667 | 1 | # | 10 | Baseline | paralle | 7 |
| # | time | 2 | 2 | 0 | 0.5 | 1 | # | 10 | Baseline | paralle | 7 |
| # | interaction | 2 | 7 | 0 | 0.222222222 | 1 | # | 10 | Baseline | paralle | 7 |
| # | disease | 4 | 2 | 0 | 0.666666667 | 1 | # | 10 | Baseline | ancom | 7 |
| # | sex | 2 | 0 | 0 | 1 | 1 | # | 10 | Baseline | ancom | 7 |
| # | time | 2 | 2 | 0 | 0.5 | 1 | # | 10 | Baseline | ancom | 7 |
| # | interaction | 0 | 0 | 2 | 0 | 0 | # | 10 | Baseline | ancom | 7 |
| # | disease | 4 | 23 | 0 | 0.148148148 | 1 | # | 10 | Baseline | aldex2 | 7 |
| # | sex | 2 | 9 | 0 | 0.181818182 | 1 | # | 10 | Baseline | aldex2 | 7 |
| # | time | 2 | 10 | 0 | 0.166666667 | 1 | # | 10 | Baseline | aldex2 | 7 |
| # | interaction | 0 | 0 | 2 | 0 | 0 | # | 10 | Baseline | aldex2 | 7 |
| # | disease | 4 | 0 | 0 | 1 | 1 | # | 10 | Small | paralle | 7 |
| # | sex | 2 | 0 | 0 | 1 | 1 | # | 10 | Small | paralle | 7 |
| # | time | 2 | 0 | 0 | 1 | 1 | # | 10 | Small | paralle | 7 |
| # | interaction | 2 | 0 | 0 | 1 | 1 | # | 10 | Small | paralle | 7 |
| # | disease | 4 | 3 | 0 | 0.571428571 | 1 | # | 10 | Small | ancom | 7 |
| # | sex | 2 | 0 | 0 | 1 | 1 | # | 10 | Small | ancom | 7 |
| # | time | 2 | 2 | 0 | 0.5 | 1 | # | 10 | Small | ancom | 7 |
| # | interaction | 0 | 0 | 2 | 0 | 0 | # | 10 | Small | ancom | 7 |
| # | disease | 4 | 3 | 0 | 0.571428571 | 1 | # | 10 | Small | aldex2 | 7 |
| # | sex | 2 | 0 | 0 | 1 | 1 | # | 10 | Small | aldex2 | 7 |
| # | time | 2 | 2 | 0 | 0.5 | 1 | # | 10 | Small | aldex2 | 7 |
| # | interaction | 0 | 0 | 2 | 0 | 0 | # | 10 | Small | aldex2 | 7 |
| # | disease | 4 | 6 | 0 | 0.4 | 1 | # | 10 | Baseline | paralle | 7 |
| # | sex | 2 | 1 | 0 | 0.666666667 | 1 | # | 10 | Baseline | paralle | 7 |
| # | time | 2 | 2 | 0 | 0.5 | 1 | # | 10 | Baseline | paralle | 7 |
| # | interaction | 2 | 2 | 0 | 0.5 | 1 | # | 10 | Baseline | paralle | 7 |
| # | disease | 4 | 3 | 0 | 0.571428571 | 1 | # | 10 | Baseline | ancom | 7 |
| # | sex | 2 | 0 | 0 | 1 | 1 | # | 10 | Baseline | ancom | 7 |
| # | time | 2 | 2 | 0 | 0.5 | 1 | # | 10 | Baseline | ancom | 7 |
| # | interaction | 0 | 0 | 2 | 0 | 0 | # | 10 | Baseline | ancom | 7 |
| # | disease | 4 | 30 | 0 | 0.117647059 | 1 | # | 10 | Baseline | aldex2 | 7 |
| # | sex | 2 | 10 | 0 | 0.166666667 | 1 | # | 10 | Baseline | aldex2 | 7 |
| # | time | 2 | 15 | 0 | 0.117647059 | 1 | # | 10 | Baseline | aldex2 | 7 |
| # | interaction | 0 | 0 | 2 | 0 | 0 | # | 10 | Baseline | aldex2 | 7 |
| # | disease | 4 | 0 | 0 | 1 | 1 | # | 10 | Small | paralle | 7 |
| # | sex | 2 | 0 | 0 | 1 | 1 | # | 10 | Small | paralle | 7 |
| # | time | 2 | 0 | 0 | 1 | 1 | # | 10 | Small | paralle | 7 |
| # | interaction | 2 | 0 | 0 | 1 | 1 | # | 10 | Small | paralle | 7 |
| # | disease | 4 | 2 | 0 | 0.666666667 | 1 | # | 10 | Small | ancom | 7 |
| # | sex | 2 | 0 | 0 | 1 | 1 | # | 10 | Small | ancom | 7 |
| # | time | 2 | 2 | 0 | 0.5 | 1 | # | 10 | Small | ancom | 7 |
| # | interaction | 0 | 0 | 2 | 0 | 0 | # | 10 | Small | ancom | 7 |
| # | disease | 4 | 3 | 0 | 0.571428571 | 1 | # | 10 | Small | aldex2 | 7 |
| # | sex | 2 | 0 | 0 | 1 | 1 | # | 10 | Small | aldex2 | 7 |
| # | time | 2 | 2 | 0 | 0.5 | 1 | # | 10 | Small | aldex2 | 7 |
| # | interaction | 0 | 0 | 2 | 0 | 0 | # | 10 | Small | aldex2 | 7 |
| # | disease | 7 | 7 | 0 | 0.5 | 1 | # | 20 | Baseline | paralle | 7 |
| # | sex | 4 | 12 | 0 | 0.25 | 1 | # | 20 | Baseline | paralle | 7 |
| # | time | 5 | 10 | 0 | 0.333333333 | 1 | # | 20 | Baseline | paralle | 7 |
| # | interaction | 4 | 5 | 0 | 0.444444444 | 1 | # | 20 | Baseline | paralle | 7 |
| # | disease | 7 | 2 | 0 | 0.777777778 | 1 | # | 20 | Baseline | ancom | 7 |
| # | sex | 4 | 0 | 0 | 1 | 1 | # | 20 | Baseline | ancom | 7 |
| # | time | 5 | 1 | 0 | 0.833333333 | 1 | # | 20 | Baseline | ancom | 7 |
| # | interaction | 0 | 0 | 4 | 0 | 0 | # | 20 | Baseline | ancom | 7 |
| # | disease | 7 | 16 | 0 | 0.304347826 | 1 | # | 20 | Baseline | aldex2 | 7 |
| # | sex | 4 | 8 | 0 | 0.333333333 | 1 | # | 20 | Baseline | aldex2 | 7 |
| # | time | 5 | 17 | 0 | 0.227272727 | 1 | # | 20 | Baseline | aldex2 | 7 |
| # | interaction | 0 | 0 | 4 | 0 | 0 | # | 20 | Baseline | aldex2 | 7 |
| # | disease | 6 | 0 | 1 | 1 | 0.86 | # | 20 | Small | paralle | 7 |
| # | sex | 4 | 0 | 0 | 1 | 1 | # | 20 | Small | paralle | 7 |
| # | time | 5 | 0 | 0 | 1 | 1 | # | 20 | Small | paralle | 7 |
| # | interaction | 1 | 0 | 3 | 1 | 0.25 | # | 20 | Small | paralle | 7 |
| # | disease | 7 | 1 | 0 | 0.875 | 1 | # | 20 | Small | ancom | 7 |
| # | sex | 4 | 0 | 0 | 1 | 1 | # | 20 | Small | ancom | 7 |
| # | time | 5 | 1 | 0 | 0.833333333 | 1 | # | 20 | Small | ancom | 7 |
| # | interaction | 0 | 0 | 4 | 0 | 0 | # | 20 | Small | ancom | 7 |
| # | disease | 7 | 1 | 0 | 0.875 | 1 | # | 20 | Small | aldex2 | 7 |
| # | sex | 4 | 0 | 0 | 1 | 1 | # | 20 | Small | aldex2 | 7 |
| # | time | 5 | 0 | 0 | 1 | 1 | # | 20 | Small | aldex2 | 7 |
| # | interaction | 0 | 0 | 4 | 0 | 0 | # | 20 | Small | aldex2 | 7 |
| # | disease | 7 | 11 | 0 | 0.388888889 | 1 | # | 20 | Baseline | paralle | 7 |
| # | sex | 4 | 8 | 0 | 0.333333333 | 1 | # | 20 | Baseline | paralle | 7 |
| # | time | 5 | 5 | 0 | 0.5 | 1 | # | 20 | Baseline | paralle | 7 |
| # | interaction | 4 | 11 | 0 | 0.266666667 | 1 | # | 20 | Baseline | paralle | 7 |
| # | disease | 7 | 2 | 0 | 0.777777778 | 1 | # | 20 | Baseline | ancom | 7 |
| # | sex | 4 | 0 | 0 | 1 | 1 | # | 20 | Baseline | ancom | 7 |
| # | time | 5 | 1 | 0 | 0.833333333 | 1 | # | 20 | Baseline | ancom | 7 |
| # | interaction | 0 | 0 | 4 | 0 | 0 | # | 20 | Baseline | ancom | 7 |
| # | disease | 7 | 35 | 0 | 0.166666667 | 1 | # | 20 | Baseline | aldex2 | 7 |
| # | sex | 4 | 11 | 0 | 0.266666667 | 1 | # | 20 | Baseline | aldex2 | 7 |
| # | time | 5 | 18 | 0 | 0.217391304 | 1 | # | 20 | Baseline | aldex2 | 7 |
| # | interaction | 0 | 0 | 4 | 0 | 0 | # | 20 | Baseline | aldex2 | 7 |
| # | disease | 7 | 0 | 0 | 1 | 1 | # | 20 | Small | paralle | 7 |
| # | sex | 4 | 0 | 0 | 1 | 1 | # | 20 | Small | paralle | 7 |
| # | time | 5 | 0 | 0 | 1 | 1 | # | 20 | Small | paralle | 7 |
| # | interaction | 4 | 0 | 0 | 1 | 1 | # | 20 | Small | paralle | 7 |
| # | disease | 7 | 2 | 0 | 0.777777778 | 1 | # | 20 | Small | ancom | 7 |
| # | sex | 4 | 0 | 0 | 1 | 1 | # | 20 | Small | ancom | 7 |
| # | time | 5 | 1 | 0 | 0.833333333 | 1 | # | 20 | Small | ancom | 7 |
| # | interaction | 0 | 0 | 4 | 0 | 0 | # | 20 | Small | ancom | 7 |
| # | disease | 7 | 2 | 0 | 0.777777778 | 1 | # | 20 | Small | aldex2 | 7 |
| # | sex | 4 | 0 | 0 | 1 | 1 | # | 20 | Small | aldex2 | 7 |
| # | time | 5 | 3 | 0 | 0.625 | 1 | # | 20 | Small | aldex2 | 7 |
| # | interaction | 0 | 0 | 4 | 0 | 0 | # | 20 | Small | aldex2 | 7 |
| # | disease | 7 | 9 | 0 | 0.4375 | 1 | # | 20 | Baseline | paralle | 7 |
| # | sex | 4 | 4 | 0 | 0.5 | 1 | # | 20 | Baseline | paralle | 7 |
| # | time | 5 | 3 | 0 | 0.625 | 1 | # | 20 | Baseline | paralle | 7 |
| # | interaction | 4 | 5 | 0 | 0.444444444 | 1 | # | 20 | Baseline | paralle | 7 |
| # | disease | 7 | 4 | 0 | 0.636363636 | 1 | # | 20 | Baseline | ancom | 7 |
| # | sex | 4 | 0 | 0 | 1 | 1 | # | 20 | Baseline | ancom | 7 |
| # | time | 5 | 3 | 0 | 0.625 | 1 | # | 20 | Baseline | ancom | 7 |
| # | interaction | 0 | 0 | 4 | 0 | 0 | # | 20 | Baseline | ancom | 7 |
| # | disease | 7 | 68 | 0 | 0.127272727 | 1 | # | 20 | Baseline | aldex2 | 7 |
| # | sex | 4 | 18 | 0 | 0.181818182 | 1 | # | 20 | Baseline | aldex2 | 7 |
| # | time | 5 | 32 | 0 | 0.135135135 | 1 | # | 20 | Baseline | aldex2 | 7 |
| # | interaction | 0 | 0 | 4 | 0 | 0 | # | 20 | Baseline | aldex2 | 7 |
| # | disease | 7 | 0 | 0 | 1 | 1 | # | 20 | Small | paralle | 7 |
| # | sex | 4 | 0 | 0 | 1 | 1 | # | 20 | Small | paralle | 7 |
| # | time | 5 | 0 | 0 | 1 | 1 | # | 20 | Small | paralle | 7 |
| # | interaction | 4 | 0 | 0 | 1 | 1 | # | 20 | Small | paralle | 7 |
| # | disease | 7 | 3 | 0 | 0.7 | 1 | # | 20 | Small | ancom | 7 |
| # | sex | 4 | 0 | 0 | 1 | 1 | # | 20 | Small | ancom | 7 |
| # | time | 5 | 3 | 0 | 0.625 | 1 | # | 20 | Small | ancom | 7 |
| # | interaction | 0 | 0 | 4 | 0 | 0 | # | 20 | Small | ancom | 7 |
| # | disease | 7 | 6 | 0 | 0.538461538 | 1 | # | 20 | Small | aldex2 | 7 |
| # | sex | 4 | 0 | 0 | 1 | 1 | # | 20 | Small | aldex2 | 7 |
| # | time | 5 | 3 | 0 | 0.625 | 1 | # | 20 | Small | aldex2 | 7 |
| # | interaction | 0 | 0 | 4 | 0 | 0 | # | 20 | Small | aldex2 | 7 |
| # | disease | 7 | 13 | 0 | 0.35 | 1 | # | 20 | Baseline | paralle | 7 |
| # | sex | 4 | 3 | 0 | 0.571428571 | 1 | # | 20 | Baseline | paralle | 7 |
| # | time | 5 | 8 | 0 | 0.384615385 | 1 | # | 20 | Baseline | paralle | 7 |
| # | interaction | 4 | 2 | 0 | 0.666666667 | 1 | # | 20 | Baseline | paralle | 7 |
| # | disease | 7 | 4 | 0 | 0.636363636 | 1 | # | 20 | Baseline | ancom | 7 |
| # | sex | 4 | 0 | 0 | 1 | 1 | # | 20 | Baseline | ancom | 7 |
| # | time | 5 | 4 | 0 | 0.555555556 | 1 | # | 20 | Baseline | ancom | 7 |
| # | interaction | 0 | 0 | 4 | 0 | 0 | # | 20 | Baseline | ancom | 7 |
| # | disease | 7 | 54 | 0 | 0.098591549 | 1 | # | 20 | Baseline | aldex2 | 7 |
| # | sex | 4 | 21 | 0 | 0.16 | 1 | # | 20 | Baseline | aldex2 | 7 |
| # | time | 5 | 60 | 0 | 0.111111111 | 1 | # | 20 | Baseline | aldex2 | 7 |
| # | interaction | 0 | 0 | 4 | 0 | 0 | # | 20 | Baseline | aldex2 | 7 |
| # | disease | 7 | 0 | 0 | 1 | 1 | # | 20 | Small | paralle | 7 |
| # | sex | 4 | 0 | 0 | 1 | 1 | # | 20 | Small | paralle | 7 |
| # | time | 5 | 0 | 0 | 1 | 1 | # | 20 | Small | paralle | 7 |
| # | interaction | 4 | 0 | 0 | 1 | 1 | # | 20 | Small | paralle | 7 |
| # | disease | 7 | 4 | 0 | 0.636363636 | 1 | # | 20 | Small | ancom | 7 |
| # | sex | 4 | 0 | 0 | 1 | 1 | # | 20 | Small | ancom | 7 |
| # | time | 5 | 4 | 0 | 0.555555556 | 1 | # | 20 | Small | ancom | 7 |
| # | interaction | 0 | 0 | 4 | 0 | 0 | # | 20 | Small | ancom | 7 |
| # | disease | 7 | 8 | 0 | 0.466666667 | 1 | # | 20 | Small | aldex2 | 7 |
| # | sex | 4 | 0 | 0 | 1 | 1 | # | 20 | Small | aldex2 | 7 |
| # | time | 5 | 4 | 0 | 0.555555556 | 1 | # | 20 | Small | aldex2 | 7 |
| # | interaction | 0 | 0 | 4 | 0 | 0 | # | 20 | Small | aldex2 | 7 |
| # | disease | 4 | 12 | 0 | 0.25 | 1 | # | 10 | Baseline | paralle | 8 |

| | | | | | | | | | | | |
|---|---|---|---|---|---|---|---|---|---|---|---|
| # | sex | 2 | 6 | 0 | 0.25 | 1 | # | 10 | Baseline | paralle | 8 |
| # | time | 2 | 10 | 0 | 0.166666667 | 1 | # | 10 | Baseline | paralle | 8 |
| # | interaction | 2 | 13 | 0 | 0.133333333 | 1 | # | 10 | Baseline | paralle | 8 |
| # | disease | 4 | 0 | 0 | 1 | 1 | # | 10 | Baseline | ancom | 8 |
| # | sex | 2 | 0 | 0 | 1 | 1 | # | 10 | Baseline | ancom | 8 |
| # | time | 2 | 1 | 0 | 0.666666667 | 1 | # | 10 | Baseline | ancom | 8 |
| # | interaction | 0 | 0 | 2 | 0 | 0 | # | 10 | Baseline | ancom | 8 |
| # | disease | 4 | 11 | 0 | 0.266666667 | 1 | # | 10 | Baseline | sided2 | 8 |
| # | sex | 2 | 4 | 0 | 0.333333333 | 1 | # | 10 | Baseline | sided2 | 8 |
| # | time | 2 | 7 | 0 | 0.222222222 | 1 | # | 10 | Baseline | sided2 | 8 |
| # | interaction | 0 | 0 | 2 | 0 | 0 | # | 10 | Baseline | sided2 | 8 |
| # | disease | 4 | 0 | 0 | 1 | 1 | # | 10 | Small | paralle | 8 |
| # | sex | 2 | 0 | 0 | 1 | 1 | # | 10 | Small | paralle | 8 |
| # | time | 2 | 0 | 0 | 1 | 1 | # | 10 | Small | paralle | 8 |
| # | interaction | 0 | 1 | 2 | 0 | 0 | # | 10 | Small | paralle | 8 |
| # | disease | 4 | 0 | 0 | 1 | 1 | # | 10 | Small | ancom | 8 |
| # | sex | 2 | 0 | 0 | 1 | 1 | # | 10 | Small | ancom | 8 |
| # | time | 2 | 0 | 0 | 1 | 1 | # | 10 | Small | ancom | 8 |
| # | interaction | 0 | 0 | 2 | 0 | 0 | # | 10 | Small | ancom | 8 |
| # | disease | 4 | 0 | 0 | 1 | 1 | # | 10 | Small | sided2 | 8 |
| # | sex | 2 | 0 | 0 | 1 | 1 | # | 10 | Small | sided2 | 8 |
| # | time | 2 | 1 | 0 | 0.666666667 | 1 | # | 10 | Small | sided2 | 8 |
| # | interaction | 0 | 0 | 2 | 0 | 0 | # | 10 | Small | sided2 | 8 |
| # | disease | 4 | 6 | 0 | 0.4 | 1 | # | 10 | Baseline | paralle | 8 |
| # | sex | 2 | 6 | 0 | 0.25 | 1 | # | 10 | Baseline | paralle | 8 |
| # | time | 2 | 2 | 0 | 0.5 | 1 | # | 10 | Baseline | paralle | 8 |
| # | interaction | 2 | 7 | 0 | 0.222222222 | 1 | # | 10 | Baseline | paralle | 8 |
| # | disease | 4 | 2 | 0 | 0.666666667 | 1 | # | 10 | Baseline | ancom | 8 |
| # | sex | 2 | 0 | 0 | 1 | 1 | # | 10 | Baseline | ancom | 8 |
| # | time | 2 | 2 | 0 | 0.5 | 1 | # | 10 | Baseline | ancom | 8 |
| # | interaction | 0 | 0 | 2 | 0 | 0 | # | 10 | Baseline | ancom | 8 |
| # | disease | 4 | 14 | 0 | 0.222222222 | 1 | # | 10 | Baseline | sided2 | 8 |
| # | sex | 2 | 2 | 0 | 0.5 | 1 | # | 10 | Baseline | sided2 | 8 |
| # | time | 2 | 6 | 0 | 0.25 | 1 | # | 10 | Baseline | sided2 | 8 |
| # | interaction | 0 | 0 | 2 | 0 | 0 | # | 10 | Baseline | sided2 | 8 |
| # | disease | 4 | 1 | 0 | 0.8 | 1 | # | 10 | Small | paralle | 8 |
| # | sex | 2 | 0 | 0 | 1 | 1 | # | 10 | Small | paralle | 8 |
| # | time | 2 | 0 | 0 | 1 | 1 | # | 10 | Small | paralle | 8 |
| # | interaction | 1 | 0 | 1 | 1 | 0.5 | # | 10 | Small | paralle | 8 |
| # | disease | 4 | 2 | 0 | 0.666666667 | 1 | # | 10 | Small | ancom | 8 |
| # | sex | 2 | 0 | 0 | 1 | 1 | # | 10 | Small | ancom | 8 |
| # | time | 2 | 2 | 0 | 0.5 | 1 | # | 10 | Small | ancom | 8 |
| # | interaction | 0 | 0 | 2 | 0 | 0 | # | 10 | Small | ancom | 8 |
| # | disease | 4 | 2 | 0 | 0.666666667 | 1 | # | 10 | Small | sided2 | 8 |
| # | sex | 2 | 0 | 0 | 1 | 1 | # | 10 | Small | sided2 | 8 |
| # | time | 2 | 2 | 0 | 0.5 | 1 | # | 10 | Small | sided2 | 8 |
| # | interaction | 0 | 0 | 2 | 0 | 0 | # | 10 | Small | sided2 | 8 |
| # | disease | 4 | 5 | 0 | 0.444444444 | 1 | # | 10 | Baseline | paralle | 8 |
| # | sex | 2 | 2 | 0 | 0.5 | 1 | # | 10 | Baseline | paralle | 8 |
| # | time | 2 | 1 | 0 | 0.666666667 | 1 | # | 10 | Baseline | paralle | 8 |
| # | interaction | 2 | 5 | 0 | 0.285714286 | 1 | # | 10 | Baseline | paralle | 8 |
| # | disease | 4 | 2 | 0 | 0.666666667 | 1 | # | 10 | Baseline | ancom | 8 |
| # | sex | 2 | 0 | 0 | 1 | 1 | # | 10 | Baseline | ancom | 8 |
| # | time | 2 | 2 | 0 | 0.5 | 1 | # | 10 | Baseline | ancom | 8 |
| # | interaction | 0 | 0 | 2 | 0 | 0 | # | 10 | Baseline | ancom | 8 |
| # | disease | 4 | 17 | 0 | 0.19047619 | 1 | # | 10 | Baseline | sided2 | 8 |
| # | sex | 2 | 7 | 0 | 0.222222222 | 1 | # | 10 | Baseline | sided2 | 8 |
| # | time | 2 | 8 | 0 | 0.2 | 1 | # | 10 | Baseline | sided2 | 8 |
| # | interaction | 0 | 0 | 2 | 0 | 0 | # | 10 | Baseline | sided2 | 8 |
| # | disease | 4 | 0 | 0 | 1 | 1 | # | 10 | Small | paralle | 8 |
| # | sex | 2 | 0 | 0 | 1 | 1 | # | 10 | Small | paralle | 8 |
| # | time | 2 | 0 | 0 | 1 | 1 | # | 10 | Small | paralle | 8 |
| # | interaction | 2 | 0 | 0 | 1 | 1 | # | 10 | Small | paralle | 8 |
| # | disease | 4 | 2 | 0 | 0.666666667 | 1 | # | 10 | Small | ancom | 8 |
| # | sex | 2 | 0 | 0 | 1 | 1 | # | 10 | Small | ancom | 8 |
| # | time | 2 | 2 | 0 | 0.5 | 1 | # | 10 | Small | ancom | 8 |
| # | interaction | 0 | 0 | 2 | 0 | 0 | # | 10 | Small | ancom | 8 |
| # | disease | 4 | 3 | 0 | 0.571428571 | 1 | # | 10 | Small | sided2 | 8 |
| # | sex | 2 | 1 | 0 | 0.666666667 | 1 | # | 10 | Small | sided2 | 8 |
| # | time | 2 | 2 | 0 | 0.5 | 1 | # | 10 | Small | sided2 | 8 |
| # | interaction | 0 | 0 | 2 | 0 | 0 | # | 10 | Small | sided2 | 8 |
| # | disease | 4 | 8 | 0 | 0.333333333 | 1 | # | 10 | Baseline | paralle | 8 |
| # | sex | 2 | 2 | 0 | 0.5 | 1 | # | 10 | Baseline | paralle | 8 |
| # | time | 2 | 2 | 0 | 0.5 | 1 | # | 10 | Baseline | paralle | 8 |
| # | interaction | 2 | 7 | 0 | 0.222222222 | 1 | # | 10 | Baseline | paralle | 8 |
| # | disease | 4 | 2 | 0 | 0.666666667 | 1 | # | 10 | Baseline | ancom | 8 |
| # | sex | 2 | 0 | 0 | 1 | 1 | # | 10 | Baseline | ancom | 8 |
| # | time | 2 | 2 | 0 | 0.5 | 1 | # | 10 | Baseline | ancom | 8 |
| # | interaction | 0 | 0 | 2 | 0 | 0 | # | 10 | Baseline | ancom | 8 |
| # | disease | 4 | 24 | 0 | 0.142857143 | 1 | # | 10 | Baseline | sided2 | 8 |
| # | sex | 2 | 9 | 0 | 0.181818182 | 1 | # | 10 | Baseline | sided2 | 8 |
| # | time | 2 | 8 | 0 | 0.2 | 1 | # | 10 | Baseline | sided2 | 8 |
| # | interaction | 0 | 0 | 2 | 0 | 0 | # | 10 | Baseline | sided2 | 8 |
| # | disease | 4 | 1 | 0 | 0.8 | 1 | # | 10 | Small | paralle | 8 |
| # | sex | 2 | 0 | 0 | 1 | 1 | # | 10 | Small | paralle | 8 |
| # | time | 2 | 0 | 0 | 1 | 1 | # | 10 | Small | paralle | 8 |
| # | interaction | 2 | 0 | 0 | 1 | 1 | # | 10 | Small | paralle | 8 |
| # | disease | 4 | 2 | 0 | 0.666666667 | 1 | # | 10 | Small | ancom | 8 |
| # | sex | 2 | 0 | 0 | 1 | 1 | # | 10 | Small | ancom | 8 |
| # | time | 2 | 2 | 0 | 0.5 | 1 | # | 10 | Small | ancom | 8 |
| # | interaction | 0 | 0 | 2 | 0 | 0 | # | 10 | Small | ancom | 8 |
| # | disease | 4 | 3 | 0 | 0.571428571 | 1 | # | 10 | Small | sided2 | 8 |
| # | sex | 2 | 0 | 0 | 1 | 1 | # | 10 | Small | sided2 | 8 |
| # | time | 2 | 2 | 0 | 0.5 | 1 | # | 10 | Small | sided2 | 8 |
| # | interaction | 0 | 0 | 2 | 0 | 0 | # | 10 | Small | sided2 | 8 |
| # | disease | 7 | 5 | 0 | 0.583333333 | 1 | # | 20 | Baseline | paralle | 8 |
| # | sex | 4 | 8 | 0 | 0.333333333 | 1 | # | 20 | Baseline | paralle | 8 |
| # | time | 5 | 7 | 0 | 0.416666667 | 1 | # | 20 | Baseline | paralle | 8 |
| # | interaction | 4 | 7 | 0 | 0.363636364 | 1 | # | 20 | Baseline | paralle | 8 |
| # | disease | 7 | 3 | 0 | 0.7 | 1 | # | 20 | Baseline | ancom | 8 |
| # | sex | 4 | 0 | 0 | 1 | 1 | # | 20 | Baseline | ancom | 8 |
| # | time | 5 | 1 | 0 | 0.833333333 | 1 | # | 20 | Baseline | ancom | 8 |
| # | interaction | 0 | 0 | 4 | 0 | 0 | # | 20 | Baseline | ancom | 8 |
| # | disease | 7 | 17 | 0 | 0.291666667 | 1 | # | 20 | Baseline | sided2 | 8 |
| # | sex | 4 | 8 | 0 | 0.333333333 | 1 | # | 20 | Baseline | sided2 | 8 |
| # | time | 5 | 18 | 0 | 0.217391304 | 1 | # | 20 | Baseline | sided2 | 8 |
| # | interaction | 0 | 0 | 4 | 0 | 0 | # | 20 | Baseline | sided2 | 8 |
| # | disease | 7 | 0 | 0 | 1 | 1 | # | 20 | Small | paralle | 8 |
| # | sex | 4 | 0 | 0 | 1 | 1 | # | 20 | Small | paralle | 8 |
| # | time | 5 | 0 | 0 | 1 | 1 | # | 20 | Small | paralle | 8 |
| # | interaction | 2 | 0 | 2 | 1 | 0.5 | # | 20 | Small | paralle | 8 |
| # | disease | 7 | 3 | 0 | 0.7 | 1 | # | 20 | Small | ancom | 8 |
| # | sex | 4 | 0 | 0 | 1 | 1 | # | 20 | Small | ancom | 8 |
| # | time | 5 | 1 | 0 | 0.833333333 | 1 | # | 20 | Small | ancom | 8 |
| # | interaction | 0 | 0 | 4 | 0 | 0 | # | 20 | Small | ancom | 8 |
| # | disease | 7 | 2 | 0 | 0.777777778 | 1 | # | 20 | Small | sided2 | 8 |
| # | sex | 4 | 0 | 0 | 1 | 1 | # | 20 | Small | sided2 | 8 |
| # | time | 5 | 0 | 0 | 1 | 1 | # | 20 | Small | sided2 | 8 |
| # | interaction | 0 | 0 | 4 | 0 | 0 | # | 20 | Small | sided2 | 8 |
| # | disease | 7 | 4 | 0 | 0.636363636 | 1 | # | 20 | Baseline | paralle | 8 |
| # | sex | 4 | 4 | 0 | 0.5 | 1 | # | 20 | Baseline | paralle | 8 |
| # | time | 5 | 9 | 0 | 0.357142857 | 1 | # | 20 | Baseline | paralle | 8 |
| # | interaction | 4 | 4 | 0 | 0.5 | 1 | # | 20 | Baseline | paralle | 8 |
| # | disease | 7 | 4 | 0 | 0.636363636 | 1 | # | 20 | Baseline | ancom | 8 |
| # | sex | 4 | 0 | 0 | 1 | 1 | # | 20 | Baseline | ancom | 8 |
| # | time | 5 | 4 | 0 | 0.555555556 | 1 | # | 20 | Baseline | ancom | 8 |
| # | interaction | 0 | 0 | 4 | 0 | 0 | # | 20 | Baseline | ancom | 8 |
| # | disease | 7 | 30 | 0 | 0.189189189 | 1 | # | 20 | Baseline | sided2 | 8 |
| # | sex | 4 | 9 | 0 | 0.307692308 | 1 | # | 20 | Baseline | sided2 | 8 |
| # | time | 5 | 31 | 0 | 0.138888889 | 1 | # | 20 | Baseline | sided2 | 8 |
| # | interaction | 0 | 0 | 4 | 0 | 0 | # | 20 | Baseline | sided2 | 8 |
| # | disease | 7 | 0 | 0 | 1 | 1 | # | 20 | Small | paralle | 8 |
| # | sex | 4 | 0 | 0 | 1 | 1 | # | 20 | Small | paralle | 8 |
| # | time | 5 | 0 | 0 | 1 | 1 | # | 20 | Small | paralle | 8 |
| # | interaction | 4 | 0 | 0 | 1 | 1 | # | 20 | Small | paralle | 8 |
| # | disease | 7 | 4 | 0 | 0.636363636 | 1 | # | 20 | Small | ancom | 8 |
| # | sex | 4 | 0 | 0 | 1 | 1 | # | 20 | Small | ancom | 8 |
| # | time | 5 | 4 | 0 | 0.555555556 | 1 | # | 20 | Small | ancom | 8 |
| # | interaction | 0 | 0 | 4 | 0 | 0 | # | 20 | Small | ancom | 8 |
| # | disease | 7 | 4 | 0 | 0.636363636 | 1 | # | 20 | Small | sided2 | 8 |
| # | sex | 4 | 0 | 0 | 1 | 1 | # | 20 | Small | sided2 | 8 |
| # | time | 5 | 4 | 0 | 0.555555556 | 1 | # | 20 | Small | sided2 | 8 |
| # | interaction | 0 | 0 | 4 | 0 | 0 | # | 20 | Small | sided2 | 8 |
| # | disease | 7 | 8 | 0 | 0.466666667 | 1 | # | 20 | Baseline | paralle | 8 |
| # | sex | 4 | 3 | 0 | 0.571428571 | 1 | # | 20 | Baseline | paralle | 8 |
| # | time | 5 | 10 | 0 | 0.333333333 | 1 | # | 20 | Baseline | paralle | 8 |
| # | interaction | 4 | 5 | 0 | 0.444444444 | 1 | # | 20 | Baseline | paralle | 8 |
| # | disease | 7 | 4 | 0 | 0.636363636 | 1 | # | 20 | Baseline | ancom | 8 |
| # | sex | 4 | 0 | 0 | 1 | 1 | # | 20 | Baseline | ancom | 8 |
| # | time | 5 | 4 | 0 | 0.555555556 | 1 | # | 20 | Baseline | ancom | 8 |
| # | interaction | 0 | 0 | 4 | 0 | 0 | # | 20 | Baseline | ancom | 8 |
| # | disease | 7 | 46 | 0 | 0.132075472 | 1 | # | 20 | Baseline | sided2 | 8 |
| # | sex | 4 | 11 | 0 | 0.266666667 | 1 | # | 20 | Baseline | sided2 | 8 |
| # | time | 5 | 36 | 0 | 0.12195122 | 1 | # | 20 | Baseline | sided2 | 8 |
| # | interaction | 0 | 0 | 4 | 0 | 0 | # | 20 | Baseline | sided2 | 8 |
| # | disease | 7 | 1 | 0 | 0.875 | 1 | # | 20 | Small | paralle | 8 |
| # | sex | 4 | 0 | 0 | 1 | 1 | # | 20 | Small | paralle | 8 |
| # | time | 5 | 0 | 0 | 1 | 1 | # | 20 | Small | paralle | 8 |
| # | interaction | 4 | 0 | 0 | 1 | 1 | # | 20 | Small | paralle | 8 |
| # | disease | 7 | 4 | 0 | 0.636363636 | 1 | # | 20 | Small | ancom | 8 |
| # | sex | 4 | 0 | 0 | 1 | 1 | # | 20 | Small | ancom | 8 |
| # | time | 5 | 4 | 0 | 0.555555556 | 1 | # | 20 | Small | ancom | 8 |
| # | interaction | 0 | 0 | 4 | 0 | 0 | # | 20 | Small | ancom | 8 |
| # | disease | 7 | 7 | 0 | 0.5 | 1 | # | 20 | Small | sided2 | 8 |
| # | sex | 4 | 0 | 0 | 1 | 1 | # | 20 | Small | sided2 | 8 |
| # | time | 5 | 4 | 0 | 0.555555556 | 1 | # | 20 | Small | sided2 | 8 |
| # | interaction | 0 | 0 | 4 | 0 | 0 | # | 20 | Small | sided2 | 8 |
| # | disease | 7 | 17 | 0 | 0.291666667 | 1 | # | 20 | Baseline | paralle | 8 |
| # | sex | 4 | 6 | 0 | 0.4 | 1 | # | 20 | Baseline | paralle | 8 |
| # | time | 5 | 4 | 0 | 0.555555556 | 1 | # | 20 | Baseline | paralle | 8 |
| # | interaction | 4 | 2 | 0 | 0.666666667 | 1 | # | 20 | Baseline | paralle | 8 |
| # | disease | 7 | 4 | 0 | 0.636363636 | 1 | # | 20 | Baseline | ancom | 8 |
| # | sex | 4 | 0 | 0 | 1 | 1 | # | 20 | Baseline | ancom | 8 |
| # | time | 5 | 4 | 0 | 0.555555556 | 1 | # | 20 | Baseline | ancom | 8 |
| # | interaction | 0 | 0 | 4 | 0 | 0 | # | 20 | Baseline | ancom | 8 |
| # | disease | 7 | 53 | 0 | 0.1 | 1 | # | 20 | Baseline | sided2 | 8 |
| # | sex | 4 | 20 | 0 | 0.166666667 | 1 | # | 20 | Baseline | sided2 | 8 |
| # | time | 5 | 67 | 0 | 0.096153846 | 1 | # | 20 | Baseline | sided2 | 8 |
| # | interaction | 0 | 0 | 4 | 0 | 0 | # | 20 | Baseline | sided2 | 8 |
| # | disease | 7 | 0 | 0 | 1 | 1 | # | 20 | Small | paralle | 8 |
| # | sex | 4 | 0 | 0 | 1 | 1 | # | 20 | Small | paralle | 8 |
| # | time | 5 | 0 | 0 | 1 | 1 | # | 20 | Small | paralle | 8 |
| # | interaction | 4 | 0 | 0 | 1 | 1 | # | 20 | Small | paralle | 8 |
| # | disease | 7 | 4 | 0 | 0.636363636 | 1 | # | 20 | Small | ancom | 8 |
| # | sex | 4 | 0 | 0 | 1 | 1 | # | 20 | Small | ancom | 8 |
| # | time | 5 | 4 | 0 | 0.555555556 | 1 | # | 20 | Small | ancom | 8 |
| # | interaction | 0 | 0 | 4 | 0 | 0 | # | 20 | Small | ancom | 8 |
| # | disease | 7 | 14 | 0 | 0.333333333 | 1 | # | 20 | Small | sided2 | 8 |
| # | sex | 4 | 0 | 0 | 1 | 1 | # | 20 | Small | sided2 | 8 |
| # | time | 5 | 4 | 0 | 0.555555556 | 1 | # | 20 | Small | sided2 | 8 |
| # | interaction | 0 | 0 | 4 | 0 | 0 | # | 20 | Small | sided2 | 8 |
| # | disease | 4 | 13 | 0 | 0.235294118 | 1 | # | 10 | Baseline | paralle | 9 |
| # | sex | 2 | 3 | 0 | 0.4 | 1 | # | 10 | Baseline | paralle | 9 |
| # | time | 2 | 6 | 0 | 0.25 | 1 | # | 10 | Baseline | paralle | 9 |
| # | interaction | 2 | 12 | 0 | 0.142857143 | 1 | # | 10 | Baseline | paralle | 9 |
| # | disease | 4 | 0 | 0 | 1 | 1 | # | 10 | Baseline | ancom | 9 |
| # | sex | 2 | 0 | 0 | 1 | 1 | # | 10 | Baseline | ancom | 9 |
| # | time | 2 | 0 | 0 | 1 | 1 | # | 10 | Baseline | ancom | 9 |
| # | interaction | 0 | 0 | 2 | 0 | 0 | # | 10 | Baseline | ancom | 9 |
| # | disease | 4 | 9 | 0 | 0.307692308 | 1 | # | 10 | Baseline | sided2 | 9 |
| # | sex | 2 | 8 | 0 | 0.2 | 1 | # | 10 | Baseline | sided2 | 9 |
| # | time | 2 | 7 | 0 | 0.222222222 | 1 | # | 10 | Baseline | sided2 | 9 |
| # | interaction | 0 | 0 | 2 | 0 | 0 | # | 10 | Baseline | sided2 | 9 |
| # | disease | 4 | 0 | 0 | 1 | 1 | # | 10 | Small | paralle | 9 |
| # | sex | 2 | 0 | 0 | 1 | 1 | # | 10 | Small | paralle | 9 |
| # | time | 2 | 1 | 0 | 0.666666667 | 1 | # | 10 | Small | paralle | 9 |
| # | interaction | 1 | 0 | 1 | 1 | 0.5 | # | 10 | Small | paralle | 9 |
| # | disease | 4 | 0 | 0 | 1 | 1 | # | 10 | Small | ancom | 9 |
| # | sex | 2 | 0 | 0 | 1 | 1 | # | 10 | Small | ancom | 9 |
| # | time | 2 | 0 | 0 | 1 | 1 | # | 10 | Small | ancom | 9 |
| # | interaction | 0 | 0 | 2 | 0 | 0 | # | 10 | Small | ancom | 9 |
| # | disease | 4 | 2 | 0 | 0.666666667 | 1 | # | 10 | Small | sided2 | 9 |
| # | sex | 2 | 0 | 0 | 1 | 1 | # | 10 | Small | sided2 | 9 |
| # | time | 2 | 0 | 0 | 1 | 1 | # | 10 | Small | sided2 | 9 |
| # | interaction | 0 | 0 | 2 | 0 | 0 | # | 10 | Small | sided2 | 9 |
| # | disease | 4 | 9 | 0 | 0.307692308 | 1 | # | 10 | Baseline | paralle | 9 |
| # | sex | 2 | 2 | 0 | 0.5 | 1 | # | 10 | Baseline | paralle | 9 |
| # | time | 2 | 6 | 0 | 0.25 | 1 | # | 10 | Baseline | paralle | 9 |
| # | interaction | 2 | 7 | 0 | 0.222222222 | 1 | # | 10 | Baseline | paralle | 9 |
| # | disease | 4 | 1 | 0 | 0.8 | 1 | # | 10 | Baseline | ancom | 9 |
| # | sex | 2 | 0 | 0 | 1 | 1 | # | 10 | Baseline | ancom | 9 |
| # | time | 2 | 1 | 0 | 0.666666667 | 1 | # | 10 | Baseline | ancom | 9 |
| # | interaction | 0 | 0 | 2 | 0 | 0 | # | 10 | Baseline | ancom | 9 |
| # | disease | 4 | 15 | 0 | 0.210526316 | 1 | # | 10 | Baseline | sided2 | 9 |
| # | sex | 2 | 6 | 0 | 0.25 | 1 | # | 10 | Baseline | sided2 | 9 |
| # | time | 2 | 8 | 0 | 0.2 | 1 | # | 10 | Baseline | sided2 | 9 |
| # | interaction | 0 | 0 | 2 | 0 | 0 | # | 10 | Baseline | sided2 | 9 |
| # | disease | 4 | 0 | 0 | 1 | 1 | # | 10 | Small | paralle | 9 |
| # | sex | 2 | 0 | 0 | 1 | 1 | # | 10 | Small | paralle | 9 |
| # | time | 2 | 0 | 0 | 1 | 1 | # | 10 | Small | paralle | 9 |
| # | interaction | 2 | 0 | 0 | 1 | 1 | # | 10 | Small | paralle | 9 |
| # | disease | 4 | 1 | 0 | 0.8 | 1 | # | 10 | Small | ancom | 9 |
| # | sex | 2 | 0 | 0 | 1 | 1 | # | 10 | Small | ancom | 9 |
| # | time | 2 | 0 | 0 | 1 | 1 | # | 10 | Small | ancom | 9 |
| # | interaction | 0 | 0 | 2 | 0 | 0 | # | 10 | Small | ancom | 9 |
| # | disease | 4 | 2 | 0 | 0.666666667 | 1 | # | 10 | Small | sided2 | 9 |
| # | sex | 2 | 0 | 0 | 1 | 1 | # | 10 | Small | sided2 | 9 |
| # | time | 2 | 1 | 0 | 0.666666667 | 1 | # | 10 | Small | sided2 | 9 |
| # | interaction | 0 | 0 | 2 | 0 | 0 | # | 10 | Small | sided2 | 9 |
| # | disease | 4 | 4 | 0 | 0.5 | 1 | # | 10 | Baseline | paralle | 9 |
| # | sex | 2 | 2 | 0 | 0.5 | 1 | # | 10 | Baseline | paralle | 9 |
| # | time | 2 | 6 | 0 | 0.25 | 1 | # | 10 | Baseline | paralle | 9 |
| # | interaction | 2 | 5 | 0 | 0.285714286 | 1 | # | 10 | Baseline | paralle | 9 |
| # | disease | 4 | 1 | 0 | 0.8 | 1 | # | 10 | Baseline | ancom | 9 |
| # | sex | 2 | 0 | 0 | 1 | 1 | # | 10 | Baseline | ancom | 9 |
| # | time | 2 | 1 | 0 | 0.666666667 | 1 | # | 10 | Baseline | ancom | 9 |
| # | interaction | 0 | 0 | 2 | 0 | 0 | # | 10 | Baseline | ancom | 9 |
| # | disease | 4 | 20 | 0 | 0.166666667 | 1 | # | 10 | Baseline | sided2 | 9 |
| # | sex | 2 | 5 | 0 | 0.285714286 | 1 | # | 10 | Baseline | sided2 | 9 |
| # | time | 2 | 13 | 0 | 0.133333333 | 1 | # | 10 | Baseline | sided2 | 9 |
| # | interaction | 0 | 0 | 2 | 0 | 0 | # | 10 | Baseline | sided2 | 9 |
| # | disease | 4 | 0 | 0 | 1 | 1 | # | 10 | Small | paralle | 9 |
| # | sex | 2 | 0 | 0 | 1 | 1 | # | 10 | Small | paralle | 9 |
| # | time | 2 | 0 | 0 | 1 | 1 | # | 10 | Small | paralle | 9 |
| # | interaction | 2 | 0 | 0 | 1 | 1 | # | 10 | Small | paralle | 9 |
| # | disease | 4 | 1 | 0 | 0.8 | 1 | # | 10 | Small | ancom | 9 |
| # | sex | 2 | 0 | 0 | 1 | 1 | # | 10 | Small | ancom | 9 |
| # | time | 2 | 1 | 0 | 0.666666667 | 1 | # | 10 | Small | ancom | 9 |
| # | interaction | 0 | 0 | 2 | 0 | 0 | # | 10 | Small | ancom | 9 |
| # | disease | 4 | 2 | 0 | 0.666666667 | 1 | # | 10 | Small | sided2 | 9 |
| # | sex | 2 | 0 | 0 | 1 | 1 | # | 10 | Small | sided2 | 9 |
| # | time | 2 | 1 | 0 | 0.666666667 | 1 | # | 10 | Small | sided2 | 9 |
| # | interaction | 0 | 0 | 2 | 0 | 0 | # | 10 | Small | sided2 | 9 |
| # | disease | 4 | 6 | 0 | 0.4 | 1 | # | 10 | Baseline | paralle | 9 |
| # | sex | 2 | 2 | 0 | 0.5 | 1 | # | 10 | Baseline | paralle | 9 |
| # | time | 2 | 3 | 0 | 0.4 | 1 | # | 10 | Baseline | paralle | 9 |
| # | interaction | 2 | 4 | 0 | 0.333333333 | 1 | # | 10 | Baseline | paralle | 9 |
| # | disease | 4 | 1 | 0 | 0.8 | 1 | # | 10 | Baseline | ancom | 9 |
| # | sex | 2 | 0 | 0 | 1 | 1 | # | 10 | Baseline | ancom | 9 |
| # | time | 2 | 2 | 0 | 0.5 | 1 | # | 10 | Baseline | ancom | 9 |
| # | interaction | 0 | 0 | 2 | 0 | 0 | # | 10 | Baseline | ancom | 9 |
| # | disease | 4 | 30 | 0 | 0.117647059 | 1 | # | 10 | Baseline | sided2 | 9 |
| # | sex | 2 | 8 | 0 | 0.2 | 1 | # | 10 | Baseline | sided2 | 9 |
| # | time | 2 | 16 | 0 | 0.111111111 | 1 | # | 10 | Baseline | sided2 | 9 |
| # | interaction | 0 | 0 | 2 | 0 | 0 | # | 10 | Baseline | sided2 | 9 |
| # | disease | 4 | 0 | 0 | 1 | 1 | # | 10 | Small | paralle | 9 |
| # | sex | 2 | 0 | 0 | 1 | 1 | # | 10 | Small | paralle | 9 |
| # | time | 2 | 0 | 0 | 1 | 1 | # | 10 | Small | paralle | 9 |
| # | interaction | 2 | 0 | 0 | 1 | 1 | # | 10 | Small | paralle | 9 |
| # | disease | 4 | 1 | 0 | 0.8 | 1 | # | 10 | Small | ancom | 9 |
| # | sex | 2 | 0 | 0 | 1 | 1 | # | 10 | Small | ancom | 9 |
| # | time | 2 | 2 | 0 | 0.5 | 1 | # | 10 | Small | ancom | 9 |
| # | interaction | 0 | 0 | 2 | 0 | 0 | # | 10 | Small | ancom | 9 |
| # | disease | 4 | 3 | 0 | 0.571428571 | 1 | # | 10 | Small | sided2 | 9 |
| # | sex | 2 | 0 | 0 | 1 | 1 | # | 10 | Small | sided2 | 9 |
| # | time | 2 | 2 | 0 | 0.5 | 1 | # | 10 | Small | sided2 | 9 |
| # | interaction | 0 | 0 | 2 | 0 | 0 | # | 10 | Small | sided2 | 9 |
| # | disease | 7 | 9 | 0 | 0.4375 | 1 | # | 20 | Baseline | paralle | 9 |
| # | sex | 4 | 10 | 0 | 0.285714286 | 1 | # | 20 | Baseline | paralle | 9 |
| # | time | 5 | 8 | 0 | 0.384615385 | 1 | # | 20 | Baseline | paralle | 9 |
| # | interaction | 4 | 10 | 0 | 0.285714286 | 1 | # | 20 | Baseline | paralle | 9 |
| # | disease | 7 | 1 | 0 | 0.875 | 1 | # | 20 | Baseline | ancom | 9 |
| # | sex | 4 | 0 | 0 | 1 | 1 | # | 20 | Baseline | ancom | 9 |
| # | time | 5 | 1 | 0 | 0.833333333 | 1 | # | 20 | Baseline | ancom | 9 |
| # | interaction | 0 | 0 | 4 | 0 | 0 | # | 20 | Baseline | ancom | 9 |
| # | disease | 7 | 16 | 0 | 0.304347826 | 1 | # | 20 | Baseline | sided2 | 9 |
| # | sex | 4 | 9 | 0 | 0.307692308 | 1 | # | 20 | Baseline | sided2 | 9 |
| # | time | 5 | 14 | 0 | 0.263157895 | 1 | # | 20 | Baseline | sided2 | 9 |
| # | interaction | 0 | 0 | 4 | 0 | 0 | # | 20 | Baseline | sided2 | 9 |
| # | disease | 7 | 0 | 0 | 1 | 1 | # | 20 | Small | paralle | 9 |
| # | sex | 4 | 0 | 0 | 1 | 1 | # | 20 | Small | paralle | 9 |
| # | time | 5 | 0 | 0 | 1 | 1 | # | 20 | Small | paralle | 9 |
| # | interaction | 2 | 0 | 2 | 1 | 0.5 | # | 20 | Small | paralle | 9 |
| # | disease | 7 | 0 | 0 | 1 | 1 | # | 20 | Small | ancom | 9 |
| # | sex | 4 | 0 | 0 | 1 | 1 | # | 20 | Small | ancom | 9 |
| # | time | 5 | 0 | 0 | 1 | 1 | # | 20 | Small | ancom | 9 |
| # | interaction | 0 | 0 | 4 | 0 | 0 | # | 20 | Small | ancom | 9 |
| # | disease | 7 | 1 | 0 | 0.875 | 1 | # | 20 | Small | sided2 | 9 |
| # | sex | 4 | 1 | 0 | 0.8 | 1 | # | 20 | Small | sided2 | 9 |
| # | time | 5 | 0 | 0 | 1 | 1 | # | 20 | Small | sided2 | 9 |
| # | interaction | 0 | 0 | 4 | 0 | 0 | # | 20 | Small | sided2 | 9 |
| # | disease | 7 | 9 | 0 | 0.4375 | 1 | # | 20 | Baseline | paralle | 9 |
| # | sex | 4 | 4 | 0 | 0.5 | 1 | # | 20 | Baseline | paralle | 9 |
| # | time | 5 | 6 | 0 | 0.454545455 | 1 | # | 20 | Baseline | paralle | 9 |
| # | interaction | 4 | 7 | 0 | 0.363636364 | 1 | # | 20 | Baseline | paralle | 9 |
| # | disease | 7 | 3 | 0 | 0.7 | 1 | # | 20 | Baseline | ancom | 9 |
| # | sex | 4 | 0 | 0 | 1 | 1 | # | 20 | Baseline | ancom | 9 |
| # | time | 5 | 3 | 0 | 0.625 | 1 | # | 20 | Baseline | ancom | 9 |
| # | interaction | 0 | 0 | 4 | 0 | 0 | # | 20 | Baseline | ancom | 9 |
| # | disease | 7 | 31 | 0 | 0.184210526 | 1 | # | 20 | Baseline | sided2 | 9 |
| # | sex | 4 | 11 | 0 | 0.266666667 | 1 | # | 20 | Baseline | sided2 | 9 |
| # | time | 5 | 23 | 0 | 0.178571429 | 1 | # | 20 | Baseline | sided2 | 9 |
| # | interaction | 0 | 0 | 4 | 0 | 0 | # | 20 | Baseline | sided2 | 9 |
| # | disease | 7 | 0 | 0 | 1 | 1 | # | 20 | Small | paralle | 9 |
| # | sex | 4 | 0 | 0 | 1 | 1 | # | 20 | Small | paralle | 9 |
| # | time | 5 | 0 | 0 | 1 | 1 | # | 20 | Small | paralle | 9 |
| # | interaction | 2 | 0 | 2 | 1 | 0.5 | # | 20 | Small | paralle | 9 |
| # | disease | 7 | 3 | 0 | 0.7 | 1 | # | 20 | Small | ancom | 9 |
| # | sex | 4 | 0 | 0 | 1 | 1 | # | 20 | Small | ancom | 9 |
| # | time | 5 | 3 | 0 | 0.625 | 1 | # | 20 | Small | ancom | 9 |
| # | interaction | 0 | 0 | 4 | 0 | 0 | # | 20 | Small | ancom | 9 |
| # | disease | 7 | 4 | 0 | 0.636363636 | 1 | # | 20 | Small | sided2 | 9 |
| # | sex | 4 | 0 | 0 | 1 | 1 | # | 20 | Small | sided2 | 9 |
| # | time | 5 | 2 | 0 | 0.714285714 | 1 | # | 20 | Small | sided2 | 9 |
| # | interaction | 0 | 0 | 4 | 0 | 0 | # | 20 | Small | sided2 | 9 |
| # | disease | 7 | 8 | 0 | 0.466666667 | 1 | # | 20 | Baseline | paralle | 9 |
| # | sex | 4 | 7 | 0 | 0.363636364 | 1 | # | 20 | Baseline | paralle | 9 |

| | | | | | | | | | | | |
|---|---|---|---|---|---|---|---|---|---|---|---|
| # | time | 5 | 9 | 0 | 0.357142857 | 1 | # | 20 | Baseline | paralle | 9 |
| # | interaction | 4 | 3 | 0 | 0.571428571 | 1 | # | 20 | Baseline | paralle | 9 |
| # | disease | 7 | 3 | 0 | 0.7 | 1 | # | 20 | Baseline | ancom | 9 |
| # | sex | 4 | 0 | 0 | 1 | 1 | # | 20 | Baseline | ancom | 9 |
| # | time | 5 | 3 | 0 | 0.625 | 1 | # | 20 | Baseline | ancom | 9 |
| # | interaction | 0 | 0 | 4 | 0 | 0 | # | 20 | Baseline | ancom | 9 |
| # | disease | 7 | 69 | 0 | 0.125 | 1 | # | 20 | Baseline | aldex2 | 9 |
| # | sex | 4 | 17 | 0 | 0.19047619 | 1 | # | 20 | Baseline | aldex2 | 9 |
| # | time | 5 | 35 | 0 | 0.125 | 1 | # | 20 | Baseline | aldex2 | 9 |
| # | interaction | 0 | 0 | 4 | 0 | 0 | # | 20 | Baseline | aldex2 | 9 |
| # | disease | 7 | 0 | 0 | 1 | 1 | # | 20 | Small | paralle | 9 |
| # | sex | 4 | 0 | 0 | 1 | 1 | # | 20 | Small | paralle | 9 |
| # | time | 5 | 0 | 0 | 1 | 1 | # | 20 | Small | paralle | 9 |
| # | interaction | 3 | 0 | 1 | 1 | 0.75 | # | 20 | Small | paralle | 9 |
| # | disease | 7 | 3 | 0 | 0.7 | 1 | # | 20 | Small | ancom | 9 |
| # | sex | 4 | 0 | 0 | 1 | 1 | # | 20 | Small | ancom | 9 |
| # | time | 5 | 3 | 0 | 0.625 | 1 | # | 20 | Small | ancom | 9 |
| # | interaction | 0 | 0 | 4 | 0 | 0 | # | 20 | Small | ancom | 9 |
| # | disease | 7 | 8 | 0 | 0.466666667 | 1 | # | 20 | Small | aldex2 | 9 |
| # | sex | 4 | 0 | 0 | 1 | 1 | # | 20 | Small | aldex2 | 9 |
| # | time | 5 | 5 | 0 | 0.5 | 1 | # | 20 | Small | aldex2 | 9 |
| # | interaction | 0 | 0 | 4 | 0 | 0 | # | 20 | Small | aldex2 | 9 |
| # | disease | 7 | 9 | 0 | 0.4375 | 1 | # | 20 | Baseline | paralle | 9 |
| # | sex | 4 | 7 | 0 | 0.363636364 | 1 | # | 20 | Baseline | paralle | 9 |
| # | time | 5 | 9 | 0 | 0.357142857 | 1 | # | 20 | Baseline | paralle | 9 |
| # | interaction | 4 | 6 | 0 | 0.4 | 1 | # | 20 | Baseline | paralle | 9 |
| # | disease | 7 | 3 | 0 | 0.7 | 1 | # | 20 | Baseline | ancom | 9 |
| # | sex | 4 | 0 | 0 | 1 | 1 | # | 20 | Baseline | ancom | 9 |
| # | time | 5 | 4 | 0 | 0.555555556 | 1 | # | 20 | Baseline | ancom | 9 |
| # | interaction | 0 | 0 | 4 | 0 | 0 | # | 20 | Baseline | ancom | 9 |
| # | disease | 7 | 59 | 0 | 0.106060606 | 1 | # | 20 | Baseline | aldex2 | 9 |
| # | sex | 4 | 22 | 0 | 0.153846154 | 1 | # | 20 | Baseline | aldex2 | 9 |
| # | time | 5 | 62 | 0 | 0.106382979 | 1 | # | 20 | Baseline | aldex2 | 9 |
| # | interaction | 0 | 0 | 4 | 0 | 0 | # | 20 | Baseline | aldex2 | 9 |
| # | disease | 7 | 0 | 0 | 1 | 1 | # | 20 | Small | paralle | 9 |
| # | sex | 4 | 0 | 0 | 1 | 1 | # | 20 | Small | paralle | 9 |
| # | time | 5 | 0 | 0 | 1 | 1 | # | 20 | Small | paralle | 9 |
| # | interaction | 4 | 0 | 0 | 1 | 1 | # | 20 | Small | paralle | 9 |
| # | disease | 7 | 3 | 0 | 0.7 | 1 | # | 20 | Small | ancom | 9 |
| # | sex | 4 | 0 | 0 | 1 | 1 | # | 20 | Small | ancom | 9 |
| # | time | 5 | 4 | 0 | 0.555555556 | 1 | # | 20 | Small | ancom | 9 |
| # | interaction | 0 | 0 | 4 | 0 | 0 | # | 20 | Small | ancom | 9 |
| # | disease | 7 | 10 | 0 | 0.411764706 | 1 | # | 20 | Small | aldex2 | 9 |
| # | sex | 4 | 1 | 0 | 0.8 | 1 | # | 20 | Small | aldex2 | 9 |
| # | time | 5 | 5 | 0 | 0.5 | 1 | # | 20 | Small | aldex2 | 9 |
| # | interaction | 0 | 0 | 4 | 0 | 0 | # | 20 | Small | aldex2 | 9 |
| # | disease | 4 | 13 | 0 | 0.235294118 | 1 | # | 10 | Baseline | paralle | 10 |
| # | sex | 2 | 9 | 0 | 0.181818182 | 1 | # | 10 | Baseline | paralle | 10 |
| # | time | 2 | 9 | 0 | 0.181818182 | 1 | # | 10 | Baseline | paralle | 10 |
| # | interaction | 2 | 10 | 0 | 0.166666667 | 1 | # | 10 | Baseline | paralle | 10 |
| # | disease | 4 | 0 | 0 | 1 | 1 | # | 10 | Baseline | ancom | 10 |
| # | sex | 2 | 0 | 0 | 1 | 1 | # | 10 | Baseline | ancom | 10 |
| # | time | 2 | 0 | 0 | 1 | 1 | # | 10 | Baseline | ancom | 10 |
| # | interaction | 0 | 0 | 2 | 0 | 0 | # | 10 | Baseline | ancom | 10 |
| # | disease | 4 | 11 | 0 | 0.266666667 | 1 | # | 10 | Baseline | aldex2 | 10 |
| # | sex | 2 | 8 | 0 | 0.2 | 1 | # | 10 | Baseline | aldex2 | 10 |
| # | time | 2 | 8 | 0 | 0.2 | 1 | # | 10 | Baseline | aldex2 | 10 |
| # | interaction | 0 | 0 | 2 | 0 | 0 | # | 10 | Baseline | aldex2 | 10 |
| # | disease | 4 | 0 | 0 | 1 | 1 | # | 10 | Small | paralle | 10 |
| # | sex | 2 | 0 | 0 | 1 | 1 | # | 10 | Small | paralle | 10 |
| # | time | 2 | 0 | 0 | 1 | 1 | # | 10 | Small | paralle | 10 |
| # | interaction | 0 | 0 | 2 | 0 | 0 | # | 10 | Small | paralle | 10 |
| # | disease | 4 | 0 | 0 | 1 | 1 | # | 10 | Small | ancom | 10 |
| # | sex | 2 | 0 | 0 | 1 | 1 | # | 10 | Small | ancom | 10 |
| # | time | 2 | 0 | 0 | 1 | 1 | # | 10 | Small | ancom | 10 |
| # | interaction | 0 | 0 | 2 | 0 | 0 | # | 10 | Small | ancom | 10 |
| # | disease | 4 | 1 | 0 | 0.8 | 1 | # | 10 | Small | aldex2 | 10 |
| # | sex | 2 | 0 | 0 | 1 | 1 | # | 10 | Small | aldex2 | 10 |
| # | time | 2 | 0 | 0 | 1 | 1 | # | 10 | Small | aldex2 | 10 |
| # | interaction | 0 | 0 | 2 | 0 | 0 | # | 10 | Small | aldex2 | 10 |
| # | disease | 4 | 8 | 0 | 0.333333333 | 1 | # | 10 | Baseline | paralle | 10 |
| # | sex | 2 | 3 | 0 | 0.4 | 1 | # | 10 | Baseline | paralle | 10 |
| # | time | 2 | 9 | 0 | 0.181818182 | 1 | # | 10 | Baseline | paralle | 10 |
| # | interaction | 2 | 8 | 0 | 0.2 | 1 | # | 10 | Baseline | paralle | 10 |
| # | disease | 4 | 1 | 0 | 0.8 | 1 | # | 10 | Baseline | ancom | 10 |
| # | sex | 2 | 0 | 0 | 1 | 1 | # | 10 | Baseline | ancom | 10 |
| # | time | 2 | 1 | 0 | 0.666666667 | 1 | # | 10 | Baseline | ancom | 10 |
| # | interaction | 0 | 0 | 2 | 0 | 0 | # | 10 | Baseline | ancom | 10 |
| # | disease | 4 | 14 | 0 | 0.222222222 | 1 | # | 10 | Baseline | aldex2 | 10 |
| # | sex | 2 | 5 | 0 | 0.285714286 | 1 | # | 10 | Baseline | aldex2 | 10 |
| # | time | 2 | 9 | 0 | 0.181818182 | 1 | # | 10 | Baseline | aldex2 | 10 |
| # | interaction | 0 | 0 | 2 | 0 | 0 | # | 10 | Baseline | aldex2 | 10 |
| # | disease | 4 | 0 | 0 | 1 | 1 | # | 10 | Small | paralle | 10 |
| # | sex | 2 | 0 | 0 | 1 | 1 | # | 10 | Small | paralle | 10 |
| # | time | 2 | 0 | 0 | 1 | 1 | # | 10 | Small | paralle | 10 |
| # | interaction | 2 | 0 | 0 | 1 | 1 | # | 10 | Small | paralle | 10 |
| # | disease | 4 | 1 | 0 | 0.8 | 1 | # | 10 | Small | ancom | 10 |
| # | sex | 2 | 0 | 0 | 1 | 1 | # | 10 | Small | ancom | 10 |
| # | time | 2 | 0 | 0 | 1 | 1 | # | 10 | Small | ancom | 10 |
| # | interaction | 0 | 0 | 2 | 0 | 0 | # | 10 | Small | ancom | 10 |
| # | disease | 4 | 1 | 0 | 0.8 | 1 | # | 10 | Small | aldex2 | 10 |
| # | sex | 2 | 0 | 0 | 1 | 1 | # | 10 | Small | aldex2 | 10 |
| # | time | 2 | 0 | 0 | 1 | 1 | # | 10 | Small | aldex2 | 10 |
| # | interaction | 0 | 0 | 2 | 0 | 0 | # | 10 | Small | aldex2 | 10 |
| # | disease | 4 | 10 | 0 | 0.285714286 | 1 | # | 10 | Baseline | paralle | 10 |
| # | sex | 2 | 5 | 0 | 0.285714286 | 1 | # | 10 | Baseline | paralle | 10 |
| # | time | 2 | 7 | 0 | 0.222222222 | 1 | # | 10 | Baseline | paralle | 10 |
| # | interaction | 2 | 9 | 0 | 0.181818182 | 1 | # | 10 | Baseline | paralle | 10 |
| # | disease | 4 | 1 | 0 | 0.8 | 1 | # | 10 | Baseline | ancom | 10 |
| # | sex | 2 | 0 | 0 | 1 | 1 | # | 10 | Baseline | ancom | 10 |
| # | time | 2 | 2 | 0 | 0.5 | 1 | # | 10 | Baseline | ancom | 10 |
| # | interaction | 0 | 0 | 2 | 0 | 0 | # | 10 | Baseline | ancom | 10 |
| # | disease | 4 | 24 | 0 | 0.142857143 | 1 | # | 10 | Baseline | aldex2 | 10 |
| # | sex | 2 | 5 | 0 | 0.285714286 | 1 | # | 10 | Baseline | aldex2 | 10 |
| # | time | 2 | 9 | 0 | 0.181818182 | 1 | # | 10 | Baseline | aldex2 | 10 |
| # | interaction | 0 | 0 | 2 | 0 | 0 | # | 10 | Baseline | aldex2 | 10 |
| # | disease | 4 | 0 | 0 | 1 | 1 | # | 10 | Small | paralle | 10 |
| # | sex | 2 | 0 | 0 | 1 | 1 | # | 10 | Small | paralle | 10 |
| # | time | 2 | 0 | 0 | 1 | 1 | # | 10 | Small | paralle | 10 |
| # | interaction | 2 | 0 | 0 | 1 | 1 | # | 10 | Small | paralle | 10 |
| # | disease | 4 | 1 | 0 | 0.8 | 1 | # | 10 | Small | ancom | 10 |
| # | sex | 2 | 0 | 0 | 1 | 1 | # | 10 | Small | ancom | 10 |
| # | time | 2 | 2 | 0 | 0.5 | 1 | # | 10 | Small | ancom | 10 |
| # | interaction | 0 | 0 | 2 | 0 | 0 | # | 10 | Small | ancom | 10 |
| # | disease | 4 | 2 | 0 | 0.666666667 | 1 | # | 10 | Small | aldex2 | 10 |
| # | sex | 2 | 0 | 0 | 1 | 1 | # | 10 | Small | aldex2 | 10 |
| # | time | 2 | 2 | 0 | 0.5 | 1 | # | 10 | Small | aldex2 | 10 |
| # | interaction | 0 | 0 | 2 | 0 | 0 | # | 10 | Small | aldex2 | 10 |
| # | disease | 4 | 6 | 0 | 0.4 | 1 | # | 10 | Baseline | paralle | 10 |
| # | sex | 2 | 5 | 0 | 0.285714286 | 1 | # | 10 | Baseline | paralle | 10 |
| # | time | 2 | 5 | 0 | 0.285714286 | 1 | # | 10 | Baseline | paralle | 10 |
| # | interaction | 2 | 4 | 0 | 0.333333333 | 1 | # | 10 | Baseline | paralle | 10 |
| # | disease | 4 | 2 | 0 | 0.666666667 | 1 | # | 10 | Baseline | ancom | 10 |
| # | sex | 2 | 0 | 0 | 1 | 1 | # | 10 | Baseline | ancom | 10 |
| # | time | 2 | 2 | 0 | 0.5 | 1 | # | 10 | Baseline | ancom | 10 |
| # | interaction | 0 | 0 | 2 | 0 | 0 | # | 10 | Baseline | ancom | 10 |
| # | disease | 4 | 32 | 0 | 0.111111111 | 1 | # | 10 | Baseline | aldex2 | 10 |
| # | sex | 2 | 4 | 0 | 0.333333333 | 1 | # | 10 | Baseline | aldex2 | 10 |
| # | time | 2 | 6 | 0 | 0.25 | 1 | # | 10 | Baseline | aldex2 | 10 |
| # | interaction | 0 | 0 | 2 | 0 | 0 | # | 10 | Baseline | aldex2 | 10 |
| # | disease | 4 | 0 | 0 | 1 | 1 | # | 10 | Small | paralle | 10 |
| # | sex | 2 | 0 | 0 | 1 | 1 | # | 10 | Small | paralle | 10 |
| # | time | 2 | 0 | 0 | 1 | 1 | # | 10 | Small | paralle | 10 |
| # | interaction | 2 | 0 | 0 | 1 | 1 | # | 10 | Small | paralle | 10 |
| # | disease | 4 | 2 | 0 | 0.666666667 | 1 | # | 10 | Small | ancom | 10 |
| # | sex | 2 | 0 | 0 | 1 | 1 | # | 10 | Small | ancom | 10 |
| # | time | 2 | 2 | 0 | 0.5 | 1 | # | 10 | Small | ancom | 10 |
| # | interaction | 0 | 0 | 2 | 0 | 0 | # | 10 | Small | ancom | 10 |
| # | disease | 4 | 2 | 0 | 0.666666667 | 1 | # | 10 | Small | aldex2 | 10 |
| # | sex | 2 | 0 | 0 | 1 | 1 | # | 10 | Small | aldex2 | 10 |
| # | time | 2 | 2 | 0 | 0.5 | 1 | # | 10 | Small | aldex2 | 10 |
| # | interaction | 0 | 0 | 2 | 0 | 0 | # | 10 | Small | aldex2 | 10 |
| # | disease | 7 | 16 | 0 | 0.304347826 | 1 | # | 20 | Baseline | paralle | 10 |
| # | sex | 4 | 12 | 0 | 0.25 | 1 | # | 20 | Baseline | paralle | 10 |
| # | time | 5 | 13 | 0 | 0.277777778 | 1 | # | 20 | Baseline | paralle | 10 |
| # | interaction | 4 | 10 | 0 | 0.285714286 | 1 | # | 20 | Baseline | paralle | 10 |
| # | disease | 7 | 2 | 0 | 0.777777778 | 1 | # | 20 | Baseline | ancom | 10 |
| # | sex | 4 | 0 | 0 | 1 | 1 | # | 20 | Baseline | ancom | 10 |
| # | time | 5 | 2 | 0 | 0.714285714 | 1 | # | 20 | Baseline | ancom | 10 |
| # | interaction | 0 | 0 | 4 | 0 | 0 | # | 20 | Baseline | ancom | 10 |
| # | disease | 7 | 22 | 0 | 0.24137931 | 1 | # | 20 | Baseline | aldex2 | 10 |
| # | sex | 4 | 7 | 0 | 0.363636364 | 1 | # | 20 | Baseline | aldex2 | 10 |
| # | time | 5 | 13 | 0 | 0.277777778 | 1 | # | 20 | Baseline | aldex2 | 10 |
| # | interaction | 0 | 0 | 4 | 0 | 0 | # | 20 | Baseline | aldex2 | 10 |
| # | disease | 7 | 0 | 0 | 1 | 1 | # | 20 | Small | paralle | 10 |
| # | sex | 4 | 3 | 0 | 0.571428571 | 1 | # | 20 | Small | paralle | 10 |
| # | time | 5 | 0 | 0 | 1 | 1 | # | 20 | Small | paralle | 10 |
| # | interaction | 2 | 1 | 2 | 0.666666667 | 0.5 | # | 20 | Small | paralle | 10 |
| # | disease | 7 | 2 | 0 | 0.777777778 | 1 | # | 20 | Small | ancom | 10 |
| # | sex | 4 | 0 | 0 | 1 | 1 | # | 20 | Small | ancom | 10 |
| # | time | 5 | 2 | 0 | 0.714285714 | 1 | # | 20 | Small | ancom | 10 |
| # | interaction | 0 | 0 | 4 | 0 | 0 | # | 20 | Small | ancom | 10 |
| # | disease | 7 | 2 | 0 | 0.777777778 | 1 | # | 20 | Small | aldex2 | 10 |
| # | sex | 4 | 0 | 0 | 1 | 1 | # | 20 | Small | aldex2 | 10 |
| # | time | 5 | 2 | 0 | 0.714285714 | 1 | # | 20 | Small | aldex2 | 10 |
| # | interaction | 0 | 0 | 4 | 0 | 0 | # | 20 | Small | aldex2 | 10 |
| # | disease | 7 | 17 | 0 | 0.291666667 | 1 | # | 20 | Baseline | paralle | 10 |
| # | sex | 4 | 9 | 0 | 0.307692308 | 1 | # | 20 | Baseline | paralle | 10 |
| # | time | 5 | 10 | 0 | 0.333333333 | 1 | # | 20 | Baseline | paralle | 10 |
| # | interaction | 4 | 10 | 0 | 0.285714286 | 1 | # | 20 | Baseline | paralle | 10 |
| # | disease | 7 | 2 | 0 | 0.777777778 | 1 | # | 20 | Baseline | ancom | 10 |
| # | sex | 4 | 0 | 0 | 1 | 1 | # | 20 | Baseline | ancom | 10 |
| # | time | 5 | 4 | 0 | 0.555555556 | 1 | # | 20 | Baseline | ancom | 10 |
| # | interaction | 0 | 0 | 4 | 0 | 0 | # | 20 | Baseline | ancom | 10 |
| # | disease | 7 | 34 | 0 | 0.170731707 | 1 | # | 20 | Baseline | aldex2 | 10 |
| # | sex | 4 | 12 | 0 | 0.25 | 1 | # | 20 | Baseline | aldex2 | 10 |
| # | time | 5 | 26 | 0 | 0.161290323 | 1 | # | 20 | Baseline | aldex2 | 10 |
| # | interaction | 0 | 0 | 4 | 0 | 0 | # | 20 | Baseline | aldex2 | 10 |
| # | disease | 7 | 0 | 0 | 1 | 1 | # | 20 | Small | paralle | 10 |
| # | sex | 4 | 1 | 0 | 0.8 | 1 | # | 20 | Small | paralle | 10 |
| # | time | 5 | 0 | 0 | 1 | 1 | # | 20 | Small | paralle | 10 |
| # | interaction | 4 | 0 | 0 | 1 | 1 | # | 20 | Small | paralle | 10 |
| # | disease | 7 | 2 | 0 | 0.777777778 | 1 | # | 20 | Small | ancom | 10 |
| # | sex | 4 | 0 | 0 | 1 | 1 | # | 20 | Small | ancom | 10 |
| # | time | 5 | 4 | 0 | 0.555555556 | 1 | # | 20 | Small | ancom | 10 |
| # | interaction | 0 | 0 | 4 | 0 | 0 | # | 20 | Small | ancom | 10 |
| # | disease | 7 | 5 | 0 | 0.583333333 | 1 | # | 20 | Small | aldex2 | 10 |
| # | sex | 4 | 1 | 0 | 0.8 | 1 | # | 20 | Small | aldex2 | 10 |
| # | time | 5 | 5 | 0 | 0.5 | 1 | # | 20 | Small | aldex2 | 10 |
| # | interaction | 0 | 0 | 4 | 0 | 0 | # | 20 | Small | aldex2 | 10 |
| # | disease | 7 | 19 | 0 | 0.269230769 | 1 | # | 20 | Baseline | paralle | 10 |
| # | sex | 4 | 9 | 0 | 0.307692308 | 1 | # | 20 | Baseline | paralle | 10 |
| # | time | 5 | 9 | 0 | 0.357142857 | 1 | # | 20 | Baseline | paralle | 10 |
| # | interaction | 4 | 12 | 0 | 0.25 | 1 | # | 20 | Baseline | paralle | 10 |
| # | disease | 7 | 2 | 0 | 0.777777778 | 1 | # | 20 | Baseline | ancom | 10 |
| # | sex | 4 | 0 | 0 | 1 | 1 | # | 20 | Baseline | ancom | 10 |
| # | time | 5 | 4 | 0 | 0.555555556 | 1 | # | 20 | Baseline | ancom | 10 |
| # | interaction | 0 | 0 | 4 | 0 | 0 | # | 20 | Baseline | ancom | 10 |
| # | disease | 7 | 68 | 0 | 0.127272727 | 1 | # | 20 | Baseline | aldex2 | 10 |
| # | sex | 4 | 21 | 0 | 0.16 | 1 | # | 20 | Baseline | aldex2 | 10 |
| # | time | 5 | 32 | 0 | 0.135135135 | 1 | # | 20 | Baseline | aldex2 | 10 |
| # | interaction | 0 | 0 | 4 | 0 | 0 | # | 20 | Baseline | aldex2 | 10 |
| # | disease | 7 | 0 | 0 | 1 | 1 | # | 20 | Small | paralle | 10 |
| # | sex | 4 | 0 | 0 | 1 | 1 | # | 20 | Small | paralle | 10 |
| # | time | 5 | 0 | 0 | 1 | 1 | # | 20 | Small | paralle | 10 |
| # | interaction | 4 | 1 | 0 | 0.8 | 1 | # | 20 | Small | paralle | 10 |
| # | disease | 7 | 2 | 0 | 0.777777778 | 1 | # | 20 | Small | ancom | 10 |
| # | sex | 4 | 0 | 0 | 1 | 1 | # | 20 | Small | ancom | 10 |
| # | time | 5 | 4 | 0 | 0.555555556 | 1 | # | 20 | Small | ancom | 10 |
| # | interaction | 0 | 0 | 4 | 0 | 0 | # | 20 | Small | ancom | 10 |
| # | disease | 7 | 5 | 0 | 0.583333333 | 1 | # | 20 | Small | aldex2 | 10 |
| # | sex | 4 | 0 | 0 | 1 | 1 | # | 20 | Small | aldex2 | 10 |
| # | time | 5 | 5 | 0 | 0.5 | 1 | # | 20 | Small | aldex2 | 10 |
| # | interaction | 0 | 0 | 4 | 0 | 0 | # | 20 | Small | aldex2 | 10 |
| # | disease | 7 | 14 | 0 | 0.333333333 | 1 | # | 20 | Baseline | paralle | 10 |
| # | sex | 4 | 7 | 0 | 0.363636364 | 1 | # | 20 | Baseline | paralle | 10 |
| # | time | 5 | 5 | 0 | 0.5 | 1 | # | 20 | Baseline | paralle | 10 |
| # | interaction | 4 | 9 | 0 | 0.307692308 | 1 | # | 20 | Baseline | paralle | 10 |
| # | disease | 7 | 4 | 0 | 0.636363636 | 1 | # | 20 | Baseline | ancom | 10 |
| # | sex | 4 | 0 | 0 | 1 | 1 | # | 20 | Baseline | ancom | 10 |
| # | time | 5 | 4 | 0 | 0.555555556 | 1 | # | 20 | Baseline | ancom | 10 |
| # | interaction | 0 | 0 | 4 | 0 | 0 | # | 20 | Baseline | ancom | 10 |
| # | disease | 7 | 56 | 0 | 0.111111111 | 1 | # | 20 | Baseline | aldex2 | 10 |
| # | sex | 4 | 21 | 0 | 0.16 | 1 | # | 20 | Baseline | aldex2 | 10 |
| # | time | 5 | 31 | 0 | 0.138888889 | 1 | # | 20 | Baseline | aldex2 | 10 |
| # | interaction | 0 | 0 | 4 | 0 | 0 | # | 20 | Baseline | aldex2 | 10 |
| # | disease | 7 | 0 | 0 | 1 | 1 | # | 20 | Small | paralle | 10 |
| # | sex | 4 | 0 | 0 | 1 | 1 | # | 20 | Small | paralle | 10 |
| # | time | 5 | 0 | 0 | 1 | 1 | # | 20 | Small | paralle | 10 |
| # | interaction | 4 | 0 | 0 | 1 | 1 | # | 20 | Small | paralle | 10 |
| # | disease | 7 | 4 | 0 | 0.636363636 | 1 | # | 20 | Small | ancom | 10 |
| # | sex | 4 | 0 | 0 | 1 | 1 | # | 20 | Small | ancom | 10 |
| # | time | 5 | 4 | 0 | 0.555555556 | 1 | # | 20 | Small | ancom | 10 |
| # | interaction | 0 | 0 | 4 | 0 | 0 | # | 20 | Small | ancom | 10 |
| # | disease | 7 | 10 | 0 | 0.411764706 | 1 | # | 20 | Small | aldex2 | 10 |
| # | sex | 4 | 1 | 0 | 0.8 | 1 | # | 20 | Small | aldex2 | 10 |
| # | time | 5 | 4 | 0 | 0.555555556 | 1 | # | 20 | Small | aldex2 | 10 |
| # | interaction | 0 | 0 | 4 | 0 | 0 | # | 20 | Small | aldex2 | 10 |
| # | disease | 4 | 5 | 0 | 0.444444444 | 1 | # | 10 | Baseline | paralle | 11 |
| # | sex | 2 | 6 | 0 | 0.25 | 1 | # | 10 | Baseline | paralle | 11 |
| # | time | 2 | 5 | 0 | 0.285714286 | 1 | # | 10 | Baseline | paralle | 11 |
| # | interaction | 2 | 4 | 0 | 0.333333333 | 1 | # | 10 | Baseline | paralle | 11 |
| # | disease | 4 | 0 | 0 | 1 | 1 | # | 10 | Baseline | ancom | 11 |
| # | sex | 2 | 0 | 0 | 1 | 1 | # | 10 | Baseline | ancom | 11 |
| # | time | 2 | 0 | 0 | 1 | 1 | # | 10 | Baseline | ancom | 11 |
| # | interaction | 0 | 0 | 2 | 0 | 0 | # | 10 | Baseline | ancom | 11 |
| # | disease | 4 | 10 | 0 | 0.285714286 | 1 | # | 10 | Baseline | aldex2 | 11 |
| # | sex | 2 | 9 | 0 | 0.181818182 | 1 | # | 10 | Baseline | aldex2 | 11 |
| # | time | 2 | 8 | 0 | 0.2 | 1 | # | 10 | Baseline | aldex2 | 11 |
| # | interaction | 0 | 0 | 2 | 0 | 0 | # | 10 | Baseline | aldex2 | 11 |
| # | disease | 4 | 0 | 0 | 1 | 1 | # | 10 | Small | paralle | 11 |
| # | sex | 2 | 1 | 0 | 0.666666667 | 1 | # | 10 | Small | paralle | 11 |
| # | time | 2 | 0 | 0 | 1 | 1 | # | 10 | Small | paralle | 11 |
| # | interaction | 2 | 0 | 0 | 1 | 1 | # | 10 | Small | paralle | 11 |
| # | disease | 4 | 0 | 0 | 1 | 1 | # | 10 | Small | ancom | 11 |
| # | sex | 2 | 0 | 0 | 1 | 1 | # | 10 | Small | ancom | 11 |
| # | time | 2 | 0 | 0 | 1 | 1 | # | 10 | Small | ancom | 11 |
| # | interaction | 0 | 0 | 2 | 0 | 0 | # | 10 | Small | ancom | 11 |
| # | disease | 4 | 0 | 0 | 1 | 1 | # | 10 | Small | aldex2 | 11 |
| # | sex | 2 | 0 | 0 | 1 | 1 | # | 10 | Small | aldex2 | 11 |
| # | time | 2 | 0 | 0 | 1 | 1 | # | 10 | Small | aldex2 | 11 |
| # | interaction | 0 | 0 | 2 | 0 | 0 | # | 10 | Small | aldex2 | 11 |
| # | disease | 4 | 5 | 0 | 0.444444444 | 1 | # | 10 | Baseline | paralle | 11 |
| # | sex | 2 | 7 | 0 | 0.222222222 | 1 | # | 10 | Baseline | paralle | 11 |
| # | time | 2 | 9 | 0 | 0.181818182 | 1 | # | 10 | Baseline | paralle | 11 |
| # | interaction | 2 | 3 | 0 | 0.4 | 1 | # | 10 | Baseline | paralle | 11 |
| # | disease | 4 | 2 | 0 | 0.666666667 | 1 | # | 10 | Baseline | ancom | 11 |
| # | sex | 2 | 0 | 0 | 1 | 1 | # | 10 | Baseline | ancom | 11 |
| # | time | 2 | 1 | 0 | 0.666666667 | 1 | # | 10 | Baseline | ancom | 11 |
| # | interaction | 0 | 0 | 2 | 0 | 0 | # | 10 | Baseline | ancom | 11 |
| # | disease | 4 | 19 | 0 | 0.173913043 | 1 | # | 10 | Baseline | aldex2 | 11 |
| # | sex | 2 | 6 | 0 | 0.25 | 1 | # | 10 | Baseline | aldex2 | 11 |
| # | time | 2 | 5 | 0 | 0.285714286 | 1 | # | 10 | Baseline | aldex2 | 11 |
| # | interaction | 0 | 0 | 2 | 0 | 0 | # | 10 | Baseline | aldex2 | 11 |
| # | disease | 4 | 0 | 0 | 1 | 1 | # | 10 | Small | paralle | 11 |
| # | sex | 2 | 0 | 0 | 1 | 1 | # | 10 | Small | paralle | 11 |
| # | time | 2 | 0 | 0 | 1 | 1 | # | 10 | Small | paralle | 11 |
| # | interaction | 2 | 0 | 0 | 1 | 1 | # | 10 | Small | paralle | 11 |
| # | disease | 4 | 1 | 0 | 0.8 | 1 | # | 10 | Small | ancom | 11 |
| # | sex | 2 | 0 | 0 | 1 | 1 | # | 10 | Small | ancom | 11 |
| # | time | 2 | 1 | 0 | 0.666666667 | 1 | # | 10 | Small | ancom | 11 |
| # | interaction | 0 | 0 | 2 | 0 | 0 | # | 10 | Small | ancom | 11 |
| # | disease | 4 | 1 | 0 | 0.8 | 1 | # | 10 | Small | aldex2 | 11 |
| # | sex | 2 | 0 | 0 | 1 | 1 | # | 10 | Small | aldex2 | 11 |
| # | time | 2 | 1 | 0 | 0.666666667 | 1 | # | 10 | Small | aldex2 | 11 |
| # | interaction | 0 | 0 | 2 | 0 | 0 | # | 10 | Small | aldex2 | 11 |
| # | disease | 4 | 5 | 0 | 0.444444444 | 1 | # | 10 | Baseline | paralle | 11 |
| # | sex | 2 | 8 | 0 | 0.2 | 1 | # | 10 | Baseline | paralle | 11 |
| # | time | 2 | 4 | 0 | 0.333333333 | 1 | # | 10 | Baseline | paralle | 11 |
| # | interaction | 2 | 4 | 0 | 0.333333333 | 1 | # | 10 | Baseline | paralle | 11 |
| # | disease | 4 | 2 | 0 | 0.666666667 | 1 | # | 10 | Baseline | ancom | 11 |
| # | sex | 2 | 0 | 0 | 1 | 1 | # | 10 | Baseline | ancom | 11 |
| # | time | 2 | 2 | 0 | 0.5 | 1 | # | 10 | Baseline | ancom | 11 |
| # | interaction | 0 | 0 | 2 | 0 | 0 | # | 10 | Baseline | ancom | 11 |
| # | disease | 4 | 23 | 0 | 0.148148148 | 1 | # | 10 | Baseline | aldex2 | 11 |
| # | sex | 2 | 7 | 0 | 0.222222222 | 1 | # | 10 | Baseline | aldex2 | 11 |
| # | time | 2 | 7 | 0 | 0.222222222 | 1 | # | 10 | Baseline | aldex2 | 11 |
| # | interaction | 0 | 0 | 2 | 0 | 0 | # | 10 | Baseline | aldex2 | 11 |
| # | disease | 4 | 0 | 0 | 1 | 1 | # | 10 | Small | paralle | 11 |
| # | sex | 2 | 0 | 0 | 1 | 1 | # | 10 | Small | paralle | 11 |
| # | time | 2 | 0 | 0 | 1 | 1 | # | 10 | Small | paralle | 11 |
| # | interaction | 2 | 0 | 0 | 1 | 1 | # | 10 | Small | paralle | 11 |
| # | disease | 4 | 2 | 0 | 0.666666667 | 1 | # | 10 | Small | ancom | 11 |
| # | sex | 2 | 0 | 0 | 1 | 1 | # | 10 | Small | ancom | 11 |
| # | time | 2 | 2 | 0 | 0.5 | 1 | # | 10 | Small | ancom | 11 |
| # | interaction | 0 | 0 | 2 | 0 | 0 | # | 10 | Small | ancom | 11 |
| # | disease | 4 | 3 | 0 | 0.571428571 | 1 | # | 10 | Small | aldex2 | 11 |
| # | sex | 2 | 0 | 0 | 1 | 1 | # | 10 | Small | aldex2 | 11 |
| # | time | 2 | 2 | 0 | 0.5 | 1 | # | 10 | Small | aldex2 | 11 |
| # | interaction | 0 | 0 | 2 | 0 | 0 | # | 10 | Small | aldex2 | 11 |
| # | disease | 4 | 4 | 0 | 0.5 | 1 | # | 10 | Baseline | paralle | 11 |
| # | sex | 2 | 3 | 0 | 0.4 | 1 | # | 10 | Baseline | paralle | 11 |
| # | time | 2 | 3 | 0 | 0.4 | 1 | # | 10 | Baseline | paralle | 11 |
| # | interaction | 2 | 3 | 0 | 0.4 | 1 | # | 10 | Baseline | paralle | 11 |
| # | disease | 4 | 2 | 0 | 0.666666667 | 1 | # | 10 | Baseline | ancom | 11 |
| # | sex | 2 | 0 | 0 | 1 | 1 | # | 10 | Baseline | ancom | 11 |
| # | time | 2 | 2 | 0 | 0.5 | 1 | # | 10 | Baseline | ancom | 11 |
| # | interaction | 0 | 0 | 2 | 0 | 0 | # | 10 | Baseline | ancom | 11 |
| # | disease | 4 | 22 | 0 | 0.153846154 | 1 | # | 10 | Baseline | aldex2 | 11 |
| # | sex | 2 | 9 | 0 | 0.181818182 | 1 | # | 10 | Baseline | aldex2 | 11 |
| # | time | 2 | 10 | 0 | 0.166666667 | 1 | # | 10 | Baseline | aldex2 | 11 |
| # | interaction | 0 | 0 | 2 | 0 | 0 | # | 10 | Baseline | aldex2 | 11 |
| # | disease | 4 | 0 | 0 | 1 | 1 | # | 10 | Small | paralle | 11 |
| # | sex | 2 | 0 | 0 | 1 | 1 | # | 10 | Small | paralle | 11 |
| # | time | 2 | 0 | 0 | 1 | 1 | # | 10 | Small | paralle | 11 |
| # | interaction | 2 | 0 | 0 | 1 | 1 | # | 10 | Small | paralle | 11 |
| # | disease | 4 | 2 | 0 | 0.666666667 | 1 | # | 10 | Small | ancom | 11 |
| # | sex | 2 | 0 | 0 | 1 | 1 | # | 10 | Small | ancom | 11 |
| # | time | 2 | 2 | 0 | 0.5 | 1 | # | 10 | Small | ancom | 11 |
| # | interaction | 0 | 0 | 2 | 0 | 0 | # | 10 | Small | ancom | 11 |
| # | disease | 4 | 2 | 0 | 0.666666667 | 1 | # | 10 | Small | aldex2 | 11 |
| # | sex | 2 | 0 | 0 | 1 | 1 | # | 10 | Small | aldex2 | 11 |
| # | time | 2 | 1 | 0 | 0.666666667 | 1 | # | 10 | Small | aldex2 | 11 |
| # | interaction | 0 | 0 | 2 | 0 | 0 | # | 10 | Small | aldex2 | 11 |
| # | disease | 7 | 12 | 0 | 0.368421053 | 1 | # | 20 | Baseline | paralle | 11 |
| # | sex | 4 | 6 | 0 | 0.4 | 1 | # | 20 | Baseline | paralle | 11 |
| # | time | 5 | 9 | 0 | 0.357142857 | 1 | # | 20 | Baseline | paralle | 11 |

| | | | | | | | | | | | |
|---|---|---|---|---|---|---|---|---|---|---|---|
| # | interaction | 4 | 11 | 0 | 0.266666667 | 1 | # | 20 | Baseline | paralle | 11 |
| # | disease | 7 | 0 | 0 | 1 | 1 | # | 20 | Baseline | ancom | 11 |
| # | sex | 4 | 0 | 0 | 1 | 1 | # | 20 | Baseline | ancom | 11 |
| # | time | 5 | 2 | 0 | 0.714285714 | 1 | # | 20 | Baseline | ancom | 11 |
| # | interaction | 0 | 0 | 4 | 0 | 0 | # | 20 | Baseline | ancom | 11 |
| # | disease | 7 | 23 | 0 | 0.233333333 | 1 | # | 20 | Baseline | aldex2 | 11 |
| # | sex | 4 | 2 | 0 | 0.666666667 | 1 | # | 20 | Baseline | aldex2 | 11 |
| # | time | 5 | 14 | 0 | 0.263157895 | 1 | # | 20 | Baseline | aldex2 | 11 |
| # | interaction | 0 | 0 | 4 | 0 | 0 | # | 20 | Baseline | aldex2 | 11 |
| # | disease | 7 | 1 | 0 | 0.875 | 1 | # | 20 | Small | paralle | 11 |
| # | sex | 4 | 0 | 0 | 1 | 1 | # | 20 | Small | paralle | 11 |
| # | time | 5 | 0 | 0 | 1 | 1 | # | 20 | Small | paralle | 11 |
| # | interaction | 2 | 1 | 2 | 0.666666667 | 0.5 | # | 20 | Small | paralle | 11 |
| # | disease | 7 | 0 | 0 | 1 | 1 | # | 20 | Small | ancom | 11 |
| # | sex | 4 | 0 | 0 | 1 | 1 | # | 20 | Small | ancom | 11 |
| # | time | 5 | 2 | 0 | 0.714285714 | 1 | # | 20 | Small | ancom | 11 |
| # | interaction | 0 | 0 | 4 | 0 | 0 | # | 20 | Small | ancom | 11 |
| # | disease | 7 | 1 | 0 | 0.875 | 1 | # | 20 | Small | aldex2 | 11 |
| # | sex | 4 | 0 | 0 | 1 | 1 | # | 20 | Small | aldex2 | 11 |
| # | time | 5 | 2 | 0 | 0.714285714 | 1 | # | 20 | Small | aldex2 | 11 |
| # | interaction | 0 | 0 | 4 | 0 | 0 | # | 20 | Small | aldex2 | 11 |
| # | disease | 7 | 15 | 0 | 0.318181818 | 1 | # | 20 | Baseline | paralle | 11 |
| # | sex | 4 | 3 | 0 | 0.571428571 | 1 | # | 20 | Baseline | paralle | 11 |
| # | time | 5 | 8 | 0 | 0.384615385 | 1 | # | 20 | Baseline | paralle | 11 |
| # | interaction | 4 | 4 | 0 | 0.5 | 1 | # | 20 | Baseline | paralle | 11 |
| # | disease | 7 | 3 | 0 | 0.7 | 1 | # | 20 | Baseline | ancom | 11 |
| # | sex | 4 | 0 | 0 | 1 | 1 | # | 20 | Baseline | ancom | 11 |
| # | time | 5 | 3 | 0 | 0.625 | 1 | # | 20 | Baseline | ancom | 11 |
| # | interaction | 0 | 0 | 4 | 0 | 0 | # | 20 | Baseline | ancom | 11 |
| # | disease | 7 | 31 | 0 | 0.184210526 | 1 | # | 20 | Baseline | aldex2 | 11 |
| # | sex | 4 | 9 | 0 | 0.307692308 | 1 | # | 20 | Baseline | aldex2 | 11 |
| # | time | 5 | 18 | 0 | 0.217391304 | 1 | # | 20 | Baseline | aldex2 | 11 |
| # | interaction | 0 | 0 | 4 | 0 | 0 | # | 20 | Baseline | aldex2 | 11 |
| # | disease | 7 | 0 | 0 | 1 | 1 | # | 20 | Small | paralle | 11 |
| # | sex | 4 | 0 | 0 | 1 | 1 | # | 20 | Small | paralle | 11 |
| # | time | 5 | 0 | 0 | 1 | 1 | # | 20 | Small | paralle | 11 |
| # | interaction | 3 | 0 | 1 | 1 | 0.75 | # | 20 | Small | paralle | 11 |
| # | disease | 7 | 3 | 0 | 0.7 | 1 | # | 20 | Small | ancom | 11 |
| # | sex | 4 | 0 | 0 | 1 | 1 | # | 20 | Small | ancom | 11 |
| # | time | 5 | 3 | 0 | 0.625 | 1 | # | 20 | Small | ancom | 11 |
| # | interaction | 0 | 0 | 4 | 0 | 0 | # | 20 | Small | ancom | 11 |
| # | disease | 7 | 4 | 0 | 0.636363636 | 1 | # | 20 | Small | aldex2 | 11 |
| # | sex | 4 | 0 | 0 | 1 | 1 | # | 20 | Small | aldex2 | 11 |
| # | time | 5 | 2 | 0 | 0.714285714 | 1 | # | 20 | Small | aldex2 | 11 |
| # | interaction | 0 | 0 | 4 | 0 | 0 | # | 20 | Small | aldex2 | 11 |
| # | disease | 7 | 11 | 0 | 0.388888889 | 1 | # | 20 | Baseline | paralle | 11 |
| # | sex | 4 | 8 | 0 | 0.333333333 | 1 | # | 20 | Baseline | paralle | 11 |
| # | time | 5 | 4 | 0 | 0.555555556 | 1 | # | 20 | Baseline | paralle | 11 |
| # | interaction | 4 | 3 | 0 | 0.571428571 | 1 | # | 20 | Baseline | paralle | 11 |
| # | disease | 7 | 3 | 0 | 0.7 | 1 | # | 20 | Baseline | ancom | 11 |
| # | sex | 4 | 0 | 0 | 1 | 1 | # | 20 | Baseline | ancom | 11 |
| # | time | 5 | 4 | 0 | 0.555555556 | 1 | # | 20 | Baseline | ancom | 11 |
| # | interaction | 0 | 0 | 4 | 0 | 0 | # | 20 | Baseline | ancom | 11 |
| # | disease | 7 | 43 | 0 | 0.14 | 1 | # | 20 | Baseline | aldex2 | 11 |
| # | sex | 4 | 18 | 0 | 0.181818182 | 1 | # | 20 | Baseline | aldex2 | 11 |
| # | time | 5 | 30 | 0 | 0.142857143 | 1 | # | 20 | Baseline | aldex2 | 11 |
| # | interaction | 0 | 0 | 4 | 0 | 0 | # | 20 | Baseline | aldex2 | 11 |
| # | disease | 7 | 0 | 0 | 1 | 1 | # | 20 | Small | paralle | 11 |
| # | sex | 4 | 0 | 0 | 1 | 1 | # | 20 | Small | paralle | 11 |
| # | time | 5 | 0 | 0 | 1 | 1 | # | 20 | Small | paralle | 11 |
| # | interaction | 4 | 0 | 0 | 1 | 1 | # | 20 | Small | paralle | 11 |
| # | disease | 7 | 3 | 0 | 0.7 | 1 | # | 20 | Small | ancom | 11 |
| # | sex | 4 | 0 | 0 | 1 | 1 | # | 20 | Small | ancom | 11 |
| # | time | 5 | 4 | 0 | 0.555555556 | 1 | # | 20 | Small | ancom | 11 |
| # | interaction | 0 | 0 | 4 | 0 | 0 | # | 20 | Small | ancom | 11 |
| # | disease | 7 | 4 | 0 | 0.636363636 | 1 | # | 20 | Small | aldex2 | 11 |
| # | sex | 4 | 0 | 0 | 1 | 1 | # | 20 | Small | aldex2 | 11 |
| # | time | 5 | 5 | 0 | 0.5 | 1 | # | 20 | Small | aldex2 | 11 |
| # | interaction | 0 | 0 | 4 | 0 | 0 | # | 20 | Small | aldex2 | 11 |
| # | disease | 7 | 14 | 0 | 0.333333333 | 1 | # | 20 | Baseline | paralle | 11 |
| # | sex | 4 | 4 | 0 | 0.5 | 1 | # | 20 | Baseline | paralle | 11 |
| # | time | 5 | 8 | 0 | 0.384615385 | 1 | # | 20 | Baseline | paralle | 11 |
| # | interaction | 4 | 2 | 0 | 0.666666667 | 1 | # | 20 | Baseline | paralle | 11 |
| # | disease | 7 | 4 | 0 | 0.636363636 | 1 | # | 20 | Baseline | ancom | 11 |
| # | sex | 4 | 0 | 0 | 1 | 1 | # | 20 | Baseline | ancom | 11 |
| # | time | 5 | 4 | 0 | 0.555555556 | 1 | # | 20 | Baseline | ancom | 11 |
| # | interaction | 0 | 0 | 4 | 0 | 0 | # | 20 | Baseline | ancom | 11 |
| # | disease | 7 | 61 | 0 | 0.102941176 | 1 | # | 20 | Baseline | aldex2 | 11 |
| # | sex | 4 | 16 | 0 | 0.2 | 1 | # | 20 | Baseline | aldex2 | 11 |
| # | time | 5 | 37 | 0 | 0.119047619 | 1 | # | 20 | Baseline | aldex2 | 11 |
| # | interaction | 0 | 0 | 4 | 0 | 0 | # | 20 | Baseline | aldex2 | 11 |
| # | disease | 7 | 1 | 0 | 0.875 | 1 | # | 20 | Small | paralle | 11 |
| # | sex | 4 | 0 | 0 | 1 | 1 | # | 20 | Small | paralle | 11 |
| # | time | 5 | 0 | 0 | 1 | 1 | # | 20 | Small | paralle | 11 |
| # | interaction | 4 | 0 | 0 | 1 | 1 | # | 20 | Small | paralle | 11 |
| # | disease | 7 | 3 | 0 | 0.7 | 1 | # | 20 | Small | ancom | 11 |
| # | sex | 4 | 0 | 0 | 1 | 1 | # | 20 | Small | ancom | 11 |
| # | time | 5 | 4 | 0 | 0.555555556 | 1 | # | 20 | Small | ancom | 11 |
| # | interaction | 0 | 0 | 4 | 0 | 0 | # | 20 | Small | ancom | 11 |
| # | disease | 7 | 7 | 0 | 0.5 | 1 | # | 20 | Small | aldex2 | 11 |
| # | sex | 4 | 0 | 0 | 1 | 1 | # | 20 | Small | aldex2 | 11 |
| # | time | 5 | 7 | 0 | 0.416666667 | 1 | # | 20 | Small | aldex2 | 11 |
| # | interaction | 0 | 0 | 4 | 0 | 0 | # | 20 | Small | aldex2 | 11 |
| # | disease | 4 | 5 | 0 | 0.444444444 | 1 | # | 10 | Baseline | paralle | 12 |
| # | sex | 2 | 9 | 0 | 0.181818182 | 1 | # | 10 | Baseline | paralle | 12 |
| # | time | 2 | 5 | 0 | 0.285714286 | 1 | # | 10 | Baseline | paralle | 12 |
| # | interaction | 2 | 5 | 0 | 0.285714286 | 1 | # | 10 | Baseline | paralle | 12 |
| # | disease | 4 | 0 | 0 | 1 | 1 | # | 10 | Baseline | ancom | 12 |
| # | sex | 2 | 0 | 0 | 1 | 1 | # | 10 | Baseline | ancom | 12 |
| # | time | 2 | 0 | 0 | 1 | 1 | # | 10 | Baseline | ancom | 12 |
| # | interaction | 0 | 0 | 2 | 0 | 0 | # | 10 | Baseline | ancom | 12 |
| # | disease | 4 | 13 | 0 | 0.235294118 | 1 | # | 10 | Baseline | aldex2 | 12 |
| # | sex | 2 | 5 | 0 | 0.285714286 | 1 | # | 10 | Baseline | aldex2 | 12 |
| # | time | 2 | 12 | 0 | 0.142857143 | 1 | # | 10 | Baseline | aldex2 | 12 |
| # | interaction | 0 | 0 | 2 | 0 | 0 | # | 10 | Baseline | aldex2 | 12 |
| # | disease | 4 | 0 | 0 | 1 | 1 | # | 10 | Small | paralle | 12 |
| # | sex | 2 | 0 | 0 | 1 | 1 | # | 10 | Small | paralle | 12 |
| # | time | 2 | 0 | 0 | 1 | 1 | # | 10 | Small | paralle | 12 |
| # | interaction | 1 | 0 | 1 | 1 | 0.5 | # | 10 | Small | paralle | 12 |
| # | disease | 4 | 0 | 0 | 1 | 1 | # | 10 | Small | ancom | 12 |
| # | sex | 2 | 0 | 0 | 1 | 1 | # | 10 | Small | ancom | 12 |
| # | time | 2 | 0 | 0 | 1 | 1 | # | 10 | Small | ancom | 12 |
| # | interaction | 0 | 0 | 2 | 0 | 0 | # | 10 | Small | ancom | 12 |
| # | disease | 4 | 0 | 0 | 1 | 1 | # | 10 | Small | aldex2 | 12 |
| # | sex | 2 | 1 | 0 | 0.666666667 | 1 | # | 10 | Small | aldex2 | 12 |
| # | time | 2 | 1 | 0 | 0.666666667 | 1 | # | 10 | Small | aldex2 | 12 |
| # | interaction | 0 | 0 | 2 | 0 | 0 | # | 10 | Small | aldex2 | 12 |
| # | disease | 4 | 6 | 0 | 0.4 | 1 | # | 10 | Baseline | paralle | 12 |
| # | sex | 2 | 4 | 0 | 0.333333333 | 1 | # | 10 | Baseline | paralle | 12 |
| # | time | 2 | 4 | 0 | 0.333333333 | 1 | # | 10 | Baseline | paralle | 12 |
| # | interaction | 2 | 7 | 0 | 0.222222222 | 1 | # | 10 | Baseline | paralle | 12 |
| # | disease | 4 | 2 | 0 | 0.666666667 | 1 | # | 10 | Baseline | ancom | 12 |
| # | sex | 2 | 0 | 0 | 1 | 1 | # | 10 | Baseline | ancom | 12 |
| # | time | 2 | 2 | 0 | 0.5 | 1 | # | 10 | Baseline | ancom | 12 |
| # | interaction | 0 | 0 | 2 | 0 | 0 | # | 10 | Baseline | ancom | 12 |
| # | disease | 4 | 19 | 0 | 0.173913043 | 1 | # | 10 | Baseline | aldex2 | 12 |
| # | sex | 2 | 3 | 0 | 0.4 | 1 | # | 10 | Baseline | aldex2 | 12 |
| # | time | 2 | 11 | 0 | 0.153846154 | 1 | # | 10 | Baseline | aldex2 | 12 |
| # | interaction | 0 | 0 | 2 | 0 | 0 | # | 10 | Baseline | aldex2 | 12 |
| # | disease | 4 | 0 | 0 | 1 | 1 | # | 10 | Small | paralle | 12 |
| # | sex | 2 | 1 | 0 | 0.666666667 | 1 | # | 10 | Small | paralle | 12 |
| # | time | 2 | 0 | 0 | 1 | 1 | # | 10 | Small | paralle | 12 |
| # | interaction | 2 | 0 | 0 | 1 | 1 | # | 10 | Small | paralle | 12 |
| # | disease | 4 | 2 | 0 | 0.666666667 | 1 | # | 10 | Small | ancom | 12 |
| # | sex | 2 | 0 | 0 | 1 | 1 | # | 10 | Small | ancom | 12 |
| # | time | 2 | 2 | 0 | 0.5 | 1 | # | 10 | Small | ancom | 12 |
| # | interaction | 0 | 0 | 2 | 0 | 0 | # | 10 | Small | ancom | 12 |
| # | disease | 4 | 4 | 0 | 0.5 | 1 | # | 10 | Small | aldex2 | 12 |
| # | sex | 2 | 1 | 0 | 0.666666667 | 1 | # | 10 | Small | aldex2 | 12 |
| # | time | 2 | 2 | 0 | 0.5 | 1 | # | 10 | Small | aldex2 | 12 |
| # | interaction | 0 | 0 | 2 | 0 | 0 | # | 10 | Small | aldex2 | 12 |
| # | disease | 4 | 6 | 0 | 0.4 | 1 | # | 10 | Baseline | paralle | 12 |
| # | sex | 2 | 8 | 0 | 0.2 | 1 | # | 10 | Baseline | paralle | 12 |
| # | time | 2 | 3 | 0 | 0.4 | 1 | # | 10 | Baseline | paralle | 12 |
| # | interaction | 2 | 5 | 0 | 0.285714286 | 1 | # | 10 | Baseline | paralle | 12 |
| # | disease | 4 | 3 | 0 | 0.571428571 | 1 | # | 10 | Baseline | ancom | 12 |
| # | sex | 2 | 0 | 0 | 1 | 1 | # | 10 | Baseline | ancom | 12 |
| # | time | 2 | 2 | 0 | 0.5 | 1 | # | 10 | Baseline | ancom | 12 |
| # | interaction | 0 | 0 | 2 | 0 | 0 | # | 10 | Baseline | ancom | 12 |
| # | disease | 4 | 19 | 0 | 0.173913043 | 1 | # | 10 | Baseline | aldex2 | 12 |
| # | sex | 2 | 3 | 0 | 0.4 | 1 | # | 10 | Baseline | aldex2 | 12 |
| # | time | 2 | 10 | 0 | 0.166666667 | 1 | # | 10 | Baseline | aldex2 | 12 |
| # | interaction | 0 | 0 | 2 | 0 | 0 | # | 10 | Baseline | aldex2 | 12 |
| # | disease | 4 | 1 | 0 | 0.8 | 1 | # | 10 | Small | paralle | 12 |
| # | sex | 2 | 0 | 0 | 1 | 1 | # | 10 | Small | paralle | 12 |
| # | time | 2 | 0 | 0 | 1 | 1 | # | 10 | Small | paralle | 12 |
| # | interaction | 2 | 0 | 0 | 1 | 1 | # | 10 | Small | paralle | 12 |
| # | disease | 4 | 2 | 0 | 0.666666667 | 1 | # | 10 | Small | ancom | 12 |
| # | sex | 2 | 0 | 0 | 1 | 1 | # | 10 | Small | ancom | 12 |
| # | time | 2 | 2 | 0 | 0.5 | 1 | # | 10 | Small | ancom | 12 |
| # | interaction | 0 | 0 | 2 | 0 | 0 | # | 10 | Small | ancom | 12 |
| # | disease | 4 | 4 | 0 | 0.5 | 1 | # | 10 | Small | aldex2 | 12 |
| # | sex | 2 | 1 | 0 | 0.666666667 | 1 | # | 10 | Small | aldex2 | 12 |
| # | time | 2 | 2 | 0 | 0.5 | 1 | # | 10 | Small | aldex2 | 12 |
| # | interaction | 0 | 0 | 2 | 0 | 0 | # | 10 | Small | aldex2 | 12 |
| # | disease | 4 | 5 | 0 | 0.444444444 | 1 | # | 10 | Baseline | paralle | 12 |
| # | sex | 2 | 6 | 0 | 0.25 | 1 | # | 10 | Baseline | paralle | 12 |
| # | time | 2 | 3 | 0 | 0.4 | 1 | # | 10 | Baseline | paralle | 12 |
| # | interaction | 2 | 3 | 0 | 0.4 | 1 | # | 10 | Baseline | paralle | 12 |
| # | disease | 4 | 2 | 0 | 0.666666667 | 1 | # | 10 | Baseline | ancom | 12 |
| # | sex | 2 | 0 | 0 | 1 | 1 | # | 10 | Baseline | ancom | 12 |
| # | time | 2 | 2 | 0 | 0.5 | 1 | # | 10 | Baseline | ancom | 12 |
| # | interaction | 0 | 0 | 2 | 0 | 0 | # | 10 | Baseline | ancom | 12 |
| # | disease | 4 | 24 | 0 | 0.142857143 | 1 | # | 10 | Baseline | aldex2 | 12 |
| # | sex | 2 | 4 | 0 | 0.333333333 | 1 | # | 10 | Baseline | aldex2 | 12 |
| # | time | 2 | 8 | 0 | 0.2 | 1 | # | 10 | Baseline | aldex2 | 12 |
| # | interaction | 0 | 0 | 2 | 0 | 0 | # | 10 | Baseline | aldex2 | 12 |
| # | disease | 4 | 1 | 0 | 0.8 | 1 | # | 10 | Small | paralle | 12 |
| # | sex | 2 | 0 | 0 | 1 | 1 | # | 10 | Small | paralle | 12 |
| # | time | 2 | 0 | 0 | 1 | 1 | # | 10 | Small | paralle | 12 |
| # | interaction | 2 | 0 | 0 | 1 | 1 | # | 10 | Small | paralle | 12 |
| # | disease | 4 | 2 | 0 | 0.666666667 | 1 | # | 10 | Small | ancom | 12 |
| # | sex | 2 | 0 | 0 | 1 | 1 | # | 10 | Small | ancom | 12 |
| # | time | 2 | 2 | 0 | 0.5 | 1 | # | 10 | Small | ancom | 12 |
| # | interaction | 0 | 0 | 2 | 0 | 0 | # | 10 | Small | ancom | 12 |
| # | disease | 4 | 4 | 0 | 0.5 | 1 | # | 10 | Small | aldex2 | 12 |
| # | sex | 2 | 1 | 0 | 0.666666667 | 1 | # | 10 | Small | aldex2 | 12 |
| # | time | 2 | 2 | 0 | 0.5 | 1 | # | 10 | Small | aldex2 | 12 |
| # | interaction | 0 | 0 | 2 | 0 | 0 | # | 10 | Small | aldex2 | 12 |
| # | disease | 7 | 9 | 0 | 0.4375 | 1 | # | 20 | Baseline | paralle | 12 |
| # | sex | 4 | 5 | 0 | 0.444444444 | 1 | # | 20 | Baseline | paralle | 12 |
| # | time | 5 | 9 | 0 | 0.357142857 | 1 | # | 20 | Baseline | paralle | 12 |
| # | interaction | 4 | 11 | 0 | 0.266666667 | 1 | # | 20 | Baseline | paralle | 12 |
| # | disease | 7 | 3 | 0 | 0.7 | 1 | # | 20 | Baseline | ancom | 12 |
| # | sex | 4 | 0 | 0 | 1 | 1 | # | 20 | Baseline | ancom | 12 |
| # | time | 5 | 2 | 0 | 0.714285714 | 1 | # | 20 | Baseline | ancom | 12 |
| # | interaction | 0 | 0 | 4 | 0 | 0 | # | 20 | Baseline | ancom | 12 |
| # | disease | 7 | 19 | 0 | 0.269230769 | 1 | # | 20 | Baseline | aldex2 | 12 |
| # | sex | 4 | 10 | 0 | 0.285714286 | 1 | # | 20 | Baseline | aldex2 | 12 |
| # | time | 5 | 16 | 0 | 0.238095238 | 1 | # | 20 | Baseline | aldex2 | 12 |
| # | interaction | 0 | 0 | 4 | 0 | 0 | # | 20 | Baseline | aldex2 | 12 |
| # | disease | 7 | 3 | 0 | 0.7 | 1 | # | 20 | Small | paralle | 12 |
| # | sex | 4 | 1 | 0 | 0.8 | 1 | # | 20 | Small | paralle | 12 |
| # | time | 5 | 0 | 0 | 1 | 1 | # | 20 | Small | paralle | 12 |
| # | interaction | 0 | 1 | 4 | 0 | 0 | # | 20 | Small | paralle | 12 |
| # | disease | 7 | 2 | 0 | 0.777777778 | 1 | # | 20 | Small | ancom | 12 |
| # | sex | 4 | 0 | 0 | 1 | 1 | # | 20 | Small | ancom | 12 |
| # | time | 5 | 1 | 0 | 0.833333333 | 1 | # | 20 | Small | ancom | 12 |
| # | interaction | 0 | 0 | 4 | 0 | 0 | # | 20 | Small | ancom | 12 |
| # | disease | 7 | 1 | 0 | 0.875 | 1 | # | 20 | Small | aldex2 | 12 |
| # | sex | 4 | 1 | 0 | 0.8 | 1 | # | 20 | Small | aldex2 | 12 |
| # | time | 5 | 1 | 0 | 0.833333333 | 1 | # | 20 | Small | aldex2 | 12 |
| # | interaction | 0 | 0 | 4 | 0 | 0 | # | 20 | Small | aldex2 | 12 |
| # | disease | 7 | 8 | 0 | 0.466666667 | 1 | # | 20 | Baseline | paralle | 12 |
| # | sex | 4 | 6 | 0 | 0.4 | 1 | # | 20 | Baseline | paralle | 12 |
| # | time | 5 | 9 | 0 | 0.357142857 | 1 | # | 20 | Baseline | paralle | 12 |
| # | interaction | 4 | 7 | 0 | 0.363636364 | 1 | # | 20 | Baseline | paralle | 12 |
| # | disease | 7 | 4 | 0 | 0.636363636 | 1 | # | 20 | Baseline | ancom | 12 |
| # | sex | 4 | 0 | 0 | 1 | 1 | # | 20 | Baseline | ancom | 12 |
| # | time | 5 | 4 | 0 | 0.555555556 | 1 | # | 20 | Baseline | ancom | 12 |
| # | interaction | 0 | 0 | 4 | 0 | 0 | # | 20 | Baseline | ancom | 12 |
| # | disease | 7 | 36 | 0 | 0.162790698 | 1 | # | 20 | Baseline | aldex2 | 12 |
| # | sex | 4 | 14 | 0 | 0.222222222 | 1 | # | 20 | Baseline | aldex2 | 12 |
| # | time | 5 | 23 | 0 | 0.178571429 | 1 | # | 20 | Baseline | aldex2 | 12 |
| # | interaction | 0 | 0 | 4 | 0 | 0 | # | 20 | Baseline | aldex2 | 12 |
| # | disease | 7 | 0 | 0 | 1 | 1 | # | 20 | Small | paralle | 12 |
| # | sex | 4 | 0 | 0 | 1 | 1 | # | 20 | Small | paralle | 12 |
| # | time | 5 | 0 | 0 | 1 | 1 | # | 20 | Small | paralle | 12 |
| # | interaction | 2 | 0 | 2 | 1 | 0.5 | # | 20 | Small | paralle | 12 |
| # | disease | 7 | 4 | 0 | 0.636363636 | 1 | # | 20 | Small | ancom | 12 |
| # | sex | 4 | 0 | 0 | 1 | 1 | # | 20 | Small | ancom | 12 |
| # | time | 5 | 3 | 0 | 0.625 | 1 | # | 20 | Small | ancom | 12 |
| # | interaction | 0 | 0 | 4 | 0 | 0 | # | 20 | Small | ancom | 12 |
| # | disease | 7 | 4 | 0 | 0.636363636 | 1 | # | 20 | Small | aldex2 | 12 |
| # | sex | 4 | 0 | 0 | 1 | 1 | # | 20 | Small | aldex2 | 12 |
| # | time | 5 | 6 | 0 | 0.454545455 | 1 | # | 20 | Small | aldex2 | 12 |
| # | interaction | 0 | 0 | 4 | 0 | 0 | # | 20 | Small | aldex2 | 12 |
| # | disease | 7 | 11 | 0 | 0.388888889 | 1 | # | 20 | Baseline | paralle | 12 |
| # | sex | 4 | 8 | 0 | 0.333333333 | 1 | # | 20 | Baseline | paralle | 12 |
| # | time | 5 | 7 | 0 | 0.416666667 | 1 | # | 20 | Baseline | paralle | 12 |
| # | interaction | 4 | 4 | 0 | 0.5 | 1 | # | 20 | Baseline | paralle | 12 |
| # | disease | 7 | 4 | 0 | 0.636363636 | 1 | # | 20 | Baseline | ancom | 12 |
| # | sex | 4 | 0 | 0 | 1 | 1 | # | 20 | Baseline | ancom | 12 |
| # | time | 5 | 4 | 0 | 0.555555556 | 1 | # | 20 | Baseline | ancom | 12 |
| # | interaction | 0 | 0 | 4 | 0 | 0 | # | 20 | Baseline | ancom | 12 |
| # | disease | 7 | 47 | 0 | 0.12962963 | 1 | # | 20 | Baseline | aldex2 | 12 |
| # | sex | 4 | 14 | 0 | 0.222222222 | 1 | # | 20 | Baseline | aldex2 | 12 |
| # | time | 5 | 32 | 0 | 0.135135135 | 1 | # | 20 | Baseline | aldex2 | 12 |
| # | interaction | 0 | 0 | 4 | 0 | 0 | # | 20 | Baseline | aldex2 | 12 |
| # | disease | 7 | 0 | 0 | 1 | 1 | # | 20 | Small | paralle | 12 |
| # | sex | 4 | 1 | 0 | 0.8 | 1 | # | 20 | Small | paralle | 12 |
| # | time | 5 | 0 | 0 | 1 | 1 | # | 20 | Small | paralle | 12 |
| # | interaction | 3 | 0 | 1 | 1 | 0.75 | # | 20 | Small | paralle | 12 |
| # | disease | 7 | 4 | 0 | 0.636363636 | 1 | # | 20 | Small | ancom | 12 |
| # | sex | 4 | 0 | 0 | 1 | 1 | # | 20 | Small | ancom | 12 |
| # | time | 5 | 4 | 0 | 0.555555556 | 1 | # | 20 | Small | ancom | 12 |
| # | interaction | 0 | 0 | 4 | 0 | 0 | # | 20 | Small | ancom | 12 |
| # | disease | 7 | 5 | 0 | 0.583333333 | 1 | # | 20 | Small | aldex2 | 12 |
| # | sex | 4 | 0 | 0 | 1 | 1 | # | 20 | Small | aldex2 | 12 |
| # | time | 5 | 7 | 0 | 0.416666667 | 1 | # | 20 | Small | aldex2 | 12 |
| # | interaction | 0 | 0 | 4 | 0 | 0 | # | 20 | Small | aldex2 | 12 |
| # | disease | 7 | 9 | 0 | 0.4375 | 1 | # | 20 | Baseline | paralle | 12 |
| # | sex | 4 | 7 | 0 | 0.363636364 | 1 | # | 20 | Baseline | paralle | 12 |
| # | time | 5 | 6 | 0 | 0.454545455 | 1 | # | 20 | Baseline | paralle | 12 |
| # | interaction | 4 | 0 | 0 | 1 | 1 | # | 20 | Baseline | paralle | 12 |
| # | disease | 7 | 4 | 0 | 0.636363636 | 1 | # | 20 | Baseline | ancom | 12 |
| # | sex | 4 | 0 | 0 | 1 | 1 | # | 20 | Baseline | ancom | 12 |
| # | time | 5 | 4 | 0 | 0.555555556 | 1 | # | 20 | Baseline | ancom | 12 |
| # | interaction | 0 | 0 | 4 | 0 | 0 | # | 20 | Baseline | ancom | 12 |
| # | disease | 7 | 60 | 0 | 0.104477612 | 1 | # | 20 | Baseline | aldex2 | 12 |
| # | sex | 4 | 18 | 0 | 0.181818182 | 1 | # | 20 | Baseline | aldex2 | 12 |
| # | time | 5 | 43 | 0 | 0.104166667 | 1 | # | 20 | Baseline | aldex2 | 12 |
| # | interaction | 0 | 0 | 4 | 0 | 0 | # | 20 | Baseline | aldex2 | 12 |
| # | disease | 7 | 0 | 0 | 1 | 1 | # | 20 | Small | paralle | 12 |
| # | sex | 4 | 0 | 0 | 1 | 1 | # | 20 | Small | paralle | 12 |
| # | time | 5 | 0 | 0 | 1 | 1 | # | 20 | Small | paralle | 12 |
| # | interaction | 4 | 0 | 0 | 1 | 1 | # | 20 | Small | paralle | 12 |
| # | disease | 7 | 4 | 0 | 0.636363636 | 1 | # | 20 | Small | ancom | 12 |
| # | sex | 4 | 0 | 0 | 1 | 1 | # | 20 | Small | ancom | 12 |
| # | time | 5 | 4 | 0 | 0.555555556 | 1 | # | 20 | Small | ancom | 12 |
| # | interaction | 0 | 0 | 4 | 0 | 0 | # | 20 | Small | ancom | 12 |
| # | disease | 7 | 9 | 0 | 0.4375 | 1 | # | 20 | Small | aldex2 | 12 |
| # | sex | 4 | 1 | 0 | 0.8 | 1 | # | 20 | Small | aldex2 | 12 |
| # | time | 5 | 8 | 0 | 0.384615385 | 1 | # | 20 | Small | aldex2 | 12 |
| # | interaction | 0 | 0 | 4 | 0 | 0 | # | 20 | Small | aldex2 | 12 |
| # | disease | 4 | 4 | 0 | 0.5 | 1 | # | 10 | Baseline | paralle | 13 |
| # | sex | 2 | 7 | 0 | 0.222222222 | 1 | # | 10 | Baseline | paralle | 13 |
| # | time | 2 | 9 | 0 | 0.181818182 | 1 | # | 10 | Baseline | paralle | 13 |
| # | interaction | 2 | 6 | 0 | 0.25 | 1 | # | 10 | Baseline | paralle | 13 |
| # | disease | 4 | 2 | 0 | 0.666666667 | 1 | # | 10 | Baseline | ancom | 13 |
| # | sex | 2 | 0 | 0 | 1 | 1 | # | 10 | Baseline | ancom | 13 |
| # | time | 2 | 0 | 0 | 1 | 1 | # | 10 | Baseline | ancom | 13 |
| # | interaction | 0 | 0 | 2 | 0 | 0 | # | 10 | Baseline | ancom | 13 |
| # | disease | 4 | 15 | 0 | 0.210526316 | 1 | # | 10 | Baseline | aldex2 | 13 |
| # | sex | 2 | 6 | 0 | 0.25 | 1 | # | 10 | Baseline | aldex2 | 13 |
| # | time | 2 | 7 | 0 | 0.222222222 | 1 | # | 10 | Baseline | aldex2 | 13 |
| # | interaction | 0 | 0 | 2 | 0 | 0 | # | 10 | Baseline | aldex2 | 13 |
| # | disease | 4 | 0 | 0 | 1 | 1 | # | 10 | Small | paralle | 13 |
| # | sex | 2 | 0 | 0 | 1 | 1 | # | 10 | Small | paralle | 13 |
| # | time | 2 | 1 | 0 | 0.666666667 | 1 | # | 10 | Small | paralle | 13 |
| # | interaction | 1 | 1 | 1 | 0.5 | 0.5 | # | 10 | Small | paralle | 13 |
| # | disease | 4 | 2 | 0 | 0.666666667 | 1 | # | 10 | Small | ancom | 13 |
| # | sex | 2 | 0 | 0 | 1 | 1 | # | 10 | Small | ancom | 13 |
| # | time | 2 | 0 | 0 | 1 | 1 | # | 10 | Small | ancom | 13 |
| # | interaction | 0 | 0 | 2 | 0 | 0 | # | 10 | Small | ancom | 13 |
| # | disease | 4 | 2 | 0 | 0.666666667 | 1 | # | 10 | Small | aldex2 | 13 |
| # | sex | 2 | 0 | 0 | 1 | 1 | # | 10 | Small | aldex2 | 13 |
| # | time | 2 | 0 | 0 | 1 | 1 | # | 10 | Small | aldex2 | 13 |
| # | interaction | 0 | 0 | 2 | 0 | 0 | # | 10 | Small | aldex2 | 13 |
| # | disease | 4 | 7 | 0 | 0.363636364 | 1 | # | 10 | Baseline | paralle | 13 |
| # | sex | 2 | 3 | 0 | 0.4 | 1 | # | 10 | Baseline | paralle | 13 |
| # | time | 2 | 5 | 0 | 0.285714286 | 1 | # | 10 | Baseline | paralle | 13 |
| # | interaction | 2 | 5 | 0 | 0.285714286 | 1 | # | 10 | Baseline | paralle | 13 |
| # | disease | 4 | 2 | 0 | 0.666666667 | 1 | # | 10 | Baseline | ancom | 13 |
| # | sex | 2 | 0 | 0 | 1 | 1 | # | 10 | Baseline | ancom | 13 |
| # | time | 2 | 2 | 0 | 0.5 | 1 | # | 10 | Baseline | ancom | 13 |
| # | interaction | 0 | 0 | 2 | 0 | 0 | # | 10 | Baseline | ancom | 13 |
| # | disease | 4 | 17 | 0 | 0.19047619 | 1 | # | 10 | Baseline | aldex2 | 13 |
| # | sex | 2 | 4 | 0 | 0.333333333 | 1 | # | 10 | Baseline | aldex2 | 13 |
| # | time | 2 | 9 | 0 | 0.181818182 | 1 | # | 10 | Baseline | aldex2 | 13 |
| # | interaction | 0 | 0 | 2 | 0 | 0 | # | 10 | Baseline | aldex2 | 13 |
| # | disease | 4 | 1 | 0 | 0.8 | 1 | # | 10 | Small | paralle | 13 |
| # | sex | 2 | 0 | 0 | 1 | 1 | # | 10 | Small | paralle | 13 |
| # | time | 2 | 0 | 0 | 1 | 1 | # | 10 | Small | paralle | 13 |
| # | interaction | 2 | 0 | 0 | 1 | 1 | # | 10 | Small | paralle | 13 |
| # | disease | 4 | 1 | 0 | 0.8 | 1 | # | 10 | Small | ancom | 13 |
| # | sex | 2 | 0 | 0 | 1 | 1 | # | 10 | Small | ancom | 13 |
| # | time | 2 | 2 | 0 | 0.5 | 1 | # | 10 | Small | ancom | 13 |
| # | interaction | 0 | 0 | 2 | 0 | 0 | # | 10 | Small | ancom | 13 |
| # | disease | 4 | 3 | 0 | 0.571428571 | 1 | # | 10 | Small | aldex2 | 13 |
| # | sex | 2 | 0 | 0 | 1 | 1 | # | 10 | Small | aldex2 | 13 |
| # | time | 2 | 2 | 0 | 0.5 | 1 | # | 10 | Small | aldex2 | 13 |
| # | interaction | 0 | 0 | 2 | 0 | 0 | # | 10 | Small | aldex2 | 13 |
| # | disease | 4 | 8 | 0 | 0.333333333 | 1 | # | 10 | Baseline | paralle | 13 |
| # | sex | 2 | 4 | 0 | 0.333333333 | 1 | # | 10 | Baseline | paralle | 13 |
| # | time | 2 | 9 | 0 | 0.181818182 | 1 | # | 10 | Baseline | paralle | 13 |
| # | interaction | 2 | 5 | 0 | 0.285714286 | 1 | # | 10 | Baseline | paralle | 13 |

| | | | | | | | | | | | |
|---|---|---|---|---|---|---|---|---|---|---|---|
| # | disease | 4 | 2 | 0 | 0.666666667 | 1 | # | 10 | Baseline | ancom | 13 |
| # | sex | 2 | 0 | 0 | 1 | 1 | # | 10 | Baseline | ancom | 13 |
| # | time | 2 | 2 | 0 | 0.5 | 1 | # | 10 | Baseline | ancom | 13 |
| # | interaction | 0 | 0 | 2 | 0 | 0 | # | 10 | Baseline | ancom | 13 |
| # | disease | 4 | 29 | 0 | 0.121212121 | 1 | # | 10 | Baseline | aldex2 | 13 |
| # | sex | 2 | 7 | 0 | 0.222222222 | 1 | # | 10 | Baseline | aldex2 | 13 |
| # | time | 2 | 14 | 0 | 0.125 | 1 | # | 10 | Baseline | aldex2 | 13 |
| # | interaction | 0 | 0 | 2 | 0 | 0 | # | 10 | Baseline | aldex2 | 13 |
| # | disease | 4 | 1 | 0 | 0.8 | 1 | # | 10 | Small | paralle | 13 |
| # | sex | 2 | 0 | 0 | 1 | 1 | # | 10 | Small | paralle | 13 |
| # | time | 2 | 0 | 0 | 1 | 1 | # | 10 | Small | paralle | 13 |
| # | interaction | 2 | 0 | 0 | 1 | 1 | # | 10 | Small | paralle | 13 |
| # | disease | 4 | 2 | 0 | 0.666666667 | 1 | # | 10 | Small | ancom | 13 |
| # | sex | 2 | 0 | 0 | 1 | 1 | # | 10 | Small | ancom | 13 |
| # | time | 2 | 1 | 0 | 0.666666667 | 1 | # | 10 | Small | ancom | 13 |
| # | interaction | 0 | 0 | 2 | 0 | 0 | # | 10 | Small | ancom | 13 |
| # | disease | 4 | 3 | 0 | 0.571428571 | 1 | # | 10 | Small | aldex2 | 13 |
| # | sex | 2 | 0 | 0 | 1 | 1 | # | 10 | Small | aldex2 | 13 |
| # | time | 2 | 2 | 0 | 0.5 | 1 | # | 10 | Small | aldex2 | 13 |
| # | interaction | 0 | 0 | 2 | 0 | 0 | # | 10 | Small | aldex2 | 13 |
| # | disease | 4 | 6 | 0 | 0.4 | 1 | # | 10 | Baseline | paralle | 13 |
| # | sex | 2 | 2 | 0 | 0.5 | 1 | # | 10 | Baseline | paralle | 13 |
| # | time | 2 | 6 | 0 | 0.25 | 1 | # | 10 | Baseline | paralle | 13 |
| # | interaction | 2 | 7 | 0 | 0.222222222 | 1 | # | 10 | Baseline | paralle | 13 |
| # | disease | 4 | 2 | 0 | 0.666666667 | 1 | # | 10 | Baseline | ancom | 13 |
| # | sex | 2 | 0 | 0 | 1 | 1 | # | 10 | Baseline | ancom | 13 |
| # | time | 2 | 2 | 0 | 0.5 | 1 | # | 10 | Baseline | ancom | 13 |
| # | interaction | 0 | 0 | 2 | 0 | 0 | # | 10 | Baseline | ancom | 13 |
| # | disease | 4 | 30 | 0 | 0.117647059 | 1 | # | 10 | Baseline | aldex2 | 13 |
| # | sex | 2 | 8 | 0 | 0.2 | 1 | # | 10 | Baseline | aldex2 | 13 |
| # | time | 2 | 13 | 0 | 0.133333333 | 1 | # | 10 | Baseline | aldex2 | 13 |
| # | interaction | 0 | 0 | 2 | 0 | 0 | # | 10 | Baseline | aldex2 | 13 |
| # | disease | 4 | 0 | 0 | 1 | 1 | # | 10 | Small | paralle | 13 |
| # | sex | 2 | 0 | 0 | 1 | 1 | # | 10 | Small | paralle | 13 |
| # | time | 2 | 0 | 0 | 1 | 1 | # | 10 | Small | paralle | 13 |
| # | interaction | 2 | 0 | 0 | 1 | 1 | # | 10 | Small | paralle | 13 |
| # | disease | 4 | 2 | 0 | 0.666666667 | 1 | # | 10 | Small | ancom | 13 |
| # | sex | 2 | 0 | 0 | 1 | 1 | # | 10 | Small | ancom | 13 |
| # | time | 2 | 2 | 0 | 0.5 | 1 | # | 10 | Small | ancom | 13 |
| # | interaction | 0 | 0 | 2 | 0 | 0 | # | 10 | Small | ancom | 13 |
| # | disease | 4 | 3 | 0 | 0.571428571 | 1 | # | 10 | Small | aldex2 | 13 |
| # | sex | 2 | 0 | 0 | 1 | 1 | # | 10 | Small | aldex2 | 13 |
| # | time | 2 | 2 | 0 | 0.5 | 1 | # | 10 | Small | aldex2 | 13 |
| # | interaction | 0 | 0 | 2 | 0 | 0 | # | 10 | Small | aldex2 | 13 |
| # | disease | 7 | 12 | 0 | 0.368421053 | 1 | # | 20 | Baseline | paralle | 13 |
| # | sex | 4 | 6 | 0 | 0.4 | 1 | # | 20 | Baseline | paralle | 13 |
| # | time | 5 | 10 | 0 | 0.333333333 | 1 | # | 20 | Baseline | paralle | 13 |
| # | interaction | 4 | 10 | 0 | 0.285714286 | 1 | # | 20 | Baseline | paralle | 13 |
| # | disease | 7 | 2 | 0 | 0.777777778 | 1 | # | 20 | Baseline | ancom | 13 |
| # | sex | 4 | 0 | 0 | 1 | 1 | # | 20 | Baseline | ancom | 13 |
| # | time | 5 | 0 | 0 | 1 | 1 | # | 20 | Baseline | ancom | 13 |
| # | interaction | 0 | 0 | 4 | 0 | 0 | # | 20 | Baseline | ancom | 13 |
| # | disease | 7 | 18 | 0 | 0.28 | 1 | # | 20 | Baseline | aldex2 | 13 |
| # | sex | 4 | 9 | 0 | 0.307692308 | 1 | # | 20 | Baseline | aldex2 | 13 |
| # | time | 5 | 13 | 0 | 0.277777778 | 1 | # | 20 | Baseline | aldex2 | 13 |
| # | interaction | 0 | 0 | 4 | 0 | 0 | # | 20 | Baseline | aldex2 | 13 |
| # | disease | 7 | 0 | 0 | 1 | 1 | # | 20 | Small | paralle | 13 |
| # | sex | 4 | 0 | 0 | 1 | 1 | # | 20 | Small | paralle | 13 |
| # | time | 5 | 0 | 0 | 1 | 1 | # | 20 | Small | paralle | 13 |
| # | interaction | 1 | 0 | 3 | 1 | 0.25 | # | 20 | Small | paralle | 13 |
| # | disease | 7 | 2 | 0 | 0.777777778 | 1 | # | 20 | Small | ancom | 13 |
| # | sex | 4 | 0 | 0 | 1 | 1 | # | 20 | Small | ancom | 13 |
| # | time | 5 | 0 | 0 | 1 | 1 | # | 20 | Small | ancom | 13 |
| # | interaction | 0 | 0 | 4 | 0 | 0 | # | 20 | Small | ancom | 13 |
| # | disease | 7 | 2 | 0 | 0.777777778 | 1 | # | 20 | Small | aldex2 | 13 |
| # | sex | 4 | 1 | 0 | 0.8 | 1 | # | 20 | Small | aldex2 | 13 |
| # | time | 5 | 1 | 0 | 0.833333333 | 1 | # | 20 | Small | aldex2 | 13 |
| # | interaction | 0 | 0 | 4 | 0 | 0 | # | 20 | Small | aldex2 | 13 |
| # | disease | 7 | 14 | 0 | 0.333333333 | 1 | # | 20 | Baseline | paralle | 13 |
| # | sex | 4 | 9 | 0 | 0.307692308 | 1 | # | 20 | Baseline | paralle | 13 |
| # | time | 5 | 10 | 0 | 0.333333333 | 1 | # | 20 | Baseline | paralle | 13 |
| # | interaction | 4 | 3 | 0 | 0.571428571 | 1 | # | 20 | Baseline | paralle | 13 |
| # | disease | 7 | 4 | 0 | 0.636363636 | 1 | # | 20 | Baseline | ancom | 13 |
| # | sex | 4 | 0 | 0 | 1 | 1 | # | 20 | Baseline | ancom | 13 |
| # | time | 5 | 3 | 0 | 0.625 | 1 | # | 20 | Baseline | ancom | 13 |
| # | interaction | 0 | 0 | 4 | 0 | 0 | # | 20 | Baseline | ancom | 13 |
| # | disease | 7 | 34 | 0 | 0.170731707 | 1 | # | 20 | Baseline | aldex2 | 13 |
| # | sex | 4 | 14 | 0 | 0.222222222 | 1 | # | 20 | Baseline | aldex2 | 13 |
| # | time | 5 | 20 | 0 | 0.2 | 1 | # | 20 | Baseline | aldex2 | 13 |
| # | interaction | 0 | 0 | 4 | 0 | 0 | # | 20 | Baseline | aldex2 | 13 |
| # | disease | 7 | 0 | 0 | 1 | 1 | # | 20 | Small | paralle | 13 |
| # | sex | 4 | 0 | 0 | 1 | 1 | # | 20 | Small | paralle | 13 |
| # | time | 5 | 0 | 0 | 1 | 1 | # | 20 | Small | paralle | 13 |
| # | interaction | 2 | 0 | 2 | 1 | 0.5 | # | 20 | Small | paralle | 13 |
| # | disease | 7 | 4 | 0 | 0.636363636 | 1 | # | 20 | Small | ancom | 13 |
| # | sex | 4 | 0 | 0 | 1 | 1 | # | 20 | Small | ancom | 13 |
| # | time | 5 | 3 | 0 | 0.625 | 1 | # | 20 | Small | ancom | 13 |
| # | interaction | 0 | 0 | 4 | 0 | 0 | # | 20 | Small | ancom | 13 |
| # | disease | 7 | 3 | 0 | 0.7 | 1 | # | 20 | Small | aldex2 | 13 |
| # | sex | 4 | 1 | 0 | 0.8 | 1 | # | 20 | Small | aldex2 | 13 |
| # | time | 5 | 2 | 0 | 0.714285714 | 1 | # | 20 | Small | aldex2 | 13 |
| # | interaction | 0 | 0 | 4 | 0 | 0 | # | 20 | Small | aldex2 | 13 |
| # | disease | 7 | 15 | 0 | 0.318181818 | 1 | # | 20 | Baseline | paralle | 13 |
| # | sex | 4 | 6 | 0 | 0.4 | 1 | # | 20 | Baseline | paralle | 13 |
| # | time | 5 | 4 | 0 | 0.555555556 | 1 | # | 20 | Baseline | paralle | 13 |
| # | interaction | 4 | 4 | 0 | 0.5 | 1 | # | 20 | Baseline | paralle | 13 |
| # | disease | 7 | 4 | 0 | 0.636363636 | 1 | # | 20 | Baseline | ancom | 13 |
| # | sex | 4 | 0 | 0 | 1 | 1 | # | 20 | Baseline | ancom | 13 |
| # | time | 5 | 4 | 0 | 0.555555556 | 1 | # | 20 | Baseline | ancom | 13 |
| # | interaction | 0 | 0 | 4 | 0 | 0 | # | 20 | Baseline | ancom | 13 |
| # | disease | 7 | 68 | 0 | 0.127272727 | 1 | # | 20 | Baseline | aldex2 | 13 |
| # | sex | 4 | 15 | 0 | 0.210526316 | 1 | # | 20 | Baseline | aldex2 | 13 |
| # | time | 5 | 30 | 0 | 0.142857143 | 1 | # | 20 | Baseline | aldex2 | 13 |
| # | interaction | 0 | 0 | 4 | 0 | 0 | # | 20 | Baseline | aldex2 | 13 |
| # | disease | 7 | 0 | 0 | 1 | 1 | # | 20 | Small | paralle | 13 |
| # | sex | 4 | 0 | 0 | 1 | 1 | # | 20 | Small | paralle | 13 |
| # | time | 5 | 0 | 0 | 1 | 1 | # | 20 | Small | paralle | 13 |
| # | interaction | 4 | 0 | 0 | 1 | 1 | # | 20 | Small | paralle | 13 |
| # | disease | 7 | 4 | 0 | 0.636363636 | 1 | # | 20 | Small | ancom | 13 |
| # | sex | 4 | 0 | 0 | 1 | 1 | # | 20 | Small | ancom | 13 |
| # | time | 5 | 4 | 0 | 0.555555556 | 1 | # | 20 | Small | ancom | 13 |
| # | interaction | 0 | 0 | 4 | 0 | 0 | # | 20 | Small | ancom | 13 |
| # | disease | 7 | 5 | 0 | 0.583333333 | 1 | # | 20 | Small | aldex2 | 13 |
| # | sex | 4 | 1 | 0 | 0.8 | 1 | # | 20 | Small | aldex2 | 13 |
| # | time | 5 | 4 | 0 | 0.555555556 | 1 | # | 20 | Small | aldex2 | 13 |
| # | interaction | 0 | 0 | 4 | 0 | 0 | # | 20 | Small | aldex2 | 13 |
| # | disease | 7 | 13 | 0 | 0.35 | 1 | # | 20 | Baseline | paralle | 13 |
| # | sex | 4 | 5 | 0 | 0.444444444 | 1 | # | 20 | Baseline | paralle | 13 |
| # | time | 5 | 6 | 0 | 0.454545455 | 1 | # | 20 | Baseline | paralle | 13 |
| # | interaction | 4 | 7 | 0 | 0.363636364 | 1 | # | 20 | Baseline | paralle | 13 |
| # | disease | 7 | 4 | 0 | 0.636363636 | 1 | # | 20 | Baseline | ancom | 13 |
| # | sex | 4 | 0 | 0 | 1 | 1 | # | 20 | Baseline | ancom | 13 |
| # | time | 5 | 4 | 0 | 0.555555556 | 1 | # | 20 | Baseline | ancom | 13 |
| # | interaction | 0 | 0 | 4 | 0 | 0 | # | 20 | Baseline | ancom | 13 |
| # | disease | 7 | 91 | 0 | 0.102941176 | 1 | # | 20 | Baseline | aldex2 | 13 |
| # | sex | 4 | 22 | 0 | 0.153846154 | 1 | # | 20 | Baseline | aldex2 | 13 |
| # | time | 5 | 37 | 0 | 0.119047619 | 1 | # | 20 | Baseline | aldex2 | 13 |
| # | interaction | 0 | 0 | 4 | 0 | 0 | # | 20 | Baseline | aldex2 | 13 |
| # | disease | 7 | 0 | 0 | 1 | 1 | # | 20 | Small | paralle | 13 |
| # | sex | 4 | 0 | 0 | 1 | 1 | # | 20 | Small | paralle | 13 |
| # | time | 5 | 0 | 0 | 1 | 1 | # | 20 | Small | paralle | 13 |
| # | interaction | 4 | 0 | 0 | 1 | 1 | # | 20 | Small | paralle | 13 |
| # | disease | 7 | 4 | 0 | 0.636363636 | 1 | # | 20 | Small | ancom | 13 |
| # | sex | 4 | 0 | 0 | 1 | 1 | # | 20 | Small | ancom | 13 |
| # | time | 5 | 4 | 0 | 0.555555556 | 1 | # | 20 | Small | ancom | 13 |
| # | interaction | 0 | 0 | 4 | 0 | 0 | # | 20 | Small | ancom | 13 |
| # | disease | 7 | 7 | 0 | 0.5 | 1 | # | 20 | Small | aldex2 | 13 |
| # | sex | 4 | 1 | 0 | 0.8 | 1 | # | 20 | Small | aldex2 | 13 |
| # | time | 5 | 5 | 0 | 0.5 | 1 | # | 20 | Small | aldex2 | 13 |
| # | interaction | 0 | 0 | 4 | 0 | 0 | # | 20 | Small | aldex2 | 13 |
| # | disease | 4 | 11 | 0 | 0.266666667 | 1 | # | 10 | Baseline | paralle | 14 |
| # | sex | 2 | 10 | 0 | 0.166666667 | 1 | # | 10 | Baseline | paralle | 14 |
| # | time | 2 | 11 | 0 | 0.153846154 | 1 | # | 10 | Baseline | paralle | 14 |
| # | interaction | 2 | 12 | 0 | 0.142857143 | 1 | # | 10 | Baseline | paralle | 14 |
| # | disease | 4 | 1 | 0 | 0.8 | 1 | # | 10 | Baseline | ancom | 14 |
| # | sex | 2 | 0 | 0 | 1 | 1 | # | 10 | Baseline | ancom | 14 |
| # | time | 2 | 0 | 0 | 1 | 1 | # | 10 | Baseline | ancom | 14 |
| # | interaction | 0 | 0 | 2 | 0 | 0 | # | 10 | Baseline | ancom | 14 |
| # | disease | 4 | 12 | 0 | 0.25 | 1 | # | 10 | Baseline | aldex2 | 14 |
| # | sex | 2 | 3 | 0 | 0.4 | 1 | # | 10 | Baseline | aldex2 | 14 |
| # | time | 2 | 6 | 0 | 0.25 | 1 | # | 10 | Baseline | aldex2 | 14 |
| # | interaction | 0 | 0 | 2 | 0 | 0 | # | 10 | Baseline | aldex2 | 14 |
| # | disease | 4 | 0 | 0 | 1 | 1 | # | 10 | Small | paralle | 14 |
| # | sex | 2 | 0 | 0 | 1 | 1 | # | 10 | Small | paralle | 14 |
| # | time | 2 | 0 | 0 | 1 | 1 | # | 10 | Small | paralle | 14 |
| # | interaction | 0 | 0 | 2 | 0 | 0 | # | 10 | Small | paralle | 14 |
| # | disease | 4 | 1 | 0 | 0.8 | 1 | # | 10 | Small | ancom | 14 |
| # | sex | 2 | 0 | 0 | 1 | 1 | # | 10 | Small | ancom | 14 |
| # | time | 2 | 0 | 0 | 1 | 1 | # | 10 | Small | ancom | 14 |
| # | interaction | 0 | 0 | 2 | 0 | 0 | # | 10 | Small | ancom | 14 |
| # | disease | 4 | 2 | 0 | 0.666666667 | 1 | # | 10 | Small | aldex2 | 14 |
| # | sex | 2 | 0 | 0 | 1 | 1 | # | 10 | Small | aldex2 | 14 |
| # | time | 2 | 0 | 0 | 1 | 1 | # | 10 | Small | aldex2 | 14 |
| # | interaction | 0 | 0 | 2 | 0 | 0 | # | 10 | Small | aldex2 | 14 |
| # | disease | 4 | 10 | 0 | 0.285714286 | 1 | # | 10 | Baseline | paralle | 14 |
| # | sex | 2 | 7 | 0 | 0.222222222 | 1 | # | 10 | Baseline | paralle | 14 |
| # | time | 2 | 6 | 0 | 0.25 | 1 | # | 10 | Baseline | paralle | 14 |
| # | interaction | 2 | 7 | 0 | 0.222222222 | 1 | # | 10 | Baseline | paralle | 14 |
| # | disease | 4 | 3 | 0 | 0.571428571 | 1 | # | 10 | Baseline | ancom | 14 |
| # | sex | 2 | 0 | 0 | 1 | 1 | # | 10 | Baseline | ancom | 14 |
| # | time | 2 | 1 | 0 | 0.666666667 | 1 | # | 10 | Baseline | ancom | 14 |
| # | interaction | 0 | 0 | 2 | 0 | 0 | # | 10 | Baseline | ancom | 14 |
| # | disease | 4 | 13 | 0 | 0.235294118 | 1 | # | 10 | Baseline | aldex2 | 14 |
| # | sex | 2 | 7 | 0 | 0.222222222 | 1 | # | 10 | Baseline | aldex2 | 14 |
| # | time | 2 | 9 | 0 | 0.181818182 | 1 | # | 10 | Baseline | aldex2 | 14 |
| # | interaction | 0 | 0 | 2 | 0 | 0 | # | 10 | Baseline | aldex2 | 14 |
| # | disease | 4 | 0 | 0 | 1 | 1 | # | 10 | Small | paralle | 14 |
| # | sex | 2 | 0 | 0 | 1 | 1 | # | 10 | Small | paralle | 14 |
| # | time | 2 | 0 | 0 | 1 | 1 | # | 10 | Small | paralle | 14 |
| # | interaction | 0 | 0 | 2 | 0 | 0 | # | 10 | Small | paralle | 14 |
| # | disease | 4 | 3 | 0 | 0.571428571 | 1 | # | 10 | Small | ancom | 14 |
| # | sex | 2 | 0 | 0 | 1 | 1 | # | 10 | Small | ancom | 14 |
| # | time | 2 | 1 | 0 | 0.666666667 | 1 | # | 10 | Small | ancom | 14 |
| # | interaction | 0 | 0 | 2 | 0 | 0 | # | 10 | Small | ancom | 14 |
| # | disease | 4 | 3 | 0 | 0.571428571 | 1 | # | 10 | Small | aldex2 | 14 |
| # | sex | 2 | 0 | 0 | 1 | 1 | # | 10 | Small | aldex2 | 14 |
| # | time | 2 | 2 | 0 | 0.5 | 1 | # | 10 | Small | aldex2 | 14 |
| # | interaction | 0 | 0 | 2 | 0 | 0 | # | 10 | Small | aldex2 | 14 |
| # | disease | 4 | 4 | 0 | 0.5 | 1 | # | 10 | Baseline | paralle | 14 |
| # | sex | 2 | 3 | 0 | 0.4 | 1 | # | 10 | Baseline | paralle | 14 |
| # | time | 2 | 2 | 0 | 0.5 | 1 | # | 10 | Baseline | paralle | 14 |
| # | interaction | 2 | 4 | 0 | 0.333333333 | 1 | # | 10 | Baseline | paralle | 14 |
| # | disease | 4 | 3 | 0 | 0.571428571 | 1 | # | 10 | Baseline | ancom | 14 |
| # | sex | 2 | 0 | 0 | 1 | 1 | # | 10 | Baseline | ancom | 14 |
| # | time | 2 | 2 | 0 | 0.5 | 1 | # | 10 | Baseline | ancom | 14 |
| # | interaction | 0 | 0 | 2 | 0 | 0 | # | 10 | Baseline | ancom | 14 |
| # | disease | 4 | 17 | 0 | 0.19047619 | 1 | # | 10 | Baseline | aldex2 | 14 |
| # | sex | 2 | 8 | 0 | 0.2 | 1 | # | 10 | Baseline | aldex2 | 14 |
| # | time | 2 | 11 | 0 | 0.153846154 | 1 | # | 10 | Baseline | aldex2 | 14 |
| # | interaction | 0 | 0 | 2 | 0 | 0 | # | 10 | Baseline | aldex2 | 14 |
| # | disease | 4 | 0 | 0 | 1 | 1 | # | 10 | Small | paralle | 14 |
| # | sex | 2 | 0 | 0 | 1 | 1 | # | 10 | Small | paralle | 14 |
| # | time | 2 | 0 | 0 | 1 | 1 | # | 10 | Small | paralle | 14 |
| # | interaction | 2 | 0 | 0 | 1 | 1 | # | 10 | Small | paralle | 14 |
| # | disease | 4 | 3 | 0 | 0.571428571 | 1 | # | 10 | Small | ancom | 14 |
| # | sex | 2 | 0 | 0 | 1 | 1 | # | 10 | Small | ancom | 14 |
| # | time | 2 | 2 | 0 | 0.5 | 1 | # | 10 | Small | ancom | 14 |
| # | interaction | 0 | 0 | 2 | 0 | 0 | # | 10 | Small | ancom | 14 |
| # | disease | 4 | 4 | 0 | 0.5 | 1 | # | 10 | Small | aldex2 | 14 |
| # | sex | 2 | 0 | 0 | 1 | 1 | # | 10 | Small | aldex2 | 14 |
| # | time | 2 | 2 | 0 | 0.5 | 1 | # | 10 | Small | aldex2 | 14 |
| # | interaction | 0 | 0 | 2 | 0 | 0 | # | 10 | Small | aldex2 | 14 |
| # | disease | 4 | 7 | 0 | 0.363636364 | 1 | # | 10 | Baseline | paralle | 14 |
| # | sex | 2 | 3 | 0 | 0.4 | 1 | # | 10 | Baseline | paralle | 14 |
| # | time | 2 | 2 | 0 | 0.5 | 1 | # | 10 | Baseline | paralle | 14 |
| # | interaction | 2 | 2 | 0 | 0.5 | 1 | # | 10 | Baseline | paralle | 14 |
| # | disease | 4 | 2 | 0 | 0.666666667 | 1 | # | 10 | Baseline | ancom | 14 |
| # | sex | 2 | 0 | 0 | 1 | 1 | # | 10 | Baseline | ancom | 14 |
| # | time | 2 | 2 | 0 | 0.5 | 1 | # | 10 | Baseline | ancom | 14 |
| # | interaction | 0 | 0 | 2 | 0 | 0 | # | 10 | Baseline | ancom | 14 |
| # | disease | 4 | 24 | 0 | 0.142857143 | 1 | # | 10 | Baseline | aldex2 | 14 |
| # | sex | 2 | 8 | 0 | 0.2 | 1 | # | 10 | Baseline | aldex2 | 14 |
| # | time | 2 | 11 | 0 | 0.153846154 | 1 | # | 10 | Baseline | aldex2 | 14 |
| # | interaction | 0 | 0 | 2 | 0 | 0 | # | 10 | Baseline | aldex2 | 14 |
| # | disease | 4 | 0 | 0 | 1 | 1 | # | 10 | Small | paralle | 14 |
| # | sex | 2 | 0 | 0 | 1 | 1 | # | 10 | Small | paralle | 14 |
| # | time | 2 | 0 | 0 | 1 | 1 | # | 10 | Small | paralle | 14 |
| # | interaction | 2 | 0 | 0 | 1 | 1 | # | 10 | Small | paralle | 14 |
| # | disease | 4 | 2 | 0 | 0.666666667 | 1 | # | 10 | Small | ancom | 14 |
| # | sex | 2 | 0 | 0 | 1 | 1 | # | 10 | Small | ancom | 14 |
| # | time | 2 | 2 | 0 | 0.5 | 1 | # | 10 | Small | ancom | 14 |
| # | interaction | 0 | 0 | 2 | 0 | 0 | # | 10 | Small | ancom | 14 |
| # | disease | 4 | 3 | 0 | 0.571428571 | 1 | # | 10 | Small | aldex2 | 14 |
| # | sex | 2 | 0 | 0 | 1 | 1 | # | 10 | Small | aldex2 | 14 |
| # | time | 2 | 2 | 0 | 0.5 | 1 | # | 10 | Small | aldex2 | 14 |
| # | interaction | 0 | 0 | 2 | 0 | 0 | # | 10 | Small | aldex2 | 14 |
| # | disease | 7 | 14 | 0 | 0.333333333 | 1 | # | 20 | Baseline | paralle | 14 |
| # | sex | 4 | 6 | 0 | 0.4 | 1 | # | 20 | Baseline | paralle | 14 |
| # | time | 5 | 5 | 0 | 0.5 | 1 | # | 20 | Baseline | paralle | 14 |
| # | interaction | 3 | 4 | 1 | 0.428571429 | 0.75 | # | 20 | Baseline | paralle | 14 |
| # | disease | 7 | 0 | 0 | 1 | 1 | # | 20 | Baseline | ancom | 14 |
| # | sex | 4 | 0 | 0 | 1 | 1 | # | 20 | Baseline | ancom | 14 |
| # | time | 5 | 1 | 0 | 0.833333333 | 1 | # | 20 | Baseline | ancom | 14 |
| # | interaction | 0 | 0 | 4 | 0 | 0 | # | 20 | Baseline | ancom | 14 |
| # | disease | 7 | 15 | 0 | 0.318181818 | 1 | # | 20 | Baseline | aldex2 | 14 |
| # | sex | 4 | 6 | 0 | 0.4 | 1 | # | 20 | Baseline | aldex2 | 14 |
| # | time | 5 | 14 | 0 | 0.263157895 | 1 | # | 20 | Baseline | aldex2 | 14 |
| # | interaction | 0 | 0 | 4 | 0 | 0 | # | 20 | Baseline | aldex2 | 14 |
| # | disease | 7 | 0 | 0 | 1 | 1 | # | 20 | Small | paralle | 14 |
| # | sex | 4 | 0 | 0 | 1 | 1 | # | 20 | Small | paralle | 14 |
| # | time | 5 | 0 | 0 | 1 | 1 | # | 20 | Small | paralle | 14 |
| # | interaction | 3 | 0 | 1 | 1 | 0.75 | # | 20 | Small | paralle | 14 |
| # | disease | 7 | 0 | 0 | 1 | 1 | # | 20 | Small | ancom | 14 |
| # | sex | 4 | 0 | 0 | 1 | 1 | # | 20 | Small | ancom | 14 |
| # | time | 5 | 1 | 0 | 0.833333333 | 1 | # | 20 | Small | ancom | 14 |
| # | interaction | 0 | 0 | 4 | 0 | 0 | # | 20 | Small | ancom | 14 |
| # | disease | 7 | 0 | 0 | 1 | 1 | # | 20 | Small | aldex2 | 14 |
| # | sex | 4 | 0 | 0 | 1 | 1 | # | 20 | Small | aldex2 | 14 |
| # | time | 5 | 1 | 0 | 0.833333333 | 1 | # | 20 | Small | aldex2 | 14 |
| # | interaction | 0 | 0 | 4 | 0 | 0 | # | 20 | Small | aldex2 | 14 |
| # | disease | 7 | 5 | 0 | 0.583333333 | 1 | # | 20 | Baseline | paralle | 14 |
| # | sex | 4 | 4 | 0 | 0.5 | 1 | # | 20 | Baseline | paralle | 14 |
| # | time | 5 | 6 | 0 | 0.454545455 | 1 | # | 20 | Baseline | paralle | 14 |
| # | interaction | 4 | 4 | 0 | 0.5 | 1 | # | 20 | Baseline | paralle | 14 |
| # | disease | 7 | 3 | 0 | 0.7 | 1 | # | 20 | Baseline | ancom | 14 |
| # | sex | 4 | 0 | 0 | 1 | 1 | # | 20 | Baseline | ancom | 14 |
| # | time | 5 | 4 | 0 | 0.555555556 | 1 | # | 20 | Baseline | ancom | 14 |
| # | interaction | 0 | 0 | 4 | 0 | 0 | # | 20 | Baseline | ancom | 14 |
| # | disease | 7 | 32 | 0 | 0.179487179 | 1 | # | 20 | Baseline | aldex2 | 14 |
| # | sex | 4 | 15 | 0 | 0.210526316 | 1 | # | 20 | Baseline | aldex2 | 14 |
| # | time | 5 | 24 | 0 | 0.172413793 | 1 | # | 20 | Baseline | aldex2 | 14 |
| # | interaction | 0 | 0 | 4 | 0 | 0 | # | 20 | Baseline | aldex2 | 14 |
| # | disease | 7 | 0 | 0 | 1 | 1 | # | 20 | Small | paralle | 14 |
| # | sex | 4 | 0 | 0 | 1 | 1 | # | 20 | Small | paralle | 14 |
| # | time | 5 | 0 | 0 | 1 | 1 | # | 20 | Small | paralle | 14 |
| # | interaction | 3 | 0 | 1 | 1 | 0.75 | # | 20 | Small | paralle | 14 |
| # | disease | 7 | 2 | 0 | 0.777777778 | 1 | # | 20 | Small | ancom | 14 |
| # | sex | 4 | 0 | 0 | 1 | 1 | # | 20 | Small | ancom | 14 |
| # | time | 5 | 4 | 0 | 0.555555556 | 1 | # | 20 | Small | ancom | 14 |
| # | interaction | 0 | 0 | 4 | 0 | 0 | # | 20 | Small | ancom | 14 |
| # | disease | 7 | 1 | 0 | 0.875 | 1 | # | 20 | Small | aldex2 | 14 |
| # | sex | 4 | 0 | 0 | 1 | 1 | # | 20 | Small | aldex2 | 14 |
| # | time | 5 | 4 | 0 | 0.555555556 | 1 | # | 20 | Small | aldex2 | 14 |
| # | interaction | 0 | 0 | 4 | 0 | 0 | # | 20 | Small | aldex2 | 14 |
| # | disease | 7 | 4 | 0 | 0.636363636 | 1 | # | 20 | Baseline | paralle | 14 |
| # | sex | 4 | 7 | 0 | 0.363636364 | 1 | # | 20 | Baseline | paralle | 14 |
| # | time | 5 | 8 | 0 | 0.384615385 | 1 | # | 20 | Baseline | paralle | 14 |
| # | interaction | 4 | 3 | 0 | 0.571428571 | 1 | # | 20 | Baseline | paralle | 14 |
| # | disease | 7 | 4 | 0 | 0.636363636 | 1 | # | 20 | Baseline | ancom | 14 |
| # | sex | 4 | 0 | 0 | 1 | 1 | # | 20 | Baseline | ancom | 14 |
| # | time | 5 | 4 | 0 | 0.555555556 | 1 | # | 20 | Baseline | ancom | 14 |
| # | interaction | 0 | 0 | 4 | 0 | 0 | # | 20 | Baseline | ancom | 14 |
| # | disease | 7 | 68 | 0 | 0.127272727 | 1 | # | 20 | Baseline | aldex2 | 14 |
| # | sex | 4 | 15 | 0 | 0.210526316 | 1 | # | 20 | Baseline | aldex2 | 14 |
| # | time | 5 | 34 | 0 | 0.128205128 | 1 | # | 20 | Baseline | aldex2 | 14 |
| # | interaction | 0 | 0 | 4 | 0 | 0 | # | 20 | Baseline | aldex2 | 14 |
| # | disease | 7 | 0 | 0 | 1 | 1 | # | 20 | Small | paralle | 14 |
| # | sex | 4 | 0 | 0 | 1 | 1 | # | 20 | Small | paralle | 14 |
| # | time | 5 | 0 | 0 | 1 | 1 | # | 20 | Small | paralle | 14 |
| # | interaction | 4 | 0 | 0 | 1 | 1 | # | 20 | Small | paralle | 14 |
| # | disease | 7 | 4 | 0 | 0.636363636 | 1 | # | 20 | Small | ancom | 14 |
| # | sex | 4 | 0 | 0 | 1 | 1 | # | 20 | Small | ancom | 14 |
| # | time | 5 | 4 | 0 | 0.555555556 | 1 | # | 20 | Small | ancom | 14 |
| # | interaction | 0 | 0 | 4 | 0 | 0 | # | 20 | Small | ancom | 14 |
| # | disease | 7 | 9 | 0 | 0.4375 | 1 | # | 20 | Small | aldex2 | 14 |
| # | sex | 4 | 1 | 0 | 0.8 | 1 | # | 20 | Small | aldex2 | 14 |
| # | time | 5 | 7 | 0 | 0.416666667 | 1 | # | 20 | Small | aldex2 | 14 |
| # | interaction | 0 | 0 | 4 | 0 | 0 | # | 20 | Small | aldex2 | 14 |
| # | disease | 7 | 10 | 0 | 0.411764706 | 1 | # | 20 | Baseline | paralle | 14 |
| # | sex | 4 | 1 | 0 | 0.8 | 1 | # | 20 | Baseline | paralle | 14 |
| # | time | 5 | 9 | 0 | 0.357142857 | 1 | # | 20 | Baseline | paralle | 14 |
| # | interaction | 4 | 3 | 0 | 0.571428571 | 1 | # | 20 | Baseline | paralle | 14 |
| # | disease | 7 | 4 | 0 | 0.636363636 | 1 | # | 20 | Baseline | ancom | 14 |
| # | sex | 4 | 0 | 0 | 1 | 1 | # | 20 | Baseline | ancom | 14 |
| # | time | 5 | 4 | 0 | 0.555555556 | 1 | # | 20 | Baseline | ancom | 14 |
| # | interaction | 0 | 0 | 4 | 0 | 0 | # | 20 | Baseline | ancom | 14 |
| # | disease | 7 | 93 | 0 | 0.1 | 1 | # | 20 | Baseline | aldex2 | 14 |
| # | sex | 4 | 19 | 0 | 0.173913043 | 1 | # | 20 | Baseline | aldex2 | 14 |
| # | time | 5 | 68 | 0 | 0.094339623 | 1 | # | 20 | Baseline | aldex2 | 14 |
| # | interaction | 0 | 0 | 4 | 0 | 0 | # | 20 | Baseline | aldex2 | 14 |
| # | disease | 7 | 0 | 0 | 1 | 1 | # | 20 | Small | paralle | 14 |
| # | sex | 4 | 0 | 0 | 1 | 1 | # | 20 | Small | paralle | 14 |
| # | time | 5 | 0 | 0 | 1 | 1 | # | 20 | Small | paralle | 14 |
| # | interaction | 4 | 0 | 0 | 1 | 1 | # | 20 | Small | paralle | 14 |
| # | disease | 7 | 4 | 0 | 0.636363636 | 1 | # | 20 | Small | ancom | 14 |
| # | sex | 4 | 0 | 0 | 1 | 1 | # | 20 | Small | ancom | 14 |
| # | time | 5 | 4 | 0 | 0.555555556 | 1 | # | 20 | Small | ancom | 14 |
| # | interaction | 0 | 0 | 4 | 0 | 0 | # | 20 | Small | ancom | 14 |
| # | disease | 7 | 12 | 0 | 0.368421053 | 1 | # | 20 | Small | aldex2 | 14 |
| # | sex | 4 | 1 | 0 | 0.8 | 1 | # | 20 | Small | aldex2 | 14 |
| # | time | 5 | 9 | 0 | 0.357142857 | 1 | # | 20 | Small | aldex2 | 14 |
| # | interaction | 0 | 0 | 4 | 0 | 0 | # | 20 | Small | aldex2 | 14 |
| # | disease | 4 | 5 | 0 | 0.444444444 | 1 | # | 10 | Baseline | paralle | 15 |
| # | sex | 2 | 4 | 0 | 0.333333333 | 1 | # | 10 | Baseline | paralle | 15 |
| # | time | 2 | 12 | 0 | 0.142857143 | 1 | # | 10 | Baseline | paralle | 15 |
| # | interaction | 2 | 7 | 0 | 0.222222222 | 1 | # | 10 | Baseline | paralle | 15 |
| # | disease | 4 | 0 | 0 | 1 | 1 | # | 10 | Baseline | ancom | 15 |

| | | | | | | | | | | | |
|---|---|---|---|---|---|---|---|---|---|---|---|
| # | sex | 2 | 0 | 0 | 1 | 1 | # | 10 | Baseline | ancom | 15 |
| # | time | 2 | 0 | 0 | 1 | 1 | # | 10 | Baseline | ancom | 15 |
| # | interaction | 0 | 0 | 2 | 0 | 0 | # | 10 | Baseline | ancom | 15 |
| # | disease | 4 | 12 | 0 | 0.25 | 1 | # | 10 | Baseline | aldex2 | 15 |
| # | sex | 2 | 5 | 0 | 0.285714286 | 1 | # | 10 | Baseline | aldex2 | 15 |
| # | time | 2 | 8 | 0 | 0.2 | 1 | # | 10 | Baseline | aldex2 | 15 |
| # | interaction | 0 | 0 | 2 | 0 | 0 | # | 10 | Baseline | aldex2 | 15 |
| # | disease | 4 | 1 | 0 | 0.8 | 1 | # | 10 | Small | paralle | 15 |
| # | sex | 2 | 0 | 0 | 1 | 1 | # | 10 | Small | paralle | 15 |
| # | time | 2 | 0 | 0 | 1 | 1 | # | 10 | Small | paralle | 15 |
| # | interaction | 1 | 0 | 1 | 1 | 0.5 | # | 10 | Small | paralle | 15 |
| # | disease | 4 | 0 | 0 | 1 | 1 | # | 10 | Small | ancom | 15 |
| # | sex | 2 | 0 | 0 | 1 | 1 | # | 10 | Small | ancom | 15 |
| # | time | 2 | 0 | 0 | 1 | 1 | # | 10 | Small | ancom | 15 |
| # | interaction | 0 | 0 | 2 | 0 | 0 | # | 10 | Small | ancom | 15 |
| # | disease | 4 | 0 | 0 | 1 | 1 | # | 10 | Small | aldex2 | 15 |
| # | sex | 2 | 0 | 0 | 1 | 1 | # | 10 | Small | aldex2 | 15 |
| # | time | 2 | 1 | 0 | 0.666666667 | 1 | # | 10 | Small | aldex2 | 15 |
| # | interaction | 0 | 0 | 2 | 0 | 0 | # | 10 | Small | aldex2 | 15 |
| # | disease | 4 | 7 | 0 | 0.363636364 | 1 | # | 10 | Baseline | paralle | 15 |
| # | sex | 2 | 7 | 0 | 0.222222222 | 1 | # | 10 | Baseline | paralle | 15 |
| # | time | 2 | 8 | 0 | 0.2 | 1 | # | 10 | Baseline | paralle | 15 |
| # | interaction | 2 | 6 | 0 | 0.25 | 1 | # | 10 | Baseline | paralle | 15 |
| # | disease | 4 | 1 | 0 | 0.8 | 1 | # | 10 | Baseline | ancom | 15 |
| # | sex | 2 | 0 | 0 | 1 | 1 | # | 10 | Baseline | ancom | 15 |
| # | time | 2 | 1 | 0 | 0.666666667 | 1 | # | 10 | Baseline | ancom | 15 |
| # | interaction | 0 | 0 | 2 | 0 | 0 | # | 10 | Baseline | ancom | 15 |
| # | disease | 4 | 19 | 0 | 0.173913043 | 1 | # | 10 | Baseline | aldex2 | 15 |
| # | sex | 2 | 4 | 0 | 0.333333333 | 1 | # | 10 | Baseline | aldex2 | 15 |
| # | time | 2 | 8 | 0 | 0.2 | 1 | # | 10 | Baseline | aldex2 | 15 |
| # | interaction | 0 | 0 | 2 | 0 | 0 | # | 10 | Baseline | aldex2 | 15 |
| # | disease | 4 | 1 | 0 | 0.8 | 1 | # | 10 | Small | paralle | 15 |
| # | sex | 2 | 0 | 0 | 1 | 1 | # | 10 | Small | paralle | 15 |
| # | time | 2 | 0 | 0 | 1 | 1 | # | 10 | Small | paralle | 15 |
| # | interaction | 1 | 0 | 1 | 1 | 0.5 | # | 10 | Small | paralle | 15 |
| # | disease | 4 | 1 | 0 | 0.8 | 1 | # | 10 | Small | ancom | 15 |
| # | sex | 2 | 0 | 0 | 1 | 1 | # | 10 | Small | ancom | 15 |
| # | time | 2 | 1 | 0 | 0.666666667 | 1 | # | 10 | Small | ancom | 15 |
| # | interaction | 0 | 0 | 2 | 0 | 0 | # | 10 | Small | ancom | 15 |
| # | disease | 4 | 1 | 0 | 0.8 | 1 | # | 10 | Small | aldex2 | 15 |
| # | sex | 2 | 0 | 0 | 1 | 1 | # | 10 | Small | aldex2 | 15 |
| # | time | 2 | 1 | 0 | 0.666666667 | 1 | # | 10 | Small | aldex2 | 15 |
| # | interaction | 0 | 0 | 2 | 0 | 0 | # | 10 | Small | aldex2 | 15 |
| # | disease | 4 | 7 | 0 | 0.363636364 | 1 | # | 10 | Baseline | paralle | 15 |
| # | sex | 2 | 6 | 0 | 0.25 | 1 | # | 10 | Baseline | paralle | 15 |
| # | time | 2 | 5 | 0 | 0.285714286 | 1 | # | 10 | Baseline | paralle | 15 |
| # | interaction | 2 | 3 | 0 | 0.4 | 1 | # | 10 | Baseline | paralle | 15 |
| # | disease | 4 | 1 | 0 | 0.8 | 1 | # | 10 | Baseline | ancom | 15 |
| # | sex | 2 | 0 | 0 | 1 | 1 | # | 10 | Baseline | ancom | 15 |
| # | time | 2 | 3 | 0 | 0.4 | 1 | # | 10 | Baseline | ancom | 15 |
| # | interaction | 0 | 0 | 2 | 0 | 0 | # | 10 | Baseline | ancom | 15 |
| # | disease | 4 | 23 | 0 | 0.148148148 | 1 | # | 10 | Baseline | aldex2 | 15 |
| # | sex | 2 | 4 | 0 | 0.333333333 | 1 | # | 10 | Baseline | aldex2 | 15 |
| # | time | 2 | 7 | 0 | 0.222222222 | 1 | # | 10 | Baseline | aldex2 | 15 |
| # | interaction | 0 | 0 | 2 | 0 | 0 | # | 10 | Baseline | aldex2 | 15 |
| # | disease | 4 | 0 | 0 | 1 | 1 | # | 10 | Small | paralle | 15 |
| # | sex | 2 | 0 | 0 | 1 | 1 | # | 10 | Small | paralle | 15 |
| # | time | 2 | 0 | 0 | 1 | 1 | # | 10 | Small | paralle | 15 |
| # | interaction | 1 | 0 | 1 | 1 | 0.5 | # | 10 | Small | paralle | 15 |
| # | disease | 4 | 1 | 0 | 0.8 | 1 | # | 10 | Small | ancom | 15 |
| # | sex | 2 | 0 | 0 | 1 | 1 | # | 10 | Small | ancom | 15 |
| # | time | 2 | 3 | 0 | 0.4 | 1 | # | 10 | Small | ancom | 15 |
| # | interaction | 0 | 0 | 2 | 0 | 0 | # | 10 | Small | ancom | 15 |
| # | disease | 4 | 1 | 0 | 0.8 | 1 | # | 10 | Small | aldex2 | 15 |
| # | sex | 2 | 0 | 0 | 1 | 1 | # | 10 | Small | aldex2 | 15 |
| # | time | 2 | 3 | 0 | 0.4 | 1 | # | 10 | Small | aldex2 | 15 |
| # | interaction | 0 | 0 | 2 | 0 | 0 | # | 10 | Small | aldex2 | 15 |
| # | disease | 4 | 8 | 0 | 0.333333333 | 1 | # | 10 | Baseline | paralle | 15 |
| # | sex | 2 | 7 | 0 | 0.222222222 | 1 | # | 10 | Baseline | paralle | 15 |
| # | time | 2 | 4 | 0 | 0.333333333 | 1 | # | 10 | Baseline | paralle | 15 |
| # | interaction | 2 | 3 | 0 | 0.4 | 1 | # | 10 | Baseline | paralle | 15 |
| # | disease | 4 | 2 | 0 | 0.666666667 | 1 | # | 10 | Baseline | ancom | 15 |
| # | sex | 2 | 0 | 0 | 1 | 1 | # | 10 | Baseline | ancom | 15 |
| # | time | 2 | 2 | 0 | 0.5 | 1 | # | 10 | Baseline | ancom | 15 |
| # | interaction | 0 | 0 | 2 | 0 | 0 | # | 10 | Baseline | ancom | 15 |
| # | disease | 4 | 27 | 0 | 0.129032258 | 1 | # | 10 | Baseline | aldex2 | 15 |
| # | sex | 2 | 6 | 0 | 0.25 | 1 | # | 10 | Baseline | aldex2 | 15 |
| # | time | 2 | 10 | 0 | 0.166666667 | 1 | # | 10 | Baseline | aldex2 | 15 |
| # | interaction | 0 | 0 | 2 | 0 | 0 | # | 10 | Baseline | aldex2 | 15 |
| # | disease | 4 | 0 | 0 | 1 | 1 | # | 10 | Small | paralle | 15 |
| # | sex | 2 | 0 | 0 | 1 | 1 | # | 10 | Small | paralle | 15 |
| # | time | 2 | 0 | 0 | 1 | 1 | # | 10 | Small | paralle | 15 |
| # | interaction | 2 | 0 | 0 | 1 | 1 | # | 10 | Small | paralle | 15 |
| # | disease | 4 | 2 | 0 | 0.666666667 | 1 | # | 10 | Small | ancom | 15 |
| # | sex | 2 | 0 | 0 | 1 | 1 | # | 10 | Small | ancom | 15 |
| # | time | 2 | 2 | 0 | 0.5 | 1 | # | 10 | Small | ancom | 15 |
| # | interaction | 0 | 0 | 2 | 0 | 0 | # | 10 | Small | ancom | 15 |
| # | disease | 4 | 2 | 0 | 0.666666667 | 1 | # | 10 | Small | aldex2 | 15 |
| # | sex | 2 | 0 | 0 | 1 | 1 | # | 10 | Small | aldex2 | 15 |
| # | time | 2 | 2 | 0 | 0.5 | 1 | # | 10 | Small | aldex2 | 15 |
| # | interaction | 0 | 0 | 2 | 0 | 0 | # | 10 | Small | aldex2 | 15 |
| # | disease | 7 | 17 | 0 | 0.291666667 | 1 | # | 20 | Baseline | paralle | 15 |
| # | sex | 4 | 8 | 0 | 0.333333333 | 1 | # | 20 | Baseline | paralle | 15 |
| # | time | 5 | 13 | 0 | 0.277777778 | 1 | # | 20 | Baseline | paralle | 15 |
| # | interaction | 4 | 16 | 0 | 0.2 | 1 | # | 20 | Baseline | paralle | 15 |
| # | disease | 7 | 3 | 0 | 0.7 | 1 | # | 20 | Baseline | ancom | 15 |
| # | sex | 4 | 0 | 0 | 1 | 1 | # | 20 | Baseline | ancom | 15 |
| # | time | 5 | 2 | 0 | 0.714285714 | 1 | # | 20 | Baseline | ancom | 15 |
| # | interaction | 0 | 0 | 4 | 0 | 0 | # | 20 | Baseline | ancom | 15 |
| # | disease | 7 | 15 | 0 | 0.318181818 | 1 | # | 20 | Baseline | aldex2 | 15 |
| # | sex | 4 | 6 | 0 | 0.4 | 1 | # | 20 | Baseline | aldex2 | 15 |
| # | time | 5 | 13 | 0 | 0.277777778 | 1 | # | 20 | Baseline | aldex2 | 15 |
| # | interaction | 0 | 0 | 4 | 0 | 0 | # | 20 | Baseline | aldex2 | 15 |
| # | disease | 7 | 1 | 0 | 0.875 | 1 | # | 20 | Small | paralle | 15 |
| # | sex | 4 | 0 | 0 | 1 | 1 | # | 20 | Small | paralle | 15 |
| # | time | 5 | 0 | 0 | 1 | 1 | # | 20 | Small | paralle | 15 |
| # | interaction | 2 | 1 | 2 | 0.666666667 | 0.5 | # | 20 | Small | paralle | 15 |
| # | disease | 7 | 2 | 0 | 0.777777778 | 1 | # | 20 | Small | ancom | 15 |
| # | sex | 4 | 0 | 0 | 1 | 1 | # | 20 | Small | ancom | 15 |
| # | time | 5 | 2 | 0 | 0.714285714 | 1 | # | 20 | Small | ancom | 15 |
| # | interaction | 0 | 0 | 4 | 0 | 0 | # | 20 | Small | ancom | 15 |
| # | disease | 7 | 3 | 0 | 0.7 | 1 | # | 20 | Small | aldex2 | 15 |
| # | sex | 4 | 0 | 0 | 1 | 1 | # | 20 | Small | aldex2 | 15 |
| # | time | 5 | 1 | 0 | 0.833333333 | 1 | # | 20 | Small | aldex2 | 15 |
| # | interaction | 0 | 0 | 4 | 0 | 0 | # | 20 | Small | aldex2 | 15 |
| # | disease | 7 | 10 | 0 | 0.411764706 | 1 | # | 20 | Baseline | paralle | 15 |
| # | sex | 4 | 11 | 0 | 0.266666667 | 1 | # | 20 | Baseline | paralle | 15 |
| # | time | 5 | 11 | 0 | 0.3125 | 1 | # | 20 | Baseline | paralle | 15 |
| # | interaction | 4 | 12 | 0 | 0.25 | 1 | # | 20 | Baseline | paralle | 15 |
| # | disease | 7 | 4 | 0 | 0.636363636 | 1 | # | 20 | Baseline | ancom | 15 |
| # | sex | 4 | 0 | 0 | 1 | 1 | # | 20 | Baseline | ancom | 15 |
| # | time | 5 | 4 | 0 | 0.555555556 | 1 | # | 20 | Baseline | ancom | 15 |
| # | interaction | 0 | 0 | 4 | 0 | 0 | # | 20 | Baseline | ancom | 15 |
| # | disease | 7 | 31 | 0 | 0.184210526 | 1 | # | 20 | Baseline | aldex2 | 15 |
| # | sex | 4 | 10 | 0 | 0.285714286 | 1 | # | 20 | Baseline | aldex2 | 15 |
| # | time | 5 | 23 | 0 | 0.178571429 | 1 | # | 20 | Baseline | aldex2 | 15 |
| # | interaction | 0 | 0 | 4 | 0 | 0 | # | 20 | Baseline | aldex2 | 15 |
| # | disease | 7 | 0 | 0 | 1 | 1 | # | 20 | Small | paralle | 15 |
| # | sex | 4 | 0 | 0 | 1 | 1 | # | 20 | Small | paralle | 15 |
| # | time | 5 | 0 | 0 | 1 | 1 | # | 20 | Small | paralle | 15 |
| # | interaction | 3 | 0 | 1 | 1 | 0.75 | # | 20 | Small | paralle | 15 |
| # | disease | 7 | 4 | 0 | 0.636363636 | 1 | # | 20 | Small | ancom | 15 |
| # | sex | 4 | 0 | 0 | 1 | 1 | # | 20 | Small | ancom | 15 |
| # | time | 5 | 3 | 0 | 0.625 | 1 | # | 20 | Small | ancom | 15 |
| # | interaction | 0 | 0 | 4 | 0 | 0 | # | 20 | Small | ancom | 15 |
| # | disease | 7 | 7 | 0 | 0.5 | 1 | # | 20 | Small | aldex2 | 15 |
| # | sex | 4 | 1 | 0 | 0.8 | 1 | # | 20 | Small | aldex2 | 15 |
| # | time | 5 | 3 | 0 | 0.625 | 1 | # | 20 | Small | aldex2 | 15 |
| # | interaction | 0 | 0 | 4 | 0 | 0 | # | 20 | Small | aldex2 | 15 |
| # | disease | 7 | 6 | 0 | 0.538461538 | 1 | # | 20 | Baseline | paralle | 15 |
| # | sex | 4 | 6 | 0 | 0.4 | 1 | # | 20 | Baseline | paralle | 15 |
| # | time | 5 | 10 | 0 | 0.333333333 | 1 | # | 20 | Baseline | paralle | 15 |
| # | interaction | 4 | 10 | 0 | 0.285714286 | 1 | # | 20 | Baseline | paralle | 15 |
| # | disease | 7 | 3 | 0 | 0.7 | 1 | # | 20 | Baseline | ancom | 15 |
| # | sex | 4 | 0 | 0 | 1 | 1 | # | 20 | Baseline | ancom | 15 |
| # | time | 5 | 4 | 0 | 0.555555556 | 1 | # | 20 | Baseline | ancom | 15 |
| # | interaction | 0 | 0 | 4 | 0 | 0 | # | 20 | Baseline | ancom | 15 |
| # | disease | 7 | 39 | 0 | 0.152173913 | 1 | # | 20 | Baseline | aldex2 | 15 |
| # | sex | 4 | 13 | 0 | 0.235294118 | 1 | # | 20 | Baseline | aldex2 | 15 |
| # | time | 5 | 30 | 0 | 0.142857143 | 1 | # | 20 | Baseline | aldex2 | 15 |
| # | interaction | 0 | 0 | 4 | 0 | 0 | # | 20 | Baseline | aldex2 | 15 |
| # | disease | 7 | 0 | 0 | 1 | 1 | # | 20 | Small | paralle | 15 |
| # | sex | 4 | 0 | 0 | 1 | 1 | # | 20 | Small | paralle | 15 |
| # | time | 5 | 0 | 0 | 1 | 1 | # | 20 | Small | paralle | 15 |
| # | interaction | 4 | 0 | 0 | 1 | 1 | # | 20 | Small | paralle | 15 |
| # | disease | 7 | 3 | 0 | 0.7 | 1 | # | 20 | Small | ancom | 15 |
| # | sex | 4 | 0 | 0 | 1 | 1 | # | 20 | Small | ancom | 15 |
| # | time | 5 | 3 | 0 | 0.625 | 1 | # | 20 | Small | ancom | 15 |
| # | interaction | 0 | 0 | 4 | 0 | 0 | # | 20 | Small | ancom | 15 |
| # | disease | 7 | 9 | 0 | 0.4375 | 1 | # | 20 | Small | aldex2 | 15 |
| # | sex | 4 | 1 | 0 | 0.8 | 1 | # | 20 | Small | aldex2 | 15 |
| # | time | 5 | 5 | 0 | 0.5 | 1 | # | 20 | Small | aldex2 | 15 |
| # | interaction | 0 | 0 | 4 | 0 | 0 | # | 20 | Small | aldex2 | 15 |
| # | disease | 7 | 7 | 0 | 0.5 | 1 | # | 20 | Baseline | paralle | 15 |
| # | sex | 4 | 5 | 0 | 0.444444444 | 1 | # | 20 | Baseline | paralle | 15 |
| # | time | 5 | 4 | 0 | 0.555555556 | 1 | # | 20 | Baseline | paralle | 15 |
| # | interaction | 4 | 5 | 0 | 0.444444444 | 1 | # | 20 | Baseline | paralle | 15 |
| # | disease | 7 | 4 | 0 | 0.636363636 | 1 | # | 20 | Baseline | ancom | 15 |
| # | sex | 4 | 0 | 0 | 1 | 1 | # | 20 | Baseline | ancom | 15 |
| # | time | 5 | 4 | 0 | 0.555555556 | 1 | # | 20 | Baseline | ancom | 15 |
| # | interaction | 0 | 0 | 4 | 0 | 0 | # | 20 | Baseline | ancom | 15 |
| # | disease | 7 | 50 | 0 | 0.122807018 | 1 | # | 20 | Baseline | aldex2 | 15 |
| # | sex | 4 | 16 | 0 | 0.2 | 1 | # | 20 | Baseline | aldex2 | 15 |
| # | time | 5 | 39 | 0 | 0.113636364 | 1 | # | 20 | Baseline | aldex2 | 15 |
| # | interaction | 0 | 0 | 4 | 0 | 0 | # | 20 | Baseline | aldex2 | 15 |
| # | disease | 7 | 0 | 0 | 1 | 1 | # | 20 | Small | paralle | 15 |
| # | sex | 4 | 0 | 0 | 1 | 1 | # | 20 | Small | paralle | 15 |
| # | time | 5 | 0 | 0 | 1 | 1 | # | 20 | Small | paralle | 15 |
| # | interaction | 4 | 0 | 0 | 1 | 1 | # | 20 | Small | paralle | 15 |
| # | disease | 7 | 4 | 0 | 0.636363636 | 1 | # | 20 | Small | ancom | 15 |
| # | sex | 4 | 0 | 0 | 1 | 1 | # | 20 | Small | ancom | 15 |
| # | time | 5 | 4 | 0 | 0.555555556 | 1 | # | 20 | Small | ancom | 15 |
| # | interaction | 0 | 0 | 4 | 0 | 0 | # | 20 | Small | ancom | 15 |
| # | disease | 7 | 10 | 0 | 0.411764706 | 1 | # | 20 | Small | aldex2 | 15 |
| # | sex | 4 | 2 | 0 | 0.666666667 | 1 | # | 20 | Small | aldex2 | 15 |
| # | time | 5 | 5 | 0 | 0.5 | 1 | # | 20 | Small | aldex2 | 15 |
| # | interaction | 0 | 0 | 4 | 0 | 0 | # | 20 | Small | aldex2 | 15 |
| # | disease | 4 | 10 | 0 | 0.285714286 | 1 | # | 10 | Baseline | paralle | 16 |
| # | sex | 2 | 12 | 0 | 0.142857143 | 1 | # | 10 | Baseline | paralle | 16 |
| # | time | 2 | 9 | 0 | 0.181818182 | 1 | # | 10 | Baseline | paralle | 16 |
| # | interaction | 2 | 10 | 0 | 0.166666667 | 1 | # | 10 | Baseline | paralle | 16 |
| # | disease | 4 | 0 | 0 | 1 | 1 | # | 10 | Baseline | ancom | 16 |
| # | sex | 2 | 0 | 0 | 1 | 1 | # | 10 | Baseline | ancom | 16 |
| # | time | 2 | 1 | 0 | 0.666666667 | 1 | # | 10 | Baseline | ancom | 16 |
| # | interaction | 0 | 0 | 2 | 0 | 0 | # | 10 | Baseline | ancom | 16 |
| # | disease | 4 | 14 | 0 | 0.222222222 | 1 | # | 10 | Baseline | aldex2 | 16 |
| # | sex | 2 | 6 | 0 | 0.25 | 1 | # | 10 | Baseline | aldex2 | 16 |
| # | time | 2 | 5 | 0 | 0.285714286 | 1 | # | 10 | Baseline | aldex2 | 16 |
| # | interaction | 0 | 0 | 2 | 0 | 0 | # | 10 | Baseline | aldex2 | 16 |
| # | disease | 4 | 0 | 0 | 1 | 1 | # | 10 | Small | paralle | 16 |
| # | sex | 2 | 0 | 0 | 1 | 1 | # | 10 | Small | paralle | 16 |
| # | time | 2 | 0 | 0 | 1 | 1 | # | 10 | Small | paralle | 16 |
| # | interaction | 1 | 1 | 1 | 0.5 | 0.5 | # | 10 | Small | paralle | 16 |
| # | disease | 4 | 0 | 0 | 1 | 1 | # | 10 | Small | ancom | 16 |
| # | sex | 2 | 0 | 0 | 1 | 1 | # | 10 | Small | ancom | 16 |
| # | time | 2 | 1 | 0 | 0.666666667 | 1 | # | 10 | Small | ancom | 16 |
| # | interaction | 0 | 0 | 2 | 0 | 0 | # | 10 | Small | ancom | 16 |
| # | disease | 4 | 1 | 0 | 0.8 | 1 | # | 10 | Small | aldex2 | 16 |
| # | sex | 2 | 1 | 0 | 0.666666667 | 1 | # | 10 | Small | aldex2 | 16 |
| # | time | 2 | 2 | 0 | 0.5 | 1 | # | 10 | Small | aldex2 | 16 |
| # | interaction | 0 | 0 | 2 | 0 | 0 | # | 10 | Small | aldex2 | 16 |
| # | disease | 4 | 10 | 0 | 0.285714286 | 1 | # | 10 | Baseline | paralle | 16 |
| # | sex | 2 | 4 | 0 | 0.333333333 | 1 | # | 10 | Baseline | paralle | 16 |
| # | time | 2 | 3 | 0 | 0.4 | 1 | # | 10 | Baseline | paralle | 16 |
| # | interaction | 2 | 6 | 0 | 0.25 | 1 | # | 10 | Baseline | paralle | 16 |
| # | disease | 4 | 1 | 0 | 0.8 | 1 | # | 10 | Baseline | ancom | 16 |
| # | sex | 2 | 0 | 0 | 1 | 1 | # | 10 | Baseline | ancom | 16 |
| # | time | 2 | 2 | 0 | 0.5 | 1 | # | 10 | Baseline | ancom | 16 |
| # | interaction | 0 | 0 | 2 | 0 | 0 | # | 10 | Baseline | ancom | 16 |
| # | disease | 4 | 16 | 0 | 0.2 | 1 | # | 10 | Baseline | aldex2 | 16 |
| # | sex | 2 | 4 | 0 | 0.333333333 | 1 | # | 10 | Baseline | aldex2 | 16 |
| # | time | 2 | 6 | 0 | 0.25 | 1 | # | 10 | Baseline | aldex2 | 16 |
| # | interaction | 0 | 0 | 2 | 0 | 0 | # | 10 | Baseline | aldex2 | 16 |
| # | disease | 4 | 0 | 0 | 1 | 1 | # | 10 | Small | paralle | 16 |
| # | sex | 2 | 0 | 0 | 1 | 1 | # | 10 | Small | paralle | 16 |
| # | time | 2 | 0 | 0 | 1 | 1 | # | 10 | Small | paralle | 16 |
| # | interaction | 2 | 0 | 0 | 1 | 1 | # | 10 | Small | paralle | 16 |
| # | disease | 4 | 1 | 0 | 0.8 | 1 | # | 10 | Small | ancom | 16 |
| # | sex | 2 | 0 | 0 | 1 | 1 | # | 10 | Small | ancom | 16 |
| # | time | 2 | 2 | 0 | 0.5 | 1 | # | 10 | Small | ancom | 16 |
| # | interaction | 0 | 0 | 2 | 0 | 0 | # | 10 | Small | ancom | 16 |
| # | disease | 4 | 1 | 0 | 0.8 | 1 | # | 10 | Small | aldex2 | 16 |
| # | sex | 2 | 1 | 0 | 0.666666667 | 1 | # | 10 | Small | aldex2 | 16 |
| # | time | 2 | 2 | 0 | 0.5 | 1 | # | 10 | Small | aldex2 | 16 |
| # | interaction | 0 | 0 | 2 | 0 | 0 | # | 10 | Small | aldex2 | 16 |
| # | disease | 4 | 8 | 0 | 0.333333333 | 1 | # | 10 | Baseline | paralle | 16 |
| # | sex | 2 | 2 | 0 | 0.5 | 1 | # | 10 | Baseline | paralle | 16 |
| # | time | 2 | 3 | 0 | 0.4 | 1 | # | 10 | Baseline | paralle | 16 |
| # | interaction | 2 | 5 | 0 | 0.285714286 | 1 | # | 10 | Baseline | paralle | 16 |
| # | disease | 4 | 2 | 0 | 0.666666667 | 1 | # | 10 | Baseline | ancom | 16 |
| # | sex | 2 | 0 | 0 | 1 | 1 | # | 10 | Baseline | ancom | 16 |
| # | time | 2 | 2 | 0 | 0.5 | 1 | # | 10 | Baseline | ancom | 16 |
| # | interaction | 0 | 0 | 2 | 0 | 0 | # | 10 | Baseline | ancom | 16 |
| # | disease | 4 | 19 | 0 | 0.173913043 | 1 | # | 10 | Baseline | aldex2 | 16 |
| # | sex | 2 | 9 | 0 | 0.181818182 | 1 | # | 10 | Baseline | aldex2 | 16 |
| # | time | 2 | 7 | 0 | 0.222222222 | 1 | # | 10 | Baseline | aldex2 | 16 |
| # | interaction | 0 | 0 | 2 | 0 | 0 | # | 10 | Baseline | aldex2 | 16 |
| # | disease | 4 | 0 | 0 | 1 | 1 | # | 10 | Small | paralle | 16 |
| # | sex | 2 | 0 | 0 | 1 | 1 | # | 10 | Small | paralle | 16 |
| # | time | 2 | 0 | 0 | 1 | 1 | # | 10 | Small | paralle | 16 |
| # | interaction | 2 | 0 | 0 | 1 | 1 | # | 10 | Small | paralle | 16 |
| # | disease | 4 | 2 | 0 | 0.666666667 | 1 | # | 10 | Small | ancom | 16 |
| # | sex | 2 | 0 | 0 | 1 | 1 | # | 10 | Small | ancom | 16 |
| # | time | 2 | 2 | 0 | 0.5 | 1 | # | 10 | Small | ancom | 16 |
| # | interaction | 0 | 0 | 2 | 0 | 0 | # | 10 | Small | ancom | 16 |
| # | disease | 4 | 1 | 0 | 0.8 | 1 | # | 10 | Small | aldex2 | 16 |
| # | sex | 2 | 1 | 0 | 0.666666667 | 1 | # | 10 | Small | aldex2 | 16 |
| # | time | 2 | 2 | 0 | 0.5 | 1 | # | 10 | Small | aldex2 | 16 |
| # | interaction | 0 | 0 | 2 | 0 | 0 | # | 10 | Small | aldex2 | 16 |
| # | disease | 4 | 5 | 0 | 0.444444444 | 1 | # | 10 | Baseline | paralle | 16 |
| # | sex | 2 | 1 | 0 | 0.666666667 | 1 | # | 10 | Baseline | paralle | 16 |
| # | time | 2 | 3 | 0 | 0.4 | 1 | # | 10 | Baseline | paralle | 16 |
| # | interaction | 2 | 3 | 0 | 0.4 | 1 | # | 10 | Baseline | paralle | 16 |
| # | disease | 4 | 2 | 0 | 0.666666667 | 1 | # | 10 | Baseline | ancom | 16 |
| # | sex | 2 | 0 | 0 | 1 | 1 | # | 10 | Baseline | ancom | 16 |
| # | time | 2 | 2 | 0 | 0.5 | 1 | # | 10 | Baseline | ancom | 16 |
| # | interaction | 0 | 0 | 2 | 0 | 0 | # | 10 | Baseline | ancom | 16 |
| # | disease | 4 | 20 | 0 | 0.166666667 | 1 | # | 10 | Baseline | aldex2 | 16 |
| # | sex | 2 | 7 | 0 | 0.222222222 | 1 | # | 10 | Baseline | aldex2 | 16 |
| # | time | 2 | 8 | 0 | 0.2 | 1 | # | 10 | Baseline | aldex2 | 16 |
| # | interaction | 0 | 0 | 2 | 0 | 0 | # | 10 | Baseline | aldex2 | 16 |
| # | disease | 4 | 0 | 0 | 1 | 1 | # | 10 | Small | paralle | 16 |
| # | sex | 2 | 0 | 0 | 1 | 1 | # | 10 | Small | paralle | 16 |
| # | time | 2 | 0 | 0 | 1 | 1 | # | 10 | Small | paralle | 16 |
| # | interaction | 2 | 0 | 0 | 1 | 1 | # | 10 | Small | paralle | 16 |
| # | disease | 4 | 2 | 0 | 0.666666667 | 1 | # | 10 | Small | ancom | 16 |
| # | sex | 2 | 0 | 0 | 1 | 1 | # | 10 | Small | ancom | 16 |
| # | time | 2 | 2 | 0 | 0.5 | 1 | # | 10 | Small | ancom | 16 |
| # | interaction | 0 | 0 | 2 | 0 | 0 | # | 10 | Small | ancom | 16 |
| # | disease | 4 | 2 | 0 | 0.666666667 | 1 | # | 10 | Small | aldex2 | 16 |
| # | sex | 2 | 1 | 0 | 0.666666667 | 1 | # | 10 | Small | aldex2 | 16 |
| # | time | 2 | 2 | 0 | 0.5 | 1 | # | 10 | Small | aldex2 | 16 |
| # | interaction | 0 | 0 | 2 | 0 | 0 | # | 10 | Small | aldex2 | 16 |
| # | disease | 7 | 16 | 0 | 0.304347826 | 1 | # | 20 | Baseline | paralle | 16 |
| # | sex | 4 | 5 | 0 | 0.444444444 | 1 | # | 20 | Baseline | paralle | 16 |
| # | time | 5 | 9 | 0 | 0.357142857 | 1 | # | 20 | Baseline | paralle | 16 |
| # | interaction | 4 | 8 | 0 | 0.333333333 | 1 | # | 20 | Baseline | paralle | 16 |
| # | disease | 7 | 1 | 0 | 0.875 | 1 | # | 20 | Baseline | ancom | 16 |
| # | sex | 4 | 0 | 0 | 1 | 1 | # | 20 | Baseline | ancom | 16 |
| # | time | 5 | 2 | 0 | 0.714285714 | 1 | # | 20 | Baseline | ancom | 16 |
| # | interaction | 0 | 0 | 4 | 0 | 0 | # | 20 | Baseline | ancom | 16 |
| # | disease | 7 | 22 | 0 | 0.24137931 | 1 | # | 20 | Baseline | aldex2 | 16 |
| # | sex | 4 | 13 | 0 | 0.235294118 | 1 | # | 20 | Baseline | aldex2 | 16 |
| # | time | 5 | 16 | 0 | 0.238095238 | 1 | # | 20 | Baseline | aldex2 | 16 |
| # | interaction | 0 | 0 | 4 | 0 | 0 | # | 20 | Baseline | aldex2 | 16 |
| # | disease | 7 | 0 | 0 | 1 | 1 | # | 20 | Small | paralle | 16 |
| # | sex | 4 | 0 | 0 | 1 | 1 | # | 20 | Small | paralle | 16 |
| # | time | 5 | 0 | 0 | 1 | 1 | # | 20 | Small | paralle | 16 |
| # | interaction | 3 | 0 | 1 | 1 | 0.75 | # | 20 | Small | paralle | 16 |
| # | disease | 7 | 0 | 0 | 1 | 1 | # | 20 | Small | ancom | 16 |
| # | sex | 4 | 0 | 0 | 1 | 1 | # | 20 | Small | ancom | 16 |
| # | time | 5 | 1 | 0 | 0.833333333 | 1 | # | 20 | Small | ancom | 16 |
| # | interaction | 0 | 0 | 4 | 0 | 0 | # | 20 | Small | ancom | 16 |
| # | disease | 7 | 2 | 0 | 0.777777778 | 1 | # | 20 | Small | aldex2 | 16 |
| # | sex | 4 | 2 | 0 | 0.666666667 | 1 | # | 20 | Small | aldex2 | 16 |
| # | time | 5 | 2 | 0 | 0.714285714 | 1 | # | 20 | Small | aldex2 | 16 |
| # | interaction | 0 | 0 | 4 | 0 | 0 | # | 20 | Small | aldex2 | 16 |
| # | disease | 7 | 6 | 0 | 0.538461538 | 1 | # | 20 | Baseline | paralle | 16 |
| # | sex | 4 | 5 | 0 | 0.444444444 | 1 | # | 20 | Baseline | paralle | 16 |
| # | time | 5 | 5 | 0 | 0.5 | 1 | # | 20 | Baseline | paralle | 16 |
| # | interaction | 4 | 1 | 0 | 0.8 | 1 | # | 20 | Baseline | paralle | 16 |
| # | disease | 7 | 3 | 0 | 0.7 | 1 | # | 20 | Baseline | ancom | 16 |
| # | sex | 4 | 0 | 0 | 1 | 1 | # | 20 | Baseline | ancom | 16 |
| # | time | 5 | 4 | 0 | 0.555555556 | 1 | # | 20 | Baseline | ancom | 16 |
| # | interaction | 0 | 0 | 4 | 0 | 0 | # | 20 | Baseline | ancom | 16 |
| # | disease | 7 | 35 | 0 | 0.166666667 | 1 | # | 20 | Baseline | aldex2 | 16 |
| # | sex | 4 | 10 | 0 | 0.285714286 | 1 | # | 20 | Baseline | aldex2 | 16 |
| # | time | 5 | 22 | 0 | 0.185185185 | 1 | # | 20 | Baseline | aldex2 | 16 |
| # | interaction | 0 | 0 | 4 | 0 | 0 | # | 20 | Baseline | aldex2 | 16 |
| # | disease | 7 | 0 | 0 | 1 | 1 | # | 20 | Small | paralle | 16 |
| # | sex | 4 | 0 | 0 | 1 | 1 | # | 20 | Small | paralle | 16 |
| # | time | 5 | 0 | 0 | 1 | 1 | # | 20 | Small | paralle | 16 |
| # | interaction | 4 | 0 | 0 | 1 | 1 | # | 20 | Small | paralle | 16 |
| # | disease | 7 | 3 | 0 | 0.7 | 1 | # | 20 | Small | ancom | 16 |
| # | sex | 4 | 0 | 0 | 1 | 1 | # | 20 | Small | ancom | 16 |
| # | time | 5 | 4 | 0 | 0.555555556 | 1 | # | 20 | Small | ancom | 16 |
| # | interaction | 0 | 0 | 4 | 0 | 0 | # | 20 | Small | ancom | 16 |
| # | disease | 7 | 4 | 0 | 0.636363636 | 1 | # | 20 | Small | aldex2 | 16 |
| # | sex | 4 | 1 | 0 | 0.8 | 1 | # | 20 | Small | aldex2 | 16 |
| # | time | 5 | 4 | 0 | 0.555555556 | 1 | # | 20 | Small | aldex2 | 16 |
| # | interaction | 0 | 0 | 4 | 0 | 0 | # | 20 | Small | aldex2 | 16 |
| # | disease | 7 | 10 | 0 | 0.411764706 | 1 | # | 20 | Baseline | paralle | 16 |
| # | sex | 4 | 3 | 0 | 0.571428571 | 1 | # | 20 | Baseline | paralle | 16 |
| # | time | 5 | 7 | 0 | 0.416666667 | 1 | # | 20 | Baseline | paralle | 16 |
| # | interaction | 4 | 3 | 0 | 0.571428571 | 1 | # | 20 | Baseline | paralle | 16 |
| # | disease | 7 | 4 | 0 | 0.636363636 | 1 | # | 20 | Baseline | ancom | 16 |
| # | sex | 4 | 0 | 0 | 1 | 1 | # | 20 | Baseline | ancom | 16 |

| | | | | | | | | | | | |
|---|---|---|---|---|---|---|---|---|---|---|---|
| # | time | 5 | 4 | 0 | 0.555555556 | 1 | # | 20 | Baseline | ancom | 16 |
| # | interaction | 0 | 0 | 4 | 0 | 0 | # | 20 | Baseline | ancom | 16 |
| # | disease | 7 | 46 | 0 | 0.132075472 | 1 | # | 20 | Baseline | aldex2 | 16 |
| # | sex | 4 | 15 | 0 | 0.210526316 | 1 | # | 20 | Baseline | aldex2 | 16 |
| # | time | 5 | 33 | 0 | 0.131578947 | 1 | # | 20 | Baseline | aldex2 | 16 |
| # | interaction | 0 | 0 | 4 | 0 | 0 | # | 20 | Baseline | aldex2 | 16 |
| # | disease | 7 | 0 | 0 | 1 | 1 | # | 20 | Small | paralle | 16 |
| # | sex | 4 | 0 | 0 | 1 | 1 | # | 20 | Small | paralle | 16 |
| # | time | 5 | 0 | 0 | 1 | 1 | # | 20 | Small | paralle | 16 |
| # | interaction | 4 | 0 | 0 | 1 | 1 | # | 20 | Small | paralle | 16 |
| # | disease | 7 | 4 | 0 | 0.636363636 | 1 | # | 20 | Small | ancom | 16 |
| # | sex | 4 | 0 | 0 | 1 | 1 | # | 20 | Small | ancom | 16 |
| # | time | 5 | 4 | 0 | 0.555555556 | 1 | # | 20 | Small | ancom | 16 |
| # | interaction | 0 | 0 | 4 | 0 | 0 | # | 20 | Small | ancom | 16 |
| # | disease | 7 | 6 | 0 | 0.538461538 | 1 | # | 20 | Small | aldex2 | 16 |
| # | sex | 4 | 0 | 0 | 1 | 1 | # | 20 | Small | aldex2 | 16 |
| # | time | 5 | 5 | 0 | 0.5 | 1 | # | 20 | Small | aldex2 | 16 |
| # | interaction | 0 | 0 | 4 | 0 | 0 | # | 20 | Small | aldex2 | 16 |
| # | disease | 7 | 14 | 0 | 0.333333333 | 1 | # | 20 | Baseline | paralle | 16 |
| # | sex | 4 | 7 | 0 | 0.363636364 | 1 | # | 20 | Baseline | paralle | 16 |
| # | time | 5 | 6 | 0 | 0.454545455 | 1 | # | 20 | Baseline | paralle | 16 |
| # | interaction | 4 | 3 | 0 | 0.571428571 | 1 | # | 20 | Baseline | paralle | 16 |
| # | disease | 7 | 5 | 0 | 0.583333333 | 1 | # | 20 | Baseline | ancom | 16 |
| # | sex | 4 | 0 | 0 | 1 | 1 | # | 20 | Baseline | ancom | 16 |
| # | time | 5 | 4 | 0 | 0.555555556 | 1 | # | 20 | Baseline | ancom | 16 |
| # | interaction | 0 | 0 | 4 | 0 | 0 | # | 20 | Baseline | ancom | 16 |
| # | disease | 7 | 56 | 0 | 0.111111111 | 1 | # | 20 | Baseline | aldex2 | 16 |
| # | sex | 4 | 23 | 0 | 0.148148148 | 1 | # | 20 | Baseline | aldex2 | 16 |
| # | time | 5 | 38 | 0 | 0.11627907 | 1 | # | 20 | Baseline | aldex2 | 16 |
| # | interaction | 0 | 0 | 4 | 0 | 0 | # | 20 | Baseline | aldex2 | 16 |
| # | disease | 7 | 0 | 0 | 1 | 1 | # | 20 | Small | paralle | 16 |
| # | sex | 4 | 0 | 0 | 1 | 1 | # | 20 | Small | paralle | 16 |
| # | time | 5 | 1 | 0 | 0.833333333 | 1 | # | 20 | Small | paralle | 16 |
| # | interaction | 4 | 0 | 0 | 1 | 1 | # | 20 | Small | paralle | 16 |
| # | disease | 7 | 4 | 0 | 0.636363636 | 1 | # | 20 | Small | ancom | 16 |
| # | sex | 4 | 0 | 0 | 1 | 1 | # | 20 | Small | ancom | 16 |
| # | time | 5 | 4 | 0 | 0.555555556 | 1 | # | 20 | Small | ancom | 16 |
| # | interaction | 0 | 0 | 4 | 0 | 0 | # | 20 | Small | ancom | 16 |
| # | disease | 7 | 10 | 0 | 0.411764706 | 1 | # | 20 | Small | aldex2 | 16 |
| # | sex | 4 | 0 | 0 | 1 | 1 | # | 20 | Small | aldex2 | 16 |
| # | time | 5 | 5 | 0 | 0.5 | 1 | # | 20 | Small | aldex2 | 16 |
| # | interaction | 0 | 0 | 4 | 0 | 0 | # | 20 | Small | aldex2 | 16 |
| # | disease | 4 | 5 | 0 | 0.444444444 | 1 | # | 10 | Baseline | paralle | 17 |
| # | sex | 2 | 7 | 0 | 0.222222222 | 1 | # | 10 | Baseline | paralle | 17 |
| # | time | 2 | 8 | 0 | 0.2 | 1 | # | 10 | Baseline | paralle | 17 |
| # | interaction | 2 | 10 | 0 | 0.166666667 | 1 | # | 10 | Baseline | paralle | 17 |
| # | disease | 4 | 0 | 0 | 1 | 1 | # | 10 | Baseline | ancom | 17 |
| # | sex | 2 | 0 | 0 | 1 | 1 | # | 10 | Baseline | ancom | 17 |
| # | time | 2 | 0 | 0 | 1 | 1 | # | 10 | Baseline | ancom | 17 |
| # | interaction | 0 | 0 | 2 | 0 | 0 | # | 10 | Baseline | ancom | 17 |
| # | disease | 4 | 9 | 0 | 0.307692308 | 1 | # | 10 | Baseline | aldex2 | 17 |
| # | sex | 2 | 5 | 0 | 0.285714286 | 1 | # | 10 | Baseline | aldex2 | 17 |
| # | time | 2 | 5 | 0 | 0.285714286 | 1 | # | 10 | Baseline | aldex2 | 17 |
| # | interaction | 0 | 0 | 2 | 0 | 0 | # | 10 | Baseline | aldex2 | 17 |
| # | disease | 4 | 0 | 0 | 1 | 1 | # | 10 | Small | paralle | 17 |
| # | sex | 2 | 0 | 0 | 1 | 1 | # | 10 | Small | paralle | 17 |
| # | time | 2 | 0 | 0 | 1 | 1 | # | 10 | Small | paralle | 17 |
| # | interaction | 1 | 2 | 1 | 0.333333333 | 0.5 | # | 10 | Small | paralle | 17 |
| # | disease | 4 | 0 | 0 | 1 | 1 | # | 10 | Small | ancom | 17 |
| # | sex | 2 | 0 | 0 | 1 | 1 | # | 10 | Small | ancom | 17 |
| # | time | 2 | 0 | 0 | 1 | 1 | # | 10 | Small | ancom | 17 |
| # | interaction | 0 | 0 | 2 | 0 | 0 | # | 10 | Small | ancom | 17 |
| # | disease | 4 | 0 | 0 | 1 | 1 | # | 10 | Small | aldex2 | 17 |
| # | sex | 2 | 0 | 0 | 1 | 1 | # | 10 | Small | aldex2 | 17 |
| # | time | 2 | 0 | 0 | 1 | 1 | # | 10 | Small | aldex2 | 17 |
| # | interaction | 0 | 0 | 2 | 0 | 0 | # | 10 | Small | aldex2 | 17 |
| # | disease | 4 | 10 | 0 | 0.285714286 | 1 | # | 10 | Baseline | paralle | 17 |
| # | sex | 2 | 3 | 0 | 0.4 | 1 | # | 10 | Baseline | paralle | 17 |
| # | time | 2 | 10 | 0 | 0.166666667 | 1 | # | 10 | Baseline | paralle | 17 |
| # | interaction | 2 | 8 | 0 | 0.2 | 1 | # | 10 | Baseline | paralle | 17 |
| # | disease | 4 | 2 | 0 | 0.666666667 | 1 | # | 10 | Baseline | ancom | 17 |
| # | sex | 2 | 0 | 0 | 1 | 1 | # | 10 | Baseline | ancom | 17 |
| # | time | 2 | 1 | 0 | 0.666666667 | 1 | # | 10 | Baseline | ancom | 17 |
| # | interaction | 0 | 0 | 2 | 0 | 0 | # | 10 | Baseline | ancom | 17 |
| # | disease | 4 | 17 | 0 | 0.19047619 | 1 | # | 10 | Baseline | aldex2 | 17 |
| # | sex | 2 | 7 | 0 | 0.222222222 | 1 | # | 10 | Baseline | aldex2 | 17 |
| # | time | 2 | 12 | 0 | 0.142857143 | 1 | # | 10 | Baseline | aldex2 | 17 |
| # | interaction | 0 | 0 | 2 | 0 | 0 | # | 10 | Baseline | aldex2 | 17 |
| # | disease | 4 | 0 | 0 | 1 | 1 | # | 10 | Small | paralle | 17 |
| # | sex | 2 | 0 | 0 | 1 | 1 | # | 10 | Small | paralle | 17 |
| # | time | 2 | 0 | 0 | 1 | 1 | # | 10 | Small | paralle | 17 |
| # | interaction | 2 | 1 | 0 | 0.666666667 | 1 | # | 10 | Small | paralle | 17 |
| # | disease | 4 | 2 | 0 | 0.666666667 | 1 | # | 10 | Small | ancom | 17 |
| # | sex | 2 | 0 | 0 | 1 | 1 | # | 10 | Small | ancom | 17 |
| # | time | 2 | 1 | 0 | 0.666666667 | 1 | # | 10 | Small | ancom | 17 |
| # | interaction | 0 | 0 | 2 | 0 | 0 | # | 10 | Small | ancom | 17 |
| # | disease | 4 | 1 | 0 | 0.8 | 1 | # | 10 | Small | aldex2 | 17 |
| # | sex | 2 | 0 | 0 | 1 | 1 | # | 10 | Small | aldex2 | 17 |
| # | time | 2 | 1 | 0 | 0.666666667 | 1 | # | 10 | Small | aldex2 | 17 |
| # | interaction | 0 | 0 | 2 | 0 | 0 | # | 10 | Small | aldex2 | 17 |
| # | disease | 4 | 5 | 0 | 0.444444444 | 1 | # | 10 | Baseline | paralle | 17 |
| # | sex | 2 | 4 | 0 | 0.333333333 | 1 | # | 10 | Baseline | paralle | 17 |
| # | time | 2 | 5 | 0 | 0.285714286 | 1 | # | 10 | Baseline | paralle | 17 |
| # | interaction | 2 | 5 | 0 | 0.285714286 | 1 | # | 10 | Baseline | paralle | 17 |
| # | disease | 4 | 2 | 0 | 0.666666667 | 1 | # | 10 | Baseline | ancom | 17 |
| # | sex | 2 | 0 | 0 | 1 | 1 | # | 10 | Baseline | ancom | 17 |
| # | time | 2 | 2 | 0 | 0.5 | 1 | # | 10 | Baseline | ancom | 17 |
| # | interaction | 0 | 0 | 2 | 0 | 0 | # | 10 | Baseline | ancom | 17 |
| # | disease | 4 | 21 | 0 | 0.16 | 1 | # | 10 | Baseline | aldex2 | 17 |
| # | sex | 2 | 8 | 0 | 0.2 | 1 | # | 10 | Baseline | aldex2 | 17 |
| # | time | 2 | 8 | 0 | 0.2 | 1 | # | 10 | Baseline | aldex2 | 17 |
| # | interaction | 0 | 0 | 2 | 0 | 0 | # | 10 | Baseline | aldex2 | 17 |
| # | disease | 4 | 0 | 0 | 1 | 1 | # | 10 | Small | paralle | 17 |
| # | sex | 2 | 0 | 0 | 1 | 1 | # | 10 | Small | paralle | 17 |
| # | time | 2 | 0 | 0 | 1 | 1 | # | 10 | Small | paralle | 17 |
| # | interaction | 1 | 0 | 1 | 1 | 0.5 | # | 10 | Small | paralle | 17 |
| # | disease | 4 | 2 | 0 | 0.666666667 | 1 | # | 10 | Small | ancom | 17 |
| # | sex | 2 | 0 | 0 | 1 | 1 | # | 10 | Small | ancom | 17 |
| # | time | 2 | 2 | 0 | 0.5 | 1 | # | 10 | Small | ancom | 17 |
| # | interaction | 0 | 0 | 2 | 0 | 0 | # | 10 | Small | ancom | 17 |
| # | disease | 4 | 2 | 0 | 0.666666667 | 1 | # | 10 | Small | aldex2 | 17 |
| # | sex | 2 | 0 | 0 | 1 | 1 | # | 10 | Small | aldex2 | 17 |
| # | time | 2 | 2 | 0 | 0.5 | 1 | # | 10 | Small | aldex2 | 17 |
| # | interaction | 0 | 0 | 2 | 0 | 0 | # | 10 | Small | aldex2 | 17 |
| # | disease | 4 | 8 | 0 | 0.333333333 | 1 | # | 10 | Baseline | paralle | 17 |
| # | sex | 2 | 2 | 0 | 0.5 | 1 | # | 10 | Baseline | paralle | 17 |
| # | time | 2 | 4 | 0 | 0.333333333 | 1 | # | 10 | Baseline | paralle | 17 |
| # | interaction | 2 | 5 | 0 | 0.285714286 | 1 | # | 10 | Baseline | paralle | 17 |
| # | disease | 4 | 2 | 0 | 0.666666667 | 1 | # | 10 | Baseline | ancom | 17 |
| # | sex | 2 | 0 | 0 | 1 | 1 | # | 10 | Baseline | ancom | 17 |
| # | time | 2 | 2 | 0 | 0.5 | 1 | # | 10 | Baseline | ancom | 17 |
| # | interaction | 0 | 0 | 2 | 0 | 0 | # | 10 | Baseline | ancom | 17 |
| # | disease | 4 | 29 | 0 | 0.121212121 | 1 | # | 10 | Baseline | aldex2 | 17 |
| # | sex | 2 | 7 | 0 | 0.222222222 | 1 | # | 10 | Baseline | aldex2 | 17 |
| # | time | 2 | 14 | 0 | 0.125 | 1 | # | 10 | Baseline | aldex2 | 17 |
| # | interaction | 0 | 0 | 2 | 0 | 0 | # | 10 | Baseline | aldex2 | 17 |
| # | disease | 4 | 0 | 0 | 1 | 1 | # | 10 | Small | paralle | 17 |
| # | sex | 2 | 0 | 0 | 1 | 1 | # | 10 | Small | paralle | 17 |
| # | time | 2 | 0 | 0 | 1 | 1 | # | 10 | Small | paralle | 17 |
| # | interaction | 1 | 0 | 1 | 1 | 0.5 | # | 10 | Small | paralle | 17 |
| # | disease | 4 | 2 | 0 | 0.666666667 | 1 | # | 10 | Small | ancom | 17 |
| # | sex | 2 | 0 | 0 | 1 | 1 | # | 10 | Small | ancom | 17 |
| # | time | 2 | 2 | 0 | 0.5 | 1 | # | 10 | Small | ancom | 17 |
| # | interaction | 0 | 0 | 2 | 0 | 0 | # | 10 | Small | ancom | 17 |
| # | disease | 4 | 2 | 0 | 0.666666667 | 1 | # | 10 | Small | aldex2 | 17 |
| # | sex | 2 | 0 | 0 | 1 | 1 | # | 10 | Small | aldex2 | 17 |
| # | time | 2 | 2 | 0 | 0.5 | 1 | # | 10 | Small | aldex2 | 17 |
| # | interaction | 0 | 0 | 2 | 0 | 0 | # | 10 | Small | aldex2 | 17 |
| # | disease | 7 | 16 | 0 | 0.304347826 | 1 | # | 20 | Baseline | paralle | 17 |
| # | sex | 4 | 11 | 0 | 0.266666667 | 1 | # | 20 | Baseline | paralle | 17 |
| # | time | 5 | 11 | 0 | 0.3125 | 1 | # | 20 | Baseline | paralle | 17 |
| # | interaction | 4 | 9 | 0 | 0.307692308 | 1 | # | 20 | Baseline | paralle | 17 |
| # | disease | 7 | 0 | 0 | 1 | 1 | # | 20 | Baseline | ancom | 17 |
| # | sex | 4 | 0 | 0 | 1 | 1 | # | 20 | Baseline | ancom | 17 |
| # | time | 5 | 1 | 0 | 0.833333333 | 1 | # | 20 | Baseline | ancom | 17 |
| # | interaction | 0 | 0 | 4 | 0 | 0 | # | 20 | Baseline | ancom | 17 |
| # | disease | 7 | 16 | 0 | 0.304347826 | 1 | # | 20 | Baseline | aldex2 | 17 |
| # | sex | 4 | 7 | 0 | 0.363636364 | 1 | # | 20 | Baseline | aldex2 | 17 |
| # | time | 5 | 10 | 0 | 0.333333333 | 1 | # | 20 | Baseline | aldex2 | 17 |
| # | interaction | 0 | 0 | 4 | 0 | 0 | # | 20 | Baseline | aldex2 | 17 |
| # | disease | 7 | 0 | 0 | 1 | 1 | # | 20 | Small | paralle | 17 |
| # | sex | 4 | 0 | 0 | 1 | 1 | # | 20 | Small | paralle | 17 |
| # | time | 5 | 0 | 0 | 1 | 1 | # | 20 | Small | paralle | 17 |
| # | interaction | 1 | 0 | 3 | 1 | 0.25 | # | 20 | Small | paralle | 17 |
| # | disease | 7 | 0 | 0 | 1 | 1 | # | 20 | Small | ancom | 17 |
| # | sex | 4 | 0 | 0 | 1 | 1 | # | 20 | Small | ancom | 17 |
| # | time | 5 | 1 | 0 | 0.833333333 | 1 | # | 20 | Small | ancom | 17 |
| # | interaction | 0 | 0 | 4 | 0 | 0 | # | 20 | Small | ancom | 17 |
| # | disease | 7 | 1 | 0 | 0.875 | 1 | # | 20 | Small | aldex2 | 17 |
| # | sex | 4 | 0 | 0 | 1 | 1 | # | 20 | Small | aldex2 | 17 |
| # | time | 5 | 2 | 0 | 0.714285714 | 1 | # | 20 | Small | aldex2 | 17 |
| # | interaction | 0 | 0 | 4 | 0 | 0 | # | 20 | Small | aldex2 | 17 |
| # | disease | 7 | 11 | 0 | 0.388888889 | 1 | # | 20 | Baseline | paralle | 17 |
| # | sex | 4 | 7 | 0 | 0.363636364 | 1 | # | 20 | Baseline | paralle | 17 |
| # | time | 5 | 8 | 0 | 0.384615385 | 1 | # | 20 | Baseline | paralle | 17 |
| # | interaction | 4 | 7 | 0 | 0.363636364 | 1 | # | 20 | Baseline | paralle | 17 |
| # | disease | 7 | 3 | 0 | 0.7 | 1 | # | 20 | Baseline | ancom | 17 |
| # | sex | 4 | 0 | 0 | 1 | 1 | # | 20 | Baseline | ancom | 17 |
| # | time | 5 | 2 | 0 | 0.714285714 | 1 | # | 20 | Baseline | ancom | 17 |
| # | interaction | 0 | 0 | 4 | 0 | 0 | # | 20 | Baseline | ancom | 17 |
| # | disease | 7 | 31 | 0 | 0.184210526 | 1 | # | 20 | Baseline | aldex2 | 17 |
| # | sex | 4 | 11 | 0 | 0.266666667 | 1 | # | 20 | Baseline | aldex2 | 17 |
| # | time | 5 | 23 | 0 | 0.178571429 | 1 | # | 20 | Baseline | aldex2 | 17 |
| # | interaction | 0 | 0 | 4 | 0 | 0 | # | 20 | Baseline | aldex2 | 17 |
| # | disease | 7 | 0 | 0 | 1 | 1 | # | 20 | Small | paralle | 17 |
| # | sex | 4 | 0 | 0 | 1 | 1 | # | 20 | Small | paralle | 17 |
| # | time | 5 | 0 | 0 | 1 | 1 | # | 20 | Small | paralle | 17 |
| # | interaction | 3 | 0 | 1 | 1 | 0.75 | # | 20 | Small | paralle | 17 |
| # | disease | 7 | 3 | 0 | 0.7 | 1 | # | 20 | Small | ancom | 17 |
| # | sex | 4 | 0 | 0 | 1 | 1 | # | 20 | Small | ancom | 17 |
| # | time | 5 | 2 | 0 | 0.714285714 | 1 | # | 20 | Small | ancom | 17 |
| # | interaction | 0 | 0 | 4 | 0 | 0 | # | 20 | Small | ancom | 17 |
| # | disease | 7 | 2 | 0 | 0.777777778 | 1 | # | 20 | Small | aldex2 | 17 |
| # | sex | 4 | 1 | 0 | 0.8 | 1 | # | 20 | Small | aldex2 | 17 |
| # | time | 5 | 3 | 0 | 0.625 | 1 | # | 20 | Small | aldex2 | 17 |
| # | interaction | 0 | 0 | 4 | 0 | 0 | # | 20 | Small | aldex2 | 17 |
| # | disease | 7 | 14 | 0 | 0.333333333 | 1 | # | 20 | Baseline | paralle | 17 |
| # | sex | 4 | 4 | 0 | 0.5 | 1 | # | 20 | Baseline | paralle | 17 |
| # | time | 5 | 3 | 0 | 0.625 | 1 | # | 20 | Baseline | paralle | 17 |
| # | interaction | 4 | 3 | 0 | 0.571428571 | 1 | # | 20 | Baseline | paralle | 17 |
| # | disease | 7 | 4 | 0 | 0.636363636 | 1 | # | 20 | Baseline | ancom | 17 |
| # | sex | 4 | 0 | 0 | 1 | 1 | # | 20 | Baseline | ancom | 17 |
| # | time | 5 | 3 | 0 | 0.625 | 1 | # | 20 | Baseline | ancom | 17 |
| # | interaction | 0 | 0 | 4 | 0 | 0 | # | 20 | Baseline | ancom | 17 |
| # | disease | 7 | 44 | 0 | 0.137254902 | 1 | # | 20 | Baseline | aldex2 | 17 |
| # | sex | 4 | 11 | 0 | 0.266666667 | 1 | # | 20 | Baseline | aldex2 | 17 |
| # | time | 5 | 26 | 0 | 0.161290323 | 1 | # | 20 | Baseline | aldex2 | 17 |
| # | interaction | 0 | 0 | 4 | 0 | 0 | # | 20 | Baseline | aldex2 | 17 |
| # | disease | 7 | 0 | 0 | 1 | 1 | # | 20 | Small | paralle | 17 |
| # | sex | 4 | 0 | 0 | 1 | 1 | # | 20 | Small | paralle | 17 |
| # | time | 5 | 0 | 0 | 1 | 1 | # | 20 | Small | paralle | 17 |
| # | interaction | 4 | 0 | 0 | 1 | 1 | # | 20 | Small | paralle | 17 |
| # | disease | 7 | 4 | 0 | 0.636363636 | 1 | # | 20 | Small | ancom | 17 |
| # | sex | 4 | 0 | 0 | 1 | 1 | # | 20 | Small | ancom | 17 |
| # | time | 5 | 3 | 0 | 0.625 | 1 | # | 20 | Small | ancom | 17 |
| # | interaction | 0 | 0 | 4 | 0 | 0 | # | 20 | Small | ancom | 17 |
| # | disease | 7 | 3 | 0 | 0.7 | 1 | # | 20 | Small | aldex2 | 17 |
| # | sex | 4 | 0 | 0 | 1 | 1 | # | 20 | Small | aldex2 | 17 |
| # | time | 5 | 5 | 0 | 0.5 | 1 | # | 20 | Small | aldex2 | 17 |
| # | interaction | 0 | 0 | 4 | 0 | 0 | # | 20 | Small | aldex2 | 17 |
| # | disease | 7 | 12 | 0 | 0.368421053 | 1 | # | 20 | Baseline | paralle | 17 |
| # | sex | 4 | 9 | 0 | 0.307692308 | 1 | # | 20 | Baseline | paralle | 17 |
| # | time | 5 | 5 | 0 | 0.5 | 1 | # | 20 | Baseline | paralle | 17 |
| # | interaction | 4 | 2 | 0 | 0.666666667 | 1 | # | 20 | Baseline | paralle | 17 |
| # | disease | 7 | 4 | 0 | 0.636363636 | 1 | # | 20 | Baseline | ancom | 17 |
| # | sex | 4 | 0 | 0 | 1 | 1 | # | 20 | Baseline | ancom | 17 |
| # | time | 5 | 3 | 0 | 0.625 | 1 | # | 20 | Baseline | ancom | 17 |
| # | interaction | 0 | 0 | 4 | 0 | 0 | # | 20 | Baseline | ancom | 17 |
| # | disease | 7 | 54 | 0 | 0.114754098 | 1 | # | 20 | Baseline | aldex2 | 17 |
| # | sex | 4 | 18 | 0 | 0.181818182 | 1 | # | 20 | Baseline | aldex2 | 17 |
| # | time | 5 | 33 | 0 | 0.131578947 | 1 | # | 20 | Baseline | aldex2 | 17 |
| # | interaction | 0 | 0 | 4 | 0 | 0 | # | 20 | Baseline | aldex2 | 17 |
| # | disease | 7 | 0 | 0 | 1 | 1 | # | 20 | Small | paralle | 17 |
| # | sex | 4 | 0 | 0 | 1 | 1 | # | 20 | Small | paralle | 17 |
| # | time | 5 | 0 | 0 | 1 | 1 | # | 20 | Small | paralle | 17 |
| # | interaction | 4 | 0 | 0 | 1 | 1 | # | 20 | Small | paralle | 17 |
| # | disease | 7 | 4 | 0 | 0.636363636 | 1 | # | 20 | Small | ancom | 17 |
| # | sex | 4 | 0 | 0 | 1 | 1 | # | 20 | Small | ancom | 17 |
| # | time | 5 | 3 | 0 | 0.625 | 1 | # | 20 | Small | ancom | 17 |
| # | interaction | 0 | 0 | 4 | 0 | 0 | # | 20 | Small | ancom | 17 |
| # | disease | 7 | 7 | 0 | 0.5 | 1 | # | 20 | Small | aldex2 | 17 |
| # | sex | 4 | 3 | 0 | 0.571428571 | 1 | # | 20 | Small | aldex2 | 17 |
| # | time | 5 | 4 | 0 | 0.555555556 | 1 | # | 20 | Small | aldex2 | 17 |
| # | interaction | 0 | 0 | 4 | 0 | 0 | # | 20 | Small | aldex2 | 17 |
| # | disease | 4 | 12 | 0 | 0.25 | 1 | # | 10 | Baseline | paralle | 18 |
| # | sex | 2 | 5 | 0 | 0.285714286 | 1 | # | 10 | Baseline | paralle | 18 |
| # | time | 2 | 10 | 0 | 0.166666667 | 1 | # | 10 | Baseline | paralle | 18 |
| # | interaction | 2 | 7 | 0 | 0.222222222 | 1 | # | 10 | Baseline | paralle | 18 |
| # | disease | 4 | 0 | 0 | 1 | 1 | # | 10 | Baseline | ancom | 18 |
| # | sex | 2 | 0 | 0 | 1 | 1 | # | 10 | Baseline | ancom | 18 |
| # | time | 2 | 0 | 0 | 1 | 1 | # | 10 | Baseline | ancom | 18 |
| # | interaction | 0 | 0 | 2 | 0 | 0 | # | 10 | Baseline | ancom | 18 |
| # | disease | 4 | 10 | 0 | 0.285714286 | 1 | # | 10 | Baseline | aldex2 | 18 |
| # | sex | 2 | 6 | 0 | 0.25 | 1 | # | 10 | Baseline | aldex2 | 18 |
| # | time | 2 | 6 | 0 | 0.25 | 1 | # | 10 | Baseline | aldex2 | 18 |
| # | interaction | 0 | 0 | 2 | 0 | 0 | # | 10 | Baseline | aldex2 | 18 |
| # | disease | 4 | 0 | 0 | 1 | 1 | # | 10 | Small | paralle | 18 |
| # | sex | 2 | 0 | 0 | 1 | 1 | # | 10 | Small | paralle | 18 |
| # | time | 2 | 0 | 0 | 1 | 1 | # | 10 | Small | paralle | 18 |
| # | interaction | 0 | 0 | 2 | 0 | 0 | # | 10 | Small | paralle | 18 |
| # | disease | 4 | 0 | 0 | 1 | 1 | # | 10 | Small | ancom | 18 |
| # | sex | 2 | 0 | 0 | 1 | 1 | # | 10 | Small | ancom | 18 |
| # | time | 2 | 0 | 0 | 1 | 1 | # | 10 | Small | ancom | 18 |
| # | interaction | 0 | 0 | 2 | 0 | 0 | # | 10 | Small | ancom | 18 |
| # | disease | 4 | 1 | 0 | 0.8 | 1 | # | 10 | Small | aldex2 | 18 |
| # | sex | 2 | 0 | 0 | 1 | 1 | # | 10 | Small | aldex2 | 18 |
| # | time | 2 | 4 | 0 | 0.333333333 | 1 | # | 10 | Small | aldex2 | 18 |
| # | interaction | 0 | 0 | 2 | 0 | 0 | # | 10 | Small | aldex2 | 18 |
| # | disease | 4 | 9 | 0 | 0.307692308 | 1 | # | 10 | Baseline | paralle | 18 |
| # | sex | 2 | 0 | 0 | 1 | 1 | # | 10 | Baseline | paralle | 18 |
| # | time | 2 | 7 | 0 | 0.222222222 | 1 | # | 10 | Baseline | paralle | 18 |
| # | interaction | 2 | 6 | 0 | 0.25 | 1 | # | 10 | Baseline | paralle | 18 |
| # | disease | 4 | 1 | 0 | 0.8 | 1 | # | 10 | Baseline | ancom | 18 |
| # | sex | 2 | 0 | 0 | 1 | 1 | # | 10 | Baseline | ancom | 18 |
| # | time | 2 | 2 | 0 | 0.5 | 1 | # | 10 | Baseline | ancom | 18 |
| # | interaction | 0 | 0 | 2 | 0 | 0 | # | 10 | Baseline | ancom | 18 |
| # | disease | 4 | 13 | 0 | 0.235294118 | 1 | # | 10 | Baseline | aldex2 | 18 |
| # | sex | 2 | 6 | 0 | 0.25 | 1 | # | 10 | Baseline | aldex2 | 18 |
| # | time | 2 | 10 | 0 | 0.166666667 | 1 | # | 10 | Baseline | aldex2 | 18 |
| # | interaction | 0 | 0 | 2 | 0 | 0 | # | 10 | Baseline | aldex2 | 18 |
| # | disease | 4 | 0 | 0 | 1 | 1 | # | 10 | Small | paralle | 18 |
| # | sex | 2 | 0 | 0 | 1 | 1 | # | 10 | Small | paralle | 18 |
| # | time | 2 | 0 | 0 | 1 | 1 | # | 10 | Small | paralle | 18 |
| # | interaction | 1 | 0 | 1 | 1 | 0.5 | # | 10 | Small | paralle | 18 |
| # | disease | 4 | 1 | 0 | 0.8 | 1 | # | 10 | Small | ancom | 18 |
| # | sex | 2 | 0 | 0 | 1 | 1 | # | 10 | Small | ancom | 18 |
| # | time | 2 | 1 | 0 | 0.666666667 | 1 | # | 10 | Small | ancom | 18 |
| # | interaction | 0 | 0 | 2 | 0 | 0 | # | 10 | Small | ancom | 18 |
| # | disease | 4 | 1 | 0 | 0.8 | 1 | # | 10 | Small | aldex2 | 18 |
| # | sex | 2 | 1 | 0 | 0.666666667 | 1 | # | 10 | Small | aldex2 | 18 |
| # | time | 2 | 5 | 0 | 0.285714286 | 1 | # | 10 | Small | aldex2 | 18 |
| # | interaction | 0 | 0 | 2 | 0 | 0 | # | 10 | Small | aldex2 | 18 |
| # | disease | 4 | 5 | 0 | 0.444444444 | 1 | # | 10 | Baseline | paralle | 18 |
| # | sex | 2 | 3 | 0 | 0.4 | 1 | # | 10 | Baseline | paralle | 18 |
| # | time | 2 | 5 | 0 | 0.285714286 | 1 | # | 10 | Baseline | paralle | 18 |
| # | interaction | 2 | 4 | 0 | 0.333333333 | 1 | # | 10 | Baseline | paralle | 18 |
| # | disease | 4 | 1 | 0 | 0.8 | 1 | # | 10 | Baseline | ancom | 18 |
| # | sex | 2 | 0 | 0 | 1 | 1 | # | 10 | Baseline | ancom | 18 |
| # | time | 2 | 2 | 0 | 0.5 | 1 | # | 10 | Baseline | ancom | 18 |
| # | interaction | 0 | 0 | 2 | 0 | 0 | # | 10 | Baseline | ancom | 18 |
| # | disease | 4 | 20 | 0 | 0.166666667 | 1 | # | 10 | Baseline | aldex2 | 18 |
| # | sex | 2 | 10 | 0 | 0.166666667 | 1 | # | 10 | Baseline | aldex2 | 18 |
| # | time | 2 | 13 | 0 | 0.133333333 | 1 | # | 10 | Baseline | aldex2 | 18 |
| # | interaction | 0 | 0 | 2 | 0 | 0 | # | 10 | Baseline | aldex2 | 18 |
| # | disease | 4 | 2 | 0 | 0.666666667 | 1 | # | 10 | Small | paralle | 18 |
| # | sex | 2 | 0 | 0 | 1 | 1 | # | 10 | Small | paralle | 18 |
| # | time | 2 | 0 | 0 | 1 | 1 | # | 10 | Small | paralle | 18 |
| # | interaction | 1 | 0 | 1 | 1 | 0.5 | # | 10 | Small | paralle | 18 |
| # | disease | 4 | 1 | 0 | 0.8 | 1 | # | 10 | Small | ancom | 18 |
| # | sex | 2 | 0 | 0 | 1 | 1 | # | 10 | Small | ancom | 18 |
| # | time | 2 | 2 | 0 | 0.5 | 1 | # | 10 | Small | ancom | 18 |
| # | interaction | 0 | 0 | 2 | 0 | 0 | # | 10 | Small | ancom | 18 |
| # | disease | 4 | 2 | 0 | 0.666666667 | 1 | # | 10 | Small | aldex2 | 18 |
| # | sex | 2 | 0 | 0 | 1 | 1 | # | 10 | Small | aldex2 | 18 |
| # | time | 2 | 4 | 0 | 0.333333333 | 1 | # | 10 | Small | aldex2 | 18 |
| # | interaction | 0 | 0 | 2 | 0 | 0 | # | 10 | Small | aldex2 | 18 |
| # | disease | 4 | 8 | 0 | 0.333333333 | 1 | # | 10 | Baseline | paralle | 18 |
| # | sex | 2 | 2 | 0 | 0.5 | 1 | # | 10 | Baseline | paralle | 18 |
| # | time | 2 | 4 | 0 | 0.333333333 | 1 | # | 10 | Baseline | paralle | 18 |
| # | interaction | 2 | 5 | 0 | 0.285714286 | 1 | # | 10 | Baseline | paralle | 18 |
| # | disease | 4 | 1 | 0 | 0.8 | 1 | # | 10 | Baseline | ancom | 18 |
| # | sex | 2 | 0 | 0 | 1 | 1 | # | 10 | Baseline | ancom | 18 |
| # | time | 2 | 2 | 0 | 0.5 | 1 | # | 10 | Baseline | ancom | 18 |
| # | interaction | 0 | 0 | 2 | 0 | 0 | # | 10 | Baseline | ancom | 18 |
| # | disease | 4 | 29 | 0 | 0.121212121 | 1 | # | 10 | Baseline | aldex2 | 18 |
| # | sex | 2 | 9 | 0 | 0.181818182 | 1 | # | 10 | Baseline | aldex2 | 18 |
| # | time | 2 | 12 | 0 | 0.142857143 | 1 | # | 10 | Baseline | aldex2 | 18 |
| # | interaction | 0 | 0 | 2 | 0 | 0 | # | 10 | Baseline | aldex2 | 18 |
| # | disease | 4 | 0 | 0 | 1 | 1 | # | 10 | Small | paralle | 18 |
| # | sex | 2 | 0 | 0 | 1 | 1 | # | 10 | Small | paralle | 18 |
| # | time | 2 | 0 | 0 | 1 | 1 | # | 10 | Small | paralle | 18 |
| # | interaction | 1 | 0 | 1 | 1 | 0.5 | # | 10 | Small | paralle | 18 |
| # | disease | 4 | 1 | 0 | 0.8 | 1 | # | 10 | Small | ancom | 18 |
| # | sex | 2 | 0 | 0 | 1 | 1 | # | 10 | Small | ancom | 18 |
| # | time | 2 | 2 | 0 | 0.5 | 1 | # | 10 | Small | ancom | 18 |
| # | interaction | 0 | 0 | 2 | 0 | 0 | # | 10 | Small | ancom | 18 |
| # | disease | 4 | 2 | 0 | 0.666666667 | 1 | # | 10 | Small | aldex2 | 18 |
| # | sex | 2 | 0 | 0 | 1 | 1 | # | 10 | Small | aldex2 | 18 |
| # | time | 2 | 2 | 0 | 0.5 | 1 | # | 10 | Small | aldex2 | 18 |
| # | interaction | 0 | 0 | 2 | 0 | 0 | # | 10 | Small | aldex2 | 18 |
| # | disease | 7 | 8 | 0 | 0.466666667 | 1 | # | 20 | Baseline | paralle | 18 |
| # | sex | 4 | 11 | 0 | 0.266666667 | 1 | # | 20 | Baseline | paralle | 18 |
| # | time | 5 | 4 | 0 | 0.555555556 | 1 | # | 20 | Baseline | paralle | 18 |
| # | interaction | 4 | 10 | 0 | 0.285714286 | 1 | # | 20 | Baseline | paralle | 18 |
| # | disease | 7 | 1 | 0 | 0.875 | 1 | # | 20 | Baseline | ancom | 18 |
| # | sex | 4 | 0 | 0 | 1 | 1 | # | 20 | Baseline | ancom | 18 |
| # | time | 5 | 1 | 0 | 0.833333333 | 1 | # | 20 | Baseline | ancom | 18 |

| | | | | | | | | | | | |
|---|---|---|---|---|---|---|---|---|---|---|---|
| # | interaction | 0 | 0 | 4 | 0 | 0 | # | 20 | Baseline | ancom | 18 |
| # | disease | 7 | 15 | 0 | 0.318181818 | 1 | # | 20 | Baseline | aldex2 | 18 |
| # | sex | 4 | 6 | 0 | 0.4 | 1 | # | 20 | Baseline | aldex2 | 18 |
| # | time | 5 | 12 | 0 | 0.294117647 | 1 | # | 20 | Baseline | aldex2 | 18 |
| # | interaction | 0 | 0 | 4 | 0 | 0 | # | 20 | Baseline | aldex2 | 18 |
| # | disease | 7 | 1 | 0 | 0.875 | 1 | # | 20 | Small | paralle | 18 |
| # | sex | 4 | 0 | 0 | 1 | 1 | # | 20 | Small | paralle | 18 |
| # | time | 4 | 0 | 1 | 1 | 0.8 | # | 20 | Small | paralle | 18 |
| # | interaction | 2 | 1 | 2 | 0.666666667 | 0.5 | # | 20 | Small | paralle | 18 |
| # | disease | 7 | 0 | 0 | 1 | 1 | # | 20 | Small | ancom | 18 |
| # | sex | 4 | 0 | 0 | 1 | 1 | # | 20 | Small | ancom | 18 |
| # | time | 5 | 0 | 0 | 1 | 1 | # | 20 | Small | ancom | 18 |
| # | interaction | 0 | 0 | 4 | 0 | 0 | # | 20 | Small | ancom | 18 |
| # | disease | 7 | 1 | 0 | 0.875 | 1 | # | 20 | Small | aldex2 | 18 |
| # | sex | 4 | 0 | 0 | 1 | 1 | # | 20 | Small | aldex2 | 18 |
| # | time | 5 | 0 | 0 | 1 | 1 | # | 20 | Small | aldex2 | 18 |
| # | interaction | 0 | 0 | 4 | 0 | 0 | # | 20 | Small | aldex2 | 18 |
| # | disease | 7 | 11 | 0 | 0.388888889 | 1 | # | 20 | Baseline | paralle | 18 |
| # | sex | 4 | 7 | 0 | 0.363636364 | 1 | # | 20 | Baseline | paralle | 18 |
| # | time | 5 | 5 | 0 | 0.5 | 1 | # | 20 | Baseline | paralle | 18 |
| # | interaction | 4 | 7 | 0 | 0.363636364 | 1 | # | 20 | Baseline | paralle | 18 |
| # | disease | 7 | 3 | 0 | 0.7 | 1 | # | 20 | Baseline | ancom | 18 |
| # | sex | 4 | 0 | 0 | 1 | 1 | # | 20 | Baseline | ancom | 18 |
| # | time | 5 | 2 | 0 | 0.714285714 | 1 | # | 20 | Baseline | ancom | 18 |
| # | interaction | 0 | 0 | 4 | 0 | 0 | # | 20 | Baseline | ancom | 18 |
| # | disease | 7 | 34 | 0 | 0.170731707 | 1 | # | 20 | Baseline | aldex2 | 18 |
| # | sex | 4 | 13 | 0 | 0.235294118 | 1 | # | 20 | Baseline | aldex2 | 18 |
| # | time | 5 | 24 | 0 | 0.172413793 | 1 | # | 20 | Baseline | aldex2 | 18 |
| # | interaction | 0 | 0 | 4 | 0 | 0 | # | 20 | Baseline | aldex2 | 18 |
| # | disease | 7 | 0 | 0 | 1 | 1 | # | 20 | Small | paralle | 18 |
| # | sex | 4 | 0 | 0 | 1 | 1 | # | 20 | Small | paralle | 18 |
| # | time | 5 | 0 | 0 | 1 | 1 | # | 20 | Small | paralle | 18 |
| # | interaction | 3 | 0 | 1 | 1 | 0.75 | # | 20 | Small | paralle | 18 |
| # | disease | 7 | 4 | 0 | 0.636363636 | 1 | # | 20 | Small | ancom | 18 |
| # | sex | 4 | 0 | 0 | 1 | 1 | # | 20 | Small | ancom | 18 |
| # | time | 5 | 2 | 0 | 0.714285714 | 1 | # | 20 | Small | ancom | 18 |
| # | interaction | 0 | 0 | 4 | 0 | 0 | # | 20 | Small | ancom | 18 |
| # | disease | 7 | 2 | 0 | 0.777777778 | 1 | # | 20 | Small | aldex2 | 18 |
| # | sex | 4 | 0 | 0 | 1 | 1 | # | 20 | Small | aldex2 | 18 |
| # | time | 5 | 2 | 0 | 0.714285714 | 1 | # | 20 | Small | aldex2 | 18 |
| # | interaction | 0 | 0 | 4 | 0 | 0 | # | 20 | Small | aldex2 | 18 |
| # | disease | 7 | 8 | 0 | 0.466666667 | 1 | # | 20 | Baseline | paralle | 18 |
| # | sex | 4 | 4 | 0 | 0.5 | 1 | # | 20 | Baseline | paralle | 18 |
| # | time | 5 | 6 | 0 | 0.454545455 | 1 | # | 20 | Baseline | paralle | 18 |
| # | interaction | 4 | 3 | 0 | 0.571428571 | 1 | # | 20 | Baseline | paralle | 18 |
| # | disease | 7 | 4 | 0 | 0.636363636 | 1 | # | 20 | Baseline | ancom | 18 |
| # | sex | 4 | 0 | 0 | 1 | 1 | # | 20 | Baseline | ancom | 18 |
| # | time | 5 | 4 | 0 | 0.555555556 | 1 | # | 20 | Baseline | ancom | 18 |
| # | interaction | 0 | 0 | 4 | 0 | 0 | # | 20 | Baseline | ancom | 18 |
| # | disease | 7 | 46 | 0 | 0.132075472 | 1 | # | 20 | Baseline | aldex2 | 18 |
| # | sex | 4 | 13 | 0 | 0.235294118 | 1 | # | 20 | Baseline | aldex2 | 18 |
| # | time | 5 | 31 | 0 | 0.138888889 | 1 | # | 20 | Baseline | aldex2 | 18 |
| # | interaction | 0 | 0 | 4 | 0 | 0 | # | 20 | Baseline | aldex2 | 18 |
| # | disease | 7 | 0 | 0 | 1 | 1 | # | 20 | Small | paralle | 18 |
| # | sex | 4 | 0 | 0 | 1 | 1 | # | 20 | Small | paralle | 18 |
| # | time | 5 | 0 | 0 | 1 | 1 | # | 20 | Small | paralle | 18 |
| # | interaction | 4 | 0 | 0 | 1 | 1 | # | 20 | Small | paralle | 18 |
| # | disease | 7 | 4 | 0 | 0.636363636 | 1 | # | 20 | Small | ancom | 18 |
| # | sex | 4 | 0 | 0 | 1 | 1 | # | 20 | Small | ancom | 18 |
| # | time | 5 | 3 | 0 | 0.625 | 1 | # | 20 | Small | ancom | 18 |
| # | interaction | 0 | 0 | 4 | 0 | 0 | # | 20 | Small | ancom | 18 |
| # | disease | 7 | 3 | 0 | 0.7 | 1 | # | 20 | Small | aldex2 | 18 |
| # | sex | 4 | 0 | 0 | 1 | 1 | # | 20 | Small | aldex2 | 18 |
| # | time | 5 | 3 | 0 | 0.625 | 1 | # | 20 | Small | aldex2 | 18 |
| # | interaction | 0 | 0 | 4 | 0 | 0 | # | 20 | Small | aldex2 | 18 |
| # | disease | 7 | 11 | 0 | 0.388888889 | 1 | # | 20 | Baseline | paralle | 18 |
| # | sex | 4 | 5 | 0 | 0.444444444 | 1 | # | 20 | Baseline | paralle | 18 |
| # | time | 5 | 7 | 0 | 0.416666667 | 1 | # | 20 | Baseline | paralle | 18 |
| # | interaction | 4 | 5 | 0 | 0.444444444 | 1 | # | 20 | Baseline | paralle | 18 |
| # | disease | 7 | 4 | 0 | 0.636363636 | 1 | # | 20 | Baseline | ancom | 18 |
| # | sex | 4 | 0 | 0 | 1 | 1 | # | 20 | Baseline | ancom | 18 |
| # | time | 5 | 4 | 0 | 0.555555556 | 1 | # | 20 | Baseline | ancom | 18 |
| # | interaction | 0 | 0 | 4 | 0 | 0 | # | 20 | Baseline | ancom | 18 |
| # | disease | 7 | 60 | 0 | 0.104477612 | 1 | # | 20 | Baseline | aldex2 | 18 |
| # | sex | 4 | 18 | 0 | 0.181818182 | 1 | # | 20 | Baseline | aldex2 | 18 |
| # | time | 5 | 38 | 0 | 0.11627907 | 1 | # | 20 | Baseline | aldex2 | 18 |
| # | interaction | 0 | 0 | 4 | 0 | 0 | # | 20 | Baseline | aldex2 | 18 |
| # | disease | 7 | 0 | 0 | 1 | 1 | # | 20 | Small | paralle | 18 |
| # | sex | 4 | 0 | 0 | 1 | 1 | # | 20 | Small | paralle | 18 |
| # | time | 5 | 0 | 0 | 1 | 1 | # | 20 | Small | paralle | 18 |
| # | interaction | 3 | 0 | 1 | 1 | 0.75 | # | 20 | Small | paralle | 18 |
| # | disease | 7 | 4 | 0 | 0.636363636 | 1 | # | 20 | Small | ancom | 18 |
| # | sex | 4 | 0 | 0 | 1 | 1 | # | 20 | Small | ancom | 18 |
| # | time | 5 | 3 | 0 | 0.625 | 1 | # | 20 | Small | ancom | 18 |
| # | interaction | 0 | 0 | 4 | 0 | 0 | # | 20 | Small | ancom | 18 |
| # | disease | 7 | 6 | 0 | 0.538461538 | 1 | # | 20 | Small | aldex2 | 18 |
| # | sex | 4 | 0 | 0 | 1 | 1 | # | 20 | Small | aldex2 | 18 |
| # | time | 5 | 3 | 0 | 0.625 | 1 | # | 20 | Small | aldex2 | 18 |
| # | interaction | 0 | 0 | 4 | 0 | 0 | # | 20 | Small | aldex2 | 18 |
| # | disease | 4 | 12 | 0 | 0.25 | 1 | # | 10 | Baseline | paralle | 19 |
| # | sex | 2 | 6 | 0 | 0.25 | 1 | # | 10 | Baseline | paralle | 19 |
| # | time | 2 | 8 | 0 | 0.2 | 1 | # | 10 | Baseline | paralle | 19 |
| # | interaction | 2 | 10 | 0 | 0.166666667 | 1 | # | 10 | Baseline | paralle | 19 |
| # | disease | 4 | 1 | 0 | 0.8 | 1 | # | 10 | Baseline | ancom | 19 |
| # | sex | 2 | 0 | 0 | 1 | 1 | # | 10 | Baseline | ancom | 19 |
| # | time | 2 | 0 | 0 | 1 | 1 | # | 10 | Baseline | ancom | 19 |
| # | interaction | 0 | 0 | 2 | 0 | 0 | # | 10 | Baseline | ancom | 19 |
| # | disease | 4 | 14 | 0 | 0.222222222 | 1 | # | 10 | Baseline | aldex2 | 19 |
| # | sex | 2 | 4 | 0 | 0.333333333 | 1 | # | 10 | Baseline | aldex2 | 19 |
| # | time | 2 | 6 | 0 | 0.25 | 1 | # | 10 | Baseline | aldex2 | 19 |
| # | interaction | 0 | 0 | 2 | 0 | 0 | # | 10 | Baseline | aldex2 | 19 |
| # | disease | 4 | 0 | 0 | 1 | 1 | # | 10 | Small | paralle | 19 |
| # | sex | 2 | 0 | 0 | 1 | 1 | # | 10 | Small | paralle | 19 |
| # | time | 2 | 0 | 0 | 1 | 1 | # | 10 | Small | paralle | 19 |
| # | interaction | 2 | 0 | 0 | 1 | 1 | # | 10 | Small | paralle | 19 |
| # | disease | 4 | 1 | 0 | 0.8 | 1 | # | 10 | Small | ancom | 19 |
| # | sex | 2 | 0 | 0 | 1 | 1 | # | 10 | Small | ancom | 19 |
| # | time | 2 | 0 | 0 | 1 | 1 | # | 10 | Small | ancom | 19 |
| # | interaction | 0 | 0 | 2 | 0 | 0 | # | 10 | Small | ancom | 19 |
| # | disease | 4 | 1 | 0 | 0.8 | 1 | # | 10 | Small | aldex2 | 19 |
| # | sex | 2 | 0 | 0 | 1 | 1 | # | 10 | Small | aldex2 | 19 |
| # | time | 2 | 0 | 0 | 1 | 1 | # | 10 | Small | aldex2 | 19 |
| # | interaction | 0 | 0 | 2 | 0 | 0 | # | 10 | Small | aldex2 | 19 |
| # | disease | 4 | 4 | 0 | 0.5 | 1 | # | 10 | Baseline | paralle | 19 |
| # | sex | 2 | 7 | 0 | 0.222222222 | 1 | # | 10 | Baseline | paralle | 19 |
| # | time | 2 | 6 | 0 | 0.25 | 1 | # | 10 | Baseline | paralle | 19 |
| # | interaction | 2 | 7 | 0 | 0.222222222 | 1 | # | 10 | Baseline | paralle | 19 |
| # | disease | 4 | 1 | 0 | 0.8 | 1 | # | 10 | Baseline | ancom | 19 |
| # | sex | 2 | 0 | 0 | 1 | 1 | # | 10 | Baseline | ancom | 19 |
| # | time | 2 | 1 | 0 | 0.666666667 | 1 | # | 10 | Baseline | ancom | 19 |
| # | interaction | 0 | 0 | 2 | 0 | 0 | # | 10 | Baseline | ancom | 19 |
| # | disease | 4 | 15 | 0 | 0.210526316 | 1 | # | 10 | Baseline | aldex2 | 19 |
| # | sex | 2 | 7 | 0 | 0.222222222 | 1 | # | 10 | Baseline | aldex2 | 19 |
| # | time | 2 | 7 | 0 | 0.222222222 | 1 | # | 10 | Baseline | aldex2 | 19 |
| # | interaction | 0 | 0 | 2 | 0 | 0 | # | 10 | Baseline | aldex2 | 19 |
| # | disease | 4 | 0 | 0 | 1 | 1 | # | 10 | Small | paralle | 19 |
| # | sex | 2 | 0 | 0 | 1 | 1 | # | 10 | Small | paralle | 19 |
| # | time | 2 | 0 | 0 | 1 | 1 | # | 10 | Small | paralle | 19 |
| # | interaction | 2 | 0 | 0 | 1 | 1 | # | 10 | Small | paralle | 19 |
| # | disease | 4 | 1 | 0 | 0.8 | 1 | # | 10 | Small | ancom | 19 |
| # | sex | 2 | 0 | 0 | 1 | 1 | # | 10 | Small | ancom | 19 |
| # | time | 2 | 1 | 0 | 0.666666667 | 1 | # | 10 | Small | ancom | 19 |
| # | interaction | 0 | 0 | 2 | 0 | 0 | # | 10 | Small | ancom | 19 |
| # | disease | 4 | 1 | 0 | 0.8 | 1 | # | 10 | Small | aldex2 | 19 |
| # | sex | 2 | 0 | 0 | 1 | 1 | # | 10 | Small | aldex2 | 19 |
| # | time | 2 | 1 | 0 | 0.666666667 | 1 | # | 10 | Small | aldex2 | 19 |
| # | interaction | 0 | 0 | 2 | 0 | 0 | # | 10 | Small | aldex2 | 19 |
| # | disease | 4 | 3 | 0 | 0.571428571 | 1 | # | 10 | Baseline | paralle | 19 |
| # | sex | 2 | 5 | 0 | 0.285714286 | 1 | # | 10 | Baseline | paralle | 19 |
| # | time | 2 | 4 | 0 | 0.333333333 | 1 | # | 10 | Baseline | paralle | 19 |
| # | interaction | 2 | 3 | 0 | 0.4 | 1 | # | 10 | Baseline | paralle | 19 |
| # | disease | 4 | 2 | 0 | 0.666666667 | 1 | # | 10 | Baseline | ancom | 19 |
| # | sex | 2 | 0 | 0 | 1 | 1 | # | 10 | Baseline | ancom | 19 |
| # | time | 2 | 2 | 0 | 0.5 | 1 | # | 10 | Baseline | ancom | 19 |
| # | interaction | 0 | 0 | 2 | 0 | 0 | # | 10 | Baseline | ancom | 19 |
| # | disease | 4 | 22 | 0 | 0.153846154 | 1 | # | 10 | Baseline | aldex2 | 19 |
| # | sex | 2 | 8 | 0 | 0.2 | 1 | # | 10 | Baseline | aldex2 | 19 |
| # | time | 2 | 11 | 0 | 0.153846154 | 1 | # | 10 | Baseline | aldex2 | 19 |
| # | interaction | 0 | 0 | 2 | 0 | 0 | # | 10 | Baseline | aldex2 | 19 |
| # | disease | 4 | 0 | 0 | 1 | 1 | # | 10 | Small | paralle | 19 |
| # | sex | 2 | 0 | 0 | 1 | 1 | # | 10 | Small | paralle | 19 |
| # | time | 2 | 0 | 0 | 1 | 1 | # | 10 | Small | paralle | 19 |
| # | interaction | 2 | 0 | 0 | 1 | 1 | # | 10 | Small | paralle | 19 |
| # | disease | 4 | 2 | 0 | 0.666666667 | 1 | # | 10 | Small | ancom | 19 |
| # | sex | 2 | 0 | 0 | 1 | 1 | # | 10 | Small | ancom | 19 |
| # | time | 2 | 2 | 0 | 0.5 | 1 | # | 10 | Small | ancom | 19 |
| # | interaction | 0 | 0 | 2 | 0 | 0 | # | 10 | Small | ancom | 19 |
| # | disease | 4 | 2 | 0 | 0.666666667 | 1 | # | 10 | Small | aldex2 | 19 |
| # | sex | 2 | 0 | 0 | 1 | 1 | # | 10 | Small | aldex2 | 19 |
| # | time | 2 | 2 | 0 | 0.5 | 1 | # | 10 | Small | aldex2 | 19 |
| # | interaction | 0 | 0 | 2 | 0 | 0 | # | 10 | Small | aldex2 | 19 |
| # | disease | 4 | 5 | 0 | 0.444444444 | 1 | # | 10 | Baseline | paralle | 19 |
| # | sex | 2 | 6 | 0 | 0.25 | 1 | # | 10 | Baseline | paralle | 19 |
| # | time | 2 | 2 | 0 | 0.5 | 1 | # | 10 | Baseline | paralle | 19 |
| # | interaction | 2 | 2 | 0 | 0.5 | 1 | # | 10 | Baseline | paralle | 19 |
| # | disease | 4 | 2 | 0 | 0.666666667 | 1 | # | 10 | Baseline | ancom | 19 |
| # | sex | 2 | 0 | 0 | 1 | 1 | # | 10 | Baseline | ancom | 19 |
| # | time | 2 | 2 | 0 | 0.5 | 1 | # | 10 | Baseline | ancom | 19 |
| # | interaction | 0 | 0 | 2 | 0 | 0 | # | 10 | Baseline | ancom | 19 |
| # | disease | 4 | 25 | 0 | 0.137931034 | 1 | # | 10 | Baseline | aldex2 | 19 |
| # | sex | 2 | 8 | 0 | 0.2 | 1 | # | 10 | Baseline | aldex2 | 19 |
| # | time | 2 | 13 | 0 | 0.133333333 | 1 | # | 10 | Baseline | aldex2 | 19 |
| # | interaction | 0 | 0 | 2 | 0 | 0 | # | 10 | Baseline | aldex2 | 19 |
| # | disease | 4 | 0 | 0 | 1 | 1 | # | 10 | Small | paralle | 19 |
| # | sex | 2 | 0 | 0 | 1 | 1 | # | 10 | Small | paralle | 19 |
| # | time | 2 | 0 | 0 | 1 | 1 | # | 10 | Small | paralle | 19 |
| # | interaction | 2 | 0 | 0 | 1 | 1 | # | 10 | Small | paralle | 19 |
| # | disease | 4 | 2 | 0 | 0.666666667 | 1 | # | 10 | Small | ancom | 19 |
| # | sex | 2 | 0 | 0 | 1 | 1 | # | 10 | Small | ancom | 19 |
| # | time | 2 | 2 | 0 | 0.5 | 1 | # | 10 | Small | ancom | 19 |
| # | interaction | 0 | 0 | 2 | 0 | 0 | # | 10 | Small | ancom | 19 |
| # | disease | 4 | 2 | 0 | 0.666666667 | 1 | # | 10 | Small | aldex2 | 19 |
| # | sex | 2 | 0 | 0 | 1 | 1 | # | 10 | Small | aldex2 | 19 |
| # | time | 2 | 3 | 0 | 0.4 | 1 | # | 10 | Small | aldex2 | 19 |
| # | interaction | 0 | 0 | 2 | 0 | 0 | # | 10 | Small | aldex2 | 19 |
| # | disease | 7 | 14 | 0 | 0.333333333 | 1 | # | 20 | Baseline | paralle | 19 |
| # | sex | 4 | 9 | 0 | 0.307692308 | 1 | # | 20 | Baseline | paralle | 19 |
| # | time | 5 | 8 | 0 | 0.384615385 | 1 | # | 20 | Baseline | paralle | 19 |
| # | interaction | 4 | 5 | 0 | 0.444444444 | 1 | # | 20 | Baseline | paralle | 19 |
| # | disease | 7 | 2 | 0 | 0.777777778 | 1 | # | 20 | Baseline | ancom | 19 |
| # | sex | 4 | 0 | 0 | 1 | 1 | # | 20 | Baseline | ancom | 19 |
| # | time | 5 | 2 | 0 | 0.714285714 | 1 | # | 20 | Baseline | ancom | 19 |
| # | interaction | 0 | 0 | 4 | 0 | 0 | # | 20 | Baseline | ancom | 19 |
| # | disease | 7 | 21 | 0 | 0.25 | 1 | # | 20 | Baseline | aldex2 | 19 |
| # | sex | 4 | 5 | 0 | 0.444444444 | 1 | # | 20 | Baseline | aldex2 | 19 |
| # | time | 5 | 12 | 0 | 0.294117647 | 1 | # | 20 | Baseline | aldex2 | 19 |
| # | interaction | 0 | 0 | 4 | 0 | 0 | # | 20 | Baseline | aldex2 | 19 |
| # | disease | 7 | 0 | 0 | 1 | 1 | # | 20 | Small | paralle | 19 |
| # | sex | 4 | 0 | 0 | 1 | 1 | # | 20 | Small | paralle | 19 |
| # | time | 5 | 0 | 0 | 1 | 1 | # | 20 | Small | paralle | 19 |
| # | interaction | 1 | 0 | 3 | 1 | 0.25 | # | 20 | Small | paralle | 19 |
| # | disease | 7 | 1 | 0 | 0.875 | 1 | # | 20 | Small | ancom | 19 |
| # | sex | 4 | 0 | 0 | 1 | 1 | # | 20 | Small | ancom | 19 |
| # | time | 5 | 0 | 0 | 1 | 1 | # | 20 | Small | ancom | 19 |
| # | interaction | 0 | 0 | 4 | 0 | 0 | # | 20 | Small | ancom | 19 |
| # | disease | 7 | 1 | 0 | 0.875 | 1 | # | 20 | Small | aldex2 | 19 |
| # | sex | 4 | 0 | 0 | 1 | 1 | # | 20 | Small | aldex2 | 19 |
| # | time | 5 | 1 | 0 | 0.833333333 | 1 | # | 20 | Small | aldex2 | 19 |
| # | interaction | 0 | 0 | 4 | 0 | 0 | # | 20 | Small | aldex2 | 19 |
| # | disease | 7 | 12 | 0 | 0.368421053 | 1 | # | 20 | Baseline | paralle | 19 |
| # | sex | 4 | 5 | 0 | 0.444444444 | 1 | # | 20 | Baseline | paralle | 19 |
| # | time | 5 | 7 | 0 | 0.416666667 | 1 | # | 20 | Baseline | paralle | 19 |
| # | interaction | 4 | 8 | 0 | 0.333333333 | 1 | # | 20 | Baseline | paralle | 19 |
| # | disease | 7 | 3 | 0 | 0.7 | 1 | # | 20 | Baseline | ancom | 19 |
| # | sex | 4 | 0 | 0 | 1 | 1 | # | 20 | Baseline | ancom | 19 |
| # | time | 5 | 3 | 0 | 0.625 | 1 | # | 20 | Baseline | ancom | 19 |
| # | interaction | 0 | 0 | 4 | 0 | 0 | # | 20 | Baseline | ancom | 19 |
| # | disease | 7 | 28 | 0 | 0.2 | 1 | # | 20 | Baseline | aldex2 | 19 |
| # | sex | 4 | 14 | 0 | 0.222222222 | 1 | # | 20 | Baseline | aldex2 | 19 |
| # | time | 5 | 24 | 0 | 0.172413793 | 1 | # | 20 | Baseline | aldex2 | 19 |
| # | interaction | 0 | 0 | 4 | 0 | 0 | # | 20 | Baseline | aldex2 | 19 |
| # | disease | 7 | 0 | 0 | 1 | 1 | # | 20 | Small | paralle | 19 |
| # | sex | 4 | 0 | 0 | 1 | 1 | # | 20 | Small | paralle | 19 |
| # | time | 5 | 0 | 0 | 1 | 1 | # | 20 | Small | paralle | 19 |
| # | interaction | 3 | 0 | 1 | 1 | 0.75 | # | 20 | Small | paralle | 19 |
| # | disease | 7 | 3 | 0 | 0.7 | 1 | # | 20 | Small | ancom | 19 |
| # | sex | 4 | 0 | 0 | 1 | 1 | # | 20 | Small | ancom | 19 |
| # | time | 5 | 2 | 0 | 0.714285714 | 1 | # | 20 | Small | ancom | 19 |
| # | interaction | 0 | 0 | 4 | 0 | 0 | # | 20 | Small | ancom | 19 |
| # | disease | 7 | 3 | 0 | 0.7 | 1 | # | 20 | Small | aldex2 | 19 |
| # | sex | 4 | 0 | 0 | 1 | 1 | # | 20 | Small | aldex2 | 19 |
| # | time | 5 | 3 | 0 | 0.625 | 1 | # | 20 | Small | aldex2 | 19 |
| # | interaction | 0 | 0 | 4 | 0 | 0 | # | 20 | Small | aldex2 | 19 |
| # | disease | 7 | 8 | 0 | 0.466666667 | 1 | # | 20 | Baseline | paralle | 19 |
| # | sex | 4 | 2 | 0 | 0.666666667 | 1 | # | 20 | Baseline | paralle | 19 |
| # | time | 5 | 6 | 0 | 0.454545455 | 1 | # | 20 | Baseline | paralle | 19 |
| # | interaction | 4 | 4 | 0 | 0.5 | 1 | # | 20 | Baseline | paralle | 19 |
| # | disease | 7 | 4 | 0 | 0.636363636 | 1 | # | 20 | Baseline | ancom | 19 |
| # | sex | 4 | 0 | 0 | 1 | 1 | # | 20 | Baseline | ancom | 19 |
| # | time | 5 | 4 | 0 | 0.555555556 | 1 | # | 20 | Baseline | ancom | 19 |
| # | interaction | 0 | 0 | 4 | 0 | 0 | # | 20 | Baseline | ancom | 19 |
| # | disease | 7 | 45 | 0 | 0.134615385 | 1 | # | 20 | Baseline | aldex2 | 19 |
| # | sex | 4 | 19 | 0 | 0.173913043 | 1 | # | 20 | Baseline | aldex2 | 19 |
| # | time | 5 | 30 | 0 | 0.142857143 | 1 | # | 20 | Baseline | aldex2 | 19 |
| # | interaction | 0 | 0 | 4 | 0 | 0 | # | 20 | Baseline | aldex2 | 19 |
| # | disease | 7 | 0 | 0 | 1 | 1 | # | 20 | Small | paralle | 19 |
| # | sex | 4 | 1 | 0 | 0.8 | 1 | # | 20 | Small | paralle | 19 |
| # | time | 5 | 0 | 0 | 1 | 1 | # | 20 | Small | paralle | 19 |
| # | interaction | 4 | 0 | 0 | 1 | 1 | # | 20 | Small | paralle | 19 |
| # | disease | 7 | 4 | 0 | 0.636363636 | 1 | # | 20 | Small | ancom | 19 |
| # | sex | 4 | 0 | 0 | 1 | 1 | # | 20 | Small | ancom | 19 |
| # | time | 5 | 4 | 0 | 0.555555556 | 1 | # | 20 | Small | ancom | 19 |
| # | interaction | 0 | 0 | 4 | 0 | 0 | # | 20 | Small | ancom | 19 |
| # | disease | 7 | 6 | 0 | 0.538461538 | 1 | # | 20 | Small | aldex2 | 19 |
| # | sex | 4 | 0 | 0 | 1 | 1 | # | 20 | Small | aldex2 | 19 |
| # | time | 5 | 3 | 0 | 0.625 | 1 | # | 20 | Small | aldex2 | 19 |
| # | interaction | 0 | 0 | 4 | 0 | 0 | # | 20 | Small | aldex2 | 19 |
| # | disease | 7 | 11 | 0 | 0.388888889 | 1 | # | 20 | Baseline | paralle | 19 |
| # | sex | 4 | 6 | 0 | 0.4 | 1 | # | 20 | Baseline | paralle | 19 |
| # | time | 5 | 8 | 0 | 0.384615385 | 1 | # | 20 | Baseline | paralle | 19 |
| # | interaction | 4 | 8 | 0 | 0.333333333 | 1 | # | 20 | Baseline | paralle | 19 |
| # | disease | 7 | 4 | 0 | 0.636363636 | 1 | # | 20 | Baseline | ancom | 19 |
| # | sex | 4 | 0 | 0 | 1 | 1 | # | 20 | Baseline | ancom | 19 |
| # | time | 5 | 4 | 0 | 0.555555556 | 1 | # | 20 | Baseline | ancom | 19 |
| # | interaction | 0 | 0 | 4 | 0 | 0 | # | 20 | Baseline | ancom | 19 |
| # | disease | 7 | 53 | 0 | 0.116666667 | 1 | # | 20 | Baseline | aldex2 | 19 |
| # | sex | 4 | 23 | 0 | 0.148148148 | 1 | # | 20 | Baseline | aldex2 | 19 |
| # | time | 5 | 43 | 0 | 0.104166667 | 1 | # | 20 | Baseline | aldex2 | 19 |
| # | interaction | 0 | 0 | 4 | 0 | 0 | # | 20 | Baseline | aldex2 | 19 |
| # | disease | 7 | 0 | 0 | 1 | 1 | # | 20 | Small | paralle | 19 |
| # | sex | 4 | 1 | 0 | 0.8 | 1 | # | 20 | Small | paralle | 19 |
| # | time | 5 | 0 | 0 | 1 | 1 | # | 20 | Small | paralle | 19 |
| # | interaction | 4 | 0 | 0 | 1 | 1 | # | 20 | Small | paralle | 19 |
| # | disease | 7 | 4 | 0 | 0.636363636 | 1 | # | 20 | Small | ancom | 19 |
| # | sex | 4 | 0 | 0 | 1 | 1 | # | 20 | Small | ancom | 19 |
| # | time | 5 | 4 | 0 | 0.555555556 | 1 | # | 20 | Small | ancom | 19 |
| # | interaction | 0 | 0 | 4 | 0 | 0 | # | 20 | Small | ancom | 19 |
| # | disease | 7 | 10 | 0 | 0.411764706 | 1 | # | 20 | Small | aldex2 | 19 |
| # | sex | 4 | 0 | 0 | 1 | 1 | # | 20 | Small | aldex2 | 19 |
| # | time | 5 | 4 | 0 | 0.555555556 | 1 | # | 20 | Small | aldex2 | 19 |
| # | interaction | 0 | 0 | 4 | 0 | 0 | # | 20 | Small | aldex2 | 19 |
| # | disease | 4 | 7 | 0 | 0.363636364 | 1 | # | 10 | Baseline | paralle | 20 |
| # | sex | 2 | 7 | 0 | 0.222222222 | 1 | # | 10 | Baseline | paralle | 20 |
| # | time | 2 | 5 | 0 | 0.285714286 | 1 | # | 10 | Baseline | paralle | 20 |
| # | interaction | 2 | 10 | 0 | 0.166666667 | 1 | # | 10 | Baseline | paralle | 20 |
| # | disease | 4 | 0 | 0 | 1 | 1 | # | 10 | Baseline | ancom | 20 |
| # | sex | 2 | 0 | 0 | 1 | 1 | # | 10 | Baseline | ancom | 20 |
| # | time | 2 | 0 | 0 | 1 | 1 | # | 10 | Baseline | ancom | 20 |
| # | interaction | 0 | 0 | 2 | 0 | 0 | # | 10 | Baseline | ancom | 20 |
| # | disease | 4 | 10 | 0 | 0.285714286 | 1 | # | 10 | Baseline | aldex2 | 20 |
| # | sex | 2 | 1 | 0 | 0.666666667 | 1 | # | 10 | Baseline | aldex2 | 20 |
| # | time | 2 | 5 | 0 | 0.285714286 | 1 | # | 10 | Baseline | aldex2 | 20 |
| # | interaction | 0 | 0 | 2 | 0 | 0 | # | 10 | Baseline | aldex2 | 20 |
| # | disease | 4 | 0 | 0 | 1 | 1 | # | 10 | Small | paralle | 20 |
| # | sex | 2 | 0 | 0 | 1 | 1 | # | 10 | Small | paralle | 20 |
| # | time | 2 | 0 | 0 | 1 | 1 | # | 10 | Small | paralle | 20 |
| # | interaction | 0 | 1 | 2 | 0 | 0 | # | 10 | Small | paralle | 20 |
| # | disease | 4 | 0 | 0 | 1 | 1 | # | 10 | Small | ancom | 20 |
| # | sex | 2 | 0 | 0 | 1 | 1 | # | 10 | Small | ancom | 20 |
| # | time | 2 | 0 | 0 | 1 | 1 | # | 10 | Small | ancom | 20 |
| # | interaction | 0 | 0 | 2 | 0 | 0 | # | 10 | Small | ancom | 20 |
| # | disease | 4 | 1 | 0 | 0.8 | 1 | # | 10 | Small | aldex2 | 20 |
| # | sex | 2 | 0 | 0 | 1 | 1 | # | 10 | Small | aldex2 | 20 |
| # | time | 2 | 1 | 0 | 0.666666667 | 1 | # | 10 | Small | aldex2 | 20 |
| # | interaction | 0 | 0 | 2 | 0 | 0 | # | 10 | Small | aldex2 | 20 |
| # | disease | 4 | 4 | 0 | 0.5 | 1 | # | 10 | Baseline | paralle | 20 |
| # | sex | 2 | 6 | 0 | 0.25 | 1 | # | 10 | Baseline | paralle | 20 |
| # | time | 2 | 5 | 0 | 0.285714286 | 1 | # | 10 | Baseline | paralle | 20 |
| # | interaction | 2 | 3 | 0 | 0.4 | 1 | # | 10 | Baseline | paralle | 20 |
| # | disease | 4 | 2 | 0 | 0.666666667 | 1 | # | 10 | Baseline | ancom | 20 |
| # | sex | 2 | 0 | 0 | 1 | 1 | # | 10 | Baseline | ancom | 20 |
| # | time | 2 | 2 | 0 | 0.5 | 1 | # | 10 | Baseline | ancom | 20 |
| # | interaction | 0 | 0 | 2 | 0 | 0 | # | 10 | Baseline | ancom | 20 |
| # | disease | 4 | 12 | 0 | 0.25 | 1 | # | 10 | Baseline | aldex2 | 20 |
| # | sex | 2 | 5 | 0 | 0.285714286 | 1 | # | 10 | Baseline | aldex2 | 20 |
| # | time | 2 | 9 | 0 | 0.181818182 | 1 | # | 10 | Baseline | aldex2 | 20 |
| # | interaction | 0 | 0 | 2 | 0 | 0 | # | 10 | Baseline | aldex2 | 20 |
| # | disease | 4 | 0 | 0 | 1 | 1 | # | 10 | Small | paralle | 20 |
| # | sex | 2 | 0 | 0 | 1 | 1 | # | 10 | Small | paralle | 20 |
| # | time | 2 | 0 | 0 | 1 | 1 | # | 10 | Small | paralle | 20 |
| # | interaction | 2 | 1 | 0 | 0.666666667 | 1 | # | 10 | Small | paralle | 20 |
| # | disease | 4 | 2 | 0 | 0.666666667 | 1 | # | 10 | Small | ancom | 20 |
| # | sex | 2 | 0 | 0 | 1 | 1 | # | 10 | Small | ancom | 20 |
| # | time | 2 | 2 | 0 | 0.5 | 1 | # | 10 | Small | ancom | 20 |
| # | interaction | 0 | 0 | 2 | 0 | 0 | # | 10 | Small | ancom | 20 |
| # | disease | 4 | 2 | 0 | 0.666666667 | 1 | # | 10 | Small | aldex2 | 20 |
| # | sex | 2 | 0 | 0 | 1 | 1 | # | 10 | Small | aldex2 | 20 |
| # | time | 2 | 2 | 0 | 0.5 | 1 | # | 10 | Small | aldex2 | 20 |
| # | interaction | 0 | 0 | 2 | 0 | 0 | # | 10 | Small | aldex2 | 20 |
| # | disease | 4 | 8 | 0 | 0.333333333 | 1 | # | 10 | Baseline | paralle | 20 |
| # | sex | 2 | 8 | 0 | 0.2 | 1 | # | 10 | Baseline | paralle | 20 |
| # | time | 2 | 1 | 0 | 0.666666667 | 1 | # | 10 | Baseline | paralle | 20 |
| # | interaction | 2 | 5 | 0 | 0.285714286 | 1 | # | 10 | Baseline | paralle | 20 |
| # | disease | 4 | 2 | 0 | 0.666666667 | 1 | # | 10 | Baseline | ancom | 20 |
| # | sex | 2 | 0 | 0 | 1 | 1 | # | 10 | Baseline | ancom | 20 |
| # | time | 2 | 2 | 0 | 0.5 | 1 | # | 10 | Baseline | ancom | 20 |
| # | interaction | 0 | 0 | 2 | 0 | 0 | # | 10 | Baseline | ancom | 20 |

| | | | | | | | | | | | |
|---|---|---|---|---|---|---|---|---|---|---|---|
| # | disease | 4 | 20 | 0 | 0.166666667 | 1 | # | 10 | Baseline | aldex2 | 20 |
| # | sex | 2 | 9 | 0 | 0.181818182 | 1 | # | 10 | Baseline | aldex2 | 20 |
| # | time | 2 | 10 | 0 | 0.166666667 | 1 | # | 10 | Baseline | aldex2 | 20 |
| # | interaction | 0 | 0 | 2 | 0 | 0 | # | 10 | Baseline | aldex2 | 20 |
| # | disease | 4 | 0 | 0 | 1 | 1 | # | 10 | Small | paralle | 20 |
| # | sex | 2 | 0 | 0 | 1 | 1 | # | 10 | Small | paralle | 20 |
| # | time | 2 | 0 | 0 | 1 | 1 | # | 10 | Small | paralle | 20 |
| # | interaction | 2 | 0 | 0 | 1 | 1 | # | 10 | Small | paralle | 20 |
| # | disease | 4 | 2 | 0 | 0.666666667 | 1 | # | 10 | Small | ancom | 20 |
| # | sex | 2 | 0 | 0 | 1 | 1 | # | 10 | Small | ancom | 20 |
| # | time | 2 | 2 | 0 | 0.5 | 1 | # | 10 | Small | ancom | 20 |
| # | interaction | 0 | 0 | 2 | 0 | 0 | # | 10 | Small | ancom | 20 |
| # | disease | 4 | 4 | 0 | 0.5 | 1 | # | 10 | Small | aldex2 | 20 |
| # | sex | 2 | 1 | 0 | 0.666666667 | 1 | # | 10 | Small | aldex2 | 20 |
| # | time | 2 | 3 | 0 | 0.4 | 1 | # | 10 | Small | aldex2 | 20 |
| # | interaction | 0 | 0 | 2 | 0 | 0 | # | 10 | Small | aldex2 | 20 |
| # | disease | 4 | 4 | 0 | 0.5 | 1 | # | 10 | Baseline | paralle | 20 |
| # | sex | 2 | 5 | 0 | 0.285714286 | 1 | # | 10 | Baseline | paralle | 20 |
| # | time | 2 | 3 | 0 | 0.4 | 1 | # | 10 | Baseline | paralle | 20 |
| # | interaction | 2 | 4 | 0 | 0.333333333 | 1 | # | 10 | Baseline | paralle | 20 |
| # | disease | 4 | 2 | 0 | 0.666666667 | 1 | # | 10 | Baseline | ancom | 20 |
| # | sex | 2 | 0 | 0 | 1 | 1 | # | 10 | Baseline | ancom | 20 |
| # | time | 2 | 2 | 0 | 0.5 | 1 | # | 10 | Baseline | ancom | 20 |
| # | interaction | 0 | 0 | 2 | 0 | 0 | # | 10 | Baseline | ancom | 20 |
| # | disease | 4 | 23 | 0 | 0.148148148 | 1 | # | 10 | Baseline | aldex2 | 20 |
| # | sex | 2 | 9 | 0 | 0.181818182 | 1 | # | 10 | Baseline | aldex2 | 20 |
| # | time | 2 | 12 | 0 | 0.142857143 | 1 | # | 10 | Baseline | aldex2 | 20 |
| # | interaction | 0 | 0 | 2 | 0 | 0 | # | 10 | Baseline | aldex2 | 20 |
| # | disease | 4 | 1 | 0 | 0.8 | 1 | # | 10 | Small | paralle | 20 |
| # | sex | 2 | 0 | 0 | 1 | 1 | # | 10 | Small | paralle | 20 |
| # | time | 2 | 0 | 0 | 1 | 1 | # | 10 | Small | paralle | 20 |
| # | interaction | 2 | 1 | 0 | 0.666666667 | 1 | # | 10 | Small | paralle | 20 |
| # | disease | 4 | 2 | 0 | 0.666666667 | 1 | # | 10 | Small | ancom | 20 |
| # | sex | 2 | 0 | 0 | 1 | 1 | # | 10 | Small | ancom | 20 |
| # | time | 2 | 2 | 0 | 0.5 | 1 | # | 10 | Small | ancom | 20 |
| # | interaction | 0 | 0 | 2 | 0 | 0 | # | 10 | Small | ancom | 20 |
| # | disease | 4 | 5 | 0 | 0.444444444 | 1 | # | 10 | Small | aldex2 | 20 |
| # | sex | 2 | 0 | 0 | 1 | 1 | # | 10 | Small | aldex2 | 20 |
| # | time | 2 | 3 | 0 | 0.4 | 1 | # | 10 | Small | aldex2 | 20 |
| # | interaction | 0 | 0 | 2 | 0 | 0 | # | 10 | Small | aldex2 | 20 |
| # | disease | 7 | 9 | 0 | 0.4375 | 1 | # | 20 | Baseline | paralle | 20 |
| # | sex | 4 | 8 | 0 | 0.333333333 | 1 | # | 20 | Baseline | paralle | 20 |
| # | time | 5 | 10 | 0 | 0.333333333 | 1 | # | 20 | Baseline | paralle | 20 |
| # | interaction | 4 | 6 | 0 | 0.4 | 1 | # | 20 | Baseline | paralle | 20 |
| # | disease | 7 | 0 | 0 | 1 | 1 | # | 20 | Baseline | ancom | 20 |
| # | sex | 4 | 0 | 0 | 1 | 1 | # | 20 | Baseline | ancom | 20 |
| # | time | 5 | 0 | 0 | 1 | 1 | # | 20 | Baseline | ancom | 20 |
| # | interaction | 0 | 0 | 4 | 0 | 0 | # | 20 | Baseline | ancom | 20 |
| # | disease | 7 | 23 | 0 | 0.233333333 | 1 | # | 20 | Baseline | aldex2 | 20 |
| # | sex | 4 | 6 | 0 | 0.4 | 1 | # | 20 | Baseline | aldex2 | 20 |
| # | time | 5 | 10 | 0 | 0.333333333 | 1 | # | 20 | Baseline | aldex2 | 20 |
| # | interaction | 0 | 0 | 4 | 0 | 0 | # | 20 | Baseline | aldex2 | 20 |
| # | disease | 7 | 0 | 0 | 1 | 1 | # | 20 | Small | paralle | 20 |
| # | sex | 4 | 0 | 0 | 1 | 1 | # | 20 | Small | paralle | 20 |
| # | time | 5 | 0 | 0 | 1 | 1 | # | 20 | Small | paralle | 20 |
| # | interaction | 0 | 0 | 4 | 0 | 0 | # | 20 | Small | paralle | 20 |
| # | disease | 7 | 0 | 0 | 1 | 1 | # | 20 | Small | ancom | 20 |
| # | sex | 4 | 0 | 0 | 1 | 1 | # | 20 | Small | ancom | 20 |
| # | time | 5 | 0 | 0 | 1 | 1 | # | 20 | Small | ancom | 20 |
| # | interaction | 0 | 0 | 4 | 0 | 0 | # | 20 | Small | ancom | 20 |
| # | disease | 7 | 0 | 0 | 1 | 1 | # | 20 | Small | aldex2 | 20 |
| # | sex | 4 | 0 | 0 | 1 | 1 | # | 20 | Small | aldex2 | 20 |
| # | time | 5 | 0 | 0 | 1 | 1 | # | 20 | Small | aldex2 | 20 |
| # | interaction | 0 | 0 | 4 | 0 | 0 | # | 20 | Small | aldex2 | 20 |
| # | disease | 7 | 9 | 0 | 0.4375 | 1 | # | 20 | Baseline | paralle | 20 |
| # | sex | 4 | 5 | 0 | 0.444444444 | 1 | # | 20 | Baseline | paralle | 20 |
| # | time | 5 | 6 | 0 | 0.454545455 | 1 | # | 20 | Baseline | paralle | 20 |
| # | interaction | 4 | 9 | 0 | 0.307692308 | 1 | # | 20 | Baseline | paralle | 20 |
| # | disease | 7 | 2 | 0 | 0.777777778 | 1 | # | 20 | Baseline | ancom | 20 |
| # | sex | 4 | 0 | 0 | 1 | 1 | # | 20 | Baseline | ancom | 20 |
| # | time | 5 | 3 | 0 | 0.625 | 1 | # | 20 | Baseline | ancom | 20 |
| # | interaction | 0 | 0 | 4 | 0 | 0 | # | 20 | Baseline | ancom | 20 |
| # | disease | 7 | 37 | 0 | 0.159090909 | 1 | # | 20 | Baseline | aldex2 | 20 |
| # | sex | 4 | 11 | 0 | 0.266666667 | 1 | # | 20 | Baseline | aldex2 | 20 |
| # | time | 5 | 21 | 0 | 0.192307692 | 1 | # | 20 | Baseline | aldex2 | 20 |
| # | interaction | 0 | 0 | 4 | 0 | 0 | # | 20 | Baseline | aldex2 | 20 |
| # | disease | 7 | 0 | 0 | 1 | 1 | # | 20 | Small | paralle | 20 |
| # | sex | 4 | 0 | 0 | 1 | 1 | # | 20 | Small | paralle | 20 |
| # | time | 5 | 0 | 0 | 1 | 1 | # | 20 | Small | paralle | 20 |
| # | interaction | 0 | 0 | 4 | 0 | 0 | # | 20 | Small | paralle | 20 |
| # | disease | 7 | 2 | 0 | 0.777777778 | 1 | # | 20 | Small | ancom | 20 |
| # | sex | 4 | 0 | 0 | 1 | 1 | # | 20 | Small | ancom | 20 |
| # | time | 5 | 3 | 0 | 0.625 | 1 | # | 20 | Small | ancom | 20 |
| # | interaction | 0 | 0 | 4 | 0 | 0 | # | 20 | Small | ancom | 20 |
| # | disease | 7 | 0 | 0 | 1 | 1 | # | 20 | Small | aldex2 | 20 |
| # | sex | 4 | 0 | 0 | 1 | 1 | # | 20 | Small | aldex2 | 20 |
| # | time | 5 | 3 | 0 | 0.625 | 1 | # | 20 | Small | aldex2 | 20 |
| # | interaction | 0 | 0 | 4 | 0 | 0 | # | 20 | Small | aldex2 | 20 |
| # | disease | 7 | 15 | 0 | 0.318181818 | 1 | # | 20 | Baseline | paralle | 20 |
| # | sex | 4 | 2 | 0 | 0.666666667 | 1 | # | 20 | Baseline | paralle | 20 |
| # | time | 5 | 8 | 0 | 0.384615385 | 1 | # | 20 | Baseline | paralle | 20 |
| # | interaction | 4 | 8 | 0 | 0.333333333 | 1 | # | 20 | Baseline | paralle | 20 |
| # | disease | 7 | 3 | 0 | 0.7 | 1 | # | 20 | Baseline | ancom | 20 |
| # | sex | 4 | 0 | 0 | 1 | 1 | # | 20 | Baseline | ancom | 20 |
| # | time | 5 | 4 | 0 | 0.555555556 | 1 | # | 20 | Baseline | ancom | 20 |
| # | interaction | 0 | 0 | 4 | 0 | 0 | # | 20 | Baseline | ancom | 20 |
| # | disease | 7 | 42 | 0 | 0.142857143 | 1 | # | 20 | Baseline | aldex2 | 20 |
| # | sex | 4 | 16 | 0 | 0.2 | 1 | # | 20 | Baseline | aldex2 | 20 |
| # | time | 5 | 31 | 0 | 0.138888889 | 1 | # | 20 | Baseline | aldex2 | 20 |
| # | interaction | 0 | 0 | 4 | 0 | 0 | # | 20 | Baseline | aldex2 | 20 |
| # | disease | 7 | 0 | 0 | 1 | 1 | # | 20 | Small | paralle | 20 |
| # | sex | 4 | 0 | 0 | 1 | 1 | # | 20 | Small | paralle | 20 |
| # | time | 5 | 0 | 0 | 1 | 1 | # | 20 | Small | paralle | 20 |
| # | interaction | 4 | 0 | 0 | 1 | 1 | # | 20 | Small | paralle | 20 |
| # | disease | 7 | 3 | 0 | 0.7 | 1 | # | 20 | Small | ancom | 20 |
| # | sex | 4 | 0 | 0 | 1 | 1 | # | 20 | Small | ancom | 20 |
| # | time | 5 | 4 | 0 | 0.555555556 | 1 | # | 20 | Small | ancom | 20 |
| # | interaction | 0 | 0 | 4 | 0 | 0 | # | 20 | Small | ancom | 20 |
| # | disease | 7 | 2 | 0 | 0.777777778 | 1 | # | 20 | Small | aldex2 | 20 |
| # | sex | 4 | 0 | 0 | 1 | 1 | # | 20 | Small | aldex2 | 20 |
| # | time | 5 | 5 | 0 | 0.5 | 1 | # | 20 | Small | aldex2 | 20 |
| # | interaction | 0 | 0 | 4 | 0 | 0 | # | 20 | Small | aldex2 | 20 |
| # | disease | 7 | 17 | 0 | 0.291666667 | 1 | # | 20 | Baseline | paralle | 20 |
| # | sex | 4 | 7 | 0 | 0.363636364 | 1 | # | 20 | Baseline | paralle | 20 |
| # | time | 5 | 5 | 0 | 0.5 | 1 | # | 20 | Baseline | paralle | 20 |
| # | interaction | 4 | 7 | 0 | 0.363636364 | 1 | # | 20 | Baseline | paralle | 20 |
| # | disease | 7 | 4 | 0 | 0.636363636 | 1 | # | 20 | Baseline | ancom | 20 |
| # | sex | 4 | 0 | 0 | 1 | 1 | # | 20 | Baseline | ancom | 20 |
| # | time | 5 | 4 | 0 | 0.555555556 | 1 | # | 20 | Baseline | ancom | 20 |
| # | interaction | 0 | 0 | 4 | 0 | 0 | # | 20 | Baseline | ancom | 20 |
| # | disease | 7 | 57 | 0 | 0.109375 | 1 | # | 20 | Baseline | aldex2 | 20 |
| # | sex | 4 | 22 | 0 | 0.153846154 | 1 | # | 20 | Baseline | aldex2 | 20 |
| # | time | 5 | 41 | 0 | 0.108695652 | 1 | # | 20 | Baseline | aldex2 | 20 |
| # | interaction | 0 | 0 | 4 | 0 | 0 | # | 20 | Baseline | aldex2 | 20 |
| # | disease | 7 | 0 | 0 | 1 | 1 | # | 20 | Small | paralle | 20 |
| # | sex | 4 | 0 | 0 | 1 | 1 | # | 20 | Small | paralle | 20 |
| # | time | 5 | 0 | 0 | 1 | 1 | # | 20 | Small | paralle | 20 |
| # | interaction | 4 | 0 | 0 | 1 | 1 | # | 20 | Small | paralle | 20 |
| # | disease | 7 | 3 | 0 | 0.7 | 1 | # | 20 | Small | ancom | 20 |
| # | sex | 4 | 0 | 0 | 1 | 1 | # | 20 | Small | ancom | 20 |
| # | time | 5 | 4 | 0 | 0.555555556 | 1 | # | 20 | Small | ancom | 20 |
| # | interaction | 0 | 0 | 4 | 0 | 0 | # | 20 | Small | ancom | 20 |
| # | disease | 7 | 7 | 0 | 0.5 | 1 | # | 20 | Small | aldex2 | 20 |
| # | sex | 4 | 1 | 0 | 0.8 | 1 | # | 20 | Small | aldex2 | 20 |
| # | time | 5 | 6 | 0 | 0.454545455 | 1 | # | 20 | Small | aldex2 | 20 |
| # | interaction | 0 | 0 | 4 | 0 | 0 | # | 20 | Small | aldex2 | 20 |
| # | disease | 4 | 15 | 0 | 0.210526316 | 1 | # | 10 | Baseline | paralle | 21 |
| # | sex | 2 | 13 | 0 | 0.133333333 | 1 | # | 10 | Baseline | paralle | 21 |
| # | time | 2 | 10 | 0 | 0.166666667 | 1 | # | 10 | Baseline | paralle | 21 |
| # | interaction | 2 | 14 | 0 | 0.125 | 1 | # | 10 | Baseline | paralle | 21 |
| # | disease | 4 | 0 | 0 | 1 | 1 | # | 10 | Baseline | ancom | 21 |
| # | sex | 2 | 0 | 0 | 1 | 1 | # | 10 | Baseline | ancom | 21 |
| # | time | 2 | 0 | 0 | 1 | 1 | # | 10 | Baseline | ancom | 21 |
| # | interaction | 0 | 0 | 2 | 0 | 0 | # | 10 | Baseline | ancom | 21 |
| # | disease | 4 | 11 | 0 | 0.266666667 | 1 | # | 10 | Baseline | aldex2 | 21 |
| # | sex | 2 | 5 | 0 | 0.285714286 | 1 | # | 10 | Baseline | aldex2 | 21 |
| # | time | 2 | 9 | 0 | 0.181818182 | 1 | # | 10 | Baseline | aldex2 | 21 |
| # | interaction | 0 | 0 | 2 | 0 | 0 | # | 10 | Baseline | aldex2 | 21 |
| # | disease | 4 | 0 | 0 | 1 | 1 | # | 10 | Small | paralle | 21 |
| # | sex | 2 | 0 | 0 | 1 | 1 | # | 10 | Small | paralle | 21 |
| # | time | 2 | 1 | 0 | 0.666666667 | 1 | # | 10 | Small | paralle | 21 |
| # | interaction | 1 | 0 | 1 | 1 | 0.5 | # | 10 | Small | paralle | 21 |
| # | disease | 4 | 0 | 0 | 1 | 1 | # | 10 | Small | ancom | 21 |
| # | sex | 2 | 0 | 0 | 1 | 1 | # | 10 | Small | ancom | 21 |
| # | time | 2 | 0 | 0 | 1 | 1 | # | 10 | Small | ancom | 21 |
| # | interaction | 0 | 0 | 2 | 0 | 0 | # | 10 | Small | ancom | 21 |
| # | disease | 4 | 1 | 0 | 0.8 | 1 | # | 10 | Small | aldex2 | 21 |
| # | sex | 2 | 0 | 0 | 1 | 1 | # | 10 | Small | aldex2 | 21 |
| # | time | 2 | 1 | 0 | 0.666666667 | 1 | # | 10 | Small | aldex2 | 21 |
| # | interaction | 0 | 0 | 2 | 0 | 0 | # | 10 | Small | aldex2 | 21 |
| # | disease | 4 | 10 | 0 | 0.285714286 | 1 | # | 10 | Baseline | paralle | 21 |
| # | sex | 2 | 6 | 0 | 0.25 | 1 | # | 10 | Baseline | paralle | 21 |
| # | time | 2 | 5 | 0 | 0.285714286 | 1 | # | 10 | Baseline | paralle | 21 |
| # | interaction | 2 | 11 | 0 | 0.153846154 | 1 | # | 10 | Baseline | paralle | 21 |
| # | disease | 4 | 1 | 0 | 0.8 | 1 | # | 10 | Baseline | ancom | 21 |
| # | sex | 2 | 0 | 0 | 1 | 1 | # | 10 | Baseline | ancom | 21 |
| # | time | 2 | 0 | 0 | 1 | 1 | # | 10 | Baseline | ancom | 21 |
| # | interaction | 0 | 0 | 2 | 0 | 0 | # | 10 | Baseline | ancom | 21 |
| # | disease | 4 | 15 | 0 | 0.210526316 | 1 | # | 10 | Baseline | aldex2 | 21 |
| # | sex | 2 | 10 | 0 | 0.166666667 | 1 | # | 10 | Baseline | aldex2 | 21 |
| # | time | 2 | 7 | 0 | 0.222222222 | 1 | # | 10 | Baseline | aldex2 | 21 |
| # | interaction | 0 | 0 | 2 | 0 | 0 | # | 10 | Baseline | aldex2 | 21 |
| # | disease | 4 | 0 | 0 | 1 | 1 | # | 10 | Small | paralle | 21 |
| # | sex | 2 | 0 | 0 | 1 | 1 | # | 10 | Small | paralle | 21 |
| # | time | 2 | 0 | 0 | 1 | 1 | # | 10 | Small | paralle | 21 |
| # | interaction | 2 | 2 | 0 | 0.5 | 1 | # | 10 | Small | paralle | 21 |
| # | disease | 4 | 1 | 0 | 0.8 | 1 | # | 10 | Small | ancom | 21 |
| # | sex | 2 | 0 | 0 | 1 | 1 | # | 10 | Small | ancom | 21 |
| # | time | 2 | 0 | 0 | 1 | 1 | # | 10 | Small | ancom | 21 |
| # | interaction | 0 | 0 | 2 | 0 | 0 | # | 10 | Small | ancom | 21 |
| # | disease | 4 | 2 | 0 | 0.666666667 | 1 | # | 10 | Small | aldex2 | 21 |
| # | sex | 2 | 0 | 0 | 1 | 1 | # | 10 | Small | aldex2 | 21 |
| # | time | 2 | 0 | 0 | 1 | 1 | # | 10 | Small | aldex2 | 21 |
| # | interaction | 0 | 0 | 2 | 0 | 0 | # | 10 | Small | aldex2 | 21 |
| # | disease | 4 | 6 | 0 | 0.4 | 1 | # | 10 | Baseline | paralle | 21 |
| # | sex | 2 | 4 | 0 | 0.333333333 | 1 | # | 10 | Baseline | paralle | 21 |
| # | time | 2 | 4 | 0 | 0.333333333 | 1 | # | 10 | Baseline | paralle | 21 |
| # | interaction | 2 | 8 | 0 | 0.2 | 1 | # | 10 | Baseline | paralle | 21 |
| # | disease | 4 | 2 | 0 | 0.666666667 | 1 | # | 10 | Baseline | ancom | 21 |
| # | sex | 2 | 0 | 0 | 1 | 1 | # | 10 | Baseline | ancom | 21 |
| # | time | 2 | 2 | 0 | 0.5 | 1 | # | 10 | Baseline | ancom | 21 |
| # | interaction | 0 | 0 | 2 | 0 | 0 | # | 10 | Baseline | ancom | 21 |
| # | disease | 4 | 15 | 0 | 0.210526316 | 1 | # | 10 | Baseline | aldex2 | 21 |
| # | sex | 2 | 4 | 0 | 0.333333333 | 1 | # | 10 | Baseline | aldex2 | 21 |
| # | time | 2 | 10 | 0 | 0.166666667 | 1 | # | 10 | Baseline | aldex2 | 21 |
| # | interaction | 0 | 0 | 2 | 0 | 0 | # | 10 | Baseline | aldex2 | 21 |
| # | disease | 4 | 0 | 0 | 1 | 1 | # | 10 | Small | paralle | 21 |
| # | sex | 2 | 0 | 0 | 1 | 1 | # | 10 | Small | paralle | 21 |
| # | time | 2 | 0 | 0 | 1 | 1 | # | 10 | Small | paralle | 21 |
| # | interaction | 2 | 1 | 0 | 0.666666667 | 1 | # | 10 | Small | paralle | 21 |
| # | disease | 4 | 2 | 0 | 0.666666667 | 1 | # | 10 | Small | ancom | 21 |
| # | sex | 2 | 0 | 0 | 1 | 1 | # | 10 | Small | ancom | 21 |
| # | time | 2 | 2 | 0 | 0.5 | 1 | # | 10 | Small | ancom | 21 |
| # | interaction | 0 | 0 | 2 | 0 | 0 | # | 10 | Small | ancom | 21 |
| # | disease | 4 | 3 | 0 | 0.571428571 | 1 | # | 10 | Small | aldex2 | 21 |
| # | sex | 2 | 0 | 0 | 1 | 1 | # | 10 | Small | aldex2 | 21 |
| # | time | 2 | 2 | 0 | 0.5 | 1 | # | 10 | Small | aldex2 | 21 |
| # | interaction | 0 | 0 | 2 | 0 | 0 | # | 10 | Small | aldex2 | 21 |
| # | disease | 4 | 10 | 0 | 0.285714286 | 1 | # | 10 | Baseline | paralle | 21 |
| # | sex | 2 | 0 | 0 | 1 | 1 | # | 10 | Baseline | paralle | 21 |
| # | time | 2 | 4 | 0 | 0.333333333 | 1 | # | 10 | Baseline | paralle | 21 |
| # | interaction | 2 | 8 | 0 | 0.2 | 1 | # | 10 | Baseline | paralle | 21 |
| # | disease | 4 | 2 | 0 | 0.666666667 | 1 | # | 10 | Baseline | ancom | 21 |
| # | sex | 2 | 0 | 0 | 1 | 1 | # | 10 | Baseline | ancom | 21 |
| # | time | 2 | 2 | 0 | 0.5 | 1 | # | 10 | Baseline | ancom | 21 |
| # | interaction | 0 | 0 | 2 | 0 | 0 | # | 10 | Baseline | ancom | 21 |
| # | disease | 4 | 19 | 0 | 0.173913043 | 1 | # | 10 | Baseline | aldex2 | 21 |
| # | sex | 2 | 2 | 0 | 0.5 | 1 | # | 10 | Baseline | aldex2 | 21 |
| # | time | 2 | 7 | 0 | 0.222222222 | 1 | # | 10 | Baseline | aldex2 | 21 |
| # | interaction | 0 | 0 | 2 | 0 | 0 | # | 10 | Baseline | aldex2 | 21 |
| # | disease | 4 | 0 | 0 | 1 | 1 | # | 10 | Small | paralle | 21 |
| # | sex | 2 | 0 | 0 | 1 | 1 | # | 10 | Small | paralle | 21 |
| # | time | 2 | 0 | 0 | 1 | 1 | # | 10 | Small | paralle | 21 |
| # | interaction | 2 | 0 | 0 | 1 | 1 | # | 10 | Small | paralle | 21 |
| # | disease | 4 | 2 | 0 | 0.666666667 | 1 | # | 10 | Small | ancom | 21 |
| # | sex | 2 | 0 | 0 | 1 | 1 | # | 10 | Small | ancom | 21 |
| # | time | 2 | 2 | 0 | 0.5 | 1 | # | 10 | Small | ancom | 21 |
| # | interaction | 0 | 0 | 2 | 0 | 0 | # | 10 | Small | ancom | 21 |
| # | disease | 4 | 4 | 0 | 0.5 | 1 | # | 10 | Small | aldex2 | 21 |
| # | sex | 2 | 0 | 0 | 1 | 1 | # | 10 | Small | aldex2 | 21 |
| # | time | 2 | 2 | 0 | 0.5 | 1 | # | 10 | Small | aldex2 | 21 |
| # | interaction | 0 | 0 | 2 | 0 | 0 | # | 10 | Small | aldex2 | 21 |
| # | disease | 7 | 11 | 0 | 0.388888889 | 1 | # | 20 | Baseline | paralle | 21 |
| # | sex | 4 | 9 | 0 | 0.307692308 | 1 | # | 20 | Baseline | paralle | 21 |
| # | time | 5 | 8 | 0 | 0.384615385 | 1 | # | 20 | Baseline | paralle | 21 |
| # | interaction | 4 | 9 | 0 | 0.307692308 | 1 | # | 20 | Baseline | paralle | 21 |
| # | disease | 7 | 3 | 0 | 0.7 | 1 | # | 20 | Baseline | ancom | 21 |
| # | sex | 4 | 0 | 0 | 1 | 1 | # | 20 | Baseline | ancom | 21 |
| # | time | 5 | 1 | 0 | 0.833333333 | 1 | # | 20 | Baseline | ancom | 21 |
| # | interaction | 0 | 0 | 4 | 0 | 0 | # | 20 | Baseline | ancom | 21 |
| # | disease | 7 | 19 | 0 | 0.269230769 | 1 | # | 20 | Baseline | aldex2 | 21 |
| # | sex | 4 | 9 | 0 | 0.307692308 | 1 | # | 20 | Baseline | aldex2 | 21 |
| # | time | 5 | 17 | 0 | 0.227272727 | 1 | # | 20 | Baseline | aldex2 | 21 |
| # | interaction | 0 | 0 | 4 | 0 | 0 | # | 20 | Baseline | aldex2 | 21 |
| # | disease | 7 | 0 | 0 | 1 | 1 | # | 20 | Small | paralle | 21 |
| # | sex | 4 | 0 | 0 | 1 | 1 | # | 20 | Small | paralle | 21 |
| # | time | 5 | 0 | 0 | 1 | 1 | # | 20 | Small | paralle | 21 |
| # | interaction | 3 | 0 | 1 | 1 | 0.75 | # | 20 | Small | paralle | 21 |
| # | disease | 7 | 2 | 0 | 0.777777778 | 1 | # | 20 | Small | ancom | 21 |
| # | sex | 4 | 1 | 0 | 0.8 | 1 | # | 20 | Small | ancom | 21 |
| # | time | 5 | 1 | 0 | 0.833333333 | 1 | # | 20 | Small | ancom | 21 |
| # | interaction | 0 | 0 | 4 | 0 | 0 | # | 20 | Small | ancom | 21 |
| # | disease | 7 | 1 | 0 | 0.875 | 1 | # | 20 | Small | aldex2 | 21 |
| # | sex | 4 | 0 | 0 | 1 | 1 | # | 20 | Small | aldex2 | 21 |
| # | time | 5 | 2 | 0 | 0.714285714 | 1 | # | 20 | Small | aldex2 | 21 |
| # | interaction | 0 | 0 | 4 | 0 | 0 | # | 20 | Small | aldex2 | 21 |
| # | disease | 7 | 4 | 0 | 0.636363636 | 1 | # | 20 | Baseline | paralle | 21 |
| # | sex | 4 | 5 | 0 | 0.444444444 | 1 | # | 20 | Baseline | paralle | 21 |
| # | time | 5 | 7 | 0 | 0.416666667 | 1 | # | 20 | Baseline | paralle | 21 |
| # | interaction | 4 | 8 | 0 | 0.333333333 | 1 | # | 20 | Baseline | paralle | 21 |
| # | disease | 7 | 4 | 0 | 0.636363636 | 1 | # | 20 | Baseline | ancom | 21 |
| # | sex | 4 | 0 | 0 | 1 | 1 | # | 20 | Baseline | ancom | 21 |
| # | time | 5 | 4 | 0 | 0.555555556 | 1 | # | 20 | Baseline | ancom | 21 |
| # | interaction | 0 | 0 | 4 | 0 | 0 | # | 20 | Baseline | ancom | 21 |
| # | disease | 7 | 35 | 0 | 0.166666667 | 1 | # | 20 | Baseline | aldex2 | 21 |
| # | sex | 4 | 16 | 0 | 0.2 | 1 | # | 20 | Baseline | aldex2 | 21 |
| # | time | 5 | 27 | 0 | 0.15625 | 1 | # | 20 | Baseline | aldex2 | 21 |
| # | interaction | 0 | 0 | 4 | 0 | 0 | # | 20 | Baseline | aldex2 | 21 |
| # | disease | 7 | 0 | 0 | 1 | 1 | # | 20 | Small | paralle | 21 |
| # | sex | 4 | 0 | 0 | 1 | 1 | # | 20 | Small | paralle | 21 |
| # | time | 5 | 0 | 0 | 1 | 1 | # | 20 | Small | paralle | 21 |
| # | interaction | 3 | 0 | 1 | 1 | 0.75 | # | 20 | Small | paralle | 21 |
| # | disease | 7 | 4 | 0 | 0.636363636 | 1 | # | 20 | Small | ancom | 21 |
| # | sex | 4 | 0 | 0 | 1 | 1 | # | 20 | Small | ancom | 21 |
| # | time | 5 | 4 | 0 | 0.555555556 | 1 | # | 20 | Small | ancom | 21 |
| # | interaction | 0 | 0 | 4 | 0 | 0 | # | 20 | Small | ancom | 21 |
| # | disease | 7 | 5 | 0 | 0.583333333 | 1 | # | 20 | Small | aldex2 | 21 |
| # | sex | 4 | 0 | 0 | 1 | 1 | # | 20 | Small | aldex2 | 21 |
| # | time | 5 | 4 | 0 | 0.555555556 | 1 | # | 20 | Small | aldex2 | 21 |
| # | interaction | 0 | 0 | 4 | 0 | 0 | # | 20 | Small | aldex2 | 21 |
| # | disease | 7 | 7 | 0 | 0.5 | 1 | # | 20 | Baseline | paralle | 21 |
| # | sex | 4 | 8 | 0 | 0.333333333 | 1 | # | 20 | Baseline | paralle | 21 |
| # | time | 5 | 2 | 0 | 0.714285714 | 1 | # | 20 | Baseline | paralle | 21 |
| # | interaction | 4 | 4 | 0 | 0.5 | 1 | # | 20 | Baseline | paralle | 21 |
| # | disease | 7 | 4 | 0 | 0.636363636 | 1 | # | 20 | Baseline | ancom | 21 |
| # | sex | 4 | 0 | 0 | 1 | 1 | # | 20 | Baseline | ancom | 21 |
| # | time | 5 | 4 | 0 | 0.555555556 | 1 | # | 20 | Baseline | ancom | 21 |
| # | interaction | 0 | 0 | 4 | 0 | 0 | # | 20 | Baseline | ancom | 21 |
| # | disease | 7 | 51 | 0 | 0.120689655 | 1 | # | 20 | Baseline | aldex2 | 21 |
| # | sex | 4 | 19 | 0 | 0.173913043 | 1 | # | 20 | Baseline | aldex2 | 21 |
| # | time | 5 | 60 | 0 | 0.111111111 | 1 | # | 20 | Baseline | aldex2 | 21 |
| # | interaction | 0 | 0 | 4 | 0 | 0 | # | 20 | Baseline | aldex2 | 21 |
| # | disease | 7 | 0 | 0 | 1 | 1 | # | 20 | Small | paralle | 21 |
| # | sex | 4 | 0 | 0 | 1 | 1 | # | 20 | Small | paralle | 21 |
| # | time | 5 | 0 | 0 | 1 | 1 | # | 20 | Small | paralle | 21 |
| # | interaction | 4 | 0 | 0 | 1 | 1 | # | 20 | Small | paralle | 21 |
| # | disease | 7 | 4 | 0 | 0.636363636 | 1 | # | 20 | Small | ancom | 21 |
| # | sex | 4 | 0 | 0 | 1 | 1 | # | 20 | Small | ancom | 21 |
| # | time | 5 | 4 | 0 | 0.555555556 | 1 | # | 20 | Small | ancom | 21 |
| # | interaction | 0 | 0 | 4 | 0 | 0 | # | 20 | Small | ancom | 21 |
| # | disease | 7 | 6 | 0 | 0.538461538 | 1 | # | 20 | Small | aldex2 | 21 |
| # | sex | 4 | 0 | 0 | 1 | 1 | # | 20 | Small | aldex2 | 21 |
| # | time | 5 | 4 | 0 | 0.555555556 | 1 | # | 20 | Small | aldex2 | 21 |
| # | interaction | 0 | 0 | 4 | 0 | 0 | # | 20 | Small | aldex2 | 21 |
| # | disease | 7 | 7 | 0 | 0.5 | 1 | # | 20 | Baseline | paralle | 21 |
| # | sex | 4 | 7 | 0 | 0.363636364 | 1 | # | 20 | Baseline | paralle | 21 |
| # | time | 5 | 5 | 0 | 0.5 | 1 | # | 20 | Baseline | paralle | 21 |
| # | interaction | 4 | 3 | 0 | 0.571428571 | 1 | # | 20 | Baseline | paralle | 21 |
| # | disease | 7 | 4 | 0 | 0.636363636 | 1 | # | 20 | Baseline | ancom | 21 |
| # | sex | 4 | 0 | 0 | 1 | 1 | # | 20 | Baseline | ancom | 21 |
| # | time | 5 | 4 | 0 | 0.555555556 | 1 | # | 20 | Baseline | ancom | 21 |
| # | interaction | 0 | 0 | 4 | 0 | 0 | # | 20 | Baseline | ancom | 21 |
| # | disease | 7 | 62 | 0 | 0.101449275 | 1 | # | 20 | Baseline | aldex2 | 21 |
| # | sex | 4 | 21 | 0 | 0.16 | 1 | # | 20 | Baseline | aldex2 | 21 |
| # | time | 5 | 60 | 0 | 0.111111111 | 1 | # | 20 | Baseline | aldex2 | 21 |
| # | interaction | 0 | 0 | 4 | 0 | 0 | # | 20 | Baseline | aldex2 | 21 |
| # | disease | 7 | 0 | 0 | 1 | 1 | # | 20 | Small | paralle | 21 |
| # | sex | 4 | 0 | 0 | 1 | 1 | # | 20 | Small | paralle | 21 |
| # | time | 5 | 0 | 0 | 1 | 1 | # | 20 | Small | paralle | 21 |
| # | interaction | 4 | 0 | 0 | 1 | 1 | # | 20 | Small | paralle | 21 |
| # | disease | 7 | 4 | 0 | 0.636363636 | 1 | # | 20 | Small | ancom | 21 |
| # | sex | 4 | 0 | 0 | 1 | 1 | # | 20 | Small | ancom | 21 |
| # | time | 5 | 4 | 0 | 0.555555556 | 1 | # | 20 | Small | ancom | 21 |
| # | interaction | 0 | 0 | 4 | 0 | 0 | # | 20 | Small | ancom | 21 |
| # | disease | 7 | 9 | 0 | 0.4375 | 1 | # | 20 | Small | aldex2 | 21 |
| # | sex | 4 | 0 | 0 | 1 | 1 | # | 20 | Small | aldex2 | 21 |
| # | time | 5 | 4 | 0 | 0.555555556 | 1 | # | 20 | Small | aldex2 | 21 |
| # | interaction | 0 | 0 | 4 | 0 | 0 | # | 20 | Small | aldex2 | 21 |
| # | disease | 4 | 5 | 0 | 0.444444444 | 1 | # | 10 | Baseline | paralle | 22 |
| # | sex | 2 | 10 | 0 | 0.166666667 | 1 | # | 10 | Baseline | paralle | 22 |
| # | time | 2 | 4 | 0 | 0.333333333 | 1 | # | 10 | Baseline | paralle | 22 |
| # | interaction | 2 | 8 | 0 | 0.2 | 1 | # | 10 | Baseline | paralle | 22 |
| # | disease | 4 | 0 | 0 | 1 | 1 | # | 10 | Baseline | ancom | 22 |
| # | sex | 2 | 0 | 0 | 1 | 1 | # | 10 | Baseline | ancom | 22 |
| # | time | 2 | 0 | 0 | 1 | 1 | # | 10 | Baseline | ancom | 22 |
| # | interaction | 0 | 0 | 2 | 0 | 0 | # | 10 | Baseline | ancom | 22 |
| # | disease | 4 | 7 | 0 | 0.363636364 | 1 | # | 10 | Baseline | aldex2 | 22 |

| | | | | | | | | | | | |
|---|---|---|---|---|---|---|---|---|---|---|---|
| # | sex | 2 | 3 | 0 | 0.4 | 1 | # | 10 | Baseline | aldex2 | 22 |
| # | time | 2 | 5 | 0 | 0.285714286 | 1 | # | 10 | Baseline | aldex2 | 22 |
| # | interaction | 0 | 0 | 2 | 0 | 0 | # | 10 | Baseline | aldex2 | 22 |
| # | disease | 4 | 0 | 0 | 1 | 1 | # | 10 | Small | paralle | 22 |
| # | sex | 2 | 0 | 0 | 1 | 1 | # | 10 | Small | paralle | 22 |
| # | time | 2 | 0 | 0 | 1 | 1 | # | 10 | Small | paralle | 22 |
| # | interaction | 2 | 0 | 0 | 1 | 1 | # | 10 | Small | paralle | 22 |
| # | disease | 4 | 0 | 0 | 1 | 1 | # | 10 | Small | ancom | 22 |
| # | sex | 2 | 0 | 0 | 1 | 1 | # | 10 | Small | ancom | 22 |
| # | time | 2 | 0 | 0 | 1 | 1 | # | 10 | Small | ancom | 22 |
| # | interaction | 0 | 0 | 2 | 0 | 0 | # | 10 | Small | ancom | 22 |
| # | disease | 4 | 0 | 0 | 1 | 1 | # | 10 | Small | aldex2 | 22 |
| # | sex | 2 | 0 | 0 | 1 | 1 | # | 10 | Small | aldex2 | 22 |
| # | time | 2 | 0 | 0 | 1 | 1 | # | 10 | Small | aldex2 | 22 |
| # | interaction | 0 | 0 | 2 | 0 | 0 | # | 10 | Small | aldex2 | 22 |
| # | disease | 4 | 8 | 0 | 0.333333333 | 1 | # | 10 | Baseline | paralle | 22 |
| # | sex | 2 | 6 | 0 | 0.25 | 1 | # | 10 | Baseline | paralle | 22 |
| # | time | 2 | 7 | 0 | 0.222222222 | 1 | # | 10 | Baseline | paralle | 22 |
| # | interaction | 2 | 5 | 0 | 0.285714286 | 1 | # | 10 | Baseline | paralle | 22 |
| # | disease | 4 | 2 | 0 | 0.666666667 | 1 | # | 10 | Baseline | ancom | 22 |
| # | sex | 2 | 0 | 0 | 1 | 1 | # | 10 | Baseline | ancom | 22 |
| # | time | 2 | 2 | 0 | 0.5 | 1 | # | 10 | Baseline | ancom | 22 |
| # | interaction | 0 | 0 | 2 | 0 | 0 | # | 10 | Baseline | ancom | 22 |
| # | disease | 4 | 13 | 0 | 0.235294118 | 1 | # | 10 | Baseline | aldex2 | 22 |
| # | sex | 2 | 4 | 0 | 0.333333333 | 1 | # | 10 | Baseline | aldex2 | 22 |
| # | time | 2 | 11 | 0 | 0.153846154 | 1 | # | 10 | Baseline | aldex2 | 22 |
| # | interaction | 0 | 0 | 2 | 0 | 0 | # | 10 | Baseline | aldex2 | 22 |
| # | disease | 4 | 0 | 0 | 1 | 1 | # | 10 | Small | paralle | 22 |
| # | sex | 2 | 1 | 0 | 0.666666667 | 1 | # | 10 | Small | paralle | 22 |
| # | time | 2 | 0 | 0 | 1 | 1 | # | 10 | Small | paralle | 22 |
| # | interaction | 2 | 0 | 0 | 1 | 1 | # | 10 | Small | paralle | 22 |
| # | disease | 4 | 2 | 0 | 0.666666667 | 1 | # | 10 | Small | ancom | 22 |
| # | sex | 2 | 0 | 0 | 1 | 1 | # | 10 | Small | ancom | 22 |
| # | time | 2 | 1 | 0 | 0.666666667 | 1 | # | 10 | Small | ancom | 22 |
| # | interaction | 0 | 0 | 2 | 0 | 0 | # | 10 | Small | ancom | 22 |
| # | disease | 4 | 2 | 0 | 0.666666667 | 1 | # | 10 | Small | aldex2 | 22 |
| # | sex | 2 | 0 | 0 | 1 | 1 | # | 10 | Small | aldex2 | 22 |
| # | time | 2 | 2 | 0 | 0.5 | 1 | # | 10 | Small | aldex2 | 22 |
| # | interaction | 0 | 0 | 2 | 0 | 0 | # | 10 | Small | aldex2 | 22 |
| # | disease | 4 | 4 | 0 | 0.5 | 1 | # | 10 | Baseline | paralle | 22 |
| # | sex | 2 | 2 | 0 | 0.5 | 1 | # | 10 | Baseline | paralle | 22 |
| # | time | 2 | 2 | 0 | 0.5 | 1 | # | 10 | Baseline | paralle | 22 |
| # | interaction | 2 | 4 | 0 | 0.333333333 | 1 | # | 10 | Baseline | paralle | 22 |
| # | disease | 4 | 2 | 0 | 0.666666667 | 1 | # | 10 | Baseline | ancom | 22 |
| # | sex | 2 | 0 | 0 | 1 | 1 | # | 10 | Baseline | ancom | 22 |
| # | time | 2 | 2 | 0 | 0.5 | 1 | # | 10 | Baseline | ancom | 22 |
| # | interaction | 0 | 0 | 2 | 0 | 0 | # | 10 | Baseline | ancom | 22 |
| # | disease | 4 | 15 | 0 | 0.210526316 | 1 | # | 10 | Baseline | aldex2 | 22 |
| # | sex | 2 | 3 | 0 | 0.4 | 1 | # | 10 | Baseline | aldex2 | 22 |
| # | time | 2 | 11 | 0 | 0.153846154 | 1 | # | 10 | Baseline | aldex2 | 22 |
| # | interaction | 0 | 0 | 2 | 0 | 0 | # | 10 | Baseline | aldex2 | 22 |
| # | disease | 4 | 0 | 0 | 1 | 1 | # | 10 | Small | paralle | 22 |
| # | sex | 2 | 0 | 0 | 1 | 1 | # | 10 | Small | paralle | 22 |
| # | time | 2 | 0 | 0 | 1 | 1 | # | 10 | Small | paralle | 22 |
| # | interaction | 2 | 0 | 0 | 1 | 1 | # | 10 | Small | paralle | 22 |
| # | disease | 4 | 2 | 0 | 0.666666667 | 1 | # | 10 | Small | ancom | 22 |
| # | sex | 2 | 0 | 0 | 1 | 1 | # | 10 | Small | ancom | 22 |
| # | time | 2 | 2 | 0 | 0.5 | 1 | # | 10 | Small | ancom | 22 |
| # | interaction | 0 | 0 | 2 | 0 | 0 | # | 10 | Small | ancom | 22 |
| # | disease | 4 | 2 | 0 | 0.666666667 | 1 | # | 10 | Small | aldex2 | 22 |
| # | sex | 2 | 0 | 0 | 1 | 1 | # | 10 | Small | aldex2 | 22 |
| # | time | 2 | 2 | 0 | 0.5 | 1 | # | 10 | Small | aldex2 | 22 |
| # | interaction | 0 | 0 | 2 | 0 | 0 | # | 10 | Small | aldex2 | 22 |
| # | disease | 4 | 2 | 0 | 0.666666667 | 1 | # | 10 | Baseline | paralle | 22 |
| # | sex | 2 | 1 | 0 | 0.666666667 | 1 | # | 10 | Baseline | paralle | 22 |
| # | time | 2 | 0 | 0 | 1 | 1 | # | 10 | Baseline | paralle | 22 |
| # | interaction | 2 | 2 | 0 | 0.5 | 1 | # | 10 | Baseline | paralle | 22 |
| # | disease | 4 | 2 | 0 | 0.666666667 | 1 | # | 10 | Baseline | ancom | 22 |
| # | sex | 2 | 0 | 0 | 1 | 1 | # | 10 | Baseline | ancom | 22 |
| # | time | 2 | 2 | 0 | 0.5 | 1 | # | 10 | Baseline | ancom | 22 |
| # | interaction | 0 | 0 | 2 | 0 | 0 | # | 10 | Baseline | ancom | 22 |
| # | disease | 4 | 18 | 0 | 0.181818182 | 1 | # | 10 | Baseline | aldex2 | 22 |
| # | sex | 2 | 7 | 0 | 0.222222222 | 1 | # | 10 | Baseline | aldex2 | 22 |
| # | time | 2 | 15 | 0 | 0.117647059 | 1 | # | 10 | Baseline | aldex2 | 22 |
| # | interaction | 0 | 0 | 2 | 0 | 0 | # | 10 | Baseline | aldex2 | 22 |
| # | disease | 4 | 0 | 0 | 1 | 1 | # | 10 | Small | paralle | 22 |
| # | sex | 2 | 0 | 0 | 1 | 1 | # | 10 | Small | paralle | 22 |
| # | time | 2 | 0 | 0 | 1 | 1 | # | 10 | Small | paralle | 22 |
| # | interaction | 2 | 0 | 0 | 1 | 1 | # | 10 | Small | paralle | 22 |
| # | disease | 4 | 2 | 0 | 0.666666667 | 1 | # | 10 | Small | ancom | 22 |
| # | sex | 2 | 0 | 0 | 1 | 1 | # | 10 | Small | ancom | 22 |
| # | time | 2 | 2 | 0 | 0.5 | 1 | # | 10 | Small | ancom | 22 |
| # | interaction | 0 | 0 | 2 | 0 | 0 | # | 10 | Small | ancom | 22 |
| # | disease | 4 | 2 | 0 | 0.666666667 | 1 | # | 10 | Small | aldex2 | 22 |
| # | sex | 2 | 0 | 0 | 1 | 1 | # | 10 | Small | aldex2 | 22 |
| # | time | 2 | 2 | 0 | 0.5 | 1 | # | 10 | Small | aldex2 | 22 |
| # | interaction | 0 | 0 | 2 | 0 | 0 | # | 10 | Small | aldex2 | 22 |
| # | disease | 7 | 6 | 0 | 0.538461538 | 1 | # | 20 | Baseline | paralle | 22 |
| # | sex | 4 | 9 | 0 | 0.307692308 | 1 | # | 20 | Baseline | paralle | 22 |
| # | time | 5 | 10 | 0 | 0.333333333 | 1 | # | 20 | Baseline | paralle | 22 |
| # | interaction | 4 | 7 | 0 | 0.363636364 | 1 | # | 20 | Baseline | paralle | 22 |
| # | disease | 7 | 1 | 0 | 0.875 | 1 | # | 20 | Baseline | ancom | 22 |
| # | sex | 4 | 0 | 0 | 1 | 1 | # | 20 | Baseline | ancom | 22 |
| # | time | 5 | 3 | 0 | 0.625 | 1 | # | 20 | Baseline | ancom | 22 |
| # | interaction | 0 | 0 | 4 | 0 | 0 | # | 20 | Baseline | ancom | 22 |
| # | disease | 7 | 18 | 0 | 0.28 | 1 | # | 20 | Baseline | aldex2 | 22 |
| # | sex | 4 | 4 | 0 | 0.5 | 1 | # | 20 | Baseline | aldex2 | 22 |
| # | time | 5 | 14 | 0 | 0.263157895 | 1 | # | 20 | Baseline | aldex2 | 22 |
| # | interaction | 0 | 0 | 4 | 0 | 0 | # | 20 | Baseline | aldex2 | 22 |
| # | disease | 7 | 0 | 0 | 1 | 1 | # | 20 | Small | paralle | 22 |
| # | sex | 4 | 0 | 0 | 1 | 1 | # | 20 | Small | paralle | 22 |
| # | time | 5 | 1 | 0 | 0.833333333 | 1 | # | 20 | Small | paralle | 22 |
| # | interaction | 1 | 0 | 3 | 1 | 0.25 | # | 20 | Small | paralle | 22 |
| # | disease | 7 | 1 | 0 | 0.875 | 1 | # | 20 | Small | ancom | 22 |
| # | sex | 4 | 0 | 0 | 1 | 1 | # | 20 | Small | ancom | 22 |
| # | time | 5 | 2 | 0 | 0.714285714 | 1 | # | 20 | Small | ancom | 22 |
| # | interaction | 0 | 0 | 4 | 0 | 0 | # | 20 | Small | ancom | 22 |
| # | disease | 7 | 1 | 0 | 0.875 | 1 | # | 20 | Small | aldex2 | 22 |
| # | sex | 4 | 0 | 0 | 1 | 1 | # | 20 | Small | aldex2 | 22 |
| # | time | 5 | 3 | 0 | 0.625 | 1 | # | 20 | Small | aldex2 | 22 |
| # | interaction | 0 | 0 | 4 | 0 | 0 | # | 20 | Small | aldex2 | 22 |
| # | disease | 7 | 5 | 0 | 0.583333333 | 1 | # | 20 | Baseline | paralle | 22 |
| # | sex | 4 | 11 | 0 | 0.266666667 | 1 | # | 20 | Baseline | paralle | 22 |
| # | time | 5 | 8 | 0 | 0.384615385 | 1 | # | 20 | Baseline | paralle | 22 |
| # | interaction | 4 | 8 | 0 | 0.333333333 | 1 | # | 20 | Baseline | paralle | 22 |
| # | disease | 7 | 4 | 0 | 0.636363636 | 1 | # | 20 | Baseline | ancom | 22 |
| # | sex | 4 | 0 | 0 | 1 | 1 | # | 20 | Baseline | ancom | 22 |
| # | time | 5 | 4 | 0 | 0.555555556 | 1 | # | 20 | Baseline | ancom | 22 |
| # | interaction | 0 | 0 | 4 | 0 | 0 | # | 20 | Baseline | ancom | 22 |
| # | disease | 7 | 33 | 0 | 0.175 | 1 | # | 20 | Baseline | aldex2 | 22 |
| # | sex | 4 | 8 | 0 | 0.333333333 | 1 | # | 20 | Baseline | aldex2 | 22 |
| # | time | 5 | 22 | 0 | 0.185185185 | 1 | # | 20 | Baseline | aldex2 | 22 |
| # | interaction | 0 | 0 | 4 | 0 | 0 | # | 20 | Baseline | aldex2 | 22 |
| # | disease | 7 | 1 | 0 | 0.875 | 1 | # | 20 | Small | paralle | 22 |
| # | sex | 4 | 0 | 0 | 1 | 1 | # | 20 | Small | paralle | 22 |
| # | time | 5 | 0 | 0 | 1 | 1 | # | 20 | Small | paralle | 22 |
| # | interaction | 3 | 1 | 1 | 0.75 | 0.75 | # | 20 | Small | paralle | 22 |
| # | disease | 7 | 4 | 0 | 0.636363636 | 1 | # | 20 | Small | ancom | 22 |
| # | sex | 4 | 0 | 0 | 1 | 1 | # | 20 | Small | ancom | 22 |
| # | time | 5 | 4 | 0 | 0.555555556 | 1 | # | 20 | Small | ancom | 22 |
| # | interaction | 0 | 0 | 4 | 0 | 0 | # | 20 | Small | ancom | 22 |
| # | disease | 7 | 3 | 0 | 0.7 | 1 | # | 20 | Small | aldex2 | 22 |
| # | sex | 4 | 0 | 0 | 1 | 1 | # | 20 | Small | aldex2 | 22 |
| # | time | 5 | 2 | 0 | 0.714285714 | 1 | # | 20 | Small | aldex2 | 22 |
| # | interaction | 0 | 0 | 4 | 0 | 0 | # | 20 | Small | aldex2 | 22 |
| # | disease | 7 | 10 | 0 | 0.411764706 | 1 | # | 20 | Baseline | paralle | 22 |
| # | sex | 4 | 9 | 0 | 0.307692308 | 1 | # | 20 | Baseline | paralle | 22 |
| # | time | 5 | 7 | 0 | 0.416666667 | 1 | # | 20 | Baseline | paralle | 22 |
| # | interaction | 4 | 7 | 0 | 0.363636364 | 1 | # | 20 | Baseline | paralle | 22 |
| # | disease | 7 | 4 | 0 | 0.636363636 | 1 | # | 20 | Baseline | ancom | 22 |
| # | sex | 4 | 0 | 0 | 1 | 1 | # | 20 | Baseline | ancom | 22 |
| # | time | 5 | 4 | 0 | 0.555555556 | 1 | # | 20 | Baseline | ancom | 22 |
| # | interaction | 0 | 0 | 4 | 0 | 0 | # | 20 | Baseline | ancom | 22 |
| # | disease | 7 | 49 | 0 | 0.125 | 1 | # | 20 | Baseline | aldex2 | 22 |
| # | sex | 4 | 16 | 0 | 0.2 | 1 | # | 20 | Baseline | aldex2 | 22 |
| # | time | 5 | 33 | 0 | 0.131578947 | 1 | # | 20 | Baseline | aldex2 | 22 |
| # | interaction | 0 | 0 | 4 | 0 | 0 | # | 20 | Baseline | aldex2 | 22 |
| # | disease | 7 | 1 | 0 | 0.875 | 1 | # | 20 | Small | paralle | 22 |
| # | sex | 4 | 1 | 0 | 0.8 | 1 | # | 20 | Small | paralle | 22 |
| # | time | 5 | 1 | 0 | 0.833333333 | 1 | # | 20 | Small | paralle | 22 |
| # | interaction | 4 | 0 | 0 | 1 | 1 | # | 20 | Small | paralle | 22 |
| # | disease | 7 | 4 | 0 | 0.636363636 | 1 | # | 20 | Small | ancom | 22 |
| # | sex | 4 | 0 | 0 | 1 | 1 | # | 20 | Small | ancom | 22 |
| # | time | 5 | 4 | 0 | 0.555555556 | 1 | # | 20 | Small | ancom | 22 |
| # | interaction | 0 | 0 | 4 | 0 | 0 | # | 20 | Small | ancom | 22 |
| # | disease | 7 | 4 | 0 | 0.636363636 | 1 | # | 20 | Small | aldex2 | 22 |
| # | sex | 4 | 0 | 0 | 1 | 1 | # | 20 | Small | aldex2 | 22 |
| # | time | 5 | 4 | 0 | 0.555555556 | 1 | # | 20 | Small | aldex2 | 22 |
| # | interaction | 0 | 0 | 4 | 0 | 0 | # | 20 | Small | aldex2 | 22 |
| # | disease | 7 | 13 | 0 | 0.35 | 1 | # | 20 | Baseline | paralle | 22 |
| # | sex | 4 | 8 | 0 | 0.333333333 | 1 | # | 20 | Baseline | paralle | 22 |
| # | time | 5 | 8 | 0 | 0.384615385 | 1 | # | 20 | Baseline | paralle | 22 |
| # | interaction | 4 | 5 | 0 | 0.444444444 | 1 | # | 20 | Baseline | paralle | 22 |
| # | disease | 7 | 4 | 0 | 0.636363636 | 1 | # | 20 | Baseline | ancom | 22 |
| # | sex | 4 | 0 | 0 | 1 | 1 | # | 20 | Baseline | ancom | 22 |
| # | time | 5 | 4 | 0 | 0.555555556 | 1 | # | 20 | Baseline | ancom | 22 |
| # | interaction | 0 | 0 | 4 | 0 | 0 | # | 20 | Baseline | ancom | 22 |
| # | disease | 7 | 64 | 0 | 0.098591549 | 1 | # | 20 | Baseline | aldex2 | 22 |
| # | sex | 4 | 17 | 0 | 0.19047619 | 1 | # | 20 | Baseline | aldex2 | 22 |
| # | time | 5 | 36 | 0 | 0.12195122 | 1 | # | 20 | Baseline | aldex2 | 22 |
| # | interaction | 0 | 0 | 4 | 0 | 0 | # | 20 | Baseline | aldex2 | 22 |
| # | disease | 7 | 0 | 0 | 1 | 1 | # | 20 | Small | paralle | 22 |
| # | sex | 4 | 1 | 0 | 0.8 | 1 | # | 20 | Small | paralle | 22 |
| # | time | 5 | 1 | 0 | 0.833333333 | 1 | # | 20 | Small | paralle | 22 |
| # | interaction | 4 | 0 | 0 | 1 | 1 | # | 20 | Small | paralle | 22 |
| # | disease | 7 | 4 | 0 | 0.636363636 | 1 | # | 20 | Small | ancom | 22 |
| # | sex | 4 | 0 | 0 | 1 | 1 | # | 20 | Small | ancom | 22 |
| # | time | 5 | 4 | 0 | 0.555555556 | 1 | # | 20 | Small | ancom | 22 |
| # | interaction | 0 | 0 | 4 | 0 | 0 | # | 20 | Small | ancom | 22 |
| # | disease | 7 | 5 | 0 | 0.583333333 | 1 | # | 20 | Small | aldex2 | 22 |
| # | sex | 4 | 0 | 0 | 1 | 1 | # | 20 | Small | aldex2 | 22 |
| # | time | 5 | 6 | 0 | 0.454545455 | 1 | # | 20 | Small | aldex2 | 22 |
| # | interaction | 0 | 0 | 4 | 0 | 0 | # | 20 | Small | aldex2 | 22 |
| # | disease | 4 | 9 | 0 | 0.307692308 | 1 | # | 10 | Baseline | paralle | 23 |
| # | sex | 2 | 7 | 0 | 0.222222222 | 1 | # | 10 | Baseline | paralle | 23 |
| # | time | 2 | 5 | 0 | 0.285714286 | 1 | # | 10 | Baseline | paralle | 23 |
| # | interaction | 2 | 11 | 0 | 0.153846154 | 1 | # | 10 | Baseline | paralle | 23 |
| # | disease | 4 | 0 | 0 | 1 | 1 | # | 10 | Baseline | ancom | 23 |
| # | sex | 2 | 0 | 0 | 1 | 1 | # | 10 | Baseline | ancom | 23 |
| # | time | 2 | 1 | 0 | 0.666666667 | 1 | # | 10 | Baseline | ancom | 23 |
| # | interaction | 0 | 0 | 2 | 0 | 0 | # | 10 | Baseline | ancom | 23 |
| # | disease | 4 | 11 | 0 | 0.266666667 | 1 | # | 10 | Baseline | aldex2 | 23 |
| # | sex | 2 | 7 | 0 | 0.222222222 | 1 | # | 10 | Baseline | aldex2 | 23 |
| # | time | 2 | 5 | 0 | 0.285714286 | 1 | # | 10 | Baseline | aldex2 | 23 |
| # | interaction | 0 | 0 | 2 | 0 | 0 | # | 10 | Baseline | aldex2 | 23 |
| # | disease | 4 | 0 | 0 | 1 | 1 | # | 10 | Small | paralle | 23 |
| # | sex | 2 | 0 | 0 | 1 | 1 | # | 10 | Small | paralle | 23 |
| # | time | 2 | 0 | 0 | 1 | 1 | # | 10 | Small | paralle | 23 |
| # | interaction | 1 | 0 | 1 | 1 | 0.5 | # | 10 | Small | paralle | 23 |
| # | disease | 4 | 0 | 0 | 1 | 1 | # | 10 | Small | ancom | 23 |
| # | sex | 2 | 0 | 0 | 1 | 1 | # | 10 | Small | ancom | 23 |
| # | time | 2 | 0 | 0 | 1 | 1 | # | 10 | Small | ancom | 23 |
| # | interaction | 0 | 0 | 2 | 0 | 0 | # | 10 | Small | ancom | 23 |
| # | disease | 4 | 1 | 0 | 0.8 | 1 | # | 10 | Small | aldex2 | 23 |
| # | sex | 2 | 0 | 0 | 1 | 1 | # | 10 | Small | aldex2 | 23 |
| # | time | 2 | 1 | 0 | 0.666666667 | 1 | # | 10 | Small | aldex2 | 23 |
| # | interaction | 0 | 0 | 2 | 0 | 0 | # | 10 | Small | aldex2 | 23 |
| # | disease | 4 | 8 | 0 | 0.333333333 | 1 | # | 10 | Baseline | paralle | 23 |
| # | sex | 2 | 4 | 0 | 0.333333333 | 1 | # | 10 | Baseline | paralle | 23 |
| # | time | 2 | 2 | 0 | 0.5 | 1 | # | 10 | Baseline | paralle | 23 |
| # | interaction | 2 | 4 | 0 | 0.333333333 | 1 | # | 10 | Baseline | paralle | 23 |
| # | disease | 4 | 2 | 0 | 0.666666667 | 1 | # | 10 | Baseline | ancom | 23 |
| # | sex | 2 | 0 | 0 | 1 | 1 | # | 10 | Baseline | ancom | 23 |
| # | time | 2 | 2 | 0 | 0.5 | 1 | # | 10 | Baseline | ancom | 23 |
| # | interaction | 0 | 0 | 2 | 0 | 0 | # | 10 | Baseline | ancom | 23 |
| # | disease | 4 | 19 | 0 | 0.173913043 | 1 | # | 10 | Baseline | aldex2 | 23 |
| # | sex | 2 | 5 | 0 | 0.285714286 | 1 | # | 10 | Baseline | aldex2 | 23 |
| # | time | 2 | 12 | 0 | 0.142857143 | 1 | # | 10 | Baseline | aldex2 | 23 |
| # | interaction | 0 | 0 | 2 | 0 | 0 | # | 10 | Baseline | aldex2 | 23 |
| # | disease | 4 | 0 | 0 | 1 | 1 | # | 10 | Small | paralle | 23 |
| # | sex | 2 | 0 | 0 | 1 | 1 | # | 10 | Small | paralle | 23 |
| # | time | 2 | 0 | 0 | 1 | 1 | # | 10 | Small | paralle | 23 |
| # | interaction | 2 | 0 | 0 | 1 | 1 | # | 10 | Small | paralle | 23 |
| # | disease | 4 | 2 | 0 | 0.666666667 | 1 | # | 10 | Small | ancom | 23 |
| # | sex | 2 | 0 | 0 | 1 | 1 | # | 10 | Small | ancom | 23 |
| # | time | 2 | 2 | 0 | 0.5 | 1 | # | 10 | Small | ancom | 23 |
| # | interaction | 0 | 0 | 2 | 0 | 0 | # | 10 | Small | ancom | 23 |
| # | disease | 4 | 2 | 0 | 0.666666667 | 1 | # | 10 | Small | aldex2 | 23 |
| # | sex | 2 | 0 | 0 | 1 | 1 | # | 10 | Small | aldex2 | 23 |
| # | time | 2 | 2 | 0 | 0.5 | 1 | # | 10 | Small | aldex2 | 23 |
| # | interaction | 0 | 0 | 2 | 0 | 0 | # | 10 | Small | aldex2 | 23 |
| # | disease | 4 | 6 | 0 | 0.4 | 1 | # | 10 | Baseline | paralle | 23 |
| # | sex | 2 | 8 | 0 | 0.2 | 1 | # | 10 | Baseline | paralle | 23 |
| # | time | 2 | 3 | 0 | 0.4 | 1 | # | 10 | Baseline | paralle | 23 |
| # | interaction | 2 | 7 | 0 | 0.222222222 | 1 | # | 10 | Baseline | paralle | 23 |
| # | disease | 4 | 2 | 0 | 0.666666667 | 1 | # | 10 | Baseline | ancom | 23 |
| # | sex | 2 | 0 | 0 | 1 | 1 | # | 10 | Baseline | ancom | 23 |
| # | time | 2 | 2 | 0 | 0.5 | 1 | # | 10 | Baseline | ancom | 23 |
| # | interaction | 0 | 0 | 2 | 0 | 0 | # | 10 | Baseline | ancom | 23 |
| # | disease | 4 | 20 | 0 | 0.166666667 | 1 | # | 10 | Baseline | aldex2 | 23 |
| # | sex | 2 | 10 | 0 | 0.166666667 | 1 | # | 10 | Baseline | aldex2 | 23 |
| # | time | 2 | 10 | 0 | 0.166666667 | 1 | # | 10 | Baseline | aldex2 | 23 |
| # | interaction | 0 | 0 | 2 | 0 | 0 | # | 10 | Baseline | aldex2 | 23 |
| # | disease | 4 | 0 | 0 | 1 | 1 | # | 10 | Small | paralle | 23 |
| # | sex | 2 | 0 | 0 | 1 | 1 | # | 10 | Small | paralle | 23 |
| # | time | 2 | 0 | 0 | 1 | 1 | # | 10 | Small | paralle | 23 |
| # | interaction | 2 | 0 | 0 | 1 | 1 | # | 10 | Small | paralle | 23 |
| # | disease | 4 | 2 | 0 | 0.666666667 | 1 | # | 10 | Small | ancom | 23 |
| # | sex | 2 | 0 | 0 | 1 | 1 | # | 10 | Small | ancom | 23 |
| # | time | 2 | 2 | 0 | 0.5 | 1 | # | 10 | Small | ancom | 23 |
| # | interaction | 0 | 0 | 2 | 0 | 0 | # | 10 | Small | ancom | 23 |
| # | disease | 4 | 4 | 0 | 0.5 | 1 | # | 10 | Small | aldex2 | 23 |
| # | sex | 2 | 0 | 0 | 1 | 1 | # | 10 | Small | aldex2 | 23 |
| # | time | 2 | 3 | 0 | 0.4 | 1 | # | 10 | Small | aldex2 | 23 |
| # | interaction | 0 | 0 | 2 | 0 | 0 | # | 10 | Small | aldex2 | 23 |
| # | disease | 4 | 5 | 0 | 0.444444444 | 1 | # | 10 | Baseline | paralle | 23 |
| # | sex | 2 | 3 | 0 | 0.4 | 1 | # | 10 | Baseline | paralle | 23 |
| # | time | 2 | 0 | 0 | 1 | 1 | # | 10 | Baseline | paralle | 23 |
| # | interaction | 2 | 3 | 0 | 0.4 | 1 | # | 10 | Baseline | paralle | 23 |
| # | disease | 4 | 2 | 0 | 0.666666667 | 1 | # | 10 | Baseline | ancom | 23 |
| # | sex | 2 | 0 | 0 | 1 | 1 | # | 10 | Baseline | ancom | 23 |
| # | time | 2 | 2 | 0 | 0.5 | 1 | # | 10 | Baseline | ancom | 23 |
| # | interaction | 0 | 0 | 2 | 0 | 0 | # | 10 | Baseline | ancom | 23 |
| # | disease | 4 | 22 | 0 | 0.153846154 | 1 | # | 10 | Baseline | aldex2 | 23 |
| # | sex | 2 | 13 | 0 | 0.133333333 | 1 | # | 10 | Baseline | aldex2 | 23 |
| # | time | 2 | 12 | 0 | 0.142857143 | 1 | # | 10 | Baseline | aldex2 | 23 |
| # | interaction | 0 | 0 | 2 | 0 | 0 | # | 10 | Baseline | aldex2 | 23 |
| # | disease | 4 | 0 | 0 | 1 | 1 | # | 10 | Small | paralle | 23 |
| # | sex | 2 | 0 | 0 | 1 | 1 | # | 10 | Small | paralle | 23 |
| # | time | 2 | 0 | 0 | 1 | 1 | # | 10 | Small | paralle | 23 |
| # | interaction | 2 | 0 | 0 | 1 | 1 | # | 10 | Small | paralle | 23 |
| # | disease | 4 | 2 | 0 | 0.666666667 | 1 | # | 10 | Small | ancom | 23 |
| # | sex | 2 | 0 | 0 | 1 | 1 | # | 10 | Small | ancom | 23 |
| # | time | 2 | 2 | 0 | 0.5 | 1 | # | 10 | Small | ancom | 23 |
| # | interaction | 0 | 0 | 2 | 0 | 0 | # | 10 | Small | ancom | 23 |
| # | disease | 4 | 5 | 0 | 0.444444444 | 1 | # | 10 | Small | aldex2 | 23 |
| # | sex | 2 | 0 | 0 | 1 | 1 | # | 10 | Small | aldex2 | 23 |
| # | time | 2 | 2 | 0 | 0.5 | 1 | # | 10 | Small | aldex2 | 23 |
| # | interaction | 0 | 0 | 2 | 0 | 0 | # | 10 | Small | aldex2 | 23 |
| # | disease | 7 | 9 | 0 | 0.4375 | 1 | # | 20 | Baseline | paralle | 23 |
| # | sex | 4 | 6 | 0 | 0.4 | 1 | # | 20 | Baseline | paralle | 23 |
| # | time | 5 | 4 | 0 | 0.555555556 | 1 | # | 20 | Baseline | paralle | 23 |
| # | interaction | 4 | 6 | 0 | 0.4 | 1 | # | 20 | Baseline | paralle | 23 |
| # | disease | 7 | 2 | 0 | 0.777777778 | 1 | # | 20 | Baseline | ancom | 23 |
| # | sex | 4 | 0 | 0 | 1 | 1 | # | 20 | Baseline | ancom | 23 |
| # | time | 5 | 0 | 0 | 1 | 1 | # | 20 | Baseline | ancom | 23 |
| # | interaction | 0 | 0 | 4 | 0 | 0 | # | 20 | Baseline | ancom | 23 |
| # | disease | 7 | 23 | 0 | 0.233333333 | 1 | # | 20 | Baseline | aldex2 | 23 |
| # | sex | 4 | 6 | 0 | 0.4 | 1 | # | 20 | Baseline | aldex2 | 23 |
| # | time | 5 | 11 | 0 | 0.3125 | 1 | # | 20 | Baseline | aldex2 | 23 |
| # | interaction | 0 | 0 | 4 | 0 | 0 | # | 20 | Baseline | aldex2 | 23 |
| # | disease | 7 | 1 | 0 | 0.875 | 1 | # | 20 | Small | paralle | 23 |
| # | sex | 4 | 0 | 0 | 1 | 1 | # | 20 | Small | paralle | 23 |
| # | time | 5 | 0 | 0 | 1 | 1 | # | 20 | Small | paralle | 23 |
| # | interaction | 0 | 0 | 4 | 0 | 0 | # | 20 | Small | paralle | 23 |
| # | disease | 7 | 3 | 0 | 0.7 | 1 | # | 20 | Small | ancom | 23 |
| # | sex | 4 | 0 | 0 | 1 | 1 | # | 20 | Small | ancom | 23 |
| # | time | 5 | 0 | 0 | 1 | 1 | # | 20 | Small | ancom | 23 |
| # | interaction | 0 | 0 | 4 | 0 | 0 | # | 20 | Small | ancom | 23 |
| # | disease | 7 | 1 | 0 | 0.875 | 1 | # | 20 | Small | aldex2 | 23 |
| # | sex | 4 | 0 | 0 | 1 | 1 | # | 20 | Small | aldex2 | 23 |
| # | time | 5 | 1 | 0 | 0.833333333 | 1 | # | 20 | Small | aldex2 | 23 |
| # | interaction | 0 | 0 | 4 | 0 | 0 | # | 20 | Small | aldex2 | 23 |
| # | disease | 7 | 9 | 0 | 0.4375 | 1 | # | 20 | Baseline | paralle | 23 |
| # | sex | 4 | 2 | 0 | 0.666666667 | 1 | # | 20 | Baseline | paralle | 23 |
| # | time | 5 | 8 | 0 | 0.384615385 | 1 | # | 20 | Baseline | paralle | 23 |
| # | interaction | 4 | 6 | 0 | 0.4 | 1 | # | 20 | Baseline | paralle | 23 |
| # | disease | 7 | 4 | 0 | 0.636363636 | 1 | # | 20 | Baseline | ancom | 23 |
| # | sex | 4 | 0 | 0 | 1 | 1 | # | 20 | Baseline | ancom | 23 |
| # | time | 5 | 4 | 0 | 0.555555556 | 1 | # | 20 | Baseline | ancom | 23 |
| # | interaction | 0 | 0 | 4 | 0 | 0 | # | 20 | Baseline | ancom | 23 |
| # | disease | 7 | 41 | 0 | 0.145833333 | 1 | # | 20 | Baseline | aldex2 | 23 |
| # | sex | 4 | 12 | 0 | 0.25 | 1 | # | 20 | Baseline | aldex2 | 23 |
| # | time | 5 | 21 | 0 | 0.192307692 | 1 | # | 20 | Baseline | aldex2 | 23 |
| # | interaction | 0 | 0 | 4 | 0 | 0 | # | 20 | Baseline | aldex2 | 23 |
| # | disease | 7 | 0 | 0 | 1 | 1 | # | 20 | Small | paralle | 23 |
| # | sex | 4 | 0 | 0 | 1 | 1 | # | 20 | Small | paralle | 23 |
| # | time | 5 | 0 | 0 | 1 | 1 | # | 20 | Small | paralle | 23 |
| # | interaction | 3 | 0 | 1 | 1 | 0.75 | # | 20 | Small | paralle | 23 |
| # | disease | 7 | 4 | 0 | 0.636363636 | 1 | # | 20 | Small | ancom | 23 |
| # | sex | 4 | 0 | 0 | 1 | 1 | # | 20 | Small | ancom | 23 |
| # | time | 5 | 3 | 0 | 0.625 | 1 | # | 20 | Small | ancom | 23 |
| # | interaction | 0 | 0 | 4 | 0 | 0 | # | 20 | Small | ancom | 23 |
| # | disease | 7 | 6 | 0 | 0.538461538 | 1 | # | 20 | Small | aldex2 | 23 |
| # | sex | 4 | 0 | 0 | 1 | 1 | # | 20 | Small | aldex2 | 23 |
| # | time | 5 | 2 | 0 | 0.714285714 | 1 | # | 20 | Small | aldex2 | 23 |
| # | interaction | 0 | 0 | 4 | 0 | 0 | # | 20 | Small | aldex2 | 23 |
| # | disease | 7 | 9 | 0 | 0.4375 | 1 | # | 20 | Baseline | paralle | 23 |
| # | sex | 4 | 5 | 0 | 0.444444444 | 1 | # | 20 | Baseline | paralle | 23 |
| # | time | 5 | 10 | 0 | 0.333333333 | 1 | # | 20 | Baseline | paralle | 23 |
| # | interaction | 4 | 4 | 0 | 0.5 | 1 | # | 20 | Baseline | paralle | 23 |
| # | disease | 7 | 4 | 0 | 0.636363636 | 1 | # | 20 | Baseline | ancom | 23 |
| # | sex | 4 | 0 | 0 | 1 | 1 | # | 20 | Baseline | ancom | 23 |
| # | time | 5 | 4 | 0 | 0.555555556 | 1 | # | 20 | Baseline | ancom | 23 |
| # | interaction | 0 | 0 | 4 | 0 | 0 | # | 20 | Baseline | ancom | 23 |
| # | disease | 7 | 55 | 0 | 0.112903226 | 1 | # | 20 | Baseline | aldex2 | 23 |
| # | sex | 4 | 17 | 0 | 0.19047619 | 1 | # | 20 | Baseline | aldex2 | 23 |

| | | | | | | | | | | | |
|---|---|---|---|---|---|---|---|---|---|---|---|
| # | time | 5 | 33 | 0 | 0.131578947 | 1 | # | 20 | Baseline | aldex2 | 23 |
| # | interaction | 0 | 0 | 4 | 0 | 0 | # | 20 | Baseline | aldex2 | 23 |
| # | disease | 7 | 0 | 0 | 1 | 1 | # | 20 | Small | paralle | 23 |
| # | sex | 4 | 0 | 0 | 1 | 1 | # | 20 | Small | paralle | 23 |
| # | time | 5 | 0 | 0 | 1 | 1 | # | 20 | Small | paralle | 23 |
| # | interaction | 4 | 0 | 0 | 1 | 1 | # | 20 | Small | paralle | 23 |
| # | disease | 7 | 4 | 0 | 0.636363636 | 1 | # | 20 | Small | ancom | 23 |
| # | sex | 4 | 0 | 0 | 1 | 1 | # | 20 | Small | ancom | 23 |
| # | time | 5 | 4 | 0 | 0.555555556 | 1 | # | 20 | Small | ancom | 23 |
| # | interaction | 0 | 0 | 4 | 0 | 0 | # | 20 | Small | ancom | 23 |
| # | disease | 7 | 8 | 0 | 0.466666667 | 1 | # | 20 | Small | aldex2 | 23 |
| # | sex | 4 | 1 | 0 | 0.8 | 1 | # | 20 | Small | aldex2 | 23 |
| # | time | 5 | 6 | 0 | 0.454545455 | 1 | # | 20 | Small | aldex2 | 23 |
| # | interaction | 0 | 0 | 4 | 0 | 0 | # | 20 | Small | aldex2 | 23 |
| # | disease | 7 | 13 | 0 | 0.35 | 1 | # | 20 | Baseline | paralle | 23 |
| # | sex | 4 | 8 | 0 | 0.333333333 | 1 | # | 20 | Baseline | paralle | 23 |
| # | time | 5 | 7 | 0 | 0.416666667 | 1 | # | 20 | Baseline | paralle | 23 |
| # | interaction | 4 | 8 | 0 | 0.333333333 | 1 | # | 20 | Baseline | paralle | 23 |
| # | disease | 7 | 4 | 0 | 0.636363636 | 1 | # | 20 | Baseline | ancom | 23 |
| # | sex | 4 | 0 | 0 | 1 | 1 | # | 20 | Baseline | ancom | 23 |
| # | time | 5 | 4 | 0 | 0.555555556 | 1 | # | 20 | Baseline | ancom | 23 |
| # | interaction | 0 | 0 | 4 | 0 | 0 | # | 20 | Baseline | ancom | 23 |
| # | disease | 7 | 70 | 0 | 0.090909091 | 1 | # | 20 | Baseline | aldex2 | 23 |
| # | sex | 4 | 24 | 0 | 0.142857143 | 1 | # | 20 | Baseline | aldex2 | 23 |
| # | time | 5 | 67 | 0 | 0.096153846 | 1 | # | 20 | Baseline | aldex2 | 23 |
| # | interaction | 0 | 0 | 4 | 0 | 0 | # | 20 | Baseline | aldex2 | 23 |
| # | disease | 7 | 0 | 0 | 1 | 1 | # | 20 | Small | paralle | 23 |
| # | sex | 4 | 0 | 0 | 1 | 1 | # | 20 | Small | paralle | 23 |
| # | time | 5 | 1 | 0 | 0.833333333 | 1 | # | 20 | Small | paralle | 23 |
| # | interaction | 4 | 0 | 0 | 1 | 1 | # | 20 | Small | paralle | 23 |
| # | disease | 7 | 4 | 0 | 0.636363636 | 1 | # | 20 | Small | ancom | 23 |
| # | sex | 4 | 0 | 0 | 1 | 1 | # | 20 | Small | ancom | 23 |
| # | time | 5 | 4 | 0 | 0.555555556 | 1 | # | 20 | Small | ancom | 23 |
| # | interaction | 0 | 0 | 4 | 0 | 0 | # | 20 | Small | ancom | 23 |
| # | disease | 7 | 12 | 0 | 0.368421053 | 1 | # | 20 | Small | aldex2 | 23 |
| # | sex | 4 | 2 | 0 | 0.666666667 | 1 | # | 20 | Small | aldex2 | 23 |
| # | time | 5 | 5 | 0 | 0.5 | 1 | # | 20 | Small | aldex2 | 23 |
| # | interaction | 0 | 0 | 4 | 0 | 0 | # | 20 | Small | aldex2 | 23 |
| # | disease | 4 | 6 | 0 | 0.4 | 1 | # | 10 | Baseline | paralle | 24 |
| # | sex | 2 | 5 | 0 | 0.285714286 | 1 | # | 10 | Baseline | paralle | 24 |
| # | time | 2 | 8 | 0 | 0.2 | 1 | # | 10 | Baseline | paralle | 24 |
| # | interaction | 2 | 15 | 0 | 0.117647059 | 1 | # | 10 | Baseline | paralle | 24 |
| # | disease | 4 | 0 | 0 | 1 | 1 | # | 10 | Baseline | ancom | 24 |
| # | sex | 2 | 0 | 0 | 1 | 1 | # | 10 | Baseline | ancom | 24 |
| # | time | 2 | 0 | 0 | 1 | 1 | # | 10 | Baseline | ancom | 24 |
| # | interaction | 0 | 0 | 2 | 0 | 0 | # | 10 | Baseline | ancom | 24 |
| # | disease | 4 | 11 | 0 | 0.266666667 | 1 | # | 10 | Baseline | aldex2 | 24 |
| # | sex | 2 | 2 | 0 | 0.5 | 1 | # | 10 | Baseline | aldex2 | 24 |
| # | time | 2 | 4 | 0 | 0.333333333 | 1 | # | 10 | Baseline | aldex2 | 24 |
| # | interaction | 0 | 0 | 2 | 0 | 0 | # | 10 | Baseline | aldex2 | 24 |
| # | disease | 4 | 0 | 0 | 1 | 1 | # | 10 | Small | paralle | 24 |
| # | sex | 2 | 0 | 0 | 1 | 1 | # | 10 | Small | paralle | 24 |
| # | time | 2 | 0 | 0 | 1 | 1 | # | 10 | Small | paralle | 24 |
| # | interaction | 1 | 0 | 1 | 1 | 0.5 | # | 10 | Small | paralle | 24 |
| # | disease | 4 | 0 | 0 | 1 | 1 | # | 10 | Small | ancom | 24 |
| # | sex | 2 | 0 | 0 | 1 | 1 | # | 10 | Small | ancom | 24 |
| # | time | 2 | 0 | 0 | 1 | 1 | # | 10 | Small | ancom | 24 |
| # | interaction | 0 | 0 | 2 | 0 | 0 | # | 10 | Small | ancom | 24 |
| # | disease | 4 | 1 | 0 | 0.8 | 1 | # | 10 | Small | aldex2 | 24 |
| # | sex | 2 | 0 | 0 | 1 | 1 | # | 10 | Small | aldex2 | 24 |
| # | time | 2 | 1 | 0 | 0.666666667 | 1 | # | 10 | Small | aldex2 | 24 |
| # | interaction | 0 | 0 | 2 | 0 | 0 | # | 10 | Small | aldex2 | 24 |
| # | disease | 4 | 4 | 0 | 0.5 | 1 | # | 10 | Baseline | paralle | 24 |
| # | sex | 2 | 6 | 0 | 0.25 | 1 | # | 10 | Baseline | paralle | 24 |
| # | time | 2 | 7 | 0 | 0.222222222 | 1 | # | 10 | Baseline | paralle | 24 |
| # | interaction | 2 | 7 | 0 | 0.222222222 | 1 | # | 10 | Baseline | paralle | 24 |
| # | disease | 4 | 2 | 0 | 0.666666667 | 1 | # | 10 | Baseline | ancom | 24 |
| # | sex | 2 | 0 | 0 | 1 | 1 | # | 10 | Baseline | ancom | 24 |
| # | time | 2 | 1 | 0 | 0.666666667 | 1 | # | 10 | Baseline | ancom | 24 |
| # | interaction | 0 | 0 | 2 | 0 | 0 | # | 10 | Baseline | ancom | 24 |
| # | disease | 4 | 12 | 0 | 0.25 | 1 | # | 10 | Baseline | aldex2 | 24 |
| # | sex | 2 | 3 | 0 | 0.4 | 1 | # | 10 | Baseline | aldex2 | 24 |
| # | time | 2 | 8 | 0 | 0.2 | 1 | # | 10 | Baseline | aldex2 | 24 |
| # | interaction | 0 | 0 | 2 | 0 | 0 | # | 10 | Baseline | aldex2 | 24 |
| # | disease | 4 | 0 | 0 | 1 | 1 | # | 10 | Small | paralle | 24 |
| # | sex | 2 | 0 | 0 | 1 | 1 | # | 10 | Small | paralle | 24 |
| # | time | 2 | 0 | 0 | 1 | 1 | # | 10 | Small | paralle | 24 |
| # | interaction | 2 | 0 | 0 | 1 | 1 | # | 10 | Small | paralle | 24 |
| # | disease | 4 | 2 | 0 | 0.666666667 | 1 | # | 10 | Small | ancom | 24 |
| # | sex | 2 | 0 | 0 | 1 | 1 | # | 10 | Small | ancom | 24 |
| # | time | 2 | 1 | 0 | 0.666666667 | 1 | # | 10 | Small | ancom | 24 |
| # | interaction | 0 | 0 | 2 | 0 | 0 | # | 10 | Small | ancom | 24 |
| # | disease | 4 | 2 | 0 | 0.666666667 | 1 | # | 10 | Small | aldex2 | 24 |
| # | sex | 2 | 0 | 0 | 1 | 1 | # | 10 | Small | aldex2 | 24 |
| # | time | 2 | 1 | 0 | 0.666666667 | 1 | # | 10 | Small | aldex2 | 24 |
| # | interaction | 0 | 0 | 2 | 0 | 0 | # | 10 | Small | aldex2 | 24 |
| # | disease | 4 | 4 | 0 | 0.5 | 1 | # | 10 | Baseline | paralle | 24 |
| # | sex | 2 | 3 | 0 | 0.4 | 1 | # | 10 | Baseline | paralle | 24 |
| # | time | 2 | 6 | 0 | 0.25 | 1 | # | 10 | Baseline | paralle | 24 |
| # | interaction | 2 | 5 | 0 | 0.285714286 | 1 | # | 10 | Baseline | paralle | 24 |
| # | disease | 4 | 2 | 0 | 0.666666667 | 1 | # | 10 | Baseline | ancom | 24 |
| # | sex | 2 | 0 | 0 | 1 | 1 | # | 10 | Baseline | ancom | 24 |
| # | time | 2 | 2 | 0 | 0.5 | 1 | # | 10 | Baseline | ancom | 24 |
| # | interaction | 0 | 0 | 2 | 0 | 0 | # | 10 | Baseline | ancom | 24 |
| # | disease | 4 | 28 | 0 | 0.125 | 1 | # | 10 | Baseline | aldex2 | 24 |
| # | sex | 2 | 2 | 0 | 0.5 | 1 | # | 10 | Baseline | aldex2 | 24 |
| # | time | 2 | 8 | 0 | 0.2 | 1 | # | 10 | Baseline | aldex2 | 24 |
| # | interaction | 0 | 0 | 2 | 0 | 0 | # | 10 | Baseline | aldex2 | 24 |
| # | disease | 4 | 0 | 0 | 1 | 1 | # | 10 | Small | paralle | 24 |
| # | sex | 2 | 0 | 0 | 1 | 1 | # | 10 | Small | paralle | 24 |
| # | time | 2 | 0 | 0 | 1 | 1 | # | 10 | Small | paralle | 24 |
| # | interaction | 2 | 0 | 0 | 1 | 1 | # | 10 | Small | paralle | 24 |
| # | disease | 4 | 2 | 0 | 0.666666667 | 1 | # | 10 | Small | ancom | 24 |
| # | sex | 2 | 0 | 0 | 1 | 1 | # | 10 | Small | ancom | 24 |
| # | time | 2 | 2 | 0 | 0.5 | 1 | # | 10 | Small | ancom | 24 |
| # | interaction | 0 | 0 | 2 | 0 | 0 | # | 10 | Small | ancom | 24 |
| # | disease | 4 | 2 | 0 | 0.666666667 | 1 | # | 10 | Small | aldex2 | 24 |
| # | sex | 2 | 0 | 0 | 1 | 1 | # | 10 | Small | aldex2 | 24 |
| # | time | 2 | 2 | 0 | 0.5 | 1 | # | 10 | Small | aldex2 | 24 |
| # | interaction | 0 | 0 | 2 | 0 | 0 | # | 10 | Small | aldex2 | 24 |
| # | disease | 4 | 3 | 0 | 0.571428571 | 1 | # | 10 | Baseline | paralle | 24 |
| # | sex | 2 | 3 | 0 | 0.4 | 1 | # | 10 | Baseline | paralle | 24 |
| # | time | 2 | 5 | 0 | 0.285714286 | 1 | # | 10 | Baseline | paralle | 24 |
| # | interaction | 2 | 4 | 0 | 0.333333333 | 1 | # | 10 | Baseline | paralle | 24 |
| # | disease | 4 | 2 | 0 | 0.666666667 | 1 | # | 10 | Baseline | ancom | 24 |
| # | sex | 2 | 0 | 0 | 1 | 1 | # | 10 | Baseline | ancom | 24 |
| # | time | 2 | 2 | 0 | 0.5 | 1 | # | 10 | Baseline | ancom | 24 |
| # | interaction | 0 | 0 | 2 | 0 | 0 | # | 10 | Baseline | ancom | 24 |
| # | disease | 4 | 24 | 0 | 0.142857143 | 1 | # | 10 | Baseline | aldex2 | 24 |
| # | sex | 2 | 6 | 0 | 0.25 | 1 | # | 10 | Baseline | aldex2 | 24 |
| # | time | 2 | 8 | 0 | 0.2 | 1 | # | 10 | Baseline | aldex2 | 24 |
| # | interaction | 0 | 0 | 2 | 0 | 0 | # | 10 | Baseline | aldex2 | 24 |
| # | disease | 4 | 0 | 0 | 1 | 1 | # | 10 | Small | paralle | 24 |
| # | sex | 2 | 0 | 0 | 1 | 1 | # | 10 | Small | paralle | 24 |
| # | time | 2 | 0 | 0 | 1 | 1 | # | 10 | Small | paralle | 24 |
| # | interaction | 2 | 0 | 0 | 1 | 1 | # | 10 | Small | paralle | 24 |
| # | disease | 4 | 2 | 0 | 0.666666667 | 1 | # | 10 | Small | ancom | 24 |
| # | sex | 2 | 0 | 0 | 1 | 1 | # | 10 | Small | ancom | 24 |
| # | time | 2 | 2 | 0 | 0.5 | 1 | # | 10 | Small | ancom | 24 |
| # | interaction | 0 | 0 | 2 | 0 | 0 | # | 10 | Small | ancom | 24 |
| # | disease | 4 | 2 | 0 | 0.666666667 | 1 | # | 10 | Small | aldex2 | 24 |
| # | sex | 2 | 0 | 0 | 1 | 1 | # | 10 | Small | aldex2 | 24 |
| # | time | 2 | 2 | 0 | 0.5 | 1 | # | 10 | Small | aldex2 | 24 |
| # | interaction | 0 | 0 | 2 | 0 | 0 | # | 10 | Small | aldex2 | 24 |
| # | disease | 7 | 14 | 0 | 0.333333333 | 1 | # | 20 | Baseline | paralle | 24 |
| # | sex | 4 | 9 | 0 | 0.307692308 | 1 | # | 20 | Baseline | paralle | 24 |
| # | time | 5 | 12 | 0 | 0.294117647 | 1 | # | 20 | Baseline | paralle | 24 |
| # | interaction | 4 | 10 | 0 | 0.285714286 | 1 | # | 20 | Baseline | paralle | 24 |
| # | disease | 7 | 1 | 0 | 0.875 | 1 | # | 20 | Baseline | ancom | 24 |
| # | sex | 4 | 0 | 0 | 1 | 1 | # | 20 | Baseline | ancom | 24 |
| # | time | 5 | 3 | 0 | 0.625 | 1 | # | 20 | Baseline | ancom | 24 |
| # | interaction | 0 | 0 | 4 | 0 | 0 | # | 20 | Baseline | ancom | 24 |
| # | disease | 7 | 27 | 0 | 0.205882353 | 1 | # | 20 | Baseline | aldex2 | 24 |
| # | sex | 4 | 7 | 0 | 0.363636364 | 1 | # | 20 | Baseline | aldex2 | 24 |
| # | time | 5 | 12 | 0 | 0.294117647 | 1 | # | 20 | Baseline | aldex2 | 24 |
| # | interaction | 0 | 0 | 4 | 0 | 0 | # | 20 | Baseline | aldex2 | 24 |
| # | disease | 7 | 2 | 0 | 0.777777778 | 1 | # | 20 | Small | paralle | 24 |
| # | sex | 4 | 0 | 0 | 1 | 1 | # | 20 | Small | paralle | 24 |
| # | time | 5 | 1 | 0 | 0.833333333 | 1 | # | 20 | Small | paralle | 24 |
| # | interaction | 3 | 0 | 1 | 1 | 0.75 | # | 20 | Small | paralle | 24 |
| # | disease | 7 | 1 | 0 | 0.875 | 1 | # | 20 | Small | ancom | 24 |
| # | sex | 4 | 0 | 0 | 1 | 1 | # | 20 | Small | ancom | 24 |
| # | time | 5 | 3 | 0 | 0.625 | 1 | # | 20 | Small | ancom | 24 |
| # | interaction | 0 | 0 | 4 | 0 | 0 | # | 20 | Small | ancom | 24 |
| # | disease | 7 | 1 | 0 | 0.875 | 1 | # | 20 | Small | aldex2 | 24 |
| # | sex | 4 | 0 | 0 | 1 | 1 | # | 20 | Small | aldex2 | 24 |
| # | time | 5 | 3 | 0 | 0.625 | 1 | # | 20 | Small | aldex2 | 24 |
| # | interaction | 0 | 0 | 4 | 0 | 0 | # | 20 | Small | aldex2 | 24 |
| # | disease | 7 | 12 | 0 | 0.368421053 | 1 | # | 20 | Baseline | paralle | 24 |
| # | sex | 4 | 6 | 0 | 0.4 | 1 | # | 20 | Baseline | paralle | 24 |
| # | time | 5 | 11 | 0 | 0.3125 | 1 | # | 20 | Baseline | paralle | 24 |
| # | interaction | 4 | 7 | 0 | 0.363636364 | 1 | # | 20 | Baseline | paralle | 24 |
| # | disease | 7 | 3 | 0 | 0.7 | 1 | # | 20 | Baseline | ancom | 24 |
| # | sex | 4 | 0 | 0 | 1 | 1 | # | 20 | Baseline | ancom | 24 |
| # | time | 5 | 3 | 0 | 0.625 | 1 | # | 20 | Baseline | ancom | 24 |
| # | interaction | 0 | 0 | 4 | 0 | 0 | # | 20 | Baseline | ancom | 24 |
| # | disease | 7 | 33 | 0 | 0.175 | 1 | # | 20 | Baseline | aldex2 | 24 |
| # | sex | 4 | 14 | 0 | 0.222222222 | 1 | # | 20 | Baseline | aldex2 | 24 |
| # | time | 5 | 21 | 0 | 0.192307692 | 1 | # | 20 | Baseline | aldex2 | 24 |
| # | interaction | 0 | 0 | 4 | 0 | 0 | # | 20 | Baseline | aldex2 | 24 |
| # | disease | 7 | 1 | 0 | 0.875 | 1 | # | 20 | Small | paralle | 24 |
| # | sex | 4 | 1 | 0 | 0.8 | 1 | # | 20 | Small | paralle | 24 |
| # | time | 5 | 0 | 0 | 1 | 1 | # | 20 | Small | paralle | 24 |
| # | interaction | 4 | 1 | 0 | 0.8 | 1 | # | 20 | Small | paralle | 24 |
| # | disease | 7 | 3 | 0 | 0.7 | 1 | # | 20 | Small | ancom | 24 |
| # | sex | 4 | 0 | 0 | 1 | 1 | # | 20 | Small | ancom | 24 |
| # | time | 5 | 3 | 0 | 0.625 | 1 | # | 20 | Small | ancom | 24 |
| # | interaction | 0 | 0 | 4 | 0 | 0 | # | 20 | Small | ancom | 24 |
| # | disease | 7 | 2 | 0 | 0.777777778 | 1 | # | 20 | Small | aldex2 | 24 |
| # | sex | 4 | 0 | 0 | 1 | 1 | # | 20 | Small | aldex2 | 24 |
| # | time | 5 | 5 | 0 | 0.5 | 1 | # | 20 | Small | aldex2 | 24 |
| # | interaction | 0 | 0 | 4 | 0 | 0 | # | 20 | Small | aldex2 | 24 |
| # | disease | 7 | 10 | 0 | 0.411764706 | 1 | # | 20 | Baseline | paralle | 24 |
| # | sex | 4 | 9 | 0 | 0.307692308 | 1 | # | 20 | Baseline | paralle | 24 |
| # | time | 5 | 8 | 0 | 0.384615385 | 1 | # | 20 | Baseline | paralle | 24 |
| # | interaction | 4 | 7 | 0 | 0.363636364 | 1 | # | 20 | Baseline | paralle | 24 |
| # | disease | 7 | 3 | 0 | 0.7 | 1 | # | 20 | Baseline | ancom | 24 |
| # | sex | 4 | 0 | 0 | 1 | 1 | # | 20 | Baseline | ancom | 24 |
| # | time | 5 | 4 | 0 | 0.555555556 | 1 | # | 20 | Baseline | ancom | 24 |
| # | interaction | 0 | 0 | 4 | 0 | 0 | # | 20 | Baseline | ancom | 24 |
| # | disease | 7 | 50 | 0 | 0.122807018 | 1 | # | 20 | Baseline | aldex2 | 24 |
| # | sex | 4 | 15 | 0 | 0.210526316 | 1 | # | 20 | Baseline | aldex2 | 24 |
| # | time | 5 | 29 | 0 | 0.147058824 | 1 | # | 20 | Baseline | aldex2 | 24 |
| # | interaction | 0 | 0 | 4 | 0 | 0 | # | 20 | Baseline | aldex2 | 24 |
| # | disease | 7 | 0 | 0 | 1 | 1 | # | 20 | Small | paralle | 24 |
| # | sex | 4 | 0 | 0 | 1 | 1 | # | 20 | Small | paralle | 24 |
| # | time | 5 | 0 | 0 | 1 | 1 | # | 20 | Small | paralle | 24 |
| # | interaction | 4 | 0 | 0 | 1 | 1 | # | 20 | Small | paralle | 24 |
| # | disease | 7 | 3 | 0 | 0.7 | 1 | # | 20 | Small | ancom | 24 |
| # | sex | 4 | 0 | 0 | 1 | 1 | # | 20 | Small | ancom | 24 |
| # | time | 5 | 4 | 0 | 0.555555556 | 1 | # | 20 | Small | ancom | 24 |
| # | interaction | 0 | 0 | 4 | 0 | 0 | # | 20 | Small | ancom | 24 |
| # | disease | 7 | 7 | 0 | 0.5 | 1 | # | 20 | Small | aldex2 | 24 |
| # | sex | 4 | 0 | 0 | 1 | 1 | # | 20 | Small | aldex2 | 24 |
| # | time | 5 | 4 | 0 | 0.555555556 | 1 | # | 20 | Small | aldex2 | 24 |
| # | interaction | 0 | 0 | 4 | 0 | 0 | # | 20 | Small | aldex2 | 24 |
| # | disease | 7 | 11 | 0 | 0.388888889 | 1 | # | 20 | Baseline | paralle | 24 |
| # | sex | 4 | 6 | 0 | 0.4 | 1 | # | 20 | Baseline | paralle | 24 |
| # | time | 5 | 7 | 0 | 0.416666667 | 1 | # | 20 | Baseline | paralle | 24 |
| # | interaction | 4 | 7 | 0 | 0.363636364 | 1 | # | 20 | Baseline | paralle | 24 |
| # | disease | 7 | 4 | 0 | 0.636363636 | 1 | # | 20 | Baseline | ancom | 24 |
| # | sex | 4 | 0 | 0 | 1 | 1 | # | 20 | Baseline | ancom | 24 |
| # | time | 5 | 4 | 0 | 0.555555556 | 1 | # | 20 | Baseline | ancom | 24 |
| # | interaction | 0 | 0 | 4 | 0 | 0 | # | 20 | Baseline | ancom | 24 |
| # | disease | 7 | 57 | 0 | 0.094594595 | 1 | # | 20 | Baseline | aldex2 | 24 |
| # | sex | 4 | 20 | 0 | 0.166666667 | 1 | # | 20 | Baseline | aldex2 | 24 |
| # | time | 5 | 60 | 0 | 0.111111111 | 1 | # | 20 | Baseline | aldex2 | 24 |
| # | interaction | 0 | 0 | 4 | 0 | 0 | # | 20 | Baseline | aldex2 | 24 |
| # | disease | 7 | 1 | 0 | 0.875 | 1 | # | 20 | Small | paralle | 24 |
| # | sex | 4 | 0 | 0 | 1 | 1 | # | 20 | Small | paralle | 24 |
| # | time | 5 | 0 | 0 | 1 | 1 | # | 20 | Small | paralle | 24 |
| # | interaction | 4 | 0 | 0 | 1 | 1 | # | 20 | Small | paralle | 24 |
| # | disease | 7 | 4 | 0 | 0.636363636 | 1 | # | 20 | Small | ancom | 24 |
| # | sex | 4 | 0 | 0 | 1 | 1 | # | 20 | Small | ancom | 24 |
| # | time | 5 | 4 | 0 | 0.555555556 | 1 | # | 20 | Small | ancom | 24 |
| # | interaction | 0 | 0 | 4 | 0 | 0 | # | 20 | Small | ancom | 24 |
| # | disease | 7 | 11 | 0 | 0.388888889 | 1 | # | 20 | Small | aldex2 | 24 |
| # | sex | 4 | 2 | 0 | 0.666666667 | 1 | # | 20 | Small | aldex2 | 24 |
| # | time | 5 | 6 | 0 | 0.454545455 | 1 | # | 20 | Small | aldex2 | 24 |
| # | interaction | 0 | 0 | 4 | 0 | 0 | # | 20 | Small | aldex2 | 24 |
| # | disease | 4 | 6 | 0 | 0.4 | 1 | # | 10 | Baseline | paralle | 25 |
| # | sex | 2 | 7 | 0 | 0.222222222 | 1 | # | 10 | Baseline | paralle | 25 |
| # | time | 2 | 10 | 0 | 0.166666667 | 1 | # | 10 | Baseline | paralle | 25 |
| # | interaction | 2 | 8 | 0 | 0.2 | 1 | # | 10 | Baseline | paralle | 25 |
| # | disease | 4 | 1 | 0 | 0.8 | 1 | # | 10 | Baseline | ancom | 25 |
| # | sex | 2 | 0 | 0 | 1 | 1 | # | 10 | Baseline | ancom | 25 |
| # | time | 2 | 0 | 0 | 1 | 1 | # | 10 | Baseline | ancom | 25 |
| # | interaction | 0 | 0 | 2 | 0 | 0 | # | 10 | Baseline | ancom | 25 |
| # | disease | 4 | 11 | 0 | 0.266666667 | 1 | # | 10 | Baseline | aldex2 | 25 |
| # | sex | 2 | 5 | 0 | 0.285714286 | 1 | # | 10 | Baseline | aldex2 | 25 |
| # | time | 2 | 12 | 0 | 0.142857143 | 1 | # | 10 | Baseline | aldex2 | 25 |
| # | interaction | 0 | 0 | 2 | 0 | 0 | # | 10 | Baseline | aldex2 | 25 |
| # | disease | 4 | 0 | 0 | 1 | 1 | # | 10 | Small | paralle | 25 |
| # | sex | 2 | 0 | 0 | 1 | 1 | # | 10 | Small | paralle | 25 |
| # | time | 2 | 0 | 0 | 1 | 1 | # | 10 | Small | paralle | 25 |
| # | interaction | 0 | 1 | 2 | 0 | 0 | # | 10 | Small | paralle | 25 |
| # | disease | 4 | 1 | 0 | 0.8 | 1 | # | 10 | Small | ancom | 25 |
| # | sex | 2 | 0 | 0 | 1 | 1 | # | 10 | Small | ancom | 25 |
| # | time | 2 | 0 | 0 | 1 | 1 | # | 10 | Small | ancom | 25 |
| # | interaction | 0 | 0 | 2 | 0 | 0 | # | 10 | Small | ancom | 25 |
| # | disease | 4 | 1 | 0 | 0.8 | 1 | # | 10 | Small | aldex2 | 25 |
| # | sex | 2 | 0 | 0 | 1 | 1 | # | 10 | Small | aldex2 | 25 |
| # | time | 2 | 1 | 0 | 0.666666667 | 1 | # | 10 | Small | aldex2 | 25 |
| # | interaction | 0 | 0 | 2 | 0 | 0 | # | 10 | Small | aldex2 | 25 |
| # | disease | 4 | 7 | 0 | 0.363636364 | 1 | # | 10 | Baseline | paralle | 25 |
| # | sex | 2 | 4 | 0 | 0.333333333 | 1 | # | 10 | Baseline | paralle | 25 |
| # | time | 2 | 6 | 0 | 0.25 | 1 | # | 10 | Baseline | paralle | 25 |
| # | interaction | 2 | 4 | 0 | 0.333333333 | 1 | # | 10 | Baseline | paralle | 25 |
| # | disease | 4 | 2 | 0 | 0.666666667 | 1 | # | 10 | Baseline | ancom | 25 |
| # | sex | 2 | 0 | 0 | 1 | 1 | # | 10 | Baseline | ancom | 25 |
| # | time | 2 | 3 | 0 | 0.4 | 1 | # | 10 | Baseline | ancom | 25 |
| # | interaction | 0 | 0 | 2 | 0 | 0 | # | 10 | Baseline | ancom | 25 |
| # | disease | 4 | 13 | 0 | 0.235294118 | 1 | # | 10 | Baseline | aldex2 | 25 |
| # | sex | 2 | 4 | 0 | 0.333333333 | 1 | # | 10 | Baseline | aldex2 | 25 |
| # | time | 2 | 10 | 0 | 0.166666667 | 1 | # | 10 | Baseline | aldex2 | 25 |
| # | interaction | 0 | 0 | 2 | 0 | 0 | # | 10 | Baseline | aldex2 | 25 |
| # | disease | 4 | 0 | 0 | 1 | 1 | # | 10 | Small | paralle | 25 |
| # | sex | 2 | 0 | 0 | 1 | 1 | # | 10 | Small | paralle | 25 |
| # | time | 2 | 0 | 0 | 1 | 1 | # | 10 | Small | paralle | 25 |
| # | interaction | 2 | 0 | 0 | 1 | 1 | # | 10 | Small | paralle | 25 |
| # | disease | 4 | 2 | 0 | 0.666666667 | 1 | # | 10 | Small | ancom | 25 |
| # | sex | 2 | 0 | 0 | 1 | 1 | # | 10 | Small | ancom | 25 |
| # | time | 2 | 3 | 0 | 0.4 | 1 | # | 10 | Small | ancom | 25 |
| # | interaction | 0 | 0 | 2 | 0 | 0 | # | 10 | Small | ancom | 25 |
| # | disease | 4 | 3 | 0 | 0.571428571 | 1 | # | 10 | Small | aldex2 | 25 |
| # | sex | 2 | 1 | 0 | 0.666666667 | 1 | # | 10 | Small | aldex2 | 25 |
| # | time | 2 | 3 | 0 | 0.4 | 1 | # | 10 | Small | aldex2 | 25 |
| # | interaction | 0 | 0 | 2 | 0 | 0 | # | 10 | Small | aldex2 | 25 |
| # | disease | 4 | 3 | 0 | 0.571428571 | 1 | # | 10 | Baseline | paralle | 25 |
| # | sex | 2 | 3 | 0 | 0.4 | 1 | # | 10 | Baseline | paralle | 25 |
| # | time | 2 | 3 | 0 | 0.4 | 1 | # | 10 | Baseline | paralle | 25 |
| # | interaction | 2 | 2 | 0 | 0.5 | 1 | # | 10 | Baseline | paralle | 25 |
| # | disease | 4 | 2 | 0 | 0.666666667 | 1 | # | 10 | Baseline | ancom | 25 |
| # | sex | 2 | 0 | 0 | 1 | 1 | # | 10 | Baseline | ancom | 25 |
| # | time | 2 | 2 | 0 | 0.5 | 1 | # | 10 | Baseline | ancom | 25 |
| # | interaction | 0 | 0 | 2 | 0 | 0 | # | 10 | Baseline | ancom | 25 |
| # | disease | 4 | 15 | 0 | 0.210526316 | 1 | # | 10 | Baseline | aldex2 | 25 |
| # | sex | 2 | 6 | 0 | 0.25 | 1 | # | 10 | Baseline | aldex2 | 25 |
| # | time | 2 | 14 | 0 | 0.125 | 1 | # | 10 | Baseline | aldex2 | 25 |
| # | interaction | 0 | 0 | 2 | 0 | 0 | # | 10 | Baseline | aldex2 | 25 |
| # | disease | 4 | 0 | 0 | 1 | 1 | # | 10 | Small | paralle | 25 |
| # | sex | 2 | 0 | 0 | 1 | 1 | # | 10 | Small | paralle | 25 |
| # | time | 2 | 0 | 0 | 1 | 1 | # | 10 | Small | paralle | 25 |
| # | interaction | 2 | 0 | 0 | 1 | 1 | # | 10 | Small | paralle | 25 |
| # | disease | 4 | 2 | 0 | 0.666666667 | 1 | # | 10 | Small | ancom | 25 |
| # | sex | 2 | 0 | 0 | 1 | 1 | # | 10 | Small | ancom | 25 |
| # | time | 2 | 2 | 0 | 0.5 | 1 | # | 10 | Small | ancom | 25 |
| # | interaction | 0 | 0 | 2 | 0 | 0 | # | 10 | Small | ancom | 25 |
| # | disease | 4 | 3 | 0 | 0.571428571 | 1 | # | 10 | Small | aldex2 | 25 |
| # | sex | 2 | 1 | 0 | 0.666666667 | 1 | # | 10 | Small | aldex2 | 25 |
| # | time | 2 | 2 | 0 | 0.5 | 1 | # | 10 | Small | aldex2 | 25 |
| # | interaction | 0 | 0 | 2 | 0 | 0 | # | 10 | Small | aldex2 | 25 |
| # | disease | 4 | 1 | 0 | 0.8 | 1 | # | 10 | Baseline | paralle | 25 |
| # | sex | 2 | 1 | 0 | 0.666666667 | 1 | # | 10 | Baseline | paralle | 25 |
| # | time | 2 | 2 | 0 | 0.5 | 1 | # | 10 | Baseline | paralle | 25 |
| # | interaction | 2 | 2 | 0 | 0.5 | 1 | # | 10 | Baseline | paralle | 25 |
| # | disease | 4 | 2 | 0 | 0.666666667 | 1 | # | 10 | Baseline | ancom | 25 |
| # | sex | 2 | 0 | 0 | 1 | 1 | # | 10 | Baseline | ancom | 25 |
| # | time | 2 | 2 | 0 | 0.5 | 1 | # | 10 | Baseline | ancom | 25 |
| # | interaction | 0 | 0 | 2 | 0 | 0 | # | 10 | Baseline | ancom | 25 |
| # | disease | 4 | 21 | 0 | 0.16 | 1 | # | 10 | Baseline | aldex2 | 25 |
| # | sex | 2 | 6 | 0 | 0.25 | 1 | # | 10 | Baseline | aldex2 | 25 |
| # | time | 2 | 19 | 0 | 0.095238095 | 1 | # | 10 | Baseline | aldex2 | 25 |
| # | interaction | 0 | 0 | 2 | 0 | 0 | # | 10 | Baseline | aldex2 | 25 |
| # | disease | 4 | 0 | 0 | 1 | 1 | # | 10 | Small | paralle | 25 |
| # | sex | 2 | 0 | 0 | 1 | 1 | # | 10 | Small | paralle | 25 |
| # | time | 2 | 0 | 0 | 1 | 1 | # | 10 | Small | paralle | 25 |
| # | interaction | 2 | 0 | 0 | 1 | 1 | # | 10 | Small | paralle | 25 |
| # | disease | 4 | 2 | 0 | 0.666666667 | 1 | # | 10 | Small | ancom | 25 |
| # | sex | 2 | 0 | 0 | 1 | 1 | # | 10 | Small | ancom | 25 |
| # | time | 2 | 2 | 0 | 0.5 | 1 | # | 10 | Small | ancom | 25 |
| # | interaction | 0 | 0 | 2 | 0 | 0 | # | 10 | Small | ancom | 25 |
| # | disease | 4 | 3 | 0 | 0.571428571 | 1 | # | 10 | Small | aldex2 | 25 |
| # | sex | 2 | 1 | 0 | 0.666666667 | 1 | # | 10 | Small | aldex2 | 25 |
| # | time | 2 | 2 | 0 | 0.5 | 1 | # | 10 | Small | aldex2 | 25 |
| # | interaction | 0 | 0 | 2 | 0 | 0 | # | 10 | Small | aldex2 | 25 |
| # | disease | 7 | 16 | 0 | 0.304347826 | 1 | # | 20 | Baseline | paralle | 25 |
| # | sex | 4 | 15 | 0 | 0.210526316 | 1 | # | 20 | Baseline | paralle | 25 |
| # | time | 5 | 10 | 0 | 0.333333333 | 1 | # | 20 | Baseline | paralle | 25 |
| # | interaction | 4 | 10 | 0 | 0.285714286 | 1 | # | 20 | Baseline | paralle | 25 |
| # | disease | 7 | 1 | 0 | 0.875 | 1 | # | 20 | Baseline | ancom | 25 |
| # | sex | 4 | 0 | 0 | 1 | 1 | # | 20 | Baseline | ancom | 25 |
| # | time | 5 | 2 | 0 | 0.714285714 | 1 | # | 20 | Baseline | ancom | 25 |
| # | interaction | 0 | 0 | 4 | 0 | 0 | # | 20 | Baseline | ancom | 25 |
| # | disease | 7 | 16 | 0 | 0.304347826 | 1 | # | 20 | Baseline | aldex2 | 25 |
| # | sex | 4 | 7 | 0 | 0.363636364 | 1 | # | 20 | Baseline | aldex2 | 25 |
| # | time | 5 | 18 | 0 | 0.217391304 | 1 | # | 20 | Baseline | aldex2 | 25 |

| | | | | | | | | | | | |
|---|---|---|---|---|---|---|---|---|---|---|---|
| # | interaction | 0 | 0 | 4 | 0 | 0 | # | 20 | Baseline | aldex2 | 25 |
| # | disease | 7 | 0 | 0 | 1 | 1 | # | 20 | Small | paralle | 25 |
| # | sex | 4 | 1 | 0 | 0.8 | 1 | # | 20 | Small | paralle | 25 |
| # | time | 5 | 0 | 0 | 1 | 1 | # | 20 | Small | paralle | 25 |
| # | interaction | 1 | 0 | 3 | 1 | 0.25 | # | 20 | Small | paralle | 25 |
| # | disease | 7 | 0 | 0 | 1 | 1 | # | 20 | Small | ancom | 25 |
| # | sex | 4 | 0 | 0 | 1 | 1 | # | 20 | Small | ancom | 25 |
| # | time | 5 | 2 | 0 | 0.714285714 | 1 | # | 20 | Small | ancom | 25 |
| # | interaction | 0 | 0 | 4 | 0 | 0 | # | 20 | Small | ancom | 25 |
| # | disease | 7 | 0 | 0 | 1 | 1 | # | 20 | Small | aldex2 | 25 |
| # | sex | 4 | 0 | 0 | 1 | 1 | # | 20 | Small | aldex2 | 25 |
| # | time | 5 | 1 | 0 | 0.833333333 | 1 | # | 20 | Small | aldex2 | 25 |
| # | interaction | 0 | 0 | 4 | 0 | 0 | # | 20 | Small | aldex2 | 25 |
| # | disease | 7 | 12 | 0 | 0.368421053 | 1 | # | 20 | Baseline | paralle | 25 |
| # | sex | 4 | 11 | 0 | 0.266666667 | 1 | # | 20 | Baseline | paralle | 25 |
| # | time | 5 | 8 | 0 | 0.384615385 | 1 | # | 20 | Baseline | paralle | 25 |
| # | interaction | 4 | 11 | 0 | 0.266666667 | 1 | # | 20 | Baseline | paralle | 25 |
| # | disease | 7 | 2 | 0 | 0.777777778 | 1 | # | 20 | Baseline | ancom | 25 |
| # | sex | 4 | 0 | 0 | 1 | 1 | # | 20 | Baseline | ancom | 25 |
| # | time | 5 | 4 | 0 | 0.555555556 | 1 | # | 20 | Baseline | ancom | 25 |
| # | interaction | 0 | 0 | 4 | 0 | 0 | # | 20 | Baseline | ancom | 25 |
| # | disease | 7 | 31 | 0 | 0.184210526 | 1 | # | 20 | Baseline | aldex2 | 25 |
| # | sex | 4 | 9 | 0 | 0.307692308 | 1 | # | 20 | Baseline | aldex2 | 25 |
| # | time | 5 | 27 | 0 | 0.15625 | 1 | # | 20 | Baseline | aldex2 | 25 |
| # | interaction | 0 | 0 | 4 | 0 | 0 | # | 20 | Baseline | aldex2 | 25 |
| # | disease | 7 | 1 | 0 | 0.875 | 1 | # | 20 | Small | paralle | 25 |
| # | sex | 4 | 0 | 0 | 1 | 1 | # | 20 | Small | paralle | 25 |
| # | time | 5 | 0 | 0 | 1 | 1 | # | 20 | Small | paralle | 25 |
| # | interaction | 4 | 0 | 0 | 1 | 1 | # | 20 | Small | paralle | 25 |
| # | disease | 7 | 2 | 0 | 0.777777778 | 1 | # | 20 | Small | ancom | 25 |
| # | sex | 4 | 0 | 0 | 1 | 1 | # | 20 | Small | ancom | 25 |
| # | time | 5 | 4 | 0 | 0.555555556 | 1 | # | 20 | Small | ancom | 25 |
| # | interaction | 0 | 0 | 4 | 0 | 0 | # | 20 | Small | ancom | 25 |
| # | disease | 7 | 3 | 0 | 0.7 | 1 | # | 20 | Small | aldex2 | 25 |
| # | sex | 4 | 0 | 0 | 1 | 1 | # | 20 | Small | aldex2 | 25 |
| # | time | 5 | 4 | 0 | 0.555555556 | 1 | # | 20 | Small | aldex2 | 25 |
| # | interaction | 0 | 0 | 4 | 0 | 0 | # | 20 | Small | aldex2 | 25 |
| # | disease | 7 | 14 | 0 | 0.333333333 | 1 | # | 20 | Baseline | paralle | 25 |
| # | sex | 4 | 7 | 0 | 0.363636364 | 1 | # | 20 | Baseline | paralle | 25 |
| # | time | 5 | 4 | 0 | 0.555555556 | 1 | # | 20 | Baseline | paralle | 25 |
| # | interaction | 4 | 12 | 0 | 0.25 | 1 | # | 20 | Baseline | paralle | 25 |
| # | disease | 7 | 4 | 0 | 0.636363636 | 1 | # | 20 | Baseline | ancom | 25 |
| # | sex | 4 | 0 | 0 | 1 | 1 | # | 20 | Baseline | ancom | 25 |
| # | time | 5 | 4 | 0 | 0.555555556 | 1 | # | 20 | Baseline | ancom | 25 |
| # | interaction | 0 | 0 | 4 | 0 | 0 | # | 20 | Baseline | ancom | 25 |
| # | disease | 7 | 40 | 0 | 0.14893617 | 1 | # | 20 | Baseline | aldex2 | 25 |
| # | sex | 4 | 14 | 0 | 0.222222222 | 1 | # | 20 | Baseline | aldex2 | 25 |
| # | time | 5 | 38 | 0 | 0.11627907 | 1 | # | 20 | Baseline | aldex2 | 25 |
| # | interaction | 0 | 0 | 4 | 0 | 0 | # | 20 | Baseline | aldex2 | 25 |
| # | disease | 7 | 0 | 0 | 1 | 1 | # | 20 | Small | paralle | 25 |
| # | sex | 4 | 0 | 0 | 1 | 1 | # | 20 | Small | paralle | 25 |
| # | time | 5 | 0 | 0 | 1 | 1 | # | 20 | Small | paralle | 25 |
| # | interaction | 4 | 0 | 0 | 1 | 1 | # | 20 | Small | paralle | 25 |
| # | disease | 7 | 4 | 0 | 0.636363636 | 1 | # | 20 | Small | ancom | 25 |
| # | sex | 4 | 0 | 0 | 1 | 1 | # | 20 | Small | ancom | 25 |
| # | time | 5 | 4 | 0 | 0.555555556 | 1 | # | 20 | Small | ancom | 25 |
| # | interaction | 0 | 0 | 4 | 0 | 0 | # | 20 | Small | ancom | 25 |
| # | disease | 7 | 6 | 0 | 0.538461538 | 1 | # | 20 | Small | aldex2 | 25 |
| # | sex | 4 | 0 | 0 | 1 | 1 | # | 20 | Small | aldex2 | 25 |
| # | time | 5 | 5 | 0 | 0.5 | 1 | # | 20 | Small | aldex2 | 25 |
| # | interaction | 0 | 0 | 4 | 0 | 0 | # | 20 | Small | aldex2 | 25 |
| # | disease | 7 | 9 | 0 | 0.4375 | 1 | # | 20 | Baseline | paralle | 25 |
| # | sex | 4 | 7 | 0 | 0.363636364 | 1 | # | 20 | Baseline | paralle | 25 |
| # | time | 5 | 7 | 0 | 0.416666667 | 1 | # | 20 | Baseline | paralle | 25 |
| # | interaction | 4 | 8 | 0 | 0.333333333 | 1 | # | 20 | Baseline | paralle | 25 |
| # | disease | 7 | 4 | 0 | 0.636363636 | 1 | # | 20 | Baseline | ancom | 25 |
| # | sex | 4 | 0 | 0 | 1 | 1 | # | 20 | Baseline | ancom | 25 |
| # | time | 5 | 4 | 0 | 0.555555556 | 1 | # | 20 | Baseline | ancom | 25 |
| # | interaction | 0 | 0 | 4 | 0 | 0 | # | 20 | Baseline | ancom | 25 |
| # | disease | 7 | 54 | 0 | 0.114754098 | 1 | # | 20 | Baseline | aldex2 | 25 |
| # | sex | 4 | 16 | 0 | 0.2 | 1 | # | 20 | Baseline | aldex2 | 25 |
| # | time | 5 | 47 | 0 | 0.096153846 | 1 | # | 20 | Baseline | aldex2 | 25 |
| # | interaction | 0 | 0 | 4 | 0 | 0 | # | 20 | Baseline | aldex2 | 25 |
| # | disease | 7 | 0 | 0 | 1 | 1 | # | 20 | Small | paralle | 25 |
| # | sex | 4 | 0 | 0 | 1 | 1 | # | 20 | Small | paralle | 25 |
| # | time | 5 | 0 | 0 | 1 | 1 | # | 20 | Small | paralle | 25 |
| # | interaction | 4 | 1 | 0 | 0.8 | 1 | # | 20 | Small | paralle | 25 |
| # | disease | 7 | 4 | 0 | 0.636363636 | 1 | # | 20 | Small | ancom | 25 |
| # | sex | 4 | 0 | 0 | 1 | 1 | # | 20 | Small | ancom | 25 |
| # | time | 5 | 4 | 0 | 0.555555556 | 1 | # | 20 | Small | ancom | 25 |
| # | interaction | 0 | 0 | 4 | 0 | 0 | # | 20 | Small | ancom | 25 |
| # | disease | 7 | 10 | 0 | 0.411764706 | 1 | # | 20 | Small | aldex2 | 25 |
| # | sex | 4 | 0 | 0 | 1 | 1 | # | 20 | Small | aldex2 | 25 |
| # | time | 5 | 7 | 0 | 0.416666667 | 1 | # | 20 | Small | aldex2 | 25 |
| # | interaction | 0 | 0 | 4 | 0 | 0 | # | 20 | Small | aldex2 | 25 |
| # | disease | 4 | 8 | 0 | 0.333333333 | 1 | # | 10 | Baseline | paralle | 26 |
| # | sex | 2 | 6 | 0 | 0.25 | 1 | # | 10 | Baseline | paralle | 26 |
| # | time | 2 | 10 | 0 | 0.166666667 | 1 | # | 10 | Baseline | paralle | 26 |
| # | interaction | 2 | 11 | 0 | 0.153846154 | 1 | # | 10 | Baseline | paralle | 26 |
| # | disease | 4 | 0 | 0 | 1 | 1 | # | 10 | Baseline | ancom | 26 |
| # | sex | 2 | 0 | 0 | 1 | 1 | # | 10 | Baseline | ancom | 26 |
| # | time | 2 | 0 | 0 | 1 | 1 | # | 10 | Baseline | ancom | 26 |
| # | interaction | 0 | 0 | 2 | 0 | 0 | # | 10 | Baseline | ancom | 26 |
| # | disease | 4 | 8 | 0 | 0.333333333 | 1 | # | 10 | Baseline | aldex2 | 26 |
| # | sex | 2 | 5 | 0 | 0.285714286 | 1 | # | 10 | Baseline | aldex2 | 26 |
| # | time | 2 | 6 | 0 | 0.25 | 1 | # | 10 | Baseline | aldex2 | 26 |
| # | interaction | 0 | 0 | 2 | 0 | 0 | # | 10 | Baseline | aldex2 | 26 |
| # | disease | 4 | 0 | 0 | 1 | 1 | # | 10 | Small | paralle | 26 |
| # | sex | 2 | 0 | 0 | 1 | 1 | # | 10 | Small | paralle | 26 |
| # | time | 2 | 0 | 0 | 1 | 1 | # | 10 | Small | paralle | 26 |
| # | interaction | 2 | 0 | 0 | 1 | 1 | # | 10 | Small | paralle | 26 |
| # | disease | 4 | 0 | 0 | 1 | 1 | # | 10 | Small | ancom | 26 |
| # | sex | 2 | 0 | 0 | 1 | 1 | # | 10 | Small | ancom | 26 |
| # | time | 2 | 0 | 0 | 1 | 1 | # | 10 | Small | ancom | 26 |
| # | interaction | 0 | 0 | 2 | 0 | 0 | # | 10 | Small | ancom | 26 |
| # | disease | 4 | 1 | 0 | 0.8 | 1 | # | 10 | Small | aldex2 | 26 |
| # | sex | 2 | 0 | 0 | 1 | 1 | # | 10 | Small | aldex2 | 26 |
| # | time | 2 | 0 | 0 | 1 | 1 | # | 10 | Small | aldex2 | 26 |
| # | interaction | 0 | 0 | 2 | 0 | 0 | # | 10 | Small | aldex2 | 26 |
| # | disease | 4 | 7 | 0 | 0.363636364 | 1 | # | 10 | Baseline | paralle | 26 |
| # | sex | 2 | 3 | 0 | 0.4 | 1 | # | 10 | Baseline | paralle | 26 |
| # | time | 2 | 7 | 0 | 0.222222222 | 1 | # | 10 | Baseline | paralle | 26 |
| # | interaction | 2 | 4 | 0 | 0.333333333 | 1 | # | 10 | Baseline | paralle | 26 |
| # | disease | 4 | 2 | 0 | 0.666666667 | 1 | # | 10 | Baseline | ancom | 26 |
| # | sex | 2 | 0 | 0 | 1 | 1 | # | 10 | Baseline | ancom | 26 |
| # | time | 2 | 0 | 0 | 1 | 1 | # | 10 | Baseline | ancom | 26 |
| # | interaction | 0 | 0 | 2 | 0 | 0 | # | 10 | Baseline | ancom | 26 |
| # | disease | 4 | 16 | 0 | 0.2 | 1 | # | 10 | Baseline | aldex2 | 26 |
| # | sex | 2 | 4 | 0 | 0.333333333 | 1 | # | 10 | Baseline | aldex2 | 26 |
| # | time | 2 | 8 | 0 | 0.2 | 1 | # | 10 | Baseline | aldex2 | 26 |
| # | interaction | 0 | 0 | 2 | 0 | 0 | # | 10 | Baseline | aldex2 | 26 |
| # | disease | 4 | 0 | 0 | 1 | 1 | # | 10 | Small | paralle | 26 |
| # | sex | 2 | 0 | 0 | 1 | 1 | # | 10 | Small | paralle | 26 |
| # | time | 2 | 0 | 0 | 1 | 1 | # | 10 | Small | paralle | 26 |
| # | interaction | 2 | 0 | 0 | 1 | 1 | # | 10 | Small | paralle | 26 |
| # | disease | 4 | 2 | 0 | 0.666666667 | 1 | # | 10 | Small | ancom | 26 |
| # | sex | 2 | 0 | 0 | 1 | 1 | # | 10 | Small | ancom | 26 |
| # | time | 2 | 0 | 0 | 1 | 1 | # | 10 | Small | ancom | 26 |
| # | interaction | 0 | 0 | 2 | 0 | 0 | # | 10 | Small | ancom | 26 |
| # | disease | 4 | 3 | 0 | 0.571428571 | 1 | # | 10 | Small | aldex2 | 26 |
| # | sex | 2 | 0 | 0 | 1 | 1 | # | 10 | Small | aldex2 | 26 |
| # | time | 2 | 1 | 0 | 0.666666667 | 1 | # | 10 | Small | aldex2 | 26 |
| # | interaction | 0 | 0 | 2 | 0 | 0 | # | 10 | Small | aldex2 | 26 |
| # | disease | 4 | 4 | 0 | 0.5 | 1 | # | 10 | Baseline | paralle | 26 |
| # | sex | 2 | 5 | 0 | 0.285714286 | 1 | # | 10 | Baseline | paralle | 26 |
| # | time | 2 | 8 | 0 | 0.2 | 1 | # | 10 | Baseline | paralle | 26 |
| # | interaction | 2 | 8 | 0 | 0.2 | 1 | # | 10 | Baseline | paralle | 26 |
| # | disease | 4 | 2 | 0 | 0.666666667 | 1 | # | 10 | Baseline | ancom | 26 |
| # | sex | 2 | 0 | 0 | 1 | 1 | # | 10 | Baseline | ancom | 26 |
| # | time | 2 | 2 | 0 | 0.5 | 1 | # | 10 | Baseline | ancom | 26 |
| # | interaction | 0 | 0 | 2 | 0 | 0 | # | 10 | Baseline | ancom | 26 |
| # | disease | 4 | 21 | 0 | 0.16 | 1 | # | 10 | Baseline | aldex2 | 26 |
| # | sex | 2 | 8 | 0 | 0.2 | 1 | # | 10 | Baseline | aldex2 | 26 |
| # | time | 2 | 8 | 0 | 0.2 | 1 | # | 10 | Baseline | aldex2 | 26 |
| # | interaction | 0 | 0 | 2 | 0 | 0 | # | 10 | Baseline | aldex2 | 26 |
| # | disease | 4 | 0 | 0 | 1 | 1 | # | 10 | Small | paralle | 26 |
| # | sex | 2 | 0 | 0 | 1 | 1 | # | 10 | Small | paralle | 26 |
| # | time | 2 | 0 | 0 | 1 | 1 | # | 10 | Small | paralle | 26 |
| # | interaction | 2 | 0 | 0 | 1 | 1 | # | 10 | Small | paralle | 26 |
| # | disease | 4 | 2 | 0 | 0.666666667 | 1 | # | 10 | Small | ancom | 26 |
| # | sex | 2 | 0 | 0 | 1 | 1 | # | 10 | Small | ancom | 26 |
| # | time | 2 | 2 | 0 | 0.5 | 1 | # | 10 | Small | ancom | 26 |
| # | interaction | 0 | 0 | 2 | 0 | 0 | # | 10 | Small | ancom | 26 |
| # | disease | 4 | 3 | 0 | 0.571428571 | 1 | # | 10 | Small | aldex2 | 26 |
| # | sex | 2 | 0 | 0 | 1 | 1 | # | 10 | Small | aldex2 | 26 |
| # | time | 2 | 2 | 0 | 0.5 | 1 | # | 10 | Small | aldex2 | 26 |
| # | interaction | 0 | 0 | 2 | 0 | 0 | # | 10 | Small | aldex2 | 26 |
| # | disease | 4 | 8 | 0 | 0.333333333 | 1 | # | 10 | Baseline | paralle | 26 |
| # | sex | 2 | 3 | 0 | 0.4 | 1 | # | 10 | Baseline | paralle | 26 |
| # | time | 2 | 4 | 0 | 0.333333333 | 1 | # | 10 | Baseline | paralle | 26 |
| # | interaction | 2 | 1 | 0 | 0.666666667 | 1 | # | 10 | Baseline | paralle | 26 |
| # | disease | 4 | 2 | 0 | 0.666666667 | 1 | # | 10 | Baseline | ancom | 26 |
| # | sex | 2 | 0 | 0 | 1 | 1 | # | 10 | Baseline | ancom | 26 |
| # | time | 2 | 2 | 0 | 0.5 | 1 | # | 10 | Baseline | ancom | 26 |
| # | interaction | 0 | 0 | 2 | 0 | 0 | # | 10 | Baseline | ancom | 26 |
| # | disease | 4 | 23 | 0 | 0.148148148 | 1 | # | 10 | Baseline | aldex2 | 26 |
| # | sex | 2 | 10 | 0 | 0.166666667 | 1 | # | 10 | Baseline | aldex2 | 26 |
| # | time | 2 | 8 | 0 | 0.2 | 1 | # | 10 | Baseline | aldex2 | 26 |
| # | interaction | 0 | 0 | 2 | 0 | 0 | # | 10 | Baseline | aldex2 | 26 |
| # | disease | 4 | 0 | 0 | 1 | 1 | # | 10 | Small | paralle | 26 |
| # | sex | 2 | 0 | 0 | 1 | 1 | # | 10 | Small | paralle | 26 |
| # | time | 2 | 0 | 0 | 1 | 1 | # | 10 | Small | paralle | 26 |
| # | interaction | 2 | 0 | 0 | 1 | 1 | # | 10 | Small | paralle | 26 |
| # | disease | 4 | 2 | 0 | 0.666666667 | 1 | # | 10 | Small | ancom | 26 |
| # | sex | 2 | 0 | 0 | 1 | 1 | # | 10 | Small | ancom | 26 |
| # | time | 2 | 2 | 0 | 0.5 | 1 | # | 10 | Small | ancom | 26 |
| # | interaction | 0 | 0 | 2 | 0 | 0 | # | 10 | Small | ancom | 26 |
| # | disease | 4 | 3 | 0 | 0.571428571 | 1 | # | 10 | Small | aldex2 | 26 |
| # | sex | 2 | 0 | 0 | 1 | 1 | # | 10 | Small | aldex2 | 26 |
| # | time | 2 | 2 | 0 | 0.5 | 1 | # | 10 | Small | aldex2 | 26 |
| # | interaction | 0 | 0 | 2 | 0 | 0 | # | 10 | Small | aldex2 | 26 |
| # | disease | 7 | 8 | 0 | 0.466666667 | 1 | # | 20 | Baseline | paralle | 26 |
| # | sex | 4 | 5 | 0 | 0.444444444 | 1 | # | 20 | Baseline | paralle | 26 |
| # | time | 5 | 8 | 0 | 0.384615385 | 1 | # | 20 | Baseline | paralle | 26 |
| # | interaction | 4 | 9 | 0 | 0.307692308 | 1 | # | 20 | Baseline | paralle | 26 |
| # | disease | 7 | 0 | 0 | 1 | 1 | # | 20 | Baseline | ancom | 26 |
| # | sex | 4 | 0 | 0 | 1 | 1 | # | 20 | Baseline | ancom | 26 |
| # | time | 5 | 1 | 0 | 0.833333333 | 1 | # | 20 | Baseline | ancom | 26 |
| # | interaction | 0 | 0 | 4 | 0 | 0 | # | 20 | Baseline | ancom | 26 |
| # | disease | 7 | 17 | 0 | 0.291666667 | 1 | # | 20 | Baseline | aldex2 | 26 |
| # | sex | 4 | 2 | 0 | 0.666666667 | 1 | # | 20 | Baseline | aldex2 | 26 |
| # | time | 5 | 15 | 0 | 0.25 | 1 | # | 20 | Baseline | aldex2 | 26 |
| # | interaction | 0 | 0 | 4 | 0 | 0 | # | 20 | Baseline | aldex2 | 26 |
| # | disease | 7 | 0 | 0 | 1 | 1 | # | 20 | Small | paralle | 26 |
| # | sex | 4 | 0 | 0 | 1 | 1 | # | 20 | Small | paralle | 26 |
| # | time | 5 | 0 | 0 | 1 | 1 | # | 20 | Small | paralle | 26 |
| # | interaction | 3 | 0 | 1 | 1 | 0.75 | # | 20 | Small | paralle | 26 |
| # | disease | 7 | 0 | 0 | 1 | 1 | # | 20 | Small | ancom | 26 |
| # | sex | 4 | 0 | 0 | 1 | 1 | # | 20 | Small | ancom | 26 |
| # | time | 5 | 0 | 0 | 1 | 1 | # | 20 | Small | ancom | 26 |
| # | interaction | 0 | 0 | 4 | 0 | 0 | # | 20 | Small | ancom | 26 |
| # | disease | 7 | 0 | 0 | 1 | 1 | # | 20 | Small | aldex2 | 26 |
| # | sex | 4 | 1 | 0 | 0.8 | 1 | # | 20 | Small | aldex2 | 26 |
| # | time | 5 | 1 | 0 | 0.833333333 | 1 | # | 20 | Small | aldex2 | 26 |
| # | interaction | 0 | 0 | 4 | 0 | 0 | # | 20 | Small | aldex2 | 26 |
| # | disease | 7 | 11 | 0 | 0.388888889 | 1 | # | 20 | Baseline | paralle | 26 |
| # | sex | 4 | 7 | 0 | 0.363636364 | 1 | # | 20 | Baseline | paralle | 26 |
| # | time | 5 | 11 | 0 | 0.3125 | 1 | # | 20 | Baseline | paralle | 26 |
| # | interaction | 4 | 9 | 0 | 0.307692308 | 1 | # | 20 | Baseline | paralle | 26 |
| # | disease | 7 | 3 | 0 | 0.7 | 1 | # | 20 | Baseline | ancom | 26 |
| # | sex | 4 | 0 | 0 | 1 | 1 | # | 20 | Baseline | ancom | 26 |
| # | time | 5 | 3 | 0 | 0.625 | 1 | # | 20 | Baseline | ancom | 26 |
| # | interaction | 0 | 0 | 4 | 0 | 0 | # | 20 | Baseline | ancom | 26 |
| # | disease | 7 | 34 | 0 | 0.170731707 | 1 | # | 20 | Baseline | aldex2 | 26 |
| # | sex | 4 | 7 | 0 | 0.363636364 | 1 | # | 20 | Baseline | aldex2 | 26 |
| # | time | 5 | 23 | 0 | 0.178571429 | 1 | # | 20 | Baseline | aldex2 | 26 |
| # | interaction | 0 | 0 | 4 | 0 | 0 | # | 20 | Baseline | aldex2 | 26 |
| # | disease | 7 | 0 | 0 | 1 | 1 | # | 20 | Small | paralle | 26 |
| # | sex | 4 | 0 | 0 | 1 | 1 | # | 20 | Small | paralle | 26 |
| # | time | 5 | 0 | 0 | 1 | 1 | # | 20 | Small | paralle | 26 |
| # | interaction | 4 | 0 | 0 | 1 | 1 | # | 20 | Small | paralle | 26 |
| # | disease | 7 | 3 | 0 | 0.7 | 1 | # | 20 | Small | ancom | 26 |
| # | sex | 4 | 0 | 0 | 1 | 1 | # | 20 | Small | ancom | 26 |
| # | time | 5 | 3 | 0 | 0.625 | 1 | # | 20 | Small | ancom | 26 |
| # | interaction | 0 | 0 | 4 | 0 | 0 | # | 20 | Small | ancom | 26 |
| # | disease | 7 | 4 | 0 | 0.636363636 | 1 | # | 20 | Small | aldex2 | 26 |
| # | sex | 4 | 0 | 0 | 1 | 1 | # | 20 | Small | aldex2 | 26 |
| # | time | 5 | 5 | 0 | 0.5 | 1 | # | 20 | Small | aldex2 | 26 |
| # | interaction | 0 | 0 | 4 | 0 | 0 | # | 20 | Small | aldex2 | 26 |
| # | disease | 7 | 14 | 0 | 0.333333333 | 1 | # | 20 | Baseline | paralle | 26 |
| # | sex | 4 | 8 | 0 | 0.333333333 | 1 | # | 20 | Baseline | paralle | 26 |
| # | time | 5 | 6 | 0 | 0.454545455 | 1 | # | 20 | Baseline | paralle | 26 |
| # | interaction | 4 | 3 | 0 | 0.571428571 | 1 | # | 20 | Baseline | paralle | 26 |
| # | disease | 7 | 4 | 0 | 0.636363636 | 1 | # | 20 | Baseline | ancom | 26 |
| # | sex | 4 | 0 | 0 | 1 | 1 | # | 20 | Baseline | ancom | 26 |
| # | time | 5 | 4 | 0 | 0.555555556 | 1 | # | 20 | Baseline | ancom | 26 |
| # | interaction | 0 | 0 | 4 | 0 | 0 | # | 20 | Baseline | ancom | 26 |
| # | disease | 7 | 44 | 0 | 0.137254902 | 1 | # | 20 | Baseline | aldex2 | 26 |
| # | sex | 4 | 10 | 0 | 0.285714286 | 1 | # | 20 | Baseline | aldex2 | 26 |
| # | time | 5 | 33 | 0 | 0.131578947 | 1 | # | 20 | Baseline | aldex2 | 26 |
| # | interaction | 0 | 0 | 4 | 0 | 0 | # | 20 | Baseline | aldex2 | 26 |
| # | disease | 7 | 0 | 0 | 1 | 1 | # | 20 | Small | paralle | 26 |
| # | sex | 4 | 0 | 0 | 1 | 1 | # | 20 | Small | paralle | 26 |
| # | time | 5 | 0 | 0 | 1 | 1 | # | 20 | Small | paralle | 26 |
| # | interaction | 4 | 0 | 0 | 1 | 1 | # | 20 | Small | paralle | 26 |
| # | disease | 7 | 4 | 0 | 0.636363636 | 1 | # | 20 | Small | ancom | 26 |
| # | sex | 4 | 0 | 0 | 1 | 1 | # | 20 | Small | ancom | 26 |
| # | time | 5 | 4 | 0 | 0.555555556 | 1 | # | 20 | Small | ancom | 26 |
| # | interaction | 0 | 0 | 4 | 0 | 0 | # | 20 | Small | ancom | 26 |
| # | disease | 7 | 5 | 0 | 0.583333333 | 1 | # | 20 | Small | aldex2 | 26 |
| # | sex | 4 | 1 | 0 | 0.8 | 1 | # | 20 | Small | aldex2 | 26 |
| # | time | 5 | 5 | 0 | 0.5 | 1 | # | 20 | Small | aldex2 | 26 |
| # | interaction | 0 | 0 | 4 | 0 | 0 | # | 20 | Small | aldex2 | 26 |
| # | disease | 7 | 10 | 0 | 0.411764706 | 1 | # | 20 | Baseline | paralle | 26 |
| # | sex | 4 | 4 | 0 | 0.5 | 1 | # | 20 | Baseline | paralle | 26 |
| # | time | 5 | 2 | 0 | 0.714285714 | 1 | # | 20 | Baseline | paralle | 26 |
| # | interaction | 4 | 4 | 0 | 0.5 | 1 | # | 20 | Baseline | paralle | 26 |
| # | disease | 7 | 4 | 0 | 0.636363636 | 1 | # | 20 | Baseline | ancom | 26 |
| # | sex | 4 | 0 | 0 | 1 | 1 | # | 20 | Baseline | ancom | 26 |
| # | time | 5 | 4 | 0 | 0.555555556 | 1 | # | 20 | Baseline | ancom | 26 |
| # | interaction | 0 | 0 | 4 | 0 | 0 | # | 20 | Baseline | ancom | 26 |
| # | disease | 7 | 57 | 0 | 0.109375 | 1 | # | 20 | Baseline | aldex2 | 26 |
| # | sex | 4 | 14 | 0 | 0.222222222 | 1 | # | 20 | Baseline | aldex2 | 26 |
| # | time | 5 | 38 | 0 | 0.11627907 | 1 | # | 20 | Baseline | aldex2 | 26 |
| # | interaction | 0 | 0 | 4 | 0 | 0 | # | 20 | Baseline | aldex2 | 26 |
| # | disease | 7 | 0 | 0 | 1 | 1 | # | 20 | Small | paralle | 26 |
| # | sex | 4 | 0 | 0 | 1 | 1 | # | 20 | Small | paralle | 26 |
| # | time | 5 | 0 | 0 | 1 | 1 | # | 20 | Small | paralle | 26 |
| # | interaction | 4 | 0 | 0 | 1 | 1 | # | 20 | Small | paralle | 26 |
| # | disease | 7 | 4 | 0 | 0.636363636 | 1 | # | 20 | Small | ancom | 26 |
| # | sex | 4 | 0 | 0 | 1 | 1 | # | 20 | Small | ancom | 26 |
| # | time | 5 | 4 | 0 | 0.555555556 | 1 | # | 20 | Small | ancom | 26 |
| # | interaction | 0 | 0 | 4 | 0 | 0 | # | 20 | Small | ancom | 26 |
| # | disease | 7 | 9 | 0 | 0.4375 | 1 | # | 20 | Small | aldex2 | 26 |
| # | sex | 4 | 0 | 0 | 1 | 1 | # | 20 | Small | aldex2 | 26 |
| # | time | 5 | 6 | 0 | 0.454545455 | 1 | # | 20 | Small | aldex2 | 26 |
| # | interaction | 0 | 0 | 4 | 0 | 0 | # | 20 | Small | aldex2 | 26 |
| # | disease | 4 | 12 | 0 | 0.25 | 1 | # | 10 | Baseline | paralle | 27 |
| # | sex | 2 | 10 | 0 | 0.166666667 | 1 | # | 10 | Baseline | paralle | 27 |
| # | time | 2 | 11 | 0 | 0.153846154 | 1 | # | 10 | Baseline | paralle | 27 |
| # | interaction | 2 | 10 | 0 | 0.166666667 | 1 | # | 10 | Baseline | paralle | 27 |
| # | disease | 4 | 0 | 0 | 1 | 1 | # | 10 | Baseline | ancom | 27 |
| # | sex | 2 | 0 | 0 | 1 | 1 | # | 10 | Baseline | ancom | 27 |
| # | time | 2 | 0 | 0 | 1 | 1 | # | 10 | Baseline | ancom | 27 |
| # | interaction | 0 | 0 | 2 | 0 | 0 | # | 10 | Baseline | ancom | 27 |
| # | disease | 4 | 10 | 0 | 0.285714286 | 1 | # | 10 | Baseline | aldex2 | 27 |
| # | sex | 2 | 3 | 0 | 0.4 | 1 | # | 10 | Baseline | aldex2 | 27 |
| # | time | 2 | 7 | 0 | 0.222222222 | 1 | # | 10 | Baseline | aldex2 | 27 |
| # | interaction | 0 | 0 | 2 | 0 | 0 | # | 10 | Baseline | aldex2 | 27 |
| # | disease | 4 | 0 | 0 | 1 | 1 | # | 10 | Small | paralle | 27 |
| # | sex | 2 | 0 | 0 | 1 | 1 | # | 10 | Small | paralle | 27 |
| # | time | 2 | 0 | 0 | 1 | 1 | # | 10 | Small | paralle | 27 |
| # | interaction | 1 | 1 | 1 | 0.5 | 0.5 | # | 10 | Small | paralle | 27 |
| # | disease | 4 | 0 | 0 | 1 | 1 | # | 10 | Small | ancom | 27 |
| # | sex | 2 | 0 | 0 | 1 | 1 | # | 10 | Small | ancom | 27 |
| # | time | 2 | 0 | 0 | 1 | 1 | # | 10 | Small | ancom | 27 |
| # | interaction | 0 | 0 | 2 | 0 | 0 | # | 10 | Small | ancom | 27 |
| # | disease | 4 | 0 | 0 | 1 | 1 | # | 10 | Small | aldex2 | 27 |
| # | sex | 2 | 0 | 0 | 1 | 1 | # | 10 | Small | aldex2 | 27 |
| # | time | 2 | 1 | 0 | 0.666666667 | 1 | # | 10 | Small | aldex2 | 27 |
| # | interaction | 0 | 0 | 2 | 0 | 0 | # | 10 | Small | aldex2 | 27 |
| # | disease | 4 | 7 | 0 | 0.363636364 | 1 | # | 10 | Baseline | paralle | 27 |
| # | sex | 2 | 6 | 0 | 0.25 | 1 | # | 10 | Baseline | paralle | 27 |
| # | time | 2 | 3 | 0 | 0.4 | 1 | # | 10 | Baseline | paralle | 27 |
| # | interaction | 2 | 9 | 0 | 0.181818182 | 1 | # | 10 | Baseline | paralle | 27 |
| # | disease | 4 | 2 | 0 | 0.666666667 | 1 | # | 10 | Baseline | ancom | 27 |
| # | sex | 2 | 0 | 0 | 1 | 1 | # | 10 | Baseline | ancom | 27 |
| # | time | 2 | 2 | 0 | 0.5 | 1 | # | 10 | Baseline | ancom | 27 |
| # | interaction | 0 | 0 | 2 | 0 | 0 | # | 10 | Baseline | ancom | 27 |
| # | disease | 4 | 15 | 0 | 0.210526316 | 1 | # | 10 | Baseline | aldex2 | 27 |
| # | sex | 2 | 6 | 0 | 0.25 | 1 | # | 10 | Baseline | aldex2 | 27 |
| # | time | 2 | 9 | 0 | 0.181818182 | 1 | # | 10 | Baseline | aldex2 | 27 |
| # | interaction | 0 | 0 | 2 | 0 | 0 | # | 10 | Baseline | aldex2 | 27 |
| # | disease | 4 | 0 | 0 | 1 | 1 | # | 10 | Small | paralle | 27 |
| # | sex | 2 | 0 | 0 | 1 | 1 | # | 10 | Small | paralle | 27 |
| # | time | 2 | 0 | 0 | 1 | 1 | # | 10 | Small | paralle | 27 |
| # | interaction | 2 | 0 | 0 | 1 | 1 | # | 10 | Small | paralle | 27 |
| # | disease | 4 | 2 | 0 | 0.666666667 | 1 | # | 10 | Small | ancom | 27 |
| # | sex | 2 | 0 | 0 | 1 | 1 | # | 10 | Small | ancom | 27 |
| # | time | 2 | 2 | 0 | 0.5 | 1 | # | 10 | Small | ancom | 27 |
| # | interaction | 0 | 0 | 2 | 0 | 0 | # | 10 | Small | ancom | 27 |
| # | disease | 4 | 2 | 0 | 0.666666667 | 1 | # | 10 | Small | aldex2 | 27 |
| # | sex | 2 | 0 | 0 | 1 | 1 | # | 10 | Small | aldex2 | 27 |
| # | time | 2 | 3 | 0 | 0.4 | 1 | # | 10 | Small | aldex2 | 27 |
| # | interaction | 0 | 0 | 2 | 0 | 0 | # | 10 | Small | aldex2 | 27 |
| # | disease | 4 | 9 | 0 | 0.307692308 | 1 | # | 10 | Baseline | paralle | 27 |
| # | sex | 2 | 7 | 0 | 0.222222222 | 1 | # | 10 | Baseline | paralle | 27 |
| # | time | 2 | 7 | 0 | 0.222222222 | 1 | # | 10 | Baseline | paralle | 27 |
| # | interaction | 2 | 5 | 0 | 0.285714286 | 1 | # | 10 | Baseline | paralle | 27 |
| # | disease | 4 | 2 | 0 | 0.666666667 | 1 | # | 10 | Baseline | ancom | 27 |
| # | sex | 2 | 0 | 0 | 1 | 1 | # | 10 | Baseline | ancom | 27 |
| # | time | 2 | 2 | 0 | 0.5 | 1 | # | 10 | Baseline | ancom | 27 |
| # | interaction | 0 | 0 | 2 | 0 | 0 | # | 10 | Baseline | ancom | 27 |
| # | disease | 4 | 28 | 0 | 0.125 | 1 | # | 10 | Baseline | aldex2 | 27 |
| # | sex | 2 | 10 | 0 | 0.166666667 | 1 | # | 10 | Baseline | aldex2 | 27 |
| # | time | 2 | 11 | 0 | 0.153846154 | 1 | # | 10 | Baseline | aldex2 | 27 |
| # | interaction | 0 | 0 | 2 | 0 | 0 | # | 10 | Baseline | aldex2 | 27 |

| | | | | | | | | | | | |
|---|---|---|---|---|---|---|---|---|---|---|---|
| # | disease | 4 | 0 | 0 | 1 | 1 | # | 10 | Small | paralle | 27 |
| # | sex | 2 | 0 | 0 | 1 | 1 | # | 10 | Small | paralle | 27 |
| # | time | 2 | 0 | 0 | 1 | 1 | # | 10 | Small | paralle | 27 |
| # | interaction | 2 | 0 | 0 | 1 | 1 | # | 10 | Small | paralle | 27 |
| # | disease | 4 | 2 | 0 | 0.666666667 | 1 | # | 10 | Small | ancom | 27 |
| # | sex | 2 | 0 | 0 | 1 | 1 | # | 10 | Small | ancom | 27 |
| # | time | 2 | 2 | 0 | 0.5 | 1 | # | 10 | Small | ancom | 27 |
| # | interaction | 0 | 0 | 2 | 0 | 0 | # | 10 | Small | ancom | 27 |
| # | disease | 4 | 3 | 0 | 0.571428571 | 1 | # | 10 | Small | aldex2 | 27 |
| # | sex | 2 | 0 | 0 | 1 | 1 | # | 10 | Small | aldex2 | 27 |
| # | time | 2 | 4 | 0 | 0.333333333 | 1 | # | 10 | Small | aldex2 | 27 |
| # | interaction | 0 | 0 | 2 | 0 | 0 | # | 10 | Small | aldex2 | 27 |
| # | disease | 4 | 4 | 0 | 0.5 | 1 | # | 10 | Baseline | paralle | 27 |
| # | sex | 2 | 4 | 0 | 0.333333333 | 1 | # | 10 | Baseline | paralle | 27 |
| # | time | 2 | 3 | 0 | 0.4 | 1 | # | 10 | Baseline | paralle | 27 |
| # | interaction | 2 | 0 | 0 | 1 | 1 | # | 10 | Baseline | paralle | 27 |
| # | disease | 4 | 3 | 0 | 0.571428571 | 1 | # | 10 | Baseline | ancom | 27 |
| # | sex | 2 | 0 | 0 | 1 | 1 | # | 10 | Baseline | ancom | 27 |
| # | time | 2 | 2 | 0 | 0.5 | 1 | # | 10 | Baseline | ancom | 27 |
| # | interaction | 0 | 0 | 2 | 0 | 0 | # | 10 | Baseline | ancom | 27 |
| # | disease | 4 | 28 | 0 | 0.125 | 1 | # | 10 | Baseline | aldex2 | 27 |
| # | sex | 2 | 8 | 0 | 0.2 | 1 | # | 10 | Baseline | aldex2 | 27 |
| # | time | 2 | 10 | 0 | 0.166666667 | 1 | # | 10 | Baseline | aldex2 | 27 |
| # | interaction | 0 | 0 | 2 | 0 | 0 | # | 10 | Baseline | aldex2 | 27 |
| # | disease | 4 | 0 | 0 | 1 | 1 | # | 10 | Small | paralle | 27 |
| # | sex | 2 | 0 | 0 | 1 | 1 | # | 10 | Small | paralle | 27 |
| # | time | 2 | 0 | 0 | 1 | 1 | # | 10 | Small | paralle | 27 |
| # | interaction | 2 | 0 | 0 | 1 | 1 | # | 10 | Small | paralle | 27 |
| # | disease | 4 | 3 | 0 | 0.571428571 | 1 | # | 10 | Small | ancom | 27 |
| # | sex | 2 | 0 | 0 | 1 | 1 | # | 10 | Small | ancom | 27 |
| # | time | 2 | 2 | 0 | 0.5 | 1 | # | 10 | Small | ancom | 27 |
| # | interaction | 0 | 0 | 2 | 0 | 0 | # | 10 | Small | ancom | 27 |
| # | disease | 4 | 3 | 0 | 0.571428571 | 1 | # | 10 | Small | aldex2 | 27 |
| # | sex | 2 | 0 | 0 | 1 | 1 | # | 10 | Small | aldex2 | 27 |
| # | time | 2 | 2 | 0 | 0.5 | 1 | # | 10 | Small | aldex2 | 27 |
| # | interaction | 0 | 0 | 2 | 0 | 0 | # | 10 | Small | aldex2 | 27 |
| # | disease | 7 | 10 | 0 | 0.411764706 | 1 | # | 20 | Baseline | paralle | 27 |
| # | sex | 4 | 11 | 0 | 0.266666667 | 1 | # | 20 | Baseline | paralle | 27 |
| # | time | 5 | 12 | 0 | 0.294117647 | 1 | # | 20 | Baseline | paralle | 27 |
| # | interaction | 4 | 5 | 0 | 0.444444444 | 1 | # | 20 | Baseline | paralle | 27 |
| # | disease | 7 | 2 | 0 | 0.777777778 | 1 | # | 20 | Baseline | ancom | 27 |
| # | sex | 4 | 0 | 0 | 1 | 1 | # | 20 | Baseline | ancom | 27 |
| # | time | 5 | 1 | 0 | 0.833333333 | 1 | # | 20 | Baseline | ancom | 27 |
| # | interaction | 0 | 0 | 4 | 0 | 0 | # | 20 | Baseline | ancom | 27 |
| # | disease | 7 | 16 | 0 | 0.304347826 | 1 | # | 20 | Baseline | aldex2 | 27 |
| # | sex | 4 | 12 | 0 | 0.25 | 1 | # | 20 | Baseline | aldex2 | 27 |
| # | time | 5 | 20 | 0 | 0.2 | 1 | # | 20 | Baseline | aldex2 | 27 |
| # | interaction | 0 | 0 | 4 | 0 | 0 | # | 20 | Baseline | aldex2 | 27 |
| # | disease | 7 | 1 | 0 | 0.875 | 1 | # | 20 | Small | paralle | 27 |
| # | sex | 4 | 2 | 0 | 0.666666667 | 1 | # | 20 | Small | paralle | 27 |
| # | time | 5 | 0 | 0 | 1 | 1 | # | 20 | Small | paralle | 27 |
| # | interaction | 2 | 2 | 2 | 0.5 | 0.5 | # | 20 | Small | paralle | 27 |
| # | disease | 7 | 1 | 0 | 0.875 | 1 | # | 20 | Small | ancom | 27 |
| # | sex | 4 | 0 | 0 | 1 | 1 | # | 20 | Small | ancom | 27 |
| # | time | 5 | 1 | 0 | 0.833333333 | 1 | # | 20 | Small | ancom | 27 |
| # | interaction | 0 | 0 | 4 | 0 | 0 | # | 20 | Small | ancom | 27 |
| # | disease | 7 | 1 | 0 | 0.875 | 1 | # | 20 | Small | aldex2 | 27 |
| # | sex | 4 | 0 | 0 | 1 | 1 | # | 20 | Small | aldex2 | 27 |
| # | time | 5 | 1 | 0 | 0.833333333 | 1 | # | 20 | Small | aldex2 | 27 |
| # | interaction | 0 | 0 | 4 | 0 | 0 | # | 20 | Small | aldex2 | 27 |
| # | disease | 7 | 15 | 0 | 0.318181818 | 1 | # | 20 | Baseline | paralle | 27 |
| # | sex | 4 | 8 | 0 | 0.333333333 | 1 | # | 20 | Baseline | paralle | 27 |
| # | time | 5 | 7 | 0 | 0.416666667 | 1 | # | 20 | Baseline | paralle | 27 |
| # | interaction | 4 | 5 | 0 | 0.444444444 | 1 | # | 20 | Baseline | paralle | 27 |
| # | disease | 7 | 3 | 0 | 0.7 | 1 | # | 20 | Baseline | ancom | 27 |
| # | sex | 4 | 0 | 0 | 1 | 1 | # | 20 | Baseline | ancom | 27 |
| # | time | 5 | 4 | 0 | 0.555555556 | 1 | # | 20 | Baseline | ancom | 27 |
| # | interaction | 0 | 0 | 4 | 0 | 0 | # | 20 | Baseline | ancom | 27 |
| # | disease | 7 | 33 | 0 | 0.175 | 1 | # | 20 | Baseline | aldex2 | 27 |
| # | sex | 4 | 8 | 0 | 0.333333333 | 1 | # | 20 | Baseline | aldex2 | 27 |
| # | time | 5 | 36 | 0 | 0.12195122 | 1 | # | 20 | Baseline | aldex2 | 27 |
| # | interaction | 0 | 0 | 4 | 0 | 0 | # | 20 | Baseline | aldex2 | 27 |
| # | disease | 7 | 0 | 0 | 1 | 1 | # | 20 | Small | paralle | 27 |
| # | sex | 4 | 0 | 0 | 1 | 1 | # | 20 | Small | paralle | 27 |
| # | time | 5 | 0 | 0 | 1 | 1 | # | 20 | Small | paralle | 27 |
| # | interaction | 3 | 0 | 1 | 1 | 0.75 | # | 20 | Small | paralle | 27 |
| # | disease | 7 | 3 | 0 | 0.7 | 1 | # | 20 | Small | ancom | 27 |
| # | sex | 4 | 0 | 0 | 1 | 1 | # | 20 | Small | ancom | 27 |
| # | time | 5 | 4 | 0 | 0.555555556 | 1 | # | 20 | Small | ancom | 27 |
| # | interaction | 0 | 0 | 4 | 0 | 0 | # | 20 | Small | ancom | 27 |
| # | disease | 7 | 2 | 0 | 0.777777778 | 1 | # | 20 | Small | aldex2 | 27 |
| # | sex | 4 | 0 | 0 | 1 | 1 | # | 20 | Small | aldex2 | 27 |
| # | time | 5 | 3 | 0 | 0.625 | 1 | # | 20 | Small | aldex2 | 27 |
| # | interaction | 0 | 0 | 4 | 0 | 0 | # | 20 | Small | aldex2 | 27 |
| # | disease | 7 | 6 | 0 | 0.538461538 | 1 | # | 20 | Baseline | paralle | 27 |
| # | sex | 4 | 8 | 0 | 0.333333333 | 1 | # | 20 | Baseline | paralle | 27 |
| # | time | 5 | 10 | 0 | 0.333333333 | 1 | # | 20 | Baseline | paralle | 27 |
| # | interaction | 4 | 5 | 0 | 0.444444444 | 1 | # | 20 | Baseline | paralle | 27 |
| # | disease | 7 | 4 | 0 | 0.636363636 | 1 | # | 20 | Baseline | ancom | 27 |
| # | sex | 4 | 0 | 0 | 1 | 1 | # | 20 | Baseline | ancom | 27 |
| # | time | 5 | 4 | 0 | 0.555555556 | 1 | # | 20 | Baseline | ancom | 27 |
| # | interaction | 0 | 0 | 4 | 0 | 0 | # | 20 | Baseline | ancom | 27 |
| # | disease | 7 | 47 | 0 | 0.12962963 | 1 | # | 20 | Baseline | aldex2 | 27 |
| # | sex | 4 | 12 | 0 | 0.25 | 1 | # | 20 | Baseline | aldex2 | 27 |
| # | time | 5 | 37 | 0 | 0.119047619 | 1 | # | 20 | Baseline | aldex2 | 27 |
| # | interaction | 0 | 0 | 4 | 0 | 0 | # | 20 | Baseline | aldex2 | 27 |
| # | disease | 7 | 0 | 0 | 1 | 1 | # | 20 | Small | paralle | 27 |
| # | sex | 4 | 0 | 0 | 1 | 1 | # | 20 | Small | paralle | 27 |
| # | time | 5 | 0 | 0 | 1 | 1 | # | 20 | Small | paralle | 27 |
| # | interaction | 4 | 0 | 0 | 1 | 1 | # | 20 | Small | paralle | 27 |
| # | disease | 7 | 4 | 0 | 0.636363636 | 1 | # | 20 | Small | ancom | 27 |
| # | sex | 4 | 0 | 0 | 1 | 1 | # | 20 | Small | ancom | 27 |
| # | time | 5 | 4 | 0 | 0.555555556 | 1 | # | 20 | Small | ancom | 27 |
| # | interaction | 0 | 0 | 4 | 0 | 0 | # | 20 | Small | ancom | 27 |
| # | disease | 7 | 7 | 0 | 0.5 | 1 | # | 20 | Small | aldex2 | 27 |
| # | sex | 4 | 0 | 0 | 1 | 1 | # | 20 | Small | aldex2 | 27 |
| # | time | 5 | 5 | 0 | 0.5 | 1 | # | 20 | Small | aldex2 | 27 |
| # | interaction | 0 | 0 | 4 | 0 | 0 | # | 20 | Small | aldex2 | 27 |
| # | disease | 7 | 12 | 0 | 0.368421053 | 1 | # | 20 | Baseline | paralle | 27 |
| # | sex | 4 | 8 | 0 | 0.333333333 | 1 | # | 20 | Baseline | paralle | 27 |
| # | time | 5 | 11 | 0 | 0.3125 | 1 | # | 20 | Baseline | paralle | 27 |
| # | interaction | 4 | 3 | 0 | 0.571428571 | 1 | # | 20 | Baseline | paralle | 27 |
| # | disease | 7 | 4 | 0 | 0.636363636 | 1 | # | 20 | Baseline | ancom | 27 |
| # | sex | 4 | 0 | 0 | 1 | 1 | # | 20 | Baseline | ancom | 27 |
| # | time | 5 | 4 | 0 | 0.555555556 | 1 | # | 20 | Baseline | ancom | 27 |
| # | interaction | 0 | 0 | 4 | 0 | 0 | # | 20 | Baseline | ancom | 27 |
| # | disease | 7 | 68 | 0 | 0.093333333 | 1 | # | 20 | Baseline | aldex2 | 27 |
| # | sex | 4 | 13 | 0 | 0.235294118 | 1 | # | 20 | Baseline | aldex2 | 27 |
| # | time | 5 | 48 | 0 | 0.094339623 | 1 | # | 20 | Baseline | aldex2 | 27 |
| # | interaction | 0 | 0 | 4 | 0 | 0 | # | 20 | Baseline | aldex2 | 27 |
| # | disease | 7 | 0 | 0 | 1 | 1 | # | 20 | Small | paralle | 27 |
| # | sex | 4 | 0 | 0 | 1 | 1 | # | 20 | Small | paralle | 27 |
| # | time | 5 | 0 | 0 | 1 | 1 | # | 20 | Small | paralle | 27 |
| # | interaction | 4 | 0 | 0 | 1 | 1 | # | 20 | Small | paralle | 27 |
| # | disease | 7 | 4 | 0 | 0.636363636 | 1 | # | 20 | Small | ancom | 27 |
| # | sex | 4 | 0 | 0 | 1 | 1 | # | 20 | Small | ancom | 27 |
| # | time | 5 | 4 | 0 | 0.555555556 | 1 | # | 20 | Small | ancom | 27 |
| # | interaction | 0 | 0 | 4 | 0 | 0 | # | 20 | Small | ancom | 27 |
| # | disease | 7 | 7 | 0 | 0.5 | 1 | # | 20 | Small | aldex2 | 27 |
| # | sex | 4 | 0 | 0 | 1 | 1 | # | 20 | Small | aldex2 | 27 |
| # | time | 5 | 9 | 0 | 0.357142857 | 1 | # | 20 | Small | aldex2 | 27 |
| # | interaction | 0 | 0 | 4 | 0 | 0 | # | 20 | Small | aldex2 | 27 |
| # | disease | 4 | 12 | 0 | 0.25 | 1 | # | 10 | Baseline | paralle | 28 |
| # | sex | 2 | 10 | 0 | 0.166666667 | 1 | # | 10 | Baseline | paralle | 28 |
| # | time | 2 | 12 | 0 | 0.142857143 | 1 | # | 10 | Baseline | paralle | 28 |
| # | interaction | 2 | 15 | 0 | 0.117647059 | 1 | # | 10 | Baseline | paralle | 28 |
| # | disease | 4 | 1 | 0 | 0.8 | 1 | # | 10 | Baseline | ancom | 28 |
| # | sex | 2 | 0 | 0 | 1 | 1 | # | 10 | Baseline | ancom | 28 |
| # | time | 2 | 0 | 0 | 1 | 1 | # | 10 | Baseline | ancom | 28 |
| # | interaction | 0 | 0 | 2 | 0 | 0 | # | 10 | Baseline | ancom | 28 |
| # | disease | 4 | 16 | 0 | 0.2 | 1 | # | 10 | Baseline | aldex2 | 28 |
| # | sex | 2 | 4 | 0 | 0.333333333 | 1 | # | 10 | Baseline | aldex2 | 28 |
| # | time | 2 | 5 | 0 | 0.285714286 | 1 | # | 10 | Baseline | aldex2 | 28 |
| # | interaction | 0 | 0 | 2 | 0 | 0 | # | 10 | Baseline | aldex2 | 28 |
| # | disease | 4 | 1 | 0 | 0.8 | 1 | # | 10 | Small | paralle | 28 |
| # | sex | 2 | 0 | 0 | 1 | 1 | # | 10 | Small | paralle | 28 |
| # | time | 2 | 1 | 0 | 0.666666667 | 1 | # | 10 | Small | paralle | 28 |
| # | interaction | 1 | 1 | 1 | 0.5 | 0.5 | # | 10 | Small | paralle | 28 |
| # | disease | 4 | 1 | 0 | 0.8 | 1 | # | 10 | Small | ancom | 28 |
| # | sex | 2 | 0 | 0 | 1 | 1 | # | 10 | Small | ancom | 28 |
| # | time | 2 | 0 | 0 | 1 | 1 | # | 10 | Small | ancom | 28 |
| # | interaction | 0 | 0 | 2 | 0 | 0 | # | 10 | Small | ancom | 28 |
| # | disease | 4 | 2 | 0 | 0.666666667 | 1 | # | 10 | Small | aldex2 | 28 |
| # | sex | 2 | 0 | 0 | 1 | 1 | # | 10 | Small | aldex2 | 28 |
| # | time | 2 | 1 | 0 | 0.666666667 | 1 | # | 10 | Small | aldex2 | 28 |
| # | interaction | 0 | 0 | 2 | 0 | 0 | # | 10 | Small | aldex2 | 28 |
| # | disease | 4 | 7 | 0 | 0.363636364 | 1 | # | 10 | Baseline | paralle | 28 |
| # | sex | 2 | 9 | 0 | 0.181818182 | 1 | # | 10 | Baseline | paralle | 28 |
| # | time | 2 | 4 | 0 | 0.333333333 | 1 | # | 10 | Baseline | paralle | 28 |
| # | interaction | 2 | 5 | 0 | 0.285714286 | 1 | # | 10 | Baseline | paralle | 28 |
| # | disease | 4 | 2 | 0 | 0.666666667 | 1 | # | 10 | Baseline | ancom | 28 |
| # | sex | 2 | 0 | 0 | 1 | 1 | # | 10 | Baseline | ancom | 28 |
| # | time | 2 | 2 | 0 | 0.5 | 1 | # | 10 | Baseline | ancom | 28 |
| # | interaction | 0 | 0 | 2 | 0 | 0 | # | 10 | Baseline | ancom | 28 |
| # | disease | 4 | 16 | 0 | 0.2 | 1 | # | 10 | Baseline | aldex2 | 28 |
| # | sex | 2 | 5 | 0 | 0.285714286 | 1 | # | 10 | Baseline | aldex2 | 28 |
| # | time | 2 | 10 | 0 | 0.166666667 | 1 | # | 10 | Baseline | aldex2 | 28 |
| # | interaction | 0 | 0 | 2 | 0 | 0 | # | 10 | Baseline | aldex2 | 28 |
| # | disease | 4 | 0 | 0 | 1 | 1 | # | 10 | Small | paralle | 28 |
| # | sex | 2 | 0 | 0 | 1 | 1 | # | 10 | Small | paralle | 28 |
| # | time | 2 | 0 | 0 | 1 | 1 | # | 10 | Small | paralle | 28 |
| # | interaction | 2 | 0 | 0 | 1 | 1 | # | 10 | Small | paralle | 28 |
| # | disease | 4 | 2 | 0 | 0.666666667 | 1 | # | 10 | Small | ancom | 28 |
| # | sex | 2 | 0 | 0 | 1 | 1 | # | 10 | Small | ancom | 28 |
| # | time | 2 | 2 | 0 | 0.5 | 1 | # | 10 | Small | ancom | 28 |
| # | interaction | 0 | 0 | 2 | 0 | 0 | # | 10 | Small | ancom | 28 |
| # | disease | 4 | 3 | 0 | 0.571428571 | 1 | # | 10 | Small | aldex2 | 28 |
| # | sex | 2 | 0 | 0 | 1 | 1 | # | 10 | Small | aldex2 | 28 |
| # | time | 2 | 3 | 0 | 0.4 | 1 | # | 10 | Small | aldex2 | 28 |
| # | interaction | 0 | 0 | 2 | 0 | 0 | # | 10 | Small | aldex2 | 28 |
| # | disease | 4 | 3 | 0 | 0.571428571 | 1 | # | 10 | Baseline | paralle | 28 |
| # | sex | 2 | 4 | 0 | 0.333333333 | 1 | # | 10 | Baseline | paralle | 28 |
| # | time | 2 | 8 | 0 | 0.2 | 1 | # | 10 | Baseline | paralle | 28 |
| # | interaction | 2 | 6 | 0 | 0.25 | 1 | # | 10 | Baseline | paralle | 28 |
| # | disease | 4 | 2 | 0 | 0.666666667 | 1 | # | 10 | Baseline | ancom | 28 |
| # | sex | 2 | 0 | 0 | 1 | 1 | # | 10 | Baseline | ancom | 28 |
| # | time | 2 | 2 | 0 | 0.5 | 1 | # | 10 | Baseline | ancom | 28 |
| # | interaction | 0 | 0 | 2 | 0 | 0 | # | 10 | Baseline | ancom | 28 |
| # | disease | 4 | 16 | 0 | 0.2 | 1 | # | 10 | Baseline | aldex2 | 28 |
| # | sex | 2 | 6 | 0 | 0.25 | 1 | # | 10 | Baseline | aldex2 | 28 |
| # | time | 2 | 9 | 0 | 0.181818182 | 1 | # | 10 | Baseline | aldex2 | 28 |
| # | interaction | 0 | 0 | 2 | 0 | 0 | # | 10 | Baseline | aldex2 | 28 |
| # | disease | 4 | 0 | 0 | 1 | 1 | # | 10 | Small | paralle | 28 |
| # | sex | 2 | 0 | 0 | 1 | 1 | # | 10 | Small | paralle | 28 |
| # | time | 2 | 0 | 0 | 1 | 1 | # | 10 | Small | paralle | 28 |
| # | interaction | 2 | 0 | 0 | 1 | 1 | # | 10 | Small | paralle | 28 |
| # | disease | 4 | 2 | 0 | 0.666666667 | 1 | # | 10 | Small | ancom | 28 |
| # | sex | 2 | 0 | 0 | 1 | 1 | # | 10 | Small | ancom | 28 |
| # | time | 2 | 2 | 0 | 0.5 | 1 | # | 10 | Small | ancom | 28 |
| # | interaction | 0 | 0 | 2 | 0 | 0 | # | 10 | Small | ancom | 28 |
| # | disease | 4 | 3 | 0 | 0.571428571 | 1 | # | 10 | Small | aldex2 | 28 |
| # | sex | 2 | 0 | 0 | 1 | 1 | # | 10 | Small | aldex2 | 28 |
| # | time | 2 | 3 | 0 | 0.4 | 1 | # | 10 | Small | aldex2 | 28 |
| # | interaction | 0 | 0 | 2 | 0 | 0 | # | 10 | Small | aldex2 | 28 |
| # | disease | 4 | 3 | 0 | 0.571428571 | 1 | # | 10 | Baseline | paralle | 28 |
| # | sex | 2 | 4 | 0 | 0.333333333 | 1 | # | 10 | Baseline | paralle | 28 |
| # | time | 2 | 7 | 0 | 0.222222222 | 1 | # | 10 | Baseline | paralle | 28 |
| # | interaction | 2 | 2 | 0 | 0.5 | 1 | # | 10 | Baseline | paralle | 28 |
| # | disease | 4 | 2 | 0 | 0.666666667 | 1 | # | 10 | Baseline | ancom | 28 |
| # | sex | 2 | 0 | 0 | 1 | 1 | # | 10 | Baseline | ancom | 28 |
| # | time | 2 | 2 | 0 | 0.5 | 1 | # | 10 | Baseline | ancom | 28 |
| # | interaction | 0 | 0 | 2 | 0 | 0 | # | 10 | Baseline | ancom | 28 |
| # | disease | 4 | 30 | 0 | 0.117647059 | 1 | # | 10 | Baseline | aldex2 | 28 |
| # | sex | 2 | 8 | 0 | 0.2 | 1 | # | 10 | Baseline | aldex2 | 28 |
| # | time | 2 | 11 | 0 | 0.153846154 | 1 | # | 10 | Baseline | aldex2 | 28 |
| # | interaction | 0 | 0 | 2 | 0 | 0 | # | 10 | Baseline | aldex2 | 28 |
| # | disease | 4 | 0 | 0 | 1 | 1 | # | 10 | Small | paralle | 28 |
| # | sex | 2 | 0 | 0 | 1 | 1 | # | 10 | Small | paralle | 28 |
| # | time | 2 | 0 | 0 | 1 | 1 | # | 10 | Small | paralle | 28 |
| # | interaction | 2 | 0 | 0 | 1 | 1 | # | 10 | Small | paralle | 28 |
| # | disease | 4 | 2 | 0 | 0.666666667 | 1 | # | 10 | Small | ancom | 28 |
| # | sex | 2 | 0 | 0 | 1 | 1 | # | 10 | Small | ancom | 28 |
| # | time | 2 | 2 | 0 | 0.5 | 1 | # | 10 | Small | ancom | 28 |
| # | interaction | 0 | 0 | 2 | 0 | 0 | # | 10 | Small | ancom | 28 |
| # | disease | 4 | 3 | 0 | 0.571428571 | 1 | # | 10 | Small | aldex2 | 28 |
| # | sex | 2 | 0 | 0 | 1 | 1 | # | 10 | Small | aldex2 | 28 |
| # | time | 2 | 2 | 0 | 0.5 | 1 | # | 10 | Small | aldex2 | 28 |
| # | interaction | 0 | 0 | 2 | 0 | 0 | # | 10 | Small | aldex2 | 28 |
| # | disease | 7 | 7 | 0 | 0.5 | 1 | # | 20 | Baseline | paralle | 28 |
| # | sex | 4 | 5 | 0 | 0.444444444 | 1 | # | 20 | Baseline | paralle | 28 |
| # | time | 5 | 6 | 0 | 0.454545455 | 1 | # | 20 | Baseline | paralle | 28 |
| # | interaction | 2 | 11 | 2 | 0.153846154 | 0.5 | # | 20 | Baseline | paralle | 28 |
| # | disease | 7 | 2 | 0 | 0.777777778 | 1 | # | 20 | Baseline | ancom | 28 |
| # | sex | 4 | 0 | 0 | 1 | 1 | # | 20 | Baseline | ancom | 28 |
| # | time | 5 | 1 | 0 | 0.833333333 | 1 | # | 20 | Baseline | ancom | 28 |
| # | interaction | 0 | 0 | 4 | 0 | 0 | # | 20 | Baseline | ancom | 28 |
| # | disease | 7 | 20 | 0 | 0.259259259 | 1 | # | 20 | Baseline | aldex2 | 28 |
| # | sex | 4 | 5 | 0 | 0.444444444 | 1 | # | 20 | Baseline | aldex2 | 28 |
| # | time | 5 | 11 | 0 | 0.3125 | 1 | # | 20 | Baseline | aldex2 | 28 |
| # | interaction | 0 | 0 | 4 | 0 | 0 | # | 20 | Baseline | aldex2 | 28 |
| # | disease | 7 | 0 | 0 | 1 | 1 | # | 20 | Small | paralle | 28 |
| # | sex | 4 | 0 | 0 | 1 | 1 | # | 20 | Small | paralle | 28 |
| # | time | 5 | 0 | 0 | 1 | 1 | # | 20 | Small | paralle | 28 |
| # | interaction | 2 | 0 | 2 | 1 | 0.5 | # | 20 | Small | paralle | 28 |
| # | disease | 7 | 0 | 0 | 1 | 1 | # | 20 | Small | ancom | 28 |
| # | sex | 4 | 0 | 0 | 1 | 1 | # | 20 | Small | ancom | 28 |
| # | time | 5 | 0 | 0 | 1 | 1 | # | 20 | Small | ancom | 28 |
| # | interaction | 0 | 0 | 4 | 0 | 0 | # | 20 | Small | ancom | 28 |
| # | disease | 7 | 2 | 0 | 0.777777778 | 1 | # | 20 | Small | aldex2 | 28 |
| # | sex | 4 | 0 | 0 | 1 | 1 | # | 20 | Small | aldex2 | 28 |
| # | time | 5 | 1 | 0 | 0.833333333 | 1 | # | 20 | Small | aldex2 | 28 |
| # | interaction | 0 | 0 | 4 | 0 | 0 | # | 20 | Small | aldex2 | 28 |
| # | disease | 7 | 8 | 0 | 0.466666667 | 1 | # | 20 | Baseline | paralle | 28 |
| # | sex | 4 | 6 | 0 | 0.4 | 1 | # | 20 | Baseline | paralle | 28 |
| # | time | 5 | 9 | 0 | 0.357142857 | 1 | # | 20 | Baseline | paralle | 28 |
| # | interaction | 4 | 10 | 0 | 0.285714286 | 1 | # | 20 | Baseline | paralle | 28 |
| # | disease | 7 | 3 | 0 | 0.7 | 1 | # | 20 | Baseline | ancom | 28 |
| # | sex | 4 | 0 | 0 | 1 | 1 | # | 20 | Baseline | ancom | 28 |
| # | time | 5 | 4 | 0 | 0.555555556 | 1 | # | 20 | Baseline | ancom | 28 |
| # | interaction | 0 | 0 | 4 | 0 | 0 | # | 20 | Baseline | ancom | 28 |
| # | disease | 7 | 31 | 0 | 0.184210526 | 1 | # | 20 | Baseline | aldex2 | 28 |
| # | sex | 4 | 10 | 0 | 0.285714286 | 1 | # | 20 | Baseline | aldex2 | 28 |
| # | time | 5 | 22 | 0 | 0.185185185 | 1 | # | 20 | Baseline | aldex2 | 28 |
| # | interaction | 0 | 0 | 4 | 0 | 0 | # | 20 | Baseline | aldex2 | 28 |
| # | disease | 7 | 0 | 0 | 1 | 1 | # | 20 | Small | paralle | 28 |
| # | sex | 4 | 0 | 0 | 1 | 1 | # | 20 | Small | paralle | 28 |
| # | time | 5 | 1 | 0 | 0.833333333 | 1 | # | 20 | Small | paralle | 28 |
| # | interaction | 2 | 0 | 2 | 1 | 0.5 | # | 20 | Small | paralle | 28 |
| # | disease | 7 | 3 | 0 | 0.7 | 1 | # | 20 | Small | ancom | 28 |
| # | sex | 4 | 0 | 0 | 1 | 1 | # | 20 | Small | ancom | 28 |
| # | time | 5 | 4 | 0 | 0.555555556 | 1 | # | 20 | Small | ancom | 28 |
| # | interaction | 0 | 0 | 4 | 0 | 0 | # | 20 | Small | ancom | 28 |
| # | disease | 7 | 5 | 0 | 0.583333333 | 1 | # | 20 | Small | aldex2 | 28 |
| # | sex | 4 | 0 | 0 | 1 | 1 | # | 20 | Small | aldex2 | 28 |
| # | time | 5 | 3 | 0 | 0.625 | 1 | # | 20 | Small | aldex2 | 28 |
| # | interaction | 0 | 0 | 4 | 0 | 0 | # | 20 | Small | aldex2 | 28 |
| # | disease | 7 | 7 | 0 | 0.5 | 1 | # | 20 | Baseline | paralle | 28 |
| # | sex | 4 | 2 | 0 | 0.666666667 | 1 | # | 20 | Baseline | paralle | 28 |
| # | time | 5 | 6 | 0 | 0.454545455 | 1 | # | 20 | Baseline | paralle | 28 |
| # | interaction | 4 | 8 | 0 | 0.333333333 | 1 | # | 20 | Baseline | paralle | 28 |
| # | disease | 7 | 3 | 0 | 0.7 | 1 | # | 20 | Baseline | ancom | 28 |
| # | sex | 4 | 0 | 0 | 1 | 1 | # | 20 | Baseline | ancom | 28 |
| # | time | 5 | 4 | 0 | 0.555555556 | 1 | # | 20 | Baseline | ancom | 28 |
| # | interaction | 0 | 0 | 4 | 0 | 0 | # | 20 | Baseline | ancom | 28 |
| # | disease | 7 | 44 | 0 | 0.137254902 | 1 | # | 20 | Baseline | aldex2 | 28 |
| # | sex | 4 | 11 | 0 | 0.266666667 | 1 | # | 20 | Baseline | aldex2 | 28 |
| # | time | 5 | 36 | 0 | 0.12195122 | 1 | # | 20 | Baseline | aldex2 | 28 |
| # | interaction | 0 | 0 | 4 | 0 | 0 | # | 20 | Baseline | aldex2 | 28 |
| # | disease | 7 | 0 | 0 | 1 | 1 | # | 20 | Small | paralle | 28 |
| # | sex | 4 | 0 | 0 | 1 | 1 | # | 20 | Small | paralle | 28 |
| # | time | 5 | 1 | 0 | 0.833333333 | 1 | # | 20 | Small | paralle | 28 |
| # | interaction | 2 | 0 | 2 | 1 | 0.5 | # | 20 | Small | paralle | 28 |
| # | disease | 7 | 3 | 0 | 0.7 | 1 | # | 20 | Small | ancom | 28 |
| # | sex | 4 | 0 | 0 | 1 | 1 | # | 20 | Small | ancom | 28 |
| # | time | 5 | 4 | 0 | 0.555555556 | 1 | # | 20 | Small | ancom | 28 |
| # | interaction | 0 | 0 | 4 | 0 | 0 | # | 20 | Small | ancom | 28 |
| # | disease | 7 | 7 | 0 | 0.5 | 1 | # | 20 | Small | aldex2 | 28 |
| # | sex | 4 | 1 | 0 | 0.8 | 1 | # | 20 | Small | aldex2 | 28 |
| # | time | 5 | 4 | 0 | 0.555555556 | 1 | # | 20 | Small | aldex2 | 28 |
| # | interaction | 0 | 0 | 4 | 0 | 0 | # | 20 | Small | aldex2 | 28 |
| # | disease | 7 | 15 | 0 | 0.318181818 | 1 | # | 20 | Baseline | paralle | 28 |
| # | sex | 4 | 4 | 0 | 0.5 | 1 | # | 20 | Baseline | paralle | 28 |
| # | time | 5 | 7 | 0 | 0.416666667 | 1 | # | 20 | Baseline | paralle | 28 |
| # | interaction | 4 | 4 | 0 | 0.5 | 1 | # | 20 | Baseline | paralle | 28 |
| # | disease | 7 | 3 | 0 | 0.7 | 1 | # | 20 | Baseline | ancom | 28 |
| # | sex | 4 | 0 | 0 | 1 | 1 | # | 20 | Baseline | ancom | 28 |
| # | time | 5 | 4 | 0 | 0.555555556 | 1 | # | 20 | Baseline | ancom | 28 |
| # | interaction | 0 | 0 | 4 | 0 | 0 | # | 20 | Baseline | ancom | 28 |
| # | disease | 7 | 55 | 0 | 0.112903226 | 1 | # | 20 | Baseline | aldex2 | 28 |
| # | sex | 4 | 15 | 0 | 0.210526316 | 1 | # | 20 | Baseline | aldex2 | 28 |
| # | time | 5 | 39 | 0 | 0.113636364 | 1 | # | 20 | Baseline | aldex2 | 28 |
| # | interaction | 0 | 0 | 4 | 0 | 0 | # | 20 | Baseline | aldex2 | 28 |
| # | disease | 7 | 0 | 0 | 1 | 1 | # | 20 | Small | paralle | 28 |
| # | sex | 4 | 0 | 0 | 1 | 1 | # | 20 | Small | paralle | 28 |
| # | time | 5 | 0 | 0 | 1 | 1 | # | 20 | Small | paralle | 28 |
| # | interaction | 4 | 0 | 0 | 1 | 1 | # | 20 | Small | paralle | 28 |
| # | disease | 7 | 3 | 0 | 0.7 | 1 | # | 20 | Small | ancom | 28 |
| # | sex | 4 | 0 | 0 | 1 | 1 | # | 20 | Small | ancom | 28 |
| # | time | 5 | 4 | 0 | 0.555555556 | 1 | # | 20 | Small | ancom | 28 |
| # | interaction | 0 | 0 | 4 | 0 | 0 | # | 20 | Small | ancom | 28 |
| # | disease | 7 | 7 | 0 | 0.5 | 1 | # | 20 | Small | aldex2 | 28 |
| # | sex | 4 | 2 | 0 | 0.666666667 | 1 | # | 20 | Small | aldex2 | 28 |
| # | time | 5 | 5 | 0 | 0.5 | 1 | # | 20 | Small | aldex2 | 28 |
| # | interaction | 0 | 0 | 4 | 0 | 0 | # | 20 | Small | aldex2 | 28 |
| # | disease | 4 | 6 | 0 | 0.4 | 1 | # | 10 | Baseline | paralle | 29 |
| # | sex | 2 | 7 | 0 | 0.222222222 | 1 | # | 10 | Baseline | paralle | 29 |
| # | time | 2 | 3 | 0 | 0.4 | 1 | # | 10 | Baseline | paralle | 29 |
| # | interaction | 2 | 8 | 0 | 0.2 | 1 | # | 10 | Baseline | paralle | 29 |
| # | disease | 4 | 0 | 0 | 1 | 1 | # | 10 | Baseline | ancom | 29 |
| # | sex | 2 | 0 | 0 | 1 | 1 | # | 10 | Baseline | ancom | 29 |
| # | time | 2 | 1 | 0 | 0.666666667 | 1 | # | 10 | Baseline | ancom | 29 |
| # | interaction | 0 | 0 | 2 | 0 | 0 | # | 10 | Baseline | ancom | 29 |
| # | disease | 4 | 12 | 0 | 0.25 | 1 | # | 10 | Baseline | aldex2 | 29 |
| # | sex | 2 | 6 | 0 | 0.25 | 1 | # | 10 | Baseline | aldex2 | 29 |
| # | time | 2 | 10 | 0 | 0.166666667 | 1 | # | 10 | Baseline | aldex2 | 29 |
| # | interaction | 0 | 0 | 2 | 0 | 0 | # | 10 | Baseline | aldex2 | 29 |
| # | disease | 4 | 1 | 0 | 0.8 | 1 | # | 10 | Small | paralle | 29 |

| | | | | | | | | | | | |
|---|---|---|---|---|---|---|---|---|---|---|---|
| # | sex | 2 | 0 | 0 | 1 | 1 | # | 10 | Small | paralle | 29 |
| # | time | 2 | 0 | 0 | 1 | 1 | # | 10 | Small | paralle | 29 |
| # | interaction | 2 | 0 | 0 | 1 | 1 | # | 10 | Small | paralle | 29 |
| # | disease | 4 | 0 | 0 | 1 | 1 | # | 10 | Small | ancom | 29 |
| # | sex | 2 | 0 | 0 | 1 | 1 | # | 10 | Small | ancom | 29 |
| # | time | 2 | 1 | 0 | 0.666666667 | 1 | # | 10 | Small | ancom | 29 |
| # | interaction | 0 | 0 | 2 | 0 | 0 | # | 10 | Small | ancom | 29 |
| # | disease | 4 | 0 | 0 | 1 | 1 | # | 10 | Small | aldex2 | 29 |
| # | sex | 2 | 1 | 0 | 0.666666667 | 1 | # | 10 | Small | aldex2 | 29 |
| # | time | 2 | 1 | 0 | 0.666666667 | 1 | # | 10 | Small | aldex2 | 29 |
| # | interaction | 0 | 0 | 2 | 0 | 0 | # | 10 | Small | aldex2 | 29 |
| # | disease | 4 | 10 | 0 | 0.285714286 | 1 | # | 10 | Baseline | paralle | 29 |
| # | sex | 2 | 5 | 0 | 0.285714286 | 1 | # | 10 | Baseline | paralle | 29 |
| # | time | 2 | 7 | 0 | 0.222222222 | 1 | # | 10 | Baseline | paralle | 29 |
| # | interaction | 2 | 7 | 0 | 0.222222222 | 1 | # | 10 | Baseline | paralle | 29 |
| # | disease | 4 | 1 | 0 | 0.8 | 1 | # | 10 | Baseline | ancom | 29 |
| # | sex | 2 | 0 | 0 | 1 | 1 | # | 10 | Baseline | ancom | 29 |
| # | time | 2 | 2 | 0 | 0.5 | 1 | # | 10 | Baseline | ancom | 29 |
| # | interaction | 0 | 0 | 2 | 0 | 0 | # | 10 | Baseline | ancom | 29 |
| # | disease | 4 | 14 | 0 | 0.222222222 | 1 | # | 10 | Baseline | aldex2 | 29 |
| # | sex | 2 | 6 | 0 | 0.25 | 1 | # | 10 | Baseline | aldex2 | 29 |
| # | time | 2 | 15 | 0 | 0.117647059 | 1 | # | 10 | Baseline | aldex2 | 29 |
| # | interaction | 0 | 0 | 2 | 0 | 0 | # | 10 | Baseline | aldex2 | 29 |
| # | disease | 4 | 0 | 0 | 1 | 1 | # | 10 | Small | paralle | 29 |
| # | sex | 2 | 0 | 0 | 1 | 1 | # | 10 | Small | paralle | 29 |
| # | time | 2 | 0 | 0 | 1 | 1 | # | 10 | Small | paralle | 29 |
| # | interaction | 2 | 0 | 0 | 1 | 1 | # | 10 | Small | paralle | 29 |
| # | disease | 4 | 1 | 0 | 0.8 | 1 | # | 10 | Small | ancom | 29 |
| # | sex | 2 | 0 | 0 | 1 | 1 | # | 10 | Small | ancom | 29 |
| # | time | 2 | 2 | 0 | 0.5 | 1 | # | 10 | Small | ancom | 29 |
| # | interaction | 0 | 0 | 2 | 0 | 0 | # | 10 | Small | ancom | 29 |
| # | disease | 4 | 1 | 0 | 0.8 | 1 | # | 10 | Small | aldex2 | 29 |
| # | sex | 2 | 0 | 0 | 1 | 1 | # | 10 | Small | aldex2 | 29 |
| # | time | 2 | 3 | 0 | 0.4 | 1 | # | 10 | Small | aldex2 | 29 |
| # | interaction | 0 | 0 | 2 | 0 | 0 | # | 10 | Small | aldex2 | 29 |
| # | disease | 4 | 4 | 0 | 0.5 | 1 | # | 10 | Baseline | paralle | 29 |
| # | sex | 2 | 7 | 0 | 0.222222222 | 1 | # | 10 | Baseline | paralle | 29 |
| # | time | 2 | 1 | 0 | 0.666666667 | 1 | # | 10 | Baseline | paralle | 29 |
| # | interaction | 2 | 5 | 0 | 0.285714286 | 1 | # | 10 | Baseline | paralle | 29 |
| # | disease | 4 | 2 | 0 | 0.666666667 | 1 | # | 10 | Baseline | ancom | 29 |
| # | sex | 2 | 0 | 0 | 1 | 1 | # | 10 | Baseline | ancom | 29 |
| # | time | 2 | 2 | 0 | 0.5 | 1 | # | 10 | Baseline | ancom | 29 |
| # | interaction | 0 | 0 | 2 | 0 | 0 | # | 10 | Baseline | ancom | 29 |
| # | disease | 4 | 20 | 0 | 0.166666667 | 1 | # | 10 | Baseline | aldex2 | 29 |
| # | sex | 2 | 7 | 0 | 0.222222222 | 1 | # | 10 | Baseline | aldex2 | 29 |
| # | time | 2 | 14 | 0 | 0.125 | 1 | # | 10 | Baseline | aldex2 | 29 |
| # | interaction | 0 | 0 | 2 | 0 | 0 | # | 10 | Baseline | aldex2 | 29 |
| # | disease | 4 | 0 | 0 | 1 | 1 | # | 10 | Small | paralle | 29 |
| # | sex | 2 | 0 | 0 | 1 | 1 | # | 10 | Small | paralle | 29 |
| # | time | 2 | 0 | 0 | 1 | 1 | # | 10 | Small | paralle | 29 |
| # | interaction | 2 | 0 | 0 | 1 | 1 | # | 10 | Small | paralle | 29 |
| # | disease | 4 | 2 | 0 | 0.666666667 | 1 | # | 10 | Small | ancom | 29 |
| # | sex | 2 | 0 | 0 | 1 | 1 | # | 10 | Small | ancom | 29 |
| # | time | 2 | 2 | 0 | 0.5 | 1 | # | 10 | Small | ancom | 29 |
| # | interaction | 0 | 0 | 2 | 0 | 0 | # | 10 | Small | ancom | 29 |
| # | disease | 4 | 3 | 0 | 0.571428571 | 1 | # | 10 | Small | aldex2 | 29 |
| # | sex | 2 | 0 | 0 | 1 | 1 | # | 10 | Small | aldex2 | 29 |
| # | time | 2 | 2 | 0 | 0.5 | 1 | # | 10 | Small | aldex2 | 29 |
| # | interaction | 0 | 0 | 2 | 0 | 0 | # | 10 | Small | aldex2 | 29 |
| # | disease | 4 | 6 | 0 | 0.4 | 1 | # | 10 | Baseline | paralle | 29 |
| # | sex | 2 | 4 | 0 | 0.333333333 | 1 | # | 10 | Baseline | paralle | 29 |
| # | time | 2 | 3 | 0 | 0.4 | 1 | # | 10 | Baseline | paralle | 29 |
| # | interaction | 2 | 6 | 0 | 0.25 | 1 | # | 10 | Baseline | paralle | 29 |
| # | disease | 4 | 2 | 0 | 0.666666667 | 1 | # | 10 | Baseline | ancom | 29 |
| # | sex | 2 | 0 | 0 | 1 | 1 | # | 10 | Baseline | ancom | 29 |
| # | time | 2 | 2 | 0 | 0.5 | 1 | # | 10 | Baseline | ancom | 29 |
| # | interaction | 0 | 0 | 2 | 0 | 0 | # | 10 | Baseline | ancom | 29 |
| # | disease | 4 | 23 | 0 | 0.148148148 | 1 | # | 10 | Baseline | aldex2 | 29 |
| # | sex | 2 | 9 | 0 | 0.181818182 | 1 | # | 10 | Baseline | aldex2 | 29 |
| # | time | 2 | 19 | 0 | 0.095238095 | 1 | # | 10 | Baseline | aldex2 | 29 |
| # | interaction | 0 | 0 | 2 | 0 | 0 | # | 10 | Baseline | aldex2 | 29 |
| # | disease | 4 | 0 | 0 | 1 | 1 | # | 10 | Small | paralle | 29 |
| # | sex | 2 | 0 | 0 | 1 | 1 | # | 10 | Small | paralle | 29 |
| # | time | 2 | 0 | 0 | 1 | 1 | # | 10 | Small | paralle | 29 |
| # | interaction | 2 | 0 | 0 | 1 | 1 | # | 10 | Small | paralle | 29 |
| # | disease | 4 | 2 | 0 | 0.666666667 | 1 | # | 10 | Small | ancom | 29 |
| # | sex | 2 | 0 | 0 | 1 | 1 | # | 10 | Small | ancom | 29 |
| # | time | 2 | 2 | 0 | 0.5 | 1 | # | 10 | Small | ancom | 29 |
| # | interaction | 0 | 0 | 2 | 0 | 0 | # | 10 | Small | ancom | 29 |
| # | disease | 4 | 2 | 0 | 0.666666667 | 1 | # | 10 | Small | aldex2 | 29 |
| # | sex | 2 | 0 | 0 | 1 | 1 | # | 10 | Small | aldex2 | 29 |
| # | time | 2 | 2 | 0 | 0.5 | 1 | # | 10 | Small | aldex2 | 29 |
| # | interaction | 0 | 0 | 2 | 0 | 0 | # | 10 | Small | aldex2 | 29 |
| # | disease | 7 | 8 | 0 | 0.466666667 | 1 | # | 20 | Baseline | paralle | 29 |
| # | sex | 4 | 8 | 0 | 0.333333333 | 1 | # | 20 | Baseline | paralle | 29 |
| # | time | 5 | 10 | 0 | 0.333333333 | 1 | # | 20 | Baseline | paralle | 29 |
| # | interaction | 4 | 11 | 0 | 0.266666667 | 1 | # | 20 | Baseline | paralle | 29 |
| # | disease | 7 | 1 | 0 | 0.875 | 1 | # | 20 | Baseline | ancom | 29 |
| # | sex | 4 | 0 | 0 | 1 | 1 | # | 20 | Baseline | ancom | 29 |
| # | time | 5 | 1 | 0 | 0.833333333 | 1 | # | 20 | Baseline | ancom | 29 |
| # | interaction | 0 | 0 | 4 | 0 | 0 | # | 20 | Baseline | ancom | 29 |
| # | disease | 7 | 17 | 0 | 0.291666667 | 1 | # | 20 | Baseline | aldex2 | 29 |
| # | sex | 4 | 7 | 0 | 0.363636364 | 1 | # | 20 | Baseline | aldex2 | 29 |
| # | time | 5 | 13 | 0 | 0.277777778 | 1 | # | 20 | Baseline | aldex2 | 29 |
| # | interaction | 0 | 0 | 4 | 0 | 0 | # | 20 | Baseline | aldex2 | 29 |
| # | disease | 7 | 0 | 0 | 1 | 1 | # | 20 | Small | paralle | 29 |
| # | sex | 4 | 0 | 0 | 1 | 1 | # | 20 | Small | paralle | 29 |
| # | time | 5 | 0 | 0 | 1 | 1 | # | 20 | Small | paralle | 29 |
| # | interaction | 2 | 0 | 2 | 1 | 0.5 | # | 20 | Small | paralle | 29 |
| # | disease | 7 | 1 | 0 | 0.875 | 1 | # | 20 | Small | ancom | 29 |
| # | sex | 4 | 0 | 0 | 1 | 1 | # | 20 | Small | ancom | 29 |
| # | time | 5 | 1 | 0 | 0.833333333 | 1 | # | 20 | Small | ancom | 29 |
| # | interaction | 0 | 0 | 4 | 0 | 0 | # | 20 | Small | ancom | 29 |
| # | disease | 7 | 1 | 0 | 0.875 | 1 | # | 20 | Small | aldex2 | 29 |
| # | sex | 4 | 1 | 0 | 0.8 | 1 | # | 20 | Small | aldex2 | 29 |
| # | time | 5 | 0 | 0 | 1 | 1 | # | 20 | Small | aldex2 | 29 |
| # | interaction | 0 | 0 | 4 | 0 | 0 | # | 20 | Small | aldex2 | 29 |
| # | disease | 7 | 6 | 0 | 0.538461538 | 1 | # | 20 | Baseline | paralle | 29 |
| # | sex | 4 | 4 | 0 | 0.5 | 1 | # | 20 | Baseline | paralle | 29 |
| # | time | 5 | 7 | 0 | 0.416666667 | 1 | # | 20 | Baseline | paralle | 29 |
| # | interaction | 4 | 6 | 0 | 0.4 | 1 | # | 20 | Baseline | paralle | 29 |
| # | disease | 7 | 4 | 0 | 0.636363636 | 1 | # | 20 | Baseline | ancom | 29 |
| # | sex | 4 | 0 | 0 | 1 | 1 | # | 20 | Baseline | ancom | 29 |
| # | time | 5 | 3 | 0 | 0.625 | 1 | # | 20 | Baseline | ancom | 29 |
| # | interaction | 0 | 0 | 4 | 0 | 0 | # | 20 | Baseline | ancom | 29 |
| # | disease | 7 | 41 | 0 | 0.145833333 | 1 | # | 20 | Baseline | aldex2 | 29 |
| # | sex | 4 | 11 | 0 | 0.266666667 | 1 | # | 20 | Baseline | aldex2 | 29 |
| # | time | 5 | 18 | 0 | 0.217391304 | 1 | # | 20 | Baseline | aldex2 | 29 |
| # | interaction | 0 | 0 | 4 | 0 | 0 | # | 20 | Baseline | aldex2 | 29 |
| # | disease | 7 | 0 | 0 | 1 | 1 | # | 20 | Small | paralle | 29 |
| # | sex | 4 | 0 | 0 | 1 | 1 | # | 20 | Small | paralle | 29 |
| # | time | 5 | 0 | 0 | 1 | 1 | # | 20 | Small | paralle | 29 |
| # | interaction | 4 | 1 | 0 | 0.8 | 1 | # | 20 | Small | paralle | 29 |
| # | disease | 7 | 3 | 0 | 0.7 | 1 | # | 20 | Small | ancom | 29 |
| # | sex | 4 | 0 | 0 | 1 | 1 | # | 20 | Small | ancom | 29 |
| # | time | 5 | 2 | 0 | 0.714285714 | 1 | # | 20 | Small | ancom | 29 |
| # | interaction | 0 | 0 | 4 | 0 | 0 | # | 20 | Small | ancom | 29 |
| # | disease | 7 | 5 | 0 | 0.583333333 | 1 | # | 20 | Small | aldex2 | 29 |
| # | sex | 4 | 0 | 0 | 1 | 1 | # | 20 | Small | aldex2 | 29 |
| # | time | 5 | 2 | 0 | 0.714285714 | 1 | # | 20 | Small | aldex2 | 29 |
| # | interaction | 0 | 0 | 4 | 0 | 0 | # | 20 | Small | aldex2 | 29 |
| # | disease | 7 | 10 | 0 | 0.411764706 | 1 | # | 20 | Baseline | paralle | 29 |
| # | sex | 4 | 8 | 0 | 0.333333333 | 1 | # | 20 | Baseline | paralle | 29 |
| # | time | 5 | 9 | 0 | 0.357142857 | 1 | # | 20 | Baseline | paralle | 29 |
| # | interaction | 4 | 8 | 0 | 0.333333333 | 1 | # | 20 | Baseline | paralle | 29 |
| # | disease | 7 | 4 | 0 | 0.636363636 | 1 | # | 20 | Baseline | ancom | 29 |
| # | sex | 4 | 0 | 0 | 1 | 1 | # | 20 | Baseline | ancom | 29 |
| # | time | 5 | 4 | 0 | 0.555555556 | 1 | # | 20 | Baseline | ancom | 29 |
| # | interaction | 0 | 0 | 4 | 0 | 0 | # | 20 | Baseline | ancom | 29 |
| # | disease | 7 | 42 | 0 | 0.142857143 | 1 | # | 20 | Baseline | aldex2 | 29 |
| # | sex | 4 | 15 | 0 | 0.210526316 | 1 | # | 20 | Baseline | aldex2 | 29 |
| # | time | 5 | 27 | 0 | 0.15625 | 1 | # | 20 | Baseline | aldex2 | 29 |
| # | interaction | 0 | 0 | 4 | 0 | 0 | # | 20 | Baseline | aldex2 | 29 |
| # | disease | 7 | 0 | 0 | 1 | 1 | # | 20 | Small | paralle | 29 |
| # | sex | 4 | 0 | 0 | 1 | 1 | # | 20 | Small | paralle | 29 |
| # | time | 5 | 0 | 0 | 1 | 1 | # | 20 | Small | paralle | 29 |
| # | interaction | 4 | 0 | 0 | 1 | 1 | # | 20 | Small | paralle | 29 |
| # | disease | 7 | 4 | 0 | 0.636363636 | 1 | # | 20 | Small | ancom | 29 |
| # | sex | 4 | 0 | 0 | 1 | 1 | # | 20 | Small | ancom | 29 |
| # | time | 5 | 4 | 0 | 0.555555556 | 1 | # | 20 | Small | ancom | 29 |
| # | interaction | 0 | 0 | 4 | 0 | 0 | # | 20 | Small | ancom | 29 |
| # | disease | 7 | 7 | 0 | 0.5 | 1 | # | 20 | Small | aldex2 | 29 |
| # | sex | 4 | 1 | 0 | 0.8 | 1 | # | 20 | Small | aldex2 | 29 |
| # | time | 5 | 4 | 0 | 0.555555556 | 1 | # | 20 | Small | aldex2 | 29 |
| # | interaction | 0 | 0 | 4 | 0 | 0 | # | 20 | Small | aldex2 | 29 |
| # | disease | 7 | 14 | 0 | 0.333333333 | 1 | # | 20 | Baseline | paralle | 29 |
| # | sex | 4 | 6 | 0 | 0.4 | 1 | # | 20 | Baseline | paralle | 29 |
| # | time | 5 | 4 | 0 | 0.555555556 | 1 | # | 20 | Baseline | paralle | 29 |
| # | interaction | 4 | 8 | 0 | 0.333333333 | 1 | # | 20 | Baseline | paralle | 29 |
| # | disease | 7 | 4 | 0 | 0.636363636 | 1 | # | 20 | Baseline | ancom | 29 |
| # | sex | 4 | 0 | 0 | 1 | 1 | # | 20 | Baseline | ancom | 29 |
| # | time | 5 | 4 | 0 | 0.555555556 | 1 | # | 20 | Baseline | ancom | 29 |
| # | interaction | 0 | 0 | 4 | 0 | 0 | # | 20 | Baseline | ancom | 29 |
| # | disease | 7 | 54 | 0 | 0.114754098 | 1 | # | 20 | Baseline | aldex2 | 29 |
| # | sex | 4 | 22 | 0 | 0.153846154 | 1 | # | 20 | Baseline | aldex2 | 29 |
| # | time | 5 | 42 | 0 | 0.106382979 | 1 | # | 20 | Baseline | aldex2 | 29 |
| # | interaction | 0 | 0 | 4 | 0 | 0 | # | 20 | Baseline | aldex2 | 29 |
| # | disease | 7 | 0 | 0 | 1 | 1 | # | 20 | Small | paralle | 29 |
| # | sex | 4 | 0 | 0 | 1 | 1 | # | 20 | Small | paralle | 29 |
| # | time | 5 | 0 | 0 | 1 | 1 | # | 20 | Small | paralle | 29 |
| # | interaction | 4 | 0 | 0 | 1 | 1 | # | 20 | Small | paralle | 29 |
| # | disease | 7 | 4 | 0 | 0.636363636 | 1 | # | 20 | Small | ancom | 29 |
| # | sex | 4 | 0 | 0 | 1 | 1 | # | 20 | Small | ancom | 29 |
| # | time | 5 | 4 | 0 | 0.555555556 | 1 | # | 20 | Small | ancom | 29 |
| # | interaction | 0 | 0 | 4 | 0 | 0 | # | 20 | Small | ancom | 29 |
| # | disease | 7 | 11 | 0 | 0.388888889 | 1 | # | 20 | Small | aldex2 | 29 |
| # | sex | 4 | 2 | 0 | 0.666666667 | 1 | # | 20 | Small | aldex2 | 29 |
| # | time | 5 | 4 | 0 | 0.555555556 | 1 | # | 20 | Small | aldex2 | 29 |
| # | interaction | 0 | 0 | 4 | 0 | 0 | # | 20 | Small | aldex2 | 29 |
| # | disease | 4 | 4 | 0 | 0.5 | 1 | # | 10 | Baseline | paralle | 30 |
| # | sex | 2 | 7 | 0 | 0.222222222 | 1 | # | 10 | Baseline | paralle | 30 |
| # | time | 2 | 6 | 0 | 0.25 | 1 | # | 10 | Baseline | paralle | 30 |
| # | interaction | 2 | 6 | 0 | 0.25 | 1 | # | 10 | Baseline | paralle | 30 |
| # | disease | 4 | 0 | 0 | 1 | 1 | # | 10 | Baseline | ancom | 30 |
| # | sex | 2 | 0 | 0 | 1 | 1 | # | 10 | Baseline | ancom | 30 |
| # | time | 2 | 0 | 0 | 1 | 1 | # | 10 | Baseline | ancom | 30 |
| # | interaction | 0 | 0 | 2 | 0 | 0 | # | 10 | Baseline | ancom | 30 |
| # | disease | 4 | 7 | 0 | 0.363636364 | 1 | # | 10 | Baseline | aldex2 | 30 |
| # | sex | 2 | 3 | 0 | 0.4 | 1 | # | 10 | Baseline | aldex2 | 30 |
| # | time | 2 | 7 | 0 | 0.222222222 | 1 | # | 10 | Baseline | aldex2 | 30 |
| # | interaction | 0 | 0 | 2 | 0 | 0 | # | 10 | Baseline | aldex2 | 30 |
| # | disease | 4 | 0 | 0 | 1 | 1 | # | 10 | Small | paralle | 30 |
| # | sex | 2 | 0 | 0 | 1 | 1 | # | 10 | Small | paralle | 30 |
| # | time | 2 | 0 | 0 | 1 | 1 | # | 10 | Small | paralle | 30 |
| # | interaction | 1 | 0 | 1 | 1 | 0.5 | # | 10 | Small | paralle | 30 |
| # | disease | 4 | 0 | 0 | 1 | 1 | # | 10 | Small | ancom | 30 |
| # | sex | 2 | 0 | 0 | 1 | 1 | # | 10 | Small | ancom | 30 |
| # | time | 2 | 0 | 0 | 1 | 1 | # | 10 | Small | ancom | 30 |
| # | interaction | 0 | 0 | 2 | 0 | 0 | # | 10 | Small | ancom | 30 |
| # | disease | 4 | 0 | 0 | 1 | 1 | # | 10 | Small | aldex2 | 30 |
| # | sex | 2 | 1 | 0 | 0.666666667 | 1 | # | 10 | Small | aldex2 | 30 |
| # | time | 2 | 1 | 0 | 0.666666667 | 1 | # | 10 | Small | aldex2 | 30 |
| # | interaction | 0 | 0 | 2 | 0 | 0 | # | 10 | Small | aldex2 | 30 |
| # | disease | 4 | 4 | 0 | 0.5 | 1 | # | 10 | Baseline | paralle | 30 |
| # | sex | 2 | 4 | 0 | 0.333333333 | 1 | # | 10 | Baseline | paralle | 30 |
| # | time | 2 | 3 | 0 | 0.4 | 1 | # | 10 | Baseline | paralle | 30 |
| # | interaction | 2 | 4 | 0 | 0.333333333 | 1 | # | 10 | Baseline | paralle | 30 |
| # | disease | 4 | 1 | 0 | 0.8 | 1 | # | 10 | Baseline | ancom | 30 |
| # | sex | 2 | 0 | 0 | 1 | 1 | # | 10 | Baseline | ancom | 30 |
| # | time | 2 | 0 | 0 | 1 | 1 | # | 10 | Baseline | ancom | 30 |
| # | interaction | 0 | 0 | 2 | 0 | 0 | # | 10 | Baseline | ancom | 30 |
| # | disease | 4 | 14 | 0 | 0.222222222 | 1 | # | 10 | Baseline | aldex2 | 30 |
| # | sex | 2 | 5 | 0 | 0.285714286 | 1 | # | 10 | Baseline | aldex2 | 30 |
| # | time | 2 | 3 | 0 | 0.4 | 1 | # | 10 | Baseline | aldex2 | 30 |
| # | interaction | 0 | 0 | 2 | 0 | 0 | # | 10 | Baseline | aldex2 | 30 |
| # | disease | 4 | 0 | 0 | 1 | 1 | # | 10 | Small | paralle | 30 |
| # | sex | 2 | 0 | 0 | 1 | 1 | # | 10 | Small | paralle | 30 |
| # | time | 2 | 0 | 0 | 1 | 1 | # | 10 | Small | paralle | 30 |
| # | interaction | 2 | 0 | 0 | 1 | 1 | # | 10 | Small | paralle | 30 |
| # | disease | 4 | 1 | 0 | 0.8 | 1 | # | 10 | Small | ancom | 30 |
| # | sex | 2 | 0 | 0 | 1 | 1 | # | 10 | Small | ancom | 30 |
| # | time | 2 | 0 | 0 | 1 | 1 | # | 10 | Small | ancom | 30 |
| # | interaction | 0 | 0 | 2 | 0 | 0 | # | 10 | Small | ancom | 30 |
| # | disease | 4 | 1 | 0 | 0.8 | 1 | # | 10 | Small | aldex2 | 30 |
| # | sex | 2 | 1 | 0 | 0.666666667 | 1 | # | 10 | Small | aldex2 | 30 |
| # | time | 2 | 1 | 0 | 0.666666667 | 1 | # | 10 | Small | aldex2 | 30 |
| # | interaction | 0 | 0 | 2 | 0 | 0 | # | 10 | Small | aldex2 | 30 |
| # | disease | 4 | 6 | 0 | 0.4 | 1 | # | 10 | Baseline | paralle | 30 |
| # | sex | 2 | 5 | 0 | 0.285714286 | 1 | # | 10 | Baseline | paralle | 30 |
| # | time | 2 | 6 | 0 | 0.25 | 1 | # | 10 | Baseline | paralle | 30 |
| # | interaction | 2 | 4 | 0 | 0.333333333 | 1 | # | 10 | Baseline | paralle | 30 |
| # | disease | 4 | 1 | 0 | 0.8 | 1 | # | 10 | Baseline | ancom | 30 |
| # | sex | 2 | 0 | 0 | 1 | 1 | # | 10 | Baseline | ancom | 30 |
| # | time | 2 | 0 | 0 | 1 | 1 | # | 10 | Baseline | ancom | 30 |
| # | interaction | 0 | 0 | 2 | 0 | 0 | # | 10 | Baseline | ancom | 30 |
| # | disease | 4 | 22 | 0 | 0.153846154 | 1 | # | 10 | Baseline | aldex2 | 30 |
| # | sex | 2 | 6 | 0 | 0.25 | 1 | # | 10 | Baseline | aldex2 | 30 |
| # | time | 2 | 8 | 0 | 0.2 | 1 | # | 10 | Baseline | aldex2 | 30 |
| # | interaction | 0 | 0 | 2 | 0 | 0 | # | 10 | Baseline | aldex2 | 30 |
| # | disease | 4 | 0 | 0 | 1 | 1 | # | 10 | Small | paralle | 30 |
| # | sex | 2 | 0 | 0 | 1 | 1 | # | 10 | Small | paralle | 30 |
| # | time | 2 | 0 | 0 | 1 | 1 | # | 10 | Small | paralle | 30 |
| # | interaction | 2 | 0 | 0 | 1 | 1 | # | 10 | Small | paralle | 30 |
| # | disease | 4 | 1 | 0 | 0.8 | 1 | # | 10 | Small | ancom | 30 |
| # | sex | 2 | 0 | 0 | 1 | 1 | # | 10 | Small | ancom | 30 |
| # | time | 2 | 0 | 0 | 1 | 1 | # | 10 | Small | ancom | 30 |
| # | interaction | 0 | 0 | 2 | 0 | 0 | # | 10 | Small | ancom | 30 |
| # | disease | 4 | 1 | 0 | 0.8 | 1 | # | 10 | Small | aldex2 | 30 |
| # | sex | 2 | 1 | 0 | 0.666666667 | 1 | # | 10 | Small | aldex2 | 30 |
| # | time | 2 | 1 | 0 | 0.666666667 | 1 | # | 10 | Small | aldex2 | 30 |
| # | interaction | 0 | 0 | 2 | 0 | 0 | # | 10 | Small | aldex2 | 30 |
| # | disease | 4 | 3 | 0 | 0.571428571 | 1 | # | 10 | Baseline | paralle | 30 |
| # | sex | 2 | 2 | 0 | 0.5 | 1 | # | 10 | Baseline | paralle | 30 |
| # | time | 2 | 7 | 0 | 0.222222222 | 1 | # | 10 | Baseline | paralle | 30 |
| # | interaction | 2 | 5 | 0 | 0.285714286 | 1 | # | 10 | Baseline | paralle | 30 |
| # | disease | 4 | 2 | 0 | 0.666666667 | 1 | # | 10 | Baseline | ancom | 30 |
| # | sex | 2 | 0 | 0 | 1 | 1 | # | 10 | Baseline | ancom | 30 |
| # | time | 2 | 1 | 0 | 0.666666667 | 1 | # | 10 | Baseline | ancom | 30 |
| # | interaction | 0 | 0 | 2 | 0 | 0 | # | 10 | Baseline | ancom | 30 |
| # | disease | 4 | 26 | 0 | 0.133333333 | 1 | # | 10 | Baseline | aldex2 | 30 |
| # | sex | 2 | 3 | 0 | 0.4 | 1 | # | 10 | Baseline | aldex2 | 30 |
| # | time | 2 | 9 | 0 | 0.181818182 | 1 | # | 10 | Baseline | aldex2 | 30 |
| # | interaction | 0 | 0 | 2 | 0 | 0 | # | 10 | Baseline | aldex2 | 30 |
| # | disease | 4 | 0 | 0 | 1 | 1 | # | 10 | Small | paralle | 30 |
| # | sex | 2 | 0 | 0 | 1 | 1 | # | 10 | Small | paralle | 30 |
| # | time | 2 | 0 | 0 | 1 | 1 | # | 10 | Small | paralle | 30 |
| # | interaction | 2 | 0 | 0 | 1 | 1 | # | 10 | Small | paralle | 30 |
| # | disease | 4 | 2 | 0 | 0.666666667 | 1 | # | 10 | Small | ancom | 30 |
| # | sex | 2 | 0 | 0 | 1 | 1 | # | 10 | Small | ancom | 30 |
| # | time | 2 | 1 | 0 | 0.666666667 | 1 | # | 10 | Small | ancom | 30 |
| # | interaction | 0 | 0 | 2 | 0 | 0 | # | 10 | Small | ancom | 30 |
| # | disease | 4 | 1 | 0 | 0.8 | 1 | # | 10 | Small | aldex2 | 30 |
| # | sex | 2 | 2 | 0 | 0.5 | 1 | # | 10 | Small | aldex2 | 30 |
| # | time | 2 | 2 | 0 | 0.5 | 1 | # | 10 | Small | aldex2 | 30 |
| # | interaction | 0 | 0 | 2 | 0 | 0 | # | 10 | Small | aldex2 | 30 |
| # | disease | 7 | 8 | 0 | 0.466666667 | 1 | # | 20 | Baseline | paralle | 30 |
| # | sex | 4 | 6 | 0 | 0.4 | 1 | # | 20 | Baseline | paralle | 30 |
| # | time | 5 | 12 | 0 | 0.294117647 | 1 | # | 20 | Baseline | paralle | 30 |
| # | interaction | 4 | 8 | 0 | 0.333333333 | 1 | # | 20 | Baseline | paralle | 30 |
| # | disease | 7 | 1 | 0 | 0.875 | 1 | # | 20 | Baseline | ancom | 30 |
| # | sex | 4 | 0 | 0 | 1 | 1 | # | 20 | Baseline | ancom | 30 |
| # | time | 5 | 0 | 0 | 1 | 1 | # | 20 | Baseline | ancom | 30 |
| # | interaction | 0 | 0 | 4 | 0 | 0 | # | 20 | Baseline | ancom | 30 |
| # | disease | 7 | 19 | 0 | 0.269230769 | 1 | # | 20 | Baseline | aldex2 | 30 |
| # | sex | 4 | 11 | 0 | 0.266666667 | 1 | # | 20 | Baseline | aldex2 | 30 |
| # | time | 5 | 17 | 0 | 0.227272727 | 1 | # | 20 | Baseline | aldex2 | 30 |
| # | interaction | 0 | 0 | 4 | 0 | 0 | # | 20 | Baseline | aldex2 | 30 |
| # | disease | 7 | 0 | 0 | 1 | 1 | # | 20 | Small | paralle | 30 |
| # | sex | 4 | 0 | 0 | 1 | 1 | # | 20 | Small | paralle | 30 |
| # | time | 5 | 0 | 0 | 1 | 1 | # | 20 | Small | paralle | 30 |
| # | interaction | 2 | 0 | 2 | 1 | 0.5 | # | 20 | Small | paralle | 30 |
| # | disease | 7 | 1 | 0 | 0.875 | 1 | # | 20 | Small | ancom | 30 |
| # | sex | 4 | 0 | 0 | 1 | 1 | # | 20 | Small | ancom | 30 |
| # | time | 5 | 1 | 0 | 0.833333333 | 1 | # | 20 | Small | ancom | 30 |
| # | interaction | 0 | 0 | 4 | 0 | 0 | # | 20 | Small | ancom | 30 |
| # | disease | 7 | 0 | 0 | 1 | 1 | # | 20 | Small | aldex2 | 30 |
| # | sex | 4 | 0 | 0 | 1 | 1 | # | 20 | Small | aldex2 | 30 |
| # | time | 5 | 0 | 0 | 1 | 1 | # | 20 | Small | aldex2 | 30 |
| # | interaction | 0 | 0 | 4 | 0 | 0 | # | 20 | Small | aldex2 | 30 |
| # | disease | 7 | 11 | 0 | 0.388888889 | 1 | # | 20 | Baseline | paralle | 30 |
| # | sex | 4 | 6 | 0 | 0.4 | 1 | # | 20 | Baseline | paralle | 30 |
| # | time | 5 | 5 | 0 | 0.5 | 1 | # | 20 | Baseline | paralle | 30 |
| # | interaction | 4 | 8 | 0 | 0.333333333 | 1 | # | 20 | Baseline | paralle | 30 |
| # | disease | 7 | 3 | 0 | 0.7 | 1 | # | 20 | Baseline | ancom | 30 |
| # | sex | 4 | 0 | 0 | 1 | 1 | # | 20 | Baseline | ancom | 30 |
| # | time | 5 | 3 | 0 | 0.625 | 1 | # | 20 | Baseline | ancom | 30 |
| # | interaction | 0 | 0 | 4 | 0 | 0 | # | 20 | Baseline | ancom | 30 |
| # | disease | 7 | 33 | 0 | 0.175 | 1 | # | 20 | Baseline | aldex2 | 30 |
| # | sex | 4 | 9 | 0 | 0.307692308 | 1 | # | 20 | Baseline | aldex2 | 30 |
| # | time | 5 | 24 | 0 | 0.172413793 | 1 | # | 20 | Baseline | aldex2 | 30 |
| # | interaction | 0 | 0 | 4 | 0 | 0 | # | 20 | Baseline | aldex2 | 30 |
| # | disease | 7 | 0 | 0 | 1 | 1 | # | 20 | Small | paralle | 30 |
| # | sex | 4 | 0 | 0 | 1 | 1 | # | 20 | Small | paralle | 30 |
| # | time | 5 | 0 | 0 | 1 | 1 | # | 20 | Small | paralle | 30 |
| # | interaction | 3 | 0 | 1 | 1 | 0.75 | # | 20 | Small | paralle | 30 |
| # | disease | 7 | 3 | 0 | 0.7 | 1 | # | 20 | Small | ancom | 30 |
| # | sex | 4 | 0 | 0 | 1 | 1 | # | 20 | Small | ancom | 30 |
| # | time | 5 | 0 | 0 | 1 | 1 | # | 20 | Small | ancom | 30 |
| # | interaction | 0 | 0 | 4 | 0 | 0 | # | 20 | Small | ancom | 30 |
| # | disease | 7 | 4 | 0 | 0.636363636 | 1 | # | 20 | Small | aldex2 | 30 |
| # | sex | 4 | 0 | 0 | 1 | 1 | # | 20 | Small | aldex2 | 30 |
| # | time | 5 | 1 | 0 | 0.833333333 | 1 | # | 20 | Small | aldex2 | 30 |
| # | interaction | 0 | 0 | 4 | 0 | 0 | # | 20 | Small | aldex2 | 30 |
| # | disease | 7 | 10 | 0 | 0.411764706 | 1 | # | 20 | Baseline | paralle | 30 |
| # | sex | 4 | 3 | 0 | 0.571428571 | 1 | # | 20 | Baseline | paralle | 30 |
| # | time | 5 | 3 | 0 | 0.625 | 1 | # | 20 | Baseline | paralle | 30 |
| # | interaction | 4 | 6 | 0 | 0.4 | 1 | # | 20 | Baseline | paralle | 30 |
| # | disease | 7 | 4 | 0 | 0.636363636 | 1 | # | 20 | Baseline | ancom | 30 |
| # | sex | 4 | 0 | 0 | 1 | 1 | # | 20 | Baseline | ancom | 30 |
| # | time | 5 | 4 | 0 | 0.555555556 | 1 | # | 20 | Baseline | ancom | 30 |
| # | interaction | 0 | 0 | 4 | 0 | 0 | # | 20 | Baseline | ancom | 30 |
| # | disease | 7 | 65 | 0 | 0.134615385 | 1 | # | 20 | Baseline | aldex2 | 30 |
| # | sex | 4 | 11 | 0 | 0.266666667 | 1 | # | 20 | Baseline | aldex2 | 30 |
| # | time | 5 | 32 | 0 | 0.135135135 | 1 | # | 20 | Baseline | aldex2 | 30 |
| # | interaction | 0 | 0 | 4 | 0 | 0 | # | 20 | Baseline | aldex2 | 30 |
| # | disease | 7 | 1 | 0 | 0.875 | 1 | # | 20 | Small | paralle | 30 |
| # | sex | 4 | 1 | 0 | 0.8 | 1 | # | 20 | Small | paralle | 30 |

| | | | | | | | | | | | |
|---|---|---|---|---|---|---|---|---|---|---|---|
| # | time | 5 | 0 | 0 | 1 | 1 | # | 20 | Small | paralle | 30 |
| # | interaction | 4 | 0 | 0 | 1 | 1 | # | 20 | Small | paralle | 30 |
| # | disease | 7 | 3 | 0 | 0.7 | 1 | # | 20 | Small | ancom | 30 |
| # | sex | 4 | 0 | 0 | 1 | 1 | # | 20 | Small | ancom | 30 |
| # | time | 5 | 4 | 0 | 0.555555556 | 1 | # | 20 | Small | ancom | 30 |
| # | interaction | 0 | 0 | 4 | 0 | 0 | # | 20 | Small | ancom | 30 |
| # | disease | 7 | 6 | 0 | 0.538461538 | 1 | # | 20 | Small | aldex2 | 30 |
| # | sex | 4 | 0 | 0 | 1 | 1 | # | 20 | Small | aldex2 | 30 |
| # | time | 5 | 2 | 0 | 0.714285714 | 1 | # | 20 | Small | aldex2 | 30 |
| # | interaction | 0 | 0 | 4 | 0 | 0 | # | 20 | Small | aldex2 | 30 |
| # | disease | 7 | 5 | 0 | 0.583333333 | 1 | # | 20 | Baseline | paralle | 30 |
| # | sex | 4 | 6 | 0 | 0.4 | 1 | # | 20 | Baseline | paralle | 30 |
| # | time | 5 | 3 | 0 | 0.625 | 1 | # | 20 | Baseline | paralle | 30 |
| # | interaction | 4 | 10 | 0 | 0.285714286 | 1 | # | 20 | Baseline | paralle | 30 |
| # | disease | 7 | 4 | 0 | 0.636363636 | 1 | # | 20 | Baseline | ancom | 30 |
| # | sex | 4 | 0 | 0 | 1 | 1 | # | 20 | Baseline | ancom | 30 |
| # | time | 5 | 4 | 0 | 0.555555556 | 1 | # | 20 | Baseline | ancom | 30 |
| # | interaction | 0 | 0 | 4 | 0 | 0 | # | 20 | Baseline | ancom | 30 |
| # | disease | 7 | 54 | 0 | 0.114754098 | 1 | # | 20 | Baseline | aldex2 | 30 |
| # | sex | 4 | 16 | 0 | 0.2 | 1 | # | 20 | Baseline | aldex2 | 30 |
| # | time | 5 | 44 | 0 | 0.102040816 | 1 | # | 20 | Baseline | aldex2 | 30 |
| # | interaction | 0 | 0 | 4 | 0 | 0 | # | 20 | Baseline | aldex2 | 30 |
| # | disease | 7 | 1 | 0 | 0.875 | 1 | # | 20 | Small | paralle | 30 |
| # | sex | 4 | 0 | 0 | 1 | 1 | # | 20 | Small | paralle | 30 |
| # | time | 5 | 0 | 0 | 1 | 1 | # | 20 | Small | paralle | 30 |
| # | interaction | 4 | 0 | 0 | 1 | 1 | # | 20 | Small | paralle | 30 |
| # | disease | 7 | 4 | 0 | 0.636363636 | 1 | # | 20 | Small | ancom | 30 |
| # | sex | 4 | 0 | 0 | 1 | 1 | # | 20 | Small | ancom | 30 |
| # | time | 5 | 4 | 0 | 0.555555556 | 1 | # | 20 | Small | ancom | 30 |
| # | interaction | 0 | 0 | 4 | 0 | 0 | # | 20 | Small | ancom | 30 |
| # | disease | 7 | 6 | 0 | 0.538461538 | 1 | # | 20 | Small | aldex2 | 30 |
| # | sex | 4 | 0 | 0 | 1 | 1 | # | 20 | Small | aldex2 | 30 |
| # | time | 5 | 3 | 0 | 0.625 | 1 | # | 20 | Small | aldex2 | 30 |
| # | interaction | 0 | 0 | 4 | 0 | 0 | # | 20 | Small | aldex2 | 30 |
| # | disease | 4 | 6 | 0 | 0.4 | 1 | # | 10 | Baseline | paralle | 31 |
| # | sex | 2 | 9 | 0 | 0.181818182 | 1 | # | 10 | Baseline | paralle | 31 |
| # | time | 2 | 12 | 0 | 0.142857143 | 1 | # | 10 | Baseline | paralle | 31 |
| # | interaction | 2 | 6 | 0 | 0.25 | 1 | # | 10 | Baseline | paralle | 31 |
| # | disease | 4 | 0 | 0 | 1 | 1 | # | 10 | Baseline | ancom | 31 |
| # | sex | 2 | 0 | 0 | 1 | 1 | # | 10 | Baseline | ancom | 31 |
| # | time | 2 | 1 | 0 | 0.666666667 | 1 | # | 10 | Baseline | ancom | 31 |
| # | interaction | 0 | 0 | 2 | 0 | 0 | # | 10 | Baseline | ancom | 31 |
| # | disease | 4 | 13 | 0 | 0.235294118 | 1 | # | 10 | Baseline | aldex2 | 31 |
| # | sex | 2 | 10 | 0 | 0.166666667 | 1 | # | 10 | Baseline | aldex2 | 31 |
| # | time | 2 | 8 | 0 | 0.2 | 1 | # | 10 | Baseline | aldex2 | 31 |
| # | interaction | 0 | 0 | 2 | 0 | 0 | # | 10 | Baseline | aldex2 | 31 |
| # | disease | 4 | 0 | 0 | 1 | 1 | # | 10 | Small | paralle | 31 |
| # | sex | 2 | 0 | 0 | 1 | 1 | # | 10 | Small | paralle | 31 |
| # | time | 2 | 0 | 0 | 1 | 1 | # | 10 | Small | paralle | 31 |
| # | interaction | 2 | 0 | 0 | 1 | 1 | # | 10 | Small | paralle | 31 |
| # | disease | 4 | 0 | 0 | 1 | 1 | # | 10 | Small | ancom | 31 |
| # | sex | 2 | 0 | 0 | 1 | 1 | # | 10 | Small | ancom | 31 |
| # | time | 2 | 1 | 0 | 0.666666667 | 1 | # | 10 | Small | ancom | 31 |
| # | interaction | 0 | 0 | 2 | 0 | 0 | # | 10 | Small | ancom | 31 |
| # | disease | 4 | 0 | 0 | 1 | 1 | # | 10 | Small | aldex2 | 31 |
| # | sex | 2 | 0 | 0 | 1 | 1 | # | 10 | Small | aldex2 | 31 |
| # | time | 2 | 1 | 0 | 0.666666667 | 1 | # | 10 | Small | aldex2 | 31 |
| # | interaction | 0 | 0 | 2 | 0 | 0 | # | 10 | Small | aldex2 | 31 |
| # | disease | 4 | 1 | 0 | 0.8 | 1 | # | 10 | Baseline | paralle | 31 |
| # | sex | 2 | 6 | 0 | 0.25 | 1 | # | 10 | Baseline | paralle | 31 |
| # | time | 2 | 7 | 0 | 0.222222222 | 1 | # | 10 | Baseline | paralle | 31 |
| # | interaction | 2 | 2 | 0 | 0.5 | 1 | # | 10 | Baseline | paralle | 31 |
| # | disease | 4 | 2 | 0 | 0.666666667 | 1 | # | 10 | Baseline | ancom | 31 |
| # | sex | 2 | 0 | 0 | 1 | 1 | # | 10 | Baseline | ancom | 31 |
| # | time | 2 | 2 | 0 | 0.5 | 1 | # | 10 | Baseline | ancom | 31 |
| # | interaction | 0 | 0 | 2 | 0 | 0 | # | 10 | Baseline | ancom | 31 |
| # | disease | 4 | 15 | 0 | 0.210526316 | 1 | # | 10 | Baseline | aldex2 | 31 |
| # | sex | 2 | 10 | 0 | 0.166666667 | 1 | # | 10 | Baseline | aldex2 | 31 |
| # | time | 2 | 11 | 0 | 0.153846154 | 1 | # | 10 | Baseline | aldex2 | 31 |
| # | interaction | 0 | 0 | 2 | 0 | 0 | # | 10 | Baseline | aldex2 | 31 |
| # | disease | 4 | 0 | 0 | 1 | 1 | # | 10 | Small | paralle | 31 |
| # | sex | 2 | 0 | 0 | 1 | 1 | # | 10 | Small | paralle | 31 |
| # | time | 2 | 0 | 0 | 1 | 1 | # | 10 | Small | paralle | 31 |
| # | interaction | 2 | 0 | 0 | 1 | 1 | # | 10 | Small | paralle | 31 |
| # | disease | 4 | 1 | 0 | 0.8 | 1 | # | 10 | Small | ancom | 31 |
| # | sex | 2 | 0 | 0 | 1 | 1 | # | 10 | Small | ancom | 31 |
| # | time | 2 | 2 | 0 | 0.5 | 1 | # | 10 | Small | ancom | 31 |
| # | interaction | 0 | 0 | 2 | 0 | 0 | # | 10 | Small | ancom | 31 |
| # | disease | 4 | 2 | 0 | 0.666666667 | 1 | # | 10 | Small | aldex2 | 31 |
| # | sex | 2 | 0 | 0 | 1 | 1 | # | 10 | Small | aldex2 | 31 |
| # | time | 2 | 1 | 0 | 0.666666667 | 1 | # | 10 | Small | aldex2 | 31 |
| # | interaction | 0 | 0 | 2 | 0 | 0 | # | 10 | Small | aldex2 | 31 |
| # | disease | 4 | 3 | 0 | 0.571428571 | 1 | # | 10 | Baseline | paralle | 31 |
| # | sex | 2 | 2 | 0 | 0.5 | 1 | # | 10 | Baseline | paralle | 31 |
| # | time | 2 | 1 | 0 | 0.666666667 | 1 | # | 10 | Baseline | paralle | 31 |
| # | interaction | 2 | 1 | 0 | 0.666666667 | 1 | # | 10 | Baseline | paralle | 31 |
| # | disease | 4 | 2 | 0 | 0.666666667 | 1 | # | 10 | Baseline | ancom | 31 |
| # | sex | 2 | 0 | 0 | 1 | 1 | # | 10 | Baseline | ancom | 31 |
| # | time | 2 | 2 | 0 | 0.5 | 1 | # | 10 | Baseline | ancom | 31 |
| # | interaction | 0 | 0 | 2 | 0 | 0 | # | 10 | Baseline | ancom | 31 |
| # | disease | 4 | 22 | 0 | 0.153846154 | 1 | # | 10 | Baseline | aldex2 | 31 |
| # | sex | 2 | 9 | 0 | 0.181818182 | 1 | # | 10 | Baseline | aldex2 | 31 |
| # | time | 2 | 12 | 0 | 0.142857143 | 1 | # | 10 | Baseline | aldex2 | 31 |
| # | interaction | 0 | 0 | 2 | 0 | 0 | # | 10 | Baseline | aldex2 | 31 |
| # | disease | 4 | 0 | 0 | 1 | 1 | # | 10 | Small | paralle | 31 |
| # | sex | 2 | 0 | 0 | 1 | 1 | # | 10 | Small | paralle | 31 |
| # | time | 2 | 0 | 0 | 1 | 1 | # | 10 | Small | paralle | 31 |
| # | interaction | 2 | 0 | 0 | 1 | 1 | # | 10 | Small | paralle | 31 |
| # | disease | 4 | 2 | 0 | 0.666666667 | 1 | # | 10 | Small | ancom | 31 |
| # | sex | 2 | 0 | 0 | 1 | 1 | # | 10 | Small | ancom | 31 |
| # | time | 2 | 2 | 0 | 0.5 | 1 | # | 10 | Small | ancom | 31 |
| # | interaction | 0 | 0 | 2 | 0 | 0 | # | 10 | Small | ancom | 31 |
| # | disease | 4 | 1 | 0 | 0.8 | 1 | # | 10 | Small | aldex2 | 31 |
| # | sex | 2 | 1 | 0 | 0.666666667 | 1 | # | 10 | Small | aldex2 | 31 |
| # | time | 2 | 4 | 0 | 0.333333333 | 1 | # | 10 | Small | aldex2 | 31 |
| # | interaction | 0 | 0 | 2 | 0 | 0 | # | 10 | Small | aldex2 | 31 |
| # | disease | 4 | 5 | 0 | 0.444444444 | 1 | # | 10 | Baseline | paralle | 31 |
| # | sex | 2 | 4 | 0 | 0.333333333 | 1 | # | 10 | Baseline | paralle | 31 |
| # | time | 2 | 4 | 0 | 0.333333333 | 1 | # | 10 | Baseline | paralle | 31 |
| # | interaction | 2 | 1 | 0 | 0.666666667 | 1 | # | 10 | Baseline | paralle | 31 |
| # | disease | 4 | 2 | 0 | 0.666666667 | 1 | # | 10 | Baseline | ancom | 31 |
| # | sex | 2 | 0 | 0 | 1 | 1 | # | 10 | Baseline | ancom | 31 |
| # | time | 2 | 2 | 0 | 0.5 | 1 | # | 10 | Baseline | ancom | 31 |
| # | interaction | 0 | 0 | 2 | 0 | 0 | # | 10 | Baseline | ancom | 31 |
| # | disease | 4 | 28 | 0 | 0.125 | 1 | # | 10 | Baseline | aldex2 | 31 |
| # | sex | 2 | 8 | 0 | 0.2 | 1 | # | 10 | Baseline | aldex2 | 31 |
| # | time | 2 | 14 | 0 | 0.125 | 1 | # | 10 | Baseline | aldex2 | 31 |
| # | interaction | 0 | 0 | 2 | 0 | 0 | # | 10 | Baseline | aldex2 | 31 |
| # | disease | 4 | 0 | 0 | 1 | 1 | # | 10 | Small | paralle | 31 |
| # | sex | 2 | 0 | 0 | 1 | 1 | # | 10 | Small | paralle | 31 |
| # | time | 2 | 0 | 0 | 1 | 1 | # | 10 | Small | paralle | 31 |
| # | interaction | 2 | 0 | 0 | 1 | 1 | # | 10 | Small | paralle | 31 |
| # | disease | 4 | 2 | 0 | 0.666666667 | 1 | # | 10 | Small | ancom | 31 |
| # | sex | 2 | 0 | 0 | 1 | 1 | # | 10 | Small | ancom | 31 |
| # | time | 2 | 2 | 0 | 0.5 | 1 | # | 10 | Small | ancom | 31 |
| # | interaction | 0 | 0 | 2 | 0 | 0 | # | 10 | Small | ancom | 31 |
| # | disease | 4 | 2 | 0 | 0.666666667 | 1 | # | 10 | Small | aldex2 | 31 |
| # | sex | 2 | 0 | 0 | 1 | 1 | # | 10 | Small | aldex2 | 31 |
| # | time | 2 | 3 | 0 | 0.4 | 1 | # | 10 | Small | aldex2 | 31 |
| # | interaction | 0 | 0 | 2 | 0 | 0 | # | 10 | Small | aldex2 | 31 |
| # | disease | 7 | 11 | 0 | 0.388888889 | 1 | # | 20 | Baseline | paralle | 31 |
| # | sex | 4 | 4 | 0 | 0.5 | 1 | # | 20 | Baseline | paralle | 31 |
| # | time | 5 | 6 | 0 | 0.454545455 | 1 | # | 20 | Baseline | paralle | 31 |
| # | interaction | 4 | 10 | 0 | 0.285714286 | 1 | # | 20 | Baseline | paralle | 31 |
| # | disease | 7 | 1 | 0 | 0.875 | 1 | # | 20 | Baseline | ancom | 31 |
| # | sex | 4 | 0 | 0 | 1 | 1 | # | 20 | Baseline | ancom | 31 |
| # | time | 5 | 2 | 0 | 0.714285714 | 1 | # | 20 | Baseline | ancom | 31 |
| # | interaction | 0 | 0 | 4 | 0 | 0 | # | 20 | Baseline | ancom | 31 |
| # | disease | 7 | 19 | 0 | 0.269230769 | 1 | # | 20 | Baseline | aldex2 | 31 |
| # | sex | 4 | 10 | 0 | 0.285714286 | 1 | # | 20 | Baseline | aldex2 | 31 |
| # | time | 5 | 14 | 0 | 0.263157895 | 1 | # | 20 | Baseline | aldex2 | 31 |
| # | interaction | 0 | 0 | 4 | 0 | 0 | # | 20 | Baseline | aldex2 | 31 |
| # | disease | 7 | 0 | 0 | 1 | 1 | # | 20 | Small | paralle | 31 |
| # | sex | 4 | 1 | 0 | 0.8 | 1 | # | 20 | Small | paralle | 31 |
| # | time | 5 | 0 | 0 | 1 | 1 | # | 20 | Small | paralle | 31 |
| # | interaction | 4 | 0 | 0 | 1 | 1 | # | 20 | Small | paralle | 31 |
| # | disease | 7 | 1 | 0 | 0.875 | 1 | # | 20 | Small | ancom | 31 |
| # | sex | 4 | 0 | 0 | 1 | 1 | # | 20 | Small | ancom | 31 |
| # | time | 5 | 2 | 0 | 0.714285714 | 1 | # | 20 | Small | ancom | 31 |
| # | interaction | 0 | 0 | 4 | 0 | 0 | # | 20 | Small | ancom | 31 |
| # | disease | 7 | 1 | 0 | 0.875 | 1 | # | 20 | Small | aldex2 | 31 |
| # | sex | 4 | 0 | 0 | 1 | 1 | # | 20 | Small | aldex2 | 31 |
| # | time | 5 | 0 | 0 | 1 | 1 | # | 20 | Small | aldex2 | 31 |
| # | interaction | 0 | 0 | 4 | 0 | 0 | # | 20 | Small | aldex2 | 31 |
| # | disease | 7 | 10 | 0 | 0.411764706 | 1 | # | 20 | Baseline | paralle | 31 |
| # | sex | 4 | 4 | 0 | 0.5 | 1 | # | 20 | Baseline | paralle | 31 |
| # | time | 5 | 3 | 0 | 0.625 | 1 | # | 20 | Baseline | paralle | 31 |
| # | interaction | 4 | 6 | 0 | 0.4 | 1 | # | 20 | Baseline | paralle | 31 |
| # | disease | 7 | 3 | 0 | 0.7 | 1 | # | 20 | Baseline | ancom | 31 |
| # | sex | 4 | 0 | 0 | 1 | 1 | # | 20 | Baseline | ancom | 31 |
| # | time | 5 | 4 | 0 | 0.555555556 | 1 | # | 20 | Baseline | ancom | 31 |
| # | interaction | 0 | 0 | 4 | 0 | 0 | # | 20 | Baseline | ancom | 31 |
| # | disease | 7 | 36 | 0 | 0.162790698 | 1 | # | 20 | Baseline | aldex2 | 31 |
| # | sex | 4 | 9 | 0 | 0.307692308 | 1 | # | 20 | Baseline | aldex2 | 31 |
| # | time | 5 | 25 | 0 | 0.166666667 | 1 | # | 20 | Baseline | aldex2 | 31 |
| # | interaction | 0 | 0 | 4 | 0 | 0 | # | 20 | Baseline | aldex2 | 31 |
| # | disease | 7 | 0 | 0 | 1 | 1 | # | 20 | Small | paralle | 31 |
| # | sex | 4 | 0 | 0 | 1 | 1 | # | 20 | Small | paralle | 31 |
| # | time | 5 | 0 | 0 | 1 | 1 | # | 20 | Small | paralle | 31 |
| # | interaction | 4 | 0 | 0 | 1 | 1 | # | 20 | Small | paralle | 31 |
| # | disease | 7 | 3 | 0 | 0.7 | 1 | # | 20 | Small | ancom | 31 |
| # | sex | 4 | 0 | 0 | 1 | 1 | # | 20 | Small | ancom | 31 |
| # | time | 5 | 4 | 0 | 0.555555556 | 1 | # | 20 | Small | ancom | 31 |
| # | interaction | 0 | 0 | 4 | 0 | 0 | # | 20 | Small | ancom | 31 |
| # | disease | 7 | 5 | 0 | 0.583333333 | 1 | # | 20 | Small | aldex2 | 31 |
| # | sex | 4 | 0 | 0 | 1 | 1 | # | 20 | Small | aldex2 | 31 |
| # | time | 5 | 5 | 0 | 0.5 | 1 | # | 20 | Small | aldex2 | 31 |
| # | interaction | 0 | 0 | 4 | 0 | 0 | # | 20 | Small | aldex2 | 31 |
| # | disease | 7 | 13 | 0 | 0.35 | 1 | # | 20 | Baseline | paralle | 31 |
| # | sex | 4 | 2 | 0 | 0.666666667 | 1 | # | 20 | Baseline | paralle | 31 |
| # | time | 5 | 6 | 0 | 0.454545455 | 1 | # | 20 | Baseline | paralle | 31 |
| # | interaction | 4 | 5 | 0 | 0.444444444 | 1 | # | 20 | Baseline | paralle | 31 |
| # | disease | 7 | 5 | 0 | 0.583333333 | 1 | # | 20 | Baseline | ancom | 31 |
| # | sex | 4 | 0 | 0 | 1 | 1 | # | 20 | Baseline | ancom | 31 |
| # | time | 5 | 4 | 0 | 0.555555556 | 1 | # | 20 | Baseline | ancom | 31 |
| # | interaction | 0 | 0 | 4 | 0 | 0 | # | 20 | Baseline | ancom | 31 |
| # | disease | 7 | 56 | 0 | 0.111111111 | 1 | # | 20 | Baseline | aldex2 | 31 |
| # | sex | 4 | 9 | 0 | 0.307692308 | 1 | # | 20 | Baseline | aldex2 | 31 |
| # | time | 5 | 28 | 0 | 0.151515152 | 1 | # | 20 | Baseline | aldex2 | 31 |
| # | interaction | 0 | 0 | 4 | 0 | 0 | # | 20 | Baseline | aldex2 | 31 |
| # | disease | 7 | 1 | 0 | 0.875 | 1 | # | 20 | Small | paralle | 31 |
| # | sex | 4 | 0 | 0 | 1 | 1 | # | 20 | Small | paralle | 31 |
| # | time | 5 | 0 | 0 | 1 | 1 | # | 20 | Small | paralle | 31 |
| # | interaction | 4 | 0 | 0 | 1 | 1 | # | 20 | Small | paralle | 31 |
| # | disease | 7 | 4 | 0 | 0.636363636 | 1 | # | 20 | Small | ancom | 31 |
| # | sex | 4 | 0 | 0 | 1 | 1 | # | 20 | Small | ancom | 31 |
| # | time | 5 | 4 | 0 | 0.555555556 | 1 | # | 20 | Small | ancom | 31 |
| # | interaction | 0 | 0 | 4 | 0 | 0 | # | 20 | Small | ancom | 31 |
| # | disease | 7 | 5 | 0 | 0.583333333 | 1 | # | 20 | Small | aldex2 | 31 |
| # | sex | 4 | 0 | 0 | 1 | 1 | # | 20 | Small | aldex2 | 31 |
| # | time | 5 | 5 | 0 | 0.5 | 1 | # | 20 | Small | aldex2 | 31 |
| # | interaction | 0 | 0 | 4 | 0 | 0 | # | 20 | Small | aldex2 | 31 |
| # | disease | 7 | 14 | 0 | 0.333333333 | 1 | # | 20 | Baseline | paralle | 31 |
| # | sex | 4 | 2 | 0 | 0.666666667 | 1 | # | 20 | Baseline | paralle | 31 |
| # | time | 5 | 4 | 0 | 0.555555556 | 1 | # | 20 | Baseline | paralle | 31 |
| # | interaction | 4 | 7 | 0 | 0.363636364 | 1 | # | 20 | Baseline | paralle | 31 |
| # | disease | 7 | 4 | 0 | 0.636363636 | 1 | # | 20 | Baseline | ancom | 31 |
| # | sex | 4 | 0 | 0 | 1 | 1 | # | 20 | Baseline | ancom | 31 |
| # | time | 5 | 4 | 0 | 0.555555556 | 1 | # | 20 | Baseline | ancom | 31 |
| # | interaction | 0 | 0 | 4 | 0 | 0 | # | 20 | Baseline | ancom | 31 |
| # | disease | 7 | 62 | 0 | 0.101449275 | 1 | # | 20 | Baseline | aldex2 | 31 |
| # | sex | 4 | 9 | 0 | 0.307692308 | 1 | # | 20 | Baseline | aldex2 | 31 |
| # | time | 5 | 39 | 0 | 0.113636364 | 1 | # | 20 | Baseline | aldex2 | 31 |
| # | interaction | 0 | 0 | 4 | 0 | 0 | # | 20 | Baseline | aldex2 | 31 |
| # | disease | 7 | 0 | 0 | 1 | 1 | # | 20 | Small | paralle | 31 |
| # | sex | 4 | 0 | 0 | 1 | 1 | # | 20 | Small | paralle | 31 |
| # | time | 5 | 0 | 0 | 1 | 1 | # | 20 | Small | paralle | 31 |
| # | interaction | 4 | 0 | 0 | 1 | 1 | # | 20 | Small | paralle | 31 |
| # | disease | 7 | 4 | 0 | 0.636363636 | 1 | # | 20 | Small | ancom | 31 |
| # | sex | 4 | 0 | 0 | 1 | 1 | # | 20 | Small | ancom | 31 |
| # | time | 5 | 4 | 0 | 0.555555556 | 1 | # | 20 | Small | ancom | 31 |
| # | interaction | 0 | 0 | 4 | 0 | 0 | # | 20 | Small | ancom | 31 |
| # | disease | 7 | 8 | 0 | 0.466666667 | 1 | # | 20 | Small | aldex2 | 31 |
| # | sex | 4 | 0 | 0 | 1 | 1 | # | 20 | Small | aldex2 | 31 |
| # | time | 5 | 6 | 0 | 0.454545455 | 1 | # | 20 | Small | aldex2 | 31 |
| # | interaction | 0 | 0 | 4 | 0 | 0 | # | 20 | Small | aldex2 | 31 |
| # | disease | 4 | 8 | 0 | 0.333333333 | 1 | # | 10 | Baseline | paralle | 32 |
| # | sex | 2 | 11 | 0 | 0.153846154 | 1 | # | 10 | Baseline | paralle | 32 |
| # | time | 2 | 11 | 0 | 0.153846154 | 1 | # | 10 | Baseline | paralle | 32 |
| # | interaction | 2 | 13 | 0 | 0.133333333 | 1 | # | 10 | Baseline | paralle | 32 |
| # | disease | 4 | 2 | 0 | 0.666666667 | 1 | # | 10 | Baseline | ancom | 32 |
| # | sex | 2 | 0 | 0 | 1 | 1 | # | 10 | Baseline | ancom | 32 |
| # | time | 2 | 0 | 0 | 1 | 1 | # | 10 | Baseline | ancom | 32 |
| # | interaction | 0 | 0 | 2 | 0 | 0 | # | 10 | Baseline | ancom | 32 |
| # | disease | 4 | 6 | 0 | 0.4 | 1 | # | 10 | Baseline | aldex2 | 32 |
| # | sex | 2 | 6 | 0 | 0.25 | 1 | # | 10 | Baseline | aldex2 | 32 |
| # | time | 2 | 7 | 0 | 0.222222222 | 1 | # | 10 | Baseline | aldex2 | 32 |
| # | interaction | 0 | 0 | 2 | 0 | 0 | # | 10 | Baseline | aldex2 | 32 |
| # | disease | 4 | 0 | 0 | 1 | 1 | # | 10 | Small | paralle | 32 |
| # | sex | 2 | 0 | 0 | 1 | 1 | # | 10 | Small | paralle | 32 |
| # | time | 2 | 0 | 0 | 1 | 1 | # | 10 | Small | paralle | 32 |
| # | interaction | 1 | 0 | 1 | 1 | 0.5 | # | 10 | Small | paralle | 32 |
| # | disease | 4 | 1 | 0 | 0.8 | 1 | # | 10 | Small | ancom | 32 |
| # | sex | 2 | 0 | 0 | 1 | 1 | # | 10 | Small | ancom | 32 |
| # | time | 2 | 0 | 0 | 1 | 1 | # | 10 | Small | ancom | 32 |
| # | interaction | 0 | 0 | 2 | 0 | 0 | # | 10 | Small | ancom | 32 |
| # | disease | 4 | 1 | 0 | 0.8 | 1 | # | 10 | Small | aldex2 | 32 |
| # | sex | 2 | 0 | 0 | 1 | 1 | # | 10 | Small | aldex2 | 32 |
| # | time | 2 | 0 | 0 | 1 | 1 | # | 10 | Small | aldex2 | 32 |
| # | interaction | 0 | 0 | 2 | 0 | 0 | # | 10 | Small | aldex2 | 32 |
| # | disease | 4 | 8 | 0 | 0.333333333 | 1 | # | 10 | Baseline | paralle | 32 |
| # | sex | 2 | 10 | 0 | 0.166666667 | 1 | # | 10 | Baseline | paralle | 32 |
| # | time | 2 | 4 | 0 | 0.333333333 | 1 | # | 10 | Baseline | paralle | 32 |
| # | interaction | 2 | 8 | 0 | 0.2 | 1 | # | 10 | Baseline | paralle | 32 |
| # | disease | 4 | 2 | 0 | 0.666666667 | 1 | # | 10 | Baseline | ancom | 32 |
| # | sex | 2 | 0 | 0 | 1 | 1 | # | 10 | Baseline | ancom | 32 |
| # | time | 2 | 0 | 0 | 1 | 1 | # | 10 | Baseline | ancom | 32 |
| # | interaction | 0 | 0 | 2 | 0 | 0 | # | 10 | Baseline | ancom | 32 |
| # | disease | 4 | 17 | 0 | 0.19047619 | 1 | # | 10 | Baseline | aldex2 | 32 |
| # | sex | 2 | 8 | 0 | 0.2 | 1 | # | 10 | Baseline | aldex2 | 32 |
| # | time | 2 | 11 | 0 | 0.153846154 | 1 | # | 10 | Baseline | aldex2 | 32 |
| # | interaction | 0 | 0 | 2 | 0 | 0 | # | 10 | Baseline | aldex2 | 32 |
| # | disease | 4 | 1 | 0 | 0.8 | 1 | # | 10 | Small | paralle | 32 |
| # | sex | 2 | 0 | 0 | 1 | 1 | # | 10 | Small | paralle | 32 |
| # | time | 2 | 0 | 0 | 1 | 1 | # | 10 | Small | paralle | 32 |
| # | interaction | 2 | 0 | 0 | 1 | 1 | # | 10 | Small | paralle | 32 |
| # | disease | 4 | 2 | 0 | 0.666666667 | 1 | # | 10 | Small | ancom | 32 |
| # | sex | 2 | 0 | 0 | 1 | 1 | # | 10 | Small | ancom | 32 |
| # | time | 2 | 0 | 0 | 1 | 1 | # | 10 | Small | ancom | 32 |
| # | interaction | 0 | 0 | 2 | 0 | 0 | # | 10 | Small | ancom | 32 |
| # | disease | 4 | 2 | 0 | 0.666666667 | 1 | # | 10 | Small | aldex2 | 32 |
| # | sex | 2 | 0 | 0 | 1 | 1 | # | 10 | Small | aldex2 | 32 |
| # | time | 2 | 0 | 0 | 1 | 1 | # | 10 | Small | aldex2 | 32 |
| # | interaction | 0 | 0 | 2 | 0 | 0 | # | 10 | Small | aldex2 | 32 |
| # | disease | 4 | 13 | 0 | 0.235294118 | 1 | # | 10 | Baseline | paralle | 32 |
| # | sex | 2 | 5 | 0 | 0.285714286 | 1 | # | 10 | Baseline | paralle | 32 |
| # | time | 2 | 4 | 0 | 0.333333333 | 1 | # | 10 | Baseline | paralle | 32 |
| # | interaction | 2 | 5 | 0 | 0.285714286 | 1 | # | 10 | Baseline | paralle | 32 |
| # | disease | 4 | 2 | 0 | 0.666666667 | 1 | # | 10 | Baseline | ancom | 32 |
| # | sex | 2 | 0 | 0 | 1 | 1 | # | 10 | Baseline | ancom | 32 |
| # | time | 2 | 2 | 0 | 0.5 | 1 | # | 10 | Baseline | ancom | 32 |
| # | interaction | 0 | 0 | 2 | 0 | 0 | # | 10 | Baseline | ancom | 32 |
| # | disease | 4 | 18 | 0 | 0.181818182 | 1 | # | 10 | Baseline | aldex2 | 32 |
| # | sex | 2 | 9 | 0 | 0.181818182 | 1 | # | 10 | Baseline | aldex2 | 32 |
| # | time | 2 | 11 | 0 | 0.153846154 | 1 | # | 10 | Baseline | aldex2 | 32 |
| # | interaction | 0 | 0 | 2 | 0 | 0 | # | 10 | Baseline | aldex2 | 32 |
| # | disease | 4 | 0 | 0 | 1 | 1 | # | 10 | Small | paralle | 32 |
| # | sex | 2 | 0 | 0 | 1 | 1 | # | 10 | Small | paralle | 32 |
| # | time | 2 | 0 | 0 | 1 | 1 | # | 10 | Small | paralle | 32 |
| # | interaction | 2 | 0 | 0 | 1 | 1 | # | 10 | Small | paralle | 32 |
| # | disease | 4 | 2 | 0 | 0.666666667 | 1 | # | 10 | Small | ancom | 32 |
| # | sex | 2 | 0 | 0 | 1 | 1 | # | 10 | Small | ancom | 32 |
| # | time | 2 | 2 | 0 | 0.5 | 1 | # | 10 | Small | ancom | 32 |
| # | interaction | 0 | 0 | 2 | 0 | 0 | # | 10 | Small | ancom | 32 |
| # | disease | 4 | 2 | 0 | 0.666666667 | 1 | # | 10 | Small | aldex2 | 32 |
| # | sex | 2 | 1 | 0 | 0.666666667 | 1 | # | 10 | Small | aldex2 | 32 |
| # | time | 2 | 2 | 0 | 0.5 | 1 | # | 10 | Small | aldex2 | 32 |
| # | interaction | 0 | 0 | 2 | 0 | 0 | # | 10 | Small | aldex2 | 32 |
| # | disease | 4 | 8 | 0 | 0.333333333 | 1 | # | 10 | Baseline | paralle | 32 |
| # | sex | 2 | 6 | 0 | 0.25 | 1 | # | 10 | Baseline | paralle | 32 |
| # | time | 2 | 1 | 0 | 0.666666667 | 1 | # | 10 | Baseline | paralle | 32 |
| # | interaction | 2 | 5 | 0 | 0.285714286 | 1 | # | 10 | Baseline | paralle | 32 |
| # | disease | 4 | 2 | 0 | 0.666666667 | 1 | # | 10 | Baseline | ancom | 32 |
| # | sex | 2 | 0 | 0 | 1 | 1 | # | 10 | Baseline | ancom | 32 |
| # | time | 2 | 2 | 0 | 0.5 | 1 | # | 10 | Baseline | ancom | 32 |
| # | interaction | 0 | 0 | 2 | 0 | 0 | # | 10 | Baseline | ancom | 32 |
| # | disease | 4 | 23 | 0 | 0.148148148 | 1 | # | 10 | Baseline | aldex2 | 32 |
| # | sex | 2 | 9 | 0 | 0.181818182 | 1 | # | 10 | Baseline | aldex2 | 32 |
| # | time | 2 | 13 | 0 | 0.133333333 | 1 | # | 10 | Baseline | aldex2 | 32 |
| # | interaction | 0 | 0 | 2 | 0 | 0 | # | 10 | Baseline | aldex2 | 32 |
| # | disease | 4 | 0 | 0 | 1 | 1 | # | 10 | Small | paralle | 32 |
| # | sex | 2 | 0 | 0 | 1 | 1 | # | 10 | Small | paralle | 32 |
| # | time | 2 | 0 | 0 | 1 | 1 | # | 10 | Small | paralle | 32 |
| # | interaction | 2 | 0 | 0 | 1 | 1 | # | 10 | Small | paralle | 32 |
| # | disease | 4 | 2 | 0 | 0.666666667 | 1 | # | 10 | Small | ancom | 32 |
| # | sex | 2 | 0 | 0 | 1 | 1 | # | 10 | Small | ancom | 32 |
| # | time | 2 | 2 | 0 | 0.5 | 1 | # | 10 | Small | ancom | 32 |
| # | interaction | 0 | 0 | 2 | 0 | 0 | # | 10 | Small | ancom | 32 |
| # | disease | 4 | 3 | 0 | 0.571428571 | 1 | # | 10 | Small | aldex2 | 32 |
| # | sex | 2 | 0 | 0 | 1 | 1 | # | 10 | Small | aldex2 | 32 |
| # | time | 2 | 2 | 0 | 0.5 | 1 | # | 10 | Small | aldex2 | 32 |
| # | interaction | 0 | 0 | 2 | 0 | 0 | # | 10 | Small | aldex2 | 32 |
| # | disease | 7 | 6 | 0 | 0.538461538 | 1 | # | 20 | Baseline | paralle | 32 |
| # | sex | 4 | 7 | 0 | 0.363636364 | 1 | # | 20 | Baseline | paralle | 32 |
| # | time | 5 | 8 | 0 | 0.384615385 | 1 | # | 20 | Baseline | paralle | 32 |
| # | interaction | 4 | 5 | 0 | 0.444444444 | 1 | # | 20 | Baseline | paralle | 32 |
| # | disease | 7 | 3 | 0 | 0.7 | 1 | # | 20 | Baseline | ancom | 32 |
| # | sex | 4 | 0 | 0 | 1 | 1 | # | 20 | Baseline | ancom | 32 |
| # | time | 5 | 2 | 0 | 0.714285714 | 1 | # | 20 | Baseline | ancom | 32 |
| # | interaction | 0 | 0 | 4 | 0 | 0 | # | 20 | Baseline | ancom | 32 |
| # | disease | 7 | 24 | 0 | 0.225806452 | 1 | # | 20 | Baseline | aldex2 | 32 |
| # | sex | 4 | 6 | 0 | 0.4 | 1 | # | 20 | Baseline | aldex2 | 32 |
| # | time | 5 | 12 | 0 | 0.294117647 | 1 | # | 20 | Baseline | aldex2 | 32 |
| # | interaction | 0 | 0 | 4 | 0 | 0 | # | 20 | Baseline | aldex2 | 32 |
| # | disease | 7 | 0 | 0 | 1 | 1 | # | 20 | Small | paralle | 32 |
| # | sex | 4 | 0 | 0 | 1 | 1 | # | 20 | Small | paralle | 32 |
| # | time | 5 | 0 | 0 | 1 | 1 | # | 20 | Small | paralle | 32 |

| | | | | | | | | | | | |
|---|---|---|---|---|---|---|---|---|---|---|---|
| # | interaction | 3 | 0 | 1 | 1 | 0.75 | # | 20 | Small | paralle | 32 |
| # | disease | 7 | 3 | 0 | 0.7 | 1 | # | 20 | Small | ancom | 32 |
| # | sex | 4 | 0 | 0 | 1 | 1 | # | 20 | Small | ancom | 32 |
| # | time | 5 | 2 | 0 | 0.714285714 | 1 | # | 20 | Small | ancom | 32 |
| # | interaction | 0 | 0 | 4 | 0 | 0 | # | 20 | Small | ancom | 32 |
| # | disease | 7 | 3 | 0 | 0.7 | 1 | # | 20 | Small | aldex2 | 32 |
| # | sex | 4 | 0 | 0 | 1 | 1 | # | 20 | Small | aldex2 | 32 |
| # | time | 5 | 4 | 0 | 0.555555556 | 1 | # | 20 | Small | aldex2 | 32 |
| # | interaction | 0 | 0 | 4 | 0 | 0 | # | 20 | Small | aldex2 | 32 |
| # | disease | 7 | 7 | 0 | 0.5 | 1 | # | 20 | Baseline | paralle | 32 |
| # | sex | 4 | 8 | 0 | 0.333333333 | 1 | # | 20 | Baseline | paralle | 32 |
| # | time | 5 | 11 | 0 | 0.3125 | 1 | # | 20 | Baseline | paralle | 32 |
| # | interaction | 4 | 6 | 0 | 0.4 | 1 | # | 20 | Baseline | paralle | 32 |
| # | disease | 7 | 4 | 0 | 0.636363636 | 1 | # | 20 | Baseline | ancom | 32 |
| # | sex | 4 | 0 | 0 | 1 | 1 | # | 20 | Baseline | ancom | 32 |
| # | time | 5 | 2 | 0 | 0.714285714 | 1 | # | 20 | Baseline | ancom | 32 |
| # | interaction | 0 | 0 | 4 | 0 | 0 | # | 20 | Baseline | ancom | 32 |
| # | disease | 7 | 32 | 0 | 0.179487179 | 1 | # | 20 | Baseline | aldex2 | 32 |
| # | sex | 4 | 12 | 0 | 0.25 | 1 | # | 20 | Baseline | aldex2 | 32 |
| # | time | 5 | 26 | 0 | 0.161290323 | 1 | # | 20 | Baseline | aldex2 | 32 |
| # | interaction | 0 | 0 | 4 | 0 | 0 | # | 20 | Baseline | aldex2 | 32 |
| # | disease | 7 | 0 | 0 | 1 | 1 | # | 20 | Small | paralle | 32 |
| # | sex | 4 | 0 | 0 | 1 | 1 | # | 20 | Small | paralle | 32 |
| # | time | 5 | 0 | 0 | 1 | 1 | # | 20 | Small | paralle | 32 |
| # | interaction | 4 | 0 | 0 | 1 | 1 | # | 20 | Small | paralle | 32 |
| # | disease | 7 | 4 | 0 | 0.636363636 | 1 | # | 20 | Small | ancom | 32 |
| # | sex | 4 | 0 | 0 | 1 | 1 | # | 20 | Small | ancom | 32 |
| # | time | 5 | 3 | 0 | 0.625 | 1 | # | 20 | Small | ancom | 32 |
| # | interaction | 0 | 0 | 4 | 0 | 0 | # | 20 | Small | ancom | 32 |
| # | disease | 7 | 4 | 0 | 0.636363636 | 1 | # | 20 | Small | aldex2 | 32 |
| # | sex | 4 | 0 | 0 | 1 | 1 | # | 20 | Small | aldex2 | 32 |
| # | time | 5 | 3 | 0 | 0.625 | 1 | # | 20 | Small | aldex2 | 32 |
| # | interaction | 0 | 0 | 4 | 0 | 0 | # | 20 | Small | aldex2 | 32 |
| # | disease | 7 | 6 | 0 | 0.538461538 | 1 | # | 20 | Baseline | paralle | 32 |
| # | sex | 4 | 6 | 0 | 0.4 | 1 | # | 20 | Baseline | paralle | 32 |
| # | time | 5 | 12 | 0 | 0.294117647 | 1 | # | 20 | Baseline | paralle | 32 |
| # | interaction | 4 | 6 | 0 | 0.4 | 1 | # | 20 | Baseline | paralle | 32 |
| # | disease | 7 | 5 | 0 | 0.583333333 | 1 | # | 20 | Baseline | ancom | 32 |
| # | sex | 4 | 0 | 0 | 1 | 1 | # | 20 | Baseline | ancom | 32 |
| # | time | 5 | 4 | 0 | 0.555555556 | 1 | # | 20 | Baseline | ancom | 32 |
| # | interaction | 0 | 0 | 4 | 0 | 0 | # | 20 | Baseline | ancom | 32 |
| # | disease | 7 | 67 | 0 | 0.12962963 | 1 | # | 20 | Baseline | aldex2 | 32 |
| # | sex | 4 | 16 | 0 | 0.2 | 1 | # | 20 | Baseline | aldex2 | 32 |
| # | time | 5 | 36 | 0 | 0.12195122 | 1 | # | 20 | Baseline | aldex2 | 32 |
| # | interaction | 0 | 0 | 4 | 0 | 0 | # | 20 | Baseline | aldex2 | 32 |
| # | disease | 7 | 1 | 0 | 0.875 | 1 | # | 20 | Small | paralle | 32 |
| # | sex | 4 | 0 | 0 | 1 | 1 | # | 20 | Small | paralle | 32 |
| # | time | 5 | 0 | 0 | 1 | 1 | # | 20 | Small | paralle | 32 |
| # | interaction | 4 | 0 | 0 | 1 | 1 | # | 20 | Small | paralle | 32 |
| # | disease | 7 | 4 | 0 | 0.636363636 | 1 | # | 20 | Small | ancom | 32 |
| # | sex | 4 | 0 | 0 | 1 | 1 | # | 20 | Small | ancom | 32 |
| # | time | 5 | 4 | 0 | 0.555555556 | 1 | # | 20 | Small | ancom | 32 |
| # | interaction | 0 | 0 | 4 | 0 | 0 | # | 20 | Small | ancom | 32 |
| # | disease | 7 | 5 | 0 | 0.583333333 | 1 | # | 20 | Small | aldex2 | 32 |
| # | sex | 4 | 0 | 0 | 1 | 1 | # | 20 | Small | aldex2 | 32 |
| # | time | 5 | 3 | 0 | 0.625 | 1 | # | 20 | Small | aldex2 | 32 |
| # | interaction | 0 | 0 | 4 | 0 | 0 | # | 20 | Small | aldex2 | 32 |
| # | disease | 7 | 5 | 0 | 0.583333333 | 1 | # | 20 | Baseline | paralle | 32 |
| # | sex | 4 | 4 | 0 | 0.5 | 1 | # | 20 | Baseline | paralle | 32 |
| # | time | 5 | 6 | 0 | 0.454545455 | 1 | # | 20 | Baseline | paralle | 32 |
| # | interaction | 4 | 5 | 0 | 0.444444444 | 1 | # | 20 | Baseline | paralle | 32 |
| # | disease | 7 | 4 | 0 | 0.636363636 | 1 | # | 20 | Baseline | ancom | 32 |
| # | sex | 4 | 0 | 0 | 1 | 1 | # | 20 | Baseline | ancom | 32 |
| # | time | 5 | 4 | 0 | 0.555555556 | 1 | # | 20 | Baseline | ancom | 32 |
| # | interaction | 0 | 0 | 4 | 0 | 0 | # | 20 | Baseline | ancom | 32 |
| # | disease | 7 | 54 | 0 | 0.114754098 | 1 | # | 20 | Baseline | aldex2 | 32 |
| # | sex | 4 | 24 | 0 | 0.142857143 | 1 | # | 20 | Baseline | aldex2 | 32 |
| # | time | 5 | 65 | 0 | 0.1 | 1 | # | 20 | Baseline | aldex2 | 32 |
| # | interaction | 0 | 0 | 4 | 0 | 0 | # | 20 | Baseline | aldex2 | 32 |
| # | disease | 7 | 0 | 0 | 1 | 1 | # | 20 | Small | paralle | 32 |
| # | sex | 4 | 0 | 0 | 1 | 1 | # | 20 | Small | paralle | 32 |
| # | time | 5 | 0 | 0 | 1 | 1 | # | 20 | Small | paralle | 32 |
| # | interaction | 4 | 0 | 0 | 1 | 1 | # | 20 | Small | paralle | 32 |
| # | disease | 7 | 4 | 0 | 0.636363636 | 1 | # | 20 | Small | ancom | 32 |
| # | sex | 4 | 0 | 0 | 1 | 1 | # | 20 | Small | ancom | 32 |
| # | time | 5 | 4 | 0 | 0.555555556 | 1 | # | 20 | Small | ancom | 32 |
| # | interaction | 0 | 0 | 4 | 0 | 0 | # | 20 | Small | ancom | 32 |
| # | disease | 7 | 8 | 0 | 0.466666667 | 1 | # | 20 | Small | aldex2 | 32 |
| # | sex | 4 | 0 | 0 | 1 | 1 | # | 20 | Small | aldex2 | 32 |
| # | time | 5 | 5 | 0 | 0.5 | 1 | # | 20 | Small | aldex2 | 32 |
| # | interaction | 0 | 0 | 4 | 0 | 0 | # | 20 | Small | aldex2 | 32 |
| # | disease | 4 | 6 | 0 | 0.4 | 1 | # | 10 | Baseline | paralle | 33 |
| # | sex | 2 | 6 | 0 | 0.25 | 1 | # | 10 | Baseline | paralle | 33 |
| # | time | 2 | 8 | 0 | 0.2 | 1 | # | 10 | Baseline | paralle | 33 |
| # | interaction | 2 | 9 | 0 | 0.181818182 | 1 | # | 10 | Baseline | paralle | 33 |
| # | disease | 4 | 1 | 0 | 0.8 | 1 | # | 10 | Baseline | ancom | 33 |
| # | sex | 2 | 0 | 0 | 1 | 1 | # | 10 | Baseline | ancom | 33 |
| # | time | 2 | 0 | 0 | 1 | 1 | # | 10 | Baseline | ancom | 33 |
| # | interaction | 0 | 0 | 2 | 0 | 0 | # | 10 | Baseline | ancom | 33 |
| # | disease | 4 | 8 | 0 | 0.333333333 | 1 | # | 10 | Baseline | aldex2 | 33 |
| # | sex | 2 | 7 | 0 | 0.222222222 | 1 | # | 10 | Baseline | aldex2 | 33 |
| # | time | 2 | 7 | 0 | 0.222222222 | 1 | # | 10 | Baseline | aldex2 | 33 |
| # | interaction | 0 | 0 | 2 | 0 | 0 | # | 10 | Baseline | aldex2 | 33 |
| # | disease | 4 | 0 | 0 | 1 | 1 | # | 10 | Small | paralle | 33 |
| # | sex | 2 | 0 | 0 | 1 | 1 | # | 10 | Small | paralle | 33 |
| # | time | 2 | 0 | 0 | 1 | 1 | # | 10 | Small | paralle | 33 |
| # | interaction | 1 | 0 | 1 | 1 | 0.5 | # | 10 | Small | paralle | 33 |
| # | disease | 4 | 1 | 0 | 0.8 | 1 | # | 10 | Small | ancom | 33 |
| # | sex | 2 | 0 | 0 | 1 | 1 | # | 10 | Small | ancom | 33 |
| # | time | 2 | 0 | 0 | 1 | 1 | # | 10 | Small | ancom | 33 |
| # | interaction | 0 | 0 | 2 | 0 | 0 | # | 10 | Small | ancom | 33 |
| # | disease | 4 | 1 | 0 | 0.8 | 1 | # | 10 | Small | aldex2 | 33 |
| # | sex | 2 | 0 | 0 | 1 | 1 | # | 10 | Small | aldex2 | 33 |
| # | time | 2 | 0 | 0 | 1 | 1 | # | 10 | Small | aldex2 | 33 |
| # | interaction | 0 | 0 | 2 | 0 | 0 | # | 10 | Small | aldex2 | 33 |
| # | disease | 4 | 5 | 0 | 0.444444444 | 1 | # | 10 | Baseline | paralle | 33 |
| # | sex | 2 | 6 | 0 | 0.25 | 1 | # | 10 | Baseline | paralle | 33 |
| # | time | 2 | 4 | 0 | 0.333333333 | 1 | # | 10 | Baseline | paralle | 33 |
| # | interaction | 2 | 6 | 0 | 0.25 | 1 | # | 10 | Baseline | paralle | 33 |
| # | disease | 4 | 2 | 0 | 0.666666667 | 1 | # | 10 | Baseline | ancom | 33 |
| # | sex | 2 | 0 | 0 | 1 | 1 | # | 10 | Baseline | ancom | 33 |
| # | time | 2 | 2 | 0 | 0.5 | 1 | # | 10 | Baseline | ancom | 33 |
| # | interaction | 0 | 0 | 2 | 0 | 0 | # | 10 | Baseline | ancom | 33 |
| # | disease | 4 | 12 | 0 | 0.25 | 1 | # | 10 | Baseline | aldex2 | 33 |
| # | sex | 2 | 10 | 0 | 0.166666667 | 1 | # | 10 | Baseline | aldex2 | 33 |
| # | time | 2 | 9 | 0 | 0.181818182 | 1 | # | 10 | Baseline | aldex2 | 33 |
| # | interaction | 0 | 0 | 2 | 0 | 0 | # | 10 | Baseline | aldex2 | 33 |
| # | disease | 4 | 0 | 0 | 1 | 1 | # | 10 | Small | paralle | 33 |
| # | sex | 2 | 0 | 0 | 1 | 1 | # | 10 | Small | paralle | 33 |
| # | time | 2 | 0 | 0 | 1 | 1 | # | 10 | Small | paralle | 33 |
| # | interaction | 1 | 0 | 1 | 1 | 0.5 | # | 10 | Small | paralle | 33 |
| # | disease | 4 | 2 | 0 | 0.666666667 | 1 | # | 10 | Small | ancom | 33 |
| # | sex | 2 | 0 | 0 | 1 | 1 | # | 10 | Small | ancom | 33 |
| # | time | 2 | 1 | 0 | 0.666666667 | 1 | # | 10 | Small | ancom | 33 |
| # | interaction | 0 | 0 | 2 | 0 | 0 | # | 10 | Small | ancom | 33 |
| # | disease | 4 | 1 | 0 | 0.8 | 1 | # | 10 | Small | aldex2 | 33 |
| # | sex | 2 | 0 | 0 | 1 | 1 | # | 10 | Small | aldex2 | 33 |
| # | time | 2 | 2 | 0 | 0.5 | 1 | # | 10 | Small | aldex2 | 33 |
| # | interaction | 0 | 0 | 2 | 0 | 0 | # | 10 | Small | aldex2 | 33 |
| # | disease | 4 | 5 | 0 | 0.444444444 | 1 | # | 10 | Baseline | paralle | 33 |
| # | sex | 2 | 5 | 0 | 0.285714286 | 1 | # | 10 | Baseline | paralle | 33 |
| # | time | 2 | 4 | 0 | 0.333333333 | 1 | # | 10 | Baseline | paralle | 33 |
| # | interaction | 2 | 6 | 0 | 0.25 | 1 | # | 10 | Baseline | paralle | 33 |
| # | disease | 4 | 2 | 0 | 0.666666667 | 1 | # | 10 | Baseline | ancom | 33 |
| # | sex | 2 | 0 | 0 | 1 | 1 | # | 10 | Baseline | ancom | 33 |
| # | time | 2 | 2 | 0 | 0.5 | 1 | # | 10 | Baseline | ancom | 33 |
| # | interaction | 0 | 0 | 2 | 0 | 0 | # | 10 | Baseline | ancom | 33 |
| # | disease | 4 | 13 | 0 | 0.235294118 | 1 | # | 10 | Baseline | aldex2 | 33 |
| # | sex | 2 | 10 | 0 | 0.166666667 | 1 | # | 10 | Baseline | aldex2 | 33 |
| # | time | 2 | 9 | 0 | 0.181818182 | 1 | # | 10 | Baseline | aldex2 | 33 |
| # | interaction | 0 | 0 | 2 | 0 | 0 | # | 10 | Baseline | aldex2 | 33 |
| # | disease | 4 | 0 | 0 | 1 | 1 | # | 10 | Small | paralle | 33 |
| # | sex | 2 | 0 | 0 | 1 | 1 | # | 10 | Small | paralle | 33 |
| # | time | 2 | 0 | 0 | 1 | 1 | # | 10 | Small | paralle | 33 |
| # | interaction | 2 | 1 | 0 | 0.666666667 | 1 | # | 10 | Small | paralle | 33 |
| # | disease | 4 | 2 | 0 | 0.666666667 | 1 | # | 10 | Small | ancom | 33 |
| # | sex | 2 | 0 | 0 | 1 | 1 | # | 10 | Small | ancom | 33 |
| # | time | 2 | 3 | 0 | 0.4 | 1 | # | 10 | Small | ancom | 33 |
| # | interaction | 0 | 0 | 2 | 0 | 0 | # | 10 | Small | ancom | 33 |
| # | disease | 4 | 2 | 0 | 0.666666667 | 1 | # | 10 | Small | aldex2 | 33 |
| # | sex | 2 | 0 | 0 | 1 | 1 | # | 10 | Small | aldex2 | 33 |
| # | time | 2 | 4 | 0 | 0.333333333 | 1 | # | 10 | Small | aldex2 | 33 |
| # | interaction | 0 | 0 | 2 | 0 | 0 | # | 10 | Small | aldex2 | 33 |
| # | disease | 4 | 5 | 0 | 0.444444444 | 1 | # | 10 | Baseline | paralle | 33 |
| # | sex | 2 | 3 | 0 | 0.4 | 1 | # | 10 | Baseline | paralle | 33 |
| # | time | 2 | 4 | 0 | 0.333333333 | 1 | # | 10 | Baseline | paralle | 33 |
| # | interaction | 2 | 2 | 0 | 0.5 | 1 | # | 10 | Baseline | paralle | 33 |
| # | disease | 4 | 2 | 0 | 0.666666667 | 1 | # | 10 | Baseline | ancom | 33 |
| # | sex | 2 | 0 | 0 | 1 | 1 | # | 10 | Baseline | ancom | 33 |
| # | time | 2 | 2 | 0 | 0.5 | 1 | # | 10 | Baseline | ancom | 33 |
| # | interaction | 0 | 0 | 2 | 0 | 0 | # | 10 | Baseline | ancom | 33 |
| # | disease | 4 | 24 | 0 | 0.142857143 | 1 | # | 10 | Baseline | aldex2 | 33 |
| # | sex | 2 | 12 | 0 | 0.142857143 | 1 | # | 10 | Baseline | aldex2 | 33 |
| # | time | 2 | 12 | 0 | 0.142857143 | 1 | # | 10 | Baseline | aldex2 | 33 |
| # | interaction | 0 | 0 | 2 | 0 | 0 | # | 10 | Baseline | aldex2 | 33 |
| # | disease | 4 | 0 | 0 | 1 | 1 | # | 10 | Small | paralle | 33 |
| # | sex | 2 | 0 | 0 | 1 | 1 | # | 10 | Small | paralle | 33 |
| # | time | 2 | 0 | 0 | 1 | 1 | # | 10 | Small | paralle | 33 |
| # | interaction | 2 | 0 | 0 | 1 | 1 | # | 10 | Small | paralle | 33 |
| # | disease | 4 | 2 | 0 | 0.666666667 | 1 | # | 10 | Small | ancom | 33 |
| # | sex | 2 | 0 | 0 | 1 | 1 | # | 10 | Small | ancom | 33 |
| # | time | 2 | 2 | 0 | 0.5 | 1 | # | 10 | Small | ancom | 33 |
| # | interaction | 0 | 0 | 2 | 0 | 0 | # | 10 | Small | ancom | 33 |
| # | disease | 4 | 2 | 0 | 0.666666667 | 1 | # | 10 | Small | aldex2 | 33 |
| # | sex | 2 | 0 | 0 | 1 | 1 | # | 10 | Small | aldex2 | 33 |
| # | time | 2 | 3 | 0 | 0.4 | 1 | # | 10 | Small | aldex2 | 33 |
| # | interaction | 0 | 0 | 2 | 0 | 0 | # | 10 | Small | aldex2 | 33 |
| # | disease | 7 | 12 | 0 | 0.368421053 | 1 | # | 20 | Baseline | paralle | 33 |
| # | sex | 4 | 4 | 0 | 0.5 | 1 | # | 20 | Baseline | paralle | 33 |
| # | time | 5 | 10 | 0 | 0.333333333 | 1 | # | 20 | Baseline | paralle | 33 |
| # | interaction | 4 | 7 | 0 | 0.363636364 | 1 | # | 20 | Baseline | paralle | 33 |
| # | disease | 7 | 3 | 0 | 0.7 | 1 | # | 20 | Baseline | ancom | 33 |
| # | sex | 4 | 0 | 0 | 1 | 1 | # | 20 | Baseline | ancom | 33 |
| # | time | 5 | 2 | 0 | 0.714285714 | 1 | # | 20 | Baseline | ancom | 33 |
| # | interaction | 0 | 0 | 4 | 0 | 0 | # | 20 | Baseline | ancom | 33 |
| # | disease | 7 | 19 | 0 | 0.269230769 | 1 | # | 20 | Baseline | aldex2 | 33 |
| # | sex | 4 | 5 | 0 | 0.444444444 | 1 | # | 20 | Baseline | aldex2 | 33 |
| # | time | 5 | 11 | 0 | 0.3125 | 1 | # | 20 | Baseline | aldex2 | 33 |
| # | interaction | 0 | 0 | 4 | 0 | 0 | # | 20 | Baseline | aldex2 | 33 |
| # | disease | 7 | 0 | 0 | 1 | 1 | # | 20 | Small | paralle | 33 |
| # | sex | 4 | 0 | 0 | 1 | 1 | # | 20 | Small | paralle | 33 |
| # | time | 5 | 0 | 0 | 1 | 1 | # | 20 | Small | paralle | 33 |
| # | interaction | 4 | 0 | 0 | 1 | 1 | # | 20 | Small | paralle | 33 |
| # | disease | 7 | 2 | 0 | 0.777777778 | 1 | # | 20 | Small | ancom | 33 |
| # | sex | 4 | 0 | 0 | 1 | 1 | # | 20 | Small | ancom | 33 |
| # | time | 5 | 3 | 0 | 0.625 | 1 | # | 20 | Small | ancom | 33 |
| # | interaction | 0 | 0 | 4 | 0 | 0 | # | 20 | Small | ancom | 33 |
| # | disease | 7 | 3 | 0 | 0.7 | 1 | # | 20 | Small | aldex2 | 33 |
| # | sex | 4 | 0 | 0 | 1 | 1 | # | 20 | Small | aldex2 | 33 |
| # | time | 5 | 3 | 0 | 0.625 | 1 | # | 20 | Small | aldex2 | 33 |
| # | interaction | 0 | 0 | 4 | 0 | 0 | # | 20 | Small | aldex2 | 33 |
| # | disease | 7 | 14 | 0 | 0.333333333 | 1 | # | 20 | Baseline | paralle | 33 |
| # | sex | 4 | 7 | 0 | 0.363636364 | 1 | # | 20 | Baseline | paralle | 33 |
| # | time | 5 | 12 | 0 | 0.294117647 | 1 | # | 20 | Baseline | paralle | 33 |
| # | interaction | 4 | 7 | 0 | 0.363636364 | 1 | # | 20 | Baseline | paralle | 33 |
| # | disease | 7 | 5 | 0 | 0.583333333 | 1 | # | 20 | Baseline | ancom | 33 |
| # | sex | 4 | 0 | 0 | 1 | 1 | # | 20 | Baseline | ancom | 33 |
| # | time | 5 | 4 | 0 | 0.555555556 | 1 | # | 20 | Baseline | ancom | 33 |
| # | interaction | 0 | 0 | 4 | 0 | 0 | # | 20 | Baseline | ancom | 33 |
| # | disease | 7 | 36 | 0 | 0.162790698 | 1 | # | 20 | Baseline | aldex2 | 33 |
| # | sex | 4 | 9 | 0 | 0.307692308 | 1 | # | 20 | Baseline | aldex2 | 33 |
| # | time | 5 | 26 | 0 | 0.161290323 | 1 | # | 20 | Baseline | aldex2 | 33 |
| # | interaction | 0 | 0 | 4 | 0 | 0 | # | 20 | Baseline | aldex2 | 33 |
| # | disease | 7 | 2 | 0 | 0.777777778 | 1 | # | 20 | Small | paralle | 33 |
| # | sex | 4 | 1 | 0 | 0.8 | 1 | # | 20 | Small | paralle | 33 |
| # | time | 5 | 0 | 0 | 1 | 1 | # | 20 | Small | paralle | 33 |
| # | interaction | 4 | 0 | 0 | 1 | 1 | # | 20 | Small | paralle | 33 |
| # | disease | 7 | 4 | 0 | 0.636363636 | 1 | # | 20 | Small | ancom | 33 |
| # | sex | 4 | 0 | 0 | 1 | 1 | # | 20 | Small | ancom | 33 |
| # | time | 5 | 4 | 0 | 0.555555556 | 1 | # | 20 | Small | ancom | 33 |
| # | interaction | 0 | 0 | 4 | 0 | 0 | # | 20 | Small | ancom | 33 |
| # | disease | 7 | 4 | 0 | 0.636363636 | 1 | # | 20 | Small | aldex2 | 33 |
| # | sex | 4 | 0 | 0 | 1 | 1 | # | 20 | Small | aldex2 | 33 |
| # | time | 5 | 4 | 0 | 0.555555556 | 1 | # | 20 | Small | aldex2 | 33 |
| # | interaction | 0 | 0 | 4 | 0 | 0 | # | 20 | Small | aldex2 | 33 |
| # | disease | 7 | 10 | 0 | 0.411764706 | 1 | # | 20 | Baseline | paralle | 33 |
| # | sex | 4 | 8 | 0 | 0.333333333 | 1 | # | 20 | Baseline | paralle | 33 |
| # | time | 5 | 10 | 0 | 0.333333333 | 1 | # | 20 | Baseline | paralle | 33 |
| # | interaction | 4 | 6 | 0 | 0.4 | 1 | # | 20 | Baseline | paralle | 33 |
| # | disease | 7 | 4 | 0 | 0.636363636 | 1 | # | 20 | Baseline | ancom | 33 |
| # | sex | 4 | 0 | 0 | 1 | 1 | # | 20 | Baseline | ancom | 33 |
| # | time | 5 | 4 | 0 | 0.555555556 | 1 | # | 20 | Baseline | ancom | 33 |
| # | interaction | 0 | 0 | 4 | 0 | 0 | # | 20 | Baseline | ancom | 33 |
| # | disease | 7 | 54 | 0 | 0.114754098 | 1 | # | 20 | Baseline | aldex2 | 33 |
| # | sex | 4 | 11 | 0 | 0.266666667 | 1 | # | 20 | Baseline | aldex2 | 33 |
| # | time | 5 | 36 | 0 | 0.12195122 | 1 | # | 20 | Baseline | aldex2 | 33 |
| # | interaction | 0 | 0 | 4 | 0 | 0 | # | 20 | Baseline | aldex2 | 33 |
| # | disease | 7 | 2 | 0 | 0.777777778 | 1 | # | 20 | Small | paralle | 33 |
| # | sex | 4 | 1 | 0 | 0.8 | 1 | # | 20 | Small | paralle | 33 |
| # | time | 5 | 1 | 0 | 0.833333333 | 1 | # | 20 | Small | paralle | 33 |
| # | interaction | 4 | 0 | 0 | 1 | 1 | # | 20 | Small | paralle | 33 |
| # | disease | 7 | 4 | 0 | 0.636363636 | 1 | # | 20 | Small | ancom | 33 |
| # | sex | 4 | 0 | 0 | 1 | 1 | # | 20 | Small | ancom | 33 |
| # | time | 5 | 4 | 0 | 0.555555556 | 1 | # | 20 | Small | ancom | 33 |
| # | interaction | 0 | 0 | 4 | 0 | 0 | # | 20 | Small | ancom | 33 |
| # | disease | 7 | 8 | 0 | 0.466666667 | 1 | # | 20 | Small | aldex2 | 33 |
| # | sex | 4 | 0 | 0 | 1 | 1 | # | 20 | Small | aldex2 | 33 |
| # | time | 5 | 4 | 0 | 0.555555556 | 1 | # | 20 | Small | aldex2 | 33 |
| # | interaction | 0 | 0 | 4 | 0 | 0 | # | 20 | Small | aldex2 | 33 |
| # | disease | 7 | 12 | 0 | 0.368421053 | 1 | # | 20 | Baseline | paralle | 33 |
| # | sex | 4 | 5 | 0 | 0.444444444 | 1 | # | 20 | Baseline | paralle | 33 |
| # | time | 5 | 2 | 0 | 0.714285714 | 1 | # | 20 | Baseline | paralle | 33 |
| # | interaction | 4 | 5 | 0 | 0.444444444 | 1 | # | 20 | Baseline | paralle | 33 |
| # | disease | 7 | 4 | 0 | 0.636363636 | 1 | # | 20 | Baseline | ancom | 33 |
| # | sex | 4 | 0 | 0 | 1 | 1 | # | 20 | Baseline | ancom | 33 |
| # | time | 5 | 4 | 0 | 0.555555556 | 1 | # | 20 | Baseline | ancom | 33 |
| # | interaction | 0 | 0 | 4 | 0 | 0 | # | 20 | Baseline | ancom | 33 |
| # | disease | 7 | 67 | 0 | 0.094594595 | 1 | # | 20 | Baseline | aldex2 | 33 |
| # | sex | 4 | 12 | 0 | 0.25 | 1 | # | 20 | Baseline | aldex2 | 33 |
| # | time | 5 | 43 | 0 | 0.104166667 | 1 | # | 20 | Baseline | aldex2 | 33 |
| # | interaction | 0 | 0 | 4 | 0 | 0 | # | 20 | Baseline | aldex2 | 33 |
| # | disease | 7 | 0 | 0 | 1 | 1 | # | 20 | Small | paralle | 33 |
| # | sex | 4 | 1 | 0 | 0.8 | 1 | # | 20 | Small | paralle | 33 |
| # | time | 5 | 0 | 0 | 1 | 1 | # | 20 | Small | paralle | 33 |
| # | interaction | 4 | 0 | 0 | 1 | 1 | # | 20 | Small | paralle | 33 |
| # | disease | 7 | 4 | 0 | 0.636363636 | 1 | # | 20 | Small | ancom | 33 |
| # | sex | 4 | 0 | 0 | 1 | 1 | # | 20 | Small | ancom | 33 |
| # | time | 5 | 4 | 0 | 0.555555556 | 1 | # | 20 | Small | ancom | 33 |
| # | interaction | 0 | 0 | 4 | 0 | 0 | # | 20 | Small | ancom | 33 |
| # | disease | 7 | 6 | 0 | 0.538461538 | 1 | # | 20 | Small | aldex2 | 33 |
| # | sex | 4 | 1 | 0 | 0.8 | 1 | # | 20 | Small | aldex2 | 33 |
| # | time | 5 | 4 | 0 | 0.555555556 | 1 | # | 20 | Small | aldex2 | 33 |
| # | interaction | 0 | 0 | 4 | 0 | 0 | # | 20 | Small | aldex2 | 33 |
| # | disease | 4 | 10 | 0 | 0.285714286 | 1 | # | 10 | Baseline | paralle | 34 |
| # | sex | 2 | 15 | 0 | 0.117647059 | 1 | # | 10 | Baseline | paralle | 34 |
| # | time | 2 | 11 | 0 | 0.153846154 | 1 | # | 10 | Baseline | paralle | 34 |
| # | interaction | 2 | 10 | 0 | 0.166666667 | 1 | # | 10 | Baseline | paralle | 34 |
| # | disease | 4 | 1 | 0 | 0.8 | 1 | # | 10 | Baseline | ancom | 34 |
| # | sex | 2 | 0 | 0 | 1 | 1 | # | 10 | Baseline | ancom | 34 |
| # | time | 2 | 0 | 0 | 1 | 1 | # | 10 | Baseline | ancom | 34 |
| # | interaction | 0 | 0 | 2 | 0 | 0 | # | 10 | Baseline | ancom | 34 |
| # | disease | 4 | 14 | 0 | 0.222222222 | 1 | # | 10 | Baseline | aldex2 | 34 |
| # | sex | 2 | 9 | 0 | 0.181818182 | 1 | # | 10 | Baseline | aldex2 | 34 |
| # | time | 2 | 12 | 0 | 0.142857143 | 1 | # | 10 | Baseline | aldex2 | 34 |
| # | interaction | 0 | 0 | 2 | 0 | 0 | # | 10 | Baseline | aldex2 | 34 |
| # | disease | 4 | 1 | 0 | 0.8 | 1 | # | 10 | Small | paralle | 34 |
| # | sex | 2 | 0 | 0 | 1 | 1 | # | 10 | Small | paralle | 34 |
| # | time | 2 | 1 | 0 | 0.666666667 | 1 | # | 10 | Small | paralle | 34 |
| # | interaction | 1 | 0 | 1 | 1 | 0.5 | # | 10 | Small | paralle | 34 |
| # | disease | 4 | 1 | 0 | 0.8 | 1 | # | 10 | Small | ancom | 34 |
| # | sex | 2 | 0 | 0 | 1 | 1 | # | 10 | Small | ancom | 34 |
| # | time | 2 | 0 | 0 | 1 | 1 | # | 10 | Small | ancom | 34 |
| # | interaction | 0 | 0 | 2 | 0 | 0 | # | 10 | Small | ancom | 34 |
| # | disease | 4 | 1 | 0 | 0.8 | 1 | # | 10 | Small | aldex2 | 34 |
| # | sex | 2 | 1 | 0 | 0.666666667 | 1 | # | 10 | Small | aldex2 | 34 |
| # | time | 2 | 1 | 0 | 0.666666667 | 1 | # | 10 | Small | aldex2 | 34 |
| # | interaction | 0 | 0 | 2 | 0 | 0 | # | 10 | Small | aldex2 | 34 |
| # | disease | 4 | 7 | 0 | 0.363636364 | 1 | # | 10 | Baseline | paralle | 34 |
| # | sex | 2 | 7 | 0 | 0.222222222 | 1 | # | 10 | Baseline | paralle | 34 |
| # | time | 2 | 6 | 0 | 0.25 | 1 | # | 10 | Baseline | paralle | 34 |
| # | interaction | 2 | 6 | 0 | 0.25 | 1 | # | 10 | Baseline | paralle | 34 |
| # | disease | 4 | 2 | 0 | 0.666666667 | 1 | # | 10 | Baseline | ancom | 34 |
| # | sex | 2 | 0 | 0 | 1 | 1 | # | 10 | Baseline | ancom | 34 |
| # | time | 2 | 2 | 0 | 0.5 | 1 | # | 10 | Baseline | ancom | 34 |
| # | interaction | 0 | 0 | 2 | 0 | 0 | # | 10 | Baseline | ancom | 34 |
| # | disease | 4 | 16 | 0 | 0.2 | 1 | # | 10 | Baseline | aldex2 | 34 |
| # | sex | 2 | 8 | 0 | 0.2 | 1 | # | 10 | Baseline | aldex2 | 34 |
| # | time | 2 | 11 | 0 | 0.153846154 | 1 | # | 10 | Baseline | aldex2 | 34 |
| # | interaction | 0 | 0 | 2 | 0 | 0 | # | 10 | Baseline | aldex2 | 34 |
| # | disease | 4 | 1 | 0 | 0.8 | 1 | # | 10 | Small | paralle | 34 |
| # | sex | 2 | 1 | 0 | 0.666666667 | 1 | # | 10 | Small | paralle | 34 |
| # | time | 2 | 1 | 0 | 0.666666667 | 1 | # | 10 | Small | paralle | 34 |
| # | interaction | 2 | 1 | 0 | 0.666666667 | 1 | # | 10 | Small | paralle | 34 |
| # | disease | 4 | 2 | 0 | 0.666666667 | 1 | # | 10 | Small | ancom | 34 |
| # | sex | 2 | 0 | 0 | 1 | 1 | # | 10 | Small | ancom | 34 |
| # | time | 2 | 1 | 0 | 0.666666667 | 1 | # | 10 | Small | ancom | 34 |
| # | interaction | 0 | 0 | 2 | 0 | 0 | # | 10 | Small | ancom | 34 |
| # | disease | 4 | 2 | 0 | 0.666666667 | 1 | # | 10 | Small | aldex2 | 34 |
| # | sex | 2 | 1 | 0 | 0.666666667 | 1 | # | 10 | Small | aldex2 | 34 |
| # | time | 2 | 2 | 0 | 0.5 | 1 | # | 10 | Small | aldex2 | 34 |
| # | interaction | 0 | 0 | 2 | 0 | 0 | # | 10 | Small | aldex2 | 34 |
| # | disease | 4 | 8 | 0 | 0.333333333 | 1 | # | 10 | Baseline | paralle | 34 |
| # | sex | 2 | 7 | 0 | 0.222222222 | 1 | # | 10 | Baseline | paralle | 34 |
| # | time | 2 | 4 | 0 | 0.333333333 | 1 | # | 10 | Baseline | paralle | 34 |
| # | interaction | 2 | 5 | 0 | 0.285714286 | 1 | # | 10 | Baseline | paralle | 34 |
| # | disease | 4 | 2 | 0 | 0.666666667 | 1 | # | 10 | Baseline | ancom | 34 |
| # | sex | 2 | 0 | 0 | 1 | 1 | # | 10 | Baseline | ancom | 34 |
| # | time | 2 | 2 | 0 | 0.5 | 1 | # | 10 | Baseline | ancom | 34 |
| # | interaction | 0 | 0 | 2 | 0 | 0 | # | 10 | Baseline | ancom | 34 |
| # | disease | 4 | 19 | 0 | 0.173913043 | 1 | # | 10 | Baseline | aldex2 | 34 |
| # | sex | 2 | 9 | 0 | 0.181818182 | 1 | # | 10 | Baseline | aldex2 | 34 |
| # | time | 2 | 14 | 0 | 0.125 | 1 | # | 10 | Baseline | aldex2 | 34 |
| # | interaction | 0 | 0 | 2 | 0 | 0 | # | 10 | Baseline | aldex2 | 34 |
| # | disease | 4 | 0 | 0 | 1 | 1 | # | 10 | Small | paralle | 34 |
| # | sex | 2 | 0 | 0 | 1 | 1 | # | 10 | Small | paralle | 34 |
| # | time | 2 | 0 | 0 | 1 | 1 | # | 10 | Small | paralle | 34 |
| # | interaction | 2 | 0 | 0 | 1 | 1 | # | 10 | Small | paralle | 34 |

| | | | | | | | | | | | |
|---|---|---|---|---|---|---|---|---|---|---|---|
| # | disease | 4 | 2 | 0 | 0.666666667 | 1 | # | 10 | Small | ancom | 34 |
| # | sex | 2 | 0 | 0 | 1 | 1 | # | 10 | Small | ancom | 34 |
| # | time | 2 | 2 | 0 | 0.5 | 1 | # | 10 | Small | ancom | 34 |
| # | interaction | 0 | 0 | 2 | 0 | 0 | # | 10 | Small | ancom | 34 |
| # | disease | 4 | 2 | 0 | 0.666666667 | 1 | # | 10 | Small | aldex2 | 34 |
| # | sex | 2 | 1 | 0 | 0.666666667 | 1 | # | 10 | Small | aldex2 | 34 |
| # | time | 2 | 3 | 0 | 0.4 | 1 | # | 10 | Small | aldex2 | 34 |
| # | interaction | 0 | 0 | 2 | 0 | 0 | # | 10 | Small | aldex2 | 34 |
| # | disease | 4 | 8 | 0 | 0.333333333 | 1 | # | 10 | Baseline | paralle | 34 |
| # | sex | 2 | 9 | 0 | 0.181818182 | 1 | # | 10 | Baseline | paralle | 34 |
| # | time | 2 | 4 | 0 | 0.333333333 | 1 | # | 10 | Baseline | paralle | 34 |
| # | interaction | 2 | 5 | 0 | 0.285714286 | 1 | # | 10 | Baseline | paralle | 34 |
| # | disease | 4 | 2 | 0 | 0.666666667 | 1 | # | 10 | Baseline | ancom | 34 |
| # | sex | 2 | 0 | 0 | 1 | 1 | # | 10 | Baseline | ancom | 34 |
| # | time | 2 | 2 | 0 | 0.5 | 1 | # | 10 | Baseline | ancom | 34 |
| # | interaction | 0 | 0 | 2 | 0 | 0 | # | 10 | Baseline | ancom | 34 |
| # | disease | 4 | 29 | 0 | 0.121212121 | 1 | # | 10 | Baseline | aldex2 | 34 |
| # | sex | 2 | 7 | 0 | 0.222222222 | 1 | # | 10 | Baseline | aldex2 | 34 |
| # | time | 2 | 15 | 0 | 0.117647059 | 1 | # | 10 | Baseline | aldex2 | 34 |
| # | interaction | 0 | 0 | 2 | 0 | 0 | # | 10 | Baseline | aldex2 | 34 |
| # | disease | 4 | 0 | 0 | 1 | 1 | # | 10 | Small | paralle | 34 |
| # | sex | 2 | 1 | 0 | 0.666666667 | 1 | # | 10 | Small | paralle | 34 |
| # | time | 2 | 0 | 0 | 1 | 1 | # | 10 | Small | paralle | 34 |
| # | interaction | 2 | 0 | 0 | 1 | 1 | # | 10 | Small | paralle | 34 |
| # | disease | 4 | 2 | 0 | 0.666666667 | 1 | # | 10 | Small | ancom | 34 |
| # | sex | 2 | 0 | 0 | 1 | 1 | # | 10 | Small | ancom | 34 |
| # | time | 2 | 2 | 0 | 0.5 | 1 | # | 10 | Small | ancom | 34 |
| # | interaction | 0 | 0 | 2 | 0 | 0 | # | 10 | Small | ancom | 34 |
| # | disease | 4 | 2 | 0 | 0.666666667 | 1 | # | 10 | Small | aldex2 | 34 |
| # | sex | 2 | 1 | 0 | 0.666666667 | 1 | # | 10 | Small | aldex2 | 34 |
| # | time | 2 | 2 | 0 | 0.5 | 1 | # | 10 | Small | aldex2 | 34 |
| # | interaction | 0 | 0 | 2 | 0 | 0 | # | 10 | Small | aldex2 | 34 |
| # | disease | 7 | 16 | 0 | 0.304347826 | 1 | # | 20 | Baseline | paralle | 34 |
| # | sex | 4 | 9 | 0 | 0.307692308 | 1 | # | 20 | Baseline | paralle | 34 |
| # | time | 5 | 11 | 0 | 0.3125 | 1 | # | 20 | Baseline | paralle | 34 |
| # | interaction | 3 | 9 | 1 | 0.25 | 0.75 | # | 20 | Baseline | paralle | 34 |
| # | disease | 7 | 0 | 0 | 1 | 1 | # | 20 | Baseline | ancom | 34 |
| # | sex | 4 | 0 | 0 | 1 | 1 | # | 20 | Baseline | ancom | 34 |
| # | time | 5 | 2 | 0 | 0.714285714 | 1 | # | 20 | Baseline | ancom | 34 |
| # | interaction | 0 | 0 | 4 | 0 | 0 | # | 20 | Baseline | ancom | 34 |
| # | disease | 7 | 17 | 0 | 0.291666667 | 1 | # | 20 | Baseline | aldex2 | 34 |
| # | sex | 4 | 8 | 0 | 0.333333333 | 1 | # | 20 | Baseline | aldex2 | 34 |
| # | time | 5 | 14 | 0 | 0.263157895 | 1 | # | 20 | Baseline | aldex2 | 34 |
| # | interaction | 0 | 0 | 4 | 0 | 0 | # | 20 | Baseline | aldex2 | 34 |
| # | disease | 7 | 0 | 0 | 1 | 1 | # | 20 | Small | paralle | 34 |
| # | sex | 4 | 1 | 0 | 0.8 | 1 | # | 20 | Small | paralle | 34 |
| # | time | 5 | 1 | 0 | 0.833333333 | 1 | # | 20 | Small | paralle | 34 |
| # | interaction | 3 | 0 | 1 | 1 | 0.75 | # | 20 | Small | paralle | 34 |
| # | disease | 7 | 0 | 0 | 1 | 1 | # | 20 | Small | ancom | 34 |
| # | sex | 4 | 0 | 0 | 1 | 1 | # | 20 | Small | ancom | 34 |
| # | time | 5 | 2 | 0 | 0.714285714 | 1 | # | 20 | Small | ancom | 34 |
| # | interaction | 0 | 0 | 4 | 0 | 0 | # | 20 | Small | ancom | 34 |
| # | disease | 7 | 0 | 0 | 1 | 1 | # | 20 | Small | aldex2 | 34 |
| # | sex | 4 | 0 | 0 | 1 | 1 | # | 20 | Small | aldex2 | 34 |
| # | time | 5 | 3 | 0 | 0.625 | 1 | # | 20 | Small | aldex2 | 34 |
| # | interaction | 0 | 0 | 4 | 0 | 0 | # | 20 | Small | aldex2 | 34 |
| # | disease | 7 | 7 | 0 | 0.5 | 1 | # | 20 | Baseline | paralle | 34 |
| # | sex | 4 | 5 | 0 | 0.444444444 | 1 | # | 20 | Baseline | paralle | 34 |
| # | time | 5 | 6 | 0 | 0.454545455 | 1 | # | 20 | Baseline | paralle | 34 |
| # | interaction | 4 | 6 | 0 | 0.4 | 1 | # | 20 | Baseline | paralle | 34 |
| # | disease | 7 | 4 | 0 | 0.636363636 | 1 | # | 20 | Baseline | ancom | 34 |
| # | sex | 4 | 0 | 0 | 1 | 1 | # | 20 | Baseline | ancom | 34 |
| # | time | 5 | 3 | 0 | 0.625 | 1 | # | 20 | Baseline | ancom | 34 |
| # | interaction | 0 | 0 | 4 | 0 | 0 | # | 20 | Baseline | ancom | 34 |
| # | disease | 7 | 37 | 0 | 0.159090909 | 1 | # | 20 | Baseline | aldex2 | 34 |
| # | sex | 4 | 7 | 0 | 0.363636364 | 1 | # | 20 | Baseline | aldex2 | 34 |
| # | time | 5 | 24 | 0 | 0.172413793 | 1 | # | 20 | Baseline | aldex2 | 34 |
| # | interaction | 0 | 0 | 4 | 0 | 0 | # | 20 | Baseline | aldex2 | 34 |
| # | disease | 7 | 0 | 0 | 1 | 1 | # | 20 | Small | paralle | 34 |
| # | sex | 4 | 0 | 0 | 1 | 1 | # | 20 | Small | paralle | 34 |
| # | time | 5 | 0 | 0 | 1 | 1 | # | 20 | Small | paralle | 34 |
| # | interaction | 2 | 0 | 2 | 1 | 0.5 | # | 20 | Small | paralle | 34 |
| # | disease | 7 | 4 | 0 | 0.636363636 | 1 | # | 20 | Small | ancom | 34 |
| # | sex | 4 | 0 | 0 | 1 | 1 | # | 20 | Small | ancom | 34 |
| # | time | 5 | 3 | 0 | 0.625 | 1 | # | 20 | Small | ancom | 34 |
| # | interaction | 0 | 0 | 4 | 0 | 0 | # | 20 | Small | ancom | 34 |
| # | disease | 7 | 3 | 0 | 0.7 | 1 | # | 20 | Small | aldex2 | 34 |
| # | sex | 4 | 1 | 0 | 0.8 | 1 | # | 20 | Small | aldex2 | 34 |
| # | time | 5 | 3 | 0 | 0.625 | 1 | # | 20 | Small | aldex2 | 34 |
| # | interaction | 0 | 0 | 4 | 0 | 0 | # | 20 | Small | aldex2 | 34 |
| # | disease | 7 | 10 | 0 | 0.411764706 | 1 | # | 20 | Baseline | paralle | 34 |
| # | sex | 4 | 8 | 0 | 0.333333333 | 1 | # | 20 | Baseline | paralle | 34 |
| # | time | 5 | 3 | 0 | 0.625 | 1 | # | 20 | Baseline | paralle | 34 |
| # | interaction | 4 | 4 | 0 | 0.5 | 1 | # | 20 | Baseline | paralle | 34 |
| # | disease | 7 | 4 | 0 | 0.636363636 | 1 | # | 20 | Baseline | ancom | 34 |
| # | sex | 4 | 0 | 0 | 1 | 1 | # | 20 | Baseline | ancom | 34 |
| # | time | 5 | 4 | 0 | 0.555555556 | 1 | # | 20 | Baseline | ancom | 34 |
| # | interaction | 0 | 0 | 4 | 0 | 0 | # | 20 | Baseline | ancom | 34 |
| # | disease | 7 | 52 | 0 | 0.118644068 | 1 | # | 20 | Baseline | aldex2 | 34 |
| # | sex | 4 | 11 | 0 | 0.266666667 | 1 | # | 20 | Baseline | aldex2 | 34 |
| # | time | 5 | 30 | 0 | 0.142857143 | 1 | # | 20 | Baseline | aldex2 | 34 |
| # | interaction | 0 | 0 | 4 | 0 | 0 | # | 20 | Baseline | aldex2 | 34 |
| # | disease | 7 | 0 | 0 | 1 | 1 | # | 20 | Small | paralle | 34 |
| # | sex | 4 | 0 | 0 | 1 | 1 | # | 20 | Small | paralle | 34 |
| # | time | 5 | 0 | 0 | 1 | 1 | # | 20 | Small | paralle | 34 |
| # | interaction | 4 | 0 | 0 | 1 | 1 | # | 20 | Small | paralle | 34 |
| # | disease | 7 | 4 | 0 | 0.636363636 | 1 | # | 20 | Small | ancom | 34 |
| # | sex | 4 | 0 | 0 | 1 | 1 | # | 20 | Small | ancom | 34 |
| # | time | 5 | 4 | 0 | 0.555555556 | 1 | # | 20 | Small | ancom | 34 |
| # | interaction | 0 | 0 | 4 | 0 | 0 | # | 20 | Small | ancom | 34 |
| # | disease | 7 | 5 | 0 | 0.583333333 | 1 | # | 20 | Small | aldex2 | 34 |
| # | sex | 4 | 0 | 0 | 1 | 1 | # | 20 | Small | aldex2 | 34 |
| # | time | 5 | 5 | 0 | 0.5 | 1 | # | 20 | Small | aldex2 | 34 |
| # | interaction | 0 | 0 | 4 | 0 | 0 | # | 20 | Small | aldex2 | 34 |
| # | disease | 7 | 14 | 0 | 0.333333333 | 1 | # | 20 | Baseline | paralle | 34 |
| # | sex | 4 | 8 | 0 | 0.333333333 | 1 | # | 20 | Baseline | paralle | 34 |
| # | time | 5 | 3 | 0 | 0.625 | 1 | # | 20 | Baseline | paralle | 34 |
| # | interaction | 4 | 3 | 0 | 0.571428571 | 1 | # | 20 | Baseline | paralle | 34 |
| # | disease | 7 | 4 | 0 | 0.636363636 | 1 | # | 20 | Baseline | ancom | 34 |
| # | sex | 4 | 0 | 0 | 1 | 1 | # | 20 | Baseline | ancom | 34 |
| # | time | 5 | 4 | 0 | 0.555555556 | 1 | # | 20 | Baseline | ancom | 34 |
| # | interaction | 0 | 0 | 4 | 0 | 0 | # | 20 | Baseline | ancom | 34 |
| # | disease | 7 | 55 | 0 | 0.112903226 | 1 | # | 20 | Baseline | aldex2 | 34 |
| # | sex | 4 | 13 | 0 | 0.235294118 | 1 | # | 20 | Baseline | aldex2 | 34 |
| # | time | 5 | 37 | 0 | 0.119047619 | 1 | # | 20 | Baseline | aldex2 | 34 |
| # | interaction | 0 | 0 | 4 | 0 | 0 | # | 20 | Baseline | aldex2 | 34 |
| # | disease | 7 | 0 | 0 | 1 | 1 | # | 20 | Small | paralle | 34 |
| # | sex | 4 | 0 | 0 | 1 | 1 | # | 20 | Small | paralle | 34 |
| # | time | 5 | 0 | 0 | 1 | 1 | # | 20 | Small | paralle | 34 |
| # | interaction | 4 | 0 | 0 | 1 | 1 | # | 20 | Small | paralle | 34 |
| # | disease | 7 | 4 | 0 | 0.636363636 | 1 | # | 20 | Small | ancom | 34 |
| # | sex | 4 | 0 | 0 | 1 | 1 | # | 20 | Small | ancom | 34 |
| # | time | 5 | 4 | 0 | 0.555555556 | 1 | # | 20 | Small | ancom | 34 |
| # | interaction | 0 | 0 | 4 | 0 | 0 | # | 20 | Small | ancom | 34 |
| # | disease | 7 | 11 | 0 | 0.388888889 | 1 | # | 20 | Small | aldex2 | 34 |
| # | sex | 4 | 1 | 0 | 0.8 | 1 | # | 20 | Small | aldex2 | 34 |
| # | time | 5 | 5 | 0 | 0.5 | 1 | # | 20 | Small | aldex2 | 34 |
| # | interaction | 0 | 0 | 4 | 0 | 0 | # | 20 | Small | aldex2 | 34 |
| # | disease | 4 | 10 | 0 | 0.285714286 | 1 | # | 10 | Baseline | paralle | 35 |
| # | sex | 2 | 11 | 0 | 0.153846154 | 1 | # | 10 | Baseline | paralle | 35 |
| # | time | 2 | 10 | 0 | 0.166666667 | 1 | # | 10 | Baseline | paralle | 35 |
| # | interaction | 2 | 15 | 0 | 0.117647059 | 1 | # | 10 | Baseline | paralle | 35 |
| # | disease | 4 | 1 | 0 | 0.8 | 1 | # | 10 | Baseline | ancom | 35 |
| # | sex | 2 | 0 | 0 | 1 | 1 | # | 10 | Baseline | ancom | 35 |
| # | time | 2 | 0 | 0 | 1 | 1 | # | 10 | Baseline | ancom | 35 |
| # | interaction | 0 | 0 | 2 | 0 | 0 | # | 10 | Baseline | ancom | 35 |
| # | disease | 4 | 15 | 0 | 0.210526316 | 1 | # | 10 | Baseline | aldex2 | 35 |
| # | sex | 2 | 1 | 0 | 0.666666667 | 1 | # | 10 | Baseline | aldex2 | 35 |
| # | time | 2 | 5 | 0 | 0.285714286 | 1 | # | 10 | Baseline | aldex2 | 35 |
| # | interaction | 0 | 0 | 2 | 0 | 0 | # | 10 | Baseline | aldex2 | 35 |
| # | disease | 4 | 0 | 0 | 1 | 1 | # | 10 | Small | paralle | 35 |
| # | sex | 2 | 0 | 0 | 1 | 1 | # | 10 | Small | paralle | 35 |
| # | time | 2 | 0 | 0 | 1 | 1 | # | 10 | Small | paralle | 35 |
| # | interaction | 0 | 0 | 2 | 0 | 0 | # | 10 | Small | paralle | 35 |
| # | disease | 4 | 1 | 0 | 0.8 | 1 | # | 10 | Small | ancom | 35 |
| # | sex | 2 | 0 | 0 | 1 | 1 | # | 10 | Small | ancom | 35 |
| # | time | 2 | 0 | 0 | 1 | 1 | # | 10 | Small | ancom | 35 |
| # | interaction | 0 | 0 | 2 | 0 | 0 | # | 10 | Small | ancom | 35 |
| # | disease | 4 | 1 | 0 | 0.8 | 1 | # | 10 | Small | aldex2 | 35 |
| # | sex | 2 | 0 | 0 | 1 | 1 | # | 10 | Small | aldex2 | 35 |
| # | time | 2 | 0 | 0 | 1 | 1 | # | 10 | Small | aldex2 | 35 |
| # | interaction | 0 | 0 | 2 | 0 | 0 | # | 10 | Small | aldex2 | 35 |
| # | disease | 4 | 7 | 0 | 0.363636364 | 1 | # | 10 | Baseline | paralle | 35 |
| # | sex | 2 | 3 | 0 | 0.4 | 1 | # | 10 | Baseline | paralle | 35 |
| # | time | 2 | 3 | 0 | 0.4 | 1 | # | 10 | Baseline | paralle | 35 |
| # | interaction | 2 | 4 | 0 | 0.333333333 | 1 | # | 10 | Baseline | paralle | 35 |
| # | disease | 4 | 2 | 0 | 0.666666667 | 1 | # | 10 | Baseline | ancom | 35 |
| # | sex | 2 | 0 | 0 | 1 | 1 | # | 10 | Baseline | ancom | 35 |
| # | time | 2 | 2 | 0 | 0.5 | 1 | # | 10 | Baseline | ancom | 35 |
| # | interaction | 0 | 0 | 2 | 0 | 0 | # | 10 | Baseline | ancom | 35 |
| # | disease | 4 | 16 | 0 | 0.2 | 1 | # | 10 | Baseline | aldex2 | 35 |
| # | sex | 2 | 5 | 0 | 0.285714286 | 1 | # | 10 | Baseline | aldex2 | 35 |
| # | time | 2 | 10 | 0 | 0.166666667 | 1 | # | 10 | Baseline | aldex2 | 35 |
| # | interaction | 0 | 0 | 2 | 0 | 0 | # | 10 | Baseline | aldex2 | 35 |
| # | disease | 4 | 0 | 0 | 1 | 1 | # | 10 | Small | paralle | 35 |
| # | sex | 2 | 0 | 0 | 1 | 1 | # | 10 | Small | paralle | 35 |
| # | time | 2 | 0 | 0 | 1 | 1 | # | 10 | Small | paralle | 35 |
| # | interaction | 2 | 0 | 0 | 1 | 1 | # | 10 | Small | paralle | 35 |
| # | disease | 4 | 2 | 0 | 0.666666667 | 1 | # | 10 | Small | ancom | 35 |
| # | sex | 2 | 0 | 0 | 1 | 1 | # | 10 | Small | ancom | 35 |
| # | time | 2 | 2 | 0 | 0.5 | 1 | # | 10 | Small | ancom | 35 |
| # | interaction | 0 | 0 | 2 | 0 | 0 | # | 10 | Small | ancom | 35 |
| # | disease | 4 | 1 | 0 | 0.8 | 1 | # | 10 | Small | aldex2 | 35 |
| # | sex | 2 | 0 | 0 | 1 | 1 | # | 10 | Small | aldex2 | 35 |
| # | time | 2 | 1 | 0 | 0.666666667 | 1 | # | 10 | Small | aldex2 | 35 |
| # | interaction | 0 | 0 | 2 | 0 | 0 | # | 10 | Small | aldex2 | 35 |
| # | disease | 4 | 3 | 0 | 0.571428571 | 1 | # | 10 | Baseline | paralle | 35 |
| # | sex | 2 | 5 | 0 | 0.285714286 | 1 | # | 10 | Baseline | paralle | 35 |
| # | time | 2 | 1 | 0 | 0.666666667 | 1 | # | 10 | Baseline | paralle | 35 |
| # | interaction | 2 | 1 | 0 | 0.666666667 | 1 | # | 10 | Baseline | paralle | 35 |
| # | disease | 4 | 2 | 0 | 0.666666667 | 1 | # | 10 | Baseline | ancom | 35 |
| # | sex | 2 | 0 | 0 | 1 | 1 | # | 10 | Baseline | ancom | 35 |
| # | time | 2 | 2 | 0 | 0.5 | 1 | # | 10 | Baseline | ancom | 35 |
| # | interaction | 0 | 0 | 2 | 0 | 0 | # | 10 | Baseline | ancom | 35 |
| # | disease | 4 | 21 | 0 | 0.16 | 1 | # | 10 | Baseline | aldex2 | 35 |
| # | sex | 2 | 5 | 0 | 0.285714286 | 1 | # | 10 | Baseline | aldex2 | 35 |
| # | time | 2 | 12 | 0 | 0.142857143 | 1 | # | 10 | Baseline | aldex2 | 35 |
| # | interaction | 0 | 0 | 2 | 0 | 0 | # | 10 | Baseline | aldex2 | 35 |
| # | disease | 4 | 0 | 0 | 1 | 1 | # | 10 | Small | paralle | 35 |
| # | sex | 2 | 0 | 0 | 1 | 1 | # | 10 | Small | paralle | 35 |
| # | time | 2 | 0 | 0 | 1 | 1 | # | 10 | Small | paralle | 35 |
| # | interaction | 2 | 0 | 0 | 1 | 1 | # | 10 | Small | paralle | 35 |
| # | disease | 4 | 2 | 0 | 0.666666667 | 1 | # | 10 | Small | ancom | 35 |
| # | sex | 2 | 0 | 0 | 1 | 1 | # | 10 | Small | ancom | 35 |
| # | time | 2 | 2 | 0 | 0.5 | 1 | # | 10 | Small | ancom | 35 |
| # | interaction | 0 | 0 | 2 | 0 | 0 | # | 10 | Small | ancom | 35 |
| # | disease | 4 | 2 | 0 | 0.666666667 | 1 | # | 10 | Small | aldex2 | 35 |
| # | sex | 2 | 0 | 0 | 1 | 1 | # | 10 | Small | aldex2 | 35 |
| # | time | 2 | 1 | 0 | 0.666666667 | 1 | # | 10 | Small | aldex2 | 35 |
| # | interaction | 0 | 0 | 2 | 0 | 0 | # | 10 | Small | aldex2 | 35 |
| # | disease | 4 | 5 | 0 | 0.444444444 | 1 | # | 10 | Baseline | paralle | 35 |
| # | sex | 2 | 2 | 0 | 0.5 | 1 | # | 10 | Baseline | paralle | 35 |
| # | time | 2 | 3 | 0 | 0.4 | 1 | # | 10 | Baseline | paralle | 35 |
| # | interaction | 2 | 2 | 0 | 0.5 | 1 | # | 10 | Baseline | paralle | 35 |
| # | disease | 4 | 2 | 0 | 0.666666667 | 1 | # | 10 | Baseline | ancom | 35 |
| # | sex | 2 | 0 | 0 | 1 | 1 | # | 10 | Baseline | ancom | 35 |
| # | time | 2 | 2 | 0 | 0.5 | 1 | # | 10 | Baseline | ancom | 35 |
| # | interaction | 0 | 0 | 2 | 0 | 0 | # | 10 | Baseline | ancom | 35 |
| # | disease | 4 | 22 | 0 | 0.153846154 | 1 | # | 10 | Baseline | aldex2 | 35 |
| # | sex | 2 | 8 | 0 | 0.2 | 1 | # | 10 | Baseline | aldex2 | 35 |
| # | time | 2 | 11 | 0 | 0.153846154 | 1 | # | 10 | Baseline | aldex2 | 35 |
| # | interaction | 0 | 0 | 2 | 0 | 0 | # | 10 | Baseline | aldex2 | 35 |
| # | disease | 4 | 0 | 0 | 1 | 1 | # | 10 | Small | paralle | 35 |
| # | sex | 2 | 0 | 0 | 1 | 1 | # | 10 | Small | paralle | 35 |
| # | time | 2 | 0 | 0 | 1 | 1 | # | 10 | Small | paralle | 35 |
| # | interaction | 2 | 0 | 0 | 1 | 1 | # | 10 | Small | paralle | 35 |
| # | disease | 4 | 2 | 0 | 0.666666667 | 1 | # | 10 | Small | ancom | 35 |
| # | sex | 2 | 0 | 0 | 1 | 1 | # | 10 | Small | ancom | 35 |
| # | time | 2 | 2 | 0 | 0.5 | 1 | # | 10 | Small | ancom | 35 |
| # | interaction | 0 | 0 | 2 | 0 | 0 | # | 10 | Small | ancom | 35 |
| # | disease | 4 | 3 | 0 | 0.571428571 | 1 | # | 10 | Small | aldex2 | 35 |
| # | sex | 2 | 0 | 0 | 1 | 1 | # | 10 | Small | aldex2 | 35 |
| # | time | 2 | 2 | 0 | 0.5 | 1 | # | 10 | Small | aldex2 | 35 |
| # | interaction | 0 | 0 | 2 | 0 | 0 | # | 10 | Small | aldex2 | 35 |
| # | disease | 7 | 5 | 0 | 0.583333333 | 1 | # | 20 | Baseline | paralle | 35 |
| # | sex | 4 | 3 | 0 | 0.571428571 | 1 | # | 20 | Baseline | paralle | 35 |
| # | time | 5 | 11 | 0 | 0.3125 | 1 | # | 20 | Baseline | paralle | 35 |
| # | interaction | 4 | 6 | 0 | 0.4 | 1 | # | 20 | Baseline | paralle | 35 |
| # | disease | 7 | 0 | 0 | 1 | 1 | # | 20 | Baseline | ancom | 35 |
| # | sex | 4 | 0 | 0 | 1 | 1 | # | 20 | Baseline | ancom | 35 |
| # | time | 5 | 0 | 0 | 1 | 1 | # | 20 | Baseline | ancom | 35 |
| # | interaction | 0 | 0 | 4 | 0 | 0 | # | 20 | Baseline | ancom | 35 |
| # | disease | 7 | 22 | 0 | 0.24137931 | 1 | # | 20 | Baseline | aldex2 | 35 |
| # | sex | 4 | 7 | 0 | 0.363636364 | 1 | # | 20 | Baseline | aldex2 | 35 |
| # | time | 5 | 17 | 0 | 0.227272727 | 1 | # | 20 | Baseline | aldex2 | 35 |
| # | interaction | 0 | 0 | 4 | 0 | 0 | # | 20 | Baseline | aldex2 | 35 |
| # | disease | 7 | 0 | 0 | 1 | 1 | # | 20 | Small | paralle | 35 |
| # | sex | 4 | 0 | 0 | 1 | 1 | # | 20 | Small | paralle | 35 |
| # | time | 5 | 0 | 0 | 1 | 1 | # | 20 | Small | paralle | 35 |
| # | interaction | 2 | 0 | 2 | 1 | 0.5 | # | 20 | Small | paralle | 35 |
| # | disease | 7 | 0 | 0 | 1 | 1 | # | 20 | Small | ancom | 35 |
| # | sex | 4 | 0 | 0 | 1 | 1 | # | 20 | Small | ancom | 35 |
| # | time | 5 | 0 | 0 | 1 | 1 | # | 20 | Small | ancom | 35 |
| # | interaction | 0 | 0 | 4 | 0 | 0 | # | 20 | Small | ancom | 35 |
| # | disease | 7 | 0 | 0 | 1 | 1 | # | 20 | Small | aldex2 | 35 |
| # | sex | 4 | 0 | 0 | 1 | 1 | # | 20 | Small | aldex2 | 35 |
| # | time | 5 | 1 | 0 | 0.833333333 | 1 | # | 20 | Small | aldex2 | 35 |
| # | interaction | 0 | 0 | 4 | 0 | 0 | # | 20 | Small | aldex2 | 35 |
| # | disease | 7 | 5 | 0 | 0.583333333 | 1 | # | 20 | Baseline | paralle | 35 |
| # | sex | 4 | 5 | 0 | 0.444444444 | 1 | # | 20 | Baseline | paralle | 35 |
| # | time | 5 | 8 | 0 | 0.384615385 | 1 | # | 20 | Baseline | paralle | 35 |
| # | interaction | 4 | 6 | 0 | 0.4 | 1 | # | 20 | Baseline | paralle | 35 |
| # | disease | 7 | 4 | 0 | 0.636363636 | 1 | # | 20 | Baseline | ancom | 35 |
| # | sex | 4 | 0 | 0 | 1 | 1 | # | 20 | Baseline | ancom | 35 |
| # | time | 5 | 3 | 0 | 0.625 | 1 | # | 20 | Baseline | ancom | 35 |
| # | interaction | 0 | 0 | 4 | 0 | 0 | # | 20 | Baseline | ancom | 35 |
| # | disease | 7 | 40 | 0 | 0.14893617 | 1 | # | 20 | Baseline | aldex2 | 35 |
| # | sex | 4 | 9 | 0 | 0.307692308 | 1 | # | 20 | Baseline | aldex2 | 35 |
| # | time | 5 | 23 | 0 | 0.178571429 | 1 | # | 20 | Baseline | aldex2 | 35 |
| # | interaction | 0 | 0 | 4 | 0 | 0 | # | 20 | Baseline | aldex2 | 35 |
| # | disease | 7 | 0 | 0 | 1 | 1 | # | 20 | Small | paralle | 35 |
| # | sex | 4 | 0 | 0 | 1 | 1 | # | 20 | Small | paralle | 35 |
| # | time | 5 | 0 | 0 | 1 | 1 | # | 20 | Small | paralle | 35 |
| # | interaction | 4 | 0 | 0 | 1 | 1 | # | 20 | Small | paralle | 35 |
| # | disease | 7 | 4 | 0 | 0.636363636 | 1 | # | 20 | Small | ancom | 35 |
| # | sex | 4 | 0 | 0 | 1 | 1 | # | 20 | Small | ancom | 35 |
| # | time | 5 | 3 | 0 | 0.625 | 1 | # | 20 | Small | ancom | 35 |
| # | interaction | 0 | 0 | 4 | 0 | 0 | # | 20 | Small | ancom | 35 |
| # | disease | 7 | 3 | 0 | 0.7 | 1 | # | 20 | Small | aldex2 | 35 |
| # | sex | 4 | 0 | 0 | 1 | 1 | # | 20 | Small | aldex2 | 35 |
| # | time | 5 | 7 | 0 | 0.416666667 | 1 | # | 20 | Small | aldex2 | 35 |
| # | interaction | 0 | 0 | 4 | 0 | 0 | # | 20 | Small | aldex2 | 35 |
| # | disease | 7 | 10 | 0 | 0.411764706 | 1 | # | 20 | Baseline | paralle | 35 |
| # | sex | 4 | 3 | 0 | 0.571428571 | 1 | # | 20 | Baseline | paralle | 35 |
| # | time | 5 | 11 | 0 | 0.3125 | 1 | # | 20 | Baseline | paralle | 35 |
| # | interaction | 4 | 6 | 0 | 0.4 | 1 | # | 20 | Baseline | paralle | 35 |
| # | disease | 7 | 4 | 0 | 0.636363636 | 1 | # | 20 | Baseline | ancom | 35 |
| # | sex | 4 | 0 | 0 | 1 | 1 | # | 20 | Baseline | ancom | 35 |
| # | time | 5 | 4 | 0 | 0.555555556 | 1 | # | 20 | Baseline | ancom | 35 |
| # | interaction | 0 | 0 | 4 | 0 | 0 | # | 20 | Baseline | ancom | 35 |
| # | disease | 7 | 52 | 0 | 0.118644068 | 1 | # | 20 | Baseline | aldex2 | 35 |
| # | sex | 4 | 13 | 0 | 0.235294118 | 1 | # | 20 | Baseline | aldex2 | 35 |
| # | time | 5 | 28 | 0 | 0.151515152 | 1 | # | 20 | Baseline | aldex2 | 35 |
| # | interaction | 0 | 0 | 4 | 0 | 0 | # | 20 | Baseline | aldex2 | 35 |
| # | disease | 7 | 0 | 0 | 1 | 1 | # | 20 | Small | paralle | 35 |
| # | sex | 4 | 0 | 0 | 1 | 1 | # | 20 | Small | paralle | 35 |
| # | time | 5 | 0 | 0 | 1 | 1 | # | 20 | Small | paralle | 35 |
| # | interaction | 4 | 0 | 0 | 1 | 1 | # | 20 | Small | paralle | 35 |
| # | disease | 7 | 4 | 0 | 0.636363636 | 1 | # | 20 | Small | ancom | 35 |
| # | sex | 4 | 0 | 0 | 1 | 1 | # | 20 | Small | ancom | 35 |
| # | time | 5 | 4 | 0 | 0.555555556 | 1 | # | 20 | Small | ancom | 35 |
| # | interaction | 0 | 0 | 4 | 0 | 0 | # | 20 | Small | ancom | 35 |
| # | disease | 7 | 4 | 0 | 0.636363636 | 1 | # | 20 | Small | aldex2 | 35 |
| # | sex | 4 | 0 | 0 | 1 | 1 | # | 20 | Small | aldex2 | 35 |
| # | time | 5 | 5 | 0 | 0.5 | 1 | # | 20 | Small | aldex2 | 35 |
| # | interaction | 0 | 0 | 4 | 0 | 0 | # | 20 | Small | aldex2 | 35 |
| # | disease | 7 | 7 | 0 | 0.5 | 1 | # | 20 | Baseline | paralle | 35 |
| # | sex | 4 | 5 | 0 | 0.444444444 | 1 | # | 20 | Baseline | paralle | 35 |
| # | time | 5 | 5 | 0 | 0.5 | 1 | # | 20 | Baseline | paralle | 35 |
| # | interaction | 4 | 1 | 0 | 0.8 | 1 | # | 20 | Baseline | paralle | 35 |
| # | disease | 7 | 4 | 0 | 0.636363636 | 1 | # | 20 | Baseline | ancom | 35 |
| # | sex | 4 | 0 | 0 | 1 | 1 | # | 20 | Baseline | ancom | 35 |
| # | time | 5 | 4 | 0 | 0.555555556 | 1 | # | 20 | Baseline | ancom | 35 |
| # | interaction | 0 | 0 | 4 | 0 | 0 | # | 20 | Baseline | ancom | 35 |
| # | disease | 7 | 66 | 0 | 0.095890411 | 1 | # | 20 | Baseline | aldex2 | 35 |
| # | sex | 4 | 17 | 0 | 0.19047619 | 1 | # | 20 | Baseline | aldex2 | 35 |
| # | time | 5 | 36 | 0 | 0.12195122 | 1 | # | 20 | Baseline | aldex2 | 35 |
| # | interaction | 0 | 0 | 4 | 0 | 0 | # | 20 | Baseline | aldex2 | 35 |
| # | disease | 7 | 0 | 0 | 1 | 1 | # | 20 | Small | paralle | 35 |
| # | sex | 4 | 0 | 0 | 1 | 1 | # | 20 | Small | paralle | 35 |
| # | time | 5 | 0 | 0 | 1 | 1 | # | 20 | Small | paralle | 35 |
| # | interaction | 4 | 0 | 0 | 1 | 1 | # | 20 | Small | paralle | 35 |
| # | disease | 7 | 4 | 0 | 0.636363636 | 1 | # | 20 | Small | ancom | 35 |
| # | sex | 4 | 0 | 0 | 1 | 1 | # | 20 | Small | ancom | 35 |
| # | time | 5 | 4 | 0 | 0.555555556 | 1 | # | 20 | Small | ancom | 35 |
| # | interaction | 0 | 0 | 4 | 0 | 0 | # | 20 | Small | ancom | 35 |
| # | disease | 7 | 6 | 0 | 0.538461538 | 1 | # | 20 | Small | aldex2 | 35 |
| # | sex | 4 | 0 | 0 | 1 | 1 | # | 20 | Small | aldex2 | 35 |
| # | time | 5 | 7 | 0 | 0.416666667 | 1 | # | 20 | Small | aldex2 | 35 |
| # | interaction | 0 | 0 | 4 | 0 | 0 | # | 20 | Small | aldex2 | 35 |
| # | disease | 4 | 1 | 0 | 0.8 | 1 | # | 10 | Baseline | paralle | 36 |
| # | sex | 2 | 6 | 0 | 0.25 | 1 | # | 10 | Baseline | paralle | 36 |
| # | time | 2 | 6 | 0 | 0.25 | 1 | # | 10 | Baseline | paralle | 36 |
| # | interaction | 2 | 9 | 0 | 0.181818182 | 1 | # | 10 | Baseline | paralle | 36 |
| # | disease | 4 | 1 | 0 | 0.8 | 1 | # | 10 | Baseline | ancom | 36 |
| # | sex | 2 | 0 | 0 | 1 | 1 | # | 10 | Baseline | ancom | 36 |
| # | time | 2 | 0 | 0 | 1 | 1 | # | 10 | Baseline | ancom | 36 |
| # | interaction | 0 | 0 | 2 | 0 | 0 | # | 10 | Baseline | ancom | 36 |
| # | disease | 4 | 11 | 0 | 0.266666667 | 1 | # | 10 | Baseline | aldex2 | 36 |
| # | sex | 2 | 7 | 0 | 0.222222222 | 1 | # | 10 | Baseline | aldex2 | 36 |
| # | time | 2 | 12 | 0 | 0.142857143 | 1 | # | 10 | Baseline | aldex2 | 36 |
| # | interaction | 0 | 0 | 2 | 0 | 0 | # | 10 | Baseline | aldex2 | 36 |
| # | disease | 4 | 0 | 0 | 1 | 1 | # | 10 | Small | paralle | 36 |
| # | sex | 2 | 0 | 0 | 1 | 1 | # | 10 | Small | paralle | 36 |
| # | time | 2 | 0 | 0 | 1 | 1 | # | 10 | Small | paralle | 36 |
| # | interaction | 1 | 0 | 1 | 1 | 0.5 | # | 10 | Small | paralle | 36 |
| # | disease | 4 | 0 | 0 | 1 | 1 | # | 10 | Small | ancom | 36 |

| | | | | | | | | | | | |
|---|---|---|---|---|---|---|---|---|---|---|---|
| # | sex | 2 | 0 | 0 | 1 | 1 | # | 10 | Small | ancom | 36 |
| # | time | 2 | 0 | 0 | 1 | 1 | # | 10 | Small | ancom | 36 |
| # | interaction | 0 | 0 | 2 | 0 | 0 | # | 10 | Small | ancom | 36 |
| # | disease | 4 | 1 | 0 | 0.8 | 1 | # | 10 | Small | aldex2 | 36 |
| # | sex | 2 | 0 | 0 | 1 | 1 | # | 10 | Small | aldex2 | 36 |
| # | time | 2 | 0 | 0 | 1 | 1 | # | 10 | Small | aldex2 | 36 |
| # | interaction | 0 | 0 | 2 | 0 | 0 | # | 10 | Small | aldex2 | 36 |
| # | disease | 4 | 4 | 0 | 0.5 | 1 | # | 10 | Baseline | paralle | 36 |
| # | sex | 2 | 6 | 0 | 0.25 | 1 | # | 10 | Baseline | paralle | 36 |
| # | time | 2 | 8 | 0 | 0.2 | 1 | # | 10 | Baseline | paralle | 36 |
| # | interaction | 2 | 3 | 0 | 0.4 | 1 | # | 10 | Baseline | paralle | 36 |
| # | disease | 4 | 1 | 0 | 0.8 | 1 | # | 10 | Baseline | ancom | 36 |
| # | sex | 2 | 0 | 0 | 1 | 1 | # | 10 | Baseline | ancom | 36 |
| # | time | 2 | 2 | 0 | 0.5 | 1 | # | 10 | Baseline | ancom | 36 |
| # | interaction | 0 | 0 | 2 | 0 | 0 | # | 10 | Baseline | ancom | 36 |
| # | disease | 4 | 13 | 0 | 0.235294118 | 1 | # | 10 | Baseline | aldex2 | 36 |
| # | sex | 2 | 6 | 0 | 0.25 | 1 | # | 10 | Baseline | aldex2 | 36 |
| # | time | 2 | 10 | 0 | 0.166666667 | 1 | # | 10 | Baseline | aldex2 | 36 |
| # | interaction | 0 | 0 | 2 | 0 | 0 | # | 10 | Baseline | aldex2 | 36 |
| # | disease | 4 | 0 | 0 | 1 | 1 | # | 10 | Small | paralle | 36 |
| # | sex | 2 | 0 | 0 | 1 | 1 | # | 10 | Small | paralle | 36 |
| # | time | 2 | 0 | 0 | 1 | 1 | # | 10 | Small | paralle | 36 |
| # | interaction | 2 | 0 | 0 | 1 | 1 | # | 10 | Small | paralle | 36 |
| # | disease | 4 | 1 | 0 | 0.8 | 1 | # | 10 | Small | ancom | 36 |
| # | sex | 2 | 0 | 0 | 1 | 1 | # | 10 | Small | ancom | 36 |
| # | time | 2 | 2 | 0 | 0.5 | 1 | # | 10 | Small | ancom | 36 |
| # | interaction | 0 | 0 | 2 | 0 | 0 | # | 10 | Small | ancom | 36 |
| # | disease | 4 | 1 | 0 | 0.8 | 1 | # | 10 | Small | aldex2 | 36 |
| # | sex | 2 | 0 | 0 | 1 | 1 | # | 10 | Small | aldex2 | 36 |
| # | time | 2 | 2 | 0 | 0.5 | 1 | # | 10 | Small | aldex2 | 36 |
| # | interaction | 0 | 0 | 2 | 0 | 0 | # | 10 | Small | aldex2 | 36 |
| # | disease | 4 | 2 | 0 | 0.666666667 | 1 | # | 10 | Baseline | paralle | 36 |
| # | sex | 2 | 4 | 0 | 0.333333333 | 1 | # | 10 | Baseline | paralle | 36 |
| # | time | 2 | 2 | 0 | 0.5 | 1 | # | 10 | Baseline | paralle | 36 |
| # | interaction | 2 | 3 | 0 | 0.4 | 1 | # | 10 | Baseline | paralle | 36 |
| # | disease | 4 | 2 | 0 | 0.666666667 | 1 | # | 10 | Baseline | ancom | 36 |
| # | sex | 2 | 0 | 0 | 1 | 1 | # | 10 | Baseline | ancom | 36 |
| # | time | 2 | 2 | 0 | 0.5 | 1 | # | 10 | Baseline | ancom | 36 |
| # | interaction | 0 | 0 | 2 | 0 | 0 | # | 10 | Baseline | ancom | 36 |
| # | disease | 4 | 17 | 0 | 0.19047619 | 1 | # | 10 | Baseline | aldex2 | 36 |
| # | sex | 2 | 5 | 0 | 0.285714286 | 1 | # | 10 | Baseline | aldex2 | 36 |
| # | time | 2 | 14 | 0 | 0.125 | 1 | # | 10 | Baseline | aldex2 | 36 |
| # | interaction | 0 | 0 | 2 | 0 | 0 | # | 10 | Baseline | aldex2 | 36 |
| # | disease | 4 | 0 | 0 | 1 | 1 | # | 10 | Small | paralle | 36 |
| # | sex | 2 | 0 | 0 | 1 | 1 | # | 10 | Small | paralle | 36 |
| # | time | 2 | 0 | 0 | 1 | 1 | # | 10 | Small | paralle | 36 |
| # | interaction | 2 | 0 | 0 | 1 | 1 | # | 10 | Small | paralle | 36 |
| # | disease | 4 | 2 | 0 | 0.666666667 | 1 | # | 10 | Small | ancom | 36 |
| # | sex | 2 | 0 | 0 | 1 | 1 | # | 10 | Small | ancom | 36 |
| # | time | 2 | 2 | 0 | 0.5 | 1 | # | 10 | Small | ancom | 36 |
| # | interaction | 0 | 0 | 2 | 0 | 0 | # | 10 | Small | ancom | 36 |
| # | disease | 4 | 2 | 0 | 0.666666667 | 1 | # | 10 | Small | aldex2 | 36 |
| # | sex | 2 | 0 | 0 | 1 | 1 | # | 10 | Small | aldex2 | 36 |
| # | time | 2 | 2 | 0 | 0.5 | 1 | # | 10 | Small | aldex2 | 36 |
| # | interaction | 0 | 0 | 2 | 0 | 0 | # | 10 | Small | aldex2 | 36 |
| # | disease | 4 | 4 | 0 | 0.5 | 1 | # | 10 | Baseline | paralle | 36 |
| # | sex | 2 | 3 | 0 | 0.4 | 1 | # | 10 | Baseline | paralle | 36 |
| # | time | 2 | 1 | 0 | 0.666666667 | 1 | # | 10 | Baseline | paralle | 36 |
| # | interaction | 2 | 1 | 0 | 0.666666667 | 1 | # | 10 | Baseline | paralle | 36 |
| # | disease | 4 | 2 | 0 | 0.666666667 | 1 | # | 10 | Baseline | ancom | 36 |
| # | sex | 2 | 0 | 0 | 1 | 1 | # | 10 | Baseline | ancom | 36 |
| # | time | 2 | 2 | 0 | 0.5 | 1 | # | 10 | Baseline | ancom | 36 |
| # | interaction | 0 | 0 | 2 | 0 | 0 | # | 10 | Baseline | ancom | 36 |
| # | disease | 4 | 28 | 0 | 0.125 | 1 | # | 10 | Baseline | aldex2 | 36 |
| # | sex | 2 | 6 | 0 | 0.25 | 1 | # | 10 | Baseline | aldex2 | 36 |
| # | time | 2 | 12 | 0 | 0.142857143 | 1 | # | 10 | Baseline | aldex2 | 36 |
| # | interaction | 0 | 0 | 2 | 0 | 0 | # | 10 | Baseline | aldex2 | 36 |
| # | disease | 4 | 0 | 0 | 1 | 1 | # | 10 | Small | paralle | 36 |
| # | sex | 2 | 0 | 0 | 1 | 1 | # | 10 | Small | paralle | 36 |
| # | time | 2 | 0 | 0 | 1 | 1 | # | 10 | Small | paralle | 36 |
| # | interaction | 2 | 0 | 0 | 1 | 1 | # | 10 | Small | paralle | 36 |
| # | disease | 4 | 2 | 0 | 0.666666667 | 1 | # | 10 | Small | ancom | 36 |
| # | sex | 2 | 0 | 0 | 1 | 1 | # | 10 | Small | ancom | 36 |
| # | time | 2 | 2 | 0 | 0.5 | 1 | # | 10 | Small | ancom | 36 |
| # | interaction | 0 | 0 | 2 | 0 | 0 | # | 10 | Small | ancom | 36 |
| # | disease | 4 | 2 | 0 | 0.666666667 | 1 | # | 10 | Small | aldex2 | 36 |
| # | sex | 2 | 0 | 0 | 1 | 1 | # | 10 | Small | aldex2 | 36 |
| # | time | 2 | 2 | 0 | 0.5 | 1 | # | 10 | Small | aldex2 | 36 |
| # | interaction | 0 | 0 | 2 | 0 | 0 | # | 10 | Small | aldex2 | 36 |
| # | disease | 7 | 6 | 0 | 0.538461538 | 1 | # | 20 | Baseline | paralle | 36 |
| # | sex | 4 | 14 | 0 | 0.222222222 | 1 | # | 20 | Baseline | paralle | 36 |
| # | time | 5 | 2 | 0 | 0.714285714 | 1 | # | 20 | Baseline | paralle | 36 |
| # | interaction | 2 | 5 | 2 | 0.285714286 | 0.5 | # | 20 | Baseline | paralle | 36 |
| # | disease | 7 | 0 | 0 | 1 | 1 | # | 20 | Baseline | ancom | 36 |
| # | sex | 4 | 0 | 0 | 1 | 1 | # | 20 | Baseline | ancom | 36 |
| # | time | 5 | 0 | 0 | 1 | 1 | # | 20 | Baseline | ancom | 36 |
| # | interaction | 0 | 0 | 4 | 0 | 0 | # | 20 | Baseline | ancom | 36 |
| # | disease | 7 | 11 | 0 | 0.388888889 | 1 | # | 20 | Baseline | aldex2 | 36 |
| # | sex | 4 | 6 | 0 | 0.4 | 1 | # | 20 | Baseline | aldex2 | 36 |
| # | time | 5 | 8 | 0 | 0.384615385 | 1 | # | 20 | Baseline | aldex2 | 36 |
| # | interaction | 0 | 0 | 4 | 0 | 0 | # | 20 | Baseline | aldex2 | 36 |
| # | disease | 7 | 0 | 0 | 1 | 1 | # | 20 | Small | paralle | 36 |
| # | sex | 4 | 0 | 0 | 1 | 1 | # | 20 | Small | paralle | 36 |
| # | time | 5 | 0 | 0 | 1 | 1 | # | 20 | Small | paralle | 36 |
| # | interaction | 1 | 0 | 3 | 1 | 0.25 | # | 20 | Small | paralle | 36 |
| # | disease | 7 | 0 | 0 | 1 | 1 | # | 20 | Small | ancom | 36 |
| # | sex | 4 | 0 | 0 | 1 | 1 | # | 20 | Small | ancom | 36 |
| # | time | 5 | 0 | 0 | 1 | 1 | # | 20 | Small | ancom | 36 |
| # | interaction | 0 | 0 | 4 | 0 | 0 | # | 20 | Small | ancom | 36 |
| # | disease | 7 | 0 | 0 | 1 | 1 | # | 20 | Small | aldex2 | 36 |
| # | sex | 4 | 0 | 0 | 1 | 1 | # | 20 | Small | aldex2 | 36 |
| # | time | 5 | 0 | 0 | 1 | 1 | # | 20 | Small | aldex2 | 36 |
| # | interaction | 0 | 0 | 4 | 0 | 0 | # | 20 | Small | aldex2 | 36 |
| # | disease | 7 | 11 | 0 | 0.388888889 | 1 | # | 20 | Baseline | paralle | 36 |
| # | sex | 4 | 8 | 0 | 0.333333333 | 1 | # | 20 | Baseline | paralle | 36 |
| # | time | 5 | 4 | 0 | 0.555555556 | 1 | # | 20 | Baseline | paralle | 36 |
| # | interaction | 4 | 5 | 0 | 0.444444444 | 1 | # | 20 | Baseline | paralle | 36 |
| # | disease | 7 | 4 | 0 | 0.636363636 | 1 | # | 20 | Baseline | ancom | 36 |
| # | sex | 4 | 0 | 0 | 1 | 1 | # | 20 | Baseline | ancom | 36 |
| # | time | 5 | 3 | 0 | 0.625 | 1 | # | 20 | Baseline | ancom | 36 |
| # | interaction | 0 | 0 | 4 | 0 | 0 | # | 20 | Baseline | ancom | 36 |
| # | disease | 7 | 28 | 0 | 0.2 | 1 | # | 20 | Baseline | aldex2 | 36 |
| # | sex | 4 | 9 | 0 | 0.307692308 | 1 | # | 20 | Baseline | aldex2 | 36 |
| # | time | 5 | 17 | 0 | 0.227272727 | 1 | # | 20 | Baseline | aldex2 | 36 |
| # | interaction | 0 | 0 | 4 | 0 | 0 | # | 20 | Baseline | aldex2 | 36 |
| # | disease | 7 | 0 | 0 | 1 | 1 | # | 20 | Small | paralle | 36 |
| # | sex | 4 | 0 | 0 | 1 | 1 | # | 20 | Small | paralle | 36 |
| # | time | 5 | 0 | 0 | 1 | 1 | # | 20 | Small | paralle | 36 |
| # | interaction | 2 | 0 | 2 | 1 | 0.5 | # | 20 | Small | paralle | 36 |
| # | disease | 7 | 3 | 0 | 0.7 | 1 | # | 20 | Small | ancom | 36 |
| # | sex | 4 | 0 | 0 | 1 | 1 | # | 20 | Small | ancom | 36 |
| # | time | 5 | 3 | 0 | 0.625 | 1 | # | 20 | Small | ancom | 36 |
| # | interaction | 0 | 0 | 4 | 0 | 0 | # | 20 | Small | ancom | 36 |
| # | disease | 7 | 2 | 0 | 0.777777778 | 1 | # | 20 | Small | aldex2 | 36 |
| # | sex | 4 | 0 | 0 | 1 | 1 | # | 20 | Small | aldex2 | 36 |
| # | time | 5 | 3 | 0 | 0.625 | 1 | # | 20 | Small | aldex2 | 36 |
| # | interaction | 0 | 0 | 4 | 0 | 0 | # | 20 | Small | aldex2 | 36 |
| # | disease | 7 | 10 | 0 | 0.411764706 | 1 | # | 20 | Baseline | paralle | 36 |
| # | sex | 4 | 8 | 0 | 0.333333333 | 1 | # | 20 | Baseline | paralle | 36 |
| # | time | 5 | 2 | 0 | 0.714285714 | 1 | # | 20 | Baseline | paralle | 36 |
| # | interaction | 4 | 2 | 0 | 0.666666667 | 1 | # | 20 | Baseline | paralle | 36 |
| # | disease | 7 | 3 | 0 | 0.7 | 1 | # | 20 | Baseline | ancom | 36 |
| # | sex | 4 | 0 | 0 | 1 | 1 | # | 20 | Baseline | ancom | 36 |
| # | time | 5 | 4 | 0 | 0.555555556 | 1 | # | 20 | Baseline | ancom | 36 |
| # | interaction | 0 | 0 | 4 | 0 | 0 | # | 20 | Baseline | ancom | 36 |
| # | disease | 7 | 42 | 0 | 0.142857143 | 1 | # | 20 | Baseline | aldex2 | 36 |
| # | sex | 4 | 12 | 0 | 0.25 | 1 | # | 20 | Baseline | aldex2 | 36 |
| # | time | 5 | 25 | 0 | 0.166666667 | 1 | # | 20 | Baseline | aldex2 | 36 |
| # | interaction | 0 | 0 | 4 | 0 | 0 | # | 20 | Baseline | aldex2 | 36 |
| # | disease | 7 | 0 | 0 | 1 | 1 | # | 20 | Small | paralle | 36 |
| # | sex | 4 | 0 | 0 | 1 | 1 | # | 20 | Small | paralle | 36 |
| # | time | 5 | 0 | 0 | 1 | 1 | # | 20 | Small | paralle | 36 |
| # | interaction | 3 | 0 | 1 | 1 | 0.75 | # | 20 | Small | paralle | 36 |
| # | disease | 7 | 3 | 0 | 0.7 | 1 | # | 20 | Small | ancom | 36 |
| # | sex | 4 | 0 | 0 | 1 | 1 | # | 20 | Small | ancom | 36 |
| # | time | 5 | 4 | 0 | 0.555555556 | 1 | # | 20 | Small | ancom | 36 |
| # | interaction | 0 | 0 | 4 | 0 | 0 | # | 20 | Small | ancom | 36 |
| # | disease | 7 | 5 | 0 | 0.583333333 | 1 | # | 20 | Small | aldex2 | 36 |
| # | sex | 4 | 0 | 0 | 1 | 1 | # | 20 | Small | aldex2 | 36 |
| # | time | 5 | 3 | 0 | 0.625 | 1 | # | 20 | Small | aldex2 | 36 |
| # | interaction | 0 | 0 | 4 | 0 | 0 | # | 20 | Small | aldex2 | 36 |
| # | disease | 7 | 15 | 0 | 0.318181818 | 1 | # | 20 | Baseline | paralle | 36 |
| # | sex | 4 | 5 | 0 | 0.444444444 | 1 | # | 20 | Baseline | paralle | 36 |
| # | time | 5 | 4 | 0 | 0.555555556 | 1 | # | 20 | Baseline | paralle | 36 |
| # | interaction | 4 | 4 | 0 | 0.5 | 1 | # | 20 | Baseline | paralle | 36 |
| # | disease | 7 | 4 | 0 | 0.636363636 | 1 | # | 20 | Baseline | ancom | 36 |
| # | sex | 4 | 0 | 0 | 1 | 1 | # | 20 | Baseline | ancom | 36 |
| # | time | 5 | 4 | 0 | 0.555555556 | 1 | # | 20 | Baseline | ancom | 36 |
| # | interaction | 0 | 0 | 4 | 0 | 0 | # | 20 | Baseline | ancom | 36 |
| # | disease | 7 | 55 | 0 | 0.112903226 | 1 | # | 20 | Baseline | aldex2 | 36 |
| # | sex | 4 | 12 | 0 | 0.25 | 1 | # | 20 | Baseline | aldex2 | 36 |
| # | time | 5 | 31 | 0 | 0.138888889 | 1 | # | 20 | Baseline | aldex2 | 36 |
| # | interaction | 0 | 0 | 4 | 0 | 0 | # | 20 | Baseline | aldex2 | 36 |
| # | disease | 7 | 0 | 0 | 1 | 1 | # | 20 | Small | paralle | 36 |
| # | sex | 4 | 0 | 0 | 1 | 1 | # | 20 | Small | paralle | 36 |
| # | time | 5 | 0 | 0 | 1 | 1 | # | 20 | Small | paralle | 36 |
| # | interaction | 4 | 0 | 0 | 1 | 1 | # | 20 | Small | paralle | 36 |
| # | disease | 7 | 4 | 0 | 0.636363636 | 1 | # | 20 | Small | ancom | 36 |
| # | sex | 4 | 0 | 0 | 1 | 1 | # | 20 | Small | ancom | 36 |
| # | time | 5 | 4 | 0 | 0.555555556 | 1 | # | 20 | Small | ancom | 36 |
| # | interaction | 0 | 0 | 4 | 0 | 0 | # | 20 | Small | ancom | 36 |
| # | disease | 7 | 8 | 0 | 0.466666667 | 1 | # | 20 | Small | aldex2 | 36 |
| # | sex | 4 | 0 | 0 | 1 | 1 | # | 20 | Small | aldex2 | 36 |
| # | time | 5 | 5 | 0 | 0.5 | 1 | # | 20 | Small | aldex2 | 36 |
| # | interaction | 0 | 0 | 4 | 0 | 0 | # | 20 | Small | aldex2 | 36 |
| # | disease | 4 | 3 | 0 | 0.571428571 | 1 | # | 10 | Baseline | paralle | 37 |
| # | sex | 2 | 2 | 0 | 0.5 | 1 | # | 10 | Baseline | paralle | 37 |
| # | time | 2 | 5 | 0 | 0.285714286 | 1 | # | 10 | Baseline | paralle | 37 |
| # | interaction | 2 | 5 | 0 | 0.285714286 | 1 | # | 10 | Baseline | paralle | 37 |
| # | disease | 4 | 1 | 0 | 0.8 | 1 | # | 10 | Baseline | ancom | 37 |
| # | sex | 2 | 0 | 0 | 1 | 1 | # | 10 | Baseline | ancom | 37 |
| # | time | 2 | 0 | 0 | 1 | 1 | # | 10 | Baseline | ancom | 37 |
| # | interaction | 0 | 0 | 2 | 0 | 0 | # | 10 | Baseline | ancom | 37 |
| # | disease | 4 | 5 | 0 | 0.444444444 | 1 | # | 10 | Baseline | aldex2 | 37 |
| # | sex | 2 | 4 | 0 | 0.333333333 | 1 | # | 10 | Baseline | aldex2 | 37 |
| # | time | 2 | 4 | 0 | 0.333333333 | 1 | # | 10 | Baseline | aldex2 | 37 |
| # | interaction | 0 | 0 | 2 | 0 | 0 | # | 10 | Baseline | aldex2 | 37 |
| # | disease | 4 | 0 | 0 | 1 | 1 | # | 10 | Small | paralle | 37 |
| # | sex | 2 | 0 | 0 | 1 | 1 | # | 10 | Small | paralle | 37 |
| # | time | 2 | 0 | 0 | 1 | 1 | # | 10 | Small | paralle | 37 |
| # | interaction | 2 | 0 | 0 | 1 | 1 | # | 10 | Small | paralle | 37 |
| # | disease | 4 | 1 | 0 | 0.8 | 1 | # | 10 | Small | ancom | 37 |
| # | sex | 2 | 0 | 0 | 1 | 1 | # | 10 | Small | ancom | 37 |
| # | time | 2 | 0 | 0 | 1 | 1 | # | 10 | Small | ancom | 37 |
| # | interaction | 0 | 0 | 2 | 0 | 0 | # | 10 | Small | ancom | 37 |
| # | disease | 4 | 1 | 0 | 0.8 | 1 | # | 10 | Small | aldex2 | 37 |
| # | sex | 2 | 0 | 0 | 1 | 1 | # | 10 | Small | aldex2 | 37 |
| # | time | 2 | 0 | 0 | 1 | 1 | # | 10 | Small | aldex2 | 37 |
| # | interaction | 0 | 0 | 2 | 0 | 0 | # | 10 | Small | aldex2 | 37 |
| # | disease | 4 | 5 | 0 | 0.444444444 | 1 | # | 10 | Baseline | paralle | 37 |
| # | sex | 2 | 6 | 0 | 0.25 | 1 | # | 10 | Baseline | paralle | 37 |
| # | time | 2 | 6 | 0 | 0.25 | 1 | # | 10 | Baseline | paralle | 37 |
| # | interaction | 2 | 7 | 0 | 0.222222222 | 1 | # | 10 | Baseline | paralle | 37 |
| # | disease | 4 | 2 | 0 | 0.666666667 | 1 | # | 10 | Baseline | ancom | 37 |
| # | sex | 2 | 0 | 0 | 1 | 1 | # | 10 | Baseline | ancom | 37 |
| # | time | 2 | 1 | 0 | 0.666666667 | 1 | # | 10 | Baseline | ancom | 37 |
| # | interaction | 0 | 0 | 2 | 0 | 0 | # | 10 | Baseline | ancom | 37 |
| # | disease | 4 | 11 | 0 | 0.266666667 | 1 | # | 10 | Baseline | aldex2 | 37 |
| # | sex | 2 | 4 | 0 | 0.333333333 | 1 | # | 10 | Baseline | aldex2 | 37 |
| # | time | 2 | 9 | 0 | 0.181818182 | 1 | # | 10 | Baseline | aldex2 | 37 |
| # | interaction | 0 | 0 | 2 | 0 | 0 | # | 10 | Baseline | aldex2 | 37 |
| # | disease | 4 | 0 | 0 | 1 | 1 | # | 10 | Small | paralle | 37 |
| # | sex | 2 | 0 | 0 | 1 | 1 | # | 10 | Small | paralle | 37 |
| # | time | 2 | 0 | 0 | 1 | 1 | # | 10 | Small | paralle | 37 |
| # | interaction | 2 | 0 | 0 | 1 | 1 | # | 10 | Small | paralle | 37 |
| # | disease | 4 | 2 | 0 | 0.666666667 | 1 | # | 10 | Small | ancom | 37 |
| # | sex | 2 | 0 | 0 | 1 | 1 | # | 10 | Small | ancom | 37 |
| # | time | 2 | 1 | 0 | 0.666666667 | 1 | # | 10 | Small | ancom | 37 |
| # | interaction | 0 | 0 | 2 | 0 | 0 | # | 10 | Small | ancom | 37 |
| # | disease | 4 | 2 | 0 | 0.666666667 | 1 | # | 10 | Small | aldex2 | 37 |
| # | sex | 2 | 0 | 0 | 1 | 1 | # | 10 | Small | aldex2 | 37 |
| # | time | 2 | 1 | 0 | 0.666666667 | 1 | # | 10 | Small | aldex2 | 37 |
| # | interaction | 0 | 0 | 2 | 0 | 0 | # | 10 | Small | aldex2 | 37 |
| # | disease | 4 | 7 | 0 | 0.363636364 | 1 | # | 10 | Baseline | paralle | 37 |
| # | sex | 2 | 3 | 0 | 0.4 | 1 | # | 10 | Baseline | paralle | 37 |
| # | time | 2 | 9 | 0 | 0.181818182 | 1 | # | 10 | Baseline | paralle | 37 |
| # | interaction | 2 | 6 | 0 | 0.25 | 1 | # | 10 | Baseline | paralle | 37 |
| # | disease | 4 | 2 | 0 | 0.666666667 | 1 | # | 10 | Baseline | ancom | 37 |
| # | sex | 2 | 0 | 0 | 1 | 1 | # | 10 | Baseline | ancom | 37 |
| # | time | 2 | 1 | 0 | 0.666666667 | 1 | # | 10 | Baseline | ancom | 37 |
| # | interaction | 0 | 0 | 2 | 0 | 0 | # | 10 | Baseline | ancom | 37 |
| # | disease | 4 | 16 | 0 | 0.2 | 1 | # | 10 | Baseline | aldex2 | 37 |
| # | sex | 2 | 3 | 0 | 0.4 | 1 | # | 10 | Baseline | aldex2 | 37 |
| # | time | 2 | 6 | 0 | 0.25 | 1 | # | 10 | Baseline | aldex2 | 37 |
| # | interaction | 0 | 0 | 2 | 0 | 0 | # | 10 | Baseline | aldex2 | 37 |
| # | disease | 4 | 0 | 0 | 1 | 1 | # | 10 | Small | paralle | 37 |
| # | sex | 2 | 0 | 0 | 1 | 1 | # | 10 | Small | paralle | 37 |
| # | time | 2 | 0 | 0 | 1 | 1 | # | 10 | Small | paralle | 37 |
| # | interaction | 2 | 1 | 0 | 0.666666667 | 1 | # | 10 | Small | paralle | 37 |
| # | disease | 4 | 2 | 0 | 0.666666667 | 1 | # | 10 | Small | ancom | 37 |
| # | sex | 2 | 0 | 0 | 1 | 1 | # | 10 | Small | ancom | 37 |
| # | time | 2 | 1 | 0 | 0.666666667 | 1 | # | 10 | Small | ancom | 37 |
| # | interaction | 0 | 0 | 2 | 0 | 0 | # | 10 | Small | ancom | 37 |
| # | disease | 4 | 2 | 0 | 0.666666667 | 1 | # | 10 | Small | aldex2 | 37 |
| # | sex | 2 | 0 | 0 | 1 | 1 | # | 10 | Small | aldex2 | 37 |
| # | time | 2 | 1 | 0 | 0.666666667 | 1 | # | 10 | Small | aldex2 | 37 |
| # | interaction | 0 | 0 | 2 | 0 | 0 | # | 10 | Small | aldex2 | 37 |
| # | disease | 4 | 7 | 0 | 0.363636364 | 1 | # | 10 | Baseline | paralle | 37 |
| # | sex | 2 | 2 | 0 | 0.5 | 1 | # | 10 | Baseline | paralle | 37 |
| # | time | 2 | 7 | 0 | 0.222222222 | 1 | # | 10 | Baseline | paralle | 37 |
| # | interaction | 2 | 6 | 0 | 0.25 | 1 | # | 10 | Baseline | paralle | 37 |
| # | disease | 4 | 2 | 0 | 0.666666667 | 1 | # | 10 | Baseline | ancom | 37 |
| # | sex | 2 | 0 | 0 | 1 | 1 | # | 10 | Baseline | ancom | 37 |
| # | time | 2 | 2 | 0 | 0.5 | 1 | # | 10 | Baseline | ancom | 37 |
| # | interaction | 0 | 0 | 2 | 0 | 0 | # | 10 | Baseline | ancom | 37 |
| # | disease | 4 | 19 | 0 | 0.173913043 | 1 | # | 10 | Baseline | aldex2 | 37 |
| # | sex | 2 | 8 | 0 | 0.2 | 1 | # | 10 | Baseline | aldex2 | 37 |
| # | time | 2 | 8 | 0 | 0.2 | 1 | # | 10 | Baseline | aldex2 | 37 |
| # | interaction | 0 | 0 | 2 | 0 | 0 | # | 10 | Baseline | aldex2 | 37 |
| # | disease | 4 | 0 | 0 | 1 | 1 | # | 10 | Small | paralle | 37 |
| # | sex | 2 | 0 | 0 | 1 | 1 | # | 10 | Small | paralle | 37 |
| # | time | 2 | 0 | 0 | 1 | 1 | # | 10 | Small | paralle | 37 |
| # | interaction | 2 | 0 | 0 | 1 | 1 | # | 10 | Small | paralle | 37 |
| # | disease | 4 | 2 | 0 | 0.666666667 | 1 | # | 10 | Small | ancom | 37 |
| # | sex | 2 | 0 | 0 | 1 | 1 | # | 10 | Small | ancom | 37 |
| # | time | 2 | 2 | 0 | 0.5 | 1 | # | 10 | Small | ancom | 37 |
| # | interaction | 0 | 0 | 2 | 0 | 0 | # | 10 | Small | ancom | 37 |
| # | disease | 4 | 3 | 0 | 0.571428571 | 1 | # | 10 | Small | aldex2 | 37 |
| # | sex | 2 | 0 | 0 | 1 | 1 | # | 10 | Small | aldex2 | 37 |
| # | time | 2 | 2 | 0 | 0.5 | 1 | # | 10 | Small | aldex2 | 37 |
| # | interaction | 0 | 0 | 2 | 0 | 0 | # | 10 | Small | aldex2 | 37 |
| # | disease | 7 | 8 | 0 | 0.466666667 | 1 | # | 20 | Baseline | paralle | 37 |
| # | sex | 4 | 12 | 0 | 0.25 | 1 | # | 20 | Baseline | paralle | 37 |
| # | time | 5 | 7 | 0 | 0.416666667 | 1 | # | 20 | Baseline | paralle | 37 |
| # | interaction | 4 | 11 | 0 | 0.266666667 | 1 | # | 20 | Baseline | paralle | 37 |
| # | disease | 7 | 3 | 0 | 0.7 | 1 | # | 20 | Baseline | ancom | 37 |
| # | sex | 4 | 0 | 0 | 1 | 1 | # | 20 | Baseline | ancom | 37 |
| # | time | 5 | 2 | 0 | 0.714285714 | 1 | # | 20 | Baseline | ancom | 37 |
| # | interaction | 0 | 0 | 4 | 0 | 0 | # | 20 | Baseline | ancom | 37 |
| # | disease | 7 | 25 | 0 | 0.21875 | 1 | # | 20 | Baseline | aldex2 | 37 |
| # | sex | 4 | 10 | 0 | 0.285714286 | 1 | # | 20 | Baseline | aldex2 | 37 |
| # | time | 5 | 13 | 0 | 0.277777778 | 1 | # | 20 | Baseline | aldex2 | 37 |
| # | interaction | 0 | 0 | 4 | 0 | 0 | # | 20 | Baseline | aldex2 | 37 |
| # | disease | 7 | 0 | 0 | 1 | 1 | # | 20 | Small | paralle | 37 |
| # | sex | 4 | 1 | 0 | 0.8 | 1 | # | 20 | Small | paralle | 37 |
| # | time | 5 | 1 | 0 | 0.833333333 | 1 | # | 20 | Small | paralle | 37 |
| # | interaction | 2 | 1 | 2 | 0.666666667 | 0.5 | # | 20 | Small | paralle | 37 |
| # | disease | 7 | 1 | 0 | 0.875 | 1 | # | 20 | Small | ancom | 37 |
| # | sex | 4 | 0 | 0 | 1 | 1 | # | 20 | Small | ancom | 37 |
| # | time | 5 | 2 | 0 | 0.714285714 | 1 | # | 20 | Small | ancom | 37 |
| # | interaction | 0 | 0 | 4 | 0 | 0 | # | 20 | Small | ancom | 37 |
| # | disease | 7 | 2 | 0 | 0.777777778 | 1 | # | 20 | Small | aldex2 | 37 |
| # | sex | 4 | 0 | 0 | 1 | 1 | # | 20 | Small | aldex2 | 37 |
| # | time | 5 | 2 | 0 | 0.714285714 | 1 | # | 20 | Small | aldex2 | 37 |
| # | interaction | 0 | 0 | 4 | 0 | 0 | # | 20 | Small | aldex2 | 37 |
| # | disease | 7 | 12 | 0 | 0.368421053 | 1 | # | 20 | Baseline | paralle | 37 |
| # | sex | 4 | 12 | 0 | 0.25 | 1 | # | 20 | Baseline | paralle | 37 |
| # | time | 5 | 10 | 0 | 0.333333333 | 1 | # | 20 | Baseline | paralle | 37 |
| # | interaction | 4 | 6 | 0 | 0.4 | 1 | # | 20 | Baseline | paralle | 37 |
| # | disease | 7 | 5 | 0 | 0.583333333 | 1 | # | 20 | Baseline | ancom | 37 |
| # | sex | 4 | 0 | 0 | 1 | 1 | # | 20 | Baseline | ancom | 37 |
| # | time | 5 | 3 | 0 | 0.625 | 1 | # | 20 | Baseline | ancom | 37 |
| # | interaction | 0 | 0 | 4 | 0 | 0 | # | 20 | Baseline | ancom | 37 |
| # | disease | 7 | 38 | 0 | 0.155555556 | 1 | # | 20 | Baseline | aldex2 | 37 |
| # | sex | 4 | 13 | 0 | 0.235294118 | 1 | # | 20 | Baseline | aldex2 | 37 |
| # | time | 5 | 24 | 0 | 0.172413793 | 1 | # | 20 | Baseline | aldex2 | 37 |
| # | interaction | 0 | 0 | 4 | 0 | 0 | # | 20 | Baseline | aldex2 | 37 |
| # | disease | 7 | 0 | 0 | 1 | 1 | # | 20 | Small | paralle | 37 |
| # | sex | 4 | 0 | 0 | 1 | 1 | # | 20 | Small | paralle | 37 |
| # | time | 5 | 0 | 0 | 1 | 1 | # | 20 | Small | paralle | 37 |
| # | interaction | 3 | 0 | 1 | 1 | 0.75 | # | 20 | Small | paralle | 37 |
| # | disease | 7 | 4 | 0 | 0.636363636 | 1 | # | 20 | Small | ancom | 37 |
| # | sex | 4 | 0 | 0 | 1 | 1 | # | 20 | Small | ancom | 37 |
| # | time | 5 | 3 | 0 | 0.625 | 1 | # | 20 | Small | ancom | 37 |
| # | interaction | 0 | 0 | 4 | 0 | 0 | # | 20 | Small | ancom | 37 |
| # | disease | 7 | 6 | 0 | 0.538461538 | 1 | # | 20 | Small | aldex2 | 37 |
| # | sex | 4 | 1 | 0 | 0.8 | 1 | # | 20 | Small | aldex2 | 37 |
| # | time | 5 | 3 | 0 | 0.625 | 1 | # | 20 | Small | aldex2 | 37 |
| # | interaction | 0 | 0 | 4 | 0 | 0 | # | 20 | Small | aldex2 | 37 |
| # | disease | 7 | 14 | 0 | 0.333333333 | 1 | # | 20 | Baseline | paralle | 37 |
| # | sex | 4 | 4 | 0 | 0.5 | 1 | # | 20 | Baseline | paralle | 37 |
| # | time | 5 | 10 | 0 | 0.333333333 | 1 | # | 20 | Baseline | paralle | 37 |
| # | interaction | 4 | 6 | 0 | 0.4 | 1 | # | 20 | Baseline | paralle | 37 |
| # | disease | 7 | 4 | 0 | 0.636363636 | 1 | # | 20 | Baseline | ancom | 37 |
| # | sex | 4 | 0 | 0 | 1 | 1 | # | 20 | Baseline | ancom | 37 |
| # | time | 5 | 4 | 0 | 0.555555556 | 1 | # | 20 | Baseline | ancom | 37 |
| # | interaction | 0 | 0 | 4 | 0 | 0 | # | 20 | Baseline | ancom | 37 |
| # | disease | 7 | 52 | 0 | 0.118644068 | 1 | # | 20 | Baseline | aldex2 | 37 |
| # | sex | 4 | 17 | 0 | 0.19047619 | 1 | # | 20 | Baseline | aldex2 | 37 |
| # | time | 5 | 34 | 0 | 0.128205128 | 1 | # | 20 | Baseline | aldex2 | 37 |
| # | interaction | 0 | 0 | 4 | 0 | 0 | # | 20 | Baseline | aldex2 | 37 |
| # | disease | 7 | 0 | 0 | 1 | 1 | # | 20 | Small | paralle | 37 |
| # | sex | 4 | 0 | 0 | 1 | 1 | # | 20 | Small | paralle | 37 |
| # | time | 5 | 1 | 0 | 0.833333333 | 1 | # | 20 | Small | paralle | 37 |
| # | interaction | 3 | 1 | 1 | 0.75 | 0.75 | # | 20 | Small | paralle | 37 |
| # | disease | 7 | 4 | 0 | 0.636363636 | 1 | # | 20 | Small | ancom | 37 |
| # | sex | 4 | 0 | 0 | 1 | 1 | # | 20 | Small | ancom | 37 |

| | | | | | | | | | | | |
|---|---|---|---|---|---|---|---|---|---|---|---|
| # | time | 5 | 4 | 0 | 0.555555556 | 1 | # | 20 | Small | ancom | 37 |
| # | interaction | 0 | 0 | 4 | 0 | 0 | # | 20 | Small | ancom | 37 |
| # | disease | 7 | 10 | 0 | 0.411764706 | 1 | # | 20 | Small | aldex2 | 37 |
| # | sex | 4 | 0 | 0 | 1 | 1 | # | 20 | Small | aldex2 | 37 |
| # | time | 5 | 5 | 0 | 0.5 | 1 | # | 20 | Small | aldex2 | 37 |
| # | interaction | 0 | 0 | 4 | 0 | 0 | # | 20 | Small | aldex2 | 37 |
| # | disease | 7 | 13 | 0 | 0.35 | 1 | # | 20 | Baseline | paralle | 37 |
| # | sex | 4 | 5 | 0 | 0.444444444 | 1 | # | 20 | Baseline | paralle | 37 |
| # | time | 5 | 6 | 0 | 0.454545455 | 1 | # | 20 | Baseline | paralle | 37 |
| # | interaction | 4 | 6 | 0 | 0.4 | 1 | # | 20 | Baseline | paralle | 37 |
| # | disease | 7 | 4 | 0 | 0.636363636 | 1 | # | 20 | Baseline | ancom | 37 |
| # | sex | 4 | 0 | 0 | 1 | 1 | # | 20 | Baseline | ancom | 37 |
| # | time | 5 | 4 | 0 | 0.555555556 | 1 | # | 20 | Baseline | ancom | 37 |
| # | interaction | 0 | 0 | 4 | 0 | 0 | # | 20 | Baseline | ancom | 37 |
| # | disease | 7 | 58 | 0 | 0.107692308 | 1 | # | 20 | Baseline | aldex2 | 37 |
| # | sex | 4 | 19 | 0 | 0.173913043 | 1 | # | 20 | Baseline | aldex2 | 37 |
| # | time | 5 | 41 | 0 | 0.108695652 | 1 | # | 20 | Baseline | aldex2 | 37 |
| # | interaction | 0 | 0 | 4 | 0 | 0 | # | 20 | Baseline | aldex2 | 37 |
| # | disease | 7 | 1 | 0 | 0.875 | 1 | # | 20 | Small | paralle | 37 |
| # | sex | 4 | 0 | 0 | 1 | 1 | # | 20 | Small | paralle | 37 |
| # | time | 5 | 0 | 0 | 1 | 1 | # | 20 | Small | paralle | 37 |
| # | interaction | 4 | 0 | 0 | 1 | 1 | # | 20 | Small | paralle | 37 |
| # | disease | 7 | 4 | 0 | 0.636363636 | 1 | # | 20 | Small | ancom | 37 |
| # | sex | 4 | 0 | 0 | 1 | 1 | # | 20 | Small | ancom | 37 |
| # | time | 5 | 4 | 0 | 0.555555556 | 1 | # | 20 | Small | ancom | 37 |
| # | interaction | 0 | 0 | 4 | 0 | 0 | # | 20 | Small | ancom | 37 |
| # | disease | 7 | 11 | 0 | 0.388888889 | 1 | # | 20 | Small | aldex2 | 37 |
| # | sex | 4 | 1 | 0 | 0.8 | 1 | # | 20 | Small | aldex2 | 37 |
| # | time | 5 | 6 | 0 | 0.454545455 | 1 | # | 20 | Small | aldex2 | 37 |
| # | interaction | 0 | 0 | 4 | 0 | 0 | # | 20 | Small | aldex2 | 37 |
| # | disease | 4 | 4 | 0 | 0.5 | 1 | # | 10 | Baseline | paralle | 38 |
| # | sex | 2 | 2 | 0 | 0.5 | 1 | # | 10 | Baseline | paralle | 38 |
| # | time | 2 | 8 | 0 | 0.2 | 1 | # | 10 | Baseline | paralle | 38 |
| # | interaction | 2 | 5 | 0 | 0.285714286 | 1 | # | 10 | Baseline | paralle | 38 |
| # | disease | 4 | 0 | 0 | 1 | 1 | # | 10 | Baseline | ancom | 38 |
| # | sex | 2 | 0 | 0 | 1 | 1 | # | 10 | Baseline | ancom | 38 |
| # | time | 2 | 1 | 0 | 0.666666667 | 1 | # | 10 | Baseline | ancom | 38 |
| # | interaction | 0 | 0 | 2 | 0 | 0 | # | 10 | Baseline | ancom | 38 |
| # | disease | 4 | 9 | 0 | 0.307692308 | 1 | # | 10 | Baseline | aldex2 | 38 |
| # | sex | 2 | 5 | 0 | 0.285714286 | 1 | # | 10 | Baseline | aldex2 | 38 |
| # | time | 2 | 4 | 0 | 0.333333333 | 1 | # | 10 | Baseline | aldex2 | 38 |
| # | interaction | 0 | 0 | 2 | 0 | 0 | # | 10 | Baseline | aldex2 | 38 |
| # | disease | 4 | 0 | 0 | 1 | 1 | # | 10 | Small | paralle | 38 |
| # | sex | 2 | 0 | 0 | 1 | 1 | # | 10 | Small | paralle | 38 |
| # | time | 2 | 0 | 0 | 1 | 1 | # | 10 | Small | paralle | 38 |
| # | interaction | 0 | 0 | 2 | 0 | 0 | # | 10 | Small | paralle | 38 |
| # | disease | 4 | 0 | 0 | 1 | 1 | # | 10 | Small | ancom | 38 |
| # | sex | 2 | 0 | 0 | 1 | 1 | # | 10 | Small | ancom | 38 |
| # | time | 2 | 1 | 0 | 0.666666667 | 1 | # | 10 | Small | ancom | 38 |
| # | interaction | 0 | 0 | 2 | 0 | 0 | # | 10 | Small | ancom | 38 |
| # | disease | 4 | 1 | 0 | 0.8 | 1 | # | 10 | Small | aldex2 | 38 |
| # | sex | 2 | 0 | 0 | 1 | 1 | # | 10 | Small | aldex2 | 38 |
| # | time | 2 | 1 | 0 | 0.666666667 | 1 | # | 10 | Small | aldex2 | 38 |
| # | interaction | 0 | 0 | 2 | 0 | 0 | # | 10 | Small | aldex2 | 38 |
| # | disease | 4 | 8 | 0 | 0.333333333 | 1 | # | 10 | Baseline | paralle | 38 |
| # | sex | 2 | 5 | 0 | 0.285714286 | 1 | # | 10 | Baseline | paralle | 38 |
| # | time | 2 | 5 | 0 | 0.285714286 | 1 | # | 10 | Baseline | paralle | 38 |
| # | interaction | 2 | 8 | 0 | 0.2 | 1 | # | 10 | Baseline | paralle | 38 |
| # | disease | 4 | 2 | 0 | 0.666666667 | 1 | # | 10 | Baseline | ancom | 38 |
| # | sex | 2 | 0 | 0 | 1 | 1 | # | 10 | Baseline | ancom | 38 |
| # | time | 2 | 2 | 0 | 0.5 | 1 | # | 10 | Baseline | ancom | 38 |
| # | interaction | 0 | 0 | 2 | 0 | 0 | # | 10 | Baseline | ancom | 38 |
| # | disease | 4 | 11 | 0 | 0.266666667 | 1 | # | 10 | Baseline | aldex2 | 38 |
| # | sex | 2 | 7 | 0 | 0.222222222 | 1 | # | 10 | Baseline | aldex2 | 38 |
| # | time | 2 | 9 | 0 | 0.181818182 | 1 | # | 10 | Baseline | aldex2 | 38 |
| # | interaction | 0 | 0 | 2 | 0 | 0 | # | 10 | Baseline | aldex2 | 38 |
| # | disease | 4 | 0 | 0 | 1 | 1 | # | 10 | Small | paralle | 38 |
| # | sex | 2 | 0 | 0 | 1 | 1 | # | 10 | Small | paralle | 38 |
| # | time | 2 | 0 | 0 | 1 | 1 | # | 10 | Small | paralle | 38 |
| # | interaction | 2 | 0 | 0 | 1 | 1 | # | 10 | Small | paralle | 38 |
| # | disease | 4 | 2 | 0 | 0.666666667 | 1 | # | 10 | Small | ancom | 38 |
| # | sex | 2 | 0 | 0 | 1 | 1 | # | 10 | Small | ancom | 38 |
| # | time | 2 | 1 | 0 | 0.666666667 | 1 | # | 10 | Small | ancom | 38 |
| # | interaction | 0 | 0 | 2 | 0 | 0 | # | 10 | Small | ancom | 38 |
| # | disease | 4 | 2 | 0 | 0.666666667 | 1 | # | 10 | Small | aldex2 | 38 |
| # | sex | 2 | 0 | 0 | 1 | 1 | # | 10 | Small | aldex2 | 38 |
| # | time | 2 | 2 | 0 | 0.5 | 1 | # | 10 | Small | aldex2 | 38 |
| # | interaction | 0 | 0 | 2 | 0 | 0 | # | 10 | Small | aldex2 | 38 |
| # | disease | 4 | 3 | 0 | 0.571428571 | 1 | # | 10 | Baseline | paralle | 38 |
| # | sex | 2 | 4 | 0 | 0.333333333 | 1 | # | 10 | Baseline | paralle | 38 |
| # | time | 2 | 2 | 0 | 0.5 | 1 | # | 10 | Baseline | paralle | 38 |
| # | interaction | 2 | 6 | 0 | 0.25 | 1 | # | 10 | Baseline | paralle | 38 |
| # | disease | 4 | 2 | 0 | 0.666666667 | 1 | # | 10 | Baseline | ancom | 38 |
| # | sex | 2 | 0 | 0 | 1 | 1 | # | 10 | Baseline | ancom | 38 |
| # | time | 2 | 2 | 0 | 0.5 | 1 | # | 10 | Baseline | ancom | 38 |
| # | interaction | 0 | 0 | 2 | 0 | 0 | # | 10 | Baseline | ancom | 38 |
| # | disease | 4 | 17 | 0 | 0.19047619 | 1 | # | 10 | Baseline | aldex2 | 38 |
| # | sex | 2 | 6 | 0 | 0.25 | 1 | # | 10 | Baseline | aldex2 | 38 |
| # | time | 2 | 9 | 0 | 0.181818182 | 1 | # | 10 | Baseline | aldex2 | 38 |
| # | interaction | 0 | 0 | 2 | 0 | 0 | # | 10 | Baseline | aldex2 | 38 |
| # | disease | 4 | 1 | 0 | 0.8 | 1 | # | 10 | Small | paralle | 38 |
| # | sex | 2 | 0 | 0 | 1 | 1 | # | 10 | Small | paralle | 38 |
| # | time | 2 | 0 | 0 | 1 | 1 | # | 10 | Small | paralle | 38 |
| # | interaction | 2 | 0 | 0 | 1 | 1 | # | 10 | Small | paralle | 38 |
| # | disease | 4 | 2 | 0 | 0.666666667 | 1 | # | 10 | Small | ancom | 38 |
| # | sex | 2 | 0 | 0 | 1 | 1 | # | 10 | Small | ancom | 38 |
| # | time | 2 | 2 | 0 | 0.5 | 1 | # | 10 | Small | ancom | 38 |
| # | interaction | 0 | 0 | 2 | 0 | 0 | # | 10 | Small | ancom | 38 |
| # | disease | 4 | 3 | 0 | 0.571428571 | 1 | # | 10 | Small | aldex2 | 38 |
| # | sex | 2 | 0 | 0 | 1 | 1 | # | 10 | Small | aldex2 | 38 |
| # | time | 2 | 2 | 0 | 0.5 | 1 | # | 10 | Small | aldex2 | 38 |
| # | interaction | 0 | 0 | 2 | 0 | 0 | # | 10 | Small | aldex2 | 38 |
| # | disease | 4 | 5 | 0 | 0.444444444 | 1 | # | 10 | Baseline | paralle | 38 |
| # | sex | 2 | 4 | 0 | 0.333333333 | 1 | # | 10 | Baseline | paralle | 38 |
| # | time | 2 | 1 | 0 | 0.666666667 | 1 | # | 10 | Baseline | paralle | 38 |
| # | interaction | 2 | 4 | 0 | 0.333333333 | 1 | # | 10 | Baseline | paralle | 38 |
| # | disease | 4 | 2 | 0 | 0.666666667 | 1 | # | 10 | Baseline | ancom | 38 |
| # | sex | 2 | 0 | 0 | 1 | 1 | # | 10 | Baseline | ancom | 38 |
| # | time | 2 | 2 | 0 | 0.5 | 1 | # | 10 | Baseline | ancom | 38 |
| # | interaction | 0 | 0 | 2 | 0 | 0 | # | 10 | Baseline | ancom | 38 |
| # | disease | 4 | 21 | 0 | 0.16 | 1 | # | 10 | Baseline | aldex2 | 38 |
| # | sex | 2 | 4 | 0 | 0.333333333 | 1 | # | 10 | Baseline | aldex2 | 38 |
| # | time | 2 | 13 | 0 | 0.133333333 | 1 | # | 10 | Baseline | aldex2 | 38 |
| # | interaction | 0 | 0 | 2 | 0 | 0 | # | 10 | Baseline | aldex2 | 38 |
| # | disease | 4 | 1 | 0 | 0.8 | 1 | # | 10 | Small | paralle | 38 |
| # | sex | 2 | 0 | 0 | 1 | 1 | # | 10 | Small | paralle | 38 |
| # | time | 2 | 0 | 0 | 1 | 1 | # | 10 | Small | paralle | 38 |
| # | interaction | 2 | 0 | 0 | 1 | 1 | # | 10 | Small | paralle | 38 |
| # | disease | 4 | 2 | 0 | 0.666666667 | 1 | # | 10 | Small | ancom | 38 |
| # | sex | 2 | 0 | 0 | 1 | 1 | # | 10 | Small | ancom | 38 |
| # | time | 2 | 2 | 0 | 0.5 | 1 | # | 10 | Small | ancom | 38 |
| # | interaction | 0 | 0 | 2 | 0 | 0 | # | 10 | Small | ancom | 38 |
| # | disease | 4 | 3 | 0 | 0.571428571 | 1 | # | 10 | Small | aldex2 | 38 |
| # | sex | 2 | 0 | 0 | 1 | 1 | # | 10 | Small | aldex2 | 38 |
| # | time | 2 | 2 | 0 | 0.5 | 1 | # | 10 | Small | aldex2 | 38 |
| # | interaction | 0 | 0 | 2 | 0 | 0 | # | 10 | Small | aldex2 | 38 |
| # | disease | 7 | 10 | 0 | 0.411764706 | 1 | # | 20 | Baseline | paralle | 38 |
| # | sex | 4 | 5 | 0 | 0.444444444 | 1 | # | 20 | Baseline | paralle | 38 |
| # | time | 5 | 6 | 0 | 0.454545455 | 1 | # | 20 | Baseline | paralle | 38 |
| # | interaction | 4 | 12 | 0 | 0.25 | 1 | # | 20 | Baseline | paralle | 38 |
| # | disease | 7 | 0 | 0 | 1 | 1 | # | 20 | Baseline | ancom | 38 |
| # | sex | 4 | 0 | 0 | 1 | 1 | # | 20 | Baseline | ancom | 38 |
| # | time | 5 | 0 | 0 | 1 | 1 | # | 20 | Baseline | ancom | 38 |
| # | interaction | 0 | 0 | 4 | 0 | 0 | # | 20 | Baseline | ancom | 38 |
| # | disease | 7 | 16 | 0 | 0.304347826 | 1 | # | 20 | Baseline | aldex2 | 38 |
| # | sex | 4 | 6 | 0 | 0.4 | 1 | # | 20 | Baseline | aldex2 | 38 |
| # | time | 5 | 15 | 0 | 0.25 | 1 | # | 20 | Baseline | aldex2 | 38 |
| # | interaction | 0 | 0 | 4 | 0 | 0 | # | 20 | Baseline | aldex2 | 38 |
| # | disease | 7 | 0 | 0 | 1 | 1 | # | 20 | Small | paralle | 38 |
| # | sex | 4 | 1 | 0 | 0.8 | 1 | # | 20 | Small | paralle | 38 |
| # | time | 5 | 1 | 0 | 0.833333333 | 1 | # | 20 | Small | paralle | 38 |
| # | interaction | 2 | 0 | 2 | 1 | 0.5 | # | 20 | Small | paralle | 38 |
| # | disease | 7 | 0 | 0 | 1 | 1 | # | 20 | Small | ancom | 38 |
| # | sex | 4 | 0 | 0 | 1 | 1 | # | 20 | Small | ancom | 38 |
| # | time | 5 | 0 | 0 | 1 | 1 | # | 20 | Small | ancom | 38 |
| # | interaction | 0 | 0 | 4 | 0 | 0 | # | 20 | Small | ancom | 38 |
| # | disease | 7 | 1 | 0 | 0.875 | 1 | # | 20 | Small | aldex2 | 38 |
| # | sex | 4 | 0 | 0 | 1 | 1 | # | 20 | Small | aldex2 | 38 |
| # | time | 5 | 0 | 0 | 1 | 1 | # | 20 | Small | aldex2 | 38 |
| # | interaction | 0 | 0 | 4 | 0 | 0 | # | 20 | Small | aldex2 | 38 |
| # | disease | 7 | 10 | 0 | 0.411764706 | 1 | # | 20 | Baseline | paralle | 38 |
| # | sex | 4 | 2 | 0 | 0.666666667 | 1 | # | 20 | Baseline | paralle | 38 |
| # | time | 5 | 7 | 0 | 0.416666667 | 1 | # | 20 | Baseline | paralle | 38 |
| # | interaction | 4 | 8 | 0 | 0.333333333 | 1 | # | 20 | Baseline | paralle | 38 |
| # | disease | 7 | 2 | 0 | 0.777777778 | 1 | # | 20 | Baseline | ancom | 38 |
| # | sex | 4 | 0 | 0 | 1 | 1 | # | 20 | Baseline | ancom | 38 |
| # | time | 5 | 2 | 0 | 0.714285714 | 1 | # | 20 | Baseline | ancom | 38 |
| # | interaction | 0 | 0 | 4 | 0 | 0 | # | 20 | Baseline | ancom | 38 |
| # | disease | 7 | 33 | 0 | 0.175 | 1 | # | 20 | Baseline | aldex2 | 38 |
| # | sex | 4 | 7 | 0 | 0.363636364 | 1 | # | 20 | Baseline | aldex2 | 38 |
| # | time | 5 | 23 | 0 | 0.178571429 | 1 | # | 20 | Baseline | aldex2 | 38 |
| # | interaction | 0 | 0 | 4 | 0 | 0 | # | 20 | Baseline | aldex2 | 38 |
| # | disease | 7 | 0 | 0 | 1 | 1 | # | 20 | Small | paralle | 38 |
| # | sex | 4 | 1 | 0 | 0.8 | 1 | # | 20 | Small | paralle | 38 |
| # | time | 5 | 0 | 0 | 1 | 1 | # | 20 | Small | paralle | 38 |
| # | interaction | 3 | 0 | 1 | 1 | 0.75 | # | 20 | Small | paralle | 38 |
| # | disease | 7 | 2 | 0 | 0.777777778 | 1 | # | 20 | Small | ancom | 38 |
| # | sex | 4 | 0 | 0 | 1 | 1 | # | 20 | Small | ancom | 38 |
| # | time | 5 | 2 | 0 | 0.714285714 | 1 | # | 20 | Small | ancom | 38 |
| # | interaction | 0 | 0 | 4 | 0 | 0 | # | 20 | Small | ancom | 38 |
| # | disease | 7 | 0 | 0 | 1 | 1 | # | 20 | Small | aldex2 | 38 |
| # | sex | 4 | 0 | 0 | 1 | 1 | # | 20 | Small | aldex2 | 38 |
| # | time | 5 | 1 | 0 | 0.833333333 | 1 | # | 20 | Small | aldex2 | 38 |
| # | interaction | 0 | 0 | 4 | 0 | 0 | # | 20 | Small | aldex2 | 38 |
| # | disease | 7 | 11 | 0 | 0.388888889 | 1 | # | 20 | Baseline | paralle | 38 |
| # | sex | 4 | 5 | 0 | 0.444444444 | 1 | # | 20 | Baseline | paralle | 38 |
| # | time | 5 | 10 | 0 | 0.333333333 | 1 | # | 20 | Baseline | paralle | 38 |
| # | interaction | 4 | 7 | 0 | 0.363636364 | 1 | # | 20 | Baseline | paralle | 38 |
| # | disease | 7 | 4 | 0 | 0.636363636 | 1 | # | 20 | Baseline | ancom | 38 |
| # | sex | 4 | 0 | 0 | 1 | 1 | # | 20 | Baseline | ancom | 38 |
| # | time | 5 | 3 | 0 | 0.625 | 1 | # | 20 | Baseline | ancom | 38 |
| # | interaction | 0 | 0 | 4 | 0 | 0 | # | 20 | Baseline | ancom | 38 |
| # | disease | 7 | 47 | 0 | 0.12962963 | 1 | # | 20 | Baseline | aldex2 | 38 |
| # | sex | 4 | 15 | 0 | 0.210526316 | 1 | # | 20 | Baseline | aldex2 | 38 |
| # | time | 5 | 36 | 0 | 0.12195122 | 1 | # | 20 | Baseline | aldex2 | 38 |
| # | interaction | 0 | 0 | 4 | 0 | 0 | # | 20 | Baseline | aldex2 | 38 |
| # | disease | 7 | 0 | 0 | 1 | 1 | # | 20 | Small | paralle | 38 |
| # | sex | 4 | 2 | 0 | 0.666666667 | 1 | # | 20 | Small | paralle | 38 |
| # | time | 5 | 0 | 0 | 1 | 1 | # | 20 | Small | paralle | 38 |
| # | interaction | 4 | 0 | 0 | 1 | 1 | # | 20 | Small | paralle | 38 |
| # | disease | 7 | 3 | 0 | 0.7 | 1 | # | 20 | Small | ancom | 38 |
| # | sex | 4 | 0 | 0 | 1 | 1 | # | 20 | Small | ancom | 38 |
| # | time | 5 | 3 | 0 | 0.625 | 1 | # | 20 | Small | ancom | 38 |
| # | interaction | 0 | 0 | 4 | 0 | 0 | # | 20 | Small | ancom | 38 |
| # | disease | 7 | 7 | 0 | 0.5 | 1 | # | 20 | Small | aldex2 | 38 |
| # | sex | 4 | 0 | 0 | 1 | 1 | # | 20 | Small | aldex2 | 38 |
| # | time | 5 | 3 | 0 | 0.625 | 1 | # | 20 | Small | aldex2 | 38 |
| # | interaction | 0 | 0 | 4 | 0 | 0 | # | 20 | Small | aldex2 | 38 |
| # | disease | 7 | 6 | 0 | 0.538461538 | 1 | # | 20 | Baseline | paralle | 38 |
| # | sex | 4 | 8 | 0 | 0.333333333 | 1 | # | 20 | Baseline | paralle | 38 |
| # | time | 5 | 7 | 0 | 0.416666667 | 1 | # | 20 | Baseline | paralle | 38 |
| # | interaction | 4 | 1 | 0 | 0.8 | 1 | # | 20 | Baseline | paralle | 38 |
| # | disease | 7 | 4 | 0 | 0.636363636 | 1 | # | 20 | Baseline | ancom | 38 |
| # | sex | 4 | 0 | 0 | 1 | 1 | # | 20 | Baseline | ancom | 38 |
| # | time | 5 | 3 | 0 | 0.625 | 1 | # | 20 | Baseline | ancom | 38 |
| # | interaction | 0 | 0 | 4 | 0 | 0 | # | 20 | Baseline | ancom | 38 |
| # | disease | 7 | 63 | 0 | 0.1 | 1 | # | 20 | Baseline | aldex2 | 38 |
| # | sex | 4 | 17 | 0 | 0.19047619 | 1 | # | 20 | Baseline | aldex2 | 38 |
| # | time | 5 | 40 | 0 | 0.111111111 | 1 | # | 20 | Baseline | aldex2 | 38 |
| # | interaction | 0 | 0 | 4 | 0 | 0 | # | 20 | Baseline | aldex2 | 38 |
| # | disease | 7 | 0 | 0 | 1 | 1 | # | 20 | Small | paralle | 38 |
| # | sex | 4 | 0 | 0 | 1 | 1 | # | 20 | Small | paralle | 38 |
| # | time | 5 | 0 | 0 | 1 | 1 | # | 20 | Small | paralle | 38 |
| # | interaction | 4 | 0 | 0 | 1 | 1 | # | 20 | Small | paralle | 38 |
| # | disease | 7 | 4 | 0 | 0.636363636 | 1 | # | 20 | Small | ancom | 38 |
| # | sex | 4 | 0 | 0 | 1 | 1 | # | 20 | Small | ancom | 38 |
| # | time | 5 | 3 | 0 | 0.625 | 1 | # | 20 | Small | ancom | 38 |
| # | interaction | 0 | 0 | 4 | 0 | 0 | # | 20 | Small | ancom | 38 |
| # | disease | 7 | 9 | 0 | 0.4375 | 1 | # | 20 | Small | aldex2 | 38 |
| # | sex | 4 | 1 | 0 | 0.8 | 1 | # | 20 | Small | aldex2 | 38 |
| # | time | 5 | 5 | 0 | 0.5 | 1 | # | 20 | Small | aldex2 | 38 |
| # | interaction | 0 | 0 | 4 | 0 | 0 | # | 20 | Small | aldex2 | 38 |
| # | disease | 4 | 8 | 0 | 0.333333333 | 1 | # | 10 | Baseline | paralle | 39 |
| # | sex | 2 | 11 | 0 | 0.153846154 | 1 | # | 10 | Baseline | paralle | 39 |
| # | time | 2 | 4 | 0 | 0.333333333 | 1 | # | 10 | Baseline | paralle | 39 |
| # | interaction | 2 | 5 | 0 | 0.285714286 | 1 | # | 10 | Baseline | paralle | 39 |
| # | disease | 4 | 1 | 0 | 0.8 | 1 | # | 10 | Baseline | ancom | 39 |
| # | sex | 2 | 0 | 0 | 1 | 1 | # | 10 | Baseline | ancom | 39 |
| # | time | 2 | 0 | 0 | 1 | 1 | # | 10 | Baseline | ancom | 39 |
| # | interaction | 0 | 0 | 2 | 0 | 0 | # | 10 | Baseline | ancom | 39 |
| # | disease | 4 | 12 | 0 | 0.25 | 1 | # | 10 | Baseline | aldex2 | 39 |
| # | sex | 2 | 2 | 0 | 0.5 | 1 | # | 10 | Baseline | aldex2 | 39 |
| # | time | 2 | 4 | 0 | 0.333333333 | 1 | # | 10 | Baseline | aldex2 | 39 |
| # | interaction | 0 | 0 | 2 | 0 | 0 | # | 10 | Baseline | aldex2 | 39 |
| # | disease | 4 | 0 | 0 | 1 | 1 | # | 10 | Small | paralle | 39 |
| # | sex | 2 | 0 | 0 | 1 | 1 | # | 10 | Small | paralle | 39 |
| # | time | 2 | 0 | 0 | 1 | 1 | # | 10 | Small | paralle | 39 |
| # | interaction | 0 | 0 | 2 | 0 | 0 | # | 10 | Small | paralle | 39 |
| # | disease | 4 | 1 | 0 | 0.8 | 1 | # | 10 | Small | ancom | 39 |
| # | sex | 2 | 0 | 0 | 1 | 1 | # | 10 | Small | ancom | 39 |
| # | time | 2 | 0 | 0 | 1 | 1 | # | 10 | Small | ancom | 39 |
| # | interaction | 0 | 0 | 2 | 0 | 0 | # | 10 | Small | ancom | 39 |
| # | disease | 4 | 1 | 0 | 0.8 | 1 | # | 10 | Small | aldex2 | 39 |
| # | sex | 2 | 0 | 0 | 1 | 1 | # | 10 | Small | aldex2 | 39 |
| # | time | 2 | 1 | 0 | 0.666666667 | 1 | # | 10 | Small | aldex2 | 39 |
| # | interaction | 0 | 0 | 2 | 0 | 0 | # | 10 | Small | aldex2 | 39 |
| # | disease | 4 | 6 | 0 | 0.4 | 1 | # | 10 | Baseline | paralle | 39 |
| # | sex | 2 | 2 | 0 | 0.5 | 1 | # | 10 | Baseline | paralle | 39 |
| # | time | 2 | 4 | 0 | 0.333333333 | 1 | # | 10 | Baseline | paralle | 39 |
| # | interaction | 2 | 2 | 0 | 0.5 | 1 | # | 10 | Baseline | paralle | 39 |
| # | disease | 4 | 1 | 0 | 0.8 | 1 | # | 10 | Baseline | ancom | 39 |
| # | sex | 2 | 0 | 0 | 1 | 1 | # | 10 | Baseline | ancom | 39 |
| # | time | 2 | 1 | 0 | 0.666666667 | 1 | # | 10 | Baseline | ancom | 39 |
| # | interaction | 0 | 0 | 2 | 0 | 0 | # | 10 | Baseline | ancom | 39 |
| # | disease | 4 | 19 | 0 | 0.173913043 | 1 | # | 10 | Baseline | aldex2 | 39 |
| # | sex | 2 | 7 | 0 | 0.222222222 | 1 | # | 10 | Baseline | aldex2 | 39 |
| # | time | 2 | 9 | 0 | 0.181818182 | 1 | # | 10 | Baseline | aldex2 | 39 |
| # | interaction | 0 | 0 | 2 | 0 | 0 | # | 10 | Baseline | aldex2 | 39 |
| # | disease | 4 | 0 | 0 | 1 | 1 | # | 10 | Small | paralle | 39 |
| # | sex | 2 | 0 | 0 | 1 | 1 | # | 10 | Small | paralle | 39 |
| # | time | 2 | 0 | 0 | 1 | 1 | # | 10 | Small | paralle | 39 |
| # | interaction | 2 | 0 | 0 | 1 | 1 | # | 10 | Small | paralle | 39 |
| # | disease | 4 | 1 | 0 | 0.8 | 1 | # | 10 | Small | ancom | 39 |
| # | sex | 2 | 0 | 0 | 1 | 1 | # | 10 | Small | ancom | 39 |
| # | time | 2 | 1 | 0 | 0.666666667 | 1 | # | 10 | Small | ancom | 39 |
| # | interaction | 0 | 0 | 2 | 0 | 0 | # | 10 | Small | ancom | 39 |
| # | disease | 4 | 1 | 0 | 0.8 | 1 | # | 10 | Small | aldex2 | 39 |
| # | sex | 2 | 0 | 0 | 1 | 1 | # | 10 | Small | aldex2 | 39 |
| # | time | 2 | 2 | 0 | 0.5 | 1 | # | 10 | Small | aldex2 | 39 |
| # | interaction | 0 | 0 | 2 | 0 | 0 | # | 10 | Small | aldex2 | 39 |
| # | disease | 4 | 6 | 0 | 0.4 | 1 | # | 10 | Baseline | paralle | 39 |
| # | sex | 2 | 3 | 0 | 0.4 | 1 | # | 10 | Baseline | paralle | 39 |
| # | time | 2 | 3 | 0 | 0.4 | 1 | # | 10 | Baseline | paralle | 39 |
| # | interaction | 2 | 4 | 0 | 0.333333333 | 1 | # | 10 | Baseline | paralle | 39 |
| # | disease | 4 | 2 | 0 | 0.666666667 | 1 | # | 10 | Baseline | ancom | 39 |
| # | sex | 2 | 0 | 0 | 1 | 1 | # | 10 | Baseline | ancom | 39 |
| # | time | 2 | 1 | 0 | 0.666666667 | 1 | # | 10 | Baseline | ancom | 39 |
| # | interaction | 0 | 0 | 2 | 0 | 0 | # | 10 | Baseline | ancom | 39 |
| # | disease | 4 | 23 | 0 | 0.148148148 | 1 | # | 10 | Baseline | aldex2 | 39 |
| # | sex | 2 | 6 | 0 | 0.25 | 1 | # | 10 | Baseline | aldex2 | 39 |
| # | time | 2 | 11 | 0 | 0.153846154 | 1 | # | 10 | Baseline | aldex2 | 39 |
| # | interaction | 0 | 0 | 2 | 0 | 0 | # | 10 | Baseline | aldex2 | 39 |
| # | disease | 4 | 0 | 0 | 1 | 1 | # | 10 | Small | paralle | 39 |
| # | sex | 2 | 0 | 0 | 1 | 1 | # | 10 | Small | paralle | 39 |
| # | time | 2 | 0 | 0 | 1 | 1 | # | 10 | Small | paralle | 39 |
| # | interaction | 2 | 1 | 0 | 0.666666667 | 1 | # | 10 | Small | paralle | 39 |
| # | disease | 4 | 2 | 0 | 0.666666667 | 1 | # | 10 | Small | ancom | 39 |
| # | sex | 2 | 0 | 0 | 1 | 1 | # | 10 | Small | ancom | 39 |
| # | time | 2 | 1 | 0 | 0.666666667 | 1 | # | 10 | Small | ancom | 39 |
| # | interaction | 0 | 0 | 2 | 0 | 0 | # | 10 | Small | ancom | 39 |
| # | disease | 4 | 2 | 0 | 0.666666667 | 1 | # | 10 | Small | aldex2 | 39 |
| # | sex | 2 | 0 | 0 | 1 | 1 | # | 10 | Small | aldex2 | 39 |
| # | time | 2 | 2 | 0 | 0.5 | 1 | # | 10 | Small | aldex2 | 39 |
| # | interaction | 0 | 0 | 2 | 0 | 0 | # | 10 | Small | aldex2 | 39 |
| # | disease | 4 | 4 | 0 | 0.5 | 1 | # | 10 | Baseline | paralle | 39 |
| # | sex | 2 | 0 | 0 | 1 | 1 | # | 10 | Baseline | paralle | 39 |
| # | time | 2 | 5 | 0 | 0.285714286 | 1 | # | 10 | Baseline | paralle | 39 |
| # | interaction | 2 | 4 | 0 | 0.333333333 | 1 | # | 10 | Baseline | paralle | 39 |
| # | disease | 4 | 2 | 0 | 0.666666667 | 1 | # | 10 | Baseline | ancom | 39 |
| # | sex | 2 | 0 | 0 | 1 | 1 | # | 10 | Baseline | ancom | 39 |
| # | time | 2 | 1 | 0 | 0.666666667 | 1 | # | 10 | Baseline | ancom | 39 |
| # | interaction | 0 | 0 | 2 | 0 | 0 | # | 10 | Baseline | ancom | 39 |
| # | disease | 4 | 25 | 0 | 0.137931034 | 1 | # | 10 | Baseline | aldex2 | 39 |
| # | sex | 2 | 8 | 0 | 0.2 | 1 | # | 10 | Baseline | aldex2 | 39 |
| # | time | 2 | 11 | 0 | 0.153846154 | 1 | # | 10 | Baseline | aldex2 | 39 |
| # | interaction | 0 | 0 | 2 | 0 | 0 | # | 10 | Baseline | aldex2 | 39 |
| # | disease | 4 | 0 | 0 | 1 | 1 | # | 10 | Small | paralle | 39 |
| # | sex | 2 | 0 | 0 | 1 | 1 | # | 10 | Small | paralle | 39 |
| # | time | 2 | 0 | 0 | 1 | 1 | # | 10 | Small | paralle | 39 |
| # | interaction | 2 | 1 | 0 | 0.666666667 | 1 | # | 10 | Small | paralle | 39 |
| # | disease | 4 | 2 | 0 | 0.666666667 | 1 | # | 10 | Small | ancom | 39 |
| # | sex | 2 | 0 | 0 | 1 | 1 | # | 10 | Small | ancom | 39 |
| # | time | 2 | 1 | 0 | 0.666666667 | 1 | # | 10 | Small | ancom | 39 |
| # | interaction | 0 | 0 | 2 | 0 | 0 | # | 10 | Small | ancom | 39 |
| # | disease | 4 | 2 | 0 | 0.666666667 | 1 | # | 10 | Small | aldex2 | 39 |
| # | sex | 2 | 0 | 0 | 1 | 1 | # | 10 | Small | aldex2 | 39 |
| # | time | 2 | 2 | 0 | 0.5 | 1 | # | 10 | Small | aldex2 | 39 |
| # | interaction | 0 | 0 | 2 | 0 | 0 | # | 10 | Small | aldex2 | 39 |
| # | disease | 7 | 13 | 0 | 0.35 | 1 | # | 20 | Baseline | paralle | 39 |
| # | sex | 4 | 9 | 0 | 0.307692308 | 1 | # | 20 | Baseline | paralle | 39 |
| # | time | 5 | 16 | 0 | 0.238095238 | 1 | # | 20 | Baseline | paralle | 39 |
| # | interaction | 4 | 13 | 0 | 0.235294118 | 1 | # | 20 | Baseline | paralle | 39 |
| # | disease | 7 | 3 | 0 | 0.7 | 1 | # | 20 | Baseline | ancom | 39 |
| # | sex | 4 | 0 | 0 | 1 | 1 | # | 20 | Baseline | ancom | 39 |
| # | time | 5 | 3 | 0 | 0.625 | 1 | # | 20 | Baseline | ancom | 39 |
| # | interaction | 0 | 0 | 4 | 0 | 0 | # | 20 | Baseline | ancom | 39 |
| # | disease | 7 | 23 | 0 | 0.233333333 | 1 | # | 20 | Baseline | aldex2 | 39 |
| # | sex | 4 | 6 | 0 | 0.4 | 1 | # | 20 | Baseline | aldex2 | 39 |
| # | time | 5 | 19 | 0 | 0.208333333 | 1 | # | 20 | Baseline | aldex2 | 39 |
| # | interaction | 0 | 0 | 4 | 0 | 0 | # | 20 | Baseline | aldex2 | 39 |
| # | disease | 7 | 0 | 0 | 1 | 1 | # | 20 | Small | paralle | 39 |
| # | sex | 4 | 0 | 0 | 1 | 1 | # | 20 | Small | paralle | 39 |
| # | time | 5 | 0 | 0 | 1 | 1 | # | 20 | Small | paralle | 39 |
| # | interaction | 1 | 0 | 3 | 1 | 0.25 | # | 20 | Small | paralle | 39 |
| # | disease | 7 | 2 | 0 | 0.777777778 | 1 | # | 20 | Small | ancom | 39 |
| # | sex | 4 | 0 | 0 | 1 | 1 | # | 20 | Small | ancom | 39 |
| # | time | 5 | 2 | 0 | 0.714285714 | 1 | # | 20 | Small | ancom | 39 |

| | | | | | | | | | | | |
|---|---|---|---|---|---|---|---|---|---|---|---|
| # | interaction | 0 | 0 | 4 | 0 | 0 | # | 20 | Small | ancom | 39 |
| # | disease | 7 | 1 | 0 | 0.875 | 1 | # | 20 | Small | aldex2 | 39 |
| # | sex | 4 | 0 | 0 | 1 | 1 | # | 20 | Small | aldex2 | 39 |
| # | time | 5 | 1 | 0 | 0.833333333 | 1 | # | 20 | Small | aldex2 | 39 |
| # | interaction | 0 | 0 | 4 | 0 | 0 | # | 20 | Small | aldex2 | 39 |
| # | disease | 7 | 10 | 0 | 0.411764706 | 1 | # | 20 | Baseline | paralle | 39 |
| # | sex | 4 | 8 | 0 | 0.333333333 | 1 | # | 20 | Baseline | paralle | 39 |
| # | time | 5 | 16 | 0 | 0.238095238 | 1 | # | 20 | Baseline | paralle | 39 |
| # | interaction | 4 | 9 | 0 | 0.307692308 | 1 | # | 20 | Baseline | paralle | 39 |
| # | disease | 7 | 3 | 0 | 0.7 | 1 | # | 20 | Baseline | ancom | 39 |
| # | sex | 4 | 0 | 0 | 1 | 1 | # | 20 | Baseline | ancom | 39 |
| # | time | 5 | 3 | 0 | 0.625 | 1 | # | 20 | Baseline | ancom | 39 |
| # | interaction | 0 | 0 | 4 | 0 | 0 | # | 20 | Baseline | ancom | 39 |
| # | disease | 7 | 34 | 0 | 0.170731707 | 1 | # | 20 | Baseline | aldex2 | 39 |
| # | sex | 4 | 11 | 0 | 0.266666667 | 1 | # | 20 | Baseline | aldex2 | 39 |
| # | time | 5 | 27 | 0 | 0.15625 | 1 | # | 20 | Baseline | aldex2 | 39 |
| # | interaction | 0 | 0 | 4 | 0 | 0 | # | 20 | Baseline | aldex2 | 39 |
| # | disease | 7 | 0 | 0 | 1 | 1 | # | 20 | Small | paralle | 39 |
| # | sex | 4 | 0 | 0 | 1 | 1 | # | 20 | Small | paralle | 39 |
| # | time | 5 | 0 | 0 | 1 | 1 | # | 20 | Small | paralle | 39 |
| # | interaction | 2 | 0 | 2 | 1 | 0.5 | # | 20 | Small | paralle | 39 |
| # | disease | 7 | 3 | 0 | 0.7 | 1 | # | 20 | Small | ancom | 39 |
| # | sex | 4 | 0 | 0 | 1 | 1 | # | 20 | Small | ancom | 39 |
| # | time | 5 | 3 | 0 | 0.625 | 1 | # | 20 | Small | ancom | 39 |
| # | interaction | 0 | 0 | 4 | 0 | 0 | # | 20 | Small | ancom | 39 |
| # | disease | 7 | 5 | 0 | 0.583333333 | 1 | # | 20 | Small | aldex2 | 39 |
| # | sex | 4 | 0 | 0 | 1 | 1 | # | 20 | Small | aldex2 | 39 |
| # | time | 5 | 5 | 0 | 0.5 | 1 | # | 20 | Small | aldex2 | 39 |
| # | interaction | 0 | 0 | 4 | 0 | 0 | # | 20 | Small | aldex2 | 39 |
| # | disease | 7 | 14 | 0 | 0.333333333 | 1 | # | 20 | Baseline | paralle | 39 |
| # | sex | 4 | 7 | 0 | 0.363636364 | 1 | # | 20 | Baseline | paralle | 39 |
| # | time | 5 | 8 | 0 | 0.384615385 | 1 | # | 20 | Baseline | paralle | 39 |
| # | interaction | 4 | 7 | 0 | 0.363636364 | 1 | # | 20 | Baseline | paralle | 39 |
| # | disease | 7 | 4 | 0 | 0.636363636 | 1 | # | 20 | Baseline | ancom | 39 |
| # | sex | 4 | 0 | 0 | 1 | 1 | # | 20 | Baseline | ancom | 39 |
| # | time | 5 | 4 | 0 | 0.555555556 | 1 | # | 20 | Baseline | ancom | 39 |
| # | interaction | 0 | 0 | 4 | 0 | 0 | # | 20 | Baseline | ancom | 39 |
| # | disease | 7 | 69 | 0 | 0.125 | 1 | # | 20 | Baseline | aldex2 | 39 |
| # | sex | 4 | 14 | 0 | 0.222222222 | 1 | # | 20 | Baseline | aldex2 | 39 |
| # | time | 5 | 33 | 0 | 0.131578947 | 1 | # | 20 | Baseline | aldex2 | 39 |
| # | interaction | 0 | 0 | 4 | 0 | 0 | # | 20 | Baseline | aldex2 | 39 |
| # | disease | 7 | 1 | 0 | 0.875 | 1 | # | 20 | Small | paralle | 39 |
| # | sex | 4 | 0 | 0 | 1 | 1 | # | 20 | Small | paralle | 39 |
| # | time | 5 | 1 | 0 | 0.833333333 | 1 | # | 20 | Small | paralle | 39 |
| # | interaction | 3 | 0 | 1 | 1 | 0.75 | # | 20 | Small | paralle | 39 |
| # | disease | 7 | 4 | 0 | 0.636363636 | 1 | # | 20 | Small | ancom | 39 |
| # | sex | 4 | 0 | 0 | 1 | 1 | # | 20 | Small | ancom | 39 |
| # | time | 5 | 4 | 0 | 0.555555556 | 1 | # | 20 | Small | ancom | 39 |
| # | interaction | 0 | 0 | 4 | 0 | 0 | # | 20 | Small | ancom | 39 |
| # | disease | 7 | 9 | 0 | 0.4375 | 1 | # | 20 | Small | aldex2 | 39 |
| # | sex | 4 | 0 | 0 | 1 | 1 | # | 20 | Small | aldex2 | 39 |
| # | time | 5 | 5 | 0 | 0.5 | 1 | # | 20 | Small | aldex2 | 39 |
| # | interaction | 0 | 0 | 4 | 0 | 0 | # | 20 | Small | aldex2 | 39 |
| # | disease | 7 | 15 | 0 | 0.318181818 | 1 | # | 20 | Baseline | paralle | 39 |
| # | sex | 4 | 7 | 0 | 0.363636364 | 1 | # | 20 | Baseline | paralle | 39 |
| # | time | 5 | 10 | 0 | 0.333333333 | 1 | # | 20 | Baseline | paralle | 39 |
| # | interaction | 4 | 3 | 0 | 0.571428571 | 1 | # | 20 | Baseline | paralle | 39 |
| # | disease | 7 | 4 | 0 | 0.636363636 | 1 | # | 20 | Baseline | ancom | 39 |
| # | sex | 4 | 0 | 0 | 1 | 1 | # | 20 | Baseline | ancom | 39 |
| # | time | 5 | 4 | 0 | 0.555555556 | 1 | # | 20 | Baseline | ancom | 39 |
| # | interaction | 0 | 0 | 4 | 0 | 0 | # | 20 | Baseline | ancom | 39 |
| # | disease | 7 | 55 | 0 | 0.112903226 | 1 | # | 20 | Baseline | aldex2 | 39 |
| # | sex | 4 | 19 | 0 | 0.173913043 | 1 | # | 20 | Baseline | aldex2 | 39 |
| # | time | 5 | 66 | 0 | 0.098039216 | 1 | # | 20 | Baseline | aldex2 | 39 |
| # | interaction | 0 | 0 | 4 | 0 | 0 | # | 20 | Baseline | aldex2 | 39 |
| # | disease | 7 | 1 | 0 | 0.875 | 1 | # | 20 | Small | paralle | 39 |
| # | sex | 4 | 0 | 0 | 1 | 1 | # | 20 | Small | paralle | 39 |
| # | time | 5 | 1 | 0 | 0.833333333 | 1 | # | 20 | Small | paralle | 39 |
| # | interaction | 4 | 0 | 0 | 1 | 1 | # | 20 | Small | paralle | 39 |
| # | disease | 7 | 4 | 0 | 0.636363636 | 1 | # | 20 | Small | ancom | 39 |
| # | sex | 4 | 0 | 0 | 1 | 1 | # | 20 | Small | ancom | 39 |
| # | time | 5 | 4 | 0 | 0.555555556 | 1 | # | 20 | Small | ancom | 39 |
| # | interaction | 0 | 0 | 4 | 0 | 0 | # | 20 | Small | ancom | 39 |
| # | disease | 7 | 14 | 0 | 0.333333333 | 1 | # | 20 | Small | aldex2 | 39 |
| # | sex | 4 | 0 | 0 | 1 | 1 | # | 20 | Small | aldex2 | 39 |
| # | time | 5 | 8 | 0 | 0.384615385 | 1 | # | 20 | Small | aldex2 | 39 |
| # | interaction | 0 | 0 | 4 | 0 | 0 | # | 20 | Small | aldex2 | 39 |
| # | disease | 4 | 9 | 0 | 0.307692308 | 1 | # | 10 | Baseline | paralle | 40 |
| # | sex | 2 | 7 | 0 | 0.222222222 | 1 | # | 10 | Baseline | paralle | 40 |
| # | time | 2 | 8 | 0 | 0.2 | 1 | # | 10 | Baseline | paralle | 40 |
| # | interaction | 2 | 12 | 0 | 0.142857143 | 1 | # | 10 | Baseline | paralle | 40 |
| # | disease | 4 | 1 | 0 | 0.8 | 1 | # | 10 | Baseline | ancom | 40 |
| # | sex | 2 | 0 | 0 | 1 | 1 | # | 10 | Baseline | ancom | 40 |
| # | time | 2 | 0 | 0 | 1 | 1 | # | 10 | Baseline | ancom | 40 |
| # | interaction | 0 | 0 | 2 | 0 | 0 | # | 10 | Baseline | ancom | 40 |
| # | disease | 4 | 8 | 0 | 0.333333333 | 1 | # | 10 | Baseline | aldex2 | 40 |
| # | sex | 2 | 3 | 0 | 0.4 | 1 | # | 10 | Baseline | aldex2 | 40 |
| # | time | 2 | 8 | 0 | 0.2 | 1 | # | 10 | Baseline | aldex2 | 40 |
| # | interaction | 0 | 0 | 2 | 0 | 0 | # | 10 | Baseline | aldex2 | 40 |
| # | disease | 4 | 0 | 0 | 1 | 1 | # | 10 | Small | paralle | 40 |
| # | sex | 2 | 0 | 0 | 1 | 1 | # | 10 | Small | paralle | 40 |
| # | time | 2 | 1 | 0 | 0.666666667 | 1 | # | 10 | Small | paralle | 40 |
| # | interaction | 2 | 2 | 0 | 0.5 | 1 | # | 10 | Small | paralle | 40 |
| # | disease | 4 | 0 | 0 | 1 | 1 | # | 10 | Small | ancom | 40 |
| # | sex | 2 | 0 | 0 | 1 | 1 | # | 10 | Small | ancom | 40 |
| # | time | 2 | 0 | 0 | 1 | 1 | # | 10 | Small | ancom | 40 |
| # | interaction | 0 | 0 | 2 | 0 | 0 | # | 10 | Small | ancom | 40 |
| # | disease | 4 | 1 | 0 | 0.8 | 1 | # | 10 | Small | aldex2 | 40 |
| # | sex | 2 | 0 | 0 | 1 | 1 | # | 10 | Small | aldex2 | 40 |
| # | time | 2 | 1 | 0 | 0.666666667 | 1 | # | 10 | Small | aldex2 | 40 |
| # | interaction | 0 | 0 | 2 | 0 | 0 | # | 10 | Small | aldex2 | 40 |
| # | disease | 4 | 6 | 0 | 0.4 | 1 | # | 10 | Baseline | paralle | 40 |
| # | sex | 2 | 7 | 0 | 0.222222222 | 1 | # | 10 | Baseline | paralle | 40 |
| # | time | 2 | 5 | 0 | 0.285714286 | 1 | # | 10 | Baseline | paralle | 40 |
| # | interaction | 2 | 8 | 0 | 0.2 | 1 | # | 10 | Baseline | paralle | 40 |
| # | disease | 4 | 2 | 0 | 0.666666667 | 1 | # | 10 | Baseline | ancom | 40 |
| # | sex | 2 | 0 | 0 | 1 | 1 | # | 10 | Baseline | ancom | 40 |
| # | time | 2 | 2 | 0 | 0.5 | 1 | # | 10 | Baseline | ancom | 40 |
| # | interaction | 0 | 0 | 2 | 0 | 0 | # | 10 | Baseline | ancom | 40 |
| # | disease | 4 | 14 | 0 | 0.222222222 | 1 | # | 10 | Baseline | aldex2 | 40 |
| # | sex | 2 | 7 | 0 | 0.222222222 | 1 | # | 10 | Baseline | aldex2 | 40 |
| # | time | 2 | 11 | 0 | 0.153846154 | 1 | # | 10 | Baseline | aldex2 | 40 |
| # | interaction | 0 | 0 | 2 | 0 | 0 | # | 10 | Baseline | aldex2 | 40 |
| # | disease | 4 | 1 | 0 | 0.8 | 1 | # | 10 | Small | paralle | 40 |
| # | sex | 2 | 0 | 0 | 1 | 1 | # | 10 | Small | paralle | 40 |
| # | time | 2 | 0 | 0 | 1 | 1 | # | 10 | Small | paralle | 40 |
| # | interaction | 2 | 0 | 0 | 1 | 1 | # | 10 | Small | paralle | 40 |
| # | disease | 4 | 2 | 0 | 0.666666667 | 1 | # | 10 | Small | ancom | 40 |
| # | sex | 2 | 0 | 0 | 1 | 1 | # | 10 | Small | ancom | 40 |
| # | time | 2 | 2 | 0 | 0.5 | 1 | # | 10 | Small | ancom | 40 |
| # | interaction | 0 | 0 | 2 | 0 | 0 | # | 10 | Small | ancom | 40 |
| # | disease | 4 | 3 | 0 | 0.571428571 | 1 | # | 10 | Small | aldex2 | 40 |
| # | sex | 2 | 0 | 0 | 1 | 1 | # | 10 | Small | aldex2 | 40 |
| # | time | 2 | 2 | 0 | 0.5 | 1 | # | 10 | Small | aldex2 | 40 |
| # | interaction | 0 | 0 | 2 | 0 | 0 | # | 10 | Small | aldex2 | 40 |
| # | disease | 4 | 7 | 0 | 0.363636364 | 1 | # | 10 | Baseline | paralle | 40 |
| # | sex | 2 | 1 | 0 | 0.666666667 | 1 | # | 10 | Baseline | paralle | 40 |
| # | time | 2 | 7 | 0 | 0.222222222 | 1 | # | 10 | Baseline | paralle | 40 |
| # | interaction | 2 | 5 | 0 | 0.285714286 | 1 | # | 10 | Baseline | paralle | 40 |
| # | disease | 4 | 2 | 0 | 0.666666667 | 1 | # | 10 | Baseline | ancom | 40 |
| # | sex | 2 | 0 | 0 | 1 | 1 | # | 10 | Baseline | ancom | 40 |
| # | time | 2 | 2 | 0 | 0.5 | 1 | # | 10 | Baseline | ancom | 40 |
| # | interaction | 0 | 0 | 2 | 0 | 0 | # | 10 | Baseline | ancom | 40 |
| # | disease | 4 | 16 | 0 | 0.2 | 1 | # | 10 | Baseline | aldex2 | 40 |
| # | sex | 2 | 7 | 0 | 0.222222222 | 1 | # | 10 | Baseline | aldex2 | 40 |
| # | time | 2 | 8 | 0 | 0.2 | 1 | # | 10 | Baseline | aldex2 | 40 |
| # | interaction | 0 | 0 | 2 | 0 | 0 | # | 10 | Baseline | aldex2 | 40 |
| # | disease | 4 | 0 | 0 | 1 | 1 | # | 10 | Small | paralle | 40 |
| # | sex | 2 | 0 | 0 | 1 | 1 | # | 10 | Small | paralle | 40 |
| # | time | 2 | 0 | 0 | 1 | 1 | # | 10 | Small | paralle | 40 |
| # | interaction | 2 | 1 | 0 | 0.666666667 | 1 | # | 10 | Small | paralle | 40 |
| # | disease | 4 | 2 | 0 | 0.666666667 | 1 | # | 10 | Small | ancom | 40 |
| # | sex | 2 | 0 | 0 | 1 | 1 | # | 10 | Small | ancom | 40 |
| # | time | 2 | 2 | 0 | 0.5 | 1 | # | 10 | Small | ancom | 40 |
| # | interaction | 0 | 0 | 2 | 0 | 0 | # | 10 | Small | ancom | 40 |
| # | disease | 4 | 2 | 0 | 0.666666667 | 1 | # | 10 | Small | aldex2 | 40 |
| # | sex | 2 | 0 | 0 | 1 | 1 | # | 10 | Small | aldex2 | 40 |
| # | time | 2 | 2 | 0 | 0.5 | 1 | # | 10 | Small | aldex2 | 40 |
| # | interaction | 0 | 0 | 2 | 0 | 0 | # | 10 | Small | aldex2 | 40 |
| # | disease | 4 | 8 | 0 | 0.333333333 | 1 | # | 10 | Baseline | paralle | 40 |
| # | sex | 2 | 4 | 0 | 0.333333333 | 1 | # | 10 | Baseline | paralle | 40 |
| # | time | 2 | 6 | 0 | 0.25 | 1 | # | 10 | Baseline | paralle | 40 |
| # | interaction | 2 | 5 | 0 | 0.285714286 | 1 | # | 10 | Baseline | paralle | 40 |
| # | disease | 4 | 2 | 0 | 0.666666667 | 1 | # | 10 | Baseline | ancom | 40 |
| # | sex | 2 | 0 | 0 | 1 | 1 | # | 10 | Baseline | ancom | 40 |
| # | time | 2 | 2 | 0 | 0.5 | 1 | # | 10 | Baseline | ancom | 40 |
| # | interaction | 0 | 0 | 2 | 0 | 0 | # | 10 | Baseline | ancom | 40 |
| # | disease | 4 | 31 | 0 | 0.114285714 | 1 | # | 10 | Baseline | aldex2 | 40 |
| # | sex | 2 | 11 | 0 | 0.153846154 | 1 | # | 10 | Baseline | aldex2 | 40 |
| # | time | 2 | 11 | 0 | 0.153846154 | 1 | # | 10 | Baseline | aldex2 | 40 |
| # | interaction | 0 | 0 | 2 | 0 | 0 | # | 10 | Baseline | aldex2 | 40 |
| # | disease | 4 | 0 | 0 | 1 | 1 | # | 10 | Small | paralle | 40 |
| # | sex | 2 | 0 | 0 | 1 | 1 | # | 10 | Small | paralle | 40 |
| # | time | 2 | 0 | 0 | 1 | 1 | # | 10 | Small | paralle | 40 |
| # | interaction | 2 | 1 | 0 | 0.666666667 | 1 | # | 10 | Small | paralle | 40 |
| # | disease | 4 | 2 | 0 | 0.666666667 | 1 | # | 10 | Small | ancom | 40 |
| # | sex | 2 | 0 | 0 | 1 | 1 | # | 10 | Small | ancom | 40 |
| # | time | 2 | 2 | 0 | 0.5 | 1 | # | 10 | Small | ancom | 40 |
| # | interaction | 0 | 0 | 2 | 0 | 0 | # | 10 | Small | ancom | 40 |
| # | disease | 4 | 4 | 0 | 0.5 | 1 | # | 10 | Small | aldex2 | 40 |
| # | sex | 2 | 0 | 0 | 1 | 1 | # | 10 | Small | aldex2 | 40 |
| # | time | 2 | 3 | 0 | 0.4 | 1 | # | 10 | Small | aldex2 | 40 |
| # | interaction | 0 | 0 | 2 | 0 | 0 | # | 10 | Small | aldex2 | 40 |
| # | disease | 7 | 10 | 0 | 0.411764706 | 1 | # | 20 | Baseline | paralle | 40 |
| # | sex | 4 | 2 | 0 | 0.666666667 | 1 | # | 20 | Baseline | paralle | 40 |
| # | time | 5 | 4 | 0 | 0.555555556 | 1 | # | 20 | Baseline | paralle | 40 |
| # | interaction | 4 | 7 | 0 | 0.363636364 | 1 | # | 20 | Baseline | paralle | 40 |
| # | disease | 7 | 2 | 0 | 0.777777778 | 1 | # | 20 | Baseline | ancom | 40 |
| # | sex | 4 | 0 | 0 | 1 | 1 | # | 20 | Baseline | ancom | 40 |
| # | time | 5 | 1 | 0 | 0.833333333 | 1 | # | 20 | Baseline | ancom | 40 |
| # | interaction | 0 | 0 | 4 | 0 | 0 | # | 20 | Baseline | ancom | 40 |
| # | disease | 7 | 21 | 0 | 0.25 | 1 | # | 20 | Baseline | aldex2 | 40 |
| # | sex | 4 | 9 | 0 | 0.307692308 | 1 | # | 20 | Baseline | aldex2 | 40 |
| # | time | 5 | 14 | 0 | 0.263157895 | 1 | # | 20 | Baseline | aldex2 | 40 |
| # | interaction | 0 | 0 | 4 | 0 | 0 | # | 20 | Baseline | aldex2 | 40 |
| # | disease | 7 | 1 | 0 | 0.875 | 1 | # | 20 | Small | paralle | 40 |
| # | sex | 4 | 1 | 0 | 0.8 | 1 | # | 20 | Small | paralle | 40 |
| # | time | 5 | 0 | 0 | 1 | 1 | # | 20 | Small | paralle | 40 |
| # | interaction | 2 | 0 | 2 | 1 | 0.5 | # | 20 | Small | paralle | 40 |
| # | disease | 7 | 1 | 0 | 0.875 | 1 | # | 20 | Small | ancom | 40 |
| # | sex | 4 | 0 | 0 | 1 | 1 | # | 20 | Small | ancom | 40 |
| # | time | 5 | 0 | 0 | 1 | 1 | # | 20 | Small | ancom | 40 |
| # | interaction | 0 | 0 | 4 | 0 | 0 | # | 20 | Small | ancom | 40 |
| # | disease | 7 | 1 | 0 | 0.875 | 1 | # | 20 | Small | aldex2 | 40 |
| # | sex | 4 | 1 | 0 | 0.8 | 1 | # | 20 | Small | aldex2 | 40 |
| # | time | 5 | 0 | 0 | 1 | 1 | # | 20 | Small | aldex2 | 40 |
| # | interaction | 0 | 0 | 4 | 0 | 0 | # | 20 | Small | aldex2 | 40 |
| # | disease | 7 | 11 | 0 | 0.388888889 | 1 | # | 20 | Baseline | paralle | 40 |
| # | sex | 4 | 3 | 0 | 0.571428571 | 1 | # | 20 | Baseline | paralle | 40 |
| # | time | 5 | 5 | 0 | 0.5 | 1 | # | 20 | Baseline | paralle | 40 |
| # | interaction | 4 | 8 | 0 | 0.333333333 | 1 | # | 20 | Baseline | paralle | 40 |
| # | disease | 7 | 2 | 0 | 0.777777778 | 1 | # | 20 | Baseline | ancom | 40 |
| # | sex | 4 | 0 | 0 | 1 | 1 | # | 20 | Baseline | ancom | 40 |
| # | time | 5 | 3 | 0 | 0.625 | 1 | # | 20 | Baseline | ancom | 40 |
| # | interaction | 0 | 0 | 4 | 0 | 0 | # | 20 | Baseline | ancom | 40 |
| # | disease | 7 | 34 | 0 | 0.170731707 | 1 | # | 20 | Baseline | aldex2 | 40 |
| # | sex | 4 | 8 | 0 | 0.333333333 | 1 | # | 20 | Baseline | aldex2 | 40 |
| # | time | 5 | 23 | 0 | 0.178571429 | 1 | # | 20 | Baseline | aldex2 | 40 |
| # | interaction | 0 | 0 | 4 | 0 | 0 | # | 20 | Baseline | aldex2 | 40 |
| # | disease | 7 | 1 | 0 | 0.875 | 1 | # | 20 | Small | paralle | 40 |
| # | sex | 4 | 0 | 0 | 1 | 1 | # | 20 | Small | paralle | 40 |
| # | time | 5 | 0 | 0 | 1 | 1 | # | 20 | Small | paralle | 40 |
| # | interaction | 3 | 0 | 1 | 1 | 0.75 | # | 20 | Small | paralle | 40 |
| # | disease | 7 | 2 | 0 | 0.777777778 | 1 | # | 20 | Small | ancom | 40 |
| # | sex | 4 | 0 | 0 | 1 | 1 | # | 20 | Small | ancom | 40 |
| # | time | 5 | 3 | 0 | 0.625 | 1 | # | 20 | Small | ancom | 40 |
| # | interaction | 0 | 0 | 4 | 0 | 0 | # | 20 | Small | ancom | 40 |
| # | disease | 7 | 4 | 0 | 0.636363636 | 1 | # | 20 | Small | aldex2 | 40 |
| # | sex | 4 | 0 | 0 | 1 | 1 | # | 20 | Small | aldex2 | 40 |
| # | time | 5 | 2 | 0 | 0.714285714 | 1 | # | 20 | Small | aldex2 | 40 |
| # | interaction | 0 | 0 | 4 | 0 | 0 | # | 20 | Small | aldex2 | 40 |
| # | disease | 7 | 13 | 0 | 0.35 | 1 | # | 20 | Baseline | paralle | 40 |
| # | sex | 4 | 1 | 0 | 0.8 | 1 | # | 20 | Baseline | paralle | 40 |
| # | time | 5 | 2 | 0 | 0.714285714 | 1 | # | 20 | Baseline | paralle | 40 |
| # | interaction | 4 | 8 | 0 | 0.333333333 | 1 | # | 20 | Baseline | paralle | 40 |
| # | disease | 7 | 3 | 0 | 0.7 | 1 | # | 20 | Baseline | ancom | 40 |
| # | sex | 4 | 0 | 0 | 1 | 1 | # | 20 | Baseline | ancom | 40 |
| # | time | 5 | 3 | 0 | 0.625 | 1 | # | 20 | Baseline | ancom | 40 |
| # | interaction | 0 | 0 | 4 | 0 | 0 | # | 20 | Baseline | ancom | 40 |
| # | disease | 7 | 65 | 0 | 0.134615385 | 1 | # | 20 | Baseline | aldex2 | 40 |
| # | sex | 4 | 8 | 0 | 0.333333333 | 1 | # | 20 | Baseline | aldex2 | 40 |
| # | time | 5 | 60 | 0 | 0.111111111 | 1 | # | 20 | Baseline | aldex2 | 40 |
| # | interaction | 0 | 0 | 4 | 0 | 0 | # | 20 | Baseline | aldex2 | 40 |
| # | disease | 7 | 1 | 0 | 0.875 | 1 | # | 20 | Small | paralle | 40 |
| # | sex | 4 | 0 | 0 | 1 | 1 | # | 20 | Small | paralle | 40 |
| # | time | 5 | 0 | 0 | 1 | 1 | # | 20 | Small | paralle | 40 |
| # | interaction | 4 | 0 | 0 | 1 | 1 | # | 20 | Small | paralle | 40 |
| # | disease | 7 | 3 | 0 | 0.7 | 1 | # | 20 | Small | ancom | 40 |
| # | sex | 4 | 0 | 0 | 1 | 1 | # | 20 | Small | ancom | 40 |
| # | time | 5 | 3 | 0 | 0.625 | 1 | # | 20 | Small | ancom | 40 |
| # | interaction | 0 | 0 | 4 | 0 | 0 | # | 20 | Small | ancom | 40 |
| # | disease | 7 | 3 | 0 | 0.7 | 1 | # | 20 | Small | aldex2 | 40 |
| # | sex | 4 | 0 | 0 | 1 | 1 | # | 20 | Small | aldex2 | 40 |
| # | time | 5 | 4 | 0 | 0.555555556 | 1 | # | 20 | Small | aldex2 | 40 |
| # | interaction | 0 | 0 | 4 | 0 | 0 | # | 20 | Small | aldex2 | 40 |
| # | disease | 7 | 11 | 0 | 0.388888889 | 1 | # | 20 | Baseline | paralle | 40 |
| # | sex | 4 | 2 | 0 | 0.666666667 | 1 | # | 20 | Baseline | paralle | 40 |
| # | time | 5 | 5 | 0 | 0.5 | 1 | # | 20 | Baseline | paralle | 40 |
| # | interaction | 4 | 9 | 0 | 0.307692308 | 1 | # | 20 | Baseline | paralle | 40 |
| # | disease | 7 | 3 | 0 | 0.7 | 1 | # | 20 | Baseline | ancom | 40 |
| # | sex | 4 | 0 | 0 | 1 | 1 | # | 20 | Baseline | ancom | 40 |
| # | time | 5 | 4 | 0 | 0.555555556 | 1 | # | 20 | Baseline | ancom | 40 |
| # | interaction | 0 | 0 | 4 | 0 | 0 | # | 20 | Baseline | ancom | 40 |
| # | disease | 7 | 52 | 0 | 0.118644068 | 1 | # | 20 | Baseline | aldex2 | 40 |
| # | sex | 4 | 16 | 0 | 0.2 | 1 | # | 20 | Baseline | aldex2 | 40 |
| # | time | 5 | 39 | 0 | 0.113636364 | 1 | # | 20 | Baseline | aldex2 | 40 |
| # | interaction | 0 | 0 | 4 | 0 | 0 | # | 20 | Baseline | aldex2 | 40 |
| # | disease | 7 | 1 | 0 | 0.875 | 1 | # | 20 | Small | paralle | 40 |
| # | sex | 4 | 0 | 0 | 1 | 1 | # | 20 | Small | paralle | 40 |
| # | time | 5 | 0 | 0 | 1 | 1 | # | 20 | Small | paralle | 40 |
| # | interaction | 4 | 0 | 0 | 1 | 1 | # | 20 | Small | paralle | 40 |
| # | disease | 7 | 3 | 0 | 0.7 | 1 | # | 20 | Small | ancom | 40 |
| # | sex | 4 | 0 | 0 | 1 | 1 | # | 20 | Small | ancom | 40 |
| # | time | 5 | 3 | 0 | 0.625 | 1 | # | 20 | Small | ancom | 40 |
| # | interaction | 0 | 0 | 4 | 0 | 0 | # | 20 | Small | ancom | 40 |
| # | disease | 7 | 6 | 0 | 0.538461538 | 1 | # | 20 | Small | aldex2 | 40 |
| # | sex | 4 | 0 | 0 | 1 | 1 | # | 20 | Small | aldex2 | 40 |
| # | time | 5 | 6 | 0 | 0.454545455 | 1 | # | 20 | Small | aldex2 | 40 |
| # | interaction | 0 | 0 | 4 | 0 | 0 | # | 20 | Small | aldex2 | 40 |
| # | disease | 4 | 7 | 0 | 0.363636364 | 1 | # | 10 | Baseline | paralle | 41 |
| # | sex | 2 | 10 | 0 | 0.166666667 | 1 | # | 10 | Baseline | paralle | 41 |
| # | time | 2 | 2 | 0 | 0.5 | 1 | # | 10 | Baseline | paralle | 41 |
| # | interaction | 2 | 10 | 0 | 0.166666667 | 1 | # | 10 | Baseline | paralle | 41 |
| # | disease | 4 | 0 | 0 | 1 | 1 | # | 10 | Baseline | ancom | 41 |
| # | sex | 2 | 0 | 0 | 1 | 1 | # | 10 | Baseline | ancom | 41 |
| # | time | 2 | 0 | 0 | 1 | 1 | # | 10 | Baseline | ancom | 41 |
| # | interaction | 0 | 0 | 2 | 0 | 0 | # | 10 | Baseline | ancom | 41 |
| # | disease | 4 | 7 | 0 | 0.363636364 | 1 | # | 10 | Baseline | aldex2 | 41 |
| # | sex | 2 | 9 | 0 | 0.181818182 | 1 | # | 10 | Baseline | aldex2 | 41 |
| # | time | 2 | 8 | 0 | 0.2 | 1 | # | 10 | Baseline | aldex2 | 41 |
| # | interaction | 0 | 0 | 2 | 0 | 0 | # | 10 | Baseline | aldex2 | 41 |
| # | disease | 4 | 1 | 0 | 0.8 | 1 | # | 10 | Small | paralle | 41 |
| # | sex | 2 | 0 | 0 | 1 | 1 | # | 10 | Small | paralle | 41 |
| # | time | 2 | 0 | 0 | 1 | 1 | # | 10 | Small | paralle | 41 |
| # | interaction | 1 | 1 | 1 | 0.5 | 0.5 | # | 10 | Small | paralle | 41 |
| # | disease | 4 | 0 | 0 | 1 | 1 | # | 10 | Small | ancom | 41 |
| # | sex | 2 | 0 | 0 | 1 | 1 | # | 10 | Small | ancom | 41 |
| # | time | 2 | 0 | 0 | 1 | 1 | # | 10 | Small | ancom | 41 |
| # | interaction | 0 | 0 | 2 | 0 | 0 | # | 10 | Small | ancom | 41 |
| # | disease | 4 | 0 | 0 | 1 | 1 | # | 10 | Small | aldex2 | 41 |
| # | sex | 2 | 0 | 0 | 1 | 1 | # | 10 | Small | aldex2 | 41 |
| # | time | 2 | 1 | 0 | 0.666666667 | 1 | # | 10 | Small | aldex2 | 41 |
| # | interaction | 0 | 0 | 2 | 0 | 0 | # | 10 | Small | aldex2 | 41 |
| # | disease | 4 | 6 | 0 | 0.4 | 1 | # | 10 | Baseline | paralle | 41 |
| # | sex | 2 | 5 | 0 | 0.285714286 | 1 | # | 10 | Baseline | paralle | 41 |
| # | time | 2 | 2 | 0 | 0.5 | 1 | # | 10 | Baseline | paralle | 41 |
| # | interaction | 2 | 5 | 0 | 0.285714286 | 1 | # | 10 | Baseline | paralle | 41 |
| # | disease | 4 | 2 | 0 | 0.666666667 | 1 | # | 10 | Baseline | ancom | 41 |
| # | sex | 2 | 0 | 0 | 1 | 1 | # | 10 | Baseline | ancom | 41 |
| # | time | 2 | 1 | 0 | 0.666666667 | 1 | # | 10 | Baseline | ancom | 41 |
| # | interaction | 0 | 0 | 2 | 0 | 0 | # | 10 | Baseline | ancom | 41 |
| # | disease | 4 | 13 | 0 | 0.235294118 | 1 | # | 10 | Baseline | aldex2 | 41 |
| # | sex | 2 | 9 | 0 | 0.181818182 | 1 | # | 10 | Baseline | aldex2 | 41 |
| # | time | 2 | 12 | 0 | 0.142857143 | 1 | # | 10 | Baseline | aldex2 | 41 |
| # | interaction | 0 | 0 | 2 | 0 | 0 | # | 10 | Baseline | aldex2 | 41 |
| # | disease | 4 | 0 | 0 | 1 | 1 | # | 10 | Small | paralle | 41 |
| # | sex | 2 | 0 | 0 | 1 | 1 | # | 10 | Small | paralle | 41 |
| # | time | 2 | 0 | 0 | 1 | 1 | # | 10 | Small | paralle | 41 |
| # | interaction | 2 | 0 | 0 | 1 | 1 | # | 10 | Small | paralle | 41 |
| # | disease | 4 | 1 | 0 | 0.8 | 1 | # | 10 | Small | ancom | 41 |
| # | sex | 2 | 0 | 0 | 1 | 1 | # | 10 | Small | ancom | 41 |
| # | time | 2 | 2 | 0 | 0.5 | 1 | # | 10 | Small | ancom | 41 |
| # | interaction | 0 | 0 | 2 | 0 | 0 | # | 10 | Small | ancom | 41 |
| # | disease | 4 | 1 | 0 | 0.8 | 1 | # | 10 | Small | aldex2 | 41 |
| # | sex | 2 | 0 | 0 | 1 | 1 | # | 10 | Small | aldex2 | 41 |
| # | time | 2 | 1 | 0 | 0.666666667 | 1 | # | 10 | Small | aldex2 | 41 |
| # | interaction | 0 | 0 | 2 | 0 | 0 | # | 10 | Small | aldex2 | 41 |
| # | disease | 4 | 4 | 0 | 0.5 | 1 | # | 10 | Baseline | paralle | 41 |
| # | sex | 2 | 3 | 0 | 0.4 | 1 | # | 10 | Baseline | paralle | 41 |
| # | time | 2 | 4 | 0 | 0.333333333 | 1 | # | 10 | Baseline | paralle | 41 |
| # | interaction | 2 | 3 | 0 | 0.4 | 1 | # | 10 | Baseline | paralle | 41 |
| # | disease | 4 | 2 | 0 | 0.666666667 | 1 | # | 10 | Baseline | ancom | 41 |
| # | sex | 2 | 0 | 0 | 1 | 1 | # | 10 | Baseline | ancom | 41 |
| # | time | 2 | 2 | 0 | 0.5 | 1 | # | 10 | Baseline | ancom | 41 |
| # | interaction | 0 | 0 | 2 | 0 | 0 | # | 10 | Baseline | ancom | 41 |
| # | disease | 4 | 11 | 0 | 0.266666667 | 1 | # | 10 | Baseline | aldex2 | 41 |
| # | sex | 2 | 8 | 0 | 0.2 | 1 | # | 10 | Baseline | aldex2 | 41 |
| # | time | 2 | 9 | 0 | 0.181818182 | 1 | # | 10 | Baseline | aldex2 | 41 |
| # | interaction | 0 | 0 | 2 | 0 | 0 | # | 10 | Baseline | aldex2 | 41 |
| # | disease | 4 | 0 | 0 | 1 | 1 | # | 10 | Small | paralle | 41 |
| # | sex | 2 | 0 | 0 | 1 | 1 | # | 10 | Small | paralle | 41 |
| # | time | 2 | 0 | 0 | 1 | 1 | # | 10 | Small | paralle | 41 |
| # | interaction | 2 | 0 | 0 | 1 | 1 | # | 10 | Small | paralle | 41 |
| # | disease | 4 | 2 | 0 | 0.666666667 | 1 | # | 10 | Small | ancom | 41 |
| # | sex | 2 | 0 | 0 | 1 | 1 | # | 10 | Small | ancom | 41 |
| # | time | 2 | 2 | 0 | 0.5 | 1 | # | 10 | Small | ancom | 41 |
| # | interaction | 0 | 0 | 2 | 0 | 0 | # | 10 | Small | ancom | 41 |

| | | | | | | | | | | | |
|---|---|---|---|---|---|---|---|---|---|---|---|
| # | disease | 4 | 2 | 0 | 0.666666667 | 1 | # | 10 | Small | aldex2 | 41 |
| # | sex | 2 | 0 | 0 | 1 | 1 | # | 10 | Small | aldex2 | 41 |
| # | time | 2 | 1 | 0 | 0.666666667 | 1 | # | 10 | Small | aldex2 | 41 |
| # | interaction | 0 | 0 | 2 | 0 | 0 | # | 10 | Small | aldex2 | 41 |
| # | disease | 4 | 4 | 0 | 0.5 | 1 | # | 10 | Baseline | paralle | 41 |
| # | sex | 2 | 2 | 0 | 0.5 | 1 | # | 10 | Baseline | paralle | 41 |
| # | time | 2 | 1 | 0 | 0.666666667 | 1 | # | 10 | Baseline | paralle | 41 |
| # | interaction | 2 | 4 | 0 | 0.333333333 | 1 | # | 10 | Baseline | paralle | 41 |
| # | disease | 4 | 2 | 0 | 0.666666667 | 1 | # | 10 | Baseline | ancom | 41 |
| # | sex | 2 | 0 | 0 | 1 | 1 | # | 10 | Baseline | ancom | 41 |
| # | time | 2 | 2 | 0 | 0.5 | 1 | # | 10 | Baseline | ancom | 41 |
| # | interaction | 0 | 0 | 2 | 0 | 0 | # | 10 | Baseline | ancom | 41 |
| # | disease | 4 | 19 | 0 | 0.173913043 | 1 | # | 10 | Baseline | aldex2 | 41 |
| # | sex | 2 | 10 | 0 | 0.166666667 | 1 | # | 10 | Baseline | aldex2 | 41 |
| # | time | 2 | 13 | 0 | 0.133333333 | 1 | # | 10 | Baseline | aldex2 | 41 |
| # | interaction | 0 | 0 | 2 | 0 | 0 | # | 10 | Baseline | aldex2 | 41 |
| # | disease | 4 | 0 | 0 | 1 | 1 | # | 10 | Small | paralle | 41 |
| # | sex | 2 | 0 | 0 | 1 | 1 | # | 10 | Small | paralle | 41 |
| # | time | 2 | 0 | 0 | 1 | 1 | # | 10 | Small | paralle | 41 |
| # | interaction | 2 | 0 | 0 | 1 | 1 | # | 10 | Small | paralle | 41 |
| # | disease | 4 | 2 | 0 | 0.666666667 | 1 | # | 10 | Small | ancom | 41 |
| # | sex | 2 | 0 | 0 | 1 | 1 | # | 10 | Small | ancom | 41 |
| # | time | 2 | 2 | 0 | 0.5 | 1 | # | 10 | Small | ancom | 41 |
| # | interaction | 0 | 0 | 2 | 0 | 0 | # | 10 | Small | ancom | 41 |
| # | disease | 4 | 3 | 0 | 0.571428571 | 1 | # | 10 | Small | aldex2 | 41 |
| # | sex | 2 | 0 | 0 | 1 | 1 | # | 10 | Small | aldex2 | 41 |
| # | time | 2 | 2 | 0 | 0.5 | 1 | # | 10 | Small | aldex2 | 41 |
| # | interaction | 0 | 0 | 2 | 0 | 0 | # | 10 | Small | aldex2 | 41 |
| # | disease | 7 | 10 | 0 | 0.411764706 | 1 | # | 20 | Baseline | paralle | 41 |
| # | sex | 4 | 10 | 0 | 0.285714286 | 1 | # | 20 | Baseline | paralle | 41 |
| # | time | 5 | 8 | 0 | 0.384615385 | 1 | # | 20 | Baseline | paralle | 41 |
| # | interaction | 4 | 9 | 0 | 0.307692308 | 1 | # | 20 | Baseline | paralle | 41 |
| # | disease | 7 | 2 | 0 | 0.777777778 | 1 | # | 20 | Baseline | ancom | 41 |
| # | sex | 4 | 0 | 0 | 1 | 1 | # | 20 | Baseline | ancom | 41 |
| # | time | 5 | 0 | 0 | 1 | 1 | # | 20 | Baseline | ancom | 41 |
| # | interaction | 0 | 0 | 4 | 0 | 0 | # | 20 | Baseline | ancom | 41 |
| # | disease | 7 | 19 | 0 | 0.269230769 | 1 | # | 20 | Baseline | aldex2 | 41 |
| # | sex | 4 | 7 | 0 | 0.363636364 | 1 | # | 20 | Baseline | aldex2 | 41 |
| # | time | 5 | 14 | 0 | 0.263157895 | 1 | # | 20 | Baseline | aldex2 | 41 |
| # | interaction | 0 | 0 | 4 | 0 | 0 | # | 20 | Baseline | aldex2 | 41 |
| # | disease | 7 | 0 | 0 | 1 | 1 | # | 20 | Small | paralle | 41 |
| # | sex | 4 | 0 | 0 | 1 | 1 | # | 20 | Small | paralle | 41 |
| # | time | 5 | 1 | 0 | 0.833333333 | 1 | # | 20 | Small | paralle | 41 |
| # | interaction | 1 | 1 | 3 | 0.5 | 0.25 | # | 20 | Small | paralle | 41 |
| # | disease | 7 | 2 | 0 | 0.777777778 | 1 | # | 20 | Small | ancom | 41 |
| # | sex | 4 | 0 | 0 | 1 | 1 | # | 20 | Small | ancom | 41 |
| # | time | 5 | 0 | 0 | 1 | 1 | # | 20 | Small | ancom | 41 |
| # | interaction | 0 | 0 | 4 | 0 | 0 | # | 20 | Small | ancom | 41 |
| # | disease | 7 | 2 | 0 | 0.777777778 | 1 | # | 20 | Small | aldex2 | 41 |
| # | sex | 4 | 0 | 0 | 1 | 1 | # | 20 | Small | aldex2 | 41 |
| # | time | 5 | 1 | 0 | 0.833333333 | 1 | # | 20 | Small | aldex2 | 41 |
| # | interaction | 0 | 0 | 4 | 0 | 0 | # | 20 | Small | aldex2 | 41 |
| # | disease | 7 | 11 | 0 | 0.388888889 | 1 | # | 20 | Baseline | paralle | 41 |
| # | sex | 4 | 7 | 0 | 0.363636364 | 1 | # | 20 | Baseline | paralle | 41 |
| # | time | 5 | 8 | 0 | 0.384615385 | 1 | # | 20 | Baseline | paralle | 41 |
| # | interaction | 4 | 8 | 0 | 0.333333333 | 1 | # | 20 | Baseline | paralle | 41 |
| # | disease | 7 | 5 | 0 | 0.583333333 | 1 | # | 20 | Baseline | ancom | 41 |
| # | sex | 4 | 0 | 0 | 1 | 1 | # | 20 | Baseline | ancom | 41 |
| # | time | 5 | 3 | 0 | 0.625 | 1 | # | 20 | Baseline | ancom | 41 |
| # | interaction | 0 | 0 | 4 | 0 | 0 | # | 20 | Baseline | ancom | 41 |
| # | disease | 7 | 37 | 0 | 0.159090909 | 1 | # | 20 | Baseline | aldex2 | 41 |
| # | sex | 4 | 8 | 0 | 0.333333333 | 1 | # | 20 | Baseline | aldex2 | 41 |
| # | time | 5 | 25 | 0 | 0.166666667 | 1 | # | 20 | Baseline | aldex2 | 41 |
| # | interaction | 0 | 0 | 4 | 0 | 0 | # | 20 | Baseline | aldex2 | 41 |
| # | disease | 7 | 0 | 0 | 1 | 1 | # | 20 | Small | paralle | 41 |
| # | sex | 4 | 0 | 0 | 1 | 1 | # | 20 | Small | paralle | 41 |
| # | time | 5 | 2 | 0 | 0.714285714 | 1 | # | 20 | Small | paralle | 41 |
| # | interaction | 4 | 0 | 0 | 1 | 1 | # | 20 | Small | paralle | 41 |
| # | disease | 7 | 5 | 0 | 0.583333333 | 1 | # | 20 | Small | ancom | 41 |
| # | sex | 4 | 0 | 0 | 1 | 1 | # | 20 | Small | ancom | 41 |
| # | time | 5 | 3 | 0 | 0.625 | 1 | # | 20 | Small | ancom | 41 |
| # | interaction | 0 | 0 | 4 | 0 | 0 | # | 20 | Small | ancom | 41 |
| # | disease | 7 | 4 | 0 | 0.636363636 | 1 | # | 20 | Small | aldex2 | 41 |
| # | sex | 4 | 0 | 0 | 1 | 1 | # | 20 | Small | aldex2 | 41 |
| # | time | 5 | 4 | 0 | 0.555555556 | 1 | # | 20 | Small | aldex2 | 41 |
| # | interaction | 0 | 0 | 4 | 0 | 0 | # | 20 | Small | aldex2 | 41 |
| # | disease | 7 | 8 | 0 | 0.466666667 | 1 | # | 20 | Baseline | paralle | 41 |
| # | sex | 4 | 5 | 0 | 0.444444444 | 1 | # | 20 | Baseline | paralle | 41 |
| # | time | 5 | 8 | 0 | 0.384615385 | 1 | # | 20 | Baseline | paralle | 41 |
| # | interaction | 4 | 4 | 0 | 0.5 | 1 | # | 20 | Baseline | paralle | 41 |
| # | disease | 7 | 5 | 0 | 0.583333333 | 1 | # | 20 | Baseline | ancom | 41 |
| # | sex | 4 | 0 | 0 | 1 | 1 | # | 20 | Baseline | ancom | 41 |
| # | time | 5 | 4 | 0 | 0.555555556 | 1 | # | 20 | Baseline | ancom | 41 |
| # | interaction | 0 | 0 | 4 | 0 | 0 | # | 20 | Baseline | ancom | 41 |
| # | disease | 7 | 50 | 0 | 0.122807018 | 1 | # | 20 | Baseline | aldex2 | 41 |
| # | sex | 4 | 14 | 0 | 0.222222222 | 1 | # | 20 | Baseline | aldex2 | 41 |
| # | time | 5 | 33 | 0 | 0.131578947 | 1 | # | 20 | Baseline | aldex2 | 41 |
| # | interaction | 0 | 0 | 4 | 0 | 0 | # | 20 | Baseline | aldex2 | 41 |
| # | disease | 7 | 0 | 0 | 1 | 1 | # | 20 | Small | paralle | 41 |
| # | sex | 4 | 0 | 0 | 1 | 1 | # | 20 | Small | paralle | 41 |
| # | time | 5 | 1 | 0 | 0.833333333 | 1 | # | 20 | Small | paralle | 41 |
| # | interaction | 4 | 0 | 0 | 1 | 1 | # | 20 | Small | paralle | 41 |
| # | disease | 7 | 5 | 0 | 0.583333333 | 1 | # | 20 | Small | ancom | 41 |
| # | sex | 4 | 0 | 0 | 1 | 1 | # | 20 | Small | ancom | 41 |
| # | time | 5 | 4 | 0 | 0.555555556 | 1 | # | 20 | Small | ancom | 41 |
| # | interaction | 0 | 0 | 4 | 0 | 0 | # | 20 | Small | ancom | 41 |
| # | disease | 7 | 6 | 0 | 0.538461538 | 1 | # | 20 | Small | aldex2 | 41 |
| # | sex | 4 | 0 | 0 | 1 | 1 | # | 20 | Small | aldex2 | 41 |
| # | time | 5 | 5 | 0 | 0.5 | 1 | # | 20 | Small | aldex2 | 41 |
| # | interaction | 0 | 0 | 4 | 0 | 0 | # | 20 | Small | aldex2 | 41 |
| # | disease | 7 | 12 | 0 | 0.368421053 | 1 | # | 20 | Baseline | paralle | 41 |
| # | sex | 4 | 5 | 0 | 0.444444444 | 1 | # | 20 | Baseline | paralle | 41 |
| # | time | 5 | 7 | 0 | 0.416666667 | 1 | # | 20 | Baseline | paralle | 41 |
| # | interaction | 4 | 2 | 0 | 0.666666667 | 1 | # | 20 | Baseline | paralle | 41 |
| # | disease | 7 | 5 | 0 | 0.583333333 | 1 | # | 20 | Baseline | ancom | 41 |
| # | sex | 4 | 0 | 0 | 1 | 1 | # | 20 | Baseline | ancom | 41 |
| # | time | 5 | 4 | 0 | 0.555555556 | 1 | # | 20 | Baseline | ancom | 41 |
| # | interaction | 0 | 0 | 4 | 0 | 0 | # | 20 | Baseline | ancom | 41 |
| # | disease | 7 | 59 | 0 | 0.106060606 | 1 | # | 20 | Baseline | aldex2 | 41 |
| # | sex | 4 | 20 | 0 | 0.166666667 | 1 | # | 20 | Baseline | aldex2 | 41 |
| # | time | 5 | 45 | 0 | 0.1 | 1 | # | 20 | Baseline | aldex2 | 41 |
| # | interaction | 0 | 0 | 4 | 0 | 0 | # | 20 | Baseline | aldex2 | 41 |
| # | disease | 7 | 0 | 0 | 1 | 1 | # | 20 | Small | paralle | 41 |
| # | sex | 4 | 0 | 0 | 1 | 1 | # | 20 | Small | paralle | 41 |
| # | time | 5 | 0 | 0 | 1 | 1 | # | 20 | Small | paralle | 41 |
| # | interaction | 4 | 0 | 0 | 1 | 1 | # | 20 | Small | paralle | 41 |
| # | disease | 7 | 5 | 0 | 0.583333333 | 1 | # | 20 | Small | ancom | 41 |
| # | sex | 4 | 0 | 0 | 1 | 1 | # | 20 | Small | ancom | 41 |
| # | time | 5 | 4 | 0 | 0.555555556 | 1 | # | 20 | Small | ancom | 41 |
| # | interaction | 0 | 0 | 4 | 0 | 0 | # | 20 | Small | ancom | 41 |
| # | disease | 7 | 8 | 0 | 0.466666667 | 1 | # | 20 | Small | aldex2 | 41 |
| # | sex | 4 | 0 | 0 | 1 | 1 | # | 20 | Small | aldex2 | 41 |
| # | time | 5 | 9 | 0 | 0.357142857 | 1 | # | 20 | Small | aldex2 | 41 |
| # | interaction | 0 | 0 | 4 | 0 | 0 | # | 20 | Small | aldex2 | 41 |
| # | disease | 4 | 11 | 0 | 0.266666667 | 1 | # | 10 | Baseline | paralle | 42 |
| # | sex | 2 | 9 | 0 | 0.181818182 | 1 | # | 10 | Baseline | paralle | 42 |
| # | time | 2 | 12 | 0 | 0.142857143 | 1 | # | 10 | Baseline | paralle | 42 |
| # | interaction | 2 | 18 | 0 | 0.1 | 1 | # | 10 | Baseline | paralle | 42 |
| # | disease | 4 | 0 | 0 | 1 | 1 | # | 10 | Baseline | ancom | 42 |
| # | sex | 2 | 0 | 0 | 1 | 1 | # | 10 | Baseline | ancom | 42 |
| # | time | 2 | 0 | 0 | 1 | 1 | # | 10 | Baseline | ancom | 42 |
| # | interaction | 0 | 0 | 2 | 0 | 0 | # | 10 | Baseline | ancom | 42 |
| # | disease | 4 | 10 | 0 | 0.285714286 | 1 | # | 10 | Baseline | aldex2 | 42 |
| # | sex | 2 | 7 | 0 | 0.222222222 | 1 | # | 10 | Baseline | aldex2 | 42 |
| # | time | 2 | 11 | 0 | 0.153846154 | 1 | # | 10 | Baseline | aldex2 | 42 |
| # | interaction | 0 | 0 | 2 | 0 | 0 | # | 10 | Baseline | aldex2 | 42 |
| # | disease | 4 | 0 | 0 | 1 | 1 | # | 10 | Small | paralle | 42 |
| # | sex | 2 | 0 | 0 | 1 | 1 | # | 10 | Small | paralle | 42 |
| # | time | 2 | 0 | 0 | 1 | 1 | # | 10 | Small | paralle | 42 |
| # | interaction | 1 | 0 | 1 | 1 | 0.5 | # | 10 | Small | paralle | 42 |
| # | disease | 4 | 0 | 0 | 1 | 1 | # | 10 | Small | ancom | 42 |
| # | sex | 2 | 0 | 0 | 1 | 1 | # | 10 | Small | ancom | 42 |
| # | time | 2 | 0 | 0 | 1 | 1 | # | 10 | Small | ancom | 42 |
| # | interaction | 0 | 0 | 2 | 0 | 0 | # | 10 | Small | ancom | 42 |
| # | disease | 4 | 1 | 0 | 0.8 | 1 | # | 10 | Small | aldex2 | 42 |
| # | sex | 2 | 0 | 0 | 1 | 1 | # | 10 | Small | aldex2 | 42 |
| # | time | 2 | 0 | 0 | 1 | 1 | # | 10 | Small | aldex2 | 42 |
| # | interaction | 0 | 0 | 2 | 0 | 0 | # | 10 | Small | aldex2 | 42 |
| # | disease | 4 | 9 | 0 | 0.307692308 | 1 | # | 10 | Baseline | paralle | 42 |
| # | sex | 2 | 6 | 0 | 0.25 | 1 | # | 10 | Baseline | paralle | 42 |
| # | time | 2 | 9 | 0 | 0.181818182 | 1 | # | 10 | Baseline | paralle | 42 |
| # | interaction | 2 | 10 | 0 | 0.166666667 | 1 | # | 10 | Baseline | paralle | 42 |
| # | disease | 4 | 2 | 0 | 0.666666667 | 1 | # | 10 | Baseline | ancom | 42 |
| # | sex | 2 | 0 | 0 | 1 | 1 | # | 10 | Baseline | ancom | 42 |
| # | time | 2 | 1 | 0 | 0.666666667 | 1 | # | 10 | Baseline | ancom | 42 |
| # | interaction | 0 | 0 | 2 | 0 | 0 | # | 10 | Baseline | ancom | 42 |
| # | disease | 4 | 16 | 0 | 0.2 | 1 | # | 10 | Baseline | aldex2 | 42 |
| # | sex | 2 | 7 | 0 | 0.222222222 | 1 | # | 10 | Baseline | aldex2 | 42 |
| # | time | 2 | 7 | 0 | 0.222222222 | 1 | # | 10 | Baseline | aldex2 | 42 |
| # | interaction | 0 | 0 | 2 | 0 | 0 | # | 10 | Baseline | aldex2 | 42 |
| # | disease | 4 | 1 | 0 | 0.8 | 1 | # | 10 | Small | paralle | 42 |
| # | sex | 2 | 0 | 0 | 1 | 1 | # | 10 | Small | paralle | 42 |
| # | time | 2 | 0 | 0 | 1 | 1 | # | 10 | Small | paralle | 42 |
| # | interaction | 2 | 0 | 0 | 1 | 1 | # | 10 | Small | paralle | 42 |
| # | disease | 4 | 2 | 0 | 0.666666667 | 1 | # | 10 | Small | ancom | 42 |
| # | sex | 2 | 0 | 0 | 1 | 1 | # | 10 | Small | ancom | 42 |
| # | time | 2 | 1 | 0 | 0.666666667 | 1 | # | 10 | Small | ancom | 42 |
| # | interaction | 0 | 0 | 2 | 0 | 0 | # | 10 | Small | ancom | 42 |
| # | disease | 4 | 2 | 0 | 0.666666667 | 1 | # | 10 | Small | aldex2 | 42 |
| # | sex | 2 | 0 | 0 | 1 | 1 | # | 10 | Small | aldex2 | 42 |
| # | time | 2 | 1 | 0 | 0.666666667 | 1 | # | 10 | Small | aldex2 | 42 |
| # | interaction | 0 | 0 | 2 | 0 | 0 | # | 10 | Small | aldex2 | 42 |
| # | disease | 4 | 7 | 0 | 0.363636364 | 1 | # | 10 | Baseline | paralle | 42 |
| # | sex | 2 | 2 | 0 | 0.5 | 1 | # | 10 | Baseline | paralle | 42 |
| # | time | 2 | 6 | 0 | 0.25 | 1 | # | 10 | Baseline | paralle | 42 |
| # | interaction | 2 | 9 | 0 | 0.181818182 | 1 | # | 10 | Baseline | paralle | 42 |
| # | disease | 4 | 2 | 0 | 0.666666667 | 1 | # | 10 | Baseline | ancom | 42 |
| # | sex | 2 | 0 | 0 | 1 | 1 | # | 10 | Baseline | ancom | 42 |
| # | time | 2 | 2 | 0 | 0.5 | 1 | # | 10 | Baseline | ancom | 42 |
| # | interaction | 0 | 0 | 2 | 0 | 0 | # | 10 | Baseline | ancom | 42 |
| # | disease | 4 | 24 | 0 | 0.142857143 | 1 | # | 10 | Baseline | aldex2 | 42 |
| # | sex | 2 | 5 | 0 | 0.285714286 | 1 | # | 10 | Baseline | aldex2 | 42 |
| # | time | 2 | 14 | 0 | 0.125 | 1 | # | 10 | Baseline | aldex2 | 42 |
| # | interaction | 0 | 0 | 2 | 0 | 0 | # | 10 | Baseline | aldex2 | 42 |
| # | disease | 4 | 0 | 0 | 1 | 1 | # | 10 | Small | paralle | 42 |
| # | sex | 2 | 0 | 0 | 1 | 1 | # | 10 | Small | paralle | 42 |
| # | time | 2 | 0 | 0 | 1 | 1 | # | 10 | Small | paralle | 42 |
| # | interaction | 2 | 0 | 0 | 1 | 1 | # | 10 | Small | paralle | 42 |
| # | disease | 4 | 2 | 0 | 0.666666667 | 1 | # | 10 | Small | ancom | 42 |
| # | sex | 2 | 0 | 0 | 1 | 1 | # | 10 | Small | ancom | 42 |
| # | time | 2 | 2 | 0 | 0.5 | 1 | # | 10 | Small | ancom | 42 |
| # | interaction | 0 | 0 | 2 | 0 | 0 | # | 10 | Small | ancom | 42 |
| # | disease | 4 | 3 | 0 | 0.571428571 | 1 | # | 10 | Small | aldex2 | 42 |
| # | sex | 2 | 1 | 0 | 0.666666667 | 1 | # | 10 | Small | aldex2 | 42 |
| # | time | 2 | 2 | 0 | 0.5 | 1 | # | 10 | Small | aldex2 | 42 |
| # | interaction | 0 | 0 | 2 | 0 | 0 | # | 10 | Small | aldex2 | 42 |
| # | disease | 4 | 5 | 0 | 0.444444444 | 1 | # | 10 | Baseline | paralle | 42 |
| # | sex | 2 | 2 | 0 | 0.5 | 1 | # | 10 | Baseline | paralle | 42 |
| # | time | 2 | 3 | 0 | 0.4 | 1 | # | 10 | Baseline | paralle | 42 |
| # | interaction | 2 | 2 | 0 | 0.5 | 1 | # | 10 | Baseline | paralle | 42 |
| # | disease | 4 | 2 | 0 | 0.666666667 | 1 | # | 10 | Baseline | ancom | 42 |
| # | sex | 2 | 0 | 0 | 1 | 1 | # | 10 | Baseline | ancom | 42 |
| # | time | 2 | 2 | 0 | 0.5 | 1 | # | 10 | Baseline | ancom | 42 |
| # | interaction | 0 | 0 | 2 | 0 | 0 | # | 10 | Baseline | ancom | 42 |
| # | disease | 4 | 27 | 0 | 0.129032258 | 1 | # | 10 | Baseline | aldex2 | 42 |
| # | sex | 2 | 5 | 0 | 0.285714286 | 1 | # | 10 | Baseline | aldex2 | 42 |
| # | time | 2 | 14 | 0 | 0.125 | 1 | # | 10 | Baseline | aldex2 | 42 |
| # | interaction | 0 | 0 | 2 | 0 | 0 | # | 10 | Baseline | aldex2 | 42 |
| # | disease | 4 | 1 | 0 | 0.8 | 1 | # | 10 | Small | paralle | 42 |
| # | sex | 2 | 0 | 0 | 1 | 1 | # | 10 | Small | paralle | 42 |
| # | time | 2 | 0 | 0 | 1 | 1 | # | 10 | Small | paralle | 42 |
| # | interaction | 2 | 0 | 0 | 1 | 1 | # | 10 | Small | paralle | 42 |
| # | disease | 4 | 2 | 0 | 0.666666667 | 1 | # | 10 | Small | ancom | 42 |
| # | sex | 2 | 0 | 0 | 1 | 1 | # | 10 | Small | ancom | 42 |
| # | time | 2 | 2 | 0 | 0.5 | 1 | # | 10 | Small | ancom | 42 |
| # | interaction | 0 | 0 | 2 | 0 | 0 | # | 10 | Small | ancom | 42 |
| # | disease | 4 | 4 | 0 | 0.5 | 1 | # | 10 | Small | aldex2 | 42 |
| # | sex | 2 | 1 | 0 | 0.666666667 | 1 | # | 10 | Small | aldex2 | 42 |
| # | time | 2 | 2 | 0 | 0.5 | 1 | # | 10 | Small | aldex2 | 42 |
| # | interaction | 0 | 0 | 2 | 0 | 0 | # | 10 | Small | aldex2 | 42 |
| # | disease | 7 | 8 | 0 | 0.466666667 | 1 | # | 20 | Baseline | paralle | 42 |
| # | sex | 4 | 6 | 0 | 0.4 | 1 | # | 20 | Baseline | paralle | 42 |
| # | time | 5 | 6 | 0 | 0.454545455 | 1 | # | 20 | Baseline | paralle | 42 |
| # | interaction | 4 | 11 | 0 | 0.266666667 | 1 | # | 20 | Baseline | paralle | 42 |
| # | disease | 7 | 2 | 0 | 0.777777778 | 1 | # | 20 | Baseline | ancom | 42 |
| # | sex | 4 | 0 | 0 | 1 | 1 | # | 20 | Baseline | ancom | 42 |
| # | time | 5 | 1 | 0 | 0.833333333 | 1 | # | 20 | Baseline | ancom | 42 |
| # | interaction | 0 | 0 | 4 | 0 | 0 | # | 20 | Baseline | ancom | 42 |
| # | disease | 7 | 22 | 0 | 0.24137931 | 1 | # | 20 | Baseline | aldex2 | 42 |
| # | sex | 4 | 5 | 0 | 0.444444444 | 1 | # | 20 | Baseline | aldex2 | 42 |
| # | time | 5 | 13 | 0 | 0.277777778 | 1 | # | 20 | Baseline | aldex2 | 42 |
| # | interaction | 0 | 0 | 4 | 0 | 0 | # | 20 | Baseline | aldex2 | 42 |
| # | disease | 7 | 0 | 0 | 1 | 1 | # | 20 | Small | paralle | 42 |
| # | sex | 4 | 0 | 0 | 1 | 1 | # | 20 | Small | paralle | 42 |
| # | time | 5 | 0 | 0 | 1 | 1 | # | 20 | Small | paralle | 42 |
| # | interaction | 1 | 0 | 3 | 1 | 0.25 | # | 20 | Small | paralle | 42 |
| # | disease | 7 | 2 | 0 | 0.777777778 | 1 | # | 20 | Small | ancom | 42 |
| # | sex | 4 | 0 | 0 | 1 | 1 | # | 20 | Small | ancom | 42 |
| # | time | 5 | 0 | 0 | 1 | 1 | # | 20 | Small | ancom | 42 |
| # | interaction | 0 | 0 | 4 | 0 | 0 | # | 20 | Small | ancom | 42 |
| # | disease | 7 | 4 | 0 | 0.636363636 | 1 | # | 20 | Small | aldex2 | 42 |
| # | sex | 4 | 0 | 0 | 1 | 1 | # | 20 | Small | aldex2 | 42 |
| # | time | 5 | 0 | 0 | 1 | 1 | # | 20 | Small | aldex2 | 42 |
| # | interaction | 0 | 0 | 4 | 0 | 0 | # | 20 | Small | aldex2 | 42 |
| # | disease | 7 | 10 | 0 | 0.411764706 | 1 | # | 20 | Baseline | paralle | 42 |
| # | sex | 4 | 8 | 0 | 0.333333333 | 1 | # | 20 | Baseline | paralle | 42 |
| # | time | 5 | 8 | 0 | 0.384615385 | 1 | # | 20 | Baseline | paralle | 42 |
| # | interaction | 4 | 9 | 0 | 0.307692308 | 1 | # | 20 | Baseline | paralle | 42 |
| # | disease | 7 | 4 | 0 | 0.636363636 | 1 | # | 20 | Baseline | ancom | 42 |
| # | sex | 4 | 0 | 0 | 1 | 1 | # | 20 | Baseline | ancom | 42 |
| # | time | 5 | 4 | 0 | 0.555555556 | 1 | # | 20 | Baseline | ancom | 42 |
| # | interaction | 0 | 0 | 4 | 0 | 0 | # | 20 | Baseline | ancom | 42 |
| # | disease | 7 | 44 | 0 | 0.137254902 | 1 | # | 20 | Baseline | aldex2 | 42 |
| # | sex | 4 | 15 | 0 | 0.210526316 | 1 | # | 20 | Baseline | aldex2 | 42 |
| # | time | 5 | 26 | 0 | 0.161290323 | 1 | # | 20 | Baseline | aldex2 | 42 |
| # | interaction | 0 | 0 | 4 | 0 | 0 | # | 20 | Baseline | aldex2 | 42 |
| # | disease | 7 | 0 | 0 | 1 | 1 | # | 20 | Small | paralle | 42 |
| # | sex | 4 | 0 | 0 | 1 | 1 | # | 20 | Small | paralle | 42 |
| # | time | 5 | 0 | 0 | 1 | 1 | # | 20 | Small | paralle | 42 |
| # | interaction | 2 | 0 | 2 | 1 | 0.5 | # | 20 | Small | paralle | 42 |
| # | disease | 7 | 4 | 0 | 0.636363636 | 1 | # | 20 | Small | ancom | 42 |
| # | sex | 4 | 0 | 0 | 1 | 1 | # | 20 | Small | ancom | 42 |
| # | time | 5 | 2 | 0 | 0.714285714 | 1 | # | 20 | Small | ancom | 42 |
| # | interaction | 0 | 0 | 4 | 0 | 0 | # | 20 | Small | ancom | 42 |
| # | disease | 7 | 5 | 0 | 0.583333333 | 1 | # | 20 | Small | aldex2 | 42 |
| # | sex | 4 | 1 | 0 | 0.8 | 1 | # | 20 | Small | aldex2 | 42 |
| # | time | 5 | 3 | 0 | 0.625 | 1 | # | 20 | Small | aldex2 | 42 |
| # | interaction | 0 | 0 | 4 | 0 | 0 | # | 20 | Small | aldex2 | 42 |
| # | disease | 7 | 11 | 0 | 0.388888889 | 1 | # | 20 | Baseline | paralle | 42 |
| # | sex | 4 | 5 | 0 | 0.444444444 | 1 | # | 20 | Baseline | paralle | 42 |
| # | time | 5 | 9 | 0 | 0.357142857 | 1 | # | 20 | Baseline | paralle | 42 |
| # | interaction | 4 | 6 | 0 | 0.4 | 1 | # | 20 | Baseline | paralle | 42 |
| # | disease | 7 | 4 | 0 | 0.636363636 | 1 | # | 20 | Baseline | ancom | 42 |
| # | sex | 4 | 0 | 0 | 1 | 1 | # | 20 | Baseline | ancom | 42 |
| # | time | 5 | 4 | 0 | 0.555555556 | 1 | # | 20 | Baseline | ancom | 42 |
| # | interaction | 0 | 0 | 4 | 0 | 0 | # | 20 | Baseline | ancom | 42 |
| # | disease | 7 | 57 | 0 | 0.109375 | 1 | # | 20 | Baseline | aldex2 | 42 |
| # | sex | 4 | 19 | 0 | 0.173913043 | 1 | # | 20 | Baseline | aldex2 | 42 |
| # | time | 5 | 38 | 0 | 0.11627907 | 1 | # | 20 | Baseline | aldex2 | 42 |
| # | interaction | 0 | 0 | 4 | 0 | 0 | # | 20 | Baseline | aldex2 | 42 |
| # | disease | 7 | 0 | 0 | 1 | 1 | # | 20 | Small | paralle | 42 |
| # | sex | 4 | 0 | 0 | 1 | 1 | # | 20 | Small | paralle | 42 |
| # | time | 5 | 0 | 0 | 1 | 1 | # | 20 | Small | paralle | 42 |
| # | interaction | 3 | 0 | 1 | 1 | 0.75 | # | 20 | Small | paralle | 42 |
| # | disease | 7 | 4 | 0 | 0.636363636 | 1 | # | 20 | Small | ancom | 42 |
| # | sex | 4 | 0 | 0 | 1 | 1 | # | 20 | Small | ancom | 42 |
| # | time | 5 | 4 | 0 | 0.555555556 | 1 | # | 20 | Small | ancom | 42 |
| # | interaction | 0 | 0 | 4 | 0 | 0 | # | 20 | Small | ancom | 42 |
| # | disease | 7 | 7 | 0 | 0.5 | 1 | # | 20 | Small | aldex2 | 42 |
| # | sex | 4 | 1 | 0 | 0.8 | 1 | # | 20 | Small | aldex2 | 42 |
| # | time | 5 | 4 | 0 | 0.555555556 | 1 | # | 20 | Small | aldex2 | 42 |
| # | interaction | 0 | 0 | 4 | 0 | 0 | # | 20 | Small | aldex2 | 42 |
| # | disease | 7 | 8 | 0 | 0.466666667 | 1 | # | 20 | Baseline | paralle | 42 |
| # | sex | 4 | 9 | 0 | 0.307692308 | 1 | # | 20 | Baseline | paralle | 42 |
| # | time | 5 | 11 | 0 | 0.3125 | 1 | # | 20 | Baseline | paralle | 42 |
| # | interaction | 4 | 5 | 0 | 0.444444444 | 1 | # | 20 | Baseline | paralle | 42 |
| # | disease | 7 | 4 | 0 | 0.636363636 | 1 | # | 20 | Baseline | ancom | 42 |
| # | sex | 4 | 0 | 0 | 1 | 1 | # | 20 | Baseline | ancom | 42 |
| # | time | 5 | 4 | 0 | 0.555555556 | 1 | # | 20 | Baseline | ancom | 42 |
| # | interaction | 0 | 0 | 4 | 0 | 0 | # | 20 | Baseline | ancom | 42 |
| # | disease | 7 | 59 | 0 | 0.106060606 | 1 | # | 20 | Baseline | aldex2 | 42 |
| # | sex | 4 | 23 | 0 | 0.148148148 | 1 | # | 20 | Baseline | aldex2 | 42 |
| # | time | 5 | 41 | 0 | 0.108695652 | 1 | # | 20 | Baseline | aldex2 | 42 |
| # | interaction | 0 | 0 | 4 | 0 | 0 | # | 20 | Baseline | aldex2 | 42 |
| # | disease | 7 | 0 | 0 | 1 | 1 | # | 20 | Small | paralle | 42 |
| # | sex | 4 | 0 | 0 | 1 | 1 | # | 20 | Small | paralle | 42 |
| # | time | 5 | 0 | 0 | 1 | 1 | # | 20 | Small | paralle | 42 |
| # | interaction | 4 | 0 | 0 | 1 | 1 | # | 20 | Small | paralle | 42 |
| # | disease | 7 | 4 | 0 | 0.636363636 | 1 | # | 20 | Small | ancom | 42 |
| # | sex | 4 | 0 | 0 | 1 | 1 | # | 20 | Small | ancom | 42 |
| # | time | 5 | 4 | 0 | 0.555555556 | 1 | # | 20 | Small | ancom | 42 |
| # | interaction | 0 | 0 | 4 | 0 | 0 | # | 20 | Small | ancom | 42 |
| # | disease | 7 | 10 | 0 | 0.411764706 | 1 | # | 20 | Small | aldex2 | 42 |
| # | sex | 4 | 3 | 0 | 0.571428571 | 1 | # | 20 | Small | aldex2 | 42 |
| # | time | 5 | 6 | 0 | 0.454545455 | 1 | # | 20 | Small | aldex2 | 42 |
| # | interaction | 0 | 0 | 4 | 0 | 0 | # | 20 | Small | aldex2 | 42 |
| # | disease | 4 | 5 | 0 | 0.444444444 | 1 | # | 10 | Baseline | paralle | 43 |
| # | sex | 2 | 7 | 0 | 0.222222222 | 1 | # | 10 | Baseline | paralle | 43 |
| # | time | 2 | 8 | 0 | 0.2 | 1 | # | 10 | Baseline | paralle | 43 |
| # | interaction | 2 | 7 | 0 | 0.222222222 | 1 | # | 10 | Baseline | paralle | 43 |
| # | disease | 4 | 0 | 0 | 1 | 1 | # | 10 | Baseline | ancom | 43 |
| # | sex | 2 | 0 | 0 | 1 | 1 | # | 10 | Baseline | ancom | 43 |
| # | time | 2 | 0 | 0 | 1 | 1 | # | 10 | Baseline | ancom | 43 |
| # | interaction | 0 | 0 | 2 | 0 | 0 | # | 10 | Baseline | ancom | 43 |
| # | disease | 4 | 14 | 0 | 0.222222222 | 1 | # | 10 | Baseline | aldex2 | 43 |
| # | sex | 2 | 4 | 0 | 0.333333333 | 1 | # | 10 | Baseline | aldex2 | 43 |
| # | time | 2 | 7 | 0 | 0.222222222 | 1 | # | 10 | Baseline | aldex2 | 43 |
| # | interaction | 0 | 0 | 2 | 0 | 0 | # | 10 | Baseline | aldex2 | 43 |
| # | disease | 4 | 1 | 0 | 0.8 | 1 | # | 10 | Small | paralle | 43 |
| # | sex | 2 | 0 | 0 | 1 | 1 | # | 10 | Small | paralle | 43 |
| # | time | 2 | 0 | 0 | 1 | 1 | # | 10 | Small | paralle | 43 |
| # | interaction | 1 | 1 | 1 | 0.5 | 0.5 | # | 10 | Small | paralle | 43 |
| # | disease | 4 | 0 | 0 | 1 | 1 | # | 10 | Small | ancom | 43 |
| # | sex | 2 | 0 | 0 | 1 | 1 | # | 10 | Small | ancom | 43 |
| # | time | 2 | 0 | 0 | 1 | 1 | # | 10 | Small | ancom | 43 |
| # | interaction | 0 | 0 | 2 | 0 | 0 | # | 10 | Small | ancom | 43 |
| # | disease | 4 | 1 | 0 | 0.8 | 1 | # | 10 | Small | aldex2 | 43 |

| | | | | | | | | | | | |
|---|---|---|---|---|---|---|---|---|---|---|---|
| # | sex | 2 | 0 | 0 | 1 | 1 | # | 10 | Small | aldex2 | 40 |
| # | time | 2 | 1 | 0 | 0.666666667 | 1 | # | 10 | Small | aldex2 | 40 |
| # | interaction | 0 | 0 | 2 | 0 | 0 | # | 10 | Small | aldex2 | 40 |
| # | disease | 4 | 6 | 0 | 0.4 | 1 | # | 10 | Baseline | paralle | 40 |
| # | sex | 2 | 3 | 0 | 0.4 | 1 | # | 10 | Baseline | paralle | 40 |
| # | time | 2 | 7 | 0 | 0.222222222 | 1 | # | 10 | Baseline | paralle | 40 |
| # | interaction | 2 | 3 | 0 | 0.4 | 1 | # | 10 | Baseline | paralle | 40 |
| # | disease | 4 | 2 | 0 | 0.666666667 | 1 | # | 10 | Baseline | ancom | 40 |
| # | sex | 2 | 0 | 0 | 1 | 1 | # | 10 | Baseline | ancom | 40 |
| # | time | 2 | 1 | 0 | 0.666666667 | 1 | # | 10 | Baseline | ancom | 40 |
| # | interaction | 0 | 0 | 2 | 0 | 0 | # | 10 | Baseline | ancom | 40 |
| # | disease | 4 | 16 | 0 | 0.2 | 1 | # | 10 | Baseline | aldex2 | 40 |
| # | sex | 2 | 10 | 0 | 0.166666667 | 1 | # | 10 | Baseline | aldex2 | 40 |
| # | time | 2 | 9 | 0 | 0.181818182 | 1 | # | 10 | Baseline | aldex2 | 40 |
| # | interaction | 0 | 0 | 2 | 0 | 0 | # | 10 | Baseline | aldex2 | 40 |
| # | disease | 4 | 0 | 0 | 1 | 1 | # | 10 | Small | paralle | 40 |
| # | sex | 2 | 0 | 0 | 1 | 1 | # | 10 | Small | paralle | 40 |
| # | time | 2 | 0 | 0 | 1 | 1 | # | 10 | Small | paralle | 40 |
| # | interaction | 2 | 0 | 0 | 1 | 1 | # | 10 | Small | paralle | 40 |
| # | disease | 4 | 2 | 0 | 0.666666667 | 1 | # | 10 | Small | ancom | 40 |
| # | sex | 2 | 0 | 0 | 1 | 1 | # | 10 | Small | ancom | 40 |
| # | time | 2 | 0 | 0 | 1 | 1 | # | 10 | Small | ancom | 40 |
| # | interaction | 0 | 0 | 2 | 0 | 0 | # | 10 | Small | ancom | 40 |
| # | disease | 4 | 2 | 0 | 0.666666667 | 1 | # | 10 | Small | aldex2 | 40 |
| # | sex | 2 | 0 | 0 | 1 | 1 | # | 10 | Small | aldex2 | 40 |
| # | time | 2 | 1 | 0 | 0.666666667 | 1 | # | 10 | Small | aldex2 | 40 |
| # | interaction | 0 | 0 | 2 | 0 | 0 | # | 10 | Small | aldex2 | 40 |
| # | disease | 4 | 4 | 0 | 0.5 | 1 | # | 10 | Baseline | paralle | 40 |
| # | sex | 2 | 3 | 0 | 0.4 | 1 | # | 10 | Baseline | paralle | 40 |
| # | time | 2 | 3 | 0 | 0.4 | 1 | # | 10 | Baseline | paralle | 40 |
| # | interaction | 2 | 3 | 0 | 0.4 | 1 | # | 10 | Baseline | paralle | 40 |
| # | disease | 4 | 2 | 0 | 0.666666667 | 1 | # | 10 | Baseline | ancom | 40 |
| # | sex | 2 | 0 | 0 | 1 | 1 | # | 10 | Baseline | ancom | 40 |
| # | time | 2 | 1 | 0 | 0.666666667 | 1 | # | 10 | Baseline | ancom | 40 |
| # | interaction | 0 | 0 | 2 | 0 | 0 | # | 10 | Baseline | ancom | 40 |
| # | disease | 4 | 23 | 0 | 0.148148148 | 1 | # | 10 | Baseline | aldex2 | 40 |
| # | sex | 2 | 7 | 0 | 0.222222222 | 1 | # | 10 | Baseline | aldex2 | 40 |
| # | time | 2 | 9 | 0 | 0.181818182 | 1 | # | 10 | Baseline | aldex2 | 40 |
| # | interaction | 0 | 0 | 2 | 0 | 0 | # | 10 | Baseline | aldex2 | 40 |
| # | disease | 4 | 0 | 0 | 1 | 1 | # | 10 | Small | paralle | 40 |
| # | sex | 2 | 0 | 0 | 1 | 1 | # | 10 | Small | paralle | 40 |
| # | time | 2 | 0 | 0 | 1 | 1 | # | 10 | Small | paralle | 40 |
| # | interaction | 2 | 0 | 0 | 1 | 1 | # | 10 | Small | paralle | 40 |
| # | disease | 4 | 2 | 0 | 0.666666667 | 1 | # | 10 | Small | ancom | 40 |
| # | sex | 2 | 0 | 0 | 1 | 1 | # | 10 | Small | ancom | 40 |
| # | time | 2 | 1 | 0 | 0.666666667 | 1 | # | 10 | Small | ancom | 40 |
| # | interaction | 0 | 0 | 2 | 0 | 0 | # | 10 | Small | ancom | 40 |
| # | disease | 4 | 2 | 0 | 0.666666667 | 1 | # | 10 | Small | aldex2 | 40 |
| # | sex | 2 | 0 | 0 | 1 | 1 | # | 10 | Small | aldex2 | 40 |
| # | time | 2 | 1 | 0 | 0.666666667 | 1 | # | 10 | Small | aldex2 | 40 |
| # | interaction | 0 | 0 | 2 | 0 | 0 | # | 10 | Small | aldex2 | 40 |
| # | disease | 4 | 2 | 0 | 0.666666667 | 1 | # | 10 | Baseline | paralle | 40 |
| # | sex | 2 | 1 | 0 | 0.666666667 | 1 | # | 10 | Baseline | paralle | 40 |
| # | time | 2 | 4 | 0 | 0.333333333 | 1 | # | 10 | Baseline | paralle | 40 |
| # | interaction | 2 | 2 | 0 | 0.5 | 1 | # | 10 | Baseline | paralle | 40 |
| # | disease | 4 | 2 | 0 | 0.666666667 | 1 | # | 10 | Baseline | ancom | 40 |
| # | sex | 2 | 0 | 0 | 1 | 1 | # | 10 | Baseline | ancom | 40 |
| # | time | 2 | 2 | 0 | 0.5 | 1 | # | 10 | Baseline | ancom | 40 |
| # | interaction | 0 | 0 | 2 | 0 | 0 | # | 10 | Baseline | ancom | 40 |
| # | disease | 4 | 26 | 0 | 0.133333333 | 1 | # | 10 | Baseline | aldex2 | 40 |
| # | sex | 2 | 11 | 0 | 0.153846154 | 1 | # | 10 | Baseline | aldex2 | 40 |
| # | time | 2 | 13 | 0 | 0.133333333 | 1 | # | 10 | Baseline | aldex2 | 40 |
| # | interaction | 0 | 0 | 2 | 0 | 0 | # | 10 | Baseline | aldex2 | 40 |
| # | disease | 4 | 0 | 0 | 1 | 1 | # | 10 | Small | paralle | 40 |
| # | sex | 2 | 0 | 0 | 1 | 1 | # | 10 | Small | paralle | 40 |
| # | time | 2 | 0 | 0 | 1 | 1 | # | 10 | Small | paralle | 40 |
| # | interaction | 2 | 0 | 0 | 1 | 1 | # | 10 | Small | paralle | 40 |
| # | disease | 4 | 2 | 0 | 0.666666667 | 1 | # | 10 | Small | ancom | 40 |
| # | sex | 2 | 0 | 0 | 1 | 1 | # | 10 | Small | ancom | 40 |
| # | time | 2 | 1 | 0 | 0.666666667 | 1 | # | 10 | Small | ancom | 40 |
| # | interaction | 0 | 0 | 2 | 0 | 0 | # | 10 | Small | ancom | 40 |
| # | disease | 4 | 3 | 0 | 0.571428571 | 1 | # | 10 | Small | aldex2 | 40 |
| # | sex | 2 | 0 | 0 | 1 | 1 | # | 10 | Small | aldex2 | 40 |
| # | time | 2 | 1 | 0 | 0.666666667 | 1 | # | 10 | Small | aldex2 | 40 |
| # | interaction | 0 | 0 | 2 | 0 | 0 | # | 10 | Small | aldex2 | 40 |
| # | disease | 7 | 8 | 0 | 0.466666667 | 1 | # | 20 | Baseline | paralle | 40 |
| # | sex | 4 | 7 | 0 | 0.363636364 | 1 | # | 20 | Baseline | paralle | 40 |
| # | time | 5 | 9 | 0 | 0.357142857 | 1 | # | 20 | Baseline | paralle | 40 |
| # | interaction | 4 | 10 | 0 | 0.285714286 | 1 | # | 20 | Baseline | paralle | 40 |
| # | disease | 7 | 3 | 0 | 0.7 | 1 | # | 20 | Baseline | ancom | 40 |
| # | sex | 4 | 0 | 0 | 1 | 1 | # | 20 | Baseline | ancom | 40 |
| # | time | 5 | 0 | 0 | 1 | 1 | # | 20 | Baseline | ancom | 40 |
| # | interaction | 0 | 0 | 4 | 0 | 0 | # | 20 | Baseline | ancom | 40 |
| # | disease | 7 | 21 | 0 | 0.25 | 1 | # | 20 | Baseline | aldex2 | 40 |
| # | sex | 4 | 8 | 0 | 0.333333333 | 1 | # | 20 | Baseline | aldex2 | 40 |
| # | time | 5 | 18 | 0 | 0.217391304 | 1 | # | 20 | Baseline | aldex2 | 40 |
| # | interaction | 0 | 0 | 4 | 0 | 0 | # | 20 | Baseline | aldex2 | 40 |
| # | disease | 7 | 1 | 0 | 0.875 | 1 | # | 20 | Small | paralle | 40 |
| # | sex | 4 | 0 | 0 | 1 | 1 | # | 20 | Small | paralle | 40 |
| # | time | 5 | 0 | 0 | 1 | 1 | # | 20 | Small | paralle | 40 |
| # | interaction | 2 | 1 | 2 | 0.666666667 | 0.5 | # | 20 | Small | paralle | 40 |
| # | disease | 7 | 1 | 0 | 0.875 | 1 | # | 20 | Small | ancom | 40 |
| # | sex | 4 | 0 | 0 | 1 | 1 | # | 20 | Small | ancom | 40 |
| # | time | 5 | 0 | 0 | 1 | 1 | # | 20 | Small | ancom | 40 |
| # | interaction | 0 | 0 | 4 | 0 | 0 | # | 20 | Small | ancom | 40 |
| # | disease | 7 | 2 | 0 | 0.777777778 | 1 | # | 20 | Small | aldex2 | 40 |
| # | sex | 4 | 0 | 0 | 1 | 1 | # | 20 | Small | aldex2 | 40 |
| # | time | 5 | 1 | 0 | 0.833333333 | 1 | # | 20 | Small | aldex2 | 40 |
| # | interaction | 0 | 0 | 4 | 0 | 0 | # | 20 | Small | aldex2 | 40 |
| # | disease | 7 | 11 | 0 | 0.388888889 | 1 | # | 20 | Baseline | paralle | 40 |
| # | sex | 4 | 6 | 0 | 0.4 | 1 | # | 20 | Baseline | paralle | 40 |
| # | time | 5 | 8 | 0 | 0.384615385 | 1 | # | 20 | Baseline | paralle | 40 |
| # | interaction | 3 | 10 | 1 | 0.230769231 | 0.75 | # | 20 | Baseline | paralle | 40 |
| # | disease | 7 | 4 | 0 | 0.636363636 | 1 | # | 20 | Baseline | ancom | 40 |
| # | sex | 4 | 0 | 0 | 1 | 1 | # | 20 | Baseline | ancom | 40 |
| # | time | 5 | 4 | 0 | 0.555555556 | 1 | # | 20 | Baseline | ancom | 40 |
| # | interaction | 0 | 0 | 4 | 0 | 0 | # | 20 | Baseline | ancom | 40 |
| # | disease | 7 | 33 | 0 | 0.175 | 1 | # | 20 | Baseline | aldex2 | 40 |
| # | sex | 4 | 13 | 0 | 0.235294118 | 1 | # | 20 | Baseline | aldex2 | 40 |
| # | time | 5 | 23 | 0 | 0.178571429 | 1 | # | 20 | Baseline | aldex2 | 40 |
| # | interaction | 0 | 0 | 4 | 0 | 0 | # | 20 | Baseline | aldex2 | 40 |
| # | disease | 7 | 1 | 0 | 0.875 | 1 | # | 20 | Small | paralle | 40 |
| # | sex | 4 | 0 | 0 | 1 | 1 | # | 20 | Small | paralle | 40 |
| # | time | 5 | 0 | 0 | 1 | 1 | # | 20 | Small | paralle | 40 |
| # | interaction | 2 | 2 | 2 | 0.5 | 0.5 | # | 20 | Small | paralle | 40 |
| # | disease | 7 | 3 | 0 | 0.7 | 1 | # | 20 | Small | ancom | 40 |
| # | sex | 4 | 0 | 0 | 1 | 1 | # | 20 | Small | ancom | 40 |
| # | time | 5 | 4 | 0 | 0.555555556 | 1 | # | 20 | Small | ancom | 40 |
| # | interaction | 0 | 0 | 4 | 0 | 0 | # | 20 | Small | ancom | 40 |
| # | disease | 7 | 4 | 0 | 0.636363636 | 1 | # | 20 | Small | aldex2 | 40 |
| # | sex | 4 | 0 | 0 | 1 | 1 | # | 20 | Small | aldex2 | 40 |
| # | time | 5 | 3 | 0 | 0.625 | 1 | # | 20 | Small | aldex2 | 40 |
| # | interaction | 0 | 0 | 4 | 0 | 0 | # | 20 | Small | aldex2 | 40 |
| # | disease | 7 | 14 | 0 | 0.333333333 | 1 | # | 20 | Baseline | paralle | 40 |
| # | sex | 4 | 6 | 0 | 0.4 | 1 | # | 20 | Baseline | paralle | 40 |
| # | time | 5 | 9 | 0 | 0.357142857 | 1 | # | 20 | Baseline | paralle | 40 |
| # | interaction | 4 | 7 | 0 | 0.363636364 | 1 | # | 20 | Baseline | paralle | 40 |
| # | disease | 7 | 4 | 0 | 0.636363636 | 1 | # | 20 | Baseline | ancom | 40 |
| # | sex | 4 | 0 | 0 | 1 | 1 | # | 20 | Baseline | ancom | 40 |
| # | time | 5 | 4 | 0 | 0.555555556 | 1 | # | 20 | Baseline | ancom | 40 |
| # | interaction | 0 | 0 | 4 | 0 | 0 | # | 20 | Baseline | ancom | 40 |
| # | disease | 7 | 50 | 0 | 0.122807018 | 1 | # | 20 | Baseline | aldex2 | 40 |
| # | sex | 4 | 14 | 0 | 0.222222222 | 1 | # | 20 | Baseline | aldex2 | 40 |
| # | time | 5 | 31 | 0 | 0.138888889 | 1 | # | 20 | Baseline | aldex2 | 40 |
| # | interaction | 0 | 0 | 4 | 0 | 0 | # | 20 | Baseline | aldex2 | 40 |
| # | disease | 7 | 1 | 0 | 0.875 | 1 | # | 20 | Small | paralle | 40 |
| # | sex | 4 | 0 | 0 | 1 | 1 | # | 20 | Small | paralle | 40 |
| # | time | 5 | 0 | 0 | 1 | 1 | # | 20 | Small | paralle | 40 |
| # | interaction | 3 | 0 | 1 | 1 | 0.75 | # | 20 | Small | paralle | 40 |
| # | disease | 7 | 4 | 0 | 0.636363636 | 1 | # | 20 | Small | ancom | 40 |
| # | sex | 4 | 0 | 0 | 1 | 1 | # | 20 | Small | ancom | 40 |
| # | time | 5 | 3 | 0 | 0.625 | 1 | # | 20 | Small | ancom | 40 |
| # | interaction | 0 | 0 | 4 | 0 | 0 | # | 20 | Small | ancom | 40 |
| # | disease | 7 | 5 | 0 | 0.583333333 | 1 | # | 20 | Small | aldex2 | 40 |
| # | sex | 4 | 0 | 0 | 1 | 1 | # | 20 | Small | aldex2 | 40 |
| # | time | 5 | 3 | 0 | 0.625 | 1 | # | 20 | Small | aldex2 | 40 |
| # | interaction | 0 | 0 | 4 | 0 | 0 | # | 20 | Small | aldex2 | 40 |
| # | disease | 7 | 10 | 0 | 0.411764706 | 1 | # | 20 | Baseline | paralle | 40 |
| # | sex | 4 | 5 | 0 | 0.444444444 | 1 | # | 20 | Baseline | paralle | 40 |
| # | time | 5 | 7 | 0 | 0.416666667 | 1 | # | 20 | Baseline | paralle | 40 |
| # | interaction | 4 | 7 | 0 | 0.363636364 | 1 | # | 20 | Baseline | paralle | 40 |
| # | disease | 7 | 4 | 0 | 0.636363636 | 1 | # | 20 | Baseline | ancom | 40 |
| # | sex | 4 | 0 | 0 | 1 | 1 | # | 20 | Baseline | ancom | 40 |
| # | time | 5 | 4 | 0 | 0.555555556 | 1 | # | 20 | Baseline | ancom | 40 |
| # | interaction | 0 | 0 | 4 | 0 | 0 | # | 20 | Baseline | ancom | 40 |
| # | disease | 7 | 60 | 0 | 0.104477612 | 1 | # | 20 | Baseline | aldex2 | 40 |
| # | sex | 4 | 24 | 0 | 0.142857143 | 1 | # | 20 | Baseline | aldex2 | 40 |
| # | time | 5 | 43 | 0 | 0.104166667 | 1 | # | 20 | Baseline | aldex2 | 40 |
| # | interaction | 0 | 0 | 4 | 0 | 0 | # | 20 | Baseline | aldex2 | 40 |
| # | disease | 7 | 1 | 0 | 0.875 | 1 | # | 20 | Small | paralle | 40 |
| # | sex | 4 | 0 | 0 | 1 | 1 | # | 20 | Small | paralle | 40 |
| # | time | 5 | 0 | 0 | 1 | 1 | # | 20 | Small | paralle | 40 |
| # | interaction | 4 | 0 | 0 | 1 | 1 | # | 20 | Small | paralle | 40 |
| # | disease | 7 | 4 | 0 | 0.636363636 | 1 | # | 20 | Small | ancom | 40 |
| # | sex | 4 | 0 | 0 | 1 | 1 | # | 20 | Small | ancom | 40 |
| # | time | 5 | 4 | 0 | 0.555555556 | 1 | # | 20 | Small | ancom | 40 |
| # | interaction | 0 | 0 | 4 | 0 | 0 | # | 20 | Small | ancom | 40 |
| # | disease | 7 | 13 | 0 | 0.35 | 1 | # | 20 | Small | aldex2 | 40 |
| # | sex | 4 | 1 | 0 | 0.8 | 1 | # | 20 | Small | aldex2 | 40 |
| # | time | 5 | 6 | 0 | 0.454545455 | 1 | # | 20 | Small | aldex2 | 40 |
| # | interaction | 0 | 0 | 4 | 0 | 0 | # | 20 | Small | aldex2 | 40 |
| # | disease | 4 | 12 | 0 | 0.25 | 1 | # | 10 | Baseline | paralle | 44 |
| # | sex | 2 | 6 | 0 | 0.25 | 1 | # | 10 | Baseline | paralle | 44 |
| # | time | 2 | 6 | 0 | 0.25 | 1 | # | 10 | Baseline | paralle | 44 |
| # | interaction | 2 | 9 | 0 | 0.181818182 | 1 | # | 10 | Baseline | paralle | 44 |
| # | disease | 4 | 1 | 0 | 0.8 | 1 | # | 10 | Baseline | ancom | 44 |
| # | sex | 2 | 0 | 0 | 1 | 1 | # | 10 | Baseline | ancom | 44 |
| # | time | 2 | 1 | 0 | 0.666666667 | 1 | # | 10 | Baseline | ancom | 44 |
| # | interaction | 0 | 0 | 2 | 0 | 0 | # | 10 | Baseline | ancom | 44 |
| # | disease | 4 | 20 | 0 | 0.166666667 | 1 | # | 10 | Baseline | aldex2 | 44 |
| # | sex | 2 | 5 | 0 | 0.285714286 | 1 | # | 10 | Baseline | aldex2 | 44 |
| # | time | 2 | 5 | 0 | 0.285714286 | 1 | # | 10 | Baseline | aldex2 | 44 |
| # | interaction | 0 | 0 | 2 | 0 | 0 | # | 10 | Baseline | aldex2 | 44 |
| # | disease | 4 | 0 | 0 | 1 | 1 | # | 10 | Small | paralle | 44 |
| # | sex | 2 | 0 | 0 | 1 | 1 | # | 10 | Small | paralle | 44 |
| # | time | 2 | 0 | 0 | 1 | 1 | # | 10 | Small | paralle | 44 |
| # | interaction | 2 | 0 | 0 | 1 | 1 | # | 10 | Small | paralle | 44 |
| # | disease | 4 | 1 | 0 | 0.8 | 1 | # | 10 | Small | ancom | 44 |
| # | sex | 2 | 0 | 0 | 1 | 1 | # | 10 | Small | ancom | 44 |
| # | time | 2 | 2 | 0 | 0.5 | 1 | # | 10 | Small | ancom | 44 |
| # | interaction | 0 | 0 | 2 | 0 | 0 | # | 10 | Small | ancom | 44 |
| # | disease | 4 | 1 | 0 | 0.8 | 1 | # | 10 | Small | aldex2 | 44 |
| # | sex | 2 | 0 | 0 | 1 | 1 | # | 10 | Small | aldex2 | 44 |
| # | time | 2 | 2 | 0 | 0.5 | 1 | # | 10 | Small | aldex2 | 44 |
| # | interaction | 0 | 0 | 2 | 0 | 0 | # | 10 | Small | aldex2 | 44 |
| # | disease | 4 | 11 | 0 | 0.266666667 | 1 | # | 10 | Baseline | paralle | 44 |
| # | sex | 2 | 5 | 0 | 0.285714286 | 1 | # | 10 | Baseline | paralle | 44 |
| # | time | 2 | 7 | 0 | 0.222222222 | 1 | # | 10 | Baseline | paralle | 44 |
| # | interaction | 2 | 7 | 0 | 0.222222222 | 1 | # | 10 | Baseline | paralle | 44 |
| # | disease | 4 | 2 | 0 | 0.666666667 | 1 | # | 10 | Baseline | ancom | 44 |
| # | sex | 2 | 0 | 0 | 1 | 1 | # | 10 | Baseline | ancom | 44 |
| # | time | 2 | 2 | 0 | 0.5 | 1 | # | 10 | Baseline | ancom | 44 |
| # | interaction | 0 | 0 | 2 | 0 | 0 | # | 10 | Baseline | ancom | 44 |
| # | disease | 4 | 22 | 0 | 0.153846154 | 1 | # | 10 | Baseline | aldex2 | 44 |
| # | sex | 2 | 8 | 0 | 0.2 | 1 | # | 10 | Baseline | aldex2 | 44 |
| # | time | 2 | 10 | 0 | 0.166666667 | 1 | # | 10 | Baseline | aldex2 | 44 |
| # | interaction | 0 | 0 | 2 | 0 | 0 | # | 10 | Baseline | aldex2 | 44 |
| # | disease | 4 | 0 | 0 | 1 | 1 | # | 10 | Small | paralle | 44 |
| # | sex | 2 | 0 | 0 | 1 | 1 | # | 10 | Small | paralle | 44 |
| # | time | 2 | 0 | 0 | 1 | 1 | # | 10 | Small | paralle | 44 |
| # | interaction | 2 | 0 | 0 | 1 | 1 | # | 10 | Small | paralle | 44 |
| # | disease | 4 | 3 | 0 | 0.571428571 | 1 | # | 10 | Small | ancom | 44 |
| # | sex | 2 | 0 | 0 | 1 | 1 | # | 10 | Small | ancom | 44 |
| # | time | 2 | 2 | 0 | 0.5 | 1 | # | 10 | Small | ancom | 44 |
| # | interaction | 0 | 0 | 2 | 0 | 0 | # | 10 | Small | ancom | 44 |
| # | disease | 4 | 3 | 0 | 0.571428571 | 1 | # | 10 | Small | aldex2 | 44 |
| # | sex | 2 | 0 | 0 | 1 | 1 | # | 10 | Small | aldex2 | 44 |
| # | time | 2 | 2 | 0 | 0.5 | 1 | # | 10 | Small | aldex2 | 44 |
| # | interaction | 0 | 0 | 2 | 0 | 0 | # | 10 | Small | aldex2 | 44 |
| # | disease | 4 | 8 | 0 | 0.333333333 | 1 | # | 10 | Baseline | paralle | 44 |
| # | sex | 2 | 9 | 0 | 0.181818182 | 1 | # | 10 | Baseline | paralle | 44 |
| # | time | 2 | 4 | 0 | 0.333333333 | 1 | # | 10 | Baseline | paralle | 44 |
| # | interaction | 2 | 10 | 0 | 0.166666667 | 1 | # | 10 | Baseline | paralle | 44 |
| # | disease | 4 | 2 | 0 | 0.666666667 | 1 | # | 10 | Baseline | ancom | 44 |
| # | sex | 2 | 0 | 0 | 1 | 1 | # | 10 | Baseline | ancom | 44 |
| # | time | 2 | 2 | 0 | 0.5 | 1 | # | 10 | Baseline | ancom | 44 |
| # | interaction | 0 | 0 | 2 | 0 | 0 | # | 10 | Baseline | ancom | 44 |
| # | disease | 4 | 25 | 0 | 0.137931034 | 1 | # | 10 | Baseline | aldex2 | 44 |
| # | sex | 2 | 11 | 0 | 0.153846154 | 1 | # | 10 | Baseline | aldex2 | 44 |
| # | time | 2 | 10 | 0 | 0.166666667 | 1 | # | 10 | Baseline | aldex2 | 44 |
| # | interaction | 0 | 0 | 2 | 0 | 0 | # | 10 | Baseline | aldex2 | 44 |
| # | disease | 4 | 0 | 0 | 1 | 1 | # | 10 | Small | paralle | 44 |
| # | sex | 2 | 0 | 0 | 1 | 1 | # | 10 | Small | paralle | 44 |
| # | time | 2 | 1 | 0 | 0.666666667 | 1 | # | 10 | Small | paralle | 44 |
| # | interaction | 2 | 1 | 0 | 0.666666667 | 1 | # | 10 | Small | paralle | 44 |
| # | disease | 4 | 2 | 0 | 0.666666667 | 1 | # | 10 | Small | ancom | 44 |
| # | sex | 2 | 0 | 0 | 1 | 1 | # | 10 | Small | ancom | 44 |
| # | time | 2 | 2 | 0 | 0.5 | 1 | # | 10 | Small | ancom | 44 |
| # | interaction | 0 | 0 | 2 | 0 | 0 | # | 10 | Small | ancom | 44 |
| # | disease | 4 | 2 | 0 | 0.666666667 | 1 | # | 10 | Small | aldex2 | 44 |
| # | sex | 2 | 1 | 0 | 0.666666667 | 1 | # | 10 | Small | aldex2 | 44 |
| # | time | 2 | 2 | 0 | 0.5 | 1 | # | 10 | Small | aldex2 | 44 |
| # | interaction | 0 | 0 | 2 | 0 | 0 | # | 10 | Small | aldex2 | 44 |
| # | disease | 4 | 6 | 0 | 0.4 | 1 | # | 10 | Baseline | paralle | 44 |
| # | sex | 2 | 3 | 0 | 0.4 | 1 | # | 10 | Baseline | paralle | 44 |
| # | time | 2 | 3 | 0 | 0.4 | 1 | # | 10 | Baseline | paralle | 44 |
| # | interaction | 2 | 3 | 0 | 0.4 | 1 | # | 10 | Baseline | paralle | 44 |
| # | disease | 4 | 2 | 0 | 0.666666667 | 1 | # | 10 | Baseline | ancom | 44 |
| # | sex | 2 | 0 | 0 | 1 | 1 | # | 10 | Baseline | ancom | 44 |
| # | time | 2 | 2 | 0 | 0.5 | 1 | # | 10 | Baseline | ancom | 44 |
| # | interaction | 0 | 0 | 2 | 0 | 0 | # | 10 | Baseline | ancom | 44 |
| # | disease | 4 | 33 | 0 | 0.108108108 | 1 | # | 10 | Baseline | aldex2 | 44 |
| # | sex | 2 | 8 | 0 | 0.2 | 1 | # | 10 | Baseline | aldex2 | 44 |
| # | time | 2 | 14 | 0 | 0.125 | 1 | # | 10 | Baseline | aldex2 | 44 |
| # | interaction | 0 | 0 | 2 | 0 | 0 | # | 10 | Baseline | aldex2 | 44 |
| # | disease | 4 | 0 | 0 | 1 | 1 | # | 10 | Small | paralle | 44 |
| # | sex | 2 | 0 | 0 | 1 | 1 | # | 10 | Small | paralle | 44 |
| # | time | 2 | 0 | 0 | 1 | 1 | # | 10 | Small | paralle | 44 |
| # | interaction | 2 | 0 | 0 | 1 | 1 | # | 10 | Small | paralle | 44 |
| # | disease | 4 | 2 | 0 | 0.666666667 | 1 | # | 10 | Small | ancom | 44 |
| # | sex | 2 | 0 | 0 | 1 | 1 | # | 10 | Small | ancom | 44 |
| # | time | 2 | 2 | 0 | 0.5 | 1 | # | 10 | Small | ancom | 44 |
| # | interaction | 0 | 0 | 2 | 0 | 0 | # | 10 | Small | ancom | 44 |
| # | disease | 4 | 3 | 0 | 0.571428571 | 1 | # | 10 | Small | aldex2 | 44 |
| # | sex | 2 | 0 | 0 | 1 | 1 | # | 10 | Small | aldex2 | 44 |
| # | time | 2 | 2 | 0 | 0.5 | 1 | # | 10 | Small | aldex2 | 44 |
| # | interaction | 0 | 0 | 2 | 0 | 0 | # | 10 | Small | aldex2 | 44 |
| # | disease | 7 | 14 | 0 | 0.333333333 | 1 | # | 20 | Baseline | paralle | 44 |
| # | sex | 4 | 15 | 0 | 0.210526316 | 1 | # | 20 | Baseline | paralle | 44 |
| # | time | 5 | 13 | 0 | 0.277777778 | 1 | # | 20 | Baseline | paralle | 44 |
| # | interaction | 4 | 5 | 0 | 0.444444444 | 1 | # | 20 | Baseline | paralle | 44 |
| # | disease | 7 | 1 | 0 | 0.875 | 1 | # | 20 | Baseline | ancom | 44 |
| # | sex | 4 | 0 | 0 | 1 | 1 | # | 20 | Baseline | ancom | 44 |
| # | time | 5 | 1 | 0 | 0.833333333 | 1 | # | 20 | Baseline | ancom | 44 |
| # | interaction | 0 | 0 | 4 | 0 | 0 | # | 20 | Baseline | ancom | 44 |
| # | disease | 7 | 22 | 0 | 0.24137931 | 1 | # | 20 | Baseline | aldex2 | 44 |
| # | sex | 4 | 9 | 0 | 0.307692308 | 1 | # | 20 | Baseline | aldex2 | 44 |
| # | time | 5 | 13 | 0 | 0.277777778 | 1 | # | 20 | Baseline | aldex2 | 44 |
| # | interaction | 0 | 0 | 4 | 0 | 0 | # | 20 | Baseline | aldex2 | 44 |
| # | disease | 7 | 0 | 0 | 1 | 1 | # | 20 | Small | paralle | 44 |
| # | sex | 4 | 0 | 0 | 1 | 1 | # | 20 | Small | paralle | 44 |
| # | time | 5 | 0 | 0 | 1 | 1 | # | 20 | Small | paralle | 44 |
| # | interaction | 1 | 0 | 3 | 1 | 0.25 | # | 20 | Small | paralle | 44 |
| # | disease | 7 | 1 | 0 | 0.875 | 1 | # | 20 | Small | ancom | 44 |
| # | sex | 4 | 0 | 0 | 1 | 1 | # | 20 | Small | ancom | 44 |
| # | time | 5 | 1 | 0 | 0.833333333 | 1 | # | 20 | Small | ancom | 44 |
| # | interaction | 0 | 0 | 4 | 0 | 0 | # | 20 | Small | ancom | 44 |
| # | disease | 7 | 2 | 0 | 0.777777778 | 1 | # | 20 | Small | aldex2 | 44 |
| # | sex | 4 | 0 | 0 | 1 | 1 | # | 20 | Small | aldex2 | 44 |
| # | time | 5 | 1 | 0 | 0.833333333 | 1 | # | 20 | Small | aldex2 | 44 |
| # | interaction | 0 | 0 | 4 | 0 | 0 | # | 20 | Small | aldex2 | 44 |
| # | disease | 7 | 8 | 0 | 0.466666667 | 1 | # | 20 | Baseline | paralle | 44 |
| # | sex | 4 | 9 | 0 | 0.307692308 | 1 | # | 20 | Baseline | paralle | 44 |
| # | time | 5 | 6 | 0 | 0.454545455 | 1 | # | 20 | Baseline | paralle | 44 |
| # | interaction | 4 | 6 | 0 | 0.4 | 1 | # | 20 | Baseline | paralle | 44 |
| # | disease | 7 | 2 | 0 | 0.777777778 | 1 | # | 20 | Baseline | ancom | 44 |
| # | sex | 4 | 0 | 0 | 1 | 1 | # | 20 | Baseline | ancom | 44 |
| # | time | 5 | 2 | 0 | 0.714285714 | 1 | # | 20 | Baseline | ancom | 44 |
| # | interaction | 0 | 0 | 4 | 0 | 0 | # | 20 | Baseline | ancom | 44 |
| # | disease | 7 | 36 | 0 | 0.162790698 | 1 | # | 20 | Baseline | aldex2 | 44 |
| # | sex | 4 | 12 | 0 | 0.25 | 1 | # | 20 | Baseline | aldex2 | 44 |
| # | time | 5 | 25 | 0 | 0.166666667 | 1 | # | 20 | Baseline | aldex2 | 44 |
| # | interaction | 0 | 0 | 4 | 0 | 0 | # | 20 | Baseline | aldex2 | 44 |
| # | disease | 7 | 0 | 0 | 1 | 1 | # | 20 | Small | paralle | 44 |
| # | sex | 4 | 0 | 0 | 1 | 1 | # | 20 | Small | paralle | 44 |
| # | time | 5 | 0 | 0 | 1 | 1 | # | 20 | Small | paralle | 44 |
| # | interaction | 4 | 0 | 0 | 1 | 1 | # | 20 | Small | paralle | 44 |
| # | disease | 7 | 2 | 0 | 0.777777778 | 1 | # | 20 | Small | ancom | 44 |
| # | sex | 4 | 0 | 0 | 1 | 1 | # | 20 | Small | ancom | 44 |
| # | time | 5 | 1 | 0 | 0.833333333 | 1 | # | 20 | Small | ancom | 44 |
| # | interaction | 0 | 0 | 4 | 0 | 0 | # | 20 | Small | ancom | 44 |
| # | disease | 7 | 3 | 0 | 0.7 | 1 | # | 20 | Small | aldex2 | 44 |
| # | sex | 4 | 0 | 0 | 1 | 1 | # | 20 | Small | aldex2 | 44 |
| # | time | 5 | 3 | 0 | 0.625 | 1 | # | 20 | Small | aldex2 | 44 |
| # | interaction | 0 | 0 | 4 | 0 | 0 | # | 20 | Small | aldex2 | 44 |
| # | disease | 7 | 14 | 0 | 0.333333333 | 1 | # | 20 | Baseline | paralle | 44 |
| # | sex | 4 | 12 | 0 | 0.25 | 1 | # | 20 | Baseline | paralle | 44 |
| # | time | 5 | 11 | 0 | 0.3125 | 1 | # | 20 | Baseline | paralle | 44 |
| # | interaction | 4 | 8 | 0 | 0.333333333 | 1 | # | 20 | Baseline | paralle | 44 |
| # | disease | 7 | 4 | 0 | 0.636363636 | 1 | # | 20 | Baseline | ancom | 44 |
| # | sex | 4 | 0 | 0 | 1 | 1 | # | 20 | Baseline | ancom | 44 |
| # | time | 5 | 2 | 0 | 0.714285714 | 1 | # | 20 | Baseline | ancom | 44 |
| # | interaction | 0 | 0 | 4 | 0 | 0 | # | 20 | Baseline | ancom | 44 |
| # | disease | 7 | 47 | 0 | 0.12962963 | 1 | # | 20 | Baseline | aldex2 | 44 |
| # | sex | 4 | 14 | 0 | 0.222222222 | 1 | # | 20 | Baseline | aldex2 | 44 |
| # | time | 5 | 38 | 0 | 0.11627907 | 1 | # | 20 | Baseline | aldex2 | 44 |
| # | interaction | 0 | 0 | 4 | 0 | 0 | # | 20 | Baseline | aldex2 | 44 |
| # | disease | 7 | 0 | 0 | 1 | 1 | # | 20 | Small | paralle | 44 |
| # | sex | 4 | 0 | 0 | 1 | 1 | # | 20 | Small | paralle | 44 |
| # | time | 5 | 0 | 0 | 1 | 1 | # | 20 | Small | paralle | 44 |
| # | interaction | 4 | 1 | 0 | 0.8 | 1 | # | 20 | Small | paralle | 44 |
| # | disease | 7 | 4 | 0 | 0.636363636 | 1 | # | 20 | Small | ancom | 44 |
| # | sex | 4 | 0 | 0 | 1 | 1 | # | 20 | Small | ancom | 44 |
| # | time | 5 | 2 | 0 | 0.714285714 | 1 | # | 20 | Small | ancom | 44 |
| # | interaction | 0 | 0 | 4 | 0 | 0 | # | 20 | Small | ancom | 44 |
| # | disease | 7 | 9 | 0 | 0.4375 | 1 | # | 20 | Small | aldex2 | 44 |
| # | sex | 4 | 1 | 0 | 0.8 | 1 | # | 20 | Small | aldex2 | 44 |

| | | | | | | | | | | | |
|---|---|---|---|---|---|---|---|---|---|---|---|
| # | time | 5 | 3 | 0 | 0.625 | 1 | # | 20 | Small | aldex2 | 44 |
| # | interaction | 0 | 0 | 4 | 0 | 0 | # | 20 | Small | aldex2 | 44 |
| # | disease | 7 | 13 | 0 | 0.35 | 1 | # | 20 | Baseline | paralle | 44 |
| # | sex | 4 | 9 | 0 | 0.307692308 | 1 | # | 20 | Baseline | paralle | 44 |
| # | time | 5 | 9 | 0 | 0.357142857 | 1 | # | 20 | Baseline | paralle | 44 |
| # | interaction | 4 | 4 | 0 | 0.5 | 1 | # | 20 | Baseline | paralle | 44 |
| # | disease | 7 | 4 | 0 | 0.636363636 | 1 | # | 20 | Baseline | ancom | 44 |
| # | sex | 4 | 0 | 0 | 1 | 1 | # | 20 | Baseline | ancom | 44 |
| # | time | 5 | 3 | 0 | 0.625 | 1 | # | 20 | Baseline | ancom | 44 |
| # | interaction | 0 | 0 | 4 | 0 | 0 | # | 20 | Baseline | ancom | 44 |
| # | disease | 7 | 51 | 0 | 0.102941176 | 1 | # | 20 | Baseline | aldex2 | 44 |
| # | sex | 4 | 18 | 0 | 0.181818182 | 1 | # | 20 | Baseline | aldex2 | 44 |
| # | time | 5 | 51 | 0 | 0.089285714 | 1 | # | 20 | Baseline | aldex2 | 44 |
| # | interaction | 0 | 0 | 4 | 0 | 0 | # | 20 | Baseline | aldex2 | 44 |
| # | disease | 7 | 0 | 0 | 1 | 1 | # | 20 | Small | paralle | 44 |
| # | sex | 4 | 1 | 0 | 0.8 | 1 | # | 20 | Small | paralle | 44 |
| # | time | 5 | 0 | 0 | 1 | 1 | # | 20 | Small | paralle | 44 |
| # | interaction | 4 | 0 | 0 | 1 | 1 | # | 20 | Small | paralle | 44 |
| # | disease | 7 | 4 | 0 | 0.636363636 | 1 | # | 20 | Small | ancom | 44 |
| # | sex | 4 | 0 | 0 | 1 | 1 | # | 20 | Small | ancom | 44 |
| # | time | 5 | 3 | 0 | 0.625 | 1 | # | 20 | Small | ancom | 44 |
| # | interaction | 0 | 0 | 4 | 0 | 0 | # | 20 | Small | ancom | 44 |
| # | disease | 7 | 8 | 0 | 0.466666667 | 1 | # | 20 | Small | aldex2 | 44 |
| # | sex | 4 | 1 | 0 | 0.8 | 1 | # | 20 | Small | aldex2 | 44 |
| # | time | 5 | 4 | 0 | 0.555555556 | 1 | # | 20 | Small | aldex2 | 44 |
| # | interaction | 0 | 0 | 4 | 0 | 0 | # | 20 | Small | aldex2 | 44 |
| # | disease | 4 | 6 | 0 | 0.4 | 1 | # | 10 | Baseline | paralle | 45 |
| # | sex | 2 | 5 | 0 | 0.285714286 | 1 | # | 10 | Baseline | paralle | 45 |
| # | time | 2 | 3 | 0 | 0.4 | 1 | # | 10 | Baseline | paralle | 45 |
| # | interaction | 2 | 7 | 0 | 0.222222222 | 1 | # | 10 | Baseline | paralle | 45 |
| # | disease | 4 | 1 | 0 | 0.8 | 1 | # | 10 | Baseline | ancom | 45 |
| # | sex | 2 | 0 | 0 | 1 | 1 | # | 10 | Baseline | ancom | 45 |
| # | time | 2 | 0 | 0 | 1 | 1 | # | 10 | Baseline | ancom | 45 |
| # | interaction | 0 | 0 | 2 | 0 | 0 | # | 10 | Baseline | ancom | 45 |
| # | disease | 4 | 11 | 0 | 0.266666667 | 1 | # | 10 | Baseline | aldex2 | 45 |
| # | sex | 2 | 8 | 0 | 0.2 | 1 | # | 10 | Baseline | aldex2 | 45 |
| # | time | 2 | 5 | 0 | 0.285714286 | 1 | # | 10 | Baseline | aldex2 | 45 |
| # | interaction | 0 | 0 | 2 | 0 | 0 | # | 10 | Baseline | aldex2 | 45 |
| # | disease | 4 | 0 | 0 | 1 | 1 | # | 10 | Small | paralle | 45 |
| # | sex | 2 | 0 | 0 | 1 | 1 | # | 10 | Small | paralle | 45 |
| # | time | 2 | 0 | 0 | 1 | 1 | # | 10 | Small | paralle | 45 |
| # | interaction | 2 | 0 | 0 | 1 | 1 | # | 10 | Small | paralle | 45 |
| # | disease | 4 | 0 | 0 | 1 | 1 | # | 10 | Small | ancom | 45 |
| # | sex | 2 | 0 | 0 | 1 | 1 | # | 10 | Small | ancom | 45 |
| # | time | 2 | 0 | 0 | 1 | 1 | # | 10 | Small | ancom | 45 |
| # | interaction | 0 | 0 | 2 | 0 | 0 | # | 10 | Small | ancom | 45 |
| # | disease | 4 | 0 | 0 | 1 | 1 | # | 10 | Small | aldex2 | 45 |
| # | sex | 2 | 0 | 0 | 1 | 1 | # | 10 | Small | aldex2 | 45 |
| # | time | 2 | 0 | 0 | 1 | 1 | # | 10 | Small | aldex2 | 45 |
| # | interaction | 0 | 0 | 2 | 0 | 0 | # | 10 | Small | aldex2 | 45 |
| # | disease | 4 | 7 | 0 | 0.363636364 | 1 | # | 10 | Baseline | paralle | 45 |
| # | sex | 2 | 6 | 0 | 0.25 | 1 | # | 10 | Baseline | paralle | 45 |
| # | time | 2 | 4 | 0 | 0.333333333 | 1 | # | 10 | Baseline | paralle | 45 |
| # | interaction | 2 | 3 | 0 | 0.4 | 1 | # | 10 | Baseline | paralle | 45 |
| # | disease | 4 | 2 | 0 | 0.666666667 | 1 | # | 10 | Baseline | ancom | 45 |
| # | sex | 2 | 0 | 0 | 1 | 1 | # | 10 | Baseline | ancom | 45 |
| # | time | 2 | 1 | 0 | 0.666666667 | 1 | # | 10 | Baseline | ancom | 45 |
| # | interaction | 0 | 0 | 2 | 0 | 0 | # | 10 | Baseline | ancom | 45 |
| # | disease | 4 | 13 | 0 | 0.235294118 | 1 | # | 10 | Baseline | aldex2 | 45 |
| # | sex | 2 | 12 | 0 | 0.142857143 | 1 | # | 10 | Baseline | aldex2 | 45 |
| # | time | 2 | 7 | 0 | 0.222222222 | 1 | # | 10 | Baseline | aldex2 | 45 |
| # | interaction | 0 | 0 | 2 | 0 | 0 | # | 10 | Baseline | aldex2 | 45 |
| # | disease | 4 | 0 | 0 | 1 | 1 | # | 10 | Small | paralle | 45 |
| # | sex | 2 | 0 | 0 | 1 | 1 | # | 10 | Small | paralle | 45 |
| # | time | 2 | 1 | 0 | 0.666666667 | 1 | # | 10 | Small | paralle | 45 |
| # | interaction | 2 | 0 | 0 | 1 | 1 | # | 10 | Small | paralle | 45 |
| # | disease | 4 | 2 | 0 | 0.666666667 | 1 | # | 10 | Small | ancom | 45 |
| # | sex | 2 | 0 | 0 | 1 | 1 | # | 10 | Small | ancom | 45 |
| # | time | 2 | 1 | 0 | 0.666666667 | 1 | # | 10 | Small | ancom | 45 |
| # | interaction | 0 | 0 | 2 | 0 | 0 | # | 10 | Small | ancom | 45 |
| # | disease | 4 | 1 | 0 | 0.8 | 1 | # | 10 | Small | aldex2 | 45 |
| # | sex | 2 | 1 | 0 | 0.666666667 | 1 | # | 10 | Small | aldex2 | 45 |
| # | time | 2 | 1 | 0 | 0.666666667 | 1 | # | 10 | Small | aldex2 | 45 |
| # | interaction | 0 | 0 | 2 | 0 | 0 | # | 10 | Small | aldex2 | 45 |
| # | disease | 4 | 4 | 0 | 0.5 | 1 | # | 10 | Baseline | paralle | 45 |
| # | sex | 2 | 3 | 0 | 0.4 | 1 | # | 10 | Baseline | paralle | 45 |
| # | time | 2 | 2 | 0 | 0.5 | 1 | # | 10 | Baseline | paralle | 45 |
| # | interaction | 2 | 3 | 0 | 0.4 | 1 | # | 10 | Baseline | paralle | 45 |
| # | disease | 4 | 2 | 0 | 0.666666667 | 1 | # | 10 | Baseline | ancom | 45 |
| # | sex | 2 | 0 | 0 | 1 | 1 | # | 10 | Baseline | ancom | 45 |
| # | time | 2 | 2 | 0 | 0.5 | 1 | # | 10 | Baseline | ancom | 45 |
| # | interaction | 0 | 0 | 2 | 0 | 0 | # | 10 | Baseline | ancom | 45 |
| # | disease | 4 | 22 | 0 | 0.153846154 | 1 | # | 10 | Baseline | aldex2 | 45 |
| # | sex | 2 | 6 | 0 | 0.25 | 1 | # | 10 | Baseline | aldex2 | 45 |
| # | time | 2 | 11 | 0 | 0.153846154 | 1 | # | 10 | Baseline | aldex2 | 45 |
| # | interaction | 0 | 0 | 2 | 0 | 0 | # | 10 | Baseline | aldex2 | 45 |
| # | disease | 4 | 0 | 0 | 1 | 1 | # | 10 | Small | paralle | 45 |
| # | sex | 2 | 0 | 0 | 1 | 1 | # | 10 | Small | paralle | 45 |
| # | time | 2 | 0 | 0 | 1 | 1 | # | 10 | Small | paralle | 45 |
| # | interaction | 2 | 0 | 0 | 1 | 1 | # | 10 | Small | paralle | 45 |
| # | disease | 4 | 2 | 0 | 0.666666667 | 1 | # | 10 | Small | ancom | 45 |
| # | sex | 2 | 0 | 0 | 1 | 1 | # | 10 | Small | ancom | 45 |
| # | time | 2 | 1 | 0 | 0.666666667 | 1 | # | 10 | Small | ancom | 45 |
| # | interaction | 0 | 0 | 2 | 0 | 0 | # | 10 | Small | ancom | 45 |
| # | disease | 4 | 2 | 0 | 0.666666667 | 1 | # | 10 | Small | aldex2 | 45 |
| # | sex | 2 | 0 | 0 | 1 | 1 | # | 10 | Small | aldex2 | 45 |
| # | time | 2 | 1 | 0 | 0.666666667 | 1 | # | 10 | Small | aldex2 | 45 |
| # | interaction | 0 | 0 | 2 | 0 | 0 | # | 10 | Small | aldex2 | 45 |
| # | disease | 4 | 3 | 0 | 0.571428571 | 1 | # | 10 | Baseline | paralle | 45 |
| # | sex | 2 | 4 | 0 | 0.333333333 | 1 | # | 10 | Baseline | paralle | 45 |
| # | time | 2 | 3 | 0 | 0.4 | 1 | # | 10 | Baseline | paralle | 45 |
| # | interaction | 2 | 3 | 0 | 0.4 | 1 | # | 10 | Baseline | paralle | 45 |
| # | disease | 4 | 2 | 0 | 0.666666667 | 1 | # | 10 | Baseline | ancom | 45 |
| # | sex | 2 | 0 | 0 | 1 | 1 | # | 10 | Baseline | ancom | 45 |
| # | time | 2 | 2 | 0 | 0.5 | 1 | # | 10 | Baseline | ancom | 45 |
| # | interaction | 0 | 0 | 2 | 0 | 0 | # | 10 | Baseline | ancom | 45 |
| # | disease | 4 | 26 | 0 | 0.133333333 | 1 | # | 10 | Baseline | aldex2 | 45 |
| # | sex | 2 | 13 | 0 | 0.133333333 | 1 | # | 10 | Baseline | aldex2 | 45 |
| # | time | 2 | 12 | 0 | 0.142857143 | 1 | # | 10 | Baseline | aldex2 | 45 |
| # | interaction | 0 | 0 | 2 | 0 | 0 | # | 10 | Baseline | aldex2 | 45 |
| # | disease | 4 | 0 | 0 | 1 | 1 | # | 10 | Small | paralle | 45 |
| # | sex | 2 | 0 | 0 | 1 | 1 | # | 10 | Small | paralle | 45 |
| # | time | 2 | 0 | 0 | 1 | 1 | # | 10 | Small | paralle | 45 |
| # | interaction | 2 | 0 | 0 | 1 | 1 | # | 10 | Small | paralle | 45 |
| # | disease | 4 | 2 | 0 | 0.666666667 | 1 | # | 10 | Small | ancom | 45 |
| # | sex | 2 | 0 | 0 | 1 | 1 | # | 10 | Small | ancom | 45 |
| # | time | 2 | 2 | 0 | 0.5 | 1 | # | 10 | Small | ancom | 45 |
| # | interaction | 0 | 0 | 2 | 0 | 0 | # | 10 | Small | ancom | 45 |
| # | disease | 4 | 3 | 0 | 0.571428571 | 1 | # | 10 | Small | aldex2 | 45 |
| # | sex | 2 | 0 | 0 | 1 | 1 | # | 10 | Small | aldex2 | 45 |
| # | time | 2 | 1 | 0 | 0.666666667 | 1 | # | 10 | Small | aldex2 | 45 |
| # | interaction | 0 | 0 | 2 | 0 | 0 | # | 10 | Small | aldex2 | 45 |
| # | disease | 7 | 7 | 0 | 0.5 | 1 | # | 20 | Baseline | paralle | 45 |
| # | sex | 4 | 6 | 0 | 0.4 | 1 | # | 20 | Baseline | paralle | 45 |
| # | time | 5 | 6 | 0 | 0.454545455 | 1 | # | 20 | Baseline | paralle | 45 |
| # | interaction | 4 | 6 | 0 | 0.4 | 1 | # | 20 | Baseline | paralle | 45 |
| # | disease | 7 | 0 | 0 | 1 | 1 | # | 20 | Baseline | ancom | 45 |
| # | sex | 4 | 0 | 0 | 1 | 1 | # | 20 | Baseline | ancom | 45 |
| # | time | 5 | 0 | 0 | 1 | 1 | # | 20 | Baseline | ancom | 45 |
| # | interaction | 0 | 0 | 4 | 0 | 0 | # | 20 | Baseline | ancom | 45 |
| # | disease | 7 | 19 | 0 | 0.269230769 | 1 | # | 20 | Baseline | aldex2 | 45 |
| # | sex | 4 | 3 | 0 | 0.571428571 | 1 | # | 20 | Baseline | aldex2 | 45 |
| # | time | 5 | 13 | 0 | 0.277777778 | 1 | # | 20 | Baseline | aldex2 | 45 |
| # | interaction | 0 | 0 | 4 | 0 | 0 | # | 20 | Baseline | aldex2 | 45 |
| # | disease | 7 | 0 | 0 | 1 | 1 | # | 20 | Small | paralle | 45 |
| # | sex | 4 | 0 | 0 | 1 | 1 | # | 20 | Small | paralle | 45 |
| # | time | 5 | 0 | 0 | 1 | 1 | # | 20 | Small | paralle | 45 |
| # | interaction | 2 | 0 | 2 | 1 | 0.5 | # | 20 | Small | paralle | 45 |
| # | disease | 7 | 1 | 0 | 0.875 | 1 | # | 20 | Small | ancom | 45 |
| # | sex | 4 | 0 | 0 | 1 | 1 | # | 20 | Small | ancom | 45 |
| # | time | 5 | 1 | 0 | 0.833333333 | 1 | # | 20 | Small | ancom | 45 |
| # | interaction | 0 | 0 | 4 | 0 | 0 | # | 20 | Small | ancom | 45 |
| # | disease | 7 | 1 | 0 | 0.875 | 1 | # | 20 | Small | aldex2 | 45 |
| # | sex | 4 | 0 | 0 | 1 | 1 | # | 20 | Small | aldex2 | 45 |
| # | time | 5 | 1 | 0 | 0.833333333 | 1 | # | 20 | Small | aldex2 | 45 |
| # | interaction | 0 | 0 | 4 | 0 | 0 | # | 20 | Small | aldex2 | 45 |
| # | disease | 7 | 4 | 0 | 0.636363636 | 1 | # | 20 | Baseline | paralle | 45 |
| # | sex | 4 | 0 | 0 | 1 | 1 | # | 20 | Baseline | paralle | 45 |
| # | time | 5 | 5 | 0 | 0.5 | 1 | # | 20 | Baseline | paralle | 45 |
| # | interaction | 4 | 4 | 0 | 0.5 | 1 | # | 20 | Baseline | paralle | 45 |
| # | disease | 7 | 4 | 0 | 0.636363636 | 1 | # | 20 | Baseline | ancom | 45 |
| # | sex | 4 | 0 | 0 | 1 | 1 | # | 20 | Baseline | ancom | 45 |
| # | time | 5 | 3 | 0 | 0.625 | 1 | # | 20 | Baseline | ancom | 45 |
| # | interaction | 0 | 0 | 4 | 0 | 0 | # | 20 | Baseline | ancom | 45 |
| # | disease | 7 | 35 | 0 | 0.166666667 | 1 | # | 20 | Baseline | aldex2 | 45 |
| # | sex | 4 | 7 | 0 | 0.363636364 | 1 | # | 20 | Baseline | aldex2 | 45 |
| # | time | 5 | 23 | 0 | 0.178571429 | 1 | # | 20 | Baseline | aldex2 | 45 |
| # | interaction | 0 | 0 | 4 | 0 | 0 | # | 20 | Baseline | aldex2 | 45 |
| # | disease | 7 | 0 | 0 | 1 | 1 | # | 20 | Small | paralle | 45 |
| # | sex | 4 | 0 | 0 | 1 | 1 | # | 20 | Small | paralle | 45 |
| # | time | 5 | 0 | 0 | 1 | 1 | # | 20 | Small | paralle | 45 |
| # | interaction | 4 | 0 | 0 | 1 | 1 | # | 20 | Small | paralle | 45 |
| # | disease | 7 | 4 | 0 | 0.636363636 | 1 | # | 20 | Small | ancom | 45 |
| # | sex | 4 | 0 | 0 | 1 | 1 | # | 20 | Small | ancom | 45 |
| # | time | 5 | 3 | 0 | 0.625 | 1 | # | 20 | Small | ancom | 45 |
| # | interaction | 0 | 0 | 4 | 0 | 0 | # | 20 | Small | ancom | 45 |
| # | disease | 7 | 6 | 0 | 0.538461538 | 1 | # | 20 | Small | aldex2 | 45 |
| # | sex | 4 | 0 | 0 | 1 | 1 | # | 20 | Small | aldex2 | 45 |
| # | time | 5 | 4 | 0 | 0.555555556 | 1 | # | 20 | Small | aldex2 | 45 |
| # | interaction | 0 | 0 | 4 | 0 | 0 | # | 20 | Small | aldex2 | 45 |
| # | disease | 7 | 11 | 0 | 0.388888889 | 1 | # | 20 | Baseline | paralle | 45 |
| # | sex | 4 | 1 | 0 | 0.8 | 1 | # | 20 | Baseline | paralle | 45 |
| # | time | 5 | 7 | 0 | 0.416666667 | 1 | # | 20 | Baseline | paralle | 45 |
| # | interaction | 4 | 3 | 0 | 0.571428571 | 1 | # | 20 | Baseline | paralle | 45 |
| # | disease | 7 | 4 | 0 | 0.636363636 | 1 | # | 20 | Baseline | ancom | 45 |
| # | sex | 4 | 0 | 0 | 1 | 1 | # | 20 | Baseline | ancom | 45 |
| # | time | 5 | 4 | 0 | 0.555555556 | 1 | # | 20 | Baseline | ancom | 45 |
| # | interaction | 0 | 0 | 4 | 0 | 0 | # | 20 | Baseline | ancom | 45 |
| # | disease | 7 | 47 | 0 | 0.12962963 | 1 | # | 20 | Baseline | aldex2 | 45 |
| # | sex | 4 | 15 | 0 | 0.210526316 | 1 | # | 20 | Baseline | aldex2 | 45 |
| # | time | 5 | 26 | 0 | 0.161290323 | 1 | # | 20 | Baseline | aldex2 | 45 |
| # | interaction | 0 | 0 | 4 | 0 | 0 | # | 20 | Baseline | aldex2 | 45 |
| # | disease | 7 | 0 | 0 | 1 | 1 | # | 20 | Small | paralle | 45 |
| # | sex | 4 | 0 | 0 | 1 | 1 | # | 20 | Small | paralle | 45 |
| # | time | 5 | 0 | 0 | 1 | 1 | # | 20 | Small | paralle | 45 |
| # | interaction | 4 | 0 | 0 | 1 | 1 | # | 20 | Small | paralle | 45 |
| # | disease | 7 | 4 | 0 | 0.636363636 | 1 | # | 20 | Small | ancom | 45 |
| # | sex | 4 | 0 | 0 | 1 | 1 | # | 20 | Small | ancom | 45 |
| # | time | 5 | 4 | 0 | 0.555555556 | 1 | # | 20 | Small | ancom | 45 |
| # | interaction | 0 | 0 | 4 | 0 | 0 | # | 20 | Small | ancom | 45 |
| # | disease | 7 | 7 | 0 | 0.5 | 1 | # | 20 | Small | aldex2 | 45 |
| # | sex | 4 | 0 | 0 | 1 | 1 | # | 20 | Small | aldex2 | 45 |
| # | time | 5 | 2 | 0 | 0.714285714 | 1 | # | 20 | Small | aldex2 | 45 |
| # | interaction | 0 | 0 | 4 | 0 | 0 | # | 20 | Small | aldex2 | 45 |
| # | disease | 7 | 7 | 0 | 0.5 | 1 | # | 20 | Baseline | paralle | 45 |
| # | sex | 4 | 2 | 0 | 0.666666667 | 1 | # | 20 | Baseline | paralle | 45 |
| # | time | 5 | 8 | 0 | 0.384615385 | 1 | # | 20 | Baseline | paralle | 45 |
| # | interaction | 4 | 0 | 0 | 1 | 1 | # | 20 | Baseline | paralle | 45 |
| # | disease | 7 | 4 | 0 | 0.636363636 | 1 | # | 20 | Baseline | ancom | 45 |
| # | sex | 4 | 0 | 0 | 1 | 1 | # | 20 | Baseline | ancom | 45 |
| # | time | 5 | 4 | 0 | 0.555555556 | 1 | # | 20 | Baseline | ancom | 45 |
| # | interaction | 0 | 0 | 4 | 0 | 0 | # | 20 | Baseline | ancom | 45 |
| # | disease | 7 | 57 | 0 | 0.094594595 | 1 | # | 20 | Baseline | aldex2 | 45 |
| # | sex | 4 | 15 | 0 | 0.210526316 | 1 | # | 20 | Baseline | aldex2 | 45 |
| # | time | 5 | 37 | 0 | 0.119047619 | 1 | # | 20 | Baseline | aldex2 | 45 |
| # | interaction | 0 | 0 | 4 | 0 | 0 | # | 20 | Baseline | aldex2 | 45 |
| # | disease | 7 | 0 | 0 | 1 | 1 | # | 20 | Small | paralle | 45 |
| # | sex | 4 | 0 | 0 | 1 | 1 | # | 20 | Small | paralle | 45 |
| # | time | 5 | 0 | 0 | 1 | 1 | # | 20 | Small | paralle | 45 |
| # | interaction | 4 | 0 | 0 | 1 | 1 | # | 20 | Small | paralle | 45 |
| # | disease | 7 | 4 | 0 | 0.636363636 | 1 | # | 20 | Small | ancom | 45 |
| # | sex | 4 | 0 | 0 | 1 | 1 | # | 20 | Small | ancom | 45 |
| # | time | 5 | 4 | 0 | 0.555555556 | 1 | # | 20 | Small | ancom | 45 |
| # | interaction | 0 | 0 | 4 | 0 | 0 | # | 20 | Small | ancom | 45 |
| # | disease | 7 | 12 | 0 | 0.368421053 | 1 | # | 20 | Small | aldex2 | 45 |
| # | sex | 4 | 1 | 0 | 0.8 | 1 | # | 20 | Small | aldex2 | 45 |
| # | time | 5 | 7 | 0 | 0.416666667 | 1 | # | 20 | Small | aldex2 | 45 |
| # | interaction | 0 | 0 | 4 | 0 | 0 | # | 20 | Small | aldex2 | 45 |
| # | disease | 4 | 9 | 0 | 0.307692308 | 1 | # | 10 | Baseline | paralle | 46 |
| # | sex | 2 | 5 | 0 | 0.285714286 | 1 | # | 10 | Baseline | paralle | 46 |
| # | time | 2 | 10 | 0 | 0.166666667 | 1 | # | 10 | Baseline | paralle | 46 |
| # | interaction | 2 | 10 | 0 | 0.166666667 | 1 | # | 10 | Baseline | paralle | 46 |
| # | disease | 4 | 2 | 0 | 0.666666667 | 1 | # | 10 | Baseline | ancom | 46 |
| # | sex | 2 | 0 | 0 | 1 | 1 | # | 10 | Baseline | ancom | 46 |
| # | time | 2 | 0 | 0 | 1 | 1 | # | 10 | Baseline | ancom | 46 |
| # | interaction | 0 | 0 | 2 | 0 | 0 | # | 10 | Baseline | ancom | 46 |
| # | disease | 4 | 8 | 0 | 0.333333333 | 1 | # | 10 | Baseline | aldex2 | 46 |
| # | sex | 2 | 5 | 0 | 0.285714286 | 1 | # | 10 | Baseline | aldex2 | 46 |
| # | time | 2 | 9 | 0 | 0.181818182 | 1 | # | 10 | Baseline | aldex2 | 46 |
| # | interaction | 0 | 0 | 2 | 0 | 0 | # | 10 | Baseline | aldex2 | 46 |
| # | disease | 4 | 0 | 0 | 1 | 1 | # | 10 | Small | paralle | 46 |
| # | sex | 2 | 0 | 0 | 1 | 1 | # | 10 | Small | paralle | 46 |
| # | time | 2 | 0 | 0 | 1 | 1 | # | 10 | Small | paralle | 46 |
| # | interaction | 0 | 0 | 2 | 0 | 0 | # | 10 | Small | paralle | 46 |
| # | disease | 4 | 2 | 0 | 0.666666667 | 1 | # | 10 | Small | ancom | 46 |
| # | sex | 2 | 0 | 0 | 1 | 1 | # | 10 | Small | ancom | 46 |
| # | time | 2 | 0 | 0 | 1 | 1 | # | 10 | Small | ancom | 46 |
| # | interaction | 0 | 0 | 2 | 0 | 0 | # | 10 | Small | ancom | 46 |
| # | disease | 4 | 2 | 0 | 0.666666667 | 1 | # | 10 | Small | aldex2 | 46 |
| # | sex | 2 | 0 | 0 | 1 | 1 | # | 10 | Small | aldex2 | 46 |
| # | time | 2 | 0 | 0 | 1 | 1 | # | 10 | Small | aldex2 | 46 |
| # | interaction | 0 | 0 | 2 | 0 | 0 | # | 10 | Small | aldex2 | 46 |
| # | disease | 4 | 6 | 0 | 0.4 | 1 | # | 10 | Baseline | paralle | 46 |
| # | sex | 2 | 3 | 0 | 0.4 | 1 | # | 10 | Baseline | paralle | 46 |
| # | time | 2 | 8 | 0 | 0.2 | 1 | # | 10 | Baseline | paralle | 46 |
| # | interaction | 2 | 3 | 0 | 0.4 | 1 | # | 10 | Baseline | paralle | 46 |
| # | disease | 4 | 2 | 0 | 0.666666667 | 1 | # | 10 | Baseline | ancom | 46 |
| # | sex | 2 | 0 | 0 | 1 | 1 | # | 10 | Baseline | ancom | 46 |
| # | time | 2 | 2 | 0 | 0.5 | 1 | # | 10 | Baseline | ancom | 46 |
| # | interaction | 0 | 0 | 2 | 0 | 0 | # | 10 | Baseline | ancom | 46 |
| # | disease | 4 | 11 | 0 | 0.266666667 | 1 | # | 10 | Baseline | aldex2 | 46 |
| # | sex | 2 | 4 | 0 | 0.333333333 | 1 | # | 10 | Baseline | aldex2 | 46 |
| # | time | 2 | 9 | 0 | 0.181818182 | 1 | # | 10 | Baseline | aldex2 | 46 |
| # | interaction | 0 | 0 | 2 | 0 | 0 | # | 10 | Baseline | aldex2 | 46 |
| # | disease | 4 | 0 | 0 | 1 | 1 | # | 10 | Small | paralle | 46 |
| # | sex | 2 | 0 | 0 | 1 | 1 | # | 10 | Small | paralle | 46 |
| # | time | 2 | 0 | 0 | 1 | 1 | # | 10 | Small | paralle | 46 |
| # | interaction | 2 | 0 | 0 | 1 | 1 | # | 10 | Small | paralle | 46 |
| # | disease | 4 | 2 | 0 | 0.666666667 | 1 | # | 10 | Small | ancom | 46 |
| # | sex | 2 | 0 | 0 | 1 | 1 | # | 10 | Small | ancom | 46 |
| # | time | 2 | 2 | 0 | 0.5 | 1 | # | 10 | Small | ancom | 46 |
| # | interaction | 0 | 0 | 2 | 0 | 0 | # | 10 | Small | ancom | 46 |
| # | disease | 4 | 1 | 0 | 0.8 | 1 | # | 10 | Small | aldex2 | 46 |
| # | sex | 2 | 0 | 0 | 1 | 1 | # | 10 | Small | aldex2 | 46 |
| # | time | 2 | 2 | 0 | 0.5 | 1 | # | 10 | Small | aldex2 | 46 |
| # | interaction | 0 | 0 | 2 | 0 | 0 | # | 10 | Small | aldex2 | 46 |
| # | disease | 4 | 5 | 0 | 0.444444444 | 1 | # | 10 | Baseline | paralle | 46 |
| # | sex | 2 | 4 | 0 | 0.333333333 | 1 | # | 10 | Baseline | paralle | 46 |
| # | time | 2 | 9 | 0 | 0.181818182 | 1 | # | 10 | Baseline | paralle | 46 |
| # | interaction | 2 | 6 | 0 | 0.25 | 1 | # | 10 | Baseline | paralle | 46 |
| # | disease | 4 | 2 | 0 | 0.666666667 | 1 | # | 10 | Baseline | ancom | 46 |
| # | sex | 2 | 0 | 0 | 1 | 1 | # | 10 | Baseline | ancom | 46 |
| # | time | 2 | 2 | 0 | 0.5 | 1 | # | 10 | Baseline | ancom | 46 |
| # | interaction | 0 | 0 | 2 | 0 | 0 | # | 10 | Baseline | ancom | 46 |
| # | disease | 4 | 18 | 0 | 0.181818182 | 1 | # | 10 | Baseline | aldex2 | 46 |
| # | sex | 2 | 7 | 0 | 0.222222222 | 1 | # | 10 | Baseline | aldex2 | 46 |
| # | time | 2 | 11 | 0 | 0.153846154 | 1 | # | 10 | Baseline | aldex2 | 46 |
| # | interaction | 0 | 0 | 2 | 0 | 0 | # | 10 | Baseline | aldex2 | 46 |
| # | disease | 4 | 0 | 0 | 1 | 1 | # | 10 | Small | paralle | 46 |
| # | sex | 2 | 0 | 0 | 1 | 1 | # | 10 | Small | paralle | 46 |
| # | time | 2 | 0 | 0 | 1 | 1 | # | 10 | Small | paralle | 46 |
| # | interaction | 2 | 0 | 0 | 1 | 1 | # | 10 | Small | paralle | 46 |
| # | disease | 4 | 2 | 0 | 0.666666667 | 1 | # | 10 | Small | ancom | 46 |
| # | sex | 2 | 0 | 0 | 1 | 1 | # | 10 | Small | ancom | 46 |
| # | time | 2 | 2 | 0 | 0.5 | 1 | # | 10 | Small | ancom | 46 |
| # | interaction | 0 | 0 | 2 | 0 | 0 | # | 10 | Small | ancom | 46 |
| # | disease | 4 | 2 | 0 | 0.666666667 | 1 | # | 10 | Small | aldex2 | 46 |
| # | sex | 2 | 0 | 0 | 1 | 1 | # | 10 | Small | aldex2 | 46 |
| # | time | 2 | 2 | 0 | 0.5 | 1 | # | 10 | Small | aldex2 | 46 |
| # | interaction | 0 | 0 | 2 | 0 | 0 | # | 10 | Small | aldex2 | 46 |
| # | disease | 4 | 5 | 0 | 0.444444444 | 1 | # | 10 | Baseline | paralle | 46 |
| # | sex | 2 | 3 | 0 | 0.4 | 1 | # | 10 | Baseline | paralle | 46 |
| # | time | 2 | 7 | 0 | 0.222222222 | 1 | # | 10 | Baseline | paralle | 46 |
| # | interaction | 2 | 3 | 0 | 0.4 | 1 | # | 10 | Baseline | paralle | 46 |
| # | disease | 4 | 2 | 0 | 0.666666667 | 1 | # | 10 | Baseline | ancom | 46 |
| # | sex | 2 | 0 | 0 | 1 | 1 | # | 10 | Baseline | ancom | 46 |
| # | time | 2 | 2 | 0 | 0.5 | 1 | # | 10 | Baseline | ancom | 46 |
| # | interaction | 0 | 0 | 2 | 0 | 0 | # | 10 | Baseline | ancom | 46 |
| # | disease | 4 | 21 | 0 | 0.16 | 1 | # | 10 | Baseline | aldex2 | 46 |
| # | sex | 2 | 6 | 0 | 0.25 | 1 | # | 10 | Baseline | aldex2 | 46 |
| # | time | 2 | 13 | 0 | 0.133333333 | 1 | # | 10 | Baseline | aldex2 | 46 |
| # | interaction | 0 | 0 | 2 | 0 | 0 | # | 10 | Baseline | aldex2 | 46 |
| # | disease | 4 | 0 | 0 | 1 | 1 | # | 10 | Small | paralle | 46 |
| # | sex | 2 | 0 | 0 | 1 | 1 | # | 10 | Small | paralle | 46 |
| # | time | 2 | 1 | 0 | 0.666666667 | 1 | # | 10 | Small | paralle | 46 |
| # | interaction | 2 | 0 | 0 | 1 | 1 | # | 10 | Small | paralle | 46 |
| # | disease | 4 | 2 | 0 | 0.666666667 | 1 | # | 10 | Small | ancom | 46 |
| # | sex | 2 | 0 | 0 | 1 | 1 | # | 10 | Small | ancom | 46 |
| # | time | 2 | 2 | 0 | 0.5 | 1 | # | 10 | Small | ancom | 46 |
| # | interaction | 0 | 0 | 2 | 0 | 0 | # | 10 | Small | ancom | 46 |
| # | disease | 4 | 2 | 0 | 0.666666667 | 1 | # | 10 | Small | aldex2 | 46 |
| # | sex | 2 | 0 | 0 | 1 | 1 | # | 10 | Small | aldex2 | 46 |
| # | time | 2 | 3 | 0 | 0.4 | 1 | # | 10 | Small | aldex2 | 46 |
| # | interaction | 0 | 0 | 2 | 0 | 0 | # | 10 | Small | aldex2 | 46 |
| # | disease | 7 | 6 | 0 | 0.538461538 | 1 | # | 20 | Baseline | paralle | 46 |
| # | sex | 4 | 12 | 0 | 0.25 | 1 | # | 20 | Baseline | paralle | 46 |
| # | time | 5 | 6 | 0 | 0.454545455 | 1 | # | 20 | Baseline | paralle | 46 |
| # | interaction | 4 | 8 | 0 | 0.333333333 | 1 | # | 20 | Baseline | paralle | 46 |
| # | disease | 7 | 2 | 0 | 0.777777778 | 1 | # | 20 | Baseline | ancom | 46 |
| # | sex | 4 | 0 | 0 | 1 | 1 | # | 20 | Baseline | ancom | 46 |
| # | time | 5 | 0 | 0 | 1 | 1 | # | 20 | Baseline | ancom | 46 |
| # | interaction | 0 | 0 | 4 | 0 | 0 | # | 20 | Baseline | ancom | 46 |
| # | disease | 7 | 27 | 0 | 0.205882353 | 1 | # | 20 | Baseline | aldex2 | 46 |
| # | sex | 4 | 7 | 0 | 0.363636364 | 1 | # | 20 | Baseline | aldex2 | 46 |
| # | time | 5 | 16 | 0 | 0.238095238 | 1 | # | 20 | Baseline | aldex2 | 46 |
| # | interaction | 0 | 0 | 4 | 0 | 0 | # | 20 | Baseline | aldex2 | 46 |
| # | disease | 7 | 0 | 0 | 1 | 1 | # | 20 | Small | paralle | 46 |
| # | sex | 4 | 0 | 0 | 1 | 1 | # | 20 | Small | paralle | 46 |
| # | time | 5 | 1 | 0 | 0.833333333 | 1 | # | 20 | Small | paralle | 46 |
| # | interaction | 2 | 0 | 2 | 1 | 0.5 | # | 20 | Small | paralle | 46 |
| # | disease | 7 | 1 | 0 | 0.875 | 1 | # | 20 | Small | ancom | 46 |
| # | sex | 4 | 0 | 0 | 1 | 1 | # | 20 | Small | ancom | 46 |
| # | time | 5 | 0 | 0 | 1 | 1 | # | 20 | Small | ancom | 46 |
| # | interaction | 0 | 0 | 4 | 0 | 0 | # | 20 | Small | ancom | 46 |
| # | disease | 7 | 2 | 0 | 0.777777778 | 1 | # | 20 | Small | aldex2 | 46 |
| # | sex | 4 | 0 | 0 | 1 | 1 | # | 20 | Small | aldex2 | 46 |
| # | time | 5 | 0 | 0 | 1 | 1 | # | 20 | Small | aldex2 | 46 |

| | | | | | | | | | | | |
|---|---|---|---|---|---|---|---|---|---|---|---|
| # | interaction | 0 | 0 | 4 | 0 | 0 | # | 20 | Small | aldex2 | 46 |
| # | disease | 7 | 8 | 0 | 0.466666667 | 1 | # | 20 | Baseline | paralle | 46 |
| # | sex | 4 | 8 | 0 | 0.333333333 | 1 | # | 20 | Baseline | paralle | 46 |
| # | time | 5 | 7 | 0 | 0.416666667 | 1 | # | 20 | Baseline | paralle | 46 |
| # | interaction | 4 | 5 | 0 | 0.444444444 | 1 | # | 20 | Baseline | paralle | 46 |
| # | disease | 7 | 4 | 0 | 0.636363636 | 1 | # | 20 | Baseline | ancom | 46 |
| # | sex | 4 | 0 | 0 | 1 | 1 | # | 20 | Baseline | ancom | 46 |
| # | time | 5 | 3 | 0 | 0.625 | 1 | # | 20 | Baseline | ancom | 46 |
| # | interaction | 0 | 0 | 4 | 0 | 0 | # | 20 | Baseline | ancom | 46 |
| # | disease | 7 | 39 | 0 | 0.152173913 | 1 | # | 20 | Baseline | aldex2 | 46 |
| # | sex | 4 | 12 | 0 | 0.25 | 1 | # | 20 | Baseline | aldex2 | 46 |
| # | time | 5 | 25 | 0 | 0.166666667 | 1 | # | 20 | Baseline | aldex2 | 46 |
| # | interaction | 0 | 0 | 4 | 0 | 0 | # | 20 | Baseline | aldex2 | 46 |
| # | disease | 7 | 0 | 0 | 1 | 1 | # | 20 | Small | paralle | 46 |
| # | sex | 4 | 0 | 0 | 1 | 1 | # | 20 | Small | paralle | 46 |
| # | time | 5 | 0 | 0 | 1 | 1 | # | 20 | Small | paralle | 46 |
| # | interaction | 3 | 0 | 1 | 1 | 0.75 | # | 20 | Small | paralle | 46 |
| # | disease | 7 | 4 | 0 | 0.636363636 | 1 | # | 20 | Small | ancom | 46 |
| # | sex | 4 | 0 | 0 | 1 | 1 | # | 20 | Small | ancom | 46 |
| # | time | 5 | 3 | 0 | 0.625 | 1 | # | 20 | Small | ancom | 46 |
| # | interaction | 0 | 0 | 4 | 0 | 0 | # | 20 | Small | ancom | 46 |
| # | disease | 7 | 6 | 0 | 0.538461538 | 1 | # | 20 | Small | aldex2 | 46 |
| # | sex | 4 | 0 | 0 | 1 | 1 | # | 20 | Small | aldex2 | 46 |
| # | time | 5 | 1 | 0 | 0.833333333 | 1 | # | 20 | Small | aldex2 | 46 |
| # | interaction | 0 | 0 | 4 | 0 | 0 | # | 20 | Small | aldex2 | 46 |
| # | disease | 7 | 9 | 0 | 0.4375 | 1 | # | 20 | Baseline | paralle | 46 |
| # | sex | 4 | 6 | 0 | 0.4 | 1 | # | 20 | Baseline | paralle | 46 |
| # | time | 5 | 10 | 0 | 0.333333333 | 1 | # | 20 | Baseline | paralle | 46 |
| # | interaction | 4 | 2 | 0 | 0.666666667 | 1 | # | 20 | Baseline | paralle | 46 |
| # | disease | 7 | 4 | 0 | 0.636363636 | 1 | # | 20 | Baseline | ancom | 46 |
| # | sex | 4 | 0 | 0 | 1 | 1 | # | 20 | Baseline | ancom | 46 |
| # | time | 5 | 4 | 0 | 0.555555556 | 1 | # | 20 | Baseline | ancom | 46 |
| # | interaction | 0 | 0 | 4 | 0 | 0 | # | 20 | Baseline | ancom | 46 |
| # | disease | 7 | 50 | 0 | 0.104477612 | 1 | # | 20 | Baseline | aldex2 | 46 |
| # | sex | 4 | 10 | 0 | 0.285714286 | 1 | # | 20 | Baseline | aldex2 | 46 |
| # | time | 5 | 34 | 0 | 0.128205128 | 1 | # | 20 | Baseline | aldex2 | 46 |
| # | interaction | 0 | 0 | 4 | 0 | 0 | # | 20 | Baseline | aldex2 | 46 |
| # | disease | 7 | 0 | 0 | 1 | 1 | # | 20 | Small | paralle | 46 |
| # | sex | 4 | 0 | 0 | 1 | 1 | # | 20 | Small | paralle | 46 |
| # | time | 5 | 0 | 0 | 1 | 1 | # | 20 | Small | paralle | 46 |
| # | interaction | 4 | 0 | 0 | 1 | 1 | # | 20 | Small | paralle | 46 |
| # | disease | 7 | 4 | 0 | 0.636363636 | 1 | # | 20 | Small | ancom | 46 |
| # | sex | 4 | 0 | 0 | 1 | 1 | # | 20 | Small | ancom | 46 |
| # | time | 5 | 4 | 0 | 0.555555556 | 1 | # | 20 | Small | ancom | 46 |
| # | interaction | 0 | 0 | 4 | 0 | 0 | # | 20 | Small | ancom | 46 |
| # | disease | 7 | 8 | 0 | 0.466666667 | 1 | # | 20 | Small | aldex2 | 46 |
| # | sex | 4 | 0 | 0 | 1 | 1 | # | 20 | Small | aldex2 | 46 |
| # | time | 5 | 5 | 0 | 0.5 | 1 | # | 20 | Small | aldex2 | 46 |
| # | interaction | 0 | 0 | 4 | 0 | 0 | # | 20 | Small | aldex2 | 46 |
| # | disease | 7 | 7 | 0 | 0.5 | 1 | # | 20 | Baseline | paralle | 46 |
| # | sex | 4 | 6 | 0 | 0.4 | 1 | # | 20 | Baseline | paralle | 46 |
| # | time | 5 | 7 | 0 | 0.416666667 | 1 | # | 20 | Baseline | paralle | 46 |
| # | interaction | 4 | 3 | 0 | 0.571428571 | 1 | # | 20 | Baseline | paralle | 46 |
| # | disease | 7 | 4 | 0 | 0.636363636 | 1 | # | 20 | Baseline | ancom | 46 |
| # | sex | 4 | 0 | 0 | 1 | 1 | # | 20 | Baseline | ancom | 46 |
| # | time | 5 | 4 | 0 | 0.555555556 | 1 | # | 20 | Baseline | ancom | 46 |
| # | interaction | 0 | 0 | 4 | 0 | 0 | # | 20 | Baseline | ancom | 46 |
| # | disease | 7 | 72 | 0 | 0.088607595 | 1 | # | 20 | Baseline | aldex2 | 46 |
| # | sex | 4 | 20 | 0 | 0.166666667 | 1 | # | 20 | Baseline | aldex2 | 46 |
| # | time | 5 | 62 | 0 | 0.106382979 | 1 | # | 20 | Baseline | aldex2 | 46 |
| # | interaction | 0 | 0 | 4 | 0 | 0 | # | 20 | Baseline | aldex2 | 46 |
| # | disease | 7 | 0 | 0 | 1 | 1 | # | 20 | Small | paralle | 46 |
| # | sex | 4 | 0 | 0 | 1 | 1 | # | 20 | Small | paralle | 46 |
| # | time | 5 | 0 | 0 | 1 | 1 | # | 20 | Small | paralle | 46 |
| # | interaction | 4 | 0 | 0 | 1 | 1 | # | 20 | Small | paralle | 46 |
| # | disease | 7 | 4 | 0 | 0.636363636 | 1 | # | 20 | Small | ancom | 46 |
| # | sex | 4 | 0 | 0 | 1 | 1 | # | 20 | Small | ancom | 46 |
| # | time | 5 | 4 | 0 | 0.555555556 | 1 | # | 20 | Small | ancom | 46 |
| # | interaction | 0 | 0 | 4 | 0 | 0 | # | 20 | Small | ancom | 46 |
| # | disease | 7 | 9 | 0 | 0.4375 | 1 | # | 20 | Small | aldex2 | 46 |
| # | sex | 4 | 1 | 0 | 0.8 | 1 | # | 20 | Small | aldex2 | 46 |
| # | time | 5 | 7 | 0 | 0.416666667 | 1 | # | 20 | Small | aldex2 | 46 |
| # | interaction | 0 | 0 | 4 | 0 | 0 | # | 20 | Small | aldex2 | 46 |
| # | disease | 4 | 12 | 0 | 0.25 | 1 | # | 10 | Baseline | paralle | 47 |
| # | sex | 2 | 6 | 0 | 0.25 | 1 | # | 10 | Baseline | paralle | 47 |
| # | time | 2 | 4 | 0 | 0.333333333 | 1 | # | 10 | Baseline | paralle | 47 |
| # | interaction | 2 | 15 | 0 | 0.117647059 | 1 | # | 10 | Baseline | paralle | 47 |
| # | disease | 4 | 1 | 0 | 0.8 | 1 | # | 10 | Baseline | ancom | 47 |
| # | sex | 2 | 0 | 0 | 1 | 1 | # | 10 | Baseline | ancom | 47 |
| # | time | 2 | 0 | 0 | 1 | 1 | # | 10 | Baseline | ancom | 47 |
| # | interaction | 0 | 0 | 2 | 0 | 0 | # | 10 | Baseline | ancom | 47 |
| # | disease | 4 | 8 | 0 | 0.333333333 | 1 | # | 10 | Baseline | aldex2 | 47 |
| # | sex | 2 | 1 | 0 | 0.666666667 | 1 | # | 10 | Baseline | aldex2 | 47 |
| # | time | 2 | 5 | 0 | 0.285714286 | 1 | # | 10 | Baseline | aldex2 | 47 |
| # | interaction | 0 | 0 | 2 | 0 | 0 | # | 10 | Baseline | aldex2 | 47 |
| # | disease | 4 | 0 | 0 | 1 | 1 | # | 10 | Small | paralle | 47 |
| # | sex | 2 | 0 | 0 | 1 | 1 | # | 10 | Small | paralle | 47 |
| # | time | 2 | 0 | 0 | 1 | 1 | # | 10 | Small | paralle | 47 |
| # | interaction | 1 | 0 | 1 | 1 | 0.5 | # | 10 | Small | paralle | 47 |
| # | disease | 4 | 1 | 0 | 0.8 | 1 | # | 10 | Small | ancom | 47 |
| # | sex | 2 | 0 | 0 | 1 | 1 | # | 10 | Small | ancom | 47 |
| # | time | 2 | 0 | 0 | 1 | 1 | # | 10 | Small | ancom | 47 |
| # | interaction | 0 | 0 | 2 | 0 | 0 | # | 10 | Small | ancom | 47 |
| # | disease | 4 | 1 | 0 | 0.8 | 1 | # | 10 | Small | aldex2 | 47 |
| # | sex | 2 | 0 | 0 | 1 | 1 | # | 10 | Small | aldex2 | 47 |
| # | time | 2 | 0 | 0 | 1 | 1 | # | 10 | Small | aldex2 | 47 |
| # | interaction | 0 | 0 | 2 | 0 | 0 | # | 10 | Small | aldex2 | 47 |
| # | disease | 4 | 3 | 0 | 0.571428571 | 1 | # | 10 | Baseline | paralle | 47 |
| # | sex | 2 | 5 | 0 | 0.285714286 | 1 | # | 10 | Baseline | paralle | 47 |
| # | time | 2 | 6 | 0 | 0.25 | 1 | # | 10 | Baseline | paralle | 47 |
| # | interaction | 2 | 7 | 0 | 0.222222222 | 1 | # | 10 | Baseline | paralle | 47 |
| # | disease | 4 | 2 | 0 | 0.666666667 | 1 | # | 10 | Baseline | ancom | 47 |
| # | sex | 2 | 0 | 0 | 1 | 1 | # | 10 | Baseline | ancom | 47 |
| # | time | 2 | 2 | 0 | 0.5 | 1 | # | 10 | Baseline | ancom | 47 |
| # | interaction | 0 | 0 | 2 | 0 | 0 | # | 10 | Baseline | ancom | 47 |
| # | disease | 4 | 9 | 0 | 0.307692308 | 1 | # | 10 | Baseline | aldex2 | 47 |
| # | sex | 2 | 6 | 0 | 0.25 | 1 | # | 10 | Baseline | aldex2 | 47 |
| # | time | 2 | 7 | 0 | 0.222222222 | 1 | # | 10 | Baseline | aldex2 | 47 |
| # | interaction | 0 | 0 | 2 | 0 | 0 | # | 10 | Baseline | aldex2 | 47 |
| # | disease | 4 | 0 | 0 | 1 | 1 | # | 10 | Small | paralle | 47 |
| # | sex | 2 | 0 | 0 | 1 | 1 | # | 10 | Small | paralle | 47 |
| # | time | 2 | 0 | 0 | 1 | 1 | # | 10 | Small | paralle | 47 |
| # | interaction | 2 | 0 | 0 | 1 | 1 | # | 10 | Small | paralle | 47 |
| # | disease | 4 | 2 | 0 | 0.666666667 | 1 | # | 10 | Small | ancom | 47 |
| # | sex | 2 | 0 | 0 | 1 | 1 | # | 10 | Small | ancom | 47 |
| # | time | 2 | 2 | 0 | 0.5 | 1 | # | 10 | Small | ancom | 47 |
| # | interaction | 0 | 0 | 2 | 0 | 0 | # | 10 | Small | ancom | 47 |
| # | disease | 4 | 3 | 0 | 0.571428571 | 1 | # | 10 | Small | aldex2 | 47 |
| # | sex | 2 | 0 | 0 | 1 | 1 | # | 10 | Small | aldex2 | 47 |
| # | time | 2 | 2 | 0 | 0.5 | 1 | # | 10 | Small | aldex2 | 47 |
| # | interaction | 0 | 0 | 2 | 0 | 0 | # | 10 | Small | aldex2 | 47 |
| # | disease | 4 | 7 | 0 | 0.363636364 | 1 | # | 10 | Baseline | paralle | 47 |
| # | sex | 2 | 2 | 0 | 0.5 | 1 | # | 10 | Baseline | paralle | 47 |
| # | time | 2 | 7 | 0 | 0.222222222 | 1 | # | 10 | Baseline | paralle | 47 |
| # | interaction | 2 | 8 | 0 | 0.2 | 1 | # | 10 | Baseline | paralle | 47 |
| # | disease | 4 | 2 | 0 | 0.666666667 | 1 | # | 10 | Baseline | ancom | 47 |
| # | sex | 2 | 0 | 0 | 1 | 1 | # | 10 | Baseline | ancom | 47 |
| # | time | 2 | 2 | 0 | 0.5 | 1 | # | 10 | Baseline | ancom | 47 |
| # | interaction | 0 | 0 | 2 | 0 | 0 | # | 10 | Baseline | ancom | 47 |
| # | disease | 4 | 18 | 0 | 0.181818182 | 1 | # | 10 | Baseline | aldex2 | 47 |
| # | sex | 2 | 4 | 0 | 0.333333333 | 1 | # | 10 | Baseline | aldex2 | 47 |
| # | time | 2 | 6 | 0 | 0.25 | 1 | # | 10 | Baseline | aldex2 | 47 |
| # | interaction | 0 | 0 | 2 | 0 | 0 | # | 10 | Baseline | aldex2 | 47 |
| # | disease | 4 | 0 | 0 | 1 | 1 | # | 10 | Small | paralle | 47 |
| # | sex | 2 | 0 | 0 | 1 | 1 | # | 10 | Small | paralle | 47 |
| # | time | 2 | 1 | 0 | 0.666666667 | 1 | # | 10 | Small | paralle | 47 |
| # | interaction | 2 | 1 | 0 | 0.666666667 | 1 | # | 10 | Small | paralle | 47 |
| # | disease | 4 | 2 | 0 | 0.666666667 | 1 | # | 10 | Small | ancom | 47 |
| # | sex | 2 | 0 | 0 | 1 | 1 | # | 10 | Small | ancom | 47 |
| # | time | 2 | 2 | 0 | 0.5 | 1 | # | 10 | Small | ancom | 47 |
| # | interaction | 0 | 0 | 2 | 0 | 0 | # | 10 | Small | ancom | 47 |
| # | disease | 4 | 3 | 0 | 0.571428571 | 1 | # | 10 | Small | aldex2 | 47 |
| # | sex | 2 | 0 | 0 | 1 | 1 | # | 10 | Small | aldex2 | 47 |
| # | time | 2 | 2 | 0 | 0.5 | 1 | # | 10 | Small | aldex2 | 47 |
| # | interaction | 0 | 0 | 2 | 0 | 0 | # | 10 | Small | aldex2 | 47 |
| # | disease | 4 | 6 | 0 | 0.4 | 1 | # | 10 | Baseline | paralle | 47 |
| # | sex | 2 | 1 | 0 | 0.666666667 | 1 | # | 10 | Baseline | paralle | 47 |
| # | time | 2 | 8 | 0 | 0.2 | 1 | # | 10 | Baseline | paralle | 47 |
| # | interaction | 2 | 11 | 0 | 0.153846154 | 1 | # | 10 | Baseline | paralle | 47 |
| # | disease | 4 | 2 | 0 | 0.666666667 | 1 | # | 10 | Baseline | ancom | 47 |
| # | sex | 2 | 0 | 0 | 1 | 1 | # | 10 | Baseline | ancom | 47 |
| # | time | 2 | 2 | 0 | 0.5 | 1 | # | 10 | Baseline | ancom | 47 |
| # | interaction | 0 | 0 | 2 | 0 | 0 | # | 10 | Baseline | ancom | 47 |
| # | disease | 4 | 20 | 0 | 0.166666667 | 1 | # | 10 | Baseline | aldex2 | 47 |
| # | sex | 2 | 6 | 0 | 0.25 | 1 | # | 10 | Baseline | aldex2 | 47 |
| # | time | 2 | 12 | 0 | 0.142857143 | 1 | # | 10 | Baseline | aldex2 | 47 |
| # | interaction | 0 | 0 | 2 | 0 | 0 | # | 10 | Baseline | aldex2 | 47 |
| # | disease | 4 | 0 | 0 | 1 | 1 | # | 10 | Small | paralle | 47 |
| # | sex | 2 | 0 | 0 | 1 | 1 | # | 10 | Small | paralle | 47 |
| # | time | 2 | 0 | 0 | 1 | 1 | # | 10 | Small | paralle | 47 |
| # | interaction | 2 | 0 | 0 | 1 | 1 | # | 10 | Small | paralle | 47 |
| # | disease | 4 | 2 | 0 | 0.666666667 | 1 | # | 10 | Small | ancom | 47 |
| # | sex | 2 | 0 | 0 | 1 | 1 | # | 10 | Small | ancom | 47 |
| # | time | 2 | 2 | 0 | 0.5 | 1 | # | 10 | Small | ancom | 47 |
| # | interaction | 0 | 0 | 2 | 0 | 0 | # | 10 | Small | ancom | 47 |
| # | disease | 4 | 2 | 0 | 0.666666667 | 1 | # | 10 | Small | aldex2 | 47 |
| # | sex | 2 | 0 | 0 | 1 | 1 | # | 10 | Small | aldex2 | 47 |
| # | time | 2 | 2 | 0 | 0.5 | 1 | # | 10 | Small | aldex2 | 47 |
| # | interaction | 0 | 0 | 2 | 0 | 0 | # | 10 | Small | aldex2 | 47 |
| # | disease | 7 | 8 | 0 | 0.466666667 | 1 | # | 20 | Baseline | paralle | 47 |
| # | sex | 4 | 7 | 0 | 0.363636364 | 1 | # | 20 | Baseline | paralle | 47 |
| # | time | 5 | 9 | 0 | 0.357142857 | 1 | # | 20 | Baseline | paralle | 47 |
| # | interaction | 4 | 8 | 0 | 0.333333333 | 1 | # | 20 | Baseline | paralle | 47 |
| # | disease | 7 | 3 | 0 | 0.7 | 1 | # | 20 | Baseline | ancom | 47 |
| # | sex | 4 | 0 | 0 | 1 | 1 | # | 20 | Baseline | ancom | 47 |
| # | time | 5 | 1 | 0 | 0.833333333 | 1 | # | 20 | Baseline | ancom | 47 |
| # | interaction | 0 | 0 | 4 | 0 | 0 | # | 20 | Baseline | ancom | 47 |
| # | disease | 7 | 24 | 0 | 0.225806452 | 1 | # | 20 | Baseline | aldex2 | 47 |
| # | sex | 4 | 3 | 0 | 0.571428571 | 1 | # | 20 | Baseline | aldex2 | 47 |
| # | time | 5 | 16 | 0 | 0.238095238 | 1 | # | 20 | Baseline | aldex2 | 47 |
| # | interaction | 0 | 0 | 4 | 0 | 0 | # | 20 | Baseline | aldex2 | 47 |
| # | disease | 7 | 0 | 0 | 1 | 1 | # | 20 | Small | paralle | 47 |
| # | sex | 4 | 0 | 0 | 1 | 1 | # | 20 | Small | paralle | 47 |
| # | time | 5 | 0 | 0 | 1 | 1 | # | 20 | Small | paralle | 47 |
| # | interaction | 3 | 0 | 1 | 1 | 0.75 | # | 20 | Small | paralle | 47 |
| # | disease | 7 | 2 | 0 | 0.777777778 | 1 | # | 20 | Small | ancom | 47 |
| # | sex | 4 | 0 | 0 | 1 | 1 | # | 20 | Small | ancom | 47 |
| # | time | 5 | 1 | 0 | 0.833333333 | 1 | # | 20 | Small | ancom | 47 |
| # | interaction | 0 | 0 | 4 | 0 | 0 | # | 20 | Small | ancom | 47 |
| # | disease | 7 | 2 | 0 | 0.777777778 | 1 | # | 20 | Small | aldex2 | 47 |
| # | sex | 4 | 1 | 0 | 0.8 | 1 | # | 20 | Small | aldex2 | 47 |
| # | time | 5 | 1 | 0 | 0.833333333 | 1 | # | 20 | Small | aldex2 | 47 |
| # | interaction | 0 | 0 | 4 | 0 | 0 | # | 20 | Small | aldex2 | 47 |
| # | disease | 7 | 17 | 0 | 0.291666667 | 1 | # | 20 | Baseline | paralle | 47 |
| # | sex | 4 | 7 | 0 | 0.363636364 | 1 | # | 20 | Baseline | paralle | 47 |
| # | time | 5 | 6 | 0 | 0.454545455 | 1 | # | 20 | Baseline | paralle | 47 |
| # | interaction | 4 | 8 | 0 | 0.333333333 | 1 | # | 20 | Baseline | paralle | 47 |
| # | disease | 7 | 3 | 0 | 0.7 | 1 | # | 20 | Baseline | ancom | 47 |
| # | sex | 4 | 0 | 0 | 1 | 1 | # | 20 | Baseline | ancom | 47 |
| # | time | 5 | 4 | 0 | 0.555555556 | 1 | # | 20 | Baseline | ancom | 47 |
| # | interaction | 0 | 0 | 4 | 0 | 0 | # | 20 | Baseline | ancom | 47 |
| # | disease | 7 | 37 | 0 | 0.159090909 | 1 | # | 20 | Baseline | aldex2 | 47 |
| # | sex | 4 | 13 | 0 | 0.235294118 | 1 | # | 20 | Baseline | aldex2 | 47 |
| # | time | 5 | 22 | 0 | 0.185185185 | 1 | # | 20 | Baseline | aldex2 | 47 |
| # | interaction | 0 | 0 | 4 | 0 | 0 | # | 20 | Baseline | aldex2 | 47 |
| # | disease | 7 | 1 | 0 | 0.875 | 1 | # | 20 | Small | paralle | 47 |
| # | sex | 4 | 0 | 0 | 1 | 1 | # | 20 | Small | paralle | 47 |
| # | time | 5 | 0 | 0 | 1 | 1 | # | 20 | Small | paralle | 47 |
| # | interaction | 4 | 0 | 0 | 1 | 1 | # | 20 | Small | paralle | 47 |
| # | disease | 7 | 3 | 0 | 0.7 | 1 | # | 20 | Small | ancom | 47 |
| # | sex | 4 | 0 | 0 | 1 | 1 | # | 20 | Small | ancom | 47 |
| # | time | 5 | 4 | 0 | 0.555555556 | 1 | # | 20 | Small | ancom | 47 |
| # | interaction | 0 | 0 | 4 | 0 | 0 | # | 20 | Small | ancom | 47 |
| # | disease | 7 | 5 | 0 | 0.583333333 | 1 | # | 20 | Small | aldex2 | 47 |
| # | sex | 4 | 0 | 0 | 1 | 1 | # | 20 | Small | aldex2 | 47 |
| # | time | 5 | 4 | 0 | 0.555555556 | 1 | # | 20 | Small | aldex2 | 47 |
| # | interaction | 0 | 0 | 4 | 0 | 0 | # | 20 | Small | aldex2 | 47 |
| # | disease | 7 | 12 | 0 | 0.368421053 | 1 | # | 20 | Baseline | paralle | 47 |
| # | sex | 4 | 6 | 0 | 0.4 | 1 | # | 20 | Baseline | paralle | 47 |
| # | time | 5 | 4 | 0 | 0.555555556 | 1 | # | 20 | Baseline | paralle | 47 |
| # | interaction | 4 | 3 | 0 | 0.571428571 | 1 | # | 20 | Baseline | paralle | 47 |
| # | disease | 7 | 3 | 0 | 0.7 | 1 | # | 20 | Baseline | ancom | 47 |
| # | sex | 4 | 0 | 0 | 1 | 1 | # | 20 | Baseline | ancom | 47 |
| # | time | 5 | 4 | 0 | 0.555555556 | 1 | # | 20 | Baseline | ancom | 47 |
| # | interaction | 0 | 0 | 4 | 0 | 0 | # | 20 | Baseline | ancom | 47 |
| # | disease | 7 | 52 | 0 | 0.118644068 | 1 | # | 20 | Baseline | aldex2 | 47 |
| # | sex | 4 | 16 | 0 | 0.2 | 1 | # | 20 | Baseline | aldex2 | 47 |
| # | time | 5 | 31 | 0 | 0.138888889 | 1 | # | 20 | Baseline | aldex2 | 47 |
| # | interaction | 0 | 0 | 4 | 0 | 0 | # | 20 | Baseline | aldex2 | 47 |
| # | disease | 7 | 1 | 0 | 0.875 | 1 | # | 20 | Small | paralle | 47 |
| # | sex | 4 | 0 | 0 | 1 | 1 | # | 20 | Small | paralle | 47 |
| # | time | 5 | 0 | 0 | 1 | 1 | # | 20 | Small | paralle | 47 |
| # | interaction | 4 | 0 | 0 | 1 | 1 | # | 20 | Small | paralle | 47 |
| # | disease | 7 | 3 | 0 | 0.7 | 1 | # | 20 | Small | ancom | 47 |
| # | sex | 4 | 0 | 0 | 1 | 1 | # | 20 | Small | ancom | 47 |
| # | time | 5 | 4 | 0 | 0.555555556 | 1 | # | 20 | Small | ancom | 47 |
| # | interaction | 0 | 0 | 4 | 0 | 0 | # | 20 | Small | ancom | 47 |
| # | disease | 7 | 8 | 0 | 0.466666667 | 1 | # | 20 | Small | aldex2 | 47 |
| # | sex | 4 | 0 | 0 | 1 | 1 | # | 20 | Small | aldex2 | 47 |
| # | time | 5 | 6 | 0 | 0.454545455 | 1 | # | 20 | Small | aldex2 | 47 |
| # | interaction | 0 | 0 | 4 | 0 | 0 | # | 20 | Small | aldex2 | 47 |
| # | disease | 7 | 8 | 0 | 0.466666667 | 1 | # | 20 | Baseline | paralle | 47 |
| # | sex | 4 | 8 | 0 | 0.333333333 | 1 | # | 20 | Baseline | paralle | 47 |
| # | time | 5 | 5 | 0 | 0.5 | 1 | # | 20 | Baseline | paralle | 47 |
| # | interaction | 4 | 2 | 0 | 0.666666667 | 1 | # | 20 | Baseline | paralle | 47 |
| # | disease | 7 | 4 | 0 | 0.636363636 | 1 | # | 20 | Baseline | ancom | 47 |
| # | sex | 4 | 0 | 0 | 1 | 1 | # | 20 | Baseline | ancom | 47 |
| # | time | 5 | 4 | 0 | 0.555555556 | 1 | # | 20 | Baseline | ancom | 47 |
| # | interaction | 0 | 0 | 4 | 0 | 0 | # | 20 | Baseline | ancom | 47 |
| # | disease | 7 | 58 | 0 | 0.093333333 | 1 | # | 20 | Baseline | aldex2 | 47 |
| # | sex | 4 | 25 | 0 | 0.137931034 | 1 | # | 20 | Baseline | aldex2 | 47 |
| # | time | 5 | 36 | 0 | 0.12195122 | 1 | # | 20 | Baseline | aldex2 | 47 |
| # | interaction | 0 | 0 | 4 | 0 | 0 | # | 20 | Baseline | aldex2 | 47 |
| # | disease | 7 | 1 | 0 | 0.875 | 1 | # | 20 | Small | paralle | 47 |
| # | sex | 4 | 0 | 0 | 1 | 1 | # | 20 | Small | paralle | 47 |
| # | time | 5 | 0 | 0 | 1 | 1 | # | 20 | Small | paralle | 47 |
| # | interaction | 4 | 0 | 0 | 1 | 1 | # | 20 | Small | paralle | 47 |
| # | disease | 7 | 4 | 0 | 0.636363636 | 1 | # | 20 | Small | ancom | 47 |
| # | sex | 4 | 0 | 0 | 1 | 1 | # | 20 | Small | ancom | 47 |
| # | time | 5 | 4 | 0 | 0.555555556 | 1 | # | 20 | Small | ancom | 47 |
| # | interaction | 0 | 0 | 4 | 0 | 0 | # | 20 | Small | ancom | 47 |
| # | disease | 7 | 9 | 0 | 0.4375 | 1 | # | 20 | Small | aldex2 | 47 |
| # | sex | 4 | 0 | 0 | 1 | 1 | # | 20 | Small | aldex2 | 47 |
| # | time | 5 | 4 | 0 | 0.555555556 | 1 | # | 20 | Small | aldex2 | 47 |
| # | interaction | 0 | 0 | 4 | 0 | 0 | # | 20 | Small | aldex2 | 47 |
| # | disease | 4 | 5 | 0 | 0.444444444 | 1 | # | 10 | Baseline | paralle | 48 |
| # | sex | 2 | 10 | 0 | 0.166666667 | 1 | # | 10 | Baseline | paralle | 48 |
| # | time | 2 | 8 | 0 | 0.2 | 1 | # | 10 | Baseline | paralle | 48 |
| # | interaction | 2 | 3 | 0 | 0.4 | 1 | # | 10 | Baseline | paralle | 48 |
| # | disease | 4 | 0 | 0 | 1 | 1 | # | 10 | Baseline | ancom | 48 |
| # | sex | 2 | 0 | 0 | 1 | 1 | # | 10 | Baseline | ancom | 48 |
| # | time | 2 | 0 | 0 | 1 | 1 | # | 10 | Baseline | ancom | 48 |
| # | interaction | 0 | 0 | 2 | 0 | 0 | # | 10 | Baseline | ancom | 48 |
| # | disease | 4 | 9 | 0 | 0.307692308 | 1 | # | 10 | Baseline | aldex2 | 48 |
| # | sex | 2 | 5 | 0 | 0.285714286 | 1 | # | 10 | Baseline | aldex2 | 48 |
| # | time | 2 | 9 | 0 | 0.181818182 | 1 | # | 10 | Baseline | aldex2 | 48 |
| # | interaction | 0 | 0 | 2 | 0 | 0 | # | 10 | Baseline | aldex2 | 48 |
| # | disease | 4 | 1 | 0 | 0.8 | 1 | # | 10 | Small | paralle | 48 |
| # | sex | 2 | 0 | 0 | 1 | 1 | # | 10 | Small | paralle | 48 |
| # | time | 2 | 0 | 0 | 1 | 1 | # | 10 | Small | paralle | 48 |
| # | interaction | 0 | 0 | 2 | 0 | 0 | # | 10 | Small | paralle | 48 |
| # | disease | 4 | 0 | 0 | 1 | 1 | # | 10 | Small | ancom | 48 |
| # | sex | 2 | 0 | 0 | 1 | 1 | # | 10 | Small | ancom | 48 |
| # | time | 2 | 0 | 0 | 1 | 1 | # | 10 | Small | ancom | 48 |
| # | interaction | 0 | 0 | 2 | 0 | 0 | # | 10 | Small | ancom | 48 |
| # | disease | 4 | 1 | 0 | 0.8 | 1 | # | 10 | Small | aldex2 | 48 |
| # | sex | 2 | 0 | 0 | 1 | 1 | # | 10 | Small | aldex2 | 48 |
| # | time | 2 | 1 | 0 | 0.666666667 | 1 | # | 10 | Small | aldex2 | 48 |
| # | interaction | 0 | 0 | 2 | 0 | 0 | # | 10 | Small | aldex2 | 48 |
| # | disease | 4 | 5 | 0 | 0.444444444 | 1 | # | 10 | Baseline | paralle | 48 |
| # | sex | 2 | 10 | 0 | 0.166666667 | 1 | # | 10 | Baseline | paralle | 48 |
| # | time | 2 | 5 | 0 | 0.285714286 | 1 | # | 10 | Baseline | paralle | 48 |
| # | interaction | 2 | 5 | 0 | 0.285714286 | 1 | # | 10 | Baseline | paralle | 48 |
| # | disease | 4 | 2 | 0 | 0.666666667 | 1 | # | 10 | Baseline | ancom | 48 |
| # | sex | 2 | 0 | 0 | 1 | 1 | # | 10 | Baseline | ancom | 48 |
| # | time | 2 | 1 | 0 | 0.666666667 | 1 | # | 10 | Baseline | ancom | 48 |
| # | interaction | 0 | 0 | 2 | 0 | 0 | # | 10 | Baseline | ancom | 48 |
| # | disease | 4 | 16 | 0 | 0.2 | 1 | # | 10 | Baseline | aldex2 | 48 |
| # | sex | 2 | 7 | 0 | 0.222222222 | 1 | # | 10 | Baseline | aldex2 | 48 |
| # | time | 2 | 11 | 0 | 0.153846154 | 1 | # | 10 | Baseline | aldex2 | 48 |
| # | interaction | 0 | 0 | 2 | 0 | 0 | # | 10 | Baseline | aldex2 | 48 |
| # | disease | 4 | 0 | 0 | 1 | 1 | # | 10 | Small | paralle | 48 |
| # | sex | 2 | 0 | 0 | 1 | 1 | # | 10 | Small | paralle | 48 |
| # | time | 2 | 0 | 0 | 1 | 1 | # | 10 | Small | paralle | 48 |
| # | interaction | 2 | 0 | 0 | 1 | 1 | # | 10 | Small | paralle | 48 |
| # | disease | 4 | 2 | 0 | 0.666666667 | 1 | # | 10 | Small | ancom | 48 |
| # | sex | 2 | 0 | 0 | 1 | 1 | # | 10 | Small | ancom | 48 |
| # | time | 2 | 1 | 0 | 0.666666667 | 1 | # | 10 | Small | ancom | 48 |
| # | interaction | 0 | 0 | 2 | 0 | 0 | # | 10 | Small | ancom | 48 |
| # | disease | 4 | 1 | 0 | 0.8 | 1 | # | 10 | Small | aldex2 | 48 |
| # | sex | 2 | 0 | 0 | 1 | 1 | # | 10 | Small | aldex2 | 48 |
| # | time | 2 | 1 | 0 | 0.666666667 | 1 | # | 10 | Small | aldex2 | 48 |
| # | interaction | 0 | 0 | 2 | 0 | 0 | # | 10 | Small | aldex2 | 48 |
| # | disease | 4 | 5 | 0 | 0.444444444 | 1 | # | 10 | Baseline | paralle | 48 |
| # | sex | 2 | 5 | 0 | 0.285714286 | 1 | # | 10 | Baseline | paralle | 48 |
| # | time | 2 | 4 | 0 | 0.333333333 | 1 | # | 10 | Baseline | paralle | 48 |
| # | interaction | 2 | 3 | 0 | 0.4 | 1 | # | 10 | Baseline | paralle | 48 |
| # | disease | 4 | 2 | 0 | 0.666666667 | 1 | # | 10 | Baseline | ancom | 48 |
| # | sex | 2 | 0 | 0 | 1 | 1 | # | 10 | Baseline | ancom | 48 |
| # | time | 2 | 2 | 0 | 0.5 | 1 | # | 10 | Baseline | ancom | 48 |
| # | interaction | 0 | 0 | 2 | 0 | 0 | # | 10 | Baseline | ancom | 48 |
| # | disease | 4 | 19 | 0 | 0.173913043 | 1 | # | 10 | Baseline | aldex2 | 48 |
| # | sex | 2 | 7 | 0 | 0.222222222 | 1 | # | 10 | Baseline | aldex2 | 48 |
| # | time | 2 | 10 | 0 | 0.166666667 | 1 | # | 10 | Baseline | aldex2 | 48 |
| # | interaction | 0 | 0 | 2 | 0 | 0 | # | 10 | Baseline | aldex2 | 48 |
| # | disease | 4 | 0 | 0 | 1 | 1 | # | 10 | Small | paralle | 48 |
| # | sex | 2 | 0 | 0 | 1 | 1 | # | 10 | Small | paralle | 48 |
| # | time | 2 | 0 | 0 | 1 | 1 | # | 10 | Small | paralle | 48 |
| # | interaction | 2 | 0 | 0 | 1 | 1 | # | 10 | Small | paralle | 48 |
| # | disease | 4 | 2 | 0 | 0.666666667 | 1 | # | 10 | Small | ancom | 48 |
| # | sex | 2 | 0 | 0 | 1 | 1 | # | 10 | Small | ancom | 48 |
| # | time | 2 | 2 | 0 | 0.5 | 1 | # | 10 | Small | ancom | 48 |
| # | interaction | 0 | 0 | 2 | 0 | 0 | # | 10 | Small | ancom | 48 |
| # | disease | 4 | 2 | 0 | 0.666666667 | 1 | # | 10 | Small | aldex2 | 48 |
| # | sex | 2 | 0 | 0 | 1 | 1 | # | 10 | Small | aldex2 | 48 |
| # | time | 2 | 1 | 0 | 0.666666667 | 1 | # | 10 | Small | aldex2 | 48 |
| # | interaction | 0 | 0 | 2 | 0 | 0 | # | 10 | Small | aldex2 | 48 |

| | | | | | | | | | | | |
|---|---|---|---|---|---|---|---|---|---|---|---|
| # | disease | 4 | 5 | 0 | 0.444444444 | 1 | # | 10 | Baseline | paralle | 48 |
| # | sex | 2 | 3 | 0 | 0.4 | 1 | # | 10 | Baseline | paralle | 48 |
| # | time | 2 | 2 | 0 | 0.5 | 1 | # | 10 | Baseline | paralle | 48 |
| # | interaction | 2 | 2 | 0 | 0.5 | 1 | # | 10 | Baseline | paralle | 48 |
| # | disease | 4 | 2 | 0 | 0.666666667 | 1 | # | 10 | Baseline | ancom | 48 |
| # | sex | 2 | 0 | 0 | 1 | 1 | # | 10 | Baseline | ancom | 48 |
| # | time | 2 | 2 | 0 | 0.5 | 1 | # | 10 | Baseline | ancom | 48 |
| # | interaction | 0 | 0 | 2 | 0 | 0 | # | 10 | Baseline | ancom | 48 |
| # | disease | 4 | 23 | 0 | 0.148148148 | 1 | # | 10 | Baseline | aldex2 | 48 |
| # | sex | 2 | 6 | 0 | 0.25 | 1 | # | 10 | Baseline | aldex2 | 48 |
| # | time | 2 | 13 | 0 | 0.133333333 | 1 | # | 10 | Baseline | aldex2 | 48 |
| # | interaction | 0 | 0 | 2 | 0 | 0 | # | 10 | Baseline | aldex2 | 48 |
| # | disease | 4 | 0 | 0 | 1 | 1 | # | 10 | Small | paralle | 48 |
| # | sex | 2 | 0 | 0 | 1 | 1 | # | 10 | Small | paralle | 48 |
| # | time | 2 | 0 | 0 | 1 | 1 | # | 10 | Small | paralle | 48 |
| # | interaction | 2 | 0 | 0 | 1 | 1 | # | 10 | Small | paralle | 48 |
| # | disease | 4 | 2 | 0 | 0.666666667 | 1 | # | 10 | Small | ancom | 48 |
| # | sex | 2 | 0 | 0 | 1 | 1 | # | 10 | Small | ancom | 48 |
| # | time | 2 | 2 | 0 | 0.5 | 1 | # | 10 | Small | ancom | 48 |
| # | interaction | 0 | 0 | 2 | 0 | 0 | # | 10 | Small | ancom | 48 |
| # | disease | 4 | 2 | 0 | 0.666666667 | 1 | # | 10 | Small | aldex2 | 48 |
| # | sex | 2 | 0 | 0 | 1 | 1 | # | 10 | Small | aldex2 | 48 |
| # | time | 2 | 2 | 0 | 0.5 | 1 | # | 10 | Small | aldex2 | 48 |
| # | interaction | 0 | 0 | 2 | 0 | 0 | # | 10 | Small | aldex2 | 48 |
| # | disease | 7 | 11 | 0 | 0.388888889 | 1 | # | 20 | Baseline | paralle | 48 |
| # | sex | 4 | 8 | 0 | 0.333333333 | 1 | # | 20 | Baseline | paralle | 48 |
| # | time | 5 | 11 | 0 | 0.3125 | 1 | # | 20 | Baseline | paralle | 48 |
| # | interaction | 4 | 9 | 0 | 0.307692308 | 1 | # | 20 | Baseline | paralle | 48 |
| # | disease | 7 | 3 | 0 | 0.7 | 1 | # | 20 | Baseline | ancom | 48 |
| # | sex | 4 | 0 | 0 | 1 | 1 | # | 20 | Baseline | ancom | 48 |
| # | time | 5 | 0 | 0 | 1 | 1 | # | 20 | Baseline | ancom | 48 |
| # | interaction | 0 | 0 | 4 | 0 | 0 | # | 20 | Baseline | ancom | 48 |
| # | disease | 7 | 24 | 0 | 0.225806452 | 1 | # | 20 | Baseline | aldex2 | 48 |
| # | sex | 4 | 4 | 0 | 0.5 | 1 | # | 20 | Baseline | aldex2 | 48 |
| # | time | 5 | 9 | 0 | 0.357142857 | 1 | # | 20 | Baseline | aldex2 | 48 |
| # | interaction | 0 | 0 | 4 | 0 | 0 | # | 20 | Baseline | aldex2 | 48 |
| # | disease | 7 | 0 | 0 | 1 | 1 | # | 20 | Small | paralle | 48 |
| # | sex | 4 | 0 | 0 | 1 | 1 | # | 20 | Small | paralle | 48 |
| # | time | 5 | 0 | 0 | 1 | 1 | # | 20 | Small | paralle | 48 |
| # | interaction | 1 | 0 | 3 | 1 | 0.25 | # | 20 | Small | paralle | 48 |
| # | disease | 7 | 2 | 0 | 0.777777778 | 1 | # | 20 | Small | ancom | 48 |
| # | sex | 4 | 0 | 0 | 1 | 1 | # | 20 | Small | ancom | 48 |
| # | time | 5 | 0 | 0 | 1 | 1 | # | 20 | Small | ancom | 48 |
| # | interaction | 0 | 0 | 4 | 0 | 0 | # | 20 | Small | ancom | 48 |
| # | disease | 7 | 2 | 0 | 0.777777778 | 1 | # | 20 | Small | aldex2 | 48 |
| # | sex | 4 | 1 | 0 | 0.8 | 1 | # | 20 | Small | aldex2 | 48 |
| # | time | 5 | 0 | 0 | 1 | 1 | # | 20 | Small | aldex2 | 48 |
| # | interaction | 0 | 0 | 4 | 0 | 0 | # | 20 | Small | aldex2 | 48 |
| # | disease | 7 | 9 | 0 | 0.4375 | 1 | # | 20 | Baseline | paralle | 48 |
| # | sex | 4 | 2 | 0 | 0.666666667 | 1 | # | 20 | Baseline | paralle | 48 |
| # | time | 5 | 12 | 0 | 0.294117647 | 1 | # | 20 | Baseline | paralle | 48 |
| # | interaction | 4 | 5 | 0 | 0.444444444 | 1 | # | 20 | Baseline | paralle | 48 |
| # | disease | 7 | 4 | 0 | 0.636363636 | 1 | # | 20 | Baseline | ancom | 48 |
| # | sex | 4 | 0 | 0 | 1 | 1 | # | 20 | Baseline | ancom | 48 |
| # | time | 5 | 2 | 0 | 0.714285714 | 1 | # | 20 | Baseline | ancom | 48 |
| # | interaction | 0 | 0 | 4 | 0 | 0 | # | 20 | Baseline | ancom | 48 |
| # | disease | 7 | 62 | 0 | 0.142857143 | 1 | # | 20 | Baseline | aldex2 | 48 |
| # | sex | 4 | 5 | 0 | 0.444444444 | 1 | # | 20 | Baseline | aldex2 | 48 |
| # | time | 5 | 24 | 0 | 0.172413793 | 1 | # | 20 | Baseline | aldex2 | 48 |
| # | interaction | 0 | 0 | 4 | 0 | 0 | # | 20 | Baseline | aldex2 | 48 |
| # | disease | 7 | 0 | 0 | 1 | 1 | # | 20 | Small | paralle | 48 |
| # | sex | 4 | 0 | 0 | 1 | 1 | # | 20 | Small | paralle | 48 |
| # | time | 5 | 0 | 0 | 1 | 1 | # | 20 | Small | paralle | 48 |
| # | interaction | 4 | 1 | 0 | 0.8 | 1 | # | 20 | Small | paralle | 48 |
| # | disease | 7 | 4 | 0 | 0.636363636 | 1 | # | 20 | Small | ancom | 48 |
| # | sex | 4 | 0 | 0 | 1 | 1 | # | 20 | Small | ancom | 48 |
| # | time | 5 | 2 | 0 | 0.714285714 | 1 | # | 20 | Small | ancom | 48 |
| # | interaction | 0 | 0 | 4 | 0 | 0 | # | 20 | Small | ancom | 48 |
| # | disease | 7 | 4 | 0 | 0.636363636 | 1 | # | 20 | Small | aldex2 | 48 |
| # | sex | 4 | 0 | 0 | 1 | 1 | # | 20 | Small | aldex2 | 48 |
| # | time | 5 | 2 | 0 | 0.714285714 | 1 | # | 20 | Small | aldex2 | 48 |
| # | interaction | 0 | 0 | 4 | 0 | 0 | # | 20 | Small | aldex2 | 48 |
| # | disease | 7 | 12 | 0 | 0.368421053 | 1 | # | 20 | Baseline | paralle | 48 |
| # | sex | 4 | 5 | 0 | 0.444444444 | 1 | # | 20 | Baseline | paralle | 48 |
| # | time | 5 | 9 | 0 | 0.357142857 | 1 | # | 20 | Baseline | paralle | 48 |
| # | interaction | 4 | 2 | 0 | 0.666666667 | 1 | # | 20 | Baseline | paralle | 48 |
| # | disease | 7 | 4 | 0 | 0.636363636 | 1 | # | 20 | Baseline | ancom | 48 |
| # | sex | 4 | 0 | 0 | 1 | 1 | # | 20 | Baseline | ancom | 48 |
| # | time | 5 | 3 | 0 | 0.625 | 1 | # | 20 | Baseline | ancom | 48 |
| # | interaction | 0 | 0 | 4 | 0 | 0 | # | 20 | Baseline | ancom | 48 |
| # | disease | 7 | 50 | 0 | 0.122807018 | 1 | # | 20 | Baseline | aldex2 | 48 |
| # | sex | 4 | 13 | 0 | 0.235294118 | 1 | # | 20 | Baseline | aldex2 | 48 |
| # | time | 5 | 30 | 0 | 0.142857143 | 1 | # | 20 | Baseline | aldex2 | 48 |
| # | interaction | 0 | 0 | 4 | 0 | 0 | # | 20 | Baseline | aldex2 | 48 |
| # | disease | 7 | 0 | 0 | 1 | 1 | # | 20 | Small | paralle | 48 |
| # | sex | 4 | 0 | 0 | 1 | 1 | # | 20 | Small | paralle | 48 |
| # | time | 5 | 0 | 0 | 1 | 1 | # | 20 | Small | paralle | 48 |
| # | interaction | 4 | 0 | 0 | 1 | 1 | # | 20 | Small | paralle | 48 |
| # | disease | 7 | 4 | 0 | 0.636363636 | 1 | # | 20 | Small | ancom | 48 |
| # | sex | 4 | 0 | 0 | 1 | 1 | # | 20 | Small | ancom | 48 |
| # | time | 5 | 4 | 0 | 0.555555556 | 1 | # | 20 | Small | ancom | 48 |
| # | interaction | 0 | 0 | 4 | 0 | 0 | # | 20 | Small | ancom | 48 |
| # | disease | 7 | 8 | 0 | 0.466666667 | 1 | # | 20 | Small | aldex2 | 48 |
| # | sex | 4 | 0 | 0 | 1 | 1 | # | 20 | Small | aldex2 | 48 |
| # | time | 5 | 4 | 0 | 0.555555556 | 1 | # | 20 | Small | aldex2 | 48 |
| # | interaction | 0 | 0 | 4 | 0 | 0 | # | 20 | Small | aldex2 | 48 |
| # | disease | 7 | 8 | 0 | 0.466666667 | 1 | # | 20 | Baseline | paralle | 48 |
| # | sex | 4 | 6 | 0 | 0.4 | 1 | # | 20 | Baseline | paralle | 48 |
| # | time | 5 | 10 | 0 | 0.333333333 | 1 | # | 20 | Baseline | paralle | 48 |
| # | interaction | 4 | 1 | 0 | 0.8 | 1 | # | 20 | Baseline | paralle | 48 |
| # | disease | 7 | 4 | 0 | 0.636363636 | 1 | # | 20 | Baseline | ancom | 48 |
| # | sex | 4 | 0 | 0 | 1 | 1 | # | 20 | Baseline | ancom | 48 |
| # | time | 5 | 4 | 0 | 0.555555556 | 1 | # | 20 | Baseline | ancom | 48 |
| # | interaction | 0 | 0 | 4 | 0 | 0 | # | 20 | Baseline | ancom | 48 |
| # | disease | 7 | 59 | 0 | 0.106060606 | 1 | # | 20 | Baseline | aldex2 | 48 |
| # | sex | 4 | 18 | 0 | 0.181818182 | 1 | # | 20 | Baseline | aldex2 | 48 |
| # | time | 5 | 66 | 0 | 0.098039216 | 1 | # | 20 | Baseline | aldex2 | 48 |
| # | interaction | 0 | 0 | 4 | 0 | 0 | # | 20 | Baseline | aldex2 | 48 |
| # | disease | 7 | 0 | 0 | 1 | 1 | # | 20 | Small | paralle | 48 |
| # | sex | 4 | 0 | 0 | 1 | 1 | # | 20 | Small | paralle | 48 |
| # | time | 5 | 0 | 0 | 1 | 1 | # | 20 | Small | paralle | 48 |
| # | interaction | 4 | 0 | 0 | 1 | 1 | # | 20 | Small | paralle | 48 |
| # | disease | 7 | 4 | 0 | 0.636363636 | 1 | # | 20 | Small | ancom | 48 |
| # | sex | 4 | 0 | 0 | 1 | 1 | # | 20 | Small | ancom | 48 |
| # | time | 5 | 4 | 0 | 0.555555556 | 1 | # | 20 | Small | ancom | 48 |
| # | interaction | 0 | 0 | 4 | 0 | 0 | # | 20 | Small | ancom | 48 |
| # | disease | 7 | 14 | 0 | 0.333333333 | 1 | # | 20 | Small | aldex2 | 48 |
| # | sex | 4 | 0 | 0 | 1 | 1 | # | 20 | Small | aldex2 | 48 |
| # | time | 5 | 7 | 0 | 0.416666667 | 1 | # | 20 | Small | aldex2 | 48 |
| # | interaction | 0 | 0 | 4 | 0 | 0 | # | 20 | Small | aldex2 | 48 |
| # | disease | 4 | 8 | 0 | 0.333333333 | 1 | # | 10 | Baseline | paralle | 49 |
| # | sex | 2 | 7 | 0 | 0.222222222 | 1 | # | 10 | Baseline | paralle | 49 |
| # | time | 2 | 5 | 0 | 0.285714286 | 1 | # | 10 | Baseline | paralle | 49 |
| # | interaction | 2 | 5 | 0 | 0.285714286 | 1 | # | 10 | Baseline | paralle | 49 |
| # | disease | 4 | 2 | 0 | 0.666666667 | 1 | # | 10 | Baseline | ancom | 49 |
| # | sex | 2 | 0 | 0 | 1 | 1 | # | 10 | Baseline | ancom | 49 |
| # | time | 2 | 0 | 0 | 1 | 1 | # | 10 | Baseline | ancom | 49 |
| # | interaction | 0 | 0 | 2 | 0 | 0 | # | 10 | Baseline | ancom | 49 |
| # | disease | 4 | 9 | 0 | 0.307692308 | 1 | # | 10 | Baseline | aldex2 | 49 |
| # | sex | 2 | 3 | 0 | 0.4 | 1 | # | 10 | Baseline | aldex2 | 49 |
| # | time | 2 | 5 | 0 | 0.285714286 | 1 | # | 10 | Baseline | aldex2 | 49 |
| # | interaction | 0 | 0 | 2 | 0 | 0 | # | 10 | Baseline | aldex2 | 49 |
| # | disease | 4 | 0 | 0 | 1 | 1 | # | 10 | Small | paralle | 49 |
| # | sex | 2 | 0 | 0 | 1 | 1 | # | 10 | Small | paralle | 49 |
| # | time | 2 | 0 | 0 | 1 | 1 | # | 10 | Small | paralle | 49 |
| # | interaction | 1 | 0 | 1 | 1 | 0.5 | # | 10 | Small | paralle | 49 |
| # | disease | 4 | 2 | 0 | 0.666666667 | 1 | # | 10 | Small | ancom | 49 |
| # | sex | 2 | 0 | 0 | 1 | 1 | # | 10 | Small | ancom | 49 |
| # | time | 2 | 0 | 0 | 1 | 1 | # | 10 | Small | ancom | 49 |
| # | interaction | 0 | 0 | 2 | 0 | 0 | # | 10 | Small | ancom | 49 |
| # | disease | 4 | 2 | 0 | 0.666666667 | 1 | # | 10 | Small | aldex2 | 49 |
| # | sex | 2 | 0 | 0 | 1 | 1 | # | 10 | Small | aldex2 | 49 |
| # | time | 2 | 1 | 0 | 0.666666667 | 1 | # | 10 | Small | aldex2 | 49 |
| # | interaction | 0 | 0 | 2 | 0 | 0 | # | 10 | Small | aldex2 | 49 |
| # | disease | 4 | 7 | 0 | 0.363636364 | 1 | # | 10 | Baseline | paralle | 49 |
| # | sex | 2 | 4 | 0 | 0.333333333 | 1 | # | 10 | Baseline | paralle | 49 |
| # | time | 2 | 7 | 0 | 0.222222222 | 1 | # | 10 | Baseline | paralle | 49 |
| # | interaction | 2 | 3 | 0 | 0.4 | 1 | # | 10 | Baseline | paralle | 49 |
| # | disease | 4 | 2 | 0 | 0.666666667 | 1 | # | 10 | Baseline | ancom | 49 |
| # | sex | 2 | 0 | 0 | 1 | 1 | # | 10 | Baseline | ancom | 49 |
| # | time | 2 | 2 | 0 | 0.5 | 1 | # | 10 | Baseline | ancom | 49 |
| # | interaction | 0 | 0 | 2 | 0 | 0 | # | 10 | Baseline | ancom | 49 |
| # | disease | 4 | 15 | 0 | 0.210526316 | 1 | # | 10 | Baseline | aldex2 | 49 |
| # | sex | 2 | 6 | 0 | 0.25 | 1 | # | 10 | Baseline | aldex2 | 49 |
| # | time | 2 | 7 | 0 | 0.222222222 | 1 | # | 10 | Baseline | aldex2 | 49 |
| # | interaction | 0 | 0 | 2 | 0 | 0 | # | 10 | Baseline | aldex2 | 49 |
| # | disease | 4 | 0 | 0 | 1 | 1 | # | 10 | Small | paralle | 49 |
| # | sex | 2 | 0 | 0 | 1 | 1 | # | 10 | Small | paralle | 49 |
| # | time | 2 | 0 | 0 | 1 | 1 | # | 10 | Small | paralle | 49 |
| # | interaction | 2 | 0 | 0 | 1 | 1 | # | 10 | Small | paralle | 49 |
| # | disease | 4 | 2 | 0 | 0.666666667 | 1 | # | 10 | Small | ancom | 49 |
| # | sex | 2 | 0 | 0 | 1 | 1 | # | 10 | Small | ancom | 49 |
| # | time | 2 | 2 | 0 | 0.5 | 1 | # | 10 | Small | ancom | 49 |
| # | interaction | 0 | 0 | 2 | 0 | 0 | # | 10 | Small | ancom | 49 |
| # | disease | 4 | 4 | 0 | 0.5 | 1 | # | 10 | Small | aldex2 | 49 |
| # | sex | 2 | 0 | 0 | 1 | 1 | # | 10 | Small | aldex2 | 49 |
| # | time | 2 | 2 | 0 | 0.5 | 1 | # | 10 | Small | aldex2 | 49 |
| # | interaction | 0 | 0 | 2 | 0 | 0 | # | 10 | Small | aldex2 | 49 |
| # | disease | 4 | 7 | 0 | 0.363636364 | 1 | # | 10 | Baseline | paralle | 49 |
| # | sex | 2 | 6 | 0 | 0.25 | 1 | # | 10 | Baseline | paralle | 49 |
| # | time | 2 | 7 | 0 | 0.222222222 | 1 | # | 10 | Baseline | paralle | 49 |
| # | interaction | 2 | 6 | 0 | 0.25 | 1 | # | 10 | Baseline | paralle | 49 |
| # | disease | 4 | 2 | 0 | 0.666666667 | 1 | # | 10 | Baseline | ancom | 49 |
| # | sex | 2 | 0 | 0 | 1 | 1 | # | 10 | Baseline | ancom | 49 |
| # | time | 2 | 2 | 0 | 0.5 | 1 | # | 10 | Baseline | ancom | 49 |
| # | interaction | 0 | 0 | 2 | 0 | 0 | # | 10 | Baseline | ancom | 49 |
| # | disease | 4 | 19 | 0 | 0.173913043 | 1 | # | 10 | Baseline | aldex2 | 49 |
| # | sex | 2 | 9 | 0 | 0.181818182 | 1 | # | 10 | Baseline | aldex2 | 49 |
| # | time | 2 | 12 | 0 | 0.142857143 | 1 | # | 10 | Baseline | aldex2 | 49 |
| # | interaction | 0 | 0 | 2 | 0 | 0 | # | 10 | Baseline | aldex2 | 49 |
| # | disease | 4 | 0 | 0 | 1 | 1 | # | 10 | Small | paralle | 49 |
| # | sex | 2 | 0 | 0 | 1 | 1 | # | 10 | Small | paralle | 49 |
| # | time | 2 | 0 | 0 | 1 | 1 | # | 10 | Small | paralle | 49 |
| # | interaction | 2 | 1 | 0 | 0.666666667 | 1 | # | 10 | Small | paralle | 49 |
| # | disease | 4 | 2 | 0 | 0.666666667 | 1 | # | 10 | Small | ancom | 49 |
| # | sex | 2 | 0 | 0 | 1 | 1 | # | 10 | Small | ancom | 49 |
| # | time | 2 | 2 | 0 | 0.5 | 1 | # | 10 | Small | ancom | 49 |
| # | interaction | 0 | 0 | 2 | 0 | 0 | # | 10 | Small | ancom | 49 |
| # | disease | 4 | 4 | 0 | 0.5 | 1 | # | 10 | Small | aldex2 | 49 |
| # | sex | 2 | 0 | 0 | 1 | 1 | # | 10 | Small | aldex2 | 49 |
| # | time | 2 | 2 | 0 | 0.5 | 1 | # | 10 | Small | aldex2 | 49 |
| # | interaction | 0 | 0 | 2 | 0 | 0 | # | 10 | Small | aldex2 | 49 |
| # | disease | 4 | 3 | 0 | 0.571428571 | 1 | # | 10 | Baseline | paralle | 49 |
| # | sex | 2 | 2 | 0 | 0.5 | 1 | # | 10 | Baseline | paralle | 49 |
| # | time | 2 | 6 | 0 | 0.25 | 1 | # | 10 | Baseline | paralle | 49 |
| # | interaction | 2 | 6 | 0 | 0.25 | 1 | # | 10 | Baseline | paralle | 49 |
| # | disease | 4 | 2 | 0 | 0.666666667 | 1 | # | 10 | Baseline | ancom | 49 |
| # | sex | 2 | 0 | 0 | 1 | 1 | # | 10 | Baseline | ancom | 49 |
| # | time | 2 | 2 | 0 | 0.5 | 1 | # | 10 | Baseline | ancom | 49 |
| # | interaction | 0 | 0 | 2 | 0 | 0 | # | 10 | Baseline | ancom | 49 |
| # | disease | 4 | 22 | 0 | 0.153846154 | 1 | # | 10 | Baseline | aldex2 | 49 |
| # | sex | 2 | 8 | 0 | 0.2 | 1 | # | 10 | Baseline | aldex2 | 49 |
| # | time | 2 | 14 | 0 | 0.125 | 1 | # | 10 | Baseline | aldex2 | 49 |
| # | interaction | 0 | 0 | 2 | 0 | 0 | # | 10 | Baseline | aldex2 | 49 |
| # | disease | 4 | 0 | 0 | 1 | 1 | # | 10 | Small | paralle | 49 |
| # | sex | 2 | 0 | 0 | 1 | 1 | # | 10 | Small | paralle | 49 |
| # | time | 2 | 0 | 0 | 1 | 1 | # | 10 | Small | paralle | 49 |
| # | interaction | 2 | 0 | 0 | 1 | 1 | # | 10 | Small | paralle | 49 |
| # | disease | 4 | 2 | 0 | 0.666666667 | 1 | # | 10 | Small | ancom | 49 |
| # | sex | 2 | 0 | 0 | 1 | 1 | # | 10 | Small | ancom | 49 |
| # | time | 2 | 2 | 0 | 0.5 | 1 | # | 10 | Small | ancom | 49 |
| # | interaction | 0 | 0 | 2 | 0 | 0 | # | 10 | Small | ancom | 49 |
| # | disease | 4 | 4 | 0 | 0.5 | 1 | # | 10 | Small | aldex2 | 49 |
| # | sex | 2 | 0 | 0 | 1 | 1 | # | 10 | Small | aldex2 | 49 |
| # | time | 2 | 2 | 0 | 0.5 | 1 | # | 10 | Small | aldex2 | 49 |
| # | interaction | 0 | 0 | 2 | 0 | 0 | # | 10 | Small | aldex2 | 49 |
| # | disease | 7 | 6 | 0 | 0.538461538 | 1 | # | 20 | Baseline | paralle | 49 |
| # | sex | 4 | 6 | 0 | 0.4 | 1 | # | 20 | Baseline | paralle | 49 |
| # | time | 5 | 8 | 0 | 0.384615385 | 1 | # | 20 | Baseline | paralle | 49 |
| # | interaction | 3 | 7 | 1 | 0.3 | 0.75 | # | 20 | Baseline | paralle | 49 |
| # | disease | 7 | 1 | 0 | 0.875 | 1 | # | 20 | Baseline | ancom | 49 |
| # | sex | 4 | 0 | 0 | 1 | 1 | # | 20 | Baseline | ancom | 49 |
| # | time | 5 | 0 | 0 | 1 | 1 | # | 20 | Baseline | ancom | 49 |
| # | interaction | 0 | 0 | 4 | 0 | 0 | # | 20 | Baseline | ancom | 49 |
| # | disease | 7 | 17 | 0 | 0.291666667 | 1 | # | 20 | Baseline | aldex2 | 49 |
| # | sex | 4 | 10 | 0 | 0.285714286 | 1 | # | 20 | Baseline | aldex2 | 49 |
| # | time | 5 | 16 | 0 | 0.238095238 | 1 | # | 20 | Baseline | aldex2 | 49 |
| # | interaction | 0 | 0 | 4 | 0 | 0 | # | 20 | Baseline | aldex2 | 49 |
| # | disease | 7 | 0 | 0 | 1 | 1 | # | 20 | Small | paralle | 49 |
| # | sex | 4 | 0 | 0 | 1 | 1 | # | 20 | Small | paralle | 49 |
| # | time | 5 | 0 | 0 | 1 | 1 | # | 20 | Small | paralle | 49 |
| # | interaction | 0 | 0 | 4 | 0 | 0 | # | 20 | Small | paralle | 49 |
| # | disease | 7 | 0 | 0 | 1 | 1 | # | 20 | Small | ancom | 49 |
| # | sex | 4 | 0 | 0 | 1 | 1 | # | 20 | Small | ancom | 49 |
| # | time | 5 | 0 | 0 | 1 | 1 | # | 20 | Small | ancom | 49 |
| # | interaction | 0 | 0 | 4 | 0 | 0 | # | 20 | Small | ancom | 49 |
| # | disease | 7 | 1 | 0 | 0.875 | 1 | # | 20 | Small | aldex2 | 49 |
| # | sex | 4 | 0 | 0 | 1 | 1 | # | 20 | Small | aldex2 | 49 |
| # | time | 5 | 0 | 0 | 1 | 1 | # | 20 | Small | aldex2 | 49 |
| # | interaction | 0 | 0 | 4 | 0 | 0 | # | 20 | Small | aldex2 | 49 |
| # | disease | 7 | 5 | 0 | 0.583333333 | 1 | # | 20 | Baseline | paralle | 49 |
| # | sex | 4 | 5 | 0 | 0.444444444 | 1 | # | 20 | Baseline | paralle | 49 |
| # | time | 5 | 5 | 0 | 0.5 | 1 | # | 20 | Baseline | paralle | 49 |
| # | interaction | 4 | 8 | 0 | 0.333333333 | 1 | # | 20 | Baseline | paralle | 49 |
| # | disease | 7 | 4 | 0 | 0.636363636 | 1 | # | 20 | Baseline | ancom | 49 |
| # | sex | 4 | 0 | 0 | 1 | 1 | # | 20 | Baseline | ancom | 49 |
| # | time | 5 | 2 | 0 | 0.714285714 | 1 | # | 20 | Baseline | ancom | 49 |
| # | interaction | 0 | 0 | 4 | 0 | 0 | # | 20 | Baseline | ancom | 49 |
| # | disease | 7 | 31 | 0 | 0.184210526 | 1 | # | 20 | Baseline | aldex2 | 49 |
| # | sex | 4 | 14 | 0 | 0.222222222 | 1 | # | 20 | Baseline | aldex2 | 49 |
| # | time | 5 | 24 | 0 | 0.172413793 | 1 | # | 20 | Baseline | aldex2 | 49 |
| # | interaction | 0 | 0 | 4 | 0 | 0 | # | 20 | Baseline | aldex2 | 49 |
| # | disease | 7 | 0 | 0 | 1 | 1 | # | 20 | Small | paralle | 49 |
| # | sex | 4 | 0 | 0 | 1 | 1 | # | 20 | Small | paralle | 49 |
| # | time | 5 | 0 | 0 | 1 | 1 | # | 20 | Small | paralle | 49 |
| # | interaction | 2 | 0 | 2 | 1 | 0.5 | # | 20 | Small | paralle | 49 |
| # | disease | 7 | 4 | 0 | 0.636363636 | 1 | # | 20 | Small | ancom | 49 |
| # | sex | 4 | 0 | 0 | 1 | 1 | # | 20 | Small | ancom | 49 |
| # | time | 5 | 2 | 0 | 0.714285714 | 1 | # | 20 | Small | ancom | 49 |
| # | interaction | 0 | 0 | 4 | 0 | 0 | # | 20 | Small | ancom | 49 |
| # | disease | 7 | 6 | 0 | 0.538461538 | 1 | # | 20 | Small | aldex2 | 49 |
| # | sex | 4 | 0 | 0 | 1 | 1 | # | 20 | Small | aldex2 | 49 |
| # | time | 5 | 2 | 0 | 0.714285714 | 1 | # | 20 | Small | aldex2 | 49 |
| # | interaction | 0 | 0 | 4 | 0 | 0 | # | 20 | Small | aldex2 | 49 |
| # | disease | 7 | 5 | 0 | 0.583333333 | 1 | # | 20 | Baseline | paralle | 49 |
| # | sex | 4 | 5 | 0 | 0.444444444 | 1 | # | 20 | Baseline | paralle | 49 |
| # | time | 5 | 9 | 0 | 0.357142857 | 1 | # | 20 | Baseline | paralle | 49 |
| # | interaction | 4 | 5 | 0 | 0.444444444 | 1 | # | 20 | Baseline | paralle | 49 |
| # | disease | 7 | 4 | 0 | 0.636363636 | 1 | # | 20 | Baseline | ancom | 49 |
| # | sex | 4 | 0 | 0 | 1 | 1 | # | 20 | Baseline | ancom | 49 |
| # | time | 5 | 4 | 0 | 0.555555556 | 1 | # | 20 | Baseline | ancom | 49 |
| # | interaction | 0 | 0 | 4 | 0 | 0 | # | 20 | Baseline | ancom | 49 |
| # | disease | 7 | 63 | 0 | 0.14 | 1 | # | 20 | Baseline | aldex2 | 49 |
| # | sex | 4 | 19 | 0 | 0.173913043 | 1 | # | 20 | Baseline | aldex2 | 49 |
| # | time | 5 | 36 | 0 | 0.12195122 | 1 | # | 20 | Baseline | aldex2 | 49 |
| # | interaction | 0 | 0 | 4 | 0 | 0 | # | 20 | Baseline | aldex2 | 49 |
| # | disease | 7 | 0 | 0 | 1 | 1 | # | 20 | Small | paralle | 49 |
| # | sex | 4 | 0 | 0 | 1 | 1 | # | 20 | Small | paralle | 49 |
| # | time | 5 | 0 | 0 | 1 | 1 | # | 20 | Small | paralle | 49 |
| # | interaction | 4 | 0 | 0 | 1 | 1 | # | 20 | Small | paralle | 49 |
| # | disease | 7 | 4 | 0 | 0.636363636 | 1 | # | 20 | Small | ancom | 49 |
| # | sex | 4 | 0 | 0 | 1 | 1 | # | 20 | Small | ancom | 49 |
| # | time | 5 | 4 | 0 | 0.555555556 | 1 | # | 20 | Small | ancom | 49 |
| # | interaction | 0 | 0 | 4 | 0 | 0 | # | 20 | Small | ancom | 49 |
| # | disease | 7 | 7 | 0 | 0.5 | 1 | # | 20 | Small | aldex2 | 49 |
| # | sex | 4 | 1 | 0 | 0.8 | 1 | # | 20 | Small | aldex2 | 49 |
| # | time | 5 | 5 | 0 | 0.5 | 1 | # | 20 | Small | aldex2 | 49 |
| # | interaction | 0 | 0 | 4 | 0 | 0 | # | 20 | Small | aldex2 | 49 |
| # | disease | 7 | 5 | 0 | 0.583333333 | 1 | # | 20 | Baseline | paralle | 49 |
| # | sex | 4 | 3 | 0 | 0.571428571 | 1 | # | 20 | Baseline | paralle | 49 |
| # | time | 5 | 7 | 0 | 0.416666667 | 1 | # | 20 | Baseline | paralle | 49 |
| # | interaction | 4 | 4 | 0 | 0.5 | 1 | # | 20 | Baseline | paralle | 49 |
| # | disease | 7 | 5 | 0 | 0.583333333 | 1 | # | 20 | Baseline | ancom | 49 |
| # | sex | 4 | 0 | 0 | 1 | 1 | # | 20 | Baseline | ancom | 49 |
| # | time | 5 | 4 | 0 | 0.555555556 | 1 | # | 20 | Baseline | ancom | 49 |
| # | interaction | 0 | 0 | 4 | 0 | 0 | # | 20 | Baseline | ancom | 49 |
| # | disease | 7 | 58 | 0 | 0.107692308 | 1 | # | 20 | Baseline | aldex2 | 49 |
| # | sex | 4 | 22 | 0 | 0.153846154 | 1 | # | 20 | Baseline | aldex2 | 49 |
| # | time | 5 | 66 | 0 | 0.098039216 | 1 | # | 20 | Baseline | aldex2 | 49 |
| # | interaction | 0 | 0 | 4 | 0 | 0 | # | 20 | Baseline | aldex2 | 49 |
| # | disease | 7 | 0 | 0 | 1 | 1 | # | 20 | Small | paralle | 49 |
| # | sex | 4 | 0 | 0 | 1 | 1 | # | 20 | Small | paralle | 49 |
| # | time | 5 | 0 | 0 | 1 | 1 | # | 20 | Small | paralle | 49 |
| # | interaction | 4 | 0 | 0 | 1 | 1 | # | 20 | Small | paralle | 49 |
| # | disease | 7 | 4 | 0 | 0.636363636 | 1 | # | 20 | Small | ancom | 49 |
| # | sex | 4 | 0 | 0 | 1 | 1 | # | 20 | Small | ancom | 49 |
| # | time | 5 | 4 | 0 | 0.555555556 | 1 | # | 20 | Small | ancom | 49 |
| # | interaction | 0 | 0 | 4 | 0 | 0 | # | 20 | Small | ancom | 49 |
| # | disease | 7 | 10 | 0 | 0.411764706 | 1 | # | 20 | Small | aldex2 | 49 |
| # | sex | 4 | 1 | 0 | 0.8 | 1 | # | 20 | Small | aldex2 | 49 |
| # | time | 5 | 5 | 0 | 0.5 | 1 | # | 20 | Small | aldex2 | 49 |
| # | interaction | 0 | 0 | 4 | 0 | 0 | # | 20 | Small | aldex2 | 49 |
| # | disease | 4 | 8 | 0 | 0.333333333 | 1 | # | 10 | Baseline | paralle | 50 |
| # | sex | 2 | 6 | 0 | 0.25 | 1 | # | 10 | Baseline | paralle | 50 |
| # | time | 2 | 11 | 0 | 0.153846154 | 1 | # | 10 | Baseline | paralle | 50 |
| # | interaction | 2 | 7 | 0 | 0.222222222 | 1 | # | 10 | Baseline | paralle | 50 |
| # | disease | 4 | 1 | 0 | 0.8 | 1 | # | 10 | Baseline | ancom | 50 |
| # | sex | 2 | 0 | 0 | 1 | 1 | # | 10 | Baseline | ancom | 50 |
| # | time | 2 | 1 | 0 | 0.666666667 | 1 | # | 10 | Baseline | ancom | 50 |
| # | interaction | 0 | 0 | 2 | 0 | 0 | # | 10 | Baseline | ancom | 50 |
| # | disease | 4 | 12 | 0 | 0.25 | 1 | # | 10 | Baseline | aldex2 | 50 |
| # | sex | 2 | 5 | 0 | 0.285714286 | 1 | # | 10 | Baseline | aldex2 | 50 |
| # | time | 2 | 7 | 0 | 0.222222222 | 1 | # | 10 | Baseline | aldex2 | 50 |
| # | interaction | 0 | 0 | 2 | 0 | 0 | # | 10 | Baseline | aldex2 | 50 |
| # | disease | 4 | 0 | 0 | 1 | 1 | # | 10 | Small | paralle | 50 |
| # | sex | 2 | 0 | 0 | 1 | 1 | # | 10 | Small | paralle | 50 |
| # | time | 2 | 0 | 0 | 1 | 1 | # | 10 | Small | paralle | 50 |
| # | interaction | 1 | 0 | 1 | 1 | 0.5 | # | 10 | Small | paralle | 50 |
| # | disease | 4 | 1 | 0 | 0.8 | 1 | # | 10 | Small | ancom | 50 |
| # | sex | 2 | 0 | 0 | 1 | 1 | # | 10 | Small | ancom | 50 |
| # | time | 2 | 0 | 0 | 1 | 1 | # | 10 | Small | ancom | 50 |
| # | interaction | 0 | 0 | 2 | 0 | 0 | # | 10 | Small | ancom | 50 |
| # | disease | 4 | 1 | 0 | 0.8 | 1 | # | 10 | Small | aldex2 | 50 |
| # | sex | 2 | 0 | 0 | 1 | 1 | # | 10 | Small | aldex2 | 50 |
| # | time | 2 | 1 | 0 | 0.666666667 | 1 | # | 10 | Small | aldex2 | 50 |
| # | interaction | 0 | 0 | 2 | 0 | 0 | # | 10 | Small | aldex2 | 50 |
| # | disease | 4 | 10 | 0 | 0.285714286 | 1 | # | 10 | Baseline | paralle | 50 |

| | | | | | | | | | | | |
|---|---|---|---|---|---|---|---|---|---|---|---|
| # | sex | 2 | 2 | 0 | 0.5 | 1 | # | 10 | Baseline | paralle | 50 |
| # | time | 2 | 4 | 0 | 0.333333333 | 1 | # | 10 | Baseline | paralle | 50 |
| # | interaction | 2 | 4 | 0 | 0.333333333 | 1 | # | 10 | Baseline | paralle | 50 |
| # | disease | 4 | 2 | 0 | 0.666666667 | 1 | # | 10 | Baseline | ancom | 50 |
| # | sex | 2 | 0 | 0 | 1 | 1 | # | 10 | Baseline | ancom | 50 |
| # | time | 2 | 2 | 0 | 0.5 | 1 | # | 10 | Baseline | ancom | 50 |
| # | interaction | 0 | 0 | 2 | 0 | 0 | # | 10 | Baseline | ancom | 50 |
| # | disease | 4 | 19 | 0 | 0.173913043 | 1 | # | 10 | Baseline | aldex2 | 50 |
| # | sex | 2 | 5 | 0 | 0.285714286 | 1 | # | 10 | Baseline | aldex2 | 50 |
| # | time | 2 | 8 | 0 | 0.2 | 1 | # | 10 | Baseline | aldex2 | 50 |
| # | interaction | 0 | 0 | 2 | 0 | 0 | # | 10 | Baseline | aldex2 | 50 |
| # | disease | 4 | 0 | 0 | 1 | 1 | # | 10 | Small | paralle | 50 |
| # | sex | 2 | 0 | 0 | 1 | 1 | # | 10 | Small | paralle | 50 |
| # | time | 2 | 0 | 0 | 1 | 1 | # | 10 | Small | paralle | 50 |
| # | interaction | 2 | 0 | 0 | 1 | 1 | # | 10 | Small | paralle | 50 |
| # | disease | 4 | 2 | 0 | 0.666666667 | 1 | # | 10 | Small | ancom | 50 |
| # | sex | 2 | 0 | 0 | 1 | 1 | # | 10 | Small | ancom | 50 |
| # | time | 2 | 2 | 0 | 0.5 | 1 | # | 10 | Small | ancom | 50 |
| # | interaction | 0 | 0 | 2 | 0 | 0 | # | 10 | Small | ancom | 50 |
| # | disease | 4 | 2 | 0 | 0.666666667 | 1 | # | 10 | Small | aldex2 | 50 |
| # | sex | 2 | 1 | 0 | 0.666666667 | 1 | # | 10 | Small | aldex2 | 50 |
| # | time | 2 | 2 | 0 | 0.5 | 1 | # | 10 | Small | aldex2 | 50 |
| # | interaction | 0 | 0 | 2 | 0 | 0 | # | 10 | Small | aldex2 | 50 |
| # | disease | 4 | 10 | 0 | 0.285714286 | 1 | # | 10 | Baseline | paralle | 50 |
| # | sex | 2 | 0 | 0 | 1 | 1 | # | 10 | Baseline | paralle | 50 |
| # | time | 2 | 4 | 0 | 0.333333333 | 1 | # | 10 | Baseline | paralle | 50 |
| # | interaction | 2 | 4 | 0 | 0.333333333 | 1 | # | 10 | Baseline | paralle | 50 |
| # | disease | 4 | 2 | 0 | 0.666666667 | 1 | # | 10 | Baseline | ancom | 50 |
| # | sex | 2 | 0 | 0 | 1 | 1 | # | 10 | Baseline | ancom | 50 |
| # | time | 2 | 2 | 0 | 0.5 | 1 | # | 10 | Baseline | ancom | 50 |
| # | interaction | 0 | 0 | 2 | 0 | 0 | # | 10 | Baseline | ancom | 50 |
| # | disease | 4 | 22 | 0 | 0.153846154 | 1 | # | 10 | Baseline | aldex2 | 50 |
| # | sex | 2 | 7 | 0 | 0.222222222 | 1 | # | 10 | Baseline | aldex2 | 50 |
| # | time | 2 | 7 | 0 | 0.222222222 | 1 | # | 10 | Baseline | aldex2 | 50 |
| # | interaction | 0 | 0 | 2 | 0 | 0 | # | 10 | Baseline | aldex2 | 50 |
| # | disease | 4 | 0 | 0 | 1 | 1 | # | 10 | Small | paralle | 50 |
| # | sex | 2 | 0 | 0 | 1 | 1 | # | 10 | Small | paralle | 50 |
| # | time | 2 | 1 | 0 | 0.666666667 | 1 | # | 10 | Small | paralle | 50 |
| # | interaction | 2 | 1 | 0 | 0.666666667 | 1 | # | 10 | Small | paralle | 50 |
| # | disease | 4 | 2 | 0 | 0.666666667 | 1 | # | 10 | Small | ancom | 50 |
| # | sex | 2 | 0 | 0 | 1 | 1 | # | 10 | Small | ancom | 50 |
| # | time | 2 | 2 | 0 | 0.5 | 1 | # | 10 | Small | ancom | 50 |
| # | interaction | 0 | 0 | 2 | 0 | 0 | # | 10 | Small | ancom | 50 |
| # | disease | 4 | 2 | 0 | 0.666666667 | 1 | # | 10 | Small | aldex2 | 50 |
| # | sex | 2 | 0 | 0 | 1 | 1 | # | 10 | Small | aldex2 | 50 |
| # | time | 2 | 2 | 0 | 0.5 | 1 | # | 10 | Small | aldex2 | 50 |
| # | interaction | 0 | 0 | 2 | 0 | 0 | # | 10 | Small | aldex2 | 50 |
| # | disease | 4 | 3 | 0 | 0.571428571 | 1 | # | 10 | Baseline | paralle | 50 |
| # | sex | 2 | 3 | 0 | 0.4 | 1 | # | 10 | Baseline | paralle | 50 |
| # | time | 2 | 3 | 0 | 0.4 | 1 | # | 10 | Baseline | paralle | 50 |
| # | interaction | 2 | 2 | 0 | 0.5 | 1 | # | 10 | Baseline | paralle | 50 |
| # | disease | 4 | 2 | 0 | 0.666666667 | 1 | # | 10 | Baseline | ancom | 50 |
| # | sex | 2 | 0 | 0 | 1 | 1 | # | 10 | Baseline | ancom | 50 |
| # | time | 2 | 2 | 0 | 0.5 | 1 | # | 10 | Baseline | ancom | 50 |
| # | interaction | 0 | 0 | 2 | 0 | 0 | # | 10 | Baseline | ancom | 50 |
| # | disease | 4 | 24 | 0 | 0.142857143 | 1 | # | 10 | Baseline | aldex2 | 50 |
| # | sex | 2 | 6 | 0 | 0.25 | 1 | # | 10 | Baseline | aldex2 | 50 |
| # | time | 2 | 8 | 0 | 0.2 | 1 | # | 10 | Baseline | aldex2 | 50 |
| # | interaction | 0 | 0 | 2 | 0 | 0 | # | 10 | Baseline | aldex2 | 50 |
| # | disease | 4 | 0 | 0 | 1 | 1 | # | 10 | Small | paralle | 50 |
| # | sex | 2 | 0 | 0 | 1 | 1 | # | 10 | Small | paralle | 50 |
| # | time | 2 | 0 | 0 | 1 | 1 | # | 10 | Small | paralle | 50 |
| # | interaction | 2 | 1 | 0 | 0.666666667 | 1 | # | 10 | Small | paralle | 50 |
| # | disease | 4 | 2 | 0 | 0.666666667 | 1 | # | 10 | Small | ancom | 50 |
| # | sex | 2 | 0 | 0 | 1 | 1 | # | 10 | Small | ancom | 50 |
| # | time | 2 | 2 | 0 | 0.5 | 1 | # | 10 | Small | ancom | 50 |
| # | interaction | 0 | 0 | 2 | 0 | 0 | # | 10 | Small | ancom | 50 |
| # | disease | 4 | 3 | 0 | 0.571428571 | 1 | # | 10 | Small | aldex2 | 50 |
| # | sex | 2 | 0 | 0 | 1 | 1 | # | 10 | Small | aldex2 | 50 |
| # | time | 2 | 2 | 0 | 0.5 | 1 | # | 10 | Small | aldex2 | 50 |
| # | interaction | 0 | 0 | 2 | 0 | 0 | # | 10 | Small | aldex2 | 50 |
| # | disease | 7 | 8 | 0 | 0.466666667 | 1 | # | 20 | Baseline | paralle | 50 |
| # | sex | 4 | 8 | 0 | 0.333333333 | 1 | # | 20 | Baseline | paralle | 50 |
| # | time | 5 | 8 | 0 | 0.384615385 | 1 | # | 20 | Baseline | paralle | 50 |
| # | interaction | 4 | 9 | 0 | 0.307692308 | 1 | # | 20 | Baseline | paralle | 50 |
| # | disease | 7 | 0 | 0 | 1 | 1 | # | 20 | Baseline | ancom | 50 |
| # | sex | 4 | 0 | 0 | 1 | 1 | # | 20 | Baseline | ancom | 50 |
| # | time | 5 | 1 | 0 | 0.833333333 | 1 | # | 20 | Baseline | ancom | 50 |
| # | interaction | 0 | 0 | 4 | 0 | 0 | # | 20 | Baseline | ancom | 50 |
| # | disease | 7 | 19 | 0 | 0.269230769 | 1 | # | 20 | Baseline | aldex2 | 50 |
| # | sex | 4 | 8 | 0 | 0.333333333 | 1 | # | 20 | Baseline | aldex2 | 50 |
| # | time | 5 | 14 | 0 | 0.263157895 | 1 | # | 20 | Baseline | aldex2 | 50 |
| # | interaction | 0 | 0 | 4 | 0 | 0 | # | 20 | Baseline | aldex2 | 50 |
| # | disease | 7 | 1 | 0 | 0.875 | 1 | # | 20 | Small | paralle | 50 |
| # | sex | 4 | 0 | 0 | 1 | 1 | # | 20 | Small | paralle | 50 |
| # | time | 5 | 0 | 0 | 1 | 1 | # | 20 | Small | paralle | 50 |
| # | interaction | 1 | 0 | 3 | 1 | 0.25 | # | 20 | Small | paralle | 50 |
| # | disease | 7 | 0 | 0 | 1 | 1 | # | 20 | Small | ancom | 50 |
| # | sex | 4 | 0 | 0 | 1 | 1 | # | 20 | Small | ancom | 50 |
| # | time | 5 | 1 | 0 | 0.833333333 | 1 | # | 20 | Small | ancom | 50 |
| # | interaction | 0 | 0 | 4 | 0 | 0 | # | 20 | Small | ancom | 50 |
| # | disease | 7 | 0 | 0 | 1 | 1 | # | 20 | Small | aldex2 | 50 |
| # | sex | 4 | 0 | 0 | 1 | 1 | # | 20 | Small | aldex2 | 50 |
| # | time | 5 | 1 | 0 | 0.833333333 | 1 | # | 20 | Small | aldex2 | 50 |
| # | interaction | 0 | 0 | 4 | 0 | 0 | # | 20 | Small | aldex2 | 50 |
| # | disease | 7 | 15 | 0 | 0.318181818 | 1 | # | 20 | Baseline | paralle | 50 |
| # | sex | 4 | 8 | 0 | 0.333333333 | 1 | # | 20 | Baseline | paralle | 50 |
| # | time | 5 | 9 | 0 | 0.357142857 | 1 | # | 20 | Baseline | paralle | 50 |
| # | interaction | 4 | 5 | 0 | 0.444444444 | 1 | # | 20 | Baseline | paralle | 50 |
| # | disease | 7 | 4 | 0 | 0.636363636 | 1 | # | 20 | Baseline | ancom | 50 |
| # | sex | 4 | 0 | 0 | 1 | 1 | # | 20 | Baseline | ancom | 50 |
| # | time | 5 | 4 | 0 | 0.555555556 | 1 | # | 20 | Baseline | ancom | 50 |
| # | interaction | 0 | 0 | 4 | 0 | 0 | # | 20 | Baseline | ancom | 50 |
| # | disease | 7 | 44 | 0 | 0.137254902 | 1 | # | 20 | Baseline | aldex2 | 50 |
| # | sex | 4 | 11 | 0 | 0.266666667 | 1 | # | 20 | Baseline | aldex2 | 50 |
| # | time | 5 | 20 | 0 | 0.2 | 1 | # | 20 | Baseline | aldex2 | 50 |
| # | interaction | 0 | 0 | 4 | 0 | 0 | # | 20 | Baseline | aldex2 | 50 |
| # | disease | 7 | 0 | 0 | 1 | 1 | # | 20 | Small | paralle | 50 |
| # | sex | 4 | 0 | 0 | 1 | 1 | # | 20 | Small | paralle | 50 |
| # | time | 5 | 0 | 0 | 1 | 1 | # | 20 | Small | paralle | 50 |
| # | interaction | 4 | 0 | 0 | 1 | 1 | # | 20 | Small | paralle | 50 |
| # | disease | 7 | 4 | 0 | 0.636363636 | 1 | # | 20 | Small | ancom | 50 |
| # | sex | 4 | 0 | 0 | 1 | 1 | # | 20 | Small | ancom | 50 |
| # | time | 5 | 4 | 0 | 0.555555556 | 1 | # | 20 | Small | ancom | 50 |
| # | interaction | 0 | 0 | 4 | 0 | 0 | # | 20 | Small | ancom | 50 |
| # | disease | 7 | 6 | 0 | 0.538461538 | 1 | # | 20 | Small | aldex2 | 50 |
| # | sex | 4 | 1 | 0 | 0.8 | 1 | # | 20 | Small | aldex2 | 50 |
| # | time | 5 | 4 | 0 | 0.555555556 | 1 | # | 20 | Small | aldex2 | 50 |
| # | interaction | 0 | 0 | 4 | 0 | 0 | # | 20 | Small | aldex2 | 50 |
| # | disease | 7 | 14 | 0 | 0.333333333 | 1 | # | 20 | Baseline | paralle | 50 |
| # | sex | 4 | 5 | 0 | 0.444444444 | 1 | # | 20 | Baseline | paralle | 50 |
| # | time | 5 | 7 | 0 | 0.416666667 | 1 | # | 20 | Baseline | paralle | 50 |
| # | interaction | 4 | 3 | 0 | 0.571428571 | 1 | # | 20 | Baseline | paralle | 50 |
| # | disease | 7 | 4 | 0 | 0.636363636 | 1 | # | 20 | Baseline | ancom | 50 |
| # | sex | 4 | 0 | 0 | 1 | 1 | # | 20 | Baseline | ancom | 50 |
| # | time | 5 | 4 | 0 | 0.555555556 | 1 | # | 20 | Baseline | ancom | 50 |
| # | interaction | 0 | 0 | 4 | 0 | 0 | # | 20 | Baseline | ancom | 50 |
| # | disease | 7 | 55 | 0 | 0.112903226 | 1 | # | 20 | Baseline | aldex2 | 50 |
| # | sex | 4 | 11 | 0 | 0.266666667 | 1 | # | 20 | Baseline | aldex2 | 50 |
| # | time | 5 | 27 | 0 | 0.15625 | 1 | # | 20 | Baseline | aldex2 | 50 |
| # | interaction | 0 | 0 | 4 | 0 | 0 | # | 20 | Baseline | aldex2 | 50 |
| # | disease | 7 | 1 | 0 | 0.875 | 1 | # | 20 | Small | paralle | 50 |
| # | sex | 4 | 0 | 0 | 1 | 1 | # | 20 | Small | paralle | 50 |
| # | time | 5 | 0 | 0 | 1 | 1 | # | 20 | Small | paralle | 50 |
| # | interaction | 4 | 0 | 0 | 1 | 1 | # | 20 | Small | paralle | 50 |
| # | disease | 7 | 4 | 0 | 0.636363636 | 1 | # | 20 | Small | ancom | 50 |
| # | sex | 4 | 0 | 0 | 1 | 1 | # | 20 | Small | ancom | 50 |
| # | time | 5 | 4 | 0 | 0.555555556 | 1 | # | 20 | Small | ancom | 50 |
| # | interaction | 0 | 0 | 4 | 0 | 0 | # | 20 | Small | ancom | 50 |
| # | disease | 7 | 9 | 0 | 0.4375 | 1 | # | 20 | Small | aldex2 | 50 |
| # | sex | 4 | 0 | 0 | 1 | 1 | # | 20 | Small | aldex2 | 50 |
| # | time | 5 | 5 | 0 | 0.5 | 1 | # | 20 | Small | aldex2 | 50 |
| # | interaction | 0 | 0 | 4 | 0 | 0 | # | 20 | Small | aldex2 | 50 |
| # | disease | 7 | 9 | 0 | 0.4375 | 1 | # | 20 | Baseline | paralle | 50 |
| # | sex | 4 | 4 | 0 | 0.5 | 1 | # | 20 | Baseline | paralle | 50 |
| # | time | 5 | 5 | 0 | 0.5 | 1 | # | 20 | Baseline | paralle | 50 |
| # | interaction | 4 | 1 | 0 | 0.8 | 1 | # | 20 | Baseline | paralle | 50 |
| # | disease | 7 | 4 | 0 | 0.636363636 | 1 | # | 20 | Baseline | ancom | 50 |
| # | sex | 4 | 0 | 0 | 1 | 1 | # | 20 | Baseline | ancom | 50 |
| # | time | 5 | 5 | 0 | 0.5 | 1 | # | 20 | Baseline | ancom | 50 |
| # | interaction | 0 | 0 | 4 | 0 | 0 | # | 20 | Baseline | ancom | 50 |
| # | disease | 7 | 70 | 0 | 0.090909091 | 1 | # | 20 | Baseline | aldex2 | 50 |
| # | sex | 4 | 15 | 0 | 0.210526316 | 1 | # | 20 | Baseline | aldex2 | 50 |
| # | time | 5 | 37 | 0 | 0.119047619 | 1 | # | 20 | Baseline | aldex2 | 50 |
| # | interaction | 0 | 0 | 4 | 0 | 0 | # | 20 | Baseline | aldex2 | 50 |
| # | disease | 7 | 0 | 0 | 1 | 1 | # | 20 | Small | paralle | 50 |
| # | sex | 4 | 0 | 0 | 1 | 1 | # | 20 | Small | paralle | 50 |
| # | time | 5 | 0 | 0 | 1 | 1 | # | 20 | Small | paralle | 50 |
| # | interaction | 4 | 0 | 0 | 1 | 1 | # | 20 | Small | paralle | 50 |
| # | disease | 7 | 4 | 0 | 0.636363636 | 1 | # | 20 | Small | ancom | 50 |
| # | sex | 4 | 0 | 0 | 1 | 1 | # | 20 | Small | ancom | 50 |
| # | time | 5 | 5 | 0 | 0.5 | 1 | # | 20 | Small | ancom | 50 |
| # | interaction | 0 | 0 | 4 | 0 | 0 | # | 20 | Small | ancom | 50 |
| # | disease | 7 | 12 | 0 | 0.368421053 | 1 | # | 20 | Small | aldex2 | 50 |
| # | sex | 4 | 0 | 0 | 1 | 1 | # | 20 | Small | aldex2 | 50 |
| # | time | 5 | 6 | 0 | 0.454545455 | 1 | # | 20 | Small | aldex2 | 50 |
| # | interaction | 0 | 0 | 4 | 0 | 0 | # | 20 | Small | aldex2 | 50 |
| # | disease | 4 | 10 | 0 | 0.285714286 | 1 | # | 10 | Baseline | paralle | 51 |
| # | sex | 2 | 2 | 0 | 0.5 | 1 | # | 10 | Baseline | paralle | 51 |
| # | time | 2 | 5 | 0 | 0.285714286 | 1 | # | 10 | Baseline | paralle | 51 |
| # | interaction | 2 | 10 | 0 | 0.166666667 | 1 | # | 10 | Baseline | paralle | 51 |
| # | disease | 4 | 1 | 0 | 0.8 | 1 | # | 10 | Baseline | ancom | 51 |
| # | sex | 2 | 0 | 0 | 1 | 1 | # | 10 | Baseline | ancom | 51 |
| # | time | 2 | 0 | 0 | 1 | 1 | # | 10 | Baseline | ancom | 51 |
| # | interaction | 0 | 0 | 2 | 0 | 0 | # | 10 | Baseline | ancom | 51 |
| # | disease | 4 | 7 | 0 | 0.363636364 | 1 | # | 10 | Baseline | aldex2 | 51 |
| # | sex | 2 | 9 | 0 | 0.181818182 | 1 | # | 10 | Baseline | aldex2 | 51 |
| # | time | 2 | 8 | 0 | 0.2 | 1 | # | 10 | Baseline | aldex2 | 51 |
| # | interaction | 0 | 0 | 2 | 0 | 0 | # | 10 | Baseline | aldex2 | 51 |
| # | disease | 4 | 0 | 0 | 1 | 1 | # | 10 | Small | paralle | 51 |
| # | sex | 2 | 0 | 0 | 1 | 1 | # | 10 | Small | paralle | 51 |
| # | time | 2 | 0 | 0 | 1 | 1 | # | 10 | Small | paralle | 51 |
| # | interaction | 2 | 1 | 0 | 0.666666667 | 1 | # | 10 | Small | paralle | 51 |
| # | disease | 4 | 0 | 0 | 1 | 1 | # | 10 | Small | ancom | 51 |
| # | sex | 2 | 0 | 0 | 1 | 1 | # | 10 | Small | ancom | 51 |
| # | time | 2 | 0 | 0 | 1 | 1 | # | 10 | Small | ancom | 51 |
| # | interaction | 0 | 0 | 2 | 0 | 0 | # | 10 | Small | ancom | 51 |
| # | disease | 4 | 1 | 0 | 0.8 | 1 | # | 10 | Small | aldex2 | 51 |
| # | sex | 2 | 0 | 0 | 1 | 1 | # | 10 | Small | aldex2 | 51 |
| # | time | 2 | 1 | 0 | 0.666666667 | 1 | # | 10 | Small | aldex2 | 51 |
| # | interaction | 0 | 0 | 2 | 0 | 0 | # | 10 | Small | aldex2 | 51 |
| # | disease | 4 | 8 | 0 | 0.333333333 | 1 | # | 10 | Baseline | paralle | 51 |
| # | sex | 2 | 7 | 0 | 0.222222222 | 1 | # | 10 | Baseline | paralle | 51 |
| # | time | 2 | 3 | 0 | 0.4 | 1 | # | 10 | Baseline | paralle | 51 |
| # | interaction | 2 | 5 | 0 | 0.285714286 | 1 | # | 10 | Baseline | paralle | 51 |
| # | disease | 4 | 2 | 0 | 0.666666667 | 1 | # | 10 | Baseline | ancom | 51 |
| # | sex | 2 | 0 | 0 | 1 | 1 | # | 10 | Baseline | ancom | 51 |
| # | time | 2 | 2 | 0 | 0.5 | 1 | # | 10 | Baseline | ancom | 51 |
| # | interaction | 0 | 0 | 2 | 0 | 0 | # | 10 | Baseline | ancom | 51 |
| # | disease | 4 | 12 | 0 | 0.25 | 1 | # | 10 | Baseline | aldex2 | 51 |
| # | sex | 2 | 9 | 0 | 0.181818182 | 1 | # | 10 | Baseline | aldex2 | 51 |
| # | time | 2 | 8 | 0 | 0.2 | 1 | # | 10 | Baseline | aldex2 | 51 |
| # | interaction | 0 | 0 | 2 | 0 | 0 | # | 10 | Baseline | aldex2 | 51 |
| # | disease | 4 | 0 | 0 | 1 | 1 | # | 10 | Small | paralle | 51 |
| # | sex | 2 | 0 | 0 | 1 | 1 | # | 10 | Small | paralle | 51 |
| # | time | 2 | 0 | 0 | 1 | 1 | # | 10 | Small | paralle | 51 |
| # | interaction | 2 | 0 | 0 | 1 | 1 | # | 10 | Small | paralle | 51 |
| # | disease | 4 | 2 | 0 | 0.666666667 | 1 | # | 10 | Small | ancom | 51 |
| # | sex | 2 | 0 | 0 | 1 | 1 | # | 10 | Small | ancom | 51 |
| # | time | 2 | 2 | 0 | 0.5 | 1 | # | 10 | Small | ancom | 51 |
| # | interaction | 0 | 0 | 2 | 0 | 0 | # | 10 | Small | ancom | 51 |
| # | disease | 4 | 1 | 0 | 0.8 | 1 | # | 10 | Small | aldex2 | 51 |
| # | sex | 2 | 0 | 0 | 1 | 1 | # | 10 | Small | aldex2 | 51 |
| # | time | 2 | 1 | 0 | 0.666666667 | 1 | # | 10 | Small | aldex2 | 51 |
| # | interaction | 0 | 0 | 2 | 0 | 0 | # | 10 | Small | aldex2 | 51 |
| # | disease | 4 | 6 | 0 | 0.4 | 1 | # | 10 | Baseline | paralle | 51 |
| # | sex | 2 | 5 | 0 | 0.285714286 | 1 | # | 10 | Baseline | paralle | 51 |
| # | time | 2 | 2 | 0 | 0.5 | 1 | # | 10 | Baseline | paralle | 51 |
| # | interaction | 2 | 3 | 0 | 0.4 | 1 | # | 10 | Baseline | paralle | 51 |
| # | disease | 4 | 2 | 0 | 0.666666667 | 1 | # | 10 | Baseline | ancom | 51 |
| # | sex | 2 | 0 | 0 | 1 | 1 | # | 10 | Baseline | ancom | 51 |
| # | time | 2 | 2 | 0 | 0.5 | 1 | # | 10 | Baseline | ancom | 51 |
| # | interaction | 0 | 0 | 2 | 0 | 0 | # | 10 | Baseline | ancom | 51 |
| # | disease | 4 | 19 | 0 | 0.173913043 | 1 | # | 10 | Baseline | aldex2 | 51 |
| # | sex | 2 | 7 | 0 | 0.222222222 | 1 | # | 10 | Baseline | aldex2 | 51 |
| # | time | 2 | 8 | 0 | 0.2 | 1 | # | 10 | Baseline | aldex2 | 51 |
| # | interaction | 0 | 0 | 2 | 0 | 0 | # | 10 | Baseline | aldex2 | 51 |
| # | disease | 4 | 0 | 0 | 1 | 1 | # | 10 | Small | paralle | 51 |
| # | sex | 2 | 0 | 0 | 1 | 1 | # | 10 | Small | paralle | 51 |
| # | time | 2 | 0 | 0 | 1 | 1 | # | 10 | Small | paralle | 51 |
| # | interaction | 2 | 0 | 0 | 1 | 1 | # | 10 | Small | paralle | 51 |
| # | disease | 4 | 2 | 0 | 0.666666667 | 1 | # | 10 | Small | ancom | 51 |
| # | sex | 2 | 0 | 0 | 1 | 1 | # | 10 | Small | ancom | 51 |
| # | time | 2 | 2 | 0 | 0.5 | 1 | # | 10 | Small | ancom | 51 |
| # | interaction | 0 | 0 | 2 | 0 | 0 | # | 10 | Small | ancom | 51 |
| # | disease | 4 | 2 | 0 | 0.666666667 | 1 | # | 10 | Small | aldex2 | 51 |
| # | sex | 2 | 0 | 0 | 1 | 1 | # | 10 | Small | aldex2 | 51 |
| # | time | 2 | 3 | 0 | 0.4 | 1 | # | 10 | Small | aldex2 | 51 |
| # | interaction | 0 | 0 | 2 | 0 | 0 | # | 10 | Small | aldex2 | 51 |
| # | disease | 4 | 4 | 0 | 0.5 | 1 | # | 10 | Baseline | paralle | 51 |
| # | sex | 2 | 3 | 0 | 0.4 | 1 | # | 10 | Baseline | paralle | 51 |
| # | time | 2 | 4 | 0 | 0.333333333 | 1 | # | 10 | Baseline | paralle | 51 |
| # | interaction | 2 | 3 | 0 | 0.4 | 1 | # | 10 | Baseline | paralle | 51 |
| # | disease | 4 | 2 | 0 | 0.666666667 | 1 | # | 10 | Baseline | ancom | 51 |
| # | sex | 2 | 0 | 0 | 1 | 1 | # | 10 | Baseline | ancom | 51 |
| # | time | 2 | 2 | 0 | 0.5 | 1 | # | 10 | Baseline | ancom | 51 |
| # | interaction | 0 | 0 | 2 | 0 | 0 | # | 10 | Baseline | ancom | 51 |
| # | disease | 4 | 24 | 0 | 0.142857143 | 1 | # | 10 | Baseline | aldex2 | 51 |
| # | sex | 2 | 12 | 0 | 0.142857143 | 1 | # | 10 | Baseline | aldex2 | 51 |
| # | time | 2 | 14 | 0 | 0.125 | 1 | # | 10 | Baseline | aldex2 | 51 |
| # | interaction | 0 | 0 | 2 | 0 | 0 | # | 10 | Baseline | aldex2 | 51 |
| # | disease | 4 | 0 | 0 | 1 | 1 | # | 10 | Small | paralle | 51 |
| # | sex | 2 | 0 | 0 | 1 | 1 | # | 10 | Small | paralle | 51 |
| # | time | 2 | 0 | 0 | 1 | 1 | # | 10 | Small | paralle | 51 |
| # | interaction | 2 | 0 | 0 | 1 | 1 | # | 10 | Small | paralle | 51 |
| # | disease | 4 | 2 | 0 | 0.666666667 | 1 | # | 10 | Small | ancom | 51 |
| # | sex | 2 | 0 | 0 | 1 | 1 | # | 10 | Small | ancom | 51 |
| # | time | 2 | 2 | 0 | 0.5 | 1 | # | 10 | Small | ancom | 51 |
| # | interaction | 0 | 0 | 2 | 0 | 0 | # | 10 | Small | ancom | 51 |
| # | disease | 4 | 3 | 0 | 0.571428571 | 1 | # | 10 | Small | aldex2 | 51 |
| # | sex | 2 | 0 | 0 | 1 | 1 | # | 10 | Small | aldex2 | 51 |
| # | time | 2 | 3 | 0 | 0.4 | 1 | # | 10 | Small | aldex2 | 51 |
| # | interaction | 0 | 0 | 2 | 0 | 0 | # | 10 | Small | aldex2 | 51 |
| # | disease | 7 | 5 | 0 | 0.583333333 | 1 | # | 20 | Baseline | paralle | 51 |
| # | sex | 4 | 8 | 0 | 0.333333333 | 1 | # | 20 | Baseline | paralle | 51 |
| # | time | 5 | 6 | 0 | 0.454545455 | 1 | # | 20 | Baseline | paralle | 51 |
| # | interaction | 4 | 6 | 0 | 0.4 | 1 | # | 20 | Baseline | paralle | 51 |
| # | disease | 7 | 1 | 0 | 0.875 | 1 | # | 20 | Baseline | ancom | 51 |
| # | sex | 4 | 0 | 0 | 1 | 1 | # | 20 | Baseline | ancom | 51 |
| # | time | 5 | 1 | 0 | 0.833333333 | 1 | # | 20 | Baseline | ancom | 51 |
| # | interaction | 0 | 0 | 4 | 0 | 0 | # | 20 | Baseline | ancom | 51 |
| # | disease | 7 | 20 | 0 | 0.259259259 | 1 | # | 20 | Baseline | aldex2 | 51 |
| # | sex | 4 | 6 | 0 | 0.4 | 1 | # | 20 | Baseline | aldex2 | 51 |
| # | time | 5 | 17 | 0 | 0.227272727 | 1 | # | 20 | Baseline | aldex2 | 51 |
| # | interaction | 0 | 0 | 4 | 0 | 0 | # | 20 | Baseline | aldex2 | 51 |
| # | disease | 7 | 1 | 0 | 0.875 | 1 | # | 20 | Small | paralle | 51 |
| # | sex | 4 | 0 | 0 | 1 | 1 | # | 20 | Small | paralle | 51 |
| # | time | 5 | 0 | 0 | 1 | 1 | # | 20 | Small | paralle | 51 |
| # | interaction | 3 | 0 | 1 | 1 | 0.75 | # | 20 | Small | paralle | 51 |
| # | disease | 7 | 1 | 0 | 0.875 | 1 | # | 20 | Small | ancom | 51 |
| # | sex | 4 | 0 | 0 | 1 | 1 | # | 20 | Small | ancom | 51 |
| # | time | 5 | 1 | 0 | 0.833333333 | 1 | # | 20 | Small | ancom | 51 |
| # | interaction | 0 | 0 | 4 | 0 | 0 | # | 20 | Small | ancom | 51 |
| # | disease | 7 | 0 | 0 | 1 | 1 | # | 20 | Small | aldex2 | 51 |
| # | sex | 4 | 0 | 0 | 1 | 1 | # | 20 | Small | aldex2 | 51 |
| # | time | 5 | 0 | 0 | 1 | 1 | # | 20 | Small | aldex2 | 51 |
| # | interaction | 0 | 0 | 4 | 0 | 0 | # | 20 | Small | aldex2 | 51 |
| # | disease | 7 | 9 | 0 | 0.4375 | 1 | # | 20 | Baseline | paralle | 51 |
| # | sex | 4 | 8 | 0 | 0.333333333 | 1 | # | 20 | Baseline | paralle | 51 |
| # | time | 5 | 3 | 0 | 0.625 | 1 | # | 20 | Baseline | paralle | 51 |
| # | interaction | 4 | 4 | 0 | 0.5 | 1 | # | 20 | Baseline | paralle | 51 |
| # | disease | 7 | 3 | 0 | 0.7 | 1 | # | 20 | Baseline | ancom | 51 |
| # | sex | 4 | 0 | 0 | 1 | 1 | # | 20 | Baseline | ancom | 51 |
| # | time | 5 | 3 | 0 | 0.625 | 1 | # | 20 | Baseline | ancom | 51 |
| # | interaction | 0 | 0 | 4 | 0 | 0 | # | 20 | Baseline | ancom | 51 |
| # | disease | 7 | 60 | 0 | 0.14893617 | 1 | # | 20 | Baseline | aldex2 | 51 |
| # | sex | 4 | 10 | 0 | 0.285714286 | 1 | # | 20 | Baseline | aldex2 | 51 |
| # | time | 5 | 25 | 0 | 0.166666667 | 1 | # | 20 | Baseline | aldex2 | 51 |
| # | interaction | 0 | 0 | 4 | 0 | 0 | # | 20 | Baseline | aldex2 | 51 |
| # | disease | 7 | 1 | 0 | 0.875 | 1 | # | 20 | Small | paralle | 51 |
| # | sex | 4 | 0 | 0 | 1 | 1 | # | 20 | Small | paralle | 51 |
| # | time | 5 | 0 | 0 | 1 | 1 | # | 20 | Small | paralle | 51 |
| # | interaction | 4 | 0 | 0 | 1 | 1 | # | 20 | Small | paralle | 51 |
| # | disease | 7 | 2 | 0 | 0.777777778 | 1 | # | 20 | Small | ancom | 51 |
| # | sex | 4 | 0 | 0 | 1 | 1 | # | 20 | Small | ancom | 51 |
| # | time | 5 | 3 | 0 | 0.625 | 1 | # | 20 | Small | ancom | 51 |
| # | interaction | 0 | 0 | 4 | 0 | 0 | # | 20 | Small | ancom | 51 |
| # | disease | 7 | 4 | 0 | 0.636363636 | 1 | # | 20 | Small | aldex2 | 51 |
| # | sex | 4 | 1 | 0 | 0.8 | 1 | # | 20 | Small | aldex2 | 51 |
| # | time | 5 | 2 | 0 | 0.714285714 | 1 | # | 20 | Small | aldex2 | 51 |
| # | interaction | 0 | 0 | 4 | 0 | 0 | # | 20 | Small | aldex2 | 51 |
| # | disease | 7 | 8 | 0 | 0.466666667 | 1 | # | 20 | Baseline | paralle | 51 |
| # | sex | 4 | 8 | 0 | 0.333333333 | 1 | # | 20 | Baseline | paralle | 51 |
| # | time | 5 | 2 | 0 | 0.714285714 | 1 | # | 20 | Baseline | paralle | 51 |
| # | interaction | 4 | 5 | 0 | 0.444444444 | 1 | # | 20 | Baseline | paralle | 51 |
| # | disease | 7 | 4 | 0 | 0.636363636 | 1 | # | 20 | Baseline | ancom | 51 |
| # | sex | 4 | 0 | 0 | 1 | 1 | # | 20 | Baseline | ancom | 51 |
| # | time | 5 | 4 | 0 | 0.555555556 | 1 | # | 20 | Baseline | ancom | 51 |
| # | interaction | 0 | 0 | 4 | 0 | 0 | # | 20 | Baseline | ancom | 51 |
| # | disease | 7 | 54 | 0 | 0.114754098 | 1 | # | 20 | Baseline | aldex2 | 51 |
| # | sex | 4 | 16 | 0 | 0.2 | 1 | # | 20 | Baseline | aldex2 | 51 |
| # | time | 5 | 33 | 0 | 0.131578947 | 1 | # | 20 | Baseline | aldex2 | 51 |
| # | interaction | 0 | 0 | 4 | 0 | 0 | # | 20 | Baseline | aldex2 | 51 |
| # | disease | 7 | 1 | 0 | 0.875 | 1 | # | 20 | Small | paralle | 51 |
| # | sex | 4 | 0 | 0 | 1 | 1 | # | 20 | Small | paralle | 51 |
| # | time | 5 | 0 | 0 | 1 | 1 | # | 20 | Small | paralle | 51 |
| # | interaction | 4 | 1 | 0 | 0.8 | 1 | # | 20 | Small | paralle | 51 |
| # | disease | 7 | 4 | 0 | 0.636363636 | 1 | # | 20 | Small | ancom | 51 |
| # | sex | 4 | 0 | 0 | 1 | 1 | # | 20 | Small | ancom | 51 |
| # | time | 5 | 4 | 0 | 0.555555556 | 1 | # | 20 | Small | ancom | 51 |
| # | interaction | 0 | 0 | 4 | 0 | 0 | # | 20 | Small | ancom | 51 |
| # | disease | 7 | 6 | 0 | 0.538461538 | 1 | # | 20 | Small | aldex2 | 51 |
| # | sex | 4 | 0 | 0 | 1 | 1 | # | 20 | Small | aldex2 | 51 |
| # | time | 5 | 5 | 0 | 0.5 | 1 | # | 20 | Small | aldex2 | 51 |
| # | interaction | 0 | 0 | 4 | 0 | 0 | # | 20 | Small | aldex2 | 51 |
| # | disease | 7 | 9 | 0 | 0.4375 | 1 | # | 20 | Baseline | paralle | 51 |
| # | sex | 4 | 8 | 0 | 0.333333333 | 1 | # | 20 | Baseline | paralle | 51 |

| | | | | | | | | | | | |
|---|---|---|---|---|---|---|---|---|---|---|---|
| # | time | 5 | 4 | 0 | 0.555555556 | 1 | # | 20 | Baseline | paralle | 51 |
| # | interaction | 4 | 7 | 0 | 0.363636364 | 1 | # | 20 | Baseline | paralle | 51 |
| # | disease | 7 | 4 | 0 | 0.636363636 | 1 | # | 20 | Baseline | ancom | 51 |
| # | sex | 4 | 0 | 0 | 1 | 1 | # | 20 | Baseline | ancom | 51 |
| # | time | 5 | 4 | 0 | 0.555555556 | 1 | # | 20 | Baseline | ancom | 51 |
| # | interaction | 0 | 0 | 4 | 0 | 0 | # | 20 | Baseline | ancom | 51 |
| # | disease | 7 | 69 | 0 | 0.092105263 | 1 | # | 20 | Baseline | aldex2 | 51 |
| # | sex | 4 | 21 | 0 | 0.16 | 1 | # | 20 | Baseline | aldex2 | 51 |
| # | time | 5 | 49 | 0 | 0.092592593 | 1 | # | 20 | Baseline | aldex2 | 51 |
| # | interaction | 0 | 0 | 4 | 0 | 0 | # | 20 | Baseline | aldex2 | 51 |
| # | disease | 7 | 1 | 0 | 0.875 | 1 | # | 20 | Small | paralle | 51 |
| # | sex | 4 | 0 | 0 | 1 | 1 | # | 20 | Small | paralle | 51 |
| # | time | 5 | 1 | 0 | 0.833333333 | 1 | # | 20 | Small | paralle | 51 |
| # | interaction | 4 | 0 | 0 | 1 | 1 | # | 20 | Small | paralle | 51 |
| # | disease | 7 | 4 | 0 | 0.636363636 | 1 | # | 20 | Small | ancom | 51 |
| # | sex | 4 | 0 | 0 | 1 | 1 | # | 20 | Small | ancom | 51 |
| # | time | 5 | 4 | 0 | 0.555555556 | 1 | # | 20 | Small | ancom | 51 |
| # | interaction | 0 | 0 | 4 | 0 | 0 | # | 20 | Small | ancom | 51 |
| # | disease | 7 | 11 | 0 | 0.388888889 | 1 | # | 20 | Small | aldex2 | 51 |
| # | sex | 4 | 1 | 0 | 0.8 | 1 | # | 20 | Small | aldex2 | 51 |
| # | time | 5 | 9 | 0 | 0.357142857 | 1 | # | 20 | Small | aldex2 | 51 |
| # | interaction | 0 | 0 | 4 | 0 | 0 | # | 20 | Small | aldex2 | 51 |
| # | disease | 4 | 2 | 0 | 0.666666667 | 1 | # | 10 | Baseline | paralle | 52 |
| # | sex | 2 | 2 | 0 | 0.5 | 1 | # | 10 | Baseline | paralle | 52 |
| # | time | 2 | 6 | 0 | 0.25 | 1 | # | 10 | Baseline | paralle | 52 |
| # | interaction | 2 | 4 | 0 | 0.333333333 | 1 | # | 10 | Baseline | paralle | 52 |
| # | disease | 4 | 1 | 0 | 0.8 | 1 | # | 10 | Baseline | ancom | 52 |
| # | sex | 2 | 0 | 0 | 1 | 1 | # | 10 | Baseline | ancom | 52 |
| # | time | 2 | 0 | 0 | 1 | 1 | # | 10 | Baseline | ancom | 52 |
| # | interaction | 0 | 0 | 2 | 0 | 0 | # | 10 | Baseline | ancom | 52 |
| # | disease | 4 | 10 | 0 | 0.285714286 | 1 | # | 10 | Baseline | aldex2 | 52 |
| # | sex | 2 | 2 | 0 | 0.5 | 1 | # | 10 | Baseline | aldex2 | 52 |
| # | time | 2 | 6 | 0 | 0.25 | 1 | # | 10 | Baseline | aldex2 | 52 |
| # | interaction | 0 | 0 | 2 | 0 | 0 | # | 10 | Baseline | aldex2 | 52 |
| # | disease | 4 | 0 | 0 | 1 | 1 | # | 10 | Small | paralle | 52 |
| # | sex | 2 | 1 | 0 | 0.666666667 | 1 | # | 10 | Small | paralle | 52 |
| # | time | 2 | 0 | 0 | 1 | 1 | # | 10 | Small | paralle | 52 |
| # | interaction | 0 | 0 | 2 | 0 | 0 | # | 10 | Small | paralle | 52 |
| # | disease | 4 | 1 | 0 | 0.8 | 1 | # | 10 | Small | ancom | 52 |
| # | sex | 2 | 0 | 0 | 1 | 1 | # | 10 | Small | ancom | 52 |
| # | time | 2 | 0 | 0 | 1 | 1 | # | 10 | Small | ancom | 52 |
| # | interaction | 0 | 0 | 2 | 0 | 0 | # | 10 | Small | ancom | 52 |
| # | disease | 4 | 1 | 0 | 0.8 | 1 | # | 10 | Small | aldex2 | 52 |
| # | sex | 2 | 0 | 0 | 1 | 1 | # | 10 | Small | aldex2 | 52 |
| # | time | 2 | 1 | 0 | 0.666666667 | 1 | # | 10 | Small | aldex2 | 52 |
| # | interaction | 0 | 0 | 2 | 0 | 0 | # | 10 | Small | aldex2 | 52 |
| # | disease | 4 | 6 | 0 | 0.4 | 1 | # | 10 | Baseline | paralle | 52 |
| # | sex | 2 | 4 | 0 | 0.333333333 | 1 | # | 10 | Baseline | paralle | 52 |
| # | time | 2 | 5 | 0 | 0.285714286 | 1 | # | 10 | Baseline | paralle | 52 |
| # | interaction | 2 | 8 | 0 | 0.2 | 1 | # | 10 | Baseline | paralle | 52 |
| # | disease | 4 | 2 | 0 | 0.666666667 | 1 | # | 10 | Baseline | ancom | 52 |
| # | sex | 2 | 0 | 0 | 1 | 1 | # | 10 | Baseline | ancom | 52 |
| # | time | 2 | 1 | 0 | 0.666666667 | 1 | # | 10 | Baseline | ancom | 52 |
| # | interaction | 0 | 0 | 2 | 0 | 0 | # | 10 | Baseline | ancom | 52 |
| # | disease | 4 | 15 | 0 | 0.210526316 | 1 | # | 10 | Baseline | aldex2 | 52 |
| # | sex | 2 | 5 | 0 | 0.285714286 | 1 | # | 10 | Baseline | aldex2 | 52 |
| # | time | 2 | 7 | 0 | 0.222222222 | 1 | # | 10 | Baseline | aldex2 | 52 |
| # | interaction | 0 | 0 | 2 | 0 | 0 | # | 10 | Baseline | aldex2 | 52 |
| # | disease | 4 | 0 | 0 | 1 | 1 | # | 10 | Small | paralle | 52 |
| # | sex | 2 | 0 | 0 | 1 | 1 | # | 10 | Small | paralle | 52 |
| # | time | 2 | 0 | 0 | 1 | 1 | # | 10 | Small | paralle | 52 |
| # | interaction | 2 | 0 | 0 | 1 | 1 | # | 10 | Small | paralle | 52 |
| # | disease | 4 | 2 | 0 | 0.666666667 | 1 | # | 10 | Small | ancom | 52 |
| # | sex | 2 | 0 | 0 | 1 | 1 | # | 10 | Small | ancom | 52 |
| # | time | 2 | 1 | 0 | 0.666666667 | 1 | # | 10 | Small | ancom | 52 |
| # | interaction | 0 | 0 | 2 | 0 | 0 | # | 10 | Small | ancom | 52 |
| # | disease | 4 | 1 | 0 | 0.8 | 1 | # | 10 | Small | aldex2 | 52 |
| # | sex | 2 | 0 | 0 | 1 | 1 | # | 10 | Small | aldex2 | 52 |
| # | time | 2 | 1 | 0 | 0.666666667 | 1 | # | 10 | Small | aldex2 | 52 |
| # | interaction | 0 | 0 | 2 | 0 | 0 | # | 10 | Small | aldex2 | 52 |
| # | disease | 4 | 2 | 0 | 0.666666667 | 1 | # | 10 | Baseline | paralle | 52 |
| # | sex | 2 | 1 | 0 | 0.666666667 | 1 | # | 10 | Baseline | paralle | 52 |
| # | time | 2 | 3 | 0 | 0.4 | 1 | # | 10 | Baseline | paralle | 52 |
| # | interaction | 2 | 5 | 0 | 0.285714286 | 1 | # | 10 | Baseline | paralle | 52 |
| # | disease | 4 | 2 | 0 | 0.666666667 | 1 | # | 10 | Baseline | ancom | 52 |
| # | sex | 2 | 0 | 0 | 1 | 1 | # | 10 | Baseline | ancom | 52 |
| # | time | 2 | 2 | 0 | 0.5 | 1 | # | 10 | Baseline | ancom | 52 |
| # | interaction | 0 | 0 | 2 | 0 | 0 | # | 10 | Baseline | ancom | 52 |
| # | disease | 4 | 18 | 0 | 0.181818182 | 1 | # | 10 | Baseline | aldex2 | 52 |
| # | sex | 2 | 5 | 0 | 0.285714286 | 1 | # | 10 | Baseline | aldex2 | 52 |
| # | time | 2 | 9 | 0 | 0.181818182 | 1 | # | 10 | Baseline | aldex2 | 52 |
| # | interaction | 0 | 0 | 2 | 0 | 0 | # | 10 | Baseline | aldex2 | 52 |
| # | disease | 4 | 0 | 0 | 1 | 1 | # | 10 | Small | paralle | 52 |
| # | sex | 2 | 0 | 0 | 1 | 1 | # | 10 | Small | paralle | 52 |
| # | time | 2 | 0 | 0 | 1 | 1 | # | 10 | Small | paralle | 52 |
| # | interaction | 2 | 0 | 0 | 1 | 1 | # | 10 | Small | paralle | 52 |
| # | disease | 4 | 2 | 0 | 0.666666667 | 1 | # | 10 | Small | ancom | 52 |
| # | sex | 2 | 0 | 0 | 1 | 1 | # | 10 | Small | ancom | 52 |
| # | time | 2 | 2 | 0 | 0.5 | 1 | # | 10 | Small | ancom | 52 |
| # | interaction | 0 | 0 | 2 | 0 | 0 | # | 10 | Small | ancom | 52 |
| # | disease | 4 | 2 | 0 | 0.666666667 | 1 | # | 10 | Small | aldex2 | 52 |
| # | sex | 2 | 0 | 0 | 1 | 1 | # | 10 | Small | aldex2 | 52 |
| # | time | 2 | 1 | 0 | 0.666666667 | 1 | # | 10 | Small | aldex2 | 52 |
| # | interaction | 0 | 0 | 2 | 0 | 0 | # | 10 | Small | aldex2 | 52 |
| # | disease | 4 | 2 | 0 | 0.666666667 | 1 | # | 10 | Baseline | paralle | 52 |
| # | sex | 2 | 1 | 0 | 0.666666667 | 1 | # | 10 | Baseline | paralle | 52 |
| # | time | 2 | 1 | 0 | 0.666666667 | 1 | # | 10 | Baseline | paralle | 52 |
| # | interaction | 2 | 3 | 0 | 0.4 | 1 | # | 10 | Baseline | paralle | 52 |
| # | disease | 4 | 2 | 0 | 0.666666667 | 1 | # | 10 | Baseline | ancom | 52 |
| # | sex | 2 | 0 | 0 | 1 | 1 | # | 10 | Baseline | ancom | 52 |
| # | time | 2 | 2 | 0 | 0.5 | 1 | # | 10 | Baseline | ancom | 52 |
| # | interaction | 0 | 0 | 2 | 0 | 0 | # | 10 | Baseline | ancom | 52 |
| # | disease | 4 | 23 | 0 | 0.148148148 | 1 | # | 10 | Baseline | aldex2 | 52 |
| # | sex | 2 | 7 | 0 | 0.222222222 | 1 | # | 10 | Baseline | aldex2 | 52 |
| # | time | 2 | 14 | 0 | 0.125 | 1 | # | 10 | Baseline | aldex2 | 52 |
| # | interaction | 0 | 0 | 2 | 0 | 0 | # | 10 | Baseline | aldex2 | 52 |
| # | disease | 4 | 0 | 0 | 1 | 1 | # | 10 | Small | paralle | 52 |
| # | sex | 2 | 0 | 0 | 1 | 1 | # | 10 | Small | paralle | 52 |
| # | time | 2 | 0 | 0 | 1 | 1 | # | 10 | Small | paralle | 52 |
| # | interaction | 2 | 1 | 0 | 0.666666667 | 1 | # | 10 | Small | paralle | 52 |
| # | disease | 4 | 2 | 0 | 0.666666667 | 1 | # | 10 | Small | ancom | 52 |
| # | sex | 2 | 0 | 0 | 1 | 1 | # | 10 | Small | ancom | 52 |
| # | time | 2 | 2 | 0 | 0.5 | 1 | # | 10 | Small | ancom | 52 |
| # | interaction | 0 | 0 | 2 | 0 | 0 | # | 10 | Small | ancom | 52 |
| # | disease | 4 | 2 | 0 | 0.666666667 | 1 | # | 10 | Small | aldex2 | 52 |
| # | sex | 2 | 0 | 0 | 1 | 1 | # | 10 | Small | aldex2 | 52 |
| # | time | 2 | 2 | 0 | 0.5 | 1 | # | 10 | Small | aldex2 | 52 |
| # | interaction | 0 | 0 | 2 | 0 | 0 | # | 10 | Small | aldex2 | 52 |
| # | disease | 7 | 17 | 0 | 0.291666667 | 1 | # | 20 | Baseline | paralle | 52 |
| # | sex | 4 | 4 | 0 | 0.5 | 1 | # | 20 | Baseline | paralle | 52 |
| # | time | 5 | 9 | 0 | 0.357142857 | 1 | # | 20 | Baseline | paralle | 52 |
| # | interaction | 4 | 11 | 0 | 0.266666667 | 1 | # | 20 | Baseline | paralle | 52 |
| # | disease | 7 | 3 | 0 | 0.7 | 1 | # | 20 | Baseline | ancom | 52 |
| # | sex | 4 | 0 | 0 | 1 | 1 | # | 20 | Baseline | ancom | 52 |
| # | time | 5 | 1 | 0 | 0.833333333 | 1 | # | 20 | Baseline | ancom | 52 |
| # | interaction | 0 | 0 | 4 | 0 | 0 | # | 20 | Baseline | ancom | 52 |
| # | disease | 7 | 18 | 0 | 0.28 | 1 | # | 20 | Baseline | aldex2 | 52 |
| # | sex | 4 | 9 | 0 | 0.307692308 | 1 | # | 20 | Baseline | aldex2 | 52 |
| # | time | 5 | 16 | 0 | 0.238095238 | 1 | # | 20 | Baseline | aldex2 | 52 |
| # | interaction | 0 | 0 | 4 | 0 | 0 | # | 20 | Baseline | aldex2 | 52 |
| # | disease | 7 | 0 | 0 | 1 | 1 | # | 20 | Small | paralle | 52 |
| # | sex | 4 | 0 | 0 | 1 | 1 | # | 20 | Small | paralle | 52 |
| # | time | 5 | 0 | 0 | 1 | 1 | # | 20 | Small | paralle | 52 |
| # | interaction | 1 | 0 | 3 | 1 | 0.25 | # | 20 | Small | paralle | 52 |
| # | disease | 7 | 3 | 0 | 0.7 | 1 | # | 20 | Small | ancom | 52 |
| # | sex | 4 | 0 | 0 | 1 | 1 | # | 20 | Small | ancom | 52 |
| # | time | 5 | 1 | 0 | 0.833333333 | 1 | # | 20 | Small | ancom | 52 |
| # | interaction | 0 | 0 | 4 | 0 | 0 | # | 20 | Small | ancom | 52 |
| # | disease | 7 | 1 | 0 | 0.875 | 1 | # | 20 | Small | aldex2 | 52 |
| # | sex | 4 | 0 | 0 | 1 | 1 | # | 20 | Small | aldex2 | 52 |
| # | time | 5 | 0 | 0 | 1 | 1 | # | 20 | Small | aldex2 | 52 |
| # | interaction | 0 | 0 | 4 | 0 | 0 | # | 20 | Small | aldex2 | 52 |
| # | disease | 7 | 8 | 0 | 0.466666667 | 1 | # | 20 | Baseline | paralle | 52 |
| # | sex | 4 | 7 | 0 | 0.363636364 | 1 | # | 20 | Baseline | paralle | 52 |
| # | time | 5 | 7 | 0 | 0.416666667 | 1 | # | 20 | Baseline | paralle | 52 |
| # | interaction | 4 | 7 | 0 | 0.363636364 | 1 | # | 20 | Baseline | paralle | 52 |
| # | disease | 7 | 3 | 0 | 0.7 | 1 | # | 20 | Baseline | ancom | 52 |
| # | sex | 4 | 0 | 0 | 1 | 1 | # | 20 | Baseline | ancom | 52 |
| # | time | 5 | 3 | 0 | 0.625 | 1 | # | 20 | Baseline | ancom | 52 |
| # | interaction | 0 | 0 | 4 | 0 | 0 | # | 20 | Baseline | ancom | 52 |
| # | disease | 7 | 28 | 0 | 0.2 | 1 | # | 20 | Baseline | aldex2 | 52 |
| # | sex | 4 | 14 | 0 | 0.222222222 | 1 | # | 20 | Baseline | aldex2 | 52 |
| # | time | 5 | 19 | 0 | 0.208333333 | 1 | # | 20 | Baseline | aldex2 | 52 |
| # | interaction | 0 | 0 | 4 | 0 | 0 | # | 20 | Baseline | aldex2 | 52 |
| # | disease | 7 | 0 | 0 | 1 | 1 | # | 20 | Small | paralle | 52 |
| # | sex | 4 | 0 | 0 | 1 | 1 | # | 20 | Small | paralle | 52 |
| # | time | 5 | 0 | 0 | 1 | 1 | # | 20 | Small | paralle | 52 |
| # | interaction | 2 | 1 | 2 | 0.666666667 | 0.5 | # | 20 | Small | paralle | 52 |
| # | disease | 7 | 3 | 0 | 0.7 | 1 | # | 20 | Small | ancom | 52 |
| # | sex | 4 | 0 | 0 | 1 | 1 | # | 20 | Small | ancom | 52 |
| # | time | 5 | 3 | 0 | 0.625 | 1 | # | 20 | Small | ancom | 52 |
| # | interaction | 0 | 0 | 4 | 0 | 0 | # | 20 | Small | ancom | 52 |
| # | disease | 7 | 4 | 0 | 0.636363636 | 1 | # | 20 | Small | aldex2 | 52 |
| # | sex | 4 | 0 | 0 | 1 | 1 | # | 20 | Small | aldex2 | 52 |
| # | time | 5 | 1 | 0 | 0.833333333 | 1 | # | 20 | Small | aldex2 | 52 |
| # | interaction | 0 | 0 | 4 | 0 | 0 | # | 20 | Small | aldex2 | 52 |
| # | disease | 7 | 10 | 0 | 0.411764706 | 1 | # | 20 | Baseline | paralle | 52 |
| # | sex | 4 | 3 | 0 | 0.571428571 | 1 | # | 20 | Baseline | paralle | 52 |
| # | time | 5 | 4 | 0 | 0.555555556 | 1 | # | 20 | Baseline | paralle | 52 |
| # | interaction | 4 | 5 | 0 | 0.444444444 | 1 | # | 20 | Baseline | paralle | 52 |
| # | disease | 7 | 4 | 0 | 0.636363636 | 1 | # | 20 | Baseline | ancom | 52 |
| # | sex | 4 | 0 | 0 | 1 | 1 | # | 20 | Baseline | ancom | 52 |
| # | time | 5 | 4 | 0 | 0.555555556 | 1 | # | 20 | Baseline | ancom | 52 |
| # | interaction | 0 | 0 | 4 | 0 | 0 | # | 20 | Baseline | ancom | 52 |
| # | disease | 7 | 35 | 0 | 0.166666667 | 1 | # | 20 | Baseline | aldex2 | 52 |
| # | sex | 4 | 22 | 0 | 0.153846154 | 1 | # | 20 | Baseline | aldex2 | 52 |
| # | time | 5 | 26 | 0 | 0.161290323 | 1 | # | 20 | Baseline | aldex2 | 52 |
| # | interaction | 0 | 0 | 4 | 0 | 0 | # | 20 | Baseline | aldex2 | 52 |
| # | disease | 7 | 0 | 0 | 1 | 1 | # | 20 | Small | paralle | 52 |
| # | sex | 4 | 1 | 0 | 0.8 | 1 | # | 20 | Small | paralle | 52 |
| # | time | 5 | 0 | 0 | 1 | 1 | # | 20 | Small | paralle | 52 |
| # | interaction | 4 | 0 | 0 | 1 | 1 | # | 20 | Small | paralle | 52 |
| # | disease | 7 | 4 | 0 | 0.636363636 | 1 | # | 20 | Small | ancom | 52 |
| # | sex | 4 | 0 | 0 | 1 | 1 | # | 20 | Small | ancom | 52 |
| # | time | 5 | 4 | 0 | 0.555555556 | 1 | # | 20 | Small | ancom | 52 |
| # | interaction | 0 | 0 | 4 | 0 | 0 | # | 20 | Small | ancom | 52 |
| # | disease | 7 | 5 | 0 | 0.583333333 | 1 | # | 20 | Small | aldex2 | 52 |
| # | sex | 4 | 2 | 0 | 0.666666667 | 1 | # | 20 | Small | aldex2 | 52 |
| # | time | 5 | 3 | 0 | 0.625 | 1 | # | 20 | Small | aldex2 | 52 |
| # | interaction | 0 | 0 | 4 | 0 | 0 | # | 20 | Small | aldex2 | 52 |
| # | disease | 7 | 8 | 0 | 0.466666667 | 1 | # | 20 | Baseline | paralle | 52 |
| # | sex | 4 | 3 | 0 | 0.571428571 | 1 | # | 20 | Baseline | paralle | 52 |
| # | time | 5 | 3 | 0 | 0.625 | 1 | # | 20 | Baseline | paralle | 52 |
| # | interaction | 4 | 4 | 0 | 0.5 | 1 | # | 20 | Baseline | paralle | 52 |
| # | disease | 7 | 4 | 0 | 0.636363636 | 1 | # | 20 | Baseline | ancom | 52 |
| # | sex | 4 | 0 | 0 | 1 | 1 | # | 20 | Baseline | ancom | 52 |
| # | time | 5 | 4 | 0 | 0.555555556 | 1 | # | 20 | Baseline | ancom | 52 |
| # | interaction | 0 | 0 | 4 | 0 | 0 | # | 20 | Baseline | ancom | 52 |
| # | disease | 7 | 56 | 0 | 0.111111111 | 1 | # | 20 | Baseline | aldex2 | 52 |
| # | sex | 4 | 22 | 0 | 0.153846154 | 1 | # | 20 | Baseline | aldex2 | 52 |
| # | time | 5 | 44 | 0 | 0.102040816 | 1 | # | 20 | Baseline | aldex2 | 52 |
| # | interaction | 0 | 0 | 4 | 0 | 0 | # | 20 | Baseline | aldex2 | 52 |
| # | disease | 7 | 0 | 0 | 1 | 1 | # | 20 | Small | paralle | 52 |
| # | sex | 4 | 0 | 0 | 1 | 1 | # | 20 | Small | paralle | 52 |
| # | time | 5 | 0 | 0 | 1 | 1 | # | 20 | Small | paralle | 52 |
| # | interaction | 4 | 0 | 0 | 1 | 1 | # | 20 | Small | paralle | 52 |
| # | disease | 7 | 4 | 0 | 0.636363636 | 1 | # | 20 | Small | ancom | 52 |
| # | sex | 4 | 0 | 0 | 1 | 1 | # | 20 | Small | ancom | 52 |
| # | time | 5 | 4 | 0 | 0.555555556 | 1 | # | 20 | Small | ancom | 52 |
| # | interaction | 0 | 0 | 4 | 0 | 0 | # | 20 | Small | ancom | 52 |
| # | disease | 7 | 6 | 0 | 0.538461538 | 1 | # | 20 | Small | aldex2 | 52 |
| # | sex | 4 | 3 | 0 | 0.571428571 | 1 | # | 20 | Small | aldex2 | 52 |
| # | time | 5 | 5 | 0 | 0.5 | 1 | # | 20 | Small | aldex2 | 52 |
| # | interaction | 0 | 0 | 4 | 0 | 0 | # | 20 | Small | aldex2 | 52 |
| # | disease | 4 | 8 | 0 | 0.333333333 | 1 | # | 10 | Baseline | paralle | 53 |
| # | sex | 2 | 7 | 0 | 0.222222222 | 1 | # | 10 | Baseline | paralle | 53 |
| # | time | 2 | 9 | 0 | 0.181818182 | 1 | # | 10 | Baseline | paralle | 53 |
| # | interaction | 2 | 7 | 0 | 0.222222222 | 1 | # | 10 | Baseline | paralle | 53 |
| # | disease | 4 | 1 | 0 | 0.8 | 1 | # | 10 | Baseline | ancom | 53 |
| # | sex | 2 | 0 | 0 | 1 | 1 | # | 10 | Baseline | ancom | 53 |
| # | time | 2 | 0 | 0 | 1 | 1 | # | 10 | Baseline | ancom | 53 |
| # | interaction | 0 | 0 | 2 | 0 | 0 | # | 10 | Baseline | ancom | 53 |
| # | disease | 4 | 14 | 0 | 0.222222222 | 1 | # | 10 | Baseline | aldex2 | 53 |
| # | sex | 2 | 4 | 0 | 0.333333333 | 1 | # | 10 | Baseline | aldex2 | 53 |
| # | time | 2 | 7 | 0 | 0.222222222 | 1 | # | 10 | Baseline | aldex2 | 53 |
| # | interaction | 0 | 0 | 2 | 0 | 0 | # | 10 | Baseline | aldex2 | 53 |
| # | disease | 4 | 0 | 0 | 1 | 1 | # | 10 | Small | paralle | 53 |
| # | sex | 2 | 1 | 0 | 0.666666667 | 1 | # | 10 | Small | paralle | 53 |
| # | time | 2 | 1 | 0 | 0.666666667 | 1 | # | 10 | Small | paralle | 53 |
| # | interaction | 2 | 0 | 0 | 1 | 1 | # | 10 | Small | paralle | 53 |
| # | disease | 4 | 1 | 0 | 0.8 | 1 | # | 10 | Small | ancom | 53 |
| # | sex | 2 | 0 | 0 | 1 | 1 | # | 10 | Small | ancom | 53 |
| # | time | 2 | 0 | 0 | 1 | 1 | # | 10 | Small | ancom | 53 |
| # | interaction | 0 | 0 | 2 | 0 | 0 | # | 10 | Small | ancom | 53 |
| # | disease | 4 | 1 | 0 | 0.8 | 1 | # | 10 | Small | aldex2 | 53 |
| # | sex | 2 | 0 | 0 | 1 | 1 | # | 10 | Small | aldex2 | 53 |
| # | time | 2 | 0 | 0 | 1 | 1 | # | 10 | Small | aldex2 | 53 |
| # | interaction | 0 | 0 | 2 | 0 | 0 | # | 10 | Small | aldex2 | 53 |
| # | disease | 4 | 10 | 0 | 0.285714286 | 1 | # | 10 | Baseline | paralle | 53 |
| # | sex | 2 | 3 | 0 | 0.4 | 1 | # | 10 | Baseline | paralle | 53 |
| # | time | 2 | 4 | 0 | 0.333333333 | 1 | # | 10 | Baseline | paralle | 53 |
| # | interaction | 2 | 6 | 0 | 0.25 | 1 | # | 10 | Baseline | paralle | 53 |
| # | disease | 4 | 2 | 0 | 0.666666667 | 1 | # | 10 | Baseline | ancom | 53 |
| # | sex | 2 | 0 | 0 | 1 | 1 | # | 10 | Baseline | ancom | 53 |
| # | time | 2 | 1 | 0 | 0.666666667 | 1 | # | 10 | Baseline | ancom | 53 |
| # | interaction | 0 | 0 | 2 | 0 | 0 | # | 10 | Baseline | ancom | 53 |
| # | disease | 4 | 16 | 0 | 0.2 | 1 | # | 10 | Baseline | aldex2 | 53 |
| # | sex | 2 | 7 | 0 | 0.222222222 | 1 | # | 10 | Baseline | aldex2 | 53 |
| # | time | 2 | 7 | 0 | 0.222222222 | 1 | # | 10 | Baseline | aldex2 | 53 |
| # | interaction | 0 | 0 | 2 | 0 | 0 | # | 10 | Baseline | aldex2 | 53 |
| # | disease | 4 | 0 | 0 | 1 | 1 | # | 10 | Small | paralle | 53 |
| # | sex | 2 | 0 | 0 | 1 | 1 | # | 10 | Small | paralle | 53 |
| # | time | 2 | 0 | 0 | 1 | 1 | # | 10 | Small | paralle | 53 |
| # | interaction | 2 | 0 | 0 | 1 | 1 | # | 10 | Small | paralle | 53 |
| # | disease | 4 | 2 | 0 | 0.666666667 | 1 | # | 10 | Small | ancom | 53 |
| # | sex | 2 | 0 | 0 | 1 | 1 | # | 10 | Small | ancom | 53 |
| # | time | 2 | 0 | 0 | 1 | 1 | # | 10 | Small | ancom | 53 |
| # | interaction | 0 | 0 | 2 | 0 | 0 | # | 10 | Small | ancom | 53 |
| # | disease | 4 | 1 | 0 | 0.8 | 1 | # | 10 | Small | aldex2 | 53 |
| # | sex | 2 | 0 | 0 | 1 | 1 | # | 10 | Small | aldex2 | 53 |
| # | time | 2 | 1 | 0 | 0.666666667 | 1 | # | 10 | Small | aldex2 | 53 |
| # | interaction | 0 | 0 | 2 | 0 | 0 | # | 10 | Small | aldex2 | 53 |
| # | disease | 4 | 3 | 0 | 0.571428571 | 1 | # | 10 | Baseline | paralle | 53 |
| # | sex | 2 | 6 | 0 | 0.25 | 1 | # | 10 | Baseline | paralle | 53 |
| # | time | 2 | 3 | 0 | 0.4 | 1 | # | 10 | Baseline | paralle | 53 |
| # | interaction | 2 | 3 | 0 | 0.4 | 1 | # | 10 | Baseline | paralle | 53 |
| # | disease | 4 | 2 | 0 | 0.666666667 | 1 | # | 10 | Baseline | ancom | 53 |
| # | sex | 2 | 0 | 0 | 1 | 1 | # | 10 | Baseline | ancom | 53 |
| # | time | 2 | 2 | 0 | 0.5 | 1 | # | 10 | Baseline | ancom | 53 |
| # | interaction | 0 | 0 | 2 | 0 | 0 | # | 10 | Baseline | ancom | 53 |
| # | disease | 4 | 29 | 0 | 0.121212121 | 1 | # | 10 | Baseline | aldex2 | 53 |
| # | sex | 2 | 6 | 0 | 0.25 | 1 | # | 10 | Baseline | aldex2 | 53 |
| # | time | 2 | 7 | 0 | 0.222222222 | 1 | # | 10 | Baseline | aldex2 | 53 |
| # | interaction | 0 | 0 | 2 | 0 | 0 | # | 10 | Baseline | aldex2 | 53 |
| # | disease | 4 | 1 | 0 | 0.8 | 1 | # | 10 | Small | paralle | 53 |
| # | sex | 2 | 0 | 0 | 1 | 1 | # | 10 | Small | paralle | 53 |
| # | time | 2 | 0 | 0 | 1 | 1 | # | 10 | Small | paralle | 53 |
| # | interaction | 2 | 0 | 0 | 1 | 1 | # | 10 | Small | paralle | 53 |
| # | disease | 4 | 2 | 0 | 0.666666667 | 1 | # | 10 | Small | ancom | 53 |
| # | sex | 2 | 0 | 0 | 1 | 1 | # | 10 | Small | ancom | 53 |
| # | time | 2 | 2 | 0 | 0.5 | 1 | # | 10 | Small | ancom | 53 |
| # | interaction | 0 | 0 | 2 | 0 | 0 | # | 10 | Small | ancom | 53 |
| # | disease | 4 | 2 | 0 | 0.666666667 | 1 | # | 10 | Small | aldex2 | 53 |
| # | sex | 2 | 0 | 0 | 1 | 1 | # | 10 | Small | aldex2 | 53 |
| # | time | 2 | 2 | 0 | 0.5 | 1 | # | 10 | Small | aldex2 | 53 |
| # | interaction | 0 | 0 | 2 | 0 | 0 | # | 10 | Small | aldex2 | 53 |
| # | disease | 4 | 6 | 0 | 0.4 | 1 | # | 10 | Baseline | paralle | 53 |
| # | sex | 2 | 4 | 0 | 0.333333333 | 1 | # | 10 | Baseline | paralle | 53 |
| # | time | 2 | 3 | 0 | 0.4 | 1 | # | 10 | Baseline | paralle | 53 |
| # | interaction | 2 | 3 | 0 | 0.4 | 1 | # | 10 | Baseline | paralle | 53 |
| # | disease | 4 | 2 | 0 | 0.666666667 | 1 | # | 10 | Baseline | ancom | 53 |
| # | sex | 2 | 0 | 0 | 1 | 1 | # | 10 | Baseline | ancom | 53 |
| # | time | 2 | 2 | 0 | 0.5 | 1 | # | 10 | Baseline | ancom | 53 |
| # | interaction | 0 | 0 | 2 | 0 | 0 | # | 10 | Baseline | ancom | 53 |
| # | disease | 4 | 34 | 0 | 0.105263158 | 1 | # | 10 | Baseline | aldex2 | 53 |
| # | sex | 2 | 7 | 0 | 0.222222222 | 1 | # | 10 | Baseline | aldex2 | 53 |
| # | time | 2 | 11 | 0 | 0.153846154 | 1 | # | 10 | Baseline | aldex2 | 53 |
| # | interaction | 0 | 0 | 2 | 0 | 0 | # | 10 | Baseline | aldex2 | 53 |
| # | disease | 4 | 0 | 0 | 1 | 1 | # | 10 | Small | paralle | 53 |
| # | sex | 2 | 0 | 0 | 1 | 1 | # | 10 | Small | paralle | 53 |
| # | time | 2 | 0 | 0 | 1 | 1 | # | 10 | Small | paralle | 53 |
| # | interaction | 2 | 0 | 0 | 1 | 1 | # | 10 | Small | paralle | 53 |
| # | disease | 4 | 2 | 0 | 0.666666667 | 1 | # | 10 | Small | ancom | 53 |
| # | sex | 2 | 0 | 0 | 1 | 1 | # | 10 | Small | ancom | 53 |
| # | time | 2 | 2 | 0 | 0.5 | 1 | # | 10 | Small | ancom | 53 |
| # | interaction | 0 | 0 | 2 | 0 | 0 | # | 10 | Small | ancom | 53 |
| # | disease | 4 | 4 | 0 | 0.5 | 1 | # | 10 | Small | aldex2 | 53 |
| # | sex | 2 | 0 | 0 | 1 | 1 | # | 10 | Small | aldex2 | 53 |
| # | time | 2 | 2 | 0 | 0.5 | 1 | # | 10 | Small | aldex2 | 53 |
| # | interaction | 0 | 0 | 2 | 0 | 0 | # | 10 | Small | aldex2 | 53 |
| # | disease | 7 | 10 | 0 | 0.411764706 | 1 | # | 20 | Baseline | paralle | 53 |
| # | sex | 4 | 12 | 0 | 0.25 | 1 | # | 20 | Baseline | paralle | 53 |
| # | time | 5 | 9 | 0 | 0.357142857 | 1 | # | 20 | Baseline | paralle | 53 |
| # | interaction | 3 | 10 | 1 | 0.230769231 | 0.75 | # | 20 | Baseline | paralle | 53 |
| # | disease | 7 | 2 | 0 | 0.777777778 | 1 | # | 20 | Baseline | ancom | 53 |
| # | sex | 4 | 0 | 0 | 1 | 1 | # | 20 | Baseline | ancom | 53 |
| # | time | 5 | 0 | 0 | 1 | 1 | # | 20 | Baseline | ancom | 53 |
| # | interaction | 0 | 0 | 4 | 0 | 0 | # | 20 | Baseline | ancom | 53 |
| # | disease | 7 | 21 | 0 | 0.25 | 1 | # | 20 | Baseline | aldex2 | 53 |
| # | sex | 4 | 7 | 0 | 0.363636364 | 1 | # | 20 | Baseline | aldex2 | 53 |
| # | time | 5 | 13 | 0 | 0.277777778 | 1 | # | 20 | Baseline | aldex2 | 53 |
| # | interaction | 0 | 0 | 4 | 0 | 0 | # | 20 | Baseline | aldex2 | 53 |
| # | disease | 7 | 0 | 0 | 1 | 1 | # | 20 | Small | paralle | 53 |
| # | sex | 4 | 2 | 0 | 0.666666667 | 1 | # | 20 | Small | paralle | 53 |
| # | time | 5 | 1 | 0 | 0.833333333 | 1 | # | 20 | Small | paralle | 53 |
| # | interaction | 3 | 0 | 1 | 1 | 0.75 | # | 20 | Small | paralle | 53 |
| # | disease | 7 | 1 | 0 | 0.875 | 1 | # | 20 | Small | ancom | 53 |
| # | sex | 4 | 0 | 0 | 1 | 1 | # | 20 | Small | ancom | 53 |
| # | time | 5 | 0 | 0 | 1 | 1 | # | 20 | Small | ancom | 53 |
| # | interaction | 0 | 0 | 4 | 0 | 0 | # | 20 | Small | ancom | 53 |
| # | disease | 7 | 0 | 0 | 1 | 1 | # | 20 | Small | aldex2 | 53 |
| # | sex | 4 | 0 | 0 | 1 | 1 | # | 20 | Small | aldex2 | 53 |
| # | time | 5 | 0 | 0 | 1 | 1 | # | 20 | Small | aldex2 | 53 |
| # | interaction | 0 | 0 | 4 | 0 | 0 | # | 20 | Small | aldex2 | 53 |
| # | disease | 7 | 6 | 0 | 0.538461538 | 1 | # | 20 | Baseline | paralle | 53 |
| # | sex | 4 | 8 | 0 | 0.333333333 | 1 | # | 20 | Baseline | paralle | 53 |
| # | time | 5 | 7 | 0 | 0.416666667 | 1 | # | 20 | Baseline | paralle | 53 |

| | | | | | | | | | | | |
|---|---|---|---|---|---|---|---|---|---|---|---|
| # | interaction | 4 | 4 | 0 | 0.5 | 1 | # | 20 | Baseline | paralle | 53 |
| # | disease | 7 | 4 | 0 | 0.636363636 | 1 | # | 20 | Baseline | ancom | 53 |
| # | sex | 4 | 0 | 0 | 1 | 1 | # | 20 | Baseline | ancom | 53 |
| # | time | 5 | 4 | 0 | 0.555555556 | 1 | # | 20 | Baseline | ancom | 53 |
| # | interaction | 0 | 0 | 4 | 0 | 0 | # | 20 | Baseline | ancom | 53 |
| # | disease | 7 | 39 | 0 | 0.152173913 | 1 | # | 20 | Baseline | aldex2 | 53 |
| # | sex | 4 | 12 | 0 | 0.25 | 1 | # | 20 | Baseline | aldex2 | 53 |
| # | time | 5 | 31 | 0 | 0.138888889 | 1 | # | 20 | Baseline | aldex2 | 53 |
| # | interaction | 0 | 0 | 4 | 0 | 0 | # | 20 | Baseline | aldex2 | 53 |
| # | disease | 7 | 0 | 0 | 1 | 1 | # | 20 | Small | paralle | 53 |
| # | sex | 4 | 0 | 0 | 1 | 1 | # | 20 | Small | paralle | 53 |
| # | time | 5 | 0 | 0 | 1 | 1 | # | 20 | Small | paralle | 53 |
| # | interaction | 3 | 0 | 1 | 1 | 0.75 | # | 20 | Small | paralle | 53 |
| # | disease | 7 | 4 | 0 | 0.636363636 | 1 | # | 20 | Small | ancom | 53 |
| # | sex | 4 | 0 | 0 | 1 | 1 | # | 20 | Small | ancom | 53 |
| # | time | 5 | 4 | 0 | 0.555555556 | 1 | # | 20 | Small | ancom | 53 |
| # | interaction | 0 | 0 | 4 | 0 | 0 | # | 20 | Small | ancom | 53 |
| # | disease | 7 | 6 | 0 | 0.538461538 | 1 | # | 20 | Small | aldex2 | 53 |
| # | sex | 4 | 1 | 0 | 0.8 | 1 | # | 20 | Small | aldex2 | 53 |
| # | time | 5 | 2 | 0 | 0.714285714 | 1 | # | 20 | Small | aldex2 | 53 |
| # | interaction | 0 | 0 | 4 | 0 | 0 | # | 20 | Small | aldex2 | 53 |
| # | disease | 7 | 11 | 0 | 0.388888889 | 1 | # | 20 | Baseline | paralle | 53 |
| # | sex | 4 | 4 | 0 | 0.5 | 1 | # | 20 | Baseline | paralle | 53 |
| # | time | 5 | 3 | 0 | 0.625 | 1 | # | 20 | Baseline | paralle | 53 |
| # | interaction | 4 | 5 | 0 | 0.444444444 | 1 | # | 20 | Baseline | paralle | 53 |
| # | disease | 7 | 4 | 0 | 0.636363636 | 1 | # | 20 | Baseline | ancom | 53 |
| # | sex | 4 | 0 | 0 | 1 | 1 | # | 20 | Baseline | ancom | 53 |
| # | time | 5 | 4 | 0 | 0.555555556 | 1 | # | 20 | Baseline | ancom | 53 |
| # | interaction | 0 | 0 | 4 | 0 | 0 | # | 20 | Baseline | ancom | 53 |
| # | disease | 7 | 51 | 0 | 0.102941176 | 1 | # | 20 | Baseline | aldex2 | 53 |
| # | sex | 4 | 13 | 0 | 0.235294118 | 1 | # | 20 | Baseline | aldex2 | 53 |
| # | time | 5 | 27 | 0 | 0.15625 | 1 | # | 20 | Baseline | aldex2 | 53 |
| # | interaction | 0 | 0 | 4 | 0 | 0 | # | 20 | Baseline | aldex2 | 53 |
| # | disease | 7 | 0 | 0 | 1 | 1 | # | 20 | Small | paralle | 53 |
| # | sex | 4 | 0 | 0 | 1 | 1 | # | 20 | Small | paralle | 53 |
| # | time | 5 | 0 | 0 | 1 | 1 | # | 20 | Small | paralle | 53 |
| # | interaction | 3 | 0 | 1 | 1 | 0.75 | # | 20 | Small | paralle | 53 |
| # | disease | 7 | 4 | 0 | 0.636363636 | 1 | # | 20 | Small | ancom | 53 |
| # | sex | 4 | 0 | 0 | 1 | 1 | # | 20 | Small | ancom | 53 |
| # | time | 5 | 4 | 0 | 0.555555556 | 1 | # | 20 | Small | ancom | 53 |
| # | interaction | 0 | 0 | 4 | 0 | 0 | # | 20 | Small | ancom | 53 |
| # | disease | 7 | 5 | 0 | 0.583333333 | 1 | # | 20 | Small | aldex2 | 53 |
| # | sex | 4 | 0 | 0 | 1 | 1 | # | 20 | Small | aldex2 | 53 |
| # | time | 5 | 3 | 0 | 0.625 | 1 | # | 20 | Small | aldex2 | 53 |
| # | interaction | 0 | 0 | 4 | 0 | 0 | # | 20 | Small | aldex2 | 53 |
| # | disease | 7 | 16 | 0 | 0.304347826 | 1 | # | 20 | Baseline | paralle | 53 |
| # | sex | 4 | 5 | 0 | 0.444444444 | 1 | # | 20 | Baseline | paralle | 53 |
| # | time | 5 | 8 | 0 | 0.384615385 | 1 | # | 20 | Baseline | paralle | 53 |
| # | interaction | 4 | 3 | 0 | 0.571428571 | 1 | # | 20 | Baseline | paralle | 53 |
| # | disease | 7 | 4 | 0 | 0.636363636 | 1 | # | 20 | Baseline | ancom | 53 |
| # | sex | 4 | 0 | 0 | 1 | 1 | # | 20 | Baseline | ancom | 53 |
| # | time | 5 | 4 | 0 | 0.555555556 | 1 | # | 20 | Baseline | ancom | 53 |
| # | interaction | 0 | 0 | 4 | 0 | 0 | # | 20 | Baseline | ancom | 53 |
| # | disease | 7 | 70 | 0 | 0.090909091 | 1 | # | 20 | Baseline | aldex2 | 53 |
| # | sex | 4 | 19 | 0 | 0.173913043 | 1 | # | 20 | Baseline | aldex2 | 53 |
| # | time | 5 | 41 | 0 | 0.108695652 | 1 | # | 20 | Baseline | aldex2 | 53 |
| # | interaction | 0 | 0 | 4 | 0 | 0 | # | 20 | Baseline | aldex2 | 53 |
| # | disease | 7 | 0 | 0 | 1 | 1 | # | 20 | Small | paralle | 53 |
| # | sex | 4 | 0 | 0 | 1 | 1 | # | 20 | Small | paralle | 53 |
| # | time | 5 | 0 | 0 | 1 | 1 | # | 20 | Small | paralle | 53 |
| # | interaction | 3 | 0 | 1 | 1 | 0.75 | # | 20 | Small | paralle | 53 |
| # | disease | 7 | 4 | 0 | 0.636363636 | 1 | # | 20 | Small | ancom | 53 |
| # | sex | 4 | 0 | 0 | 1 | 1 | # | 20 | Small | ancom | 53 |
| # | time | 5 | 4 | 0 | 0.555555556 | 1 | # | 20 | Small | ancom | 53 |
| # | interaction | 0 | 0 | 4 | 0 | 0 | # | 20 | Small | ancom | 53 |
| # | disease | 7 | 10 | 0 | 0.411764706 | 1 | # | 20 | Small | aldex2 | 53 |
| # | sex | 4 | 1 | 0 | 0.8 | 1 | # | 20 | Small | aldex2 | 53 |
| # | time | 5 | 5 | 0 | 0.5 | 1 | # | 20 | Small | aldex2 | 53 |
| # | interaction | 0 | 0 | 4 | 0 | 0 | # | 20 | Small | aldex2 | 53 |
| # | disease | 4 | 7 | 0 | 0.363636364 | 1 | # | 10 | Baseline | paralle | 54 |
| # | sex | 2 | 8 | 0 | 0.2 | 1 | # | 10 | Baseline | paralle | 54 |
| # | time | 2 | 9 | 0 | 0.181818182 | 1 | # | 10 | Baseline | paralle | 54 |
| # | interaction | 2 | 6 | 0 | 0.25 | 1 | # | 10 | Baseline | paralle | 54 |
| # | disease | 4 | 2 | 0 | 0.666666667 | 1 | # | 10 | Baseline | ancom | 54 |
| # | sex | 2 | 0 | 0 | 1 | 1 | # | 10 | Baseline | ancom | 54 |
| # | time | 2 | 0 | 0 | 1 | 1 | # | 10 | Baseline | ancom | 54 |
| # | interaction | 0 | 0 | 2 | 0 | 0 | # | 10 | Baseline | ancom | 54 |
| # | disease | 4 | 11 | 0 | 0.266666667 | 1 | # | 10 | Baseline | aldex2 | 54 |
| # | sex | 2 | 3 | 0 | 0.4 | 1 | # | 10 | Baseline | aldex2 | 54 |
| # | time | 2 | 4 | 0 | 0.333333333 | 1 | # | 10 | Baseline | aldex2 | 54 |
| # | interaction | 0 | 0 | 2 | 0 | 0 | # | 10 | Baseline | aldex2 | 54 |
| # | disease | 4 | 0 | 0 | 1 | 1 | # | 10 | Small | paralle | 54 |
| # | sex | 2 | 1 | 0 | 0.666666667 | 1 | # | 10 | Small | paralle | 54 |
| # | time | 2 | 0 | 0 | 1 | 1 | # | 10 | Small | paralle | 54 |
| # | interaction | 0 | 0 | 2 | 0 | 0 | # | 10 | Small | paralle | 54 |
| # | disease | 4 | 1 | 0 | 0.8 | 1 | # | 10 | Small | ancom | 54 |
| # | sex | 2 | 0 | 0 | 1 | 1 | # | 10 | Small | ancom | 54 |
| # | time | 2 | 0 | 0 | 1 | 1 | # | 10 | Small | ancom | 54 |
| # | interaction | 0 | 0 | 2 | 0 | 0 | # | 10 | Small | ancom | 54 |
| # | disease | 4 | 2 | 0 | 0.666666667 | 1 | # | 10 | Small | aldex2 | 54 |
| # | sex | 2 | 0 | 0 | 1 | 1 | # | 10 | Small | aldex2 | 54 |
| # | time | 2 | 1 | 0 | 0.666666667 | 1 | # | 10 | Small | aldex2 | 54 |
| # | interaction | 0 | 0 | 2 | 0 | 0 | # | 10 | Small | aldex2 | 54 |
| # | disease | 4 | 6 | 0 | 0.4 | 1 | # | 10 | Baseline | paralle | 54 |
| # | sex | 2 | 9 | 0 | 0.181818182 | 1 | # | 10 | Baseline | paralle | 54 |
| # | time | 2 | 5 | 0 | 0.285714286 | 1 | # | 10 | Baseline | paralle | 54 |
| # | interaction | 2 | 9 | 0 | 0.181818182 | 1 | # | 10 | Baseline | paralle | 54 |
| # | disease | 4 | 2 | 0 | 0.666666667 | 1 | # | 10 | Baseline | ancom | 54 |
| # | sex | 2 | 0 | 0 | 1 | 1 | # | 10 | Baseline | ancom | 54 |
| # | time | 2 | 1 | 0 | 0.666666667 | 1 | # | 10 | Baseline | ancom | 54 |
| # | interaction | 0 | 0 | 2 | 0 | 0 | # | 10 | Baseline | ancom | 54 |
| # | disease | 4 | 15 | 0 | 0.210526316 | 1 | # | 10 | Baseline | aldex2 | 54 |
| # | sex | 2 | 4 | 0 | 0.333333333 | 1 | # | 10 | Baseline | aldex2 | 54 |
| # | time | 2 | 9 | 0 | 0.181818182 | 1 | # | 10 | Baseline | aldex2 | 54 |
| # | interaction | 0 | 0 | 2 | 0 | 0 | # | 10 | Baseline | aldex2 | 54 |
| # | disease | 4 | 0 | 0 | 1 | 1 | # | 10 | Small | paralle | 54 |
| # | sex | 2 | 1 | 0 | 0.666666667 | 1 | # | 10 | Small | paralle | 54 |
| # | time | 2 | 0 | 0 | 1 | 1 | # | 10 | Small | paralle | 54 |
| # | interaction | 2 | 0 | 0 | 1 | 1 | # | 10 | Small | paralle | 54 |
| # | disease | 4 | 2 | 0 | 0.666666667 | 1 | # | 10 | Small | ancom | 54 |
| # | sex | 2 | 0 | 0 | 1 | 1 | # | 10 | Small | ancom | 54 |
| # | time | 2 | 1 | 0 | 0.666666667 | 1 | # | 10 | Small | ancom | 54 |
| # | interaction | 0 | 0 | 2 | 0 | 0 | # | 10 | Small | ancom | 54 |
| # | disease | 4 | 1 | 0 | 0.8 | 1 | # | 10 | Small | aldex2 | 54 |
| # | sex | 2 | 0 | 0 | 1 | 1 | # | 10 | Small | aldex2 | 54 |
| # | time | 2 | 1 | 0 | 0.666666667 | 1 | # | 10 | Small | aldex2 | 54 |
| # | interaction | 0 | 0 | 2 | 0 | 0 | # | 10 | Small | aldex2 | 54 |
| # | disease | 4 | 1 | 0 | 0.8 | 1 | # | 10 | Baseline | paralle | 54 |
| # | sex | 2 | 4 | 0 | 0.333333333 | 1 | # | 10 | Baseline | paralle | 54 |
| # | time | 2 | 5 | 0 | 0.285714286 | 1 | # | 10 | Baseline | paralle | 54 |
| # | interaction | 2 | 5 | 0 | 0.285714286 | 1 | # | 10 | Baseline | paralle | 54 |
| # | disease | 4 | 2 | 0 | 0.666666667 | 1 | # | 10 | Baseline | ancom | 54 |
| # | sex | 2 | 0 | 0 | 1 | 1 | # | 10 | Baseline | ancom | 54 |
| # | time | 2 | 2 | 0 | 0.5 | 1 | # | 10 | Baseline | ancom | 54 |
| # | interaction | 0 | 0 | 2 | 0 | 0 | # | 10 | Baseline | ancom | 54 |
| # | disease | 4 | 19 | 0 | 0.173913043 | 1 | # | 10 | Baseline | aldex2 | 54 |
| # | sex | 2 | 7 | 0 | 0.222222222 | 1 | # | 10 | Baseline | aldex2 | 54 |
| # | time | 2 | 10 | 0 | 0.166666667 | 1 | # | 10 | Baseline | aldex2 | 54 |
| # | interaction | 0 | 0 | 2 | 0 | 0 | # | 10 | Baseline | aldex2 | 54 |
| # | disease | 4 | 0 | 0 | 1 | 1 | # | 10 | Small | paralle | 54 |
| # | sex | 2 | 1 | 0 | 0.666666667 | 1 | # | 10 | Small | paralle | 54 |
| # | time | 2 | 0 | 0 | 1 | 1 | # | 10 | Small | paralle | 54 |
| # | interaction | 2 | 0 | 0 | 1 | 1 | # | 10 | Small | paralle | 54 |
| # | disease | 4 | 2 | 0 | 0.666666667 | 1 | # | 10 | Small | ancom | 54 |
| # | sex | 2 | 0 | 0 | 1 | 1 | # | 10 | Small | ancom | 54 |
| # | time | 2 | 1 | 0 | 0.666666667 | 1 | # | 10 | Small | ancom | 54 |
| # | interaction | 0 | 0 | 2 | 0 | 0 | # | 10 | Small | ancom | 54 |
| # | disease | 4 | 3 | 0 | 0.571428571 | 1 | # | 10 | Small | aldex2 | 54 |
| # | sex | 2 | 0 | 0 | 1 | 1 | # | 10 | Small | aldex2 | 54 |
| # | time | 2 | 1 | 0 | 0.666666667 | 1 | # | 10 | Small | aldex2 | 54 |
| # | interaction | 0 | 0 | 2 | 0 | 0 | # | 10 | Small | aldex2 | 54 |
| # | disease | 4 | 0 | 0 | 1 | 1 | # | 10 | Baseline | paralle | 54 |
| # | sex | 2 | 4 | 0 | 0.333333333 | 1 | # | 10 | Baseline | paralle | 54 |
| # | time | 2 | 2 | 0 | 0.5 | 1 | # | 10 | Baseline | paralle | 54 |
| # | interaction | 2 | 5 | 0 | 0.285714286 | 1 | # | 10 | Baseline | paralle | 54 |
| # | disease | 4 | 2 | 0 | 0.666666667 | 1 | # | 10 | Baseline | ancom | 54 |
| # | sex | 2 | 0 | 0 | 1 | 1 | # | 10 | Baseline | ancom | 54 |
| # | time | 2 | 2 | 0 | 0.5 | 1 | # | 10 | Baseline | ancom | 54 |
| # | interaction | 0 | 0 | 2 | 0 | 0 | # | 10 | Baseline | ancom | 54 |
| # | disease | 4 | 23 | 0 | 0.148148148 | 1 | # | 10 | Baseline | aldex2 | 54 |
| # | sex | 2 | 6 | 0 | 0.25 | 1 | # | 10 | Baseline | aldex2 | 54 |
| # | time | 2 | 8 | 0 | 0.2 | 1 | # | 10 | Baseline | aldex2 | 54 |
| # | interaction | 0 | 0 | 2 | 0 | 0 | # | 10 | Baseline | aldex2 | 54 |
| # | disease | 4 | 0 | 0 | 1 | 1 | # | 10 | Small | paralle | 54 |
| # | sex | 2 | 0 | 0 | 1 | 1 | # | 10 | Small | paralle | 54 |
| # | time | 2 | 0 | 0 | 1 | 1 | # | 10 | Small | paralle | 54 |
| # | interaction | 2 | 0 | 0 | 1 | 1 | # | 10 | Small | paralle | 54 |
| # | disease | 4 | 2 | 0 | 0.666666667 | 1 | # | 10 | Small | ancom | 54 |
| # | sex | 2 | 0 | 0 | 1 | 1 | # | 10 | Small | ancom | 54 |
| # | time | 2 | 2 | 0 | 0.5 | 1 | # | 10 | Small | ancom | 54 |
| # | interaction | 0 | 0 | 2 | 0 | 0 | # | 10 | Small | ancom | 54 |
| # | disease | 4 | 3 | 0 | 0.571428571 | 1 | # | 10 | Small | aldex2 | 54 |
| # | sex | 2 | 0 | 0 | 1 | 1 | # | 10 | Small | aldex2 | 54 |
| # | time | 2 | 2 | 0 | 0.5 | 1 | # | 10 | Small | aldex2 | 54 |
| # | interaction | 0 | 0 | 2 | 0 | 0 | # | 10 | Small | aldex2 | 54 |
| # | disease | 7 | 9 | 0 | 0.4375 | 1 | # | 20 | Baseline | paralle | 54 |
| # | sex | 4 | 4 | 0 | 0.5 | 1 | # | 20 | Baseline | paralle | 54 |
| # | time | 5 | 10 | 0 | 0.333333333 | 1 | # | 20 | Baseline | paralle | 54 |
| # | interaction | 4 | 11 | 0 | 0.266666667 | 1 | # | 20 | Baseline | paralle | 54 |
| # | disease | 7 | 0 | 0 | 1 | 1 | # | 20 | Baseline | ancom | 54 |
| # | sex | 4 | 0 | 0 | 1 | 1 | # | 20 | Baseline | ancom | 54 |
| # | time | 5 | 0 | 0 | 1 | 1 | # | 20 | Baseline | ancom | 54 |
| # | interaction | 0 | 0 | 4 | 0 | 0 | # | 20 | Baseline | ancom | 54 |
| # | disease | 7 | 19 | 0 | 0.269230769 | 1 | # | 20 | Baseline | aldex2 | 54 |
| # | sex | 4 | 4 | 0 | 0.5 | 1 | # | 20 | Baseline | aldex2 | 54 |
| # | time | 5 | 18 | 0 | 0.217391304 | 1 | # | 20 | Baseline | aldex2 | 54 |
| # | interaction | 0 | 0 | 4 | 0 | 0 | # | 20 | Baseline | aldex2 | 54 |
| # | disease | 7 | 0 | 0 | 1 | 1 | # | 20 | Small | paralle | 54 |
| # | sex | 4 | 0 | 0 | 1 | 1 | # | 20 | Small | paralle | 54 |
| # | time | 5 | 0 | 0 | 1 | 1 | # | 20 | Small | paralle | 54 |
| # | interaction | 2 | 0 | 2 | 1 | 0.5 | # | 20 | Small | paralle | 54 |
| # | disease | 7 | 0 | 0 | 1 | 1 | # | 20 | Small | ancom | 54 |
| # | sex | 4 | 0 | 0 | 1 | 1 | # | 20 | Small | ancom | 54 |
| # | time | 5 | 0 | 0 | 1 | 1 | # | 20 | Small | ancom | 54 |
| # | interaction | 0 | 0 | 4 | 0 | 0 | # | 20 | Small | ancom | 54 |
| # | disease | 7 | 2 | 0 | 0.777777778 | 1 | # | 20 | Small | aldex2 | 54 |
| # | sex | 4 | 0 | 0 | 1 | 1 | # | 20 | Small | aldex2 | 54 |
| # | time | 5 | 1 | 0 | 0.833333333 | 1 | # | 20 | Small | aldex2 | 54 |
| # | interaction | 0 | 0 | 4 | 0 | 0 | # | 20 | Small | aldex2 | 54 |
| # | disease | 7 | 9 | 0 | 0.4375 | 1 | # | 20 | Baseline | paralle | 54 |
| # | sex | 4 | 5 | 0 | 0.444444444 | 1 | # | 20 | Baseline | paralle | 54 |
| # | time | 5 | 10 | 0 | 0.333333333 | 1 | # | 20 | Baseline | paralle | 54 |
| # | interaction | 4 | 7 | 0 | 0.363636364 | 1 | # | 20 | Baseline | paralle | 54 |
| # | disease | 7 | 1 | 0 | 0.875 | 1 | # | 20 | Baseline | ancom | 54 |
| # | sex | 4 | 0 | 0 | 1 | 1 | # | 20 | Baseline | ancom | 54 |
| # | time | 5 | 4 | 0 | 0.555555556 | 1 | # | 20 | Baseline | ancom | 54 |
| # | interaction | 0 | 0 | 4 | 0 | 0 | # | 20 | Baseline | ancom | 54 |
| # | disease | 7 | 30 | 0 | 0.189189189 | 1 | # | 20 | Baseline | aldex2 | 54 |
| # | sex | 4 | 9 | 0 | 0.307692308 | 1 | # | 20 | Baseline | aldex2 | 54 |
| # | time | 5 | 27 | 0 | 0.15625 | 1 | # | 20 | Baseline | aldex2 | 54 |
| # | interaction | 0 | 0 | 4 | 0 | 0 | # | 20 | Baseline | aldex2 | 54 |
| # | disease | 7 | 0 | 0 | 1 | 1 | # | 20 | Small | paralle | 54 |
| # | sex | 4 | 0 | 0 | 1 | 1 | # | 20 | Small | paralle | 54 |
| # | time | 5 | 0 | 0 | 1 | 1 | # | 20 | Small | paralle | 54 |
| # | interaction | 3 | 0 | 1 | 1 | 0.75 | # | 20 | Small | paralle | 54 |
| # | disease | 7 | 2 | 0 | 0.777777778 | 1 | # | 20 | Small | ancom | 54 |
| # | sex | 4 | 0 | 0 | 1 | 1 | # | 20 | Small | ancom | 54 |
| # | time | 5 | 3 | 0 | 0.625 | 1 | # | 20 | Small | ancom | 54 |
| # | interaction | 0 | 0 | 4 | 0 | 0 | # | 20 | Small | ancom | 54 |
| # | disease | 7 | 3 | 0 | 0.7 | 1 | # | 20 | Small | aldex2 | 54 |
| # | sex | 4 | 0 | 0 | 1 | 1 | # | 20 | Small | aldex2 | 54 |
| # | time | 5 | 1 | 0 | 0.833333333 | 1 | # | 20 | Small | aldex2 | 54 |
| # | interaction | 0 | 0 | 4 | 0 | 0 | # | 20 | Small | aldex2 | 54 |
| # | disease | 7 | 12 | 0 | 0.368421053 | 1 | # | 20 | Baseline | paralle | 54 |
| # | sex | 4 | 6 | 0 | 0.4 | 1 | # | 20 | Baseline | paralle | 54 |
| # | time | 5 | 10 | 0 | 0.333333333 | 1 | # | 20 | Baseline | paralle | 54 |
| # | interaction | 4 | 4 | 0 | 0.5 | 1 | # | 20 | Baseline | paralle | 54 |
| # | disease | 7 | 4 | 0 | 0.636363636 | 1 | # | 20 | Baseline | ancom | 54 |
| # | sex | 4 | 0 | 0 | 1 | 1 | # | 20 | Baseline | ancom | 54 |
| # | time | 5 | 4 | 0 | 0.555555556 | 1 | # | 20 | Baseline | ancom | 54 |
| # | interaction | 0 | 0 | 4 | 0 | 0 | # | 20 | Baseline | ancom | 54 |
| # | disease | 7 | 49 | 0 | 0.125 | 1 | # | 20 | Baseline | aldex2 | 54 |
| # | sex | 4 | 11 | 0 | 0.266666667 | 1 | # | 20 | Baseline | aldex2 | 54 |
| # | time | 5 | 30 | 0 | 0.142857143 | 1 | # | 20 | Baseline | aldex2 | 54 |
| # | interaction | 0 | 0 | 4 | 0 | 0 | # | 20 | Baseline | aldex2 | 54 |
| # | disease | 7 | 0 | 0 | 1 | 1 | # | 20 | Small | paralle | 54 |
| # | sex | 4 | 0 | 0 | 1 | 1 | # | 20 | Small | paralle | 54 |
| # | time | 5 | 0 | 0 | 1 | 1 | # | 20 | Small | paralle | 54 |
| # | interaction | 4 | 0 | 0 | 1 | 1 | # | 20 | Small | paralle | 54 |
| # | disease | 7 | 4 | 0 | 0.636363636 | 1 | # | 20 | Small | ancom | 54 |
| # | sex | 4 | 0 | 0 | 1 | 1 | # | 20 | Small | ancom | 54 |
| # | time | 5 | 4 | 0 | 0.555555556 | 1 | # | 20 | Small | ancom | 54 |
| # | interaction | 0 | 0 | 4 | 0 | 0 | # | 20 | Small | ancom | 54 |
| # | disease | 7 | 3 | 0 | 0.7 | 1 | # | 20 | Small | aldex2 | 54 |
| # | sex | 4 | 0 | 0 | 1 | 1 | # | 20 | Small | aldex2 | 54 |
| # | time | 5 | 7 | 0 | 0.416666667 | 1 | # | 20 | Small | aldex2 | 54 |
| # | interaction | 0 | 0 | 4 | 0 | 0 | # | 20 | Small | aldex2 | 54 |
| # | disease | 7 | 14 | 0 | 0.333333333 | 1 | # | 20 | Baseline | paralle | 54 |
| # | sex | 4 | 3 | 0 | 0.571428571 | 1 | # | 20 | Baseline | paralle | 54 |
| # | time | 5 | 5 | 0 | 0.5 | 1 | # | 20 | Baseline | paralle | 54 |
| # | interaction | 4 | 6 | 0 | 0.4 | 1 | # | 20 | Baseline | paralle | 54 |
| # | disease | 7 | 4 | 0 | 0.636363636 | 1 | # | 20 | Baseline | ancom | 54 |
| # | sex | 4 | 0 | 0 | 1 | 1 | # | 20 | Baseline | ancom | 54 |
| # | time | 5 | 4 | 0 | 0.555555556 | 1 | # | 20 | Baseline | ancom | 54 |
| # | interaction | 0 | 0 | 4 | 0 | 0 | # | 20 | Baseline | ancom | 54 |
| # | disease | 7 | 60 | 0 | 0.104477612 | 1 | # | 20 | Baseline | aldex2 | 54 |
| # | sex | 4 | 16 | 0 | 0.2 | 1 | # | 20 | Baseline | aldex2 | 54 |
| # | time | 5 | 46 | 0 | 0.098039216 | 1 | # | 20 | Baseline | aldex2 | 54 |
| # | interaction | 0 | 0 | 4 | 0 | 0 | # | 20 | Baseline | aldex2 | 54 |
| # | disease | 7 | 0 | 0 | 1 | 1 | # | 20 | Small | paralle | 54 |
| # | sex | 4 | 0 | 0 | 1 | 1 | # | 20 | Small | paralle | 54 |
| # | time | 5 | 0 | 0 | 1 | 1 | # | 20 | Small | paralle | 54 |
| # | interaction | 4 | 0 | 0 | 1 | 1 | # | 20 | Small | paralle | 54 |
| # | disease | 7 | 4 | 0 | 0.636363636 | 1 | # | 20 | Small | ancom | 54 |
| # | sex | 4 | 0 | 0 | 1 | 1 | # | 20 | Small | ancom | 54 |
| # | time | 5 | 4 | 0 | 0.555555556 | 1 | # | 20 | Small | ancom | 54 |
| # | interaction | 0 | 0 | 4 | 0 | 0 | # | 20 | Small | ancom | 54 |
| # | disease | 7 | 7 | 0 | 0.5 | 1 | # | 20 | Small | aldex2 | 54 |
| # | sex | 4 | 0 | 0 | 1 | 1 | # | 20 | Small | aldex2 | 54 |
| # | time | 5 | 7 | 0 | 0.416666667 | 1 | # | 20 | Small | aldex2 | 54 |
| # | interaction | 0 | 0 | 4 | 0 | 0 | # | 20 | Small | aldex2 | 54 |
| # | disease | 4 | 5 | 0 | 0.444444444 | 1 | # | 10 | Baseline | paralle | 55 |
| # | sex | 2 | 7 | 0 | 0.222222222 | 1 | # | 10 | Baseline | paralle | 55 |
| # | time | 2 | 8 | 0 | 0.2 | 1 | # | 10 | Baseline | paralle | 55 |
| # | interaction | 2 | 5 | 0 | 0.285714286 | 1 | # | 10 | Baseline | paralle | 55 |
| # | disease | 4 | 0 | 0 | 1 | 1 | # | 10 | Baseline | ancom | 55 |
| # | sex | 2 | 0 | 0 | 1 | 1 | # | 10 | Baseline | ancom | 55 |
| # | time | 2 | 0 | 0 | 1 | 1 | # | 10 | Baseline | ancom | 55 |
| # | interaction | 0 | 0 | 2 | 0 | 0 | # | 10 | Baseline | ancom | 55 |
| # | disease | 4 | 6 | 0 | 0.4 | 1 | # | 10 | Baseline | aldex2 | 55 |
| # | sex | 2 | 8 | 0 | 0.2 | 1 | # | 10 | Baseline | aldex2 | 55 |
| # | time | 2 | 7 | 0 | 0.222222222 | 1 | # | 10 | Baseline | aldex2 | 55 |
| # | interaction | 0 | 0 | 2 | 0 | 0 | # | 10 | Baseline | aldex2 | 55 |
| # | disease | 4 | 0 | 0 | 1 | 1 | # | 10 | Small | paralle | 55 |
| # | sex | 2 | 0 | 0 | 1 | 1 | # | 10 | Small | paralle | 55 |
| # | time | 2 | 0 | 0 | 1 | 1 | # | 10 | Small | paralle | 55 |
| # | interaction | 1 | 0 | 1 | 1 | 0.5 | # | 10 | Small | paralle | 55 |
| # | disease | 4 | 0 | 0 | 1 | 1 | # | 10 | Small | ancom | 55 |
| # | sex | 2 | 0 | 0 | 1 | 1 | # | 10 | Small | ancom | 55 |
| # | time | 2 | 0 | 0 | 1 | 1 | # | 10 | Small | ancom | 55 |
| # | interaction | 0 | 0 | 2 | 0 | 0 | # | 10 | Small | ancom | 55 |
| # | disease | 4 | 0 | 0 | 1 | 1 | # | 10 | Small | aldex2 | 55 |
| # | sex | 2 | 0 | 0 | 1 | 1 | # | 10 | Small | aldex2 | 55 |
| # | time | 2 | 1 | 0 | 0.666666667 | 1 | # | 10 | Small | aldex2 | 55 |
| # | interaction | 0 | 0 | 2 | 0 | 0 | # | 10 | Small | aldex2 | 55 |
| # | disease | 4 | 6 | 0 | 0.4 | 1 | # | 10 | Baseline | paralle | 55 |
| # | sex | 2 | 5 | 0 | 0.285714286 | 1 | # | 10 | Baseline | paralle | 55 |
| # | time | 2 | 3 | 0 | 0.4 | 1 | # | 10 | Baseline | paralle | 55 |
| # | interaction | 2 | 3 | 0 | 0.4 | 1 | # | 10 | Baseline | paralle | 55 |
| # | disease | 4 | 1 | 0 | 0.8 | 1 | # | 10 | Baseline | ancom | 55 |
| # | sex | 2 | 0 | 0 | 1 | 1 | # | 10 | Baseline | ancom | 55 |
| # | time | 2 | 0 | 0 | 1 | 1 | # | 10 | Baseline | ancom | 55 |
| # | interaction | 0 | 0 | 2 | 0 | 0 | # | 10 | Baseline | ancom | 55 |
| # | disease | 4 | 14 | 0 | 0.222222222 | 1 | # | 10 | Baseline | aldex2 | 55 |
| # | sex | 2 | 9 | 0 | 0.181818182 | 1 | # | 10 | Baseline | aldex2 | 55 |
| # | time | 2 | 13 | 0 | 0.133333333 | 1 | # | 10 | Baseline | aldex2 | 55 |
| # | interaction | 0 | 0 | 2 | 0 | 0 | # | 10 | Baseline | aldex2 | 55 |
| # | disease | 4 | 0 | 0 | 1 | 1 | # | 10 | Small | paralle | 55 |
| # | sex | 2 | 0 | 0 | 1 | 1 | # | 10 | Small | paralle | 55 |
| # | time | 2 | 0 | 0 | 1 | 1 | # | 10 | Small | paralle | 55 |
| # | interaction | 1 | 0 | 1 | 1 | 0.5 | # | 10 | Small | paralle | 55 |
| # | disease | 4 | 1 | 0 | 0.8 | 1 | # | 10 | Small | ancom | 55 |
| # | sex | 2 | 0 | 0 | 1 | 1 | # | 10 | Small | ancom | 55 |
| # | time | 2 | 0 | 0 | 1 | 1 | # | 10 | Small | ancom | 55 |
| # | interaction | 0 | 0 | 2 | 0 | 0 | # | 10 | Small | ancom | 55 |
| # | disease | 4 | 0 | 0 | 1 | 1 | # | 10 | Small | aldex2 | 55 |
| # | sex | 2 | 0 | 0 | 1 | 1 | # | 10 | Small | aldex2 | 55 |
| # | time | 2 | 1 | 0 | 0.666666667 | 1 | # | 10 | Small | aldex2 | 55 |
| # | interaction | 0 | 0 | 2 | 0 | 0 | # | 10 | Small | aldex2 | 55 |
| # | disease | 4 | 4 | 0 | 0.5 | 1 | # | 10 | Baseline | paralle | 55 |
| # | sex | 2 | 4 | 0 | 0.333333333 | 1 | # | 10 | Baseline | paralle | 55 |
| # | time | 2 | 5 | 0 | 0.285714286 | 1 | # | 10 | Baseline | paralle | 55 |
| # | interaction | 2 | 3 | 0 | 0.4 | 1 | # | 10 | Baseline | paralle | 55 |
| # | disease | 4 | 1 | 0 | 0.8 | 1 | # | 10 | Baseline | ancom | 55 |
| # | sex | 2 | 0 | 0 | 1 | 1 | # | 10 | Baseline | ancom | 55 |
| # | time | 2 | 2 | 0 | 0.5 | 1 | # | 10 | Baseline | ancom | 55 |
| # | interaction | 0 | 0 | 2 | 0 | 0 | # | 10 | Baseline | ancom | 55 |
| # | disease | 4 | 23 | 0 | 0.148148148 | 1 | # | 10 | Baseline | aldex2 | 55 |
| # | sex | 2 | 9 | 0 | 0.181818182 | 1 | # | 10 | Baseline | aldex2 | 55 |
| # | time | 2 | 10 | 0 | 0.166666667 | 1 | # | 10 | Baseline | aldex2 | 55 |
| # | interaction | 0 | 0 | 2 | 0 | 0 | # | 10 | Baseline | aldex2 | 55 |
| # | disease | 4 | 0 | 0 | 1 | 1 | # | 10 | Small | paralle | 55 |
| # | sex | 2 | 0 | 0 | 1 | 1 | # | 10 | Small | paralle | 55 |
| # | time | 2 | 0 | 0 | 1 | 1 | # | 10 | Small | paralle | 55 |
| # | interaction | 2 | 0 | 0 | 1 | 1 | # | 10 | Small | paralle | 55 |
| # | disease | 4 | 1 | 0 | 0.8 | 1 | # | 10 | Small | ancom | 55 |
| # | sex | 2 | 0 | 0 | 1 | 1 | # | 10 | Small | ancom | 55 |
| # | time | 2 | 2 | 0 | 0.5 | 1 | # | 10 | Small | ancom | 55 |
| # | interaction | 0 | 0 | 2 | 0 | 0 | # | 10 | Small | ancom | 55 |
| # | disease | 4 | 1 | 0 | 0.8 | 1 | # | 10 | Small | aldex2 | 55 |
| # | sex | 2 | 1 | 0 | 0.666666667 | 1 | # | 10 | Small | aldex2 | 55 |
| # | time | 2 | 3 | 0 | 0.4 | 1 | # | 10 | Small | aldex2 | 55 |
| # | interaction | 0 | 0 | 2 | 0 | 0 | # | 10 | Small | aldex2 | 55 |
| # | disease | 4 | 6 | 0 | 0.4 | 1 | # | 10 | Baseline | paralle | 55 |
| # | sex | 2 | 3 | 0 | 0.4 | 1 | # | 10 | Baseline | paralle | 55 |
| # | time | 2 | 4 | 0 | 0.333333333 | 1 | # | 10 | Baseline | paralle | 55 |
| # | interaction | 2 | 3 | 0 | 0.4 | 1 | # | 10 | Baseline | paralle | 55 |

| | | | | | | | | | | | |
|---|---|---|---|---|---|---|---|---|---|---|---|
| # | disease | 4 | 2 | 0 | 0.666666667 | 1 | # | 10 | Baseline | ancom | 55 |
| # | sex | 2 | 0 | 0 | 1 | 1 | # | 10 | Baseline | ancom | 55 |
| # | time | 2 | 2 | 0 | 0.5 | 1 | # | 10 | Baseline | ancom | 55 |
| # | interaction | 0 | 0 | 2 | 0 | 0 | # | 10 | Baseline | ancom | 55 |
| # | disease | 4 | 24 | 0 | 0.142857143 | 1 | # | 10 | Baseline | aldex2 | 55 |
| # | sex | 2 | 12 | 0 | 0.142857143 | 1 | # | 10 | Baseline | aldex2 | 55 |
| # | time | 2 | 11 | 0 | 0.153846154 | 1 | # | 10 | Baseline | aldex2 | 55 |
| # | interaction | 0 | 0 | 2 | 0 | 0 | # | 10 | Baseline | aldex2 | 55 |
| # | disease | 4 | 0 | 0 | 1 | 1 | # | 10 | Small | paralle | 55 |
| # | sex | 2 | 0 | 0 | 1 | 1 | # | 10 | Small | paralle | 55 |
| # | time | 2 | 0 | 0 | 1 | 1 | # | 10 | Small | paralle | 55 |
| # | interaction | 2 | 0 | 0 | 1 | 1 | # | 10 | Small | paralle | 55 |
| # | disease | 4 | 2 | 0 | 0.666666667 | 1 | # | 10 | Small | ancom | 55 |
| # | sex | 2 | 0 | 0 | 1 | 1 | # | 10 | Small | ancom | 55 |
| # | time | 2 | 2 | 0 | 0.5 | 1 | # | 10 | Small | ancom | 55 |
| # | interaction | 0 | 0 | 2 | 0 | 0 | # | 10 | Small | ancom | 55 |
| # | disease | 4 | 1 | 0 | 0.8 | 1 | # | 10 | Small | aldex2 | 55 |
| # | sex | 2 | 0 | 0 | 1 | 1 | # | 10 | Small | aldex2 | 55 |
| # | time | 2 | 3 | 0 | 0.4 | 1 | # | 10 | Small | aldex2 | 55 |
| # | interaction | 0 | 0 | 2 | 0 | 0 | # | 10 | Small | aldex2 | 55 |
| # | disease | 7 | 10 | 0 | 0.411764706 | 1 | # | 20 | Baseline | paralle | 55 |
| # | sex | 4 | 9 | 0 | 0.307692308 | 1 | # | 20 | Baseline | paralle | 55 |
| # | time | 5 | 6 | 0 | 0.454545455 | 1 | # | 20 | Baseline | paralle | 55 |
| # | interaction | 4 | 7 | 0 | 0.363636364 | 1 | # | 20 | Baseline | paralle | 55 |
| # | disease | 7 | 1 | 0 | 0.875 | 1 | # | 20 | Baseline | ancom | 55 |
| # | sex | 4 | 0 | 0 | 1 | 1 | # | 20 | Baseline | ancom | 55 |
| # | time | 5 | 4 | 0 | 0.555555556 | 1 | # | 20 | Baseline | ancom | 55 |
| # | interaction | 0 | 0 | 4 | 0 | 0 | # | 20 | Baseline | ancom | 55 |
| # | disease | 7 | 18 | 0 | 0.28 | 1 | # | 20 | Baseline | aldex2 | 55 |
| # | sex | 4 | 9 | 0 | 0.307692308 | 1 | # | 20 | Baseline | aldex2 | 55 |
| # | time | 5 | 16 | 0 | 0.238095238 | 1 | # | 20 | Baseline | aldex2 | 55 |
| # | interaction | 0 | 0 | 4 | 0 | 0 | # | 20 | Baseline | aldex2 | 55 |
| # | disease | 7 | 0 | 0 | 1 | 1 | # | 20 | Small | paralle | 55 |
| # | sex | 4 | 0 | 0 | 1 | 1 | # | 20 | Small | paralle | 55 |
| # | time | 5 | 0 | 0 | 1 | 1 | # | 20 | Small | paralle | 55 |
| # | interaction | 4 | 0 | 0 | 1 | 1 | # | 20 | Small | paralle | 55 |
| # | disease | 7 | 1 | 0 | 0.875 | 1 | # | 20 | Small | ancom | 55 |
| # | sex | 4 | 0 | 0 | 1 | 1 | # | 20 | Small | ancom | 55 |
| # | time | 5 | 2 | 0 | 0.714285714 | 1 | # | 20 | Small | ancom | 55 |
| # | interaction | 0 | 0 | 4 | 0 | 0 | # | 20 | Small | ancom | 55 |
| # | disease | 7 | 1 | 0 | 0.875 | 1 | # | 20 | Small | aldex2 | 55 |
| # | sex | 4 | 1 | 0 | 0.8 | 1 | # | 20 | Small | aldex2 | 55 |
| # | time | 5 | 2 | 0 | 0.714285714 | 1 | # | 20 | Small | aldex2 | 55 |
| # | interaction | 0 | 0 | 4 | 0 | 0 | # | 20 | Small | aldex2 | 55 |
| # | disease | 7 | 15 | 0 | 0.318181818 | 1 | # | 20 | Baseline | paralle | 55 |
| # | sex | 4 | 5 | 0 | 0.444444444 | 1 | # | 20 | Baseline | paralle | 55 |
| # | time | 5 | 8 | 0 | 0.384615385 | 1 | # | 20 | Baseline | paralle | 55 |
| # | interaction | 4 | 6 | 0 | 0.4 | 1 | # | 20 | Baseline | paralle | 55 |
| # | disease | 7 | 4 | 0 | 0.636363636 | 1 | # | 20 | Baseline | ancom | 55 |
| # | sex | 4 | 0 | 0 | 1 | 1 | # | 20 | Baseline | ancom | 55 |
| # | time | 5 | 4 | 0 | 0.555555556 | 1 | # | 20 | Baseline | ancom | 55 |
| # | interaction | 0 | 0 | 4 | 0 | 0 | # | 20 | Baseline | ancom | 55 |
| # | disease | 7 | 32 | 0 | 0.179487179 | 1 | # | 20 | Baseline | aldex2 | 55 |
| # | sex | 4 | 12 | 0 | 0.25 | 1 | # | 20 | Baseline | aldex2 | 55 |
| # | time | 5 | 18 | 0 | 0.217391304 | 1 | # | 20 | Baseline | aldex2 | 55 |
| # | interaction | 0 | 0 | 4 | 0 | 0 | # | 20 | Baseline | aldex2 | 55 |
| # | disease | 7 | 1 | 0 | 0.875 | 1 | # | 20 | Small | paralle | 55 |
| # | sex | 4 | 0 | 0 | 1 | 1 | # | 20 | Small | paralle | 55 |
| # | time | 5 | 0 | 0 | 1 | 1 | # | 20 | Small | paralle | 55 |
| # | interaction | 4 | 1 | 0 | 0.8 | 1 | # | 20 | Small | paralle | 55 |
| # | disease | 7 | 3 | 0 | 0.7 | 1 | # | 20 | Small | ancom | 55 |
| # | sex | 4 | 0 | 0 | 1 | 1 | # | 20 | Small | ancom | 55 |
| # | time | 5 | 4 | 0 | 0.555555556 | 1 | # | 20 | Small | ancom | 55 |
| # | interaction | 0 | 0 | 4 | 0 | 0 | # | 20 | Small | ancom | 55 |
| # | disease | 7 | 5 | 0 | 0.583333333 | 1 | # | 20 | Small | aldex2 | 55 |
| # | sex | 4 | 2 | 0 | 0.666666667 | 1 | # | 20 | Small | aldex2 | 55 |
| # | time | 5 | 4 | 0 | 0.555555556 | 1 | # | 20 | Small | aldex2 | 55 |
| # | interaction | 0 | 0 | 4 | 0 | 0 | # | 20 | Small | aldex2 | 55 |
| # | disease | 7 | 9 | 0 | 0.4375 | 1 | # | 20 | Baseline | paralle | 55 |
| # | sex | 4 | 4 | 0 | 0.5 | 1 | # | 20 | Baseline | paralle | 55 |
| # | time | 5 | 6 | 0 | 0.454545455 | 1 | # | 20 | Baseline | paralle | 55 |
| # | interaction | 4 | 6 | 0 | 0.4 | 1 | # | 20 | Baseline | paralle | 55 |
| # | disease | 7 | 3 | 0 | 0.7 | 1 | # | 20 | Baseline | ancom | 55 |
| # | sex | 4 | 0 | 0 | 1 | 1 | # | 20 | Baseline | ancom | 55 |
| # | time | 5 | 4 | 0 | 0.555555556 | 1 | # | 20 | Baseline | ancom | 55 |
| # | interaction | 0 | 0 | 4 | 0 | 0 | # | 20 | Baseline | ancom | 55 |
| # | disease | 7 | 50 | 0 | 0.122807018 | 1 | # | 20 | Baseline | aldex2 | 55 |
| # | sex | 4 | 16 | 0 | 0.2 | 1 | # | 20 | Baseline | aldex2 | 55 |
| # | time | 5 | 23 | 0 | 0.178571429 | 1 | # | 20 | Baseline | aldex2 | 55 |
| # | interaction | 0 | 0 | 4 | 0 | 0 | # | 20 | Baseline | aldex2 | 55 |
| # | disease | 7 | 0 | 0 | 1 | 1 | # | 20 | Small | paralle | 55 |
| # | sex | 4 | 0 | 0 | 1 | 1 | # | 20 | Small | paralle | 55 |
| # | time | 5 | 0 | 0 | 1 | 1 | # | 20 | Small | paralle | 55 |
| # | interaction | 4 | 0 | 0 | 1 | 1 | # | 20 | Small | paralle | 55 |
| # | disease | 7 | 2 | 0 | 0.777777778 | 1 | # | 20 | Small | ancom | 55 |
| # | sex | 4 | 0 | 0 | 1 | 1 | # | 20 | Small | ancom | 55 |
| # | time | 5 | 4 | 0 | 0.555555556 | 1 | # | 20 | Small | ancom | 55 |
| # | interaction | 0 | 0 | 4 | 0 | 0 | # | 20 | Small | ancom | 55 |
| # | disease | 7 | 7 | 0 | 0.5 | 1 | # | 20 | Small | aldex2 | 55 |
| # | sex | 4 | 0 | 0 | 1 | 1 | # | 20 | Small | aldex2 | 55 |
| # | time | 5 | 3 | 0 | 0.625 | 1 | # | 20 | Small | aldex2 | 55 |
| # | interaction | 0 | 0 | 4 | 0 | 0 | # | 20 | Small | aldex2 | 55 |
| # | disease | 7 | 7 | 0 | 0.5 | 1 | # | 20 | Baseline | paralle | 55 |
| # | sex | 4 | 3 | 0 | 0.571428571 | 1 | # | 20 | Baseline | paralle | 55 |
| # | time | 5 | 5 | 0 | 0.5 | 1 | # | 20 | Baseline | paralle | 55 |
| # | interaction | 4 | 7 | 0 | 0.363636364 | 1 | # | 20 | Baseline | paralle | 55 |
| # | disease | 7 | 4 | 0 | 0.636363636 | 1 | # | 20 | Baseline | ancom | 55 |
| # | sex | 4 | 0 | 0 | 1 | 1 | # | 20 | Baseline | ancom | 55 |
| # | time | 5 | 4 | 0 | 0.555555556 | 1 | # | 20 | Baseline | ancom | 55 |
| # | interaction | 0 | 0 | 4 | 0 | 0 | # | 20 | Baseline | ancom | 55 |
| # | disease | 7 | 65 | 0 | 0.097222222 | 1 | # | 20 | Baseline | aldex2 | 55 |
| # | sex | 4 | 18 | 0 | 0.181818182 | 1 | # | 20 | Baseline | aldex2 | 55 |
| # | time | 5 | 38 | 0 | 0.11627907 | 1 | # | 20 | Baseline | aldex2 | 55 |
| # | interaction | 0 | 0 | 4 | 0 | 0 | # | 20 | Baseline | aldex2 | 55 |
| # | disease | 7 | 0 | 0 | 1 | 1 | # | 20 | Small | paralle | 55 |
| # | sex | 4 | 0 | 0 | 1 | 1 | # | 20 | Small | paralle | 55 |
| # | time | 5 | 0 | 0 | 1 | 1 | # | 20 | Small | paralle | 55 |
| # | interaction | 4 | 0 | 0 | 1 | 1 | # | 20 | Small | paralle | 55 |
| # | disease | 7 | 4 | 0 | 0.636363636 | 1 | # | 20 | Small | ancom | 55 |
| # | sex | 4 | 0 | 0 | 1 | 1 | # | 20 | Small | ancom | 55 |
| # | time | 5 | 4 | 0 | 0.555555556 | 1 | # | 20 | Small | ancom | 55 |
| # | interaction | 0 | 0 | 4 | 0 | 0 | # | 20 | Small | ancom | 55 |
| # | disease | 7 | 11 | 0 | 0.388888889 | 1 | # | 20 | Small | aldex2 | 55 |
| # | sex | 4 | 0 | 0 | 1 | 1 | # | 20 | Small | aldex2 | 55 |
| # | time | 5 | 6 | 0 | 0.454545455 | 1 | # | 20 | Small | aldex2 | 55 |
| # | interaction | 0 | 0 | 4 | 0 | 0 | # | 20 | Small | aldex2 | 55 |
| # | disease | 4 | 11 | 0 | 0.266666667 | 1 | # | 10 | Baseline | paralle | 56 |
| # | sex | 2 | 10 | 0 | 0.166666667 | 1 | # | 10 | Baseline | paralle | 56 |
| # | time | 2 | 5 | 0 | 0.285714286 | 1 | # | 10 | Baseline | paralle | 56 |
| # | interaction | 2 | 16 | 0 | 0.111111111 | 1 | # | 10 | Baseline | paralle | 56 |
| # | disease | 4 | 0 | 0 | 1 | 1 | # | 10 | Baseline | ancom | 56 |
| # | sex | 2 | 0 | 0 | 1 | 1 | # | 10 | Baseline | ancom | 56 |
| # | time | 2 | 0 | 0 | 1 | 1 | # | 10 | Baseline | ancom | 56 |
| # | interaction | 0 | 0 | 2 | 0 | 0 | # | 10 | Baseline | ancom | 56 |
| # | disease | 4 | 10 | 0 | 0.285714286 | 1 | # | 10 | Baseline | aldex2 | 56 |
| # | sex | 2 | 10 | 0 | 0.166666667 | 1 | # | 10 | Baseline | aldex2 | 56 |
| # | time | 2 | 5 | 0 | 0.285714286 | 1 | # | 10 | Baseline | aldex2 | 56 |
| # | interaction | 0 | 0 | 2 | 0 | 0 | # | 10 | Baseline | aldex2 | 56 |
| # | disease | 4 | 0 | 0 | 1 | 1 | # | 10 | Small | paralle | 56 |
| # | sex | 2 | 0 | 0 | 1 | 1 | # | 10 | Small | paralle | 56 |
| # | time | 2 | 1 | 0 | 0.666666667 | 1 | # | 10 | Small | paralle | 56 |
| # | interaction | 2 | 0 | 0 | 1 | 1 | # | 10 | Small | paralle | 56 |
| # | disease | 4 | 0 | 0 | 1 | 1 | # | 10 | Small | ancom | 56 |
| # | sex | 2 | 0 | 0 | 1 | 1 | # | 10 | Small | ancom | 56 |
| # | time | 2 | 0 | 0 | 1 | 1 | # | 10 | Small | ancom | 56 |
| # | interaction | 0 | 0 | 2 | 0 | 0 | # | 10 | Small | ancom | 56 |
| # | disease | 4 | 1 | 0 | 0.8 | 1 | # | 10 | Small | aldex2 | 56 |
| # | sex | 2 | 0 | 0 | 1 | 1 | # | 10 | Small | aldex2 | 56 |
| # | time | 2 | 0 | 0 | 1 | 1 | # | 10 | Small | aldex2 | 56 |
| # | interaction | 0 | 0 | 2 | 0 | 0 | # | 10 | Small | aldex2 | 56 |
| # | disease | 4 | 3 | 0 | 0.571428571 | 1 | # | 10 | Baseline | paralle | 56 |
| # | sex | 2 | 4 | 0 | 0.333333333 | 1 | # | 10 | Baseline | paralle | 56 |
| # | time | 2 | 5 | 0 | 0.285714286 | 1 | # | 10 | Baseline | paralle | 56 |
| # | interaction | 2 | 3 | 0 | 0.4 | 1 | # | 10 | Baseline | paralle | 56 |
| # | disease | 4 | 1 | 0 | 0.8 | 1 | # | 10 | Baseline | ancom | 56 |
| # | sex | 2 | 0 | 0 | 1 | 1 | # | 10 | Baseline | ancom | 56 |
| # | time | 2 | 1 | 0 | 0.666666667 | 1 | # | 10 | Baseline | ancom | 56 |
| # | interaction | 0 | 0 | 2 | 0 | 0 | # | 10 | Baseline | ancom | 56 |
| # | disease | 4 | 19 | 0 | 0.173913043 | 1 | # | 10 | Baseline | aldex2 | 56 |
| # | sex | 2 | 5 | 0 | 0.285714286 | 1 | # | 10 | Baseline | aldex2 | 56 |
| # | time | 2 | 8 | 0 | 0.2 | 1 | # | 10 | Baseline | aldex2 | 56 |
| # | interaction | 0 | 0 | 2 | 0 | 0 | # | 10 | Baseline | aldex2 | 56 |
| # | disease | 4 | 0 | 0 | 1 | 1 | # | 10 | Small | paralle | 56 |
| # | sex | 2 | 0 | 0 | 1 | 1 | # | 10 | Small | paralle | 56 |
| # | time | 2 | 0 | 0 | 1 | 1 | # | 10 | Small | paralle | 56 |
| # | interaction | 2 | 0 | 0 | 1 | 1 | # | 10 | Small | paralle | 56 |
| # | disease | 4 | 1 | 0 | 0.8 | 1 | # | 10 | Small | ancom | 56 |
| # | sex | 2 | 0 | 0 | 1 | 1 | # | 10 | Small | ancom | 56 |
| # | time | 2 | 1 | 0 | 0.666666667 | 1 | # | 10 | Small | ancom | 56 |
| # | interaction | 0 | 0 | 2 | 0 | 0 | # | 10 | Small | ancom | 56 |
| # | disease | 4 | 1 | 0 | 0.8 | 1 | # | 10 | Small | aldex2 | 56 |
| # | sex | 2 | 0 | 0 | 1 | 1 | # | 10 | Small | aldex2 | 56 |
| # | time | 2 | 1 | 0 | 0.666666667 | 1 | # | 10 | Small | aldex2 | 56 |
| # | interaction | 0 | 0 | 2 | 0 | 0 | # | 10 | Small | aldex2 | 56 |
| # | disease | 4 | 4 | 0 | 0.5 | 1 | # | 10 | Baseline | paralle | 56 |
| # | sex | 2 | 6 | 0 | 0.25 | 1 | # | 10 | Baseline | paralle | 56 |
| # | time | 2 | 2 | 0 | 0.5 | 1 | # | 10 | Baseline | paralle | 56 |
| # | interaction | 2 | 5 | 0 | 0.285714286 | 1 | # | 10 | Baseline | paralle | 56 |
| # | disease | 4 | 2 | 0 | 0.666666667 | 1 | # | 10 | Baseline | ancom | 56 |
| # | sex | 2 | 0 | 0 | 1 | 1 | # | 10 | Baseline | ancom | 56 |
| # | time | 2 | 1 | 0 | 0.666666667 | 1 | # | 10 | Baseline | ancom | 56 |
| # | interaction | 0 | 0 | 2 | 0 | 0 | # | 10 | Baseline | ancom | 56 |
| # | disease | 4 | 21 | 0 | 0.16 | 1 | # | 10 | Baseline | aldex2 | 56 |
| # | sex | 2 | 5 | 0 | 0.285714286 | 1 | # | 10 | Baseline | aldex2 | 56 |
| # | time | 2 | 12 | 0 | 0.142857143 | 1 | # | 10 | Baseline | aldex2 | 56 |
| # | interaction | 0 | 0 | 2 | 0 | 0 | # | 10 | Baseline | aldex2 | 56 |
| # | disease | 4 | 0 | 0 | 1 | 1 | # | 10 | Small | paralle | 56 |
| # | sex | 2 | 0 | 0 | 1 | 1 | # | 10 | Small | paralle | 56 |
| # | time | 2 | 0 | 0 | 1 | 1 | # | 10 | Small | paralle | 56 |
| # | interaction | 2 | 0 | 0 | 1 | 1 | # | 10 | Small | paralle | 56 |
| # | disease | 4 | 1 | 0 | 0.8 | 1 | # | 10 | Small | ancom | 56 |
| # | sex | 2 | 0 | 0 | 1 | 1 | # | 10 | Small | ancom | 56 |
| # | time | 2 | 1 | 0 | 0.666666667 | 1 | # | 10 | Small | ancom | 56 |
| # | interaction | 0 | 0 | 2 | 0 | 0 | # | 10 | Small | ancom | 56 |
| # | disease | 4 | 2 | 0 | 0.666666667 | 1 | # | 10 | Small | aldex2 | 56 |
| # | sex | 2 | 0 | 0 | 1 | 1 | # | 10 | Small | aldex2 | 56 |
| # | time | 2 | 1 | 0 | 0.666666667 | 1 | # | 10 | Small | aldex2 | 56 |
| # | interaction | 0 | 0 | 2 | 0 | 0 | # | 10 | Small | aldex2 | 56 |
| # | disease | 4 | 6 | 0 | 0.4 | 1 | # | 10 | Baseline | paralle | 56 |
| # | sex | 2 | 8 | 0 | 0.2 | 1 | # | 10 | Baseline | paralle | 56 |
| # | time | 2 | 6 | 0 | 0.25 | 1 | # | 10 | Baseline | paralle | 56 |
| # | interaction | 2 | 7 | 0 | 0.222222222 | 1 | # | 10 | Baseline | paralle | 56 |
| # | disease | 4 | 2 | 0 | 0.666666667 | 1 | # | 10 | Baseline | ancom | 56 |
| # | sex | 2 | 0 | 0 | 1 | 1 | # | 10 | Baseline | ancom | 56 |
| # | time | 2 | 1 | 0 | 0.666666667 | 1 | # | 10 | Baseline | ancom | 56 |
| # | interaction | 0 | 0 | 2 | 0 | 0 | # | 10 | Baseline | ancom | 56 |
| # | disease | 4 | 27 | 0 | 0.129032258 | 1 | # | 10 | Baseline | aldex2 | 56 |
| # | sex | 2 | 4 | 0 | 0.333333333 | 1 | # | 10 | Baseline | aldex2 | 56 |
| # | time | 2 | 13 | 0 | 0.133333333 | 1 | # | 10 | Baseline | aldex2 | 56 |
| # | interaction | 0 | 0 | 2 | 0 | 0 | # | 10 | Baseline | aldex2 | 56 |
| # | disease | 4 | 0 | 0 | 1 | 1 | # | 10 | Small | paralle | 56 |
| # | sex | 2 | 0 | 0 | 1 | 1 | # | 10 | Small | paralle | 56 |
| # | time | 2 | 0 | 0 | 1 | 1 | # | 10 | Small | paralle | 56 |
| # | interaction | 2 | 0 | 0 | 1 | 1 | # | 10 | Small | paralle | 56 |
| # | disease | 4 | 2 | 0 | 0.666666667 | 1 | # | 10 | Small | ancom | 56 |
| # | sex | 2 | 0 | 0 | 1 | 1 | # | 10 | Small | ancom | 56 |
| # | time | 2 | 1 | 0 | 0.666666667 | 1 | # | 10 | Small | ancom | 56 |
| # | interaction | 0 | 0 | 2 | 0 | 0 | # | 10 | Small | ancom | 56 |
| # | disease | 4 | 1 | 0 | 0.8 | 1 | # | 10 | Small | aldex2 | 56 |
| # | sex | 2 | 0 | 0 | 1 | 1 | # | 10 | Small | aldex2 | 56 |
| # | time | 2 | 2 | 0 | 0.5 | 1 | # | 10 | Small | aldex2 | 56 |
| # | interaction | 0 | 0 | 2 | 0 | 0 | # | 10 | Small | aldex2 | 56 |
| # | disease | 7 | 10 | 0 | 0.411764706 | 1 | # | 20 | Baseline | paralle | 56 |
| # | sex | 4 | 9 | 0 | 0.307692308 | 1 | # | 20 | Baseline | paralle | 56 |
| # | time | 5 | 9 | 0 | 0.357142857 | 1 | # | 20 | Baseline | paralle | 56 |
| # | interaction | 3 | 9 | 1 | 0.25 | 0.75 | # | 20 | Baseline | paralle | 56 |
| # | disease | 7 | 1 | 0 | 0.875 | 1 | # | 20 | Baseline | ancom | 56 |
| # | sex | 4 | 0 | 0 | 1 | 1 | # | 20 | Baseline | ancom | 56 |
| # | time | 5 | 1 | 0 | 0.833333333 | 1 | # | 20 | Baseline | ancom | 56 |
| # | interaction | 0 | 0 | 4 | 0 | 0 | # | 20 | Baseline | ancom | 56 |
| # | disease | 7 | 18 | 0 | 0.28 | 1 | # | 20 | Baseline | aldex2 | 56 |
| # | sex | 4 | 4 | 0 | 0.5 | 1 | # | 20 | Baseline | aldex2 | 56 |
| # | time | 5 | 15 | 0 | 0.25 | 1 | # | 20 | Baseline | aldex2 | 56 |
| # | interaction | 0 | 0 | 4 | 0 | 0 | # | 20 | Baseline | aldex2 | 56 |
| # | disease | 7 | 0 | 0 | 1 | 1 | # | 20 | Small | paralle | 56 |
| # | sex | 4 | 2 | 0 | 0.666666667 | 1 | # | 20 | Small | paralle | 56 |
| # | time | 5 | 0 | 0 | 1 | 1 | # | 20 | Small | paralle | 56 |
| # | interaction | 0 | 0 | 4 | 0 | 0 | # | 20 | Small | paralle | 56 |
| # | disease | 7 | 0 | 0 | 1 | 1 | # | 20 | Small | ancom | 56 |
| # | sex | 4 | 0 | 0 | 1 | 1 | # | 20 | Small | ancom | 56 |
| # | time | 5 | 1 | 0 | 0.833333333 | 1 | # | 20 | Small | ancom | 56 |
| # | interaction | 0 | 0 | 4 | 0 | 0 | # | 20 | Small | ancom | 56 |
| # | disease | 7 | 0 | 0 | 1 | 1 | # | 20 | Small | aldex2 | 56 |
| # | sex | 4 | 0 | 0 | 1 | 1 | # | 20 | Small | aldex2 | 56 |
| # | time | 5 | 1 | 0 | 0.833333333 | 1 | # | 20 | Small | aldex2 | 56 |
| # | interaction | 0 | 0 | 4 | 0 | 0 | # | 20 | Small | aldex2 | 56 |
| # | disease | 7 | 11 | 0 | 0.388888889 | 1 | # | 20 | Baseline | paralle | 56 |
| # | sex | 4 | 12 | 0 | 0.25 | 1 | # | 20 | Baseline | paralle | 56 |
| # | time | 5 | 7 | 0 | 0.416666667 | 1 | # | 20 | Baseline | paralle | 56 |
| # | interaction | 4 | 6 | 0 | 0.4 | 1 | # | 20 | Baseline | paralle | 56 |
| # | disease | 7 | 2 | 0 | 0.777777778 | 1 | # | 20 | Baseline | ancom | 56 |
| # | sex | 4 | 0 | 0 | 1 | 1 | # | 20 | Baseline | ancom | 56 |
| # | time | 5 | 2 | 0 | 0.714285714 | 1 | # | 20 | Baseline | ancom | 56 |
| # | interaction | 0 | 0 | 4 | 0 | 0 | # | 20 | Baseline | ancom | 56 |
| # | disease | 7 | 37 | 0 | 0.159090909 | 1 | # | 20 | Baseline | aldex2 | 56 |
| # | sex | 4 | 16 | 0 | 0.2 | 1 | # | 20 | Baseline | aldex2 | 56 |
| # | time | 5 | 27 | 0 | 0.15625 | 1 | # | 20 | Baseline | aldex2 | 56 |
| # | interaction | 0 | 0 | 4 | 0 | 0 | # | 20 | Baseline | aldex2 | 56 |
| # | disease | 7 | 0 | 0 | 1 | 1 | # | 20 | Small | paralle | 56 |
| # | sex | 4 | 1 | 0 | 0.8 | 1 | # | 20 | Small | paralle | 56 |
| # | time | 5 | 1 | 0 | 0.833333333 | 1 | # | 20 | Small | paralle | 56 |
| # | interaction | 2 | 1 | 2 | 0.666666667 | 0.5 | # | 20 | Small | paralle | 56 |
| # | disease | 7 | 2 | 0 | 0.777777778 | 1 | # | 20 | Small | ancom | 56 |
| # | sex | 4 | 0 | 0 | 1 | 1 | # | 20 | Small | ancom | 56 |
| # | time | 5 | 2 | 0 | 0.714285714 | 1 | # | 20 | Small | ancom | 56 |
| # | interaction | 0 | 0 | 4 | 0 | 0 | # | 20 | Small | ancom | 56 |
| # | disease | 7 | 2 | 0 | 0.777777778 | 1 | # | 20 | Small | aldex2 | 56 |
| # | sex | 4 | 0 | 0 | 1 | 1 | # | 20 | Small | aldex2 | 56 |
| # | time | 5 | 2 | 0 | 0.714285714 | 1 | # | 20 | Small | aldex2 | 56 |
| # | interaction | 0 | 0 | 4 | 0 | 0 | # | 20 | Small | aldex2 | 56 |
| # | disease | 7 | 12 | 0 | 0.368421053 | 1 | # | 20 | Baseline | paralle | 56 |
| # | sex | 4 | 11 | 0 | 0.266666667 | 1 | # | 20 | Baseline | paralle | 56 |
| # | time | 5 | 13 | 0 | 0.277777778 | 1 | # | 20 | Baseline | paralle | 56 |
| # | interaction | 4 | 8 | 0 | 0.333333333 | 1 | # | 20 | Baseline | paralle | 56 |
| # | disease | 7 | 4 | 0 | 0.636363636 | 1 | # | 20 | Baseline | ancom | 56 |
| # | sex | 4 | 0 | 0 | 1 | 1 | # | 20 | Baseline | ancom | 56 |
| # | time | 5 | 4 | 0 | 0.555555556 | 1 | # | 20 | Baseline | ancom | 56 |
| # | interaction | 0 | 0 | 4 | 0 | 0 | # | 20 | Baseline | ancom | 56 |
| # | disease | 7 | 48 | 0 | 0.127272727 | 1 | # | 20 | Baseline | aldex2 | 56 |
| # | sex | 4 | 11 | 0 | 0.266666667 | 1 | # | 20 | Baseline | aldex2 | 56 |
| # | time | 5 | 32 | 0 | 0.135135135 | 1 | # | 20 | Baseline | aldex2 | 56 |
| # | interaction | 0 | 0 | 4 | 0 | 0 | # | 20 | Baseline | aldex2 | 56 |
| # | disease | 7 | 1 | 0 | 0.875 | 1 | # | 20 | Small | paralle | 56 |
| # | sex | 4 | 1 | 0 | 0.8 | 1 | # | 20 | Small | paralle | 56 |
| # | time | 5 | 1 | 0 | 0.833333333 | 1 | # | 20 | Small | paralle | 56 |
| # | interaction | 2 | 0 | 2 | 1 | 0.5 | # | 20 | Small | paralle | 56 |
| # | disease | 7 | 4 | 0 | 0.636363636 | 1 | # | 20 | Small | ancom | 56 |
| # | sex | 4 | 0 | 0 | 1 | 1 | # | 20 | Small | ancom | 56 |
| # | time | 5 | 4 | 0 | 0.555555556 | 1 | # | 20 | Small | ancom | 56 |
| # | interaction | 0 | 0 | 4 | 0 | 0 | # | 20 | Small | ancom | 56 |
| # | disease | 7 | 4 | 0 | 0.636363636 | 1 | # | 20 | Small | aldex2 | 56 |
| # | sex | 4 | 1 | 0 | 0.8 | 1 | # | 20 | Small | aldex2 | 56 |
| # | time | 5 | 2 | 0 | 0.714285714 | 1 | # | 20 | Small | aldex2 | 56 |
| # | interaction | 0 | 0 | 4 | 0 | 0 | # | 20 | Small | aldex2 | 56 |
| # | disease | 7 | 10 | 0 | 0.411764706 | 1 | # | 20 | Baseline | paralle | 56 |
| # | sex | 4 | 6 | 0 | 0.4 | 1 | # | 20 | Baseline | paralle | 56 |
| # | time | 5 | 5 | 0 | 0.5 | 1 | # | 20 | Baseline | paralle | 56 |
| # | interaction | 4 | 5 | 0 | 0.444444444 | 1 | # | 20 | Baseline | paralle | 56 |
| # | disease | 7 | 4 | 0 | 0.636363636 | 1 | # | 20 | Baseline | ancom | 56 |
| # | sex | 4 | 0 | 0 | 1 | 1 | # | 20 | Baseline | ancom | 56 |
| # | time | 5 | 4 | 0 | 0.555555556 | 1 | # | 20 | Baseline | ancom | 56 |
| # | interaction | 0 | 0 | 4 | 0 | 0 | # | 20 | Baseline | ancom | 56 |
| # | disease | 7 | 61 | 0 | 0.102941176 | 1 | # | 20 | Baseline | aldex2 | 56 |
| # | sex | 4 | 17 | 0 | 0.19047619 | 1 | # | 20 | Baseline | aldex2 | 56 |
| # | time | 5 | 39 | 0 | 0.113636364 | 1 | # | 20 | Baseline | aldex2 | 56 |
| # | interaction | 0 | 0 | 4 | 0 | 0 | # | 20 | Baseline | aldex2 | 56 |
| # | disease | 7 | 0 | 0 | 1 | 1 | # | 20 | Small | paralle | 56 |
| # | sex | 4 | 0 | 0 | 1 | 1 | # | 20 | Small | paralle | 56 |
| # | time | 5 | 1 | 0 | 0.833333333 | 1 | # | 20 | Small | paralle | 56 |
| # | interaction | 3 | 0 | 1 | 1 | 0.75 | # | 20 | Small | paralle | 56 |
| # | disease | 7 | 4 | 0 | 0.636363636 | 1 | # | 20 | Small | ancom | 56 |
| # | sex | 4 | 0 | 0 | 1 | 1 | # | 20 | Small | ancom | 56 |
| # | time | 5 | 4 | 0 | 0.555555556 | 1 | # | 20 | Small | ancom | 56 |
| # | interaction | 0 | 0 | 4 | 0 | 0 | # | 20 | Small | ancom | 56 |
| # | disease | 7 | 7 | 0 | 0.5 | 1 | # | 20 | Small | aldex2 | 56 |
| # | sex | 4 | 0 | 0 | 1 | 1 | # | 20 | Small | aldex2 | 56 |
| # | time | 5 | 5 | 0 | 0.5 | 1 | # | 20 | Small | aldex2 | 56 |
| # | interaction | 0 | 0 | 4 | 0 | 0 | # | 20 | Small | aldex2 | 56 |
| # | disease | 4 | 13 | 0 | 0.235294118 | 1 | # | 10 | Baseline | paralle | 57 |
| # | sex | 2 | 8 | 0 | 0.2 | 1 | # | 10 | Baseline | paralle | 57 |
| # | time | 2 | 14 | 0 | 0.125 | 1 | # | 10 | Baseline | paralle | 57 |
| # | interaction | 1 | 5 | 1 | 0.166666667 | 0.5 | # | 10 | Baseline | paralle | 57 |
| # | disease | 4 | 0 | 0 | 1 | 1 | # | 10 | Baseline | ancom | 57 |
| # | sex | 2 | 0 | 0 | 1 | 1 | # | 10 | Baseline | ancom | 57 |
| # | time | 2 | 0 | 0 | 1 | 1 | # | 10 | Baseline | ancom | 57 |
| # | interaction | 0 | 0 | 2 | 0 | 0 | # | 10 | Baseline | ancom | 57 |
| # | disease | 4 | 10 | 0 | 0.285714286 | 1 | # | 10 | Baseline | aldex2 | 57 |
| # | sex | 2 | 3 | 0 | 0.4 | 1 | # | 10 | Baseline | aldex2 | 57 |
| # | time | 2 | 8 | 0 | 0.2 | 1 | # | 10 | Baseline | aldex2 | 57 |
| # | interaction | 0 | 0 | 2 | 0 | 0 | # | 10 | Baseline | aldex2 | 57 |
| # | disease | 4 | 0 | 0 | 1 | 1 | # | 10 | Small | paralle | 57 |
| # | sex | 2 | 0 | 0 | 1 | 1 | # | 10 | Small | paralle | 57 |
| # | time | 2 | 0 | 0 | 1 | 1 | # | 10 | Small | paralle | 57 |
| # | interaction | 1 | 0 | 1 | 1 | 0.5 | # | 10 | Small | paralle | 57 |
| # | disease | 4 | 0 | 0 | 1 | 1 | # | 10 | Small | ancom | 57 |
| # | sex | 2 | 0 | 0 | 1 | 1 | # | 10 | Small | ancom | 57 |
| # | time | 2 | 0 | 0 | 1 | 1 | # | 10 | Small | ancom | 57 |
| # | interaction | 0 | 0 | 2 | 0 | 0 | # | 10 | Small | ancom | 57 |
| # | disease | 4 | 1 | 0 | 0.8 | 1 | # | 10 | Small | aldex2 | 57 |
| # | sex | 2 | 0 | 0 | 1 | 1 | # | 10 | Small | aldex2 | 57 |
| # | time | 2 | 0 | 0 | 1 | 1 | # | 10 | Small | aldex2 | 57 |
| # | interaction | 0 | 0 | 2 | 0 | 0 | # | 10 | Small | aldex2 | 57 |
| # | disease | 4 | 8 | 0 | 0.333333333 | 1 | # | 10 | Baseline | paralle | 57 |
| # | sex | 2 | 5 | 0 | 0.285714286 | 1 | # | 10 | Baseline | paralle | 57 |
| # | time | 2 | 5 | 0 | 0.285714286 | 1 | # | 10 | Baseline | paralle | 57 |
| # | interaction | 2 | 2 | 0 | 0.5 | 1 | # | 10 | Baseline | paralle | 57 |
| # | disease | 4 | 0 | 0 | 1 | 1 | # | 10 | Baseline | ancom | 57 |

| | | | | | | | | | | | |
|---|---|---|---|---|---|---|---|---|---|---|---|
| # | sex | 2 | 0 | 0 | 1 | 1 | # | 10 | Baseline | ancom | 57 |
| # | time | 2 | 1 | 0 | 0.666666667 | 1 | # | 10 | Baseline | ancom | 57 |
| # | interaction | 0 | 0 | 2 | 0 | 0 | # | 10 | Baseline | ancom | 57 |
| # | disease | 4 | 17 | 0 | 0.19047619 | 1 | # | 10 | Baseline | aldex2 | 57 |
| # | sex | 2 | 5 | 0 | 0.285714286 | 1 | # | 10 | Baseline | aldex2 | 57 |
| # | time | 2 | 10 | 0 | 0.166666667 | 1 | # | 10 | Baseline | aldex2 | 57 |
| # | interaction | 0 | 0 | 2 | 0 | 0 | # | 10 | Baseline | aldex2 | 57 |
| # | disease | 4 | 1 | 0 | 0.8 | 1 | # | 10 | Small | paralle | 57 |
| # | sex | 2 | 0 | 0 | 1 | 1 | # | 10 | Small | paralle | 57 |
| # | time | 2 | 0 | 0 | 1 | 1 | # | 10 | Small | paralle | 57 |
| # | interaction | 1 | 0 | 1 | 1 | 0.5 | # | 10 | Small | paralle | 57 |
| # | disease | 4 | 0 | 0 | 1 | 1 | # | 10 | Small | ancom | 57 |
| # | sex | 2 | 0 | 0 | 1 | 1 | # | 10 | Small | ancom | 57 |
| # | time | 2 | 1 | 0 | 0.666666667 | 1 | # | 10 | Small | ancom | 57 |
| # | interaction | 0 | 0 | 2 | 0 | 0 | # | 10 | Small | ancom | 57 |
| # | disease | 4 | 0 | 0 | 1 | 1 | # | 10 | Small | aldex2 | 57 |
| # | sex | 2 | 1 | 0 | 0.666666667 | 1 | # | 10 | Small | aldex2 | 57 |
| # | time | 2 | 1 | 0 | 0.666666667 | 1 | # | 10 | Small | aldex2 | 57 |
| # | interaction | 0 | 0 | 2 | 0 | 0 | # | 10 | Small | aldex2 | 57 |
| # | disease | 4 | 8 | 0 | 0.333333333 | 1 | # | 10 | Baseline | paralle | 57 |
| # | sex | 2 | 5 | 0 | 0.285714286 | 1 | # | 10 | Baseline | paralle | 57 |
| # | time | 2 | 4 | 0 | 0.333333333 | 1 | # | 10 | Baseline | paralle | 57 |
| # | interaction | 2 | 2 | 0 | 0.5 | 1 | # | 10 | Baseline | paralle | 57 |
| # | disease | 4 | 2 | 0 | 0.666666667 | 1 | # | 10 | Baseline | ancom | 57 |
| # | sex | 2 | 0 | 0 | 1 | 1 | # | 10 | Baseline | ancom | 57 |
| # | time | 2 | 1 | 0 | 0.666666667 | 1 | # | 10 | Baseline | ancom | 57 |
| # | interaction | 0 | 0 | 2 | 0 | 0 | # | 10 | Baseline | ancom | 57 |
| # | disease | 4 | 21 | 0 | 0.16 | 1 | # | 10 | Baseline | aldex2 | 57 |
| # | sex | 2 | 6 | 0 | 0.25 | 1 | # | 10 | Baseline | aldex2 | 57 |
| # | time | 2 | 12 | 0 | 0.142857143 | 1 | # | 10 | Baseline | aldex2 | 57 |
| # | interaction | 0 | 0 | 2 | 0 | 0 | # | 10 | Baseline | aldex2 | 57 |
| # | disease | 4 | 0 | 0 | 1 | 1 | # | 10 | Small | paralle | 57 |
| # | sex | 2 | 0 | 0 | 1 | 1 | # | 10 | Small | paralle | 57 |
| # | time | 2 | 0 | 0 | 1 | 1 | # | 10 | Small | paralle | 57 |
| # | interaction | 2 | 0 | 0 | 1 | 1 | # | 10 | Small | paralle | 57 |
| # | disease | 4 | 2 | 0 | 0.666666667 | 1 | # | 10 | Small | ancom | 57 |
| # | sex | 2 | 0 | 0 | 1 | 1 | # | 10 | Small | ancom | 57 |
| # | time | 2 | 1 | 0 | 0.666666667 | 1 | # | 10 | Small | ancom | 57 |
| # | interaction | 0 | 0 | 2 | 0 | 0 | # | 10 | Small | ancom | 57 |
| # | disease | 4 | 1 | 0 | 0.8 | 1 | # | 10 | Small | aldex2 | 57 |
| # | sex | 2 | 0 | 0 | 1 | 1 | # | 10 | Small | aldex2 | 57 |
| # | time | 2 | 1 | 0 | 0.666666667 | 1 | # | 10 | Small | aldex2 | 57 |
| # | interaction | 0 | 0 | 2 | 0 | 0 | # | 10 | Small | aldex2 | 57 |
| # | disease | 4 | 3 | 0 | 0.571428571 | 1 | # | 10 | Baseline | paralle | 57 |
| # | sex | 2 | 8 | 0 | 0.2 | 1 | # | 10 | Baseline | paralle | 57 |
| # | time | 2 | 3 | 0 | 0.4 | 1 | # | 10 | Baseline | paralle | 57 |
| # | interaction | 2 | 1 | 0 | 0.666666667 | 1 | # | 10 | Baseline | paralle | 57 |
| # | disease | 4 | 2 | 0 | 0.666666667 | 1 | # | 10 | Baseline | ancom | 57 |
| # | sex | 2 | 0 | 0 | 1 | 1 | # | 10 | Baseline | ancom | 57 |
| # | time | 2 | 2 | 0 | 0.5 | 1 | # | 10 | Baseline | ancom | 57 |
| # | interaction | 0 | 0 | 2 | 0 | 0 | # | 10 | Baseline | ancom | 57 |
| # | disease | 4 | 27 | 0 | 0.129032258 | 1 | # | 10 | Baseline | aldex2 | 57 |
| # | sex | 2 | 8 | 0 | 0.2 | 1 | # | 10 | Baseline | aldex2 | 57 |
| # | time | 2 | 12 | 0 | 0.142857143 | 1 | # | 10 | Baseline | aldex2 | 57 |
| # | interaction | 0 | 0 | 2 | 0 | 0 | # | 10 | Baseline | aldex2 | 57 |
| # | disease | 4 | 0 | 0 | 1 | 1 | # | 10 | Small | paralle | 57 |
| # | sex | 2 | 0 | 0 | 1 | 1 | # | 10 | Small | paralle | 57 |
| # | time | 2 | 0 | 0 | 1 | 1 | # | 10 | Small | paralle | 57 |
| # | interaction | 2 | 0 | 0 | 1 | 1 | # | 10 | Small | paralle | 57 |
| # | disease | 4 | 2 | 0 | 0.666666667 | 1 | # | 10 | Small | ancom | 57 |
| # | sex | 2 | 0 | 0 | 1 | 1 | # | 10 | Small | ancom | 57 |
| # | time | 2 | 1 | 0 | 0.666666667 | 1 | # | 10 | Small | ancom | 57 |
| # | interaction | 0 | 0 | 2 | 0 | 0 | # | 10 | Small | ancom | 57 |
| # | disease | 4 | 2 | 0 | 0.666666667 | 1 | # | 10 | Small | aldex2 | 57 |
| # | sex | 2 | 0 | 0 | 1 | 1 | # | 10 | Small | aldex2 | 57 |
| # | time | 2 | 2 | 0 | 0.5 | 1 | # | 10 | Small | aldex2 | 57 |
| # | interaction | 0 | 0 | 2 | 0 | 0 | # | 10 | Small | aldex2 | 57 |
| # | disease | 7 | 7 | 0 | 0.5 | 1 | # | 20 | Baseline | paralle | 57 |
| # | sex | 4 | 8 | 0 | 0.333333333 | 1 | # | 20 | Baseline | paralle | 57 |
| # | time | 5 | 10 | 0 | 0.333333333 | 1 | # | 20 | Baseline | paralle | 57 |
| # | interaction | 4 | 15 | 0 | 0.210526316 | 1 | # | 20 | Baseline | paralle | 57 |
| # | disease | 7 | 1 | 0 | 0.875 | 1 | # | 20 | Baseline | ancom | 57 |
| # | sex | 4 | 0 | 0 | 1 | 1 | # | 20 | Baseline | ancom | 57 |
| # | time | 5 | 3 | 0 | 0.625 | 1 | # | 20 | Baseline | ancom | 57 |
| # | interaction | 0 | 0 | 4 | 0 | 0 | # | 20 | Baseline | ancom | 57 |
| # | disease | 7 | 13 | 0 | 0.35 | 1 | # | 20 | Baseline | aldex2 | 57 |
| # | sex | 4 | 4 | 0 | 0.5 | 1 | # | 20 | Baseline | aldex2 | 57 |
| # | time | 5 | 18 | 0 | 0.217391304 | 1 | # | 20 | Baseline | aldex2 | 57 |
| # | interaction | 0 | 0 | 4 | 0 | 0 | # | 20 | Baseline | aldex2 | 57 |
| # | disease | 7 | 0 | 0 | 1 | 1 | # | 20 | Small | paralle | 57 |
| # | sex | 4 | 0 | 0 | 1 | 1 | # | 20 | Small | paralle | 57 |
| # | time | 5 | 0 | 0 | 1 | 1 | # | 20 | Small | paralle | 57 |
| # | interaction | 0 | 0 | 4 | 0 | 0 | # | 20 | Small | paralle | 57 |
| # | disease | 7 | 1 | 0 | 0.875 | 1 | # | 20 | Small | ancom | 57 |
| # | sex | 4 | 0 | 0 | 1 | 1 | # | 20 | Small | ancom | 57 |
| # | time | 5 | 2 | 0 | 0.714285714 | 1 | # | 20 | Small | ancom | 57 |
| # | interaction | 0 | 0 | 4 | 0 | 0 | # | 20 | Small | ancom | 57 |
| # | disease | 7 | 1 | 0 | 0.875 | 1 | # | 20 | Small | aldex2 | 57 |
| # | sex | 4 | 0 | 0 | 1 | 1 | # | 20 | Small | aldex2 | 57 |
| # | time | 5 | 3 | 0 | 0.625 | 1 | # | 20 | Small | aldex2 | 57 |
| # | interaction | 0 | 0 | 4 | 0 | 0 | # | 20 | Small | aldex2 | 57 |
| # | disease | 7 | 11 | 0 | 0.388888889 | 1 | # | 20 | Baseline | paralle | 57 |
| # | sex | 4 | 5 | 0 | 0.444444444 | 1 | # | 20 | Baseline | paralle | 57 |
| # | time | 5 | 9 | 0 | 0.357142857 | 1 | # | 20 | Baseline | paralle | 57 |
| # | interaction | 4 | 6 | 0 | 0.4 | 1 | # | 20 | Baseline | paralle | 57 |
| # | disease | 7 | 3 | 0 | 0.7 | 1 | # | 20 | Baseline | ancom | 57 |
| # | sex | 4 | 0 | 0 | 1 | 1 | # | 20 | Baseline | ancom | 57 |
| # | time | 5 | 4 | 0 | 0.555555556 | 1 | # | 20 | Baseline | ancom | 57 |
| # | interaction | 0 | 0 | 4 | 0 | 0 | # | 20 | Baseline | ancom | 57 |
| # | disease | 7 | 28 | 0 | 0.2 | 1 | # | 20 | Baseline | aldex2 | 57 |
| # | sex | 4 | 9 | 0 | 0.307692308 | 1 | # | 20 | Baseline | aldex2 | 57 |
| # | time | 5 | 27 | 0 | 0.15625 | 1 | # | 20 | Baseline | aldex2 | 57 |
| # | interaction | 0 | 0 | 4 | 0 | 0 | # | 20 | Baseline | aldex2 | 57 |
| # | disease | 7 | 0 | 0 | 1 | 1 | # | 20 | Small | paralle | 57 |
| # | sex | 4 | 0 | 0 | 1 | 1 | # | 20 | Small | paralle | 57 |
| # | time | 5 | 0 | 0 | 1 | 1 | # | 20 | Small | paralle | 57 |
| # | interaction | 3 | 0 | 1 | 1 | 0.75 | # | 20 | Small | paralle | 57 |
| # | disease | 7 | 3 | 0 | 0.7 | 1 | # | 20 | Small | ancom | 57 |
| # | sex | 4 | 0 | 0 | 1 | 1 | # | 20 | Small | ancom | 57 |
| # | time | 5 | 4 | 0 | 0.555555556 | 1 | # | 20 | Small | ancom | 57 |
| # | interaction | 0 | 0 | 4 | 0 | 0 | # | 20 | Small | ancom | 57 |
| # | disease | 7 | 3 | 0 | 0.7 | 1 | # | 20 | Small | aldex2 | 57 |
| # | sex | 4 | 0 | 0 | 1 | 1 | # | 20 | Small | aldex2 | 57 |
| # | time | 5 | 5 | 0 | 0.5 | 1 | # | 20 | Small | aldex2 | 57 |
| # | interaction | 0 | 0 | 4 | 0 | 0 | # | 20 | Small | aldex2 | 57 |
| # | disease | 7 | 9 | 0 | 0.4375 | 1 | # | 20 | Baseline | paralle | 57 |
| # | sex | 4 | 4 | 0 | 0.5 | 1 | # | 20 | Baseline | paralle | 57 |
| # | time | 5 | 6 | 0 | 0.454545455 | 1 | # | 20 | Baseline | paralle | 57 |
| # | interaction | 4 | 1 | 0 | 0.8 | 1 | # | 20 | Baseline | paralle | 57 |
| # | disease | 7 | 4 | 0 | 0.636363636 | 1 | # | 20 | Baseline | ancom | 57 |
| # | sex | 4 | 0 | 0 | 1 | 1 | # | 20 | Baseline | ancom | 57 |
| # | time | 5 | 4 | 0 | 0.555555556 | 1 | # | 20 | Baseline | ancom | 57 |
| # | interaction | 0 | 0 | 4 | 0 | 0 | # | 20 | Baseline | ancom | 57 |
| # | disease | 7 | 47 | 0 | 0.12962963 | 1 | # | 20 | Baseline | aldex2 | 57 |
| # | sex | 4 | 12 | 0 | 0.25 | 1 | # | 20 | Baseline | aldex2 | 57 |
| # | time | 5 | 30 | 0 | 0.142857143 | 1 | # | 20 | Baseline | aldex2 | 57 |
| # | interaction | 0 | 0 | 4 | 0 | 0 | # | 20 | Baseline | aldex2 | 57 |
| # | disease | 7 | 0 | 0 | 1 | 1 | # | 20 | Small | paralle | 57 |
| # | sex | 4 | 0 | 0 | 1 | 1 | # | 20 | Small | paralle | 57 |
| # | time | 5 | 0 | 0 | 1 | 1 | # | 20 | Small | paralle | 57 |
| # | interaction | 4 | 0 | 0 | 1 | 1 | # | 20 | Small | paralle | 57 |
| # | disease | 7 | 4 | 0 | 0.636363636 | 1 | # | 20 | Small | ancom | 57 |
| # | sex | 4 | 0 | 0 | 1 | 1 | # | 20 | Small | ancom | 57 |
| # | time | 5 | 4 | 0 | 0.555555556 | 1 | # | 20 | Small | ancom | 57 |
| # | interaction | 0 | 0 | 4 | 0 | 0 | # | 20 | Small | ancom | 57 |
| # | disease | 7 | 4 | 0 | 0.636363636 | 1 | # | 20 | Small | aldex2 | 57 |
| # | sex | 4 | 1 | 0 | 0.8 | 1 | # | 20 | Small | aldex2 | 57 |
| # | time | 5 | 6 | 0 | 0.454545455 | 1 | # | 20 | Small | aldex2 | 57 |
| # | interaction | 0 | 0 | 4 | 0 | 0 | # | 20 | Small | aldex2 | 57 |
| # | disease | 7 | 11 | 0 | 0.388888889 | 1 | # | 20 | Baseline | paralle | 57 |
| # | sex | 4 | 7 | 0 | 0.363636364 | 1 | # | 20 | Baseline | paralle | 57 |
| # | time | 5 | 12 | 0 | 0.294117647 | 1 | # | 20 | Baseline | paralle | 57 |
| # | interaction | 4 | 3 | 0 | 0.571428571 | 1 | # | 20 | Baseline | paralle | 57 |
| # | disease | 7 | 4 | 0 | 0.636363636 | 1 | # | 20 | Baseline | ancom | 57 |
| # | sex | 4 | 0 | 0 | 1 | 1 | # | 20 | Baseline | ancom | 57 |
| # | time | 5 | 4 | 0 | 0.555555556 | 1 | # | 20 | Baseline | ancom | 57 |
| # | interaction | 0 | 0 | 4 | 0 | 0 | # | 20 | Baseline | ancom | 57 |
| # | disease | 7 | 58 | 0 | 0.107692308 | 1 | # | 20 | Baseline | aldex2 | 57 |
| # | sex | 4 | 18 | 0 | 0.181818182 | 1 | # | 20 | Baseline | aldex2 | 57 |
| # | time | 5 | 37 | 0 | 0.119047619 | 1 | # | 20 | Baseline | aldex2 | 57 |
| # | interaction | 0 | 0 | 4 | 0 | 0 | # | 20 | Baseline | aldex2 | 57 |
| # | disease | 7 | 0 | 0 | 1 | 1 | # | 20 | Small | paralle | 57 |
| # | sex | 4 | 0 | 0 | 1 | 1 | # | 20 | Small | paralle | 57 |
| # | time | 5 | 0 | 0 | 1 | 1 | # | 20 | Small | paralle | 57 |
| # | interaction | 4 | 0 | 0 | 1 | 1 | # | 20 | Small | paralle | 57 |
| # | disease | 7 | 4 | 0 | 0.636363636 | 1 | # | 20 | Small | ancom | 57 |
| # | sex | 4 | 0 | 0 | 1 | 1 | # | 20 | Small | ancom | 57 |
| # | time | 5 | 4 | 0 | 0.555555556 | 1 | # | 20 | Small | ancom | 57 |
| # | interaction | 0 | 0 | 4 | 0 | 0 | # | 20 | Small | ancom | 57 |
| # | disease | 7 | 7 | 0 | 0.5 | 1 | # | 20 | Small | aldex2 | 57 |
| # | sex | 4 | 1 | 0 | 0.8 | 1 | # | 20 | Small | aldex2 | 57 |
| # | time | 5 | 8 | 0 | 0.384615385 | 1 | # | 20 | Small | aldex2 | 57 |
| # | interaction | 0 | 0 | 4 | 0 | 0 | # | 20 | Small | aldex2 | 57 |
| # | disease | 4 | 15 | 0 | 0.210526316 | 1 | # | 10 | Baseline | paralle | 58 |
| # | sex | 2 | 10 | 0 | 0.166666667 | 1 | # | 10 | Baseline | paralle | 58 |
| # | time | 2 | 11 | 0 | 0.153846154 | 1 | # | 10 | Baseline | paralle | 58 |
| # | interaction | 2 | 11 | 0 | 0.153846154 | 1 | # | 10 | Baseline | paralle | 58 |
| # | disease | 4 | 0 | 0 | 1 | 1 | # | 10 | Baseline | ancom | 58 |
| # | sex | 2 | 0 | 0 | 1 | 1 | # | 10 | Baseline | ancom | 58 |
| # | time | 2 | 0 | 0 | 1 | 1 | # | 10 | Baseline | ancom | 58 |
| # | interaction | 0 | 0 | 2 | 0 | 0 | # | 10 | Baseline | ancom | 58 |
| # | disease | 4 | 13 | 0 | 0.235294118 | 1 | # | 10 | Baseline | aldex2 | 58 |
| # | sex | 2 | 3 | 0 | 0.4 | 1 | # | 10 | Baseline | aldex2 | 58 |
| # | time | 2 | 5 | 0 | 0.285714286 | 1 | # | 10 | Baseline | aldex2 | 58 |
| # | interaction | 0 | 0 | 2 | 0 | 0 | # | 10 | Baseline | aldex2 | 58 |
| # | disease | 4 | 0 | 0 | 1 | 1 | # | 10 | Small | paralle | 58 |
| # | sex | 2 | 1 | 0 | 0.666666667 | 1 | # | 10 | Small | paralle | 58 |
| # | time | 2 | 0 | 0 | 1 | 1 | # | 10 | Small | paralle | 58 |
| # | interaction | 1 | 0 | 1 | 1 | 0.5 | # | 10 | Small | paralle | 58 |
| # | disease | 4 | 0 | 0 | 1 | 1 | # | 10 | Small | ancom | 58 |
| # | sex | 2 | 0 | 0 | 1 | 1 | # | 10 | Small | ancom | 58 |
| # | time | 2 | 0 | 0 | 1 | 1 | # | 10 | Small | ancom | 58 |
| # | interaction | 0 | 0 | 2 | 0 | 0 | # | 10 | Small | ancom | 58 |
| # | disease | 4 | 0 | 0 | 1 | 1 | # | 10 | Small | aldex2 | 58 |
| # | sex | 2 | 0 | 0 | 1 | 1 | # | 10 | Small | aldex2 | 58 |
| # | time | 2 | 0 | 0 | 1 | 1 | # | 10 | Small | aldex2 | 58 |
| # | interaction | 0 | 0 | 2 | 0 | 0 | # | 10 | Small | aldex2 | 58 |
| # | disease | 4 | 11 | 0 | 0.266666667 | 1 | # | 10 | Baseline | paralle | 58 |
| # | sex | 2 | 1 | 0 | 0.666666667 | 1 | # | 10 | Baseline | paralle | 58 |
| # | time | 2 | 7 | 0 | 0.222222222 | 1 | # | 10 | Baseline | paralle | 58 |
| # | interaction | 2 | 9 | 0 | 0.181818182 | 1 | # | 10 | Baseline | paralle | 58 |
| # | disease | 4 | 2 | 0 | 0.666666667 | 1 | # | 10 | Baseline | ancom | 58 |
| # | sex | 2 | 0 | 0 | 1 | 1 | # | 10 | Baseline | ancom | 58 |
| # | time | 2 | 1 | 0 | 0.666666667 | 1 | # | 10 | Baseline | ancom | 58 |
| # | interaction | 0 | 0 | 2 | 0 | 0 | # | 10 | Baseline | ancom | 58 |
| # | disease | 4 | 14 | 0 | 0.222222222 | 1 | # | 10 | Baseline | aldex2 | 58 |
| # | sex | 2 | 5 | 0 | 0.285714286 | 1 | # | 10 | Baseline | aldex2 | 58 |
| # | time | 2 | 6 | 0 | 0.25 | 1 | # | 10 | Baseline | aldex2 | 58 |
| # | interaction | 0 | 0 | 2 | 0 | 0 | # | 10 | Baseline | aldex2 | 58 |
| # | disease | 4 | 0 | 0 | 1 | 1 | # | 10 | Small | paralle | 58 |
| # | sex | 2 | 0 | 0 | 1 | 1 | # | 10 | Small | paralle | 58 |
| # | time | 2 | 0 | 0 | 1 | 1 | # | 10 | Small | paralle | 58 |
| # | interaction | 2 | 1 | 0 | 0.666666667 | 1 | # | 10 | Small | paralle | 58 |
| # | disease | 4 | 1 | 0 | 0.8 | 1 | # | 10 | Small | ancom | 58 |
| # | sex | 2 | 0 | 0 | 1 | 1 | # | 10 | Small | ancom | 58 |
| # | time | 2 | 1 | 0 | 0.666666667 | 1 | # | 10 | Small | ancom | 58 |
| # | interaction | 0 | 0 | 2 | 0 | 0 | # | 10 | Small | ancom | 58 |
| # | disease | 4 | 1 | 0 | 0.8 | 1 | # | 10 | Small | aldex2 | 58 |
| # | sex | 2 | 0 | 0 | 1 | 1 | # | 10 | Small | aldex2 | 58 |
| # | time | 2 | 1 | 0 | 0.666666667 | 1 | # | 10 | Small | aldex2 | 58 |
| # | interaction | 0 | 0 | 2 | 0 | 0 | # | 10 | Small | aldex2 | 58 |
| # | disease | 4 | 8 | 0 | 0.333333333 | 1 | # | 10 | Baseline | paralle | 58 |
| # | sex | 2 | 1 | 0 | 0.666666667 | 1 | # | 10 | Baseline | paralle | 58 |
| # | time | 2 | 8 | 0 | 0.2 | 1 | # | 10 | Baseline | paralle | 58 |
| # | interaction | 2 | 7 | 0 | 0.222222222 | 1 | # | 10 | Baseline | paralle | 58 |
| # | disease | 4 | 2 | 0 | 0.666666667 | 1 | # | 10 | Baseline | ancom | 58 |
| # | sex | 2 | 0 | 0 | 1 | 1 | # | 10 | Baseline | ancom | 58 |
| # | time | 2 | 1 | 0 | 0.666666667 | 1 | # | 10 | Baseline | ancom | 58 |
| # | interaction | 0 | 0 | 2 | 0 | 0 | # | 10 | Baseline | ancom | 58 |
| # | disease | 4 | 20 | 0 | 0.166666667 | 1 | # | 10 | Baseline | aldex2 | 58 |
| # | sex | 2 | 14 | 0 | 0.125 | 1 | # | 10 | Baseline | aldex2 | 58 |
| # | time | 2 | 8 | 0 | 0.2 | 1 | # | 10 | Baseline | aldex2 | 58 |
| # | interaction | 0 | 0 | 2 | 0 | 0 | # | 10 | Baseline | aldex2 | 58 |
| # | disease | 4 | 0 | 0 | 1 | 1 | # | 10 | Small | paralle | 58 |
| # | sex | 2 | 0 | 0 | 1 | 1 | # | 10 | Small | paralle | 58 |
| # | time | 2 | 0 | 0 | 1 | 1 | # | 10 | Small | paralle | 58 |
| # | interaction | 2 | 0 | 0 | 1 | 1 | # | 10 | Small | paralle | 58 |
| # | disease | 4 | 2 | 0 | 0.666666667 | 1 | # | 10 | Small | ancom | 58 |
| # | sex | 2 | 0 | 0 | 1 | 1 | # | 10 | Small | ancom | 58 |
| # | time | 2 | 1 | 0 | 0.666666667 | 1 | # | 10 | Small | ancom | 58 |
| # | interaction | 0 | 0 | 2 | 0 | 0 | # | 10 | Small | ancom | 58 |
| # | disease | 4 | 3 | 0 | 0.571428571 | 1 | # | 10 | Small | aldex2 | 58 |
| # | sex | 2 | 0 | 0 | 1 | 1 | # | 10 | Small | aldex2 | 58 |
| # | time | 2 | 1 | 0 | 0.666666667 | 1 | # | 10 | Small | aldex2 | 58 |
| # | interaction | 0 | 0 | 2 | 0 | 0 | # | 10 | Small | aldex2 | 58 |
| # | disease | 4 | 5 | 0 | 0.444444444 | 1 | # | 10 | Baseline | paralle | 58 |
| # | sex | 2 | 1 | 0 | 0.666666667 | 1 | # | 10 | Baseline | paralle | 58 |
| # | time | 2 | 5 | 0 | 0.285714286 | 1 | # | 10 | Baseline | paralle | 58 |
| # | interaction | 2 | 5 | 0 | 0.285714286 | 1 | # | 10 | Baseline | paralle | 58 |
| # | disease | 4 | 3 | 0 | 0.571428571 | 1 | # | 10 | Baseline | ancom | 58 |
| # | sex | 2 | 0 | 0 | 1 | 1 | # | 10 | Baseline | ancom | 58 |
| # | time | 2 | 2 | 0 | 0.5 | 1 | # | 10 | Baseline | ancom | 58 |
| # | interaction | 0 | 0 | 2 | 0 | 0 | # | 10 | Baseline | ancom | 58 |
| # | disease | 4 | 24 | 0 | 0.142857143 | 1 | # | 10 | Baseline | aldex2 | 58 |
| # | sex | 2 | 12 | 0 | 0.142857143 | 1 | # | 10 | Baseline | aldex2 | 58 |
| # | time | 2 | 13 | 0 | 0.133333333 | 1 | # | 10 | Baseline | aldex2 | 58 |
| # | interaction | 0 | 0 | 2 | 0 | 0 | # | 10 | Baseline | aldex2 | 58 |
| # | disease | 4 | 1 | 0 | 0.8 | 1 | # | 10 | Small | paralle | 58 |
| # | sex | 2 | 0 | 0 | 1 | 1 | # | 10 | Small | paralle | 58 |
| # | time | 2 | 0 | 0 | 1 | 1 | # | 10 | Small | paralle | 58 |
| # | interaction | 2 | 0 | 0 | 1 | 1 | # | 10 | Small | paralle | 58 |
| # | disease | 4 | 2 | 0 | 0.666666667 | 1 | # | 10 | Small | ancom | 58 |
| # | sex | 2 | 0 | 0 | 1 | 1 | # | 10 | Small | ancom | 58 |
| # | time | 2 | 1 | 0 | 0.666666667 | 1 | # | 10 | Small | ancom | 58 |
| # | interaction | 0 | 0 | 2 | 0 | 0 | # | 10 | Small | ancom | 58 |
| # | disease | 4 | 4 | 0 | 0.5 | 1 | # | 10 | Small | aldex2 | 58 |
| # | sex | 2 | 0 | 0 | 1 | 1 | # | 10 | Small | aldex2 | 58 |
| # | time | 2 | 1 | 0 | 0.666666667 | 1 | # | 10 | Small | aldex2 | 58 |
| # | interaction | 0 | 0 | 2 | 0 | 0 | # | 10 | Small | aldex2 | 58 |
| # | disease | 7 | 7 | 0 | 0.5 | 1 | # | 20 | Baseline | paralle | 58 |
| # | sex | 4 | 10 | 0 | 0.285714286 | 1 | # | 20 | Baseline | paralle | 58 |
| # | time | 5 | 12 | 0 | 0.294117647 | 1 | # | 20 | Baseline | paralle | 58 |
| # | interaction | 4 | 8 | 0 | 0.333333333 | 1 | # | 20 | Baseline | paralle | 58 |
| # | disease | 7 | 2 | 0 | 0.777777778 | 1 | # | 20 | Baseline | ancom | 58 |
| # | sex | 4 | 0 | 0 | 1 | 1 | # | 20 | Baseline | ancom | 58 |
| # | time | 5 | 3 | 0 | 0.625 | 1 | # | 20 | Baseline | ancom | 58 |
| # | interaction | 0 | 0 | 4 | 0 | 0 | # | 20 | Baseline | ancom | 58 |
| # | disease | 7 | 19 | 0 | 0.269230769 | 1 | # | 20 | Baseline | aldex2 | 58 |
| # | sex | 4 | 8 | 0 | 0.333333333 | 1 | # | 20 | Baseline | aldex2 | 58 |
| # | time | 5 | 14 | 0 | 0.263157895 | 1 | # | 20 | Baseline | aldex2 | 58 |
| # | interaction | 0 | 0 | 4 | 0 | 0 | # | 20 | Baseline | aldex2 | 58 |
| # | disease | 7 | 1 | 0 | 0.875 | 1 | # | 20 | Small | paralle | 58 |
| # | sex | 4 | 0 | 0 | 1 | 1 | # | 20 | Small | paralle | 58 |
| # | time | 5 | 0 | 0 | 1 | 1 | # | 20 | Small | paralle | 58 |
| # | interaction | 2 | 1 | 2 | 0.666666667 | 0.5 | # | 20 | Small | paralle | 58 |
| # | disease | 7 | 2 | 0 | 0.777777778 | 1 | # | 20 | Small | ancom | 58 |
| # | sex | 4 | 0 | 0 | 1 | 1 | # | 20 | Small | ancom | 58 |
| # | time | 5 | 3 | 0 | 0.625 | 1 | # | 20 | Small | ancom | 58 |
| # | interaction | 0 | 0 | 4 | 0 | 0 | # | 20 | Small | ancom | 58 |
| # | disease | 7 | 3 | 0 | 0.7 | 1 | # | 20 | Small | aldex2 | 58 |
| # | sex | 4 | 0 | 0 | 1 | 1 | # | 20 | Small | aldex2 | 58 |
| # | time | 5 | 2 | 0 | 0.714285714 | 1 | # | 20 | Small | aldex2 | 58 |
| # | interaction | 0 | 0 | 4 | 0 | 0 | # | 20 | Small | aldex2 | 58 |
| # | disease | 7 | 12 | 0 | 0.368421053 | 1 | # | 20 | Baseline | paralle | 58 |
| # | sex | 4 | 8 | 0 | 0.333333333 | 1 | # | 20 | Baseline | paralle | 58 |
| # | time | 5 | 13 | 0 | 0.277777778 | 1 | # | 20 | Baseline | paralle | 58 |
| # | interaction | 4 | 8 | 0 | 0.333333333 | 1 | # | 20 | Baseline | paralle | 58 |
| # | disease | 7 | 4 | 0 | 0.636363636 | 1 | # | 20 | Baseline | ancom | 58 |
| # | sex | 4 | 0 | 0 | 1 | 1 | # | 20 | Baseline | ancom | 58 |
| # | time | 5 | 4 | 0 | 0.555555556 | 1 | # | 20 | Baseline | ancom | 58 |
| # | interaction | 0 | 0 | 4 | 0 | 0 | # | 20 | Baseline | ancom | 58 |
| # | disease | 7 | 33 | 0 | 0.175 | 1 | # | 20 | Baseline | aldex2 | 58 |
| # | sex | 4 | 10 | 0 | 0.285714286 | 1 | # | 20 | Baseline | aldex2 | 58 |
| # | time | 5 | 31 | 0 | 0.138888889 | 1 | # | 20 | Baseline | aldex2 | 58 |
| # | interaction | 0 | 0 | 4 | 0 | 0 | # | 20 | Baseline | aldex2 | 58 |
| # | disease | 7 | 0 | 0 | 1 | 1 | # | 20 | Small | paralle | 58 |
| # | sex | 4 | 1 | 0 | 0.8 | 1 | # | 20 | Small | paralle | 58 |
| # | time | 5 | 0 | 0 | 1 | 1 | # | 20 | Small | paralle | 58 |
| # | interaction | 3 | 1 | 1 | 0.75 | 0.75 | # | 20 | Small | paralle | 58 |
| # | disease | 7 | 4 | 0 | 0.636363636 | 1 | # | 20 | Small | ancom | 58 |
| # | sex | 4 | 0 | 0 | 1 | 1 | # | 20 | Small | ancom | 58 |
| # | time | 5 | 4 | 0 | 0.555555556 | 1 | # | 20 | Small | ancom | 58 |
| # | interaction | 0 | 0 | 4 | 0 | 0 | # | 20 | Small | ancom | 58 |
| # | disease | 7 | 4 | 0 | 0.636363636 | 1 | # | 20 | Small | aldex2 | 58 |
| # | sex | 4 | 1 | 0 | 0.8 | 1 | # | 20 | Small | aldex2 | 58 |
| # | time | 5 | 3 | 0 | 0.625 | 1 | # | 20 | Small | aldex2 | 58 |
| # | interaction | 0 | 0 | 4 | 0 | 0 | # | 20 | Small | aldex2 | 58 |
| # | disease | 7 | 8 | 0 | 0.466666667 | 1 | # | 20 | Baseline | paralle | 58 |
| # | sex | 4 | 4 | 0 | 0.5 | 1 | # | 20 | Baseline | paralle | 58 |
| # | time | 5 | 5 | 0 | 0.5 | 1 | # | 20 | Baseline | paralle | 58 |
| # | interaction | 4 | 5 | 0 | 0.444444444 | 1 | # | 20 | Baseline | paralle | 58 |
| # | disease | 7 | 4 | 0 | 0.636363636 | 1 | # | 20 | Baseline | ancom | 58 |
| # | sex | 4 | 0 | 0 | 1 | 1 | # | 20 | Baseline | ancom | 58 |
| # | time | 5 | 4 | 0 | 0.555555556 | 1 | # | 20 | Baseline | ancom | 58 |
| # | interaction | 0 | 0 | 4 | 0 | 0 | # | 20 | Baseline | ancom | 58 |
| # | disease | 7 | 47 | 0 | 0.12962963 | 1 | # | 20 | Baseline | aldex2 | 58 |
| # | sex | 4 | 12 | 0 | 0.25 | 1 | # | 20 | Baseline | aldex2 | 58 |
| # | time | 5 | 38 | 0 | 0.11627907 | 1 | # | 20 | Baseline | aldex2 | 58 |
| # | interaction | 0 | 0 | 4 | 0 | 0 | # | 20 | Baseline | aldex2 | 58 |
| # | disease | 7 | 1 | 0 | 0.875 | 1 | # | 20 | Small | paralle | 58 |
| # | sex | 4 | 1 | 0 | 0.8 | 1 | # | 20 | Small | paralle | 58 |
| # | time | 5 | 0 | 0 | 1 | 1 | # | 20 | Small | paralle | 58 |
| # | interaction | 4 | 0 | 0 | 1 | 1 | # | 20 | Small | paralle | 58 |
| # | disease | 7 | 4 | 0 | 0.636363636 | 1 | # | 20 | Small | ancom | 58 |
| # | sex | 4 | 0 | 0 | 1 | 1 | # | 20 | Small | ancom | 58 |
| # | time | 5 | 4 | 0 | 0.555555556 | 1 | # | 20 | Small | ancom | 58 |
| # | interaction | 0 | 0 | 4 | 0 | 0 | # | 20 | Small | ancom | 58 |
| # | disease | 7 | 9 | 0 | 0.4375 | 1 | # | 20 | Small | aldex2 | 58 |
| # | sex | 4 | 1 | 0 | 0.8 | 1 | # | 20 | Small | aldex2 | 58 |
| # | time | 5 | 6 | 0 | 0.454545455 | 1 | # | 20 | Small | aldex2 | 58 |
| # | interaction | 0 | 0 | 4 | 0 | 0 | # | 20 | Small | aldex2 | 58 |
| # | disease | 7 | 8 | 0 | 0.466666667 | 1 | # | 20 | Baseline | paralle | 58 |
| # | sex | 4 | 4 | 0 | 0.5 | 1 | # | 20 | Baseline | paralle | 58 |
| # | time | 5 | 7 | 0 | 0.416666667 | 1 | # | 20 | Baseline | paralle | 58 |
| # | interaction | 4 | 4 | 0 | 0.5 | 1 | # | 20 | Baseline | paralle | 58 |
| # | disease | 7 | 4 | 0 | 0.636363636 | 1 | # | 20 | Baseline | ancom | 58 |
| # | sex | 4 | 0 | 0 | 1 | 1 | # | 20 | Baseline | ancom | 58 |

| | | | | | | | | | | | |
|---|---|---|---|---|---|---|---|---|---|---|---|
| # | time | 5 | 4 | 0 | 0.555555556 | 1 | # | 20 | Baseline | ancom | 58 |
| # | interaction | 0 | 0 | 4 | 0 | 0 | # | 20 | Baseline | ancom | 58 |
| # | disease | 7 | 53 | 0 | 0.116666667 | 1 | # | 20 | Baseline | sided2 | 58 |
| # | sex | 4 | 15 | 0 | 0.210526316 | 1 | # | 20 | Baseline | sided2 | 58 |
| # | time | 5 | 69 | 0 | 0.092592593 | 1 | # | 20 | Baseline | sided2 | 58 |
| # | interaction | 0 | 0 | 4 | 0 | 0 | # | 20 | Baseline | sided2 | 58 |
| # | disease | 7 | 1 | 0 | 0.875 | 1 | # | 20 | Small | paralle | 58 |
| # | sex | 4 | 0 | 0 | 1 | 1 | # | 20 | Small | paralle | 58 |
| # | time | 5 | 0 | 0 | 1 | 1 | # | 20 | Small | paralle | 58 |
| # | interaction | 4 | 0 | 0 | 1 | 1 | # | 20 | Small | paralle | 58 |
| # | disease | 7 | 4 | 0 | 0.636363636 | 1 | # | 20 | Small | ancom | 58 |
| # | sex | 4 | 0 | 0 | 1 | 1 | # | 20 | Small | ancom | 58 |
| # | time | 5 | 4 | 0 | 0.555555556 | 1 | # | 20 | Small | ancom | 58 |
| # | interaction | 0 | 0 | 4 | 0 | 0 | # | 20 | Small | ancom | 58 |
| # | disease | 7 | 10 | 0 | 0.411764706 | 1 | # | 20 | Small | sided2 | 58 |
| # | sex | 4 | 1 | 0 | 0.8 | 1 | # | 20 | Small | sided2 | 58 |
| # | time | 5 | 9 | 0 | 0.357142857 | 1 | # | 20 | Small | sided2 | 58 |
| # | interaction | 0 | 0 | 4 | 0 | 0 | # | 20 | Small | sided2 | 58 |
| # | disease | 4 | 6 | 0 | 0.4 | 1 | # | 10 | Baseline | paralle | 59 |
| # | sex | 2 | 10 | 0 | 0.166666667 | 1 | # | 10 | Baseline | paralle | 59 |
| # | time | 2 | 11 | 0 | 0.153846154 | 1 | # | 10 | Baseline | paralle | 59 |
| # | interaction | 2 | 8 | 0 | 0.2 | 1 | # | 10 | Baseline | paralle | 59 |
| # | disease | 4 | 0 | 0 | 1 | 1 | # | 10 | Baseline | ancom | 59 |
| # | sex | 2 | 0 | 0 | 1 | 1 | # | 10 | Baseline | ancom | 59 |
| # | time | 2 | 0 | 0 | 1 | 1 | # | 10 | Baseline | ancom | 59 |
| # | interaction | 0 | 0 | 2 | 0 | 0 | # | 10 | Baseline | ancom | 59 |
| # | disease | 4 | 19 | 0 | 0.173913043 | 1 | # | 10 | Baseline | sided2 | 59 |
| # | sex | 2 | 4 | 0 | 0.333333333 | 1 | # | 10 | Baseline | sided2 | 59 |
| # | time | 2 | 12 | 0 | 0.142857143 | 1 | # | 10 | Baseline | sided2 | 59 |
| # | interaction | 0 | 0 | 2 | 0 | 0 | # | 10 | Baseline | sided2 | 59 |
| # | disease | 4 | 1 | 0 | 0.8 | 1 | # | 10 | Small | paralle | 59 |
| # | sex | 2 | 0 | 0 | 1 | 1 | # | 10 | Small | paralle | 59 |
| # | time | 2 | 0 | 0 | 1 | 1 | # | 10 | Small | paralle | 59 |
| # | interaction | 1 | 1 | 1 | 0.5 | 0.5 | # | 10 | Small | paralle | 59 |
| # | disease | 4 | 0 | 0 | 1 | 1 | # | 10 | Small | ancom | 59 |
| # | sex | 2 | 0 | 0 | 1 | 1 | # | 10 | Small | ancom | 59 |
| # | time | 2 | 0 | 0 | 1 | 1 | # | 10 | Small | ancom | 59 |
| # | interaction | 0 | 0 | 2 | 0 | 0 | # | 10 | Small | ancom | 59 |
| # | disease | 4 | 2 | 0 | 0.666666667 | 1 | # | 10 | Small | sided2 | 59 |
| # | sex | 2 | 0 | 0 | 1 | 1 | # | 10 | Small | sided2 | 59 |
| # | time | 2 | 1 | 0 | 0.666666667 | 1 | # | 10 | Small | sided2 | 59 |
| # | interaction | 0 | 0 | 2 | 0 | 0 | # | 10 | Small | sided2 | 59 |
| # | disease | 4 | 5 | 0 | 0.444444444 | 1 | # | 10 | Baseline | paralle | 59 |
| # | sex | 2 | 4 | 0 | 0.333333333 | 1 | # | 10 | Baseline | paralle | 59 |
| # | time | 2 | 4 | 0 | 0.333333333 | 1 | # | 10 | Baseline | paralle | 59 |
| # | interaction | 2 | 4 | 0 | 0.333333333 | 1 | # | 10 | Baseline | paralle | 59 |
| # | disease | 4 | 2 | 0 | 0.666666667 | 1 | # | 10 | Baseline | ancom | 59 |
| # | sex | 2 | 0 | 0 | 1 | 1 | # | 10 | Baseline | ancom | 59 |
| # | time | 2 | 2 | 0 | 0.5 | 1 | # | 10 | Baseline | ancom | 59 |
| # | interaction | 0 | 0 | 2 | 0 | 0 | # | 10 | Baseline | ancom | 59 |
| # | disease | 4 | 22 | 0 | 0.153846154 | 1 | # | 10 | Baseline | sided2 | 59 |
| # | sex | 2 | 9 | 0 | 0.181818182 | 1 | # | 10 | Baseline | sided2 | 59 |
| # | time | 2 | 13 | 0 | 0.133333333 | 1 | # | 10 | Baseline | sided2 | 59 |
| # | interaction | 0 | 0 | 2 | 0 | 0 | # | 10 | Baseline | sided2 | 59 |
| # | disease | 4 | 1 | 0 | 0.8 | 1 | # | 10 | Small | paralle | 59 |
| # | sex | 2 | 0 | 0 | 1 | 1 | # | 10 | Small | paralle | 59 |
| # | time | 2 | 0 | 0 | 1 | 1 | # | 10 | Small | paralle | 59 |
| # | interaction | 2 | 0 | 0 | 1 | 1 | # | 10 | Small | paralle | 59 |
| # | disease | 4 | 2 | 0 | 0.666666667 | 1 | # | 10 | Small | ancom | 59 |
| # | sex | 2 | 0 | 0 | 1 | 1 | # | 10 | Small | ancom | 59 |
| # | time | 2 | 2 | 0 | 0.5 | 1 | # | 10 | Small | ancom | 59 |
| # | interaction | 0 | 0 | 2 | 0 | 0 | # | 10 | Small | ancom | 59 |
| # | disease | 4 | 3 | 0 | 0.571428571 | 1 | # | 10 | Small | sided2 | 59 |
| # | sex | 2 | 0 | 0 | 1 | 1 | # | 10 | Small | sided2 | 59 |
| # | time | 2 | 2 | 0 | 0.5 | 1 | # | 10 | Small | sided2 | 59 |
| # | interaction | 0 | 0 | 2 | 0 | 0 | # | 10 | Small | sided2 | 59 |
| # | disease | 4 | 6 | 0 | 0.4 | 1 | # | 10 | Baseline | paralle | 59 |
| # | sex | 2 | 1 | 0 | 0.666666667 | 1 | # | 10 | Baseline | paralle | 59 |
| # | time | 2 | 1 | 0 | 0.666666667 | 1 | # | 10 | Baseline | paralle | 59 |
| # | interaction | 2 | 6 | 0 | 0.25 | 1 | # | 10 | Baseline | paralle | 59 |
| # | disease | 4 | 2 | 0 | 0.666666667 | 1 | # | 10 | Baseline | ancom | 59 |
| # | sex | 2 | 0 | 0 | 1 | 1 | # | 10 | Baseline | ancom | 59 |
| # | time | 2 | 2 | 0 | 0.5 | 1 | # | 10 | Baseline | ancom | 59 |
| # | interaction | 0 | 0 | 2 | 0 | 0 | # | 10 | Baseline | ancom | 59 |
| # | disease | 4 | 24 | 0 | 0.142857143 | 1 | # | 10 | Baseline | sided2 | 59 |
| # | sex | 2 | 6 | 0 | 0.25 | 1 | # | 10 | Baseline | sided2 | 59 |
| # | time | 2 | 14 | 0 | 0.125 | 1 | # | 10 | Baseline | sided2 | 59 |
| # | interaction | 0 | 0 | 2 | 0 | 0 | # | 10 | Baseline | sided2 | 59 |
| # | disease | 4 | 0 | 0 | 1 | 1 | # | 10 | Small | paralle | 59 |
| # | sex | 2 | 0 | 0 | 1 | 1 | # | 10 | Small | paralle | 59 |
| # | time | 2 | 0 | 0 | 1 | 1 | # | 10 | Small | paralle | 59 |
| # | interaction | 2 | 0 | 0 | 1 | 1 | # | 10 | Small | paralle | 59 |
| # | disease | 4 | 2 | 0 | 0.666666667 | 1 | # | 10 | Small | ancom | 59 |
| # | sex | 2 | 0 | 0 | 1 | 1 | # | 10 | Small | ancom | 59 |
| # | time | 2 | 2 | 0 | 0.5 | 1 | # | 10 | Small | ancom | 59 |
| # | interaction | 0 | 0 | 2 | 0 | 0 | # | 10 | Small | ancom | 59 |
| # | disease | 4 | 4 | 0 | 0.5 | 1 | # | 10 | Small | sided2 | 59 |
| # | sex | 2 | 0 | 0 | 1 | 1 | # | 10 | Small | sided2 | 59 |
| # | time | 2 | 2 | 0 | 0.5 | 1 | # | 10 | Small | sided2 | 59 |
| # | interaction | 0 | 0 | 2 | 0 | 0 | # | 10 | Small | sided2 | 59 |
| # | disease | 4 | 7 | 0 | 0.363636364 | 1 | # | 10 | Baseline | paralle | 59 |
| # | sex | 2 | 0 | 0 | 1 | 1 | # | 10 | Baseline | paralle | 59 |
| # | time | 2 | 2 | 0 | 0.5 | 1 | # | 10 | Baseline | paralle | 59 |
| # | interaction | 2 | 4 | 0 | 0.333333333 | 1 | # | 10 | Baseline | paralle | 59 |
| # | disease | 4 | 2 | 0 | 0.666666667 | 1 | # | 10 | Baseline | ancom | 59 |
| # | sex | 2 | 0 | 0 | 1 | 1 | # | 10 | Baseline | ancom | 59 |
| # | time | 2 | 2 | 0 | 0.5 | 1 | # | 10 | Baseline | ancom | 59 |
| # | interaction | 0 | 0 | 2 | 0 | 0 | # | 10 | Baseline | ancom | 59 |
| # | disease | 4 | 25 | 0 | 0.137931034 | 1 | # | 10 | Baseline | sided2 | 59 |
| # | sex | 2 | 5 | 0 | 0.285714286 | 1 | # | 10 | Baseline | sided2 | 59 |
| # | time | 2 | 16 | 0 | 0.111111111 | 1 | # | 10 | Baseline | sided2 | 59 |
| # | interaction | 0 | 0 | 2 | 0 | 0 | # | 10 | Baseline | sided2 | 59 |
| # | disease | 4 | 0 | 0 | 1 | 1 | # | 10 | Small | paralle | 59 |
| # | sex | 2 | 0 | 0 | 1 | 1 | # | 10 | Small | paralle | 59 |
| # | time | 2 | 0 | 0 | 1 | 1 | # | 10 | Small | paralle | 59 |
| # | interaction | 2 | 0 | 0 | 1 | 1 | # | 10 | Small | paralle | 59 |
| # | disease | 4 | 2 | 0 | 0.666666667 | 1 | # | 10 | Small | ancom | 59 |
| # | sex | 2 | 0 | 0 | 1 | 1 | # | 10 | Small | ancom | 59 |
| # | time | 2 | 2 | 0 | 0.5 | 1 | # | 10 | Small | ancom | 59 |
| # | interaction | 0 | 0 | 2 | 0 | 0 | # | 10 | Small | ancom | 59 |
| # | disease | 4 | 5 | 0 | 0.444444444 | 1 | # | 10 | Small | sided2 | 59 |
| # | sex | 2 | 0 | 0 | 1 | 1 | # | 10 | Small | sided2 | 59 |
| # | time | 2 | 3 | 0 | 0.4 | 1 | # | 10 | Small | sided2 | 59 |
| # | interaction | 0 | 0 | 2 | 0 | 0 | # | 10 | Small | sided2 | 59 |
| # | disease | 7 | 6 | 0 | 0.538461538 | 1 | # | 20 | Baseline | paralle | 59 |
| # | sex | 4 | 6 | 0 | 0.4 | 1 | # | 20 | Baseline | paralle | 59 |
| # | time | 5 | 6 | 0 | 0.454545455 | 1 | # | 20 | Baseline | paralle | 59 |
| # | interaction | 3 | 7 | 1 | 0.3 | 0.75 | # | 20 | Baseline | paralle | 59 |
| # | disease | 7 | 1 | 0 | 0.875 | 1 | # | 20 | Baseline | ancom | 59 |
| # | sex | 4 | 0 | 0 | 1 | 1 | # | 20 | Baseline | ancom | 59 |
| # | time | 5 | 0 | 0 | 1 | 1 | # | 20 | Baseline | ancom | 59 |
| # | interaction | 0 | 0 | 4 | 0 | 0 | # | 20 | Baseline | ancom | 59 |
| # | disease | 7 | 25 | 0 | 0.21875 | 1 | # | 20 | Baseline | sided2 | 59 |
| # | sex | 4 | 2 | 0 | 0.666666667 | 1 | # | 20 | Baseline | sided2 | 59 |
| # | time | 5 | 16 | 0 | 0.238095238 | 1 | # | 20 | Baseline | sided2 | 59 |
| # | interaction | 0 | 0 | 4 | 0 | 0 | # | 20 | Baseline | sided2 | 59 |
| # | disease | 7 | 0 | 0 | 1 | 1 | # | 20 | Small | paralle | 59 |
| # | sex | 4 | 0 | 0 | 1 | 1 | # | 20 | Small | paralle | 59 |
| # | time | 5 | 0 | 0 | 1 | 1 | # | 20 | Small | paralle | 59 |
| # | interaction | 0 | 0 | 4 | 0 | 0 | # | 20 | Small | paralle | 59 |
| # | disease | 7 | 1 | 0 | 0.875 | 1 | # | 20 | Small | ancom | 59 |
| # | sex | 4 | 0 | 0 | 1 | 1 | # | 20 | Small | ancom | 59 |
| # | time | 5 | 0 | 0 | 1 | 1 | # | 20 | Small | ancom | 59 |
| # | interaction | 0 | 0 | 4 | 0 | 0 | # | 20 | Small | ancom | 59 |
| # | disease | 7 | 1 | 0 | 0.875 | 1 | # | 20 | Small | sided2 | 59 |
| # | sex | 4 | 0 | 0 | 1 | 1 | # | 20 | Small | sided2 | 59 |
| # | time | 5 | 1 | 0 | 0.833333333 | 1 | # | 20 | Small | sided2 | 59 |
| # | interaction | 0 | 0 | 4 | 0 | 0 | # | 20 | Small | sided2 | 59 |
| # | disease | 7 | 6 | 0 | 0.538461538 | 1 | # | 20 | Baseline | paralle | 59 |
| # | sex | 4 | 6 | 0 | 0.4 | 1 | # | 20 | Baseline | paralle | 59 |
| # | time | 5 | 5 | 0 | 0.5 | 1 | # | 20 | Baseline | paralle | 59 |
| # | interaction | 4 | 5 | 0 | 0.444444444 | 1 | # | 20 | Baseline | paralle | 59 |
| # | disease | 7 | 4 | 0 | 0.636363636 | 1 | # | 20 | Baseline | ancom | 59 |
| # | sex | 4 | 0 | 0 | 1 | 1 | # | 20 | Baseline | ancom | 59 |
| # | time | 5 | 1 | 0 | 0.833333333 | 1 | # | 20 | Baseline | ancom | 59 |
| # | interaction | 0 | 0 | 4 | 0 | 0 | # | 20 | Baseline | ancom | 59 |
| # | disease | 7 | 64 | 0 | 0.137254902 | 1 | # | 20 | Baseline | sided2 | 59 |
| # | sex | 4 | 11 | 0 | 0.266666667 | 1 | # | 20 | Baseline | sided2 | 59 |
| # | time | 5 | 23 | 0 | 0.178571429 | 1 | # | 20 | Baseline | sided2 | 59 |
| # | interaction | 0 | 0 | 4 | 0 | 0 | # | 20 | Baseline | sided2 | 59 |
| # | disease | 7 | 0 | 0 | 1 | 1 | # | 20 | Small | paralle | 59 |
| # | sex | 4 | 0 | 0 | 1 | 1 | # | 20 | Small | paralle | 59 |
| # | time | 5 | 0 | 0 | 1 | 1 | # | 20 | Small | paralle | 59 |
| # | interaction | 4 | 0 | 0 | 1 | 1 | # | 20 | Small | paralle | 59 |
| # | disease | 7 | 3 | 0 | 0.7 | 1 | # | 20 | Small | ancom | 59 |
| # | sex | 4 | 0 | 0 | 1 | 1 | # | 20 | Small | ancom | 59 |
| # | time | 5 | 0 | 0 | 1 | 1 | # | 20 | Small | ancom | 59 |
| # | interaction | 0 | 0 | 4 | 0 | 0 | # | 20 | Small | ancom | 59 |
| # | disease | 7 | 1 | 0 | 0.875 | 1 | # | 20 | Small | sided2 | 59 |
| # | sex | 4 | 0 | 0 | 1 | 1 | # | 20 | Small | sided2 | 59 |
| # | time | 5 | 1 | 0 | 0.833333333 | 1 | # | 20 | Small | sided2 | 59 |
| # | interaction | 0 | 0 | 4 | 0 | 0 | # | 20 | Small | sided2 | 59 |
| # | disease | 7 | 10 | 0 | 0.411764706 | 1 | # | 20 | Baseline | paralle | 59 |
| # | sex | 4 | 4 | 0 | 0.5 | 1 | # | 20 | Baseline | paralle | 59 |
| # | time | 5 | 4 | 0 | 0.555555556 | 1 | # | 20 | Baseline | paralle | 59 |
| # | interaction | 4 | 4 | 0 | 0.5 | 1 | # | 20 | Baseline | paralle | 59 |
| # | disease | 7 | 5 | 0 | 0.583333333 | 1 | # | 20 | Baseline | ancom | 59 |
| # | sex | 4 | 0 | 0 | 1 | 1 | # | 20 | Baseline | ancom | 59 |
| # | time | 5 | 3 | 0 | 0.625 | 1 | # | 20 | Baseline | ancom | 59 |
| # | interaction | 0 | 0 | 4 | 0 | 0 | # | 20 | Baseline | ancom | 59 |
| # | disease | 7 | 56 | 0 | 0.111111111 | 1 | # | 20 | Baseline | sided2 | 59 |
| # | sex | 4 | 20 | 0 | 0.166666667 | 1 | # | 20 | Baseline | sided2 | 59 |
| # | time | 5 | 33 | 0 | 0.131578947 | 1 | # | 20 | Baseline | sided2 | 59 |
| # | interaction | 0 | 0 | 4 | 0 | 0 | # | 20 | Baseline | sided2 | 59 |
| # | disease | 7 | 0 | 0 | 1 | 1 | # | 20 | Small | paralle | 59 |
| # | sex | 4 | 0 | 0 | 1 | 1 | # | 20 | Small | paralle | 59 |
| # | time | 5 | 0 | 0 | 1 | 1 | # | 20 | Small | paralle | 59 |
| # | interaction | 4 | 0 | 0 | 1 | 1 | # | 20 | Small | paralle | 59 |
| # | disease | 7 | 4 | 0 | 0.636363636 | 1 | # | 20 | Small | ancom | 59 |
| # | sex | 4 | 0 | 0 | 1 | 1 | # | 20 | Small | ancom | 59 |
| # | time | 5 | 3 | 0 | 0.625 | 1 | # | 20 | Small | ancom | 59 |
| # | interaction | 0 | 0 | 4 | 0 | 0 | # | 20 | Small | ancom | 59 |
| # | disease | 7 | 6 | 0 | 0.538461538 | 1 | # | 20 | Small | sided2 | 59 |
| # | sex | 4 | 0 | 0 | 1 | 1 | # | 20 | Small | sided2 | 59 |
| # | time | 5 | 3 | 0 | 0.625 | 1 | # | 20 | Small | sided2 | 59 |
| # | interaction | 0 | 0 | 4 | 0 | 0 | # | 20 | Small | sided2 | 59 |
| # | disease | 7 | 6 | 0 | 0.538461538 | 1 | # | 20 | Baseline | paralle | 59 |
| # | sex | 4 | 3 | 0 | 0.571428571 | 1 | # | 20 | Baseline | paralle | 59 |
| # | time | 5 | 8 | 0 | 0.384615385 | 1 | # | 20 | Baseline | paralle | 59 |
| # | interaction | 4 | 6 | 0 | 0.4 | 1 | # | 20 | Baseline | paralle | 59 |
| # | disease | 7 | 4 | 0 | 0.636363636 | 1 | # | 20 | Baseline | ancom | 59 |
| # | sex | 4 | 0 | 0 | 1 | 1 | # | 20 | Baseline | ancom | 59 |
| # | time | 5 | 4 | 0 | 0.555555556 | 1 | # | 20 | Baseline | ancom | 59 |
| # | interaction | 0 | 0 | 4 | 0 | 0 | # | 20 | Baseline | ancom | 59 |
| # | disease | 7 | 82 | 0 | 0.101449275 | 1 | # | 20 | Baseline | sided2 | 59 |
| # | sex | 4 | 22 | 0 | 0.153846154 | 1 | # | 20 | Baseline | sided2 | 59 |
| # | time | 5 | 39 | 0 | 0.113636364 | 1 | # | 20 | Baseline | sided2 | 59 |
| # | interaction | 0 | 0 | 4 | 0 | 0 | # | 20 | Baseline | sided2 | 59 |
| # | disease | 7 | 0 | 0 | 1 | 1 | # | 20 | Small | paralle | 59 |
| # | sex | 4 | 0 | 0 | 1 | 1 | # | 20 | Small | paralle | 59 |
| # | time | 5 | 0 | 0 | 1 | 1 | # | 20 | Small | paralle | 59 |
| # | interaction | 4 | 0 | 0 | 1 | 1 | # | 20 | Small | paralle | 59 |
| # | disease | 7 | 4 | 0 | 0.636363636 | 1 | # | 20 | Small | ancom | 59 |
| # | sex | 4 | 0 | 0 | 1 | 1 | # | 20 | Small | ancom | 59 |
| # | time | 5 | 4 | 0 | 0.555555556 | 1 | # | 20 | Small | ancom | 59 |
| # | interaction | 0 | 0 | 4 | 0 | 0 | # | 20 | Small | ancom | 59 |
| # | disease | 7 | 10 | 0 | 0.411764706 | 1 | # | 20 | Small | sided2 | 59 |
| # | sex | 4 | 0 | 0 | 1 | 1 | # | 20 | Small | sided2 | 59 |
| # | time | 5 | 5 | 0 | 0.5 | 1 | # | 20 | Small | sided2 | 59 |
| # | interaction | 0 | 0 | 4 | 0 | 0 | # | 20 | Small | sided2 | 59 |
| # | disease | 4 | 9 | 0 | 0.307692308 | 1 | # | 10 | Baseline | paralle | 59 |
| # | sex | 2 | 3 | 0 | 0.4 | 1 | # | 10 | Baseline | paralle | 60 |
| # | time | 2 | 4 | 0 | 0.333333333 | 1 | # | 10 | Baseline | paralle | 60 |
| # | interaction | 2 | 6 | 0 | 0.25 | 1 | # | 10 | Baseline | paralle | 60 |
| # | disease | 4 | 1 | 0 | 0.8 | 1 | # | 10 | Baseline | ancom | 60 |
| # | sex | 2 | 0 | 0 | 1 | 1 | # | 10 | Baseline | ancom | 60 |
| # | time | 2 | 2 | 0 | 0.5 | 1 | # | 10 | Baseline | ancom | 60 |
| # | interaction | 0 | 0 | 2 | 0 | 0 | # | 10 | Baseline | ancom | 60 |
| # | disease | 4 | 14 | 0 | 0.222222222 | 1 | # | 10 | Baseline | sided2 | 60 |
| # | sex | 2 | 7 | 0 | 0.222222222 | 1 | # | 10 | Baseline | sided2 | 60 |
| # | time | 2 | 7 | 0 | 0.222222222 | 1 | # | 10 | Baseline | sided2 | 60 |
| # | interaction | 0 | 0 | 2 | 0 | 0 | # | 10 | Baseline | sided2 | 60 |
| # | disease | 4 | 0 | 0 | 1 | 1 | # | 10 | Small | paralle | 60 |
| # | sex | 2 | 0 | 0 | 1 | 1 | # | 10 | Small | paralle | 60 |
| # | time | 1 | 0 | 1 | 1 | 0.5 | # | 10 | Small | paralle | 60 |
| # | interaction | 1 | 0 | 1 | 1 | 0.5 | # | 10 | Small | paralle | 60 |
| # | disease | 4 | 1 | 0 | 0.8 | 1 | # | 10 | Small | ancom | 60 |
| # | sex | 2 | 0 | 0 | 1 | 1 | # | 10 | Small | ancom | 60 |
| # | time | 2 | 1 | 0 | 0.666666667 | 1 | # | 10 | Small | ancom | 60 |
| # | interaction | 0 | 0 | 2 | 0 | 0 | # | 10 | Small | ancom | 60 |
| # | disease | 4 | 2 | 0 | 0.666666667 | 1 | # | 10 | Small | sided2 | 60 |
| # | sex | 2 | 0 | 0 | 1 | 1 | # | 10 | Small | sided2 | 60 |
| # | time | 2 | 2 | 0 | 0.5 | 1 | # | 10 | Small | sided2 | 60 |
| # | interaction | 0 | 0 | 2 | 0 | 0 | # | 10 | Small | sided2 | 60 |
| # | disease | 4 | 7 | 0 | 0.363636364 | 1 | # | 10 | Baseline | paralle | 60 |
| # | sex | 2 | 2 | 0 | 0.5 | 1 | # | 10 | Baseline | paralle | 60 |
| # | time | 2 | 4 | 0 | 0.333333333 | 1 | # | 10 | Baseline | paralle | 60 |
| # | interaction | 2 | 5 | 0 | 0.285714286 | 1 | # | 10 | Baseline | paralle | 60 |
| # | disease | 4 | 2 | 0 | 0.666666667 | 1 | # | 10 | Baseline | ancom | 60 |
| # | sex | 2 | 0 | 0 | 1 | 1 | # | 10 | Baseline | ancom | 60 |
| # | time | 2 | 1 | 0 | 0.666666667 | 1 | # | 10 | Baseline | ancom | 60 |
| # | interaction | 0 | 0 | 2 | 0 | 0 | # | 10 | Baseline | ancom | 60 |
| # | disease | 4 | 12 | 0 | 0.25 | 1 | # | 10 | Baseline | sided2 | 60 |
| # | sex | 2 | 6 | 0 | 0.25 | 1 | # | 10 | Baseline | sided2 | 60 |
| # | time | 2 | 10 | 0 | 0.166666667 | 1 | # | 10 | Baseline | sided2 | 60 |
| # | interaction | 0 | 0 | 2 | 0 | 0 | # | 10 | Baseline | sided2 | 60 |
| # | disease | 4 | 0 | 0 | 1 | 1 | # | 10 | Small | paralle | 60 |
| # | sex | 2 | 0 | 0 | 1 | 1 | # | 10 | Small | paralle | 60 |
| # | time | 2 | 0 | 0 | 1 | 1 | # | 10 | Small | paralle | 60 |
| # | interaction | 2 | 0 | 0 | 1 | 1 | # | 10 | Small | paralle | 60 |
| # | disease | 4 | 2 | 0 | 0.666666667 | 1 | # | 10 | Small | ancom | 60 |
| # | sex | 2 | 0 | 0 | 1 | 1 | # | 10 | Small | ancom | 60 |
| # | time | 2 | 1 | 0 | 0.666666667 | 1 | # | 10 | Small | ancom | 60 |
| # | interaction | 0 | 0 | 2 | 0 | 0 | # | 10 | Small | ancom | 60 |
| # | disease | 4 | 2 | 0 | 0.666666667 | 1 | # | 10 | Small | sided2 | 60 |
| # | sex | 2 | 0 | 0 | 1 | 1 | # | 10 | Small | sided2 | 60 |
| # | time | 2 | 3 | 0 | 0.4 | 1 | # | 10 | Small | sided2 | 60 |
| # | interaction | 0 | 0 | 2 | 0 | 0 | # | 10 | Small | sided2 | 60 |
| # | disease | 4 | 5 | 0 | 0.444444444 | 1 | # | 10 | Baseline | paralle | 60 |
| # | sex | 2 | 3 | 0 | 0.4 | 1 | # | 10 | Baseline | paralle | 60 |
| # | time | 2 | 4 | 0 | 0.333333333 | 1 | # | 10 | Baseline | paralle | 60 |
| # | interaction | 2 | 1 | 0 | 0.666666667 | 1 | # | 10 | Baseline | paralle | 60 |
| # | disease | 4 | 2 | 0 | 0.666666667 | 1 | # | 10 | Baseline | ancom | 60 |
| # | sex | 2 | 0 | 0 | 1 | 1 | # | 10 | Baseline | ancom | 60 |
| # | time | 2 | 2 | 0 | 0.5 | 1 | # | 10 | Baseline | ancom | 60 |
| # | interaction | 0 | 0 | 2 | 0 | 0 | # | 10 | Baseline | ancom | 60 |
| # | disease | 4 | 15 | 0 | 0.210526316 | 1 | # | 10 | Baseline | sided2 | 60 |
| # | sex | 2 | 5 | 0 | 0.285714286 | 1 | # | 10 | Baseline | sided2 | 60 |
| # | time | 2 | 14 | 0 | 0.125 | 1 | # | 10 | Baseline | sided2 | 60 |
| # | interaction | 0 | 0 | 2 | 0 | 0 | # | 10 | Baseline | sided2 | 60 |
| # | disease | 4 | 0 | 0 | 1 | 1 | # | 10 | Small | paralle | 60 |
| # | sex | 2 | 0 | 0 | 1 | 1 | # | 10 | Small | paralle | 60 |
| # | time | 2 | 0 | 0 | 1 | 1 | # | 10 | Small | paralle | 60 |
| # | interaction | 2 | 0 | 0 | 1 | 1 | # | 10 | Small | paralle | 60 |
| # | disease | 4 | 2 | 0 | 0.666666667 | 1 | # | 10 | Small | ancom | 60 |
| # | sex | 2 | 0 | 0 | 1 | 1 | # | 10 | Small | ancom | 60 |
| # | time | 2 | 2 | 0 | 0.5 | 1 | # | 10 | Small | ancom | 60 |
| # | interaction | 0 | 0 | 2 | 0 | 0 | # | 10 | Small | ancom | 60 |
| # | disease | 4 | 2 | 0 | 0.666666667 | 1 | # | 10 | Small | sided2 | 60 |
| # | sex | 2 | 0 | 0 | 1 | 1 | # | 10 | Small | sided2 | 60 |
| # | time | 2 | 2 | 0 | 0.5 | 1 | # | 10 | Small | sided2 | 60 |
| # | interaction | 0 | 0 | 2 | 0 | 0 | # | 10 | Small | sided2 | 60 |
| # | disease | 4 | 3 | 0 | 0.571428571 | 1 | # | 10 | Baseline | paralle | 60 |
| # | sex | 2 | 4 | 0 | 0.333333333 | 1 | # | 10 | Baseline | paralle | 60 |
| # | time | 2 | 1 | 0 | 0.666666667 | 1 | # | 10 | Baseline | paralle | 60 |
| # | interaction | 2 | 3 | 0 | 0.4 | 1 | # | 10 | Baseline | paralle | 60 |
| # | disease | 4 | 2 | 0 | 0.666666667 | 1 | # | 10 | Baseline | ancom | 60 |
| # | sex | 2 | 0 | 0 | 1 | 1 | # | 10 | Baseline | ancom | 60 |
| # | time | 2 | 2 | 0 | 0.5 | 1 | # | 10 | Baseline | ancom | 60 |
| # | interaction | 0 | 0 | 2 | 0 | 0 | # | 10 | Baseline | ancom | 60 |
| # | disease | 4 | 20 | 0 | 0.166666667 | 1 | # | 10 | Baseline | sided2 | 60 |
| # | sex | 2 | 8 | 0 | 0.2 | 1 | # | 10 | Baseline | sided2 | 60 |
| # | time | 2 | 13 | 0 | 0.133333333 | 1 | # | 10 | Baseline | sided2 | 60 |
| # | interaction | 0 | 0 | 2 | 0 | 0 | # | 10 | Baseline | sided2 | 60 |
| # | disease | 4 | 0 | 0 | 1 | 1 | # | 10 | Small | paralle | 60 |
| # | sex | 2 | 0 | 0 | 1 | 1 | # | 10 | Small | paralle | 60 |
| # | time | 2 | 0 | 0 | 1 | 1 | # | 10 | Small | paralle | 60 |
| # | interaction | 2 | 0 | 0 | 1 | 1 | # | 10 | Small | paralle | 60 |
| # | disease | 4 | 2 | 0 | 0.666666667 | 1 | # | 10 | Small | ancom | 60 |
| # | sex | 2 | 0 | 0 | 1 | 1 | # | 10 | Small | ancom | 60 |
| # | time | 2 | 2 | 0 | 0.5 | 1 | # | 10 | Small | ancom | 60 |
| # | interaction | 0 | 0 | 2 | 0 | 0 | # | 10 | Small | ancom | 60 |
| # | disease | 4 | 2 | 0 | 0.666666667 | 1 | # | 10 | Small | sided2 | 60 |
| # | sex | 2 | 0 | 0 | 1 | 1 | # | 10 | Small | sided2 | 60 |
| # | time | 2 | 2 | 0 | 0.5 | 1 | # | 10 | Small | sided2 | 60 |
| # | interaction | 0 | 0 | 2 | 0 | 0 | # | 10 | Small | sided2 | 60 |
| # | disease | 7 | 12 | 0 | 0.368421053 | 1 | # | 20 | Baseline | paralle | 60 |
| # | sex | 4 | 5 | 0 | 0.444444444 | 1 | # | 20 | Baseline | paralle | 60 |
| # | time | 5 | 9 | 0 | 0.357142857 | 1 | # | 20 | Baseline | paralle | 60 |
| # | interaction | 4 | 7 | 0 | 0.363636364 | 1 | # | 20 | Baseline | paralle | 60 |
| # | disease | 7 | 1 | 0 | 0.875 | 1 | # | 20 | Baseline | ancom | 60 |
| # | sex | 4 | 0 | 0 | 1 | 1 | # | 20 | Baseline | ancom | 60 |
| # | time | 5 | 0 | 0 | 1 | 1 | # | 20 | Baseline | ancom | 60 |
| # | interaction | 0 | 0 | 4 | 0 | 0 | # | 20 | Baseline | ancom | 60 |
| # | disease | 7 | 20 | 0 | 0.259259259 | 1 | # | 20 | Baseline | sided2 | 60 |
| # | sex | 4 | 9 | 0 | 0.307692308 | 1 | # | 20 | Baseline | sided2 | 60 |
| # | time | 5 | 14 | 0 | 0.263157895 | 1 | # | 20 | Baseline | sided2 | 60 |
| # | interaction | 0 | 0 | 4 | 0 | 0 | # | 20 | Baseline | sided2 | 60 |
| # | disease | 7 | 0 | 0 | 1 | 1 | # | 20 | Small | paralle | 60 |
| # | sex | 4 | 0 | 0 | 1 | 1 | # | 20 | Small | paralle | 60 |
| # | time | 5 | 0 | 0 | 1 | 1 | # | 20 | Small | paralle | 60 |
| # | interaction | 1 | 0 | 3 | 1 | 0.25 | # | 20 | Small | paralle | 60 |
| # | disease | 7 | 1 | 0 | 0.875 | 1 | # | 20 | Small | ancom | 60 |
| # | sex | 4 | 0 | 0 | 1 | 1 | # | 20 | Small | ancom | 60 |
| # | time | 5 | 0 | 0 | 1 | 1 | # | 20 | Small | ancom | 60 |
| # | interaction | 0 | 0 | 4 | 0 | 0 | # | 20 | Small | ancom | 60 |
| # | disease | 7 | 0 | 0 | 1 | 1 | # | 20 | Small | sided2 | 60 |
| # | sex | 4 | 0 | 0 | 1 | 1 | # | 20 | Small | sided2 | 60 |
| # | time | 5 | 2 | 0 | 0.714285714 | 1 | # | 20 | Small | sided2 | 60 |
| # | interaction | 0 | 0 | 4 | 0 | 0 | # | 20 | Small | sided2 | 60 |
| # | disease | 7 | 8 | 0 | 0.466666667 | 1 | # | 20 | Baseline | paralle | 60 |
| # | sex | 4 | 5 | 0 | 0.444444444 | 1 | # | 20 | Baseline | paralle | 60 |
| # | time | 5 | 8 | 0 | 0.384615385 | 1 | # | 20 | Baseline | paralle | 60 |
| # | interaction | 4 | 2 | 0 | 0.666666667 | 1 | # | 20 | Baseline | paralle | 60 |
| # | disease | 7 | 3 | 0 | 0.7 | 1 | # | 20 | Baseline | ancom | 60 |
| # | sex | 4 | 0 | 0 | 1 | 1 | # | 20 | Baseline | ancom | 60 |
| # | time | 5 | 3 | 0 | 0.625 | 1 | # | 20 | Baseline | ancom | 60 |

| | | | | | | | | | | | |
|---|---|---|---|---|---|---|---|---|---|---|---|
| # | interaction | 0 | 0 | 4 | 0 | 0 | # | 20 | Baseline | ancom | 60 |
| # | disease | 7 | 33 | 0 | 0.175 | 1 | # | 20 | Baseline | sides2 | 60 |
| # | sex | 4 | 13 | 0 | 0.235294118 | 1 | # | 20 | Baseline | sides2 | 60 |
| # | time | 5 | 24 | 0 | 0.172413793 | 1 | # | 20 | Baseline | sides2 | 60 |
| # | interaction | 0 | 0 | 4 | 0 | 0 | # | 20 | Baseline | sides2 | 60 |
| # | disease | 7 | 0 | 0 | 1 | 1 | # | 20 | Small | paralle | 60 |
| # | sex | 4 | 0 | 0 | 1 | 1 | # | 20 | Small | paralle | 60 |
| # | time | 5 | 0 | 0 | 1 | 1 | # | 20 | Small | paralle | 60 |
| # | interaction | 3 | 0 | 1 | 1 | 0.75 | # | 20 | Small | paralle | 60 |
| # | disease | 7 | 3 | 0 | 0.7 | 1 | # | 20 | Small | ancom | 60 |
| # | sex | 4 | 0 | 0 | 1 | 1 | # | 20 | Small | ancom | 60 |
| # | time | 5 | 2 | 0 | 0.714285714 | 1 | # | 20 | Small | ancom | 60 |
| # | interaction | 0 | 0 | 4 | 0 | 0 | # | 20 | Small | ancom | 60 |
| # | disease | 7 | 2 | 0 | 0.777777778 | 1 | # | 20 | Small | sides2 | 60 |
| # | sex | 4 | 0 | 0 | 1 | 1 | # | 20 | Small | sides2 | 60 |
| # | time | 5 | 2 | 0 | 0.714285714 | 1 | # | 20 | Small | sides2 | 60 |
| # | interaction | 0 | 0 | 4 | 0 | 0 | # | 20 | Small | sides2 | 60 |
| # | disease | 7 | 11 | 0 | 0.388888889 | 1 | # | 20 | Baseline | paralle | 60 |
| # | sex | 4 | 4 | 0 | 0.5 | 1 | # | 20 | Baseline | paralle | 60 |
| # | time | 5 | 11 | 0 | 0.3125 | 1 | # | 20 | Baseline | paralle | 60 |
| # | interaction | 4 | 2 | 0 | 0.666666667 | 1 | # | 20 | Baseline | paralle | 60 |
| # | disease | 7 | 4 | 0 | 0.636363636 | 1 | # | 20 | Baseline | ancom | 60 |
| # | sex | 4 | 0 | 0 | 1 | 1 | # | 20 | Baseline | ancom | 60 |
| # | time | 5 | 3 | 0 | 0.625 | 1 | # | 20 | Baseline | ancom | 60 |
| # | interaction | 0 | 0 | 4 | 0 | 0 | # | 20 | Baseline | ancom | 60 |
| # | disease | 7 | 51 | 0 | 0.120689655 | 1 | # | 20 | Baseline | sides2 | 60 |
| # | sex | 4 | 20 | 0 | 0.166666667 | 1 | # | 20 | Baseline | sides2 | 60 |
| # | time | 5 | 34 | 0 | 0.128205128 | 1 | # | 20 | Baseline | sides2 | 60 |
| # | interaction | 0 | 0 | 4 | 0 | 0 | # | 20 | Baseline | sides2 | 60 |
| # | disease | 7 | 0 | 0 | 1 | 1 | # | 20 | Small | paralle | 60 |
| # | sex | 4 | 0 | 0 | 1 | 1 | # | 20 | Small | paralle | 60 |
| # | time | 5 | 0 | 0 | 1 | 1 | # | 20 | Small | paralle | 60 |
| # | interaction | 4 | 0 | 0 | 1 | 1 | # | 20 | Small | paralle | 60 |
| # | disease | 7 | 3 | 0 | 0.7 | 1 | # | 20 | Small | ancom | 60 |
| # | sex | 4 | 0 | 0 | 1 | 1 | # | 20 | Small | ancom | 60 |
| # | time | 5 | 3 | 0 | 0.625 | 1 | # | 20 | Small | ancom | 60 |
| # | interaction | 0 | 0 | 4 | 0 | 0 | # | 20 | Small | ancom | 60 |
| # | disease | 7 | 4 | 0 | 0.636363636 | 1 | # | 20 | Small | sides2 | 60 |
| # | sex | 4 | 0 | 0 | 1 | 1 | # | 20 | Small | sides2 | 60 |
| # | time | 5 | 2 | 0 | 0.714285714 | 1 | # | 20 | Small | sides2 | 60 |
| # | interaction | 0 | 0 | 4 | 0 | 0 | # | 20 | Small | sides2 | 60 |
| # | disease | 7 | 12 | 0 | 0.368421053 | 1 | # | 20 | Baseline | paralle | 60 |
| # | sex | 4 | 4 | 0 | 0.5 | 1 | # | 20 | Baseline | paralle | 60 |
| # | time | 5 | 6 | 0 | 0.454545455 | 1 | # | 20 | Baseline | paralle | 60 |
| # | interaction | 4 | 1 | 0 | 0.8 | 1 | # | 20 | Baseline | paralle | 60 |
| # | disease | 7 | 4 | 0 | 0.636363636 | 1 | # | 20 | Baseline | ancom | 60 |
| # | sex | 4 | 0 | 0 | 1 | 1 | # | 20 | Baseline | ancom | 60 |
| # | time | 5 | 3 | 0 | 0.625 | 1 | # | 20 | Baseline | ancom | 60 |
| # | interaction | 0 | 0 | 4 | 0 | 0 | # | 20 | Baseline | ancom | 60 |
| # | disease | 7 | 59 | 0 | 0.106060606 | 1 | # | 20 | Baseline | sides2 | 60 |
| # | sex | 4 | 27 | 0 | 0.129032258 | 1 | # | 20 | Baseline | sides2 | 60 |
| # | time | 5 | 60 | 0 | 0.111111111 | 1 | # | 20 | Baseline | sides2 | 60 |
| # | interaction | 0 | 0 | 4 | 0 | 0 | # | 20 | Baseline | sides2 | 60 |
| # | disease | 7 | 0 | 0 | 1 | 1 | # | 20 | Small | paralle | 60 |
| # | sex | 4 | 0 | 0 | 1 | 1 | # | 20 | Small | paralle | 60 |
| # | time | 5 | 0 | 0 | 1 | 1 | # | 20 | Small | paralle | 60 |
| # | interaction | 4 | 0 | 0 | 1 | 1 | # | 20 | Small | paralle | 60 |
| # | disease | 7 | 4 | 0 | 0.636363636 | 1 | # | 20 | Small | ancom | 60 |
| # | sex | 4 | 0 | 0 | 1 | 1 | # | 20 | Small | ancom | 60 |
| # | time | 5 | 3 | 0 | 0.625 | 1 | # | 20 | Small | ancom | 60 |
| # | interaction | 0 | 0 | 4 | 0 | 0 | # | 20 | Small | ancom | 60 |
| # | disease | 7 | 8 | 0 | 0.466666667 | 1 | # | 20 | Small | sides2 | 60 |
| # | sex | 4 | 0 | 0 | 1 | 1 | # | 20 | Small | sides2 | 60 |
| # | time | 5 | 4 | 0 | 0.555555556 | 1 | # | 20 | Small | sides2 | 60 |
| # | interaction | 0 | 0 | 4 | 0 | 0 | # | 20 | Small | sides2 | 60 |
| # | disease | 4 | 10 | 0 | 0.285714286 | 1 | # | 10 | Baseline | paralle | 61 |
| # | sex | 2 | 6 | 0 | 0.25 | 1 | # | 10 | Baseline | paralle | 61 |
| # | time | 2 | 11 | 0 | 0.153846154 | 1 | # | 10 | Baseline | paralle | 61 |
| # | interaction | 1 | 13 | 1 | 0.071428571 | 0.5 | # | 10 | Baseline | paralle | 61 |
| # | disease | 4 | 1 | 0 | 0.8 | 1 | # | 10 | Baseline | ancom | 61 |
| # | sex | 2 | 0 | 0 | 1 | 1 | # | 10 | Baseline | ancom | 61 |
| # | time | 2 | 0 | 0 | 1 | 1 | # | 10 | Baseline | ancom | 61 |
| # | interaction | 0 | 0 | 2 | 0 | 0 | # | 10 | Baseline | ancom | 61 |
| # | disease | 4 | 15 | 0 | 0.210526316 | 1 | # | 10 | Baseline | sides2 | 61 |
| # | sex | 2 | 3 | 0 | 0.4 | 1 | # | 10 | Baseline | sides2 | 61 |
| # | time | 2 | 7 | 0 | 0.222222222 | 1 | # | 10 | Baseline | sides2 | 61 |
| # | interaction | 0 | 0 | 2 | 0 | 0 | # | 10 | Baseline | sides2 | 61 |
| # | disease | 4 | 1 | 0 | 0.8 | 1 | # | 10 | Small | paralle | 61 |
| # | sex | 2 | 0 | 0 | 1 | 1 | # | 10 | Small | paralle | 61 |
| # | time | 2 | 3 | 0 | 0.4 | 1 | # | 10 | Small | paralle | 61 |
| # | interaction | 1 | 1 | 1 | 0.5 | 0.5 | # | 10 | Small | paralle | 61 |
| # | disease | 4 | 1 | 0 | 0.8 | 1 | # | 10 | Small | ancom | 61 |
| # | sex | 2 | 0 | 0 | 1 | 1 | # | 10 | Small | ancom | 61 |
| # | time | 2 | 0 | 0 | 1 | 1 | # | 10 | Small | ancom | 61 |
| # | interaction | 0 | 0 | 2 | 0 | 0 | # | 10 | Small | ancom | 61 |
| # | disease | 4 | 1 | 0 | 0.8 | 1 | # | 10 | Small | sides2 | 61 |
| # | sex | 2 | 0 | 0 | 1 | 1 | # | 10 | Small | sides2 | 61 |
| # | time | 2 | 1 | 0 | 0.666666667 | 1 | # | 10 | Small | sides2 | 61 |
| # | interaction | 0 | 0 | 2 | 0 | 0 | # | 10 | Small | sides2 | 61 |
| # | disease | 4 | 11 | 0 | 0.266666667 | 1 | # | 10 | Baseline | paralle | 61 |
| # | sex | 2 | 7 | 0 | 0.222222222 | 1 | # | 10 | Baseline | paralle | 61 |
| # | time | 2 | 8 | 0 | 0.2 | 1 | # | 10 | Baseline | paralle | 61 |
| # | interaction | 2 | 12 | 0 | 0.142857143 | 1 | # | 10 | Baseline | paralle | 61 |
| # | disease | 4 | 2 | 0 | 0.666666667 | 1 | # | 10 | Baseline | ancom | 61 |
| # | sex | 2 | 0 | 0 | 1 | 1 | # | 10 | Baseline | ancom | 61 |
| # | time | 2 | 1 | 0 | 0.666666667 | 1 | # | 10 | Baseline | ancom | 61 |
| # | interaction | 0 | 0 | 2 | 0 | 0 | # | 10 | Baseline | ancom | 61 |
| # | disease | 4 | 17 | 0 | 0.19047619 | 1 | # | 10 | Baseline | sides2 | 61 |
| # | sex | 2 | 8 | 0 | 0.2 | 1 | # | 10 | Baseline | sides2 | 61 |
| # | time | 2 | 9 | 0 | 0.181818182 | 1 | # | 10 | Baseline | sides2 | 61 |
| # | interaction | 0 | 0 | 2 | 0 | 0 | # | 10 | Baseline | sides2 | 61 |
| # | disease | 4 | 1 | 0 | 0.8 | 1 | # | 10 | Small | paralle | 61 |
| # | sex | 2 | 0 | 0 | 1 | 1 | # | 10 | Small | paralle | 61 |
| # | time | 2 | 1 | 0 | 0.666666667 | 1 | # | 10 | Small | paralle | 61 |
| # | interaction | 1 | 0 | 1 | 1 | 0.5 | # | 10 | Small | paralle | 61 |
| # | disease | 4 | 2 | 0 | 0.666666667 | 1 | # | 10 | Small | ancom | 61 |
| # | sex | 2 | 0 | 0 | 1 | 1 | # | 10 | Small | ancom | 61 |
| # | time | 2 | 1 | 0 | 0.666666667 | 1 | # | 10 | Small | ancom | 61 |
| # | interaction | 0 | 0 | 2 | 0 | 0 | # | 10 | Small | ancom | 61 |
| # | disease | 4 | 2 | 0 | 0.666666667 | 1 | # | 10 | Small | sides2 | 61 |
| # | sex | 2 | 0 | 0 | 1 | 1 | # | 10 | Small | sides2 | 61 |
| # | time | 2 | 2 | 0 | 0.5 | 1 | # | 10 | Small | sides2 | 61 |
| # | interaction | 0 | 0 | 2 | 0 | 0 | # | 10 | Small | sides2 | 61 |
| # | disease | 4 | 5 | 0 | 0.444444444 | 1 | # | 10 | Baseline | paralle | 61 |
| # | sex | 2 | 5 | 0 | 0.285714286 | 1 | # | 10 | Baseline | paralle | 61 |
| # | time | 2 | 10 | 0 | 0.166666667 | 1 | # | 10 | Baseline | paralle | 61 |
| # | interaction | 2 | 11 | 0 | 0.153846154 | 1 | # | 10 | Baseline | paralle | 61 |
| # | disease | 4 | 2 | 0 | 0.666666667 | 1 | # | 10 | Baseline | ancom | 61 |
| # | sex | 2 | 0 | 0 | 1 | 1 | # | 10 | Baseline | ancom | 61 |
| # | time | 2 | 2 | 0 | 0.5 | 1 | # | 10 | Baseline | ancom | 61 |
| # | interaction | 0 | 0 | 2 | 0 | 0 | # | 10 | Baseline | ancom | 61 |
| # | disease | 4 | 22 | 0 | 0.153846154 | 1 | # | 10 | Baseline | sides2 | 61 |
| # | sex | 2 | 5 | 0 | 0.285714286 | 1 | # | 10 | Baseline | sides2 | 61 |
| # | time | 2 | 14 | 0 | 0.125 | 1 | # | 10 | Baseline | sides2 | 61 |
| # | interaction | 0 | 0 | 2 | 0 | 0 | # | 10 | Baseline | sides2 | 61 |
| # | disease | 4 | 0 | 0 | 1 | 1 | # | 10 | Small | paralle | 61 |
| # | sex | 2 | 0 | 0 | 1 | 1 | # | 10 | Small | paralle | 61 |
| # | time | 2 | 2 | 0 | 0.5 | 1 | # | 10 | Small | paralle | 61 |
| # | interaction | 2 | 0 | 0 | 1 | 1 | # | 10 | Small | paralle | 61 |
| # | disease | 4 | 2 | 0 | 0.666666667 | 1 | # | 10 | Small | ancom | 61 |
| # | sex | 2 | 0 | 0 | 1 | 1 | # | 10 | Small | ancom | 61 |
| # | time | 2 | 2 | 0 | 0.5 | 1 | # | 10 | Small | ancom | 61 |
| # | interaction | 0 | 0 | 2 | 0 | 0 | # | 10 | Small | ancom | 61 |
| # | disease | 4 | 2 | 0 | 0.666666667 | 1 | # | 10 | Small | sides2 | 61 |
| # | sex | 2 | 0 | 0 | 1 | 1 | # | 10 | Small | sides2 | 61 |
| # | time | 2 | 2 | 0 | 0.5 | 1 | # | 10 | Small | sides2 | 61 |
| # | interaction | 0 | 0 | 2 | 0 | 0 | # | 10 | Small | sides2 | 61 |
| # | disease | 4 | 3 | 0 | 0.571428571 | 1 | # | 10 | Baseline | paralle | 61 |
| # | sex | 2 | 6 | 0 | 0.25 | 1 | # | 10 | Baseline | paralle | 61 |
| # | time | 2 | 7 | 0 | 0.222222222 | 1 | # | 10 | Baseline | paralle | 61 |
| # | interaction | 2 | 7 | 0 | 0.222222222 | 1 | # | 10 | Baseline | paralle | 61 |
| # | disease | 4 | 2 | 0 | 0.666666667 | 1 | # | 10 | Baseline | ancom | 61 |
| # | sex | 2 | 0 | 0 | 1 | 1 | # | 10 | Baseline | ancom | 61 |
| # | time | 2 | 2 | 0 | 0.5 | 1 | # | 10 | Baseline | ancom | 61 |
| # | interaction | 0 | 0 | 2 | 0 | 0 | # | 10 | Baseline | ancom | 61 |
| # | disease | 4 | 31 | 0 | 0.114285714 | 1 | # | 10 | Baseline | sides2 | 61 |
| # | sex | 2 | 5 | 0 | 0.285714286 | 1 | # | 10 | Baseline | sides2 | 61 |
| # | time | 2 | 12 | 0 | 0.142857143 | 1 | # | 10 | Baseline | sides2 | 61 |
| # | interaction | 0 | 0 | 2 | 0 | 0 | # | 10 | Baseline | sides2 | 61 |
| # | disease | 4 | 0 | 0 | 1 | 1 | # | 10 | Small | paralle | 61 |
| # | sex | 2 | 0 | 0 | 1 | 1 | # | 10 | Small | paralle | 61 |
| # | time | 2 | 0 | 0 | 1 | 1 | # | 10 | Small | paralle | 61 |
| # | interaction | 2 | 0 | 0 | 1 | 1 | # | 10 | Small | paralle | 61 |
| # | disease | 4 | 2 | 0 | 0.666666667 | 1 | # | 10 | Small | ancom | 61 |
| # | sex | 2 | 0 | 0 | 1 | 1 | # | 10 | Small | ancom | 61 |
| # | time | 2 | 2 | 0 | 0.5 | 1 | # | 10 | Small | ancom | 61 |
| # | interaction | 0 | 0 | 2 | 0 | 0 | # | 10 | Small | ancom | 61 |
| # | disease | 4 | 5 | 0 | 0.444444444 | 1 | # | 10 | Small | sides2 | 61 |
| # | sex | 2 | 0 | 0 | 1 | 1 | # | 10 | Small | sides2 | 61 |
| # | time | 2 | 2 | 0 | 0.5 | 1 | # | 10 | Small | sides2 | 61 |
| # | interaction | 0 | 0 | 2 | 0 | 0 | # | 10 | Small | sides2 | 61 |
| # | disease | 7 | 10 | 0 | 0.411764706 | 1 | # | 20 | Baseline | paralle | 61 |
| # | sex | 4 | 8 | 0 | 0.333333333 | 1 | # | 20 | Baseline | paralle | 61 |
| # | time | 5 | 8 | 0 | 0.384615385 | 1 | # | 20 | Baseline | paralle | 61 |
| # | interaction | 4 | 7 | 0 | 0.363636364 | 1 | # | 20 | Baseline | paralle | 61 |
| # | disease | 7 | 2 | 0 | 0.777777778 | 1 | # | 20 | Baseline | ancom | 61 |
| # | sex | 4 | 0 | 0 | 1 | 1 | # | 20 | Baseline | ancom | 61 |
| # | time | 5 | 1 | 0 | 0.833333333 | 1 | # | 20 | Baseline | ancom | 61 |
| # | interaction | 0 | 0 | 4 | 0 | 0 | # | 20 | Baseline | ancom | 61 |
| # | disease | 7 | 22 | 0 | 0.24137931 | 1 | # | 20 | Baseline | sides2 | 61 |
| # | sex | 4 | 3 | 0 | 0.571428571 | 1 | # | 20 | Baseline | sides2 | 61 |
| # | time | 5 | 12 | 0 | 0.294117647 | 1 | # | 20 | Baseline | sides2 | 61 |
| # | interaction | 0 | 0 | 4 | 0 | 0 | # | 20 | Baseline | sides2 | 61 |
| # | disease | 7 | 0 | 0 | 1 | 1 | # | 20 | Small | paralle | 61 |
| # | sex | 4 | 0 | 0 | 1 | 1 | # | 20 | Small | paralle | 61 |
| # | time | 5 | 0 | 0 | 1 | 1 | # | 20 | Small | paralle | 61 |
| # | interaction | 3 | 0 | 1 | 1 | 0.75 | # | 20 | Small | paralle | 61 |
| # | disease | 7 | 2 | 0 | 0.777777778 | 1 | # | 20 | Small | ancom | 61 |
| # | sex | 4 | 0 | 0 | 1 | 1 | # | 20 | Small | ancom | 61 |
| # | time | 5 | 1 | 0 | 0.833333333 | 1 | # | 20 | Small | ancom | 61 |
| # | interaction | 0 | 0 | 4 | 0 | 0 | # | 20 | Small | ancom | 61 |
| # | disease | 7 | 1 | 0 | 0.875 | 1 | # | 20 | Small | sides2 | 61 |
| # | sex | 4 | 0 | 0 | 1 | 1 | # | 20 | Small | sides2 | 61 |
| # | time | 5 | 1 | 0 | 0.833333333 | 1 | # | 20 | Small | sides2 | 61 |
| # | interaction | 0 | 0 | 4 | 0 | 0 | # | 20 | Small | sides2 | 61 |
| # | disease | 7 | 9 | 0 | 0.4375 | 1 | # | 20 | Baseline | paralle | 61 |
| # | sex | 4 | 3 | 0 | 0.571428571 | 1 | # | 20 | Baseline | paralle | 61 |
| # | time | 5 | 10 | 0 | 0.333333333 | 1 | # | 20 | Baseline | paralle | 61 |
| # | interaction | 4 | 5 | 0 | 0.444444444 | 1 | # | 20 | Baseline | paralle | 61 |
| # | disease | 7 | 3 | 0 | 0.7 | 1 | # | 20 | Baseline | ancom | 61 |
| # | sex | 4 | 0 | 0 | 1 | 1 | # | 20 | Baseline | ancom | 61 |
| # | time | 5 | 4 | 0 | 0.555555556 | 1 | # | 20 | Baseline | ancom | 61 |
| # | interaction | 0 | 0 | 4 | 0 | 0 | # | 20 | Baseline | ancom | 61 |
| # | disease | 7 | 35 | 0 | 0.166666667 | 1 | # | 20 | Baseline | sides2 | 61 |
| # | sex | 4 | 5 | 0 | 0.444444444 | 1 | # | 20 | Baseline | sides2 | 61 |
| # | time | 5 | 25 | 0 | 0.166666667 | 1 | # | 20 | Baseline | sides2 | 61 |
| # | interaction | 0 | 0 | 4 | 0 | 0 | # | 20 | Baseline | sides2 | 61 |
| # | disease | 7 | 0 | 0 | 1 | 1 | # | 20 | Small | paralle | 61 |
| # | sex | 4 | 0 | 0 | 1 | 1 | # | 20 | Small | paralle | 61 |
| # | time | 5 | 0 | 0 | 1 | 1 | # | 20 | Small | paralle | 61 |
| # | interaction | 3 | 0 | 1 | 1 | 0.75 | # | 20 | Small | paralle | 61 |
| # | disease | 7 | 3 | 0 | 0.7 | 1 | # | 20 | Small | ancom | 61 |
| # | sex | 4 | 0 | 0 | 1 | 1 | # | 20 | Small | ancom | 61 |
| # | time | 5 | 4 | 0 | 0.555555556 | 1 | # | 20 | Small | ancom | 61 |
| # | interaction | 0 | 0 | 4 | 0 | 0 | # | 20 | Small | ancom | 61 |
| # | disease | 7 | 4 | 0 | 0.636363636 | 1 | # | 20 | Small | sides2 | 61 |
| # | sex | 4 | 0 | 0 | 1 | 1 | # | 20 | Small | sides2 | 61 |
| # | time | 5 | 4 | 0 | 0.555555556 | 1 | # | 20 | Small | sides2 | 61 |
| # | interaction | 0 | 0 | 4 | 0 | 0 | # | 20 | Small | sides2 | 61 |
| # | disease | 7 | 7 | 0 | 0.5 | 1 | # | 20 | Baseline | paralle | 61 |
| # | sex | 4 | 1 | 0 | 0.8 | 1 | # | 20 | Baseline | paralle | 61 |
| # | time | 5 | 7 | 0 | 0.416666667 | 1 | # | 20 | Baseline | paralle | 61 |
| # | interaction | 4 | 1 | 0 | 0.8 | 1 | # | 20 | Baseline | paralle | 61 |
| # | disease | 7 | 3 | 0 | 0.7 | 1 | # | 20 | Baseline | ancom | 61 |
| # | sex | 4 | 0 | 0 | 1 | 1 | # | 20 | Baseline | ancom | 61 |
| # | time | 5 | 4 | 0 | 0.555555556 | 1 | # | 20 | Baseline | ancom | 61 |
| # | interaction | 0 | 0 | 4 | 0 | 0 | # | 20 | Baseline | ancom | 61 |
| # | disease | 7 | 68 | 0 | 0.127272727 | 1 | # | 20 | Baseline | sides2 | 61 |
| # | sex | 4 | 12 | 0 | 0.25 | 1 | # | 20 | Baseline | sides2 | 61 |
| # | time | 5 | 26 | 0 | 0.161290323 | 1 | # | 20 | Baseline | sides2 | 61 |
| # | interaction | 0 | 0 | 4 | 0 | 0 | # | 20 | Baseline | sides2 | 61 |
| # | disease | 7 | 0 | 0 | 1 | 1 | # | 20 | Small | paralle | 61 |
| # | sex | 4 | 0 | 0 | 1 | 1 | # | 20 | Small | paralle | 61 |
| # | time | 5 | 0 | 0 | 1 | 1 | # | 20 | Small | paralle | 61 |
| # | interaction | 4 | 0 | 0 | 1 | 1 | # | 20 | Small | paralle | 61 |
| # | disease | 7 | 3 | 0 | 0.7 | 1 | # | 20 | Small | ancom | 61 |
| # | sex | 4 | 0 | 0 | 1 | 1 | # | 20 | Small | ancom | 61 |
| # | time | 5 | 4 | 0 | 0.555555556 | 1 | # | 20 | Small | ancom | 61 |
| # | interaction | 0 | 0 | 4 | 0 | 0 | # | 20 | Small | ancom | 61 |
| # | disease | 7 | 6 | 0 | 0.538461538 | 1 | # | 20 | Small | sides2 | 61 |
| # | sex | 4 | 0 | 0 | 1 | 1 | # | 20 | Small | sides2 | 61 |
| # | time | 5 | 5 | 0 | 0.5 | 1 | # | 20 | Small | sides2 | 61 |
| # | interaction | 0 | 0 | 4 | 0 | 0 | # | 20 | Small | sides2 | 61 |
| # | disease | 7 | 13 | 0 | 0.35 | 1 | # | 20 | Baseline | paralle | 61 |
| # | sex | 4 | 2 | 0 | 0.666666667 | 1 | # | 20 | Baseline | paralle | 61 |
| # | time | 5 | 10 | 0 | 0.333333333 | 1 | # | 20 | Baseline | paralle | 61 |
| # | interaction | 4 | 5 | 0 | 0.444444444 | 1 | # | 20 | Baseline | paralle | 61 |
| # | disease | 7 | 4 | 0 | 0.636363636 | 1 | # | 20 | Baseline | ancom | 61 |
| # | sex | 4 | 0 | 0 | 1 | 1 | # | 20 | Baseline | ancom | 61 |
| # | time | 5 | 4 | 0 | 0.555555556 | 1 | # | 20 | Baseline | ancom | 61 |
| # | interaction | 0 | 0 | 4 | 0 | 0 | # | 20 | Baseline | ancom | 61 |
| # | disease | 7 | 57 | 0 | 0.094594595 | 1 | # | 20 | Baseline | sides2 | 61 |
| # | sex | 4 | 14 | 0 | 0.222222222 | 1 | # | 20 | Baseline | sides2 | 61 |
| # | time | 5 | 33 | 0 | 0.131578947 | 1 | # | 20 | Baseline | sides2 | 61 |
| # | interaction | 0 | 0 | 4 | 0 | 0 | # | 20 | Baseline | sides2 | 61 |
| # | disease | 7 | 0 | 0 | 1 | 1 | # | 20 | Small | paralle | 61 |
| # | sex | 4 | 0 | 0 | 1 | 1 | # | 20 | Small | paralle | 61 |
| # | time | 5 | 0 | 0 | 1 | 1 | # | 20 | Small | paralle | 61 |
| # | interaction | 4 | 0 | 0 | 1 | 1 | # | 20 | Small | paralle | 61 |
| # | disease | 7 | 4 | 0 | 0.636363636 | 1 | # | 20 | Small | ancom | 61 |
| # | sex | 4 | 0 | 0 | 1 | 1 | # | 20 | Small | ancom | 61 |
| # | time | 5 | 4 | 0 | 0.555555556 | 1 | # | 20 | Small | ancom | 61 |
| # | interaction | 0 | 0 | 4 | 0 | 0 | # | 20 | Small | ancom | 61 |
| # | disease | 7 | 10 | 0 | 0.411764706 | 1 | # | 20 | Small | sides2 | 61 |
| # | sex | 4 | 0 | 0 | 1 | 1 | # | 20 | Small | sides2 | 61 |
| # | time | 5 | 6 | 0 | 0.454545455 | 1 | # | 20 | Small | sides2 | 61 |
| # | interaction | 0 | 0 | 4 | 0 | 0 | # | 20 | Small | sides2 | 61 |
| # | disease | 4 | 11 | 0 | 0.266666667 | 1 | # | 10 | Baseline | paralle | 62 |
| # | sex | 2 | 5 | 0 | 0.285714286 | 1 | # | 10 | Baseline | paralle | 62 |
| # | time | 2 | 8 | 0 | 0.2 | 1 | # | 10 | Baseline | paralle | 62 |
| # | interaction | 2 | 14 | 0 | 0.125 | 1 | # | 10 | Baseline | paralle | 62 |
| # | disease | 4 | 0 | 0 | 1 | 1 | # | 10 | Baseline | ancom | 62 |
| # | sex | 2 | 0 | 0 | 1 | 1 | # | 10 | Baseline | ancom | 62 |
| # | time | 2 | 0 | 0 | 1 | 1 | # | 10 | Baseline | ancom | 62 |
| # | interaction | 0 | 0 | 2 | 0 | 0 | # | 10 | Baseline | ancom | 62 |
| # | disease | 4 | 10 | 0 | 0.285714286 | 1 | # | 10 | Baseline | sides2 | 62 |
| # | sex | 2 | 4 | 0 | 0.333333333 | 1 | # | 10 | Baseline | sides2 | 62 |
| # | time | 2 | 5 | 0 | 0.285714286 | 1 | # | 10 | Baseline | sides2 | 62 |
| # | interaction | 0 | 0 | 2 | 0 | 0 | # | 10 | Baseline | sides2 | 62 |
| # | disease | 4 | 0 | 0 | 1 | 1 | # | 10 | Small | paralle | 62 |
| # | sex | 2 | 0 | 0 | 1 | 1 | # | 10 | Small | paralle | 62 |
| # | time | 2 | 0 | 0 | 1 | 1 | # | 10 | Small | paralle | 62 |
| # | interaction | 1 | 0 | 1 | 1 | 0.5 | # | 10 | Small | paralle | 62 |
| # | disease | 4 | 0 | 0 | 1 | 1 | # | 10 | Small | ancom | 62 |
| # | sex | 2 | 0 | 0 | 1 | 1 | # | 10 | Small | ancom | 62 |
| # | time | 2 | 0 | 0 | 1 | 1 | # | 10 | Small | ancom | 62 |
| # | interaction | 0 | 0 | 2 | 0 | 0 | # | 10 | Small | ancom | 62 |
| # | disease | 4 | 0 | 0 | 1 | 1 | # | 10 | Small | sides2 | 62 |
| # | sex | 2 | 0 | 0 | 1 | 1 | # | 10 | Small | sides2 | 62 |
| # | time | 2 | 1 | 0 | 0.666666667 | 1 | # | 10 | Small | sides2 | 62 |
| # | interaction | 0 | 0 | 2 | 0 | 0 | # | 10 | Small | sides2 | 62 |
| # | disease | 4 | 4 | 0 | 0.5 | 1 | # | 10 | Baseline | paralle | 62 |
| # | sex | 2 | 5 | 0 | 0.285714286 | 1 | # | 10 | Baseline | paralle | 62 |
| # | time | 2 | 8 | 0 | 0.2 | 1 | # | 10 | Baseline | paralle | 62 |
| # | interaction | 2 | 8 | 0 | 0.2 | 1 | # | 10 | Baseline | paralle | 62 |
| # | disease | 4 | 1 | 0 | 0.8 | 1 | # | 10 | Baseline | ancom | 62 |
| # | sex | 2 | 0 | 0 | 1 | 1 | # | 10 | Baseline | ancom | 62 |
| # | time | 2 | 2 | 0 | 0.5 | 1 | # | 10 | Baseline | ancom | 62 |
| # | interaction | 0 | 0 | 2 | 0 | 0 | # | 10 | Baseline | ancom | 62 |
| # | disease | 4 | 11 | 0 | 0.266666667 | 1 | # | 10 | Baseline | sides2 | 62 |
| # | sex | 2 | 10 | 0 | 0.166666667 | 1 | # | 10 | Baseline | sides2 | 62 |
| # | time | 2 | 10 | 0 | 0.166666667 | 1 | # | 10 | Baseline | sides2 | 62 |
| # | interaction | 0 | 0 | 2 | 0 | 0 | # | 10 | Baseline | sides2 | 62 |
| # | disease | 4 | 1 | 0 | 0.8 | 1 | # | 10 | Small | paralle | 62 |
| # | sex | 2 | 0 | 0 | 1 | 1 | # | 10 | Small | paralle | 62 |
| # | time | 2 | 0 | 0 | 1 | 1 | # | 10 | Small | paralle | 62 |
| # | interaction | 2 | 0 | 0 | 1 | 1 | # | 10 | Small | paralle | 62 |
| # | disease | 4 | 1 | 0 | 0.8 | 1 | # | 10 | Small | ancom | 62 |
| # | sex | 2 | 0 | 0 | 1 | 1 | # | 10 | Small | ancom | 62 |
| # | time | 2 | 2 | 0 | 0.5 | 1 | # | 10 | Small | ancom | 62 |
| # | interaction | 0 | 0 | 2 | 0 | 0 | # | 10 | Small | ancom | 62 |
| # | disease | 4 | 1 | 0 | 0.8 | 1 | # | 10 | Small | sides2 | 62 |
| # | sex | 2 | 1 | 0 | 0.666666667 | 1 | # | 10 | Small | sides2 | 62 |
| # | time | 2 | 3 | 0 | 0.4 | 1 | # | 10 | Small | sides2 | 62 |
| # | interaction | 0 | 0 | 2 | 0 | 0 | # | 10 | Small | sides2 | 62 |
| # | disease | 4 | 8 | 0 | 0.333333333 | 1 | # | 10 | Baseline | paralle | 62 |
| # | sex | 2 | 2 | 0 | 0.5 | 1 | # | 10 | Baseline | paralle | 62 |
| # | time | 2 | 4 | 0 | 0.333333333 | 1 | # | 10 | Baseline | paralle | 62 |
| # | interaction | 2 | 7 | 0 | 0.222222222 | 1 | # | 10 | Baseline | paralle | 62 |
| # | disease | 4 | 2 | 0 | 0.666666667 | 1 | # | 10 | Baseline | ancom | 62 |
| # | sex | 2 | 0 | 0 | 1 | 1 | # | 10 | Baseline | ancom | 62 |
| # | time | 2 | 2 | 0 | 0.5 | 1 | # | 10 | Baseline | ancom | 62 |
| # | interaction | 0 | 0 | 2 | 0 | 0 | # | 10 | Baseline | ancom | 62 |
| # | disease | 4 | 19 | 0 | 0.173913043 | 1 | # | 10 | Baseline | sides2 | 62 |
| # | sex | 2 | 13 | 0 | 0.133333333 | 1 | # | 10 | Baseline | sides2 | 62 |
| # | time | 2 | 10 | 0 | 0.166666667 | 1 | # | 10 | Baseline | sides2 | 62 |
| # | interaction | 0 | 0 | 2 | 0 | 0 | # | 10 | Baseline | sides2 | 62 |
| # | disease | 4 | 1 | 0 | 0.8 | 1 | # | 10 | Small | paralle | 62 |
| # | sex | 2 | 0 | 0 | 1 | 1 | # | 10 | Small | paralle | 62 |
| # | time | 2 | 0 | 0 | 1 | 1 | # | 10 | Small | paralle | 62 |
| # | interaction | 2 | 0 | 0 | 1 | 1 | # | 10 | Small | paralle | 62 |
| # | disease | 4 | 2 | 0 | 0.666666667 | 1 | # | 10 | Small | ancom | 62 |
| # | sex | 2 | 0 | 0 | 1 | 1 | # | 10 | Small | ancom | 62 |
| # | time | 2 | 2 | 0 | 0.5 | 1 | # | 10 | Small | ancom | 62 |
| # | interaction | 0 | 0 | 2 | 0 | 0 | # | 10 | Small | ancom | 62 |
| # | disease | 4 | 2 | 0 | 0.666666667 | 1 | # | 10 | Small | sides2 | 62 |
| # | sex | 2 | 0 | 0 | 1 | 1 | # | 10 | Small | sides2 | 62 |
| # | time | 2 | 3 | 0 | 0.4 | 1 | # | 10 | Small | sides2 | 62 |
| # | interaction | 0 | 0 | 2 | 0 | 0 | # | 10 | Small | sides2 | 62 |
| # | disease | 4 | 7 | 0 | 0.363636364 | 1 | # | 10 | Baseline | paralle | 62 |
| # | sex | 2 | 0 | 0 | 1 | 1 | # | 10 | Baseline | paralle | 62 |
| # | time | 2 | 2 | 0 | 0.5 | 1 | # | 10 | Baseline | paralle | 62 |
| # | interaction | 2 | 12 | 0 | 0.142857143 | 1 | # | 10 | Baseline | paralle | 62 |
| # | disease | 4 | 2 | 0 | 0.666666667 | 1 | # | 10 | Baseline | ancom | 62 |
| # | sex | 2 | 0 | 0 | 1 | 1 | # | 10 | Baseline | ancom | 62 |
| # | time | 2 | 2 | 0 | 0.5 | 1 | # | 10 | Baseline | ancom | 62 |
| # | interaction | 0 | 0 | 2 | 0 | 0 | # | 10 | Baseline | ancom | 62 |

| | | | | | | | | | | | |
|---|---|---|---|---|---|---|---|---|---|---|---|
| # | disease | 4 | 19 | 0 | 0.173913043 | 1 | # | 10 | Baseline | aldex2 | 62 |
| # | sex | 2 | 8 | 0 | 0.2 | 1 | # | 10 | Baseline | aldex2 | 62 |
| # | time | 2 | 16 | 0 | 0.111111111 | 1 | # | 10 | Baseline | aldex2 | 62 |
| # | interaction | 0 | 0 | 2 | 0 | 0 | # | 10 | Baseline | aldex2 | 62 |
| # | disease | 4 | 2 | 0 | 0.666666667 | 1 | # | 10 | Small | paralle | 62 |
| # | sex | 2 | 0 | 0 | 1 | 1 | # | 10 | Small | paralle | 62 |
| # | time | 2 | 0 | 0 | 1 | 1 | # | 10 | Small | paralle | 62 |
| # | interaction | 2 | 0 | 0 | 1 | 1 | # | 10 | Small | paralle | 62 |
| # | disease | 4 | 2 | 0 | 0.666666667 | 1 | # | 10 | Small | ancom | 62 |
| # | sex | 2 | 0 | 0 | 1 | 1 | # | 10 | Small | ancom | 62 |
| # | time | 2 | 2 | 0 | 0.5 | 1 | # | 10 | Small | ancom | 62 |
| # | interaction | 0 | 0 | 2 | 0 | 0 | # | 10 | Small | ancom | 62 |
| # | disease | 4 | 2 | 0 | 0.666666667 | 1 | # | 10 | Small | aldex2 | 62 |
| # | sex | 2 | 0 | 0 | 1 | 1 | # | 10 | Small | aldex2 | 62 |
| # | time | 2 | 2 | 0 | 0.5 | 1 | # | 10 | Small | aldex2 | 62 |
| # | interaction | 0 | 0 | 2 | 0 | 0 | # | 10 | Small | aldex2 | 62 |
| # | disease | 7 | 12 | 0 | 0.368421053 | 1 | # | 20 | Baseline | paralle | 62 |
| # | sex | 4 | 7 | 0 | 0.363636364 | 1 | # | 20 | Baseline | paralle | 62 |
| # | time | 5 | 4 | 0 | 0.555555556 | 1 | # | 20 | Baseline | paralle | 62 |
| # | interaction | 4 | 6 | 0 | 0.4 | 1 | # | 20 | Baseline | paralle | 62 |
| # | disease | 7 | 2 | 0 | 0.777777778 | 1 | # | 20 | Baseline | ancom | 62 |
| # | sex | 4 | 0 | 0 | 1 | 1 | # | 20 | Baseline | ancom | 62 |
| # | time | 5 | 1 | 0 | 0.833333333 | 1 | # | 20 | Baseline | ancom | 62 |
| # | interaction | 0 | 0 | 4 | 0 | 0 | # | 20 | Baseline | ancom | 62 |
| # | disease | 7 | 19 | 0 | 0.269230769 | 1 | # | 20 | Baseline | aldex2 | 62 |
| # | sex | 4 | 10 | 0 | 0.285714286 | 1 | # | 20 | Baseline | aldex2 | 62 |
| # | time | 5 | 20 | 0 | 0.2 | 1 | # | 20 | Baseline | aldex2 | 62 |
| # | interaction | 0 | 0 | 4 | 0 | 0 | # | 20 | Baseline | aldex2 | 62 |
| # | disease | 7 | 1 | 0 | 0.875 | 1 | # | 20 | Small | paralle | 62 |
| # | sex | 4 | 0 | 0 | 1 | 1 | # | 20 | Small | paralle | 62 |
| # | time | 5 | 0 | 0 | 1 | 1 | # | 20 | Small | paralle | 62 |
| # | interaction | 1 | 0 | 3 | 1 | 0.25 | # | 20 | Small | paralle | 62 |
| # | disease | 7 | 2 | 0 | 0.777777778 | 1 | # | 20 | Small | ancom | 62 |
| # | sex | 4 | 0 | 0 | 1 | 1 | # | 20 | Small | ancom | 62 |
| # | time | 5 | 0 | 0 | 1 | 1 | # | 20 | Small | ancom | 62 |
| # | interaction | 0 | 0 | 4 | 0 | 0 | # | 20 | Small | ancom | 62 |
| # | disease | 7 | 1 | 0 | 0.875 | 1 | # | 20 | Small | aldex2 | 62 |
| # | sex | 4 | 0 | 0 | 1 | 1 | # | 20 | Small | aldex2 | 62 |
| # | time | 5 | 2 | 0 | 0.714285714 | 1 | # | 20 | Small | aldex2 | 62 |
| # | interaction | 0 | 0 | 4 | 0 | 0 | # | 20 | Small | aldex2 | 62 |
| # | disease | 7 | 9 | 0 | 0.4375 | 1 | # | 20 | Baseline | paralle | 62 |
| # | sex | 4 | 4 | 0 | 0.5 | 1 | # | 20 | Baseline | paralle | 62 |
| # | time | 5 | 7 | 0 | 0.416666667 | 1 | # | 20 | Baseline | paralle | 62 |
| # | interaction | 4 | 5 | 0 | 0.444444444 | 1 | # | 20 | Baseline | paralle | 62 |
| # | disease | 7 | 3 | 0 | 0.7 | 1 | # | 20 | Baseline | ancom | 62 |
| # | sex | 4 | 0 | 0 | 1 | 1 | # | 20 | Baseline | ancom | 62 |
| # | time | 5 | 4 | 0 | 0.555555556 | 1 | # | 20 | Baseline | ancom | 62 |
| # | interaction | 0 | 0 | 4 | 0 | 0 | # | 20 | Baseline | ancom | 62 |
| # | disease | 7 | 46 | 0 | 0.132075472 | 1 | # | 20 | Baseline | aldex2 | 62 |
| # | sex | 4 | 10 | 0 | 0.285714286 | 1 | # | 20 | Baseline | aldex2 | 62 |
| # | time | 5 | 23 | 0 | 0.178571429 | 1 | # | 20 | Baseline | aldex2 | 62 |
| # | interaction | 0 | 0 | 4 | 0 | 0 | # | 20 | Baseline | aldex2 | 62 |
| # | disease | 7 | 0 | 0 | 1 | 1 | # | 20 | Small | paralle | 62 |
| # | sex | 4 | 0 | 0 | 1 | 1 | # | 20 | Small | paralle | 62 |
| # | time | 5 | 0 | 0 | 1 | 1 | # | 20 | Small | paralle | 62 |
| # | interaction | 4 | 0 | 0 | 1 | 1 | # | 20 | Small | paralle | 62 |
| # | disease | 7 | 3 | 0 | 0.7 | 1 | # | 20 | Small | ancom | 62 |
| # | sex | 4 | 0 | 0 | 1 | 1 | # | 20 | Small | ancom | 62 |
| # | time | 5 | 3 | 0 | 0.625 | 1 | # | 20 | Small | ancom | 62 |
| # | interaction | 0 | 0 | 4 | 0 | 0 | # | 20 | Small | ancom | 62 |
| # | disease | 7 | 6 | 0 | 0.538461538 | 1 | # | 20 | Small | aldex2 | 62 |
| # | sex | 4 | 0 | 0 | 1 | 1 | # | 20 | Small | aldex2 | 62 |
| # | time | 5 | 2 | 0 | 0.714285714 | 1 | # | 20 | Small | aldex2 | 62 |
| # | interaction | 0 | 0 | 4 | 0 | 0 | # | 20 | Small | aldex2 | 62 |
| # | disease | 7 | 6 | 0 | 0.538461538 | 1 | # | 20 | Baseline | paralle | 62 |
| # | sex | 4 | 4 | 0 | 0.5 | 1 | # | 20 | Baseline | paralle | 62 |
| # | time | 5 | 2 | 0 | 0.714285714 | 1 | # | 20 | Baseline | paralle | 62 |
| # | interaction | 4 | 6 | 0 | 0.4 | 1 | # | 20 | Baseline | paralle | 62 |
| # | disease | 7 | 4 | 0 | 0.636363636 | 1 | # | 20 | Baseline | ancom | 62 |
| # | sex | 4 | 0 | 0 | 1 | 1 | # | 20 | Baseline | ancom | 62 |
| # | time | 5 | 4 | 0 | 0.555555556 | 1 | # | 20 | Baseline | ancom | 62 |
| # | interaction | 0 | 0 | 4 | 0 | 0 | # | 20 | Baseline | ancom | 62 |
| # | disease | 7 | 50 | 0 | 0.104477612 | 1 | # | 20 | Baseline | aldex2 | 62 |
| # | sex | 4 | 14 | 0 | 0.222222222 | 1 | # | 20 | Baseline | aldex2 | 62 |
| # | time | 5 | 31 | 0 | 0.138888889 | 1 | # | 20 | Baseline | aldex2 | 62 |
| # | interaction | 0 | 0 | 4 | 0 | 0 | # | 20 | Baseline | aldex2 | 62 |
| # | disease | 7 | 2 | 0 | 0.777777778 | 1 | # | 20 | Small | paralle | 62 |
| # | sex | 4 | 0 | 0 | 1 | 1 | # | 20 | Small | paralle | 62 |
| # | time | 5 | 0 | 0 | 1 | 1 | # | 20 | Small | paralle | 62 |
| # | interaction | 4 | 0 | 0 | 1 | 1 | # | 20 | Small | paralle | 62 |
| # | disease | 7 | 4 | 0 | 0.636363636 | 1 | # | 20 | Small | ancom | 62 |
| # | sex | 4 | 0 | 0 | 1 | 1 | # | 20 | Small | ancom | 62 |
| # | time | 5 | 4 | 0 | 0.555555556 | 1 | # | 20 | Small | ancom | 62 |
| # | interaction | 0 | 0 | 4 | 0 | 0 | # | 20 | Small | ancom | 62 |
| # | disease | 7 | 9 | 0 | 0.4375 | 1 | # | 20 | Small | aldex2 | 62 |
| # | sex | 4 | 1 | 0 | 0.8 | 1 | # | 20 | Small | aldex2 | 62 |
| # | time | 5 | 4 | 0 | 0.555555556 | 1 | # | 20 | Small | aldex2 | 62 |
| # | interaction | 0 | 0 | 4 | 0 | 0 | # | 20 | Small | aldex2 | 62 |
| # | disease | 7 | 11 | 0 | 0.388888889 | 1 | # | 20 | Baseline | paralle | 62 |
| # | sex | 4 | 5 | 0 | 0.444444444 | 1 | # | 20 | Baseline | paralle | 62 |
| # | time | 5 | 3 | 0 | 0.625 | 1 | # | 20 | Baseline | paralle | 62 |
| # | interaction | 4 | 4 | 0 | 0.5 | 1 | # | 20 | Baseline | paralle | 62 |
| # | disease | 7 | 4 | 0 | 0.636363636 | 1 | # | 20 | Baseline | ancom | 62 |
| # | sex | 4 | 0 | 0 | 1 | 1 | # | 20 | Baseline | ancom | 62 |
| # | time | 5 | 4 | 0 | 0.555555556 | 1 | # | 20 | Baseline | ancom | 62 |
| # | interaction | 0 | 0 | 4 | 0 | 0 | # | 20 | Baseline | ancom | 62 |
| # | disease | 7 | 78 | 0 | 0.082352941 | 1 | # | 20 | Baseline | aldex2 | 62 |
| # | sex | 4 | 18 | 0 | 0.181818182 | 1 | # | 20 | Baseline | aldex2 | 62 |
| # | time | 5 | 38 | 0 | 0.11627907 | 1 | # | 20 | Baseline | aldex2 | 62 |
| # | interaction | 0 | 0 | 4 | 0 | 0 | # | 20 | Baseline | aldex2 | 62 |
| # | disease | 7 | 0 | 0 | 1 | 1 | # | 20 | Small | paralle | 62 |
| # | sex | 4 | 0 | 0 | 1 | 1 | # | 20 | Small | paralle | 62 |
| # | time | 5 | 0 | 0 | 1 | 1 | # | 20 | Small | paralle | 62 |
| # | interaction | 4 | 0 | 0 | 1 | 1 | # | 20 | Small | paralle | 62 |
| # | disease | 7 | 4 | 0 | 0.636363636 | 1 | # | 20 | Small | ancom | 62 |
| # | sex | 4 | 0 | 0 | 1 | 1 | # | 20 | Small | ancom | 62 |
| # | time | 5 | 4 | 0 | 0.555555556 | 1 | # | 20 | Small | ancom | 62 |
| # | interaction | 0 | 0 | 4 | 0 | 0 | # | 20 | Small | ancom | 62 |
| # | disease | 7 | 13 | 0 | 0.35 | 1 | # | 20 | Small | aldex2 | 62 |
| # | sex | 4 | 0 | 0 | 1 | 1 | # | 20 | Small | aldex2 | 62 |
| # | time | 5 | 6 | 0 | 0.454545455 | 1 | # | 20 | Small | aldex2 | 62 |
| # | interaction | 0 | 0 | 4 | 0 | 0 | # | 20 | Small | aldex2 | 62 |
| # | disease | 4 | 8 | 0 | 0.333333333 | 1 | # | 10 | Baseline | paralle | 63 |
| # | sex | 2 | 9 | 0 | 0.181818182 | 1 | # | 10 | Baseline | paralle | 63 |
| # | time | 2 | 10 | 0 | 0.166666667 | 1 | # | 10 | Baseline | paralle | 63 |
| # | interaction | 2 | 11 | 0 | 0.153846154 | 1 | # | 10 | Baseline | paralle | 63 |
| # | disease | 4 | 1 | 0 | 0.8 | 1 | # | 10 | Baseline | ancom | 63 |
| # | sex | 2 | 0 | 0 | 1 | 1 | # | 10 | Baseline | ancom | 63 |
| # | time | 2 | 2 | 0 | 0.5 | 1 | # | 10 | Baseline | ancom | 63 |
| # | interaction | 0 | 0 | 2 | 0 | 0 | # | 10 | Baseline | ancom | 63 |
| # | disease | 4 | 9 | 0 | 0.307692308 | 1 | # | 10 | Baseline | aldex2 | 63 |
| # | sex | 2 | 4 | 0 | 0.333333333 | 1 | # | 10 | Baseline | aldex2 | 63 |
| # | time | 2 | 7 | 0 | 0.222222222 | 1 | # | 10 | Baseline | aldex2 | 63 |
| # | interaction | 0 | 0 | 2 | 0 | 0 | # | 10 | Baseline | aldex2 | 63 |
| # | disease | 4 | 0 | 0 | 1 | 1 | # | 10 | Small | paralle | 63 |
| # | sex | 2 | 0 | 0 | 1 | 1 | # | 10 | Small | paralle | 63 |
| # | time | 2 | 1 | 0 | 0.666666667 | 1 | # | 10 | Small | paralle | 63 |
| # | interaction | 1 | 0 | 1 | 1 | 0.5 | # | 10 | Small | paralle | 63 |
| # | disease | 4 | 1 | 0 | 0.8 | 1 | # | 10 | Small | ancom | 63 |
| # | sex | 2 | 0 | 0 | 1 | 1 | # | 10 | Small | ancom | 63 |
| # | time | 2 | 1 | 0 | 0.666666667 | 1 | # | 10 | Small | ancom | 63 |
| # | interaction | 0 | 0 | 2 | 0 | 0 | # | 10 | Small | ancom | 63 |
| # | disease | 4 | 2 | 0 | 0.666666667 | 1 | # | 10 | Small | aldex2 | 63 |
| # | sex | 2 | 0 | 0 | 1 | 1 | # | 10 | Small | aldex2 | 63 |
| # | time | 2 | 0 | 0 | 1 | 1 | # | 10 | Small | aldex2 | 63 |
| # | interaction | 0 | 0 | 2 | 0 | 0 | # | 10 | Small | aldex2 | 63 |
| # | disease | 4 | 10 | 0 | 0.285714286 | 1 | # | 10 | Baseline | paralle | 63 |
| # | sex | 2 | 7 | 0 | 0.222222222 | 1 | # | 10 | Baseline | paralle | 63 |
| # | time | 2 | 3 | 0 | 0.4 | 1 | # | 10 | Baseline | paralle | 63 |
| # | interaction | 2 | 7 | 0 | 0.222222222 | 1 | # | 10 | Baseline | paralle | 63 |
| # | disease | 4 | 2 | 0 | 0.666666667 | 1 | # | 10 | Baseline | ancom | 63 |
| # | sex | 2 | 0 | 0 | 1 | 1 | # | 10 | Baseline | ancom | 63 |
| # | time | 2 | 2 | 0 | 0.5 | 1 | # | 10 | Baseline | ancom | 63 |
| # | interaction | 0 | 0 | 2 | 0 | 0 | # | 10 | Baseline | ancom | 63 |
| # | disease | 4 | 18 | 0 | 0.181818182 | 1 | # | 10 | Baseline | aldex2 | 63 |
| # | sex | 2 | 4 | 0 | 0.333333333 | 1 | # | 10 | Baseline | aldex2 | 63 |
| # | time | 2 | 9 | 0 | 0.181818182 | 1 | # | 10 | Baseline | aldex2 | 63 |
| # | interaction | 0 | 0 | 2 | 0 | 0 | # | 10 | Baseline | aldex2 | 63 |
| # | disease | 4 | 1 | 0 | 0.8 | 1 | # | 10 | Small | paralle | 63 |
| # | sex | 2 | 0 | 0 | 1 | 1 | # | 10 | Small | paralle | 63 |
| # | time | 2 | 0 | 0 | 1 | 1 | # | 10 | Small | paralle | 63 |
| # | interaction | 2 | 0 | 0 | 1 | 1 | # | 10 | Small | paralle | 63 |
| # | disease | 4 | 2 | 0 | 0.666666667 | 1 | # | 10 | Small | ancom | 63 |
| # | sex | 2 | 0 | 0 | 1 | 1 | # | 10 | Small | ancom | 63 |
| # | time | 2 | 2 | 0 | 0.5 | 1 | # | 10 | Small | ancom | 63 |
| # | interaction | 0 | 0 | 2 | 0 | 0 | # | 10 | Small | ancom | 63 |
| # | disease | 4 | 2 | 0 | 0.666666667 | 1 | # | 10 | Small | aldex2 | 63 |
| # | sex | 2 | 0 | 0 | 1 | 1 | # | 10 | Small | aldex2 | 63 |
| # | time | 2 | 2 | 0 | 0.5 | 1 | # | 10 | Small | aldex2 | 63 |
| # | interaction | 0 | 0 | 2 | 0 | 0 | # | 10 | Small | aldex2 | 63 |
| # | disease | 4 | 7 | 0 | 0.363636364 | 1 | # | 10 | Baseline | paralle | 63 |
| # | sex | 2 | 4 | 0 | 0.333333333 | 1 | # | 10 | Baseline | paralle | 63 |
| # | time | 2 | 6 | 0 | 0.25 | 1 | # | 10 | Baseline | paralle | 63 |
| # | interaction | 2 | 7 | 0 | 0.222222222 | 1 | # | 10 | Baseline | paralle | 63 |
| # | disease | 4 | 2 | 0 | 0.666666667 | 1 | # | 10 | Baseline | ancom | 63 |
| # | sex | 2 | 0 | 0 | 1 | 1 | # | 10 | Baseline | ancom | 63 |
| # | time | 2 | 2 | 0 | 0.5 | 1 | # | 10 | Baseline | ancom | 63 |
| # | interaction | 0 | 0 | 2 | 0 | 0 | # | 10 | Baseline | ancom | 63 |
| # | disease | 4 | 26 | 0 | 0.133333333 | 1 | # | 10 | Baseline | aldex2 | 63 |
| # | sex | 2 | 9 | 0 | 0.181818182 | 1 | # | 10 | Baseline | aldex2 | 63 |
| # | time | 2 | 14 | 0 | 0.125 | 1 | # | 10 | Baseline | aldex2 | 63 |
| # | interaction | 0 | 0 | 2 | 0 | 0 | # | 10 | Baseline | aldex2 | 63 |
| # | disease | 4 | 0 | 0 | 1 | 1 | # | 10 | Small | paralle | 63 |
| # | sex | 2 | 0 | 0 | 1 | 1 | # | 10 | Small | paralle | 63 |
| # | time | 2 | 0 | 0 | 1 | 1 | # | 10 | Small | paralle | 63 |
| # | interaction | 2 | 0 | 0 | 1 | 1 | # | 10 | Small | paralle | 63 |
| # | disease | 4 | 2 | 0 | 0.666666667 | 1 | # | 10 | Small | ancom | 63 |
| # | sex | 2 | 0 | 0 | 1 | 1 | # | 10 | Small | ancom | 63 |
| # | time | 2 | 2 | 0 | 0.5 | 1 | # | 10 | Small | ancom | 63 |
| # | interaction | 0 | 0 | 2 | 0 | 0 | # | 10 | Small | ancom | 63 |
| # | disease | 4 | 4 | 0 | 0.5 | 1 | # | 10 | Small | aldex2 | 63 |
| # | sex | 2 | 0 | 0 | 1 | 1 | # | 10 | Small | aldex2 | 63 |
| # | time | 2 | 2 | 0 | 0.5 | 1 | # | 10 | Small | aldex2 | 63 |
| # | interaction | 0 | 0 | 2 | 0 | 0 | # | 10 | Small | aldex2 | 63 |
| # | disease | 4 | 6 | 0 | 0.4 | 1 | # | 10 | Baseline | paralle | 63 |
| # | sex | 2 | 1 | 0 | 0.666666667 | 1 | # | 10 | Baseline | paralle | 63 |
| # | time | 2 | 9 | 0 | 0.181818182 | 1 | # | 10 | Baseline | paralle | 63 |
| # | interaction | 2 | 8 | 0 | 0.2 | 1 | # | 10 | Baseline | paralle | 63 |
| # | disease | 4 | 2 | 0 | 0.666666667 | 1 | # | 10 | Baseline | ancom | 63 |
| # | sex | 2 | 0 | 0 | 1 | 1 | # | 10 | Baseline | ancom | 63 |
| # | time | 2 | 2 | 0 | 0.5 | 1 | # | 10 | Baseline | ancom | 63 |
| # | interaction | 0 | 0 | 2 | 0 | 0 | # | 10 | Baseline | ancom | 63 |
| # | disease | 4 | 29 | 0 | 0.121212121 | 1 | # | 10 | Baseline | aldex2 | 63 |
| # | sex | 2 | 8 | 0 | 0.2 | 1 | # | 10 | Baseline | aldex2 | 63 |
| # | time | 2 | 12 | 0 | 0.142857143 | 1 | # | 10 | Baseline | aldex2 | 63 |
| # | interaction | 0 | 0 | 2 | 0 | 0 | # | 10 | Baseline | aldex2 | 63 |
| # | disease | 4 | 1 | 0 | 0.8 | 1 | # | 10 | Small | paralle | 63 |
| # | sex | 2 | 0 | 0 | 1 | 1 | # | 10 | Small | paralle | 63 |
| # | time | 2 | 0 | 0 | 1 | 1 | # | 10 | Small | paralle | 63 |
| # | interaction | 2 | 0 | 0 | 1 | 1 | # | 10 | Small | paralle | 63 |
| # | disease | 4 | 2 | 0 | 0.666666667 | 1 | # | 10 | Small | ancom | 63 |
| # | sex | 2 | 0 | 0 | 1 | 1 | # | 10 | Small | ancom | 63 |
| # | time | 2 | 2 | 0 | 0.5 | 1 | # | 10 | Small | ancom | 63 |
| # | interaction | 0 | 0 | 2 | 0 | 0 | # | 10 | Small | ancom | 63 |
| # | disease | 4 | 3 | 0 | 0.571428571 | 1 | # | 10 | Small | aldex2 | 63 |
| # | sex | 2 | 0 | 0 | 1 | 1 | # | 10 | Small | aldex2 | 63 |
| # | time | 2 | 2 | 0 | 0.5 | 1 | # | 10 | Small | aldex2 | 63 |
| # | interaction | 0 | 0 | 2 | 0 | 0 | # | 10 | Small | aldex2 | 63 |
| # | disease | 7 | 8 | 0 | 0.466666667 | 1 | # | 20 | Baseline | paralle | 63 |
| # | sex | 4 | 8 | 0 | 0.333333333 | 1 | # | 20 | Baseline | paralle | 63 |
| # | time | 5 | 6 | 0 | 0.454545455 | 1 | # | 20 | Baseline | paralle | 63 |
| # | interaction | 4 | 7 | 0 | 0.363636364 | 1 | # | 20 | Baseline | paralle | 63 |
| # | disease | 7 | 1 | 0 | 0.875 | 1 | # | 20 | Baseline | ancom | 63 |
| # | sex | 4 | 0 | 0 | 1 | 1 | # | 20 | Baseline | ancom | 63 |
| # | time | 5 | 2 | 0 | 0.714285714 | 1 | # | 20 | Baseline | ancom | 63 |
| # | interaction | 0 | 0 | 4 | 0 | 0 | # | 20 | Baseline | ancom | 63 |
| # | disease | 7 | 26 | 0 | 0.212121212 | 1 | # | 20 | Baseline | aldex2 | 63 |
| # | sex | 4 | 4 | 0 | 0.5 | 1 | # | 20 | Baseline | aldex2 | 63 |
| # | time | 5 | 14 | 0 | 0.263157895 | 1 | # | 20 | Baseline | aldex2 | 63 |
| # | interaction | 0 | 0 | 4 | 0 | 0 | # | 20 | Baseline | aldex2 | 63 |
| # | disease | 7 | 0 | 0 | 1 | 1 | # | 20 | Small | paralle | 63 |
| # | sex | 4 | 0 | 0 | 1 | 1 | # | 20 | Small | paralle | 63 |
| # | time | 5 | 1 | 0 | 0.833333333 | 1 | # | 20 | Small | paralle | 63 |
| # | interaction | 1 | 0 | 3 | 1 | 0.25 | # | 20 | Small | paralle | 63 |
| # | disease | 7 | 2 | 0 | 0.777777778 | 1 | # | 20 | Small | ancom | 63 |
| # | sex | 4 | 0 | 0 | 1 | 1 | # | 20 | Small | ancom | 63 |
| # | time | 5 | 2 | 0 | 0.714285714 | 1 | # | 20 | Small | ancom | 63 |
| # | interaction | 0 | 0 | 4 | 0 | 0 | # | 20 | Small | ancom | 63 |
| # | disease | 7 | 1 | 0 | 0.875 | 1 | # | 20 | Small | aldex2 | 63 |
| # | sex | 4 | 0 | 0 | 1 | 1 | # | 20 | Small | aldex2 | 63 |
| # | time | 5 | 3 | 0 | 0.625 | 1 | # | 20 | Small | aldex2 | 63 |
| # | interaction | 0 | 0 | 4 | 0 | 0 | # | 20 | Small | aldex2 | 63 |
| # | disease | 7 | 9 | 0 | 0.4375 | 1 | # | 20 | Baseline | paralle | 63 |
| # | sex | 4 | 3 | 0 | 0.571428571 | 1 | # | 20 | Baseline | paralle | 63 |
| # | time | 5 | 8 | 0 | 0.384615385 | 1 | # | 20 | Baseline | paralle | 63 |
| # | interaction | 4 | 5 | 0 | 0.444444444 | 1 | # | 20 | Baseline | paralle | 63 |
| # | disease | 7 | 4 | 0 | 0.636363636 | 1 | # | 20 | Baseline | ancom | 63 |
| # | sex | 4 | 0 | 0 | 1 | 1 | # | 20 | Baseline | ancom | 63 |
| # | time | 5 | 4 | 0 | 0.555555556 | 1 | # | 20 | Baseline | ancom | 63 |
| # | interaction | 0 | 0 | 4 | 0 | 0 | # | 20 | Baseline | ancom | 63 |
| # | disease | 7 | 38 | 0 | 0.155555556 | 1 | # | 20 | Baseline | aldex2 | 63 |
| # | sex | 4 | 7 | 0 | 0.363636364 | 1 | # | 20 | Baseline | aldex2 | 63 |
| # | time | 5 | 23 | 0 | 0.178571429 | 1 | # | 20 | Baseline | aldex2 | 63 |
| # | interaction | 0 | 0 | 4 | 0 | 0 | # | 20 | Baseline | aldex2 | 63 |
| # | disease | 7 | 0 | 0 | 1 | 1 | # | 20 | Small | paralle | 63 |
| # | sex | 4 | 0 | 0 | 1 | 1 | # | 20 | Small | paralle | 63 |
| # | time | 5 | 1 | 0 | 0.833333333 | 1 | # | 20 | Small | paralle | 63 |
| # | interaction | 4 | 0 | 0 | 1 | 1 | # | 20 | Small | paralle | 63 |
| # | disease | 7 | 4 | 0 | 0.636363636 | 1 | # | 20 | Small | ancom | 63 |
| # | sex | 4 | 0 | 0 | 1 | 1 | # | 20 | Small | ancom | 63 |
| # | time | 5 | 4 | 0 | 0.555555556 | 1 | # | 20 | Small | ancom | 63 |
| # | interaction | 0 | 0 | 4 | 0 | 0 | # | 20 | Small | ancom | 63 |
| # | disease | 7 | 3 | 0 | 0.7 | 1 | # | 20 | Small | aldex2 | 63 |
| # | sex | 4 | 0 | 0 | 1 | 1 | # | 20 | Small | aldex2 | 63 |
| # | time | 5 | 4 | 0 | 0.555555556 | 1 | # | 20 | Small | aldex2 | 63 |
| # | interaction | 0 | 0 | 4 | 0 | 0 | # | 20 | Small | aldex2 | 63 |
| # | disease | 7 | 9 | 0 | 0.4375 | 1 | # | 20 | Baseline | paralle | 63 |
| # | sex | 4 | 5 | 0 | 0.444444444 | 1 | # | 20 | Baseline | paralle | 63 |
| # | time | 5 | 8 | 0 | 0.384615385 | 1 | # | 20 | Baseline | paralle | 63 |
| # | interaction | 4 | 3 | 0 | 0.571428571 | 1 | # | 20 | Baseline | paralle | 63 |
| # | disease | 7 | 4 | 0 | 0.636363636 | 1 | # | 20 | Baseline | ancom | 63 |
| # | sex | 4 | 0 | 0 | 1 | 1 | # | 20 | Baseline | ancom | 63 |
| # | time | 5 | 4 | 0 | 0.555555556 | 1 | # | 20 | Baseline | ancom | 63 |
| # | interaction | 0 | 0 | 4 | 0 | 0 | # | 20 | Baseline | ancom | 63 |
| # | disease | 7 | 50 | 0 | 0.122807018 | 1 | # | 20 | Baseline | aldex2 | 63 |
| # | sex | 4 | 13 | 0 | 0.235294118 | 1 | # | 20 | Baseline | aldex2 | 63 |
| # | time | 5 | 29 | 0 | 0.147058824 | 1 | # | 20 | Baseline | aldex2 | 63 |
| # | interaction | 0 | 0 | 4 | 0 | 0 | # | 20 | Baseline | aldex2 | 63 |
| # | disease | 7 | 1 | 0 | 0.875 | 1 | # | 20 | Small | paralle | 63 |
| # | sex | 4 | 0 | 0 | 1 | 1 | # | 20 | Small | paralle | 63 |
| # | time | 5 | 1 | 0 | 0.833333333 | 1 | # | 20 | Small | paralle | 63 |
| # | interaction | 4 | 0 | 0 | 1 | 1 | # | 20 | Small | paralle | 63 |
| # | disease | 7 | 4 | 0 | 0.636363636 | 1 | # | 20 | Small | ancom | 63 |
| # | sex | 4 | 0 | 0 | 1 | 1 | # | 20 | Small | ancom | 63 |
| # | time | 5 | 4 | 0 | 0.555555556 | 1 | # | 20 | Small | ancom | 63 |
| # | interaction | 0 | 0 | 4 | 0 | 0 | # | 20 | Small | ancom | 63 |
| # | disease | 7 | 8 | 0 | 0.466666667 | 1 | # | 20 | Small | aldex2 | 63 |
| # | sex | 4 | 1 | 0 | 0.8 | 1 | # | 20 | Small | aldex2 | 63 |
| # | time | 5 | 4 | 0 | 0.555555556 | 1 | # | 20 | Small | aldex2 | 63 |
| # | interaction | 0 | 0 | 4 | 0 | 0 | # | 20 | Small | aldex2 | 63 |
| # | disease | 7 | 11 | 0 | 0.388888889 | 1 | # | 20 | Baseline | paralle | 63 |
| # | sex | 4 | 3 | 0 | 0.571428571 | 1 | # | 20 | Baseline | paralle | 63 |
| # | time | 5 | 6 | 0 | 0.454545455 | 1 | # | 20 | Baseline | paralle | 63 |
| # | interaction | 4 | 5 | 0 | 0.444444444 | 1 | # | 20 | Baseline | paralle | 63 |
| # | disease | 7 | 4 | 0 | 0.636363636 | 1 | # | 20 | Baseline | ancom | 63 |
| # | sex | 4 | 0 | 0 | 1 | 1 | # | 20 | Baseline | ancom | 63 |
| # | time | 5 | 4 | 0 | 0.555555556 | 1 | # | 20 | Baseline | ancom | 63 |
| # | interaction | 0 | 0 | 4 | 0 | 0 | # | 20 | Baseline | ancom | 63 |
| # | disease | 7 | 65 | 0 | 0.097222222 | 1 | # | 20 | Baseline | aldex2 | 63 |
| # | sex | 4 | 18 | 0 | 0.181818182 | 1 | # | 20 | Baseline | aldex2 | 63 |
| # | time | 5 | 37 | 0 | 0.119047619 | 1 | # | 20 | Baseline | aldex2 | 63 |
| # | interaction | 0 | 0 | 4 | 0 | 0 | # | 20 | Baseline | aldex2 | 63 |
| # | disease | 7 | 1 | 0 | 0.875 | 1 | # | 20 | Small | paralle | 63 |
| # | sex | 4 | 0 | 0 | 1 | 1 | # | 20 | Small | paralle | 63 |
| # | time | 5 | 1 | 0 | 0.833333333 | 1 | # | 20 | Small | paralle | 63 |
| # | interaction | 4 | 1 | 0 | 0.8 | 1 | # | 20 | Small | paralle | 63 |
| # | disease | 7 | 4 | 0 | 0.636363636 | 1 | # | 20 | Small | ancom | 63 |
| # | sex | 4 | 0 | 0 | 1 | 1 | # | 20 | Small | ancom | 63 |
| # | time | 5 | 4 | 0 | 0.555555556 | 1 | # | 20 | Small | ancom | 63 |
| # | interaction | 0 | 0 | 4 | 0 | 0 | # | 20 | Small | ancom | 63 |
| # | disease | 7 | 10 | 0 | 0.411764706 | 1 | # | 20 | Small | aldex2 | 63 |
| # | sex | 4 | 0 | 0 | 1 | 1 | # | 20 | Small | aldex2 | 63 |
| # | time | 5 | 4 | 0 | 0.555555556 | 1 | # | 20 | Small | aldex2 | 63 |
| # | interaction | 0 | 0 | 4 | 0 | 0 | # | 20 | Small | aldex2 | 63 |
| # | disease | 4 | 9 | 0 | 0.307692308 | 1 | # | 10 | Baseline | paralle | 64 |
| # | sex | 2 | 11 | 0 | 0.153846154 | 1 | # | 10 | Baseline | paralle | 64 |
| # | time | 2 | 11 | 0 | 0.153846154 | 1 | # | 10 | Baseline | paralle | 64 |
| # | interaction | 2 | 8 | 0 | 0.2 | 1 | # | 10 | Baseline | paralle | 64 |
| # | disease | 4 | 0 | 0 | 1 | 1 | # | 10 | Baseline | ancom | 64 |
| # | sex | 2 | 0 | 0 | 1 | 1 | # | 10 | Baseline | ancom | 64 |
| # | time | 2 | 1 | 0 | 0.666666667 | 1 | # | 10 | Baseline | ancom | 64 |
| # | interaction | 0 | 0 | 2 | 0 | 0 | # | 10 | Baseline | ancom | 64 |
| # | disease | 4 | 12 | 0 | 0.25 | 1 | # | 10 | Baseline | aldex2 | 64 |
| # | sex | 2 | 5 | 0 | 0.285714286 | 1 | # | 10 | Baseline | aldex2 | 64 |
| # | time | 2 | 7 | 0 | 0.222222222 | 1 | # | 10 | Baseline | aldex2 | 64 |
| # | interaction | 0 | 0 | 2 | 0 | 0 | # | 10 | Baseline | aldex2 | 64 |
| # | disease | 4 | 0 | 0 | 1 | 1 | # | 10 | Small | paralle | 64 |
| # | sex | 2 | 0 | 0 | 1 | 1 | # | 10 | Small | paralle | 64 |
| # | time | 2 | 0 | 0 | 1 | 1 | # | 10 | Small | paralle | 64 |
| # | interaction | 1 | 0 | 1 | 1 | 0.5 | # | 10 | Small | paralle | 64 |
| # | disease | 4 | 0 | 0 | 1 | 1 | # | 10 | Small | ancom | 64 |
| # | sex | 2 | 0 | 0 | 1 | 1 | # | 10 | Small | ancom | 64 |
| # | time | 2 | 1 | 0 | 0.666666667 | 1 | # | 10 | Small | ancom | 64 |
| # | interaction | 0 | 0 | 2 | 0 | 0 | # | 10 | Small | ancom | 64 |
| # | disease | 4 | 0 | 0 | 1 | 1 | # | 10 | Small | aldex2 | 64 |
| # | sex | 2 | 0 | 0 | 1 | 1 | # | 10 | Small | aldex2 | 64 |
| # | time | 2 | 1 | 0 | 0.666666667 | 1 | # | 10 | Small | aldex2 | 64 |
| # | interaction | 0 | 0 | 2 | 0 | 0 | # | 10 | Small | aldex2 | 64 |
| # | disease | 4 | 8 | 0 | 0.333333333 | 1 | # | 10 | Baseline | paralle | 64 |
| # | sex | 2 | 7 | 0 | 0.222222222 | 1 | # | 10 | Baseline | paralle | 64 |
| # | time | 2 | 9 | 0 | 0.181818182 | 1 | # | 10 | Baseline | paralle | 64 |
| # | interaction | 2 | 7 | 0 | 0.222222222 | 1 | # | 10 | Baseline | paralle | 64 |
| # | disease | 4 | 2 | 0 | 0.666666667 | 1 | # | 10 | Baseline | ancom | 64 |
| # | sex | 2 | 0 | 0 | 1 | 1 | # | 10 | Baseline | ancom | 64 |
| # | time | 2 | 1 | 0 | 0.666666667 | 1 | # | 10 | Baseline | ancom | 64 |
| # | interaction | 0 | 0 | 2 | 0 | 0 | # | 10 | Baseline | ancom | 64 |
| # | disease | 4 | 16 | 0 | 0.2 | 1 | # | 10 | Baseline | aldex2 | 64 |

| | | | | | | | | | | | |
|---|---|---|---|---|---|---|---|---|---|---|---|
| # | sex | 2 | 8 | 0 | 0.2 | 1 | # | 10 | Baseline | aldex2 | 64 |
| # | time | 2 | 12 | 0 | 0.142857143 | 1 | # | 10 | Baseline | aldex2 | 64 |
| # | interaction | 0 | 0 | 2 | 0 | 0 | # | 10 | Baseline | aldex2 | 64 |
| # | disease | 4 | 0 | 0 | 1 | 1 | # | 10 | Small | paralle | 64 |
| # | sex | 2 | 0 | 0 | 1 | 1 | # | 10 | Small | paralle | 64 |
| # | time | 2 | 0 | 0 | 1 | 1 | # | 10 | Small | paralle | 64 |
| # | interaction | 2 | 0 | 0 | 1 | 1 | # | 10 | Small | paralle | 64 |
| # | disease | 4 | 2 | 0 | 0.666666667 | 1 | # | 10 | Small | ancom | 64 |
| # | sex | 2 | 0 | 0 | 1 | 1 | # | 10 | Small | ancom | 64 |
| # | time | 2 | 1 | 0 | 0.666666667 | 1 | # | 10 | Small | ancom | 64 |
| # | interaction | 0 | 0 | 2 | 0 | 0 | # | 10 | Small | ancom | 64 |
| # | disease | 4 | 2 | 0 | 0.666666667 | 1 | # | 10 | Small | aldex2 | 64 |
| # | sex | 2 | 1 | 0 | 0.666666667 | 1 | # | 10 | Small | aldex2 | 64 |
| # | time | 2 | 3 | 0 | 0.4 | 1 | # | 10 | Small | aldex2 | 64 |
| # | interaction | 0 | 0 | 2 | 0 | 0 | # | 10 | Small | aldex2 | 64 |
| # | disease | 4 | 6 | 0 | 0.4 | 1 | # | 10 | Baseline | paralle | 64 |
| # | sex | 2 | 8 | 0 | 0.2 | 1 | # | 10 | Baseline | paralle | 64 |
| # | time | 2 | 6 | 0 | 0.25 | 1 | # | 10 | Baseline | paralle | 64 |
| # | interaction | 2 | 7 | 0 | 0.222222222 | 1 | # | 10 | Baseline | paralle | 64 |
| # | disease | 4 | 2 | 0 | 0.666666667 | 1 | # | 10 | Baseline | ancom | 64 |
| # | sex | 2 | 0 | 0 | 1 | 1 | # | 10 | Baseline | ancom | 64 |
| # | time | 2 | 2 | 0 | 0.5 | 1 | # | 10 | Baseline | ancom | 64 |
| # | interaction | 0 | 0 | 2 | 0 | 0 | # | 10 | Baseline | ancom | 64 |
| # | disease | 4 | 18 | 0 | 0.181818182 | 1 | # | 10 | Baseline | aldex2 | 64 |
| # | sex | 2 | 11 | 0 | 0.153846154 | 1 | # | 10 | Baseline | aldex2 | 64 |
| # | time | 2 | 12 | 0 | 0.142857143 | 1 | # | 10 | Baseline | aldex2 | 64 |
| # | interaction | 0 | 0 | 2 | 0 | 0 | # | 10 | Baseline | aldex2 | 64 |
| # | disease | 4 | 0 | 0 | 1 | 1 | # | 10 | Small | paralle | 64 |
| # | sex | 2 | 0 | 0 | 1 | 1 | # | 10 | Small | paralle | 64 |
| # | time | 2 | 0 | 0 | 1 | 1 | # | 10 | Small | paralle | 64 |
| # | interaction | 2 | 0 | 0 | 1 | 1 | # | 10 | Small | paralle | 64 |
| # | disease | 4 | 2 | 0 | 0.666666667 | 1 | # | 10 | Small | ancom | 64 |
| # | sex | 2 | 0 | 0 | 1 | 1 | # | 10 | Small | ancom | 64 |
| # | time | 2 | 2 | 0 | 0.5 | 1 | # | 10 | Small | ancom | 64 |
| # | interaction | 0 | 0 | 2 | 0 | 0 | # | 10 | Small | ancom | 64 |
| # | disease | 4 | 3 | 0 | 0.571428571 | 1 | # | 10 | Small | aldex2 | 64 |
| # | sex | 2 | 1 | 0 | 0.666666667 | 1 | # | 10 | Small | aldex2 | 64 |
| # | time | 2 | 3 | 0 | 0.4 | 1 | # | 10 | Small | aldex2 | 64 |
| # | interaction | 0 | 0 | 2 | 0 | 0 | # | 10 | Small | aldex2 | 64 |
| # | disease | 4 | 7 | 0 | 0.363636364 | 1 | # | 10 | Baseline | paralle | 64 |
| # | sex | 2 | 3 | 0 | 0.4 | 1 | # | 10 | Baseline | paralle | 64 |
| # | time | 2 | 4 | 0 | 0.333333333 | 1 | # | 10 | Baseline | paralle | 64 |
| # | interaction | 2 | 4 | 0 | 0.333333333 | 1 | # | 10 | Baseline | paralle | 64 |
| # | disease | 4 | 2 | 0 | 0.666666667 | 1 | # | 10 | Baseline | ancom | 64 |
| # | sex | 2 | 0 | 0 | 1 | 1 | # | 10 | Baseline | ancom | 64 |
| # | time | 2 | 2 | 0 | 0.5 | 1 | # | 10 | Baseline | ancom | 64 |
| # | interaction | 0 | 0 | 2 | 0 | 0 | # | 10 | Baseline | ancom | 64 |
| # | disease | 4 | 26 | 0 | 0.133333333 | 1 | # | 10 | Baseline | aldex2 | 64 |
| # | sex | 2 | 10 | 0 | 0.166666667 | 1 | # | 10 | Baseline | aldex2 | 64 |
| # | time | 2 | 17 | 0 | 0.105263158 | 1 | # | 10 | Baseline | aldex2 | 64 |
| # | interaction | 0 | 0 | 2 | 0 | 0 | # | 10 | Baseline | aldex2 | 64 |
| # | disease | 4 | 0 | 0 | 1 | 1 | # | 10 | Small | paralle | 64 |
| # | sex | 2 | 0 | 0 | 1 | 1 | # | 10 | Small | paralle | 64 |
| # | time | 2 | 0 | 0 | 1 | 1 | # | 10 | Small | paralle | 64 |
| # | interaction | 2 | 0 | 0 | 1 | 1 | # | 10 | Small | paralle | 64 |
| # | disease | 4 | 2 | 0 | 0.666666667 | 1 | # | 10 | Small | ancom | 64 |
| # | sex | 2 | 0 | 0 | 1 | 1 | # | 10 | Small | ancom | 64 |
| # | time | 2 | 2 | 0 | 0.5 | 1 | # | 10 | Small | ancom | 64 |
| # | interaction | 0 | 0 | 2 | 0 | 0 | # | 10 | Small | ancom | 64 |
| # | disease | 4 | 3 | 0 | 0.571428571 | 1 | # | 10 | Small | aldex2 | 64 |
| # | sex | 2 | 1 | 0 | 0.666666667 | 1 | # | 10 | Small | aldex2 | 64 |
| # | time | 2 | 4 | 0 | 0.333333333 | 1 | # | 10 | Small | aldex2 | 64 |
| # | interaction | 0 | 0 | 2 | 0 | 0 | # | 10 | Small | aldex2 | 64 |
| # | disease | 7 | 3 | 0 | 0.7 | 1 | # | 20 | Baseline | paralle | 64 |
| # | sex | 4 | 9 | 0 | 0.307692308 | 1 | # | 20 | Baseline | paralle | 64 |
| # | time | 5 | 8 | 0 | 0.384615385 | 1 | # | 20 | Baseline | paralle | 64 |
| # | interaction | 4 | 6 | 0 | 0.4 | 1 | # | 20 | Baseline | paralle | 64 |
| # | disease | 7 | 1 | 0 | 0.875 | 1 | # | 20 | Baseline | ancom | 64 |
| # | sex | 4 | 0 | 0 | 1 | 1 | # | 20 | Baseline | ancom | 64 |
| # | time | 5 | 1 | 0 | 0.833333333 | 1 | # | 20 | Baseline | ancom | 64 |
| # | interaction | 0 | 0 | 4 | 0 | 0 | # | 20 | Baseline | ancom | 64 |
| # | disease | 7 | 21 | 0 | 0.25 | 1 | # | 20 | Baseline | aldex2 | 64 |
| # | sex | 4 | 10 | 0 | 0.285714286 | 1 | # | 20 | Baseline | aldex2 | 64 |
| # | time | 5 | 11 | 0 | 0.3125 | 1 | # | 20 | Baseline | aldex2 | 64 |
| # | interaction | 0 | 0 | 4 | 0 | 0 | # | 20 | Baseline | aldex2 | 64 |
| # | disease | 7 | 1 | 0 | 0.875 | 1 | # | 20 | Small | paralle | 64 |
| # | sex | 4 | 0 | 0 | 1 | 1 | # | 20 | Small | paralle | 64 |
| # | time | 5 | 0 | 0 | 1 | 1 | # | 20 | Small | paralle | 64 |
| # | interaction | 3 | 1 | 1 | 0.75 | 0.75 | # | 20 | Small | paralle | 64 |
| # | disease | 7 | 1 | 0 | 0.875 | 1 | # | 20 | Small | ancom | 64 |
| # | sex | 4 | 0 | 0 | 1 | 1 | # | 20 | Small | ancom | 64 |
| # | time | 5 | 1 | 0 | 0.833333333 | 1 | # | 20 | Small | ancom | 64 |
| # | interaction | 0 | 0 | 4 | 0 | 0 | # | 20 | Small | ancom | 64 |
| # | disease | 7 | 0 | 0 | 1 | 1 | # | 20 | Small | aldex2 | 64 |
| # | sex | 4 | 0 | 0 | 1 | 1 | # | 20 | Small | aldex2 | 64 |
| # | time | 5 | 0 | 0 | 1 | 1 | # | 20 | Small | aldex2 | 64 |
| # | interaction | 0 | 0 | 4 | 0 | 0 | # | 20 | Small | aldex2 | 64 |
| # | disease | 7 | 9 | 0 | 0.4375 | 1 | # | 20 | Baseline | paralle | 64 |
| # | sex | 4 | 13 | 0 | 0.235294118 | 1 | # | 20 | Baseline | paralle | 64 |
| # | time | 5 | 5 | 0 | 0.5 | 1 | # | 20 | Baseline | paralle | 64 |
| # | interaction | 4 | 5 | 0 | 0.444444444 | 1 | # | 20 | Baseline | paralle | 64 |
| # | disease | 7 | 2 | 0 | 0.777777778 | 1 | # | 20 | Baseline | ancom | 64 |
| # | sex | 4 | 0 | 0 | 1 | 1 | # | 20 | Baseline | ancom | 64 |
| # | time | 5 | 1 | 0 | 0.833333333 | 1 | # | 20 | Baseline | ancom | 64 |
| # | interaction | 0 | 0 | 4 | 0 | 0 | # | 20 | Baseline | ancom | 64 |
| # | disease | 7 | 40 | 0 | 0.14893617 | 1 | # | 20 | Baseline | aldex2 | 64 |
| # | sex | 4 | 12 | 0 | 0.25 | 1 | # | 20 | Baseline | aldex2 | 64 |
| # | time | 5 | 24 | 0 | 0.172413793 | 1 | # | 20 | Baseline | aldex2 | 64 |
| # | interaction | 0 | 0 | 4 | 0 | 0 | # | 20 | Baseline | aldex2 | 64 |
| # | disease | 7 | 1 | 0 | 0.875 | 1 | # | 20 | Small | paralle | 64 |
| # | sex | 4 | 0 | 0 | 1 | 1 | # | 20 | Small | paralle | 64 |
| # | time | 5 | 0 | 0 | 1 | 1 | # | 20 | Small | paralle | 64 |
| # | interaction | 4 | 1 | 0 | 0.8 | 1 | # | 20 | Small | paralle | 64 |
| # | disease | 7 | 2 | 0 | 0.777777778 | 1 | # | 20 | Small | ancom | 64 |
| # | sex | 4 | 0 | 0 | 1 | 1 | # | 20 | Small | ancom | 64 |
| # | time | 5 | 1 | 0 | 0.833333333 | 1 | # | 20 | Small | ancom | 64 |
| # | interaction | 0 | 0 | 4 | 0 | 0 | # | 20 | Small | ancom | 64 |
| # | disease | 7 | 2 | 0 | 0.777777778 | 1 | # | 20 | Small | aldex2 | 64 |
| # | sex | 4 | 1 | 0 | 0.8 | 1 | # | 20 | Small | aldex2 | 64 |
| # | time | 5 | 1 | 0 | 0.833333333 | 1 | # | 20 | Small | aldex2 | 64 |
| # | interaction | 0 | 0 | 4 | 0 | 0 | # | 20 | Small | aldex2 | 64 |
| # | disease | 7 | 8 | 0 | 0.466666667 | 1 | # | 20 | Baseline | paralle | 64 |
| # | sex | 4 | 3 | 0 | 0.571428571 | 1 | # | 20 | Baseline | paralle | 64 |
| # | time | 5 | 10 | 0 | 0.333333333 | 1 | # | 20 | Baseline | paralle | 64 |
| # | interaction | 4 | 3 | 0 | 0.571428571 | 1 | # | 20 | Baseline | paralle | 64 |
| # | disease | 7 | 4 | 0 | 0.636363636 | 1 | # | 20 | Baseline | ancom | 64 |
| # | sex | 4 | 0 | 0 | 1 | 1 | # | 20 | Baseline | ancom | 64 |
| # | time | 5 | 4 | 0 | 0.555555556 | 1 | # | 20 | Baseline | ancom | 64 |
| # | interaction | 0 | 0 | 4 | 0 | 0 | # | 20 | Baseline | ancom | 64 |
| # | disease | 7 | 52 | 0 | 0.118644068 | 1 | # | 20 | Baseline | aldex2 | 64 |
| # | sex | 4 | 14 | 0 | 0.222222222 | 1 | # | 20 | Baseline | aldex2 | 64 |
| # | time | 5 | 35 | 0 | 0.125 | 1 | # | 20 | Baseline | aldex2 | 64 |
| # | interaction | 0 | 0 | 4 | 0 | 0 | # | 20 | Baseline | aldex2 | 64 |
| # | disease | 7 | 0 | 0 | 1 | 1 | # | 20 | Small | paralle | 64 |
| # | sex | 4 | 0 | 0 | 1 | 1 | # | 20 | Small | paralle | 64 |
| # | time | 5 | 0 | 0 | 1 | 1 | # | 20 | Small | paralle | 64 |
| # | interaction | 4 | 1 | 0 | 0.8 | 1 | # | 20 | Small | paralle | 64 |
| # | disease | 7 | 3 | 0 | 0.7 | 1 | # | 20 | Small | ancom | 64 |
| # | sex | 4 | 0 | 0 | 1 | 1 | # | 20 | Small | ancom | 64 |
| # | time | 5 | 4 | 0 | 0.555555556 | 1 | # | 20 | Small | ancom | 64 |
| # | interaction | 0 | 0 | 4 | 0 | 0 | # | 20 | Small | ancom | 64 |
| # | disease | 7 | 6 | 0 | 0.538461538 | 1 | # | 20 | Small | aldex2 | 64 |
| # | sex | 4 | 2 | 0 | 0.666666667 | 1 | # | 20 | Small | aldex2 | 64 |
| # | time | 5 | 5 | 0 | 0.5 | 1 | # | 20 | Small | aldex2 | 64 |
| # | interaction | 0 | 0 | 4 | 0 | 0 | # | 20 | Small | aldex2 | 64 |
| # | disease | 7 | 8 | 0 | 0.466666667 | 1 | # | 20 | Baseline | paralle | 64 |
| # | sex | 4 | 2 | 0 | 0.666666667 | 1 | # | 20 | Baseline | paralle | 64 |
| # | time | 5 | 8 | 0 | 0.384615385 | 1 | # | 20 | Baseline | paralle | 64 |
| # | interaction | 4 | 3 | 0 | 0.571428571 | 1 | # | 20 | Baseline | paralle | 64 |
| # | disease | 7 | 4 | 0 | 0.636363636 | 1 | # | 20 | Baseline | ancom | 64 |
| # | sex | 4 | 0 | 0 | 1 | 1 | # | 20 | Baseline | ancom | 64 |
| # | time | 5 | 4 | 0 | 0.555555556 | 1 | # | 20 | Baseline | ancom | 64 |
| # | interaction | 0 | 0 | 4 | 0 | 0 | # | 20 | Baseline | ancom | 64 |
| # | disease | 7 | 61 | 0 | 0.102941176 | 1 | # | 20 | Baseline | aldex2 | 64 |
| # | sex | 4 | 17 | 0 | 0.19047619 | 1 | # | 20 | Baseline | aldex2 | 64 |
| # | time | 5 | 35 | 0 | 0.125 | 1 | # | 20 | Baseline | aldex2 | 64 |
| # | interaction | 0 | 0 | 4 | 0 | 0 | # | 20 | Baseline | aldex2 | 64 |
| # | disease | 7 | 0 | 0 | 1 | 1 | # | 20 | Small | paralle | 64 |
| # | sex | 4 | 0 | 0 | 1 | 1 | # | 20 | Small | paralle | 64 |
| # | time | 5 | 0 | 0 | 1 | 1 | # | 20 | Small | paralle | 64 |
| # | interaction | 4 | 0 | 0 | 1 | 1 | # | 20 | Small | paralle | 64 |
| # | disease | 7 | 4 | 0 | 0.636363636 | 1 | # | 20 | Small | ancom | 64 |
| # | sex | 4 | 0 | 0 | 1 | 1 | # | 20 | Small | ancom | 64 |
| # | time | 5 | 4 | 0 | 0.555555556 | 1 | # | 20 | Small | ancom | 64 |
| # | interaction | 0 | 0 | 4 | 0 | 0 | # | 20 | Small | ancom | 64 |
| # | disease | 7 | 10 | 0 | 0.411764706 | 1 | # | 20 | Small | aldex2 | 64 |
| # | sex | 4 | 2 | 0 | 0.666666667 | 1 | # | 20 | Small | aldex2 | 64 |
| # | time | 5 | 3 | 0 | 0.625 | 1 | # | 20 | Small | aldex2 | 64 |
| # | interaction | 0 | 0 | 4 | 0 | 0 | # | 20 | Small | aldex2 | 64 |
| # | disease | 4 | 10 | 0 | 0.285714286 | 1 | # | 10 | Baseline | paralle | 65 |
| # | sex | 2 | 4 | 0 | 0.333333333 | 1 | # | 10 | Baseline | paralle | 65 |
| # | time | 2 | 8 | 0 | 0.2 | 1 | # | 10 | Baseline | paralle | 65 |
| # | interaction | 2 | 7 | 0 | 0.222222222 | 1 | # | 10 | Baseline | paralle | 65 |
| # | disease | 4 | 1 | 0 | 0.8 | 1 | # | 10 | Baseline | ancom | 65 |
| # | sex | 2 | 0 | 0 | 1 | 1 | # | 10 | Baseline | ancom | 65 |
| # | time | 2 | 0 | 0 | 1 | 1 | # | 10 | Baseline | ancom | 65 |
| # | interaction | 0 | 0 | 2 | 0 | 0 | # | 10 | Baseline | ancom | 65 |
| # | disease | 4 | 10 | 0 | 0.285714286 | 1 | # | 10 | Baseline | aldex2 | 65 |
| # | sex | 2 | 3 | 0 | 0.4 | 1 | # | 10 | Baseline | aldex2 | 65 |
| # | time | 2 | 3 | 0 | 0.4 | 1 | # | 10 | Baseline | aldex2 | 65 |
| # | interaction | 0 | 0 | 2 | 0 | 0 | # | 10 | Baseline | aldex2 | 65 |
| # | disease | 4 | 0 | 0 | 1 | 1 | # | 10 | Small | paralle | 65 |
| # | sex | 2 | 0 | 0 | 1 | 1 | # | 10 | Small | paralle | 65 |
| # | time | 2 | 0 | 0 | 1 | 1 | # | 10 | Small | paralle | 65 |
| # | interaction | 1 | 0 | 1 | 1 | 0.5 | # | 10 | Small | paralle | 65 |
| # | disease | 4 | 1 | 0 | 0.8 | 1 | # | 10 | Small | ancom | 65 |
| # | sex | 2 | 0 | 0 | 1 | 1 | # | 10 | Small | ancom | 65 |
| # | time | 2 | 0 | 0 | 1 | 1 | # | 10 | Small | ancom | 65 |
| # | interaction | 0 | 0 | 2 | 0 | 0 | # | 10 | Small | ancom | 65 |
| # | disease | 4 | 2 | 0 | 0.666666667 | 1 | # | 10 | Small | aldex2 | 65 |
| # | sex | 2 | 0 | 0 | 1 | 1 | # | 10 | Small | aldex2 | 65 |
| # | time | 2 | 0 | 0 | 1 | 1 | # | 10 | Small | aldex2 | 65 |
| # | interaction | 0 | 0 | 2 | 0 | 0 | # | 10 | Small | aldex2 | 65 |
| # | disease | 4 | 2 | 0 | 0.666666667 | 1 | # | 10 | Baseline | paralle | 65 |
| # | sex | 2 | 2 | 0 | 0.5 | 1 | # | 10 | Baseline | paralle | 65 |
| # | time | 2 | 3 | 0 | 0.4 | 1 | # | 10 | Baseline | paralle | 65 |
| # | interaction | 2 | 9 | 0 | 0.181818182 | 1 | # | 10 | Baseline | paralle | 65 |
| # | disease | 4 | 2 | 0 | 0.666666667 | 1 | # | 10 | Baseline | ancom | 65 |
| # | sex | 2 | 0 | 0 | 1 | 1 | # | 10 | Baseline | ancom | 65 |
| # | time | 2 | 1 | 0 | 0.666666667 | 1 | # | 10 | Baseline | ancom | 65 |
| # | interaction | 0 | 0 | 2 | 0 | 0 | # | 10 | Baseline | ancom | 65 |
| # | disease | 4 | 16 | 0 | 0.2 | 1 | # | 10 | Baseline | aldex2 | 65 |
| # | sex | 2 | 6 | 0 | 0.25 | 1 | # | 10 | Baseline | aldex2 | 65 |
| # | time | 2 | 10 | 0 | 0.166666667 | 1 | # | 10 | Baseline | aldex2 | 65 |
| # | interaction | 0 | 0 | 2 | 0 | 0 | # | 10 | Baseline | aldex2 | 65 |
| # | disease | 4 | 0 | 0 | 1 | 1 | # | 10 | Small | paralle | 65 |
| # | sex | 2 | 0 | 0 | 1 | 1 | # | 10 | Small | paralle | 65 |
| # | time | 2 | 0 | 0 | 1 | 1 | # | 10 | Small | paralle | 65 |
| # | interaction | 2 | 0 | 0 | 1 | 1 | # | 10 | Small | paralle | 65 |
| # | disease | 4 | 2 | 0 | 0.666666667 | 1 | # | 10 | Small | ancom | 65 |
| # | sex | 2 | 0 | 0 | 1 | 1 | # | 10 | Small | ancom | 65 |
| # | time | 2 | 1 | 0 | 0.666666667 | 1 | # | 10 | Small | ancom | 65 |
| # | interaction | 0 | 0 | 2 | 0 | 0 | # | 10 | Small | ancom | 65 |
| # | disease | 4 | 5 | 0 | 0.444444444 | 1 | # | 10 | Small | aldex2 | 65 |
| # | sex | 2 | 0 | 0 | 1 | 1 | # | 10 | Small | aldex2 | 65 |
| # | time | 2 | 2 | 0 | 0.5 | 1 | # | 10 | Small | aldex2 | 65 |
| # | interaction | 0 | 0 | 2 | 0 | 0 | # | 10 | Small | aldex2 | 65 |
| # | disease | 4 | 3 | 0 | 0.571428571 | 1 | # | 10 | Baseline | paralle | 65 |
| # | sex | 2 | 4 | 0 | 0.333333333 | 1 | # | 10 | Baseline | paralle | 65 |
| # | time | 2 | 4 | 0 | 0.333333333 | 1 | # | 10 | Baseline | paralle | 65 |
| # | interaction | 2 | 3 | 0 | 0.4 | 1 | # | 10 | Baseline | paralle | 65 |
| # | disease | 4 | 2 | 0 | 0.666666667 | 1 | # | 10 | Baseline | ancom | 65 |
| # | sex | 2 | 0 | 0 | 1 | 1 | # | 10 | Baseline | ancom | 65 |
| # | time | 2 | 2 | 0 | 0.5 | 1 | # | 10 | Baseline | ancom | 65 |
| # | interaction | 0 | 0 | 2 | 0 | 0 | # | 10 | Baseline | ancom | 65 |
| # | disease | 4 | 20 | 0 | 0.166666667 | 1 | # | 10 | Baseline | aldex2 | 65 |
| # | sex | 2 | 6 | 0 | 0.25 | 1 | # | 10 | Baseline | aldex2 | 65 |
| # | time | 2 | 15 | 0 | 0.117647059 | 1 | # | 10 | Baseline | aldex2 | 65 |
| # | interaction | 0 | 0 | 2 | 0 | 0 | # | 10 | Baseline | aldex2 | 65 |
| # | disease | 4 | 0 | 0 | 1 | 1 | # | 10 | Small | paralle | 65 |
| # | sex | 2 | 0 | 0 | 1 | 1 | # | 10 | Small | paralle | 65 |
| # | time | 2 | 0 | 0 | 1 | 1 | # | 10 | Small | paralle | 65 |
| # | interaction | 2 | 0 | 0 | 1 | 1 | # | 10 | Small | paralle | 65 |
| # | disease | 4 | 2 | 0 | 0.666666667 | 1 | # | 10 | Small | ancom | 65 |
| # | sex | 2 | 0 | 0 | 1 | 1 | # | 10 | Small | ancom | 65 |
| # | time | 2 | 2 | 0 | 0.5 | 1 | # | 10 | Small | ancom | 65 |
| # | interaction | 0 | 0 | 2 | 0 | 0 | # | 10 | Small | ancom | 65 |
| # | disease | 4 | 2 | 0 | 0.666666667 | 1 | # | 10 | Small | aldex2 | 65 |
| # | sex | 2 | 0 | 0 | 1 | 1 | # | 10 | Small | aldex2 | 65 |
| # | time | 2 | 2 | 0 | 0.5 | 1 | # | 10 | Small | aldex2 | 65 |
| # | interaction | 0 | 0 | 2 | 0 | 0 | # | 10 | Small | aldex2 | 65 |
| # | disease | 4 | 4 | 0 | 0.5 | 1 | # | 10 | Baseline | paralle | 65 |
| # | sex | 2 | 3 | 0 | 0.4 | 1 | # | 10 | Baseline | paralle | 65 |
| # | time | 2 | 6 | 0 | 0.25 | 1 | # | 10 | Baseline | paralle | 65 |
| # | interaction | 2 | 4 | 0 | 0.333333333 | 1 | # | 10 | Baseline | paralle | 65 |
| # | disease | 4 | 2 | 0 | 0.666666667 | 1 | # | 10 | Baseline | ancom | 65 |
| # | sex | 2 | 0 | 0 | 1 | 1 | # | 10 | Baseline | ancom | 65 |
| # | time | 2 | 2 | 0 | 0.5 | 1 | # | 10 | Baseline | ancom | 65 |
| # | interaction | 0 | 0 | 2 | 0 | 0 | # | 10 | Baseline | ancom | 65 |
| # | disease | 4 | 23 | 0 | 0.148148148 | 1 | # | 10 | Baseline | aldex2 | 65 |
| # | sex | 2 | 7 | 0 | 0.222222222 | 1 | # | 10 | Baseline | aldex2 | 65 |
| # | time | 2 | 15 | 0 | 0.117647059 | 1 | # | 10 | Baseline | aldex2 | 65 |
| # | interaction | 0 | 0 | 2 | 0 | 0 | # | 10 | Baseline | aldex2 | 65 |
| # | disease | 4 | 0 | 0 | 1 | 1 | # | 10 | Small | paralle | 65 |
| # | sex | 2 | 0 | 0 | 1 | 1 | # | 10 | Small | paralle | 65 |
| # | time | 2 | 0 | 0 | 1 | 1 | # | 10 | Small | paralle | 65 |
| # | interaction | 2 | 0 | 0 | 1 | 1 | # | 10 | Small | paralle | 65 |
| # | disease | 4 | 2 | 0 | 0.666666667 | 1 | # | 10 | Small | ancom | 65 |
| # | sex | 2 | 0 | 0 | 1 | 1 | # | 10 | Small | ancom | 65 |
| # | time | 2 | 2 | 0 | 0.5 | 1 | # | 10 | Small | ancom | 65 |
| # | interaction | 0 | 0 | 2 | 0 | 0 | # | 10 | Small | ancom | 65 |
| # | disease | 4 | 5 | 0 | 0.444444444 | 1 | # | 10 | Small | aldex2 | 65 |
| # | sex | 2 | 0 | 0 | 1 | 1 | # | 10 | Small | aldex2 | 65 |
| # | time | 2 | 2 | 0 | 0.5 | 1 | # | 10 | Small | aldex2 | 65 |
| # | interaction | 0 | 0 | 2 | 0 | 0 | # | 10 | Small | aldex2 | 65 |
| # | disease | 7 | 7 | 0 | 0.5 | 1 | # | 20 | Baseline | paralle | 65 |
| # | sex | 4 | 9 | 0 | 0.307692308 | 1 | # | 20 | Baseline | paralle | 65 |
| # | time | 5 | 9 | 0 | 0.357142857 | 1 | # | 20 | Baseline | paralle | 65 |
| # | interaction | 4 | 5 | 0 | 0.444444444 | 1 | # | 20 | Baseline | paralle | 65 |
| # | disease | 7 | 2 | 0 | 0.777777778 | 1 | # | 20 | Baseline | ancom | 65 |
| # | sex | 4 | 0 | 0 | 1 | 1 | # | 20 | Baseline | ancom | 65 |
| # | time | 5 | 0 | 0 | 1 | 1 | # | 20 | Baseline | ancom | 65 |
| # | interaction | 0 | 0 | 4 | 0 | 0 | # | 20 | Baseline | ancom | 65 |
| # | disease | 7 | 20 | 0 | 0.259259259 | 1 | # | 20 | Baseline | aldex2 | 65 |
| # | sex | 4 | 10 | 0 | 0.285714286 | 1 | # | 20 | Baseline | aldex2 | 65 |
| # | time | 5 | 16 | 0 | 0.238095238 | 1 | # | 20 | Baseline | aldex2 | 65 |
| # | interaction | 0 | 0 | 4 | 0 | 0 | # | 20 | Baseline | aldex2 | 65 |
| # | disease | 7 | 0 | 0 | 1 | 1 | # | 20 | Small | paralle | 65 |
| # | sex | 4 | 1 | 0 | 0.8 | 1 | # | 20 | Small | paralle | 65 |
| # | time | 5 | 0 | 0 | 1 | 1 | # | 20 | Small | paralle | 65 |
| # | interaction | 2 | 0 | 2 | 1 | 0.5 | # | 20 | Small | paralle | 65 |
| # | disease | 7 | 2 | 0 | 0.777777778 | 1 | # | 20 | Small | ancom | 65 |
| # | sex | 4 | 0 | 0 | 1 | 1 | # | 20 | Small | ancom | 65 |
| # | time | 5 | 0 | 0 | 1 | 1 | # | 20 | Small | ancom | 65 |
| # | interaction | 0 | 0 | 4 | 0 | 0 | # | 20 | Small | ancom | 65 |
| # | disease | 7 | 2 | 0 | 0.777777778 | 1 | # | 20 | Small | aldex2 | 65 |
| # | sex | 4 | 0 | 0 | 1 | 1 | # | 20 | Small | aldex2 | 65 |
| # | time | 5 | 2 | 0 | 0.714285714 | 1 | # | 20 | Small | aldex2 | 65 |
| # | interaction | 0 | 0 | 4 | 0 | 0 | # | 20 | Small | aldex2 | 65 |
| # | disease | 7 | 9 | 0 | 0.4375 | 1 | # | 20 | Baseline | paralle | 65 |
| # | sex | 4 | 12 | 0 | 0.25 | 1 | # | 20 | Baseline | paralle | 65 |
| # | time | 5 | 12 | 0 | 0.294117647 | 1 | # | 20 | Baseline | paralle | 65 |
| # | interaction | 4 | 4 | 0 | 0.5 | 1 | # | 20 | Baseline | paralle | 65 |
| # | disease | 7 | 4 | 0 | 0.636363636 | 1 | # | 20 | Baseline | ancom | 65 |
| # | sex | 4 | 0 | 0 | 1 | 1 | # | 20 | Baseline | ancom | 65 |
| # | time | 5 | 3 | 0 | 0.625 | 1 | # | 20 | Baseline | ancom | 65 |
| # | interaction | 0 | 0 | 4 | 0 | 0 | # | 20 | Baseline | ancom | 65 |
| # | disease | 7 | 33 | 0 | 0.175 | 1 | # | 20 | Baseline | aldex2 | 65 |
| # | sex | 4 | 18 | 0 | 0.181818182 | 1 | # | 20 | Baseline | aldex2 | 65 |
| # | time | 5 | 21 | 0 | 0.192307692 | 1 | # | 20 | Baseline | aldex2 | 65 |
| # | interaction | 0 | 0 | 4 | 0 | 0 | # | 20 | Baseline | aldex2 | 65 |
| # | disease | 7 | 0 | 0 | 1 | 1 | # | 20 | Small | paralle | 65 |
| # | sex | 4 | 1 | 0 | 0.8 | 1 | # | 20 | Small | paralle | 65 |
| # | time | 5 | 0 | 0 | 1 | 1 | # | 20 | Small | paralle | 65 |
| # | interaction | 4 | 0 | 0 | 1 | 1 | # | 20 | Small | paralle | 65 |
| # | disease | 7 | 4 | 0 | 0.636363636 | 1 | # | 20 | Small | ancom | 65 |
| # | sex | 4 | 0 | 0 | 1 | 1 | # | 20 | Small | ancom | 65 |
| # | time | 5 | 4 | 0 | 0.555555556 | 1 | # | 20 | Small | ancom | 65 |
| # | interaction | 0 | 0 | 4 | 0 | 0 | # | 20 | Small | ancom | 65 |
| # | disease | 7 | 5 | 0 | 0.583333333 | 1 | # | 20 | Small | aldex2 | 65 |
| # | sex | 4 | 0 | 0 | 1 | 1 | # | 20 | Small | aldex2 | 65 |
| # | time | 5 | 4 | 0 | 0.555555556 | 1 | # | 20 | Small | aldex2 | 65 |
| # | interaction | 0 | 0 | 4 | 0 | 0 | # | 20 | Small | aldex2 | 65 |
| # | disease | 7 | 10 | 0 | 0.411764706 | 1 | # | 20 | Baseline | paralle | 65 |
| # | sex | 4 | 7 | 0 | 0.363636364 | 1 | # | 20 | Baseline | paralle | 65 |
| # | time | 5 | 6 | 0 | 0.454545455 | 1 | # | 20 | Baseline | paralle | 65 |
| # | interaction | 4 | 5 | 0 | 0.444444444 | 1 | # | 20 | Baseline | paralle | 65 |
| # | disease | 7 | 4 | 0 | 0.636363636 | 1 | # | 20 | Baseline | ancom | 65 |
| # | sex | 4 | 0 | 0 | 1 | 1 | # | 20 | Baseline | ancom | 65 |
| # | time | 5 | 4 | 0 | 0.555555556 | 1 | # | 20 | Baseline | ancom | 65 |
| # | interaction | 0 | 0 | 4 | 0 | 0 | # | 20 | Baseline | ancom | 65 |
| # | disease | 7 | 44 | 0 | 0.137254902 | 1 | # | 20 | Baseline | aldex2 | 65 |
| # | sex | 4 | 17 | 0 | 0.19047619 | 1 | # | 20 | Baseline | aldex2 | 65 |
| # | time | 5 | 27 | 0 | 0.15625 | 1 | # | 20 | Baseline | aldex2 | 65 |
| # | interaction | 0 | 0 | 4 | 0 | 0 | # | 20 | Baseline | aldex2 | 65 |
| # | disease | 7 | 0 | 0 | 1 | 1 | # | 20 | Small | paralle | 65 |
| # | sex | 4 | 1 | 0 | 0.8 | 1 | # | 20 | Small | paralle | 65 |
| # | time | 5 | 1 | 0 | 0.833333333 | 1 | # | 20 | Small | paralle | 65 |
| # | interaction | 4 | 0 | 0 | 1 | 1 | # | 20 | Small | paralle | 65 |
| # | disease | 7 | 4 | 0 | 0.636363636 | 1 | # | 20 | Small | ancom | 65 |
| # | sex | 4 | 0 | 0 | 1 | 1 | # | 20 | Small | ancom | 65 |
| # | time | 5 | 4 | 0 | 0.555555556 | 1 | # | 20 | Small | ancom | 65 |
| # | interaction | 0 | 0 | 4 | 0 | 0 | # | 20 | Small | ancom | 65 |
| # | disease | 7 | 5 | 0 | 0.583333333 | 1 | # | 20 | Small | aldex2 | 65 |
| # | sex | 4 | 0 | 0 | 1 | 1 | # | 20 | Small | aldex2 | 65 |
| # | time | 5 | 5 | 0 | 0.5 | 1 | # | 20 | Small | aldex2 | 65 |
| # | interaction | 0 | 0 | 4 | 0 | 0 | # | 20 | Small | aldex2 | 65 |
| # | disease | 7 | 9 | 0 | 0.4375 | 1 | # | 20 | Baseline | paralle | 65 |
| # | sex | 4 | 8 | 0 | 0.333333333 | 1 | # | 20 | Baseline | paralle | 65 |
| # | time | 5 | 2 | 0 | 0.714285714 | 1 | # | 20 | Baseline | paralle | 65 |
| # | interaction | 4 | 7 | 0 | 0.363636364 | 1 | # | 20 | Baseline | paralle | 65 |
| # | disease | 7 | 4 | 0 | 0.636363636 | 1 | # | 20 | Baseline | ancom | 65 |
| # | sex | 4 | 0 | 0 | 1 | 1 | # | 20 | Baseline | ancom | 65 |
| # | time | 5 | 4 | 0 | 0.555555556 | 1 | # | 20 | Baseline | ancom | 65 |
| # | interaction | 0 | 0 | 4 | 0 | 0 | # | 20 | Baseline | ancom | 65 |
| # | disease | 7 | 60 | 0 | 0.104477612 | 1 | # | 20 | Baseline | aldex2 | 65 |
| # | sex | 4 | 21 | 0 | 0.16 | 1 | # | 20 | Baseline | aldex2 | 65 |

| | | | | | | | | | | | |
|---|---|---|---|---|---|---|---|---|---|---|---|
| # | time | 5 | 34 | 0 | 0.128205128 | 1 | # | 20 | Baseline | aldex2 | 65 |
| # | interaction | 0 | 0 | 4 | 0 | 0 | # | 20 | Baseline | aldex2 | 65 |
| # | disease | 7 | 0 | 0 | 1 | 1 | # | 20 | Small | paralle | 65 |
| # | sex | 4 | 0 | 0 | 1 | 1 | # | 20 | Small | paralle | 65 |
| # | time | 5 | 0 | 0 | 1 | 1 | # | 20 | Small | paralle | 65 |
| # | interaction | 4 | 0 | 0 | 1 | 1 | # | 20 | Small | paralle | 65 |
| # | disease | 7 | 4 | 0 | 0.636363636 | 1 | # | 20 | Small | ancom | 65 |
| # | sex | 4 | 0 | 0 | 1 | 1 | # | 20 | Small | ancom | 65 |
| # | time | 5 | 4 | 0 | 0.555555556 | 1 | # | 20 | Small | ancom | 65 |
| # | interaction | 0 | 0 | 4 | 0 | 0 | # | 20 | Small | ancom | 65 |
| # | disease | 7 | 6 | 0 | 0.538461538 | 1 | # | 20 | Small | aldex2 | 65 |
| # | sex | 4 | 0 | 0 | 1 | 1 | # | 20 | Small | aldex2 | 65 |
| # | time | 5 | 6 | 0 | 0.454545455 | 1 | # | 20 | Small | aldex2 | 65 |
| # | interaction | 0 | 0 | 4 | 0 | 0 | # | 20 | Small | aldex2 | 65 |
| # | disease | 4 | 7 | 0 | 0.363636364 | 1 | # | 10 | Baseline | paralle | 66 |
| # | sex | 2 | 4 | 0 | 0.333333333 | 1 | # | 10 | Baseline | paralle | 66 |
| # | time | 2 | 4 | 0 | 0.333333333 | 1 | # | 10 | Baseline | paralle | 66 |
| # | interaction | 2 | 10 | 0 | 0.166666667 | 1 | # | 10 | Baseline | paralle | 66 |
| # | disease | 4 | 0 | 0 | 1 | 1 | # | 10 | Baseline | ancom | 66 |
| # | sex | 2 | 0 | 0 | 1 | 1 | # | 10 | Baseline | ancom | 66 |
| # | time | 2 | 0 | 0 | 1 | 1 | # | 10 | Baseline | ancom | 66 |
| # | interaction | 0 | 0 | 2 | 0 | 0 | # | 10 | Baseline | ancom | 66 |
| # | disease | 4 | 12 | 0 | 0.25 | 1 | # | 10 | Baseline | aldex2 | 66 |
| # | sex | 2 | 4 | 0 | 0.333333333 | 1 | # | 10 | Baseline | aldex2 | 66 |
| # | time | 2 | 4 | 0 | 0.333333333 | 1 | # | 10 | Baseline | aldex2 | 66 |
| # | interaction | 0 | 0 | 2 | 0 | 0 | # | 10 | Baseline | aldex2 | 66 |
| # | disease | 4 | 1 | 0 | 0.8 | 1 | # | 10 | Small | paralle | 66 |
| # | sex | 2 | 0 | 0 | 1 | 1 | # | 10 | Small | paralle | 66 |
| # | time | 2 | 0 | 0 | 1 | 1 | # | 10 | Small | paralle | 66 |
| # | interaction | 1 | 0 | 1 | 1 | 0.5 | # | 10 | Small | paralle | 66 |
| # | disease | 4 | 0 | 0 | 1 | 1 | # | 10 | Small | ancom | 66 |
| # | sex | 2 | 0 | 0 | 1 | 1 | # | 10 | Small | ancom | 66 |
| # | time | 2 | 0 | 0 | 1 | 1 | # | 10 | Small | ancom | 66 |
| # | interaction | 0 | 0 | 2 | 0 | 0 | # | 10 | Small | ancom | 66 |
| # | disease | 4 | 2 | 0 | 0.666666667 | 1 | # | 10 | Small | aldex2 | 66 |
| # | sex | 2 | 0 | 0 | 1 | 1 | # | 10 | Small | aldex2 | 66 |
| # | time | 2 | 1 | 0 | 0.666666667 | 1 | # | 10 | Small | aldex2 | 66 |
| # | interaction | 0 | 0 | 2 | 0 | 0 | # | 10 | Small | aldex2 | 66 |
| # | disease | 4 | 6 | 0 | 0.4 | 1 | # | 10 | Baseline | paralle | 66 |
| # | sex | 2 | 8 | 0 | 0.2 | 1 | # | 10 | Baseline | paralle | 66 |
| # | time | 2 | 6 | 0 | 0.25 | 1 | # | 10 | Baseline | paralle | 66 |
| # | interaction | 2 | 5 | 0 | 0.285714286 | 1 | # | 10 | Baseline | paralle | 66 |
| # | disease | 4 | 2 | 0 | 0.666666667 | 1 | # | 10 | Baseline | ancom | 66 |
| # | sex | 2 | 0 | 0 | 1 | 1 | # | 10 | Baseline | ancom | 66 |
| # | time | 2 | 2 | 0 | 0.5 | 1 | # | 10 | Baseline | ancom | 66 |
| # | interaction | 0 | 0 | 2 | 0 | 0 | # | 10 | Baseline | ancom | 66 |
| # | disease | 4 | 19 | 0 | 0.173913043 | 1 | # | 10 | Baseline | aldex2 | 66 |
| # | sex | 2 | 8 | 0 | 0.2 | 1 | # | 10 | Baseline | aldex2 | 66 |
| # | time | 2 | 9 | 0 | 0.181818182 | 1 | # | 10 | Baseline | aldex2 | 66 |
| # | interaction | 0 | 0 | 2 | 0 | 0 | # | 10 | Baseline | aldex2 | 66 |
| # | disease | 4 | 0 | 0 | 1 | 1 | # | 10 | Small | paralle | 66 |
| # | sex | 2 | 0 | 0 | 1 | 1 | # | 10 | Small | paralle | 66 |
| # | time | 2 | 1 | 0 | 0.666666667 | 1 | # | 10 | Small | paralle | 66 |
| # | interaction | 1 | 0 | 1 | 1 | 0.5 | # | 10 | Small | paralle | 66 |
| # | disease | 4 | 2 | 0 | 0.666666667 | 1 | # | 10 | Small | ancom | 66 |
| # | sex | 2 | 0 | 0 | 1 | 1 | # | 10 | Small | ancom | 66 |
| # | time | 2 | 2 | 0 | 0.5 | 1 | # | 10 | Small | ancom | 66 |
| # | interaction | 0 | 0 | 2 | 0 | 0 | # | 10 | Small | ancom | 66 |
| # | disease | 4 | 3 | 0 | 0.571428571 | 1 | # | 10 | Small | aldex2 | 66 |
| # | sex | 2 | 0 | 0 | 1 | 1 | # | 10 | Small | aldex2 | 66 |
| # | time | 2 | 2 | 0 | 0.5 | 1 | # | 10 | Small | aldex2 | 66 |
| # | interaction | 0 | 0 | 2 | 0 | 0 | # | 10 | Small | aldex2 | 66 |
| # | disease | 4 | 5 | 0 | 0.444444444 | 1 | # | 10 | Baseline | paralle | 66 |
| # | sex | 2 | 7 | 0 | 0.222222222 | 1 | # | 10 | Baseline | paralle | 66 |
| # | time | 2 | 3 | 0 | 0.4 | 1 | # | 10 | Baseline | paralle | 66 |
| # | interaction | 2 | 6 | 0 | 0.25 | 1 | # | 10 | Baseline | paralle | 66 |
| # | disease | 4 | 2 | 0 | 0.666666667 | 1 | # | 10 | Baseline | ancom | 66 |
| # | sex | 2 | 0 | 0 | 1 | 1 | # | 10 | Baseline | ancom | 66 |
| # | time | 2 | 2 | 0 | 0.5 | 1 | # | 10 | Baseline | ancom | 66 |
| # | interaction | 0 | 0 | 2 | 0 | 0 | # | 10 | Baseline | ancom | 66 |
| # | disease | 4 | 19 | 0 | 0.173913043 | 1 | # | 10 | Baseline | aldex2 | 66 |
| # | sex | 2 | 14 | 0 | 0.125 | 1 | # | 10 | Baseline | aldex2 | 66 |
| # | time | 2 | 11 | 0 | 0.153846154 | 1 | # | 10 | Baseline | aldex2 | 66 |
| # | interaction | 0 | 0 | 2 | 0 | 0 | # | 10 | Baseline | aldex2 | 66 |
| # | disease | 4 | 0 | 0 | 1 | 1 | # | 10 | Small | paralle | 66 |
| # | sex | 2 | 0 | 0 | 1 | 1 | # | 10 | Small | paralle | 66 |
| # | time | 2 | 0 | 0 | 1 | 1 | # | 10 | Small | paralle | 66 |
| # | interaction | 2 | 0 | 0 | 1 | 1 | # | 10 | Small | paralle | 66 |
| # | disease | 4 | 2 | 0 | 0.666666667 | 1 | # | 10 | Small | ancom | 66 |
| # | sex | 2 | 0 | 0 | 1 | 1 | # | 10 | Small | ancom | 66 |
| # | time | 2 | 2 | 0 | 0.5 | 1 | # | 10 | Small | ancom | 66 |
| # | interaction | 0 | 0 | 2 | 0 | 0 | # | 10 | Small | ancom | 66 |
| # | disease | 4 | 3 | 0 | 0.571428571 | 1 | # | 10 | Small | aldex2 | 66 |
| # | sex | 2 | 1 | 0 | 0.666666667 | 1 | # | 10 | Small | aldex2 | 66 |
| # | time | 2 | 2 | 0 | 0.5 | 1 | # | 10 | Small | aldex2 | 66 |
| # | interaction | 0 | 0 | 2 | 0 | 0 | # | 10 | Small | aldex2 | 66 |
| # | disease | 4 | 5 | 0 | 0.444444444 | 1 | # | 10 | Baseline | paralle | 66 |
| # | sex | 2 | 6 | 0 | 0.25 | 1 | # | 10 | Baseline | paralle | 66 |
| # | time | 2 | 4 | 0 | 0.333333333 | 1 | # | 10 | Baseline | paralle | 66 |
| # | interaction | 2 | 4 | 0 | 0.333333333 | 1 | # | 10 | Baseline | paralle | 66 |
| # | disease | 4 | 2 | 0 | 0.666666667 | 1 | # | 10 | Baseline | ancom | 66 |
| # | sex | 2 | 0 | 0 | 1 | 1 | # | 10 | Baseline | ancom | 66 |
| # | time | 2 | 2 | 0 | 0.5 | 1 | # | 10 | Baseline | ancom | 66 |
| # | interaction | 0 | 0 | 2 | 0 | 0 | # | 10 | Baseline | ancom | 66 |
| # | disease | 4 | 26 | 0 | 0.133333333 | 1 | # | 10 | Baseline | aldex2 | 66 |
| # | sex | 2 | 9 | 0 | 0.181818182 | 1 | # | 10 | Baseline | aldex2 | 66 |
| # | time | 2 | 10 | 0 | 0.166666667 | 1 | # | 10 | Baseline | aldex2 | 66 |
| # | interaction | 0 | 0 | 2 | 0 | 0 | # | 10 | Baseline | aldex2 | 66 |
| # | disease | 4 | 1 | 0 | 0.8 | 1 | # | 10 | Small | paralle | 66 |
| # | sex | 2 | 0 | 0 | 1 | 1 | # | 10 | Small | paralle | 66 |
| # | time | 2 | 0 | 0 | 1 | 1 | # | 10 | Small | paralle | 66 |
| # | interaction | 2 | 0 | 0 | 1 | 1 | # | 10 | Small | paralle | 66 |
| # | disease | 4 | 2 | 0 | 0.666666667 | 1 | # | 10 | Small | ancom | 66 |
| # | sex | 2 | 0 | 0 | 1 | 1 | # | 10 | Small | ancom | 66 |
| # | time | 2 | 2 | 0 | 0.5 | 1 | # | 10 | Small | ancom | 66 |
| # | interaction | 0 | 0 | 2 | 0 | 0 | # | 10 | Small | ancom | 66 |
| # | disease | 4 | 3 | 0 | 0.571428571 | 1 | # | 10 | Small | aldex2 | 66 |
| # | sex | 2 | 0 | 0 | 1 | 1 | # | 10 | Small | aldex2 | 66 |
| # | time | 2 | 3 | 0 | 0.4 | 1 | # | 10 | Small | aldex2 | 66 |
| # | interaction | 0 | 0 | 2 | 0 | 0 | # | 10 | Small | aldex2 | 66 |
| # | disease | 7 | 15 | 0 | 0.318181818 | 1 | # | 20 | Baseline | paralle | 66 |
| # | sex | 4 | 7 | 0 | 0.363636364 | 1 | # | 20 | Baseline | paralle | 66 |
| # | time | 5 | 12 | 0 | 0.294117647 | 1 | # | 20 | Baseline | paralle | 66 |
| # | interaction | 4 | 7 | 0 | 0.363636364 | 1 | # | 20 | Baseline | paralle | 66 |
| # | disease | 7 | 3 | 0 | 0.7 | 1 | # | 20 | Baseline | ancom | 66 |
| # | sex | 4 | 0 | 0 | 1 | 1 | # | 20 | Baseline | ancom | 66 |
| # | time | 5 | 1 | 0 | 0.833333333 | 1 | # | 20 | Baseline | ancom | 66 |
| # | interaction | 0 | 0 | 4 | 0 | 0 | # | 20 | Baseline | ancom | 66 |
| # | disease | 7 | 18 | 0 | 0.28 | 1 | # | 20 | Baseline | aldex2 | 66 |
| # | sex | 4 | 7 | 0 | 0.363636364 | 1 | # | 20 | Baseline | aldex2 | 66 |
| # | time | 5 | 25 | 0 | 0.166666667 | 1 | # | 20 | Baseline | aldex2 | 66 |
| # | interaction | 0 | 0 | 4 | 0 | 0 | # | 20 | Baseline | aldex2 | 66 |
| # | disease | 7 | 0 | 0 | 1 | 1 | # | 20 | Small | paralle | 66 |
| # | sex | 4 | 0 | 0 | 1 | 1 | # | 20 | Small | paralle | 66 |
| # | time | 5 | 0 | 0 | 1 | 1 | # | 20 | Small | paralle | 66 |
| # | interaction | 1 | 0 | 3 | 1 | 0.25 | # | 20 | Small | paralle | 66 |
| # | disease | 7 | 2 | 0 | 0.777777778 | 1 | # | 20 | Small | ancom | 66 |
| # | sex | 4 | 0 | 0 | 1 | 1 | # | 20 | Small | ancom | 66 |
| # | time | 5 | 0 | 0 | 1 | 1 | # | 20 | Small | ancom | 66 |
| # | interaction | 0 | 0 | 4 | 0 | 0 | # | 20 | Small | ancom | 66 |
| # | disease | 7 | 1 | 0 | 0.875 | 1 | # | 20 | Small | aldex2 | 66 |
| # | sex | 4 | 0 | 0 | 1 | 1 | # | 20 | Small | aldex2 | 66 |
| # | time | 5 | 1 | 0 | 0.833333333 | 1 | # | 20 | Small | aldex2 | 66 |
| # | interaction | 0 | 0 | 4 | 0 | 0 | # | 20 | Small | aldex2 | 66 |
| # | disease | 7 | 8 | 0 | 0.466666667 | 1 | # | 20 | Baseline | paralle | 66 |
| # | sex | 4 | 5 | 0 | 0.444444444 | 1 | # | 20 | Baseline | paralle | 66 |
| # | time | 5 | 14 | 0 | 0.263157895 | 1 | # | 20 | Baseline | paralle | 66 |
| # | interaction | 4 | 7 | 0 | 0.363636364 | 1 | # | 20 | Baseline | paralle | 66 |
| # | disease | 7 | 4 | 0 | 0.636363636 | 1 | # | 20 | Baseline | ancom | 66 |
| # | sex | 4 | 0 | 0 | 1 | 1 | # | 20 | Baseline | ancom | 66 |
| # | time | 5 | 3 | 0 | 0.625 | 1 | # | 20 | Baseline | ancom | 66 |
| # | interaction | 0 | 0 | 4 | 0 | 0 | # | 20 | Baseline | ancom | 66 |
| # | disease | 7 | 28 | 0 | 0.2 | 1 | # | 20 | Baseline | aldex2 | 66 |
| # | sex | 4 | 13 | 0 | 0.235294118 | 1 | # | 20 | Baseline | aldex2 | 66 |
| # | time | 5 | 31 | 0 | 0.138888889 | 1 | # | 20 | Baseline | aldex2 | 66 |
| # | interaction | 0 | 0 | 4 | 0 | 0 | # | 20 | Baseline | aldex2 | 66 |
| # | disease | 7 | 0 | 0 | 1 | 1 | # | 20 | Small | paralle | 66 |
| # | sex | 4 | 0 | 0 | 1 | 1 | # | 20 | Small | paralle | 66 |
| # | time | 5 | 0 | 0 | 1 | 1 | # | 20 | Small | paralle | 66 |
| # | interaction | 3 | 0 | 1 | 1 | 0.75 | # | 20 | Small | paralle | 66 |
| # | disease | 7 | 4 | 0 | 0.636363636 | 1 | # | 20 | Small | ancom | 66 |
| # | sex | 4 | 0 | 0 | 1 | 1 | # | 20 | Small | ancom | 66 |
| # | time | 5 | 2 | 0 | 0.714285714 | 1 | # | 20 | Small | ancom | 66 |
| # | interaction | 0 | 0 | 4 | 0 | 0 | # | 20 | Small | ancom | 66 |
| # | disease | 7 | 3 | 0 | 0.7 | 1 | # | 20 | Small | aldex2 | 66 |
| # | sex | 4 | 0 | 0 | 1 | 1 | # | 20 | Small | aldex2 | 66 |
| # | time | 5 | 3 | 0 | 0.625 | 1 | # | 20 | Small | aldex2 | 66 |
| # | interaction | 0 | 0 | 4 | 0 | 0 | # | 20 | Small | aldex2 | 66 |
| # | disease | 7 | 9 | 0 | 0.4375 | 1 | # | 20 | Baseline | paralle | 66 |
| # | sex | 4 | 8 | 0 | 0.333333333 | 1 | # | 20 | Baseline | paralle | 66 |
| # | time | 5 | 8 | 0 | 0.384615385 | 1 | # | 20 | Baseline | paralle | 66 |
| # | interaction | 4 | 5 | 0 | 0.444444444 | 1 | # | 20 | Baseline | paralle | 66 |
| # | disease | 7 | 4 | 0 | 0.636363636 | 1 | # | 20 | Baseline | ancom | 66 |
| # | sex | 4 | 0 | 0 | 1 | 1 | # | 20 | Baseline | ancom | 66 |
| # | time | 5 | 4 | 0 | 0.555555556 | 1 | # | 20 | Baseline | ancom | 66 |
| # | interaction | 0 | 0 | 4 | 0 | 0 | # | 20 | Baseline | ancom | 66 |
| # | disease | 7 | 37 | 0 | 0.159090909 | 1 | # | 20 | Baseline | aldex2 | 66 |
| # | sex | 4 | 15 | 0 | 0.210526316 | 1 | # | 20 | Baseline | aldex2 | 66 |
| # | time | 5 | 39 | 0 | 0.113636364 | 1 | # | 20 | Baseline | aldex2 | 66 |
| # | interaction | 0 | 0 | 4 | 0 | 0 | # | 20 | Baseline | aldex2 | 66 |
| # | disease | 7 | 0 | 0 | 1 | 1 | # | 20 | Small | paralle | 66 |
| # | sex | 4 | 0 | 0 | 1 | 1 | # | 20 | Small | paralle | 66 |
| # | time | 5 | 0 | 0 | 1 | 1 | # | 20 | Small | paralle | 66 |
| # | interaction | 3 | 0 | 1 | 1 | 0.75 | # | 20 | Small | paralle | 66 |
| # | disease | 7 | 4 | 0 | 0.636363636 | 1 | # | 20 | Small | ancom | 66 |
| # | sex | 4 | 0 | 0 | 1 | 1 | # | 20 | Small | ancom | 66 |
| # | time | 5 | 4 | 0 | 0.555555556 | 1 | # | 20 | Small | ancom | 66 |
| # | interaction | 0 | 0 | 4 | 0 | 0 | # | 20 | Small | ancom | 66 |
| # | disease | 7 | 7 | 0 | 0.5 | 1 | # | 20 | Small | aldex2 | 66 |
| # | sex | 4 | 0 | 0 | 1 | 1 | # | 20 | Small | aldex2 | 66 |
| # | time | 5 | 6 | 0 | 0.454545455 | 1 | # | 20 | Small | aldex2 | 66 |
| # | interaction | 0 | 0 | 4 | 0 | 0 | # | 20 | Small | aldex2 | 66 |
| # | disease | 7 | 7 | 0 | 0.5 | 1 | # | 20 | Baseline | paralle | 66 |
| # | sex | 4 | 6 | 0 | 0.4 | 1 | # | 20 | Baseline | paralle | 66 |
| # | time | 5 | 6 | 0 | 0.454545455 | 1 | # | 20 | Baseline | paralle | 66 |
| # | interaction | 4 | 5 | 0 | 0.444444444 | 1 | # | 20 | Baseline | paralle | 66 |
| # | disease | 7 | 4 | 0 | 0.636363636 | 1 | # | 20 | Baseline | ancom | 66 |
| # | sex | 4 | 0 | 0 | 1 | 1 | # | 20 | Baseline | ancom | 66 |
| # | time | 5 | 4 | 0 | 0.555555556 | 1 | # | 20 | Baseline | ancom | 66 |
| # | interaction | 0 | 0 | 4 | 0 | 0 | # | 20 | Baseline | ancom | 66 |
| # | disease | 7 | 62 | 0 | 0.142857143 | 1 | # | 20 | Baseline | aldex2 | 66 |
| # | sex | 4 | 20 | 0 | 0.166666667 | 1 | # | 20 | Baseline | aldex2 | 66 |
| # | time | 5 | 47 | 0 | 0.096153846 | 1 | # | 20 | Baseline | aldex2 | 66 |
| # | interaction | 0 | 0 | 4 | 0 | 0 | # | 20 | Baseline | aldex2 | 66 |
| # | disease | 7 | 0 | 0 | 1 | 1 | # | 20 | Small | paralle | 66 |
| # | sex | 4 | 0 | 0 | 1 | 1 | # | 20 | Small | paralle | 66 |
| # | time | 5 | 0 | 0 | 1 | 1 | # | 20 | Small | paralle | 66 |
| # | interaction | 4 | 0 | 0 | 1 | 1 | # | 20 | Small | paralle | 66 |
| # | disease | 7 | 4 | 0 | 0.636363636 | 1 | # | 20 | Small | ancom | 66 |
| # | sex | 4 | 0 | 0 | 1 | 1 | # | 20 | Small | ancom | 66 |
| # | time | 5 | 4 | 0 | 0.555555556 | 1 | # | 20 | Small | ancom | 66 |
| # | interaction | 0 | 0 | 4 | 0 | 0 | # | 20 | Small | ancom | 66 |
| # | disease | 7 | 5 | 0 | 0.583333333 | 1 | # | 20 | Small | aldex2 | 66 |
| # | sex | 4 | 0 | 0 | 1 | 1 | # | 20 | Small | aldex2 | 66 |
| # | time | 5 | 8 | 0 | 0.384615385 | 1 | # | 20 | Small | aldex2 | 66 |
| # | interaction | 0 | 0 | 4 | 0 | 0 | # | 20 | Small | aldex2 | 66 |
| # | disease | 4 | 8 | 0 | 0.333333333 | 1 | # | 10 | Baseline | paralle | 67 |
| # | sex | 2 | 4 | 0 | 0.333333333 | 1 | # | 10 | Baseline | paralle | 67 |
| # | time | 2 | 5 | 0 | 0.285714286 | 1 | # | 10 | Baseline | paralle | 67 |
| # | interaction | 2 | 5 | 0 | 0.285714286 | 1 | # | 10 | Baseline | paralle | 67 |
| # | disease | 4 | 0 | 0 | 1 | 1 | # | 10 | Baseline | ancom | 67 |
| # | sex | 2 | 0 | 0 | 1 | 1 | # | 10 | Baseline | ancom | 67 |
| # | time | 2 | 0 | 0 | 1 | 1 | # | 10 | Baseline | ancom | 67 |
| # | interaction | 0 | 0 | 2 | 0 | 0 | # | 10 | Baseline | ancom | 67 |
| # | disease | 4 | 9 | 0 | 0.307692308 | 1 | # | 10 | Baseline | aldex2 | 67 |
| # | sex | 2 | 4 | 0 | 0.333333333 | 1 | # | 10 | Baseline | aldex2 | 67 |
| # | time | 2 | 8 | 0 | 0.2 | 1 | # | 10 | Baseline | aldex2 | 67 |
| # | interaction | 0 | 0 | 2 | 0 | 0 | # | 10 | Baseline | aldex2 | 67 |
| # | disease | 4 | 1 | 0 | 0.8 | 1 | # | 10 | Small | paralle | 67 |
| # | sex | 2 | 0 | 0 | 1 | 1 | # | 10 | Small | paralle | 67 |
| # | time | 2 | 1 | 0 | 0.666666667 | 1 | # | 10 | Small | paralle | 67 |
| # | interaction | 2 | 1 | 0 | 0.666666667 | 1 | # | 10 | Small | paralle | 67 |
| # | disease | 4 | 0 | 0 | 1 | 1 | # | 10 | Small | ancom | 67 |
| # | sex | 2 | 0 | 0 | 1 | 1 | # | 10 | Small | ancom | 67 |
| # | time | 2 | 0 | 0 | 1 | 1 | # | 10 | Small | ancom | 67 |
| # | interaction | 0 | 0 | 2 | 0 | 0 | # | 10 | Small | ancom | 67 |
| # | disease | 4 | 1 | 0 | 0.8 | 1 | # | 10 | Small | aldex2 | 67 |
| # | sex | 2 | 0 | 0 | 1 | 1 | # | 10 | Small | aldex2 | 67 |
| # | time | 2 | 1 | 0 | 0.666666667 | 1 | # | 10 | Small | aldex2 | 67 |
| # | interaction | 0 | 0 | 2 | 0 | 0 | # | 10 | Small | aldex2 | 67 |
| # | disease | 4 | 4 | 0 | 0.5 | 1 | # | 10 | Baseline | paralle | 67 |
| # | sex | 2 | 6 | 0 | 0.25 | 1 | # | 10 | Baseline | paralle | 67 |
| # | time | 2 | 5 | 0 | 0.285714286 | 1 | # | 10 | Baseline | paralle | 67 |
| # | interaction | 2 | 5 | 0 | 0.285714286 | 1 | # | 10 | Baseline | paralle | 67 |
| # | disease | 4 | 1 | 0 | 0.8 | 1 | # | 10 | Baseline | ancom | 67 |
| # | sex | 2 | 0 | 0 | 1 | 1 | # | 10 | Baseline | ancom | 67 |
| # | time | 2 | 1 | 0 | 0.666666667 | 1 | # | 10 | Baseline | ancom | 67 |
| # | interaction | 0 | 0 | 2 | 0 | 0 | # | 10 | Baseline | ancom | 67 |
| # | disease | 4 | 19 | 0 | 0.173913043 | 1 | # | 10 | Baseline | aldex2 | 67 |
| # | sex | 2 | 8 | 0 | 0.2 | 1 | # | 10 | Baseline | aldex2 | 67 |
| # | time | 2 | 8 | 0 | 0.2 | 1 | # | 10 | Baseline | aldex2 | 67 |
| # | interaction | 0 | 0 | 2 | 0 | 0 | # | 10 | Baseline | aldex2 | 67 |
| # | disease | 4 | 0 | 0 | 1 | 1 | # | 10 | Small | paralle | 67 |
| # | sex | 2 | 0 | 0 | 1 | 1 | # | 10 | Small | paralle | 67 |
| # | time | 2 | 0 | 0 | 1 | 1 | # | 10 | Small | paralle | 67 |
| # | interaction | 2 | 0 | 0 | 1 | 1 | # | 10 | Small | paralle | 67 |
| # | disease | 4 | 1 | 0 | 0.8 | 1 | # | 10 | Small | ancom | 67 |
| # | sex | 2 | 0 | 0 | 1 | 1 | # | 10 | Small | ancom | 67 |
| # | time | 2 | 1 | 0 | 0.666666667 | 1 | # | 10 | Small | ancom | 67 |
| # | interaction | 0 | 0 | 2 | 0 | 0 | # | 10 | Small | ancom | 67 |
| # | disease | 4 | 2 | 0 | 0.666666667 | 1 | # | 10 | Small | aldex2 | 67 |
| # | sex | 2 | 0 | 0 | 1 | 1 | # | 10 | Small | aldex2 | 67 |
| # | time | 2 | 1 | 0 | 0.666666667 | 1 | # | 10 | Small | aldex2 | 67 |
| # | interaction | 0 | 0 | 2 | 0 | 0 | # | 10 | Small | aldex2 | 67 |
| # | disease | 4 | 8 | 0 | 0.333333333 | 1 | # | 10 | Baseline | paralle | 67 |
| # | sex | 2 | 2 | 0 | 0.5 | 1 | # | 10 | Baseline | paralle | 67 |
| # | time | 2 | 5 | 0 | 0.285714286 | 1 | # | 10 | Baseline | paralle | 67 |
| # | interaction | 2 | 1 | 0 | 0.666666667 | 1 | # | 10 | Baseline | paralle | 67 |
| # | disease | 4 | 2 | 0 | 0.666666667 | 1 | # | 10 | Baseline | ancom | 67 |
| # | sex | 2 | 0 | 0 | 1 | 1 | # | 10 | Baseline | ancom | 67 |
| # | time | 2 | 2 | 0 | 0.5 | 1 | # | 10 | Baseline | ancom | 67 |
| # | interaction | 0 | 0 | 2 | 0 | 0 | # | 10 | Baseline | ancom | 67 |
| # | disease | 4 | 20 | 0 | 0.166666667 | 1 | # | 10 | Baseline | aldex2 | 67 |
| # | sex | 2 | 3 | 0 | 0.4 | 1 | # | 10 | Baseline | aldex2 | 67 |
| # | time | 2 | 10 | 0 | 0.166666667 | 1 | # | 10 | Baseline | aldex2 | 67 |
| # | interaction | 0 | 0 | 2 | 0 | 0 | # | 10 | Baseline | aldex2 | 67 |
| # | disease | 4 | 0 | 0 | 1 | 1 | # | 10 | Small | paralle | 67 |
| # | sex | 2 | 0 | 0 | 1 | 1 | # | 10 | Small | paralle | 67 |
| # | time | 2 | 0 | 0 | 1 | 1 | # | 10 | Small | paralle | 67 |
| # | interaction | 2 | 0 | 0 | 1 | 1 | # | 10 | Small | paralle | 67 |
| # | disease | 4 | 2 | 0 | 0.666666667 | 1 | # | 10 | Small | ancom | 67 |
| # | sex | 2 | 0 | 0 | 1 | 1 | # | 10 | Small | ancom | 67 |
| # | time | 2 | 2 | 0 | 0.5 | 1 | # | 10 | Small | ancom | 67 |
| # | interaction | 0 | 0 | 2 | 0 | 0 | # | 10 | Small | ancom | 67 |
| # | disease | 4 | 2 | 0 | 0.666666667 | 1 | # | 10 | Small | aldex2 | 67 |
| # | sex | 2 | 0 | 0 | 1 | 1 | # | 10 | Small | aldex2 | 67 |
| # | time | 2 | 2 | 0 | 0.5 | 1 | # | 10 | Small | aldex2 | 67 |
| # | interaction | 0 | 0 | 2 | 0 | 0 | # | 10 | Small | aldex2 | 67 |
| # | disease | 4 | 7 | 0 | 0.363636364 | 1 | # | 10 | Baseline | paralle | 67 |
| # | sex | 2 | 0 | 0 | 1 | 1 | # | 10 | Baseline | paralle | 67 |
| # | time | 2 | 4 | 0 | 0.333333333 | 1 | # | 10 | Baseline | paralle | 67 |
| # | interaction | 2 | 1 | 0 | 0.666666667 | 1 | # | 10 | Baseline | paralle | 67 |
| # | disease | 4 | 2 | 0 | 0.666666667 | 1 | # | 10 | Baseline | ancom | 67 |
| # | sex | 2 | 0 | 0 | 1 | 1 | # | 10 | Baseline | ancom | 67 |
| # | time | 2 | 2 | 0 | 0.5 | 1 | # | 10 | Baseline | ancom | 67 |
| # | interaction | 0 | 0 | 2 | 0 | 0 | # | 10 | Baseline | ancom | 67 |
| # | disease | 4 | 24 | 0 | 0.142857143 | 1 | # | 10 | Baseline | aldex2 | 67 |
| # | sex | 2 | 7 | 0 | 0.222222222 | 1 | # | 10 | Baseline | aldex2 | 67 |
| # | time | 2 | 9 | 0 | 0.181818182 | 1 | # | 10 | Baseline | aldex2 | 67 |
| # | interaction | 0 | 0 | 2 | 0 | 0 | # | 10 | Baseline | aldex2 | 67 |
| # | disease | 4 | 0 | 0 | 1 | 1 | # | 10 | Small | paralle | 67 |
| # | sex | 2 | 0 | 0 | 1 | 1 | # | 10 | Small | paralle | 67 |
| # | time | 2 | 0 | 0 | 1 | 1 | # | 10 | Small | paralle | 67 |
| # | interaction | 2 | 0 | 0 | 1 | 1 | # | 10 | Small | paralle | 67 |
| # | disease | 4 | 2 | 0 | 0.666666667 | 1 | # | 10 | Small | ancom | 67 |
| # | sex | 2 | 0 | 0 | 1 | 1 | # | 10 | Small | ancom | 67 |
| # | time | 2 | 2 | 0 | 0.5 | 1 | # | 10 | Small | ancom | 67 |
| # | interaction | 0 | 0 | 2 | 0 | 0 | # | 10 | Small | ancom | 67 |
| # | disease | 4 | 3 | 0 | 0.571428571 | 1 | # | 10 | Small | aldex2 | 67 |
| # | sex | 2 | 0 | 0 | 1 | 1 | # | 10 | Small | aldex2 | 67 |
| # | time | 2 | 2 | 0 | 0.5 | 1 | # | 10 | Small | aldex2 | 67 |
| # | interaction | 0 | 0 | 2 | 0 | 0 | # | 10 | Small | aldex2 | 67 |
| # | disease | 7 | 11 | 0 | 0.388888889 | 1 | # | 20 | Baseline | paralle | 67 |
| # | sex | 4 | 12 | 0 | 0.25 | 1 | # | 20 | Baseline | paralle | 67 |
| # | time | 5 | 11 | 0 | 0.3125 | 1 | # | 20 | Baseline | paralle | 67 |
| # | interaction | 4 | 9 | 0 | 0.307692308 | 1 | # | 20 | Baseline | paralle | 67 |
| # | disease | 7 | 2 | 0 | 0.777777778 | 1 | # | 20 | Baseline | ancom | 67 |
| # | sex | 4 | 0 | 0 | 1 | 1 | # | 20 | Baseline | ancom | 67 |
| # | time | 5 | 2 | 0 | 0.714285714 | 1 | # | 20 | Baseline | ancom | 67 |
| # | interaction | 0 | 0 | 4 | 0 | 0 | # | 20 | Baseline | ancom | 67 |
| # | disease | 7 | 23 | 0 | 0.233333333 | 1 | # | 20 | Baseline | aldex2 | 67 |
| # | sex | 4 | 4 | 0 | 0.5 | 1 | # | 20 | Baseline | aldex2 | 67 |
| # | time | 5 | 13 | 0 | 0.277777778 | 1 | # | 20 | Baseline | aldex2 | 67 |
| # | interaction | 0 | 0 | 4 | 0 | 0 | # | 20 | Baseline | aldex2 | 67 |
| # | disease | 7 | 2 | 0 | 0.777777778 | 1 | # | 20 | Small | paralle | 67 |
| # | sex | 4 | 0 | 0 | 1 | 1 | # | 20 | Small | paralle | 67 |
| # | time | 5 | 1 | 0 | 0.833333333 | 1 | # | 20 | Small | paralle | 67 |
| # | interaction | 1 | 1 | 3 | 0.5 | 0.25 | # | 20 | Small | paralle | 67 |
| # | disease | 7 | 2 | 0 | 0.777777778 | 1 | # | 20 | Small | ancom | 67 |
| # | sex | 4 | 0 | 0 | 1 | 1 | # | 20 | Small | ancom | 67 |
| # | time | 5 | 1 | 0 | 0.833333333 | 1 | # | 20 | Small | ancom | 67 |
| # | interaction | 0 | 0 | 4 | 0 | 0 | # | 20 | Small | ancom | 67 |
| # | disease | 7 | 1 | 0 | 0.875 | 1 | # | 20 | Small | aldex2 | 67 |
| # | sex | 4 | 0 | 0 | 1 | 1 | # | 20 | Small | aldex2 | 67 |
| # | time | 5 | 2 | 0 | 0.714285714 | 1 | # | 20 | Small | aldex2 | 67 |
| # | interaction | 0 | 0 | 4 | 0 | 0 | # | 20 | Small | aldex2 | 67 |
| # | disease | 7 | 10 | 0 | 0.411764706 | 1 | # | 20 | Baseline | paralle | 67 |
| # | sex | 4 | 7 | 0 | 0.363636364 | 1 | # | 20 | Baseline | paralle | 67 |
| # | time | 5 | 4 | 0 | 0.555555556 | 1 | # | 20 | Baseline | paralle | 67 |
| # | interaction | 4 | 2 | 0 | 0.666666667 | 1 | # | 20 | Baseline | paralle | 67 |
| # | disease | 7 | 4 | 0 | 0.636363636 | 1 | # | 20 | Baseline | ancom | 67 |
| # | sex | 4 | 0 | 0 | 1 | 1 | # | 20 | Baseline | ancom | 67 |
| # | time | 5 | 3 | 0 | 0.625 | 1 | # | 20 | Baseline | ancom | 67 |
| # | interaction | 0 | 0 | 4 | 0 | 0 | # | 20 | Baseline | ancom | 67 |
| # | disease | 7 | 37 | 0 | 0.159090909 | 1 | # | 20 | Baseline | aldex2 | 67 |
| # | sex | 4 | 14 | 0 | 0.222222222 | 1 | # | 20 | Baseline | aldex2 | 67 |
| # | time | 5 | 18 | 0 | 0.217391304 | 1 | # | 20 | Baseline | aldex2 | 67 |

| | | | | | | | | | | | |
|---|---|---|---|---|---|---|---|---|---|---|---|
| # | interaction | 0 | 0 | 4 | 0 | 0 | # | 20 | Baseline | aldex2 | 67 |
| # | disease | 7 | 1 | 0 | 0.875 | 1 | # | 20 | Small | paralle | 67 |
| # | sex | 4 | 0 | 0 | 1 | 1 | # | 20 | Small | paralle | 67 |
| # | time | 5 | 0 | 0 | 1 | 1 | # | 20 | Small | paralle | 67 |
| # | interaction | 4 | 0 | 0 | 1 | 1 | # | 20 | Small | paralle | 67 |
| # | disease | 7 | 4 | 0 | 0.636363636 | 1 | # | 20 | Small | ancom | 67 |
| # | sex | 4 | 0 | 0 | 1 | 1 | # | 20 | Small | ancom | 67 |
| # | time | 5 | 2 | 0 | 0.714285714 | 1 | # | 20 | Small | ancom | 67 |
| # | interaction | 0 | 0 | 4 | 0 | 0 | # | 20 | Small | ancom | 67 |
| # | disease | 7 | 3 | 0 | 0.7 | 1 | # | 20 | Small | aldex2 | 67 |
| # | sex | 4 | 2 | 0 | 0.666666667 | 1 | # | 20 | Small | aldex2 | 67 |
| # | time | 5 | 2 | 0 | 0.714285714 | 1 | # | 20 | Small | aldex2 | 67 |
| # | interaction | 0 | 0 | 4 | 0 | 0 | # | 20 | Small | aldex2 | 67 |
| # | disease | 7 | 6 | 0 | 0.538461538 | 1 | # | 20 | Baseline | paralle | 67 |
| # | sex | 4 | 5 | 0 | 0.444444444 | 1 | # | 20 | Baseline | paralle | 67 |
| # | time | 5 | 8 | 0 | 0.384615385 | 1 | # | 20 | Baseline | paralle | 67 |
| # | interaction | 4 | 4 | 0 | 0.5 | 1 | # | 20 | Baseline | paralle | 67 |
| # | disease | 7 | 4 | 0 | 0.636363636 | 1 | # | 20 | Baseline | ancom | 67 |
| # | sex | 4 | 0 | 0 | 1 | 1 | # | 20 | Baseline | ancom | 67 |
| # | time | 5 | 4 | 0 | 0.555555556 | 1 | # | 20 | Baseline | ancom | 67 |
| # | interaction | 0 | 0 | 4 | 0 | 0 | # | 20 | Baseline | ancom | 67 |
| # | disease | 7 | 67 | 0 | 0.12962963 | 1 | # | 20 | Baseline | aldex2 | 67 |
| # | sex | 4 | 14 | 0 | 0.222222222 | 1 | # | 20 | Baseline | aldex2 | 67 |
| # | time | 5 | 28 | 0 | 0.151515152 | 1 | # | 20 | Baseline | aldex2 | 67 |
| # | interaction | 0 | 0 | 4 | 0 | 0 | # | 20 | Baseline | aldex2 | 67 |
| # | disease | 7 | 0 | 0 | 1 | 1 | # | 20 | Small | paralle | 67 |
| # | sex | 4 | 0 | 0 | 1 | 1 | # | 20 | Small | paralle | 67 |
| # | time | 5 | 0 | 0 | 1 | 1 | # | 20 | Small | paralle | 67 |
| # | interaction | 4 | 0 | 0 | 1 | 1 | # | 20 | Small | paralle | 67 |
| # | disease | 7 | 4 | 0 | 0.636363636 | 1 | # | 20 | Small | ancom | 67 |
| # | sex | 4 | 0 | 0 | 1 | 1 | # | 20 | Small | ancom | 67 |
| # | time | 5 | 4 | 0 | 0.555555556 | 1 | # | 20 | Small | ancom | 67 |
| # | interaction | 0 | 0 | 4 | 0 | 0 | # | 20 | Small | ancom | 67 |
| # | disease | 7 | 6 | 0 | 0.538461538 | 1 | # | 20 | Small | aldex2 | 67 |
| # | sex | 4 | 3 | 0 | 0.571428571 | 1 | # | 20 | Small | aldex2 | 67 |
| # | time | 5 | 4 | 0 | 0.555555556 | 1 | # | 20 | Small | aldex2 | 67 |
| # | interaction | 0 | 0 | 4 | 0 | 0 | # | 20 | Small | aldex2 | 67 |
| # | disease | 7 | 8 | 0 | 0.466666667 | 1 | # | 20 | Baseline | paralle | 67 |
| # | sex | 4 | 6 | 0 | 0.4 | 1 | # | 20 | Baseline | paralle | 67 |
| # | time | 5 | 7 | 0 | 0.416666667 | 1 | # | 20 | Baseline | paralle | 67 |
| # | interaction | 4 | 3 | 0 | 0.571428571 | 1 | # | 20 | Baseline | paralle | 67 |
| # | disease | 7 | 4 | 0 | 0.636363636 | 1 | # | 20 | Baseline | ancom | 67 |
| # | sex | 4 | 0 | 0 | 1 | 1 | # | 20 | Baseline | ancom | 67 |
| # | time | 5 | 4 | 0 | 0.555555556 | 1 | # | 20 | Baseline | ancom | 67 |
| # | interaction | 0 | 0 | 4 | 0 | 0 | # | 20 | Baseline | ancom | 67 |
| # | disease | 7 | 58 | 0 | 0.107692308 | 1 | # | 20 | Baseline | aldex2 | 67 |
| # | sex | 4 | 14 | 0 | 0.222222222 | 1 | # | 20 | Baseline | aldex2 | 67 |
| # | time | 5 | 33 | 0 | 0.131578947 | 1 | # | 20 | Baseline | aldex2 | 67 |
| # | interaction | 0 | 0 | 4 | 0 | 0 | # | 20 | Baseline | aldex2 | 67 |
| # | disease | 7 | 0 | 0 | 1 | 1 | # | 20 | Small | paralle | 67 |
| # | sex | 4 | 0 | 0 | 1 | 1 | # | 20 | Small | paralle | 67 |
| # | time | 5 | 0 | 0 | 1 | 1 | # | 20 | Small | paralle | 67 |
| # | interaction | 4 | 0 | 0 | 1 | 1 | # | 20 | Small | paralle | 67 |
| # | disease | 7 | 4 | 0 | 0.636363636 | 1 | # | 20 | Small | ancom | 67 |
| # | sex | 4 | 0 | 0 | 1 | 1 | # | 20 | Small | ancom | 67 |
| # | time | 5 | 4 | 0 | 0.555555556 | 1 | # | 20 | Small | ancom | 67 |
| # | interaction | 0 | 0 | 4 | 0 | 0 | # | 20 | Small | ancom | 67 |
| # | disease | 7 | 6 | 0 | 0.538461538 | 1 | # | 20 | Small | aldex2 | 67 |
| # | sex | 4 | 3 | 0 | 0.571428571 | 1 | # | 20 | Small | aldex2 | 67 |
| # | time | 5 | 4 | 0 | 0.555555556 | 1 | # | 20 | Small | aldex2 | 67 |
| # | interaction | 0 | 0 | 4 | 0 | 0 | # | 20 | Small | aldex2 | 67 |
| # | disease | 4 | 8 | 0 | 0.333333333 | 1 | # | 10 | Baseline | paralle | 68 |
| # | sex | 2 | 6 | 0 | 0.25 | 1 | # | 10 | Baseline | paralle | 68 |
| # | time | 2 | 5 | 0 | 0.285714286 | 1 | # | 10 | Baseline | paralle | 68 |
| # | interaction | 2 | 5 | 0 | 0.285714286 | 1 | # | 10 | Baseline | paralle | 68 |
| # | disease | 4 | 0 | 0 | 1 | 1 | # | 10 | Baseline | ancom | 68 |
| # | sex | 2 | 0 | 0 | 1 | 1 | # | 10 | Baseline | ancom | 68 |
| # | time | 2 | 1 | 0 | 0.666666667 | 1 | # | 10 | Baseline | ancom | 68 |
| # | interaction | 0 | 0 | 2 | 0 | 0 | # | 10 | Baseline | ancom | 68 |
| # | disease | 4 | 8 | 0 | 0.333333333 | 1 | # | 10 | Baseline | aldex2 | 68 |
| # | sex | 2 | 3 | 0 | 0.4 | 1 | # | 10 | Baseline | aldex2 | 68 |
| # | time | 2 | 7 | 0 | 0.222222222 | 1 | # | 10 | Baseline | aldex2 | 68 |
| # | interaction | 0 | 0 | 2 | 0 | 0 | # | 10 | Baseline | aldex2 | 68 |
| # | disease | 4 | 0 | 0 | 1 | 1 | # | 10 | Small | paralle | 68 |
| # | sex | 2 | 0 | 0 | 1 | 1 | # | 10 | Small | paralle | 68 |
| # | time | 2 | 0 | 0 | 1 | 1 | # | 10 | Small | paralle | 68 |
| # | interaction | 1 | 1 | 1 | 0.5 | 0.5 | # | 10 | Small | paralle | 68 |
| # | disease | 4 | 0 | 0 | 1 | 1 | # | 10 | Small | ancom | 68 |
| # | sex | 2 | 0 | 0 | 1 | 1 | # | 10 | Small | ancom | 68 |
| # | time | 2 | 0 | 0 | 1 | 1 | # | 10 | Small | ancom | 68 |
| # | interaction | 0 | 0 | 2 | 0 | 0 | # | 10 | Small | ancom | 68 |
| # | disease | 4 | 0 | 0 | 1 | 1 | # | 10 | Small | aldex2 | 68 |
| # | sex | 2 | 1 | 0 | 0.666666667 | 1 | # | 10 | Small | aldex2 | 68 |
| # | time | 2 | 2 | 0 | 0.5 | 1 | # | 10 | Small | aldex2 | 68 |
| # | interaction | 0 | 0 | 2 | 0 | 0 | # | 10 | Small | aldex2 | 68 |
| # | disease | 4 | 8 | 0 | 0.333333333 | 1 | # | 10 | Baseline | paralle | 68 |
| # | sex | 2 | 4 | 0 | 0.333333333 | 1 | # | 10 | Baseline | paralle | 68 |
| # | time | 2 | 5 | 0 | 0.285714286 | 1 | # | 10 | Baseline | paralle | 68 |
| # | interaction | 2 | 4 | 0 | 0.333333333 | 1 | # | 10 | Baseline | paralle | 68 |
| # | disease | 4 | 2 | 0 | 0.666666667 | 1 | # | 10 | Baseline | ancom | 68 |
| # | sex | 2 | 0 | 0 | 1 | 1 | # | 10 | Baseline | ancom | 68 |
| # | time | 2 | 2 | 0 | 0.5 | 1 | # | 10 | Baseline | ancom | 68 |
| # | interaction | 0 | 0 | 2 | 0 | 0 | # | 10 | Baseline | ancom | 68 |
| # | disease | 4 | 20 | 0 | 0.166666667 | 1 | # | 10 | Baseline | aldex2 | 68 |
| # | sex | 2 | 3 | 0 | 0.4 | 1 | # | 10 | Baseline | aldex2 | 68 |
| # | time | 2 | 7 | 0 | 0.222222222 | 1 | # | 10 | Baseline | aldex2 | 68 |
| # | interaction | 0 | 0 | 2 | 0 | 0 | # | 10 | Baseline | aldex2 | 68 |
| # | disease | 4 | 0 | 0 | 1 | 1 | # | 10 | Small | paralle | 68 |
| # | sex | 2 | 0 | 0 | 1 | 1 | # | 10 | Small | paralle | 68 |
| # | time | 2 | 0 | 0 | 1 | 1 | # | 10 | Small | paralle | 68 |
| # | interaction | 2 | 0 | 0 | 1 | 1 | # | 10 | Small | paralle | 68 |
| # | disease | 4 | 2 | 0 | 0.666666667 | 1 | # | 10 | Small | ancom | 68 |
| # | sex | 2 | 0 | 0 | 1 | 1 | # | 10 | Small | ancom | 68 |
| # | time | 2 | 2 | 0 | 0.5 | 1 | # | 10 | Small | ancom | 68 |
| # | interaction | 0 | 0 | 2 | 0 | 0 | # | 10 | Small | ancom | 68 |
| # | disease | 4 | 2 | 0 | 0.666666667 | 1 | # | 10 | Small | aldex2 | 68 |
| # | sex | 2 | 0 | 0 | 1 | 1 | # | 10 | Small | aldex2 | 68 |
| # | time | 2 | 2 | 0 | 0.5 | 1 | # | 10 | Small | aldex2 | 68 |
| # | interaction | 0 | 0 | 2 | 0 | 0 | # | 10 | Small | aldex2 | 68 |
| # | disease | 4 | 5 | 0 | 0.444444444 | 1 | # | 10 | Baseline | paralle | 68 |
| # | sex | 2 | 3 | 0 | 0.4 | 1 | # | 10 | Baseline | paralle | 68 |
| # | time | 2 | 6 | 0 | 0.25 | 1 | # | 10 | Baseline | paralle | 68 |
| # | interaction | 2 | 2 | 0 | 0.5 | 1 | # | 10 | Baseline | paralle | 68 |
| # | disease | 4 | 2 | 0 | 0.666666667 | 1 | # | 10 | Baseline | ancom | 68 |
| # | sex | 2 | 0 | 0 | 1 | 1 | # | 10 | Baseline | ancom | 68 |
| # | time | 2 | 2 | 0 | 0.5 | 1 | # | 10 | Baseline | ancom | 68 |
| # | interaction | 0 | 0 | 2 | 0 | 0 | # | 10 | Baseline | ancom | 68 |
| # | disease | 4 | 22 | 0 | 0.153846154 | 1 | # | 10 | Baseline | aldex2 | 68 |
| # | sex | 2 | 6 | 0 | 0.25 | 1 | # | 10 | Baseline | aldex2 | 68 |
| # | time | 2 | 7 | 0 | 0.222222222 | 1 | # | 10 | Baseline | aldex2 | 68 |
| # | interaction | 0 | 0 | 2 | 0 | 0 | # | 10 | Baseline | aldex2 | 68 |
| # | disease | 4 | 0 | 0 | 1 | 1 | # | 10 | Small | paralle | 68 |
| # | sex | 2 | 0 | 0 | 1 | 1 | # | 10 | Small | paralle | 68 |
| # | time | 2 | 0 | 0 | 1 | 1 | # | 10 | Small | paralle | 68 |
| # | interaction | 2 | 0 | 0 | 1 | 1 | # | 10 | Small | paralle | 68 |
| # | disease | 4 | 2 | 0 | 0.666666667 | 1 | # | 10 | Small | ancom | 68 |
| # | sex | 2 | 0 | 0 | 1 | 1 | # | 10 | Small | ancom | 68 |
| # | time | 2 | 2 | 0 | 0.5 | 1 | # | 10 | Small | ancom | 68 |
| # | interaction | 0 | 0 | 2 | 0 | 0 | # | 10 | Small | ancom | 68 |
| # | disease | 4 | 4 | 0 | 0.5 | 1 | # | 10 | Small | aldex2 | 68 |
| # | sex | 2 | 0 | 0 | 1 | 1 | # | 10 | Small | aldex2 | 68 |
| # | time | 2 | 2 | 0 | 0.5 | 1 | # | 10 | Small | aldex2 | 68 |
| # | interaction | 0 | 0 | 2 | 0 | 0 | # | 10 | Small | aldex2 | 68 |
| # | disease | 4 | 5 | 0 | 0.444444444 | 1 | # | 10 | Baseline | paralle | 68 |
| # | sex | 2 | 2 | 0 | 0.5 | 1 | # | 10 | Baseline | paralle | 68 |
| # | time | 2 | 3 | 0 | 0.4 | 1 | # | 10 | Baseline | paralle | 68 |
| # | interaction | 2 | 3 | 0 | 0.4 | 1 | # | 10 | Baseline | paralle | 68 |
| # | disease | 4 | 2 | 0 | 0.666666667 | 1 | # | 10 | Baseline | ancom | 68 |
| # | sex | 2 | 0 | 0 | 1 | 1 | # | 10 | Baseline | ancom | 68 |
| # | time | 2 | 2 | 0 | 0.5 | 1 | # | 10 | Baseline | ancom | 68 |
| # | interaction | 0 | 0 | 2 | 0 | 0 | # | 10 | Baseline | ancom | 68 |
| # | disease | 4 | 29 | 0 | 0.121212121 | 1 | # | 10 | Baseline | aldex2 | 68 |
| # | sex | 2 | 5 | 0 | 0.285714286 | 1 | # | 10 | Baseline | aldex2 | 68 |
| # | time | 2 | 8 | 0 | 0.2 | 1 | # | 10 | Baseline | aldex2 | 68 |
| # | interaction | 0 | 0 | 2 | 0 | 0 | # | 10 | Baseline | aldex2 | 68 |
| # | disease | 4 | 0 | 0 | 1 | 1 | # | 10 | Small | paralle | 68 |
| # | sex | 2 | 0 | 0 | 1 | 1 | # | 10 | Small | paralle | 68 |
| # | time | 2 | 0 | 0 | 1 | 1 | # | 10 | Small | paralle | 68 |
| # | interaction | 2 | 0 | 0 | 1 | 1 | # | 10 | Small | paralle | 68 |
| # | disease | 4 | 2 | 0 | 0.666666667 | 1 | # | 10 | Small | ancom | 68 |
| # | sex | 2 | 0 | 0 | 1 | 1 | # | 10 | Small | ancom | 68 |
| # | time | 2 | 2 | 0 | 0.5 | 1 | # | 10 | Small | ancom | 68 |
| # | interaction | 0 | 0 | 2 | 0 | 0 | # | 10 | Small | ancom | 68 |
| # | disease | 4 | 5 | 0 | 0.444444444 | 1 | # | 10 | Small | aldex2 | 68 |
| # | sex | 2 | 0 | 0 | 1 | 1 | # | 10 | Small | aldex2 | 68 |
| # | time | 2 | 2 | 0 | 0.5 | 1 | # | 10 | Small | aldex2 | 68 |
| # | interaction | 0 | 0 | 2 | 0 | 0 | # | 10 | Small | aldex2 | 68 |
| # | disease | 7 | 12 | 0 | 0.368421053 | 1 | # | 20 | Baseline | paralle | 68 |
| # | sex | 4 | 7 | 0 | 0.363636364 | 1 | # | 20 | Baseline | paralle | 68 |
| # | time | 5 | 6 | 0 | 0.454545455 | 1 | # | 20 | Baseline | paralle | 68 |
| # | interaction | 4 | 9 | 0 | 0.307692308 | 1 | # | 20 | Baseline | paralle | 68 |
| # | disease | 7 | 1 | 0 | 0.875 | 1 | # | 20 | Baseline | ancom | 68 |
| # | sex | 4 | 0 | 0 | 1 | 1 | # | 20 | Baseline | ancom | 68 |
| # | time | 5 | 3 | 0 | 0.625 | 1 | # | 20 | Baseline | ancom | 68 |
| # | interaction | 0 | 0 | 4 | 0 | 0 | # | 20 | Baseline | ancom | 68 |
| # | disease | 7 | 23 | 0 | 0.233333333 | 1 | # | 20 | Baseline | aldex2 | 68 |
| # | sex | 4 | 8 | 0 | 0.333333333 | 1 | # | 20 | Baseline | aldex2 | 68 |
| # | time | 5 | 11 | 0 | 0.3125 | 1 | # | 20 | Baseline | aldex2 | 68 |
| # | interaction | 0 | 0 | 4 | 0 | 0 | # | 20 | Baseline | aldex2 | 68 |
| # | disease | 7 | 2 | 0 | 0.777777778 | 1 | # | 20 | Small | paralle | 68 |
| # | sex | 4 | 1 | 0 | 0.8 | 1 | # | 20 | Small | paralle | 68 |
| # | time | 5 | 0 | 0 | 1 | 1 | # | 20 | Small | paralle | 68 |
| # | interaction | 1 | 0 | 3 | 1 | 0.25 | # | 20 | Small | paralle | 68 |
| # | disease | 7 | 0 | 0 | 1 | 1 | # | 20 | Small | ancom | 68 |
| # | sex | 4 | 0 | 0 | 1 | 1 | # | 20 | Small | ancom | 68 |
| # | time | 5 | 3 | 0 | 0.625 | 1 | # | 20 | Small | ancom | 68 |
| # | interaction | 0 | 0 | 4 | 0 | 0 | # | 20 | Small | ancom | 68 |
| # | disease | 7 | 0 | 0 | 1 | 1 | # | 20 | Small | aldex2 | 68 |
| # | sex | 4 | 0 | 0 | 1 | 1 | # | 20 | Small | aldex2 | 68 |
| # | time | 5 | 1 | 0 | 0.833333333 | 1 | # | 20 | Small | aldex2 | 68 |
| # | interaction | 0 | 0 | 4 | 0 | 0 | # | 20 | Small | aldex2 | 68 |
| # | disease | 7 | 7 | 0 | 0.5 | 1 | # | 20 | Baseline | paralle | 68 |
| # | sex | 4 | 8 | 0 | 0.333333333 | 1 | # | 20 | Baseline | paralle | 68 |
| # | time | 5 | 8 | 0 | 0.384615385 | 1 | # | 20 | Baseline | paralle | 68 |
| # | interaction | 4 | 8 | 0 | 0.333333333 | 1 | # | 20 | Baseline | paralle | 68 |
| # | disease | 7 | 2 | 0 | 0.777777778 | 1 | # | 20 | Baseline | ancom | 68 |
| # | sex | 4 | 0 | 0 | 1 | 1 | # | 20 | Baseline | ancom | 68 |
| # | time | 5 | 3 | 0 | 0.625 | 1 | # | 20 | Baseline | ancom | 68 |
| # | interaction | 0 | 0 | 4 | 0 | 0 | # | 20 | Baseline | ancom | 68 |
| # | disease | 7 | 34 | 0 | 0.170731707 | 1 | # | 20 | Baseline | aldex2 | 68 |
| # | sex | 4 | 10 | 0 | 0.285714286 | 1 | # | 20 | Baseline | aldex2 | 68 |
| # | time | 5 | 19 | 0 | 0.208333333 | 1 | # | 20 | Baseline | aldex2 | 68 |
| # | interaction | 0 | 0 | 4 | 0 | 0 | # | 20 | Baseline | aldex2 | 68 |
| # | disease | 7 | 0 | 0 | 1 | 1 | # | 20 | Small | paralle | 68 |
| # | sex | 4 | 2 | 0 | 0.666666667 | 1 | # | 20 | Small | paralle | 68 |
| # | time | 5 | 0 | 0 | 1 | 1 | # | 20 | Small | paralle | 68 |
| # | interaction | 3 | 0 | 1 | 1 | 0.75 | # | 20 | Small | paralle | 68 |
| # | disease | 7 | 2 | 0 | 0.777777778 | 1 | # | 20 | Small | ancom | 68 |
| # | sex | 4 | 0 | 0 | 1 | 1 | # | 20 | Small | ancom | 68 |
| # | time | 5 | 3 | 0 | 0.625 | 1 | # | 20 | Small | ancom | 68 |
| # | interaction | 0 | 0 | 4 | 0 | 0 | # | 20 | Small | ancom | 68 |
| # | disease | 7 | 1 | 0 | 0.875 | 1 | # | 20 | Small | aldex2 | 68 |
| # | sex | 4 | 1 | 0 | 0.8 | 1 | # | 20 | Small | aldex2 | 68 |
| # | time | 5 | 2 | 0 | 0.714285714 | 1 | # | 20 | Small | aldex2 | 68 |
| # | interaction | 0 | 0 | 4 | 0 | 0 | # | 20 | Small | aldex2 | 68 |
| # | disease | 7 | 8 | 0 | 0.466666667 | 1 | # | 20 | Baseline | paralle | 68 |
| # | sex | 4 | 6 | 0 | 0.4 | 1 | # | 20 | Baseline | paralle | 68 |
| # | time | 5 | 11 | 0 | 0.3125 | 1 | # | 20 | Baseline | paralle | 68 |
| # | interaction | 4 | 3 | 0 | 0.571428571 | 1 | # | 20 | Baseline | paralle | 68 |
| # | disease | 7 | 4 | 0 | 0.636363636 | 1 | # | 20 | Baseline | ancom | 68 |
| # | sex | 4 | 0 | 0 | 1 | 1 | # | 20 | Baseline | ancom | 68 |
| # | time | 5 | 4 | 0 | 0.555555556 | 1 | # | 20 | Baseline | ancom | 68 |
| # | interaction | 0 | 0 | 4 | 0 | 0 | # | 20 | Baseline | ancom | 68 |
| # | disease | 7 | 50 | 0 | 0.122807018 | 1 | # | 20 | Baseline | aldex2 | 68 |
| # | sex | 4 | 14 | 0 | 0.222222222 | 1 | # | 20 | Baseline | aldex2 | 68 |
| # | time | 5 | 33 | 0 | 0.131578947 | 1 | # | 20 | Baseline | aldex2 | 68 |
| # | interaction | 0 | 0 | 4 | 0 | 0 | # | 20 | Baseline | aldex2 | 68 |
| # | disease | 7 | 0 | 0 | 1 | 1 | # | 20 | Small | paralle | 68 |
| # | sex | 4 | 1 | 0 | 0.8 | 1 | # | 20 | Small | paralle | 68 |
| # | time | 5 | 0 | 0 | 1 | 1 | # | 20 | Small | paralle | 68 |
| # | interaction | 4 | 0 | 0 | 1 | 1 | # | 20 | Small | paralle | 68 |
| # | disease | 7 | 4 | 0 | 0.636363636 | 1 | # | 20 | Small | ancom | 68 |
| # | sex | 4 | 0 | 0 | 1 | 1 | # | 20 | Small | ancom | 68 |
| # | time | 5 | 4 | 0 | 0.555555556 | 1 | # | 20 | Small | ancom | 68 |
| # | interaction | 0 | 0 | 4 | 0 | 0 | # | 20 | Small | ancom | 68 |
| # | disease | 7 | 7 | 0 | 0.5 | 1 | # | 20 | Small | aldex2 | 68 |
| # | sex | 4 | 0 | 0 | 1 | 1 | # | 20 | Small | aldex2 | 68 |
| # | time | 5 | 3 | 0 | 0.625 | 1 | # | 20 | Small | aldex2 | 68 |
| # | interaction | 0 | 0 | 4 | 0 | 0 | # | 20 | Small | aldex2 | 68 |
| # | disease | 7 | 12 | 0 | 0.368421053 | 1 | # | 20 | Baseline | paralle | 68 |
| # | sex | 4 | 6 | 0 | 0.4 | 1 | # | 20 | Baseline | paralle | 68 |
| # | time | 5 | 14 | 0 | 0.263157895 | 1 | # | 20 | Baseline | paralle | 68 |
| # | interaction | 4 | 4 | 0 | 0.5 | 1 | # | 20 | Baseline | paralle | 68 |
| # | disease | 7 | 4 | 0 | 0.636363636 | 1 | # | 20 | Baseline | ancom | 68 |
| # | sex | 4 | 0 | 0 | 1 | 1 | # | 20 | Baseline | ancom | 68 |
| # | time | 5 | 4 | 0 | 0.555555556 | 1 | # | 20 | Baseline | ancom | 68 |
| # | interaction | 0 | 0 | 4 | 0 | 0 | # | 20 | Baseline | ancom | 68 |
| # | disease | 7 | 54 | 0 | 0.098591549 | 1 | # | 20 | Baseline | aldex2 | 68 |
| # | sex | 4 | 19 | 0 | 0.173913043 | 1 | # | 20 | Baseline | aldex2 | 68 |
| # | time | 5 | 39 | 0 | 0.113636364 | 1 | # | 20 | Baseline | aldex2 | 68 |
| # | interaction | 0 | 0 | 4 | 0 | 0 | # | 20 | Baseline | aldex2 | 68 |
| # | disease | 7 | 0 | 0 | 1 | 1 | # | 20 | Small | paralle | 68 |
| # | sex | 4 | 1 | 0 | 0.8 | 1 | # | 20 | Small | paralle | 68 |
| # | time | 5 | 0 | 0 | 1 | 1 | # | 20 | Small | paralle | 68 |
| # | interaction | 4 | 0 | 0 | 1 | 1 | # | 20 | Small | paralle | 68 |
| # | disease | 7 | 4 | 0 | 0.636363636 | 1 | # | 20 | Small | ancom | 68 |
| # | sex | 4 | 0 | 0 | 1 | 1 | # | 20 | Small | ancom | 68 |
| # | time | 5 | 4 | 0 | 0.555555556 | 1 | # | 20 | Small | ancom | 68 |
| # | interaction | 0 | 0 | 4 | 0 | 0 | # | 20 | Small | ancom | 68 |
| # | disease | 7 | 6 | 0 | 0.538461538 | 1 | # | 20 | Small | aldex2 | 68 |
| # | sex | 4 | 1 | 0 | 0.8 | 1 | # | 20 | Small | aldex2 | 68 |
| # | time | 5 | 5 | 0 | 0.5 | 1 | # | 20 | Small | aldex2 | 68 |
| # | interaction | 0 | 0 | 4 | 0 | 0 | # | 20 | Small | aldex2 | 68 |
| # | disease | 4 | 6 | 0 | 0.4 | 1 | # | 10 | Baseline | paralle | 69 |
| # | sex | 2 | 7 | 0 | 0.222222222 | 1 | # | 10 | Baseline | paralle | 69 |
| # | time | 2 | 7 | 0 | 0.222222222 | 1 | # | 10 | Baseline | paralle | 69 |
| # | interaction | 2 | 11 | 0 | 0.153846154 | 1 | # | 10 | Baseline | paralle | 69 |
| # | disease | 4 | 0 | 0 | 1 | 1 | # | 10 | Baseline | ancom | 69 |
| # | sex | 2 | 0 | 0 | 1 | 1 | # | 10 | Baseline | ancom | 69 |
| # | time | 2 | 0 | 0 | 1 | 1 | # | 10 | Baseline | ancom | 69 |
| # | interaction | 0 | 0 | 2 | 0 | 0 | # | 10 | Baseline | ancom | 69 |
| # | disease | 4 | 14 | 0 | 0.222222222 | 1 | # | 10 | Baseline | aldex2 | 69 |
| # | sex | 2 | 2 | 0 | 0.5 | 1 | # | 10 | Baseline | aldex2 | 69 |
| # | time | 2 | 7 | 0 | 0.222222222 | 1 | # | 10 | Baseline | aldex2 | 69 |
| # | interaction | 0 | 0 | 2 | 0 | 0 | # | 10 | Baseline | aldex2 | 69 |
| # | disease | 4 | 0 | 0 | 1 | 1 | # | 10 | Small | paralle | 69 |
| # | sex | 2 | 0 | 0 | 1 | 1 | # | 10 | Small | paralle | 69 |
| # | time | 2 | 0 | 0 | 1 | 1 | # | 10 | Small | paralle | 69 |
| # | interaction | 1 | 0 | 1 | 1 | 0.5 | # | 10 | Small | paralle | 69 |
| # | disease | 4 | 0 | 0 | 1 | 1 | # | 10 | Small | ancom | 69 |
| # | sex | 2 | 0 | 0 | 1 | 1 | # | 10 | Small | ancom | 69 |
| # | time | 2 | 0 | 0 | 1 | 1 | # | 10 | Small | ancom | 69 |
| # | interaction | 0 | 0 | 2 | 0 | 0 | # | 10 | Small | ancom | 69 |
| # | disease | 4 | 0 | 0 | 1 | 1 | # | 10 | Small | aldex2 | 69 |
| # | sex | 2 | 0 | 0 | 1 | 1 | # | 10 | Small | aldex2 | 69 |
| # | time | 2 | 0 | 0 | 1 | 1 | # | 10 | Small | aldex2 | 69 |
| # | interaction | 0 | 0 | 2 | 0 | 0 | # | 10 | Small | aldex2 | 69 |
| # | disease | 4 | 5 | 0 | 0.444444444 | 1 | # | 10 | Baseline | paralle | 69 |
| # | sex | 2 | 4 | 0 | 0.333333333 | 1 | # | 10 | Baseline | paralle | 69 |
| # | time | 2 | 7 | 0 | 0.222222222 | 1 | # | 10 | Baseline | paralle | 69 |
| # | interaction | 2 | 7 | 0 | 0.222222222 | 1 | # | 10 | Baseline | paralle | 69 |
| # | disease | 4 | 1 | 0 | 0.8 | 1 | # | 10 | Baseline | ancom | 69 |
| # | sex | 2 | 0 | 0 | 1 | 1 | # | 10 | Baseline | ancom | 69 |
| # | time | 2 | 2 | 0 | 0.5 | 1 | # | 10 | Baseline | ancom | 69 |
| # | interaction | 0 | 0 | 2 | 0 | 0 | # | 10 | Baseline | ancom | 69 |
| # | disease | 4 | 15 | 0 | 0.210526316 | 1 | # | 10 | Baseline | aldex2 | 69 |
| # | sex | 2 | 11 | 0 | 0.153846154 | 1 | # | 10 | Baseline | aldex2 | 69 |
| # | time | 2 | 9 | 0 | 0.181818182 | 1 | # | 10 | Baseline | aldex2 | 69 |
| # | interaction | 0 | 0 | 2 | 0 | 0 | # | 10 | Baseline | aldex2 | 69 |
| # | disease | 4 | 0 | 0 | 1 | 1 | # | 10 | Small | paralle | 69 |
| # | sex | 2 | 0 | 0 | 1 | 1 | # | 10 | Small | paralle | 69 |
| # | time | 2 | 0 | 0 | 1 | 1 | # | 10 | Small | paralle | 69 |
| # | interaction | 1 | 0 | 1 | 1 | 0.5 | # | 10 | Small | paralle | 69 |
| # | disease | 4 | 1 | 0 | 0.8 | 1 | # | 10 | Small | ancom | 69 |
| # | sex | 2 | 0 | 0 | 1 | 1 | # | 10 | Small | ancom | 69 |
| # | time | 2 | 2 | 0 | 0.5 | 1 | # | 10 | Small | ancom | 69 |
| # | interaction | 0 | 0 | 2 | 0 | 0 | # | 10 | Small | ancom | 69 |
| # | disease | 4 | 3 | 0 | 0.571428571 | 1 | # | 10 | Small | aldex2 | 69 |
| # | sex | 2 | 1 | 0 | 0.666666667 | 1 | # | 10 | Small | aldex2 | 69 |
| # | time | 2 | 2 | 0 | 0.5 | 1 | # | 10 | Small | aldex2 | 69 |
| # | interaction | 0 | 0 | 2 | 0 | 0 | # | 10 | Small | aldex2 | 69 |
| # | disease | 4 | 8 | 0 | 0.333333333 | 1 | # | 10 | Baseline | paralle | 69 |
| # | sex | 2 | 5 | 0 | 0.285714286 | 1 | # | 10 | Baseline | paralle | 69 |
| # | time | 2 | 5 | 0 | 0.285714286 | 1 | # | 10 | Baseline | paralle | 69 |
| # | interaction | 2 | 6 | 0 | 0.25 | 1 | # | 10 | Baseline | paralle | 69 |
| # | disease | 4 | 2 | 0 | 0.666666667 | 1 | # | 10 | Baseline | ancom | 69 |
| # | sex | 2 | 0 | 0 | 1 | 1 | # | 10 | Baseline | ancom | 69 |
| # | time | 2 | 2 | 0 | 0.5 | 1 | # | 10 | Baseline | ancom | 69 |
| # | interaction | 0 | 0 | 2 | 0 | 0 | # | 10 | Baseline | ancom | 69 |
| # | disease | 4 | 25 | 0 | 0.137931034 | 1 | # | 10 | Baseline | aldex2 | 69 |
| # | sex | 2 | 10 | 0 | 0.166666667 | 1 | # | 10 | Baseline | aldex2 | 69 |
| # | time | 2 | 14 | 0 | 0.125 | 1 | # | 10 | Baseline | aldex2 | 69 |
| # | interaction | 0 | 0 | 2 | 0 | 0 | # | 10 | Baseline | aldex2 | 69 |
| # | disease | 4 | 0 | 0 | 1 | 1 | # | 10 | Small | paralle | 69 |
| # | sex | 2 | 0 | 0 | 1 | 1 | # | 10 | Small | paralle | 69 |
| # | time | 2 | 0 | 0 | 1 | 1 | # | 10 | Small | paralle | 69 |
| # | interaction | 2 | 0 | 0 | 1 | 1 | # | 10 | Small | paralle | 69 |
| # | disease | 4 | 2 | 0 | 0.666666667 | 1 | # | 10 | Small | ancom | 69 |
| # | sex | 2 | 0 | 0 | 1 | 1 | # | 10 | Small | ancom | 69 |
| # | time | 2 | 2 | 0 | 0.5 | 1 | # | 10 | Small | ancom | 69 |
| # | interaction | 0 | 0 | 2 | 0 | 0 | # | 10 | Small | ancom | 69 |
| # | disease | 4 | 4 | 0 | 0.5 | 1 | # | 10 | Small | aldex2 | 69 |
| # | sex | 2 | 0 | 0 | 1 | 1 | # | 10 | Small | aldex2 | 69 |
| # | time | 2 | 2 | 0 | 0.5 | 1 | # | 10 | Small | aldex2 | 69 |
| # | interaction | 0 | 0 | 2 | 0 | 0 | # | 10 | Small | aldex2 | 69 |
| # | disease | 4 | 7 | 0 | 0.363636364 | 1 | # | 10 | Baseline | paralle | 69 |
| # | sex | 2 | 7 | 0 | 0.222222222 | 1 | # | 10 | Baseline | paralle | 69 |
| # | time | 2 | 2 | 0 | 0.5 | 1 | # | 10 | Baseline | paralle | 69 |
| # | interaction | 2 | 4 | 0 | 0.333333333 | 1 | # | 10 | Baseline | paralle | 69 |
| # | disease | 4 | 2 | 0 | 0.666666667 | 1 | # | 10 | Baseline | ancom | 69 |
| # | sex | 2 | 0 | 0 | 1 | 1 | # | 10 | Baseline | ancom | 69 |
| # | time | 2 | 2 | 0 | 0.5 | 1 | # | 10 | Baseline | ancom | 69 |
| # | interaction | 0 | 0 | 2 | 0 | 0 | # | 10 | Baseline | ancom | 69 |
| # | disease | 4 | 30 | 0 | 0.117647059 | 1 | # | 10 | Baseline | aldex2 | 69 |
| # | sex | 2 | 15 | 0 | 0.117647059 | 1 | # | 10 | Baseline | aldex2 | 69 |
| # | time | 2 | 15 | 0 | 0.117647059 | 1 | # | 10 | Baseline | aldex2 | 69 |
| # | interaction | 0 | 0 | 2 | 0 | 0 | # | 10 | Baseline | aldex2 | 69 |

| | | | | | | | | | | | |
|---|---|---|---|---|---|---|---|---|---|---|---|
| # | disease | 4 | 0 | 0 | 1 | 1 | # | 10 | Small | paralle | 69 |
| # | sex | 2 | 0 | 0 | 1 | 1 | # | 10 | Small | paralle | 69 |
| # | time | 2 | 0 | 0 | 1 | 1 | # | 10 | Small | paralle | 69 |
| # | interaction | 2 | 0 | 0 | 1 | 1 | # | 10 | Small | paralle | 69 |
| # | disease | 4 | 2 | 0 | 0.666666667 | 1 | # | 10 | Small | ancom | 69 |
| # | sex | 2 | 0 | 0 | 1 | 1 | # | 10 | Small | ancom | 69 |
| # | time | 2 | 2 | 0 | 0.5 | 1 | # | 10 | Small | ancom | 69 |
| # | interaction | 0 | 0 | 2 | 0 | 0 | # | 10 | Small | ancom | 69 |
| # | disease | 4 | 3 | 0 | 0.571428571 | 1 | # | 10 | Small | aldex2 | 69 |
| # | sex | 2 | 0 | 0 | 1 | 1 | # | 10 | Small | aldex2 | 69 |
| # | time | 2 | 2 | 0 | 0.5 | 1 | # | 10 | Small | aldex2 | 69 |
| # | interaction | 0 | 0 | 2 | 0 | 0 | # | 10 | Small | aldex2 | 69 |
| # | disease | 7 | 6 | 0 | 0.538461538 | 1 | # | 20 | Baseline | paralle | 69 |
| # | sex | 4 | 15 | 0 | 0.210526316 | 1 | # | 20 | Baseline | paralle | 69 |
| # | time | 5 | 10 | 0 | 0.333333333 | 1 | # | 20 | Baseline | paralle | 69 |
| # | interaction | 3 | 9 | 1 | 0.25 | 0.75 | # | 20 | Baseline | paralle | 69 |
| # | disease | 7 | 2 | 0 | 0.777777778 | 1 | # | 20 | Baseline | ancom | 69 |
| # | sex | 4 | 0 | 0 | 1 | 1 | # | 20 | Baseline | ancom | 69 |
| # | time | 5 | 1 | 0 | 0.833333333 | 1 | # | 20 | Baseline | ancom | 69 |
| # | interaction | 0 | 0 | 4 | 0 | 0 | # | 20 | Baseline | ancom | 69 |
| # | disease | 7 | 17 | 0 | 0.291666667 | 1 | # | 20 | Baseline | aldex2 | 69 |
| # | sex | 4 | 6 | 0 | 0.4 | 1 | # | 20 | Baseline | aldex2 | 69 |
| # | time | 5 | 11 | 0 | 0.3125 | 1 | # | 20 | Baseline | aldex2 | 69 |
| # | interaction | 0 | 0 | 4 | 0 | 0 | # | 20 | Baseline | aldex2 | 69 |
| # | disease | 7 | 0 | 0 | 1 | 1 | # | 20 | Small | paralle | 69 |
| # | sex | 4 | 0 | 0 | 1 | 1 | # | 20 | Small | paralle | 69 |
| # | time | 5 | 0 | 0 | 1 | 1 | # | 20 | Small | paralle | 69 |
| # | interaction | 1 | 0 | 3 | 1 | 0.25 | # | 20 | Small | paralle | 69 |
| # | disease | 7 | 2 | 0 | 0.777777778 | 1 | # | 20 | Small | ancom | 69 |
| # | sex | 4 | 0 | 0 | 1 | 1 | # | 20 | Small | ancom | 69 |
| # | time | 5 | 0 | 0 | 1 | 1 | # | 20 | Small | ancom | 69 |
| # | interaction | 0 | 0 | 4 | 0 | 0 | # | 20 | Small | ancom | 69 |
| # | disease | 7 | 1 | 0 | 0.875 | 1 | # | 20 | Small | aldex2 | 69 |
| # | sex | 4 | 0 | 0 | 1 | 1 | # | 20 | Small | aldex2 | 69 |
| # | time | 5 | 1 | 0 | 0.833333333 | 1 | # | 20 | Small | aldex2 | 69 |
| # | interaction | 0 | 0 | 4 | 0 | 0 | # | 20 | Small | aldex2 | 69 |
| # | disease | 7 | 8 | 0 | 0.466666667 | 1 | # | 20 | Baseline | paralle | 69 |
| # | sex | 4 | 8 | 0 | 0.333333333 | 1 | # | 20 | Baseline | paralle | 69 |
| # | time | 5 | 6 | 0 | 0.454545455 | 1 | # | 20 | Baseline | paralle | 69 |
| # | interaction | 4 | 4 | 0 | 0.5 | 1 | # | 20 | Baseline | paralle | 69 |
| # | disease | 7 | 3 | 0 | 0.7 | 1 | # | 20 | Baseline | ancom | 69 |
| # | sex | 4 | 0 | 0 | 1 | 1 | # | 20 | Baseline | ancom | 69 |
| # | time | 5 | 3 | 0 | 0.625 | 1 | # | 20 | Baseline | ancom | 69 |
| # | interaction | 0 | 0 | 4 | 0 | 0 | # | 20 | Baseline | ancom | 69 |
| # | disease | 7 | 35 | 0 | 0.166666667 | 1 | # | 20 | Baseline | aldex2 | 69 |
| # | sex | 4 | 11 | 0 | 0.266666667 | 1 | # | 20 | Baseline | aldex2 | 69 |
| # | time | 5 | 19 | 0 | 0.208333333 | 1 | # | 20 | Baseline | aldex2 | 69 |
| # | interaction | 0 | 0 | 4 | 0 | 0 | # | 20 | Baseline | aldex2 | 69 |
| # | disease | 7 | 0 | 0 | 1 | 1 | # | 20 | Small | paralle | 69 |
| # | sex | 4 | 0 | 0 | 1 | 1 | # | 20 | Small | paralle | 69 |
| # | time | 5 | 0 | 0 | 1 | 1 | # | 20 | Small | paralle | 69 |
| # | interaction | 2 | 0 | 2 | 1 | 0.5 | # | 20 | Small | paralle | 69 |
| # | disease | 7 | 3 | 0 | 0.7 | 1 | # | 20 | Small | ancom | 69 |
| # | sex | 4 | 0 | 0 | 1 | 1 | # | 20 | Small | ancom | 69 |
| # | time | 5 | 3 | 0 | 0.625 | 1 | # | 20 | Small | ancom | 69 |
| # | interaction | 0 | 0 | 4 | 0 | 0 | # | 20 | Small | ancom | 69 |
| # | disease | 7 | 2 | 0 | 0.777777778 | 1 | # | 20 | Small | aldex2 | 69 |
| # | sex | 4 | 1 | 0 | 0.8 | 1 | # | 20 | Small | aldex2 | 69 |
| # | time | 5 | 2 | 0 | 0.714285714 | 1 | # | 20 | Small | aldex2 | 69 |
| # | interaction | 0 | 0 | 4 | 0 | 0 | # | 20 | Small | aldex2 | 69 |
| # | disease | 7 | 9 | 0 | 0.4375 | 1 | # | 20 | Baseline | paralle | 69 |
| # | sex | 4 | 6 | 0 | 0.4 | 1 | # | 20 | Baseline | paralle | 69 |
| # | time | 5 | 6 | 0 | 0.454545455 | 1 | # | 20 | Baseline | paralle | 69 |
| # | interaction | 4 | 1 | 0 | 0.8 | 1 | # | 20 | Baseline | paralle | 69 |
| # | disease | 7 | 3 | 0 | 0.7 | 1 | # | 20 | Baseline | ancom | 69 |
| # | sex | 4 | 0 | 0 | 1 | 1 | # | 20 | Baseline | ancom | 69 |
| # | time | 5 | 3 | 0 | 0.625 | 1 | # | 20 | Baseline | ancom | 69 |
| # | interaction | 0 | 0 | 4 | 0 | 0 | # | 20 | Baseline | ancom | 69 |
| # | disease | 7 | 51 | 0 | 0.120689655 | 1 | # | 20 | Baseline | aldex2 | 69 |
| # | sex | 4 | 18 | 0 | 0.181818182 | 1 | # | 20 | Baseline | aldex2 | 69 |
| # | time | 5 | 28 | 0 | 0.151515152 | 1 | # | 20 | Baseline | aldex2 | 69 |
| # | interaction | 0 | 0 | 4 | 0 | 0 | # | 20 | Baseline | aldex2 | 69 |
| # | disease | 7 | 0 | 0 | 1 | 1 | # | 20 | Small | paralle | 69 |
| # | sex | 4 | 0 | 0 | 1 | 1 | # | 20 | Small | paralle | 69 |
| # | time | 5 | 0 | 0 | 1 | 1 | # | 20 | Small | paralle | 69 |
| # | interaction | 3 | 0 | 1 | 1 | 0.75 | # | 20 | Small | paralle | 69 |
| # | disease | 7 | 3 | 0 | 0.7 | 1 | # | 20 | Small | ancom | 69 |
| # | sex | 4 | 0 | 0 | 1 | 1 | # | 20 | Small | ancom | 69 |
| # | time | 5 | 3 | 0 | 0.625 | 1 | # | 20 | Small | ancom | 69 |
| # | interaction | 0 | 0 | 4 | 0 | 0 | # | 20 | Small | ancom | 69 |
| # | disease | 7 | 8 | 0 | 0.466666667 | 1 | # | 20 | Small | aldex2 | 69 |
| # | sex | 4 | 0 | 0 | 1 | 1 | # | 20 | Small | aldex2 | 69 |
| # | time | 5 | 3 | 0 | 0.625 | 1 | # | 20 | Small | aldex2 | 69 |
| # | interaction | 0 | 0 | 4 | 0 | 0 | # | 20 | Small | aldex2 | 69 |
| # | disease | 7 | 10 | 0 | 0.411764706 | 1 | # | 20 | Baseline | paralle | 69 |
| # | sex | 4 | 3 | 0 | 0.571428571 | 1 | # | 20 | Baseline | paralle | 69 |
| # | time | 5 | 8 | 0 | 0.384615385 | 1 | # | 20 | Baseline | paralle | 69 |
| # | interaction | 4 | 5 | 0 | 0.444444444 | 1 | # | 20 | Baseline | paralle | 69 |
| # | disease | 7 | 4 | 0 | 0.636363636 | 1 | # | 20 | Baseline | ancom | 69 |
| # | sex | 4 | 0 | 0 | 1 | 1 | # | 20 | Baseline | ancom | 69 |
| # | time | 5 | 3 | 0 | 0.625 | 1 | # | 20 | Baseline | ancom | 69 |
| # | interaction | 0 | 0 | 4 | 0 | 0 | # | 20 | Baseline | ancom | 69 |
| # | disease | 7 | 62 | 0 | 0.101449275 | 1 | # | 20 | Baseline | aldex2 | 69 |
| # | sex | 4 | 18 | 0 | 0.181818182 | 1 | # | 20 | Baseline | aldex2 | 69 |
| # | time | 5 | 39 | 0 | 0.113636364 | 1 | # | 20 | Baseline | aldex2 | 69 |
| # | interaction | 0 | 0 | 4 | 0 | 0 | # | 20 | Baseline | aldex2 | 69 |
| # | disease | 7 | 0 | 0 | 1 | 1 | # | 20 | Small | paralle | 69 |
| # | sex | 4 | 0 | 0 | 1 | 1 | # | 20 | Small | paralle | 69 |
| # | time | 5 | 0 | 0 | 1 | 1 | # | 20 | Small | paralle | 69 |
| # | interaction | 4 | 0 | 0 | 1 | 1 | # | 20 | Small | paralle | 69 |
| # | disease | 7 | 3 | 0 | 0.7 | 1 | # | 20 | Small | ancom | 69 |
| # | sex | 4 | 0 | 0 | 1 | 1 | # | 20 | Small | ancom | 69 |
| # | time | 5 | 3 | 0 | 0.625 | 1 | # | 20 | Small | ancom | 69 |
| # | interaction | 0 | 0 | 4 | 0 | 0 | # | 20 | Small | ancom | 69 |
| # | disease | 7 | 11 | 0 | 0.388888889 | 1 | # | 20 | Small | aldex2 | 69 |
| # | sex | 4 | 0 | 0 | 1 | 1 | # | 20 | Small | aldex2 | 69 |
| # | time | 5 | 4 | 0 | 0.555555556 | 1 | # | 20 | Small | aldex2 | 69 |
| # | interaction | 0 | 0 | 4 | 0 | 0 | # | 20 | Small | aldex2 | 69 |
| # | disease | 4 | 10 | 0 | 0.285714286 | 1 | # | 10 | Baseline | paralle | 70 |
| # | sex | 2 | 12 | 0 | 0.142857143 | 1 | # | 10 | Baseline | paralle | 70 |
| # | time | 2 | 9 | 0 | 0.181818182 | 1 | # | 10 | Baseline | paralle | 70 |
| # | interaction | 2 | 6 | 0 | 0.25 | 1 | # | 10 | Baseline | paralle | 70 |
| # | disease | 4 | 1 | 0 | 0.8 | 1 | # | 10 | Baseline | ancom | 70 |
| # | sex | 2 | 0 | 0 | 1 | 1 | # | 10 | Baseline | ancom | 70 |
| # | time | 2 | 0 | 0 | 1 | 1 | # | 10 | Baseline | ancom | 70 |
| # | interaction | 0 | 0 | 2 | 0 | 0 | # | 10 | Baseline | ancom | 70 |
| # | disease | 4 | 10 | 0 | 0.285714286 | 1 | # | 10 | Baseline | aldex2 | 70 |
| # | sex | 2 | 9 | 0 | 0.181818182 | 1 | # | 10 | Baseline | aldex2 | 70 |
| # | time | 2 | 6 | 0 | 0.25 | 1 | # | 10 | Baseline | aldex2 | 70 |
| # | interaction | 0 | 0 | 2 | 0 | 0 | # | 10 | Baseline | aldex2 | 70 |
| # | disease | 4 | 1 | 0 | 0.8 | 1 | # | 10 | Small | paralle | 70 |
| # | sex | 2 | 0 | 0 | 1 | 1 | # | 10 | Small | paralle | 70 |
| # | time | 2 | 0 | 0 | 1 | 1 | # | 10 | Small | paralle | 70 |
| # | interaction | 1 | 0 | 1 | 1 | 0.5 | # | 10 | Small | paralle | 70 |
| # | disease | 4 | 0 | 0 | 1 | 1 | # | 10 | Small | ancom | 70 |
| # | sex | 2 | 0 | 0 | 1 | 1 | # | 10 | Small | ancom | 70 |
| # | time | 2 | 0 | 0 | 1 | 1 | # | 10 | Small | ancom | 70 |
| # | interaction | 0 | 0 | 2 | 0 | 0 | # | 10 | Small | ancom | 70 |
| # | disease | 4 | 1 | 0 | 0.8 | 1 | # | 10 | Small | aldex2 | 70 |
| # | sex | 2 | 0 | 0 | 1 | 1 | # | 10 | Small | aldex2 | 70 |
| # | time | 2 | 0 | 0 | 1 | 1 | # | 10 | Small | aldex2 | 70 |
| # | interaction | 0 | 0 | 2 | 0 | 0 | # | 10 | Small | aldex2 | 70 |
| # | disease | 4 | 8 | 0 | 0.333333333 | 1 | # | 10 | Baseline | paralle | 70 |
| # | sex | 2 | 4 | 0 | 0.333333333 | 1 | # | 10 | Baseline | paralle | 70 |
| # | time | 2 | 7 | 0 | 0.222222222 | 1 | # | 10 | Baseline | paralle | 70 |
| # | interaction | 2 | 2 | 0 | 0.5 | 1 | # | 10 | Baseline | paralle | 70 |
| # | disease | 4 | 1 | 0 | 0.8 | 1 | # | 10 | Baseline | ancom | 70 |
| # | sex | 2 | 0 | 0 | 1 | 1 | # | 10 | Baseline | ancom | 70 |
| # | time | 2 | 2 | 0 | 0.5 | 1 | # | 10 | Baseline | ancom | 70 |
| # | interaction | 0 | 0 | 2 | 0 | 0 | # | 10 | Baseline | ancom | 70 |
| # | disease | 4 | 11 | 0 | 0.266666667 | 1 | # | 10 | Baseline | aldex2 | 70 |
| # | sex | 2 | 4 | 0 | 0.333333333 | 1 | # | 10 | Baseline | aldex2 | 70 |
| # | time | 2 | 6 | 0 | 0.25 | 1 | # | 10 | Baseline | aldex2 | 70 |
| # | interaction | 0 | 0 | 2 | 0 | 0 | # | 10 | Baseline | aldex2 | 70 |
| # | disease | 4 | 0 | 0 | 1 | 1 | # | 10 | Small | paralle | 70 |
| # | sex | 2 | 0 | 0 | 1 | 1 | # | 10 | Small | paralle | 70 |
| # | time | 2 | 0 | 0 | 1 | 1 | # | 10 | Small | paralle | 70 |
| # | interaction | 2 | 0 | 0 | 1 | 1 | # | 10 | Small | paralle | 70 |
| # | disease | 4 | 1 | 0 | 0.8 | 1 | # | 10 | Small | ancom | 70 |
| # | sex | 2 | 0 | 0 | 1 | 1 | # | 10 | Small | ancom | 70 |
| # | time | 2 | 2 | 0 | 0.5 | 1 | # | 10 | Small | ancom | 70 |
| # | interaction | 0 | 0 | 2 | 0 | 0 | # | 10 | Small | ancom | 70 |
| # | disease | 4 | 1 | 0 | 0.8 | 1 | # | 10 | Small | aldex2 | 70 |
| # | sex | 2 | 0 | 0 | 1 | 1 | # | 10 | Small | aldex2 | 70 |
| # | time | 2 | 2 | 0 | 0.5 | 1 | # | 10 | Small | aldex2 | 70 |
| # | interaction | 0 | 0 | 2 | 0 | 0 | # | 10 | Small | aldex2 | 70 |
| # | disease | 4 | 5 | 0 | 0.444444444 | 1 | # | 10 | Baseline | paralle | 70 |
| # | sex | 2 | 3 | 0 | 0.4 | 1 | # | 10 | Baseline | paralle | 70 |
| # | time | 2 | 1 | 0 | 0.666666667 | 1 | # | 10 | Baseline | paralle | 70 |
| # | interaction | 2 | 2 | 0 | 0.5 | 1 | # | 10 | Baseline | paralle | 70 |
| # | disease | 4 | 2 | 0 | 0.666666667 | 1 | # | 10 | Baseline | ancom | 70 |
| # | sex | 2 | 0 | 0 | 1 | 1 | # | 10 | Baseline | ancom | 70 |
| # | time | 2 | 2 | 0 | 0.5 | 1 | # | 10 | Baseline | ancom | 70 |
| # | interaction | 0 | 0 | 2 | 0 | 0 | # | 10 | Baseline | ancom | 70 |
| # | disease | 4 | 15 | 0 | 0.210526316 | 1 | # | 10 | Baseline | aldex2 | 70 |
| # | sex | 2 | 6 | 0 | 0.25 | 1 | # | 10 | Baseline | aldex2 | 70 |
| # | time | 2 | 3 | 0 | 0.4 | 1 | # | 10 | Baseline | aldex2 | 70 |
| # | interaction | 0 | 0 | 2 | 0 | 0 | # | 10 | Baseline | aldex2 | 70 |
| # | disease | 4 | 0 | 0 | 1 | 1 | # | 10 | Small | paralle | 70 |
| # | sex | 2 | 0 | 0 | 1 | 1 | # | 10 | Small | paralle | 70 |
| # | time | 2 | 0 | 0 | 1 | 1 | # | 10 | Small | paralle | 70 |
| # | interaction | 2 | 0 | 0 | 1 | 1 | # | 10 | Small | paralle | 70 |
| # | disease | 4 | 2 | 0 | 0.666666667 | 1 | # | 10 | Small | ancom | 70 |
| # | sex | 2 | 0 | 0 | 1 | 1 | # | 10 | Small | ancom | 70 |
| # | time | 2 | 2 | 0 | 0.5 | 1 | # | 10 | Small | ancom | 70 |
| # | interaction | 0 | 0 | 2 | 0 | 0 | # | 10 | Small | ancom | 70 |
| # | disease | 4 | 2 | 0 | 0.666666667 | 1 | # | 10 | Small | aldex2 | 70 |
| # | sex | 2 | 0 | 0 | 1 | 1 | # | 10 | Small | aldex2 | 70 |
| # | time | 2 | 2 | 0 | 0.5 | 1 | # | 10 | Small | aldex2 | 70 |
| # | interaction | 0 | 0 | 2 | 0 | 0 | # | 10 | Small | aldex2 | 70 |
| # | disease | 4 | 8 | 0 | 0.333333333 | 1 | # | 10 | Baseline | paralle | 70 |
| # | sex | 2 | 3 | 0 | 0.4 | 1 | # | 10 | Baseline | paralle | 70 |
| # | time | 2 | 3 | 0 | 0.4 | 1 | # | 10 | Baseline | paralle | 70 |
| # | interaction | 2 | 3 | 0 | 0.4 | 1 | # | 10 | Baseline | paralle | 70 |
| # | disease | 4 | 2 | 0 | 0.666666667 | 1 | # | 10 | Baseline | ancom | 70 |
| # | sex | 2 | 1 | 0 | 0.666666667 | 1 | # | 10 | Baseline | ancom | 70 |
| # | time | 2 | 2 | 0 | 0.5 | 1 | # | 10 | Baseline | ancom | 70 |
| # | interaction | 0 | 0 | 2 | 0 | 0 | # | 10 | Baseline | ancom | 70 |
| # | disease | 4 | 27 | 0 | 0.129032258 | 1 | # | 10 | Baseline | aldex2 | 70 |
| # | sex | 2 | 8 | 0 | 0.2 | 1 | # | 10 | Baseline | aldex2 | 70 |
| # | time | 2 | 4 | 0 | 0.333333333 | 1 | # | 10 | Baseline | aldex2 | 70 |
| # | interaction | 0 | 0 | 2 | 0 | 0 | # | 10 | Baseline | aldex2 | 70 |
| # | disease | 4 | 0 | 0 | 1 | 1 | # | 10 | Small | paralle | 70 |
| # | sex | 2 | 0 | 0 | 1 | 1 | # | 10 | Small | paralle | 70 |
| # | time | 2 | 0 | 0 | 1 | 1 | # | 10 | Small | paralle | 70 |
| # | interaction | 2 | 0 | 0 | 1 | 1 | # | 10 | Small | paralle | 70 |
| # | disease | 4 | 2 | 0 | 0.666666667 | 1 | # | 10 | Small | ancom | 70 |
| # | sex | 2 | 1 | 0 | 0.666666667 | 1 | # | 10 | Small | ancom | 70 |
| # | time | 2 | 2 | 0 | 0.5 | 1 | # | 10 | Small | ancom | 70 |
| # | interaction | 0 | 0 | 2 | 0 | 0 | # | 10 | Small | ancom | 70 |
| # | disease | 4 | 2 | 0 | 0.666666667 | 1 | # | 10 | Small | aldex2 | 70 |
| # | sex | 2 | 1 | 0 | 0.666666667 | 1 | # | 10 | Small | aldex2 | 70 |
| # | time | 2 | 2 | 0 | 0.5 | 1 | # | 10 | Small | aldex2 | 70 |
| # | interaction | 0 | 0 | 2 | 0 | 0 | # | 10 | Small | aldex2 | 70 |
| # | disease | 7 | 13 | 0 | 0.35 | 1 | # | 20 | Baseline | paralle | 70 |
| # | sex | 4 | 8 | 0 | 0.333333333 | 1 | # | 20 | Baseline | paralle | 70 |
| # | time | 5 | 11 | 0 | 0.3125 | 1 | # | 20 | Baseline | paralle | 70 |
| # | interaction | 4 | 4 | 0 | 0.5 | 1 | # | 20 | Baseline | paralle | 70 |
| # | disease | 7 | 3 | 0 | 0.7 | 1 | # | 20 | Baseline | ancom | 70 |
| # | sex | 4 | 0 | 0 | 1 | 1 | # | 20 | Baseline | ancom | 70 |
| # | time | 5 | 3 | 0 | 0.625 | 1 | # | 20 | Baseline | ancom | 70 |
| # | interaction | 0 | 0 | 4 | 0 | 0 | # | 20 | Baseline | ancom | 70 |
| # | disease | 7 | 27 | 0 | 0.205882353 | 1 | # | 20 | Baseline | aldex2 | 70 |
| # | sex | 4 | 13 | 0 | 0.235294118 | 1 | # | 20 | Baseline | aldex2 | 70 |
| # | time | 5 | 8 | 0 | 0.384615385 | 1 | # | 20 | Baseline | aldex2 | 70 |
| # | interaction | 0 | 0 | 4 | 0 | 0 | # | 20 | Baseline | aldex2 | 70 |
| # | disease | 7 | 1 | 0 | 0.875 | 1 | # | 20 | Small | paralle | 70 |
| # | sex | 4 | 0 | 0 | 1 | 1 | # | 20 | Small | paralle | 70 |
| # | time | 5 | 0 | 0 | 1 | 1 | # | 20 | Small | paralle | 70 |
| # | interaction | 4 | 0 | 0 | 1 | 1 | # | 20 | Small | paralle | 70 |
| # | disease | 7 | 1 | 0 | 0.875 | 1 | # | 20 | Small | ancom | 70 |
| # | sex | 4 | 0 | 0 | 1 | 1 | # | 20 | Small | ancom | 70 |
| # | time | 5 | 2 | 0 | 0.714285714 | 1 | # | 20 | Small | ancom | 70 |
| # | interaction | 0 | 0 | 4 | 0 | 0 | # | 20 | Small | ancom | 70 |
| # | disease | 7 | 4 | 0 | 0.636363636 | 1 | # | 20 | Small | aldex2 | 70 |
| # | sex | 4 | 0 | 0 | 1 | 1 | # | 20 | Small | aldex2 | 70 |
| # | time | 5 | 2 | 0 | 0.714285714 | 1 | # | 20 | Small | aldex2 | 70 |
| # | interaction | 0 | 0 | 4 | 0 | 0 | # | 20 | Small | aldex2 | 70 |
| # | disease | 7 | 6 | 0 | 0.538461538 | 1 | # | 20 | Baseline | paralle | 70 |
| # | sex | 4 | 7 | 0 | 0.363636364 | 1 | # | 20 | Baseline | paralle | 70 |
| # | time | 5 | 8 | 0 | 0.384615385 | 1 | # | 20 | Baseline | paralle | 70 |
| # | interaction | 4 | 6 | 0 | 0.4 | 1 | # | 20 | Baseline | paralle | 70 |
| # | disease | 7 | 3 | 0 | 0.7 | 1 | # | 20 | Baseline | ancom | 70 |
| # | sex | 4 | 0 | 0 | 1 | 1 | # | 20 | Baseline | ancom | 70 |
| # | time | 5 | 4 | 0 | 0.555555556 | 1 | # | 20 | Baseline | ancom | 70 |
| # | interaction | 0 | 0 | 4 | 0 | 0 | # | 20 | Baseline | ancom | 70 |
| # | disease | 7 | 42 | 0 | 0.142857143 | 1 | # | 20 | Baseline | aldex2 | 70 |
| # | sex | 4 | 11 | 0 | 0.266666667 | 1 | # | 20 | Baseline | aldex2 | 70 |
| # | time | 5 | 23 | 0 | 0.178571429 | 1 | # | 20 | Baseline | aldex2 | 70 |
| # | interaction | 0 | 0 | 4 | 0 | 0 | # | 20 | Baseline | aldex2 | 70 |
| # | disease | 7 | 0 | 0 | 1 | 1 | # | 20 | Small | paralle | 70 |
| # | sex | 4 | 0 | 0 | 1 | 1 | # | 20 | Small | paralle | 70 |
| # | time | 5 | 0 | 0 | 1 | 1 | # | 20 | Small | paralle | 70 |
| # | interaction | 4 | 0 | 0 | 1 | 1 | # | 20 | Small | paralle | 70 |
| # | disease | 7 | 3 | 0 | 0.7 | 1 | # | 20 | Small | ancom | 70 |
| # | sex | 4 | 0 | 0 | 1 | 1 | # | 20 | Small | ancom | 70 |
| # | time | 5 | 4 | 0 | 0.555555556 | 1 | # | 20 | Small | ancom | 70 |
| # | interaction | 0 | 0 | 4 | 0 | 0 | # | 20 | Small | ancom | 70 |
| # | disease | 7 | 4 | 0 | 0.636363636 | 1 | # | 20 | Small | aldex2 | 70 |
| # | sex | 4 | 0 | 0 | 1 | 1 | # | 20 | Small | aldex2 | 70 |
| # | time | 5 | 3 | 0 | 0.625 | 1 | # | 20 | Small | aldex2 | 70 |
| # | interaction | 0 | 0 | 4 | 0 | 0 | # | 20 | Small | aldex2 | 70 |
| # | disease | 7 | 10 | 0 | 0.411764706 | 1 | # | 20 | Baseline | paralle | 70 |
| # | sex | 4 | 6 | 0 | 0.4 | 1 | # | 20 | Baseline | paralle | 70 |
| # | time | 5 | 3 | 0 | 0.625 | 1 | # | 20 | Baseline | paralle | 70 |
| # | interaction | 4 | 7 | 0 | 0.363636364 | 1 | # | 20 | Baseline | paralle | 70 |
| # | disease | 7 | 4 | 0 | 0.636363636 | 1 | # | 20 | Baseline | ancom | 70 |
| # | sex | 4 | 0 | 0 | 1 | 1 | # | 20 | Baseline | ancom | 70 |
| # | time | 5 | 4 | 0 | 0.555555556 | 1 | # | 20 | Baseline | ancom | 70 |
| # | interaction | 0 | 0 | 4 | 0 | 0 | # | 20 | Baseline | ancom | 70 |
| # | disease | 7 | 61 | 0 | 0.102941176 | 1 | # | 20 | Baseline | aldex2 | 70 |
| # | sex | 4 | 21 | 0 | 0.16 | 1 | # | 20 | Baseline | aldex2 | 70 |
| # | time | 5 | 41 | 0 | 0.108695652 | 1 | # | 20 | Baseline | aldex2 | 70 |
| # | interaction | 0 | 0 | 4 | 0 | 0 | # | 20 | Baseline | aldex2 | 70 |
| # | disease | 7 | 0 | 0 | 1 | 1 | # | 20 | Small | paralle | 70 |
| # | sex | 4 | 0 | 0 | 1 | 1 | # | 20 | Small | paralle | 70 |
| # | time | 5 | 1 | 0 | 0.833333333 | 1 | # | 20 | Small | paralle | 70 |
| # | interaction | 4 | 0 | 0 | 1 | 1 | # | 20 | Small | paralle | 70 |
| # | disease | 7 | 4 | 0 | 0.636363636 | 1 | # | 20 | Small | ancom | 70 |
| # | sex | 4 | 0 | 0 | 1 | 1 | # | 20 | Small | ancom | 70 |
| # | time | 5 | 4 | 0 | 0.555555556 | 1 | # | 20 | Small | ancom | 70 |
| # | interaction | 0 | 0 | 4 | 0 | 0 | # | 20 | Small | ancom | 70 |
| # | disease | 7 | 5 | 0 | 0.583333333 | 1 | # | 20 | Small | aldex2 | 70 |
| # | sex | 4 | 0 | 0 | 1 | 1 | # | 20 | Small | aldex2 | 70 |
| # | time | 5 | 5 | 0 | 0.5 | 1 | # | 20 | Small | aldex2 | 70 |
| # | interaction | 0 | 0 | 4 | 0 | 0 | # | 20 | Small | aldex2 | 70 |
| # | disease | 7 | 6 | 0 | 0.538461538 | 1 | # | 20 | Baseline | paralle | 70 |
| # | sex | 4 | 2 | 0 | 0.666666667 | 1 | # | 20 | Baseline | paralle | 70 |
| # | time | 5 | 5 | 0 | 0.5 | 1 | # | 20 | Baseline | paralle | 70 |
| # | interaction | 4 | 5 | 0 | 0.444444444 | 1 | # | 20 | Baseline | paralle | 70 |
| # | disease | 7 | 4 | 0 | 0.636363636 | 1 | # | 20 | Baseline | ancom | 70 |
| # | sex | 4 | 0 | 0 | 1 | 1 | # | 20 | Baseline | ancom | 70 |
| # | time | 5 | 4 | 0 | 0.555555556 | 1 | # | 20 | Baseline | ancom | 70 |
| # | interaction | 0 | 0 | 4 | 0 | 0 | # | 20 | Baseline | ancom | 70 |
| # | disease | 7 | 69 | 0 | 0.092105263 | 1 | # | 20 | Baseline | aldex2 | 70 |
| # | sex | 4 | 24 | 0 | 0.142857143 | 1 | # | 20 | Baseline | aldex2 | 70 |
| # | time | 5 | 40 | 0 | 0.111111111 | 1 | # | 20 | Baseline | aldex2 | 70 |
| # | interaction | 0 | 0 | 4 | 0 | 0 | # | 20 | Baseline | aldex2 | 70 |
| # | disease | 7 | 0 | 0 | 1 | 1 | # | 20 | Small | paralle | 70 |
| # | sex | 4 | 0 | 0 | 1 | 1 | # | 20 | Small | paralle | 70 |
| # | time | 5 | 0 | 0 | 1 | 1 | # | 20 | Small | paralle | 70 |
| # | interaction | 4 | 0 | 0 | 1 | 1 | # | 20 | Small | paralle | 70 |
| # | disease | 7 | 4 | 0 | 0.636363636 | 1 | # | 20 | Small | ancom | 70 |
| # | sex | 4 | 0 | 0 | 1 | 1 | # | 20 | Small | ancom | 70 |
| # | time | 5 | 4 | 0 | 0.555555556 | 1 | # | 20 | Small | ancom | 70 |
| # | interaction | 0 | 0 | 4 | 0 | 0 | # | 20 | Small | ancom | 70 |
| # | disease | 7 | 11 | 0 | 0.388888889 | 1 | # | 20 | Small | aldex2 | 70 |
| # | sex | 4 | 0 | 0 | 1 | 1 | # | 20 | Small | aldex2 | 70 |
| # | time | 5 | 6 | 0 | 0.454545455 | 1 | # | 20 | Small | aldex2 | 70 |
| # | interaction | 0 | 0 | 4 | 0 | 0 | # | 20 | Small | aldex2 | 70 |
| # | disease | 4 | 7 | 0 | 0.363636364 | 1 | # | 10 | Baseline | paralle | 71 |
| # | sex | 2 | 7 | 0 | 0.222222222 | 1 | # | 10 | Baseline | paralle | 71 |
| # | time | 2 | 6 | 0 | 0.25 | 1 | # | 10 | Baseline | paralle | 71 |
| # | interaction | 2 | 7 | 0 | 0.222222222 | 1 | # | 10 | Baseline | paralle | 71 |
| # | disease | 4 | 1 | 0 | 0.8 | 1 | # | 10 | Baseline | ancom | 71 |
| # | sex | 2 | 0 | 0 | 1 | 1 | # | 10 | Baseline | ancom | 71 |
| # | time | 2 | 1 | 0 | 0.666666667 | 1 | # | 10 | Baseline | ancom | 71 |
| # | interaction | 0 | 0 | 2 | 0 | 0 | # | 10 | Baseline | ancom | 71 |
| # | disease | 4 | 8 | 0 | 0.333333333 | 1 | # | 10 | Baseline | aldex2 | 71 |
| # | sex | 2 | 5 | 0 | 0.285714286 | 1 | # | 10 | Baseline | aldex2 | 71 |
| # | time | 2 | 6 | 0 | 0.25 | 1 | # | 10 | Baseline | aldex2 | 71 |
| # | interaction | 0 | 0 | 2 | 0 | 0 | # | 10 | Baseline | aldex2 | 71 |
| # | disease | 4 | 1 | 0 | 0.8 | 1 | # | 10 | Small | paralle | 71 |
| # | sex | 2 | 0 | 0 | 1 | 1 | # | 10 | Small | paralle | 71 |
| # | time | 2 | 0 | 0 | 1 | 1 | # | 10 | Small | paralle | 71 |
| # | interaction | 1 | 0 | 1 | 1 | 0.5 | # | 10 | Small | paralle | 71 |
| # | disease | 4 | 1 | 0 | 0.8 | 1 | # | 10 | Small | ancom | 71 |
| # | sex | 2 | 0 | 0 | 1 | 1 | # | 10 | Small | ancom | 71 |
| # | time | 2 | 1 | 0 | 0.666666667 | 1 | # | 10 | Small | ancom | 71 |
| # | interaction | 0 | 0 | 2 | 0 | 0 | # | 10 | Small | ancom | 71 |
| # | disease | 4 | 1 | 0 | 0.8 | 1 | # | 10 | Small | aldex2 | 71 |
| # | sex | 2 | 0 | 0 | 1 | 1 | # | 10 | Small | aldex2 | 71 |
| # | time | 2 | 2 | 0 | 0.5 | 1 | # | 10 | Small | aldex2 | 71 |
| # | interaction | 0 | 0 | 2 | 0 | 0 | # | 10 | Small | aldex2 | 71 |
| # | disease | 4 | 8 | 0 | 0.333333333 | 1 | # | 10 | Baseline | paralle | 71 |
| # | sex | 2 | 3 | 0 | 0.4 | 1 | # | 10 | Baseline | paralle | 71 |
| # | time | 2 | 6 | 0 | 0.25 | 1 | # | 10 | Baseline | paralle | 71 |
| # | interaction | 2 | 7 | 0 | 0.222222222 | 1 | # | 10 | Baseline | paralle | 71 |
| # | disease | 4 | 2 | 0 | 0.666666667 | 1 | # | 10 | Baseline | ancom | 71 |
| # | sex | 2 | 0 | 0 | 1 | 1 | # | 10 | Baseline | ancom | 71 |
| # | time | 2 | 2 | 0 | 0.5 | 1 | # | 10 | Baseline | ancom | 71 |
| # | interaction | 0 | 0 | 2 | 0 | 0 | # | 10 | Baseline | ancom | 71 |
| # | disease | 4 | 14 | 0 | 0.222222222 | 1 | # | 10 | Baseline | aldex2 | 71 |
| # | sex | 2 | 6 | 0 | 0.25 | 1 | # | 10 | Baseline | aldex2 | 71 |
| # | time | 2 | 6 | 0 | 0.25 | 1 | # | 10 | Baseline | aldex2 | 71 |
| # | interaction | 0 | 0 | 2 | 0 | 0 | # | 10 | Baseline | aldex2 | 71 |
| # | disease | 4 | 0 | 0 | 1 | 1 | # | 10 | Small | paralle | 71 |

| | | | | | | | | | | | |
|---|---|---|---|---|---|---|---|---|---|---|---|
| # | sex | 2 | 0 | 0 | 1 | 1 | # | 10 | Small | paralle | 71 |
| # | time | 2 | 0 | 0 | 1 | 1 | # | 10 | Small | paralle | 71 |
| # | interaction | 1 | 0 | 1 | 1 | 0.5 | # | 10 | Small | paralle | 71 |
| # | disease | 4 | 3 | 0 | 0.571428571 | 1 | # | 10 | Small | ancom | 71 |
| # | sex | 2 | 0 | 0 | 1 | 1 | # | 10 | Small | ancom | 71 |
| # | time | 2 | 2 | 0 | 0.5 | 1 | # | 10 | Small | ancom | 71 |
| # | interaction | 0 | 0 | 2 | 0 | 0 | # | 10 | Small | ancom | 71 |
| # | disease | 4 | 3 | 0 | 0.571428571 | 1 | # | 10 | Small | aldex2 | 71 |
| # | sex | 2 | 0 | 0 | 1 | 1 | # | 10 | Small | aldex2 | 71 |
| # | time | 2 | 1 | 0 | 0.666666667 | 1 | # | 10 | Small | aldex2 | 71 |
| # | interaction | 0 | 0 | 2 | 0 | 0 | # | 10 | Small | aldex2 | 71 |
| # | disease | 4 | 10 | 0 | 0.285714286 | 1 | # | 10 | Baseline | paralle | 71 |
| # | sex | 2 | 6 | 0 | 0.25 | 1 | # | 10 | Baseline | paralle | 71 |
| # | time | 2 | 5 | 0 | 0.285714286 | 1 | # | 10 | Baseline | paralle | 71 |
| # | interaction | 2 | 5 | 0 | 0.285714286 | 1 | # | 10 | Baseline | paralle | 71 |
| # | disease | 4 | 2 | 0 | 0.666666667 | 1 | # | 10 | Baseline | ancom | 71 |
| # | sex | 2 | 0 | 0 | 1 | 1 | # | 10 | Baseline | ancom | 71 |
| # | time | 2 | 2 | 0 | 0.5 | 1 | # | 10 | Baseline | ancom | 71 |
| # | interaction | 0 | 0 | 2 | 0 | 0 | # | 10 | Baseline | ancom | 71 |
| # | disease | 4 | 20 | 0 | 0.166666667 | 1 | # | 10 | Baseline | aldex2 | 71 |
| # | sex | 2 | 6 | 0 | 0.25 | 1 | # | 10 | Baseline | aldex2 | 71 |
| # | time | 2 | 12 | 0 | 0.142857143 | 1 | # | 10 | Baseline | aldex2 | 71 |
| # | interaction | 0 | 0 | 2 | 0 | 0 | # | 10 | Baseline | aldex2 | 71 |
| # | disease | 4 | 0 | 0 | 1 | 1 | # | 10 | Small | paralle | 71 |
| # | sex | 2 | 0 | 0 | 1 | 1 | # | 10 | Small | paralle | 71 |
| # | time | 2 | 0 | 0 | 1 | 1 | # | 10 | Small | paralle | 71 |
| # | interaction | 2 | 0 | 0 | 1 | 1 | # | 10 | Small | paralle | 71 |
| # | disease | 4 | 2 | 0 | 0.666666667 | 1 | # | 10 | Small | ancom | 71 |
| # | sex | 2 | 0 | 0 | 1 | 1 | # | 10 | Small | ancom | 71 |
| # | time | 2 | 2 | 0 | 0.5 | 1 | # | 10 | Small | ancom | 71 |
| # | interaction | 0 | 0 | 2 | 0 | 0 | # | 10 | Small | ancom | 71 |
| # | disease | 4 | 4 | 0 | 0.5 | 1 | # | 10 | Small | aldex2 | 71 |
| # | sex | 2 | 0 | 0 | 1 | 1 | # | 10 | Small | aldex2 | 71 |
| # | time | 2 | 3 | 0 | 0.4 | 1 | # | 10 | Small | aldex2 | 71 |
| # | interaction | 0 | 0 | 2 | 0 | 0 | # | 10 | Small | aldex2 | 71 |
| # | disease | 4 | 5 | 0 | 0.444444444 | 1 | # | 10 | Baseline | paralle | 71 |
| # | sex | 2 | 2 | 0 | 0.5 | 1 | # | 10 | Baseline | paralle | 71 |
| # | time | 2 | 5 | 0 | 0.285714286 | 1 | # | 10 | Baseline | paralle | 71 |
| # | interaction | 2 | 3 | 0 | 0.4 | 1 | # | 10 | Baseline | paralle | 71 |
| # | disease | 4 | 2 | 0 | 0.666666667 | 1 | # | 10 | Baseline | ancom | 71 |
| # | sex | 2 | 0 | 0 | 1 | 1 | # | 10 | Baseline | ancom | 71 |
| # | time | 2 | 2 | 0 | 0.5 | 1 | # | 10 | Baseline | ancom | 71 |
| # | interaction | 0 | 0 | 2 | 0 | 0 | # | 10 | Baseline | ancom | 71 |
| # | disease | 4 | 26 | 0 | 0.133333333 | 1 | # | 10 | Baseline | aldex2 | 71 |
| # | sex | 2 | 8 | 0 | 0.2 | 1 | # | 10 | Baseline | aldex2 | 71 |
| # | time | 2 | 13 | 0 | 0.133333333 | 1 | # | 10 | Baseline | aldex2 | 71 |
| # | interaction | 0 | 0 | 2 | 0 | 0 | # | 10 | Baseline | aldex2 | 71 |
| # | disease | 4 | 0 | 0 | 1 | 1 | # | 10 | Small | paralle | 71 |
| # | sex | 2 | 0 | 0 | 1 | 1 | # | 10 | Small | paralle | 71 |
| # | time | 2 | 0 | 0 | 1 | 1 | # | 10 | Small | paralle | 71 |
| # | interaction | 2 | 0 | 0 | 1 | 1 | # | 10 | Small | paralle | 71 |
| # | disease | 4 | 2 | 0 | 0.666666667 | 1 | # | 10 | Small | ancom | 71 |
| # | sex | 2 | 0 | 0 | 1 | 1 | # | 10 | Small | ancom | 71 |
| # | time | 2 | 2 | 0 | 0.5 | 1 | # | 10 | Small | ancom | 71 |
| # | interaction | 0 | 0 | 2 | 0 | 0 | # | 10 | Small | ancom | 71 |
| # | disease | 4 | 5 | 0 | 0.444444444 | 1 | # | 10 | Small | aldex2 | 71 |
| # | sex | 2 | 0 | 0 | 1 | 1 | # | 10 | Small | aldex2 | 71 |
| # | time | 2 | 3 | 0 | 0.4 | 1 | # | 10 | Small | aldex2 | 71 |
| # | interaction | 0 | 0 | 2 | 0 | 0 | # | 10 | Small | aldex2 | 71 |
| # | disease | 7 | 9 | 0 | 0.4375 | 1 | # | 20 | Baseline | paralle | 71 |
| # | sex | 4 | 5 | 0 | 0.444444444 | 1 | # | 20 | Baseline | paralle | 71 |
| # | time | 5 | 4 | 0 | 0.555555556 | 1 | # | 20 | Baseline | paralle | 71 |
| # | interaction | 4 | 7 | 0 | 0.363636364 | 1 | # | 20 | Baseline | paralle | 71 |
| # | disease | 7 | 2 | 0 | 0.777777778 | 1 | # | 20 | Baseline | ancom | 71 |
| # | sex | 4 | 0 | 0 | 1 | 1 | # | 20 | Baseline | ancom | 71 |
| # | time | 5 | 0 | 0 | 1 | 1 | # | 20 | Baseline | ancom | 71 |
| # | interaction | 0 | 0 | 4 | 0 | 0 | # | 20 | Baseline | ancom | 71 |
| # | disease | 7 | 24 | 0 | 0.225806452 | 1 | # | 20 | Baseline | aldex2 | 71 |
| # | sex | 4 | 1 | 0 | 0.8 | 1 | # | 20 | Baseline | aldex2 | 71 |
| # | time | 5 | 10 | 0 | 0.333333333 | 1 | # | 20 | Baseline | aldex2 | 71 |
| # | interaction | 0 | 0 | 4 | 0 | 0 | # | 20 | Baseline | aldex2 | 71 |
| # | disease | 7 | 0 | 0 | 1 | 1 | # | 20 | Small | paralle | 71 |
| # | sex | 4 | 0 | 0 | 1 | 1 | # | 20 | Small | paralle | 71 |
| # | time | 5 | 0 | 0 | 1 | 1 | # | 20 | Small | paralle | 71 |
| # | interaction | 0 | 0 | 4 | 0 | 0 | # | 20 | Small | paralle | 71 |
| # | disease | 7 | 2 | 0 | 0.777777778 | 1 | # | 20 | Small | ancom | 71 |
| # | sex | 4 | 0 | 0 | 1 | 1 | # | 20 | Small | ancom | 71 |
| # | time | 5 | 0 | 0 | 1 | 1 | # | 20 | Small | ancom | 71 |
| # | interaction | 0 | 0 | 4 | 0 | 0 | # | 20 | Small | ancom | 71 |
| # | disease | 7 | 4 | 0 | 0.636363636 | 1 | # | 20 | Small | aldex2 | 71 |
| # | sex | 4 | 0 | 0 | 1 | 1 | # | 20 | Small | aldex2 | 71 |
| # | time | 5 | 1 | 0 | 0.833333333 | 1 | # | 20 | Small | aldex2 | 71 |
| # | interaction | 0 | 0 | 4 | 0 | 0 | # | 20 | Small | aldex2 | 71 |
| # | disease | 7 | 8 | 0 | 0.466666667 | 1 | # | 20 | Baseline | paralle | 71 |
| # | sex | 4 | 6 | 0 | 0.4 | 1 | # | 20 | Baseline | paralle | 71 |
| # | time | 5 | 4 | 0 | 0.555555556 | 1 | # | 20 | Baseline | paralle | 71 |
| # | interaction | 4 | 6 | 0 | 0.4 | 1 | # | 20 | Baseline | paralle | 71 |
| # | disease | 7 | 3 | 0 | 0.7 | 1 | # | 20 | Baseline | ancom | 71 |
| # | sex | 4 | 0 | 0 | 1 | 1 | # | 20 | Baseline | ancom | 71 |
| # | time | 5 | 2 | 0 | 0.714285714 | 1 | # | 20 | Baseline | ancom | 71 |
| # | interaction | 0 | 0 | 4 | 0 | 0 | # | 20 | Baseline | ancom | 71 |
| # | disease | 7 | 37 | 0 | 0.159090909 | 1 | # | 20 | Baseline | aldex2 | 71 |
| # | sex | 4 | 6 | 0 | 0.4 | 1 | # | 20 | Baseline | aldex2 | 71 |
| # | time | 5 | 16 | 0 | 0.238095238 | 1 | # | 20 | Baseline | aldex2 | 71 |
| # | interaction | 0 | 0 | 4 | 0 | 0 | # | 20 | Baseline | aldex2 | 71 |
| # | disease | 7 | 0 | 0 | 1 | 1 | # | 20 | Small | paralle | 71 |
| # | sex | 4 | 0 | 0 | 1 | 1 | # | 20 | Small | paralle | 71 |
| # | time | 5 | 0 | 0 | 1 | 1 | # | 20 | Small | paralle | 71 |
| # | interaction | 4 | 0 | 0 | 1 | 1 | # | 20 | Small | paralle | 71 |
| # | disease | 7 | 3 | 0 | 0.7 | 1 | # | 20 | Small | ancom | 71 |
| # | sex | 4 | 0 | 0 | 1 | 1 | # | 20 | Small | ancom | 71 |
| # | time | 5 | 2 | 0 | 0.714285714 | 1 | # | 20 | Small | ancom | 71 |
| # | interaction | 0 | 0 | 4 | 0 | 0 | # | 20 | Small | ancom | 71 |
| # | disease | 7 | 6 | 0 | 0.538461538 | 1 | # | 20 | Small | aldex2 | 71 |
| # | sex | 4 | 0 | 0 | 1 | 1 | # | 20 | Small | aldex2 | 71 |
| # | time | 5 | 2 | 0 | 0.714285714 | 1 | # | 20 | Small | aldex2 | 71 |
| # | interaction | 0 | 0 | 4 | 0 | 0 | # | 20 | Small | aldex2 | 71 |
| # | disease | 7 | 17 | 0 | 0.291666667 | 1 | # | 20 | Baseline | paralle | 71 |
| # | sex | 4 | 7 | 0 | 0.363636364 | 1 | # | 20 | Baseline | paralle | 71 |
| # | time | 5 | 10 | 0 | 0.333333333 | 1 | # | 20 | Baseline | paralle | 71 |
| # | interaction | 4 | 5 | 0 | 0.444444444 | 1 | # | 20 | Baseline | paralle | 71 |
| # | disease | 7 | 4 | 0 | 0.636363636 | 1 | # | 20 | Baseline | ancom | 71 |
| # | sex | 4 | 0 | 0 | 1 | 1 | # | 20 | Baseline | ancom | 71 |
| # | time | 5 | 3 | 0 | 0.625 | 1 | # | 20 | Baseline | ancom | 71 |
| # | interaction | 0 | 0 | 4 | 0 | 0 | # | 20 | Baseline | ancom | 71 |
| # | disease | 7 | 51 | 0 | 0.120689655 | 1 | # | 20 | Baseline | aldex2 | 71 |
| # | sex | 4 | 12 | 0 | 0.25 | 1 | # | 20 | Baseline | aldex2 | 71 |
| # | time | 5 | 24 | 0 | 0.172413793 | 1 | # | 20 | Baseline | aldex2 | 71 |
| # | interaction | 0 | 0 | 4 | 0 | 0 | # | 20 | Baseline | aldex2 | 71 |
| # | disease | 7 | 0 | 0 | 1 | 1 | # | 20 | Small | paralle | 71 |
| # | sex | 4 | 0 | 0 | 1 | 1 | # | 20 | Small | paralle | 71 |
| # | time | 5 | 0 | 0 | 1 | 1 | # | 20 | Small | paralle | 71 |
| # | interaction | 3 | 0 | 1 | 1 | 0.75 | # | 20 | Small | paralle | 71 |
| # | disease | 7 | 4 | 0 | 0.636363636 | 1 | # | 20 | Small | ancom | 71 |
| # | sex | 4 | 0 | 0 | 1 | 1 | # | 20 | Small | ancom | 71 |
| # | time | 5 | 3 | 0 | 0.625 | 1 | # | 20 | Small | ancom | 71 |
| # | interaction | 0 | 0 | 4 | 0 | 0 | # | 20 | Small | ancom | 71 |
| # | disease | 7 | 8 | 0 | 0.466666667 | 1 | # | 20 | Small | aldex2 | 71 |
| # | sex | 4 | 1 | 0 | 0.8 | 1 | # | 20 | Small | aldex2 | 71 |
| # | time | 5 | 3 | 0 | 0.625 | 1 | # | 20 | Small | aldex2 | 71 |
| # | interaction | 0 | 0 | 4 | 0 | 0 | # | 20 | Small | aldex2 | 71 |
| # | disease | 7 | 13 | 0 | 0.35 | 1 | # | 20 | Baseline | paralle | 71 |
| # | sex | 4 | 4 | 0 | 0.5 | 1 | # | 20 | Baseline | paralle | 71 |
| # | time | 5 | 11 | 0 | 0.3125 | 1 | # | 20 | Baseline | paralle | 71 |
| # | interaction | 4 | 4 | 0 | 0.5 | 1 | # | 20 | Baseline | paralle | 71 |
| # | disease | 7 | 4 | 0 | 0.636363636 | 1 | # | 20 | Baseline | ancom | 71 |
| # | sex | 4 | 0 | 0 | 1 | 1 | # | 20 | Baseline | ancom | 71 |
| # | time | 5 | 4 | 0 | 0.555555556 | 1 | # | 20 | Baseline | ancom | 71 |
| # | interaction | 0 | 0 | 4 | 0 | 0 | # | 20 | Baseline | ancom | 71 |
| # | disease | 7 | 66 | 0 | 0.095890411 | 1 | # | 20 | Baseline | aldex2 | 71 |
| # | sex | 4 | 18 | 0 | 0.181818182 | 1 | # | 20 | Baseline | aldex2 | 71 |
| # | time | 5 | 30 | 0 | 0.142857143 | 1 | # | 20 | Baseline | aldex2 | 71 |
| # | interaction | 0 | 0 | 4 | 0 | 0 | # | 20 | Baseline | aldex2 | 71 |
| # | disease | 7 | 1 | 0 | 0.875 | 1 | # | 20 | Small | paralle | 71 |
| # | sex | 4 | 0 | 0 | 1 | 1 | # | 20 | Small | paralle | 71 |
| # | time | 5 | 0 | 0 | 1 | 1 | # | 20 | Small | paralle | 71 |
| # | interaction | 4 | 0 | 0 | 1 | 1 | # | 20 | Small | paralle | 71 |
| # | disease | 7 | 4 | 0 | 0.636363636 | 1 | # | 20 | Small | ancom | 71 |
| # | sex | 4 | 0 | 0 | 1 | 1 | # | 20 | Small | ancom | 71 |
| # | time | 5 | 4 | 0 | 0.555555556 | 1 | # | 20 | Small | ancom | 71 |
| # | interaction | 0 | 0 | 4 | 0 | 0 | # | 20 | Small | ancom | 71 |
| # | disease | 7 | 8 | 0 | 0.466666667 | 1 | # | 20 | Small | aldex2 | 71 |
| # | sex | 4 | 1 | 0 | 0.8 | 1 | # | 20 | Small | aldex2 | 71 |
| # | time | 5 | 4 | 0 | 0.555555556 | 1 | # | 20 | Small | aldex2 | 71 |
| # | interaction | 0 | 0 | 4 | 0 | 0 | # | 20 | Small | aldex2 | 71 |
| # | disease | 4 | 6 | 0 | 0.4 | 1 | # | 10 | Baseline | paralle | 72 |
| # | sex | 2 | 10 | 0 | 0.166666667 | 1 | # | 10 | Baseline | paralle | 72 |
| # | time | 2 | 10 | 0 | 0.166666667 | 1 | # | 10 | Baseline | paralle | 72 |
| # | interaction | 2 | 9 | 0 | 0.181818182 | 1 | # | 10 | Baseline | paralle | 72 |
| # | disease | 4 | 0 | 0 | 1 | 1 | # | 10 | Baseline | ancom | 72 |
| # | sex | 2 | 0 | 0 | 1 | 1 | # | 10 | Baseline | ancom | 72 |
| # | time | 2 | 1 | 0 | 0.666666667 | 1 | # | 10 | Baseline | ancom | 72 |
| # | interaction | 0 | 0 | 2 | 0 | 0 | # | 10 | Baseline | ancom | 72 |
| # | disease | 4 | 8 | 0 | 0.333333333 | 1 | # | 10 | Baseline | aldex2 | 72 |
| # | sex | 2 | 9 | 0 | 0.181818182 | 1 | # | 10 | Baseline | aldex2 | 72 |
| # | time | 2 | 6 | 0 | 0.25 | 1 | # | 10 | Baseline | aldex2 | 72 |
| # | interaction | 0 | 0 | 2 | 0 | 0 | # | 10 | Baseline | aldex2 | 72 |
| # | disease | 4 | 0 | 0 | 1 | 1 | # | 10 | Small | paralle | 72 |
| # | sex | 2 | 0 | 0 | 1 | 1 | # | 10 | Small | paralle | 72 |
| # | time | 2 | 1 | 0 | 0.666666667 | 1 | # | 10 | Small | paralle | 72 |
| # | interaction | 2 | 2 | 0 | 0.5 | 1 | # | 10 | Small | paralle | 72 |
| # | disease | 4 | 0 | 0 | 1 | 1 | # | 10 | Small | ancom | 72 |
| # | sex | 2 | 0 | 0 | 1 | 1 | # | 10 | Small | ancom | 72 |
| # | time | 2 | 1 | 0 | 0.666666667 | 1 | # | 10 | Small | ancom | 72 |
| # | interaction | 0 | 0 | 2 | 0 | 0 | # | 10 | Small | ancom | 72 |
| # | disease | 4 | 0 | 0 | 1 | 1 | # | 10 | Small | aldex2 | 72 |
| # | sex | 2 | 1 | 0 | 0.666666667 | 1 | # | 10 | Small | aldex2 | 72 |
| # | time | 2 | 1 | 0 | 0.666666667 | 1 | # | 10 | Small | aldex2 | 72 |
| # | interaction | 0 | 0 | 2 | 0 | 0 | # | 10 | Small | aldex2 | 72 |
| # | disease | 4 | 10 | 0 | 0.285714286 | 1 | # | 10 | Baseline | paralle | 72 |
| # | sex | 2 | 10 | 0 | 0.166666667 | 1 | # | 10 | Baseline | paralle | 72 |
| # | time | 2 | 7 | 0 | 0.222222222 | 1 | # | 10 | Baseline | paralle | 72 |
| # | interaction | 2 | 7 | 0 | 0.222222222 | 1 | # | 10 | Baseline | paralle | 72 |
| # | disease | 4 | 0 | 0 | 1 | 1 | # | 10 | Baseline | ancom | 72 |
| # | sex | 2 | 0 | 0 | 1 | 1 | # | 10 | Baseline | ancom | 72 |
| # | time | 2 | 1 | 0 | 0.666666667 | 1 | # | 10 | Baseline | ancom | 72 |
| # | interaction | 0 | 0 | 2 | 0 | 0 | # | 10 | Baseline | ancom | 72 |
| # | disease | 4 | 14 | 0 | 0.222222222 | 1 | # | 10 | Baseline | aldex2 | 72 |
| # | sex | 2 | 15 | 0 | 0.117647059 | 1 | # | 10 | Baseline | aldex2 | 72 |
| # | time | 2 | 7 | 0 | 0.222222222 | 1 | # | 10 | Baseline | aldex2 | 72 |
| # | interaction | 0 | 0 | 2 | 0 | 0 | # | 10 | Baseline | aldex2 | 72 |
| # | disease | 4 | 0 | 0 | 1 | 1 | # | 10 | Small | paralle | 72 |
| # | sex | 2 | 0 | 0 | 1 | 1 | # | 10 | Small | paralle | 72 |
| # | time | 2 | 0 | 0 | 1 | 1 | # | 10 | Small | paralle | 72 |
| # | interaction | 1 | 0 | 1 | 1 | 0.5 | # | 10 | Small | paralle | 72 |
| # | disease | 4 | 0 | 0 | 1 | 1 | # | 10 | Small | ancom | 72 |
| # | sex | 2 | 0 | 0 | 1 | 1 | # | 10 | Small | ancom | 72 |
| # | time | 2 | 1 | 0 | 0.666666667 | 1 | # | 10 | Small | ancom | 72 |
| # | interaction | 0 | 0 | 2 | 0 | 0 | # | 10 | Small | ancom | 72 |
| # | disease | 4 | 0 | 0 | 1 | 1 | # | 10 | Small | aldex2 | 72 |
| # | sex | 2 | 2 | 0 | 0.5 | 1 | # | 10 | Small | aldex2 | 72 |
| # | time | 2 | 1 | 0 | 0.666666667 | 1 | # | 10 | Small | aldex2 | 72 |
| # | interaction | 0 | 0 | 2 | 0 | 0 | # | 10 | Small | aldex2 | 72 |
| # | disease | 4 | 4 | 0 | 0.5 | 1 | # | 10 | Baseline | paralle | 72 |
| # | sex | 2 | 8 | 0 | 0.2 | 1 | # | 10 | Baseline | paralle | 72 |
| # | time | 2 | 4 | 0 | 0.333333333 | 1 | # | 10 | Baseline | paralle | 72 |
| # | interaction | 2 | 4 | 0 | 0.333333333 | 1 | # | 10 | Baseline | paralle | 72 |
| # | disease | 4 | 0 | 0 | 1 | 1 | # | 10 | Baseline | ancom | 72 |
| # | sex | 2 | 0 | 0 | 1 | 1 | # | 10 | Baseline | ancom | 72 |
| # | time | 2 | 2 | 0 | 0.5 | 1 | # | 10 | Baseline | ancom | 72 |
| # | interaction | 0 | 0 | 2 | 0 | 0 | # | 10 | Baseline | ancom | 72 |
| # | disease | 4 | 18 | 0 | 0.181818182 | 1 | # | 10 | Baseline | aldex2 | 72 |
| # | sex | 2 | 7 | 0 | 0.222222222 | 1 | # | 10 | Baseline | aldex2 | 72 |
| # | time | 2 | 9 | 0 | 0.181818182 | 1 | # | 10 | Baseline | aldex2 | 72 |
| # | interaction | 0 | 0 | 2 | 0 | 0 | # | 10 | Baseline | aldex2 | 72 |
| # | disease | 4 | 0 | 0 | 1 | 1 | # | 10 | Small | paralle | 72 |
| # | sex | 2 | 0 | 0 | 1 | 1 | # | 10 | Small | paralle | 72 |
| # | time | 2 | 0 | 0 | 1 | 1 | # | 10 | Small | paralle | 72 |
| # | interaction | 2 | 0 | 0 | 1 | 1 | # | 10 | Small | paralle | 72 |
| # | disease | 4 | 0 | 0 | 1 | 1 | # | 10 | Small | ancom | 72 |
| # | sex | 2 | 0 | 0 | 1 | 1 | # | 10 | Small | ancom | 72 |
| # | time | 2 | 1 | 0 | 0.666666667 | 1 | # | 10 | Small | ancom | 72 |
| # | interaction | 0 | 0 | 2 | 0 | 0 | # | 10 | Small | ancom | 72 |
| # | disease | 4 | 1 | 0 | 0.8 | 1 | # | 10 | Small | aldex2 | 72 |
| # | sex | 2 | 1 | 0 | 0.666666667 | 1 | # | 10 | Small | aldex2 | 72 |
| # | time | 2 | 2 | 0 | 0.5 | 1 | # | 10 | Small | aldex2 | 72 |
| # | interaction | 0 | 0 | 2 | 0 | 0 | # | 10 | Small | aldex2 | 72 |
| # | disease | 4 | 6 | 0 | 0.4 | 1 | # | 10 | Baseline | paralle | 72 |
| # | sex | 2 | 4 | 0 | 0.333333333 | 1 | # | 10 | Baseline | paralle | 72 |
| # | time | 2 | 3 | 0 | 0.4 | 1 | # | 10 | Baseline | paralle | 72 |
| # | interaction | 2 | 3 | 0 | 0.4 | 1 | # | 10 | Baseline | paralle | 72 |
| # | disease | 4 | 2 | 0 | 0.666666667 | 1 | # | 10 | Baseline | ancom | 72 |
| # | sex | 2 | 0 | 0 | 1 | 1 | # | 10 | Baseline | ancom | 72 |
| # | time | 2 | 2 | 0 | 0.5 | 1 | # | 10 | Baseline | ancom | 72 |
| # | interaction | 0 | 0 | 2 | 0 | 0 | # | 10 | Baseline | ancom | 72 |
| # | disease | 4 | 23 | 0 | 0.148148148 | 1 | # | 10 | Baseline | aldex2 | 72 |
| # | sex | 2 | 5 | 0 | 0.285714286 | 1 | # | 10 | Baseline | aldex2 | 72 |
| # | time | 2 | 10 | 0 | 0.166666667 | 1 | # | 10 | Baseline | aldex2 | 72 |
| # | interaction | 0 | 0 | 2 | 0 | 0 | # | 10 | Baseline | aldex2 | 72 |
| # | disease | 4 | 0 | 0 | 1 | 1 | # | 10 | Small | paralle | 72 |
| # | sex | 2 | 0 | 0 | 1 | 1 | # | 10 | Small | paralle | 72 |
| # | time | 2 | 0 | 0 | 1 | 1 | # | 10 | Small | paralle | 72 |
| # | interaction | 2 | 0 | 0 | 1 | 1 | # | 10 | Small | paralle | 72 |
| # | disease | 4 | 1 | 0 | 0.8 | 1 | # | 10 | Small | ancom | 72 |
| # | sex | 2 | 0 | 0 | 1 | 1 | # | 10 | Small | ancom | 72 |
| # | time | 2 | 2 | 0 | 0.5 | 1 | # | 10 | Small | ancom | 72 |
| # | interaction | 0 | 0 | 2 | 0 | 0 | # | 10 | Small | ancom | 72 |
| # | disease | 4 | 5 | 0 | 0.444444444 | 1 | # | 10 | Small | aldex2 | 72 |
| # | sex | 2 | 0 | 0 | 1 | 1 | # | 10 | Small | aldex2 | 72 |
| # | time | 2 | 2 | 0 | 0.5 | 1 | # | 10 | Small | aldex2 | 72 |
| # | interaction | 0 | 0 | 2 | 0 | 0 | # | 10 | Small | aldex2 | 72 |
| # | disease | 7 | 9 | 0 | 0.4375 | 1 | # | 20 | Baseline | paralle | 72 |
| # | sex | 4 | 10 | 0 | 0.285714286 | 1 | # | 20 | Baseline | paralle | 72 |
| # | time | 5 | 9 | 0 | 0.357142857 | 1 | # | 20 | Baseline | paralle | 72 |
| # | interaction | 3 | 6 | 1 | 0.333333333 | 0.75 | # | 20 | Baseline | paralle | 72 |
| # | disease | 7 | 2 | 0 | 0.777777778 | 1 | # | 20 | Baseline | ancom | 72 |
| # | sex | 4 | 0 | 0 | 1 | 1 | # | 20 | Baseline | ancom | 72 |
| # | time | 5 | 1 | 0 | 0.833333333 | 1 | # | 20 | Baseline | ancom | 72 |
| # | interaction | 0 | 0 | 4 | 0 | 0 | # | 20 | Baseline | ancom | 72 |
| # | disease | 7 | 21 | 0 | 0.25 | 1 | # | 20 | Baseline | aldex2 | 72 |
| # | sex | 4 | 10 | 0 | 0.285714286 | 1 | # | 20 | Baseline | aldex2 | 72 |
| # | time | 5 | 11 | 0 | 0.3125 | 1 | # | 20 | Baseline | aldex2 | 72 |
| # | interaction | 0 | 0 | 4 | 0 | 0 | # | 20 | Baseline | aldex2 | 72 |
| # | disease | 7 | 0 | 0 | 1 | 1 | # | 20 | Small | paralle | 72 |
| # | sex | 4 | 0 | 0 | 1 | 1 | # | 20 | Small | paralle | 72 |
| # | time | 5 | 0 | 0 | 1 | 1 | # | 20 | Small | paralle | 72 |
| # | interaction | 0 | 0 | 4 | 0 | 0 | # | 20 | Small | paralle | 72 |
| # | disease | 7 | 2 | 0 | 0.777777778 | 1 | # | 20 | Small | ancom | 72 |
| # | sex | 4 | 0 | 0 | 1 | 1 | # | 20 | Small | ancom | 72 |
| # | time | 5 | 1 | 0 | 0.833333333 | 1 | # | 20 | Small | ancom | 72 |
| # | interaction | 0 | 0 | 4 | 0 | 0 | # | 20 | Small | ancom | 72 |
| # | disease | 7 | 1 | 0 | 0.875 | 1 | # | 20 | Small | aldex2 | 72 |
| # | sex | 4 | 0 | 0 | 1 | 1 | # | 20 | Small | aldex2 | 72 |
| # | time | 5 | 1 | 0 | 0.833333333 | 1 | # | 20 | Small | aldex2 | 72 |
| # | interaction | 0 | 0 | 4 | 0 | 0 | # | 20 | Small | aldex2 | 72 |
| # | disease | 7 | 8 | 0 | 0.466666667 | 1 | # | 20 | Baseline | paralle | 72 |
| # | sex | 4 | 3 | 0 | 0.571428571 | 1 | # | 20 | Baseline | paralle | 72 |
| # | time | 5 | 10 | 0 | 0.333333333 | 1 | # | 20 | Baseline | paralle | 72 |
| # | interaction | 4 | 6 | 0 | 0.4 | 1 | # | 20 | Baseline | paralle | 72 |
| # | disease | 7 | 3 | 0 | 0.7 | 1 | # | 20 | Baseline | ancom | 72 |
| # | sex | 4 | 0 | 0 | 1 | 1 | # | 20 | Baseline | ancom | 72 |
| # | time | 5 | 1 | 0 | 0.833333333 | 1 | # | 20 | Baseline | ancom | 72 |
| # | interaction | 0 | 0 | 4 | 0 | 0 | # | 20 | Baseline | ancom | 72 |
| # | disease | 7 | 35 | 0 | 0.166666667 | 1 | # | 20 | Baseline | aldex2 | 72 |
| # | sex | 4 | 10 | 0 | 0.285714286 | 1 | # | 20 | Baseline | aldex2 | 72 |
| # | time | 5 | 24 | 0 | 0.172413793 | 1 | # | 20 | Baseline | aldex2 | 72 |
| # | interaction | 0 | 0 | 4 | 0 | 0 | # | 20 | Baseline | aldex2 | 72 |
| # | disease | 7 | 0 | 0 | 1 | 1 | # | 20 | Small | paralle | 72 |
| # | sex | 4 | 0 | 0 | 1 | 1 | # | 20 | Small | paralle | 72 |
| # | time | 5 | 0 | 0 | 1 | 1 | # | 20 | Small | paralle | 72 |
| # | interaction | 2 | 0 | 2 | 1 | 0.5 | # | 20 | Small | paralle | 72 |
| # | disease | 7 | 2 | 0 | 0.777777778 | 1 | # | 20 | Small | ancom | 72 |
| # | sex | 4 | 0 | 0 | 1 | 1 | # | 20 | Small | ancom | 72 |
| # | time | 5 | 1 | 0 | 0.833333333 | 1 | # | 20 | Small | ancom | 72 |
| # | interaction | 0 | 0 | 4 | 0 | 0 | # | 20 | Small | ancom | 72 |
| # | disease | 7 | 2 | 0 | 0.777777778 | 1 | # | 20 | Small | aldex2 | 72 |
| # | sex | 4 | 1 | 0 | 0.8 | 1 | # | 20 | Small | aldex2 | 72 |
| # | time | 5 | 1 | 0 | 0.833333333 | 1 | # | 20 | Small | aldex2 | 72 |
| # | interaction | 0 | 0 | 4 | 0 | 0 | # | 20 | Small | aldex2 | 72 |
| # | disease | 7 | 10 | 0 | 0.411764706 | 1 | # | 20 | Baseline | paralle | 72 |
| # | sex | 4 | 3 | 0 | 0.571428571 | 1 | # | 20 | Baseline | paralle | 72 |
| # | time | 5 | 9 | 0 | 0.357142857 | 1 | # | 20 | Baseline | paralle | 72 |
| # | interaction | 4 | 4 | 0 | 0.5 | 1 | # | 20 | Baseline | paralle | 72 |
| # | disease | 7 | 3 | 0 | 0.7 | 1 | # | 20 | Baseline | ancom | 72 |
| # | sex | 4 | 0 | 0 | 1 | 1 | # | 20 | Baseline | ancom | 72 |
| # | time | 5 | 4 | 0 | 0.555555556 | 1 | # | 20 | Baseline | ancom | 72 |
| # | interaction | 0 | 0 | 4 | 0 | 0 | # | 20 | Baseline | ancom | 72 |
| # | disease | 7 | 56 | 0 | 0.111111111 | 1 | # | 20 | Baseline | aldex2 | 72 |
| # | sex | 4 | 16 | 0 | 0.2 | 1 | # | 20 | Baseline | aldex2 | 72 |
| # | time | 5 | 34 | 0 | 0.128205128 | 1 | # | 20 | Baseline | aldex2 | 72 |
| # | interaction | 0 | 0 | 4 | 0 | 0 | # | 20 | Baseline | aldex2 | 72 |
| # | disease | 7 | 0 | 0 | 1 | 1 | # | 20 | Small | paralle | 72 |
| # | sex | 4 | 0 | 0 | 1 | 1 | # | 20 | Small | paralle | 72 |
| # | time | 5 | 0 | 0 | 1 | 1 | # | 20 | Small | paralle | 72 |
| # | interaction | 2 | 0 | 2 | 1 | 0.5 | # | 20 | Small | paralle | 72 |
| # | disease | 7 | 3 | 0 | 0.7 | 1 | # | 20 | Small | ancom | 72 |
| # | sex | 4 | 0 | 0 | 1 | 1 | # | 20 | Small | ancom | 72 |
| # | time | 5 | 3 | 0 | 0.625 | 1 | # | 20 | Small | ancom | 72 |
| # | interaction | 0 | 0 | 4 | 0 | 0 | # | 20 | Small | ancom | 72 |
| # | disease | 7 | 5 | 0 | 0.583333333 | 1 | # | 20 | Small | aldex2 | 72 |
| # | sex | 4 | 0 | 0 | 1 | 1 | # | 20 | Small | aldex2 | 72 |
| # | time | 5 | 5 | 0 | 0.5 | 1 | # | 20 | Small | aldex2 | 72 |
| # | interaction | 0 | 0 | 4 | 0 | 0 | # | 20 | Small | aldex2 | 72 |
| # | disease | 7 | 8 | 0 | 0.466666667 | 1 | # | 20 | Baseline | paralle | 72 |
| # | sex | 4 | 5 | 0 | 0.444444444 | 1 | # | 20 | Baseline | paralle | 72 |
| # | time | 5 | 10 | 0 | 0.333333333 | 1 | # | 20 | Baseline | paralle | 72 |
| # | interaction | 4 | 2 | 0 | 0.666666667 | 1 | # | 20 | Baseline | paralle | 72 |
| # | disease | 7 | 4 | 0 | 0.636363636 | 1 | # | 20 | Baseline | ancom | 72 |
| # | sex | 4 | 0 | 0 | 1 | 1 | # | 20 | Baseline | ancom | 72 |
| # | time | 5 | 4 | 0 | 0.555555556 | 1 | # | 20 | Baseline | ancom | 72 |
| # | interaction | 0 | 0 | 4 | 0 | 0 | # | 20 | Baseline | ancom | 72 |
| # | disease | 7 | 58 | 0 | 0.093333333 | 1 | # | 20 | Baseline | aldex2 | 72 |
| # | sex | 4 | 15 | 0 | 0.210526316 | 1 | # | 20 | Baseline | aldex2 | 72 |
| # | time | 5 | 62 | 0 | 0.106382979 | 1 | # | 20 | Baseline | aldex2 | 72 |
| # | interaction | 0 | 0 | 4 | 0 | 0 | # | 20 | Baseline | aldex2 | 72 |
| # | disease | 7 | 0 | 0 | 1 | 1 | # | 20 | Small | paralle | 72 |
| # | sex | 4 | 0 | 0 | 1 | 1 | # | 20 | Small | paralle | 72 |

| | | | | | | | | | | | |
|---|---|---|---|---|---|---|---|---|---|---|---|
| # | time | 5 | 0 | 0 | 1 | 1 | # | 20 | Small | paralle | 72 |
| # | interaction | 3 | 0 | 1 | 1 | 0.75 | # | 20 | Small | paralle | 72 |
| # | disease | 7 | 4 | 0 | 0.636363636 | 1 | # | 20 | Small | ancom | 72 |
| # | sex | 4 | 0 | 0 | 1 | 1 | # | 20 | Small | ancom | 72 |
| # | time | 5 | 3 | 0 | 0.625 | 1 | # | 20 | Small | ancom | 72 |
| # | interaction | 0 | 0 | 4 | 0 | 0 | # | 20 | Small | ancom | 72 |
| # | disease | 7 | 9 | 0 | 0.4375 | 1 | # | 20 | Small | aldex2 | 72 |
| # | sex | 4 | 1 | 0 | 0.8 | 1 | # | 20 | Small | aldex2 | 72 |
| # | time | 5 | 4 | 0 | 0.555555556 | 1 | # | 20 | Small | aldex2 | 72 |
| # | interaction | 0 | 0 | 4 | 0 | 0 | # | 20 | Small | aldex2 | 72 |
| # | disease | 4 | 6 | 0 | 0.4 | 1 | # | 10 | Baseline | paralle | 73 |
| # | sex | 2 | 13 | 0 | 0.133333333 | 1 | # | 10 | Baseline | paralle | 73 |
| # | time | 2 | 8 | 0 | 0.2 | 1 | # | 10 | Baseline | paralle | 73 |
| # | interaction | 2 | 7 | 0 | 0.222222222 | 1 | # | 10 | Baseline | paralle | 73 |
| # | disease | 4 | 1 | 0 | 0.8 | 1 | # | 10 | Baseline | ancom | 73 |
| # | sex | 2 | 0 | 0 | 1 | 1 | # | 10 | Baseline | ancom | 73 |
| # | time | 2 | 1 | 0 | 0.666666667 | 1 | # | 10 | Baseline | ancom | 73 |
| # | interaction | 0 | 0 | 2 | 0 | 0 | # | 10 | Baseline | ancom | 73 |
| # | disease | 4 | 11 | 0 | 0.266666667 | 1 | # | 10 | Baseline | aldex2 | 73 |
| # | sex | 2 | 6 | 0 | 0.25 | 1 | # | 10 | Baseline | aldex2 | 73 |
| # | time | 2 | 12 | 0 | 0.142857143 | 1 | # | 10 | Baseline | aldex2 | 73 |
| # | interaction | 0 | 0 | 2 | 0 | 0 | # | 10 | Baseline | aldex2 | 73 |
| # | disease | 4 | 1 | 0 | 0.8 | 1 | # | 10 | Small | paralle | 73 |
| # | sex | 2 | 0 | 0 | 1 | 1 | # | 10 | Small | paralle | 73 |
| # | time | 2 | 0 | 0 | 1 | 1 | # | 10 | Small | paralle | 73 |
| # | interaction | 0 | 0 | 2 | 0 | 0 | # | 10 | Small | paralle | 73 |
| # | disease | 4 | 1 | 0 | 0.8 | 1 | # | 10 | Small | ancom | 73 |
| # | sex | 2 | 0 | 0 | 1 | 1 | # | 10 | Small | ancom | 73 |
| # | time | 2 | 1 | 0 | 0.666666667 | 1 | # | 10 | Small | ancom | 73 |
| # | interaction | 0 | 0 | 2 | 0 | 0 | # | 10 | Small | ancom | 73 |
| # | disease | 4 | 2 | 0 | 0.666666667 | 1 | # | 10 | Small | aldex2 | 73 |
| # | sex | 2 | 0 | 0 | 1 | 1 | # | 10 | Small | aldex2 | 73 |
| # | time | 2 | 1 | 0 | 0.666666667 | 1 | # | 10 | Small | aldex2 | 73 |
| # | interaction | 0 | 0 | 2 | 0 | 0 | # | 10 | Small | aldex2 | 73 |
| # | disease | 4 | 8 | 0 | 0.333333333 | 1 | # | 10 | Baseline | paralle | 73 |
| # | sex | 2 | 7 | 0 | 0.222222222 | 1 | # | 10 | Baseline | paralle | 73 |
| # | time | 2 | 1 | 0 | 0.666666667 | 1 | # | 10 | Baseline | paralle | 73 |
| # | interaction | 2 | 6 | 0 | 0.25 | 1 | # | 10 | Baseline | paralle | 73 |
| # | disease | 4 | 2 | 0 | 0.666666667 | 1 | # | 10 | Baseline | ancom | 73 |
| # | sex | 2 | 0 | 0 | 1 | 1 | # | 10 | Baseline | ancom | 73 |
| # | time | 2 | 1 | 0 | 0.666666667 | 1 | # | 10 | Baseline | ancom | 73 |
| # | interaction | 0 | 0 | 2 | 0 | 0 | # | 10 | Baseline | ancom | 73 |
| # | disease | 4 | 20 | 0 | 0.166666667 | 1 | # | 10 | Baseline | aldex2 | 73 |
| # | sex | 2 | 8 | 0 | 0.2 | 1 | # | 10 | Baseline | aldex2 | 73 |
| # | time | 2 | 11 | 0 | 0.153846154 | 1 | # | 10 | Baseline | aldex2 | 73 |
| # | interaction | 0 | 0 | 2 | 0 | 0 | # | 10 | Baseline | aldex2 | 73 |
| # | disease | 4 | 1 | 0 | 0.8 | 1 | # | 10 | Small | paralle | 73 |
| # | sex | 2 | 0 | 0 | 1 | 1 | # | 10 | Small | paralle | 73 |
| # | time | 2 | 0 | 0 | 1 | 1 | # | 10 | Small | paralle | 73 |
| # | interaction | 1 | 0 | 1 | 1 | 0.5 | # | 10 | Small | paralle | 73 |
| # | disease | 4 | 2 | 0 | 0.666666667 | 1 | # | 10 | Small | ancom | 73 |
| # | sex | 2 | 0 | 0 | 1 | 1 | # | 10 | Small | ancom | 73 |
| # | time | 2 | 1 | 0 | 0.666666667 | 1 | # | 10 | Small | ancom | 73 |
| # | interaction | 0 | 0 | 2 | 0 | 0 | # | 10 | Small | ancom | 73 |
| # | disease | 4 | 2 | 0 | 0.666666667 | 1 | # | 10 | Small | aldex2 | 73 |
| # | sex | 2 | 0 | 0 | 1 | 1 | # | 10 | Small | aldex2 | 73 |
| # | time | 2 | 1 | 0 | 0.666666667 | 1 | # | 10 | Small | aldex2 | 73 |
| # | interaction | 0 | 0 | 2 | 0 | 0 | # | 10 | Small | aldex2 | 73 |
| # | disease | 4 | 5 | 0 | 0.444444444 | 1 | # | 10 | Baseline | paralle | 73 |
| # | sex | 2 | 6 | 0 | 0.25 | 1 | # | 10 | Baseline | paralle | 73 |
| # | time | 2 | 1 | 0 | 0.666666667 | 1 | # | 10 | Baseline | paralle | 73 |
| # | interaction | 2 | 4 | 0 | 0.333333333 | 1 | # | 10 | Baseline | paralle | 73 |
| # | disease | 4 | 2 | 0 | 0.666666667 | 1 | # | 10 | Baseline | ancom | 73 |
| # | sex | 2 | 0 | 0 | 1 | 1 | # | 10 | Baseline | ancom | 73 |
| # | time | 2 | 2 | 0 | 0.5 | 1 | # | 10 | Baseline | ancom | 73 |
| # | interaction | 0 | 0 | 2 | 0 | 0 | # | 10 | Baseline | ancom | 73 |
| # | disease | 4 | 24 | 0 | 0.142857143 | 1 | # | 10 | Baseline | aldex2 | 73 |
| # | sex | 2 | 7 | 0 | 0.222222222 | 1 | # | 10 | Baseline | aldex2 | 73 |
| # | time | 2 | 11 | 0 | 0.153846154 | 1 | # | 10 | Baseline | aldex2 | 73 |
| # | interaction | 0 | 0 | 2 | 0 | 0 | # | 10 | Baseline | aldex2 | 73 |
| # | disease | 4 | 0 | 0 | 1 | 1 | # | 10 | Small | paralle | 73 |
| # | sex | 2 | 0 | 0 | 1 | 1 | # | 10 | Small | paralle | 73 |
| # | time | 2 | 0 | 0 | 1 | 1 | # | 10 | Small | paralle | 73 |
| # | interaction | 2 | 0 | 0 | 1 | 1 | # | 10 | Small | paralle | 73 |
| # | disease | 4 | 2 | 0 | 0.666666667 | 1 | # | 10 | Small | ancom | 73 |
| # | sex | 2 | 0 | 0 | 1 | 1 | # | 10 | Small | ancom | 73 |
| # | time | 2 | 2 | 0 | 0.5 | 1 | # | 10 | Small | ancom | 73 |
| # | interaction | 0 | 0 | 2 | 0 | 0 | # | 10 | Small | ancom | 73 |
| # | disease | 4 | 3 | 0 | 0.571428571 | 1 | # | 10 | Small | aldex2 | 73 |
| # | sex | 2 | 0 | 0 | 1 | 1 | # | 10 | Small | aldex2 | 73 |
| # | time | 2 | 2 | 0 | 0.5 | 1 | # | 10 | Small | aldex2 | 73 |
| # | interaction | 0 | 0 | 2 | 0 | 0 | # | 10 | Small | aldex2 | 73 |
| # | disease | 4 | 7 | 0 | 0.363636364 | 1 | # | 10 | Baseline | paralle | 73 |
| # | sex | 2 | 5 | 0 | 0.285714286 | 1 | # | 10 | Baseline | paralle | 73 |
| # | time | 2 | 3 | 0 | 0.4 | 1 | # | 10 | Baseline | paralle | 73 |
| # | interaction | 2 | 8 | 0 | 0.2 | 1 | # | 10 | Baseline | paralle | 73 |
| # | disease | 4 | 2 | 0 | 0.666666667 | 1 | # | 10 | Baseline | ancom | 73 |
| # | sex | 2 | 0 | 0 | 1 | 1 | # | 10 | Baseline | ancom | 73 |
| # | time | 2 | 2 | 0 | 0.5 | 1 | # | 10 | Baseline | ancom | 73 |
| # | interaction | 0 | 0 | 2 | 0 | 0 | # | 10 | Baseline | ancom | 73 |
| # | disease | 4 | 33 | 0 | 0.108108108 | 1 | # | 10 | Baseline | aldex2 | 73 |
| # | sex | 2 | 7 | 0 | 0.222222222 | 1 | # | 10 | Baseline | aldex2 | 73 |
| # | time | 2 | 16 | 0 | 0.111111111 | 1 | # | 10 | Baseline | aldex2 | 73 |
| # | interaction | 0 | 0 | 2 | 0 | 0 | # | 10 | Baseline | aldex2 | 73 |
| # | disease | 4 | 0 | 0 | 1 | 1 | # | 10 | Small | paralle | 73 |
| # | sex | 2 | 0 | 0 | 1 | 1 | # | 10 | Small | paralle | 73 |
| # | time | 2 | 0 | 0 | 1 | 1 | # | 10 | Small | paralle | 73 |
| # | interaction | 2 | 1 | 0 | 0.666666667 | 1 | # | 10 | Small | paralle | 73 |
| # | disease | 4 | 2 | 0 | 0.666666667 | 1 | # | 10 | Small | ancom | 73 |
| # | sex | 2 | 0 | 0 | 1 | 1 | # | 10 | Small | ancom | 73 |
| # | time | 2 | 2 | 0 | 0.5 | 1 | # | 10 | Small | ancom | 73 |
| # | interaction | 0 | 0 | 2 | 0 | 0 | # | 10 | Small | ancom | 73 |
| # | disease | 4 | 4 | 0 | 0.5 | 1 | # | 10 | Small | aldex2 | 73 |
| # | sex | 2 | 0 | 0 | 1 | 1 | # | 10 | Small | aldex2 | 73 |
| # | time | 2 | 2 | 0 | 0.5 | 1 | # | 10 | Small | aldex2 | 73 |
| # | interaction | 0 | 0 | 2 | 0 | 0 | # | 10 | Small | aldex2 | 73 |
| # | disease | 7 | 13 | 0 | 0.35 | 1 | # | 20 | Baseline | paralle | 73 |
| # | sex | 4 | 8 | 0 | 0.333333333 | 1 | # | 20 | Baseline | paralle | 73 |
| # | time | 5 | 7 | 0 | 0.416666667 | 1 | # | 20 | Baseline | paralle | 73 |
| # | interaction | 4 | 11 | 0 | 0.266666667 | 1 | # | 20 | Baseline | paralle | 73 |
| # | disease | 7 | 1 | 0 | 0.875 | 1 | # | 20 | Baseline | ancom | 73 |
| # | sex | 4 | 0 | 0 | 1 | 1 | # | 20 | Baseline | ancom | 73 |
| # | time | 5 | 1 | 0 | 0.833333333 | 1 | # | 20 | Baseline | ancom | 73 |
| # | interaction | 0 | 0 | 4 | 0 | 0 | # | 20 | Baseline | ancom | 73 |
| # | disease | 7 | 21 | 0 | 0.25 | 1 | # | 20 | Baseline | aldex2 | 73 |
| # | sex | 4 | 7 | 0 | 0.363636364 | 1 | # | 20 | Baseline | aldex2 | 73 |
| # | time | 5 | 16 | 0 | 0.238095238 | 1 | # | 20 | Baseline | aldex2 | 73 |
| # | interaction | 0 | 0 | 4 | 0 | 0 | # | 20 | Baseline | aldex2 | 73 |
| # | disease | 7 | 0 | 0 | 1 | 1 | # | 20 | Small | paralle | 73 |
| # | sex | 4 | 1 | 0 | 0.8 | 1 | # | 20 | Small | paralle | 73 |
| # | time | 5 | 1 | 0 | 0.833333333 | 1 | # | 20 | Small | paralle | 73 |
| # | interaction | 2 | 0 | 2 | 1 | 0.5 | # | 20 | Small | paralle | 73 |
| # | disease | 7 | 1 | 0 | 0.875 | 1 | # | 20 | Small | ancom | 73 |
| # | sex | 4 | 0 | 0 | 1 | 1 | # | 20 | Small | ancom | 73 |
| # | time | 5 | 1 | 0 | 0.833333333 | 1 | # | 20 | Small | ancom | 73 |
| # | interaction | 0 | 0 | 4 | 0 | 0 | # | 20 | Small | ancom | 73 |
| # | disease | 7 | 2 | 0 | 0.777777778 | 1 | # | 20 | Small | aldex2 | 73 |
| # | sex | 4 | 0 | 0 | 1 | 1 | # | 20 | Small | aldex2 | 73 |
| # | time | 5 | 2 | 0 | 0.714285714 | 1 | # | 20 | Small | aldex2 | 73 |
| # | interaction | 0 | 0 | 4 | 0 | 0 | # | 20 | Small | aldex2 | 73 |
| # | disease | 7 | 6 | 0 | 0.538461538 | 1 | # | 20 | Baseline | paralle | 73 |
| # | sex | 4 | 7 | 0 | 0.363636364 | 1 | # | 20 | Baseline | paralle | 73 |
| # | time | 5 | 11 | 0 | 0.3125 | 1 | # | 20 | Baseline | paralle | 73 |
| # | interaction | 4 | 6 | 0 | 0.4 | 1 | # | 20 | Baseline | paralle | 73 |
| # | disease | 7 | 4 | 0 | 0.636363636 | 1 | # | 20 | Baseline | ancom | 73 |
| # | sex | 4 | 0 | 0 | 1 | 1 | # | 20 | Baseline | ancom | 73 |
| # | time | 5 | 4 | 0 | 0.555555556 | 1 | # | 20 | Baseline | ancom | 73 |
| # | interaction | 0 | 0 | 4 | 0 | 0 | # | 20 | Baseline | ancom | 73 |
| # | disease | 7 | 33 | 0 | 0.175 | 1 | # | 20 | Baseline | aldex2 | 73 |
| # | sex | 4 | 10 | 0 | 0.285714286 | 1 | # | 20 | Baseline | aldex2 | 73 |
| # | time | 5 | 25 | 0 | 0.166666667 | 1 | # | 20 | Baseline | aldex2 | 73 |
| # | interaction | 0 | 0 | 4 | 0 | 0 | # | 20 | Baseline | aldex2 | 73 |
| # | disease | 7 | 0 | 0 | 1 | 1 | # | 20 | Small | paralle | 73 |
| # | sex | 4 | 0 | 0 | 1 | 1 | # | 20 | Small | paralle | 73 |
| # | time | 5 | 0 | 0 | 1 | 1 | # | 20 | Small | paralle | 73 |
| # | interaction | 4 | 0 | 0 | 1 | 1 | # | 20 | Small | paralle | 73 |
| # | disease | 7 | 4 | 0 | 0.636363636 | 1 | # | 20 | Small | ancom | 73 |
| # | sex | 4 | 0 | 0 | 1 | 1 | # | 20 | Small | ancom | 73 |
| # | time | 5 | 4 | 0 | 0.555555556 | 1 | # | 20 | Small | ancom | 73 |
| # | interaction | 0 | 0 | 4 | 0 | 0 | # | 20 | Small | ancom | 73 |
| # | disease | 7 | 3 | 0 | 0.7 | 1 | # | 20 | Small | aldex2 | 73 |
| # | sex | 4 | 0 | 0 | 1 | 1 | # | 20 | Small | aldex2 | 73 |
| # | time | 5 | 5 | 0 | 0.5 | 1 | # | 20 | Small | aldex2 | 73 |
| # | interaction | 0 | 0 | 4 | 0 | 0 | # | 20 | Small | aldex2 | 73 |
| # | disease | 7 | 10 | 0 | 0.411764706 | 1 | # | 20 | Baseline | paralle | 73 |
| # | sex | 4 | 10 | 0 | 0.285714286 | 1 | # | 20 | Baseline | paralle | 73 |
| # | time | 5 | 11 | 0 | 0.3125 | 1 | # | 20 | Baseline | paralle | 73 |
| # | interaction | 4 | 3 | 0 | 0.571428571 | 1 | # | 20 | Baseline | paralle | 73 |
| # | disease | 7 | 4 | 0 | 0.636363636 | 1 | # | 20 | Baseline | ancom | 73 |
| # | sex | 4 | 0 | 0 | 1 | 1 | # | 20 | Baseline | ancom | 73 |
| # | time | 5 | 4 | 0 | 0.555555556 | 1 | # | 20 | Baseline | ancom | 73 |
| # | interaction | 0 | 0 | 4 | 0 | 0 | # | 20 | Baseline | ancom | 73 |
| # | disease | 7 | 67 | 0 | 0.12962963 | 1 | # | 20 | Baseline | aldex2 | 73 |
| # | sex | 4 | 16 | 0 | 0.2 | 1 | # | 20 | Baseline | aldex2 | 73 |
| # | time | 5 | 32 | 0 | 0.135135135 | 1 | # | 20 | Baseline | aldex2 | 73 |
| # | interaction | 0 | 0 | 4 | 0 | 0 | # | 20 | Baseline | aldex2 | 73 |
| # | disease | 7 | 1 | 0 | 0.875 | 1 | # | 20 | Small | paralle | 73 |
| # | sex | 4 | 0 | 0 | 1 | 1 | # | 20 | Small | paralle | 73 |
| # | time | 5 | 1 | 0 | 0.833333333 | 1 | # | 20 | Small | paralle | 73 |
| # | interaction | 4 | 0 | 0 | 1 | 1 | # | 20 | Small | paralle | 73 |
| # | disease | 7 | 4 | 0 | 0.636363636 | 1 | # | 20 | Small | ancom | 73 |
| # | sex | 4 | 0 | 0 | 1 | 1 | # | 20 | Small | ancom | 73 |
| # | time | 5 | 4 | 0 | 0.555555556 | 1 | # | 20 | Small | ancom | 73 |
| # | interaction | 0 | 0 | 4 | 0 | 0 | # | 20 | Small | ancom | 73 |
| # | disease | 7 | 7 | 0 | 0.5 | 1 | # | 20 | Small | aldex2 | 73 |
| # | sex | 4 | 2 | 0 | 0.666666667 | 1 | # | 20 | Small | aldex2 | 73 |
| # | time | 5 | 6 | 0 | 0.454545455 | 1 | # | 20 | Small | aldex2 | 73 |
| # | interaction | 0 | 0 | 4 | 0 | 0 | # | 20 | Small | aldex2 | 73 |
| # | disease | 7 | 11 | 0 | 0.388888889 | 1 | # | 20 | Baseline | paralle | 73 |
| # | sex | 4 | 5 | 0 | 0.444444444 | 1 | # | 20 | Baseline | paralle | 73 |
| # | time | 5 | 8 | 0 | 0.384615385 | 1 | # | 20 | Baseline | paralle | 73 |
| # | interaction | 4 | 4 | 0 | 0.5 | 1 | # | 20 | Baseline | paralle | 73 |
| # | disease | 7 | 4 | 0 | 0.636363636 | 1 | # | 20 | Baseline | ancom | 73 |
| # | sex | 4 | 0 | 0 | 1 | 1 | # | 20 | Baseline | ancom | 73 |
| # | time | 5 | 4 | 0 | 0.555555556 | 1 | # | 20 | Baseline | ancom | 73 |
| # | interaction | 0 | 0 | 4 | 0 | 0 | # | 20 | Baseline | ancom | 73 |
| # | disease | 7 | 56 | 0 | 0.111111111 | 1 | # | 20 | Baseline | aldex2 | 73 |
| # | sex | 4 | 20 | 0 | 0.166666667 | 1 | # | 20 | Baseline | aldex2 | 73 |
| # | time | 5 | 41 | 0 | 0.108695652 | 1 | # | 20 | Baseline | aldex2 | 73 |
| # | interaction | 0 | 0 | 4 | 0 | 0 | # | 20 | Baseline | aldex2 | 73 |
| # | disease | 7 | 0 | 0 | 1 | 1 | # | 20 | Small | paralle | 73 |
| # | sex | 4 | 0 | 0 | 1 | 1 | # | 20 | Small | paralle | 73 |
| # | time | 5 | 0 | 0 | 1 | 1 | # | 20 | Small | paralle | 73 |
| # | interaction | 4 | 0 | 0 | 1 | 1 | # | 20 | Small | paralle | 73 |
| # | disease | 7 | 4 | 0 | 0.636363636 | 1 | # | 20 | Small | ancom | 73 |
| # | sex | 4 | 0 | 0 | 1 | 1 | # | 20 | Small | ancom | 73 |
| # | time | 5 | 4 | 0 | 0.555555556 | 1 | # | 20 | Small | ancom | 73 |
| # | interaction | 0 | 0 | 4 | 0 | 0 | # | 20 | Small | ancom | 73 |
| # | disease | 7 | 9 | 0 | 0.4375 | 1 | # | 20 | Small | aldex2 | 73 |
| # | sex | 4 | 2 | 0 | 0.666666667 | 1 | # | 20 | Small | aldex2 | 73 |
| # | time | 5 | 9 | 0 | 0.357142857 | 1 | # | 20 | Small | aldex2 | 73 |
| # | interaction | 0 | 0 | 4 | 0 | 0 | # | 20 | Small | aldex2 | 73 |
| # | disease | 4 | 11 | 0 | 0.266666667 | 1 | # | 10 | Baseline | paralle | 74 |
| # | sex | 2 | 7 | 0 | 0.222222222 | 1 | # | 10 | Baseline | paralle | 74 |
| # | time | 2 | 10 | 0 | 0.166666667 | 1 | # | 10 | Baseline | paralle | 74 |
| # | interaction | 2 | 7 | 0 | 0.222222222 | 1 | # | 10 | Baseline | paralle | 74 |
| # | disease | 4 | 0 | 0 | 1 | 1 | # | 10 | Baseline | ancom | 74 |
| # | sex | 2 | 0 | 0 | 1 | 1 | # | 10 | Baseline | ancom | 74 |
| # | time | 2 | 0 | 0 | 1 | 1 | # | 10 | Baseline | ancom | 74 |
| # | interaction | 0 | 0 | 2 | 0 | 0 | # | 10 | Baseline | ancom | 74 |
| # | disease | 4 | 12 | 0 | 0.25 | 1 | # | 10 | Baseline | aldex2 | 74 |
| # | sex | 2 | 6 | 0 | 0.25 | 1 | # | 10 | Baseline | aldex2 | 74 |
| # | time | 2 | 10 | 0 | 0.166666667 | 1 | # | 10 | Baseline | aldex2 | 74 |
| # | interaction | 0 | 0 | 2 | 0 | 0 | # | 10 | Baseline | aldex2 | 74 |
| # | disease | 4 | 0 | 0 | 1 | 1 | # | 10 | Small | paralle | 74 |
| # | sex | 2 | 0 | 0 | 1 | 1 | # | 10 | Small | paralle | 74 |
| # | time | 2 | 1 | 0 | 0.666666667 | 1 | # | 10 | Small | paralle | 74 |
| # | interaction | 2 | 0 | 0 | 1 | 1 | # | 10 | Small | paralle | 74 |
| # | disease | 4 | 0 | 0 | 1 | 1 | # | 10 | Small | ancom | 74 |
| # | sex | 2 | 0 | 0 | 1 | 1 | # | 10 | Small | ancom | 74 |
| # | time | 2 | 0 | 0 | 1 | 1 | # | 10 | Small | ancom | 74 |
| # | interaction | 0 | 0 | 2 | 0 | 0 | # | 10 | Small | ancom | 74 |
| # | disease | 4 | 0 | 0 | 1 | 1 | # | 10 | Small | aldex2 | 74 |
| # | sex | 2 | 0 | 0 | 1 | 1 | # | 10 | Small | aldex2 | 74 |
| # | time | 2 | 0 | 0 | 1 | 1 | # | 10 | Small | aldex2 | 74 |
| # | interaction | 0 | 0 | 2 | 0 | 0 | # | 10 | Small | aldex2 | 74 |
| # | disease | 4 | 11 | 0 | 0.266666667 | 1 | # | 10 | Baseline | paralle | 74 |
| # | sex | 2 | 9 | 0 | 0.181818182 | 1 | # | 10 | Baseline | paralle | 74 |
| # | time | 2 | 9 | 0 | 0.181818182 | 1 | # | 10 | Baseline | paralle | 74 |
| # | interaction | 2 | 7 | 0 | 0.222222222 | 1 | # | 10 | Baseline | paralle | 74 |
| # | disease | 4 | 2 | 0 | 0.666666667 | 1 | # | 10 | Baseline | ancom | 74 |
| # | sex | 2 | 0 | 0 | 1 | 1 | # | 10 | Baseline | ancom | 74 |
| # | time | 2 | 2 | 0 | 0.5 | 1 | # | 10 | Baseline | ancom | 74 |
| # | interaction | 0 | 0 | 2 | 0 | 0 | # | 10 | Baseline | ancom | 74 |
| # | disease | 4 | 19 | 0 | 0.173913043 | 1 | # | 10 | Baseline | aldex2 | 74 |
| # | sex | 2 | 7 | 0 | 0.222222222 | 1 | # | 10 | Baseline | aldex2 | 74 |
| # | time | 2 | 11 | 0 | 0.153846154 | 1 | # | 10 | Baseline | aldex2 | 74 |
| # | interaction | 0 | 0 | 2 | 0 | 0 | # | 10 | Baseline | aldex2 | 74 |
| # | disease | 4 | 0 | 0 | 1 | 1 | # | 10 | Small | paralle | 74 |
| # | sex | 2 | 0 | 0 | 1 | 1 | # | 10 | Small | paralle | 74 |
| # | time | 2 | 0 | 0 | 1 | 1 | # | 10 | Small | paralle | 74 |
| # | interaction | 2 | 0 | 0 | 1 | 1 | # | 10 | Small | paralle | 74 |
| # | disease | 4 | 2 | 0 | 0.666666667 | 1 | # | 10 | Small | ancom | 74 |
| # | sex | 2 | 0 | 0 | 1 | 1 | # | 10 | Small | ancom | 74 |
| # | time | 2 | 2 | 0 | 0.5 | 1 | # | 10 | Small | ancom | 74 |
| # | interaction | 0 | 0 | 2 | 0 | 0 | # | 10 | Small | ancom | 74 |
| # | disease | 4 | 2 | 0 | 0.666666667 | 1 | # | 10 | Small | aldex2 | 74 |
| # | sex | 2 | 0 | 0 | 1 | 1 | # | 10 | Small | aldex2 | 74 |
| # | time | 2 | 2 | 0 | 0.5 | 1 | # | 10 | Small | aldex2 | 74 |
| # | interaction | 0 | 0 | 2 | 0 | 0 | # | 10 | Small | aldex2 | 74 |
| # | disease | 4 | 7 | 0 | 0.363636364 | 1 | # | 10 | Baseline | paralle | 74 |
| # | sex | 2 | 5 | 0 | 0.285714286 | 1 | # | 10 | Baseline | paralle | 74 |
| # | time | 2 | 3 | 0 | 0.4 | 1 | # | 10 | Baseline | paralle | 74 |
| # | interaction | 2 | 4 | 0 | 0.333333333 | 1 | # | 10 | Baseline | paralle | 74 |
| # | disease | 4 | 3 | 0 | 0.571428571 | 1 | # | 10 | Baseline | ancom | 74 |
| # | sex | 2 | 0 | 0 | 1 | 1 | # | 10 | Baseline | ancom | 74 |
| # | time | 2 | 2 | 0 | 0.5 | 1 | # | 10 | Baseline | ancom | 74 |
| # | interaction | 0 | 0 | 2 | 0 | 0 | # | 10 | Baseline | ancom | 74 |
| # | disease | 4 | 22 | 0 | 0.153846154 | 1 | # | 10 | Baseline | aldex2 | 74 |
| # | sex | 2 | 11 | 0 | 0.153846154 | 1 | # | 10 | Baseline | aldex2 | 74 |
| # | time | 2 | 13 | 0 | 0.133333333 | 1 | # | 10 | Baseline | aldex2 | 74 |
| # | interaction | 0 | 0 | 2 | 0 | 0 | # | 10 | Baseline | aldex2 | 74 |
| # | disease | 4 | 0 | 0 | 1 | 1 | # | 10 | Small | paralle | 74 |
| # | sex | 2 | 0 | 0 | 1 | 1 | # | 10 | Small | paralle | 74 |
| # | time | 2 | 0 | 0 | 1 | 1 | # | 10 | Small | paralle | 74 |
| # | interaction | 2 | 0 | 0 | 1 | 1 | # | 10 | Small | paralle | 74 |
| # | disease | 4 | 3 | 0 | 0.571428571 | 1 | # | 10 | Small | ancom | 74 |
| # | sex | 2 | 0 | 0 | 1 | 1 | # | 10 | Small | ancom | 74 |
| # | time | 2 | 2 | 0 | 0.5 | 1 | # | 10 | Small | ancom | 74 |
| # | interaction | 0 | 0 | 2 | 0 | 0 | # | 10 | Small | ancom | 74 |
| # | disease | 4 | 3 | 0 | 0.571428571 | 1 | # | 10 | Small | aldex2 | 74 |
| # | sex | 2 | 0 | 0 | 1 | 1 | # | 10 | Small | aldex2 | 74 |
| # | time | 2 | 2 | 0 | 0.5 | 1 | # | 10 | Small | aldex2 | 74 |
| # | interaction | 0 | 0 | 2 | 0 | 0 | # | 10 | Small | aldex2 | 74 |
| # | disease | 4 | 6 | 0 | 0.4 | 1 | # | 10 | Baseline | paralle | 74 |
| # | sex | 2 | 6 | 0 | 0.25 | 1 | # | 10 | Baseline | paralle | 74 |
| # | time | 2 | 5 | 0 | 0.285714286 | 1 | # | 10 | Baseline | paralle | 74 |
| # | interaction | 2 | 5 | 0 | 0.285714286 | 1 | # | 10 | Baseline | paralle | 74 |
| # | disease | 4 | 2 | 0 | 0.666666667 | 1 | # | 10 | Baseline | ancom | 74 |
| # | sex | 2 | 0 | 0 | 1 | 1 | # | 10 | Baseline | ancom | 74 |
| # | time | 2 | 2 | 0 | 0.5 | 1 | # | 10 | Baseline | ancom | 74 |
| # | interaction | 0 | 0 | 2 | 0 | 0 | # | 10 | Baseline | ancom | 74 |
| # | disease | 4 | 25 | 0 | 0.137931034 | 1 | # | 10 | Baseline | aldex2 | 74 |
| # | sex | 2 | 12 | 0 | 0.142857143 | 1 | # | 10 | Baseline | aldex2 | 74 |
| # | time | 2 | 15 | 0 | 0.117647059 | 1 | # | 10 | Baseline | aldex2 | 74 |
| # | interaction | 0 | 0 | 2 | 0 | 0 | # | 10 | Baseline | aldex2 | 74 |
| # | disease | 4 | 0 | 0 | 1 | 1 | # | 10 | Small | paralle | 74 |
| # | sex | 2 | 0 | 0 | 1 | 1 | # | 10 | Small | paralle | 74 |
| # | time | 2 | 0 | 0 | 1 | 1 | # | 10 | Small | paralle | 74 |
| # | interaction | 2 | 0 | 0 | 1 | 1 | # | 10 | Small | paralle | 74 |
| # | disease | 4 | 2 | 0 | 0.666666667 | 1 | # | 10 | Small | ancom | 74 |
| # | sex | 2 | 0 | 0 | 1 | 1 | # | 10 | Small | ancom | 74 |
| # | time | 2 | 2 | 0 | 0.5 | 1 | # | 10 | Small | ancom | 74 |
| # | interaction | 0 | 0 | 2 | 0 | 0 | # | 10 | Small | ancom | 74 |
| # | disease | 4 | 4 | 0 | 0.5 | 1 | # | 10 | Small | aldex2 | 74 |
| # | sex | 2 | 1 | 0 | 0.666666667 | 1 | # | 10 | Small | aldex2 | 74 |
| # | time | 2 | 3 | 0 | 0.4 | 1 | # | 10 | Small | aldex2 | 74 |
| # | interaction | 0 | 0 | 2 | 0 | 0 | # | 10 | Small | aldex2 | 74 |
| # | disease | 7 | 10 | 0 | 0.411764706 | 1 | # | 20 | Baseline | paralle | 74 |
| # | sex | 4 | 4 | 0 | 0.5 | 1 | # | 20 | Baseline | paralle | 74 |
| # | time | 5 | 9 | 0 | 0.357142857 | 1 | # | 20 | Baseline | paralle | 74 |
| # | interaction | 2 | 2 | 2 | 0.5 | 0.5 | # | 20 | Baseline | paralle | 74 |
| # | disease | 7 | 4 | 0 | 0.636363636 | 1 | # | 20 | Baseline | ancom | 74 |
| # | sex | 4 | 0 | 0 | 1 | 1 | # | 20 | Baseline | ancom | 74 |
| # | time | 5 | 2 | 0 | 0.714285714 | 1 | # | 20 | Baseline | ancom | 74 |
| # | interaction | 0 | 0 | 4 | 0 | 0 | # | 20 | Baseline | ancom | 74 |
| # | disease | 7 | 23 | 0 | 0.233333333 | 1 | # | 20 | Baseline | aldex2 | 74 |
| # | sex | 4 | 4 | 0 | 0.5 | 1 | # | 20 | Baseline | aldex2 | 74 |
| # | time | 5 | 11 | 0 | 0.3125 | 1 | # | 20 | Baseline | aldex2 | 74 |
| # | interaction | 0 | 0 | 4 | 0 | 0 | # | 20 | Baseline | aldex2 | 74 |
| # | disease | 7 | 0 | 0 | 1 | 1 | # | 20 | Small | paralle | 74 |
| # | sex | 4 | 0 | 0 | 1 | 1 | # | 20 | Small | paralle | 74 |
| # | time | 5 | 1 | 0 | 0.833333333 | 1 | # | 20 | Small | paralle | 74 |
| # | interaction | 2 | 0 | 2 | 1 | 0.5 | # | 20 | Small | paralle | 74 |
| # | disease | 7 | 4 | 0 | 0.636363636 | 1 | # | 20 | Small | ancom | 74 |
| # | sex | 4 | 0 | 0 | 1 | 1 | # | 20 | Small | ancom | 74 |
| # | time | 5 | 1 | 0 | 0.833333333 | 1 | # | 20 | Small | ancom | 74 |
| # | interaction | 0 | 0 | 4 | 0 | 0 | # | 20 | Small | ancom | 74 |
| # | disease | 7 | 1 | 0 | 0.875 | 1 | # | 20 | Small | aldex2 | 74 |
| # | sex | 4 | 0 | 0 | 1 | 1 | # | 20 | Small | aldex2 | 74 |
| # | time | 5 | 2 | 0 | 0.714285714 | 1 | # | 20 | Small | aldex2 | 74 |
| # | interaction | 0 | 0 | 4 | 0 | 0 | # | 20 | Small | aldex2 | 74 |
| # | disease | 7 | 3 | 0 | 0.7 | 1 | # | 20 | Baseline | paralle | 74 |
| # | sex | 4 | 5 | 0 | 0.444444444 | 1 | # | 20 | Baseline | paralle | 74 |
| # | time | 5 | 8 | 0 | 0.384615385 | 1 | # | 20 | Baseline | paralle | 74 |
| # | interaction | 4 | 3 | 0 | 0.571428571 | 1 | # | 20 | Baseline | paralle | 74 |
| # | disease | 7 | 4 | 0 | 0.636363636 | 1 | # | 20 | Baseline | ancom | 74 |
| # | sex | 4 | 0 | 0 | 1 | 1 | # | 20 | Baseline | ancom | 74 |
| # | time | 5 | 3 | 0 | 0.625 | 1 | # | 20 | Baseline | ancom | 74 |
| # | interaction | 0 | 0 | 4 | 0 | 0 | # | 20 | Baseline | ancom | 74 |
| # | disease | 7 | 29 | 0 | 0.194444444 | 1 | # | 20 | Baseline | aldex2 | 74 |
| # | sex | 4 | 7 | 0 | 0.363636364 | 1 | # | 20 | Baseline | aldex2 | 74 |
| # | time | 5 | 26 | 0 | 0.161290323 | 1 | # | 20 | Baseline | aldex2 | 74 |
| # | interaction | 0 | 0 | 4 | 0 | 0 | # | 20 | Baseline | aldex2 | 74 |
| # | disease | 7 | 0 | 0 | 1 | 1 | # | 20 | Small | paralle | 74 |
| # | sex | 4 | 0 | 0 | 1 | 1 | # | 20 | Small | paralle | 74 |
| # | time | 5 | 0 | 0 | 1 | 1 | # | 20 | Small | paralle | 74 |

| | | | | | | | | | | | |
|---|---|---|---|---|---|---|---|---|---|---|---|
| # | interaction | 3 | 0 | 1 | 1 | 0.75 | # | 20 | Small | paralle | 74 |
| # | disease | 7 | 4 | 0 | 0.636363636 | 1 | # | 20 | Small | ancom | 74 |
| # | sex | 4 | 0 | 0 | 1 | 1 | # | 20 | Small | ancom | 74 |
| # | time | 5 | 2 | 0 | 0.714285714 | 1 | # | 20 | Small | ancom | 74 |
| # | interaction | 0 | 0 | 4 | 0 | 0 | # | 20 | Small | ancom | 74 |
| # | disease | 7 | 5 | 0 | 0.583333333 | 1 | # | 20 | Small | aldex2 | 74 |
| # | sex | 4 | 0 | 0 | 1 | 1 | # | 20 | Small | aldex2 | 74 |
| # | time | 5 | 3 | 0 | 0.625 | 1 | # | 20 | Small | aldex2 | 74 |
| # | interaction | 0 | 0 | 4 | 0 | 0 | # | 20 | Small | aldex2 | 74 |
| # | disease | 7 | 8 | 0 | 0.466666667 | 1 | # | 20 | Baseline | paralle | 74 |
| # | sex | 4 | 6 | 0 | 0.4 | 1 | # | 20 | Baseline | paralle | 74 |
| # | time | 5 | 9 | 0 | 0.357142857 | 1 | # | 20 | Baseline | paralle | 74 |
| # | interaction | 4 | 2 | 0 | 0.666666667 | 1 | # | 20 | Baseline | paralle | 74 |
| # | disease | 7 | 4 | 0 | 0.636363636 | 1 | # | 20 | Baseline | ancom | 74 |
| # | sex | 4 | 0 | 0 | 1 | 1 | # | 20 | Baseline | ancom | 74 |
| # | time | 5 | 4 | 0 | 0.555555556 | 1 | # | 20 | Baseline | ancom | 74 |
| # | interaction | 0 | 0 | 4 | 0 | 0 | # | 20 | Baseline | ancom | 74 |
| # | disease | 7 | 44 | 0 | 0.137254902 | 1 | # | 20 | Baseline | aldex2 | 74 |
| # | sex | 4 | 10 | 0 | 0.285714286 | 1 | # | 20 | Baseline | aldex2 | 74 |
| # | time | 5 | 35 | 0 | 0.125 | 1 | # | 20 | Baseline | aldex2 | 74 |
| # | interaction | 0 | 0 | 4 | 0 | 0 | # | 20 | Baseline | aldex2 | 74 |
| # | disease | 7 | 0 | 0 | 1 | 1 | # | 20 | Small | paralle | 74 |
| # | sex | 4 | 0 | 0 | 1 | 1 | # | 20 | Small | paralle | 74 |
| # | time | 5 | 0 | 0 | 1 | 1 | # | 20 | Small | paralle | 74 |
| # | interaction | 4 | 0 | 0 | 1 | 1 | # | 20 | Small | paralle | 74 |
| # | disease | 7 | 4 | 0 | 0.636363636 | 1 | # | 20 | Small | ancom | 74 |
| # | sex | 4 | 0 | 0 | 1 | 1 | # | 20 | Small | ancom | 74 |
| # | time | 5 | 4 | 0 | 0.555555556 | 1 | # | 20 | Small | ancom | 74 |
| # | interaction | 0 | 0 | 4 | 0 | 0 | # | 20 | Small | ancom | 74 |
| # | disease | 7 | 7 | 0 | 0.5 | 1 | # | 20 | Small | aldex2 | 74 |
| # | sex | 4 | 0 | 0 | 1 | 1 | # | 20 | Small | aldex2 | 74 |
| # | time | 5 | 7 | 0 | 0.416666667 | 1 | # | 20 | Small | aldex2 | 74 |
| # | interaction | 0 | 0 | 4 | 0 | 0 | # | 20 | Small | aldex2 | 74 |
| # | disease | 7 | 11 | 0 | 0.388888889 | 1 | # | 20 | Baseline | paralle | 74 |
| # | sex | 4 | 6 | 0 | 0.4 | 1 | # | 20 | Baseline | paralle | 74 |
| # | time | 5 | 15 | 0 | 0.25 | 1 | # | 20 | Baseline | paralle | 74 |
| # | interaction | 4 | 2 | 0 | 0.666666667 | 1 | # | 20 | Baseline | paralle | 74 |
| # | disease | 7 | 4 | 0 | 0.636363636 | 1 | # | 20 | Baseline | ancom | 74 |
| # | sex | 4 | 0 | 0 | 1 | 1 | # | 20 | Baseline | ancom | 74 |
| # | time | 5 | 4 | 0 | 0.555555556 | 1 | # | 20 | Baseline | ancom | 74 |
| # | interaction | 0 | 0 | 4 | 0 | 0 | # | 20 | Baseline | ancom | 74 |
| # | disease | 7 | 56 | 0 | 0.111111111 | 1 | # | 20 | Baseline | aldex2 | 74 |
| # | sex | 4 | 20 | 0 | 0.166666667 | 1 | # | 20 | Baseline | aldex2 | 74 |
| # | time | 5 | 62 | 0 | 0.106382979 | 1 | # | 20 | Baseline | aldex2 | 74 |
| # | interaction | 0 | 0 | 4 | 0 | 0 | # | 20 | Baseline | aldex2 | 74 |
| # | disease | 7 | 0 | 0 | 1 | 1 | # | 20 | Small | paralle | 74 |
| # | sex | 4 | 0 | 0 | 1 | 1 | # | 20 | Small | paralle | 74 |
| # | time | 5 | 0 | 0 | 1 | 1 | # | 20 | Small | paralle | 74 |
| # | interaction | 4 | 0 | 0 | 1 | 1 | # | 20 | Small | paralle | 74 |
| # | disease | 7 | 4 | 0 | 0.636363636 | 1 | # | 20 | Small | ancom | 74 |
| # | sex | 4 | 0 | 0 | 1 | 1 | # | 20 | Small | ancom | 74 |
| # | time | 5 | 4 | 0 | 0.555555556 | 1 | # | 20 | Small | ancom | 74 |
| # | interaction | 0 | 0 | 4 | 0 | 0 | # | 20 | Small | ancom | 74 |
| # | disease | 7 | 5 | 0 | 0.583333333 | 1 | # | 20 | Small | aldex2 | 74 |
| # | sex | 4 | 0 | 0 | 1 | 1 | # | 20 | Small | aldex2 | 74 |
| # | time | 5 | 9 | 0 | 0.357142857 | 1 | # | 20 | Small | aldex2 | 74 |
| # | interaction | 0 | 0 | 4 | 0 | 0 | # | 20 | Small | aldex2 | 74 |
| # | disease | 4 | 10 | 0 | 0.285714286 | 1 | # | 10 | Baseline | paralle | 75 |
| # | sex | 2 | 5 | 0 | 0.285714286 | 1 | # | 10 | Baseline | paralle | 75 |
| # | time | 2 | 7 | 0 | 0.222222222 | 1 | # | 10 | Baseline | paralle | 75 |
| # | interaction | 2 | 8 | 0 | 0.2 | 1 | # | 10 | Baseline | paralle | 75 |
| # | disease | 4 | 0 | 0 | 1 | 1 | # | 10 | Baseline | ancom | 75 |
| # | sex | 2 | 0 | 0 | 1 | 1 | # | 10 | Baseline | ancom | 75 |
| # | time | 2 | 0 | 0 | 1 | 1 | # | 10 | Baseline | ancom | 75 |
| # | interaction | 0 | 0 | 2 | 0 | 0 | # | 10 | Baseline | ancom | 75 |
| # | disease | 4 | 9 | 0 | 0.307692308 | 1 | # | 10 | Baseline | aldex2 | 75 |
| # | sex | 2 | 4 | 0 | 0.333333333 | 1 | # | 10 | Baseline | aldex2 | 75 |
| # | time | 2 | 6 | 0 | 0.25 | 1 | # | 10 | Baseline | aldex2 | 75 |
| # | interaction | 0 | 0 | 2 | 0 | 0 | # | 10 | Baseline | aldex2 | 75 |
| # | disease | 4 | 0 | 0 | 1 | 1 | # | 10 | Small | paralle | 75 |
| # | sex | 2 | 0 | 0 | 1 | 1 | # | 10 | Small | paralle | 75 |
| # | time | 2 | 0 | 0 | 1 | 1 | # | 10 | Small | paralle | 75 |
| # | interaction | 2 | 0 | 0 | 1 | 1 | # | 10 | Small | paralle | 75 |
| # | disease | 4 | 0 | 0 | 1 | 1 | # | 10 | Small | ancom | 75 |
| # | sex | 2 | 0 | 0 | 1 | 1 | # | 10 | Small | ancom | 75 |
| # | time | 2 | 0 | 0 | 1 | 1 | # | 10 | Small | ancom | 75 |
| # | interaction | 0 | 0 | 2 | 0 | 0 | # | 10 | Small | ancom | 75 |
| # | disease | 4 | 0 | 0 | 1 | 1 | # | 10 | Small | aldex2 | 75 |
| # | sex | 2 | 0 | 0 | 1 | 1 | # | 10 | Small | aldex2 | 75 |
| # | time | 2 | 0 | 0 | 1 | 1 | # | 10 | Small | aldex2 | 75 |
| # | interaction | 0 | 0 | 2 | 0 | 0 | # | 10 | Small | aldex2 | 75 |
| # | disease | 4 | 12 | 0 | 0.25 | 1 | # | 10 | Baseline | paralle | 75 |
| # | sex | 2 | 7 | 0 | 0.222222222 | 1 | # | 10 | Baseline | paralle | 75 |
| # | time | 2 | 8 | 0 | 0.2 | 1 | # | 10 | Baseline | paralle | 75 |
| # | interaction | 2 | 11 | 0 | 0.153846154 | 1 | # | 10 | Baseline | paralle | 75 |
| # | disease | 4 | 1 | 0 | 0.8 | 1 | # | 10 | Baseline | ancom | 75 |
| # | sex | 2 | 0 | 0 | 1 | 1 | # | 10 | Baseline | ancom | 75 |
| # | time | 2 | 2 | 0 | 0.5 | 1 | # | 10 | Baseline | ancom | 75 |
| # | interaction | 0 | 0 | 2 | 0 | 0 | # | 10 | Baseline | ancom | 75 |
| # | disease | 4 | 15 | 0 | 0.210526316 | 1 | # | 10 | Baseline | aldex2 | 75 |
| # | sex | 2 | 7 | 0 | 0.222222222 | 1 | # | 10 | Baseline | aldex2 | 75 |
| # | time | 2 | 10 | 0 | 0.166666667 | 1 | # | 10 | Baseline | aldex2 | 75 |
| # | interaction | 0 | 0 | 2 | 0 | 0 | # | 10 | Baseline | aldex2 | 75 |
| # | disease | 4 | 0 | 0 | 1 | 1 | # | 10 | Small | paralle | 75 |
| # | sex | 2 | 0 | 0 | 1 | 1 | # | 10 | Small | paralle | 75 |
| # | time | 2 | 0 | 0 | 1 | 1 | # | 10 | Small | paralle | 75 |
| # | interaction | 2 | 0 | 0 | 1 | 1 | # | 10 | Small | paralle | 75 |
| # | disease | 4 | 1 | 0 | 0.8 | 1 | # | 10 | Small | ancom | 75 |
| # | sex | 2 | 0 | 0 | 1 | 1 | # | 10 | Small | ancom | 75 |
| # | time | 2 | 2 | 0 | 0.5 | 1 | # | 10 | Small | ancom | 75 |
| # | interaction | 0 | 0 | 2 | 0 | 0 | # | 10 | Small | ancom | 75 |
| # | disease | 4 | 2 | 0 | 0.666666667 | 1 | # | 10 | Small | aldex2 | 75 |
| # | sex | 2 | 0 | 0 | 1 | 1 | # | 10 | Small | aldex2 | 75 |
| # | time | 2 | 1 | 0 | 0.666666667 | 1 | # | 10 | Small | aldex2 | 75 |
| # | interaction | 0 | 0 | 2 | 0 | 0 | # | 10 | Small | aldex2 | 75 |
| # | disease | 4 | 8 | 0 | 0.333333333 | 1 | # | 10 | Baseline | paralle | 75 |
| # | sex | 2 | 6 | 0 | 0.25 | 1 | # | 10 | Baseline | paralle | 75 |
| # | time | 2 | 4 | 0 | 0.333333333 | 1 | # | 10 | Baseline | paralle | 75 |
| # | interaction | 2 | 5 | 0 | 0.285714286 | 1 | # | 10 | Baseline | paralle | 75 |
| # | disease | 4 | 1 | 0 | 0.8 | 1 | # | 10 | Baseline | ancom | 75 |
| # | sex | 2 | 0 | 0 | 1 | 1 | # | 10 | Baseline | ancom | 75 |
| # | time | 2 | 2 | 0 | 0.5 | 1 | # | 10 | Baseline | ancom | 75 |
| # | interaction | 0 | 0 | 2 | 0 | 0 | # | 10 | Baseline | ancom | 75 |
| # | disease | 4 | 16 | 0 | 0.2 | 1 | # | 10 | Baseline | aldex2 | 75 |
| # | sex | 2 | 6 | 0 | 0.25 | 1 | # | 10 | Baseline | aldex2 | 75 |
| # | time | 2 | 9 | 0 | 0.181818182 | 1 | # | 10 | Baseline | aldex2 | 75 |
| # | interaction | 0 | 0 | 2 | 0 | 0 | # | 10 | Baseline | aldex2 | 75 |
| # | disease | 4 | 0 | 0 | 1 | 1 | # | 10 | Small | paralle | 75 |
| # | sex | 2 | 0 | 0 | 1 | 1 | # | 10 | Small | paralle | 75 |
| # | time | 2 | 0 | 0 | 1 | 1 | # | 10 | Small | paralle | 75 |
| # | interaction | 2 | 0 | 0 | 1 | 1 | # | 10 | Small | paralle | 75 |
| # | disease | 4 | 1 | 0 | 0.8 | 1 | # | 10 | Small | ancom | 75 |
| # | sex | 2 | 0 | 0 | 1 | 1 | # | 10 | Small | ancom | 75 |
| # | time | 2 | 2 | 0 | 0.5 | 1 | # | 10 | Small | ancom | 75 |
| # | interaction | 0 | 0 | 2 | 0 | 0 | # | 10 | Small | ancom | 75 |
| # | disease | 4 | 1 | 0 | 0.8 | 1 | # | 10 | Small | aldex2 | 75 |
| # | sex | 2 | 0 | 0 | 1 | 1 | # | 10 | Small | aldex2 | 75 |
| # | time | 2 | 2 | 0 | 0.5 | 1 | # | 10 | Small | aldex2 | 75 |
| # | interaction | 0 | 0 | 2 | 0 | 0 | # | 10 | Small | aldex2 | 75 |
| # | disease | 4 | 6 | 0 | 0.4 | 1 | # | 10 | Baseline | paralle | 75 |
| # | sex | 2 | 2 | 0 | 0.5 | 1 | # | 10 | Baseline | paralle | 75 |
| # | time | 2 | 5 | 0 | 0.285714286 | 1 | # | 10 | Baseline | paralle | 75 |
| # | interaction | 2 | 3 | 0 | 0.4 | 1 | # | 10 | Baseline | paralle | 75 |
| # | disease | 4 | 1 | 0 | 0.8 | 1 | # | 10 | Baseline | ancom | 75 |
| # | sex | 2 | 0 | 0 | 1 | 1 | # | 10 | Baseline | ancom | 75 |
| # | time | 2 | 2 | 0 | 0.5 | 1 | # | 10 | Baseline | ancom | 75 |
| # | interaction | 0 | 0 | 2 | 0 | 0 | # | 10 | Baseline | ancom | 75 |
| # | disease | 4 | 21 | 0 | 0.16 | 1 | # | 10 | Baseline | aldex2 | 75 |
| # | sex | 2 | 9 | 0 | 0.181818182 | 1 | # | 10 | Baseline | aldex2 | 75 |
| # | time | 2 | 10 | 0 | 0.166666667 | 1 | # | 10 | Baseline | aldex2 | 75 |
| # | interaction | 0 | 0 | 2 | 0 | 0 | # | 10 | Baseline | aldex2 | 75 |
| # | disease | 4 | 0 | 0 | 1 | 1 | # | 10 | Small | paralle | 75 |
| # | sex | 2 | 0 | 0 | 1 | 1 | # | 10 | Small | paralle | 75 |
| # | time | 2 | 0 | 0 | 1 | 1 | # | 10 | Small | paralle | 75 |
| # | interaction | 2 | 0 | 0 | 1 | 1 | # | 10 | Small | paralle | 75 |
| # | disease | 4 | 1 | 0 | 0.8 | 1 | # | 10 | Small | ancom | 75 |
| # | sex | 2 | 0 | 0 | 1 | 1 | # | 10 | Small | ancom | 75 |
| # | time | 2 | 2 | 0 | 0.5 | 1 | # | 10 | Small | ancom | 75 |
| # | interaction | 0 | 0 | 2 | 0 | 0 | # | 10 | Small | ancom | 75 |
| # | disease | 4 | 2 | 0 | 0.666666667 | 1 | # | 10 | Small | aldex2 | 75 |
| # | sex | 2 | 0 | 0 | 1 | 1 | # | 10 | Small | aldex2 | 75 |
| # | time | 2 | 3 | 0 | 0.4 | 1 | # | 10 | Small | aldex2 | 75 |
| # | interaction | 0 | 0 | 2 | 0 | 0 | # | 10 | Small | aldex2 | 75 |
| # | disease | 7 | 9 | 0 | 0.4375 | 1 | # | 20 | Baseline | paralle | 75 |
| # | sex | 4 | 8 | 0 | 0.333333333 | 1 | # | 20 | Baseline | paralle | 75 |
| # | time | 5 | 13 | 0 | 0.277777778 | 1 | # | 20 | Baseline | paralle | 75 |
| # | interaction | 3 | 11 | 1 | 0.214285714 | 0.75 | # | 20 | Baseline | paralle | 75 |
| # | disease | 7 | 1 | 0 | 0.875 | 1 | # | 20 | Baseline | ancom | 75 |
| # | sex | 4 | 0 | 0 | 1 | 1 | # | 20 | Baseline | ancom | 75 |
| # | time | 5 | 1 | 0 | 0.833333333 | 1 | # | 20 | Baseline | ancom | 75 |
| # | interaction | 0 | 0 | 4 | 0 | 0 | # | 20 | Baseline | ancom | 75 |
| # | disease | 7 | 20 | 0 | 0.259259259 | 1 | # | 20 | Baseline | aldex2 | 75 |
| # | sex | 4 | 7 | 0 | 0.363636364 | 1 | # | 20 | Baseline | aldex2 | 75 |
| # | time | 5 | 13 | 0 | 0.277777778 | 1 | # | 20 | Baseline | aldex2 | 75 |
| # | interaction | 0 | 0 | 4 | 0 | 0 | # | 20 | Baseline | aldex2 | 75 |
| # | disease | 7 | 0 | 0 | 1 | 1 | # | 20 | Small | paralle | 75 |
| # | sex | 4 | 0 | 0 | 1 | 1 | # | 20 | Small | paralle | 75 |
| # | time | 5 | 0 | 0 | 1 | 1 | # | 20 | Small | paralle | 75 |
| # | interaction | 0 | 0 | 4 | 0 | 0 | # | 20 | Small | paralle | 75 |
| # | disease | 7 | 1 | 0 | 0.875 | 1 | # | 20 | Small | ancom | 75 |
| # | sex | 4 | 0 | 0 | 1 | 1 | # | 20 | Small | ancom | 75 |
| # | time | 5 | 1 | 0 | 0.833333333 | 1 | # | 20 | Small | ancom | 75 |
| # | interaction | 0 | 0 | 4 | 0 | 0 | # | 20 | Small | ancom | 75 |
| # | disease | 7 | 1 | 0 | 0.875 | 1 | # | 20 | Small | aldex2 | 75 |
| # | sex | 4 | 0 | 0 | 1 | 1 | # | 20 | Small | aldex2 | 75 |
| # | time | 5 | 1 | 0 | 0.833333333 | 1 | # | 20 | Small | aldex2 | 75 |
| # | interaction | 0 | 0 | 4 | 0 | 0 | # | 20 | Small | aldex2 | 75 |
| # | disease | 7 | 18 | 0 | 0.28 | 1 | # | 20 | Baseline | paralle | 75 |
| # | sex | 4 | 5 | 0 | 0.444444444 | 1 | # | 20 | Baseline | paralle | 75 |
| # | time | 5 | 8 | 0 | 0.384615385 | 1 | # | 20 | Baseline | paralle | 75 |
| # | interaction | 4 | 7 | 0 | 0.363636364 | 1 | # | 20 | Baseline | paralle | 75 |
| # | disease | 7 | 3 | 0 | 0.7 | 1 | # | 20 | Baseline | ancom | 75 |
| # | sex | 4 | 0 | 0 | 1 | 1 | # | 20 | Baseline | ancom | 75 |
| # | time | 5 | 3 | 0 | 0.625 | 1 | # | 20 | Baseline | ancom | 75 |
| # | interaction | 0 | 0 | 4 | 0 | 0 | # | 20 | Baseline | ancom | 75 |
| # | disease | 7 | 35 | 0 | 0.166666667 | 1 | # | 20 | Baseline | aldex2 | 75 |
| # | sex | 4 | 9 | 0 | 0.307692308 | 1 | # | 20 | Baseline | aldex2 | 75 |
| # | time | 5 | 20 | 0 | 0.2 | 1 | # | 20 | Baseline | aldex2 | 75 |
| # | interaction | 0 | 0 | 4 | 0 | 0 | # | 20 | Baseline | aldex2 | 75 |
| # | disease | 7 | 0 | 0 | 1 | 1 | # | 20 | Small | paralle | 75 |
| # | sex | 4 | 0 | 0 | 1 | 1 | # | 20 | Small | paralle | 75 |
| # | time | 5 | 0 | 0 | 1 | 1 | # | 20 | Small | paralle | 75 |
| # | interaction | 2 | 0 | 2 | 1 | 0.5 | # | 20 | Small | paralle | 75 |
| # | disease | 7 | 2 | 0 | 0.777777778 | 1 | # | 20 | Small | ancom | 75 |
| # | sex | 4 | 0 | 0 | 1 | 1 | # | 20 | Small | ancom | 75 |
| # | time | 5 | 3 | 0 | 0.625 | 1 | # | 20 | Small | ancom | 75 |
| # | interaction | 0 | 0 | 4 | 0 | 0 | # | 20 | Small | ancom | 75 |
| # | disease | 7 | 4 | 0 | 0.636363636 | 1 | # | 20 | Small | aldex2 | 75 |
| # | sex | 4 | 0 | 0 | 1 | 1 | # | 20 | Small | aldex2 | 75 |
| # | time | 5 | 5 | 0 | 0.5 | 1 | # | 20 | Small | aldex2 | 75 |
| # | interaction | 0 | 0 | 4 | 0 | 0 | # | 20 | Small | aldex2 | 75 |
| # | disease | 7 | 10 | 0 | 0.411764706 | 1 | # | 20 | Baseline | paralle | 75 |
| # | sex | 4 | 9 | 0 | 0.307692308 | 1 | # | 20 | Baseline | paralle | 75 |
| # | time | 5 | 8 | 0 | 0.384615385 | 1 | # | 20 | Baseline | paralle | 75 |
| # | interaction | 4 | 6 | 0 | 0.4 | 1 | # | 20 | Baseline | paralle | 75 |
| # | disease | 7 | 3 | 0 | 0.7 | 1 | # | 20 | Baseline | ancom | 75 |
| # | sex | 4 | 0 | 0 | 1 | 1 | # | 20 | Baseline | ancom | 75 |
| # | time | 5 | 4 | 0 | 0.555555556 | 1 | # | 20 | Baseline | ancom | 75 |
| # | interaction | 0 | 0 | 4 | 0 | 0 | # | 20 | Baseline | ancom | 75 |
| # | disease | 7 | 47 | 0 | 0.12962963 | 1 | # | 20 | Baseline | aldex2 | 75 |
| # | sex | 4 | 12 | 0 | 0.25 | 1 | # | 20 | Baseline | aldex2 | 75 |
| # | time | 5 | 28 | 0 | 0.151515152 | 1 | # | 20 | Baseline | aldex2 | 75 |
| # | interaction | 0 | 0 | 4 | 0 | 0 | # | 20 | Baseline | aldex2 | 75 |
| # | disease | 7 | 0 | 0 | 1 | 1 | # | 20 | Small | paralle | 75 |
| # | sex | 4 | 0 | 0 | 1 | 1 | # | 20 | Small | paralle | 75 |
| # | time | 5 | 0 | 0 | 1 | 1 | # | 20 | Small | paralle | 75 |
| # | interaction | 4 | 0 | 0 | 1 | 1 | # | 20 | Small | paralle | 75 |
| # | disease | 7 | 3 | 0 | 0.7 | 1 | # | 20 | Small | ancom | 75 |
| # | sex | 4 | 0 | 0 | 1 | 1 | # | 20 | Small | ancom | 75 |
| # | time | 5 | 4 | 0 | 0.555555556 | 1 | # | 20 | Small | ancom | 75 |
| # | interaction | 0 | 0 | 4 | 0 | 0 | # | 20 | Small | ancom | 75 |
| # | disease | 7 | 6 | 0 | 0.538461538 | 1 | # | 20 | Small | aldex2 | 75 |
| # | sex | 4 | 0 | 0 | 1 | 1 | # | 20 | Small | aldex2 | 75 |
| # | time | 5 | 5 | 0 | 0.5 | 1 | # | 20 | Small | aldex2 | 75 |
| # | interaction | 0 | 0 | 4 | 0 | 0 | # | 20 | Small | aldex2 | 75 |
| # | disease | 7 | 13 | 0 | 0.35 | 1 | # | 20 | Baseline | paralle | 75 |
| # | sex | 4 | 9 | 0 | 0.307692308 | 1 | # | 20 | Baseline | paralle | 75 |
| # | time | 5 | 7 | 0 | 0.416666667 | 1 | # | 20 | Baseline | paralle | 75 |
| # | interaction | 4 | 8 | 0 | 0.333333333 | 1 | # | 20 | Baseline | paralle | 75 |
| # | disease | 7 | 4 | 0 | 0.636363636 | 1 | # | 20 | Baseline | ancom | 75 |
| # | sex | 4 | 0 | 0 | 1 | 1 | # | 20 | Baseline | ancom | 75 |
| # | time | 5 | 4 | 0 | 0.555555556 | 1 | # | 20 | Baseline | ancom | 75 |
| # | interaction | 0 | 0 | 4 | 0 | 0 | # | 20 | Baseline | ancom | 75 |
| # | disease | 7 | 90 | 0 | 0.104477612 | 1 | # | 20 | Baseline | aldex2 | 75 |
| # | sex | 4 | 17 | 0 | 0.19047619 | 1 | # | 20 | Baseline | aldex2 | 75 |
| # | time | 5 | 35 | 0 | 0.125 | 1 | # | 20 | Baseline | aldex2 | 75 |
| # | interaction | 0 | 0 | 4 | 0 | 0 | # | 20 | Baseline | aldex2 | 75 |
| # | disease | 7 | 0 | 0 | 1 | 1 | # | 20 | Small | paralle | 75 |
| # | sex | 4 | 0 | 0 | 1 | 1 | # | 20 | Small | paralle | 75 |
| # | time | 5 | 0 | 0 | 1 | 1 | # | 20 | Small | paralle | 75 |
| # | interaction | 4 | 0 | 0 | 1 | 1 | # | 20 | Small | paralle | 75 |
| # | disease | 7 | 4 | 0 | 0.636363636 | 1 | # | 20 | Small | ancom | 75 |
| # | sex | 4 | 0 | 0 | 1 | 1 | # | 20 | Small | ancom | 75 |
| # | time | 5 | 4 | 0 | 0.555555556 | 1 | # | 20 | Small | ancom | 75 |
| # | interaction | 0 | 0 | 4 | 0 | 0 | # | 20 | Small | ancom | 75 |
| # | disease | 7 | 10 | 0 | 0.411764706 | 1 | # | 20 | Small | aldex2 | 75 |
| # | sex | 4 | 1 | 0 | 0.8 | 1 | # | 20 | Small | aldex2 | 75 |
| # | time | 5 | 5 | 0 | 0.5 | 1 | # | 20 | Small | aldex2 | 75 |
| # | interaction | 0 | 0 | 4 | 0 | 0 | # | 20 | Small | aldex2 | 75 |
| # | disease | 4 | 8 | 0 | 0.333333333 | 1 | # | 10 | Baseline | paralle | 76 |
| # | sex | 2 | 7 | 0 | 0.222222222 | 1 | # | 10 | Baseline | paralle | 76 |
| # | time | 2 | 6 | 0 | 0.25 | 1 | # | 10 | Baseline | paralle | 76 |
| # | interaction | 2 | 2 | 0 | 0.5 | 1 | # | 10 | Baseline | paralle | 76 |
| # | disease | 4 | 0 | 0 | 1 | 1 | # | 10 | Baseline | ancom | 76 |
| # | sex | 2 | 0 | 0 | 1 | 1 | # | 10 | Baseline | ancom | 76 |
| # | time | 2 | 0 | 0 | 1 | 1 | # | 10 | Baseline | ancom | 76 |
| # | interaction | 0 | 0 | 2 | 0 | 0 | # | 10 | Baseline | ancom | 76 |
| # | disease | 4 | 8 | 0 | 0.333333333 | 1 | # | 10 | Baseline | aldex2 | 76 |
| # | sex | 2 | 6 | 0 | 0.25 | 1 | # | 10 | Baseline | aldex2 | 76 |
| # | time | 2 | 7 | 0 | 0.222222222 | 1 | # | 10 | Baseline | aldex2 | 76 |
| # | interaction | 0 | 0 | 2 | 0 | 0 | # | 10 | Baseline | aldex2 | 76 |
| # | disease | 4 | 0 | 0 | 1 | 1 | # | 10 | Small | paralle | 76 |
| # | sex | 2 | 2 | 0 | 0.5 | 1 | # | 10 | Small | paralle | 76 |
| # | time | 2 | 0 | 0 | 1 | 1 | # | 10 | Small | paralle | 76 |
| # | interaction | 1 | 0 | 1 | 1 | 0.5 | # | 10 | Small | paralle | 76 |
| # | disease | 4 | 0 | 0 | 1 | 1 | # | 10 | Small | ancom | 76 |
| # | sex | 2 | 0 | 0 | 1 | 1 | # | 10 | Small | ancom | 76 |
| # | time | 2 | 0 | 0 | 1 | 1 | # | 10 | Small | ancom | 76 |
| # | interaction | 0 | 0 | 2 | 0 | 0 | # | 10 | Small | ancom | 76 |
| # | disease | 4 | 0 | 0 | 1 | 1 | # | 10 | Small | aldex2 | 76 |
| # | sex | 2 | 0 | 0 | 1 | 1 | # | 10 | Small | aldex2 | 76 |
| # | time | 2 | 0 | 0 | 1 | 1 | # | 10 | Small | aldex2 | 76 |
| # | interaction | 0 | 0 | 2 | 0 | 0 | # | 10 | Small | aldex2 | 76 |
| # | disease | 4 | 11 | 0 | 0.266666667 | 1 | # | 10 | Baseline | paralle | 76 |
| # | sex | 2 | 9 | 0 | 0.181818182 | 1 | # | 10 | Baseline | paralle | 76 |
| # | time | 2 | 7 | 0 | 0.222222222 | 1 | # | 10 | Baseline | paralle | 76 |
| # | interaction | 2 | 5 | 0 | 0.285714286 | 1 | # | 10 | Baseline | paralle | 76 |
| # | disease | 4 | 2 | 0 | 0.666666667 | 1 | # | 10 | Baseline | ancom | 76 |
| # | sex | 2 | 0 | 0 | 1 | 1 | # | 10 | Baseline | ancom | 76 |
| # | time | 2 | 1 | 0 | 0.666666667 | 1 | # | 10 | Baseline | ancom | 76 |
| # | interaction | 0 | 0 | 2 | 0 | 0 | # | 10 | Baseline | ancom | 76 |
| # | disease | 4 | 11 | 0 | 0.266666667 | 1 | # | 10 | Baseline | aldex2 | 76 |
| # | sex | 2 | 9 | 0 | 0.181818182 | 1 | # | 10 | Baseline | aldex2 | 76 |
| # | time | 2 | 8 | 0 | 0.2 | 1 | # | 10 | Baseline | aldex2 | 76 |
| # | interaction | 0 | 0 | 2 | 0 | 0 | # | 10 | Baseline | aldex2 | 76 |
| # | disease | 4 | 0 | 0 | 1 | 1 | # | 10 | Small | paralle | 76 |
| # | sex | 2 | 1 | 0 | 0.666666667 | 1 | # | 10 | Small | paralle | 76 |
| # | time | 2 | 0 | 0 | 1 | 1 | # | 10 | Small | paralle | 76 |
| # | interaction | 2 | 0 | 0 | 1 | 1 | # | 10 | Small | paralle | 76 |
| # | disease | 4 | 2 | 0 | 0.666666667 | 1 | # | 10 | Small | ancom | 76 |
| # | sex | 2 | 0 | 0 | 1 | 1 | # | 10 | Small | ancom | 76 |
| # | time | 2 | 1 | 0 | 0.666666667 | 1 | # | 10 | Small | ancom | 76 |
| # | interaction | 0 | 0 | 2 | 0 | 0 | # | 10 | Small | ancom | 76 |
| # | disease | 4 | 3 | 0 | 0.571428571 | 1 | # | 10 | Small | aldex2 | 76 |
| # | sex | 2 | 0 | 0 | 1 | 1 | # | 10 | Small | aldex2 | 76 |
| # | time | 2 | 2 | 0 | 0.5 | 1 | # | 10 | Small | aldex2 | 76 |
| # | interaction | 0 | 0 | 2 | 0 | 0 | # | 10 | Small | aldex2 | 76 |
| # | disease | 4 | 8 | 0 | 0.333333333 | 1 | # | 10 | Baseline | paralle | 76 |
| # | sex | 2 | 7 | 0 | 0.222222222 | 1 | # | 10 | Baseline | paralle | 76 |
| # | time | 2 | 2 | 0 | 0.5 | 1 | # | 10 | Baseline | paralle | 76 |
| # | interaction | 2 | 2 | 0 | 0.5 | 1 | # | 10 | Baseline | paralle | 76 |
| # | disease | 4 | 3 | 0 | 0.571428571 | 1 | # | 10 | Baseline | ancom | 76 |
| # | sex | 2 | 0 | 0 | 1 | 1 | # | 10 | Baseline | ancom | 76 |
| # | time | 2 | 2 | 0 | 0.5 | 1 | # | 10 | Baseline | ancom | 76 |
| # | interaction | 0 | 0 | 2 | 0 | 0 | # | 10 | Baseline | ancom | 76 |
| # | disease | 4 | 20 | 0 | 0.166666667 | 1 | # | 10 | Baseline | aldex2 | 76 |
| # | sex | 2 | 10 | 0 | 0.166666667 | 1 | # | 10 | Baseline | aldex2 | 76 |
| # | time | 2 | 7 | 0 | 0.222222222 | 1 | # | 10 | Baseline | aldex2 | 76 |
| # | interaction | 0 | 0 | 2 | 0 | 0 | # | 10 | Baseline | aldex2 | 76 |
| # | disease | 4 | 0 | 0 | 1 | 1 | # | 10 | Small | paralle | 76 |
| # | sex | 2 | 1 | 0 | 0.666666667 | 1 | # | 10 | Small | paralle | 76 |
| # | time | 2 | 0 | 0 | 1 | 1 | # | 10 | Small | paralle | 76 |
| # | interaction | 2 | 0 | 0 | 1 | 1 | # | 10 | Small | paralle | 76 |
| # | disease | 4 | 2 | 0 | 0.666666667 | 1 | # | 10 | Small | ancom | 76 |
| # | sex | 2 | 0 | 0 | 1 | 1 | # | 10 | Small | ancom | 76 |
| # | time | 2 | 1 | 0 | 0.666666667 | 1 | # | 10 | Small | ancom | 76 |
| # | interaction | 0 | 0 | 2 | 0 | 0 | # | 10 | Small | ancom | 76 |
| # | disease | 4 | 3 | 0 | 0.571428571 | 1 | # | 10 | Small | aldex2 | 76 |
| # | sex | 2 | 0 | 0 | 1 | 1 | # | 10 | Small | aldex2 | 76 |
| # | time | 2 | 2 | 0 | 0.5 | 1 | # | 10 | Small | aldex2 | 76 |
| # | interaction | 0 | 0 | 2 | 0 | 0 | # | 10 | Small | aldex2 | 76 |
| # | disease | 4 | 9 | 0 | 0.307692308 | 1 | # | 10 | Baseline | paralle | 76 |
| # | sex | 2 | 9 | 0 | 0.181818182 | 1 | # | 10 | Baseline | paralle | 76 |
| # | time | 2 | 2 | 0 | 0.5 | 1 | # | 10 | Baseline | paralle | 76 |
| # | interaction | 2 | 2 | 0 | 0.5 | 1 | # | 10 | Baseline | paralle | 76 |
| # | disease | 4 | 2 | 0 | 0.666666667 | 1 | # | 10 | Baseline | ancom | 76 |
| # | sex | 2 | 0 | 0 | 1 | 1 | # | 10 | Baseline | ancom | 76 |
| # | time | 2 | 2 | 0 | 0.5 | 1 | # | 10 | Baseline | ancom | 76 |
| # | interaction | 0 | 0 | 2 | 0 | 0 | # | 10 | Baseline | ancom | 76 |
| # | disease | 4 | 19 | 0 | 0.173913043 | 1 | # | 10 | Baseline | aldex2 | 76 |
| # | sex | 2 | 14 | 0 | 0.125 | 1 | # | 10 | Baseline | aldex2 | 76 |
| # | time | 2 | 11 | 0 | 0.153846154 | 1 | # | 10 | Baseline | aldex2 | 76 |
| # | interaction | 0 | 0 | 2 | 0 | 0 | # | 10 | Baseline | aldex2 | 76 |
| # | disease | 4 | 0 | 0 | 1 | 1 | # | 10 | Small | paralle | 76 |
| # | sex | 2 | 1 | 0 | 0.666666667 | 1 | # | 10 | Small | paralle | 76 |
| # | time | 2 | 0 | 0 | 1 | 1 | # | 10 | Small | paralle | 76 |
| # | interaction | 2 | 0 | 0 | 1 | 1 | # | 10 | Small | paralle | 76 |

| | | | | | | | | | | | |
|---|---|---|---|---|---|---|---|---|---|---|---|
| # | disease | 4 | 2 | 0 | 0.666666667 | 1 | # | 10 | Small | ancom | 76 |
| # | sex | 2 | 0 | 0 | 1 | 1 | # | 10 | Small | ancom | 76 |
| # | time | 2 | 2 | 0 | 0.5 | 1 | # | 10 | Small | ancom | 76 |
| # | interaction | 0 | 0 | 2 | 0 | 0 | # | 10 | Small | ancom | 76 |
| # | disease | 4 | 4 | 0 | 0.5 | 1 | # | 10 | Small | aldex2 | 76 |
| # | sex | 2 | 0 | 0 | 1 | 1 | # | 10 | Small | aldex2 | 76 |
| # | time | 2 | 2 | 0 | 0.5 | 1 | # | 10 | Small | aldex2 | 76 |
| # | interaction | 0 | 0 | 2 | 0 | 0 | # | 10 | Small | aldex2 | 76 |
| # | disease | 7 | 5 | 0 | 0.583333333 | 1 | # | 20 | Baseline | paralle | 76 |
| # | sex | 4 | 10 | 0 | 0.285714286 | 1 | # | 20 | Baseline | paralle | 76 |
| # | time | 5 | 8 | 0 | 0.384615385 | 1 | # | 20 | Baseline | paralle | 76 |
| # | interaction | 4 | 5 | 0 | 0.444444444 | 1 | # | 20 | Baseline | paralle | 76 |
| # | disease | 7 | 1 | 0 | 0.875 | 1 | # | 20 | Baseline | ancom | 76 |
| # | sex | 4 | 0 | 0 | 1 | 1 | # | 20 | Baseline | ancom | 76 |
| # | time | 5 | 2 | 0 | 0.714285714 | 1 | # | 20 | Baseline | ancom | 76 |
| # | interaction | 0 | 0 | 4 | 0 | 0 | # | 20 | Baseline | ancom | 76 |
| # | disease | 7 | 20 | 0 | 0.259259259 | 1 | # | 20 | Baseline | aldex2 | 76 |
| # | sex | 4 | 7 | 0 | 0.363636364 | 1 | # | 20 | Baseline | aldex2 | 76 |
| # | time | 5 | 14 | 0 | 0.263157895 | 1 | # | 20 | Baseline | aldex2 | 76 |
| # | interaction | 0 | 0 | 4 | 0 | 0 | # | 20 | Baseline | aldex2 | 76 |
| # | disease | 7 | 1 | 0 | 0.875 | 1 | # | 20 | Small | paralle | 76 |
| # | sex | 4 | 1 | 0 | 0.8 | 1 | # | 20 | Small | paralle | 76 |
| # | time | 5 | 1 | 0 | 0.833333333 | 1 | # | 20 | Small | paralle | 76 |
| # | interaction | 3 | 1 | 1 | 0.75 | 0.75 | # | 20 | Small | paralle | 76 |
| # | disease | 7 | 0 | 0 | 1 | 1 | # | 20 | Small | ancom | 76 |
| # | sex | 4 | 0 | 0 | 1 | 1 | # | 20 | Small | ancom | 76 |
| # | time | 5 | 2 | 0 | 0.714285714 | 1 | # | 20 | Small | ancom | 76 |
| # | interaction | 0 | 0 | 4 | 0 | 0 | # | 20 | Small | ancom | 76 |
| # | disease | 7 | 0 | 0 | 1 | 1 | # | 20 | Small | aldex2 | 76 |
| # | sex | 4 | 0 | 0 | 1 | 1 | # | 20 | Small | aldex2 | 76 |
| # | time | 5 | 2 | 0 | 0.714285714 | 1 | # | 20 | Small | aldex2 | 76 |
| # | interaction | 0 | 0 | 4 | 0 | 0 | # | 20 | Small | aldex2 | 76 |
| # | disease | 7 | 10 | 0 | 0.411764706 | 1 | # | 20 | Baseline | paralle | 76 |
| # | sex | 4 | 8 | 0 | 0.333333333 | 1 | # | 20 | Baseline | paralle | 76 |
| # | time | 5 | 7 | 0 | 0.416666667 | 1 | # | 20 | Baseline | paralle | 76 |
| # | interaction | 4 | 5 | 0 | 0.444444444 | 1 | # | 20 | Baseline | paralle | 76 |
| # | disease | 7 | 4 | 0 | 0.636363636 | 1 | # | 20 | Baseline | ancom | 76 |
| # | sex | 4 | 0 | 0 | 1 | 1 | # | 20 | Baseline | ancom | 76 |
| # | time | 5 | 4 | 0 | 0.555555556 | 1 | # | 20 | Baseline | ancom | 76 |
| # | interaction | 0 | 0 | 4 | 0 | 0 | # | 20 | Baseline | ancom | 76 |
| # | disease | 7 | 36 | 0 | 0.162790698 | 1 | # | 20 | Baseline | aldex2 | 76 |
| # | sex | 4 | 12 | 0 | 0.25 | 1 | # | 20 | Baseline | aldex2 | 76 |
| # | time | 5 | 24 | 0 | 0.172413793 | 1 | # | 20 | Baseline | aldex2 | 76 |
| # | interaction | 0 | 0 | 4 | 0 | 0 | # | 20 | Baseline | aldex2 | 76 |
| # | disease | 7 | 0 | 0 | 1 | 1 | # | 20 | Small | paralle | 76 |
| # | sex | 4 | 0 | 0 | 1 | 1 | # | 20 | Small | paralle | 76 |
| # | time | 5 | 0 | 0 | 1 | 1 | # | 20 | Small | paralle | 76 |
| # | interaction | 3 | 0 | 1 | 1 | 0.75 | # | 20 | Small | paralle | 76 |
| # | disease | 7 | 3 | 0 | 0.7 | 1 | # | 20 | Small | ancom | 76 |
| # | sex | 4 | 0 | 0 | 1 | 1 | # | 20 | Small | ancom | 76 |
| # | time | 5 | 3 | 0 | 0.625 | 1 | # | 20 | Small | ancom | 76 |
| # | interaction | 0 | 0 | 4 | 0 | 0 | # | 20 | Small | ancom | 76 |
| # | disease | 7 | 1 | 0 | 0.875 | 1 | # | 20 | Small | aldex2 | 76 |
| # | sex | 4 | 0 | 0 | 1 | 1 | # | 20 | Small | aldex2 | 76 |
| # | time | 5 | 2 | 0 | 0.714285714 | 1 | # | 20 | Small | aldex2 | 76 |
| # | interaction | 0 | 0 | 4 | 0 | 0 | # | 20 | Small | aldex2 | 76 |
| # | disease | 7 | 8 | 0 | 0.466666667 | 1 | # | 20 | Baseline | paralle | 76 |
| # | sex | 4 | 9 | 0 | 0.307692308 | 1 | # | 20 | Baseline | paralle | 76 |
| # | time | 5 | 5 | 0 | 0.5 | 1 | # | 20 | Baseline | paralle | 76 |
| # | interaction | 4 | 3 | 0 | 0.571428571 | 1 | # | 20 | Baseline | paralle | 76 |
| # | disease | 7 | 4 | 0 | 0.636363636 | 1 | # | 20 | Baseline | ancom | 76 |
| # | sex | 4 | 0 | 0 | 1 | 1 | # | 20 | Baseline | ancom | 76 |
| # | time | 5 | 4 | 0 | 0.555555556 | 1 | # | 20 | Baseline | ancom | 76 |
| # | interaction | 0 | 0 | 4 | 0 | 0 | # | 20 | Baseline | ancom | 76 |
| # | disease | 7 | 53 | 0 | 0.116666667 | 1 | # | 20 | Baseline | aldex2 | 76 |
| # | sex | 4 | 19 | 0 | 0.173913043 | 1 | # | 20 | Baseline | aldex2 | 76 |
| # | time | 5 | 28 | 0 | 0.151515152 | 1 | # | 20 | Baseline | aldex2 | 76 |
| # | interaction | 0 | 0 | 4 | 0 | 0 | # | 20 | Baseline | aldex2 | 76 |
| # | disease | 7 | 0 | 0 | 1 | 1 | # | 20 | Small | paralle | 76 |
| # | sex | 4 | 1 | 0 | 0.8 | 1 | # | 20 | Small | paralle | 76 |
| # | time | 5 | 0 | 0 | 1 | 1 | # | 20 | Small | paralle | 76 |
| # | interaction | 4 | 0 | 0 | 1 | 1 | # | 20 | Small | paralle | 76 |
| # | disease | 7 | 4 | 0 | 0.636363636 | 1 | # | 20 | Small | ancom | 76 |
| # | sex | 4 | 0 | 0 | 1 | 1 | # | 20 | Small | ancom | 76 |
| # | time | 5 | 3 | 0 | 0.625 | 1 | # | 20 | Small | ancom | 76 |
| # | interaction | 0 | 0 | 4 | 0 | 0 | # | 20 | Small | ancom | 76 |
| # | disease | 7 | 5 | 0 | 0.583333333 | 1 | # | 20 | Small | aldex2 | 76 |
| # | sex | 4 | 1 | 0 | 0.8 | 1 | # | 20 | Small | aldex2 | 76 |
| # | time | 5 | 3 | 0 | 0.625 | 1 | # | 20 | Small | aldex2 | 76 |
| # | interaction | 0 | 0 | 4 | 0 | 0 | # | 20 | Small | aldex2 | 76 |
| # | disease | 7 | 7 | 0 | 0.5 | 1 | # | 20 | Baseline | paralle | 76 |
| # | sex | 4 | 7 | 0 | 0.363636364 | 1 | # | 20 | Baseline | paralle | 76 |
| # | time | 5 | 6 | 0 | 0.454545455 | 1 | # | 20 | Baseline | paralle | 76 |
| # | interaction | 4 | 5 | 0 | 0.444444444 | 1 | # | 20 | Baseline | paralle | 76 |
| # | disease | 7 | 4 | 0 | 0.636363636 | 1 | # | 20 | Baseline | ancom | 76 |
| # | sex | 4 | 0 | 0 | 1 | 1 | # | 20 | Baseline | ancom | 76 |
| # | time | 5 | 4 | 0 | 0.555555556 | 1 | # | 20 | Baseline | ancom | 76 |
| # | interaction | 0 | 0 | 4 | 0 | 0 | # | 20 | Baseline | ancom | 76 |
| # | disease | 7 | 13 | 0 | 0.1 | 1 | # | 20 | Baseline | aldex2 | 76 |
| # | sex | 4 | 19 | 0 | 0.173913043 | 1 | # | 20 | Baseline | aldex2 | 76 |
| # | time | 5 | 35 | 0 | 0.125 | 1 | # | 20 | Baseline | aldex2 | 76 |
| # | interaction | 0 | 0 | 4 | 0 | 0 | # | 20 | Baseline | aldex2 | 76 |
| # | disease | 7 | 0 | 0 | 1 | 1 | # | 20 | Small | paralle | 76 |
| # | sex | 4 | 0 | 0 | 1 | 1 | # | 20 | Small | paralle | 76 |
| # | time | 5 | 0 | 0 | 1 | 1 | # | 20 | Small | paralle | 76 |
| # | interaction | 4 | 0 | 0 | 1 | 1 | # | 20 | Small | paralle | 76 |
| # | disease | 7 | 4 | 0 | 0.636363636 | 1 | # | 20 | Small | ancom | 76 |
| # | sex | 4 | 0 | 0 | 1 | 1 | # | 20 | Small | ancom | 76 |
| # | time | 5 | 4 | 0 | 0.555555556 | 1 | # | 20 | Small | ancom | 76 |
| # | interaction | 0 | 0 | 4 | 0 | 0 | # | 20 | Small | ancom | 76 |
| # | disease | 7 | 8 | 0 | 0.466666667 | 1 | # | 20 | Small | aldex2 | 76 |
| # | sex | 4 | 2 | 0 | 0.666666667 | 1 | # | 20 | Small | aldex2 | 76 |
| # | time | 5 | 6 | 0 | 0.454545455 | 1 | # | 20 | Small | aldex2 | 76 |
| # | interaction | 0 | 0 | 4 | 0 | 0 | # | 20 | Small | aldex2 | 76 |
| # | disease | 4 | 8 | 0 | 0.333333333 | 1 | # | 10 | Baseline | paralle | 77 |
| # | sex | 2 | 11 | 0 | 0.153846154 | 1 | # | 10 | Baseline | paralle | 77 |
| # | time | 2 | 7 | 0 | 0.222222222 | 1 | # | 10 | Baseline | paralle | 77 |
| # | interaction | 2 | 7 | 0 | 0.222222222 | 1 | # | 10 | Baseline | paralle | 77 |
| # | disease | 4 | 1 | 0 | 0.8 | 1 | # | 10 | Baseline | ancom | 77 |
| # | sex | 2 | 0 | 0 | 1 | 1 | # | 10 | Baseline | ancom | 77 |
| # | time | 2 | 0 | 0 | 1 | 1 | # | 10 | Baseline | ancom | 77 |
| # | interaction | 0 | 0 | 2 | 0 | 0 | # | 10 | Baseline | ancom | 77 |
| # | disease | 4 | 10 | 0 | 0.285714286 | 1 | # | 10 | Baseline | aldex2 | 77 |
| # | sex | 2 | 4 | 0 | 0.333333333 | 1 | # | 10 | Baseline | aldex2 | 77 |
| # | time | 2 | 2 | 0 | 0.5 | 1 | # | 10 | Baseline | aldex2 | 77 |
| # | interaction | 0 | 0 | 2 | 0 | 0 | # | 10 | Baseline | aldex2 | 77 |
| # | disease | 4 | 1 | 0 | 0.8 | 1 | # | 10 | Small | paralle | 77 |
| # | sex | 2 | 0 | 0 | 1 | 1 | # | 10 | Small | paralle | 77 |
| # | time | 2 | 0 | 0 | 1 | 1 | # | 10 | Small | paralle | 77 |
| # | interaction | 1 | 0 | 1 | 1 | 0.5 | # | 10 | Small | paralle | 77 |
| # | disease | 4 | 1 | 0 | 0.8 | 1 | # | 10 | Small | ancom | 77 |
| # | sex | 2 | 0 | 0 | 1 | 1 | # | 10 | Small | ancom | 77 |
| # | time | 2 | 0 | 0 | 1 | 1 | # | 10 | Small | ancom | 77 |
| # | interaction | 0 | 0 | 2 | 0 | 0 | # | 10 | Small | ancom | 77 |
| # | disease | 4 | 3 | 0 | 0.571428571 | 1 | # | 10 | Small | aldex2 | 77 |
| # | sex | 2 | 0 | 0 | 1 | 1 | # | 10 | Small | aldex2 | 77 |
| # | time | 2 | 0 | 0 | 1 | 1 | # | 10 | Small | aldex2 | 77 |
| # | interaction | 0 | 0 | 2 | 0 | 0 | # | 10 | Small | aldex2 | 77 |
| # | disease | 4 | 9 | 0 | 0.307692308 | 1 | # | 10 | Baseline | paralle | 77 |
| # | sex | 2 | 8 | 0 | 0.2 | 1 | # | 10 | Baseline | paralle | 77 |
| # | time | 2 | 7 | 0 | 0.222222222 | 1 | # | 10 | Baseline | paralle | 77 |
| # | interaction | 2 | 7 | 0 | 0.222222222 | 1 | # | 10 | Baseline | paralle | 77 |
| # | disease | 4 | 2 | 0 | 0.666666667 | 1 | # | 10 | Baseline | ancom | 77 |
| # | sex | 2 | 0 | 0 | 1 | 1 | # | 10 | Baseline | ancom | 77 |
| # | time | 2 | 2 | 0 | 0.5 | 1 | # | 10 | Baseline | ancom | 77 |
| # | interaction | 0 | 0 | 2 | 0 | 0 | # | 10 | Baseline | ancom | 77 |
| # | disease | 4 | 16 | 0 | 0.2 | 1 | # | 10 | Baseline | aldex2 | 77 |
| # | sex | 2 | 3 | 0 | 0.4 | 1 | # | 10 | Baseline | aldex2 | 77 |
| # | time | 2 | 6 | 0 | 0.25 | 1 | # | 10 | Baseline | aldex2 | 77 |
| # | interaction | 0 | 0 | 2 | 0 | 0 | # | 10 | Baseline | aldex2 | 77 |
| # | disease | 4 | 1 | 0 | 0.8 | 1 | # | 10 | Small | paralle | 77 |
| # | sex | 2 | 0 | 0 | 1 | 1 | # | 10 | Small | paralle | 77 |
| # | time | 2 | 0 | 0 | 1 | 1 | # | 10 | Small | paralle | 77 |
| # | interaction | 2 | 0 | 0 | 1 | 1 | # | 10 | Small | paralle | 77 |
| # | disease | 4 | 2 | 0 | 0.666666667 | 1 | # | 10 | Small | ancom | 77 |
| # | sex | 2 | 0 | 0 | 1 | 1 | # | 10 | Small | ancom | 77 |
| # | time | 2 | 2 | 0 | 0.5 | 1 | # | 10 | Small | ancom | 77 |
| # | interaction | 0 | 0 | 2 | 0 | 0 | # | 10 | Small | ancom | 77 |
| # | disease | 4 | 2 | 0 | 0.666666667 | 1 | # | 10 | Small | aldex2 | 77 |
| # | sex | 2 | 0 | 0 | 1 | 1 | # | 10 | Small | aldex2 | 77 |
| # | time | 2 | 2 | 0 | 0.5 | 1 | # | 10 | Small | aldex2 | 77 |
| # | interaction | 0 | 0 | 2 | 0 | 0 | # | 10 | Small | aldex2 | 77 |
| # | disease | 4 | 5 | 0 | 0.444444444 | 1 | # | 10 | Baseline | paralle | 77 |
| # | sex | 2 | 8 | 0 | 0.2 | 1 | # | 10 | Baseline | paralle | 77 |
| # | time | 2 | 8 | 0 | 0.2 | 1 | # | 10 | Baseline | paralle | 77 |
| # | interaction | 2 | 9 | 0 | 0.181818182 | 1 | # | 10 | Baseline | paralle | 77 |
| # | disease | 4 | 2 | 0 | 0.666666667 | 1 | # | 10 | Baseline | ancom | 77 |
| # | sex | 2 | 0 | 0 | 1 | 1 | # | 10 | Baseline | ancom | 77 |
| # | time | 2 | 2 | 0 | 0.5 | 1 | # | 10 | Baseline | ancom | 77 |
| # | interaction | 0 | 0 | 2 | 0 | 0 | # | 10 | Baseline | ancom | 77 |
| # | disease | 4 | 19 | 0 | 0.173913043 | 1 | # | 10 | Baseline | aldex2 | 77 |
| # | sex | 2 | 5 | 0 | 0.285714286 | 1 | # | 10 | Baseline | aldex2 | 77 |
| # | time | 2 | 5 | 0 | 0.285714286 | 1 | # | 10 | Baseline | aldex2 | 77 |
| # | interaction | 0 | 0 | 2 | 0 | 0 | # | 10 | Baseline | aldex2 | 77 |
| # | disease | 4 | 1 | 0 | 0.8 | 1 | # | 10 | Small | paralle | 77 |
| # | sex | 2 | 0 | 0 | 1 | 1 | # | 10 | Small | paralle | 77 |
| # | time | 2 | 0 | 0 | 1 | 1 | # | 10 | Small | paralle | 77 |
| # | interaction | 2 | 0 | 0 | 1 | 1 | # | 10 | Small | paralle | 77 |
| # | disease | 4 | 2 | 0 | 0.666666667 | 1 | # | 10 | Small | ancom | 77 |
| # | sex | 2 | 0 | 0 | 1 | 1 | # | 10 | Small | ancom | 77 |
| # | time | 2 | 2 | 0 | 0.5 | 1 | # | 10 | Small | ancom | 77 |
| # | interaction | 0 | 0 | 2 | 0 | 0 | # | 10 | Small | ancom | 77 |
| # | disease | 4 | 3 | 0 | 0.571428571 | 1 | # | 10 | Small | aldex2 | 77 |
| # | sex | 2 | 0 | 0 | 1 | 1 | # | 10 | Small | aldex2 | 77 |
| # | time | 2 | 3 | 0 | 0.4 | 1 | # | 10 | Small | aldex2 | 77 |
| # | interaction | 0 | 0 | 2 | 0 | 0 | # | 10 | Small | aldex2 | 77 |
| # | disease | 4 | 6 | 0 | 0.4 | 1 | # | 10 | Baseline | paralle | 77 |
| # | sex | 2 | 7 | 0 | 0.222222222 | 1 | # | 10 | Baseline | paralle | 77 |
| # | time | 2 | 4 | 0 | 0.333333333 | 1 | # | 10 | Baseline | paralle | 77 |
| # | interaction | 2 | 4 | 0 | 0.333333333 | 1 | # | 10 | Baseline | paralle | 77 |
| # | disease | 4 | 2 | 0 | 0.666666667 | 1 | # | 10 | Baseline | ancom | 77 |
| # | sex | 2 | 0 | 0 | 1 | 1 | # | 10 | Baseline | ancom | 77 |
| # | time | 2 | 2 | 0 | 0.5 | 1 | # | 10 | Baseline | ancom | 77 |
| # | interaction | 0 | 0 | 2 | 0 | 0 | # | 10 | Baseline | ancom | 77 |
| # | disease | 4 | 24 | 0 | 0.142857143 | 1 | # | 10 | Baseline | aldex2 | 77 |
| # | sex | 2 | 6 | 0 | 0.25 | 1 | # | 10 | Baseline | aldex2 | 77 |
| # | time | 2 | 12 | 0 | 0.142857143 | 1 | # | 10 | Baseline | aldex2 | 77 |
| # | interaction | 0 | 0 | 2 | 0 | 0 | # | 10 | Baseline | aldex2 | 77 |
| # | disease | 4 | 0 | 0 | 1 | 1 | # | 10 | Small | paralle | 77 |
| # | sex | 2 | 0 | 0 | 1 | 1 | # | 10 | Small | paralle | 77 |
| # | time | 2 | 0 | 0 | 1 | 1 | # | 10 | Small | paralle | 77 |
| # | interaction | 2 | 0 | 0 | 1 | 1 | # | 10 | Small | paralle | 77 |
| # | disease | 4 | 2 | 0 | 0.666666667 | 1 | # | 10 | Small | ancom | 77 |
| # | sex | 2 | 0 | 0 | 1 | 1 | # | 10 | Small | ancom | 77 |
| # | time | 2 | 2 | 0 | 0.5 | 1 | # | 10 | Small | ancom | 77 |
| # | interaction | 0 | 0 | 2 | 0 | 0 | # | 10 | Small | ancom | 77 |
| # | disease | 4 | 5 | 0 | 0.444444444 | 1 | # | 10 | Small | aldex2 | 77 |
| # | sex | 2 | 0 | 0 | 1 | 1 | # | 10 | Small | aldex2 | 77 |
| # | time | 2 | 4 | 0 | 0.333333333 | 1 | # | 10 | Small | aldex2 | 77 |
| # | interaction | 0 | 0 | 2 | 0 | 0 | # | 10 | Small | aldex2 | 77 |
| # | disease | 7 | 4 | 0 | 0.636363636 | 1 | # | 20 | Baseline | paralle | 77 |
| # | sex | 4 | 10 | 0 | 0.285714286 | 1 | # | 20 | Baseline | paralle | 77 |
| # | time | 5 | 11 | 0 | 0.3125 | 1 | # | 20 | Baseline | paralle | 77 |
| # | interaction | 4 | 10 | 0 | 0.285714286 | 1 | # | 20 | Baseline | paralle | 77 |
| # | disease | 7 | 0 | 0 | 1 | 1 | # | 20 | Baseline | ancom | 77 |
| # | sex | 4 | 0 | 0 | 1 | 1 | # | 20 | Baseline | ancom | 77 |
| # | time | 5 | 0 | 0 | 1 | 1 | # | 20 | Baseline | ancom | 77 |
| # | interaction | 0 | 0 | 4 | 0 | 0 | # | 20 | Baseline | ancom | 77 |
| # | disease | 7 | 19 | 0 | 0.269230769 | 1 | # | 20 | Baseline | aldex2 | 77 |
| # | sex | 4 | 8 | 0 | 0.333333333 | 1 | # | 20 | Baseline | aldex2 | 77 |
| # | time | 5 | 16 | 0 | 0.238095238 | 1 | # | 20 | Baseline | aldex2 | 77 |
| # | interaction | 0 | 0 | 4 | 0 | 0 | # | 20 | Baseline | aldex2 | 77 |
| # | disease | 7 | 0 | 0 | 1 | 1 | # | 20 | Small | paralle | 77 |
| # | sex | 4 | 0 | 0 | 1 | 1 | # | 20 | Small | paralle | 77 |
| # | time | 5 | 1 | 0 | 0.833333333 | 1 | # | 20 | Small | paralle | 77 |
| # | interaction | 2 | 0 | 2 | 1 | 0.5 | # | 20 | Small | paralle | 77 |
| # | disease | 7 | 0 | 0 | 1 | 1 | # | 20 | Small | ancom | 77 |
| # | sex | 4 | 0 | 0 | 1 | 1 | # | 20 | Small | ancom | 77 |
| # | time | 5 | 0 | 0 | 1 | 1 | # | 20 | Small | ancom | 77 |
| # | interaction | 0 | 0 | 4 | 0 | 0 | # | 20 | Small | ancom | 77 |
| # | disease | 7 | 0 | 0 | 1 | 1 | # | 20 | Small | aldex2 | 77 |
| # | sex | 4 | 0 | 0 | 1 | 1 | # | 20 | Small | aldex2 | 77 |
| # | time | 5 | 1 | 0 | 0.833333333 | 1 | # | 20 | Small | aldex2 | 77 |
| # | interaction | 0 | 0 | 4 | 0 | 0 | # | 20 | Small | aldex2 | 77 |
| # | disease | 7 | 7 | 0 | 0.5 | 1 | # | 20 | Baseline | paralle | 77 |
| # | sex | 4 | 5 | 0 | 0.444444444 | 1 | # | 20 | Baseline | paralle | 77 |
| # | time | 5 | 6 | 0 | 0.454545455 | 1 | # | 20 | Baseline | paralle | 77 |
| # | interaction | 4 | 4 | 0 | 0.5 | 1 | # | 20 | Baseline | paralle | 77 |
| # | disease | 7 | 4 | 0 | 0.636363636 | 1 | # | 20 | Baseline | ancom | 77 |
| # | sex | 4 | 0 | 0 | 1 | 1 | # | 20 | Baseline | ancom | 77 |
| # | time | 5 | 3 | 0 | 0.625 | 1 | # | 20 | Baseline | ancom | 77 |
| # | interaction | 0 | 0 | 4 | 0 | 0 | # | 20 | Baseline | ancom | 77 |
| # | disease | 7 | 40 | 0 | 0.14893617 | 1 | # | 20 | Baseline | aldex2 | 77 |
| # | sex | 4 | 8 | 0 | 0.333333333 | 1 | # | 20 | Baseline | aldex2 | 77 |
| # | time | 5 | 26 | 0 | 0.161290323 | 1 | # | 20 | Baseline | aldex2 | 77 |
| # | interaction | 0 | 0 | 4 | 0 | 0 | # | 20 | Baseline | aldex2 | 77 |
| # | disease | 7 | 0 | 0 | 1 | 1 | # | 20 | Small | paralle | 77 |
| # | sex | 4 | 0 | 0 | 1 | 1 | # | 20 | Small | paralle | 77 |
| # | time | 5 | 0 | 0 | 1 | 1 | # | 20 | Small | paralle | 77 |
| # | interaction | 4 | 0 | 0 | 1 | 1 | # | 20 | Small | paralle | 77 |
| # | disease | 7 | 3 | 0 | 0.7 | 1 | # | 20 | Small | ancom | 77 |
| # | sex | 4 | 0 | 0 | 1 | 1 | # | 20 | Small | ancom | 77 |
| # | time | 5 | 3 | 0 | 0.625 | 1 | # | 20 | Small | ancom | 77 |
| # | interaction | 0 | 0 | 4 | 0 | 0 | # | 20 | Small | ancom | 77 |
| # | disease | 7 | 3 | 0 | 0.7 | 1 | # | 20 | Small | aldex2 | 77 |
| # | sex | 4 | 0 | 0 | 1 | 1 | # | 20 | Small | aldex2 | 77 |
| # | time | 5 | 3 | 0 | 0.625 | 1 | # | 20 | Small | aldex2 | 77 |
| # | interaction | 0 | 0 | 4 | 0 | 0 | # | 20 | Small | aldex2 | 77 |
| # | disease | 7 | 14 | 0 | 0.333333333 | 1 | # | 20 | Baseline | paralle | 77 |
| # | sex | 4 | 6 | 0 | 0.4 | 1 | # | 20 | Baseline | paralle | 77 |
| # | time | 5 | 9 | 0 | 0.357142857 | 1 | # | 20 | Baseline | paralle | 77 |
| # | interaction | 4 | 6 | 0 | 0.4 | 1 | # | 20 | Baseline | paralle | 77 |
| # | disease | 7 | 4 | 0 | 0.636363636 | 1 | # | 20 | Baseline | ancom | 77 |
| # | sex | 4 | 0 | 0 | 1 | 1 | # | 20 | Baseline | ancom | 77 |
| # | time | 5 | 3 | 0 | 0.625 | 1 | # | 20 | Baseline | ancom | 77 |
| # | interaction | 0 | 0 | 4 | 0 | 0 | # | 20 | Baseline | ancom | 77 |
| # | disease | 7 | 55 | 0 | 0.112903226 | 1 | # | 20 | Baseline | aldex2 | 77 |
| # | sex | 4 | 15 | 0 | 0.210526316 | 1 | # | 20 | Baseline | aldex2 | 77 |
| # | time | 5 | 35 | 0 | 0.125 | 1 | # | 20 | Baseline | aldex2 | 77 |
| # | interaction | 0 | 0 | 4 | 0 | 0 | # | 20 | Baseline | aldex2 | 77 |
| # | disease | 7 | 0 | 0 | 1 | 1 | # | 20 | Small | paralle | 77 |
| # | sex | 4 | 0 | 0 | 1 | 1 | # | 20 | Small | paralle | 77 |
| # | time | 5 | 0 | 0 | 1 | 1 | # | 20 | Small | paralle | 77 |
| # | interaction | 4 | 0 | 0 | 1 | 1 | # | 20 | Small | paralle | 77 |
| # | disease | 7 | 4 | 0 | 0.636363636 | 1 | # | 20 | Small | ancom | 77 |
| # | sex | 4 | 0 | 0 | 1 | 1 | # | 20 | Small | ancom | 77 |
| # | time | 5 | 3 | 0 | 0.625 | 1 | # | 20 | Small | ancom | 77 |
| # | interaction | 0 | 0 | 4 | 0 | 0 | # | 20 | Small | ancom | 77 |
| # | disease | 7 | 8 | 0 | 0.466666667 | 1 | # | 20 | Small | aldex2 | 77 |
| # | sex | 4 | 1 | 0 | 0.8 | 1 | # | 20 | Small | aldex2 | 77 |
| # | time | 5 | 5 | 0 | 0.5 | 1 | # | 20 | Small | aldex2 | 77 |
| # | interaction | 0 | 0 | 4 | 0 | 0 | # | 20 | Small | aldex2 | 77 |
| # | disease | 7 | 9 | 0 | 0.4375 | 1 | # | 20 | Baseline | paralle | 77 |
| # | sex | 4 | 7 | 0 | 0.363636364 | 1 | # | 20 | Baseline | paralle | 77 |
| # | time | 5 | 8 | 0 | 0.384615385 | 1 | # | 20 | Baseline | paralle | 77 |
| # | interaction | 4 | 5 | 0 | 0.444444444 | 1 | # | 20 | Baseline | paralle | 77 |
| # | disease | 7 | 4 | 0 | 0.636363636 | 1 | # | 20 | Baseline | ancom | 77 |
| # | sex | 4 | 0 | 0 | 1 | 1 | # | 20 | Baseline | ancom | 77 |
| # | time | 5 | 4 | 0 | 0.555555556 | 1 | # | 20 | Baseline | ancom | 77 |
| # | interaction | 0 | 0 | 4 | 0 | 0 | # | 20 | Baseline | ancom | 77 |
| # | disease | 7 | 57 | 0 | 0.094594595 | 1 | # | 20 | Baseline | aldex2 | 77 |
| # | sex | 4 | 20 | 0 | 0.166666667 | 1 | # | 20 | Baseline | aldex2 | 77 |
| # | time | 5 | 46 | 0 | 0.098039216 | 1 | # | 20 | Baseline | aldex2 | 77 |
| # | interaction | 0 | 0 | 4 | 0 | 0 | # | 20 | Baseline | aldex2 | 77 |
| # | disease | 7 | 0 | 0 | 1 | 1 | # | 20 | Small | paralle | 77 |
| # | sex | 4 | 0 | 0 | 1 | 1 | # | 20 | Small | paralle | 77 |
| # | time | 5 | 0 | 0 | 1 | 1 | # | 20 | Small | paralle | 77 |
| # | interaction | 4 | 0 | 0 | 1 | 1 | # | 20 | Small | paralle | 77 |
| # | disease | 7 | 4 | 0 | 0.636363636 | 1 | # | 20 | Small | ancom | 77 |
| # | sex | 4 | 0 | 0 | 1 | 1 | # | 20 | Small | ancom | 77 |
| # | time | 5 | 4 | 0 | 0.555555556 | 1 | # | 20 | Small | ancom | 77 |
| # | interaction | 0 | 0 | 4 | 0 | 0 | # | 20 | Small | ancom | 77 |
| # | disease | 7 | 12 | 0 | 0.368421053 | 1 | # | 20 | Small | aldex2 | 77 |
| # | sex | 4 | 1 | 0 | 0.8 | 1 | # | 20 | Small | aldex2 | 77 |
| # | time | 5 | 8 | 0 | 0.384615385 | 1 | # | 20 | Small | aldex2 | 77 |
| # | interaction | 0 | 0 | 4 | 0 | 0 | # | 20 | Small | aldex2 | 77 |
| # | disease | 4 | 4 | 0 | 0.5 | 1 | # | 10 | Baseline | paralle | 78 |
| # | sex | 2 | 10 | 0 | 0.166666667 | 1 | # | 10 | Baseline | paralle | 78 |
| # | time | 2 | 6 | 0 | 0.25 | 1 | # | 10 | Baseline | paralle | 78 |
| # | interaction | 2 | 11 | 0 | 0.153846154 | 1 | # | 10 | Baseline | paralle | 78 |
| # | disease | 4 | 1 | 0 | 0.8 | 1 | # | 10 | Baseline | ancom | 78 |
| # | sex | 2 | 0 | 0 | 1 | 1 | # | 10 | Baseline | ancom | 78 |
| # | time | 2 | 1 | 0 | 0.666666667 | 1 | # | 10 | Baseline | ancom | 78 |
| # | interaction | 0 | 0 | 2 | 0 | 0 | # | 10 | Baseline | ancom | 78 |
| # | disease | 4 | 8 | 0 | 0.333333333 | 1 | # | 10 | Baseline | aldex2 | 78 |
| # | sex | 2 | 9 | 0 | 0.181818182 | 1 | # | 10 | Baseline | aldex2 | 78 |
| # | time | 2 | 4 | 0 | 0.333333333 | 1 | # | 10 | Baseline | aldex2 | 78 |
| # | interaction | 0 | 0 | 2 | 0 | 0 | # | 10 | Baseline | aldex2 | 78 |
| # | disease | 4 | 0 | 0 | 1 | 1 | # | 10 | Small | paralle | 78 |
| # | sex | 2 | 0 | 0 | 1 | 1 | # | 10 | Small | paralle | 78 |
| # | time | 2 | 0 | 0 | 1 | 1 | # | 10 | Small | paralle | 78 |
| # | interaction | 1 | 0 | 1 | 1 | 0.5 | # | 10 | Small | paralle | 78 |
| # | disease | 4 | 1 | 0 | 0.8 | 1 | # | 10 | Small | ancom | 78 |
| # | sex | 2 | 0 | 0 | 1 | 1 | # | 10 | Small | ancom | 78 |
| # | time | 2 | 1 | 0 | 0.666666667 | 1 | # | 10 | Small | ancom | 78 |
| # | interaction | 0 | 0 | 2 | 0 | 0 | # | 10 | Small | ancom | 78 |
| # | disease | 4 | 1 | 0 | 0.8 | 1 | # | 10 | Small | aldex2 | 78 |
| # | sex | 2 | 1 | 0 | 0.666666667 | 1 | # | 10 | Small | aldex2 | 78 |
| # | time | 2 | 1 | 0 | 0.666666667 | 1 | # | 10 | Small | aldex2 | 78 |
| # | interaction | 0 | 0 | 2 | 0 | 0 | # | 10 | Small | aldex2 | 78 |
| # | disease | 4 | 3 | 0 | 0.571428571 | 1 | # | 10 | Baseline | paralle | 78 |
| # | sex | 2 | 7 | 0 | 0.222222222 | 1 | # | 10 | Baseline | paralle | 78 |
| # | time | 2 | 6 | 0 | 0.25 | 1 | # | 10 | Baseline | paralle | 78 |
| # | interaction | 2 | 7 | 0 | 0.222222222 | 1 | # | 10 | Baseline | paralle | 78 |
| # | disease | 4 | 2 | 0 | 0.666666667 | 1 | # | 10 | Baseline | ancom | 78 |
| # | sex | 2 | 0 | 0 | 1 | 1 | # | 10 | Baseline | ancom | 78 |
| # | time | 2 | 2 | 0 | 0.5 | 1 | # | 10 | Baseline | ancom | 78 |
| # | interaction | 0 | 0 | 2 | 0 | 0 | # | 10 | Baseline | ancom | 78 |
| # | disease | 4 | 19 | 0 | 0.173913043 | 1 | # | 10 | Baseline | aldex2 | 78 |
| # | sex | 2 | 9 | 0 | 0.181818182 | 1 | # | 10 | Baseline | aldex2 | 78 |
| # | time | 2 | 11 | 0 | 0.153846154 | 1 | # | 10 | Baseline | aldex2 | 78 |
| # | interaction | 0 | 0 | 2 | 0 | 0 | # | 10 | Baseline | aldex2 | 78 |
| # | disease | 4 | 0 | 0 | 1 | 1 | # | 10 | Small | paralle | 78 |
| # | sex | 2 | 0 | 0 | 1 | 1 | # | 10 | Small | paralle | 78 |
| # | time | 2 | 0 | 0 | 1 | 1 | # | 10 | Small | paralle | 78 |
| # | interaction | 2 | 0 | 0 | 1 | 1 | # | 10 | Small | paralle | 78 |
| # | disease | 4 | 2 | 0 | 0.666666667 | 1 | # | 10 | Small | ancom | 78 |

| | | | | | | | | | | | |
|---|---|---|---|---|---|---|---|---|---|---|---|
| # | sex | 2 | 0 | 0 | 1 | 1 | # | 10 | Small | ancom | 78 |
| # | time | 2 | 2 | 0 | 0.5 | 1 | # | 10 | Small | ancom | 78 |
| # | interaction | 0 | 0 | 2 | 0 | 0 | # | 10 | Small | ancom | 78 |
| # | disease | 4 | 3 | 0 | 0.571428571 | 1 | # | 10 | Small | aldex2 | 78 |
| # | sex | 2 | 0 | 0 | 1 | 1 | # | 10 | Small | aldex2 | 78 |
| # | time | 2 | 2 | 0 | 0.5 | 1 | # | 10 | Small | aldex2 | 78 |
| # | interaction | 0 | 0 | 2 | 0 | 0 | # | 10 | Small | aldex2 | 78 |
| # | disease | 4 | 3 | 0 | 0.571428571 | 1 | # | 10 | Baseline | paralle | 78 |
| # | sex | 2 | 2 | 0 | 0.5 | 1 | # | 10 | Baseline | paralle | 78 |
| # | time | 2 | 6 | 0 | 0.25 | 1 | # | 10 | Baseline | paralle | 78 |
| # | interaction | 2 | 2 | 0 | 0.5 | 1 | # | 10 | Baseline | paralle | 78 |
| # | disease | 4 | 2 | 0 | 0.666666667 | 1 | # | 10 | Baseline | ancom | 78 |
| # | sex | 2 | 0 | 0 | 1 | 1 | # | 10 | Baseline | ancom | 78 |
| # | time | 2 | 2 | 0 | 0.5 | 1 | # | 10 | Baseline | ancom | 78 |
| # | interaction | 0 | 0 | 2 | 0 | 0 | # | 10 | Baseline | ancom | 78 |
| # | disease | 4 | 26 | 0 | 0.133333333 | 1 | # | 10 | Baseline | aldex2 | 78 |
| # | sex | 2 | 4 | 0 | 0.333333333 | 1 | # | 10 | Baseline | aldex2 | 78 |
| # | time | 2 | 10 | 0 | 0.166666667 | 1 | # | 10 | Baseline | aldex2 | 78 |
| # | interaction | 0 | 0 | 2 | 0 | 0 | # | 10 | Baseline | aldex2 | 78 |
| # | disease | 4 | 0 | 0 | 1 | 1 | # | 10 | Small | paralle | 78 |
| # | sex | 2 | 0 | 0 | 1 | 1 | # | 10 | Small | paralle | 78 |
| # | time | 2 | 0 | 0 | 1 | 1 | # | 10 | Small | paralle | 78 |
| # | interaction | 2 | 0 | 0 | 1 | 1 | # | 10 | Small | paralle | 78 |
| # | disease | 4 | 2 | 0 | 0.666666667 | 1 | # | 10 | Small | ancom | 78 |
| # | sex | 2 | 0 | 0 | 1 | 1 | # | 10 | Small | ancom | 78 |
| # | time | 2 | 2 | 0 | 0.5 | 1 | # | 10 | Small | ancom | 78 |
| # | interaction | 0 | 0 | 2 | 0 | 0 | # | 10 | Small | ancom | 78 |
| # | disease | 4 | 2 | 0 | 0.666666667 | 1 | # | 10 | Small | aldex2 | 78 |
| # | sex | 2 | 0 | 0 | 1 | 1 | # | 10 | Small | aldex2 | 78 |
| # | time | 2 | 2 | 0 | 0.5 | 1 | # | 10 | Small | aldex2 | 78 |
| # | interaction | 0 | 0 | 2 | 0 | 0 | # | 10 | Small | aldex2 | 78 |
| # | disease | 4 | 5 | 0 | 0.444444444 | 1 | # | 10 | Baseline | paralle | 78 |
| # | sex | 2 | 1 | 0 | 0.666666667 | 1 | # | 10 | Baseline | paralle | 78 |
| # | time | 2 | 3 | 0 | 0.4 | 1 | # | 10 | Baseline | paralle | 78 |
| # | interaction | 2 | 4 | 0 | 0.333333333 | 1 | # | 10 | Baseline | paralle | 78 |
| # | disease | 4 | 2 | 0 | 0.666666667 | 1 | # | 10 | Baseline | ancom | 78 |
| # | sex | 2 | 0 | 0 | 1 | 1 | # | 10 | Baseline | ancom | 78 |
| # | time | 2 | 2 | 0 | 0.5 | 1 | # | 10 | Baseline | ancom | 78 |
| # | interaction | 0 | 0 | 2 | 0 | 0 | # | 10 | Baseline | ancom | 78 |
| # | disease | 4 | 29 | 0 | 0.121212121 | 1 | # | 10 | Baseline | aldex2 | 78 |
| # | sex | 2 | 11 | 0 | 0.153846154 | 1 | # | 10 | Baseline | aldex2 | 78 |
| # | time | 2 | 14 | 0 | 0.125 | 1 | # | 10 | Baseline | aldex2 | 78 |
| # | interaction | 0 | 0 | 2 | 0 | 0 | # | 10 | Baseline | aldex2 | 78 |
| # | disease | 4 | 0 | 0 | 1 | 1 | # | 10 | Small | paralle | 78 |
| # | sex | 2 | 0 | 0 | 1 | 1 | # | 10 | Small | paralle | 78 |
| # | time | 2 | 0 | 0 | 1 | 1 | # | 10 | Small | paralle | 78 |
| # | interaction | 2 | 0 | 0 | 1 | 1 | # | 10 | Small | paralle | 78 |
| # | disease | 4 | 2 | 0 | 0.666666667 | 1 | # | 10 | Small | ancom | 78 |
| # | sex | 2 | 0 | 0 | 1 | 1 | # | 10 | Small | ancom | 78 |
| # | time | 2 | 2 | 0 | 0.5 | 1 | # | 10 | Small | ancom | 78 |
| # | interaction | 0 | 0 | 2 | 0 | 0 | # | 10 | Small | ancom | 78 |
| # | disease | 4 | 2 | 0 | 0.666666667 | 1 | # | 10 | Small | aldex2 | 78 |
| # | sex | 2 | 0 | 0 | 1 | 1 | # | 10 | Small | aldex2 | 78 |
| # | time | 2 | 2 | 0 | 0.5 | 1 | # | 10 | Small | aldex2 | 78 |
| # | interaction | 0 | 0 | 2 | 0 | 0 | # | 10 | Small | aldex2 | 78 |
| # | disease | 7 | 10 | 0 | 0.411764706 | 1 | # | 20 | Baseline | paralle | 78 |
| # | sex | 4 | 11 | 0 | 0.266666667 | 1 | # | 20 | Baseline | paralle | 78 |
| # | time | 5 | 8 | 0 | 0.384615385 | 1 | # | 20 | Baseline | paralle | 78 |
| # | interaction | 4 | 9 | 0 | 0.307692308 | 1 | # | 20 | Baseline | paralle | 78 |
| # | disease | 7 | 3 | 0 | 0.7 | 1 | # | 20 | Baseline | ancom | 78 |
| # | sex | 4 | 0 | 0 | 1 | 1 | # | 20 | Baseline | ancom | 78 |
| # | time | 5 | 2 | 0 | 0.714285714 | 1 | # | 20 | Baseline | ancom | 78 |
| # | interaction | 0 | 0 | 4 | 0 | 0 | # | 20 | Baseline | ancom | 78 |
| # | disease | 7 | 18 | 0 | 0.28 | 1 | # | 20 | Baseline | aldex2 | 78 |
| # | sex | 4 | 9 | 0 | 0.307692308 | 1 | # | 20 | Baseline | aldex2 | 78 |
| # | time | 5 | 17 | 0 | 0.227272727 | 1 | # | 20 | Baseline | aldex2 | 78 |
| # | interaction | 0 | 0 | 4 | 0 | 0 | # | 20 | Baseline | aldex2 | 78 |
| # | disease | 7 | 0 | 0 | 1 | 1 | # | 20 | Small | paralle | 78 |
| # | sex | 4 | 0 | 0 | 1 | 1 | # | 20 | Small | paralle | 78 |
| # | time | 5 | 0 | 0 | 1 | 1 | # | 20 | Small | paralle | 78 |
| # | interaction | 3 | 0 | 1 | 1 | 0.75 | # | 20 | Small | paralle | 78 |
| # | disease | 7 | 3 | 0 | 0.7 | 1 | # | 20 | Small | ancom | 78 |
| # | sex | 4 | 0 | 0 | 1 | 1 | # | 20 | Small | ancom | 78 |
| # | time | 5 | 1 | 0 | 0.833333333 | 1 | # | 20 | Small | ancom | 78 |
| # | interaction | 0 | 0 | 4 | 0 | 0 | # | 20 | Small | ancom | 78 |
| # | disease | 7 | 1 | 0 | 0.875 | 1 | # | 20 | Small | aldex2 | 78 |
| # | sex | 4 | 1 | 0 | 0.8 | 1 | # | 20 | Small | aldex2 | 78 |
| # | time | 5 | 0 | 0 | 1 | 1 | # | 20 | Small | aldex2 | 78 |
| # | interaction | 0 | 0 | 4 | 0 | 0 | # | 20 | Small | aldex2 | 78 |
| # | disease | 7 | 7 | 0 | 0.5 | 1 | # | 20 | Baseline | paralle | 78 |
| # | sex | 4 | 5 | 0 | 0.444444444 | 1 | # | 20 | Baseline | paralle | 78 |
| # | time | 5 | 7 | 0 | 0.416666667 | 1 | # | 20 | Baseline | paralle | 78 |
| # | interaction | 4 | 7 | 0 | 0.363636364 | 1 | # | 20 | Baseline | paralle | 78 |
| # | disease | 7 | 4 | 0 | 0.636363636 | 1 | # | 20 | Baseline | ancom | 78 |
| # | sex | 4 | 0 | 0 | 1 | 1 | # | 20 | Baseline | ancom | 78 |
| # | time | 5 | 4 | 0 | 0.555555556 | 1 | # | 20 | Baseline | ancom | 78 |
| # | interaction | 0 | 0 | 4 | 0 | 0 | # | 20 | Baseline | ancom | 78 |
| # | disease | 7 | 36 | 0 | 0.162790698 | 1 | # | 20 | Baseline | aldex2 | 78 |
| # | sex | 4 | 13 | 0 | 0.235294118 | 1 | # | 20 | Baseline | aldex2 | 78 |
| # | time | 5 | 27 | 0 | 0.15625 | 1 | # | 20 | Baseline | aldex2 | 78 |
| # | interaction | 0 | 0 | 4 | 0 | 0 | # | 20 | Baseline | aldex2 | 78 |
| # | disease | 7 | 0 | 0 | 1 | 1 | # | 20 | Small | paralle | 78 |
| # | sex | 4 | 0 | 0 | 1 | 1 | # | 20 | Small | paralle | 78 |
| # | time | 5 | 0 | 0 | 1 | 1 | # | 20 | Small | paralle | 78 |
| # | interaction | 3 | 0 | 1 | 1 | 0.75 | # | 20 | Small | paralle | 78 |
| # | disease | 7 | 4 | 0 | 0.636363636 | 1 | # | 20 | Small | ancom | 78 |
| # | sex | 4 | 0 | 0 | 1 | 1 | # | 20 | Small | ancom | 78 |
| # | time | 5 | 4 | 0 | 0.555555556 | 1 | # | 20 | Small | ancom | 78 |
| # | interaction | 0 | 0 | 4 | 0 | 0 | # | 20 | Small | ancom | 78 |
| # | disease | 7 | 3 | 0 | 0.7 | 1 | # | 20 | Small | aldex2 | 78 |
| # | sex | 4 | 1 | 0 | 0.8 | 1 | # | 20 | Small | aldex2 | 78 |
| # | time | 5 | 5 | 0 | 0.5 | 1 | # | 20 | Small | aldex2 | 78 |
| # | interaction | 0 | 0 | 4 | 0 | 0 | # | 20 | Small | aldex2 | 78 |
| # | disease | 7 | 10 | 0 | 0.411764706 | 1 | # | 20 | Baseline | paralle | 78 |
| # | sex | 4 | 8 | 0 | 0.333333333 | 1 | # | 20 | Baseline | paralle | 78 |
| # | time | 5 | 9 | 0 | 0.357142857 | 1 | # | 20 | Baseline | paralle | 78 |
| # | interaction | 4 | 4 | 0 | 0.5 | 1 | # | 20 | Baseline | paralle | 78 |
| # | disease | 7 | 4 | 0 | 0.636363636 | 1 | # | 20 | Baseline | ancom | 78 |
| # | sex | 4 | 0 | 0 | 1 | 1 | # | 20 | Baseline | ancom | 78 |
| # | time | 5 | 4 | 0 | 0.555555556 | 1 | # | 20 | Baseline | ancom | 78 |
| # | interaction | 0 | 0 | 4 | 0 | 0 | # | 20 | Baseline | ancom | 78 |
| # | disease | 7 | 56 | 0 | 0.111111111 | 1 | # | 20 | Baseline | aldex2 | 78 |
| # | sex | 4 | 20 | 0 | 0.166666667 | 1 | # | 20 | Baseline | aldex2 | 78 |
| # | time | 5 | 38 | 0 | 0.11627907 | 1 | # | 20 | Baseline | aldex2 | 78 |
| # | interaction | 0 | 0 | 4 | 0 | 0 | # | 20 | Baseline | aldex2 | 78 |
| # | disease | 7 | 0 | 0 | 1 | 1 | # | 20 | Small | paralle | 78 |
| # | sex | 4 | 0 | 0 | 1 | 1 | # | 20 | Small | paralle | 78 |
| # | time | 5 | 0 | 0 | 1 | 1 | # | 20 | Small | paralle | 78 |
| # | interaction | 4 | 0 | 0 | 1 | 1 | # | 20 | Small | paralle | 78 |
| # | disease | 7 | 4 | 0 | 0.636363636 | 1 | # | 20 | Small | ancom | 78 |
| # | sex | 4 | 0 | 0 | 1 | 1 | # | 20 | Small | ancom | 78 |
| # | time | 5 | 4 | 0 | 0.555555556 | 1 | # | 20 | Small | ancom | 78 |
| # | interaction | 0 | 0 | 4 | 0 | 0 | # | 20 | Small | ancom | 78 |
| # | disease | 7 | 7 | 0 | 0.5 | 1 | # | 20 | Small | aldex2 | 78 |
| # | sex | 4 | 1 | 0 | 0.8 | 1 | # | 20 | Small | aldex2 | 78 |
| # | time | 5 | 5 | 0 | 0.5 | 1 | # | 20 | Small | aldex2 | 78 |
| # | interaction | 0 | 0 | 4 | 0 | 0 | # | 20 | Small | aldex2 | 78 |
| # | disease | 7 | 9 | 0 | 0.4375 | 1 | # | 20 | Baseline | paralle | 78 |
| # | sex | 4 | 7 | 0 | 0.363636364 | 1 | # | 20 | Baseline | paralle | 78 |
| # | time | 5 | 8 | 0 | 0.384615385 | 1 | # | 20 | Baseline | paralle | 78 |
| # | interaction | 4 | 3 | 0 | 0.571428571 | 1 | # | 20 | Baseline | paralle | 78 |
| # | disease | 7 | 4 | 0 | 0.636363636 | 1 | # | 20 | Baseline | ancom | 78 |
| # | sex | 4 | 0 | 0 | 1 | 1 | # | 20 | Baseline | ancom | 78 |
| # | time | 5 | 4 | 0 | 0.555555556 | 1 | # | 20 | Baseline | ancom | 78 |
| # | interaction | 0 | 0 | 4 | 0 | 0 | # | 20 | Baseline | ancom | 78 |
| # | disease | 7 | 89 | 0 | 0.092105263 | 1 | # | 20 | Baseline | aldex2 | 78 |
| # | sex | 4 | 30 | 0 | 0.117647059 | 1 | # | 20 | Baseline | aldex2 | 78 |
| # | time | 5 | 66 | 0 | 0.098039216 | 1 | # | 20 | Baseline | aldex2 | 78 |
| # | interaction | 0 | 0 | 4 | 0 | 0 | # | 20 | Baseline | aldex2 | 78 |
| # | disease | 7 | 0 | 0 | 1 | 1 | # | 20 | Small | paralle | 78 |
| # | sex | 4 | 0 | 0 | 1 | 1 | # | 20 | Small | paralle | 78 |
| # | time | 5 | 0 | 0 | 1 | 1 | # | 20 | Small | paralle | 78 |
| # | interaction | 4 | 0 | 0 | 1 | 1 | # | 20 | Small | paralle | 78 |
| # | disease | 7 | 4 | 0 | 0.636363636 | 1 | # | 20 | Small | ancom | 78 |
| # | sex | 4 | 0 | 0 | 1 | 1 | # | 20 | Small | ancom | 78 |
| # | time | 5 | 4 | 0 | 0.555555556 | 1 | # | 20 | Small | ancom | 78 |
| # | interaction | 0 | 0 | 4 | 0 | 0 | # | 20 | Small | ancom | 78 |
| # | disease | 7 | 9 | 0 | 0.4375 | 1 | # | 20 | Small | aldex2 | 78 |
| # | sex | 4 | 2 | 0 | 0.666666667 | 1 | # | 20 | Small | aldex2 | 78 |
| # | time | 5 | 6 | 0 | 0.454545455 | 1 | # | 20 | Small | aldex2 | 78 |
| # | interaction | 0 | 0 | 4 | 0 | 0 | # | 20 | Small | aldex2 | 78 |
| # | disease | 4 | 9 | 0 | 0.307692308 | 1 | # | 10 | Baseline | paralle | 79 |
| # | sex | 2 | 6 | 0 | 0.25 | 1 | # | 10 | Baseline | paralle | 79 |
| # | time | 2 | 4 | 0 | 0.333333333 | 1 | # | 10 | Baseline | paralle | 79 |
| # | interaction | 2 | 6 | 0 | 0.25 | 1 | # | 10 | Baseline | paralle | 79 |
| # | disease | 4 | 0 | 0 | 1 | 1 | # | 10 | Baseline | ancom | 79 |
| # | sex | 2 | 0 | 0 | 1 | 1 | # | 10 | Baseline | ancom | 79 |
| # | time | 2 | 2 | 0 | 0.5 | 1 | # | 10 | Baseline | ancom | 79 |
| # | interaction | 0 | 0 | 2 | 0 | 0 | # | 10 | Baseline | ancom | 79 |
| # | disease | 4 | 12 | 0 | 0.25 | 1 | # | 10 | Baseline | aldex2 | 79 |
| # | sex | 2 | 4 | 0 | 0.333333333 | 1 | # | 10 | Baseline | aldex2 | 79 |
| # | time | 2 | 7 | 0 | 0.222222222 | 1 | # | 10 | Baseline | aldex2 | 79 |
| # | interaction | 0 | 0 | 2 | 0 | 0 | # | 10 | Baseline | aldex2 | 79 |
| # | disease | 4 | 0 | 0 | 1 | 1 | # | 10 | Small | paralle | 79 |
| # | sex | 2 | 0 | 0 | 1 | 1 | # | 10 | Small | paralle | 79 |
| # | time | 2 | 0 | 0 | 1 | 1 | # | 10 | Small | paralle | 79 |
| # | interaction | 1 | 0 | 1 | 1 | 0.5 | # | 10 | Small | paralle | 79 |
| # | disease | 4 | 0 | 0 | 1 | 1 | # | 10 | Small | ancom | 79 |
| # | sex | 2 | 0 | 0 | 1 | 1 | # | 10 | Small | ancom | 79 |
| # | time | 2 | 0 | 0 | 1 | 1 | # | 10 | Small | ancom | 79 |
| # | interaction | 0 | 0 | 2 | 0 | 0 | # | 10 | Small | ancom | 79 |
| # | disease | 4 | 0 | 0 | 1 | 1 | # | 10 | Small | aldex2 | 79 |
| # | sex | 2 | 0 | 0 | 1 | 1 | # | 10 | Small | aldex2 | 79 |
| # | time | 2 | 1 | 0 | 0.666666667 | 1 | # | 10 | Small | aldex2 | 79 |
| # | interaction | 0 | 0 | 2 | 0 | 0 | # | 10 | Small | aldex2 | 79 |
| # | disease | 4 | 4 | 0 | 0.5 | 1 | # | 10 | Baseline | paralle | 79 |
| # | sex | 2 | 3 | 0 | 0.4 | 1 | # | 10 | Baseline | paralle | 79 |
| # | time | 2 | 6 | 0 | 0.25 | 1 | # | 10 | Baseline | paralle | 79 |
| # | interaction | 2 | 6 | 0 | 0.25 | 1 | # | 10 | Baseline | paralle | 79 |
| # | disease | 4 | 1 | 0 | 0.8 | 1 | # | 10 | Baseline | ancom | 79 |
| # | sex | 2 | 0 | 0 | 1 | 1 | # | 10 | Baseline | ancom | 79 |
| # | time | 2 | 2 | 0 | 0.5 | 1 | # | 10 | Baseline | ancom | 79 |
| # | interaction | 0 | 0 | 2 | 0 | 0 | # | 10 | Baseline | ancom | 79 |
| # | disease | 4 | 18 | 0 | 0.181818182 | 1 | # | 10 | Baseline | aldex2 | 79 |
| # | sex | 2 | 4 | 0 | 0.333333333 | 1 | # | 10 | Baseline | aldex2 | 79 |
| # | time | 2 | 8 | 0 | 0.2 | 1 | # | 10 | Baseline | aldex2 | 79 |
| # | interaction | 0 | 0 | 2 | 0 | 0 | # | 10 | Baseline | aldex2 | 79 |
| # | disease | 4 | 0 | 0 | 1 | 1 | # | 10 | Small | paralle | 79 |
| # | sex | 2 | 0 | 0 | 1 | 1 | # | 10 | Small | paralle | 79 |
| # | time | 2 | 0 | 0 | 1 | 1 | # | 10 | Small | paralle | 79 |
| # | interaction | 2 | 0 | 0 | 1 | 1 | # | 10 | Small | paralle | 79 |
| # | disease | 4 | 1 | 0 | 0.8 | 1 | # | 10 | Small | ancom | 79 |
| # | sex | 2 | 0 | 0 | 1 | 1 | # | 10 | Small | ancom | 79 |
| # | time | 2 | 2 | 0 | 0.5 | 1 | # | 10 | Small | ancom | 79 |
| # | interaction | 0 | 0 | 2 | 0 | 0 | # | 10 | Small | ancom | 79 |
| # | disease | 4 | 1 | 0 | 0.8 | 1 | # | 10 | Small | aldex2 | 79 |
| # | sex | 2 | 0 | 0 | 1 | 1 | # | 10 | Small | aldex2 | 79 |
| # | time | 2 | 1 | 0 | 0.666666667 | 1 | # | 10 | Small | aldex2 | 79 |
| # | interaction | 0 | 0 | 2 | 0 | 0 | # | 10 | Small | aldex2 | 79 |
| # | disease | 4 | 5 | 0 | 0.444444444 | 1 | # | 10 | Baseline | paralle | 79 |
| # | sex | 2 | 2 | 0 | 0.5 | 1 | # | 10 | Baseline | paralle | 79 |
| # | time | 2 | 4 | 0 | 0.333333333 | 1 | # | 10 | Baseline | paralle | 79 |
| # | interaction | 2 | 2 | 0 | 0.5 | 1 | # | 10 | Baseline | paralle | 79 |
| # | disease | 4 | 2 | 0 | 0.666666667 | 1 | # | 10 | Baseline | ancom | 79 |
| # | sex | 2 | 0 | 0 | 1 | 1 | # | 10 | Baseline | ancom | 79 |
| # | time | 2 | 2 | 0 | 0.5 | 1 | # | 10 | Baseline | ancom | 79 |
| # | interaction | 0 | 0 | 2 | 0 | 0 | # | 10 | Baseline | ancom | 79 |
| # | disease | 4 | 24 | 0 | 0.142857143 | 1 | # | 10 | Baseline | aldex2 | 79 |
| # | sex | 2 | 8 | 0 | 0.2 | 1 | # | 10 | Baseline | aldex2 | 79 |
| # | time | 2 | 7 | 0 | 0.222222222 | 1 | # | 10 | Baseline | aldex2 | 79 |
| # | interaction | 0 | 0 | 2 | 0 | 0 | # | 10 | Baseline | aldex2 | 79 |
| # | disease | 4 | 0 | 0 | 1 | 1 | # | 10 | Small | paralle | 79 |
| # | sex | 2 | 0 | 0 | 1 | 1 | # | 10 | Small | paralle | 79 |
| # | time | 2 | 0 | 0 | 1 | 1 | # | 10 | Small | paralle | 79 |
| # | interaction | 2 | 0 | 0 | 1 | 1 | # | 10 | Small | paralle | 79 |
| # | disease | 4 | 2 | 0 | 0.666666667 | 1 | # | 10 | Small | ancom | 79 |
| # | sex | 2 | 0 | 0 | 1 | 1 | # | 10 | Small | ancom | 79 |
| # | time | 2 | 2 | 0 | 0.5 | 1 | # | 10 | Small | ancom | 79 |
| # | interaction | 0 | 0 | 2 | 0 | 0 | # | 10 | Small | ancom | 79 |
| # | disease | 4 | 4 | 0 | 0.5 | 1 | # | 10 | Small | aldex2 | 79 |
| # | sex | 2 | 0 | 0 | 1 | 1 | # | 10 | Small | aldex2 | 79 |
| # | time | 2 | 2 | 0 | 0.5 | 1 | # | 10 | Small | aldex2 | 79 |
| # | interaction | 0 | 0 | 2 | 0 | 0 | # | 10 | Small | aldex2 | 79 |
| # | disease | 4 | 3 | 0 | 0.571428571 | 1 | # | 10 | Baseline | paralle | 79 |
| # | sex | 2 | 4 | 0 | 0.333333333 | 1 | # | 10 | Baseline | paralle | 79 |
| # | time | 2 | 5 | 0 | 0.285714286 | 1 | # | 10 | Baseline | paralle | 79 |
| # | interaction | 2 | 3 | 0 | 0.4 | 1 | # | 10 | Baseline | paralle | 79 |
| # | disease | 4 | 2 | 0 | 0.666666667 | 1 | # | 10 | Baseline | ancom | 79 |
| # | sex | 2 | 0 | 0 | 1 | 1 | # | 10 | Baseline | ancom | 79 |
| # | time | 2 | 2 | 0 | 0.5 | 1 | # | 10 | Baseline | ancom | 79 |
| # | interaction | 0 | 0 | 2 | 0 | 0 | # | 10 | Baseline | ancom | 79 |
| # | disease | 4 | 35 | 0 | 0.102564103 | 1 | # | 10 | Baseline | aldex2 | 79 |
| # | sex | 2 | 9 | 0 | 0.181818182 | 1 | # | 10 | Baseline | aldex2 | 79 |
| # | time | 2 | 11 | 0 | 0.153846154 | 1 | # | 10 | Baseline | aldex2 | 79 |
| # | interaction | 0 | 0 | 2 | 0 | 0 | # | 10 | Baseline | aldex2 | 79 |
| # | disease | 4 | 0 | 0 | 1 | 1 | # | 10 | Small | paralle | 79 |
| # | sex | 2 | 0 | 0 | 1 | 1 | # | 10 | Small | paralle | 79 |
| # | time | 2 | 0 | 0 | 1 | 1 | # | 10 | Small | paralle | 79 |
| # | interaction | 2 | 0 | 0 | 1 | 1 | # | 10 | Small | paralle | 79 |
| # | disease | 4 | 2 | 0 | 0.666666667 | 1 | # | 10 | Small | ancom | 79 |
| # | sex | 2 | 0 | 0 | 1 | 1 | # | 10 | Small | ancom | 79 |
| # | time | 2 | 2 | 0 | 0.5 | 1 | # | 10 | Small | ancom | 79 |
| # | interaction | 0 | 0 | 2 | 0 | 0 | # | 10 | Small | ancom | 79 |
| # | disease | 4 | 5 | 0 | 0.444444444 | 1 | # | 10 | Small | aldex2 | 79 |
| # | sex | 2 | 0 | 0 | 1 | 1 | # | 10 | Small | aldex2 | 79 |
| # | time | 2 | 2 | 0 | 0.5 | 1 | # | 10 | Small | aldex2 | 79 |
| # | interaction | 0 | 0 | 2 | 0 | 0 | # | 10 | Small | aldex2 | 79 |
| # | disease | 7 | 7 | 0 | 0.5 | 1 | # | 20 | Baseline | paralle | 79 |
| # | sex | 4 | 7 | 0 | 0.363636364 | 1 | # | 20 | Baseline | paralle | 79 |
| # | time | 5 | 12 | 0 | 0.294117647 | 1 | # | 20 | Baseline | paralle | 79 |
| # | interaction | 4 | 10 | 0 | 0.285714286 | 1 | # | 20 | Baseline | paralle | 79 |
| # | disease | 7 | 1 | 0 | 0.875 | 1 | # | 20 | Baseline | ancom | 79 |
| # | sex | 4 | 0 | 0 | 1 | 1 | # | 20 | Baseline | ancom | 79 |
| # | time | 5 | 2 | 0 | 0.714285714 | 1 | # | 20 | Baseline | ancom | 79 |
| # | interaction | 0 | 0 | 4 | 0 | 0 | # | 20 | Baseline | ancom | 79 |
| # | disease | 7 | 23 | 0 | 0.233333333 | 1 | # | 20 | Baseline | aldex2 | 79 |
| # | sex | 4 | 9 | 0 | 0.307692308 | 1 | # | 20 | Baseline | aldex2 | 79 |
| # | time | 5 | 19 | 0 | 0.208333333 | 1 | # | 20 | Baseline | aldex2 | 79 |
| # | interaction | 0 | 0 | 4 | 0 | 0 | # | 20 | Baseline | aldex2 | 79 |
| # | disease | 7 | 0 | 0 | 1 | 1 | # | 20 | Small | paralle | 79 |
| # | sex | 4 | 0 | 0 | 1 | 1 | # | 20 | Small | paralle | 79 |
| # | time | 5 | 0 | 0 | 1 | 1 | # | 20 | Small | paralle | 79 |
| # | interaction | 3 | 0 | 1 | 1 | 0.75 | # | 20 | Small | paralle | 79 |
| # | disease | 7 | 1 | 0 | 0.875 | 1 | # | 20 | Small | ancom | 79 |
| # | sex | 4 | 0 | 0 | 1 | 1 | # | 20 | Small | ancom | 79 |
| # | time | 5 | 1 | 0 | 0.833333333 | 1 | # | 20 | Small | ancom | 79 |
| # | interaction | 0 | 0 | 4 | 0 | 0 | # | 20 | Small | ancom | 79 |
| # | disease | 7 | 2 | 0 | 0.777777778 | 1 | # | 20 | Small | aldex2 | 79 |
| # | sex | 4 | 0 | 0 | 1 | 1 | # | 20 | Small | aldex2 | 79 |
| # | time | 5 | 1 | 0 | 0.833333333 | 1 | # | 20 | Small | aldex2 | 79 |
| # | interaction | 0 | 0 | 4 | 0 | 0 | # | 20 | Small | aldex2 | 79 |
| # | disease | 7 | 5 | 0 | 0.583333333 | 1 | # | 20 | Baseline | paralle | 79 |
| # | sex | 4 | 8 | 0 | 0.333333333 | 1 | # | 20 | Baseline | paralle | 79 |
| # | time | 5 | 12 | 0 | 0.294117647 | 1 | # | 20 | Baseline | paralle | 79 |
| # | interaction | 4 | 4 | 0 | 0.5 | 1 | # | 20 | Baseline | paralle | 79 |
| # | disease | 7 | 4 | 0 | 0.636363636 | 1 | # | 20 | Baseline | ancom | 79 |
| # | sex | 4 | 0 | 0 | 1 | 1 | # | 20 | Baseline | ancom | 79 |
| # | time | 5 | 5 | 0 | 0.5 | 1 | # | 20 | Baseline | ancom | 79 |
| # | interaction | 0 | 0 | 4 | 0 | 0 | # | 20 | Baseline | ancom | 79 |
| # | disease | 7 | 37 | 0 | 0.159090909 | 1 | # | 20 | Baseline | aldex2 | 79 |
| # | sex | 4 | 14 | 0 | 0.222222222 | 1 | # | 20 | Baseline | aldex2 | 79 |
| # | time | 5 | 30 | 0 | 0.142857143 | 1 | # | 20 | Baseline | aldex2 | 79 |
| # | interaction | 0 | 0 | 4 | 0 | 0 | # | 20 | Baseline | aldex2 | 79 |
| # | disease | 7 | 0 | 0 | 1 | 1 | # | 20 | Small | paralle | 79 |
| # | sex | 4 | 0 | 0 | 1 | 1 | # | 20 | Small | paralle | 79 |
| # | time | 5 | 0 | 0 | 1 | 1 | # | 20 | Small | paralle | 79 |
| # | interaction | 3 | 0 | 1 | 1 | 0.75 | # | 20 | Small | paralle | 79 |
| # | disease | 7 | 4 | 0 | 0.636363636 | 1 | # | 20 | Small | ancom | 79 |
| # | sex | 4 | 0 | 0 | 1 | 1 | # | 20 | Small | ancom | 79 |
| # | time | 5 | 4 | 0 | 0.555555556 | 1 | # | 20 | Small | ancom | 79 |
| # | interaction | 0 | 0 | 4 | 0 | 0 | # | 20 | Small | ancom | 79 |
| # | disease | 7 | 5 | 0 | 0.583333333 | 1 | # | 20 | Small | aldex2 | 79 |
| # | sex | 4 | 0 | 0 | 1 | 1 | # | 20 | Small | aldex2 | 79 |
| # | time | 5 | 4 | 0 | 0.555555556 | 1 | # | 20 | Small | aldex2 | 79 |
| # | interaction | 0 | 0 | 4 | 0 | 0 | # | 20 | Small | aldex2 | 79 |
| # | disease | 7 | 3 | 0 | 0.7 | 1 | # | 20 | Baseline | paralle | 79 |
| # | sex | 4 | 8 | 0 | 0.333333333 | 1 | # | 20 | Baseline | paralle | 79 |
| # | time | 5 | 11 | 0 | 0.3125 | 1 | # | 20 | Baseline | paralle | 79 |
| # | interaction | 4 | 6 | 0 | 0.4 | 1 | # | 20 | Baseline | paralle | 79 |
| # | disease | 7 | 4 | 0 | 0.636363636 | 1 | # | 20 | Baseline | ancom | 79 |
| # | sex | 4 | 0 | 0 | 1 | 1 | # | 20 | Baseline | ancom | 79 |
| # | time | 5 | 4 | 0 | 0.555555556 | 1 | # | 20 | Baseline | ancom | 79 |
| # | interaction | 0 | 0 | 4 | 0 | 0 | # | 20 | Baseline | ancom | 79 |
| # | disease | 7 | 53 | 0 | 0.116666667 | 1 | # | 20 | Baseline | aldex2 | 79 |
| # | sex | 4 | 24 | 0 | 0.142857143 | 1 | # | 20 | Baseline | aldex2 | 79 |
| # | time | 5 | 35 | 0 | 0.125 | 1 | # | 20 | Baseline | aldex2 | 79 |
| # | interaction | 0 | 0 | 4 | 0 | 0 | # | 20 | Baseline | aldex2 | 79 |
| # | disease | 7 | 0 | 0 | 1 | 1 | # | 20 | Small | paralle | 79 |
| # | sex | 4 | 0 | 0 | 1 | 1 | # | 20 | Small | paralle | 79 |
| # | time | 5 | 0 | 0 | 1 | 1 | # | 20 | Small | paralle | 79 |
| # | interaction | 4 | 0 | 0 | 1 | 1 | # | 20 | Small | paralle | 79 |
| # | disease | 7 | 4 | 0 | 0.636363636 | 1 | # | 20 | Small | ancom | 79 |
| # | sex | 4 | 0 | 0 | 1 | 1 | # | 20 | Small | ancom | 79 |
| # | time | 5 | 4 | 0 | 0.555555556 | 1 | # | 20 | Small | ancom | 79 |
| # | interaction | 0 | 0 | 4 | 0 | 0 | # | 20 | Small | ancom | 79 |
| # | disease | 7 | 7 | 0 | 0.5 | 1 | # | 20 | Small | aldex2 | 79 |
| # | sex | 4 | 0 | 0 | 1 | 1 | # | 20 | Small | aldex2 | 79 |
| # | time | 5 | 8 | 0 | 0.384615385 | 1 | # | 20 | Small | aldex2 | 79 |
| # | interaction | 0 | 0 | 4 | 0 | 0 | # | 20 | Small | aldex2 | 79 |
| # | disease | 7 | 8 | 0 | 0.466666667 | 1 | # | 20 | Baseline | paralle | 79 |
| # | sex | 4 | 7 | 0 | 0.363636364 | 1 | # | 20 | Baseline | paralle | 79 |
| # | time | 5 | 9 | 0 | 0.357142857 | 1 | # | 20 | Baseline | paralle | 79 |
| # | interaction | 4 | 4 | 0 | 0.5 | 1 | # | 20 | Baseline | paralle | 79 |
| # | disease | 7 | 4 | 0 | 0.636363636 | 1 | # | 20 | Baseline | ancom | 79 |
| # | sex | 4 | 0 | 0 | 1 | 1 | # | 20 | Baseline | ancom | 79 |
| # | time | 5 | 4 | 0 | 0.555555556 | 1 | # | 20 | Baseline | ancom | 79 |
| # | interaction | 0 | 0 | 4 | 0 | 0 | # | 20 | Baseline | ancom | 79 |
| # | disease | 7 | 65 | 0 | 0.097222222 | 1 | # | 20 | Baseline | aldex2 | 79 |
| # | sex | 4 | 24 | 0 | 0.142857143 | 1 | # | 20 | Baseline | aldex2 | 79 |
| # | time | 5 | 66 | 0 | 0.098039216 | 1 | # | 20 | Baseline | aldex2 | 79 |
| # | interaction | 0 | 0 | 4 | 0 | 0 | # | 20 | Baseline | aldex2 | 79 |
| # | disease | 7 | 0 | 0 | 1 | 1 | # | 20 | Small | paralle | 79 |
| # | sex | 4 | 0 | 0 | 1 | 1 | # | 20 | Small | paralle | 79 |
| # | time | 5 | 0 | 0 | 1 | 1 | # | 20 | Small | paralle | 79 |
| # | interaction | 4 | 0 | 0 | 1 | 1 | # | 20 | Small | paralle | 79 |
| # | disease | 7 | 4 | 0 | 0.636363636 | 1 | # | 20 | Small | ancom | 79 |
| # | sex | 4 | 0 | 0 | 1 | 1 | # | 20 | Small | ancom | 79 |

| | | | | | | | | | | | |
|---|---|---|---|---|---|---|---|---|---|---|---|
| # | time | 5 | 4 | 0 | 0.555555556 | 1 | # | 20 | Small | ancom | 79 |
| # | interaction | 0 | 0 | 4 | 0 | 0 | # | 20 | Small | ancom | 79 |
| # | disease | 7 | 9 | 0 | 0.4375 | 1 | # | 20 | Small | aldex2 | 79 |
| # | sex | 4 | 0 | 0 | 1 | 1 | # | 20 | Small | aldex2 | 79 |
| # | time | 5 | 7 | 0 | 0.416666667 | 1 | # | 20 | Small | aldex2 | 79 |
| # | interaction | 0 | 0 | 4 | 0 | 0 | # | 20 | Small | aldex2 | 79 |
| # | disease | 4 | 8 | 0 | 0.333333333 | 1 | # | 10 | Baseline | paralle | 80 |
| # | sex | 2 | 5 | 0 | 0.285714286 | 1 | # | 10 | Baseline | paralle | 80 |
| # | time | 2 | 7 | 0 | 0.222222222 | 1 | # | 10 | Baseline | paralle | 80 |
| # | interaction | 2 | 10 | 0 | 0.166666667 | 1 | # | 10 | Baseline | paralle | 80 |
| # | disease | 4 | 0 | 0 | 1 | 1 | # | 10 | Baseline | ancom | 80 |
| # | sex | 2 | 0 | 0 | 1 | 1 | # | 10 | Baseline | ancom | 80 |
| # | time | 2 | 0 | 0 | 1 | 1 | # | 10 | Baseline | ancom | 80 |
| # | interaction | 0 | 0 | 2 | 0 | 0 | # | 10 | Baseline | ancom | 80 |
| # | disease | 4 | 9 | 0 | 0.307692308 | 1 | # | 10 | Baseline | aldex2 | 80 |
| # | sex | 2 | 3 | 0 | 0.4 | 1 | # | 10 | Baseline | aldex2 | 80 |
| # | time | 2 | 2 | 0 | 0.5 | 1 | # | 10 | Baseline | aldex2 | 80 |
| # | interaction | 0 | 0 | 2 | 0 | 0 | # | 10 | Baseline | aldex2 | 80 |
| # | disease | 4 | 1 | 0 | 0.8 | 1 | # | 10 | Small | paralle | 80 |
| # | sex | 2 | 0 | 0 | 1 | 1 | # | 10 | Small | paralle | 80 |
| # | time | 2 | 0 | 0 | 1 | 1 | # | 10 | Small | paralle | 80 |
| # | interaction | 0 | 0 | 2 | 0 | 0 | # | 10 | Small | paralle | 80 |
| # | disease | 4 | 0 | 0 | 1 | 1 | # | 10 | Small | ancom | 80 |
| # | sex | 2 | 0 | 0 | 1 | 1 | # | 10 | Small | ancom | 80 |
| # | time | 2 | 0 | 0 | 1 | 1 | # | 10 | Small | ancom | 80 |
| # | interaction | 0 | 0 | 2 | 0 | 0 | # | 10 | Small | ancom | 80 |
| # | disease | 4 | 1 | 0 | 0.8 | 1 | # | 10 | Small | aldex2 | 80 |
| # | sex | 2 | 0 | 0 | 1 | 1 | # | 10 | Small | aldex2 | 80 |
| # | time | 2 | 2 | 0 | 0.5 | 1 | # | 10 | Small | aldex2 | 80 |
| # | interaction | 0 | 0 | 2 | 0 | 0 | # | 10 | Small | aldex2 | 80 |
| # | disease | 4 | 6 | 0 | 0.4 | 1 | # | 10 | Baseline | paralle | 80 |
| # | sex | 2 | 6 | 0 | 0.25 | 1 | # | 10 | Baseline | paralle | 80 |
| # | time | 2 | 4 | 0 | 0.333333333 | 1 | # | 10 | Baseline | paralle | 80 |
| # | interaction | 2 | 5 | 0 | 0.285714286 | 1 | # | 10 | Baseline | paralle | 80 |
| # | disease | 4 | 1 | 0 | 0.8 | 1 | # | 10 | Baseline | ancom | 80 |
| # | sex | 2 | 0 | 0 | 1 | 1 | # | 10 | Baseline | ancom | 80 |
| # | time | 2 | 2 | 0 | 0.5 | 1 | # | 10 | Baseline | ancom | 80 |
| # | interaction | 0 | 0 | 2 | 0 | 0 | # | 10 | Baseline | ancom | 80 |
| # | disease | 4 | 15 | 0 | 0.210526316 | 1 | # | 10 | Baseline | aldex2 | 80 |
| # | sex | 2 | 3 | 0 | 0.4 | 1 | # | 10 | Baseline | aldex2 | 80 |
| # | time | 2 | 6 | 0 | 0.25 | 1 | # | 10 | Baseline | aldex2 | 80 |
| # | interaction | 0 | 0 | 2 | 0 | 0 | # | 10 | Baseline | aldex2 | 80 |
| # | disease | 4 | 0 | 0 | 1 | 1 | # | 10 | Small | paralle | 80 |
| # | sex | 2 | 0 | 0 | 1 | 1 | # | 10 | Small | paralle | 80 |
| # | time | 2 | 0 | 0 | 1 | 1 | # | 10 | Small | paralle | 80 |
| # | interaction | 2 | 0 | 0 | 1 | 1 | # | 10 | Small | paralle | 80 |
| # | disease | 4 | 1 | 0 | 0.8 | 1 | # | 10 | Small | ancom | 80 |
| # | sex | 2 | 0 | 0 | 1 | 1 | # | 10 | Small | ancom | 80 |
| # | time | 2 | 2 | 0 | 0.5 | 1 | # | 10 | Small | ancom | 80 |
| # | interaction | 0 | 0 | 2 | 0 | 0 | # | 10 | Small | ancom | 80 |
| # | disease | 4 | 2 | 0 | 0.666666667 | 1 | # | 10 | Small | aldex2 | 80 |
| # | sex | 2 | 0 | 0 | 1 | 1 | # | 10 | Small | aldex2 | 80 |
| # | time | 2 | 2 | 0 | 0.5 | 1 | # | 10 | Small | aldex2 | 80 |
| # | interaction | 0 | 0 | 2 | 0 | 0 | # | 10 | Small | aldex2 | 80 |
| # | disease | 4 | 5 | 0 | 0.444444444 | 1 | # | 10 | Baseline | paralle | 80 |
| # | sex | 2 | 6 | 0 | 0.25 | 1 | # | 10 | Baseline | paralle | 80 |
| # | time | 2 | 5 | 0 | 0.285714286 | 1 | # | 10 | Baseline | paralle | 80 |
| # | interaction | 2 | 3 | 0 | 0.4 | 1 | # | 10 | Baseline | paralle | 80 |
| # | disease | 4 | 2 | 0 | 0.666666667 | 1 | # | 10 | Baseline | ancom | 80 |
| # | sex | 2 | 0 | 0 | 1 | 1 | # | 10 | Baseline | ancom | 80 |
| # | time | 2 | 2 | 0 | 0.5 | 1 | # | 10 | Baseline | ancom | 80 |
| # | interaction | 0 | 0 | 2 | 0 | 0 | # | 10 | Baseline | ancom | 80 |
| # | disease | 4 | 19 | 0 | 0.173913043 | 1 | # | 10 | Baseline | aldex2 | 80 |
| # | sex | 2 | 6 | 0 | 0.25 | 1 | # | 10 | Baseline | aldex2 | 80 |
| # | time | 2 | 7 | 0 | 0.222222222 | 1 | # | 10 | Baseline | aldex2 | 80 |
| # | interaction | 0 | 0 | 2 | 0 | 0 | # | 10 | Baseline | aldex2 | 80 |
| # | disease | 4 | 0 | 0 | 1 | 1 | # | 10 | Small | paralle | 80 |
| # | sex | 2 | 0 | 0 | 1 | 1 | # | 10 | Small | paralle | 80 |
| # | time | 2 | 0 | 0 | 1 | 1 | # | 10 | Small | paralle | 80 |
| # | interaction | 2 | 0 | 0 | 1 | 1 | # | 10 | Small | paralle | 80 |
| # | disease | 4 | 2 | 0 | 0.666666667 | 1 | # | 10 | Small | ancom | 80 |
| # | sex | 2 | 0 | 0 | 1 | 1 | # | 10 | Small | ancom | 80 |
| # | time | 2 | 2 | 0 | 0.5 | 1 | # | 10 | Small | ancom | 80 |
| # | interaction | 0 | 0 | 2 | 0 | 0 | # | 10 | Small | ancom | 80 |
| # | disease | 4 | 2 | 0 | 0.666666667 | 1 | # | 10 | Small | aldex2 | 80 |
| # | sex | 2 | 0 | 0 | 1 | 1 | # | 10 | Small | aldex2 | 80 |
| # | time | 2 | 2 | 0 | 0.5 | 1 | # | 10 | Small | aldex2 | 80 |
| # | interaction | 0 | 0 | 2 | 0 | 0 | # | 10 | Small | aldex2 | 80 |
| # | disease | 4 | 6 | 0 | 0.4 | 1 | # | 10 | Baseline | paralle | 80 |
| # | sex | 2 | 1 | 0 | 0.666666667 | 1 | # | 10 | Baseline | paralle | 80 |
| # | time | 2 | 2 | 0 | 0.5 | 1 | # | 10 | Baseline | paralle | 80 |
| # | interaction | 2 | 1 | 0 | 0.666666667 | 1 | # | 10 | Baseline | paralle | 80 |
| # | disease | 4 | 2 | 0 | 0.666666667 | 1 | # | 10 | Baseline | ancom | 80 |
| # | sex | 2 | 0 | 0 | 1 | 1 | # | 10 | Baseline | ancom | 80 |
| # | time | 2 | 2 | 0 | 0.5 | 1 | # | 10 | Baseline | ancom | 80 |
| # | interaction | 0 | 0 | 2 | 0 | 0 | # | 10 | Baseline | ancom | 80 |
| # | disease | 4 | 27 | 0 | 0.129032258 | 1 | # | 10 | Baseline | aldex2 | 80 |
| # | sex | 2 | 3 | 0 | 0.4 | 1 | # | 10 | Baseline | aldex2 | 80 |
| # | time | 2 | 11 | 0 | 0.153846154 | 1 | # | 10 | Baseline | aldex2 | 80 |
| # | interaction | 0 | 0 | 2 | 0 | 0 | # | 10 | Baseline | aldex2 | 80 |
| # | disease | 4 | 0 | 0 | 1 | 1 | # | 10 | Small | paralle | 80 |
| # | sex | 2 | 0 | 0 | 1 | 1 | # | 10 | Small | paralle | 80 |
| # | time | 2 | 0 | 0 | 1 | 1 | # | 10 | Small | paralle | 80 |
| # | interaction | 2 | 0 | 0 | 1 | 1 | # | 10 | Small | paralle | 80 |
| # | disease | 4 | 2 | 0 | 0.666666667 | 1 | # | 10 | Small | ancom | 80 |
| # | sex | 2 | 0 | 0 | 1 | 1 | # | 10 | Small | ancom | 80 |
| # | time | 2 | 2 | 0 | 0.5 | 1 | # | 10 | Small | ancom | 80 |
| # | interaction | 0 | 0 | 2 | 0 | 0 | # | 10 | Small | ancom | 80 |
| # | disease | 4 | 4 | 0 | 0.5 | 1 | # | 10 | Small | aldex2 | 80 |
| # | sex | 2 | 0 | 0 | 1 | 1 | # | 10 | Small | aldex2 | 80 |
| # | time | 2 | 2 | 0 | 0.5 | 1 | # | 10 | Small | aldex2 | 80 |
| # | interaction | 0 | 0 | 2 | 0 | 0 | # | 10 | Small | aldex2 | 80 |
| # | disease | 7 | 10 | 0 | 0.411764706 | 1 | # | 20 | Baseline | paralle | 80 |
| # | sex | 4 | 9 | 0 | 0.307692308 | 1 | # | 20 | Baseline | paralle | 80 |
| # | time | 5 | 7 | 0 | 0.416666667 | 1 | # | 20 | Baseline | paralle | 80 |
| # | interaction | 4 | 8 | 0 | 0.333333333 | 1 | # | 20 | Baseline | paralle | 80 |
| # | disease | 7 | 2 | 0 | 0.777777778 | 1 | # | 20 | Baseline | ancom | 80 |
| # | sex | 4 | 0 | 0 | 1 | 1 | # | 20 | Baseline | ancom | 80 |
| # | time | 5 | 0 | 0 | 1 | 1 | # | 20 | Baseline | ancom | 80 |
| # | interaction | 0 | 0 | 4 | 0 | 0 | # | 20 | Baseline | ancom | 80 |
| # | disease | 7 | 22 | 0 | 0.24137931 | 1 | # | 20 | Baseline | aldex2 | 80 |
| # | sex | 4 | 4 | 0 | 0.5 | 1 | # | 20 | Baseline | aldex2 | 80 |
| # | time | 5 | 9 | 0 | 0.357142857 | 1 | # | 20 | Baseline | aldex2 | 80 |
| # | interaction | 0 | 0 | 4 | 0 | 0 | # | 20 | Baseline | aldex2 | 80 |
| # | disease | 7 | 0 | 0 | 1 | 1 | # | 20 | Small | paralle | 80 |
| # | sex | 4 | 0 | 0 | 1 | 1 | # | 20 | Small | paralle | 80 |
| # | time | 5 | 0 | 0 | 1 | 1 | # | 20 | Small | paralle | 80 |
| # | interaction | 1 | 0 | 3 | 1 | 0.25 | # | 20 | Small | paralle | 80 |
| # | disease | 7 | 0 | 0 | 1 | 1 | # | 20 | Small | ancom | 80 |
| # | sex | 4 | 0 | 0 | 1 | 1 | # | 20 | Small | ancom | 80 |
| # | time | 5 | 1 | 0 | 0.833333333 | 1 | # | 20 | Small | ancom | 80 |
| # | interaction | 0 | 0 | 4 | 0 | 0 | # | 20 | Small | ancom | 80 |
| # | disease | 7 | 1 | 0 | 0.875 | 1 | # | 20 | Small | aldex2 | 80 |
| # | sex | 4 | 0 | 0 | 1 | 1 | # | 20 | Small | aldex2 | 80 |
| # | time | 5 | 2 | 0 | 0.714285714 | 1 | # | 20 | Small | aldex2 | 80 |
| # | interaction | 0 | 0 | 4 | 0 | 0 | # | 20 | Small | aldex2 | 80 |
| # | disease | 7 | 11 | 0 | 0.388888889 | 1 | # | 20 | Baseline | paralle | 80 |
| # | sex | 4 | 8 | 0 | 0.333333333 | 1 | # | 20 | Baseline | paralle | 80 |
| # | time | 5 | 5 | 0 | 0.5 | 1 | # | 20 | Baseline | paralle | 80 |
| # | interaction | 4 | 6 | 0 | 0.4 | 1 | # | 20 | Baseline | paralle | 80 |
| # | disease | 7 | 2 | 0 | 0.777777778 | 1 | # | 20 | Baseline | ancom | 80 |
| # | sex | 4 | 0 | 0 | 1 | 1 | # | 20 | Baseline | ancom | 80 |
| # | time | 5 | 2 | 0 | 0.714285714 | 1 | # | 20 | Baseline | ancom | 80 |
| # | interaction | 0 | 0 | 4 | 0 | 0 | # | 20 | Baseline | ancom | 80 |
| # | disease | 7 | 35 | 0 | 0.166666667 | 1 | # | 20 | Baseline | aldex2 | 80 |
| # | sex | 4 | 7 | 0 | 0.363636364 | 1 | # | 20 | Baseline | aldex2 | 80 |
| # | time | 5 | 20 | 0 | 0.2 | 1 | # | 20 | Baseline | aldex2 | 80 |
| # | interaction | 0 | 0 | 4 | 0 | 0 | # | 20 | Baseline | aldex2 | 80 |
| # | disease | 7 | 1 | 0 | 0.875 | 1 | # | 20 | Small | paralle | 80 |
| # | sex | 4 | 0 | 0 | 1 | 1 | # | 20 | Small | paralle | 80 |
| # | time | 5 | 0 | 0 | 1 | 1 | # | 20 | Small | paralle | 80 |
| # | interaction | 4 | 0 | 0 | 1 | 1 | # | 20 | Small | paralle | 80 |
| # | disease | 7 | 2 | 0 | 0.777777778 | 1 | # | 20 | Small | ancom | 80 |
| # | sex | 4 | 0 | 0 | 1 | 1 | # | 20 | Small | ancom | 80 |
| # | time | 5 | 2 | 0 | 0.714285714 | 1 | # | 20 | Small | ancom | 80 |
| # | interaction | 0 | 0 | 4 | 0 | 0 | # | 20 | Small | ancom | 80 |
| # | disease | 7 | 7 | 0 | 0.5 | 1 | # | 20 | Small | aldex2 | 80 |
| # | sex | 4 | 0 | 0 | 1 | 1 | # | 20 | Small | aldex2 | 80 |
| # | time | 5 | 1 | 0 | 0.833333333 | 1 | # | 20 | Small | aldex2 | 80 |
| # | interaction | 0 | 0 | 4 | 0 | 0 | # | 20 | Small | aldex2 | 80 |
| # | disease | 7 | 16 | 0 | 0.304347826 | 1 | # | 20 | Baseline | paralle | 80 |
| # | sex | 4 | 5 | 0 | 0.444444444 | 1 | # | 20 | Baseline | paralle | 80 |
| # | time | 5 | 6 | 0 | 0.454545455 | 1 | # | 20 | Baseline | paralle | 80 |
| # | interaction | 4 | 7 | 0 | 0.363636364 | 1 | # | 20 | Baseline | paralle | 80 |
| # | disease | 7 | 4 | 0 | 0.636363636 | 1 | # | 20 | Baseline | ancom | 80 |
| # | sex | 4 | 0 | 0 | 1 | 1 | # | 20 | Baseline | ancom | 80 |
| # | time | 5 | 3 | 0 | 0.625 | 1 | # | 20 | Baseline | ancom | 80 |
| # | interaction | 0 | 0 | 4 | 0 | 0 | # | 20 | Baseline | ancom | 80 |
| # | disease | 7 | 40 | 0 | 0.14893617 | 1 | # | 20 | Baseline | aldex2 | 80 |
| # | sex | 4 | 17 | 0 | 0.19047619 | 1 | # | 20 | Baseline | aldex2 | 80 |
| # | time | 5 | 31 | 0 | 0.138888889 | 1 | # | 20 | Baseline | aldex2 | 80 |
| # | interaction | 0 | 0 | 4 | 0 | 0 | # | 20 | Baseline | aldex2 | 80 |
| # | disease | 7 | 0 | 0 | 1 | 1 | # | 20 | Small | paralle | 80 |
| # | sex | 4 | 0 | 0 | 1 | 1 | # | 20 | Small | paralle | 80 |
| # | time | 5 | 0 | 0 | 1 | 1 | # | 20 | Small | paralle | 80 |
| # | interaction | 4 | 0 | 0 | 1 | 1 | # | 20 | Small | paralle | 80 |
| # | disease | 7 | 4 | 0 | 0.636363636 | 1 | # | 20 | Small | ancom | 80 |
| # | sex | 4 | 0 | 0 | 1 | 1 | # | 20 | Small | ancom | 80 |
| # | time | 5 | 3 | 0 | 0.625 | 1 | # | 20 | Small | ancom | 80 |
| # | interaction | 0 | 0 | 4 | 0 | 0 | # | 20 | Small | ancom | 80 |
| # | disease | 7 | 12 | 0 | 0.368421053 | 1 | # | 20 | Small | aldex2 | 80 |
| # | sex | 4 | 0 | 0 | 1 | 1 | # | 20 | Small | aldex2 | 80 |
| # | time | 5 | 5 | 0 | 0.5 | 1 | # | 20 | Small | aldex2 | 80 |
| # | interaction | 0 | 0 | 4 | 0 | 0 | # | 20 | Small | aldex2 | 80 |
| # | disease | 7 | 7 | 0 | 0.5 | 1 | # | 20 | Baseline | paralle | 80 |
| # | sex | 4 | 4 | 0 | 0.5 | 1 | # | 20 | Baseline | paralle | 80 |
| # | time | 5 | 5 | 0 | 0.5 | 1 | # | 20 | Baseline | paralle | 80 |
| # | interaction | 4 | 6 | 0 | 0.4 | 1 | # | 20 | Baseline | paralle | 80 |
| # | disease | 7 | 4 | 0 | 0.636363636 | 1 | # | 20 | Baseline | ancom | 80 |
| # | sex | 4 | 0 | 0 | 1 | 1 | # | 20 | Baseline | ancom | 80 |
| # | time | 5 | 4 | 0 | 0.555555556 | 1 | # | 20 | Baseline | ancom | 80 |
| # | interaction | 0 | 0 | 4 | 0 | 0 | # | 20 | Baseline | ancom | 80 |
| # | disease | 7 | 50 | 0 | 0.104477612 | 1 | # | 20 | Baseline | aldex2 | 80 |
| # | sex | 4 | 20 | 0 | 0.166666667 | 1 | # | 20 | Baseline | aldex2 | 80 |
| # | time | 5 | 41 | 0 | 0.108695652 | 1 | # | 20 | Baseline | aldex2 | 80 |
| # | interaction | 0 | 0 | 4 | 0 | 0 | # | 20 | Baseline | aldex2 | 80 |
| # | disease | 7 | 0 | 0 | 1 | 1 | # | 20 | Small | paralle | 80 |
| # | sex | 4 | 0 | 0 | 1 | 1 | # | 20 | Small | paralle | 80 |
| # | time | 5 | 0 | 0 | 1 | 1 | # | 20 | Small | paralle | 80 |
| # | interaction | 4 | 0 | 0 | 1 | 1 | # | 20 | Small | paralle | 80 |
| # | disease | 7 | 4 | 0 | 0.636363636 | 1 | # | 20 | Small | ancom | 80 |
| # | sex | 4 | 0 | 0 | 1 | 1 | # | 20 | Small | ancom | 80 |
| # | time | 5 | 4 | 0 | 0.555555556 | 1 | # | 20 | Small | ancom | 80 |
| # | interaction | 0 | 0 | 4 | 0 | 0 | # | 20 | Small | ancom | 80 |
| # | disease | 7 | 13 | 0 | 0.35 | 1 | # | 20 | Small | aldex2 | 80 |
| # | sex | 4 | 0 | 0 | 1 | 1 | # | 20 | Small | aldex2 | 80 |
| # | time | 5 | 6 | 0 | 0.454545455 | 1 | # | 20 | Small | aldex2 | 80 |
| # | interaction | 0 | 0 | 4 | 0 | 0 | # | 20 | Small | aldex2 | 80 |
| # | disease | 4 | 11 | 0 | 0.266666667 | 1 | # | 10 | Baseline | paralle | 81 |
| # | sex | 2 | 11 | 0 | 0.153846154 | 1 | # | 10 | Baseline | paralle | 81 |
| # | time | 2 | 5 | 0 | 0.285714286 | 1 | # | 10 | Baseline | paralle | 81 |
| # | interaction | 2 | 8 | 0 | 0.2 | 1 | # | 10 | Baseline | paralle | 81 |
| # | disease | 4 | 2 | 0 | 0.666666667 | 1 | # | 10 | Baseline | ancom | 81 |
| # | sex | 2 | 0 | 0 | 1 | 1 | # | 10 | Baseline | ancom | 81 |
| # | time | 2 | 0 | 0 | 1 | 1 | # | 10 | Baseline | ancom | 81 |
| # | interaction | 0 | 0 | 2 | 0 | 0 | # | 10 | Baseline | ancom | 81 |
| # | disease | 4 | 6 | 0 | 0.4 | 1 | # | 10 | Baseline | aldex2 | 81 |
| # | sex | 2 | 1 | 0 | 0.666666667 | 1 | # | 10 | Baseline | aldex2 | 81 |
| # | time | 2 | 10 | 0 | 0.166666667 | 1 | # | 10 | Baseline | aldex2 | 81 |
| # | interaction | 0 | 0 | 2 | 0 | 0 | # | 10 | Baseline | aldex2 | 81 |
| # | disease | 4 | 0 | 0 | 1 | 1 | # | 10 | Small | paralle | 81 |
| # | sex | 2 | 0 | 0 | 1 | 1 | # | 10 | Small | paralle | 81 |
| # | time | 2 | 0 | 0 | 1 | 1 | # | 10 | Small | paralle | 81 |
| # | interaction | 2 | 0 | 0 | 1 | 1 | # | 10 | Small | paralle | 81 |
| # | disease | 4 | 2 | 0 | 0.666666667 | 1 | # | 10 | Small | ancom | 81 |
| # | sex | 2 | 0 | 0 | 1 | 1 | # | 10 | Small | ancom | 81 |
| # | time | 2 | 0 | 0 | 1 | 1 | # | 10 | Small | ancom | 81 |
| # | interaction | 0 | 0 | 2 | 0 | 0 | # | 10 | Small | ancom | 81 |
| # | disease | 4 | 2 | 0 | 0.666666667 | 1 | # | 10 | Small | aldex2 | 81 |
| # | sex | 2 | 0 | 0 | 1 | 1 | # | 10 | Small | aldex2 | 81 |
| # | time | 2 | 3 | 0 | 0.4 | 1 | # | 10 | Small | aldex2 | 81 |
| # | interaction | 0 | 0 | 2 | 0 | 0 | # | 10 | Small | aldex2 | 81 |
| # | disease | 4 | 6 | 0 | 0.4 | 1 | # | 10 | Baseline | paralle | 81 |
| # | sex | 2 | 6 | 0 | 0.25 | 1 | # | 10 | Baseline | paralle | 81 |
| # | time | 2 | 4 | 0 | 0.333333333 | 1 | # | 10 | Baseline | paralle | 81 |
| # | interaction | 2 | 4 | 0 | 0.333333333 | 1 | # | 10 | Baseline | paralle | 81 |
| # | disease | 4 | 2 | 0 | 0.666666667 | 1 | # | 10 | Baseline | ancom | 81 |
| # | sex | 2 | 0 | 0 | 1 | 1 | # | 10 | Baseline | ancom | 81 |
| # | time | 2 | 2 | 0 | 0.5 | 1 | # | 10 | Baseline | ancom | 81 |
| # | interaction | 0 | 0 | 2 | 0 | 0 | # | 10 | Baseline | ancom | 81 |
| # | disease | 4 | 10 | 0 | 0.285714286 | 1 | # | 10 | Baseline | aldex2 | 81 |
| # | sex | 2 | 6 | 0 | 0.25 | 1 | # | 10 | Baseline | aldex2 | 81 |
| # | time | 2 | 10 | 0 | 0.166666667 | 1 | # | 10 | Baseline | aldex2 | 81 |
| # | interaction | 0 | 0 | 2 | 0 | 0 | # | 10 | Baseline | aldex2 | 81 |
| # | disease | 4 | 0 | 0 | 1 | 1 | # | 10 | Small | paralle | 81 |
| # | sex | 2 | 0 | 0 | 1 | 1 | # | 10 | Small | paralle | 81 |
| # | time | 2 | 2 | 0 | 0.5 | 1 | # | 10 | Small | paralle | 81 |
| # | interaction | 2 | 2 | 0 | 0.5 | 1 | # | 10 | Small | paralle | 81 |
| # | disease | 4 | 2 | 0 | 0.666666667 | 1 | # | 10 | Small | ancom | 81 |
| # | sex | 2 | 0 | 0 | 1 | 1 | # | 10 | Small | ancom | 81 |
| # | time | 2 | 2 | 0 | 0.5 | 1 | # | 10 | Small | ancom | 81 |
| # | interaction | 0 | 0 | 2 | 0 | 0 | # | 10 | Small | ancom | 81 |
| # | disease | 4 | 2 | 0 | 0.666666667 | 1 | # | 10 | Small | aldex2 | 81 |
| # | sex | 2 | 0 | 0 | 1 | 1 | # | 10 | Small | aldex2 | 81 |
| # | time | 2 | 2 | 0 | 0.5 | 1 | # | 10 | Small | aldex2 | 81 |
| # | interaction | 0 | 0 | 2 | 0 | 0 | # | 10 | Small | aldex2 | 81 |
| # | disease | 4 | 7 | 0 | 0.363636364 | 1 | # | 10 | Baseline | paralle | 81 |
| # | sex | 2 | 3 | 0 | 0.4 | 1 | # | 10 | Baseline | paralle | 81 |
| # | time | 2 | 3 | 0 | 0.4 | 1 | # | 10 | Baseline | paralle | 81 |
| # | interaction | 2 | 4 | 0 | 0.333333333 | 1 | # | 10 | Baseline | paralle | 81 |
| # | disease | 4 | 2 | 0 | 0.666666667 | 1 | # | 10 | Baseline | ancom | 81 |
| # | sex | 2 | 0 | 0 | 1 | 1 | # | 10 | Baseline | ancom | 81 |
| # | time | 2 | 2 | 0 | 0.5 | 1 | # | 10 | Baseline | ancom | 81 |
| # | interaction | 0 | 0 | 2 | 0 | 0 | # | 10 | Baseline | ancom | 81 |
| # | disease | 4 | 19 | 0 | 0.173913043 | 1 | # | 10 | Baseline | aldex2 | 81 |
| # | sex | 2 | 5 | 0 | 0.285714286 | 1 | # | 10 | Baseline | aldex2 | 81 |
| # | time | 2 | 10 | 0 | 0.166666667 | 1 | # | 10 | Baseline | aldex2 | 81 |
| # | interaction | 0 | 0 | 2 | 0 | 0 | # | 10 | Baseline | aldex2 | 81 |
| # | disease | 4 | 0 | 0 | 1 | 1 | # | 10 | Small | paralle | 81 |
| # | sex | 2 | 0 | 0 | 1 | 1 | # | 10 | Small | paralle | 81 |
| # | time | 2 | 2 | 0 | 0.5 | 1 | # | 10 | Small | paralle | 81 |
| # | interaction | 2 | 1 | 0 | 0.666666667 | 1 | # | 10 | Small | paralle | 81 |
| # | disease | 4 | 2 | 0 | 0.666666667 | 1 | # | 10 | Small | ancom | 81 |
| # | sex | 2 | 0 | 0 | 1 | 1 | # | 10 | Small | ancom | 81 |
| # | time | 2 | 2 | 0 | 0.5 | 1 | # | 10 | Small | ancom | 81 |
| # | interaction | 0 | 0 | 2 | 0 | 0 | # | 10 | Small | ancom | 81 |
| # | disease | 4 | 2 | 0 | 0.666666667 | 1 | # | 10 | Small | aldex2 | 81 |
| # | sex | 2 | 0 | 0 | 1 | 1 | # | 10 | Small | aldex2 | 81 |
| # | time | 2 | 2 | 0 | 0.5 | 1 | # | 10 | Small | aldex2 | 81 |
| # | interaction | 0 | 0 | 2 | 0 | 0 | # | 10 | Small | aldex2 | 81 |
| # | disease | 4 | 5 | 0 | 0.444444444 | 1 | # | 10 | Baseline | paralle | 81 |
| # | sex | 2 | 3 | 0 | 0.4 | 1 | # | 10 | Baseline | paralle | 81 |
| # | time | 2 | 4 | 0 | 0.333333333 | 1 | # | 10 | Baseline | paralle | 81 |
| # | interaction | 2 | 7 | 0 | 0.222222222 | 1 | # | 10 | Baseline | paralle | 81 |
| # | disease | 4 | 2 | 0 | 0.666666667 | 1 | # | 10 | Baseline | ancom | 81 |
| # | sex | 2 | 0 | 0 | 1 | 1 | # | 10 | Baseline | ancom | 81 |
| # | time | 2 | 2 | 0 | 0.5 | 1 | # | 10 | Baseline | ancom | 81 |
| # | interaction | 0 | 0 | 2 | 0 | 0 | # | 10 | Baseline | ancom | 81 |
| # | disease | 4 | 20 | 0 | 0.166666667 | 1 | # | 10 | Baseline | aldex2 | 81 |
| # | sex | 2 | 6 | 0 | 0.25 | 1 | # | 10 | Baseline | aldex2 | 81 |
| # | time | 2 | 14 | 0 | 0.125 | 1 | # | 10 | Baseline | aldex2 | 81 |
| # | interaction | 0 | 0 | 2 | 0 | 0 | # | 10 | Baseline | aldex2 | 81 |
| # | disease | 4 | 0 | 0 | 1 | 1 | # | 10 | Small | paralle | 81 |
| # | sex | 2 | 0 | 0 | 1 | 1 | # | 10 | Small | paralle | 81 |
| # | time | 2 | 1 | 0 | 0.666666667 | 1 | # | 10 | Small | paralle | 81 |
| # | interaction | 2 | 0 | 0 | 1 | 1 | # | 10 | Small | paralle | 81 |
| # | disease | 4 | 2 | 0 | 0.666666667 | 1 | # | 10 | Small | ancom | 81 |
| # | sex | 2 | 0 | 0 | 1 | 1 | # | 10 | Small | ancom | 81 |
| # | time | 2 | 2 | 0 | 0.5 | 1 | # | 10 | Small | ancom | 81 |
| # | interaction | 0 | 0 | 2 | 0 | 0 | # | 10 | Small | ancom | 81 |
| # | disease | 4 | 2 | 0 | 0.666666667 | 1 | # | 10 | Small | aldex2 | 81 |
| # | sex | 2 | 0 | 0 | 1 | 1 | # | 10 | Small | aldex2 | 81 |
| # | time | 2 | 2 | 0 | 0.5 | 1 | # | 10 | Small | aldex2 | 81 |
| # | interaction | 0 | 0 | 2 | 0 | 0 | # | 10 | Small | aldex2 | 81 |
| # | disease | 7 | 11 | 0 | 0.388888889 | 1 | # | 20 | Baseline | paralle | 81 |
| # | sex | 4 | 8 | 0 | 0.333333333 | 1 | # | 20 | Baseline | paralle | 81 |
| # | time | 5 | 6 | 0 | 0.454545455 | 1 | # | 20 | Baseline | paralle | 81 |
| # | interaction | 4 | 3 | 0 | 0.571428571 | 1 | # | 20 | Baseline | paralle | 81 |
| # | disease | 7 | 3 | 0 | 0.7 | 1 | # | 20 | Baseline | ancom | 81 |
| # | sex | 4 | 0 | 0 | 1 | 1 | # | 20 | Baseline | ancom | 81 |
| # | time | 5 | 1 | 0 | 0.833333333 | 1 | # | 20 | Baseline | ancom | 81 |
| # | interaction | 0 | 0 | 4 | 0 | 0 | # | 20 | Baseline | ancom | 81 |
| # | disease | 7 | 16 | 0 | 0.304347826 | 1 | # | 20 | Baseline | aldex2 | 81 |
| # | sex | 4 | 8 | 0 | 0.333333333 | 1 | # | 20 | Baseline | aldex2 | 81 |
| # | time | 5 | 17 | 0 | 0.227272727 | 1 | # | 20 | Baseline | aldex2 | 81 |
| # | interaction | 0 | 0 | 4 | 0 | 0 | # | 20 | Baseline | aldex2 | 81 |
| # | disease | 7 | 1 | 0 | 0.875 | 1 | # | 20 | Small | paralle | 81 |
| # | sex | 4 | 0 | 0 | 1 | 1 | # | 20 | Small | paralle | 81 |
| # | time | 5 | 0 | 0 | 1 | 1 | # | 20 | Small | paralle | 81 |
| # | interaction | 3 | 0 | 1 | 1 | 0.75 | # | 20 | Small | paralle | 81 |
| # | disease | 7 | 2 | 0 | 0.777777778 | 1 | # | 20 | Small | ancom | 81 |
| # | sex | 4 | 0 | 0 | 1 | 1 | # | 20 | Small | ancom | 81 |
| # | time | 5 | 0 | 0 | 1 | 1 | # | 20 | Small | ancom | 81 |
| # | interaction | 0 | 0 | 4 | 0 | 0 | # | 20 | Small | ancom | 81 |
| # | disease | 7 | 0 | 0 | 1 | 1 | # | 20 | Small | aldex2 | 81 |
| # | sex | 4 | 0 | 0 | 1 | 1 | # | 20 | Small | aldex2 | 81 |
| # | time | 5 | 1 | 0 | 0.833333333 | 1 | # | 20 | Small | aldex2 | 81 |
| # | interaction | 0 | 0 | 4 | 0 | 0 | # | 20 | Small | aldex2 | 81 |
| # | disease | 7 | 13 | 0 | 0.35 | 1 | # | 20 | Baseline | paralle | 81 |
| # | sex | 4 | 7 | 0 | 0.363636364 | 1 | # | 20 | Baseline | paralle | 81 |
| # | time | 5 | 4 | 0 | 0.555555556 | 1 | # | 20 | Baseline | paralle | 81 |
| # | interaction | 4 | 6 | 0 | 0.4 | 1 | # | 20 | Baseline | paralle | 81 |
| # | disease | 7 | 3 | 0 | 0.7 | 1 | # | 20 | Baseline | ancom | 81 |
| # | sex | 4 | 0 | 0 | 1 | 1 | # | 20 | Baseline | ancom | 81 |
| # | time | 5 | 4 | 0 | 0.555555556 | 1 | # | 20 | Baseline | ancom | 81 |
| # | interaction | 0 | 0 | 4 | 0 | 0 | # | 20 | Baseline | ancom | 81 |
| # | disease | 7 | 34 | 0 | 0.170731707 | 1 | # | 20 | Baseline | aldex2 | 81 |
| # | sex | 4 | 16 | 0 | 0.2 | 1 | # | 20 | Baseline | aldex2 | 81 |
| # | time | 5 | 23 | 0 | 0.178571429 | 1 | # | 20 | Baseline | aldex2 | 81 |
| # | interaction | 0 | 0 | 4 | 0 | 0 | # | 20 | Baseline | aldex2 | 81 |
| # | disease | 7 | 0 | 0 | 1 | 1 | # | 20 | Small | paralle | 81 |
| # | sex | 4 | 1 | 0 | 0.8 | 1 | # | 20 | Small | paralle | 81 |
| # | time | 5 | 0 | 0 | 1 | 1 | # | 20 | Small | paralle | 81 |
| # | interaction | 4 | 0 | 0 | 1 | 1 | # | 20 | Small | paralle | 81 |
| # | disease | 7 | 3 | 0 | 0.7 | 1 | # | 20 | Small | ancom | 81 |
| # | sex | 4 | 0 | 0 | 1 | 1 | # | 20 | Small | ancom | 81 |
| # | time | 5 | 3 | 0 | 0.625 | 1 | # | 20 | Small | ancom | 81 |

| | | | | | | | | | | | |
|---|---|---|---|---|---|---|---|---|---|---|---|
| # | interaction | 0 | 0 | 4 | 0 | 0 | # | 20 | Small | ancom | 01 |
| # | disease | 7 | 3 | 0 | 0.7 | 1 | # | 20 | Small | aldex2 | 01 |
| # | sex | 4 | 1 | 0 | 0.8 | 1 | # | 20 | Small | aldex2 | 01 |
| # | time | 5 | 2 | 0 | 0.714285714 | 1 | # | 20 | Small | aldex2 | 01 |
| # | interaction | 0 | 0 | 4 | 0 | 0 | # | 20 | Small | aldex2 | 01 |
| # | disease | 7 | 14 | 0 | 0.333333333 | 1 | # | 20 | Baseline | paralle | 01 |
| # | sex | 4 | 8 | 0 | 0.333333333 | 1 | # | 20 | Baseline | paralle | 01 |
| # | time | 5 | 4 | 0 | 0.555555556 | 1 | # | 20 | Baseline | paralle | 01 |
| # | interaction | 4 | 3 | 0 | 0.571428571 | 1 | # | 20 | Baseline | paralle | 01 |
| # | disease | 7 | 4 | 0 | 0.636363636 | 1 | # | 20 | Baseline | ancom | 01 |
| # | sex | 4 | 0 | 0 | 1 | 1 | # | 20 | Baseline | ancom | 01 |
| # | time | 5 | 4 | 0 | 0.555555556 | 1 | # | 20 | Baseline | ancom | 01 |
| # | interaction | 0 | 0 | 4 | 0 | 0 | # | 20 | Baseline | ancom | 01 |
| # | disease | 7 | 52 | 0 | 0.118644068 | 1 | # | 20 | Baseline | aldex2 | 01 |
| # | sex | 4 | 20 | 0 | 0.166666667 | 1 | # | 20 | Baseline | aldex2 | 01 |
| # | time | 5 | 37 | 0 | 0.119047619 | 1 | # | 20 | Baseline | aldex2 | 01 |
| # | interaction | 0 | 0 | 4 | 0 | 0 | # | 20 | Baseline | aldex2 | 01 |
| # | disease | 7 | 0 | 0 | 1 | 1 | # | 20 | Small | paralle | 01 |
| # | sex | 4 | 0 | 0 | 1 | 1 | # | 20 | Small | paralle | 01 |
| # | time | 5 | 0 | 0 | 1 | 1 | # | 20 | Small | paralle | 01 |
| # | interaction | 4 | 0 | 0 | 1 | 1 | # | 20 | Small | paralle | 01 |
| # | disease | 7 | 3 | 0 | 0.7 | 1 | # | 20 | Small | ancom | 01 |
| # | sex | 4 | 0 | 0 | 1 | 1 | # | 20 | Small | ancom | 01 |
| # | time | 5 | 4 | 0 | 0.555555556 | 1 | # | 20 | Small | ancom | 01 |
| # | interaction | 0 | 0 | 4 | 0 | 0 | # | 20 | Small | ancom | 01 |
| # | disease | 7 | 8 | 0 | 0.466666667 | 1 | # | 20 | Small | aldex2 | 01 |
| # | sex | 4 | 1 | 0 | 0.8 | 1 | # | 20 | Small | aldex2 | 01 |
| # | time | 5 | 6 | 0 | 0.454545455 | 1 | # | 20 | Small | aldex2 | 01 |
| # | interaction | 0 | 0 | 4 | 0 | 0 | # | 20 | Small | aldex2 | 01 |
| # | disease | 7 | 14 | 0 | 0.333333333 | 1 | # | 20 | Baseline | paralle | 01 |
| # | sex | 4 | 9 | 0 | 0.307692308 | 1 | # | 20 | Baseline | paralle | 01 |
| # | time | 5 | 3 | 0 | 0.625 | 1 | # | 20 | Baseline | paralle | 01 |
| # | interaction | 4 | 2 | 0 | 0.666666667 | 1 | # | 20 | Baseline | paralle | 01 |
| # | disease | 7 | 4 | 0 | 0.636363636 | 1 | # | 20 | Baseline | ancom | 01 |
| # | sex | 4 | 0 | 0 | 1 | 1 | # | 20 | Baseline | ancom | 01 |
| # | time | 5 | 4 | 0 | 0.555555556 | 1 | # | 20 | Baseline | ancom | 01 |
| # | interaction | 0 | 0 | 4 | 0 | 0 | # | 20 | Baseline | ancom | 01 |
| # | disease | 7 | 61 | 0 | 0.102941176 | 1 | # | 20 | Baseline | aldex2 | 01 |
| # | sex | 4 | 26 | 0 | 0.133333333 | 1 | # | 20 | Baseline | aldex2 | 01 |
| # | time | 5 | 41 | 0 | 0.108695652 | 1 | # | 20 | Baseline | aldex2 | 01 |
| # | interaction | 0 | 0 | 4 | 0 | 0 | # | 20 | Baseline | aldex2 | 01 |
| # | disease | 7 | 0 | 0 | 1 | 1 | # | 20 | Small | paralle | 01 |
| # | sex | 4 | 0 | 0 | 1 | 1 | # | 20 | Small | paralle | 01 |
| # | time | 5 | 1 | 0 | 0.833333333 | 1 | # | 20 | Small | paralle | 01 |
| # | interaction | 4 | 0 | 0 | 1 | 1 | # | 20 | Small | paralle | 01 |
| # | disease | 7 | 4 | 0 | 0.636363636 | 1 | # | 20 | Small | ancom | 01 |
| # | sex | 4 | 0 | 0 | 1 | 1 | # | 20 | Small | ancom | 01 |
| # | time | 5 | 4 | 0 | 0.555555556 | 1 | # | 20 | Small | ancom | 01 |
| # | interaction | 0 | 0 | 4 | 0 | 0 | # | 20 | Small | ancom | 01 |
| # | disease | 7 | 9 | 0 | 0.4375 | 1 | # | 20 | Small | aldex2 | 01 |
| # | sex | 4 | 1 | 0 | 0.8 | 1 | # | 20 | Small | aldex2 | 01 |
| # | time | 5 | 7 | 0 | 0.416666667 | 1 | # | 20 | Small | aldex2 | 01 |
| # | interaction | 0 | 0 | 4 | 0 | 0 | # | 20 | Small | aldex2 | 01 |
| # | disease | 4 | 7 | 0 | 0.363636364 | 1 | # | 10 | Baseline | paralle | 02 |
| # | sex | 2 | 14 | 0 | 0.125 | 1 | # | 10 | Baseline | paralle | 02 |
| # | time | 2 | 11 | 0 | 0.153846154 | 1 | # | 10 | Baseline | paralle | 02 |
| # | interaction | 2 | 12 | 0 | 0.142857143 | 1 | # | 10 | Baseline | paralle | 02 |
| # | disease | 4 | 0 | 0 | 1 | 1 | # | 10 | Baseline | ancom | 02 |
| # | sex | 2 | 0 | 0 | 1 | 1 | # | 10 | Baseline | ancom | 02 |
| # | time | 2 | 0 | 0 | 1 | 1 | # | 10 | Baseline | ancom | 02 |
| # | interaction | 0 | 0 | 2 | 0 | 0 | # | 10 | Baseline | ancom | 02 |
| # | disease | 4 | 9 | 0 | 0.307692308 | 1 | # | 10 | Baseline | aldex2 | 02 |
| # | sex | 2 | 9 | 0 | 0.181818182 | 1 | # | 10 | Baseline | aldex2 | 02 |
| # | time | 2 | 7 | 0 | 0.222222222 | 1 | # | 10 | Baseline | aldex2 | 02 |
| # | interaction | 0 | 0 | 2 | 0 | 0 | # | 10 | Baseline | aldex2 | 02 |
| # | disease | 4 | 0 | 0 | 1 | 1 | # | 10 | Small | paralle | 02 |
| # | sex | 2 | 0 | 0 | 1 | 1 | # | 10 | Small | paralle | 02 |
| # | time | 2 | 0 | 0 | 1 | 1 | # | 10 | Small | paralle | 02 |
| # | interaction | 0 | 0 | 2 | 0 | 0 | # | 10 | Small | paralle | 02 |
| # | disease | 4 | 0 | 0 | 1 | 1 | # | 10 | Small | ancom | 02 |
| # | sex | 2 | 0 | 0 | 1 | 1 | # | 10 | Small | ancom | 02 |
| # | time | 2 | 0 | 0 | 1 | 1 | # | 10 | Small | ancom | 02 |
| # | interaction | 0 | 0 | 2 | 0 | 0 | # | 10 | Small | ancom | 02 |
| # | disease | 4 | 0 | 0 | 1 | 1 | # | 10 | Small | aldex2 | 02 |
| # | sex | 2 | 0 | 0 | 1 | 1 | # | 10 | Small | aldex2 | 02 |
| # | time | 2 | 1 | 0 | 0.666666667 | 1 | # | 10 | Small | aldex2 | 02 |
| # | interaction | 0 | 0 | 2 | 0 | 0 | # | 10 | Small | aldex2 | 02 |
| # | disease | 4 | 8 | 0 | 0.333333333 | 1 | # | 10 | Baseline | paralle | 02 |
| # | sex | 2 | 7 | 0 | 0.222222222 | 1 | # | 10 | Baseline | paralle | 02 |
| # | time | 2 | 4 | 0 | 0.333333333 | 1 | # | 10 | Baseline | paralle | 02 |
| # | interaction | 2 | 9 | 0 | 0.181818182 | 1 | # | 10 | Baseline | paralle | 02 |
| # | disease | 4 | 1 | 0 | 0.8 | 1 | # | 10 | Baseline | ancom | 02 |
| # | sex | 2 | 0 | 0 | 1 | 1 | # | 10 | Baseline | ancom | 02 |
| # | time | 2 | 2 | 0 | 0.5 | 1 | # | 10 | Baseline | ancom | 02 |
| # | interaction | 0 | 0 | 2 | 0 | 0 | # | 10 | Baseline | ancom | 02 |
| # | disease | 4 | 14 | 0 | 0.222222222 | 1 | # | 10 | Baseline | aldex2 | 02 |
| # | sex | 2 | 7 | 0 | 0.222222222 | 1 | # | 10 | Baseline | aldex2 | 02 |
| # | time | 2 | 10 | 0 | 0.166666667 | 1 | # | 10 | Baseline | aldex2 | 02 |
| # | interaction | 0 | 0 | 2 | 0 | 0 | # | 10 | Baseline | aldex2 | 02 |
| # | disease | 4 | 0 | 0 | 1 | 1 | # | 10 | Small | paralle | 02 |
| # | sex | 2 | 0 | 0 | 1 | 1 | # | 10 | Small | paralle | 02 |
| # | time | 2 | 0 | 0 | 1 | 1 | # | 10 | Small | paralle | 02 |
| # | interaction | 2 | 0 | 0 | 1 | 1 | # | 10 | Small | paralle | 02 |
| # | disease | 4 | 1 | 0 | 0.8 | 1 | # | 10 | Small | ancom | 02 |
| # | sex | 2 | 0 | 0 | 1 | 1 | # | 10 | Small | ancom | 02 |
| # | time | 2 | 2 | 0 | 0.5 | 1 | # | 10 | Small | ancom | 02 |
| # | interaction | 0 | 0 | 2 | 0 | 0 | # | 10 | Small | ancom | 02 |
| # | disease | 4 | 2 | 0 | 0.666666667 | 1 | # | 10 | Small | aldex2 | 02 |
| # | sex | 2 | 0 | 0 | 1 | 1 | # | 10 | Small | aldex2 | 02 |
| # | time | 2 | 3 | 0 | 0.4 | 1 | # | 10 | Small | aldex2 | 02 |
| # | interaction | 0 | 0 | 2 | 0 | 0 | # | 10 | Small | aldex2 | 02 |
| # | disease | 4 | 5 | 0 | 0.444444444 | 1 | # | 10 | Baseline | paralle | 02 |
| # | sex | 2 | 3 | 0 | 0.4 | 1 | # | 10 | Baseline | paralle | 02 |
| # | time | 2 | 4 | 0 | 0.333333333 | 1 | # | 10 | Baseline | paralle | 02 |
| # | interaction | 2 | 9 | 0 | 0.181818182 | 1 | # | 10 | Baseline | paralle | 02 |
| # | disease | 4 | 1 | 0 | 0.8 | 1 | # | 10 | Baseline | ancom | 02 |
| # | sex | 2 | 0 | 0 | 1 | 1 | # | 10 | Baseline | ancom | 02 |
| # | time | 2 | 2 | 0 | 0.5 | 1 | # | 10 | Baseline | ancom | 02 |
| # | interaction | 0 | 0 | 2 | 0 | 0 | # | 10 | Baseline | ancom | 02 |
| # | disease | 4 | 17 | 0 | 0.19047619 | 1 | # | 10 | Baseline | aldex2 | 02 |
| # | sex | 2 | 8 | 0 | 0.2 | 1 | # | 10 | Baseline | aldex2 | 02 |
| # | time | 2 | 12 | 0 | 0.142857143 | 1 | # | 10 | Baseline | aldex2 | 02 |
| # | interaction | 0 | 0 | 2 | 0 | 0 | # | 10 | Baseline | aldex2 | 02 |
| # | disease | 4 | 0 | 0 | 1 | 1 | # | 10 | Small | paralle | 02 |
| # | sex | 2 | 0 | 0 | 1 | 1 | # | 10 | Small | paralle | 02 |
| # | time | 2 | 0 | 0 | 1 | 1 | # | 10 | Small | paralle | 02 |
| # | interaction | 2 | 0 | 0 | 1 | 1 | # | 10 | Small | paralle | 02 |
| # | disease | 4 | 1 | 0 | 0.8 | 1 | # | 10 | Small | ancom | 02 |
| # | sex | 2 | 0 | 0 | 1 | 1 | # | 10 | Small | ancom | 02 |
| # | time | 2 | 3 | 0 | 0.4 | 1 | # | 10 | Small | ancom | 02 |
| # | interaction | 0 | 0 | 2 | 0 | 0 | # | 10 | Small | ancom | 02 |
| # | disease | 4 | 2 | 0 | 0.666666667 | 1 | # | 10 | Small | aldex2 | 02 |
| # | sex | 2 | 0 | 0 | 1 | 1 | # | 10 | Small | aldex2 | 02 |
| # | time | 2 | 5 | 0 | 0.285714286 | 1 | # | 10 | Small | aldex2 | 02 |
| # | interaction | 0 | 0 | 2 | 0 | 0 | # | 10 | Small | aldex2 | 02 |
| # | disease | 4 | 5 | 0 | 0.444444444 | 1 | # | 10 | Baseline | paralle | 02 |
| # | sex | 2 | 6 | 0 | 0.25 | 1 | # | 10 | Baseline | paralle | 02 |
| # | time | 2 | 5 | 0 | 0.285714286 | 1 | # | 10 | Baseline | paralle | 02 |
| # | interaction | 2 | 5 | 0 | 0.285714286 | 1 | # | 10 | Baseline | paralle | 02 |
| # | disease | 4 | 2 | 0 | 0.666666667 | 1 | # | 10 | Baseline | ancom | 02 |
| # | sex | 2 | 0 | 0 | 1 | 1 | # | 10 | Baseline | ancom | 02 |
| # | time | 2 | 2 | 0 | 0.5 | 1 | # | 10 | Baseline | ancom | 02 |
| # | interaction | 0 | 0 | 2 | 0 | 0 | # | 10 | Baseline | ancom | 02 |
| # | disease | 4 | 23 | 0 | 0.148148148 | 1 | # | 10 | Baseline | aldex2 | 02 |
| # | sex | 2 | 10 | 0 | 0.166666667 | 1 | # | 10 | Baseline | aldex2 | 02 |
| # | time | 2 | 9 | 0 | 0.181818182 | 1 | # | 10 | Baseline | aldex2 | 02 |
| # | interaction | 0 | 0 | 2 | 0 | 0 | # | 10 | Baseline | aldex2 | 02 |
| # | disease | 4 | 0 | 0 | 1 | 1 | # | 10 | Small | paralle | 02 |
| # | sex | 2 | 0 | 0 | 1 | 1 | # | 10 | Small | paralle | 02 |
| # | time | 2 | 0 | 0 | 1 | 1 | # | 10 | Small | paralle | 02 |
| # | interaction | 2 | 0 | 0 | 1 | 1 | # | 10 | Small | paralle | 02 |
| # | disease | 4 | 2 | 0 | 0.666666667 | 1 | # | 10 | Small | ancom | 02 |
| # | sex | 2 | 0 | 0 | 1 | 1 | # | 10 | Small | ancom | 02 |
| # | time | 2 | 2 | 0 | 0.5 | 1 | # | 10 | Small | ancom | 02 |
| # | interaction | 0 | 0 | 2 | 0 | 0 | # | 10 | Small | ancom | 02 |
| # | disease | 4 | 3 | 0 | 0.571428571 | 1 | # | 10 | Small | aldex2 | 02 |
| # | sex | 2 | 0 | 0 | 1 | 1 | # | 10 | Small | aldex2 | 02 |
| # | time | 2 | 5 | 0 | 0.285714286 | 1 | # | 10 | Small | aldex2 | 02 |
| # | interaction | 0 | 0 | 2 | 0 | 0 | # | 10 | Small | aldex2 | 02 |
| # | disease | 7 | 8 | 0 | 0.466666667 | 1 | # | 20 | Baseline | paralle | 02 |
| # | sex | 4 | 6 | 0 | 0.4 | 1 | # | 20 | Baseline | paralle | 02 |
| # | time | 5 | 10 | 0 | 0.333333333 | 1 | # | 20 | Baseline | paralle | 02 |
| # | interaction | 4 | 6 | 0 | 0.4 | 1 | # | 20 | Baseline | paralle | 02 |
| # | disease | 7 | 0 | 0 | 1 | 1 | # | 20 | Baseline | ancom | 02 |
| # | sex | 4 | 0 | 0 | 1 | 1 | # | 20 | Baseline | ancom | 02 |
| # | time | 5 | 0 | 0 | 1 | 1 | # | 20 | Baseline | ancom | 02 |
| # | interaction | 0 | 0 | 4 | 0 | 0 | # | 20 | Baseline | ancom | 02 |
| # | disease | 7 | 14 | 0 | 0.333333333 | 1 | # | 20 | Baseline | aldex2 | 02 |
| # | sex | 4 | 9 | 0 | 0.307692308 | 1 | # | 20 | Baseline | aldex2 | 02 |
| # | time | 5 | 9 | 0 | 0.357142857 | 1 | # | 20 | Baseline | aldex2 | 02 |
| # | interaction | 0 | 0 | 4 | 0 | 0 | # | 20 | Baseline | aldex2 | 02 |
| # | disease | 7 | 0 | 0 | 1 | 1 | # | 20 | Small | paralle | 02 |
| # | sex | 4 | 0 | 0 | 1 | 1 | # | 20 | Small | paralle | 02 |
| # | time | 5 | 0 | 0 | 1 | 1 | # | 20 | Small | paralle | 02 |
| # | interaction | 1 | 0 | 3 | 1 | 0.25 | # | 20 | Small | paralle | 02 |
| # | disease | 7 | 0 | 0 | 1 | 1 | # | 20 | Small | ancom | 02 |
| # | sex | 4 | 0 | 0 | 1 | 1 | # | 20 | Small | ancom | 02 |
| # | time | 5 | 0 | 0 | 1 | 1 | # | 20 | Small | ancom | 02 |
| # | interaction | 0 | 0 | 4 | 0 | 0 | # | 20 | Small | ancom | 02 |
| # | disease | 7 | 1 | 0 | 0.875 | 1 | # | 20 | Small | aldex2 | 02 |
| # | sex | 4 | 1 | 0 | 0.8 | 1 | # | 20 | Small | aldex2 | 02 |
| # | time | 5 | 1 | 0 | 0.833333333 | 1 | # | 20 | Small | aldex2 | 02 |
| # | interaction | 0 | 0 | 4 | 0 | 0 | # | 20 | Small | aldex2 | 02 |
| # | disease | 7 | 12 | 0 | 0.368421053 | 1 | # | 20 | Baseline | paralle | 02 |
| # | sex | 4 | 3 | 0 | 0.571428571 | 1 | # | 20 | Baseline | paralle | 02 |
| # | time | 5 | 7 | 0 | 0.416666667 | 1 | # | 20 | Baseline | paralle | 02 |
| # | interaction | 4 | 8 | 0 | 0.333333333 | 1 | # | 20 | Baseline | paralle | 02 |
| # | disease | 7 | 2 | 0 | 0.777777778 | 1 | # | 20 | Baseline | ancom | 02 |
| # | sex | 4 | 0 | 0 | 1 | 1 | # | 20 | Baseline | ancom | 02 |
| # | time | 5 | 3 | 0 | 0.625 | 1 | # | 20 | Baseline | ancom | 02 |
| # | interaction | 0 | 0 | 4 | 0 | 0 | # | 20 | Baseline | ancom | 02 |
| # | disease | 7 | 34 | 0 | 0.170731707 | 1 | # | 20 | Baseline | aldex2 | 02 |
| # | sex | 4 | 14 | 0 | 0.222222222 | 1 | # | 20 | Baseline | aldex2 | 02 |
| # | time | 5 | 25 | 0 | 0.166666667 | 1 | # | 20 | Baseline | aldex2 | 02 |
| # | interaction | 0 | 0 | 4 | 0 | 0 | # | 20 | Baseline | aldex2 | 02 |
| # | disease | 7 | 1 | 0 | 0.875 | 1 | # | 20 | Small | paralle | 02 |
| # | sex | 4 | 0 | 0 | 1 | 1 | # | 20 | Small | paralle | 02 |
| # | time | 5 | 1 | 0 | 0.833333333 | 1 | # | 20 | Small | paralle | 02 |
| # | interaction | 3 | 0 | 1 | 1 | 0.75 | # | 20 | Small | paralle | 02 |
| # | disease | 7 | 1 | 0 | 0.875 | 1 | # | 20 | Small | ancom | 02 |
| # | sex | 4 | 0 | 0 | 1 | 1 | # | 20 | Small | ancom | 02 |
| # | time | 5 | 2 | 0 | 0.714285714 | 1 | # | 20 | Small | ancom | 02 |
| # | interaction | 0 | 0 | 4 | 0 | 0 | # | 20 | Small | ancom | 02 |
| # | disease | 7 | 2 | 0 | 0.777777778 | 1 | # | 20 | Small | aldex2 | 02 |
| # | sex | 4 | 1 | 0 | 0.8 | 1 | # | 20 | Small | aldex2 | 02 |
| # | time | 5 | 2 | 0 | 0.714285714 | 1 | # | 20 | Small | aldex2 | 02 |
| # | interaction | 0 | 0 | 4 | 0 | 0 | # | 20 | Small | aldex2 | 02 |
| # | disease | 7 | 10 | 0 | 0.411764706 | 1 | # | 20 | Baseline | paralle | 02 |
| # | sex | 4 | 5 | 0 | 0.444444444 | 1 | # | 20 | Baseline | paralle | 02 |
| # | time | 5 | 7 | 0 | 0.416666667 | 1 | # | 20 | Baseline | paralle | 02 |
| # | interaction | 4 | 1 | 0 | 0.8 | 1 | # | 20 | Baseline | paralle | 02 |
| # | disease | 7 | 4 | 0 | 0.636363636 | 1 | # | 20 | Baseline | ancom | 02 |
| # | sex | 4 | 0 | 0 | 1 | 1 | # | 20 | Baseline | ancom | 02 |
| # | time | 5 | 4 | 0 | 0.555555556 | 1 | # | 20 | Baseline | ancom | 02 |
| # | interaction | 0 | 0 | 4 | 0 | 0 | # | 20 | Baseline | ancom | 02 |
| # | disease | 7 | 47 | 0 | 0.12962963 | 1 | # | 20 | Baseline | aldex2 | 02 |
| # | sex | 4 | 17 | 0 | 0.19047619 | 1 | # | 20 | Baseline | aldex2 | 02 |
| # | time | 5 | 34 | 0 | 0.128205128 | 1 | # | 20 | Baseline | aldex2 | 02 |
| # | interaction | 0 | 0 | 4 | 0 | 0 | # | 20 | Baseline | aldex2 | 02 |
| # | disease | 7 | 0 | 0 | 1 | 1 | # | 20 | Small | paralle | 02 |
| # | sex | 4 | 0 | 0 | 1 | 1 | # | 20 | Small | paralle | 02 |
| # | time | 5 | 0 | 0 | 1 | 1 | # | 20 | Small | paralle | 02 |
| # | interaction | 4 | 0 | 0 | 1 | 1 | # | 20 | Small | paralle | 02 |
| # | disease | 7 | 3 | 0 | 0.7 | 1 | # | 20 | Small | ancom | 02 |
| # | sex | 4 | 0 | 0 | 1 | 1 | # | 20 | Small | ancom | 02 |
| # | time | 5 | 4 | 0 | 0.555555556 | 1 | # | 20 | Small | ancom | 02 |
| # | interaction | 0 | 0 | 4 | 0 | 0 | # | 20 | Small | ancom | 02 |
| # | disease | 7 | 4 | 0 | 0.636363636 | 1 | # | 20 | Small | aldex2 | 02 |
| # | sex | 4 | 1 | 0 | 0.8 | 1 | # | 20 | Small | aldex2 | 02 |
| # | time | 5 | 3 | 0 | 0.625 | 1 | # | 20 | Small | aldex2 | 02 |
| # | interaction | 0 | 0 | 4 | 0 | 0 | # | 20 | Small | aldex2 | 02 |
| # | disease | 7 | 7 | 0 | 0.5 | 1 | # | 20 | Baseline | paralle | 02 |
| # | sex | 4 | 1 | 0 | 0.8 | 1 | # | 20 | Baseline | paralle | 02 |
| # | time | 5 | 8 | 0 | 0.384615385 | 1 | # | 20 | Baseline | paralle | 02 |
| # | interaction | 4 | 0 | 0 | 1 | 1 | # | 20 | Baseline | paralle | 02 |
| # | disease | 7 | 4 | 0 | 0.636363636 | 1 | # | 20 | Baseline | ancom | 02 |
| # | sex | 4 | 0 | 0 | 1 | 1 | # | 20 | Baseline | ancom | 02 |
| # | time | 5 | 4 | 0 | 0.555555556 | 1 | # | 20 | Baseline | ancom | 02 |
| # | interaction | 0 | 0 | 4 | 0 | 0 | # | 20 | Baseline | ancom | 02 |
| # | disease | 7 | 63 | 0 | 0.1 | 1 | # | 20 | Baseline | aldex2 | 02 |
| # | sex | 4 | 15 | 0 | 0.210526316 | 1 | # | 20 | Baseline | aldex2 | 02 |
| # | time | 5 | 44 | 0 | 0.102040816 | 1 | # | 20 | Baseline | aldex2 | 02 |
| # | interaction | 0 | 0 | 4 | 0 | 0 | # | 20 | Baseline | aldex2 | 02 |
| # | disease | 7 | 0 | 0 | 1 | 1 | # | 20 | Small | paralle | 02 |
| # | sex | 4 | 0 | 0 | 1 | 1 | # | 20 | Small | paralle | 02 |
| # | time | 5 | 0 | 0 | 1 | 1 | # | 20 | Small | paralle | 02 |
| # | interaction | 4 | 0 | 0 | 1 | 1 | # | 20 | Small | paralle | 02 |
| # | disease | 7 | 4 | 0 | 0.636363636 | 1 | # | 20 | Small | ancom | 02 |
| # | sex | 4 | 0 | 0 | 1 | 1 | # | 20 | Small | ancom | 02 |
| # | time | 5 | 4 | 0 | 0.555555556 | 1 | # | 20 | Small | ancom | 02 |
| # | interaction | 0 | 0 | 4 | 0 | 0 | # | 20 | Small | ancom | 02 |
| # | disease | 7 | 9 | 0 | 0.4375 | 1 | # | 20 | Small | aldex2 | 02 |
| # | sex | 4 | 2 | 0 | 0.666666667 | 1 | # | 20 | Small | aldex2 | 02 |
| # | time | 5 | 4 | 0 | 0.555555556 | 1 | # | 20 | Small | aldex2 | 02 |
| # | interaction | 0 | 0 | 4 | 0 | 0 | # | 20 | Small | aldex2 | 02 |
| # | disease | 4 | 12 | 0 | 0.25 | 1 | # | 10 | Baseline | paralle | 03 |
| # | sex | 2 | 7 | 0 | 0.222222222 | 1 | # | 10 | Baseline | paralle | 03 |
| # | time | 2 | 10 | 0 | 0.166666667 | 1 | # | 10 | Baseline | paralle | 03 |
| # | interaction | 2 | 16 | 0 | 0.111111111 | 1 | # | 10 | Baseline | paralle | 03 |
| # | disease | 4 | 1 | 0 | 0.8 | 1 | # | 10 | Baseline | ancom | 03 |
| # | sex | 2 | 0 | 0 | 1 | 1 | # | 10 | Baseline | ancom | 03 |
| # | time | 2 | 1 | 0 | 0.666666667 | 1 | # | 10 | Baseline | ancom | 03 |
| # | interaction | 0 | 0 | 2 | 0 | 0 | # | 10 | Baseline | ancom | 03 |
| # | disease | 4 | 11 | 0 | 0.266666667 | 1 | # | 10 | Baseline | aldex2 | 03 |
| # | sex | 2 | 4 | 0 | 0.333333333 | 1 | # | 10 | Baseline | aldex2 | 03 |
| # | time | 2 | 9 | 0 | 0.181818182 | 1 | # | 10 | Baseline | aldex2 | 03 |
| # | interaction | 0 | 0 | 2 | 0 | 0 | # | 10 | Baseline | aldex2 | 03 |
| # | disease | 4 | 0 | 0 | 1 | 1 | # | 10 | Small | paralle | 03 |
| # | sex | 2 | 0 | 0 | 1 | 1 | # | 10 | Small | paralle | 03 |
| # | time | 2 | 0 | 0 | 1 | 1 | # | 10 | Small | paralle | 03 |
| # | interaction | 2 | 1 | 0 | 0.666666667 | 1 | # | 10 | Small | paralle | 03 |
| # | disease | 4 | 1 | 0 | 0.8 | 1 | # | 10 | Small | ancom | 03 |
| # | sex | 2 | 0 | 0 | 1 | 1 | # | 10 | Small | ancom | 03 |
| # | time | 2 | 1 | 0 | 0.666666667 | 1 | # | 10 | Small | ancom | 03 |
| # | interaction | 0 | 0 | 2 | 0 | 0 | # | 10 | Small | ancom | 03 |
| # | disease | 4 | 1 | 0 | 0.8 | 1 | # | 10 | Small | aldex2 | 03 |
| # | sex | 2 | 0 | 0 | 1 | 1 | # | 10 | Small | aldex2 | 03 |
| # | time | 2 | 2 | 0 | 0.5 | 1 | # | 10 | Small | aldex2 | 03 |
| # | interaction | 0 | 0 | 2 | 0 | 0 | # | 10 | Small | aldex2 | 03 |
| # | disease | 4 | 10 | 0 | 0.285714286 | 1 | # | 10 | Baseline | paralle | 03 |
| # | sex | 2 | 4 | 0 | 0.333333333 | 1 | # | 10 | Baseline | paralle | 03 |
| # | time | 2 | 6 | 0 | 0.25 | 1 | # | 10 | Baseline | paralle | 03 |
| # | interaction | 2 | 6 | 0 | 0.25 | 1 | # | 10 | Baseline | paralle | 03 |
| # | disease | 4 | 2 | 0 | 0.666666667 | 1 | # | 10 | Baseline | ancom | 03 |
| # | sex | 2 | 0 | 0 | 1 | 1 | # | 10 | Baseline | ancom | 03 |
| # | time | 2 | 2 | 0 | 0.5 | 1 | # | 10 | Baseline | ancom | 03 |
| # | interaction | 0 | 0 | 2 | 0 | 0 | # | 10 | Baseline | ancom | 03 |
| # | disease | 4 | 13 | 0 | 0.235294118 | 1 | # | 10 | Baseline | aldex2 | 03 |
| # | sex | 2 | 2 | 0 | 0.5 | 1 | # | 10 | Baseline | aldex2 | 03 |
| # | time | 2 | 9 | 0 | 0.181818182 | 1 | # | 10 | Baseline | aldex2 | 03 |
| # | interaction | 0 | 0 | 2 | 0 | 0 | # | 10 | Baseline | aldex2 | 03 |
| # | disease | 4 | 0 | 0 | 1 | 1 | # | 10 | Small | paralle | 03 |
| # | sex | 2 | 0 | 0 | 1 | 1 | # | 10 | Small | paralle | 03 |
| # | time | 2 | 0 | 0 | 1 | 1 | # | 10 | Small | paralle | 03 |
| # | interaction | 2 | 0 | 0 | 1 | 1 | # | 10 | Small | paralle | 03 |
| # | disease | 4 | 2 | 0 | 0.666666667 | 1 | # | 10 | Small | ancom | 03 |
| # | sex | 2 | 0 | 0 | 1 | 1 | # | 10 | Small | ancom | 03 |
| # | time | 2 | 2 | 0 | 0.5 | 1 | # | 10 | Small | ancom | 03 |
| # | interaction | 0 | 0 | 2 | 0 | 0 | # | 10 | Small | ancom | 03 |
| # | disease | 4 | 3 | 0 | 0.571428571 | 1 | # | 10 | Small | aldex2 | 03 |
| # | sex | 2 | 0 | 0 | 1 | 1 | # | 10 | Small | aldex2 | 03 |
| # | time | 2 | 2 | 0 | 0.5 | 1 | # | 10 | Small | aldex2 | 03 |
| # | interaction | 0 | 0 | 2 | 0 | 0 | # | 10 | Small | aldex2 | 03 |
| # | disease | 4 | 8 | 0 | 0.333333333 | 1 | # | 10 | Baseline | paralle | 03 |
| # | sex | 2 | 5 | 0 | 0.285714286 | 1 | # | 10 | Baseline | paralle | 03 |
| # | time | 2 | 7 | 0 | 0.222222222 | 1 | # | 10 | Baseline | paralle | 03 |
| # | interaction | 2 | 3 | 0 | 0.4 | 1 | # | 10 | Baseline | paralle | 03 |
| # | disease | 4 | 2 | 0 | 0.666666667 | 1 | # | 10 | Baseline | ancom | 03 |
| # | sex | 2 | 0 | 0 | 1 | 1 | # | 10 | Baseline | ancom | 03 |
| # | time | 2 | 2 | 0 | 0.5 | 1 | # | 10 | Baseline | ancom | 03 |
| # | interaction | 0 | 0 | 2 | 0 | 0 | # | 10 | Baseline | ancom | 03 |
| # | disease | 4 | 18 | 0 | 0.181818182 | 1 | # | 10 | Baseline | aldex2 | 03 |
| # | sex | 2 | 5 | 0 | 0.285714286 | 1 | # | 10 | Baseline | aldex2 | 03 |
| # | time | 2 | 10 | 0 | 0.166666667 | 1 | # | 10 | Baseline | aldex2 | 03 |
| # | interaction | 0 | 0 | 2 | 0 | 0 | # | 10 | Baseline | aldex2 | 03 |
| # | disease | 4 | 0 | 0 | 1 | 1 | # | 10 | Small | paralle | 03 |
| # | sex | 2 | 0 | 0 | 1 | 1 | # | 10 | Small | paralle | 03 |
| # | time | 2 | 0 | 0 | 1 | 1 | # | 10 | Small | paralle | 03 |
| # | interaction | 2 | 0 | 0 | 1 | 1 | # | 10 | Small | paralle | 03 |
| # | disease | 4 | 2 | 0 | 0.666666667 | 1 | # | 10 | Small | ancom | 03 |
| # | sex | 2 | 0 | 0 | 1 | 1 | # | 10 | Small | ancom | 03 |
| # | time | 2 | 2 | 0 | 0.5 | 1 | # | 10 | Small | ancom | 03 |
| # | interaction | 0 | 0 | 2 | 0 | 0 | # | 10 | Small | ancom | 03 |
| # | disease | 4 | 4 | 0 | 0.5 | 1 | # | 10 | Small | aldex2 | 03 |
| # | sex | 2 | 0 | 0 | 1 | 1 | # | 10 | Small | aldex2 | 03 |
| # | time | 2 | 2 | 0 | 0.5 | 1 | # | 10 | Small | aldex2 | 03 |
| # | interaction | 0 | 0 | 2 | 0 | 0 | # | 10 | Small | aldex2 | 03 |
| # | disease | 4 | 10 | 0 | 0.285714286 | 1 | # | 10 | Baseline | paralle | 03 |
| # | sex | 2 | 1 | 0 | 0.666666667 | 1 | # | 10 | Baseline | paralle | 03 |
| # | time | 2 | 2 | 0 | 0.5 | 1 | # | 10 | Baseline | paralle | 03 |
| # | interaction | 2 | 5 | 0 | 0.285714286 | 1 | # | 10 | Baseline | paralle | 03 |
| # | disease | 4 | 2 | 0 | 0.666666667 | 1 | # | 10 | Baseline | ancom | 03 |
| # | sex | 2 | 0 | 0 | 1 | 1 | # | 10 | Baseline | ancom | 03 |
| # | time | 2 | 2 | 0 | 0.5 | 1 | # | 10 | Baseline | ancom | 03 |
| # | interaction | 0 | 0 | 2 | 0 | 0 | # | 10 | Baseline | ancom | 03 |
| # | disease | 4 | 29 | 0 | 0.121212121 | 1 | # | 10 | Baseline | aldex2 | 03 |
| # | sex | 2 | 4 | 0 | 0.333333333 | 1 | # | 10 | Baseline | aldex2 | 03 |
| # | time | 2 | 13 | 0 | 0.133333333 | 1 | # | 10 | Baseline | aldex2 | 03 |
| # | interaction | 0 | 0 | 2 | 0 | 0 | # | 10 | Baseline | aldex2 | 03 |
| # | disease | 4 | 0 | 0 | 1 | 1 | # | 10 | Small | paralle | 03 |
| # | sex | 2 | 0 | 0 | 1 | 1 | # | 10 | Small | paralle | 03 |
| # | time | 2 | 0 | 0 | 1 | 1 | # | 10 | Small | paralle | 03 |
| # | interaction | 2 | 0 | 0 | 1 | 1 | # | 10 | Small | paralle | 03 |
| # | disease | 4 | 2 | 0 | 0.666666667 | 1 | # | 10 | Small | ancom | 03 |
| # | sex | 2 | 0 | 0 | 1 | 1 | # | 10 | Small | ancom | 03 |
| # | time | 2 | 2 | 0 | 0.5 | 1 | # | 10 | Small | ancom | 03 |
| # | interaction | 0 | 0 | 2 | 0 | 0 | # | 10 | Small | ancom | 03 |

| | | | | | | | | | | | |
|---|---|---|---|---|---|---|---|---|---|---|---|
| # | disease | 4 | 3 | 0 | 0.571428571 | 1 | # | 10 | Small | sides2 | 83 |
| # | sex | 2 | 0 | 0 | 1 | 1 | # | 10 | Small | sides2 | 83 |
| # | time | 2 | 2 | 0 | 0.5 | 1 | # | 10 | Small | sides2 | 83 |
| # | interaction | 0 | 0 | 2 | 0 | 0 | # | 10 | Small | sides2 | 83 |
| # | disease | 7 | 8 | 0 | 0.466666667 | 1 | # | 20 | Baseline | paralle | 83 |
| # | sex | 4 | 7 | 0 | 0.363636364 | 1 | # | 20 | Baseline | paralle | 83 |
| # | time | 5 | 10 | 0 | 0.333333333 | 1 | # | 20 | Baseline | paralle | 83 |
| # | interaction | 3 | 4 | 1 | 0.428571429 | 0.75 | # | 20 | Baseline | paralle | 83 |
| # | disease | 7 | 2 | 0 | 0.777777778 | 1 | # | 20 | Baseline | ancom | 83 |
| # | sex | 4 | 0 | 0 | 1 | 1 | # | 20 | Baseline | ancom | 83 |
| # | time | 5 | 1 | 0 | 0.833333333 | 1 | # | 20 | Baseline | ancom | 83 |
| # | interaction | 0 | 0 | 4 | 0 | 0 | # | 20 | Baseline | ancom | 83 |
| # | disease | 7 | 28 | 0 | 0.2 | 1 | # | 20 | Baseline | sides2 | 83 |
| # | sex | 4 | 4 | 0 | 0.5 | 1 | # | 20 | Baseline | sides2 | 83 |
| # | time | 5 | 16 | 0 | 0.238095238 | 1 | # | 20 | Baseline | sides2 | 83 |
| # | interaction | 0 | 0 | 4 | 0 | 0 | # | 20 | Baseline | sides2 | 83 |
| # | disease | 7 | 0 | 0 | 1 | 1 | # | 20 | Small | paralle | 83 |
| # | sex | 4 | 0 | 0 | 1 | 1 | # | 20 | Small | paralle | 83 |
| # | time | 5 | 0 | 0 | 1 | 1 | # | 20 | Small | paralle | 83 |
| # | interaction | 1 | 0 | 3 | 1 | 0.25 | # | 20 | Small | paralle | 83 |
| # | disease | 7 | 2 | 0 | 0.777777778 | 1 | # | 20 | Small | ancom | 83 |
| # | sex | 4 | 0 | 0 | 1 | 1 | # | 20 | Small | ancom | 83 |
| # | time | 5 | 1 | 0 | 0.833333333 | 1 | # | 20 | Small | ancom | 83 |
| # | interaction | 0 | 0 | 4 | 0 | 0 | # | 20 | Small | ancom | 83 |
| # | disease | 7 | 2 | 0 | 0.777777778 | 1 | # | 20 | Small | sides2 | 83 |
| # | sex | 4 | 0 | 0 | 1 | 1 | # | 20 | Small | sides2 | 83 |
| # | time | 5 | 1 | 0 | 0.833333333 | 1 | # | 20 | Small | sides2 | 83 |
| # | interaction | 0 | 0 | 4 | 0 | 0 | # | 20 | Small | sides2 | 83 |
| # | disease | 7 | 14 | 0 | 0.333333333 | 1 | # | 20 | Baseline | paralle | 83 |
| # | sex | 4 | 9 | 0 | 0.307692308 | 1 | # | 20 | Baseline | paralle | 83 |
| # | time | 5 | 5 | 0 | 0.5 | 1 | # | 20 | Baseline | paralle | 83 |
| # | interaction | 4 | 5 | 0 | 0.444444444 | 1 | # | 20 | Baseline | paralle | 83 |
| # | disease | 7 | 3 | 0 | 0.7 | 1 | # | 20 | Baseline | ancom | 83 |
| # | sex | 4 | 0 | 0 | 1 | 1 | # | 20 | Baseline | ancom | 83 |
| # | time | 5 | 3 | 0 | 0.625 | 1 | # | 20 | Baseline | ancom | 83 |
| # | interaction | 0 | 0 | 4 | 0 | 0 | # | 20 | Baseline | ancom | 83 |
| # | disease | 7 | 41 | 0 | 0.145833333 | 1 | # | 20 | Baseline | sides2 | 83 |
| # | sex | 4 | 11 | 0 | 0.266666667 | 1 | # | 20 | Baseline | sides2 | 83 |
| # | time | 5 | 17 | 0 | 0.227272727 | 1 | # | 20 | Baseline | sides2 | 83 |
| # | interaction | 0 | 0 | 4 | 0 | 0 | # | 20 | Baseline | sides2 | 83 |
| # | disease | 7 | 0 | 0 | 1 | 1 | # | 20 | Small | paralle | 83 |
| # | sex | 4 | 0 | 0 | 1 | 1 | # | 20 | Small | paralle | 83 |
| # | time | 5 | 1 | 0 | 0.833333333 | 1 | # | 20 | Small | paralle | 83 |
| # | interaction | 3 | 0 | 1 | 1 | 0.75 | # | 20 | Small | paralle | 83 |
| # | disease | 7 | 3 | 0 | 0.7 | 1 | # | 20 | Small | ancom | 83 |
| # | sex | 4 | 0 | 0 | 1 | 1 | # | 20 | Small | ancom | 83 |
| # | time | 5 | 3 | 0 | 0.625 | 1 | # | 20 | Small | ancom | 83 |
| # | interaction | 0 | 0 | 4 | 0 | 0 | # | 20 | Small | ancom | 83 |
| # | disease | 7 | 2 | 0 | 0.777777778 | 1 | # | 20 | Small | sides2 | 83 |
| # | sex | 4 | 0 | 0 | 1 | 1 | # | 20 | Small | sides2 | 83 |
| # | time | 5 | 3 | 0 | 0.625 | 1 | # | 20 | Small | sides2 | 83 |
| # | interaction | 0 | 0 | 4 | 0 | 0 | # | 20 | Small | sides2 | 83 |
| # | disease | 7 | 11 | 0 | 0.388888889 | 1 | # | 20 | Baseline | paralle | 83 |
| # | sex | 4 | 4 | 0 | 0.5 | 1 | # | 20 | Baseline | paralle | 83 |
| # | time | 5 | 6 | 0 | 0.454545455 | 1 | # | 20 | Baseline | paralle | 83 |
| # | interaction | 4 | 8 | 0 | 0.333333333 | 1 | # | 20 | Baseline | paralle | 83 |
| # | disease | 7 | 3 | 0 | 0.7 | 1 | # | 20 | Baseline | ancom | 83 |
| # | sex | 4 | 0 | 0 | 1 | 1 | # | 20 | Baseline | ancom | 83 |
| # | time | 5 | 4 | 0 | 0.555555556 | 1 | # | 20 | Baseline | ancom | 83 |
| # | interaction | 0 | 0 | 4 | 0 | 0 | # | 20 | Baseline | ancom | 83 |
| # | disease | 7 | 55 | 0 | 0.112903226 | 1 | # | 20 | Baseline | sides2 | 83 |
| # | sex | 4 | 16 | 0 | 0.2 | 1 | # | 20 | Baseline | sides2 | 83 |
| # | time | 5 | 32 | 0 | 0.135135135 | 1 | # | 20 | Baseline | sides2 | 83 |
| # | interaction | 0 | 0 | 4 | 0 | 0 | # | 20 | Baseline | sides2 | 83 |
| # | disease | 7 | 0 | 0 | 1 | 1 | # | 20 | Small | paralle | 83 |
| # | sex | 4 | 0 | 0 | 1 | 1 | # | 20 | Small | paralle | 83 |
| # | time | 5 | 1 | 0 | 0.833333333 | 1 | # | 20 | Small | paralle | 83 |
| # | interaction | 4 | 0 | 0 | 1 | 1 | # | 20 | Small | paralle | 83 |
| # | disease | 7 | 3 | 0 | 0.7 | 1 | # | 20 | Small | ancom | 83 |
| # | sex | 4 | 0 | 0 | 1 | 1 | # | 20 | Small | ancom | 83 |
| # | time | 5 | 4 | 0 | 0.555555556 | 1 | # | 20 | Small | ancom | 83 |
| # | interaction | 0 | 0 | 4 | 0 | 0 | # | 20 | Small | ancom | 83 |
| # | disease | 7 | 7 | 0 | 0.5 | 1 | # | 20 | Small | sides2 | 83 |
| # | sex | 4 | 0 | 0 | 1 | 1 | # | 20 | Small | sides2 | 83 |
| # | time | 5 | 5 | 0 | 0.5 | 1 | # | 20 | Small | sides2 | 83 |
| # | interaction | 0 | 0 | 4 | 0 | 0 | # | 20 | Small | sides2 | 83 |
| # | disease | 7 | 13 | 0 | 0.35 | 1 | # | 20 | Baseline | paralle | 83 |
| # | sex | 4 | 6 | 0 | 0.4 | 1 | # | 20 | Baseline | paralle | 83 |
| # | time | 5 | 6 | 0 | 0.454545455 | 1 | # | 20 | Baseline | paralle | 83 |
| # | interaction | 4 | 7 | 0 | 0.363636364 | 1 | # | 20 | Baseline | paralle | 83 |
| # | disease | 7 | 4 | 0 | 0.636363636 | 1 | # | 20 | Baseline | ancom | 83 |
| # | sex | 4 | 0 | 0 | 1 | 1 | # | 20 | Baseline | ancom | 83 |
| # | time | 5 | 4 | 0 | 0.555555556 | 1 | # | 20 | Baseline | ancom | 83 |
| # | interaction | 0 | 0 | 4 | 0 | 0 | # | 20 | Baseline | ancom | 83 |
| # | disease | 7 | 35 | 0 | 0.097222222 | 1 | # | 20 | Baseline | sides2 | 83 |
| # | sex | 4 | 14 | 0 | 0.222222222 | 1 | # | 20 | Baseline | sides2 | 83 |
| # | time | 5 | 37 | 0 | 0.119047619 | 1 | # | 20 | Baseline | sides2 | 83 |
| # | interaction | 0 | 0 | 4 | 0 | 0 | # | 20 | Baseline | sides2 | 83 |
| # | disease | 7 | 0 | 0 | 1 | 1 | # | 20 | Small | paralle | 83 |
| # | sex | 4 | 1 | 0 | 0.8 | 1 | # | 20 | Small | paralle | 83 |
| # | time | 5 | 1 | 0 | 0.833333333 | 1 | # | 20 | Small | paralle | 83 |
| # | interaction | 4 | 0 | 0 | 1 | 1 | # | 20 | Small | paralle | 83 |
| # | disease | 7 | 3 | 0 | 0.7 | 1 | # | 20 | Small | ancom | 83 |
| # | sex | 4 | 0 | 0 | 1 | 1 | # | 20 | Small | ancom | 83 |
| # | time | 5 | 4 | 0 | 0.555555556 | 1 | # | 20 | Small | ancom | 83 |
| # | interaction | 0 | 0 | 4 | 0 | 0 | # | 20 | Small | ancom | 83 |
| # | disease | 7 | 11 | 0 | 0.388888889 | 1 | # | 20 | Small | sides2 | 83 |
| # | sex | 4 | 0 | 0 | 1 | 1 | # | 20 | Small | sides2 | 83 |
| # | time | 5 | 6 | 0 | 0.454545455 | 1 | # | 20 | Small | sides2 | 83 |
| # | interaction | 0 | 0 | 4 | 0 | 0 | # | 20 | Small | sides2 | 83 |
| # | disease | 4 | 13 | 0 | 0.235294118 | 1 | # | 10 | Baseline | paralle | 84 |
| # | sex | 2 | 4 | 0 | 0.333333333 | 1 | # | 10 | Baseline | paralle | 84 |
| # | time | 2 | 8 | 0 | 0.2 | 1 | # | 10 | Baseline | paralle | 84 |
| # | interaction | 2 | 3 | 0 | 0.4 | 1 | # | 10 | Baseline | paralle | 84 |
| # | disease | 4 | 0 | 0 | 1 | 1 | # | 10 | Baseline | ancom | 84 |
| # | sex | 2 | 0 | 0 | 1 | 1 | # | 10 | Baseline | ancom | 84 |
| # | time | 2 | 0 | 0 | 1 | 1 | # | 10 | Baseline | ancom | 84 |
| # | interaction | 0 | 0 | 2 | 0 | 0 | # | 10 | Baseline | ancom | 84 |
| # | disease | 4 | 12 | 0 | 0.25 | 1 | # | 10 | Baseline | sides2 | 84 |
| # | sex | 2 | 5 | 0 | 0.285714286 | 1 | # | 10 | Baseline | sides2 | 84 |
| # | time | 2 | 7 | 0 | 0.222222222 | 1 | # | 10 | Baseline | sides2 | 84 |
| # | interaction | 0 | 0 | 2 | 0 | 0 | # | 10 | Baseline | sides2 | 84 |
| # | disease | 4 | 1 | 0 | 0.8 | 1 | # | 10 | Small | paralle | 84 |
| # | sex | 2 | 0 | 0 | 1 | 1 | # | 10 | Small | paralle | 84 |
| # | time | 2 | 0 | 0 | 1 | 1 | # | 10 | Small | paralle | 84 |
| # | interaction | 1 | 0 | 1 | 1 | 0.5 | # | 10 | Small | paralle | 84 |
| # | disease | 4 | 0 | 0 | 1 | 1 | # | 10 | Small | ancom | 84 |
| # | sex | 2 | 0 | 0 | 1 | 1 | # | 10 | Small | ancom | 84 |
| # | time | 2 | 0 | 0 | 1 | 1 | # | 10 | Small | ancom | 84 |
| # | interaction | 0 | 0 | 2 | 0 | 0 | # | 10 | Small | ancom | 84 |
| # | disease | 4 | 0 | 0 | 1 | 1 | # | 10 | Small | sides2 | 84 |
| # | sex | 2 | 0 | 0 | 1 | 1 | # | 10 | Small | sides2 | 84 |
| # | time | 2 | 1 | 0 | 0.666666667 | 1 | # | 10 | Small | sides2 | 84 |
| # | interaction | 0 | 0 | 2 | 0 | 0 | # | 10 | Small | sides2 | 84 |
| # | disease | 4 | 8 | 0 | 0.333333333 | 1 | # | 10 | Baseline | paralle | 84 |
| # | sex | 2 | 3 | 0 | 0.4 | 1 | # | 10 | Baseline | paralle | 84 |
| # | time | 2 | 4 | 0 | 0.333333333 | 1 | # | 10 | Baseline | paralle | 84 |
| # | interaction | 2 | 9 | 0 | 0.181818182 | 1 | # | 10 | Baseline | paralle | 84 |
| # | disease | 4 | 0 | 0 | 1 | 1 | # | 10 | Baseline | ancom | 84 |
| # | sex | 2 | 0 | 0 | 1 | 1 | # | 10 | Baseline | ancom | 84 |
| # | time | 2 | 0 | 0 | 1 | 1 | # | 10 | Baseline | ancom | 84 |
| # | interaction | 0 | 0 | 2 | 0 | 0 | # | 10 | Baseline | ancom | 84 |
| # | disease | 4 | 14 | 0 | 0.222222222 | 1 | # | 10 | Baseline | sides2 | 84 |
| # | sex | 2 | 5 | 0 | 0.285714286 | 1 | # | 10 | Baseline | sides2 | 84 |
| # | time | 2 | 7 | 0 | 0.222222222 | 1 | # | 10 | Baseline | sides2 | 84 |
| # | interaction | 0 | 0 | 2 | 0 | 0 | # | 10 | Baseline | sides2 | 84 |
| # | disease | 4 | 0 | 0 | 1 | 1 | # | 10 | Small | paralle | 84 |
| # | sex | 2 | 0 | 0 | 1 | 1 | # | 10 | Small | paralle | 84 |
| # | time | 2 | 0 | 0 | 1 | 1 | # | 10 | Small | paralle | 84 |
| # | interaction | 2 | 0 | 0 | 1 | 1 | # | 10 | Small | paralle | 84 |
| # | disease | 4 | 0 | 0 | 1 | 1 | # | 10 | Small | ancom | 84 |
| # | sex | 2 | 0 | 0 | 1 | 1 | # | 10 | Small | ancom | 84 |
| # | time | 2 | 0 | 0 | 1 | 1 | # | 10 | Small | ancom | 84 |
| # | interaction | 0 | 0 | 2 | 0 | 0 | # | 10 | Small | ancom | 84 |
| # | disease | 4 | 0 | 0 | 1 | 1 | # | 10 | Small | sides2 | 84 |
| # | sex | 2 | 0 | 0 | 1 | 1 | # | 10 | Small | sides2 | 84 |
| # | time | 2 | 2 | 0 | 0.5 | 1 | # | 10 | Small | sides2 | 84 |
| # | interaction | 0 | 0 | 2 | 0 | 0 | # | 10 | Small | sides2 | 84 |
| # | disease | 4 | 3 | 0 | 0.571428571 | 1 | # | 10 | Baseline | paralle | 84 |
| # | sex | 2 | 5 | 0 | 0.285714286 | 1 | # | 10 | Baseline | paralle | 84 |
| # | time | 2 | 6 | 0 | 0.25 | 1 | # | 10 | Baseline | paralle | 84 |
| # | interaction | 2 | 7 | 0 | 0.222222222 | 1 | # | 10 | Baseline | paralle | 84 |
| # | disease | 4 | 2 | 0 | 0.666666667 | 1 | # | 10 | Baseline | ancom | 84 |
| # | sex | 2 | 0 | 0 | 1 | 1 | # | 10 | Baseline | ancom | 84 |
| # | time | 2 | 2 | 0 | 0.5 | 1 | # | 10 | Baseline | ancom | 84 |
| # | interaction | 0 | 0 | 2 | 0 | 0 | # | 10 | Baseline | ancom | 84 |
| # | disease | 4 | 14 | 0 | 0.222222222 | 1 | # | 10 | Baseline | sides2 | 84 |
| # | sex | 2 | 8 | 0 | 0.2 | 1 | # | 10 | Baseline | sides2 | 84 |
| # | time | 2 | 9 | 0 | 0.181818182 | 1 | # | 10 | Baseline | sides2 | 84 |
| # | interaction | 0 | 0 | 2 | 0 | 0 | # | 10 | Baseline | sides2 | 84 |
| # | disease | 4 | 0 | 0 | 1 | 1 | # | 10 | Small | paralle | 84 |
| # | sex | 2 | 0 | 0 | 1 | 1 | # | 10 | Small | paralle | 84 |
| # | time | 2 | 1 | 0 | 0.666666667 | 1 | # | 10 | Small | paralle | 84 |
| # | interaction | 2 | 0 | 0 | 1 | 1 | # | 10 | Small | paralle | 84 |
| # | disease | 4 | 2 | 0 | 0.666666667 | 1 | # | 10 | Small | ancom | 84 |
| # | sex | 2 | 0 | 0 | 1 | 1 | # | 10 | Small | ancom | 84 |
| # | time | 2 | 2 | 0 | 0.5 | 1 | # | 10 | Small | ancom | 84 |
| # | interaction | 0 | 0 | 2 | 0 | 0 | # | 10 | Small | ancom | 84 |
| # | disease | 4 | 2 | 0 | 0.666666667 | 1 | # | 10 | Small | sides2 | 84 |
| # | sex | 2 | 0 | 0 | 1 | 1 | # | 10 | Small | sides2 | 84 |
| # | time | 2 | 2 | 0 | 0.5 | 1 | # | 10 | Small | sides2 | 84 |
| # | interaction | 0 | 0 | 2 | 0 | 0 | # | 10 | Small | sides2 | 84 |
| # | disease | 4 | 4 | 0 | 0.5 | 1 | # | 10 | Baseline | paralle | 84 |
| # | sex | 2 | 1 | 0 | 0.666666667 | 1 | # | 10 | Baseline | paralle | 84 |
| # | time | 2 | 3 | 0 | 0.4 | 1 | # | 10 | Baseline | paralle | 84 |
| # | interaction | 2 | 3 | 0 | 0.4 | 1 | # | 10 | Baseline | paralle | 84 |
| # | disease | 4 | 2 | 0 | 0.666666667 | 1 | # | 10 | Baseline | ancom | 84 |
| # | sex | 2 | 0 | 0 | 1 | 1 | # | 10 | Baseline | ancom | 84 |
| # | time | 2 | 2 | 0 | 0.5 | 1 | # | 10 | Baseline | ancom | 84 |
| # | interaction | 0 | 0 | 2 | 0 | 0 | # | 10 | Baseline | ancom | 84 |
| # | disease | 4 | 19 | 0 | 0.173913043 | 1 | # | 10 | Baseline | sides2 | 84 |
| # | sex | 2 | 6 | 0 | 0.25 | 1 | # | 10 | Baseline | sides2 | 84 |
| # | time | 2 | 14 | 0 | 0.125 | 1 | # | 10 | Baseline | sides2 | 84 |
| # | interaction | 0 | 0 | 2 | 0 | 0 | # | 10 | Baseline | sides2 | 84 |
| # | disease | 4 | 0 | 0 | 1 | 1 | # | 10 | Small | paralle | 84 |
| # | sex | 2 | 0 | 0 | 1 | 1 | # | 10 | Small | paralle | 84 |
| # | time | 2 | 0 | 0 | 1 | 1 | # | 10 | Small | paralle | 84 |
| # | interaction | 2 | 0 | 0 | 1 | 1 | # | 10 | Small | paralle | 84 |
| # | disease | 4 | 2 | 0 | 0.666666667 | 1 | # | 10 | Small | ancom | 84 |
| # | sex | 2 | 0 | 0 | 1 | 1 | # | 10 | Small | ancom | 84 |
| # | time | 2 | 2 | 0 | 0.5 | 1 | # | 10 | Small | ancom | 84 |
| # | interaction | 0 | 0 | 2 | 0 | 0 | # | 10 | Small | ancom | 84 |
| # | disease | 4 | 2 | 0 | 0.666666667 | 1 | # | 10 | Small | sides2 | 84 |
| # | sex | 2 | 0 | 0 | 1 | 1 | # | 10 | Small | sides2 | 84 |
| # | time | 2 | 2 | 0 | 0.5 | 1 | # | 10 | Small | sides2 | 84 |
| # | interaction | 0 | 0 | 2 | 0 | 0 | # | 10 | Small | sides2 | 84 |
| # | disease | 7 | 11 | 0 | 0.388888889 | 1 | # | 20 | Baseline | paralle | 84 |
| # | sex | 4 | 9 | 0 | 0.307692308 | 1 | # | 20 | Baseline | paralle | 84 |
| # | time | 5 | 5 | 0 | 0.5 | 1 | # | 20 | Baseline | paralle | 84 |
| # | interaction | 4 | 12 | 0 | 0.25 | 1 | # | 20 | Baseline | paralle | 84 |
| # | disease | 7 | 1 | 0 | 0.875 | 1 | # | 20 | Baseline | ancom | 84 |
| # | sex | 4 | 0 | 0 | 1 | 1 | # | 20 | Baseline | ancom | 84 |
| # | time | 5 | 1 | 0 | 0.833333333 | 1 | # | 20 | Baseline | ancom | 84 |
| # | interaction | 0 | 0 | 4 | 0 | 0 | # | 20 | Baseline | ancom | 84 |
| # | disease | 7 | 23 | 0 | 0.233333333 | 1 | # | 20 | Baseline | sides2 | 84 |
| # | sex | 4 | 9 | 0 | 0.307692308 | 1 | # | 20 | Baseline | sides2 | 84 |
| # | time | 5 | 13 | 0 | 0.277777778 | 1 | # | 20 | Baseline | sides2 | 84 |
| # | interaction | 0 | 0 | 4 | 0 | 0 | # | 20 | Baseline | sides2 | 84 |
| # | disease | 7 | 0 | 0 | 1 | 1 | # | 20 | Small | paralle | 84 |
| # | sex | 4 | 1 | 0 | 0.8 | 1 | # | 20 | Small | paralle | 84 |
| # | time | 5 | 0 | 0 | 1 | 1 | # | 20 | Small | paralle | 84 |
| # | interaction | 2 | 0 | 2 | 1 | 0.5 | # | 20 | Small | paralle | 84 |
| # | disease | 7 | 1 | 0 | 0.875 | 1 | # | 20 | Small | ancom | 84 |
| # | sex | 4 | 0 | 0 | 1 | 1 | # | 20 | Small | ancom | 84 |
| # | time | 5 | 1 | 0 | 0.833333333 | 1 | # | 20 | Small | ancom | 84 |
| # | interaction | 0 | 0 | 4 | 0 | 0 | # | 20 | Small | ancom | 84 |
| # | disease | 7 | 2 | 0 | 0.777777778 | 1 | # | 20 | Small | sides2 | 84 |
| # | sex | 4 | 0 | 0 | 1 | 1 | # | 20 | Small | sides2 | 84 |
| # | time | 5 | 1 | 0 | 0.833333333 | 1 | # | 20 | Small | sides2 | 84 |
| # | interaction | 0 | 0 | 4 | 0 | 0 | # | 20 | Small | sides2 | 84 |
| # | disease | 7 | 8 | 0 | 0.466666667 | 1 | # | 20 | Baseline | paralle | 84 |
| # | sex | 4 | 4 | 0 | 0.5 | 1 | # | 20 | Baseline | paralle | 84 |
| # | time | 5 | 11 | 0 | 0.3125 | 1 | # | 20 | Baseline | paralle | 84 |
| # | interaction | 4 | 8 | 0 | 0.333333333 | 1 | # | 20 | Baseline | paralle | 84 |
| # | disease | 7 | 4 | 0 | 0.636363636 | 1 | # | 20 | Baseline | ancom | 84 |
| # | sex | 4 | 0 | 0 | 1 | 1 | # | 20 | Baseline | ancom | 84 |
| # | time | 5 | 2 | 0 | 0.714285714 | 1 | # | 20 | Baseline | ancom | 84 |
| # | interaction | 0 | 0 | 4 | 0 | 0 | # | 20 | Baseline | ancom | 84 |
| # | disease | 7 | 33 | 0 | 0.175 | 1 | # | 20 | Baseline | sides2 | 84 |
| # | sex | 4 | 9 | 0 | 0.307692308 | 1 | # | 20 | Baseline | sides2 | 84 |
| # | time | 5 | 19 | 0 | 0.208333333 | 1 | # | 20 | Baseline | sides2 | 84 |
| # | interaction | 0 | 0 | 4 | 0 | 0 | # | 20 | Baseline | sides2 | 84 |
| # | disease | 7 | 0 | 0 | 1 | 1 | # | 20 | Small | paralle | 84 |
| # | sex | 4 | 0 | 0 | 1 | 1 | # | 20 | Small | paralle | 84 |
| # | time | 5 | 0 | 0 | 1 | 1 | # | 20 | Small | paralle | 84 |
| # | interaction | 3 | 0 | 1 | 1 | 0.75 | # | 20 | Small | paralle | 84 |
| # | disease | 7 | 4 | 0 | 0.636363636 | 1 | # | 20 | Small | ancom | 84 |
| # | sex | 4 | 0 | 0 | 1 | 1 | # | 20 | Small | ancom | 84 |
| # | time | 5 | 2 | 0 | 0.714285714 | 1 | # | 20 | Small | ancom | 84 |
| # | interaction | 0 | 0 | 4 | 0 | 0 | # | 20 | Small | ancom | 84 |
| # | disease | 7 | 4 | 0 | 0.636363636 | 1 | # | 20 | Small | sides2 | 84 |
| # | sex | 4 | 0 | 0 | 1 | 1 | # | 20 | Small | sides2 | 84 |
| # | time | 5 | 2 | 0 | 0.714285714 | 1 | # | 20 | Small | sides2 | 84 |
| # | interaction | 0 | 0 | 4 | 0 | 0 | # | 20 | Small | sides2 | 84 |
| # | disease | 7 | 13 | 0 | 0.35 | 1 | # | 20 | Baseline | paralle | 84 |
| # | sex | 4 | 3 | 0 | 0.571428571 | 1 | # | 20 | Baseline | paralle | 84 |
| # | time | 5 | 11 | 0 | 0.3125 | 1 | # | 20 | Baseline | paralle | 84 |
| # | interaction | 4 | 7 | 0 | 0.363636364 | 1 | # | 20 | Baseline | paralle | 84 |
| # | disease | 7 | 4 | 0 | 0.636363636 | 1 | # | 20 | Baseline | ancom | 84 |
| # | sex | 4 | 0 | 0 | 1 | 1 | # | 20 | Baseline | ancom | 84 |
| # | time | 5 | 3 | 0 | 0.625 | 1 | # | 20 | Baseline | ancom | 84 |
| # | interaction | 0 | 0 | 4 | 0 | 0 | # | 20 | Baseline | ancom | 84 |
| # | disease | 7 | 50 | 0 | 0.122807018 | 1 | # | 20 | Baseline | sides2 | 84 |
| # | sex | 4 | 11 | 0 | 0.266666667 | 1 | # | 20 | Baseline | sides2 | 84 |
| # | time | 5 | 27 | 0 | 0.15625 | 1 | # | 20 | Baseline | sides2 | 84 |
| # | interaction | 0 | 0 | 4 | 0 | 0 | # | 20 | Baseline | sides2 | 84 |
| # | disease | 7 | 0 | 0 | 1 | 1 | # | 20 | Small | paralle | 84 |
| # | sex | 4 | 0 | 0 | 1 | 1 | # | 20 | Small | paralle | 84 |
| # | time | 5 | 0 | 0 | 1 | 1 | # | 20 | Small | paralle | 84 |
| # | interaction | 4 | 0 | 0 | 1 | 1 | # | 20 | Small | paralle | 84 |
| # | disease | 7 | 4 | 0 | 0.636363636 | 1 | # | 20 | Small | ancom | 84 |
| # | sex | 4 | 0 | 0 | 1 | 1 | # | 20 | Small | ancom | 84 |
| # | time | 5 | 2 | 0 | 0.714285714 | 1 | # | 20 | Small | ancom | 84 |
| # | interaction | 0 | 0 | 4 | 0 | 0 | # | 20 | Small | ancom | 84 |
| # | disease | 7 | 6 | 0 | 0.538461538 | 1 | # | 20 | Small | sides2 | 84 |
| # | sex | 4 | 0 | 0 | 1 | 1 | # | 20 | Small | sides2 | 84 |
| # | time | 5 | 4 | 0 | 0.555555556 | 1 | # | 20 | Small | sides2 | 84 |
| # | interaction | 0 | 0 | 4 | 0 | 0 | # | 20 | Small | sides2 | 84 |
| # | disease | 7 | 13 | 0 | 0.35 | 1 | # | 20 | Baseline | paralle | 84 |
| # | sex | 4 | 3 | 0 | 0.571428571 | 1 | # | 20 | Baseline | paralle | 84 |
| # | time | 5 | 10 | 0 | 0.333333333 | 1 | # | 20 | Baseline | paralle | 84 |
| # | interaction | 4 | 6 | 0 | 0.4 | 1 | # | 20 | Baseline | paralle | 84 |
| # | disease | 7 | 4 | 0 | 0.636363636 | 1 | # | 20 | Baseline | ancom | 84 |
| # | sex | 4 | 0 | 0 | 1 | 1 | # | 20 | Baseline | ancom | 84 |
| # | time | 5 | 3 | 0 | 0.625 | 1 | # | 20 | Baseline | ancom | 84 |
| # | interaction | 0 | 0 | 4 | 0 | 0 | # | 20 | Baseline | ancom | 84 |
| # | disease | 7 | 92 | 0 | 0.101449275 | 1 | # | 20 | Baseline | sides2 | 84 |
| # | sex | 4 | 12 | 0 | 0.25 | 1 | # | 20 | Baseline | sides2 | 84 |
| # | time | 5 | 35 | 0 | 0.125 | 1 | # | 20 | Baseline | sides2 | 84 |
| # | interaction | 0 | 0 | 4 | 0 | 0 | # | 20 | Baseline | sides2 | 84 |
| # | disease | 7 | 1 | 0 | 0.875 | 1 | # | 20 | Small | paralle | 84 |
| # | sex | 4 | 0 | 0 | 1 | 1 | # | 20 | Small | paralle | 84 |
| # | time | 5 | 1 | 0 | 0.833333333 | 1 | # | 20 | Small | paralle | 84 |
| # | interaction | 4 | 0 | 0 | 1 | 1 | # | 20 | Small | paralle | 84 |
| # | disease | 7 | 4 | 0 | 0.636363636 | 1 | # | 20 | Small | ancom | 84 |
| # | sex | 4 | 0 | 0 | 1 | 1 | # | 20 | Small | ancom | 84 |
| # | time | 5 | 2 | 0 | 0.714285714 | 1 | # | 20 | Small | ancom | 84 |
| # | interaction | 0 | 0 | 4 | 0 | 0 | # | 20 | Small | ancom | 84 |
| # | disease | 7 | 11 | 0 | 0.388888889 | 1 | # | 20 | Small | sides2 | 84 |
| # | sex | 4 | 0 | 0 | 1 | 1 | # | 20 | Small | sides2 | 84 |
| # | time | 5 | 3 | 0 | 0.625 | 1 | # | 20 | Small | sides2 | 84 |
| # | interaction | 0 | 0 | 4 | 0 | 0 | # | 20 | Small | sides2 | 84 |
| # | disease | 4 | 9 | 0 | 0.307692308 | 1 | # | 10 | Baseline | paralle | 85 |
| # | sex | 2 | 7 | 0 | 0.222222222 | 1 | # | 10 | Baseline | paralle | 85 |
| # | time | 2 | 6 | 0 | 0.25 | 1 | # | 10 | Baseline | paralle | 85 |
| # | interaction | 1 | 3 | 1 | 0.25 | 0.5 | # | 10 | Baseline | paralle | 85 |
| # | disease | 4 | 0 | 0 | 1 | 1 | # | 10 | Baseline | ancom | 85 |
| # | sex | 2 | 0 | 0 | 1 | 1 | # | 10 | Baseline | ancom | 85 |
| # | time | 2 | 0 | 0 | 1 | 1 | # | 10 | Baseline | ancom | 85 |
| # | interaction | 0 | 0 | 2 | 0 | 0 | # | 10 | Baseline | ancom | 85 |
| # | disease | 4 | 6 | 0 | 0.4 | 1 | # | 10 | Baseline | sides2 | 85 |
| # | sex | 2 | 4 | 0 | 0.333333333 | 1 | # | 10 | Baseline | sides2 | 85 |
| # | time | 2 | 8 | 0 | 0.2 | 1 | # | 10 | Baseline | sides2 | 85 |
| # | interaction | 0 | 0 | 2 | 0 | 0 | # | 10 | Baseline | sides2 | 85 |
| # | disease | 4 | 0 | 0 | 1 | 1 | # | 10 | Small | paralle | 85 |
| # | sex | 2 | 0 | 0 | 1 | 1 | # | 10 | Small | paralle | 85 |
| # | time | 2 | 0 | 0 | 1 | 1 | # | 10 | Small | paralle | 85 |
| # | interaction | 0 | 0 | 2 | 0 | 0 | # | 10 | Small | paralle | 85 |
| # | disease | 4 | 0 | 0 | 1 | 1 | # | 10 | Small | ancom | 85 |
| # | sex | 2 | 0 | 0 | 1 | 1 | # | 10 | Small | ancom | 85 |
| # | time | 2 | 0 | 0 | 1 | 1 | # | 10 | Small | ancom | 85 |
| # | interaction | 0 | 0 | 2 | 0 | 0 | # | 10 | Small | ancom | 85 |
| # | disease | 4 | 0 | 0 | 1 | 1 | # | 10 | Small | sides2 | 85 |
| # | sex | 2 | 0 | 0 | 1 | 1 | # | 10 | Small | sides2 | 85 |
| # | time | 2 | 0 | 0 | 1 | 1 | # | 10 | Small | sides2 | 85 |
| # | interaction | 0 | 0 | 2 | 0 | 0 | # | 10 | Small | sides2 | 85 |
| # | disease | 4 | 4 | 0 | 0.5 | 1 | # | 10 | Baseline | paralle | 85 |
| # | sex | 2 | 3 | 0 | 0.4 | 1 | # | 10 | Baseline | paralle | 85 |
| # | time | 2 | 6 | 0 | 0.25 | 1 | # | 10 | Baseline | paralle | 85 |
| # | interaction | 2 | 2 | 0 | 0.5 | 1 | # | 10 | Baseline | paralle | 85 |
| # | disease | 4 | 2 | 0 | 0.666666667 | 1 | # | 10 | Baseline | ancom | 85 |
| # | sex | 2 | 0 | 0 | 1 | 1 | # | 10 | Baseline | ancom | 85 |
| # | time | 2 | 0 | 0 | 1 | 1 | # | 10 | Baseline | ancom | 85 |
| # | interaction | 0 | 0 | 2 | 0 | 0 | # | 10 | Baseline | ancom | 85 |
| # | disease | 4 | 16 | 0 | 0.2 | 1 | # | 10 | Baseline | sides2 | 85 |
| # | sex | 2 | 4 | 0 | 0.333333333 | 1 | # | 10 | Baseline | sides2 | 85 |
| # | time | 2 | 10 | 0 | 0.166666667 | 1 | # | 10 | Baseline | sides2 | 85 |
| # | interaction | 0 | 0 | 2 | 0 | 0 | # | 10 | Baseline | sides2 | 85 |
| # | disease | 4 | 0 | 0 | 1 | 1 | # | 10 | Small | paralle | 85 |
| # | sex | 2 | 0 | 0 | 1 | 1 | # | 10 | Small | paralle | 85 |
| # | time | 2 | 1 | 0 | 0.666666667 | 1 | # | 10 | Small | paralle | 85 |
| # | interaction | 0 | 1 | 2 | 0 | 0 | # | 10 | Small | paralle | 85 |
| # | disease | 4 | 1 | 0 | 0.8 | 1 | # | 10 | Small | ancom | 85 |
| # | sex | 2 | 0 | 0 | 1 | 1 | # | 10 | Small | ancom | 85 |
| # | time | 2 | 0 | 0 | 1 | 1 | # | 10 | Small | ancom | 85 |
| # | interaction | 0 | 0 | 2 | 0 | 0 | # | 10 | Small | ancom | 85 |
| # | disease | 4 | 1 | 0 | 0.8 | 1 | # | 10 | Small | sides2 | 85 |

| | | | | | | | | | | | |
|---|---|---|---|---|---|---|---|---|---|---|---|
| # | sex | 2 | 0 | 0 | 1 | 1 | # | 10 | Small | aldex2 | 85 |
| # | time | 2 | 0 | 0 | 1 | 1 | # | 10 | Small | aldex2 | 85 |
| # | interaction | 0 | 0 | 2 | 0 | 0 | # | 10 | Small | aldex2 | 85 |
| # | disease | 4 | 1 | 0 | 0.8 | 1 | # | 10 | Baseline | paralle | 85 |
| # | sex | 2 | 3 | 0 | 0.4 | 1 | # | 10 | Baseline | paralle | 85 |
| # | time | 2 | 5 | 0 | 0.285714286 | 1 | # | 10 | Baseline | paralle | 85 |
| # | interaction | 2 | 4 | 0 | 0.333333333 | 1 | # | 10 | Baseline | paralle | 85 |
| # | disease | 4 | 2 | 0 | 0.666666667 | 1 | # | 10 | Baseline | ancom | 85 |
| # | sex | 2 | 0 | 0 | 1 | 1 | # | 10 | Baseline | ancom | 85 |
| # | time | 2 | 1 | 0 | 0.666666667 | 1 | # | 10 | Baseline | ancom | 85 |
| # | interaction | 0 | 0 | 2 | 0 | 0 | # | 10 | Baseline | ancom | 85 |
| # | disease | 4 | 18 | 0 | 0.181818182 | 1 | # | 10 | Baseline | aldex2 | 85 |
| # | sex | 2 | 7 | 0 | 0.222222222 | 1 | # | 10 | Baseline | aldex2 | 85 |
| # | time | 2 | 12 | 0 | 0.142857143 | 1 | # | 10 | Baseline | aldex2 | 85 |
| # | interaction | 0 | 0 | 2 | 0 | 0 | # | 10 | Baseline | aldex2 | 85 |
| # | disease | 4 | 0 | 0 | 1 | 1 | # | 10 | Small | paralle | 85 |
| # | sex | 2 | 0 | 0 | 1 | 1 | # | 10 | Small | paralle | 85 |
| # | time | 2 | 0 | 0 | 1 | 1 | # | 10 | Small | paralle | 85 |
| # | interaction | 2 | 0 | 0 | 1 | 1 | # | 10 | Small | paralle | 85 |
| # | disease | 4 | 2 | 0 | 0.666666667 | 1 | # | 10 | Small | ancom | 85 |
| # | sex | 2 | 0 | 0 | 1 | 1 | # | 10 | Small | ancom | 85 |
| # | time | 2 | 1 | 0 | 0.666666667 | 1 | # | 10 | Small | ancom | 85 |
| # | interaction | 0 | 0 | 2 | 0 | 0 | # | 10 | Small | ancom | 85 |
| # | disease | 4 | 1 | 0 | 0.8 | 1 | # | 10 | Small | aldex2 | 85 |
| # | sex | 2 | 0 | 0 | 1 | 1 | # | 10 | Small | aldex2 | 85 |
| # | time | 2 | 2 | 0 | 0.5 | 1 | # | 10 | Small | aldex2 | 85 |
| # | interaction | 0 | 0 | 2 | 0 | 0 | # | 10 | Small | aldex2 | 85 |
| # | disease | 4 | 5 | 0 | 0.444444444 | 1 | # | 10 | Baseline | paralle | 85 |
| # | sex | 2 | 3 | 0 | 0.4 | 1 | # | 10 | Baseline | paralle | 85 |
| # | time | 2 | 3 | 0 | 0.4 | 1 | # | 10 | Baseline | paralle | 85 |
| # | interaction | 2 | 1 | 0 | 0.666666667 | 1 | # | 10 | Baseline | paralle | 85 |
| # | disease | 4 | 2 | 0 | 0.666666667 | 1 | # | 10 | Baseline | ancom | 85 |
| # | sex | 2 | 0 | 0 | 1 | 1 | # | 10 | Baseline | ancom | 85 |
| # | time | 2 | 2 | 0 | 0.5 | 1 | # | 10 | Baseline | ancom | 85 |
| # | interaction | 0 | 0 | 2 | 0 | 0 | # | 10 | Baseline | ancom | 85 |
| # | disease | 4 | 18 | 0 | 0.181818182 | 1 | # | 10 | Baseline | aldex2 | 85 |
| # | sex | 2 | 7 | 0 | 0.222222222 | 1 | # | 10 | Baseline | aldex2 | 85 |
| # | time | 2 | 16 | 0 | 0.111111111 | 1 | # | 10 | Baseline | aldex2 | 85 |
| # | interaction | 0 | 0 | 2 | 0 | 0 | # | 10 | Baseline | aldex2 | 85 |
| # | disease | 4 | 0 | 0 | 1 | 1 | # | 10 | Small | paralle | 85 |
| # | sex | 2 | 0 | 0 | 1 | 1 | # | 10 | Small | paralle | 85 |
| # | time | 2 | 0 | 0 | 1 | 1 | # | 10 | Small | paralle | 85 |
| # | interaction | 2 | 0 | 0 | 1 | 1 | # | 10 | Small | paralle | 85 |
| # | disease | 4 | 2 | 0 | 0.666666667 | 1 | # | 10 | Small | ancom | 85 |
| # | sex | 2 | 0 | 0 | 1 | 1 | # | 10 | Small | ancom | 85 |
| # | time | 2 | 2 | 0 | 0.5 | 1 | # | 10 | Small | ancom | 85 |
| # | interaction | 0 | 0 | 2 | 0 | 0 | # | 10 | Small | ancom | 85 |
| # | disease | 4 | 2 | 0 | 0.666666667 | 1 | # | 10 | Small | aldex2 | 85 |
| # | sex | 2 | 0 | 0 | 1 | 1 | # | 10 | Small | aldex2 | 85 |
| # | time | 2 | 2 | 0 | 0.5 | 1 | # | 10 | Small | aldex2 | 85 |
| # | interaction | 0 | 0 | 2 | 0 | 0 | # | 10 | Small | aldex2 | 85 |
| # | disease | 7 | 9 | 0 | 0.4375 | 1 | # | 20 | Baseline | paralle | 85 |
| # | sex | 4 | 5 | 0 | 0.444444444 | 1 | # | 20 | Baseline | paralle | 85 |
| # | time | 5 | 3 | 0 | 0.625 | 1 | # | 20 | Baseline | paralle | 85 |
| # | interaction | 4 | 6 | 0 | 0.4 | 1 | # | 20 | Baseline | paralle | 85 |
| # | disease | 7 | 1 | 0 | 0.875 | 1 | # | 20 | Baseline | ancom | 85 |
| # | sex | 4 | 0 | 0 | 1 | 1 | # | 20 | Baseline | ancom | 85 |
| # | time | 5 | 2 | 0 | 0.714285714 | 1 | # | 20 | Baseline | ancom | 85 |
| # | interaction | 0 | 0 | 4 | 0 | 0 | # | 20 | Baseline | ancom | 85 |
| # | disease | 7 | 15 | 0 | 0.318181818 | 1 | # | 20 | Baseline | aldex2 | 85 |
| # | sex | 4 | 8 | 0 | 0.333333333 | 1 | # | 20 | Baseline | aldex2 | 85 |
| # | time | 5 | 19 | 0 | 0.208333333 | 1 | # | 20 | Baseline | aldex2 | 85 |
| # | interaction | 0 | 0 | 4 | 0 | 0 | # | 20 | Baseline | aldex2 | 85 |
| # | disease | 7 | 0 | 0 | 1 | 1 | # | 20 | Small | paralle | 85 |
| # | sex | 4 | 0 | 0 | 1 | 1 | # | 20 | Small | paralle | 85 |
| # | time | 5 | 0 | 0 | 1 | 1 | # | 20 | Small | paralle | 85 |
| # | interaction | 0 | 0 | 4 | 0 | 0 | # | 20 | Small | paralle | 85 |
| # | disease | 7 | 1 | 0 | 0.875 | 1 | # | 20 | Small | ancom | 85 |
| # | sex | 4 | 0 | 0 | 1 | 1 | # | 20 | Small | ancom | 85 |
| # | time | 5 | 2 | 0 | 0.714285714 | 1 | # | 20 | Small | ancom | 85 |
| # | interaction | 0 | 0 | 4 | 0 | 0 | # | 20 | Small | ancom | 85 |
| # | disease | 7 | 0 | 0 | 1 | 1 | # | 20 | Small | aldex2 | 85 |
| # | sex | 4 | 0 | 0 | 1 | 1 | # | 20 | Small | aldex2 | 85 |
| # | time | 5 | 2 | 0 | 0.714285714 | 1 | # | 20 | Small | aldex2 | 85 |
| # | interaction | 0 | 0 | 4 | 0 | 0 | # | 20 | Small | aldex2 | 85 |
| # | disease | 7 | 9 | 0 | 0.4375 | 1 | # | 20 | Baseline | paralle | 85 |
| # | sex | 4 | 7 | 0 | 0.363636364 | 1 | # | 20 | Baseline | paralle | 85 |
| # | time | 5 | 1 | 0 | 0.833333333 | 1 | # | 20 | Baseline | paralle | 85 |
| # | interaction | 4 | 4 | 0 | 0.5 | 1 | # | 20 | Baseline | paralle | 85 |
| # | disease | 7 | 4 | 0 | 0.636363636 | 1 | # | 20 | Baseline | ancom | 85 |
| # | sex | 4 | 0 | 0 | 1 | 1 | # | 20 | Baseline | ancom | 85 |
| # | time | 5 | 4 | 0 | 0.555555556 | 1 | # | 20 | Baseline | ancom | 85 |
| # | interaction | 0 | 0 | 4 | 0 | 0 | # | 20 | Baseline | ancom | 85 |
| # | disease | 7 | 29 | 0 | 0.194444444 | 1 | # | 20 | Baseline | aldex2 | 85 |
| # | sex | 4 | 13 | 0 | 0.235294118 | 1 | # | 20 | Baseline | aldex2 | 85 |
| # | time | 5 | 28 | 0 | 0.151515152 | 1 | # | 20 | Baseline | aldex2 | 85 |
| # | interaction | 0 | 0 | 4 | 0 | 0 | # | 20 | Baseline | aldex2 | 85 |
| # | disease | 7 | 0 | 0 | 1 | 1 | # | 20 | Small | paralle | 85 |
| # | sex | 4 | 0 | 0 | 1 | 1 | # | 20 | Small | paralle | 85 |
| # | time | 5 | 0 | 0 | 1 | 1 | # | 20 | Small | paralle | 85 |
| # | interaction | 4 | 0 | 0 | 1 | 1 | # | 20 | Small | paralle | 85 |
| # | disease | 7 | 4 | 0 | 0.636363636 | 1 | # | 20 | Small | ancom | 85 |
| # | sex | 4 | 0 | 0 | 1 | 1 | # | 20 | Small | ancom | 85 |
| # | time | 5 | 4 | 0 | 0.555555556 | 1 | # | 20 | Small | ancom | 85 |
| # | interaction | 0 | 0 | 4 | 0 | 0 | # | 20 | Small | ancom | 85 |
| # | disease | 7 | 2 | 0 | 0.777777778 | 1 | # | 20 | Small | aldex2 | 85 |
| # | sex | 4 | 0 | 0 | 1 | 1 | # | 20 | Small | aldex2 | 85 |
| # | time | 5 | 3 | 0 | 0.625 | 1 | # | 20 | Small | aldex2 | 85 |
| # | interaction | 0 | 0 | 4 | 0 | 0 | # | 20 | Small | aldex2 | 85 |
| # | disease | 7 | 9 | 0 | 0.4375 | 1 | # | 20 | Baseline | paralle | 85 |
| # | sex | 4 | 4 | 0 | 0.5 | 1 | # | 20 | Baseline | paralle | 85 |
| # | time | 5 | 7 | 0 | 0.416666667 | 1 | # | 20 | Baseline | paralle | 85 |
| # | interaction | 4 | 4 | 0 | 0.5 | 1 | # | 20 | Baseline | paralle | 85 |
| # | disease | 7 | 4 | 0 | 0.636363636 | 1 | # | 20 | Baseline | ancom | 85 |
| # | sex | 4 | 0 | 0 | 1 | 1 | # | 20 | Baseline | ancom | 85 |
| # | time | 5 | 4 | 0 | 0.555555556 | 1 | # | 20 | Baseline | ancom | 85 |
| # | interaction | 0 | 0 | 4 | 0 | 0 | # | 20 | Baseline | ancom | 85 |
| # | disease | 7 | 67 | 0 | 0.12962963 | 1 | # | 20 | Baseline | aldex2 | 85 |
| # | sex | 4 | 17 | 0 | 0.19047619 | 1 | # | 20 | Baseline | aldex2 | 85 |
| # | time | 5 | 36 | 0 | 0.12195122 | 1 | # | 20 | Baseline | aldex2 | 85 |
| # | interaction | 0 | 0 | 4 | 0 | 0 | # | 20 | Baseline | aldex2 | 85 |
| # | disease | 7 | 0 | 0 | 1 | 1 | # | 20 | Small | paralle | 85 |
| # | sex | 4 | 0 | 0 | 1 | 1 | # | 20 | Small | paralle | 85 |
| # | time | 5 | 0 | 0 | 1 | 1 | # | 20 | Small | paralle | 85 |
| # | interaction | 4 | 0 | 0 | 1 | 1 | # | 20 | Small | paralle | 85 |
| # | disease | 7 | 4 | 0 | 0.636363636 | 1 | # | 20 | Small | ancom | 85 |
| # | sex | 4 | 0 | 0 | 1 | 1 | # | 20 | Small | ancom | 85 |
| # | time | 5 | 4 | 0 | 0.555555556 | 1 | # | 20 | Small | ancom | 85 |
| # | interaction | 0 | 0 | 4 | 0 | 0 | # | 20 | Small | ancom | 85 |
| # | disease | 7 | 7 | 0 | 0.5 | 1 | # | 20 | Small | aldex2 | 85 |
| # | sex | 4 | 0 | 0 | 1 | 1 | # | 20 | Small | aldex2 | 85 |
| # | time | 5 | 5 | 0 | 0.5 | 1 | # | 20 | Small | aldex2 | 85 |
| # | interaction | 0 | 0 | 4 | 0 | 0 | # | 20 | Small | aldex2 | 85 |
| # | disease | 7 | 6 | 0 | 0.538461538 | 1 | # | 20 | Baseline | paralle | 85 |
| # | sex | 4 | 2 | 0 | 0.666666667 | 1 | # | 20 | Baseline | paralle | 85 |
| # | time | 5 | 2 | 0 | 0.714285714 | 1 | # | 20 | Baseline | paralle | 85 |
| # | interaction | 4 | 2 | 0 | 0.666666667 | 1 | # | 20 | Baseline | paralle | 85 |
| # | disease | 7 | 4 | 0 | 0.636363636 | 1 | # | 20 | Baseline | ancom | 85 |
| # | sex | 4 | 0 | 0 | 1 | 1 | # | 20 | Baseline | ancom | 85 |
| # | time | 5 | 4 | 0 | 0.555555556 | 1 | # | 20 | Baseline | ancom | 85 |
| # | interaction | 0 | 0 | 4 | 0 | 0 | # | 20 | Baseline | ancom | 85 |
| # | disease | 7 | 91 | 0 | 0.102941176 | 1 | # | 20 | Baseline | aldex2 | 85 |
| # | sex | 4 | 18 | 0 | 0.181818182 | 1 | # | 20 | Baseline | aldex2 | 85 |
| # | time | 5 | 62 | 0 | 0.106382979 | 1 | # | 20 | Baseline | aldex2 | 85 |
| # | interaction | 0 | 0 | 4 | 0 | 0 | # | 20 | Baseline | aldex2 | 85 |
| # | disease | 7 | 1 | 0 | 0.875 | 1 | # | 20 | Small | paralle | 85 |
| # | sex | 4 | 0 | 0 | 1 | 1 | # | 20 | Small | paralle | 85 |
| # | time | 5 | 0 | 0 | 1 | 1 | # | 20 | Small | paralle | 85 |
| # | interaction | 4 | 0 | 0 | 1 | 1 | # | 20 | Small | paralle | 85 |
| # | disease | 7 | 4 | 0 | 0.636363636 | 1 | # | 20 | Small | ancom | 85 |
| # | sex | 4 | 0 | 0 | 1 | 1 | # | 20 | Small | ancom | 85 |
| # | time | 5 | 4 | 0 | 0.555555556 | 1 | # | 20 | Small | ancom | 85 |
| # | interaction | 0 | 0 | 4 | 0 | 0 | # | 20 | Small | ancom | 85 |
| # | disease | 7 | 7 | 0 | 0.5 | 1 | # | 20 | Small | aldex2 | 85 |
| # | sex | 4 | 0 | 0 | 1 | 1 | # | 20 | Small | aldex2 | 85 |
| # | time | 5 | 4 | 0 | 0.555555556 | 1 | # | 20 | Small | aldex2 | 85 |
| # | interaction | 0 | 0 | 4 | 0 | 0 | # | 20 | Small | aldex2 | 85 |
| # | disease | 4 | 6 | 0 | 0.4 | 1 | # | 10 | Baseline | paralle | 86 |
| # | sex | 2 | 13 | 0 | 0.133333333 | 1 | # | 10 | Baseline | paralle | 86 |
| # | time | 2 | 9 | 0 | 0.181818182 | 1 | # | 10 | Baseline | paralle | 86 |
| # | interaction | 2 | 7 | 0 | 0.222222222 | 1 | # | 10 | Baseline | paralle | 86 |
| # | disease | 4 | 0 | 0 | 1 | 1 | # | 10 | Baseline | ancom | 86 |
| # | sex | 2 | 0 | 0 | 1 | 1 | # | 10 | Baseline | ancom | 86 |
| # | time | 2 | 0 | 0 | 1 | 1 | # | 10 | Baseline | ancom | 86 |
| # | interaction | 0 | 0 | 2 | 0 | 0 | # | 10 | Baseline | ancom | 86 |
| # | disease | 4 | 12 | 0 | 0.25 | 1 | # | 10 | Baseline | aldex2 | 86 |
| # | sex | 2 | 7 | 0 | 0.222222222 | 1 | # | 10 | Baseline | aldex2 | 86 |
| # | time | 2 | 10 | 0 | 0.166666667 | 1 | # | 10 | Baseline | aldex2 | 86 |
| # | interaction | 0 | 0 | 2 | 0 | 0 | # | 10 | Baseline | aldex2 | 86 |
| # | disease | 4 | 0 | 0 | 1 | 1 | # | 10 | Small | paralle | 86 |
| # | sex | 2 | 0 | 0 | 1 | 1 | # | 10 | Small | paralle | 86 |
| # | time | 2 | 0 | 0 | 1 | 1 | # | 10 | Small | paralle | 86 |
| # | interaction | 2 | 0 | 0 | 1 | 1 | # | 10 | Small | paralle | 86 |
| # | disease | 4 | 0 | 0 | 1 | 1 | # | 10 | Small | ancom | 86 |
| # | sex | 2 | 0 | 0 | 1 | 1 | # | 10 | Small | ancom | 86 |
| # | time | 2 | 0 | 0 | 1 | 1 | # | 10 | Small | ancom | 86 |
| # | interaction | 0 | 0 | 2 | 0 | 0 | # | 10 | Small | ancom | 86 |
| # | disease | 4 | 1 | 0 | 0.8 | 1 | # | 10 | Small | aldex2 | 86 |
| # | sex | 2 | 0 | 0 | 1 | 1 | # | 10 | Small | aldex2 | 86 |
| # | time | 2 | 0 | 0 | 1 | 1 | # | 10 | Small | aldex2 | 86 |
| # | interaction | 0 | 0 | 2 | 0 | 0 | # | 10 | Small | aldex2 | 86 |
| # | disease | 4 | 6 | 0 | 0.4 | 1 | # | 10 | Baseline | paralle | 86 |
| # | sex | 2 | 10 | 0 | 0.166666667 | 1 | # | 10 | Baseline | paralle | 86 |
| # | time | 2 | 9 | 0 | 0.181818182 | 1 | # | 10 | Baseline | paralle | 86 |
| # | interaction | 2 | 6 | 0 | 0.25 | 1 | # | 10 | Baseline | paralle | 86 |
| # | disease | 4 | 2 | 0 | 0.666666667 | 1 | # | 10 | Baseline | ancom | 86 |
| # | sex | 2 | 0 | 0 | 1 | 1 | # | 10 | Baseline | ancom | 86 |
| # | time | 2 | 2 | 0 | 0.5 | 1 | # | 10 | Baseline | ancom | 86 |
| # | interaction | 0 | 0 | 2 | 0 | 0 | # | 10 | Baseline | ancom | 86 |
| # | disease | 4 | 18 | 0 | 0.181818182 | 1 | # | 10 | Baseline | aldex2 | 86 |
| # | sex | 2 | 5 | 0 | 0.285714286 | 1 | # | 10 | Baseline | aldex2 | 86 |
| # | time | 2 | 7 | 0 | 0.222222222 | 1 | # | 10 | Baseline | aldex2 | 86 |
| # | interaction | 0 | 0 | 2 | 0 | 0 | # | 10 | Baseline | aldex2 | 86 |
| # | disease | 4 | 1 | 0 | 0.8 | 1 | # | 10 | Small | paralle | 86 |
| # | sex | 2 | 0 | 0 | 1 | 1 | # | 10 | Small | paralle | 86 |
| # | time | 2 | 0 | 0 | 1 | 1 | # | 10 | Small | paralle | 86 |
| # | interaction | 2 | 1 | 0 | 0.666666667 | 1 | # | 10 | Small | paralle | 86 |
| # | disease | 4 | 1 | 0 | 0.8 | 1 | # | 10 | Small | ancom | 86 |
| # | sex | 2 | 0 | 0 | 1 | 1 | # | 10 | Small | ancom | 86 |
| # | time | 2 | 2 | 0 | 0.5 | 1 | # | 10 | Small | ancom | 86 |
| # | interaction | 0 | 0 | 2 | 0 | 0 | # | 10 | Small | ancom | 86 |
| # | disease | 4 | 1 | 0 | 0.8 | 1 | # | 10 | Small | aldex2 | 86 |
| # | sex | 2 | 1 | 0 | 0.666666667 | 1 | # | 10 | Small | aldex2 | 86 |
| # | time | 2 | 2 | 0 | 0.5 | 1 | # | 10 | Small | aldex2 | 86 |
| # | interaction | 0 | 0 | 2 | 0 | 0 | # | 10 | Small | aldex2 | 86 |
| # | disease | 4 | 8 | 0 | 0.333333333 | 1 | # | 10 | Baseline | paralle | 86 |
| # | sex | 2 | 5 | 0 | 0.285714286 | 1 | # | 10 | Baseline | paralle | 86 |
| # | time | 2 | 6 | 0 | 0.25 | 1 | # | 10 | Baseline | paralle | 86 |
| # | interaction | 2 | 4 | 0 | 0.333333333 | 1 | # | 10 | Baseline | paralle | 86 |
| # | disease | 4 | 2 | 0 | 0.666666667 | 1 | # | 10 | Baseline | ancom | 86 |
| # | sex | 2 | 0 | 0 | 1 | 1 | # | 10 | Baseline | ancom | 86 |
| # | time | 2 | 2 | 0 | 0.5 | 1 | # | 10 | Baseline | ancom | 86 |
| # | interaction | 0 | 0 | 2 | 0 | 0 | # | 10 | Baseline | ancom | 86 |
| # | disease | 4 | 18 | 0 | 0.181818182 | 1 | # | 10 | Baseline | aldex2 | 86 |
| # | sex | 2 | 6 | 0 | 0.25 | 1 | # | 10 | Baseline | aldex2 | 86 |
| # | time | 2 | 7 | 0 | 0.222222222 | 1 | # | 10 | Baseline | aldex2 | 86 |
| # | interaction | 0 | 0 | 2 | 0 | 0 | # | 10 | Baseline | aldex2 | 86 |
| # | disease | 4 | 0 | 0 | 1 | 1 | # | 10 | Small | paralle | 86 |
| # | sex | 2 | 0 | 0 | 1 | 1 | # | 10 | Small | paralle | 86 |
| # | time | 2 | 0 | 0 | 1 | 1 | # | 10 | Small | paralle | 86 |
| # | interaction | 2 | 0 | 0 | 1 | 1 | # | 10 | Small | paralle | 86 |
| # | disease | 4 | 2 | 0 | 0.666666667 | 1 | # | 10 | Small | ancom | 86 |
| # | sex | 2 | 0 | 0 | 1 | 1 | # | 10 | Small | ancom | 86 |
| # | time | 2 | 2 | 0 | 0.5 | 1 | # | 10 | Small | ancom | 86 |
| # | interaction | 0 | 0 | 2 | 0 | 0 | # | 10 | Small | ancom | 86 |
| # | disease | 4 | 2 | 0 | 0.666666667 | 1 | # | 10 | Small | aldex2 | 86 |
| # | sex | 2 | 0 | 0 | 1 | 1 | # | 10 | Small | aldex2 | 86 |
| # | time | 2 | 2 | 0 | 0.5 | 1 | # | 10 | Small | aldex2 | 86 |
| # | interaction | 0 | 0 | 2 | 0 | 0 | # | 10 | Small | aldex2 | 86 |
| # | disease | 4 | 6 | 0 | 0.4 | 1 | # | 10 | Baseline | paralle | 86 |
| # | sex | 2 | 6 | 0 | 0.25 | 1 | # | 10 | Baseline | paralle | 86 |
| # | time | 2 | 3 | 0 | 0.4 | 1 | # | 10 | Baseline | paralle | 86 |
| # | interaction | 2 | 4 | 0 | 0.333333333 | 1 | # | 10 | Baseline | paralle | 86 |
| # | disease | 4 | 2 | 0 | 0.666666667 | 1 | # | 10 | Baseline | ancom | 86 |
| # | sex | 2 | 0 | 0 | 1 | 1 | # | 10 | Baseline | ancom | 86 |
| # | time | 2 | 2 | 0 | 0.5 | 1 | # | 10 | Baseline | ancom | 86 |
| # | interaction | 0 | 0 | 2 | 0 | 0 | # | 10 | Baseline | ancom | 86 |
| # | disease | 4 | 24 | 0 | 0.142857143 | 1 | # | 10 | Baseline | aldex2 | 86 |
| # | sex | 2 | 9 | 0 | 0.181818182 | 1 | # | 10 | Baseline | aldex2 | 86 |
| # | time | 2 | 9 | 0 | 0.181818182 | 1 | # | 10 | Baseline | aldex2 | 86 |
| # | interaction | 0 | 0 | 2 | 0 | 0 | # | 10 | Baseline | aldex2 | 86 |
| # | disease | 4 | 1 | 0 | 0.8 | 1 | # | 10 | Small | paralle | 86 |
| # | sex | 2 | 0 | 0 | 1 | 1 | # | 10 | Small | paralle | 86 |
| # | time | 2 | 0 | 0 | 1 | 1 | # | 10 | Small | paralle | 86 |
| # | interaction | 2 | 0 | 0 | 1 | 1 | # | 10 | Small | paralle | 86 |
| # | disease | 4 | 2 | 0 | 0.666666667 | 1 | # | 10 | Small | ancom | 86 |
| # | sex | 2 | 0 | 0 | 1 | 1 | # | 10 | Small | ancom | 86 |
| # | time | 2 | 2 | 0 | 0.5 | 1 | # | 10 | Small | ancom | 86 |
| # | interaction | 0 | 0 | 2 | 0 | 0 | # | 10 | Small | ancom | 86 |
| # | disease | 4 | 5 | 0 | 0.444444444 | 1 | # | 10 | Small | aldex2 | 86 |
| # | sex | 2 | 1 | 0 | 0.666666667 | 1 | # | 10 | Small | aldex2 | 86 |
| # | time | 2 | 2 | 0 | 0.5 | 1 | # | 10 | Small | aldex2 | 86 |
| # | interaction | 0 | 0 | 2 | 0 | 0 | # | 10 | Small | aldex2 | 86 |
| # | disease | 7 | 11 | 0 | 0.388888889 | 1 | # | 20 | Baseline | paralle | 86 |
| # | sex | 4 | 5 | 0 | 0.444444444 | 1 | # | 20 | Baseline | paralle | 86 |
| # | time | 5 | 11 | 0 | 0.3125 | 1 | # | 20 | Baseline | paralle | 86 |
| # | interaction | 3 | 9 | 1 | 0.25 | 0.75 | # | 20 | Baseline | paralle | 86 |
| # | disease | 7 | 2 | 0 | 0.777777778 | 1 | # | 20 | Baseline | ancom | 86 |
| # | sex | 4 | 0 | 0 | 1 | 1 | # | 20 | Baseline | ancom | 86 |
| # | time | 5 | 1 | 0 | 0.833333333 | 1 | # | 20 | Baseline | ancom | 86 |
| # | interaction | 0 | 0 | 4 | 0 | 0 | # | 20 | Baseline | ancom | 86 |
| # | disease | 7 | 28 | 0 | 0.2 | 1 | # | 20 | Baseline | aldex2 | 86 |
| # | sex | 4 | 3 | 0 | 0.571428571 | 1 | # | 20 | Baseline | aldex2 | 86 |
| # | time | 5 | 19 | 0 | 0.208333333 | 1 | # | 20 | Baseline | aldex2 | 86 |
| # | interaction | 0 | 0 | 4 | 0 | 0 | # | 20 | Baseline | aldex2 | 86 |
| # | disease | 7 | 1 | 0 | 0.875 | 1 | # | 20 | Small | paralle | 86 |
| # | sex | 4 | 0 | 0 | 1 | 1 | # | 20 | Small | paralle | 86 |
| # | time | 5 | 0 | 0 | 1 | 1 | # | 20 | Small | paralle | 86 |
| # | interaction | 0 | 0 | 4 | 0 | 0 | # | 20 | Small | paralle | 86 |
| # | disease | 7 | 1 | 0 | 0.875 | 1 | # | 20 | Small | ancom | 86 |
| # | sex | 4 | 0 | 0 | 1 | 1 | # | 20 | Small | ancom | 86 |
| # | time | 5 | 1 | 0 | 0.833333333 | 1 | # | 20 | Small | ancom | 86 |
| # | interaction | 0 | 0 | 4 | 0 | 0 | # | 20 | Small | ancom | 86 |
| # | disease | 7 | 1 | 0 | 0.875 | 1 | # | 20 | Small | aldex2 | 86 |
| # | sex | 4 | 0 | 0 | 1 | 1 | # | 20 | Small | aldex2 | 86 |
| # | time | 5 | 1 | 0 | 0.833333333 | 1 | # | 20 | Small | aldex2 | 86 |
| # | interaction | 0 | 0 | 4 | 0 | 0 | # | 20 | Small | aldex2 | 86 |
| # | disease | 7 | 12 | 0 | 0.368421053 | 1 | # | 20 | Baseline | paralle | 86 |
| # | sex | 4 | 4 | 0 | 0.5 | 1 | # | 20 | Baseline | paralle | 86 |
| # | time | 5 | 2 | 0 | 0.714285714 | 1 | # | 20 | Baseline | paralle | 86 |
| # | interaction | 4 | 3 | 0 | 0.571428571 | 1 | # | 20 | Baseline | paralle | 86 |
| # | disease | 7 | 3 | 0 | 0.7 | 1 | # | 20 | Baseline | ancom | 86 |
| # | sex | 4 | 0 | 0 | 1 | 1 | # | 20 | Baseline | ancom | 86 |
| # | time | 5 | 3 | 0 | 0.625 | 1 | # | 20 | Baseline | ancom | 86 |
| # | interaction | 0 | 0 | 4 | 0 | 0 | # | 20 | Baseline | ancom | 86 |
| # | disease | 7 | 36 | 0 | 0.162790698 | 1 | # | 20 | Baseline | aldex2 | 86 |
| # | sex | 4 | 11 | 0 | 0.266666667 | 1 | # | 20 | Baseline | aldex2 | 86 |
| # | time | 5 | 21 | 0 | 0.192307692 | 1 | # | 20 | Baseline | aldex2 | 86 |
| # | interaction | 0 | 0 | 4 | 0 | 0 | # | 20 | Baseline | aldex2 | 86 |
| # | disease | 7 | 0 | 0 | 1 | 1 | # | 20 | Small | paralle | 86 |
| # | sex | 4 | 0 | 0 | 1 | 1 | # | 20 | Small | paralle | 86 |
| # | time | 5 | 0 | 0 | 1 | 1 | # | 20 | Small | paralle | 86 |
| # | interaction | 3 | 0 | 1 | 1 | 0.75 | # | 20 | Small | paralle | 86 |
| # | disease | 7 | 3 | 0 | 0.7 | 1 | # | 20 | Small | ancom | 86 |
| # | sex | 4 | 0 | 0 | 1 | 1 | # | 20 | Small | ancom | 86 |
| # | time | 5 | 3 | 0 | 0.625 | 1 | # | 20 | Small | ancom | 86 |
| # | interaction | 0 | 0 | 4 | 0 | 0 | # | 20 | Small | ancom | 86 |
| # | disease | 7 | 2 | 0 | 0.777777778 | 1 | # | 20 | Small | aldex2 | 86 |
| # | sex | 4 | 0 | 0 | 1 | 1 | # | 20 | Small | aldex2 | 86 |
| # | time | 5 | 5 | 0 | 0.5 | 1 | # | 20 | Small | aldex2 | 86 |
| # | interaction | 0 | 0 | 4 | 0 | 0 | # | 20 | Small | aldex2 | 86 |
| # | disease | 7 | 9 | 0 | 0.4375 | 1 | # | 20 | Baseline | paralle | 86 |
| # | sex | 4 | 5 | 0 | 0.444444444 | 1 | # | 20 | Baseline | paralle | 86 |
| # | time | 5 | 2 | 0 | 0.714285714 | 1 | # | 20 | Baseline | paralle | 86 |
| # | interaction | 4 | 3 | 0 | 0.571428571 | 1 | # | 20 | Baseline | paralle | 86 |
| # | disease | 7 | 4 | 0 | 0.636363636 | 1 | # | 20 | Baseline | ancom | 86 |
| # | sex | 4 | 0 | 0 | 1 | 1 | # | 20 | Baseline | ancom | 86 |
| # | time | 5 | 4 | 0 | 0.555555556 | 1 | # | 20 | Baseline | ancom | 86 |
| # | interaction | 0 | 0 | 4 | 0 | 0 | # | 20 | Baseline | ancom | 86 |
| # | disease | 7 | 52 | 0 | 0.118644068 | 1 | # | 20 | Baseline | aldex2 | 86 |
| # | sex | 4 | 12 | 0 | 0.25 | 1 | # | 20 | Baseline | aldex2 | 86 |
| # | time | 5 | 28 | 0 | 0.151515152 | 1 | # | 20 | Baseline | aldex2 | 86 |
| # | interaction | 0 | 0 | 4 | 0 | 0 | # | 20 | Baseline | aldex2 | 86 |
| # | disease | 7 | 0 | 0 | 1 | 1 | # | 20 | Small | paralle | 86 |
| # | sex | 4 | 0 | 0 | 1 | 1 | # | 20 | Small | paralle | 86 |
| # | time | 5 | 1 | 0 | 0.833333333 | 1 | # | 20 | Small | paralle | 86 |
| # | interaction | 4 | 0 | 0 | 1 | 1 | # | 20 | Small | paralle | 86 |
| # | disease | 7 | 3 | 0 | 0.7 | 1 | # | 20 | Small | ancom | 86 |
| # | sex | 4 | 0 | 0 | 1 | 1 | # | 20 | Small | ancom | 86 |
| # | time | 5 | 4 | 0 | 0.555555556 | 1 | # | 20 | Small | ancom | 86 |
| # | interaction | 0 | 0 | 4 | 0 | 0 | # | 20 | Small | ancom | 86 |
| # | disease | 7 | 5 | 0 | 0.583333333 | 1 | # | 20 | Small | aldex2 | 86 |
| # | sex | 4 | 0 | 0 | 1 | 1 | # | 20 | Small | aldex2 | 86 |
| # | time | 5 | 4 | 0 | 0.555555556 | 1 | # | 20 | Small | aldex2 | 86 |
| # | interaction | 0 | 0 | 4 | 0 | 0 | # | 20 | Small | aldex2 | 86 |
| # | disease | 7 | 6 | 0 | 0.538461538 | 1 | # | 20 | Baseline | paralle | 86 |
| # | sex | 4 | 7 | 0 | 0.363636364 | 1 | # | 20 | Baseline | paralle | 86 |
| # | time | 5 | 4 | 0 | 0.555555556 | 1 | # | 20 | Baseline | paralle | 86 |
| # | interaction | 4 | 3 | 0 | 0.571428571 | 1 | # | 20 | Baseline | paralle | 86 |
| # | disease | 7 | 4 | 0 | 0.636363636 | 1 | # | 20 | Baseline | ancom | 86 |
| # | sex | 4 | 0 | 0 | 1 | 1 | # | 20 | Baseline | ancom | 86 |
| # | time | 5 | 4 | 0 | 0.555555556 | 1 | # | 20 | Baseline | ancom | 86 |
| # | interaction | 0 | 0 | 4 | 0 | 0 | # | 20 | Baseline | ancom | 86 |
| # | disease | 7 | 85 | 0 | 0.097222222 | 1 | # | 20 | Baseline | aldex2 | 86 |
| # | sex | 4 | 19 | 0 | 0.173913043 | 1 | # | 20 | Baseline | aldex2 | 86 |
| # | time | 5 | 33 | 0 | 0.131578947 | 1 | # | 20 | Baseline | aldex2 | 86 |
| # | interaction | 0 | 0 | 4 | 0 | 0 | # | 20 | Baseline | aldex2 | 86 |
| # | disease | 7 | 0 | 0 | 1 | 1 | # | 20 | Small | paralle | 86 |
| # | sex | 4 | 0 | 0 | 1 | 1 | # | 20 | Small | paralle | 86 |
| # | time | 5 | 0 | 0 | 1 | 1 | # | 20 | Small | paralle | 86 |
| # | interaction | 4 | 0 | 0 | 1 | 1 | # | 20 | Small | paralle | 86 |
| # | disease | 7 | 4 | 0 | 0.636363636 | 1 | # | 20 | Small | ancom | 86 |
| # | sex | 4 | 0 | 0 | 1 | 1 | # | 20 | Small | ancom | 86 |
| # | time | 5 | 4 | 0 | 0.555555556 | 1 | # | 20 | Small | ancom | 86 |
| # | interaction | 0 | 0 | 4 | 0 | 0 | # | 20 | Small | ancom | 86 |
| # | disease | 7 | 10 | 0 | 0.411764706 | 1 | # | 20 | Small | aldex2 | 86 |
| # | sex | 4 | 0 | 0 | 1 | 1 | # | 20 | Small | aldex2 | 86 |

| | | | | | | | | | | | |
|---|---|---|---|---|---|---|---|---|---|---|---|
| # | time | 5 | 6 | 0 | 0.454545455 | 1 | # | 20 | Small | sided2 | 86 |
| # | interaction | 0 | 0 | 4 | 0 | 0 | # | 20 | Small | sided2 | 86 |
| # | disease | 4 | 9 | 0 | 0.307692308 | 1 | # | 10 | Baseline | paralle | 87 |
| # | sex | 2 | 9 | 0 | 0.181818182 | 1 | # | 10 | Baseline | paralle | 87 |
| # | time | 2 | 9 | 0 | 0.181818182 | 1 | # | 10 | Baseline | paralle | 87 |
| # | interaction | 2 | 11 | 0 | 0.153846154 | 1 | # | 10 | Baseline | paralle | 87 |
| # | disease | 4 | 2 | 0 | 0.666666667 | 1 | # | 10 | Baseline | ancom | 87 |
| # | sex | 2 | 0 | 0 | 1 | 1 | # | 10 | Baseline | ancom | 87 |
| # | time | 2 | 1 | 0 | 0.666666667 | 1 | # | 10 | Baseline | ancom | 87 |
| # | interaction | 0 | 0 | 2 | 0 | 0 | # | 10 | Baseline | ancom | 87 |
| # | disease | 4 | 8 | 0 | 0.333333333 | 1 | # | 10 | Baseline | sided2 | 87 |
| # | sex | 2 | 1 | 0 | 0.666666667 | 1 | # | 10 | Baseline | sided2 | 87 |
| # | time | 2 | 5 | 0 | 0.285714286 | 1 | # | 10 | Baseline | sided2 | 87 |
| # | interaction | 0 | 0 | 2 | 0 | 0 | # | 10 | Baseline | sided2 | 87 |
| # | disease | 4 | 0 | 0 | 1 | 1 | # | 10 | Small | paralle | 87 |
| # | sex | 2 | 1 | 0 | 0.666666667 | 1 | # | 10 | Small | paralle | 87 |
| # | time | 2 | 0 | 0 | 1 | 1 | # | 10 | Small | paralle | 87 |
| # | interaction | 2 | 0 | 0 | 1 | 1 | # | 10 | Small | paralle | 87 |
| # | disease | 4 | 1 | 0 | 0.8 | 1 | # | 10 | Small | ancom | 87 |
| # | sex | 2 | 0 | 0 | 1 | 1 | # | 10 | Small | ancom | 87 |
| # | time | 2 | 1 | 0 | 0.666666667 | 1 | # | 10 | Small | ancom | 87 |
| # | interaction | 0 | 0 | 2 | 0 | 0 | # | 10 | Small | ancom | 87 |
| # | disease | 4 | 2 | 0 | 0.666666667 | 1 | # | 10 | Small | sided2 | 87 |
| # | sex | 2 | 0 | 0 | 1 | 1 | # | 10 | Small | sided2 | 87 |
| # | time | 2 | 1 | 0 | 0.666666667 | 1 | # | 10 | Small | sided2 | 87 |
| # | interaction | 0 | 0 | 2 | 0 | 0 | # | 10 | Small | sided2 | 87 |
| # | disease | 4 | 7 | 0 | 0.363636364 | 1 | # | 10 | Baseline | paralle | 87 |
| # | sex | 2 | 9 | 0 | 0.181818182 | 1 | # | 10 | Baseline | paralle | 87 |
| # | time | 2 | 2 | 0 | 0.5 | 1 | # | 10 | Baseline | paralle | 87 |
| # | interaction | 2 | 6 | 0 | 0.25 | 1 | # | 10 | Baseline | paralle | 87 |
| # | disease | 4 | 1 | 0 | 0.8 | 1 | # | 10 | Baseline | ancom | 87 |
| # | sex | 2 | 0 | 0 | 1 | 1 | # | 10 | Baseline | ancom | 87 |
| # | time | 2 | 1 | 0 | 0.666666667 | 1 | # | 10 | Baseline | ancom | 87 |
| # | interaction | 0 | 0 | 2 | 0 | 0 | # | 10 | Baseline | ancom | 87 |
| # | disease | 4 | 13 | 0 | 0.235294118 | 1 | # | 10 | Baseline | sided2 | 87 |
| # | sex | 2 | 4 | 0 | 0.333333333 | 1 | # | 10 | Baseline | sided2 | 87 |
| # | time | 2 | 14 | 0 | 0.125 | 1 | # | 10 | Baseline | sided2 | 87 |
| # | interaction | 0 | 0 | 2 | 0 | 0 | # | 10 | Baseline | sided2 | 87 |
| # | disease | 4 | 1 | 0 | 0.8 | 1 | # | 10 | Small | paralle | 87 |
| # | sex | 2 | 0 | 0 | 1 | 1 | # | 10 | Small | paralle | 87 |
| # | time | 2 | 0 | 0 | 1 | 1 | # | 10 | Small | paralle | 87 |
| # | interaction | 2 | 0 | 0 | 1 | 1 | # | 10 | Small | paralle | 87 |
| # | disease | 4 | 1 | 0 | 0.8 | 1 | # | 10 | Small | ancom | 87 |
| # | sex | 2 | 0 | 0 | 1 | 1 | # | 10 | Small | ancom | 87 |
| # | time | 2 | 2 | 0 | 0.5 | 1 | # | 10 | Small | ancom | 87 |
| # | interaction | 0 | 0 | 2 | 0 | 0 | # | 10 | Small | ancom | 87 |
| # | disease | 4 | 3 | 0 | 0.571428571 | 1 | # | 10 | Small | sided2 | 87 |
| # | sex | 2 | 1 | 0 | 0.666666667 | 1 | # | 10 | Small | sided2 | 87 |
| # | time | 2 | 3 | 0 | 0.4 | 1 | # | 10 | Small | sided2 | 87 |
| # | interaction | 0 | 0 | 2 | 0 | 0 | # | 10 | Small | sided2 | 87 |
| # | disease | 4 | 5 | 0 | 0.444444444 | 1 | # | 10 | Baseline | paralle | 87 |
| # | sex | 2 | 5 | 0 | 0.285714286 | 1 | # | 10 | Baseline | paralle | 87 |
| # | time | 2 | 2 | 0 | 0.5 | 1 | # | 10 | Baseline | paralle | 87 |
| # | interaction | 2 | 5 | 0 | 0.285714286 | 1 | # | 10 | Baseline | paralle | 87 |
| # | disease | 4 | 1 | 0 | 0.8 | 1 | # | 10 | Baseline | ancom | 87 |
| # | sex | 2 | 0 | 0 | 1 | 1 | # | 10 | Baseline | ancom | 87 |
| # | time | 2 | 2 | 0 | 0.5 | 1 | # | 10 | Baseline | ancom | 87 |
| # | interaction | 0 | 0 | 2 | 0 | 0 | # | 10 | Baseline | ancom | 87 |
| # | disease | 4 | 19 | 0 | 0.173913043 | 1 | # | 10 | Baseline | sided2 | 87 |
| # | sex | 2 | 10 | 0 | 0.166666667 | 1 | # | 10 | Baseline | sided2 | 87 |
| # | time | 2 | 14 | 0 | 0.125 | 1 | # | 10 | Baseline | sided2 | 87 |
| # | interaction | 0 | 0 | 2 | 0 | 0 | # | 10 | Baseline | sided2 | 87 |
| # | disease | 4 | 0 | 0 | 1 | 1 | # | 10 | Small | paralle | 87 |
| # | sex | 2 | 0 | 0 | 1 | 1 | # | 10 | Small | paralle | 87 |
| # | time | 2 | 0 | 0 | 1 | 1 | # | 10 | Small | paralle | 87 |
| # | interaction | 2 | 0 | 0 | 1 | 1 | # | 10 | Small | paralle | 87 |
| # | disease | 4 | 1 | 0 | 0.8 | 1 | # | 10 | Small | ancom | 87 |
| # | sex | 2 | 0 | 0 | 1 | 1 | # | 10 | Small | ancom | 87 |
| # | time | 2 | 2 | 0 | 0.5 | 1 | # | 10 | Small | ancom | 87 |
| # | interaction | 0 | 0 | 2 | 0 | 0 | # | 10 | Small | ancom | 87 |
| # | disease | 4 | 2 | 0 | 0.666666667 | 1 | # | 10 | Small | sided2 | 87 |
| # | sex | 2 | 0 | 0 | 1 | 1 | # | 10 | Small | sided2 | 87 |
| # | time | 2 | 2 | 0 | 0.5 | 1 | # | 10 | Small | sided2 | 87 |
| # | interaction | 0 | 0 | 2 | 0 | 0 | # | 10 | Small | sided2 | 87 |
| # | disease | 4 | 4 | 0 | 0.5 | 1 | # | 10 | Baseline | paralle | 87 |
| # | sex | 2 | 2 | 0 | 0.5 | 1 | # | 10 | Baseline | paralle | 87 |
| # | time | 2 | 2 | 0 | 0.5 | 1 | # | 10 | Baseline | paralle | 87 |
| # | interaction | 2 | 3 | 0 | 0.4 | 1 | # | 10 | Baseline | paralle | 87 |
| # | disease | 4 | 2 | 0 | 0.666666667 | 1 | # | 10 | Baseline | ancom | 87 |
| # | sex | 2 | 0 | 0 | 1 | 1 | # | 10 | Baseline | ancom | 87 |
| # | time | 2 | 2 | 0 | 0.5 | 1 | # | 10 | Baseline | ancom | 87 |
| # | interaction | 0 | 0 | 2 | 0 | 0 | # | 10 | Baseline | ancom | 87 |
| # | disease | 4 | 21 | 0 | 0.16 | 1 | # | 10 | Baseline | sided2 | 87 |
| # | sex | 2 | 14 | 0 | 0.125 | 1 | # | 10 | Baseline | sided2 | 87 |
| # | time | 2 | 18 | 0 | 0.1 | 1 | # | 10 | Baseline | sided2 | 87 |
| # | interaction | 0 | 0 | 2 | 0 | 0 | # | 10 | Baseline | sided2 | 87 |
| # | disease | 4 | 0 | 0 | 1 | 1 | # | 10 | Small | paralle | 87 |
| # | sex | 2 | 0 | 0 | 1 | 1 | # | 10 | Small | paralle | 87 |
| # | time | 2 | 0 | 0 | 1 | 1 | # | 10 | Small | paralle | 87 |
| # | interaction | 2 | 0 | 0 | 1 | 1 | # | 10 | Small | paralle | 87 |
| # | disease | 4 | 2 | 0 | 0.666666667 | 1 | # | 10 | Small | ancom | 87 |
| # | sex | 2 | 0 | 0 | 1 | 1 | # | 10 | Small | ancom | 87 |
| # | time | 2 | 2 | 0 | 0.5 | 1 | # | 10 | Small | ancom | 87 |
| # | interaction | 0 | 0 | 2 | 0 | 0 | # | 10 | Small | ancom | 87 |
| # | disease | 4 | 2 | 0 | 0.666666667 | 1 | # | 10 | Small | sided2 | 87 |
| # | sex | 2 | 0 | 0 | 1 | 1 | # | 10 | Small | sided2 | 87 |
| # | time | 2 | 2 | 0 | 0.5 | 1 | # | 10 | Small | sided2 | 87 |
| # | interaction | 0 | 0 | 2 | 0 | 0 | # | 10 | Small | sided2 | 87 |
| # | disease | 7 | 8 | 0 | 0.466666667 | 1 | # | 20 | Baseline | paralle | 87 |
| # | sex | 4 | 7 | 0 | 0.363636364 | 1 | # | 20 | Baseline | paralle | 87 |
| # | time | 5 | 11 | 0 | 0.3125 | 1 | # | 20 | Baseline | paralle | 87 |
| # | interaction | 4 | 6 | 0 | 0.4 | 1 | # | 20 | Baseline | paralle | 87 |
| # | disease | 7 | 0 | 0 | 1 | 1 | # | 20 | Baseline | ancom | 87 |
| # | sex | 4 | 0 | 0 | 1 | 1 | # | 20 | Baseline | ancom | 87 |
| # | time | 5 | 2 | 0 | 0.714285714 | 1 | # | 20 | Baseline | ancom | 87 |
| # | interaction | 0 | 0 | 4 | 0 | 0 | # | 20 | Baseline | ancom | 87 |
| # | disease | 7 | 25 | 0 | 0.21875 | 1 | # | 20 | Baseline | sided2 | 87 |
| # | sex | 4 | 6 | 0 | 0.4 | 1 | # | 20 | Baseline | sided2 | 87 |
| # | time | 5 | 12 | 0 | 0.294117647 | 1 | # | 20 | Baseline | sided2 | 87 |
| # | interaction | 0 | 0 | 4 | 0 | 0 | # | 20 | Baseline | sided2 | 87 |
| # | disease | 7 | 1 | 0 | 0.875 | 1 | # | 20 | Small | paralle | 87 |
| # | sex | 4 | 0 | 0 | 1 | 1 | # | 20 | Small | paralle | 87 |
| # | time | 5 | 1 | 0 | 0.833333333 | 1 | # | 20 | Small | paralle | 87 |
| # | interaction | 2 | 2 | 2 | 0.5 | 0.5 | # | 20 | Small | paralle | 87 |
| # | disease | 7 | 0 | 0 | 1 | 1 | # | 20 | Small | ancom | 87 |
| # | sex | 4 | 0 | 0 | 1 | 1 | # | 20 | Small | ancom | 87 |
| # | time | 5 | 0 | 0 | 1 | 1 | # | 20 | Small | ancom | 87 |
| # | interaction | 0 | 0 | 4 | 0 | 0 | # | 20 | Small | ancom | 87 |
| # | disease | 7 | 0 | 0 | 1 | 1 | # | 20 | Small | sided2 | 87 |
| # | sex | 4 | 0 | 0 | 1 | 1 | # | 20 | Small | sided2 | 87 |
| # | time | 5 | 3 | 0 | 0.625 | 1 | # | 20 | Small | sided2 | 87 |
| # | interaction | 0 | 0 | 4 | 0 | 0 | # | 20 | Small | sided2 | 87 |
| # | disease | 7 | 6 | 0 | 0.538461538 | 1 | # | 20 | Baseline | paralle | 87 |
| # | sex | 4 | 3 | 0 | 0.571428571 | 1 | # | 20 | Baseline | paralle | 87 |
| # | time | 5 | 10 | 0 | 0.333333333 | 1 | # | 20 | Baseline | paralle | 87 |
| # | interaction | 4 | 8 | 0 | 0.333333333 | 1 | # | 20 | Baseline | paralle | 87 |
| # | disease | 7 | 2 | 0 | 0.777777778 | 1 | # | 20 | Baseline | ancom | 87 |
| # | sex | 4 | 0 | 0 | 1 | 1 | # | 20 | Baseline | ancom | 87 |
| # | time | 5 | 3 | 0 | 0.625 | 1 | # | 20 | Baseline | ancom | 87 |
| # | interaction | 0 | 0 | 4 | 0 | 0 | # | 20 | Baseline | ancom | 87 |
| # | disease | 7 | 38 | 0 | 0.155555556 | 1 | # | 20 | Baseline | sided2 | 87 |
| # | sex | 4 | 5 | 0 | 0.444444444 | 1 | # | 20 | Baseline | sided2 | 87 |
| # | time | 5 | 21 | 0 | 0.192307692 | 1 | # | 20 | Baseline | sided2 | 87 |
| # | interaction | 0 | 0 | 4 | 0 | 0 | # | 20 | Baseline | sided2 | 87 |
| # | disease | 7 | 0 | 0 | 1 | 1 | # | 20 | Small | paralle | 87 |
| # | sex | 4 | 0 | 0 | 1 | 1 | # | 20 | Small | paralle | 87 |
| # | time | 5 | 0 | 0 | 1 | 1 | # | 20 | Small | paralle | 87 |
| # | interaction | 4 | 1 | 0 | 0.8 | 1 | # | 20 | Small | paralle | 87 |
| # | disease | 7 | 2 | 0 | 0.777777778 | 1 | # | 20 | Small | ancom | 87 |
| # | sex | 4 | 0 | 0 | 1 | 1 | # | 20 | Small | ancom | 87 |
| # | time | 5 | 3 | 0 | 0.625 | 1 | # | 20 | Small | ancom | 87 |
| # | interaction | 0 | 0 | 4 | 0 | 0 | # | 20 | Small | ancom | 87 |
| # | disease | 7 | 3 | 0 | 0.7 | 1 | # | 20 | Small | sided2 | 87 |
| # | sex | 4 | 0 | 0 | 1 | 1 | # | 20 | Small | sided2 | 87 |
| # | time | 5 | 3 | 0 | 0.625 | 1 | # | 20 | Small | sided2 | 87 |
| # | interaction | 0 | 0 | 4 | 0 | 0 | # | 20 | Small | sided2 | 87 |
| # | disease | 7 | 9 | 0 | 0.4375 | 1 | # | 20 | Baseline | paralle | 87 |
| # | sex | 4 | 5 | 0 | 0.444444444 | 1 | # | 20 | Baseline | paralle | 87 |
| # | time | 5 | 9 | 0 | 0.357142857 | 1 | # | 20 | Baseline | paralle | 87 |
| # | interaction | 4 | 3 | 0 | 0.571428571 | 1 | # | 20 | Baseline | paralle | 87 |
| # | disease | 7 | 3 | 0 | 0.7 | 1 | # | 20 | Baseline | ancom | 87 |
| # | sex | 4 | 0 | 0 | 1 | 1 | # | 20 | Baseline | ancom | 87 |
| # | time | 5 | 4 | 0 | 0.555555556 | 1 | # | 20 | Baseline | ancom | 87 |
| # | interaction | 0 | 0 | 4 | 0 | 0 | # | 20 | Baseline | ancom | 87 |
| # | disease | 7 | 46 | 0 | 0.132075472 | 1 | # | 20 | Baseline | sided2 | 87 |
| # | sex | 4 | 10 | 0 | 0.285714286 | 1 | # | 20 | Baseline | sided2 | 87 |
| # | time | 5 | 33 | 0 | 0.131578947 | 1 | # | 20 | Baseline | sided2 | 87 |
| # | interaction | 0 | 0 | 4 | 0 | 0 | # | 20 | Baseline | sided2 | 87 |
| # | disease | 7 | 0 | 0 | 1 | 1 | # | 20 | Small | paralle | 87 |
| # | sex | 4 | 0 | 0 | 1 | 1 | # | 20 | Small | paralle | 87 |
| # | time | 5 | 1 | 0 | 0.833333333 | 1 | # | 20 | Small | paralle | 87 |
| # | interaction | 4 | 0 | 0 | 1 | 1 | # | 20 | Small | paralle | 87 |
| # | disease | 7 | 3 | 0 | 0.7 | 1 | # | 20 | Small | ancom | 87 |
| # | sex | 4 | 0 | 0 | 1 | 1 | # | 20 | Small | ancom | 87 |
| # | time | 5 | 4 | 0 | 0.555555556 | 1 | # | 20 | Small | ancom | 87 |
| # | interaction | 0 | 0 | 4 | 0 | 0 | # | 20 | Small | ancom | 87 |
| # | disease | 7 | 6 | 0 | 0.538461538 | 1 | # | 20 | Small | sided2 | 87 |
| # | sex | 4 | 0 | 0 | 1 | 1 | # | 20 | Small | sided2 | 87 |
| # | time | 5 | 2 | 0 | 0.714285714 | 1 | # | 20 | Small | sided2 | 87 |
| # | interaction | 0 | 0 | 4 | 0 | 0 | # | 20 | Small | sided2 | 87 |
| # | disease | 7 | 12 | 0 | 0.368421053 | 1 | # | 20 | Baseline | paralle | 87 |
| # | sex | 4 | 2 | 0 | 0.666666667 | 1 | # | 20 | Baseline | paralle | 87 |
| # | time | 5 | 11 | 0 | 0.3125 | 1 | # | 20 | Baseline | paralle | 87 |
| # | interaction | 4 | 3 | 0 | 0.571428571 | 1 | # | 20 | Baseline | paralle | 87 |
| # | disease | 7 | 4 | 0 | 0.636363636 | 1 | # | 20 | Baseline | ancom | 87 |
| # | sex | 4 | 0 | 0 | 1 | 1 | # | 20 | Baseline | ancom | 87 |
| # | time | 5 | 4 | 0 | 0.555555556 | 1 | # | 20 | Baseline | ancom | 87 |
| # | interaction | 0 | 0 | 4 | 0 | 0 | # | 20 | Baseline | ancom | 87 |
| # | disease | 7 | 58 | 0 | 0.107692308 | 1 | # | 20 | Baseline | sided2 | 87 |
| # | sex | 4 | 15 | 0 | 0.210526316 | 1 | # | 20 | Baseline | sided2 | 87 |
| # | time | 5 | 38 | 0 | 0.11627907 | 1 | # | 20 | Baseline | sided2 | 87 |
| # | interaction | 0 | 0 | 4 | 0 | 0 | # | 20 | Baseline | sided2 | 87 |
| # | disease | 7 | 0 | 0 | 1 | 1 | # | 20 | Small | paralle | 87 |
| # | sex | 4 | 0 | 0 | 1 | 1 | # | 20 | Small | paralle | 87 |
| # | time | 5 | 0 | 0 | 1 | 1 | # | 20 | Small | paralle | 87 |
| # | interaction | 4 | 0 | 0 | 1 | 1 | # | 20 | Small | paralle | 87 |
| # | disease | 7 | 4 | 0 | 0.636363636 | 1 | # | 20 | Small | ancom | 87 |
| # | sex | 4 | 0 | 0 | 1 | 1 | # | 20 | Small | ancom | 87 |
| # | time | 5 | 4 | 0 | 0.555555556 | 1 | # | 20 | Small | ancom | 87 |
| # | interaction | 0 | 0 | 4 | 0 | 0 | # | 20 | Small | ancom | 87 |
| # | disease | 7 | 9 | 0 | 0.4375 | 1 | # | 20 | Small | sided2 | 87 |
| # | sex | 4 | 1 | 0 | 0.8 | 1 | # | 20 | Small | sided2 | 87 |
| # | time | 5 | 7 | 0 | 0.416666667 | 1 | # | 20 | Small | sided2 | 87 |
| # | interaction | 0 | 0 | 4 | 0 | 0 | # | 20 | Small | sided2 | 87 |
| # | disease | 4 | 13 | 0 | 0.235294118 | 1 | # | 10 | Baseline | paralle | 88 |
| # | sex | 2 | 7 | 0 | 0.222222222 | 1 | # | 10 | Baseline | paralle | 88 |
| # | time | 2 | 6 | 0 | 0.25 | 1 | # | 10 | Baseline | paralle | 88 |
| # | interaction | 2 | 13 | 0 | 0.133333333 | 1 | # | 10 | Baseline | paralle | 88 |
| # | disease | 4 | 0 | 0 | 1 | 1 | # | 10 | Baseline | ancom | 88 |
| # | sex | 2 | 0 | 0 | 1 | 1 | # | 10 | Baseline | ancom | 88 |
| # | time | 2 | 1 | 0 | 0.666666667 | 1 | # | 10 | Baseline | ancom | 88 |
| # | interaction | 0 | 0 | 2 | 0 | 0 | # | 10 | Baseline | ancom | 88 |
| # | disease | 4 | 10 | 0 | 0.285714286 | 1 | # | 10 | Baseline | sided2 | 88 |
| # | sex | 2 | 4 | 0 | 0.333333333 | 1 | # | 10 | Baseline | sided2 | 88 |
| # | time | 2 | 8 | 0 | 0.2 | 1 | # | 10 | Baseline | sided2 | 88 |
| # | interaction | 0 | 0 | 2 | 0 | 0 | # | 10 | Baseline | sided2 | 88 |
| # | disease | 4 | 1 | 0 | 0.8 | 1 | # | 10 | Small | paralle | 88 |
| # | sex | 2 | 0 | 0 | 1 | 1 | # | 10 | Small | paralle | 88 |
| # | time | 2 | 0 | 0 | 1 | 1 | # | 10 | Small | paralle | 88 |
| # | interaction | 0 | 0 | 2 | 0 | 0 | # | 10 | Small | paralle | 88 |
| # | disease | 4 | 0 | 0 | 1 | 1 | # | 10 | Small | ancom | 88 |
| # | sex | 2 | 0 | 0 | 1 | 1 | # | 10 | Small | ancom | 88 |
| # | time | 2 | 1 | 0 | 0.666666667 | 1 | # | 10 | Small | ancom | 88 |
| # | interaction | 0 | 0 | 2 | 0 | 0 | # | 10 | Small | ancom | 88 |
| # | disease | 4 | 0 | 0 | 1 | 1 | # | 10 | Small | sided2 | 88 |
| # | sex | 2 | 0 | 0 | 1 | 1 | # | 10 | Small | sided2 | 88 |
| # | time | 2 | 1 | 0 | 0.666666667 | 1 | # | 10 | Small | sided2 | 88 |
| # | interaction | 0 | 0 | 2 | 0 | 0 | # | 10 | Small | sided2 | 88 |
| # | disease | 4 | 5 | 0 | 0.444444444 | 1 | # | 10 | Baseline | paralle | 88 |
| # | sex | 2 | 8 | 0 | 0.2 | 1 | # | 10 | Baseline | paralle | 88 |
| # | time | 2 | 4 | 0 | 0.333333333 | 1 | # | 10 | Baseline | paralle | 88 |
| # | interaction | 2 | 4 | 0 | 0.333333333 | 1 | # | 10 | Baseline | paralle | 88 |
| # | disease | 4 | 2 | 0 | 0.666666667 | 1 | # | 10 | Baseline | ancom | 88 |
| # | sex | 2 | 0 | 0 | 1 | 1 | # | 10 | Baseline | ancom | 88 |
| # | time | 2 | 1 | 0 | 0.666666667 | 1 | # | 10 | Baseline | ancom | 88 |
| # | interaction | 0 | 0 | 2 | 0 | 0 | # | 10 | Baseline | ancom | 88 |
| # | disease | 4 | 18 | 0 | 0.181818182 | 1 | # | 10 | Baseline | sided2 | 88 |
| # | sex | 2 | 8 | 0 | 0.2 | 1 | # | 10 | Baseline | sided2 | 88 |
| # | time | 2 | 10 | 0 | 0.166666667 | 1 | # | 10 | Baseline | sided2 | 88 |
| # | interaction | 0 | 0 | 2 | 0 | 0 | # | 10 | Baseline | sided2 | 88 |
| # | disease | 4 | 0 | 0 | 1 | 1 | # | 10 | Small | paralle | 88 |
| # | sex | 2 | 0 | 0 | 1 | 1 | # | 10 | Small | paralle | 88 |
| # | time | 2 | 0 | 0 | 1 | 1 | # | 10 | Small | paralle | 88 |
| # | interaction | 1 | 0 | 1 | 1 | 0.5 | # | 10 | Small | paralle | 88 |
| # | disease | 4 | 1 | 0 | 0.8 | 1 | # | 10 | Small | ancom | 88 |
| # | sex | 2 | 0 | 0 | 1 | 1 | # | 10 | Small | ancom | 88 |
| # | time | 2 | 0 | 0 | 1 | 1 | # | 10 | Small | ancom | 88 |
| # | interaction | 0 | 0 | 2 | 0 | 0 | # | 10 | Small | ancom | 88 |
| # | disease | 4 | 2 | 0 | 0.666666667 | 1 | # | 10 | Small | sided2 | 88 |
| # | sex | 2 | 0 | 0 | 1 | 1 | # | 10 | Small | sided2 | 88 |
| # | time | 2 | 1 | 0 | 0.666666667 | 1 | # | 10 | Small | sided2 | 88 |
| # | interaction | 0 | 0 | 2 | 0 | 0 | # | 10 | Small | sided2 | 88 |
| # | disease | 4 | 8 | 0 | 0.333333333 | 1 | # | 10 | Baseline | paralle | 88 |
| # | sex | 2 | 8 | 0 | 0.2 | 1 | # | 10 | Baseline | paralle | 88 |
| # | time | 2 | 6 | 0 | 0.25 | 1 | # | 10 | Baseline | paralle | 88 |
| # | interaction | 2 | 5 | 0 | 0.285714286 | 1 | # | 10 | Baseline | paralle | 88 |
| # | disease | 4 | 2 | 0 | 0.666666667 | 1 | # | 10 | Baseline | ancom | 88 |
| # | sex | 2 | 0 | 0 | 1 | 1 | # | 10 | Baseline | ancom | 88 |
| # | time | 2 | 2 | 0 | 0.5 | 1 | # | 10 | Baseline | ancom | 88 |
| # | interaction | 0 | 0 | 2 | 0 | 0 | # | 10 | Baseline | ancom | 88 |
| # | disease | 4 | 25 | 0 | 0.137931034 | 1 | # | 10 | Baseline | sided2 | 88 |
| # | sex | 2 | 3 | 0 | 0.4 | 1 | # | 10 | Baseline | sided2 | 88 |
| # | time | 2 | 10 | 0 | 0.166666667 | 1 | # | 10 | Baseline | sided2 | 88 |
| # | interaction | 0 | 0 | 2 | 0 | 0 | # | 10 | Baseline | sided2 | 88 |
| # | disease | 4 | 0 | 0 | 1 | 1 | # | 10 | Small | paralle | 88 |
| # | sex | 2 | 0 | 0 | 1 | 1 | # | 10 | Small | paralle | 88 |
| # | time | 2 | 0 | 0 | 1 | 1 | # | 10 | Small | paralle | 88 |
| # | interaction | 1 | 0 | 1 | 1 | 0.5 | # | 10 | Small | paralle | 88 |
| # | disease | 4 | 2 | 0 | 0.666666667 | 1 | # | 10 | Small | ancom | 88 |
| # | sex | 2 | 0 | 0 | 1 | 1 | # | 10 | Small | ancom | 88 |
| # | time | 2 | 2 | 0 | 0.5 | 1 | # | 10 | Small | ancom | 88 |
| # | interaction | 0 | 0 | 2 | 0 | 0 | # | 10 | Small | ancom | 88 |
| # | disease | 4 | 2 | 0 | 0.666666667 | 1 | # | 10 | Small | sided2 | 88 |
| # | sex | 2 | 0 | 0 | 1 | 1 | # | 10 | Small | sided2 | 88 |
| # | time | 2 | 2 | 0 | 0.5 | 1 | # | 10 | Small | sided2 | 88 |
| # | interaction | 0 | 0 | 2 | 0 | 0 | # | 10 | Small | sided2 | 88 |
| # | disease | 4 | 8 | 0 | 0.333333333 | 1 | # | 10 | Baseline | paralle | 88 |
| # | sex | 2 | 6 | 0 | 0.25 | 1 | # | 10 | Baseline | paralle | 88 |
| # | time | 2 | 5 | 0 | 0.285714286 | 1 | # | 10 | Baseline | paralle | 88 |
| # | interaction | 2 | 3 | 0 | 0.4 | 1 | # | 10 | Baseline | paralle | 88 |
| # | disease | 4 | 2 | 0 | 0.666666667 | 1 | # | 10 | Baseline | ancom | 88 |
| # | sex | 2 | 0 | 0 | 1 | 1 | # | 10 | Baseline | ancom | 88 |
| # | time | 2 | 2 | 0 | 0.5 | 1 | # | 10 | Baseline | ancom | 88 |
| # | interaction | 0 | 0 | 2 | 0 | 0 | # | 10 | Baseline | ancom | 88 |
| # | disease | 4 | 32 | 0 | 0.111111111 | 1 | # | 10 | Baseline | sided2 | 88 |
| # | sex | 2 | 4 | 0 | 0.333333333 | 1 | # | 10 | Baseline | sided2 | 88 |
| # | time | 2 | 8 | 0 | 0.2 | 1 | # | 10 | Baseline | sided2 | 88 |
| # | interaction | 0 | 0 | 2 | 0 | 0 | # | 10 | Baseline | sided2 | 88 |
| # | disease | 4 | 0 | 0 | 1 | 1 | # | 10 | Small | paralle | 88 |
| # | sex | 2 | 0 | 0 | 1 | 1 | # | 10 | Small | paralle | 88 |
| # | time | 2 | 0 | 0 | 1 | 1 | # | 10 | Small | paralle | 88 |
| # | interaction | 2 | 0 | 0 | 1 | 1 | # | 10 | Small | paralle | 88 |
| # | disease | 4 | 2 | 0 | 0.666666667 | 1 | # | 10 | Small | ancom | 88 |
| # | sex | 2 | 0 | 0 | 1 | 1 | # | 10 | Small | ancom | 88 |
| # | time | 2 | 2 | 0 | 0.5 | 1 | # | 10 | Small | ancom | 88 |
| # | interaction | 0 | 0 | 2 | 0 | 0 | # | 10 | Small | ancom | 88 |
| # | disease | 4 | 2 | 0 | 0.666666667 | 1 | # | 10 | Small | sided2 | 88 |
| # | sex | 2 | 0 | 0 | 1 | 1 | # | 10 | Small | sided2 | 88 |
| # | time | 2 | 2 | 0 | 0.5 | 1 | # | 10 | Small | sided2 | 88 |
| # | interaction | 0 | 0 | 2 | 0 | 0 | # | 10 | Small | sided2 | 88 |
| # | disease | 7 | 12 | 0 | 0.368421053 | 1 | # | 20 | Baseline | paralle | 88 |
| # | sex | 4 | 9 | 0 | 0.307692308 | 1 | # | 20 | Baseline | paralle | 88 |
| # | time | 5 | 9 | 0 | 0.357142857 | 1 | # | 20 | Baseline | paralle | 88 |
| # | interaction | 3 | 10 | 1 | 0.230769231 | 0.75 | # | 20 | Baseline | paralle | 88 |
| # | disease | 7 | 1 | 0 | 0.875 | 1 | # | 20 | Baseline | ancom | 88 |
| # | sex | 4 | 0 | 0 | 1 | 1 | # | 20 | Baseline | ancom | 88 |
| # | time | 5 | 0 | 0 | 1 | 1 | # | 20 | Baseline | ancom | 88 |
| # | interaction | 0 | 0 | 4 | 0 | 0 | # | 20 | Baseline | ancom | 88 |
| # | disease | 7 | 25 | 0 | 0.21875 | 1 | # | 20 | Baseline | sided2 | 88 |
| # | sex | 4 | 12 | 0 | 0.25 | 1 | # | 20 | Baseline | sided2 | 88 |
| # | time | 5 | 12 | 0 | 0.294117647 | 1 | # | 20 | Baseline | sided2 | 88 |
| # | interaction | 0 | 0 | 4 | 0 | 0 | # | 20 | Baseline | sided2 | 88 |
| # | disease | 7 | 0 | 0 | 1 | 1 | # | 20 | Small | paralle | 88 |
| # | sex | 4 | 2 | 0 | 0.666666667 | 1 | # | 20 | Small | paralle | 88 |
| # | time | 5 | 0 | 0 | 1 | 1 | # | 20 | Small | paralle | 88 |
| # | interaction | 2 | 0 | 2 | 1 | 0.5 | # | 20 | Small | paralle | 88 |
| # | disease | 7 | 0 | 0 | 1 | 1 | # | 20 | Small | ancom | 88 |
| # | sex | 4 | 0 | 0 | 1 | 1 | # | 20 | Small | ancom | 88 |
| # | time | 5 | 0 | 0 | 1 | 1 | # | 20 | Small | ancom | 88 |
| # | interaction | 0 | 0 | 4 | 0 | 0 | # | 20 | Small | ancom | 88 |
| # | disease | 7 | 0 | 0 | 1 | 1 | # | 20 | Small | sided2 | 88 |
| # | sex | 4 | 0 | 0 | 1 | 1 | # | 20 | Small | sided2 | 88 |
| # | time | 5 | 3 | 0 | 0.625 | 1 | # | 20 | Small | sided2 | 88 |
| # | interaction | 0 | 0 | 4 | 0 | 0 | # | 20 | Small | sided2 | 88 |
| # | disease | 7 | 14 | 0 | 0.333333333 | 1 | # | 20 | Baseline | paralle | 88 |
| # | sex | 4 | 9 | 0 | 0.307692308 | 1 | # | 20 | Baseline | paralle | 88 |
| # | time | 5 | 6 | 0 | 0.454545455 | 1 | # | 20 | Baseline | paralle | 88 |
| # | interaction | 4 | 6 | 0 | 0.4 | 1 | # | 20 | Baseline | paralle | 88 |
| # | disease | 7 | 4 | 0 | 0.636363636 | 1 | # | 20 | Baseline | ancom | 88 |
| # | sex | 4 | 0 | 0 | 1 | 1 | # | 20 | Baseline | ancom | 88 |
| # | time | 5 | 3 | 0 | 0.625 | 1 | # | 20 | Baseline | ancom | 88 |
| # | interaction | 0 | 0 | 4 | 0 | 0 | # | 20 | Baseline | ancom | 88 |
| # | disease | 7 | 39 | 0 | 0.152173913 | 1 | # | 20 | Baseline | sided2 | 88 |
| # | sex | 4 | 11 | 0 | 0.266666667 | 1 | # | 20 | Baseline | sided2 | 88 |
| # | time | 5 | 25 | 0 | 0.166666667 | 1 | # | 20 | Baseline | sided2 | 88 |
| # | interaction | 0 | 0 | 4 | 0 | 0 | # | 20 | Baseline | sided2 | 88 |
| # | disease | 7 | 0 | 0 | 1 | 1 | # | 20 | Small | paralle | 88 |
| # | sex | 4 | 1 | 0 | 0.8 | 1 | # | 20 | Small | paralle | 88 |
| # | time | 5 | 0 | 0 | 1 | 1 | # | 20 | Small | paralle | 88 |
| # | interaction | 3 | 0 | 1 | 1 | 0.75 | # | 20 | Small | paralle | 88 |
| # | disease | 7 | 2 | 0 | 0.777777778 | 1 | # | 20 | Small | ancom | 88 |
| # | sex | 4 | 0 | 0 | 1 | 1 | # | 20 | Small | ancom | 88 |
| # | time | 5 | 3 | 0 | 0.625 | 1 | # | 20 | Small | ancom | 88 |
| # | interaction | 0 | 0 | 4 | 0 | 0 | # | 20 | Small | ancom | 88 |
| # | disease | 7 | 4 | 0 | 0.636363636 | 1 | # | 20 | Small | sided2 | 88 |
| # | sex | 4 | 0 | 0 | 1 | 1 | # | 20 | Small | sided2 | 88 |
| # | time | 5 | 4 | 0 | 0.555555556 | 1 | # | 20 | Small | sided2 | 88 |

| | | | | | | | | | | | |
|---|---|---|---|---|---|---|---|---|---|---|---|
| # | interaction | 0 | 0 | 4 | 0 | 0 | # | 20 | Small | aldex2 | 88 |
| # | disease | 7 | 12 | 0 | 0.368421053 | 1 | # | 20 | Baseline | paralle | 88 |
| # | sex | 4 | 10 | 0 | 0.285714286 | 1 | # | 20 | Baseline | paralle | 88 |
| # | time | 5 | 3 | 0 | 0.625 | 1 | # | 20 | Baseline | paralle | 88 |
| # | interaction | 4 | 3 | 0 | 0.571428571 | 1 | # | 20 | Baseline | paralle | 88 |
| # | disease | 7 | 4 | 0 | 0.636363636 | 1 | # | 20 | Baseline | ancom | 88 |
| # | sex | 4 | 0 | 0 | 1 | 1 | # | 20 | Baseline | ancom | 88 |
| # | time | 5 | 3 | 0 | 0.625 | 1 | # | 20 | Baseline | ancom | 88 |
| # | interaction | 0 | 0 | 4 | 0 | 0 | # | 20 | Baseline | ancom | 88 |
| # | disease | 7 | 49 | 0 | 0.125 | 1 | # | 20 | Baseline | aldex2 | 88 |
| # | sex | 4 | 21 | 0 | 0.16 | 1 | # | 20 | Baseline | aldex2 | 88 |
| # | time | 5 | 31 | 0 | 0.138888889 | 1 | # | 20 | Baseline | aldex2 | 88 |
| # | interaction | 0 | 0 | 4 | 0 | 0 | # | 20 | Baseline | aldex2 | 88 |
| # | disease | 7 | 1 | 0 | 0.875 | 1 | # | 20 | Small | paralle | 88 |
| # | sex | 4 | 0 | 0 | 1 | 1 | # | 20 | Small | paralle | 88 |
| # | time | 5 | 0 | 0 | 1 | 1 | # | 20 | Small | paralle | 88 |
| # | interaction | 4 | 0 | 0 | 1 | 1 | # | 20 | Small | paralle | 88 |
| # | disease | 7 | 4 | 0 | 0.636363636 | 1 | # | 20 | Small | ancom | 88 |
| # | sex | 4 | 0 | 0 | 1 | 1 | # | 20 | Small | ancom | 88 |
| # | time | 5 | 3 | 0 | 0.625 | 1 | # | 20 | Small | ancom | 88 |
| # | interaction | 0 | 0 | 4 | 0 | 0 | # | 20 | Small | ancom | 88 |
| # | disease | 7 | 7 | 0 | 0.5 | 1 | # | 20 | Small | aldex2 | 88 |
| # | sex | 4 | 0 | 0 | 1 | 1 | # | 20 | Small | aldex2 | 88 |
| # | time | 5 | 5 | 0 | 0.5 | 1 | # | 20 | Small | aldex2 | 88 |
| # | interaction | 0 | 0 | 4 | 0 | 0 | # | 20 | Small | aldex2 | 88 |
| # | disease | 7 | 11 | 0 | 0.388888889 | 1 | # | 20 | Baseline | paralle | 88 |
| # | sex | 4 | 6 | 0 | 0.4 | 1 | # | 20 | Baseline | paralle | 88 |
| # | time | 5 | 5 | 0 | 0.5 | 1 | # | 20 | Baseline | paralle | 88 |
| # | interaction | 4 | 1 | 0 | 0.8 | 1 | # | 20 | Baseline | paralle | 88 |
| # | disease | 7 | 4 | 0 | 0.636363636 | 1 | # | 20 | Baseline | ancom | 88 |
| # | sex | 4 | 0 | 0 | 1 | 1 | # | 20 | Baseline | ancom | 88 |
| # | time | 5 | 4 | 0 | 0.555555556 | 1 | # | 20 | Baseline | ancom | 88 |
| # | interaction | 0 | 0 | 4 | 0 | 0 | # | 20 | Baseline | ancom | 88 |
| # | disease | 7 | 61 | 0 | 0.102941176 | 1 | # | 20 | Baseline | aldex2 | 88 |
| # | sex | 4 | 27 | 0 | 0.129032258 | 1 | # | 20 | Baseline | aldex2 | 88 |
| # | time | 5 | 39 | 0 | 0.113636364 | 1 | # | 20 | Baseline | aldex2 | 88 |
| # | interaction | 0 | 0 | 4 | 0 | 0 | # | 20 | Baseline | aldex2 | 88 |
| # | disease | 7 | 0 | 0 | 1 | 1 | # | 20 | Small | paralle | 88 |
| # | sex | 4 | 0 | 0 | 1 | 1 | # | 20 | Small | paralle | 88 |
| # | time | 5 | 0 | 0 | 1 | 1 | # | 20 | Small | paralle | 88 |
| # | interaction | 4 | 0 | 0 | 1 | 1 | # | 20 | Small | paralle | 88 |
| # | disease | 7 | 4 | 0 | 0.636363636 | 1 | # | 20 | Small | ancom | 88 |
| # | sex | 4 | 0 | 0 | 1 | 1 | # | 20 | Small | ancom | 88 |
| # | time | 5 | 4 | 0 | 0.555555556 | 1 | # | 20 | Small | ancom | 88 |
| # | interaction | 0 | 0 | 4 | 0 | 0 | # | 20 | Small | ancom | 88 |
| # | disease | 7 | 6 | 0 | 0.538461538 | 1 | # | 20 | Small | aldex2 | 88 |
| # | sex | 4 | 0 | 0 | 1 | 1 | # | 20 | Small | aldex2 | 88 |
| # | time | 5 | 8 | 0 | 0.384615385 | 1 | # | 20 | Small | aldex2 | 88 |
| # | interaction | 0 | 0 | 4 | 0 | 0 | # | 20 | Small | aldex2 | 88 |
| # | disease | 4 | 10 | 0 | 0.285714286 | 1 | # | 10 | Baseline | paralle | 89 |
| # | sex | 2 | 9 | 0 | 0.181818182 | 1 | # | 10 | Baseline | paralle | 89 |
| # | time | 2 | 7 | 0 | 0.222222222 | 1 | # | 10 | Baseline | paralle | 89 |
| # | interaction | 2 | 5 | 0 | 0.285714286 | 1 | # | 10 | Baseline | paralle | 89 |
| # | disease | 4 | 0 | 0 | 1 | 1 | # | 10 | Baseline | ancom | 89 |
| # | sex | 2 | 0 | 0 | 1 | 1 | # | 10 | Baseline | ancom | 89 |
| # | time | 2 | 0 | 0 | 1 | 1 | # | 10 | Baseline | ancom | 89 |
| # | interaction | 0 | 0 | 2 | 0 | 0 | # | 10 | Baseline | ancom | 89 |
| # | disease | 4 | 14 | 0 | 0.222222222 | 1 | # | 10 | Baseline | aldex2 | 89 |
| # | sex | 2 | 5 | 0 | 0.285714286 | 1 | # | 10 | Baseline | aldex2 | 89 |
| # | time | 2 | 9 | 0 | 0.181818182 | 1 | # | 10 | Baseline | aldex2 | 89 |
| # | interaction | 0 | 0 | 2 | 0 | 0 | # | 10 | Baseline | aldex2 | 89 |
| # | disease | 4 | 0 | 0 | 1 | 1 | # | 10 | Small | paralle | 89 |
| # | sex | 2 | 0 | 0 | 1 | 1 | # | 10 | Small | paralle | 89 |
| # | time | 2 | 0 | 0 | 1 | 1 | # | 10 | Small | paralle | 89 |
| # | interaction | 1 | 1 | 1 | 0.5 | 0.5 | # | 10 | Small | paralle | 89 |
| # | disease | 4 | 0 | 0 | 1 | 1 | # | 10 | Small | ancom | 89 |
| # | sex | 2 | 0 | 0 | 1 | 1 | # | 10 | Small | ancom | 89 |
| # | time | 2 | 0 | 0 | 1 | 1 | # | 10 | Small | ancom | 89 |
| # | interaction | 0 | 0 | 2 | 0 | 0 | # | 10 | Small | ancom | 89 |
| # | disease | 4 | 0 | 0 | 1 | 1 | # | 10 | Small | aldex2 | 89 |
| # | sex | 2 | 0 | 0 | 1 | 1 | # | 10 | Small | aldex2 | 89 |
| # | time | 2 | 0 | 0 | 1 | 1 | # | 10 | Small | aldex2 | 89 |
| # | interaction | 0 | 0 | 2 | 0 | 0 | # | 10 | Small | aldex2 | 89 |
| # | disease | 4 | 8 | 0 | 0.333333333 | 1 | # | 10 | Baseline | paralle | 89 |
| # | sex | 2 | 6 | 0 | 0.25 | 1 | # | 10 | Baseline | paralle | 89 |
| # | time | 2 | 5 | 0 | 0.285714286 | 1 | # | 10 | Baseline | paralle | 89 |
| # | interaction | 2 | 8 | 0 | 0.2 | 1 | # | 10 | Baseline | paralle | 89 |
| # | disease | 4 | 1 | 0 | 0.8 | 1 | # | 10 | Baseline | ancom | 89 |
| # | sex | 2 | 0 | 0 | 1 | 1 | # | 10 | Baseline | ancom | 89 |
| # | time | 2 | 1 | 0 | 0.666666667 | 1 | # | 10 | Baseline | ancom | 89 |
| # | interaction | 0 | 0 | 2 | 0 | 0 | # | 10 | Baseline | ancom | 89 |
| # | disease | 4 | 18 | 0 | 0.181818182 | 1 | # | 10 | Baseline | aldex2 | 89 |
| # | sex | 2 | 6 | 0 | 0.25 | 1 | # | 10 | Baseline | aldex2 | 89 |
| # | time | 2 | 9 | 0 | 0.181818182 | 1 | # | 10 | Baseline | aldex2 | 89 |
| # | interaction | 0 | 0 | 2 | 0 | 0 | # | 10 | Baseline | aldex2 | 89 |
| # | disease | 4 | 1 | 0 | 0.8 | 1 | # | 10 | Small | paralle | 89 |
| # | sex | 2 | 0 | 0 | 1 | 1 | # | 10 | Small | paralle | 89 |
| # | time | 2 | 0 | 0 | 1 | 1 | # | 10 | Small | paralle | 89 |
| # | interaction | 1 | 1 | 1 | 0.5 | 0.5 | # | 10 | Small | paralle | 89 |
| # | disease | 4 | 1 | 0 | 0.8 | 1 | # | 10 | Small | ancom | 89 |
| # | sex | 2 | 0 | 0 | 1 | 1 | # | 10 | Small | ancom | 89 |
| # | time | 2 | 1 | 0 | 0.666666667 | 1 | # | 10 | Small | ancom | 89 |
| # | interaction | 0 | 0 | 2 | 0 | 0 | # | 10 | Small | ancom | 89 |
| # | disease | 4 | 2 | 0 | 0.666666667 | 1 | # | 10 | Small | aldex2 | 89 |
| # | sex | 2 | 0 | 0 | 1 | 1 | # | 10 | Small | aldex2 | 89 |
| # | time | 2 | 2 | 0 | 0.5 | 1 | # | 10 | Small | aldex2 | 89 |
| # | interaction | 0 | 0 | 2 | 0 | 0 | # | 10 | Small | aldex2 | 89 |
| # | disease | 4 | 6 | 0 | 0.4 | 1 | # | 10 | Baseline | paralle | 89 |
| # | sex | 2 | 4 | 0 | 0.333333333 | 1 | # | 10 | Baseline | paralle | 89 |
| # | time | 2 | 1 | 0 | 0.666666667 | 1 | # | 10 | Baseline | paralle | 89 |
| # | interaction | 2 | 4 | 0 | 0.333333333 | 1 | # | 10 | Baseline | paralle | 89 |
| # | disease | 4 | 2 | 0 | 0.666666667 | 1 | # | 10 | Baseline | ancom | 89 |
| # | sex | 2 | 0 | 0 | 1 | 1 | # | 10 | Baseline | ancom | 89 |
| # | time | 2 | 1 | 0 | 0.666666667 | 1 | # | 10 | Baseline | ancom | 89 |
| # | interaction | 0 | 0 | 2 | 0 | 0 | # | 10 | Baseline | ancom | 89 |
| # | disease | 4 | 21 | 0 | 0.16 | 1 | # | 10 | Baseline | aldex2 | 89 |
| # | sex | 2 | 12 | 0 | 0.142857143 | 1 | # | 10 | Baseline | aldex2 | 89 |
| # | time | 2 | 8 | 0 | 0.2 | 1 | # | 10 | Baseline | aldex2 | 89 |
| # | interaction | 0 | 0 | 2 | 0 | 0 | # | 10 | Baseline | aldex2 | 89 |
| # | disease | 4 | 0 | 0 | 1 | 1 | # | 10 | Small | paralle | 89 |
| # | sex | 2 | 0 | 0 | 1 | 1 | # | 10 | Small | paralle | 89 |
| # | time | 2 | 0 | 0 | 1 | 1 | # | 10 | Small | paralle | 89 |
| # | interaction | 2 | 0 | 0 | 1 | 1 | # | 10 | Small | paralle | 89 |
| # | disease | 4 | 2 | 0 | 0.666666667 | 1 | # | 10 | Small | ancom | 89 |
| # | sex | 2 | 0 | 0 | 1 | 1 | # | 10 | Small | ancom | 89 |
| # | time | 2 | 1 | 0 | 0.666666667 | 1 | # | 10 | Small | ancom | 89 |
| # | interaction | 0 | 0 | 2 | 0 | 0 | # | 10 | Small | ancom | 89 |
| # | disease | 4 | 2 | 0 | 0.666666667 | 1 | # | 10 | Small | aldex2 | 89 |
| # | sex | 2 | 1 | 0 | 0.666666667 | 1 | # | 10 | Small | aldex2 | 89 |
| # | time | 2 | 2 | 0 | 0.5 | 1 | # | 10 | Small | aldex2 | 89 |
| # | interaction | 0 | 0 | 2 | 0 | 0 | # | 10 | Small | aldex2 | 89 |
| # | disease | 4 | 5 | 0 | 0.444444444 | 1 | # | 10 | Baseline | paralle | 89 |
| # | sex | 2 | 5 | 0 | 0.285714286 | 1 | # | 10 | Baseline | paralle | 89 |
| # | time | 2 | 0 | 0 | 1 | 1 | # | 10 | Baseline | paralle | 89 |
| # | interaction | 2 | 2 | 0 | 0.5 | 1 | # | 10 | Baseline | paralle | 89 |
| # | disease | 4 | 2 | 0 | 0.666666667 | 1 | # | 10 | Baseline | ancom | 89 |
| # | sex | 2 | 0 | 0 | 1 | 1 | # | 10 | Baseline | ancom | 89 |
| # | time | 2 | 2 | 0 | 0.5 | 1 | # | 10 | Baseline | ancom | 89 |
| # | interaction | 0 | 0 | 2 | 0 | 0 | # | 10 | Baseline | ancom | 89 |
| # | disease | 4 | 27 | 0 | 0.129032258 | 1 | # | 10 | Baseline | aldex2 | 89 |
| # | sex | 2 | 11 | 0 | 0.153846154 | 1 | # | 10 | Baseline | aldex2 | 89 |
| # | time | 2 | 10 | 0 | 0.166666667 | 1 | # | 10 | Baseline | aldex2 | 89 |
| # | interaction | 0 | 0 | 2 | 0 | 0 | # | 10 | Baseline | aldex2 | 89 |
| # | disease | 4 | 0 | 0 | 1 | 1 | # | 10 | Small | paralle | 89 |
| # | sex | 2 | 0 | 0 | 1 | 1 | # | 10 | Small | paralle | 89 |
| # | time | 2 | 0 | 0 | 1 | 1 | # | 10 | Small | paralle | 89 |
| # | interaction | 2 | 0 | 0 | 1 | 1 | # | 10 | Small | paralle | 89 |
| # | disease | 4 | 2 | 0 | 0.666666667 | 1 | # | 10 | Small | ancom | 89 |
| # | sex | 2 | 0 | 0 | 1 | 1 | # | 10 | Small | ancom | 89 |
| # | time | 2 | 2 | 0 | 0.5 | 1 | # | 10 | Small | ancom | 89 |
| # | interaction | 0 | 0 | 2 | 0 | 0 | # | 10 | Small | ancom | 89 |
| # | disease | 4 | 2 | 0 | 0.666666667 | 1 | # | 10 | Small | aldex2 | 89 |
| # | sex | 2 | 0 | 0 | 1 | 1 | # | 10 | Small | aldex2 | 89 |
| # | time | 2 | 2 | 0 | 0.5 | 1 | # | 10 | Small | aldex2 | 89 |
| # | interaction | 0 | 0 | 2 | 0 | 0 | # | 10 | Small | aldex2 | 89 |
| # | disease | 7 | 13 | 0 | 0.35 | 1 | # | 20 | Baseline | paralle | 89 |
| # | sex | 4 | 8 | 0 | 0.333333333 | 1 | # | 20 | Baseline | paralle | 89 |
| # | time | 5 | 6 | 0 | 0.454545455 | 1 | # | 20 | Baseline | paralle | 89 |
| # | interaction | 4 | 8 | 0 | 0.333333333 | 1 | # | 20 | Baseline | paralle | 89 |
| # | disease | 7 | 0 | 0 | 1 | 1 | # | 20 | Baseline | ancom | 89 |
| # | sex | 4 | 0 | 0 | 1 | 1 | # | 20 | Baseline | ancom | 89 |
| # | time | 5 | 1 | 0 | 0.833333333 | 1 | # | 20 | Baseline | ancom | 89 |
| # | interaction | 0 | 0 | 4 | 0 | 0 | # | 20 | Baseline | ancom | 89 |
| # | disease | 7 | 14 | 0 | 0.333333333 | 1 | # | 20 | Baseline | aldex2 | 89 |
| # | sex | 4 | 8 | 0 | 0.333333333 | 1 | # | 20 | Baseline | aldex2 | 89 |
| # | time | 5 | 8 | 0 | 0.384615385 | 1 | # | 20 | Baseline | aldex2 | 89 |
| # | interaction | 0 | 0 | 4 | 0 | 0 | # | 20 | Baseline | aldex2 | 89 |
| # | disease | 7 | 2 | 0 | 0.777777778 | 1 | # | 20 | Small | paralle | 89 |
| # | sex | 4 | 0 | 0 | 1 | 1 | # | 20 | Small | paralle | 89 |
| # | time | 5 | 0 | 0 | 1 | 1 | # | 20 | Small | paralle | 89 |
| # | interaction | 2 | 1 | 2 | 0.666666667 | 0.5 | # | 20 | Small | paralle | 89 |
| # | disease | 7 | 0 | 0 | 1 | 1 | # | 20 | Small | ancom | 89 |
| # | sex | 4 | 0 | 0 | 1 | 1 | # | 20 | Small | ancom | 89 |
| # | time | 5 | 1 | 0 | 0.833333333 | 1 | # | 20 | Small | ancom | 89 |
| # | interaction | 0 | 0 | 4 | 0 | 0 | # | 20 | Small | ancom | 89 |
| # | disease | 7 | 0 | 0 | 1 | 1 | # | 20 | Small | aldex2 | 89 |
| # | sex | 4 | 0 | 0 | 1 | 1 | # | 20 | Small | aldex2 | 89 |
| # | time | 5 | 2 | 0 | 0.714285714 | 1 | # | 20 | Small | aldex2 | 89 |
| # | interaction | 0 | 0 | 4 | 0 | 0 | # | 20 | Small | aldex2 | 89 |
| # | disease | 7 | 14 | 0 | 0.333333333 | 1 | # | 20 | Baseline | paralle | 89 |
| # | sex | 4 | 7 | 0 | 0.363636364 | 1 | # | 20 | Baseline | paralle | 89 |
| # | time | 5 | 5 | 0 | 0.5 | 1 | # | 20 | Baseline | paralle | 89 |
| # | interaction | 4 | 6 | 0 | 0.4 | 1 | # | 20 | Baseline | paralle | 89 |
| # | disease | 7 | 2 | 0 | 0.777777778 | 1 | # | 20 | Baseline | ancom | 89 |
| # | sex | 4 | 0 | 0 | 1 | 1 | # | 20 | Baseline | ancom | 89 |
| # | time | 5 | 3 | 0 | 0.625 | 1 | # | 20 | Baseline | ancom | 89 |
| # | interaction | 0 | 0 | 4 | 0 | 0 | # | 20 | Baseline | ancom | 89 |
| # | disease | 7 | 35 | 0 | 0.166666667 | 1 | # | 20 | Baseline | aldex2 | 89 |
| # | sex | 4 | 12 | 0 | 0.25 | 1 | # | 20 | Baseline | aldex2 | 89 |
| # | time | 5 | 25 | 0 | 0.166666667 | 1 | # | 20 | Baseline | aldex2 | 89 |
| # | interaction | 0 | 0 | 4 | 0 | 0 | # | 20 | Baseline | aldex2 | 89 |
| # | disease | 7 | 0 | 0 | 1 | 1 | # | 20 | Small | paralle | 89 |
| # | sex | 4 | 0 | 0 | 1 | 1 | # | 20 | Small | paralle | 89 |
| # | time | 5 | 0 | 0 | 1 | 1 | # | 20 | Small | paralle | 89 |
| # | interaction | 2 | 0 | 2 | 1 | 0.5 | # | 20 | Small | paralle | 89 |
| # | disease | 7 | 2 | 0 | 0.777777778 | 1 | # | 20 | Small | ancom | 89 |
| # | sex | 4 | 0 | 0 | 1 | 1 | # | 20 | Small | ancom | 89 |
| # | time | 5 | 1 | 0 | 0.833333333 | 1 | # | 20 | Small | ancom | 89 |
| # | interaction | 0 | 0 | 4 | 0 | 0 | # | 20 | Small | ancom | 89 |
| # | disease | 7 | 2 | 0 | 0.777777778 | 1 | # | 20 | Small | aldex2 | 89 |
| # | sex | 4 | 0 | 0 | 1 | 1 | # | 20 | Small | aldex2 | 89 |
| # | time | 5 | 2 | 0 | 0.714285714 | 1 | # | 20 | Small | aldex2 | 89 |
| # | interaction | 0 | 0 | 4 | 0 | 0 | # | 20 | Small | aldex2 | 89 |
| # | disease | 7 | 17 | 0 | 0.291666667 | 1 | # | 20 | Baseline | paralle | 89 |
| # | sex | 4 | 5 | 0 | 0.444444444 | 1 | # | 20 | Baseline | paralle | 89 |
| # | time | 5 | 5 | 0 | 0.5 | 1 | # | 20 | Baseline | paralle | 89 |
| # | interaction | 4 | 8 | 0 | 0.333333333 | 1 | # | 20 | Baseline | paralle | 89 |
| # | disease | 7 | 4 | 0 | 0.636363636 | 1 | # | 20 | Baseline | ancom | 89 |
| # | sex | 4 | 0 | 0 | 1 | 1 | # | 20 | Baseline | ancom | 89 |
| # | time | 5 | 3 | 0 | 0.625 | 1 | # | 20 | Baseline | ancom | 89 |
| # | interaction | 0 | 0 | 4 | 0 | 0 | # | 20 | Baseline | ancom | 89 |
| # | disease | 7 | 46 | 0 | 0.132075472 | 1 | # | 20 | Baseline | aldex2 | 89 |
| # | sex | 4 | 13 | 0 | 0.235294118 | 1 | # | 20 | Baseline | aldex2 | 89 |
| # | time | 5 | 37 | 0 | 0.119047619 | 1 | # | 20 | Baseline | aldex2 | 89 |
| # | interaction | 0 | 0 | 4 | 0 | 0 | # | 20 | Baseline | aldex2 | 89 |
| # | disease | 7 | 0 | 0 | 1 | 1 | # | 20 | Small | paralle | 89 |
| # | sex | 4 | 0 | 0 | 1 | 1 | # | 20 | Small | paralle | 89 |
| # | time | 5 | 0 | 0 | 1 | 1 | # | 20 | Small | paralle | 89 |
| # | interaction | 3 | 0 | 1 | 1 | 0.75 | # | 20 | Small | paralle | 89 |
| # | disease | 7 | 4 | 0 | 0.636363636 | 1 | # | 20 | Small | ancom | 89 |
| # | sex | 4 | 0 | 0 | 1 | 1 | # | 20 | Small | ancom | 89 |
| # | time | 5 | 3 | 0 | 0.625 | 1 | # | 20 | Small | ancom | 89 |
| # | interaction | 0 | 0 | 4 | 0 | 0 | # | 20 | Small | ancom | 89 |
| # | disease | 7 | 6 | 0 | 0.538461538 | 1 | # | 20 | Small | aldex2 | 89 |
| # | sex | 4 | 0 | 0 | 1 | 1 | # | 20 | Small | aldex2 | 89 |
| # | time | 5 | 3 | 0 | 0.625 | 1 | # | 20 | Small | aldex2 | 89 |
| # | interaction | 0 | 0 | 4 | 0 | 0 | # | 20 | Small | aldex2 | 89 |
| # | disease | 7 | 18 | 0 | 0.28 | 1 | # | 20 | Baseline | paralle | 89 |
| # | sex | 4 | 6 | 0 | 0.4 | 1 | # | 20 | Baseline | paralle | 89 |
| # | time | 5 | 7 | 0 | 0.416666667 | 1 | # | 20 | Baseline | paralle | 89 |
| # | interaction | 4 | 4 | 0 | 0.5 | 1 | # | 20 | Baseline | paralle | 89 |
| # | disease | 7 | 4 | 0 | 0.636363636 | 1 | # | 20 | Baseline | ancom | 89 |
| # | sex | 4 | 0 | 0 | 1 | 1 | # | 20 | Baseline | ancom | 89 |
| # | time | 5 | 4 | 0 | 0.555555556 | 1 | # | 20 | Baseline | ancom | 89 |
| # | interaction | 0 | 0 | 4 | 0 | 0 | # | 20 | Baseline | ancom | 89 |
| # | disease | 7 | 67 | 0 | 0.094594595 | 1 | # | 20 | Baseline | aldex2 | 89 |
| # | sex | 4 | 18 | 0 | 0.181818182 | 1 | # | 20 | Baseline | aldex2 | 89 |
| # | time | 5 | 46 | 0 | 0.098039216 | 1 | # | 20 | Baseline | aldex2 | 89 |
| # | interaction | 0 | 0 | 4 | 0 | 0 | # | 20 | Baseline | aldex2 | 89 |
| # | disease | 7 | 0 | 0 | 1 | 1 | # | 20 | Small | paralle | 89 |
| # | sex | 4 | 0 | 0 | 1 | 1 | # | 20 | Small | paralle | 89 |
| # | time | 5 | 0 | 0 | 1 | 1 | # | 20 | Small | paralle | 89 |
| # | interaction | 4 | 0 | 0 | 1 | 1 | # | 20 | Small | paralle | 89 |
| # | disease | 7 | 4 | 0 | 0.636363636 | 1 | # | 20 | Small | ancom | 89 |
| # | sex | 4 | 0 | 0 | 1 | 1 | # | 20 | Small | ancom | 89 |
| # | time | 5 | 4 | 0 | 0.555555556 | 1 | # | 20 | Small | ancom | 89 |
| # | interaction | 0 | 0 | 4 | 0 | 0 | # | 20 | Small | ancom | 89 |
| # | disease | 7 | 7 | 0 | 0.5 | 1 | # | 20 | Small | aldex2 | 89 |
| # | sex | 4 | 1 | 0 | 0.8 | 1 | # | 20 | Small | aldex2 | 89 |
| # | time | 5 | 8 | 0 | 0.384615385 | 1 | # | 20 | Small | aldex2 | 89 |
| # | interaction | 0 | 0 | 4 | 0 | 0 | # | 20 | Small | aldex2 | 89 |
| # | disease | 4 | 5 | 0 | 0.444444444 | 1 | # | 10 | Baseline | paralle | 90 |
| # | sex | 2 | 5 | 0 | 0.285714286 | 1 | # | 10 | Baseline | paralle | 90 |
| # | time | 2 | 7 | 0 | 0.222222222 | 1 | # | 10 | Baseline | paralle | 90 |
| # | interaction | 2 | 9 | 0 | 0.181818182 | 1 | # | 10 | Baseline | paralle | 90 |
| # | disease | 4 | 1 | 0 | 0.8 | 1 | # | 10 | Baseline | ancom | 90 |
| # | sex | 2 | 1 | 0 | 0.666666667 | 1 | # | 10 | Baseline | ancom | 90 |
| # | time | 2 | 1 | 0 | 0.666666667 | 1 | # | 10 | Baseline | ancom | 90 |
| # | interaction | 0 | 0 | 2 | 0 | 0 | # | 10 | Baseline | ancom | 90 |
| # | disease | 4 | 10 | 0 | 0.285714286 | 1 | # | 10 | Baseline | aldex2 | 90 |
| # | sex | 2 | 3 | 0 | 0.4 | 1 | # | 10 | Baseline | aldex2 | 90 |
| # | time | 2 | 8 | 0 | 0.2 | 1 | # | 10 | Baseline | aldex2 | 90 |
| # | interaction | 0 | 0 | 2 | 0 | 0 | # | 10 | Baseline | aldex2 | 90 |
| # | disease | 4 | 0 | 0 | 1 | 1 | # | 10 | Small | paralle | 90 |
| # | sex | 2 | 0 | 0 | 1 | 1 | # | 10 | Small | paralle | 90 |
| # | time | 2 | 1 | 0 | 0.666666667 | 1 | # | 10 | Small | paralle | 90 |
| # | interaction | 1 | 0 | 1 | 1 | 0.5 | # | 10 | Small | paralle | 90 |
| # | disease | 4 | 0 | 0 | 1 | 1 | # | 10 | Small | ancom | 90 |
| # | sex | 2 | 1 | 0 | 0.666666667 | 1 | # | 10 | Small | ancom | 90 |
| # | time | 2 | 0 | 0 | 1 | 1 | # | 10 | Small | ancom | 90 |
| # | interaction | 0 | 0 | 2 | 0 | 0 | # | 10 | Small | ancom | 90 |
| # | disease | 4 | 0 | 0 | 1 | 1 | # | 10 | Small | aldex2 | 90 |
| # | sex | 2 | 1 | 0 | 0.666666667 | 1 | # | 10 | Small | aldex2 | 90 |
| # | time | 2 | 1 | 0 | 0.666666667 | 1 | # | 10 | Small | aldex2 | 90 |
| # | interaction | 0 | 0 | 2 | 0 | 0 | # | 10 | Small | aldex2 | 90 |
| # | disease | 4 | 9 | 0 | 0.307692308 | 1 | # | 10 | Baseline | paralle | 90 |
| # | sex | 2 | 6 | 0 | 0.25 | 1 | # | 10 | Baseline | paralle | 90 |
| # | time | 2 | 4 | 0 | 0.333333333 | 1 | # | 10 | Baseline | paralle | 90 |
| # | interaction | 2 | 5 | 0 | 0.285714286 | 1 | # | 10 | Baseline | paralle | 90 |
| # | disease | 4 | 1 | 0 | 0.8 | 1 | # | 10 | Baseline | ancom | 90 |
| # | sex | 2 | 0 | 0 | 1 | 1 | # | 10 | Baseline | ancom | 90 |
| # | time | 2 | 2 | 0 | 0.5 | 1 | # | 10 | Baseline | ancom | 90 |
| # | interaction | 0 | 0 | 2 | 0 | 0 | # | 10 | Baseline | ancom | 90 |
| # | disease | 4 | 18 | 0 | 0.181818182 | 1 | # | 10 | Baseline | aldex2 | 90 |
| # | sex | 2 | 4 | 0 | 0.333333333 | 1 | # | 10 | Baseline | aldex2 | 90 |
| # | time | 2 | 7 | 0 | 0.222222222 | 1 | # | 10 | Baseline | aldex2 | 90 |
| # | interaction | 0 | 0 | 2 | 0 | 0 | # | 10 | Baseline | aldex2 | 90 |
| # | disease | 4 | 0 | 0 | 1 | 1 | # | 10 | Small | paralle | 90 |
| # | sex | 2 | 0 | 0 | 1 | 1 | # | 10 | Small | paralle | 90 |
| # | time | 2 | 0 | 0 | 1 | 1 | # | 10 | Small | paralle | 90 |
| # | interaction | 2 | 1 | 0 | 0.666666667 | 1 | # | 10 | Small | paralle | 90 |
| # | disease | 4 | 0 | 0 | 1 | 1 | # | 10 | Small | ancom | 90 |
| # | sex | 2 | 0 | 0 | 1 | 1 | # | 10 | Small | ancom | 90 |
| # | time | 2 | 2 | 0 | 0.5 | 1 | # | 10 | Small | ancom | 90 |
| # | interaction | 0 | 0 | 2 | 0 | 0 | # | 10 | Small | ancom | 90 |
| # | disease | 4 | 0 | 0 | 1 | 1 | # | 10 | Small | aldex2 | 90 |
| # | sex | 2 | 0 | 0 | 1 | 1 | # | 10 | Small | aldex2 | 90 |
| # | time | 2 | 2 | 0 | 0.5 | 1 | # | 10 | Small | aldex2 | 90 |
| # | interaction | 0 | 0 | 2 | 0 | 0 | # | 10 | Small | aldex2 | 90 |
| # | disease | 4 | 5 | 0 | 0.444444444 | 1 | # | 10 | Baseline | paralle | 90 |
| # | sex | 2 | 3 | 0 | 0.4 | 1 | # | 10 | Baseline | paralle | 90 |
| # | time | 2 | 4 | 0 | 0.333333333 | 1 | # | 10 | Baseline | paralle | 90 |
| # | interaction | 2 | 7 | 0 | 0.222222222 | 1 | # | 10 | Baseline | paralle | 90 |
| # | disease | 4 | 2 | 0 | 0.666666667 | 1 | # | 10 | Baseline | ancom | 90 |
| # | sex | 2 | 0 | 0 | 1 | 1 | # | 10 | Baseline | ancom | 90 |
| # | time | 2 | 2 | 0 | 0.5 | 1 | # | 10 | Baseline | ancom | 90 |
| # | interaction | 0 | 0 | 2 | 0 | 0 | # | 10 | Baseline | ancom | 90 |
| # | disease | 4 | 23 | 0 | 0.148148148 | 1 | # | 10 | Baseline | aldex2 | 90 |
| # | sex | 2 | 2 | 0 | 0.5 | 1 | # | 10 | Baseline | aldex2 | 90 |
| # | time | 2 | 10 | 0 | 0.166666667 | 1 | # | 10 | Baseline | aldex2 | 90 |
| # | interaction | 0 | 0 | 2 | 0 | 0 | # | 10 | Baseline | aldex2 | 90 |
| # | disease | 4 | 0 | 0 | 1 | 1 | # | 10 | Small | paralle | 90 |
| # | sex | 2 | 0 | 0 | 1 | 1 | # | 10 | Small | paralle | 90 |
| # | time | 2 | 1 | 0 | 0.666666667 | 1 | # | 10 | Small | paralle | 90 |
| # | interaction | 2 | 0 | 0 | 1 | 1 | # | 10 | Small | paralle | 90 |
| # | disease | 4 | 2 | 0 | 0.666666667 | 1 | # | 10 | Small | ancom | 90 |
| # | sex | 2 | 0 | 0 | 1 | 1 | # | 10 | Small | ancom | 90 |
| # | time | 2 | 2 | 0 | 0.5 | 1 | # | 10 | Small | ancom | 90 |
| # | interaction | 0 | 0 | 2 | 0 | 0 | # | 10 | Small | ancom | 90 |
| # | disease | 4 | 2 | 0 | 0.666666667 | 1 | # | 10 | Small | aldex2 | 90 |
| # | sex | 2 | 0 | 0 | 1 | 1 | # | 10 | Small | aldex2 | 90 |
| # | time | 2 | 2 | 0 | 0.5 | 1 | # | 10 | Small | aldex2 | 90 |
| # | interaction | 0 | 0 | 2 | 0 | 0 | # | 10 | Small | aldex2 | 90 |
| # | disease | 4 | 7 | 0 | 0.363636364 | 1 | # | 10 | Baseline | paralle | 90 |
| # | sex | 2 | 6 | 0 | 0.25 | 1 | # | 10 | Baseline | paralle | 90 |
| # | time | 2 | 2 | 0 | 0.5 | 1 | # | 10 | Baseline | paralle | 90 |
| # | interaction | 2 | 6 | 0 | 0.25 | 1 | # | 10 | Baseline | paralle | 90 |
| # | disease | 4 | 2 | 0 | 0.666666667 | 1 | # | 10 | Baseline | ancom | 90 |
| # | sex | 2 | 0 | 0 | 1 | 1 | # | 10 | Baseline | ancom | 90 |
| # | time | 2 | 2 | 0 | 0.5 | 1 | # | 10 | Baseline | ancom | 90 |
| # | interaction | 0 | 0 | 2 | 0 | 0 | # | 10 | Baseline | ancom | 90 |
| # | disease | 4 | 30 | 0 | 0.117647059 | 1 | # | 10 | Baseline | aldex2 | 90 |
| # | sex | 2 | 6 | 0 | 0.25 | 1 | # | 10 | Baseline | aldex2 | 90 |
| # | time | 2 | 13 | 0 | 0.133333333 | 1 | # | 10 | Baseline | aldex2 | 90 |
| # | interaction | 0 | 0 | 2 | 0 | 0 | # | 10 | Baseline | aldex2 | 90 |
| # | disease | 4 | 0 | 0 | 1 | 1 | # | 10 | Small | paralle | 90 |
| # | sex | 2 | 0 | 0 | 1 | 1 | # | 10 | Small | paralle | 90 |
| # | time | 2 | 1 | 0 | 0.666666667 | 1 | # | 10 | Small | paralle | 90 |
| # | interaction | 2 | 0 | 0 | 1 | 1 | # | 10 | Small | paralle | 90 |
| # | disease | 4 | 1 | 0 | 0.8 | 1 | # | 10 | Small | ancom | 90 |
| # | sex | 2 | 0 | 0 | 1 | 1 | # | 10 | Small | ancom | 90 |
| # | time | 2 | 2 | 0 | 0.5 | 1 | # | 10 | Small | ancom | 90 |
| # | interaction | 0 | 0 | 2 | 0 | 0 | # | 10 | Small | ancom | 90 |
| # | disease | 4 | 1 | 0 | 0.8 | 1 | # | 10 | Small | aldex2 | 90 |
| # | sex | 2 | 0 | 0 | 1 | 1 | # | 10 | Small | aldex2 | 90 |
| # | time | 2 | 2 | 0 | 0.5 | 1 | # | 10 | Small | aldex2 | 90 |
| # | interaction | 0 | 0 | 2 | 0 | 0 | # | 10 | Small | aldex2 | 90 |

| | | | | | | | | | | | |
|---|---|---|---|---|---|---|---|---|---|---|---|
| # | disease | 7 | 13 | 0 | 0.35 | 1 | # | 20 | Baseline | paralle | 90 |
| # | sex | 4 | 10 | 0 | 0.285714286 | 1 | # | 20 | Baseline | paralle | 90 |
| # | time | 5 | 7 | 0 | 0.416666667 | 1 | # | 20 | Baseline | paralle | 90 |
| # | interaction | 4 | 6 | 0 | 0.4 | 1 | # | 20 | Baseline | paralle | 90 |
| # | disease | 7 | 1 | 0 | 0.875 | 1 | # | 20 | Baseline | ancom | 90 |
| # | sex | 4 | 0 | 0 | 1 | 1 | # | 20 | Baseline | ancom | 90 |
| # | time | 5 | 0 | 0 | 1 | 1 | # | 20 | Baseline | ancom | 90 |
| # | interaction | 0 | 0 | 4 | 0 | 0 | # | 20 | Baseline | ancom | 90 |
| # | disease | 7 | 23 | 0 | 0.233333333 | 1 | # | 20 | Baseline | aldex2 | 90 |
| # | sex | 4 | 10 | 0 | 0.285714286 | 1 | # | 20 | Baseline | aldex2 | 90 |
| # | time | 5 | 14 | 0 | 0.263157895 | 1 | # | 20 | Baseline | aldex2 | 90 |
| # | interaction | 0 | 0 | 4 | 0 | 0 | # | 20 | Baseline | aldex2 | 90 |
| # | disease | 7 | 0 | 0 | 1 | 1 | # | 20 | Small | paralle | 90 |
| # | sex | 4 | 1 | 0 | 0.8 | 1 | # | 20 | Small | paralle | 90 |
| # | time | 5 | 0 | 0 | 1 | 1 | # | 20 | Small | paralle | 90 |
| # | interaction | 1 | 1 | 3 | 0.5 | 0.25 | # | 20 | Small | paralle | 90 |
| # | disease | 7 | 0 | 0 | 1 | 1 | # | 20 | Small | ancom | 90 |
| # | sex | 4 | 0 | 0 | 1 | 1 | # | 20 | Small | ancom | 90 |
| # | time | 5 | 0 | 0 | 1 | 1 | # | 20 | Small | ancom | 90 |
| # | interaction | 0 | 0 | 4 | 0 | 0 | # | 20 | Small | ancom | 90 |
| # | disease | 7 | 2 | 0 | 0.777777778 | 1 | # | 20 | Small | aldex2 | 90 |
| # | sex | 4 | 1 | 0 | 0.8 | 1 | # | 20 | Small | aldex2 | 90 |
| # | time | 5 | 0 | 0 | 1 | 1 | # | 20 | Small | aldex2 | 90 |
| # | interaction | 0 | 0 | 4 | 0 | 0 | # | 20 | Small | aldex2 | 90 |
| # | disease | 7 | 5 | 0 | 0.583333333 | 1 | # | 20 | Baseline | paralle | 90 |
| # | sex | 4 | 4 | 0 | 0.5 | 1 | # | 20 | Baseline | paralle | 90 |
| # | time | 5 | 10 | 0 | 0.333333333 | 1 | # | 20 | Baseline | paralle | 90 |
| # | interaction | 4 | 7 | 0 | 0.363636364 | 1 | # | 20 | Baseline | paralle | 90 |
| # | disease | 7 | 4 | 0 | 0.636363636 | 1 | # | 20 | Baseline | ancom | 90 |
| # | sex | 4 | 0 | 0 | 1 | 1 | # | 20 | Baseline | ancom | 90 |
| # | time | 5 | 3 | 0 | 0.625 | 1 | # | 20 | Baseline | ancom | 90 |
| # | interaction | 0 | 0 | 4 | 0 | 0 | # | 20 | Baseline | ancom | 90 |
| # | disease | 7 | 39 | 0 | 0.152173913 | 1 | # | 20 | Baseline | aldex2 | 90 |
| # | sex | 4 | 9 | 0 | 0.307692308 | 1 | # | 20 | Baseline | aldex2 | 90 |
| # | time | 5 | 22 | 0 | 0.185185185 | 1 | # | 20 | Baseline | aldex2 | 90 |
| # | interaction | 0 | 0 | 4 | 0 | 0 | # | 20 | Baseline | aldex2 | 90 |
| # | disease | 7 | 0 | 0 | 1 | 1 | # | 20 | Small | paralle | 90 |
| # | sex | 4 | 0 | 0 | 1 | 1 | # | 20 | Small | paralle | 90 |
| # | time | 5 | 1 | 0 | 0.833333333 | 1 | # | 20 | Small | paralle | 90 |
| # | interaction | 4 | 0 | 0 | 1 | 1 | # | 20 | Small | paralle | 90 |
| # | disease | 7 | 4 | 0 | 0.636363636 | 1 | # | 20 | Small | ancom | 90 |
| # | sex | 4 | 0 | 0 | 1 | 1 | # | 20 | Small | ancom | 90 |
| # | time | 5 | 3 | 0 | 0.625 | 1 | # | 20 | Small | ancom | 90 |
| # | interaction | 0 | 0 | 4 | 0 | 0 | # | 20 | Small | ancom | 90 |
| # | disease | 7 | 2 | 0 | 0.777777778 | 1 | # | 20 | Small | aldex2 | 90 |
| # | sex | 4 | 0 | 0 | 1 | 1 | # | 20 | Small | aldex2 | 90 |
| # | time | 5 | 2 | 0 | 0.714285714 | 1 | # | 20 | Small | aldex2 | 90 |
| # | interaction | 0 | 0 | 4 | 0 | 0 | # | 20 | Small | aldex2 | 90 |
| # | disease | 7 | 7 | 0 | 0.5 | 1 | # | 20 | Baseline | paralle | 90 |
| # | sex | 4 | 5 | 0 | 0.444444444 | 1 | # | 20 | Baseline | paralle | 90 |
| # | time | 5 | 11 | 0 | 0.3125 | 1 | # | 20 | Baseline | paralle | 90 |
| # | interaction | 4 | 7 | 0 | 0.363636364 | 1 | # | 20 | Baseline | paralle | 90 |
| # | disease | 7 | 4 | 0 | 0.636363636 | 1 | # | 20 | Baseline | ancom | 90 |
| # | sex | 4 | 0 | 0 | 1 | 1 | # | 20 | Baseline | ancom | 90 |
| # | time | 5 | 4 | 0 | 0.555555556 | 1 | # | 20 | Baseline | ancom | 90 |
| # | interaction | 0 | 0 | 4 | 0 | 0 | # | 20 | Baseline | ancom | 90 |
| # | disease | 7 | 50 | 0 | 0.122807018 | 1 | # | 20 | Baseline | aldex2 | 90 |
| # | sex | 4 | 12 | 0 | 0.25 | 1 | # | 20 | Baseline | aldex2 | 90 |
| # | time | 5 | 33 | 0 | 0.131578947 | 1 | # | 20 | Baseline | aldex2 | 90 |
| # | interaction | 0 | 0 | 4 | 0 | 0 | # | 20 | Baseline | aldex2 | 90 |
| # | disease | 7 | 1 | 0 | 0.875 | 1 | # | 20 | Small | paralle | 90 |
| # | sex | 4 | 0 | 0 | 1 | 1 | # | 20 | Small | paralle | 90 |
| # | time | 5 | 0 | 0 | 1 | 1 | # | 20 | Small | paralle | 90 |
| # | interaction | 4 | 0 | 0 | 1 | 1 | # | 20 | Small | paralle | 90 |
| # | disease | 7 | 4 | 0 | 0.636363636 | 1 | # | 20 | Small | ancom | 90 |
| # | sex | 4 | 0 | 0 | 1 | 1 | # | 20 | Small | ancom | 90 |
| # | time | 5 | 3 | 0 | 0.625 | 1 | # | 20 | Small | ancom | 90 |
| # | interaction | 0 | 0 | 4 | 0 | 0 | # | 20 | Small | ancom | 90 |
| # | disease | 7 | 9 | 0 | 0.4375 | 1 | # | 20 | Small | aldex2 | 90 |
| # | sex | 4 | 0 | 0 | 1 | 1 | # | 20 | Small | aldex2 | 90 |
| # | time | 5 | 4 | 0 | 0.555555556 | 1 | # | 20 | Small | aldex2 | 90 |
| # | interaction | 0 | 0 | 4 | 0 | 0 | # | 20 | Small | aldex2 | 90 |
| # | disease | 7 | 9 | 0 | 0.4375 | 1 | # | 20 | Baseline | paralle | 90 |
| # | sex | 4 | 6 | 0 | 0.4 | 1 | # | 20 | Baseline | paralle | 90 |
| # | time | 5 | 13 | 0 | 0.277777778 | 1 | # | 20 | Baseline | paralle | 90 |
| # | interaction | 4 | 6 | 0 | 0.4 | 1 | # | 20 | Baseline | paralle | 90 |
| # | disease | 7 | 4 | 0 | 0.636363636 | 1 | # | 20 | Baseline | ancom | 90 |
| # | sex | 4 | 0 | 0 | 1 | 1 | # | 20 | Baseline | ancom | 90 |
| # | time | 5 | 4 | 0 | 0.555555556 | 1 | # | 20 | Baseline | ancom | 90 |
| # | interaction | 0 | 0 | 4 | 0 | 0 | # | 20 | Baseline | ancom | 90 |
| # | disease | 7 | 57 | 0 | 0.109375 | 1 | # | 20 | Baseline | aldex2 | 90 |
| # | sex | 4 | 15 | 0 | 0.210526316 | 1 | # | 20 | Baseline | aldex2 | 90 |
| # | time | 5 | 44 | 0 | 0.102040816 | 1 | # | 20 | Baseline | aldex2 | 90 |
| # | interaction | 0 | 0 | 4 | 0 | 0 | # | 20 | Baseline | aldex2 | 90 |
| # | disease | 7 | 1 | 0 | 0.875 | 1 | # | 20 | Small | paralle | 90 |
| # | sex | 4 | 0 | 0 | 1 | 1 | # | 20 | Small | paralle | 90 |
| # | time | 5 | 0 | 0 | 1 | 1 | # | 20 | Small | paralle | 90 |
| # | interaction | 4 | 0 | 0 | 1 | 1 | # | 20 | Small | paralle | 90 |
| # | disease | 7 | 4 | 0 | 0.636363636 | 1 | # | 20 | Small | ancom | 90 |
| # | sex | 4 | 0 | 0 | 1 | 1 | # | 20 | Small | ancom | 90 |
| # | time | 5 | 4 | 0 | 0.555555556 | 1 | # | 20 | Small | ancom | 90 |
| # | interaction | 0 | 0 | 4 | 0 | 0 | # | 20 | Small | ancom | 90 |
| # | disease | 7 | 13 | 0 | 0.35 | 1 | # | 20 | Small | aldex2 | 90 |
| # | sex | 4 | 1 | 0 | 0.8 | 1 | # | 20 | Small | aldex2 | 90 |
| # | time | 5 | 5 | 0 | 0.5 | 1 | # | 20 | Small | aldex2 | 90 |
| # | interaction | 0 | 0 | 4 | 0 | 0 | # | 20 | Small | aldex2 | 90 |
| # | disease | 4 | 3 | 0 | 0.571428571 | 1 | # | 10 | Baseline | paralle | 91 |
| # | sex | 2 | 5 | 0 | 0.285714286 | 1 | # | 10 | Baseline | paralle | 91 |
| # | time | 2 | 5 | 0 | 0.285714286 | 1 | # | 10 | Baseline | paralle | 91 |
| # | interaction | 2 | 3 | 0 | 0.4 | 1 | # | 10 | Baseline | paralle | 91 |
| # | disease | 4 | 0 | 0 | 1 | 1 | # | 10 | Baseline | ancom | 91 |
| # | sex | 2 | 0 | 0 | 1 | 1 | # | 10 | Baseline | ancom | 91 |
| # | time | 2 | 0 | 0 | 1 | 1 | # | 10 | Baseline | ancom | 91 |
| # | interaction | 0 | 0 | 2 | 0 | 0 | # | 10 | Baseline | ancom | 91 |
| # | disease | 4 | 5 | 0 | 0.444444444 | 1 | # | 10 | Baseline | aldex2 | 91 |
| # | sex | 2 | 5 | 0 | 0.285714286 | 1 | # | 10 | Baseline | aldex2 | 91 |
| # | time | 2 | 9 | 0 | 0.181818182 | 1 | # | 10 | Baseline | aldex2 | 91 |
| # | interaction | 0 | 0 | 2 | 0 | 0 | # | 10 | Baseline | aldex2 | 91 |
| # | disease | 4 | 0 | 0 | 1 | 1 | # | 10 | Small | paralle | 91 |
| # | sex | 2 | 0 | 0 | 1 | 1 | # | 10 | Small | paralle | 91 |
| # | time | 2 | 0 | 0 | 1 | 1 | # | 10 | Small | paralle | 91 |
| # | interaction | 1 | 0 | 1 | 1 | 0.5 | # | 10 | Small | paralle | 91 |
| # | disease | 4 | 0 | 0 | 1 | 1 | # | 10 | Small | ancom | 91 |
| # | sex | 2 | 0 | 0 | 1 | 1 | # | 10 | Small | ancom | 91 |
| # | time | 2 | 0 | 0 | 1 | 1 | # | 10 | Small | ancom | 91 |
| # | interaction | 0 | 0 | 2 | 0 | 0 | # | 10 | Small | ancom | 91 |
| # | disease | 4 | 0 | 0 | 1 | 1 | # | 10 | Small | aldex2 | 91 |
| # | sex | 2 | 1 | 0 | 0.666666667 | 1 | # | 10 | Small | aldex2 | 91 |
| # | time | 2 | 1 | 0 | 0.666666667 | 1 | # | 10 | Small | aldex2 | 91 |
| # | interaction | 0 | 0 | 2 | 0 | 0 | # | 10 | Small | aldex2 | 91 |
| # | disease | 4 | 2 | 0 | 0.666666667 | 1 | # | 10 | Baseline | paralle | 91 |
| # | sex | 2 | 4 | 0 | 0.333333333 | 1 | # | 10 | Baseline | paralle | 91 |
| # | time | 2 | 5 | 0 | 0.285714286 | 1 | # | 10 | Baseline | paralle | 91 |
| # | interaction | 2 | 1 | 0 | 0.666666667 | 1 | # | 10 | Baseline | paralle | 91 |
| # | disease | 4 | 2 | 0 | 0.666666667 | 1 | # | 10 | Baseline | ancom | 91 |
| # | sex | 2 | 0 | 0 | 1 | 1 | # | 10 | Baseline | ancom | 91 |
| # | time | 2 | 2 | 0 | 0.5 | 1 | # | 10 | Baseline | ancom | 91 |
| # | interaction | 0 | 0 | 2 | 0 | 0 | # | 10 | Baseline | ancom | 91 |
| # | disease | 4 | 16 | 0 | 0.2 | 1 | # | 10 | Baseline | aldex2 | 91 |
| # | sex | 2 | 4 | 0 | 0.333333333 | 1 | # | 10 | Baseline | aldex2 | 91 |
| # | time | 2 | 10 | 0 | 0.166666667 | 1 | # | 10 | Baseline | aldex2 | 91 |
| # | interaction | 0 | 0 | 2 | 0 | 0 | # | 10 | Baseline | aldex2 | 91 |
| # | disease | 4 | 0 | 0 | 1 | 1 | # | 10 | Small | paralle | 91 |
| # | sex | 2 | 0 | 0 | 1 | 1 | # | 10 | Small | paralle | 91 |
| # | time | 2 | 0 | 0 | 1 | 1 | # | 10 | Small | paralle | 91 |
| # | interaction | 2 | 0 | 0 | 1 | 1 | # | 10 | Small | paralle | 91 |
| # | disease | 4 | 2 | 0 | 0.666666667 | 1 | # | 10 | Small | ancom | 91 |
| # | sex | 2 | 0 | 0 | 1 | 1 | # | 10 | Small | ancom | 91 |
| # | time | 2 | 2 | 0 | 0.5 | 1 | # | 10 | Small | ancom | 91 |
| # | interaction | 0 | 0 | 2 | 0 | 0 | # | 10 | Small | ancom | 91 |
| # | disease | 4 | 1 | 0 | 0.8 | 1 | # | 10 | Small | aldex2 | 91 |
| # | sex | 2 | 0 | 0 | 1 | 1 | # | 10 | Small | aldex2 | 91 |
| # | time | 2 | 3 | 0 | 0.4 | 1 | # | 10 | Small | aldex2 | 91 |
| # | interaction | 0 | 0 | 2 | 0 | 0 | # | 10 | Small | aldex2 | 91 |
| # | disease | 4 | 5 | 0 | 0.444444444 | 1 | # | 10 | Baseline | paralle | 91 |
| # | sex | 2 | 1 | 0 | 0.666666667 | 1 | # | 10 | Baseline | paralle | 91 |
| # | time | 2 | 3 | 0 | 0.4 | 1 | # | 10 | Baseline | paralle | 91 |
| # | interaction | 2 | 2 | 0 | 0.5 | 1 | # | 10 | Baseline | paralle | 91 |
| # | disease | 4 | 2 | 0 | 0.666666667 | 1 | # | 10 | Baseline | ancom | 91 |
| # | sex | 2 | 0 | 0 | 1 | 1 | # | 10 | Baseline | ancom | 91 |
| # | time | 2 | 2 | 0 | 0.5 | 1 | # | 10 | Baseline | ancom | 91 |
| # | interaction | 0 | 0 | 2 | 0 | 0 | # | 10 | Baseline | ancom | 91 |
| # | disease | 4 | 20 | 0 | 0.166666667 | 1 | # | 10 | Baseline | aldex2 | 91 |
| # | sex | 2 | 5 | 0 | 0.285714286 | 1 | # | 10 | Baseline | aldex2 | 91 |
| # | time | 2 | 13 | 0 | 0.133333333 | 1 | # | 10 | Baseline | aldex2 | 91 |
| # | interaction | 0 | 0 | 2 | 0 | 0 | # | 10 | Baseline | aldex2 | 91 |
| # | disease | 4 | 0 | 0 | 1 | 1 | # | 10 | Small | paralle | 91 |
| # | sex | 2 | 0 | 0 | 1 | 1 | # | 10 | Small | paralle | 91 |
| # | time | 2 | 0 | 0 | 1 | 1 | # | 10 | Small | paralle | 91 |
| # | interaction | 2 | 0 | 0 | 1 | 1 | # | 10 | Small | paralle | 91 |
| # | disease | 4 | 2 | 0 | 0.666666667 | 1 | # | 10 | Small | ancom | 91 |
| # | sex | 2 | 0 | 0 | 1 | 1 | # | 10 | Small | ancom | 91 |
| # | time | 2 | 2 | 0 | 0.5 | 1 | # | 10 | Small | ancom | 91 |
| # | interaction | 0 | 0 | 2 | 0 | 0 | # | 10 | Small | ancom | 91 |
| # | disease | 4 | 3 | 0 | 0.571428571 | 1 | # | 10 | Small | aldex2 | 91 |
| # | sex | 2 | 0 | 0 | 1 | 1 | # | 10 | Small | aldex2 | 91 |
| # | time | 2 | 3 | 0 | 0.4 | 1 | # | 10 | Small | aldex2 | 91 |
| # | interaction | 0 | 0 | 2 | 0 | 0 | # | 10 | Small | aldex2 | 91 |
| # | disease | 4 | 5 | 0 | 0.444444444 | 1 | # | 10 | Baseline | paralle | 91 |
| # | sex | 2 | 1 | 0 | 0.666666667 | 1 | # | 10 | Baseline | paralle | 91 |
| # | time | 2 | 4 | 0 | 0.333333333 | 1 | # | 10 | Baseline | paralle | 91 |
| # | interaction | 2 | 2 | 0 | 0.5 | 1 | # | 10 | Baseline | paralle | 91 |
| # | disease | 4 | 2 | 0 | 0.666666667 | 1 | # | 10 | Baseline | ancom | 91 |
| # | sex | 2 | 0 | 0 | 1 | 1 | # | 10 | Baseline | ancom | 91 |
| # | time | 2 | 2 | 0 | 0.5 | 1 | # | 10 | Baseline | ancom | 91 |
| # | interaction | 0 | 0 | 2 | 0 | 0 | # | 10 | Baseline | ancom | 91 |
| # | disease | 4 | 26 | 0 | 0.133333333 | 1 | # | 10 | Baseline | aldex2 | 91 |
| # | sex | 2 | 6 | 0 | 0.25 | 1 | # | 10 | Baseline | aldex2 | 91 |
| # | time | 2 | 11 | 0 | 0.153846154 | 1 | # | 10 | Baseline | aldex2 | 91 |
| # | interaction | 0 | 0 | 2 | 0 | 0 | # | 10 | Baseline | aldex2 | 91 |
| # | disease | 4 | 0 | 0 | 1 | 1 | # | 10 | Small | paralle | 91 |
| # | sex | 2 | 0 | 0 | 1 | 1 | # | 10 | Small | paralle | 91 |
| # | time | 2 | 0 | 0 | 1 | 1 | # | 10 | Small | paralle | 91 |
| # | interaction | 2 | 0 | 0 | 1 | 1 | # | 10 | Small | paralle | 91 |
| # | disease | 4 | 2 | 0 | 0.666666667 | 1 | # | 10 | Small | ancom | 91 |
| # | sex | 2 | 0 | 0 | 1 | 1 | # | 10 | Small | ancom | 91 |
| # | time | 2 | 2 | 0 | 0.5 | 1 | # | 10 | Small | ancom | 91 |
| # | interaction | 0 | 0 | 2 | 0 | 0 | # | 10 | Small | ancom | 91 |
| # | disease | 4 | 2 | 0 | 0.666666667 | 1 | # | 10 | Small | aldex2 | 91 |
| # | sex | 2 | 0 | 0 | 1 | 1 | # | 10 | Small | aldex2 | 91 |
| # | time | 2 | 2 | 0 | 0.5 | 1 | # | 10 | Small | aldex2 | 91 |
| # | interaction | 0 | 0 | 2 | 0 | 0 | # | 10 | Small | aldex2 | 91 |
| # | disease | 7 | 10 | 0 | 0.411764706 | 1 | # | 20 | Baseline | paralle | 91 |
| # | sex | 4 | 9 | 0 | 0.307692308 | 1 | # | 20 | Baseline | paralle | 91 |
| # | time | 5 | 6 | 0 | 0.454545455 | 1 | # | 20 | Baseline | paralle | 91 |
| # | interaction | 4 | 6 | 0 | 0.4 | 1 | # | 20 | Baseline | paralle | 91 |
| # | disease | 7 | 1 | 0 | 0.875 | 1 | # | 20 | Baseline | ancom | 91 |
| # | sex | 4 | 0 | 0 | 1 | 1 | # | 20 | Baseline | ancom | 91 |
| # | time | 5 | 2 | 0 | 0.714285714 | 1 | # | 20 | Baseline | ancom | 91 |
| # | interaction | 0 | 0 | 4 | 0 | 0 | # | 20 | Baseline | ancom | 91 |
| # | disease | 7 | 19 | 0 | 0.269230769 | 1 | # | 20 | Baseline | aldex2 | 91 |
| # | sex | 4 | 6 | 0 | 0.4 | 1 | # | 20 | Baseline | aldex2 | 91 |
| # | time | 5 | 12 | 0 | 0.294117647 | 1 | # | 20 | Baseline | aldex2 | 91 |
| # | interaction | 0 | 0 | 4 | 0 | 0 | # | 20 | Baseline | aldex2 | 91 |
| # | disease | 7 | 0 | 0 | 1 | 1 | # | 20 | Small | paralle | 91 |
| # | sex | 4 | 1 | 0 | 0.8 | 1 | # | 20 | Small | paralle | 91 |
| # | time | 5 | 0 | 0 | 1 | 1 | # | 20 | Small | paralle | 91 |
| # | interaction | 0 | 0 | 4 | 0 | 0 | # | 20 | Small | paralle | 91 |
| # | disease | 7 | 1 | 0 | 0.875 | 1 | # | 20 | Small | ancom | 91 |
| # | sex | 4 | 0 | 0 | 1 | 1 | # | 20 | Small | ancom | 91 |
| # | time | 5 | 2 | 0 | 0.714285714 | 1 | # | 20 | Small | ancom | 91 |
| # | interaction | 0 | 0 | 4 | 0 | 0 | # | 20 | Small | ancom | 91 |
| # | disease | 7 | 1 | 0 | 0.875 | 1 | # | 20 | Small | aldex2 | 91 |
| # | sex | 4 | 0 | 0 | 1 | 1 | # | 20 | Small | aldex2 | 91 |
| # | time | 5 | 1 | 0 | 0.833333333 | 1 | # | 20 | Small | aldex2 | 91 |
| # | interaction | 0 | 0 | 4 | 0 | 0 | # | 20 | Small | aldex2 | 91 |
| # | disease | 7 | 6 | 0 | 0.538461538 | 1 | # | 20 | Baseline | paralle | 91 |
| # | sex | 4 | 6 | 0 | 0.4 | 1 | # | 20 | Baseline | paralle | 91 |
| # | time | 5 | 7 | 0 | 0.416666667 | 1 | # | 20 | Baseline | paralle | 91 |
| # | interaction | 4 | 5 | 0 | 0.444444444 | 1 | # | 20 | Baseline | paralle | 91 |
| # | disease | 7 | 3 | 0 | 0.7 | 1 | # | 20 | Baseline | ancom | 91 |
| # | sex | 4 | 0 | 0 | 1 | 1 | # | 20 | Baseline | ancom | 91 |
| # | time | 5 | 3 | 0 | 0.625 | 1 | # | 20 | Baseline | ancom | 91 |
| # | interaction | 0 | 0 | 4 | 0 | 0 | # | 20 | Baseline | ancom | 91 |
| # | disease | 7 | 30 | 0 | 0.189189189 | 1 | # | 20 | Baseline | aldex2 | 91 |
| # | sex | 4 | 6 | 0 | 0.4 | 1 | # | 20 | Baseline | aldex2 | 91 |
| # | time | 5 | 20 | 0 | 0.2 | 1 | # | 20 | Baseline | aldex2 | 91 |
| # | interaction | 0 | 0 | 4 | 0 | 0 | # | 20 | Baseline | aldex2 | 91 |
| # | disease | 7 | 0 | 0 | 1 | 1 | # | 20 | Small | paralle | 91 |
| # | sex | 4 | 0 | 0 | 1 | 1 | # | 20 | Small | paralle | 91 |
| # | time | 5 | 0 | 0 | 1 | 1 | # | 20 | Small | paralle | 91 |
| # | interaction | 4 | 0 | 0 | 1 | 1 | # | 20 | Small | paralle | 91 |
| # | disease | 7 | 3 | 0 | 0.7 | 1 | # | 20 | Small | ancom | 91 |
| # | sex | 4 | 0 | 0 | 1 | 1 | # | 20 | Small | ancom | 91 |
| # | time | 5 | 3 | 0 | 0.625 | 1 | # | 20 | Small | ancom | 91 |
| # | interaction | 0 | 0 | 4 | 0 | 0 | # | 20 | Small | ancom | 91 |
| # | disease | 7 | 1 | 0 | 0.875 | 1 | # | 20 | Small | aldex2 | 91 |
| # | sex | 4 | 0 | 0 | 1 | 1 | # | 20 | Small | aldex2 | 91 |
| # | time | 5 | 3 | 0 | 0.625 | 1 | # | 20 | Small | aldex2 | 91 |
| # | interaction | 0 | 0 | 4 | 0 | 0 | # | 20 | Small | aldex2 | 91 |
| # | disease | 7 | 8 | 0 | 0.466666667 | 1 | # | 20 | Baseline | paralle | 91 |
| # | sex | 4 | 11 | 0 | 0.266666667 | 1 | # | 20 | Baseline | paralle | 91 |
| # | time | 5 | 7 | 0 | 0.416666667 | 1 | # | 20 | Baseline | paralle | 91 |
| # | interaction | 4 | 4 | 0 | 0.5 | 1 | # | 20 | Baseline | paralle | 91 |
| # | disease | 7 | 4 | 0 | 0.636363636 | 1 | # | 20 | Baseline | ancom | 91 |
| # | sex | 4 | 0 | 0 | 1 | 1 | # | 20 | Baseline | ancom | 91 |
| # | time | 5 | 3 | 0 | 0.625 | 1 | # | 20 | Baseline | ancom | 91 |
| # | interaction | 0 | 0 | 4 | 0 | 0 | # | 20 | Baseline | ancom | 91 |
| # | disease | 7 | 53 | 0 | 0.116666667 | 1 | # | 20 | Baseline | aldex2 | 91 |
| # | sex | 4 | 14 | 0 | 0.222222222 | 1 | # | 20 | Baseline | aldex2 | 91 |
| # | time | 5 | 29 | 0 | 0.147058824 | 1 | # | 20 | Baseline | aldex2 | 91 |
| # | interaction | 0 | 0 | 4 | 0 | 0 | # | 20 | Baseline | aldex2 | 91 |
| # | disease | 7 | 0 | 0 | 1 | 1 | # | 20 | Small | paralle | 91 |
| # | sex | 4 | 0 | 0 | 1 | 1 | # | 20 | Small | paralle | 91 |
| # | time | 5 | 0 | 0 | 1 | 1 | # | 20 | Small | paralle | 91 |
| # | interaction | 4 | 0 | 0 | 1 | 1 | # | 20 | Small | paralle | 91 |
| # | disease | 7 | 4 | 0 | 0.636363636 | 1 | # | 20 | Small | ancom | 91 |
| # | sex | 4 | 0 | 0 | 1 | 1 | # | 20 | Small | ancom | 91 |
| # | time | 5 | 3 | 0 | 0.625 | 1 | # | 20 | Small | ancom | 91 |
| # | interaction | 0 | 0 | 4 | 0 | 0 | # | 20 | Small | ancom | 91 |
| # | disease | 7 | 6 | 0 | 0.538461538 | 1 | # | 20 | Small | aldex2 | 91 |
| # | sex | 4 | 0 | 0 | 1 | 1 | # | 20 | Small | aldex2 | 91 |
| # | time | 5 | 5 | 0 | 0.5 | 1 | # | 20 | Small | aldex2 | 91 |
| # | interaction | 0 | 0 | 4 | 0 | 0 | # | 20 | Small | aldex2 | 91 |
| # | disease | 7 | 12 | 0 | 0.368421053 | 1 | # | 20 | Baseline | paralle | 91 |
| # | sex | 4 | 7 | 0 | 0.363636364 | 1 | # | 20 | Baseline | paralle | 91 |
| # | time | 5 | 8 | 0 | 0.384615385 | 1 | # | 20 | Baseline | paralle | 91 |
| # | interaction | 4 | 2 | 0 | 0.666666667 | 1 | # | 20 | Baseline | paralle | 91 |
| # | disease | 7 | 4 | 0 | 0.636363636 | 1 | # | 20 | Baseline | ancom | 91 |
| # | sex | 4 | 0 | 0 | 1 | 1 | # | 20 | Baseline | ancom | 91 |
| # | time | 5 | 4 | 0 | 0.555555556 | 1 | # | 20 | Baseline | ancom | 91 |
| # | interaction | 0 | 0 | 4 | 0 | 0 | # | 20 | Baseline | ancom | 91 |
| # | disease | 7 | 85 | 0 | 0.097222222 | 1 | # | 20 | Baseline | aldex2 | 91 |
| # | sex | 4 | 18 | 0 | 0.181818182 | 1 | # | 20 | Baseline | aldex2 | 91 |
| # | time | 5 | 39 | 0 | 0.113636364 | 1 | # | 20 | Baseline | aldex2 | 91 |
| # | interaction | 0 | 0 | 4 | 0 | 0 | # | 20 | Baseline | aldex2 | 91 |
| # | disease | 7 | 0 | 0 | 1 | 1 | # | 20 | Small | paralle | 91 |
| # | sex | 4 | 0 | 0 | 1 | 1 | # | 20 | Small | paralle | 91 |
| # | time | 5 | 0 | 0 | 1 | 1 | # | 20 | Small | paralle | 91 |
| # | interaction | 4 | 0 | 0 | 1 | 1 | # | 20 | Small | paralle | 91 |
| # | disease | 7 | 4 | 0 | 0.636363636 | 1 | # | 20 | Small | ancom | 91 |
| # | sex | 4 | 0 | 0 | 1 | 1 | # | 20 | Small | ancom | 91 |
| # | time | 5 | 4 | 0 | 0.555555556 | 1 | # | 20 | Small | ancom | 91 |
| # | interaction | 0 | 0 | 4 | 0 | 0 | # | 20 | Small | ancom | 91 |
| # | disease | 7 | 11 | 0 | 0.388888889 | 1 | # | 20 | Small | aldex2 | 91 |
| # | sex | 4 | 0 | 0 | 1 | 1 | # | 20 | Small | aldex2 | 91 |
| # | time | 5 | 6 | 0 | 0.454545455 | 1 | # | 20 | Small | aldex2 | 91 |
| # | interaction | 0 | 0 | 4 | 0 | 0 | # | 20 | Small | aldex2 | 91 |
| # | disease | 4 | 9 | 0 | 0.307692308 | 1 | # | 10 | Baseline | paralle | 92 |
| # | sex | 2 | 7 | 0 | 0.222222222 | 1 | # | 10 | Baseline | paralle | 92 |
| # | time | 2 | 12 | 0 | 0.142857143 | 1 | # | 10 | Baseline | paralle | 92 |
| # | interaction | 1 | 6 | 1 | 0.142857143 | 0.5 | # | 10 | Baseline | paralle | 92 |
| # | disease | 4 | 1 | 0 | 0.8 | 1 | # | 10 | Baseline | ancom | 92 |
| # | sex | 2 | 0 | 0 | 1 | 1 | # | 10 | Baseline | ancom | 92 |
| # | time | 2 | 2 | 0 | 0.5 | 1 | # | 10 | Baseline | ancom | 92 |
| # | interaction | 0 | 0 | 2 | 0 | 0 | # | 10 | Baseline | ancom | 92 |
| # | disease | 4 | 7 | 0 | 0.363636364 | 1 | # | 10 | Baseline | aldex2 | 92 |
| # | sex | 2 | 3 | 0 | 0.4 | 1 | # | 10 | Baseline | aldex2 | 92 |
| # | time | 2 | 11 | 0 | 0.153846154 | 1 | # | 10 | Baseline | aldex2 | 92 |
| # | interaction | 0 | 0 | 2 | 0 | 0 | # | 10 | Baseline | aldex2 | 92 |
| # | disease | 4 | 0 | 0 | 1 | 1 | # | 10 | Small | paralle | 92 |
| # | sex | 2 | 1 | 0 | 0.666666667 | 1 | # | 10 | Small | paralle | 92 |
| # | time | 2 | 0 | 0 | 1 | 1 | # | 10 | Small | paralle | 92 |
| # | interaction | 1 | 0 | 1 | 1 | 0.5 | # | 10 | Small | paralle | 92 |
| # | disease | 4 | 0 | 0 | 1 | 1 | # | 10 | Small | ancom | 92 |
| # | sex | 2 | 0 | 0 | 1 | 1 | # | 10 | Small | ancom | 92 |
| # | time | 2 | 2 | 0 | 0.5 | 1 | # | 10 | Small | ancom | 92 |
| # | interaction | 0 | 0 | 2 | 0 | 0 | # | 10 | Small | ancom | 92 |
| # | disease | 4 | 0 | 0 | 1 | 1 | # | 10 | Small | aldex2 | 92 |
| # | sex | 2 | 0 | 0 | 1 | 1 | # | 10 | Small | aldex2 | 92 |
| # | time | 2 | 2 | 0 | 0.5 | 1 | # | 10 | Small | aldex2 | 92 |
| # | interaction | 0 | 0 | 2 | 0 | 0 | # | 10 | Small | aldex2 | 92 |
| # | disease | 4 | 7 | 0 | 0.363636364 | 1 | # | 10 | Baseline | paralle | 92 |
| # | sex | 2 | 4 | 0 | 0.333333333 | 1 | # | 10 | Baseline | paralle | 92 |
| # | time | 2 | 3 | 0 | 0.4 | 1 | # | 10 | Baseline | paralle | 92 |
| # | interaction | 2 | 4 | 0 | 0.333333333 | 1 | # | 10 | Baseline | paralle | 92 |
| # | disease | 4 | 2 | 0 | 0.666666667 | 1 | # | 10 | Baseline | ancom | 92 |
| # | sex | 2 | 0 | 0 | 1 | 1 | # | 10 | Baseline | ancom | 92 |
| # | time | 2 | 2 | 0 | 0.5 | 1 | # | 10 | Baseline | ancom | 92 |
| # | interaction | 0 | 0 | 2 | 0 | 0 | # | 10 | Baseline | ancom | 92 |
| # | disease | 4 | 14 | 0 | 0.222222222 | 1 | # | 10 | Baseline | aldex2 | 92 |
| # | sex | 2 | 7 | 0 | 0.222222222 | 1 | # | 10 | Baseline | aldex2 | 92 |
| # | time | 2 | 12 | 0 | 0.142857143 | 1 | # | 10 | Baseline | aldex2 | 92 |
| # | interaction | 0 | 0 | 2 | 0 | 0 | # | 10 | Baseline | aldex2 | 92 |
| # | disease | 4 | 2 | 0 | 0.666666667 | 1 | # | 10 | Small | paralle | 92 |
| # | sex | 2 | 0 | 0 | 1 | 1 | # | 10 | Small | paralle | 92 |
| # | time | 2 | 0 | 0 | 1 | 1 | # | 10 | Small | paralle | 92 |
| # | interaction | 2 | 0 | 0 | 1 | 1 | # | 10 | Small | paralle | 92 |
| # | disease | 4 | 2 | 0 | 0.666666667 | 1 | # | 10 | Small | ancom | 92 |
| # | sex | 2 | 0 | 0 | 1 | 1 | # | 10 | Small | ancom | 92 |
| # | time | 2 | 2 | 0 | 0.5 | 1 | # | 10 | Small | ancom | 92 |
| # | interaction | 0 | 0 | 2 | 0 | 0 | # | 10 | Small | ancom | 92 |
| # | disease | 4 | 3 | 0 | 0.571428571 | 1 | # | 10 | Small | aldex2 | 92 |
| # | sex | 2 | 0 | 0 | 1 | 1 | # | 10 | Small | aldex2 | 92 |
| # | time | 2 | 2 | 0 | 0.5 | 1 | # | 10 | Small | aldex2 | 92 |
| # | interaction | 0 | 0 | 2 | 0 | 0 | # | 10 | Small | aldex2 | 92 |
| # | disease | 4 | 4 | 0 | 0.5 | 1 | # | 10 | Baseline | paralle | 92 |

| | | | | | | | | | | | |
|---|---|---|---|---|---|---|---|---|---|---|---|
| # | sex | 2 | 4 | 0 | 0.333333333 | 1 | # | 10 | Baseline | paralle | 92 |
| # | time | 2 | 2 | 0 | 0.5 | 1 | # | 10 | Baseline | paralle | 92 |
| # | interaction | 2 | 6 | 0 | 0.25 | 1 | # | 10 | Baseline | paralle | 92 |
| # | disease | 4 | 2 | 0 | 0.666666667 | 1 | # | 10 | Baseline | ancom | 92 |
| # | sex | 2 | 0 | 0 | 1 | 1 | # | 10 | Baseline | ancom | 92 |
| # | time | 2 | 2 | 0 | 0.5 | 1 | # | 10 | Baseline | ancom | 92 |
| # | interaction | 0 | 0 | 2 | 0 | 0 | # | 10 | Baseline | ancom | 92 |
| # | disease | 4 | 18 | 0 | 0.181818182 | 1 | # | 10 | Baseline | aldex2 | 92 |
| # | sex | 2 | 6 | 0 | 0.25 | 1 | # | 10 | Baseline | aldex2 | 92 |
| # | time | 2 | 16 | 0 | 0.111111111 | 1 | # | 10 | Baseline | aldex2 | 92 |
| # | interaction | 0 | 0 | 2 | 0 | 0 | # | 10 | Baseline | aldex2 | 92 |
| # | disease | 4 | 0 | 0 | 1 | 1 | # | 10 | Small | paralle | 92 |
| # | sex | 2 | 0 | 0 | 1 | 1 | # | 10 | Small | paralle | 92 |
| # | time | 2 | 0 | 0 | 1 | 1 | # | 10 | Small | paralle | 92 |
| # | interaction | 2 | 0 | 0 | 1 | 1 | # | 10 | Small | paralle | 92 |
| # | disease | 4 | 2 | 0 | 0.666666667 | 1 | # | 10 | Small | ancom | 92 |
| # | sex | 2 | 0 | 0 | 1 | 1 | # | 10 | Small | ancom | 92 |
| # | time | 2 | 2 | 0 | 0.5 | 1 | # | 10 | Small | ancom | 92 |
| # | interaction | 0 | 0 | 2 | 0 | 0 | # | 10 | Small | ancom | 92 |
| # | disease | 4 | 2 | 0 | 0.666666667 | 1 | # | 10 | Small | aldex2 | 92 |
| # | sex | 2 | 0 | 0 | 1 | 1 | # | 10 | Small | aldex2 | 92 |
| # | time | 2 | 2 | 0 | 0.5 | 1 | # | 10 | Small | aldex2 | 92 |
| # | interaction | 0 | 0 | 2 | 0 | 0 | # | 10 | Small | aldex2 | 92 |
| # | disease | 4 | 3 | 0 | 0.571428571 | 1 | # | 10 | Baseline | paralle | 92 |
| # | sex | 2 | 4 | 0 | 0.333333333 | 1 | # | 10 | Baseline | paralle | 92 |
| # | time | 2 | 2 | 0 | 0.5 | 1 | # | 10 | Baseline | paralle | 92 |
| # | interaction | 2 | 3 | 0 | 0.4 | 1 | # | 10 | Baseline | paralle | 92 |
| # | disease | 4 | 2 | 0 | 0.666666667 | 1 | # | 10 | Baseline | ancom | 92 |
| # | sex | 2 | 0 | 0 | 1 | 1 | # | 10 | Baseline | ancom | 92 |
| # | time | 2 | 2 | 0 | 0.5 | 1 | # | 10 | Baseline | ancom | 92 |
| # | interaction | 0 | 0 | 2 | 0 | 0 | # | 10 | Baseline | ancom | 92 |
| # | disease | 4 | 20 | 0 | 0.166666667 | 1 | # | 10 | Baseline | aldex2 | 92 |
| # | sex | 2 | 5 | 0 | 0.285714286 | 1 | # | 10 | Baseline | aldex2 | 92 |
| # | time | 2 | 14 | 0 | 0.125 | 1 | # | 10 | Baseline | aldex2 | 92 |
| # | interaction | 0 | 0 | 2 | 0 | 0 | # | 10 | Baseline | aldex2 | 92 |
| # | disease | 4 | 0 | 0 | 1 | 1 | # | 10 | Small | paralle | 92 |
| # | sex | 2 | 0 | 0 | 1 | 1 | # | 10 | Small | paralle | 92 |
| # | time | 2 | 0 | 0 | 1 | 1 | # | 10 | Small | paralle | 92 |
| # | interaction | 2 | 0 | 0 | 1 | 1 | # | 10 | Small | paralle | 92 |
| # | disease | 4 | 2 | 0 | 0.666666667 | 1 | # | 10 | Small | ancom | 92 |
| # | sex | 2 | 0 | 0 | 1 | 1 | # | 10 | Small | ancom | 92 |
| # | time | 2 | 2 | 0 | 0.5 | 1 | # | 10 | Small | ancom | 92 |
| # | interaction | 0 | 0 | 2 | 0 | 0 | # | 10 | Small | ancom | 92 |
| # | disease | 4 | 2 | 0 | 0.666666667 | 1 | # | 10 | Small | aldex2 | 92 |
| # | sex | 2 | 0 | 0 | 1 | 1 | # | 10 | Small | aldex2 | 92 |
| # | time | 2 | 2 | 0 | 0.5 | 1 | # | 10 | Small | aldex2 | 92 |
| # | interaction | 0 | 0 | 2 | 0 | 0 | # | 10 | Small | aldex2 | 92 |
| # | disease | 7 | 10 | 0 | 0.411764706 | 1 | # | 20 | Baseline | paralle | 92 |
| # | sex | 4 | 7 | 0 | 0.363636364 | 1 | # | 20 | Baseline | paralle | 92 |
| # | time | 5 | 9 | 0 | 0.357142857 | 1 | # | 20 | Baseline | paralle | 92 |
| # | interaction | 4 | 7 | 0 | 0.363636364 | 1 | # | 20 | Baseline | paralle | 92 |
| # | disease | 7 | 3 | 0 | 0.7 | 1 | # | 20 | Baseline | ancom | 92 |
| # | sex | 4 | 0 | 0 | 1 | 1 | # | 20 | Baseline | ancom | 92 |
| # | time | 5 | 2 | 0 | 0.714285714 | 1 | # | 20 | Baseline | ancom | 92 |
| # | interaction | 0 | 0 | 4 | 0 | 0 | # | 20 | Baseline | ancom | 92 |
| # | disease | 7 | 19 | 0 | 0.269230769 | 1 | # | 20 | Baseline | aldex2 | 92 |
| # | sex | 4 | 10 | 0 | 0.285714286 | 1 | # | 20 | Baseline | aldex2 | 92 |
| # | time | 5 | 17 | 0 | 0.227272727 | 1 | # | 20 | Baseline | aldex2 | 92 |
| # | interaction | 0 | 0 | 4 | 0 | 0 | # | 20 | Baseline | aldex2 | 92 |
| # | disease | 7 | 0 | 0 | 1 | 1 | # | 20 | Small | paralle | 92 |
| # | sex | 4 | 1 | 0 | 0.8 | 1 | # | 20 | Small | paralle | 92 |
| # | time | 5 | 0 | 0 | 1 | 1 | # | 20 | Small | paralle | 92 |
| # | interaction | 1 | 0 | 3 | 1 | 0.25 | # | 20 | Small | paralle | 92 |
| # | disease | 7 | 2 | 0 | 0.777777778 | 1 | # | 20 | Small | ancom | 92 |
| # | sex | 4 | 0 | 0 | 1 | 1 | # | 20 | Small | ancom | 92 |
| # | time | 5 | 2 | 0 | 0.714285714 | 1 | # | 20 | Small | ancom | 92 |
| # | interaction | 0 | 0 | 4 | 0 | 0 | # | 20 | Small | ancom | 92 |
| # | disease | 7 | 3 | 0 | 0.7 | 1 | # | 20 | Small | aldex2 | 92 |
| # | sex | 4 | 1 | 0 | 0.8 | 1 | # | 20 | Small | aldex2 | 92 |
| # | time | 5 | 0 | 0 | 1 | 1 | # | 20 | Small | aldex2 | 92 |
| # | interaction | 0 | 0 | 4 | 0 | 0 | # | 20 | Small | aldex2 | 92 |
| # | disease | 7 | 7 | 0 | 0.5 | 1 | # | 20 | Baseline | paralle | 92 |
| # | sex | 4 | 5 | 0 | 0.444444444 | 1 | # | 20 | Baseline | paralle | 92 |
| # | time | 5 | 6 | 0 | 0.454545455 | 1 | # | 20 | Baseline | paralle | 92 |
| # | interaction | 4 | 10 | 0 | 0.285714286 | 1 | # | 20 | Baseline | paralle | 92 |
| # | disease | 7 | 4 | 0 | 0.636363636 | 1 | # | 20 | Baseline | ancom | 92 |
| # | sex | 4 | 0 | 0 | 1 | 1 | # | 20 | Baseline | ancom | 92 |
| # | time | 5 | 4 | 0 | 0.555555556 | 1 | # | 20 | Baseline | ancom | 92 |
| # | interaction | 0 | 0 | 4 | 0 | 0 | # | 20 | Baseline | ancom | 92 |
| # | disease | 7 | 34 | 0 | 0.170731707 | 1 | # | 20 | Baseline | aldex2 | 92 |
| # | sex | 4 | 14 | 0 | 0.222222222 | 1 | # | 20 | Baseline | aldex2 | 92 |
| # | time | 5 | 27 | 0 | 0.15625 | 1 | # | 20 | Baseline | aldex2 | 92 |
| # | interaction | 0 | 0 | 4 | 0 | 0 | # | 20 | Baseline | aldex2 | 92 |
| # | disease | 7 | 0 | 0 | 1 | 1 | # | 20 | Small | paralle | 92 |
| # | sex | 4 | 0 | 0 | 1 | 1 | # | 20 | Small | paralle | 92 |
| # | time | 5 | 0 | 0 | 1 | 1 | # | 20 | Small | paralle | 92 |
| # | interaction | 4 | 0 | 0 | 1 | 1 | # | 20 | Small | paralle | 92 |
| # | disease | 7 | 4 | 0 | 0.636363636 | 1 | # | 20 | Small | ancom | 92 |
| # | sex | 4 | 0 | 0 | 1 | 1 | # | 20 | Small | ancom | 92 |
| # | time | 5 | 4 | 0 | 0.555555556 | 1 | # | 20 | Small | ancom | 92 |
| # | interaction | 0 | 0 | 4 | 0 | 0 | # | 20 | Small | ancom | 92 |
| # | disease | 7 | 2 | 0 | 0.777777778 | 1 | # | 20 | Small | aldex2 | 92 |
| # | sex | 4 | 0 | 0 | 1 | 1 | # | 20 | Small | aldex2 | 92 |
| # | time | 5 | 5 | 0 | 0.5 | 1 | # | 20 | Small | aldex2 | 92 |
| # | interaction | 0 | 0 | 4 | 0 | 0 | # | 20 | Small | aldex2 | 92 |
| # | disease | 7 | 6 | 0 | 0.538461538 | 1 | # | 20 | Baseline | paralle | 92 |
| # | sex | 4 | 3 | 0 | 0.571428571 | 1 | # | 20 | Baseline | paralle | 92 |
| # | time | 5 | 4 | 0 | 0.555555556 | 1 | # | 20 | Baseline | paralle | 92 |
| # | interaction | 4 | 4 | 0 | 0.5 | 1 | # | 20 | Baseline | paralle | 92 |
| # | disease | 7 | 4 | 0 | 0.636363636 | 1 | # | 20 | Baseline | ancom | 92 |
| # | sex | 4 | 0 | 0 | 1 | 1 | # | 20 | Baseline | ancom | 92 |
| # | time | 5 | 4 | 0 | 0.555555556 | 1 | # | 20 | Baseline | ancom | 92 |
| # | interaction | 0 | 0 | 4 | 0 | 0 | # | 20 | Baseline | ancom | 92 |
| # | disease | 7 | 51 | 0 | 0.120689655 | 1 | # | 20 | Baseline | aldex2 | 92 |
| # | sex | 4 | 18 | 0 | 0.181818182 | 1 | # | 20 | Baseline | aldex2 | 92 |
| # | time | 5 | 28 | 0 | 0.151515152 | 1 | # | 20 | Baseline | aldex2 | 92 |
| # | interaction | 0 | 0 | 4 | 0 | 0 | # | 20 | Baseline | aldex2 | 92 |
| # | disease | 7 | 1 | 0 | 0.875 | 1 | # | 20 | Small | paralle | 92 |
| # | sex | 4 | 0 | 0 | 1 | 1 | # | 20 | Small | paralle | 92 |
| # | time | 5 | 0 | 0 | 1 | 1 | # | 20 | Small | paralle | 92 |
| # | interaction | 4 | 0 | 0 | 1 | 1 | # | 20 | Small | paralle | 92 |
| # | disease | 7 | 4 | 0 | 0.636363636 | 1 | # | 20 | Small | ancom | 92 |
| # | sex | 4 | 0 | 0 | 1 | 1 | # | 20 | Small | ancom | 92 |
| # | time | 5 | 4 | 0 | 0.555555556 | 1 | # | 20 | Small | ancom | 92 |
| # | interaction | 0 | 0 | 4 | 0 | 0 | # | 20 | Small | ancom | 92 |
| # | disease | 7 | 6 | 0 | 0.538461538 | 1 | # | 20 | Small | aldex2 | 92 |
| # | sex | 4 | 0 | 0 | 1 | 1 | # | 20 | Small | aldex2 | 92 |
| # | time | 5 | 9 | 0 | 0.357142857 | 1 | # | 20 | Small | aldex2 | 92 |
| # | interaction | 0 | 0 | 4 | 0 | 0 | # | 20 | Small | aldex2 | 92 |
| # | disease | 7 | 7 | 0 | 0.5 | 1 | # | 20 | Baseline | paralle | 92 |
| # | sex | 4 | 6 | 0 | 0.4 | 1 | # | 20 | Baseline | paralle | 92 |
| # | time | 5 | 4 | 0 | 0.555555556 | 1 | # | 20 | Baseline | paralle | 92 |
| # | interaction | 4 | 6 | 0 | 0.4 | 1 | # | 20 | Baseline | paralle | 92 |
| # | disease | 7 | 4 | 0 | 0.636363636 | 1 | # | 20 | Baseline | ancom | 92 |
| # | sex | 4 | 0 | 0 | 1 | 1 | # | 20 | Baseline | ancom | 92 |
| # | time | 5 | 5 | 0 | 0.5 | 1 | # | 20 | Baseline | ancom | 92 |
| # | interaction | 0 | 0 | 4 | 0 | 0 | # | 20 | Baseline | ancom | 92 |
| # | disease | 7 | 85 | 0 | 0.097222222 | 1 | # | 20 | Baseline | aldex2 | 92 |
| # | sex | 4 | 20 | 0 | 0.166666667 | 1 | # | 20 | Baseline | aldex2 | 92 |
| # | time | 5 | 35 | 0 | 0.125 | 1 | # | 20 | Baseline | aldex2 | 92 |
| # | interaction | 0 | 0 | 4 | 0 | 0 | # | 20 | Baseline | aldex2 | 92 |
| # | disease | 7 | 0 | 0 | 1 | 1 | # | 20 | Small | paralle | 92 |
| # | sex | 4 | 0 | 0 | 1 | 1 | # | 20 | Small | paralle | 92 |
| # | time | 5 | 0 | 0 | 1 | 1 | # | 20 | Small | paralle | 92 |
| # | interaction | 4 | 0 | 0 | 1 | 1 | # | 20 | Small | paralle | 92 |
| # | disease | 7 | 4 | 0 | 0.636363636 | 1 | # | 20 | Small | ancom | 92 |
| # | sex | 4 | 0 | 0 | 1 | 1 | # | 20 | Small | ancom | 92 |
| # | time | 5 | 4 | 0 | 0.555555556 | 1 | # | 20 | Small | ancom | 92 |
| # | interaction | 0 | 0 | 4 | 0 | 0 | # | 20 | Small | ancom | 92 |
| # | disease | 7 | 8 | 0 | 0.466666667 | 1 | # | 20 | Small | aldex2 | 92 |
| # | sex | 4 | 1 | 0 | 0.8 | 1 | # | 20 | Small | aldex2 | 92 |
| # | time | 5 | 7 | 0 | 0.416666667 | 1 | # | 20 | Small | aldex2 | 92 |
| # | interaction | 0 | 0 | 4 | 0 | 0 | # | 20 | Small | aldex2 | 92 |
| # | disease | 4 | 8 | 0 | 0.333333333 | 1 | # | 10 | Baseline | paralle | 93 |
| # | sex | 2 | 8 | 0 | 0.2 | 1 | # | 10 | Baseline | paralle | 93 |
| # | time | 2 | 11 | 0 | 0.153846154 | 1 | # | 10 | Baseline | paralle | 93 |
| # | interaction | 2 | 7 | 0 | 0.222222222 | 1 | # | 10 | Baseline | paralle | 93 |
| # | disease | 4 | 0 | 0 | 1 | 1 | # | 10 | Baseline | ancom | 93 |
| # | sex | 2 | 0 | 0 | 1 | 1 | # | 10 | Baseline | ancom | 93 |
| # | time | 2 | 0 | 0 | 1 | 1 | # | 10 | Baseline | ancom | 93 |
| # | interaction | 0 | 0 | 2 | 0 | 0 | # | 10 | Baseline | ancom | 93 |
| # | disease | 4 | 7 | 0 | 0.363636364 | 1 | # | 10 | Baseline | aldex2 | 93 |
| # | sex | 2 | 3 | 0 | 0.4 | 1 | # | 10 | Baseline | aldex2 | 93 |
| # | time | 2 | 3 | 0 | 0.4 | 1 | # | 10 | Baseline | aldex2 | 93 |
| # | interaction | 0 | 0 | 2 | 0 | 0 | # | 10 | Baseline | aldex2 | 93 |
| # | disease | 4 | 0 | 0 | 1 | 1 | # | 10 | Small | paralle | 93 |
| # | sex | 2 | 0 | 0 | 1 | 1 | # | 10 | Small | paralle | 93 |
| # | time | 2 | 0 | 0 | 1 | 1 | # | 10 | Small | paralle | 93 |
| # | interaction | 1 | 0 | 1 | 1 | 0.5 | # | 10 | Small | paralle | 93 |
| # | disease | 4 | 0 | 0 | 1 | 1 | # | 10 | Small | ancom | 93 |
| # | sex | 2 | 0 | 0 | 1 | 1 | # | 10 | Small | ancom | 93 |
| # | time | 2 | 0 | 0 | 1 | 1 | # | 10 | Small | ancom | 93 |
| # | interaction | 0 | 0 | 2 | 0 | 0 | # | 10 | Small | ancom | 93 |
| # | disease | 4 | 1 | 0 | 0.8 | 1 | # | 10 | Small | aldex2 | 93 |
| # | sex | 2 | 0 | 0 | 1 | 1 | # | 10 | Small | aldex2 | 93 |
| # | time | 2 | 0 | 0 | 1 | 1 | # | 10 | Small | aldex2 | 93 |
| # | interaction | 0 | 0 | 2 | 0 | 0 | # | 10 | Small | aldex2 | 93 |
| # | disease | 4 | 8 | 0 | 0.333333333 | 1 | # | 10 | Baseline | paralle | 93 |
| # | sex | 2 | 4 | 0 | 0.333333333 | 1 | # | 10 | Baseline | paralle | 93 |
| # | time | 2 | 3 | 0 | 0.4 | 1 | # | 10 | Baseline | paralle | 93 |
| # | interaction | 2 | 7 | 0 | 0.222222222 | 1 | # | 10 | Baseline | paralle | 93 |
| # | disease | 4 | 2 | 0 | 0.666666667 | 1 | # | 10 | Baseline | ancom | 93 |
| # | sex | 2 | 0 | 0 | 1 | 1 | # | 10 | Baseline | ancom | 93 |
| # | time | 2 | 2 | 0 | 0.5 | 1 | # | 10 | Baseline | ancom | 93 |
| # | interaction | 0 | 0 | 2 | 0 | 0 | # | 10 | Baseline | ancom | 93 |
| # | disease | 4 | 16 | 0 | 0.2 | 1 | # | 10 | Baseline | aldex2 | 93 |
| # | sex | 2 | 2 | 0 | 0.5 | 1 | # | 10 | Baseline | aldex2 | 93 |
| # | time | 2 | 5 | 0 | 0.285714286 | 1 | # | 10 | Baseline | aldex2 | 93 |
| # | interaction | 0 | 0 | 2 | 0 | 0 | # | 10 | Baseline | aldex2 | 93 |
| # | disease | 4 | 0 | 0 | 1 | 1 | # | 10 | Small | paralle | 93 |
| # | sex | 2 | 0 | 0 | 1 | 1 | # | 10 | Small | paralle | 93 |
| # | time | 2 | 0 | 0 | 1 | 1 | # | 10 | Small | paralle | 93 |
| # | interaction | 2 | 0 | 0 | 1 | 1 | # | 10 | Small | paralle | 93 |
| # | disease | 4 | 2 | 0 | 0.666666667 | 1 | # | 10 | Small | ancom | 93 |
| # | sex | 2 | 0 | 0 | 1 | 1 | # | 10 | Small | ancom | 93 |
| # | time | 2 | 1 | 0 | 0.666666667 | 1 | # | 10 | Small | ancom | 93 |
| # | interaction | 0 | 0 | 2 | 0 | 0 | # | 10 | Small | ancom | 93 |
| # | disease | 4 | 3 | 0 | 0.571428571 | 1 | # | 10 | Small | aldex2 | 93 |
| # | sex | 2 | 0 | 0 | 1 | 1 | # | 10 | Small | aldex2 | 93 |
| # | time | 2 | 1 | 0 | 0.666666667 | 1 | # | 10 | Small | aldex2 | 93 |
| # | interaction | 0 | 0 | 2 | 0 | 0 | # | 10 | Small | aldex2 | 93 |
| # | disease | 4 | 5 | 0 | 0.444444444 | 1 | # | 10 | Baseline | paralle | 93 |
| # | sex | 2 | 4 | 0 | 0.333333333 | 1 | # | 10 | Baseline | paralle | 93 |
| # | time | 2 | 7 | 0 | 0.222222222 | 1 | # | 10 | Baseline | paralle | 93 |
| # | interaction | 2 | 7 | 0 | 0.222222222 | 1 | # | 10 | Baseline | paralle | 93 |
| # | disease | 4 | 2 | 0 | 0.666666667 | 1 | # | 10 | Baseline | ancom | 93 |
| # | sex | 2 | 0 | 0 | 1 | 1 | # | 10 | Baseline | ancom | 93 |
| # | time | 2 | 2 | 0 | 0.5 | 1 | # | 10 | Baseline | ancom | 93 |
| # | interaction | 0 | 0 | 2 | 0 | 0 | # | 10 | Baseline | ancom | 93 |
| # | disease | 4 | 22 | 0 | 0.153846154 | 1 | # | 10 | Baseline | aldex2 | 93 |
| # | sex | 2 | 10 | 0 | 0.166666667 | 1 | # | 10 | Baseline | aldex2 | 93 |
| # | time | 2 | 8 | 0 | 0.2 | 1 | # | 10 | Baseline | aldex2 | 93 |
| # | interaction | 0 | 0 | 2 | 0 | 0 | # | 10 | Baseline | aldex2 | 93 |
| # | disease | 4 | 0 | 0 | 1 | 1 | # | 10 | Small | paralle | 93 |
| # | sex | 2 | 0 | 0 | 1 | 1 | # | 10 | Small | paralle | 93 |
| # | time | 2 | 0 | 0 | 1 | 1 | # | 10 | Small | paralle | 93 |
| # | interaction | 2 | 0 | 0 | 1 | 1 | # | 10 | Small | paralle | 93 |
| # | disease | 4 | 2 | 0 | 0.666666667 | 1 | # | 10 | Small | ancom | 93 |
| # | sex | 2 | 0 | 0 | 1 | 1 | # | 10 | Small | ancom | 93 |
| # | time | 2 | 2 | 0 | 0.5 | 1 | # | 10 | Small | ancom | 93 |
| # | interaction | 0 | 0 | 2 | 0 | 0 | # | 10 | Small | ancom | 93 |
| # | disease | 4 | 3 | 0 | 0.571428571 | 1 | # | 10 | Small | aldex2 | 93 |
| # | sex | 2 | 0 | 0 | 1 | 1 | # | 10 | Small | aldex2 | 93 |
| # | time | 2 | 2 | 0 | 0.5 | 1 | # | 10 | Small | aldex2 | 93 |
| # | interaction | 0 | 0 | 2 | 0 | 0 | # | 10 | Small | aldex2 | 93 |
| # | disease | 4 | 5 | 0 | 0.444444444 | 1 | # | 10 | Baseline | paralle | 93 |
| # | sex | 2 | 3 | 0 | 0.4 | 1 | # | 10 | Baseline | paralle | 93 |
| # | time | 2 | 5 | 0 | 0.285714286 | 1 | # | 10 | Baseline | paralle | 93 |
| # | interaction | 2 | 8 | 0 | 0.2 | 1 | # | 10 | Baseline | paralle | 93 |
| # | disease | 4 | 2 | 0 | 0.666666667 | 1 | # | 10 | Baseline | ancom | 93 |
| # | sex | 2 | 0 | 0 | 1 | 1 | # | 10 | Baseline | ancom | 93 |
| # | time | 2 | 2 | 0 | 0.5 | 1 | # | 10 | Baseline | ancom | 93 |
| # | interaction | 0 | 0 | 2 | 0 | 0 | # | 10 | Baseline | ancom | 93 |
| # | disease | 4 | 22 | 0 | 0.153846154 | 1 | # | 10 | Baseline | aldex2 | 93 |
| # | sex | 2 | 9 | 0 | 0.181818182 | 1 | # | 10 | Baseline | aldex2 | 93 |
| # | time | 2 | 11 | 0 | 0.153846154 | 1 | # | 10 | Baseline | aldex2 | 93 |
| # | interaction | 0 | 0 | 2 | 0 | 0 | # | 10 | Baseline | aldex2 | 93 |
| # | disease | 4 | 0 | 0 | 1 | 1 | # | 10 | Small | paralle | 93 |
| # | sex | 2 | 0 | 0 | 1 | 1 | # | 10 | Small | paralle | 93 |
| # | time | 2 | 0 | 0 | 1 | 1 | # | 10 | Small | paralle | 93 |
| # | interaction | 2 | 0 | 0 | 1 | 1 | # | 10 | Small | paralle | 93 |
| # | disease | 4 | 2 | 0 | 0.666666667 | 1 | # | 10 | Small | ancom | 93 |
| # | sex | 2 | 0 | 0 | 1 | 1 | # | 10 | Small | ancom | 93 |
| # | time | 2 | 2 | 0 | 0.5 | 1 | # | 10 | Small | ancom | 93 |
| # | interaction | 0 | 0 | 2 | 0 | 0 | # | 10 | Small | ancom | 93 |
| # | disease | 4 | 3 | 0 | 0.571428571 | 1 | # | 10 | Small | aldex2 | 93 |
| # | sex | 2 | 0 | 0 | 1 | 1 | # | 10 | Small | aldex2 | 93 |
| # | time | 2 | 2 | 0 | 0.5 | 1 | # | 10 | Small | aldex2 | 93 |
| # | interaction | 0 | 0 | 2 | 0 | 0 | # | 10 | Small | aldex2 | 93 |
| # | disease | 7 | 8 | 0 | 0.466666667 | 1 | # | 20 | Baseline | paralle | 93 |
| # | sex | 4 | 8 | 0 | 0.333333333 | 1 | # | 20 | Baseline | paralle | 93 |
| # | time | 5 | 12 | 0 | 0.294117647 | 1 | # | 20 | Baseline | paralle | 93 |
| # | interaction | 4 | 9 | 0 | 0.307692308 | 1 | # | 20 | Baseline | paralle | 93 |
| # | disease | 7 | 0 | 0 | 1 | 1 | # | 20 | Baseline | ancom | 93 |
| # | sex | 4 | 0 | 0 | 1 | 1 | # | 20 | Baseline | ancom | 93 |
| # | time | 5 | 2 | 0 | 0.714285714 | 1 | # | 20 | Baseline | ancom | 93 |
| # | interaction | 0 | 0 | 4 | 0 | 0 | # | 20 | Baseline | ancom | 93 |
| # | disease | 7 | 13 | 0 | 0.35 | 1 | # | 20 | Baseline | aldex2 | 93 |
| # | sex | 4 | 5 | 0 | 0.444444444 | 1 | # | 20 | Baseline | aldex2 | 93 |
| # | time | 5 | 17 | 0 | 0.227272727 | 1 | # | 20 | Baseline | aldex2 | 93 |
| # | interaction | 0 | 0 | 4 | 0 | 0 | # | 20 | Baseline | aldex2 | 93 |
| # | disease | 7 | 0 | 0 | 1 | 1 | # | 20 | Small | paralle | 93 |
| # | sex | 4 | 0 | 0 | 1 | 1 | # | 20 | Small | paralle | 93 |
| # | time | 5 | 0 | 0 | 1 | 1 | # | 20 | Small | paralle | 93 |
| # | interaction | 2 | 0 | 2 | 1 | 0.5 | # | 20 | Small | paralle | 93 |
| # | disease | 7 | 0 | 0 | 1 | 1 | # | 20 | Small | ancom | 93 |
| # | sex | 4 | 0 | 0 | 1 | 1 | # | 20 | Small | ancom | 93 |
| # | time | 5 | 0 | 0 | 1 | 1 | # | 20 | Small | ancom | 93 |
| # | interaction | 0 | 0 | 4 | 0 | 0 | # | 20 | Small | ancom | 93 |
| # | disease | 7 | 0 | 0 | 1 | 1 | # | 20 | Small | aldex2 | 93 |
| # | sex | 4 | 0 | 0 | 1 | 1 | # | 20 | Small | aldex2 | 93 |
| # | time | 5 | 1 | 0 | 0.833333333 | 1 | # | 20 | Small | aldex2 | 93 |
| # | interaction | 0 | 0 | 4 | 0 | 0 | # | 20 | Small | aldex2 | 93 |
| # | disease | 7 | 9 | 0 | 0.4375 | 1 | # | 20 | Baseline | paralle | 93 |
| # | sex | 4 | 8 | 0 | 0.333333333 | 1 | # | 20 | Baseline | paralle | 93 |
| # | time | 5 | 9 | 0 | 0.357142857 | 1 | # | 20 | Baseline | paralle | 93 |
| # | interaction | 4 | 6 | 0 | 0.4 | 1 | # | 20 | Baseline | paralle | 93 |
| # | disease | 7 | 3 | 0 | 0.7 | 1 | # | 20 | Baseline | ancom | 93 |
| # | sex | 4 | 0 | 0 | 1 | 1 | # | 20 | Baseline | ancom | 93 |
| # | time | 5 | 4 | 0 | 0.555555556 | 1 | # | 20 | Baseline | ancom | 93 |
| # | interaction | 0 | 0 | 4 | 0 | 0 | # | 20 | Baseline | ancom | 93 |
| # | disease | 7 | 28 | 0 | 0.2 | 1 | # | 20 | Baseline | aldex2 | 93 |
| # | sex | 4 | 12 | 0 | 0.25 | 1 | # | 20 | Baseline | aldex2 | 93 |
| # | time | 5 | 28 | 0 | 0.151515152 | 1 | # | 20 | Baseline | aldex2 | 93 |
| # | interaction | 0 | 0 | 4 | 0 | 0 | # | 20 | Baseline | aldex2 | 93 |
| # | disease | 7 | 0 | 0 | 1 | 1 | # | 20 | Small | paralle | 93 |
| # | sex | 4 | 0 | 0 | 1 | 1 | # | 20 | Small | paralle | 93 |
| # | time | 5 | 0 | 0 | 1 | 1 | # | 20 | Small | paralle | 93 |
| # | interaction | 4 | 0 | 0 | 1 | 1 | # | 20 | Small | paralle | 93 |
| # | disease | 7 | 3 | 0 | 0.7 | 1 | # | 20 | Small | ancom | 93 |
| # | sex | 4 | 0 | 0 | 1 | 1 | # | 20 | Small | ancom | 93 |
| # | time | 5 | 4 | 0 | 0.555555556 | 1 | # | 20 | Small | ancom | 93 |
| # | interaction | 0 | 0 | 4 | 0 | 0 | # | 20 | Small | ancom | 93 |
| # | disease | 7 | 3 | 0 | 0.7 | 1 | # | 20 | Small | aldex2 | 93 |
| # | sex | 4 | 0 | 0 | 1 | 1 | # | 20 | Small | aldex2 | 93 |
| # | time | 5 | 3 | 0 | 0.625 | 1 | # | 20 | Small | aldex2 | 93 |
| # | interaction | 0 | 0 | 4 | 0 | 0 | # | 20 | Small | aldex2 | 93 |
| # | disease | 7 | 10 | 0 | 0.411764706 | 1 | # | 20 | Baseline | paralle | 93 |
| # | sex | 4 | 6 | 0 | 0.4 | 1 | # | 20 | Baseline | paralle | 93 |
| # | time | 5 | 10 | 0 | 0.333333333 | 1 | # | 20 | Baseline | paralle | 93 |
| # | interaction | 4 | 7 | 0 | 0.363636364 | 1 | # | 20 | Baseline | paralle | 93 |
| # | disease | 7 | 4 | 0 | 0.636363636 | 1 | # | 20 | Baseline | ancom | 93 |
| # | sex | 4 | 0 | 0 | 1 | 1 | # | 20 | Baseline | ancom | 93 |
| # | time | 5 | 4 | 0 | 0.555555556 | 1 | # | 20 | Baseline | ancom | 93 |
| # | interaction | 0 | 0 | 4 | 0 | 0 | # | 20 | Baseline | ancom | 93 |
| # | disease | 7 | 64 | 0 | 0.137254902 | 1 | # | 20 | Baseline | aldex2 | 93 |
| # | sex | 4 | 17 | 0 | 0.19047619 | 1 | # | 20 | Baseline | aldex2 | 93 |
| # | time | 5 | 33 | 0 | 0.131578947 | 1 | # | 20 | Baseline | aldex2 | 93 |
| # | interaction | 0 | 0 | 4 | 0 | 0 | # | 20 | Baseline | aldex2 | 93 |
| # | disease | 7 | 0 | 0 | 1 | 1 | # | 20 | Small | paralle | 93 |
| # | sex | 4 | 0 | 0 | 1 | 1 | # | 20 | Small | paralle | 93 |
| # | time | 5 | 0 | 0 | 1 | 1 | # | 20 | Small | paralle | 93 |
| # | interaction | 4 | 0 | 0 | 1 | 1 | # | 20 | Small | paralle | 93 |
| # | disease | 7 | 4 | 0 | 0.636363636 | 1 | # | 20 | Small | ancom | 93 |
| # | sex | 4 | 0 | 0 | 1 | 1 | # | 20 | Small | ancom | 93 |
| # | time | 5 | 4 | 0 | 0.555555556 | 1 | # | 20 | Small | ancom | 93 |
| # | interaction | 0 | 0 | 4 | 0 | 0 | # | 20 | Small | ancom | 93 |
| # | disease | 7 | 5 | 0 | 0.583333333 | 1 | # | 20 | Small | aldex2 | 93 |
| # | sex | 4 | 0 | 0 | 1 | 1 | # | 20 | Small | aldex2 | 93 |
| # | time | 5 | 4 | 0 | 0.555555556 | 1 | # | 20 | Small | aldex2 | 93 |
| # | interaction | 0 | 0 | 4 | 0 | 0 | # | 20 | Small | aldex2 | 93 |
| # | disease | 7 | 12 | 0 | 0.368421053 | 1 | # | 20 | Baseline | paralle | 93 |
| # | sex | 4 | 7 | 0 | 0.363636364 | 1 | # | 20 | Baseline | paralle | 93 |
| # | time | 5 | 4 | 0 | 0.555555556 | 1 | # | 20 | Baseline | paralle | 93 |
| # | interaction | 4 | 6 | 0 | 0.4 | 1 | # | 20 | Baseline | paralle | 93 |
| # | disease | 7 | 4 | 0 | 0.636363636 | 1 | # | 20 | Baseline | ancom | 93 |
| # | sex | 4 | 0 | 0 | 1 | 1 | # | 20 | Baseline | ancom | 93 |
| # | time | 5 | 4 | 0 | 0.555555556 | 1 | # | 20 | Baseline | ancom | 93 |
| # | interaction | 0 | 0 | 4 | 0 | 0 | # | 20 | Baseline | ancom | 93 |
| # | disease | 7 | 55 | 0 | 0.112903226 | 1 | # | 20 | Baseline | aldex2 | 93 |
| # | sex | 4 | 16 | 0 | 0.2 | 1 | # | 20 | Baseline | aldex2 | 93 |
| # | time | 5 | 41 | 0 | 0.108695652 | 1 | # | 20 | Baseline | aldex2 | 93 |
| # | interaction | 0 | 0 | 4 | 0 | 0 | # | 20 | Baseline | aldex2 | 93 |
| # | disease | 7 | 0 | 0 | 1 | 1 | # | 20 | Small | paralle | 93 |
| # | sex | 4 | 0 | 0 | 1 | 1 | # | 20 | Small | paralle | 93 |
| # | time | 5 | 0 | 0 | 1 | 1 | # | 20 | Small | paralle | 93 |
| # | interaction | 4 | 0 | 0 | 1 | 1 | # | 20 | Small | paralle | 93 |
| # | disease | 7 | 4 | 0 | 0.636363636 | 1 | # | 20 | Small | ancom | 93 |
| # | sex | 4 | 0 | 0 | 1 | 1 | # | 20 | Small | ancom | 93 |
| # | time | 5 | 4 | 0 | 0.555555556 | 1 | # | 20 | Small | ancom | 93 |
| # | interaction | 0 | 0 | 4 | 0 | 0 | # | 20 | Small | ancom | 93 |
| # | disease | 7 | 8 | 0 | 0.466666667 | 1 | # | 20 | Small | aldex2 | 93 |
| # | sex | 4 | 1 | 0 | 0.8 | 1 | # | 20 | Small | aldex2 | 93 |
| # | time | 5 | 5 | 0 | 0.5 | 1 | # | 20 | Small | aldex2 | 93 |
| # | interaction | 0 | 0 | 4 | 0 | 0 | # | 20 | Small | aldex2 | 93 |
| # | disease | 4 | 5 | 0 | 0.444444444 | 1 | # | 10 | Baseline | paralle | 94 |
| # | sex | 2 | 7 | 0 | 0.222222222 | 1 | # | 10 | Baseline | paralle | 94 |

| | | | | | | | | | | | |
|---|---|---|---|---|---|---|---|---|---|---|---|
| # | time | 2 | 7 | 0 | 0.222222222 | 1 | # | 10 | Baseline | paralle | 94 |
| # | interaction | 2 | 8 | 0 | 0.2 | 1 | # | 10 | Baseline | paralle | 94 |
| # | disease | 4 | 1 | 0 | 0.8 | 1 | # | 10 | Baseline | ancom | 94 |
| # | sex | 2 | 0 | 0 | 1 | 1 | # | 10 | Baseline | ancom | 94 |
| # | time | 2 | 0 | 0 | 1 | 1 | # | 10 | Baseline | ancom | 94 |
| # | interaction | 0 | 0 | 2 | 0 | 0 | # | 10 | Baseline | ancom | 94 |
| # | disease | 4 | 7 | 0 | 0.363636364 | 1 | # | 10 | Baseline | aldex2 | 94 |
| # | sex | 2 | 5 | 0 | 0.285714286 | 1 | # | 10 | Baseline | aldex2 | 94 |
| # | time | 2 | 7 | 0 | 0.222222222 | 1 | # | 10 | Baseline | aldex2 | 94 |
| # | interaction | 0 | 0 | 2 | 0 | 0 | # | 10 | Baseline | aldex2 | 94 |
| # | disease | 4 | 0 | 0 | 1 | 1 | # | 10 | Small | paralle | 94 |
| # | sex | 2 | 0 | 0 | 1 | 1 | # | 10 | Small | paralle | 94 |
| # | time | 2 | 0 | 0 | 1 | 1 | # | 10 | Small | paralle | 94 |
| # | interaction | 2 | 0 | 0 | 1 | 1 | # | 10 | Small | paralle | 94 |
| # | disease | 4 | 0 | 0 | 1 | 1 | # | 10 | Small | ancom | 94 |
| # | sex | 2 | 0 | 0 | 1 | 1 | # | 10 | Small | ancom | 94 |
| # | time | 2 | 1 | 0 | 0.666666667 | 1 | # | 10 | Small | ancom | 94 |
| # | interaction | 0 | 0 | 2 | 0 | 0 | # | 10 | Small | ancom | 94 |
| # | disease | 4 | 2 | 0 | 0.666666667 | 1 | # | 10 | Small | aldex2 | 94 |
| # | sex | 2 | 1 | 0 | 0.666666667 | 1 | # | 10 | Small | aldex2 | 94 |
| # | time | 2 | 1 | 0 | 0.666666667 | 1 | # | 10 | Small | aldex2 | 94 |
| # | interaction | 0 | 0 | 2 | 0 | 0 | # | 10 | Small | aldex2 | 94 |
| # | disease | 4 | 5 | 0 | 0.444444444 | 1 | # | 10 | Baseline | paralle | 94 |
| # | sex | 2 | 6 | 0 | 0.25 | 1 | # | 10 | Baseline | paralle | 94 |
| # | time | 2 | 11 | 0 | 0.153846154 | 1 | # | 10 | Baseline | paralle | 94 |
| # | interaction | 2 | 9 | 0 | 0.181818182 | 1 | # | 10 | Baseline | paralle | 94 |
| # | disease | 4 | 2 | 0 | 0.666666667 | 1 | # | 10 | Baseline | ancom | 94 |
| # | sex | 2 | 0 | 0 | 1 | 1 | # | 10 | Baseline | ancom | 94 |
| # | time | 2 | 2 | 0 | 0.5 | 1 | # | 10 | Baseline | ancom | 94 |
| # | interaction | 0 | 0 | 2 | 0 | 0 | # | 10 | Baseline | ancom | 94 |
| # | disease | 4 | 19 | 0 | 0.173913043 | 1 | # | 10 | Baseline | aldex2 | 94 |
| # | sex | 2 | 7 | 0 | 0.222222222 | 1 | # | 10 | Baseline | aldex2 | 94 |
| # | time | 2 | 12 | 0 | 0.142857143 | 1 | # | 10 | Baseline | aldex2 | 94 |
| # | interaction | 0 | 0 | 2 | 0 | 0 | # | 10 | Baseline | aldex2 | 94 |
| # | disease | 4 | 0 | 0 | 1 | 1 | # | 10 | Small | paralle | 94 |
| # | sex | 2 | 0 | 0 | 1 | 1 | # | 10 | Small | paralle | 94 |
| # | time | 2 | 0 | 0 | 1 | 1 | # | 10 | Small | paralle | 94 |
| # | interaction | 2 | 0 | 0 | 1 | 1 | # | 10 | Small | paralle | 94 |
| # | disease | 4 | 2 | 0 | 0.666666667 | 1 | # | 10 | Small | ancom | 94 |
| # | sex | 2 | 0 | 0 | 1 | 1 | # | 10 | Small | ancom | 94 |
| # | time | 2 | 2 | 0 | 0.5 | 1 | # | 10 | Small | ancom | 94 |
| # | interaction | 0 | 0 | 2 | 0 | 0 | # | 10 | Small | ancom | 94 |
| # | disease | 4 | 2 | 0 | 0.666666667 | 1 | # | 10 | Small | aldex2 | 94 |
| # | sex | 2 | 0 | 0 | 1 | 1 | # | 10 | Small | aldex2 | 94 |
| # | time | 2 | 3 | 0 | 0.4 | 1 | # | 10 | Small | aldex2 | 94 |
| # | interaction | 0 | 0 | 2 | 0 | 0 | # | 10 | Small | aldex2 | 94 |
| # | disease | 4 | 3 | 0 | 0.571428571 | 1 | # | 10 | Baseline | paralle | 94 |
| # | sex | 2 | 4 | 0 | 0.333333333 | 1 | # | 10 | Baseline | paralle | 94 |
| # | time | 2 | 5 | 0 | 0.285714286 | 1 | # | 10 | Baseline | paralle | 94 |
| # | interaction | 2 | 4 | 0 | 0.333333333 | 1 | # | 10 | Baseline | paralle | 94 |
| # | disease | 4 | 2 | 0 | 0.666666667 | 1 | # | 10 | Baseline | ancom | 94 |
| # | sex | 2 | 0 | 0 | 1 | 1 | # | 10 | Baseline | ancom | 94 |
| # | time | 2 | 2 | 0 | 0.5 | 1 | # | 10 | Baseline | ancom | 94 |
| # | interaction | 0 | 0 | 2 | 0 | 0 | # | 10 | Baseline | ancom | 94 |
| # | disease | 4 | 21 | 0 | 0.16 | 1 | # | 10 | Baseline | aldex2 | 94 |
| # | sex | 2 | 8 | 0 | 0.2 | 1 | # | 10 | Baseline | aldex2 | 94 |
| # | time | 2 | 11 | 0 | 0.153846154 | 1 | # | 10 | Baseline | aldex2 | 94 |
| # | interaction | 0 | 0 | 2 | 0 | 0 | # | 10 | Baseline | aldex2 | 94 |
| # | disease | 4 | 0 | 0 | 1 | 1 | # | 10 | Small | paralle | 94 |
| # | sex | 2 | 0 | 0 | 1 | 1 | # | 10 | Small | paralle | 94 |
| # | time | 2 | 0 | 0 | 1 | 1 | # | 10 | Small | paralle | 94 |
| # | interaction | 2 | 0 | 0 | 1 | 1 | # | 10 | Small | paralle | 94 |
| # | disease | 4 | 2 | 0 | 0.666666667 | 1 | # | 10 | Small | ancom | 94 |
| # | sex | 2 | 0 | 0 | 1 | 1 | # | 10 | Small | ancom | 94 |
| # | time | 2 | 2 | 0 | 0.5 | 1 | # | 10 | Small | ancom | 94 |
| # | interaction | 0 | 0 | 2 | 0 | 0 | # | 10 | Small | ancom | 94 |
| # | disease | 4 | 5 | 0 | 0.444444444 | 1 | # | 10 | Small | aldex2 | 94 |
| # | sex | 2 | 0 | 0 | 1 | 1 | # | 10 | Small | aldex2 | 94 |
| # | time | 2 | 3 | 0 | 0.4 | 1 | # | 10 | Small | aldex2 | 94 |
| # | interaction | 0 | 0 | 2 | 0 | 0 | # | 10 | Small | aldex2 | 94 |
| # | disease | 4 | 9 | 0 | 0.307692308 | 1 | # | 10 | Baseline | paralle | 94 |
| # | sex | 2 | 2 | 0 | 0.5 | 1 | # | 10 | Baseline | paralle | 94 |
| # | time | 2 | 2 | 0 | 0.5 | 1 | # | 10 | Baseline | paralle | 94 |
| # | interaction | 2 | 5 | 0 | 0.285714286 | 1 | # | 10 | Baseline | paralle | 94 |
| # | disease | 4 | 2 | 0 | 0.666666667 | 1 | # | 10 | Baseline | ancom | 94 |
| # | sex | 2 | 0 | 0 | 1 | 1 | # | 10 | Baseline | ancom | 94 |
| # | time | 2 | 2 | 0 | 0.5 | 1 | # | 10 | Baseline | ancom | 94 |
| # | interaction | 0 | 0 | 2 | 0 | 0 | # | 10 | Baseline | ancom | 94 |
| # | disease | 4 | 25 | 0 | 0.137931034 | 1 | # | 10 | Baseline | aldex2 | 94 |
| # | sex | 2 | 6 | 0 | 0.25 | 1 | # | 10 | Baseline | aldex2 | 94 |
| # | time | 2 | 10 | 0 | 0.166666667 | 1 | # | 10 | Baseline | aldex2 | 94 |
| # | interaction | 0 | 0 | 2 | 0 | 0 | # | 10 | Baseline | aldex2 | 94 |
| # | disease | 4 | 0 | 0 | 1 | 1 | # | 10 | Small | paralle | 94 |
| # | sex | 2 | 0 | 0 | 1 | 1 | # | 10 | Small | paralle | 94 |
| # | time | 2 | 0 | 0 | 1 | 1 | # | 10 | Small | paralle | 94 |
| # | interaction | 2 | 0 | 0 | 1 | 1 | # | 10 | Small | paralle | 94 |
| # | disease | 4 | 2 | 0 | 0.666666667 | 1 | # | 10 | Small | ancom | 94 |
| # | sex | 2 | 0 | 0 | 1 | 1 | # | 10 | Small | ancom | 94 |
| # | time | 2 | 2 | 0 | 0.5 | 1 | # | 10 | Small | ancom | 94 |
| # | interaction | 0 | 0 | 2 | 0 | 0 | # | 10 | Small | ancom | 94 |
| # | disease | 4 | 6 | 0 | 0.4 | 1 | # | 10 | Small | aldex2 | 94 |
| # | sex | 2 | 0 | 0 | 1 | 1 | # | 10 | Small | aldex2 | 94 |
| # | time | 2 | 2 | 0 | 0.5 | 1 | # | 10 | Small | aldex2 | 94 |
| # | interaction | 0 | 0 | 2 | 0 | 0 | # | 10 | Small | aldex2 | 94 |
| # | disease | 7 | 17 | 0 | 0.291666667 | 1 | # | 20 | Baseline | paralle | 94 |
| # | sex | 4 | 8 | 0 | 0.333333333 | 1 | # | 20 | Baseline | paralle | 94 |
| # | time | 5 | 9 | 0 | 0.357142857 | 1 | # | 20 | Baseline | paralle | 94 |
| # | interaction | 3 | 9 | 1 | 0.25 | 0.75 | # | 20 | Baseline | paralle | 94 |
| # | disease | 7 | 2 | 0 | 0.777777778 | 1 | # | 20 | Baseline | ancom | 94 |
| # | sex | 4 | 0 | 0 | 1 | 1 | # | 20 | Baseline | ancom | 94 |
| # | time | 5 | 0 | 0 | 1 | 1 | # | 20 | Baseline | ancom | 94 |
| # | interaction | 0 | 0 | 4 | 0 | 0 | # | 20 | Baseline | ancom | 94 |
| # | disease | 7 | 24 | 0 | 0.225806452 | 1 | # | 20 | Baseline | aldex2 | 94 |
| # | sex | 4 | 7 | 0 | 0.363636364 | 1 | # | 20 | Baseline | aldex2 | 94 |
| # | time | 5 | 13 | 0 | 0.277777778 | 1 | # | 20 | Baseline | aldex2 | 94 |
| # | interaction | 0 | 0 | 4 | 0 | 0 | # | 20 | Baseline | aldex2 | 94 |
| # | disease | 7 | 0 | 0 | 1 | 1 | # | 20 | Small | paralle | 94 |
| # | sex | 4 | 0 | 0 | 1 | 1 | # | 20 | Small | paralle | 94 |
| # | time | 5 | 0 | 0 | 1 | 1 | # | 20 | Small | paralle | 94 |
| # | interaction | 1 | 0 | 3 | 1 | 0.25 | # | 20 | Small | paralle | 94 |
| # | disease | 7 | 1 | 0 | 0.875 | 1 | # | 20 | Small | ancom | 94 |
| # | sex | 4 | 0 | 0 | 1 | 1 | # | 20 | Small | ancom | 94 |
| # | time | 5 | 0 | 0 | 1 | 1 | # | 20 | Small | ancom | 94 |
| # | interaction | 0 | 0 | 4 | 0 | 0 | # | 20 | Small | ancom | 94 |
| # | disease | 7 | 0 | 0 | 1 | 1 | # | 20 | Small | aldex2 | 94 |
| # | sex | 4 | 0 | 0 | 1 | 1 | # | 20 | Small | aldex2 | 94 |
| # | time | 5 | 0 | 0 | 1 | 1 | # | 20 | Small | aldex2 | 94 |
| # | interaction | 0 | 0 | 4 | 0 | 0 | # | 20 | Small | aldex2 | 94 |
| # | disease | 7 | 12 | 0 | 0.368421053 | 1 | # | 20 | Baseline | paralle | 94 |
| # | sex | 4 | 6 | 0 | 0.4 | 1 | # | 20 | Baseline | paralle | 94 |
| # | time | 5 | 12 | 0 | 0.294117647 | 1 | # | 20 | Baseline | paralle | 94 |
| # | interaction | 4 | 8 | 0 | 0.333333333 | 1 | # | 20 | Baseline | paralle | 94 |
| # | disease | 7 | 3 | 0 | 0.7 | 1 | # | 20 | Baseline | ancom | 94 |
| # | sex | 4 | 0 | 0 | 1 | 1 | # | 20 | Baseline | ancom | 94 |
| # | time | 5 | 2 | 0 | 0.714285714 | 1 | # | 20 | Baseline | ancom | 94 |
| # | interaction | 0 | 0 | 4 | 0 | 0 | # | 20 | Baseline | ancom | 94 |
| # | disease | 7 | 40 | 0 | 0.14893617 | 1 | # | 20 | Baseline | aldex2 | 94 |
| # | sex | 4 | 13 | 0 | 0.235294118 | 1 | # | 20 | Baseline | aldex2 | 94 |
| # | time | 5 | 22 | 0 | 0.185185185 | 1 | # | 20 | Baseline | aldex2 | 94 |
| # | interaction | 0 | 0 | 4 | 0 | 0 | # | 20 | Baseline | aldex2 | 94 |
| # | disease | 7 | 1 | 0 | 0.875 | 1 | # | 20 | Small | paralle | 94 |
| # | sex | 4 | 0 | 0 | 1 | 1 | # | 20 | Small | paralle | 94 |
| # | time | 5 | 0 | 0 | 1 | 1 | # | 20 | Small | paralle | 94 |
| # | interaction | 2 | 0 | 2 | 1 | 0.5 | # | 20 | Small | paralle | 94 |
| # | disease | 7 | 3 | 0 | 0.7 | 1 | # | 20 | Small | ancom | 94 |
| # | sex | 4 | 0 | 0 | 1 | 1 | # | 20 | Small | ancom | 94 |
| # | time | 5 | 2 | 0 | 0.714285714 | 1 | # | 20 | Small | ancom | 94 |
| # | interaction | 0 | 0 | 4 | 0 | 0 | # | 20 | Small | ancom | 94 |
| # | disease | 7 | 6 | 0 | 0.538461538 | 1 | # | 20 | Small | aldex2 | 94 |
| # | sex | 4 | 0 | 0 | 1 | 1 | # | 20 | Small | aldex2 | 94 |
| # | time | 5 | 2 | 0 | 0.714285714 | 1 | # | 20 | Small | aldex2 | 94 |
| # | interaction | 0 | 0 | 4 | 0 | 0 | # | 20 | Small | aldex2 | 94 |
| # | disease | 7 | 11 | 0 | 0.388888889 | 1 | # | 20 | Baseline | paralle | 94 |
| # | sex | 4 | 4 | 0 | 0.5 | 1 | # | 20 | Baseline | paralle | 94 |
| # | time | 5 | 11 | 0 | 0.3125 | 1 | # | 20 | Baseline | paralle | 94 |
| # | interaction | 4 | 2 | 0 | 0.666666667 | 1 | # | 20 | Baseline | paralle | 94 |
| # | disease | 7 | 3 | 0 | 0.7 | 1 | # | 20 | Baseline | ancom | 94 |
| # | sex | 4 | 0 | 0 | 1 | 1 | # | 20 | Baseline | ancom | 94 |
| # | time | 5 | 3 | 0 | 0.625 | 1 | # | 20 | Baseline | ancom | 94 |
| # | interaction | 0 | 0 | 4 | 0 | 0 | # | 20 | Baseline | ancom | 94 |
| # | disease | 7 | 53 | 0 | 0.116666667 | 1 | # | 20 | Baseline | aldex2 | 94 |
| # | sex | 4 | 17 | 0 | 0.19047619 | 1 | # | 20 | Baseline | aldex2 | 94 |
| # | time | 5 | 27 | 0 | 0.15625 | 1 | # | 20 | Baseline | aldex2 | 94 |
| # | interaction | 0 | 0 | 4 | 0 | 0 | # | 20 | Baseline | aldex2 | 94 |
| # | disease | 7 | 0 | 0 | 1 | 1 | # | 20 | Small | paralle | 94 |
| # | sex | 4 | 0 | 0 | 1 | 1 | # | 20 | Small | paralle | 94 |
| # | time | 5 | 0 | 0 | 1 | 1 | # | 20 | Small | paralle | 94 |
| # | interaction | 3 | 0 | 1 | 1 | 0.75 | # | 20 | Small | paralle | 94 |
| # | disease | 7 | 3 | 0 | 0.7 | 1 | # | 20 | Small | ancom | 94 |
| # | sex | 4 | 0 | 0 | 1 | 1 | # | 20 | Small | ancom | 94 |
| # | time | 5 | 3 | 0 | 0.625 | 1 | # | 20 | Small | ancom | 94 |
| # | interaction | 0 | 0 | 4 | 0 | 0 | # | 20 | Small | ancom | 94 |
| # | disease | 7 | 10 | 0 | 0.411764706 | 1 | # | 20 | Small | aldex2 | 94 |
| # | sex | 4 | 0 | 0 | 1 | 1 | # | 20 | Small | aldex2 | 94 |
| # | time | 5 | 3 | 0 | 0.625 | 1 | # | 20 | Small | aldex2 | 94 |
| # | interaction | 0 | 0 | 4 | 0 | 0 | # | 20 | Small | aldex2 | 94 |
| # | disease | 7 | 8 | 0 | 0.466666667 | 1 | # | 20 | Baseline | paralle | 94 |
| # | sex | 4 | 5 | 0 | 0.444444444 | 1 | # | 20 | Baseline | paralle | 94 |
| # | time | 5 | 10 | 0 | 0.333333333 | 1 | # | 20 | Baseline | paralle | 94 |
| # | interaction | 4 | 1 | 0 | 0.8 | 1 | # | 20 | Baseline | paralle | 94 |
| # | disease | 7 | 4 | 0 | 0.636363636 | 1 | # | 20 | Baseline | ancom | 94 |
| # | sex | 4 | 0 | 0 | 1 | 1 | # | 20 | Baseline | ancom | 94 |
| # | time | 5 | 4 | 0 | 0.555555556 | 1 | # | 20 | Baseline | ancom | 94 |
| # | interaction | 0 | 0 | 4 | 0 | 0 | # | 20 | Baseline | ancom | 94 |
| # | disease | 7 | 65 | 0 | 0.097222222 | 1 | # | 20 | Baseline | aldex2 | 94 |
| # | sex | 4 | 19 | 0 | 0.173913043 | 1 | # | 20 | Baseline | aldex2 | 94 |
| # | time | 5 | 35 | 0 | 0.125 | 1 | # | 20 | Baseline | aldex2 | 94 |
| # | interaction | 0 | 0 | 4 | 0 | 0 | # | 20 | Baseline | aldex2 | 94 |
| # | disease | 7 | 0 | 0 | 1 | 1 | # | 20 | Small | paralle | 94 |
| # | sex | 4 | 0 | 0 | 1 | 1 | # | 20 | Small | paralle | 94 |
| # | time | 5 | 0 | 0 | 1 | 1 | # | 20 | Small | paralle | 94 |
| # | interaction | 4 | 0 | 0 | 1 | 1 | # | 20 | Small | paralle | 94 |
| # | disease | 7 | 3 | 0 | 0.7 | 1 | # | 20 | Small | ancom | 94 |
| # | sex | 4 | 0 | 0 | 1 | 1 | # | 20 | Small | ancom | 94 |
| # | time | 5 | 4 | 0 | 0.555555556 | 1 | # | 20 | Small | ancom | 94 |
| # | interaction | 0 | 0 | 4 | 0 | 0 | # | 20 | Small | ancom | 94 |
| # | disease | 7 | 12 | 0 | 0.368421053 | 1 | # | 20 | Small | aldex2 | 94 |
| # | sex | 4 | 0 | 0 | 1 | 1 | # | 20 | Small | aldex2 | 94 |
| # | time | 5 | 7 | 0 | 0.416666667 | 1 | # | 20 | Small | aldex2 | 94 |
| # | interaction | 0 | 0 | 4 | 0 | 0 | # | 20 | Small | aldex2 | 94 |
| # | disease | 4 | 10 | 0 | 0.285714286 | 1 | # | 10 | Baseline | paralle | 95 |
| # | sex | 2 | 7 | 0 | 0.222222222 | 1 | # | 10 | Baseline | paralle | 95 |
| # | time | 2 | 7 | 0 | 0.222222222 | 1 | # | 10 | Baseline | paralle | 95 |
| # | interaction | 2 | 8 | 0 | 0.2 | 1 | # | 10 | Baseline | paralle | 95 |
| # | disease | 4 | 1 | 0 | 0.8 | 1 | # | 10 | Baseline | ancom | 95 |
| # | sex | 2 | 0 | 0 | 1 | 1 | # | 10 | Baseline | ancom | 95 |
| # | time | 2 | 1 | 0 | 0.666666667 | 1 | # | 10 | Baseline | ancom | 95 |
| # | interaction | 0 | 0 | 2 | 0 | 0 | # | 10 | Baseline | ancom | 95 |
| # | disease | 4 | 8 | 0 | 0.333333333 | 1 | # | 10 | Baseline | aldex2 | 95 |
| # | sex | 2 | 5 | 0 | 0.285714286 | 1 | # | 10 | Baseline | aldex2 | 95 |
| # | time | 2 | 6 | 0 | 0.25 | 1 | # | 10 | Baseline | aldex2 | 95 |
| # | interaction | 0 | 0 | 2 | 0 | 0 | # | 10 | Baseline | aldex2 | 95 |
| # | disease | 4 | 0 | 0 | 1 | 1 | # | 10 | Small | paralle | 95 |
| # | sex | 2 | 0 | 0 | 1 | 1 | # | 10 | Small | paralle | 95 |
| # | time | 2 | 0 | 0 | 1 | 1 | # | 10 | Small | paralle | 95 |
| # | interaction | 1 | 1 | 1 | 0.5 | 0.5 | # | 10 | Small | paralle | 95 |
| # | disease | 4 | 1 | 0 | 0.8 | 1 | # | 10 | Small | ancom | 95 |
| # | sex | 2 | 0 | 0 | 1 | 1 | # | 10 | Small | ancom | 95 |
| # | time | 2 | 1 | 0 | 0.666666667 | 1 | # | 10 | Small | ancom | 95 |
| # | interaction | 0 | 0 | 2 | 0 | 0 | # | 10 | Small | ancom | 95 |
| # | disease | 4 | 1 | 0 | 0.8 | 1 | # | 10 | Small | aldex2 | 95 |
| # | sex | 2 | 0 | 0 | 1 | 1 | # | 10 | Small | aldex2 | 95 |
| # | time | 2 | 1 | 0 | 0.666666667 | 1 | # | 10 | Small | aldex2 | 95 |
| # | interaction | 0 | 0 | 2 | 0 | 0 | # | 10 | Small | aldex2 | 95 |
| # | disease | 4 | 9 | 0 | 0.307692308 | 1 | # | 10 | Baseline | paralle | 95 |
| # | sex | 2 | 8 | 0 | 0.2 | 1 | # | 10 | Baseline | paralle | 95 |
| # | time | 2 | 6 | 0 | 0.25 | 1 | # | 10 | Baseline | paralle | 95 |
| # | interaction | 2 | 8 | 0 | 0.2 | 1 | # | 10 | Baseline | paralle | 95 |
| # | disease | 4 | 2 | 0 | 0.666666667 | 1 | # | 10 | Baseline | ancom | 95 |
| # | sex | 2 | 0 | 0 | 1 | 1 | # | 10 | Baseline | ancom | 95 |
| # | time | 2 | 2 | 0 | 0.5 | 1 | # | 10 | Baseline | ancom | 95 |
| # | interaction | 0 | 0 | 2 | 0 | 0 | # | 10 | Baseline | ancom | 95 |
| # | disease | 4 | 14 | 0 | 0.222222222 | 1 | # | 10 | Baseline | aldex2 | 95 |
| # | sex | 2 | 9 | 0 | 0.181818182 | 1 | # | 10 | Baseline | aldex2 | 95 |
| # | time | 2 | 8 | 0 | 0.2 | 1 | # | 10 | Baseline | aldex2 | 95 |
| # | interaction | 0 | 0 | 2 | 0 | 0 | # | 10 | Baseline | aldex2 | 95 |
| # | disease | 4 | 0 | 0 | 1 | 1 | # | 10 | Small | paralle | 95 |
| # | sex | 2 | 0 | 0 | 1 | 1 | # | 10 | Small | paralle | 95 |
| # | time | 2 | 0 | 0 | 1 | 1 | # | 10 | Small | paralle | 95 |
| # | interaction | 2 | 0 | 0 | 1 | 1 | # | 10 | Small | paralle | 95 |
| # | disease | 4 | 2 | 0 | 0.666666667 | 1 | # | 10 | Small | ancom | 95 |
| # | sex | 2 | 0 | 0 | 1 | 1 | # | 10 | Small | ancom | 95 |
| # | time | 2 | 2 | 0 | 0.5 | 1 | # | 10 | Small | ancom | 95 |
| # | interaction | 0 | 0 | 2 | 0 | 0 | # | 10 | Small | ancom | 95 |
| # | disease | 4 | 2 | 0 | 0.666666667 | 1 | # | 10 | Small | aldex2 | 95 |
| # | sex | 2 | 0 | 0 | 1 | 1 | # | 10 | Small | aldex2 | 95 |
| # | time | 2 | 2 | 0 | 0.5 | 1 | # | 10 | Small | aldex2 | 95 |
| # | interaction | 0 | 0 | 2 | 0 | 0 | # | 10 | Small | aldex2 | 95 |
| # | disease | 4 | 7 | 0 | 0.363636364 | 1 | # | 10 | Baseline | paralle | 95 |
| # | sex | 2 | 5 | 0 | 0.285714286 | 1 | # | 10 | Baseline | paralle | 95 |
| # | time | 2 | 3 | 0 | 0.4 | 1 | # | 10 | Baseline | paralle | 95 |
| # | interaction | 2 | 3 | 0 | 0.4 | 1 | # | 10 | Baseline | paralle | 95 |
| # | disease | 4 | 2 | 0 | 0.666666667 | 1 | # | 10 | Baseline | ancom | 95 |
| # | sex | 2 | 0 | 0 | 1 | 1 | # | 10 | Baseline | ancom | 95 |
| # | time | 2 | 2 | 0 | 0.5 | 1 | # | 10 | Baseline | ancom | 95 |
| # | interaction | 0 | 0 | 2 | 0 | 0 | # | 10 | Baseline | ancom | 95 |
| # | disease | 4 | 20 | 0 | 0.166666667 | 1 | # | 10 | Baseline | aldex2 | 95 |
| # | sex | 2 | 7 | 0 | 0.222222222 | 1 | # | 10 | Baseline | aldex2 | 95 |
| # | time | 2 | 10 | 0 | 0.166666667 | 1 | # | 10 | Baseline | aldex2 | 95 |
| # | interaction | 0 | 0 | 2 | 0 | 0 | # | 10 | Baseline | aldex2 | 95 |
| # | disease | 4 | 0 | 0 | 1 | 1 | # | 10 | Small | paralle | 95 |
| # | sex | 2 | 0 | 0 | 1 | 1 | # | 10 | Small | paralle | 95 |
| # | time | 2 | 0 | 0 | 1 | 1 | # | 10 | Small | paralle | 95 |
| # | interaction | 2 | 0 | 0 | 1 | 1 | # | 10 | Small | paralle | 95 |
| # | disease | 4 | 2 | 0 | 0.666666667 | 1 | # | 10 | Small | ancom | 95 |
| # | sex | 2 | 0 | 0 | 1 | 1 | # | 10 | Small | ancom | 95 |
| # | time | 2 | 2 | 0 | 0.5 | 1 | # | 10 | Small | ancom | 95 |
| # | interaction | 0 | 0 | 2 | 0 | 0 | # | 10 | Small | ancom | 95 |
| # | disease | 4 | 2 | 0 | 0.666666667 | 1 | # | 10 | Small | aldex2 | 95 |
| # | sex | 2 | 0 | 0 | 1 | 1 | # | 10 | Small | aldex2 | 95 |
| # | time | 2 | 2 | 0 | 0.5 | 1 | # | 10 | Small | aldex2 | 95 |
| # | interaction | 0 | 0 | 2 | 0 | 0 | # | 10 | Small | aldex2 | 95 |
| # | disease | 4 | 5 | 0 | 0.444444444 | 1 | # | 10 | Baseline | paralle | 95 |
| # | sex | 2 | 4 | 0 | 0.333333333 | 1 | # | 10 | Baseline | paralle | 95 |
| # | time | 2 | 4 | 0 | 0.333333333 | 1 | # | 10 | Baseline | paralle | 95 |
| # | interaction | 2 | 2 | 0 | 0.5 | 1 | # | 10 | Baseline | paralle | 95 |
| # | disease | 4 | 2 | 0 | 0.666666667 | 1 | # | 10 | Baseline | ancom | 95 |
| # | sex | 2 | 0 | 0 | 1 | 1 | # | 10 | Baseline | ancom | 95 |
| # | time | 2 | 2 | 0 | 0.5 | 1 | # | 10 | Baseline | ancom | 95 |
| # | interaction | 0 | 0 | 2 | 0 | 0 | # | 10 | Baseline | ancom | 95 |
| # | disease | 4 | 32 | 0 | 0.111111111 | 1 | # | 10 | Baseline | aldex2 | 95 |
| # | sex | 2 | 7 | 0 | 0.222222222 | 1 | # | 10 | Baseline | aldex2 | 95 |
| # | time | 2 | 9 | 0 | 0.181818182 | 1 | # | 10 | Baseline | aldex2 | 95 |
| # | interaction | 0 | 0 | 2 | 0 | 0 | # | 10 | Baseline | aldex2 | 95 |
| # | disease | 4 | 0 | 0 | 1 | 1 | # | 10 | Small | paralle | 95 |
| # | sex | 2 | 0 | 0 | 1 | 1 | # | 10 | Small | paralle | 95 |
| # | time | 2 | 0 | 0 | 1 | 1 | # | 10 | Small | paralle | 95 |
| # | interaction | 2 | 0 | 0 | 1 | 1 | # | 10 | Small | paralle | 95 |
| # | disease | 4 | 2 | 0 | 0.666666667 | 1 | # | 10 | Small | ancom | 95 |
| # | sex | 2 | 0 | 0 | 1 | 1 | # | 10 | Small | ancom | 95 |
| # | time | 2 | 2 | 0 | 0.5 | 1 | # | 10 | Small | ancom | 95 |
| # | interaction | 0 | 0 | 2 | 0 | 0 | # | 10 | Small | ancom | 95 |
| # | disease | 4 | 3 | 0 | 0.571428571 | 1 | # | 10 | Small | aldex2 | 95 |
| # | sex | 2 | 0 | 0 | 1 | 1 | # | 10 | Small | aldex2 | 95 |
| # | time | 2 | 2 | 0 | 0.5 | 1 | # | 10 | Small | aldex2 | 95 |
| # | interaction | 0 | 0 | 2 | 0 | 0 | # | 10 | Small | aldex2 | 95 |
| # | disease | 7 | 6 | 0 | 0.538461538 | 1 | # | 20 | Baseline | paralle | 95 |
| # | sex | 4 | 7 | 0 | 0.363636364 | 1 | # | 20 | Baseline | paralle | 95 |
| # | time | 5 | 9 | 0 | 0.357142857 | 1 | # | 20 | Baseline | paralle | 95 |
| # | interaction | 4 | 3 | 0 | 0.571428571 | 1 | # | 20 | Baseline | paralle | 95 |
| # | disease | 7 | 3 | 0 | 0.7 | 1 | # | 20 | Baseline | ancom | 95 |
| # | sex | 4 | 0 | 0 | 1 | 1 | # | 20 | Baseline | ancom | 95 |
| # | time | 5 | 0 | 0 | 1 | 1 | # | 20 | Baseline | ancom | 95 |
| # | interaction | 0 | 0 | 4 | 0 | 0 | # | 20 | Baseline | ancom | 95 |
| # | disease | 7 | 21 | 0 | 0.25 | 1 | # | 20 | Baseline | aldex2 | 95 |
| # | sex | 4 | 7 | 0 | 0.363636364 | 1 | # | 20 | Baseline | aldex2 | 95 |
| # | time | 5 | 14 | 0 | 0.263157895 | 1 | # | 20 | Baseline | aldex2 | 95 |
| # | interaction | 0 | 0 | 4 | 0 | 0 | # | 20 | Baseline | aldex2 | 95 |
| # | disease | 7 | 0 | 0 | 1 | 1 | # | 20 | Small | paralle | 95 |
| # | sex | 4 | 0 | 0 | 1 | 1 | # | 20 | Small | paralle | 95 |
| # | time | 5 | 0 | 0 | 1 | 1 | # | 20 | Small | paralle | 95 |
| # | interaction | 2 | 0 | 2 | 1 | 0.5 | # | 20 | Small | paralle | 95 |
| # | disease | 7 | 3 | 0 | 0.7 | 1 | # | 20 | Small | ancom | 95 |
| # | sex | 4 | 0 | 0 | 1 | 1 | # | 20 | Small | ancom | 95 |
| # | time | 5 | 0 | 0 | 1 | 1 | # | 20 | Small | ancom | 95 |
| # | interaction | 0 | 0 | 4 | 0 | 0 | # | 20 | Small | ancom | 95 |
| # | disease | 7 | 2 | 0 | 0.777777778 | 1 | # | 20 | Small | aldex2 | 95 |
| # | sex | 4 | 0 | 0 | 1 | 1 | # | 20 | Small | aldex2 | 95 |
| # | time | 5 | 0 | 0 | 1 | 1 | # | 20 | Small | aldex2 | 95 |
| # | interaction | 0 | 0 | 4 | 0 | 0 | # | 20 | Small | aldex2 | 95 |
| # | disease | 7 | 8 | 0 | 0.466666667 | 1 | # | 20 | Baseline | paralle | 95 |
| # | sex | 4 | 4 | 0 | 0.5 | 1 | # | 20 | Baseline | paralle | 95 |
| # | time | 5 | 1 | 0 | 0.833333333 | 1 | # | 20 | Baseline | paralle | 95 |
| # | interaction | 4 | 4 | 0 | 0.5 | 1 | # | 20 | Baseline | paralle | 95 |
| # | disease | 7 | 4 | 0 | 0.636363636 | 1 | # | 20 | Baseline | ancom | 95 |
| # | sex | 4 | 0 | 0 | 1 | 1 | # | 20 | Baseline | ancom | 95 |
| # | time | 5 | 2 | 0 | 0.714285714 | 1 | # | 20 | Baseline | ancom | 95 |
| # | interaction | 0 | 0 | 4 | 0 | 0 | # | 20 | Baseline | ancom | 95 |
| # | disease | 7 | 29 | 0 | 0.194444444 | 1 | # | 20 | Baseline | aldex2 | 95 |
| # | sex | 4 | 7 | 0 | 0.363636364 | 1 | # | 20 | Baseline | aldex2 | 95 |
| # | time | 5 | 22 | 0 | 0.185185185 | 1 | # | 20 | Baseline | aldex2 | 95 |
| # | interaction | 0 | 0 | 4 | 0 | 0 | # | 20 | Baseline | aldex2 | 95 |
| # | disease | 7 | 0 | 0 | 1 | 1 | # | 20 | Small | paralle | 95 |
| # | sex | 4 | 0 | 0 | 1 | 1 | # | 20 | Small | paralle | 95 |
| # | time | 5 | 0 | 0 | 1 | 1 | # | 20 | Small | paralle | 95 |
| # | interaction | 3 | 0 | 1 | 1 | 0.75 | # | 20 | Small | paralle | 95 |
| # | disease | 7 | 4 | 0 | 0.636363636 | 1 | # | 20 | Small | ancom | 95 |
| # | sex | 4 | 0 | 0 | 1 | 1 | # | 20 | Small | ancom | 95 |
| # | time | 5 | 1 | 0 | 0.833333333 | 1 | # | 20 | Small | ancom | 95 |
| # | interaction | 0 | 0 | 4 | 0 | 0 | # | 20 | Small | ancom | 95 |
| # | disease | 7 | 4 | 0 | 0.636363636 | 1 | # | 20 | Small | aldex2 | 95 |
| # | sex | 4 | 0 | 0 | 1 | 1 | # | 20 | Small | aldex2 | 95 |
| # | time | 5 | 2 | 0 | 0.714285714 | 1 | # | 20 | Small | aldex2 | 95 |
| # | interaction | 0 | 0 | 4 | 0 | 0 | # | 20 | Small | aldex2 | 95 |
| # | disease | 7 | 6 | 0 | 0.538461538 | 1 | # | 20 | Baseline | paralle | 95 |
| # | sex | 4 | 7 | 0 | 0.363636364 | 1 | # | 20 | Baseline | paralle | 95 |
| # | time | 5 | 5 | 0 | 0.5 | 1 | # | 20 | Baseline | paralle | 95 |

| | | | | | | | | | | | |
|---|---|---|---|---|---|---|---|---|---|---|---|
| # | interaction | 4 | 3 | 0 | 0.571428571 | 1 | # | 20 | Baseline | paralle | 95 |
| # | disease | 7 | 4 | 0 | 0.636363636 | 1 | # | 20 | Baseline | ancom | 95 |
| # | sex | 4 | 0 | 0 | 1 | 1 | # | 20 | Baseline | ancom | 95 |
| # | time | 5 | 4 | 0 | 0.555555556 | 1 | # | 20 | Baseline | ancom | 95 |
| # | interaction | 0 | 0 | 4 | 0 | 0 | # | 20 | Baseline | ancom | 95 |
| # | disease | 7 | 48 | 0 | 0.127272727 | 1 | # | 20 | Baseline | aldex2 | 95 |
| # | sex | 4 | 10 | 0 | 0.285714286 | 1 | # | 20 | Baseline | aldex2 | 95 |
| # | time | 5 | 36 | 0 | 0.12195122 | 1 | # | 20 | Baseline | aldex2 | 95 |
| # | interaction | 0 | 0 | 4 | 0 | 0 | # | 20 | Baseline | aldex2 | 95 |
| # | disease | 7 | 0 | 0 | 1 | 1 | # | 20 | Small | paralle | 95 |
| # | sex | 4 | 0 | 0 | 1 | 1 | # | 20 | Small | paralle | 95 |
| # | time | 5 | 0 | 0 | 1 | 1 | # | 20 | Small | paralle | 95 |
| # | interaction | 3 | 0 | 1 | 1 | 0.75 | # | 20 | Small | paralle | 95 |
| # | disease | 7 | 4 | 0 | 0.636363636 | 1 | # | 20 | Small | ancom | 95 |
| # | sex | 4 | 0 | 0 | 1 | 1 | # | 20 | Small | ancom | 95 |
| # | time | 5 | 4 | 0 | 0.555555556 | 1 | # | 20 | Small | ancom | 95 |
| # | interaction | 0 | 0 | 4 | 0 | 0 | # | 20 | Small | ancom | 95 |
| # | disease | 7 | 3 | 0 | 0.7 | 1 | # | 20 | Small | aldex2 | 95 |
| # | sex | 4 | 1 | 0 | 0.8 | 1 | # | 20 | Small | aldex2 | 95 |
| # | time | 5 | 3 | 0 | 0.625 | 1 | # | 20 | Small | aldex2 | 95 |
| # | interaction | 0 | 0 | 4 | 0 | 0 | # | 20 | Small | aldex2 | 95 |
| # | disease | 7 | 6 | 0 | 0.538461538 | 1 | # | 20 | Baseline | paralle | 95 |
| # | sex | 4 | 3 | 0 | 0.571428571 | 1 | # | 20 | Baseline | paralle | 95 |
| # | time | 5 | 7 | 0 | 0.416666667 | 1 | # | 20 | Baseline | paralle | 95 |
| # | interaction | 4 | 6 | 0 | 0.4 | 1 | # | 20 | Baseline | paralle | 95 |
| # | disease | 7 | 4 | 0 | 0.636363636 | 1 | # | 20 | Baseline | ancom | 95 |
| # | sex | 4 | 0 | 0 | 1 | 1 | # | 20 | Baseline | ancom | 95 |
| # | time | 5 | 4 | 0 | 0.555555556 | 1 | # | 20 | Baseline | ancom | 95 |
| # | interaction | 0 | 0 | 4 | 0 | 0 | # | 20 | Baseline | ancom | 95 |
| # | disease | 7 | 56 | 0 | 0.111111111 | 1 | # | 20 | Baseline | aldex2 | 95 |
| # | sex | 4 | 13 | 0 | 0.235294118 | 1 | # | 20 | Baseline | aldex2 | 95 |
| # | time | 5 | 41 | 0 | 0.108695652 | 1 | # | 20 | Baseline | aldex2 | 95 |
| # | interaction | 0 | 0 | 4 | 0 | 0 | # | 20 | Baseline | aldex2 | 95 |
| # | disease | 7 | 0 | 0 | 1 | 1 | # | 20 | Small | paralle | 95 |
| # | sex | 4 | 0 | 0 | 1 | 1 | # | 20 | Small | paralle | 95 |
| # | time | 5 | 0 | 0 | 1 | 1 | # | 20 | Small | paralle | 95 |
| # | interaction | 3 | 0 | 1 | 1 | 0.75 | # | 20 | Small | paralle | 95 |
| # | disease | 7 | 4 | 0 | 0.636363636 | 1 | # | 20 | Small | ancom | 95 |
| # | sex | 4 | 0 | 0 | 1 | 1 | # | 20 | Small | ancom | 95 |
| # | time | 5 | 4 | 0 | 0.555555556 | 1 | # | 20 | Small | ancom | 95 |
| # | interaction | 0 | 0 | 4 | 0 | 0 | # | 20 | Small | ancom | 95 |
| # | disease | 7 | 9 | 0 | 0.4375 | 1 | # | 20 | Small | aldex2 | 95 |
| # | sex | 4 | 0 | 0 | 1 | 1 | # | 20 | Small | aldex2 | 95 |
| # | time | 5 | 6 | 0 | 0.454545455 | 1 | # | 20 | Small | aldex2 | 95 |
| # | interaction | 0 | 0 | 4 | 0 | 0 | # | 20 | Small | aldex2 | 95 |
| # | disease | 4 | 4 | 0 | 0.5 | 1 | # | 10 | Baseline | paralle | 96 |
| # | sex | 2 | 8 | 0 | 0.2 | 1 | # | 10 | Baseline | paralle | 96 |
| # | time | 2 | 6 | 0 | 0.25 | 1 | # | 10 | Baseline | paralle | 96 |
| # | interaction | 2 | 6 | 0 | 0.25 | 1 | # | 10 | Baseline | paralle | 96 |
| # | disease | 4 | 1 | 0 | 0.8 | 1 | # | 10 | Baseline | ancom | 96 |
| # | sex | 2 | 0 | 0 | 1 | 1 | # | 10 | Baseline | ancom | 96 |
| # | time | 2 | 1 | 0 | 0.666666667 | 1 | # | 10 | Baseline | ancom | 96 |
| # | interaction | 0 | 0 | 2 | 0 | 0 | # | 10 | Baseline | ancom | 96 |
| # | disease | 4 | 6 | 0 | 0.4 | 1 | # | 10 | Baseline | aldex2 | 96 |
| # | sex | 2 | 6 | 0 | 0.25 | 1 | # | 10 | Baseline | aldex2 | 96 |
| # | time | 2 | 7 | 0 | 0.222222222 | 1 | # | 10 | Baseline | aldex2 | 96 |
| # | interaction | 0 | 0 | 2 | 0 | 0 | # | 10 | Baseline | aldex2 | 96 |
| # | disease | 4 | 0 | 0 | 1 | 1 | # | 10 | Small | paralle | 96 |
| # | sex | 2 | 0 | 0 | 1 | 1 | # | 10 | Small | paralle | 96 |
| # | time | 2 | 0 | 0 | 1 | 1 | # | 10 | Small | paralle | 96 |
| # | interaction | 1 | 0 | 1 | 1 | 0.5 | # | 10 | Small | paralle | 96 |
| # | disease | 4 | 0 | 0 | 1 | 1 | # | 10 | Small | ancom | 96 |
| # | sex | 2 | 0 | 0 | 1 | 1 | # | 10 | Small | ancom | 96 |
| # | time | 2 | 1 | 0 | 0.666666667 | 1 | # | 10 | Small | ancom | 96 |
| # | interaction | 0 | 0 | 2 | 0 | 0 | # | 10 | Small | ancom | 96 |
| # | disease | 4 | 0 | 0 | 1 | 1 | # | 10 | Small | aldex2 | 96 |
| # | sex | 2 | 0 | 0 | 1 | 1 | # | 10 | Small | aldex2 | 96 |
| # | time | 2 | 1 | 0 | 0.666666667 | 1 | # | 10 | Small | aldex2 | 96 |
| # | interaction | 0 | 0 | 2 | 0 | 0 | # | 10 | Small | aldex2 | 96 |
| # | disease | 4 | 4 | 0 | 0.5 | 1 | # | 10 | Baseline | paralle | 96 |
| # | sex | 2 | 4 | 0 | 0.333333333 | 1 | # | 10 | Baseline | paralle | 96 |
| # | time | 2 | 3 | 0 | 0.4 | 1 | # | 10 | Baseline | paralle | 96 |
| # | interaction | 2 | 3 | 0 | 0.4 | 1 | # | 10 | Baseline | paralle | 96 |
| # | disease | 4 | 2 | 0 | 0.666666667 | 1 | # | 10 | Baseline | ancom | 96 |
| # | sex | 2 | 0 | 0 | 1 | 1 | # | 10 | Baseline | ancom | 96 |
| # | time | 2 | 1 | 0 | 0.666666667 | 1 | # | 10 | Baseline | ancom | 96 |
| # | interaction | 0 | 0 | 2 | 0 | 0 | # | 10 | Baseline | ancom | 96 |
| # | disease | 4 | 12 | 0 | 0.25 | 1 | # | 10 | Baseline | aldex2 | 96 |
| # | sex | 2 | 4 | 0 | 0.333333333 | 1 | # | 10 | Baseline | aldex2 | 96 |
| # | time | 2 | 12 | 0 | 0.142857143 | 1 | # | 10 | Baseline | aldex2 | 96 |
| # | interaction | 0 | 0 | 2 | 0 | 0 | # | 10 | Baseline | aldex2 | 96 |
| # | disease | 4 | 0 | 0 | 1 | 1 | # | 10 | Small | paralle | 96 |
| # | sex | 2 | 0 | 0 | 1 | 1 | # | 10 | Small | paralle | 96 |
| # | time | 2 | 0 | 0 | 1 | 1 | # | 10 | Small | paralle | 96 |
| # | interaction | 2 | 0 | 0 | 1 | 1 | # | 10 | Small | paralle | 96 |
| # | disease | 4 | 2 | 0 | 0.666666667 | 1 | # | 10 | Small | ancom | 96 |
| # | sex | 2 | 0 | 0 | 1 | 1 | # | 10 | Small | ancom | 96 |
| # | time | 2 | 1 | 0 | 0.666666667 | 1 | # | 10 | Small | ancom | 96 |
| # | interaction | 0 | 0 | 2 | 0 | 0 | # | 10 | Small | ancom | 96 |
| # | disease | 4 | 2 | 0 | 0.666666667 | 1 | # | 10 | Small | aldex2 | 96 |
| # | sex | 2 | 0 | 0 | 1 | 1 | # | 10 | Small | aldex2 | 96 |
| # | time | 2 | 1 | 0 | 0.666666667 | 1 | # | 10 | Small | aldex2 | 96 |
| # | interaction | 0 | 0 | 2 | 0 | 0 | # | 10 | Small | aldex2 | 96 |
| # | disease | 4 | 4 | 0 | 0.5 | 1 | # | 10 | Baseline | paralle | 96 |
| # | sex | 2 | 2 | 0 | 0.5 | 1 | # | 10 | Baseline | paralle | 96 |
| # | time | 2 | 3 | 0 | 0.4 | 1 | # | 10 | Baseline | paralle | 96 |
| # | interaction | 2 | 2 | 0 | 0.5 | 1 | # | 10 | Baseline | paralle | 96 |
| # | disease | 4 | 2 | 0 | 0.666666667 | 1 | # | 10 | Baseline | ancom | 96 |
| # | sex | 2 | 0 | 0 | 1 | 1 | # | 10 | Baseline | ancom | 96 |
| # | time | 2 | 2 | 0 | 0.5 | 1 | # | 10 | Baseline | ancom | 96 |
| # | interaction | 0 | 0 | 2 | 0 | 0 | # | 10 | Baseline | ancom | 96 |
| # | disease | 4 | 13 | 0 | 0.235294118 | 1 | # | 10 | Baseline | aldex2 | 96 |
| # | sex | 2 | 5 | 0 | 0.285714286 | 1 | # | 10 | Baseline | aldex2 | 96 |
| # | time | 2 | 13 | 0 | 0.133333333 | 1 | # | 10 | Baseline | aldex2 | 96 |
| # | interaction | 0 | 0 | 2 | 0 | 0 | # | 10 | Baseline | aldex2 | 96 |
| # | disease | 4 | 0 | 0 | 1 | 1 | # | 10 | Small | paralle | 96 |
| # | sex | 2 | 0 | 0 | 1 | 1 | # | 10 | Small | paralle | 96 |
| # | time | 2 | 0 | 0 | 1 | 1 | # | 10 | Small | paralle | 96 |
| # | interaction | 2 | 0 | 0 | 1 | 1 | # | 10 | Small | paralle | 96 |
| # | disease | 4 | 2 | 0 | 0.666666667 | 1 | # | 10 | Small | ancom | 96 |
| # | sex | 2 | 0 | 0 | 1 | 1 | # | 10 | Small | ancom | 96 |
| # | time | 2 | 1 | 0 | 0.666666667 | 1 | # | 10 | Small | ancom | 96 |
| # | interaction | 0 | 0 | 2 | 0 | 0 | # | 10 | Small | ancom | 96 |
| # | disease | 4 | 2 | 0 | 0.666666667 | 1 | # | 10 | Small | aldex2 | 96 |
| # | sex | 2 | 0 | 0 | 1 | 1 | # | 10 | Small | aldex2 | 96 |
| # | time | 2 | 2 | 0 | 0.5 | 1 | # | 10 | Small | aldex2 | 96 |
| # | interaction | 0 | 0 | 2 | 0 | 0 | # | 10 | Small | aldex2 | 96 |
| # | disease | 4 | 3 | 0 | 0.571428571 | 1 | # | 10 | Baseline | paralle | 96 |
| # | sex | 2 | 5 | 0 | 0.285714286 | 1 | # | 10 | Baseline | paralle | 96 |
| # | time | 2 | 4 | 0 | 0.333333333 | 1 | # | 10 | Baseline | paralle | 96 |
| # | interaction | 2 | 0 | 0 | 1 | 1 | # | 10 | Baseline | paralle | 96 |
| # | disease | 4 | 2 | 0 | 0.666666667 | 1 | # | 10 | Baseline | ancom | 96 |
| # | sex | 2 | 0 | 0 | 1 | 1 | # | 10 | Baseline | ancom | 96 |
| # | time | 2 | 2 | 0 | 0.5 | 1 | # | 10 | Baseline | ancom | 96 |
| # | interaction | 0 | 0 | 2 | 0 | 0 | # | 10 | Baseline | ancom | 96 |
| # | disease | 4 | 18 | 0 | 0.181818182 | 1 | # | 10 | Baseline | aldex2 | 96 |
| # | sex | 2 | 6 | 0 | 0.25 | 1 | # | 10 | Baseline | aldex2 | 96 |
| # | time | 2 | 13 | 0 | 0.133333333 | 1 | # | 10 | Baseline | aldex2 | 96 |
| # | interaction | 0 | 0 | 2 | 0 | 0 | # | 10 | Baseline | aldex2 | 96 |
| # | disease | 4 | 0 | 0 | 1 | 1 | # | 10 | Small | paralle | 96 |
| # | sex | 2 | 0 | 0 | 1 | 1 | # | 10 | Small | paralle | 96 |
| # | time | 2 | 0 | 0 | 1 | 1 | # | 10 | Small | paralle | 96 |
| # | interaction | 2 | 0 | 0 | 1 | 1 | # | 10 | Small | paralle | 96 |
| # | disease | 4 | 2 | 0 | 0.666666667 | 1 | # | 10 | Small | ancom | 96 |
| # | sex | 2 | 0 | 0 | 1 | 1 | # | 10 | Small | ancom | 96 |
| # | time | 2 | 2 | 0 | 0.5 | 1 | # | 10 | Small | ancom | 96 |
| # | interaction | 0 | 0 | 2 | 0 | 0 | # | 10 | Small | ancom | 96 |
| # | disease | 4 | 3 | 0 | 0.571428571 | 1 | # | 10 | Small | aldex2 | 96 |
| # | sex | 2 | 0 | 0 | 1 | 1 | # | 10 | Small | aldex2 | 96 |
| # | time | 2 | 2 | 0 | 0.5 | 1 | # | 10 | Small | aldex2 | 96 |
| # | interaction | 0 | 0 | 2 | 0 | 0 | # | 10 | Small | aldex2 | 96 |
| # | disease | 7 | 7 | 0 | 0.5 | 1 | # | 20 | Baseline | paralle | 96 |
| # | sex | 4 | 10 | 0 | 0.285714286 | 1 | # | 20 | Baseline | paralle | 96 |
| # | time | 5 | 12 | 0 | 0.294117647 | 1 | # | 20 | Baseline | paralle | 96 |
| # | interaction | 4 | 12 | 0 | 0.25 | 1 | # | 20 | Baseline | paralle | 96 |
| # | disease | 7 | 1 | 0 | 0.875 | 1 | # | 20 | Baseline | ancom | 96 |
| # | sex | 4 | 0 | 0 | 1 | 1 | # | 20 | Baseline | ancom | 96 |
| # | time | 5 | 2 | 0 | 0.714285714 | 1 | # | 20 | Baseline | ancom | 96 |
| # | interaction | 0 | 0 | 4 | 0 | 0 | # | 20 | Baseline | ancom | 96 |
| # | disease | 7 | 18 | 0 | 0.28 | 1 | # | 20 | Baseline | aldex2 | 96 |
| # | sex | 4 | 5 | 0 | 0.444444444 | 1 | # | 20 | Baseline | aldex2 | 96 |
| # | time | 5 | 16 | 0 | 0.238095238 | 1 | # | 20 | Baseline | aldex2 | 96 |
| # | interaction | 0 | 0 | 4 | 0 | 0 | # | 20 | Baseline | aldex2 | 96 |
| # | disease | 7 | 0 | 0 | 1 | 1 | # | 20 | Small | paralle | 96 |
| # | sex | 4 | 0 | 0 | 1 | 1 | # | 20 | Small | paralle | 96 |
| # | time | 5 | 0 | 0 | 1 | 1 | # | 20 | Small | paralle | 96 |
| # | interaction | 1 | 0 | 3 | 1 | 0.25 | # | 20 | Small | paralle | 96 |
| # | disease | 7 | 1 | 0 | 0.875 | 1 | # | 20 | Small | ancom | 96 |
| # | sex | 4 | 0 | 0 | 1 | 1 | # | 20 | Small | ancom | 96 |
| # | time | 5 | 3 | 0 | 0.625 | 1 | # | 20 | Small | ancom | 96 |
| # | interaction | 0 | 0 | 4 | 0 | 0 | # | 20 | Small | ancom | 96 |
| # | disease | 7 | 1 | 0 | 0.875 | 1 | # | 20 | Small | aldex2 | 96 |
| # | sex | 4 | 0 | 0 | 1 | 1 | # | 20 | Small | aldex2 | 96 |
| # | time | 5 | 2 | 0 | 0.714285714 | 1 | # | 20 | Small | aldex2 | 96 |
| # | interaction | 0 | 0 | 4 | 0 | 0 | # | 20 | Small | aldex2 | 96 |
| # | disease | 7 | 8 | 0 | 0.466666667 | 1 | # | 20 | Baseline | paralle | 96 |
| # | sex | 4 | 8 | 0 | 0.333333333 | 1 | # | 20 | Baseline | paralle | 96 |
| # | time | 5 | 9 | 0 | 0.357142857 | 1 | # | 20 | Baseline | paralle | 96 |
| # | interaction | 4 | 11 | 0 | 0.266666667 | 1 | # | 20 | Baseline | paralle | 96 |
| # | disease | 7 | 2 | 0 | 0.777777778 | 1 | # | 20 | Baseline | ancom | 96 |
| # | sex | 4 | 0 | 0 | 1 | 1 | # | 20 | Baseline | ancom | 96 |
| # | time | 5 | 3 | 0 | 0.625 | 1 | # | 20 | Baseline | ancom | 96 |
| # | interaction | 0 | 0 | 4 | 0 | 0 | # | 20 | Baseline | ancom | 96 |
| # | disease | 7 | 37 | 0 | 0.159090909 | 1 | # | 20 | Baseline | aldex2 | 96 |
| # | sex | 4 | 13 | 0 | 0.235294118 | 1 | # | 20 | Baseline | aldex2 | 96 |
| # | time | 5 | 21 | 0 | 0.192307692 | 1 | # | 20 | Baseline | aldex2 | 96 |
| # | interaction | 0 | 0 | 4 | 0 | 0 | # | 20 | Baseline | aldex2 | 96 |
| # | disease | 7 | 0 | 0 | 1 | 1 | # | 20 | Small | paralle | 96 |
| # | sex | 4 | 0 | 0 | 1 | 1 | # | 20 | Small | paralle | 96 |
| # | time | 5 | 0 | 0 | 1 | 1 | # | 20 | Small | paralle | 96 |
| # | interaction | 3 | 0 | 1 | 1 | 0.75 | # | 20 | Small | paralle | 96 |
| # | disease | 7 | 2 | 0 | 0.777777778 | 1 | # | 20 | Small | ancom | 96 |
| # | sex | 4 | 0 | 0 | 1 | 1 | # | 20 | Small | ancom | 96 |
| # | time | 5 | 3 | 0 | 0.625 | 1 | # | 20 | Small | ancom | 96 |
| # | interaction | 0 | 0 | 4 | 0 | 0 | # | 20 | Small | ancom | 96 |
| # | disease | 7 | 1 | 0 | 0.875 | 1 | # | 20 | Small | aldex2 | 96 |
| # | sex | 4 | 0 | 0 | 1 | 1 | # | 20 | Small | aldex2 | 96 |
| # | time | 5 | 2 | 0 | 0.714285714 | 1 | # | 20 | Small | aldex2 | 96 |
| # | interaction | 0 | 0 | 4 | 0 | 0 | # | 20 | Small | aldex2 | 96 |
| # | disease | 7 | 6 | 0 | 0.538461538 | 1 | # | 20 | Baseline | paralle | 96 |
| # | sex | 4 | 5 | 0 | 0.444444444 | 1 | # | 20 | Baseline | paralle | 96 |
| # | time | 5 | 10 | 0 | 0.333333333 | 1 | # | 20 | Baseline | paralle | 96 |
| # | interaction | 4 | 5 | 0 | 0.444444444 | 1 | # | 20 | Baseline | paralle | 96 |
| # | disease | 7 | 3 | 0 | 0.7 | 1 | # | 20 | Baseline | ancom | 96 |
| # | sex | 4 | 0 | 0 | 1 | 1 | # | 20 | Baseline | ancom | 96 |
| # | time | 5 | 4 | 0 | 0.555555556 | 1 | # | 20 | Baseline | ancom | 96 |
| # | interaction | 0 | 0 | 4 | 0 | 0 | # | 20 | Baseline | ancom | 96 |
| # | disease | 7 | 50 | 0 | 0.122807018 | 1 | # | 20 | Baseline | aldex2 | 96 |
| # | sex | 4 | 18 | 0 | 0.181818182 | 1 | # | 20 | Baseline | aldex2 | 96 |
| # | time | 5 | 33 | 0 | 0.131578947 | 1 | # | 20 | Baseline | aldex2 | 96 |
| # | interaction | 0 | 0 | 4 | 0 | 0 | # | 20 | Baseline | aldex2 | 96 |
| # | disease | 7 | 0 | 0 | 1 | 1 | # | 20 | Small | paralle | 96 |
| # | sex | 4 | 0 | 0 | 1 | 1 | # | 20 | Small | paralle | 96 |
| # | time | 5 | 0 | 0 | 1 | 1 | # | 20 | Small | paralle | 96 |
| # | interaction | 3 | 0 | 1 | 1 | 0.75 | # | 20 | Small | paralle | 96 |
| # | disease | 7 | 3 | 0 | 0.7 | 1 | # | 20 | Small | ancom | 96 |
| # | sex | 4 | 0 | 0 | 1 | 1 | # | 20 | Small | ancom | 96 |
| # | time | 5 | 4 | 0 | 0.555555556 | 1 | # | 20 | Small | ancom | 96 |
| # | interaction | 0 | 0 | 4 | 0 | 0 | # | 20 | Small | ancom | 96 |
| # | disease | 7 | 3 | 0 | 0.7 | 1 | # | 20 | Small | aldex2 | 96 |
| # | sex | 4 | 0 | 0 | 1 | 1 | # | 20 | Small | aldex2 | 96 |
| # | time | 5 | 4 | 0 | 0.555555556 | 1 | # | 20 | Small | aldex2 | 96 |
| # | interaction | 0 | 0 | 4 | 0 | 0 | # | 20 | Small | aldex2 | 96 |
| # | disease | 7 | 13 | 0 | 0.35 | 1 | # | 20 | Baseline | paralle | 96 |
| # | sex | 4 | 7 | 0 | 0.363636364 | 1 | # | 20 | Baseline | paralle | 96 |
| # | time | 5 | 8 | 0 | 0.384615385 | 1 | # | 20 | Baseline | paralle | 96 |
| # | interaction | 4 | 12 | 0 | 0.25 | 1 | # | 20 | Baseline | paralle | 96 |
| # | disease | 7 | 3 | 0 | 0.7 | 1 | # | 20 | Baseline | ancom | 96 |
| # | sex | 4 | 0 | 0 | 1 | 1 | # | 20 | Baseline | ancom | 96 |
| # | time | 5 | 4 | 0 | 0.555555556 | 1 | # | 20 | Baseline | ancom | 96 |
| # | interaction | 0 | 0 | 4 | 0 | 0 | # | 20 | Baseline | ancom | 96 |
| # | disease | 7 | 53 | 0 | 0.116666667 | 1 | # | 20 | Baseline | aldex2 | 96 |
| # | sex | 4 | 23 | 0 | 0.148148148 | 1 | # | 20 | Baseline | aldex2 | 96 |
| # | time | 5 | 50 | 0 | 0.090909091 | 1 | # | 20 | Baseline | aldex2 | 96 |
| # | interaction | 0 | 0 | 4 | 0 | 0 | # | 20 | Baseline | aldex2 | 96 |
| # | disease | 7 | 1 | 0 | 0.875 | 1 | # | 20 | Small | paralle | 96 |
| # | sex | 4 | 0 | 0 | 1 | 1 | # | 20 | Small | paralle | 96 |
| # | time | 5 | 0 | 0 | 1 | 1 | # | 20 | Small | paralle | 96 |
| # | interaction | 4 | 0 | 0 | 1 | 1 | # | 20 | Small | paralle | 96 |
| # | disease | 7 | 3 | 0 | 0.7 | 1 | # | 20 | Small | ancom | 96 |
| # | sex | 4 | 0 | 0 | 1 | 1 | # | 20 | Small | ancom | 96 |
| # | time | 5 | 4 | 0 | 0.555555556 | 1 | # | 20 | Small | ancom | 96 |
| # | interaction | 0 | 0 | 4 | 0 | 0 | # | 20 | Small | ancom | 96 |
| # | disease | 7 | 9 | 0 | 0.4375 | 1 | # | 20 | Small | aldex2 | 96 |
| # | sex | 4 | 0 | 0 | 1 | 1 | # | 20 | Small | aldex2 | 96 |
| # | time | 5 | 9 | 0 | 0.357142857 | 1 | # | 20 | Small | aldex2 | 96 |
| # | interaction | 0 | 0 | 4 | 0 | 0 | # | 20 | Small | aldex2 | 96 |
| # | disease | 4 | 12 | 0 | 0.25 | 1 | # | 10 | Baseline | paralle | 97 |
| # | sex | 2 | 12 | 0 | 0.142857143 | 1 | # | 10 | Baseline | paralle | 97 |
| # | time | 2 | 5 | 0 | 0.285714286 | 1 | # | 10 | Baseline | paralle | 97 |
| # | interaction | 2 | 8 | 0 | 0.2 | 1 | # | 10 | Baseline | paralle | 97 |
| # | disease | 4 | 0 | 0 | 1 | 1 | # | 10 | Baseline | ancom | 97 |
| # | sex | 2 | 0 | 0 | 1 | 1 | # | 10 | Baseline | ancom | 97 |
| # | time | 2 | 1 | 0 | 0.666666667 | 1 | # | 10 | Baseline | ancom | 97 |
| # | interaction | 0 | 0 | 2 | 0 | 0 | # | 10 | Baseline | ancom | 97 |
| # | disease | 4 | 10 | 0 | 0.285714286 | 1 | # | 10 | Baseline | aldex2 | 97 |
| # | sex | 2 | 8 | 0 | 0.2 | 1 | # | 10 | Baseline | aldex2 | 97 |
| # | time | 2 | 10 | 0 | 0.166666667 | 1 | # | 10 | Baseline | aldex2 | 97 |
| # | interaction | 0 | 0 | 2 | 0 | 0 | # | 10 | Baseline | aldex2 | 97 |
| # | disease | 4 | 0 | 0 | 1 | 1 | # | 10 | Small | paralle | 97 |
| # | sex | 2 | 1 | 0 | 0.666666667 | 1 | # | 10 | Small | paralle | 97 |
| # | time | 2 | 0 | 0 | 1 | 1 | # | 10 | Small | paralle | 97 |
| # | interaction | 2 | 0 | 0 | 1 | 1 | # | 10 | Small | paralle | 97 |
| # | disease | 4 | 0 | 0 | 1 | 1 | # | 10 | Small | ancom | 97 |
| # | sex | 2 | 0 | 0 | 1 | 1 | # | 10 | Small | ancom | 97 |
| # | time | 2 | 0 | 0 | 1 | 1 | # | 10 | Small | ancom | 97 |
| # | interaction | 0 | 0 | 2 | 0 | 0 | # | 10 | Small | ancom | 97 |
| # | disease | 4 | 2 | 0 | 0.666666667 | 1 | # | 10 | Small | aldex2 | 97 |
| # | sex | 2 | 0 | 0 | 1 | 1 | # | 10 | Small | aldex2 | 97 |
| # | time | 2 | 1 | 0 | 0.666666667 | 1 | # | 10 | Small | aldex2 | 97 |
| # | interaction | 0 | 0 | 2 | 0 | 0 | # | 10 | Small | aldex2 | 97 |
| # | disease | 4 | 9 | 0 | 0.307692308 | 1 | # | 10 | Baseline | paralle | 97 |
| # | sex | 2 | 8 | 0 | 0.2 | 1 | # | 10 | Baseline | paralle | 97 |
| # | time | 2 | 4 | 0 | 0.333333333 | 1 | # | 10 | Baseline | paralle | 97 |
| # | interaction | 2 | 4 | 0 | 0.333333333 | 1 | # | 10 | Baseline | paralle | 97 |
| # | disease | 4 | 2 | 0 | 0.666666667 | 1 | # | 10 | Baseline | ancom | 97 |
| # | sex | 2 | 0 | 0 | 1 | 1 | # | 10 | Baseline | ancom | 97 |
| # | time | 2 | 2 | 0 | 0.5 | 1 | # | 10 | Baseline | ancom | 97 |
| # | interaction | 0 | 0 | 2 | 0 | 0 | # | 10 | Baseline | ancom | 97 |
| # | disease | 4 | 16 | 0 | 0.2 | 1 | # | 10 | Baseline | aldex2 | 97 |
| # | sex | 2 | 9 | 0 | 0.181818182 | 1 | # | 10 | Baseline | aldex2 | 97 |
| # | time | 2 | 10 | 0 | 0.166666667 | 1 | # | 10 | Baseline | aldex2 | 97 |
| # | interaction | 0 | 0 | 2 | 0 | 0 | # | 10 | Baseline | aldex2 | 97 |
| # | disease | 4 | 0 | 0 | 1 | 1 | # | 10 | Small | paralle | 97 |
| # | sex | 2 | 0 | 0 | 1 | 1 | # | 10 | Small | paralle | 97 |
| # | time | 2 | 0 | 0 | 1 | 1 | # | 10 | Small | paralle | 97 |
| # | interaction | 2 | 0 | 0 | 1 | 1 | # | 10 | Small | paralle | 97 |
| # | disease | 4 | 2 | 0 | 0.666666667 | 1 | # | 10 | Small | ancom | 97 |
| # | sex | 2 | 0 | 0 | 1 | 1 | # | 10 | Small | ancom | 97 |
| # | time | 2 | 2 | 0 | 0.5 | 1 | # | 10 | Small | ancom | 97 |
| # | interaction | 0 | 0 | 2 | 0 | 0 | # | 10 | Small | ancom | 97 |
| # | disease | 4 | 2 | 0 | 0.666666667 | 1 | # | 10 | Small | aldex2 | 97 |
| # | sex | 2 | 0 | 0 | 1 | 1 | # | 10 | Small | aldex2 | 97 |
| # | time | 2 | 2 | 0 | 0.5 | 1 | # | 10 | Small | aldex2 | 97 |
| # | interaction | 0 | 0 | 2 | 0 | 0 | # | 10 | Small | aldex2 | 97 |
| # | disease | 4 | 8 | 0 | 0.333333333 | 1 | # | 10 | Baseline | paralle | 97 |
| # | sex | 2 | 8 | 0 | 0.2 | 1 | # | 10 | Baseline | paralle | 97 |
| # | time | 2 | 5 | 0 | 0.285714286 | 1 | # | 10 | Baseline | paralle | 97 |
| # | interaction | 2 | 1 | 0 | 0.666666667 | 1 | # | 10 | Baseline | paralle | 97 |
| # | disease | 4 | 2 | 0 | 0.666666667 | 1 | # | 10 | Baseline | ancom | 97 |
| # | sex | 2 | 0 | 0 | 1 | 1 | # | 10 | Baseline | ancom | 97 |
| # | time | 2 | 2 | 0 | 0.5 | 1 | # | 10 | Baseline | ancom | 97 |
| # | interaction | 0 | 0 | 2 | 0 | 0 | # | 10 | Baseline | ancom | 97 |
| # | disease | 4 | 18 | 0 | 0.181818182 | 1 | # | 10 | Baseline | aldex2 | 97 |
| # | sex | 2 | 8 | 0 | 0.2 | 1 | # | 10 | Baseline | aldex2 | 97 |
| # | time | 2 | 13 | 0 | 0.133333333 | 1 | # | 10 | Baseline | aldex2 | 97 |
| # | interaction | 0 | 0 | 2 | 0 | 0 | # | 10 | Baseline | aldex2 | 97 |
| # | disease | 4 | 0 | 0 | 1 | 1 | # | 10 | Small | paralle | 97 |
| # | sex | 2 | 0 | 0 | 1 | 1 | # | 10 | Small | paralle | 97 |
| # | time | 2 | 0 | 0 | 1 | 1 | # | 10 | Small | paralle | 97 |
| # | interaction | 2 | 1 | 0 | 0.666666667 | 1 | # | 10 | Small | paralle | 97 |
| # | disease | 4 | 2 | 0 | 0.666666667 | 1 | # | 10 | Small | ancom | 97 |
| # | sex | 2 | 0 | 0 | 1 | 1 | # | 10 | Small | ancom | 97 |
| # | time | 2 | 2 | 0 | 0.5 | 1 | # | 10 | Small | ancom | 97 |
| # | interaction | 0 | 0 | 2 | 0 | 0 | # | 10 | Small | ancom | 97 |
| # | disease | 4 | 2 | 0 | 0.666666667 | 1 | # | 10 | Small | aldex2 | 97 |
| # | sex | 2 | 0 | 0 | 1 | 1 | # | 10 | Small | aldex2 | 97 |
| # | time | 2 | 2 | 0 | 0.5 | 1 | # | 10 | Small | aldex2 | 97 |
| # | interaction | 0 | 0 | 2 | 0 | 0 | # | 10 | Small | aldex2 | 97 |
| # | disease | 4 | 10 | 0 | 0.285714286 | 1 | # | 10 | Baseline | paralle | 97 |
| # | sex | 2 | 7 | 0 | 0.222222222 | 1 | # | 10 | Baseline | paralle | 97 |
| # | time | 2 | 5 | 0 | 0.285714286 | 1 | # | 10 | Baseline | paralle | 97 |
| # | interaction | 2 | 3 | 0 | 0.4 | 1 | # | 10 | Baseline | paralle | 97 |
| # | disease | 4 | 2 | 0 | 0.666666667 | 1 | # | 10 | Baseline | ancom | 97 |
| # | sex | 2 | 0 | 0 | 1 | 1 | # | 10 | Baseline | ancom | 97 |
| # | time | 2 | 2 | 0 | 0.5 | 1 | # | 10 | Baseline | ancom | 97 |
| # | interaction | 0 | 0 | 2 | 0 | 0 | # | 10 | Baseline | ancom | 97 |
| # | disease | 4 | 19 | 0 | 0.173913043 | 1 | # | 10 | Baseline | aldex2 | 97 |
| # | sex | 2 | 9 | 0 | 0.181818182 | 1 | # | 10 | Baseline | aldex2 | 97 |
| # | time | 2 | 12 | 0 | 0.142857143 | 1 | # | 10 | Baseline | aldex2 | 97 |
| # | interaction | 0 | 0 | 2 | 0 | 0 | # | 10 | Baseline | aldex2 | 97 |
| # | disease | 4 | 0 | 0 | 1 | 1 | # | 10 | Small | paralle | 97 |
| # | sex | 2 | 1 | 0 | 0.666666667 | 1 | # | 10 | Small | paralle | 97 |
| # | time | 2 | 0 | 0 | 1 | 1 | # | 10 | Small | paralle | 97 |
| # | interaction | 2 | 0 | 0 | 1 | 1 | # | 10 | Small | paralle | 97 |
| # | disease | 4 | 2 | 0 | 0.666666667 | 1 | # | 10 | Small | ancom | 97 |
| # | sex | 2 | 0 | 0 | 1 | 1 | # | 10 | Small | ancom | 97 |
| # | time | 2 | 2 | 0 | 0.5 | 1 | # | 10 | Small | ancom | 97 |
| # | interaction | 0 | 0 | 2 | 0 | 0 | # | 10 | Small | ancom | 97 |
| # | disease | 4 | 2 | 0 | 0.666666667 | 1 | # | 10 | Small | aldex2 | 97 |
| # | sex | 2 | 0 | 0 | 1 | 1 | # | 10 | Small | aldex2 | 97 |
| # | time | 2 | 3 | 0 | 0.4 | 1 | # | 10 | Small | aldex2 | 97 |
| # | interaction | 0 | 0 | 2 | 0 | 0 | # | 10 | Small | aldex2 | 97 |
| # | disease | 7 | 12 | 0 | 0.368421053 | 1 | # | 20 | Baseline | paralle | 97 |
| # | sex | 4 | 6 | 0 | 0.4 | 1 | # | 20 | Baseline | paralle | 97 |
| # | time | 5 | 9 | 0 | 0.357142857 | 1 | # | 20 | Baseline | paralle | 97 |
| # | interaction | 4 | 7 | 0 | 0.363636364 | 1 | # | 20 | Baseline | paralle | 97 |

| | | | | | | | | | | | |
|---|---|---|---|---|---|---|---|---|---|---|---|
| # | disease | 7 | 0 | 0 | 1 | 1 | # | 20 | Baseline | ancom | 97 |
| # | sex | 4 | 0 | 0 | 1 | 1 | # | 20 | Baseline | ancom | 97 |
| # | time | 5 | 0 | 0 | 1 | 1 | # | 20 | Baseline | ancom | 97 |
| # | interaction | 0 | 0 | 4 | 0 | 0 | # | 20 | Baseline | ancom | 97 |
| # | disease | 7 | 22 | 0 | 0.24137931 | 1 | # | 20 | Baseline | aldex2 | 97 |
| # | sex | 4 | 12 | 0 | 0.25 | 1 | # | 20 | Baseline | aldex2 | 97 |
| # | time | 5 | 17 | 0 | 0.227272727 | 1 | # | 20 | Baseline | aldex2 | 97 |
| # | interaction | 0 | 0 | 4 | 0 | 0 | # | 20 | Baseline | aldex2 | 97 |
| # | disease | 7 | 0 | 0 | 1 | 1 | # | 20 | Small | paralla | 97 |
| # | sex | 4 | 0 | 0 | 1 | 1 | # | 20 | Small | paralla | 97 |
| # | time | 5 | 2 | 0 | 0.714285714 | 1 | # | 20 | Small | paralla | 97 |
| # | interaction | 1 | 0 | 3 | 1 | 0.25 | # | 20 | Small | paralla | 97 |
| # | disease | 7 | 0 | 0 | 1 | 1 | # | 20 | Small | ancom | 97 |
| # | sex | 4 | 0 | 0 | 1 | 1 | # | 20 | Small | ancom | 97 |
| # | time | 5 | 0 | 0 | 1 | 1 | # | 20 | Small | ancom | 97 |
| # | interaction | 0 | 0 | 4 | 0 | 0 | # | 20 | Small | ancom | 97 |
| # | disease | 7 | 0 | 0 | 1 | 1 | # | 20 | Small | aldex2 | 97 |
| # | sex | 4 | 0 | 0 | 1 | 1 | # | 20 | Small | aldex2 | 97 |
| # | time | 5 | 2 | 0 | 0.714285714 | 1 | # | 20 | Small | aldex2 | 97 |
| # | interaction | 0 | 0 | 4 | 0 | 0 | # | 20 | Small | aldex2 | 97 |
| # | disease | 7 | 18 | 0 | 0.28 | 1 | # | 20 | Baseline | paralla | 97 |
| # | sex | 4 | 5 | 0 | 0.444444444 | 1 | # | 20 | Baseline | paralla | 97 |
| # | time | 5 | 11 | 0 | 0.3125 | 1 | # | 20 | Baseline | paralla | 97 |
| # | interaction | 4 | 9 | 0 | 0.307692308 | 1 | # | 20 | Baseline | paralla | 97 |
| # | disease | 7 | 2 | 0 | 0.777777778 | 1 | # | 20 | Baseline | ancom | 97 |
| # | sex | 4 | 0 | 0 | 1 | 1 | # | 20 | Baseline | ancom | 97 |
| # | time | 5 | 3 | 0 | 0.625 | 1 | # | 20 | Baseline | ancom | 97 |
| # | interaction | 0 | 0 | 4 | 0 | 0 | # | 20 | Baseline | ancom | 97 |
| # | disease | 7 | 35 | 0 | 0.166666667 | 1 | # | 20 | Baseline | aldex2 | 97 |
| # | sex | 4 | 13 | 0 | 0.235294118 | 1 | # | 20 | Baseline | aldex2 | 97 |
| # | time | 5 | 23 | 0 | 0.178571429 | 1 | # | 20 | Baseline | aldex2 | 97 |
| # | interaction | 0 | 0 | 4 | 0 | 0 | # | 20 | Baseline | aldex2 | 97 |
| # | disease | 7 | 0 | 0 | 1 | 1 | # | 20 | Small | paralla | 97 |
| # | sex | 4 | 0 | 0 | 1 | 1 | # | 20 | Small | paralla | 97 |
| # | time | 5 | 0 | 0 | 1 | 1 | # | 20 | Small | paralla | 97 |
| # | interaction | 2 | 0 | 2 | 1 | 0.5 | # | 20 | Small | paralla | 97 |
| # | disease | 7 | 0 | 0 | 1 | 1 | # | 20 | Small | ancom | 97 |
| # | sex | 4 | 0 | 0 | 1 | 1 | # | 20 | Small | ancom | 97 |
| # | time | 5 | 4 | 0 | 0.555555556 | 1 | # | 20 | Small | ancom | 97 |
| # | interaction | 0 | 0 | 4 | 0 | 0 | # | 20 | Small | ancom | 97 |
| # | disease | 7 | 3 | 0 | 0.7 | 1 | # | 20 | Small | aldex2 | 97 |
| # | sex | 4 | 0 | 0 | 1 | 1 | # | 20 | Small | aldex2 | 97 |
| # | time | 5 | 3 | 0 | 0.625 | 1 | # | 20 | Small | aldex2 | 97 |
| # | interaction | 0 | 0 | 4 | 0 | 0 | # | 20 | Small | aldex2 | 97 |
| # | disease | 7 | 15 | 0 | 0.318181818 | 1 | # | 20 | Baseline | paralla | 97 |
| # | sex | 4 | 9 | 0 | 0.307692308 | 1 | # | 20 | Baseline | paralla | 97 |
| # | time | 5 | 7 | 0 | 0.416666667 | 1 | # | 20 | Baseline | paralla | 97 |
| # | interaction | 4 | 6 | 0 | 0.4 | 1 | # | 20 | Baseline | paralla | 97 |
| # | disease | 7 | 3 | 0 | 0.7 | 1 | # | 20 | Baseline | ancom | 97 |
| # | sex | 4 | 0 | 0 | 1 | 1 | # | 20 | Baseline | ancom | 97 |
| # | time | 5 | 4 | 0 | 0.555555556 | 1 | # | 20 | Baseline | ancom | 97 |
| # | interaction | 0 | 0 | 4 | 0 | 0 | # | 20 | Baseline | ancom | 97 |
| # | disease | 7 | 52 | 0 | 0.118644068 | 1 | # | 20 | Baseline | aldex2 | 97 |
| # | sex | 4 | 19 | 0 | 0.173913043 | 1 | # | 20 | Baseline | aldex2 | 97 |
| # | time | 5 | 29 | 0 | 0.147058824 | 1 | # | 20 | Baseline | aldex2 | 97 |
| # | interaction | 0 | 0 | 4 | 0 | 0 | # | 20 | Baseline | aldex2 | 97 |
| # | disease | 7 | 0 | 0 | 1 | 1 | # | 20 | Small | paralla | 97 |
| # | sex | 4 | 0 | 0 | 1 | 1 | # | 20 | Small | paralla | 97 |
| # | time | 5 | 0 | 0 | 1 | 1 | # | 20 | Small | paralla | 97 |
| # | interaction | 3 | 0 | 1 | 1 | 0.75 | # | 20 | Small | paralla | 97 |
| # | disease | 7 | 2 | 0 | 0.777777778 | 1 | # | 20 | Small | ancom | 97 |
| # | sex | 4 | 0 | 0 | 1 | 1 | # | 20 | Small | ancom | 97 |
| # | time | 5 | 4 | 0 | 0.555555556 | 1 | # | 20 | Small | ancom | 97 |
| # | interaction | 0 | 0 | 4 | 0 | 0 | # | 20 | Small | ancom | 97 |
| # | disease | 7 | 6 | 0 | 0.538461538 | 1 | # | 20 | Small | aldex2 | 97 |
| # | sex | 4 | 0 | 0 | 1 | 1 | # | 20 | Small | aldex2 | 97 |
| # | time | 5 | 5 | 0 | 0.5 | 1 | # | 20 | Small | aldex2 | 97 |
| # | interaction | 0 | 0 | 4 | 0 | 0 | # | 20 | Small | aldex2 | 97 |
| # | disease | 7 | 20 | 0 | 0.259259259 | 1 | # | 20 | Baseline | paralla | 97 |
| # | sex | 4 | 6 | 0 | 0.4 | 1 | # | 20 | Baseline | paralla | 97 |
| # | time | 5 | 9 | 0 | 0.357142857 | 1 | # | 20 | Baseline | paralla | 97 |
| # | interaction | 4 | 5 | 0 | 0.444444444 | 1 | # | 20 | Baseline | paralla | 97 |
| # | disease | 7 | 3 | 0 | 0.7 | 1 | # | 20 | Baseline | ancom | 97 |
| # | sex | 4 | 0 | 0 | 1 | 1 | # | 20 | Baseline | ancom | 97 |
| # | time | 5 | 4 | 0 | 0.555555556 | 1 | # | 20 | Baseline | ancom | 97 |
| # | interaction | 0 | 0 | 4 | 0 | 0 | # | 20 | Baseline | ancom | 97 |
| # | disease | 7 | 63 | 0 | 0.1 | 1 | # | 20 | Baseline | aldex2 | 97 |
| # | sex | 4 | 23 | 0 | 0.148148148 | 1 | # | 20 | Baseline | aldex2 | 97 |
| # | time | 5 | 40 | 0 | 0.111111111 | 1 | # | 20 | Baseline | aldex2 | 97 |
| # | interaction | 0 | 0 | 4 | 0 | 0 | # | 20 | Baseline | aldex2 | 97 |
| # | disease | 7 | 0 | 0 | 1 | 1 | # | 20 | Small | paralla | 97 |
| # | sex | 4 | 0 | 0 | 1 | 1 | # | 20 | Small | paralla | 97 |
| # | time | 5 | 0 | 0 | 1 | 1 | # | 20 | Small | paralla | 97 |
| # | interaction | 3 | 0 | 1 | 1 | 0.75 | # | 20 | Small | paralla | 97 |
| # | disease | 7 | 3 | 0 | 0.7 | 1 | # | 20 | Small | ancom | 97 |
| # | sex | 4 | 0 | 0 | 1 | 1 | # | 20 | Small | ancom | 97 |
| # | time | 5 | 4 | 0 | 0.555555556 | 1 | # | 20 | Small | ancom | 97 |
| # | interaction | 0 | 0 | 4 | 0 | 0 | # | 20 | Small | ancom | 97 |
| # | disease | 7 | 9 | 0 | 0.4375 | 1 | # | 20 | Small | aldex2 | 97 |
| # | sex | 4 | 0 | 0 | 1 | 1 | # | 20 | Small | aldex2 | 97 |
| # | time | 5 | 7 | 0 | 0.416666667 | 1 | # | 20 | Small | aldex2 | 97 |
| # | interaction | 0 | 0 | 4 | 0 | 0 | # | 20 | Small | aldex2 | 97 |
| # | disease | 4 | 7 | 0 | 0.363636364 | 1 | # | 10 | Baseline | paralla | 98 |
| # | sex | 2 | 5 | 0 | 0.285714286 | 1 | # | 10 | Baseline | paralla | 98 |
| # | time | 2 | 9 | 0 | 0.181818182 | 1 | # | 10 | Baseline | paralla | 98 |
| # | interaction | 2 | 11 | 0 | 0.153846154 | 1 | # | 10 | Baseline | paralla | 98 |
| # | disease | 4 | 0 | 0 | 1 | 1 | # | 10 | Baseline | ancom | 98 |
| # | sex | 2 | 0 | 0 | 1 | 1 | # | 10 | Baseline | ancom | 98 |
| # | time | 2 | 0 | 0 | 1 | 1 | # | 10 | Baseline | ancom | 98 |
| # | interaction | 0 | 0 | 2 | 0 | 0 | # | 10 | Baseline | ancom | 98 |
| # | disease | 4 | 11 | 0 | 0.266666667 | 1 | # | 10 | Baseline | aldex2 | 98 |
| # | sex | 2 | 8 | 0 | 0.2 | 1 | # | 10 | Baseline | aldex2 | 98 |
| # | time | 2 | 8 | 0 | 0.2 | 1 | # | 10 | Baseline | aldex2 | 98 |
| # | interaction | 0 | 0 | 2 | 0 | 0 | # | 10 | Baseline | aldex2 | 98 |
| # | disease | 4 | 0 | 0 | 1 | 1 | # | 10 | Small | paralla | 98 |
| # | sex | 2 | 0 | 0 | 1 | 1 | # | 10 | Small | paralla | 98 |
| # | time | 2 | 0 | 0 | 1 | 1 | # | 10 | Small | paralla | 98 |
| # | interaction | 1 | 1 | 1 | 0.5 | 0.5 | # | 10 | Small | paralla | 98 |
| # | disease | 4 | 0 | 0 | 1 | 1 | # | 10 | Small | ancom | 98 |
| # | sex | 2 | 0 | 0 | 1 | 1 | # | 10 | Small | ancom | 98 |
| # | time | 2 | 0 | 0 | 1 | 1 | # | 10 | Small | ancom | 98 |
| # | interaction | 0 | 0 | 2 | 0 | 0 | # | 10 | Small | ancom | 98 |
| # | disease | 4 | 1 | 0 | 0.8 | 1 | # | 10 | Small | aldex2 | 98 |
| # | sex | 2 | 0 | 0 | 1 | 1 | # | 10 | Small | aldex2 | 98 |
| # | time | 2 | 0 | 0 | 1 | 1 | # | 10 | Small | aldex2 | 98 |
| # | interaction | 0 | 0 | 2 | 0 | 0 | # | 10 | Small | aldex2 | 98 |
| # | disease | 4 | 7 | 0 | 0.363636364 | 1 | # | 10 | Baseline | paralla | 98 |
| # | sex | 2 | 4 | 0 | 0.333333333 | 1 | # | 10 | Baseline | paralla | 98 |
| # | time | 2 | 8 | 0 | 0.2 | 1 | # | 10 | Baseline | paralla | 98 |
| # | interaction | 2 | 4 | 0 | 0.333333333 | 1 | # | 10 | Baseline | paralla | 98 |
| # | disease | 4 | 2 | 0 | 0.666666667 | 1 | # | 10 | Baseline | ancom | 98 |
| # | sex | 2 | 0 | 0 | 1 | 1 | # | 10 | Baseline | ancom | 98 |
| # | time | 2 | 2 | 0 | 0.5 | 1 | # | 10 | Baseline | ancom | 98 |
| # | interaction | 0 | 0 | 2 | 0 | 0 | # | 10 | Baseline | ancom | 98 |
| # | disease | 4 | 14 | 0 | 0.222222222 | 1 | # | 10 | Baseline | aldex2 | 98 |
| # | sex | 2 | 7 | 0 | 0.222222222 | 1 | # | 10 | Baseline | aldex2 | 98 |
| # | time | 2 | 11 | 0 | 0.153846154 | 1 | # | 10 | Baseline | aldex2 | 98 |
| # | interaction | 0 | 0 | 2 | 0 | 0 | # | 10 | Baseline | aldex2 | 98 |
| # | disease | 4 | 0 | 0 | 1 | 1 | # | 10 | Small | paralla | 98 |
| # | sex | 2 | 0 | 0 | 1 | 1 | # | 10 | Small | paralla | 98 |
| # | time | 2 | 2 | 0 | 0.5 | 1 | # | 10 | Small | paralla | 98 |
| # | interaction | 2 | 1 | 0 | 0.666666667 | 1 | # | 10 | Small | paralla | 98 |
| # | disease | 4 | 2 | 0 | 0.666666667 | 1 | # | 10 | Small | ancom | 98 |
| # | sex | 2 | 0 | 0 | 1 | 1 | # | 10 | Small | ancom | 98 |
| # | time | 2 | 2 | 0 | 0.5 | 1 | # | 10 | Small | ancom | 98 |
| # | interaction | 0 | 0 | 2 | 0 | 0 | # | 10 | Small | ancom | 98 |
| # | disease | 4 | 1 | 0 | 0.8 | 1 | # | 10 | Small | aldex2 | 98 |
| # | sex | 2 | 0 | 0 | 1 | 1 | # | 10 | Small | aldex2 | 98 |
| # | time | 2 | 2 | 0 | 0.5 | 1 | # | 10 | Small | aldex2 | 98 |
| # | interaction | 0 | 0 | 2 | 0 | 0 | # | 10 | Small | aldex2 | 98 |
| # | disease | 4 | 2 | 0 | 0.666666667 | 1 | # | 10 | Baseline | paralla | 98 |
| # | sex | 2 | 6 | 0 | 0.25 | 1 | # | 10 | Baseline | paralla | 98 |
| # | time | 2 | 6 | 0 | 0.25 | 1 | # | 10 | Baseline | paralla | 98 |
| # | interaction | 2 | 7 | 0 | 0.222222222 | 1 | # | 10 | Baseline | paralla | 98 |
| # | disease | 4 | 2 | 0 | 0.666666667 | 1 | # | 10 | Baseline | ancom | 98 |
| # | sex | 2 | 0 | 0 | 1 | 1 | # | 10 | Baseline | ancom | 98 |
| # | time | 2 | 2 | 0 | 0.5 | 1 | # | 10 | Baseline | ancom | 98 |
| # | interaction | 0 | 0 | 2 | 0 | 0 | # | 10 | Baseline | ancom | 98 |
| # | disease | 4 | 19 | 0 | 0.173913043 | 1 | # | 10 | Baseline | aldex2 | 98 |
| # | sex | 2 | 8 | 0 | 0.2 | 1 | # | 10 | Baseline | aldex2 | 98 |
| # | time | 2 | 7 | 0 | 0.222222222 | 1 | # | 10 | Baseline | aldex2 | 98 |
| # | interaction | 0 | 0 | 2 | 0 | 0 | # | 10 | Baseline | aldex2 | 98 |
| # | disease | 4 | 0 | 0 | 1 | 1 | # | 10 | Small | paralla | 98 |
| # | sex | 2 | 0 | 0 | 1 | 1 | # | 10 | Small | paralla | 98 |
| # | time | 2 | 1 | 0 | 0.666666667 | 1 | # | 10 | Small | paralla | 98 |
| # | interaction | 2 | 1 | 0 | 0.666666667 | 1 | # | 10 | Small | paralla | 98 |
| # | disease | 4 | 2 | 0 | 0.666666667 | 1 | # | 10 | Small | ancom | 98 |
| # | sex | 2 | 0 | 0 | 1 | 1 | # | 10 | Small | ancom | 98 |
| # | time | 2 | 2 | 0 | 0.5 | 1 | # | 10 | Small | ancom | 98 |
| # | interaction | 0 | 0 | 2 | 0 | 0 | # | 10 | Small | ancom | 98 |
| # | disease | 4 | 2 | 0 | 0.666666667 | 1 | # | 10 | Small | aldex2 | 98 |
| # | sex | 2 | 0 | 0 | 1 | 1 | # | 10 | Small | aldex2 | 98 |
| # | time | 2 | 2 | 0 | 0.5 | 1 | # | 10 | Small | aldex2 | 98 |
| # | interaction | 0 | 0 | 2 | 0 | 0 | # | 10 | Small | aldex2 | 98 |
| # | disease | 4 | 4 | 0 | 0.5 | 1 | # | 10 | Baseline | paralla | 98 |
| # | sex | 2 | 3 | 0 | 0.4 | 1 | # | 10 | Baseline | paralla | 98 |
| # | time | 2 | 4 | 0 | 0.333333333 | 1 | # | 10 | Baseline | paralla | 98 |
| # | interaction | 2 | 5 | 0 | 0.285714286 | 1 | # | 10 | Baseline | paralla | 98 |
| # | disease | 4 | 2 | 0 | 0.666666667 | 1 | # | 10 | Baseline | ancom | 98 |
| # | sex | 2 | 0 | 0 | 1 | 1 | # | 10 | Baseline | ancom | 98 |
| # | time | 2 | 2 | 0 | 0.5 | 1 | # | 10 | Baseline | ancom | 98 |
| # | interaction | 0 | 0 | 2 | 0 | 0 | # | 10 | Baseline | ancom | 98 |
| # | disease | 4 | 24 | 0 | 0.142857143 | 1 | # | 10 | Baseline | aldex2 | 98 |
| # | sex | 2 | 8 | 0 | 0.2 | 1 | # | 10 | Baseline | aldex2 | 98 |
| # | time | 2 | 14 | 0 | 0.125 | 1 | # | 10 | Baseline | aldex2 | 98 |
| # | interaction | 0 | 0 | 2 | 0 | 0 | # | 10 | Baseline | aldex2 | 98 |
| # | disease | 4 | 0 | 0 | 1 | 1 | # | 10 | Small | paralla | 98 |
| # | sex | 2 | 0 | 0 | 1 | 1 | # | 10 | Small | paralla | 98 |
| # | time | 2 | 1 | 0 | 0.666666667 | 1 | # | 10 | Small | paralla | 98 |
| # | interaction | 2 | 1 | 0 | 0.666666667 | 1 | # | 10 | Small | paralla | 98 |
| # | disease | 4 | 2 | 0 | 0.666666667 | 1 | # | 10 | Small | ancom | 98 |
| # | sex | 2 | 0 | 0 | 1 | 1 | # | 10 | Small | ancom | 98 |
| # | time | 2 | 2 | 0 | 0.5 | 1 | # | 10 | Small | ancom | 98 |
| # | interaction | 0 | 0 | 2 | 0 | 0 | # | 10 | Small | ancom | 98 |
| # | disease | 4 | 2 | 0 | 0.666666667 | 1 | # | 10 | Small | aldex2 | 98 |
| # | sex | 2 | 0 | 0 | 1 | 1 | # | 10 | Small | aldex2 | 98 |
| # | time | 2 | 2 | 0 | 0.5 | 1 | # | 10 | Small | aldex2 | 98 |
| # | interaction | 0 | 0 | 2 | 0 | 0 | # | 10 | Small | aldex2 | 98 |
| # | disease | 7 | 14 | 0 | 0.333333333 | 1 | # | 20 | Baseline | paralla | 98 |
| # | sex | 4 | 10 | 0 | 0.285714286 | 1 | # | 20 | Baseline | paralla | 98 |
| # | time | 5 | 3 | 0 | 0.625 | 1 | # | 20 | Baseline | paralla | 98 |
| # | interaction | 4 | 5 | 0 | 0.444444444 | 1 | # | 20 | Baseline | paralla | 98 |
| # | disease | 7 | 0 | 0 | 1 | 1 | # | 20 | Baseline | ancom | 98 |
| # | sex | 4 | 0 | 0 | 1 | 1 | # | 20 | Baseline | ancom | 98 |
| # | time | 5 | 0 | 0 | 1 | 1 | # | 20 | Baseline | ancom | 98 |
| # | interaction | 0 | 0 | 4 | 0 | 0 | # | 20 | Baseline | ancom | 98 |
| # | disease | 7 | 13 | 0 | 0.35 | 1 | # | 20 | Baseline | aldex2 | 98 |
| # | sex | 4 | 5 | 0 | 0.444444444 | 1 | # | 20 | Baseline | aldex2 | 98 |
| # | time | 5 | 17 | 0 | 0.227272727 | 1 | # | 20 | Baseline | aldex2 | 98 |
| # | interaction | 0 | 0 | 4 | 0 | 0 | # | 20 | Baseline | aldex2 | 98 |
| # | disease | 7 | 1 | 0 | 0.875 | 1 | # | 20 | Small | paralla | 98 |
| # | sex | 4 | 0 | 0 | 1 | 1 | # | 20 | Small | paralla | 98 |
| # | time | 5 | 0 | 0 | 1 | 1 | # | 20 | Small | paralla | 98 |
| # | interaction | 2 | 0 | 2 | 1 | 0.5 | # | 20 | Small | paralla | 98 |
| # | disease | 7 | 0 | 0 | 1 | 1 | # | 20 | Small | ancom | 98 |
| # | sex | 4 | 0 | 0 | 1 | 1 | # | 20 | Small | ancom | 98 |
| # | time | 5 | 1 | 0 | 0.833333333 | 1 | # | 20 | Small | ancom | 98 |
| # | interaction | 0 | 0 | 4 | 0 | 0 | # | 20 | Small | ancom | 98 |
| # | disease | 7 | 1 | 0 | 0.875 | 1 | # | 20 | Small | aldex2 | 98 |
| # | sex | 4 | 0 | 0 | 1 | 1 | # | 20 | Small | aldex2 | 98 |
| # | time | 5 | 1 | 0 | 0.833333333 | 1 | # | 20 | Small | aldex2 | 98 |
| # | interaction | 0 | 0 | 4 | 0 | 0 | # | 20 | Small | aldex2 | 98 |
| # | disease | 7 | 13 | 0 | 0.35 | 1 | # | 20 | Baseline | paralla | 98 |
| # | sex | 4 | 7 | 0 | 0.363636364 | 1 | # | 20 | Baseline | paralla | 98 |
| # | time | 5 | 4 | 0 | 0.555555556 | 1 | # | 20 | Baseline | paralla | 98 |
| # | interaction | 4 | 6 | 0 | 0.4 | 1 | # | 20 | Baseline | paralla | 98 |
| # | disease | 7 | 0 | 0 | 1 | 1 | # | 20 | Baseline | ancom | 98 |
| # | sex | 4 | 0 | 0 | 1 | 1 | # | 20 | Baseline | ancom | 98 |
| # | time | 5 | 3 | 0 | 0.625 | 1 | # | 20 | Baseline | ancom | 98 |
| # | interaction | 0 | 0 | 4 | 0 | 0 | # | 20 | Baseline | ancom | 98 |
| # | disease | 7 | 41 | 0 | 0.145833333 | 1 | # | 20 | Baseline | aldex2 | 98 |
| # | sex | 4 | 7 | 0 | 0.363636364 | 1 | # | 20 | Baseline | aldex2 | 98 |
| # | time | 5 | 21 | 0 | 0.192307692 | 1 | # | 20 | Baseline | aldex2 | 98 |
| # | interaction | 0 | 0 | 4 | 0 | 0 | # | 20 | Baseline | aldex2 | 98 |
| # | disease | 7 | 0 | 0 | 1 | 1 | # | 20 | Small | paralla | 98 |
| # | sex | 4 | 0 | 0 | 1 | 1 | # | 20 | Small | paralla | 98 |
| # | time | 5 | 0 | 0 | 1 | 1 | # | 20 | Small | paralla | 98 |
| # | interaction | 4 | 0 | 0 | 1 | 1 | # | 20 | Small | paralla | 98 |
| # | disease | 7 | 0 | 0 | 1 | 1 | # | 20 | Small | ancom | 98 |
| # | sex | 4 | 0 | 0 | 1 | 1 | # | 20 | Small | ancom | 98 |
| # | time | 5 | 3 | 0 | 0.625 | 1 | # | 20 | Small | ancom | 98 |
| # | interaction | 0 | 0 | 4 | 0 | 0 | # | 20 | Small | ancom | 98 |
| # | disease | 7 | 2 | 0 | 0.777777778 | 1 | # | 20 | Small | aldex2 | 98 |
| # | sex | 4 | 1 | 0 | 0.8 | 1 | # | 20 | Small | aldex2 | 98 |
| # | time | 5 | 5 | 0 | 0.5 | 1 | # | 20 | Small | aldex2 | 98 |
| # | interaction | 0 | 0 | 4 | 0 | 0 | # | 20 | Small | aldex2 | 98 |
| # | disease | 7 | 12 | 0 | 0.368421053 | 1 | # | 20 | Baseline | paralla | 98 |
| # | sex | 4 | 4 | 0 | 0.5 | 1 | # | 20 | Baseline | paralla | 98 |
| # | time | 5 | 4 | 0 | 0.555555556 | 1 | # | 20 | Baseline | paralla | 98 |
| # | interaction | 4 | 6 | 0 | 0.4 | 1 | # | 20 | Baseline | paralla | 98 |
| # | disease | 7 | 4 | 0 | 0.636363636 | 1 | # | 20 | Baseline | ancom | 98 |
| # | sex | 4 | 0 | 0 | 1 | 1 | # | 20 | Baseline | ancom | 98 |
| # | time | 5 | 4 | 0 | 0.555555556 | 1 | # | 20 | Baseline | ancom | 98 |
| # | interaction | 0 | 0 | 4 | 0 | 0 | # | 20 | Baseline | ancom | 98 |
| # | disease | 7 | 57 | 0 | 0.109375 | 1 | # | 20 | Baseline | aldex2 | 98 |
| # | sex | 4 | 13 | 0 | 0.235294118 | 1 | # | 20 | Baseline | aldex2 | 98 |
| # | time | 5 | 35 | 0 | 0.125 | 1 | # | 20 | Baseline | aldex2 | 98 |
| # | interaction | 0 | 0 | 4 | 0 | 0 | # | 20 | Baseline | aldex2 | 98 |
| # | disease | 7 | 0 | 0 | 1 | 1 | # | 20 | Small | paralla | 98 |
| # | sex | 4 | 0 | 0 | 1 | 1 | # | 20 | Small | paralla | 98 |
| # | time | 5 | 0 | 0 | 1 | 1 | # | 20 | Small | paralla | 98 |
| # | interaction | 4 | 0 | 0 | 1 | 1 | # | 20 | Small | paralla | 98 |
| # | disease | 7 | 4 | 0 | 0.636363636 | 1 | # | 20 | Small | ancom | 98 |
| # | sex | 4 | 0 | 0 | 1 | 1 | # | 20 | Small | ancom | 98 |
| # | time | 5 | 3 | 0 | 0.625 | 1 | # | 20 | Small | ancom | 98 |
| # | interaction | 0 | 0 | 4 | 0 | 0 | # | 20 | Small | ancom | 98 |
| # | disease | 7 | 3 | 0 | 0.7 | 1 | # | 20 | Small | aldex2 | 98 |
| # | sex | 4 | 1 | 0 | 0.8 | 1 | # | 20 | Small | aldex2 | 98 |
| # | time | 5 | 7 | 0 | 0.416666667 | 1 | # | 20 | Small | aldex2 | 98 |
| # | interaction | 0 | 0 | 4 | 0 | 0 | # | 20 | Small | aldex2 | 98 |
| # | disease | 7 | 16 | 0 | 0.304347826 | 1 | # | 20 | Baseline | paralla | 98 |
| # | sex | 4 | 4 | 0 | 0.5 | 1 | # | 20 | Baseline | paralla | 98 |
| # | time | 5 | 4 | 0 | 0.555555556 | 1 | # | 20 | Baseline | paralla | 98 |
| # | interaction | 4 | 9 | 0 | 0.307692308 | 1 | # | 20 | Baseline | paralla | 98 |
| # | disease | 7 | 4 | 0 | 0.636363636 | 1 | # | 20 | Baseline | ancom | 98 |
| # | sex | 4 | 0 | 0 | 1 | 1 | # | 20 | Baseline | ancom | 98 |
| # | time | 5 | 4 | 0 | 0.555555556 | 1 | # | 20 | Baseline | ancom | 98 |
| # | interaction | 0 | 0 | 4 | 0 | 0 | # | 20 | Baseline | ancom | 98 |
| # | disease | 7 | 71 | 0 | 0.08974359 | 1 | # | 20 | Baseline | aldex2 | 98 |
| # | sex | 4 | 17 | 0 | 0.19047619 | 1 | # | 20 | Baseline | aldex2 | 98 |
| # | time | 5 | 39 | 0 | 0.113636364 | 1 | # | 20 | Baseline | aldex2 | 98 |
| # | interaction | 0 | 0 | 4 | 0 | 0 | # | 20 | Baseline | aldex2 | 98 |
| # | disease | 7 | 0 | 0 | 1 | 1 | # | 20 | Small | paralla | 98 |
| # | sex | 4 | 0 | 0 | 1 | 1 | # | 20 | Small | paralla | 98 |
| # | time | 5 | 0 | 0 | 1 | 1 | # | 20 | Small | paralla | 98 |
| # | interaction | 4 | 0 | 0 | 1 | 1 | # | 20 | Small | paralla | 98 |
| # | disease | 7 | 4 | 0 | 0.636363636 | 1 | # | 20 | Small | ancom | 98 |
| # | sex | 4 | 0 | 0 | 1 | 1 | # | 20 | Small | ancom | 98 |
| # | time | 5 | 4 | 0 | 0.555555556 | 1 | # | 20 | Small | ancom | 98 |
| # | interaction | 0 | 0 | 4 | 0 | 0 | # | 20 | Small | ancom | 98 |
| # | disease | 7 | 7 | 0 | 0.5 | 1 | # | 20 | Small | aldex2 | 98 |
| # | sex | 4 | 1 | 0 | 0.8 | 1 | # | 20 | Small | aldex2 | 98 |
| # | time | 5 | 6 | 0 | 0.454545455 | 1 | # | 20 | Small | aldex2 | 98 |
| # | interaction | 0 | 0 | 4 | 0 | 0 | # | 20 | Small | aldex2 | 98 |
| # | disease | 4 | 14 | 0 | 0.222222222 | 1 | # | 10 | Baseline | paralla | 99 |
| # | sex | 2 | 5 | 0 | 0.285714286 | 1 | # | 10 | Baseline | paralla | 99 |
| # | time | 2 | 3 | 0 | 0.4 | 1 | # | 10 | Baseline | paralla | 99 |
| # | interaction | 2 | 9 | 0 | 0.181818182 | 1 | # | 10 | Baseline | paralla | 99 |
| # | disease | 4 | 0 | 0 | 1 | 1 | # | 10 | Baseline | ancom | 99 |
| # | sex | 2 | 0 | 0 | 1 | 1 | # | 10 | Baseline | ancom | 99 |
| # | time | 2 | 0 | 0 | 1 | 1 | # | 10 | Baseline | ancom | 99 |
| # | interaction | 0 | 0 | 2 | 0 | 0 | # | 10 | Baseline | ancom | 99 |
| # | disease | 4 | 8 | 0 | 0.333333333 | 1 | # | 10 | Baseline | aldex2 | 99 |
| # | sex | 2 | 4 | 0 | 0.333333333 | 1 | # | 10 | Baseline | aldex2 | 99 |
| # | time | 2 | 7 | 0 | 0.222222222 | 1 | # | 10 | Baseline | aldex2 | 99 |
| # | interaction | 0 | 0 | 2 | 0 | 0 | # | 10 | Baseline | aldex2 | 99 |
| # | disease | 4 | 0 | 0 | 1 | 1 | # | 10 | Small | paralla | 99 |
| # | sex | 2 | 0 | 0 | 1 | 1 | # | 10 | Small | paralla | 99 |
| # | time | 2 | 0 | 0 | 1 | 1 | # | 10 | Small | paralla | 99 |
| # | interaction | 1 | 0 | 1 | 1 | 0.5 | # | 10 | Small | paralla | 99 |
| # | disease | 4 | 0 | 0 | 1 | 1 | # | 10 | Small | ancom | 99 |
| # | sex | 2 | 0 | 0 | 1 | 1 | # | 10 | Small | ancom | 99 |
| # | time | 2 | 0 | 0 | 1 | 1 | # | 10 | Small | ancom | 99 |
| # | interaction | 0 | 0 | 2 | 0 | 0 | # | 10 | Small | ancom | 99 |
| # | disease | 4 | 0 | 0 | 1 | 1 | # | 10 | Small | aldex2 | 99 |
| # | sex | 2 | 0 | 0 | 1 | 1 | # | 10 | Small | aldex2 | 99 |
| # | time | 2 | 1 | 0 | 0.666666667 | 1 | # | 10 | Small | aldex2 | 99 |
| # | interaction | 0 | 0 | 2 | 0 | 0 | # | 10 | Small | aldex2 | 99 |
| # | disease | 4 | 12 | 0 | 0.25 | 1 | # | 10 | Baseline | paralla | 99 |
| # | sex | 2 | 9 | 0 | 0.181818182 | 1 | # | 10 | Baseline | paralla | 99 |
| # | time | 2 | 4 | 0 | 0.333333333 | 1 | # | 10 | Baseline | paralla | 99 |
| # | interaction | 2 | 6 | 0 | 0.25 | 1 | # | 10 | Baseline | paralla | 99 |
| # | disease | 4 | 1 | 0 | 0.8 | 1 | # | 10 | Baseline | ancom | 99 |
| # | sex | 2 | 0 | 0 | 1 | 1 | # | 10 | Baseline | ancom | 99 |
| # | time | 2 | 1 | 0 | 0.666666667 | 1 | # | 10 | Baseline | ancom | 99 |
| # | interaction | 0 | 0 | 2 | 0 | 0 | # | 10 | Baseline | ancom | 99 |
| # | disease | 4 | 13 | 0 | 0.235294118 | 1 | # | 10 | Baseline | aldex2 | 99 |
| # | sex | 2 | 6 | 0 | 0.25 | 1 | # | 10 | Baseline | aldex2 | 99 |
| # | time | 2 | 10 | 0 | 0.166666667 | 1 | # | 10 | Baseline | aldex2 | 99 |
| # | interaction | 0 | 0 | 2 | 0 | 0 | # | 10 | Baseline | aldex2 | 99 |
| # | disease | 4 | 0 | 0 | 1 | 1 | # | 10 | Small | paralla | 99 |
| # | sex | 2 | 0 | 0 | 1 | 1 | # | 10 | Small | paralla | 99 |
| # | time | 2 | 0 | 0 | 1 | 1 | # | 10 | Small | paralla | 99 |
| # | interaction | 2 | 0 | 0 | 1 | 1 | # | 10 | Small | paralla | 99 |
| # | disease | 4 | 1 | 0 | 0.8 | 1 | # | 10 | Small | ancom | 99 |
| # | sex | 2 | 0 | 0 | 1 | 1 | # | 10 | Small | ancom | 99 |
| # | time | 2 | 1 | 0 | 0.666666667 | 1 | # | 10 | Small | ancom | 99 |
| # | interaction | 0 | 0 | 2 | 0 | 0 | # | 10 | Small | ancom | 99 |
| # | disease | 4 | 2 | 0 | 0.666666667 | 1 | # | 10 | Small | aldex2 | 99 |
| # | sex | 2 | 0 | 0 | 1 | 1 | # | 10 | Small | aldex2 | 99 |
| # | time | 2 | 2 | 0 | 0.5 | 1 | # | 10 | Small | aldex2 | 99 |
| # | interaction | 0 | 0 | 2 | 0 | 0 | # | 10 | Small | aldex2 | 99 |
| # | disease | 4 | 5 | 0 | 0.444444444 | 1 | # | 10 | Baseline | paralla | 99 |
| # | sex | 2 | 4 | 0 | 0.333333333 | 1 | # | 10 | Baseline | paralla | 99 |
| # | time | 2 | 1 | 0 | 0.666666667 | 1 | # | 10 | Baseline | paralla | 99 |
| # | interaction | 2 | 7 | 0 | 0.222222222 | 1 | # | 10 | Baseline | paralla | 99 |
| # | disease | 4 | 2 | 0 | 0.666666667 | 1 | # | 10 | Baseline | ancom | 99 |

| | | | | | | | | | | | |
|---|---|---|---|---|---|---|---|---|---|---|---|
| # | sex | 2 | 0 | 0 | 1 | 1 | # | 10 | Baseline | ancom | 99 |
| # | time | 2 | 2 | 0 | 0.5 | 1 | # | 10 | Baseline | ancom | 99 |
| # | interaction | 0 | 0 | 2 | 0 | 0 | # | 10 | Baseline | ancom | 99 |
| # | disease | 4 | 25 | 0 | 0.137931034 | 1 | # | 10 | Baseline | sided2 | 99 |
| # | sex | 2 | 7 | 0 | 0.222222222 | 1 | # | 10 | Baseline | sided2 | 99 |
| # | time | 2 | 9 | 0 | 0.181818182 | 1 | # | 10 | Baseline | sided2 | 99 |
| # | interaction | 0 | 0 | 2 | 0 | 0 | # | 10 | Baseline | sided2 | 99 |
| # | disease | 4 | 0 | 0 | 1 | 1 | # | 10 | Small | paralle | 99 |
| # | sex | 2 | 0 | 0 | 1 | 1 | # | 10 | Small | paralle | 99 |
| # | time | 2 | 0 | 0 | 1 | 1 | # | 10 | Small | paralle | 99 |
| # | interaction | 2 | 0 | 0 | 1 | 1 | # | 10 | Small | paralle | 99 |
| # | disease | 4 | 2 | 0 | 0.666666667 | 1 | # | 10 | Small | ancom | 99 |
| # | sex | 2 | 0 | 0 | 1 | 1 | # | 10 | Small | ancom | 99 |
| # | time | 2 | 2 | 0 | 0.5 | 1 | # | 10 | Small | ancom | 99 |
| # | interaction | 0 | 0 | 2 | 0 | 0 | # | 10 | Small | ancom | 99 |
| # | disease | 4 | 6 | 0 | 0.4 | 1 | # | 10 | Small | sided2 | 99 |
| # | sex | 2 | 1 | 0 | 0.666666667 | 1 | # | 10 | Small | sided2 | 99 |
| # | time | 2 | 3 | 0 | 0.4 | 1 | # | 10 | Small | sided2 | 99 |
| # | interaction | 0 | 0 | 2 | 0 | 0 | # | 10 | Small | sided2 | 99 |
| # | disease | 4 | 7 | 0 | 0.363636364 | 1 | # | 10 | Baseline | paralle | 99 |
| # | sex | 2 | 0 | 0 | 1 | 1 | # | 10 | Baseline | paralle | 99 |
| # | time | 2 | 1 | 0 | 0.666666667 | 1 | # | 10 | Baseline | paralle | 99 |
| # | interaction | 2 | 5 | 0 | 0.285714286 | 1 | # | 10 | Baseline | paralle | 99 |
| # | disease | 4 | 2 | 0 | 0.666666667 | 1 | # | 10 | Baseline | ancom | 99 |
| # | sex | 2 | 0 | 0 | 1 | 1 | # | 10 | Baseline | ancom | 99 |
| # | time | 2 | 2 | 0 | 0.5 | 1 | # | 10 | Baseline | ancom | 99 |
| # | interaction | 0 | 0 | 2 | 0 | 0 | # | 10 | Baseline | ancom | 99 |
| # | disease | 4 | 29 | 0 | 0.121212121 | 1 | # | 10 | Baseline | sided2 | 99 |
| # | sex | 2 | 6 | 0 | 0.25 | 1 | # | 10 | Baseline | sided2 | 99 |
| # | time | 2 | 8 | 0 | 0.2 | 1 | # | 10 | Baseline | sided2 | 99 |
| # | interaction | 0 | 0 | 2 | 0 | 0 | # | 10 | Baseline | sided2 | 99 |
| # | disease | 4 | 1 | 0 | 0.8 | 1 | # | 10 | Small | paralle | 99 |
| # | sex | 2 | 0 | 0 | 1 | 1 | # | 10 | Small | paralle | 99 |
| # | time | 2 | 0 | 0 | 1 | 1 | # | 10 | Small | paralle | 99 |
| # | interaction | 2 | 0 | 0 | 1 | 1 | # | 10 | Small | paralle | 99 |
| # | disease | 4 | 2 | 0 | 0.666666667 | 1 | # | 10 | Small | ancom | 99 |
| # | sex | 2 | 0 | 0 | 1 | 1 | # | 10 | Small | ancom | 99 |
| # | time | 2 | 2 | 0 | 0.5 | 1 | # | 10 | Small | ancom | 99 |
| # | interaction | 0 | 0 | 2 | 0 | 0 | # | 10 | Small | ancom | 99 |
| # | disease | 4 | 3 | 0 | 0.571428571 | 1 | # | 10 | Small | sided2 | 99 |
| # | sex | 2 | 1 | 0 | 0.666666667 | 1 | # | 10 | Small | sided2 | 99 |
| # | time | 2 | 4 | 0 | 0.333333333 | 1 | # | 10 | Small | sided2 | 99 |
| # | interaction | 0 | 0 | 2 | 0 | 0 | # | 10 | Small | sided2 | 99 |
| # | disease | 7 | 10 | 0 | 0.411764706 | 1 | # | 20 | Baseline | paralle | 99 |
| # | sex | 4 | 15 | 0 | 0.210526316 | 1 | # | 20 | Baseline | paralle | 99 |
| # | time | 5 | 8 | 0 | 0.384615385 | 1 | # | 20 | Baseline | paralle | 99 |
| # | interaction | 3 | 9 | 1 | 0.25 | 0.75 | # | 20 | Baseline | paralle | 99 |
| # | disease | 7 | 1 | 0 | 0.875 | 1 | # | 20 | Baseline | ancom | 99 |
| # | sex | 4 | 0 | 0 | 1 | 1 | # | 20 | Baseline | ancom | 99 |
| # | time | 5 | 1 | 0 | 0.833333333 | 1 | # | 20 | Baseline | ancom | 99 |
| # | interaction | 0 | 0 | 4 | 0 | 0 | # | 20 | Baseline | ancom | 99 |
| # | disease | 7 | 21 | 0 | 0.25 | 1 | # | 20 | Baseline | sided2 | 99 |
| # | sex | 4 | 7 | 0 | 0.363636364 | 1 | # | 20 | Baseline | sided2 | 99 |
| # | time | 5 | 17 | 0 | 0.227272727 | 1 | # | 20 | Baseline | sided2 | 99 |
| # | interaction | 0 | 0 | 4 | 0 | 0 | # | 20 | Baseline | sided2 | 99 |
| # | disease | 7 | 0 | 0 | 1 | 1 | # | 20 | Small | paralle | 99 |
| # | sex | 4 | 1 | 0 | 0.8 | 1 | # | 20 | Small | paralle | 99 |
| # | time | 5 | 0 | 0 | 1 | 1 | # | 20 | Small | paralle | 99 |
| # | interaction | 0 | 0 | 4 | 0 | 0 | # | 20 | Small | paralle | 99 |
| # | disease | 7 | 1 | 0 | 0.875 | 1 | # | 20 | Small | ancom | 99 |
| # | sex | 4 | 0 | 0 | 1 | 1 | # | 20 | Small | ancom | 99 |
| # | time | 5 | 1 | 0 | 0.833333333 | 1 | # | 20 | Small | ancom | 99 |
| # | interaction | 0 | 0 | 4 | 0 | 0 | # | 20 | Small | ancom | 99 |
| # | disease | 7 | 2 | 0 | 0.777777778 | 1 | # | 20 | Small | sided2 | 99 |
| # | sex | 4 | 0 | 0 | 1 | 1 | # | 20 | Small | sided2 | 99 |
| # | time | 5 | 2 | 0 | 0.714285714 | 1 | # | 20 | Small | sided2 | 99 |
| # | interaction | 0 | 0 | 4 | 0 | 0 | # | 20 | Small | sided2 | 99 |
| # | disease | 7 | 8 | 0 | 0.466666667 | 1 | # | 20 | Baseline | paralle | 99 |
| # | sex | 4 | 16 | 0 | 0.2 | 1 | # | 20 | Baseline | paralle | 99 |
| # | time | 5 | 12 | 0 | 0.294117647 | 1 | # | 20 | Baseline | paralle | 99 |
| # | interaction | 4 | 5 | 0 | 0.444444444 | 1 | # | 20 | Baseline | paralle | 99 |
| # | disease | 7 | 4 | 0 | 0.636363636 | 1 | # | 20 | Baseline | ancom | 99 |
| # | sex | 4 | 0 | 0 | 1 | 1 | # | 20 | Baseline | ancom | 99 |
| # | time | 5 | 4 | 0 | 0.555555556 | 1 | # | 20 | Baseline | ancom | 99 |
| # | interaction | 0 | 0 | 4 | 0 | 0 | # | 20 | Baseline | ancom | 99 |
| # | disease | 7 | 34 | 0 | 0.170731707 | 1 | # | 20 | Baseline | sided2 | 99 |
| # | sex | 4 | 17 | 0 | 0.19047619 | 1 | # | 20 | Baseline | sided2 | 99 |
| # | time | 5 | 23 | 0 | 0.178571429 | 1 | # | 20 | Baseline | sided2 | 99 |
| # | interaction | 0 | 0 | 4 | 0 | 0 | # | 20 | Baseline | sided2 | 99 |
| # | disease | 7 | 0 | 0 | 1 | 1 | # | 20 | Small | paralle | 99 |
| # | sex | 4 | 1 | 0 | 0.8 | 1 | # | 20 | Small | paralle | 99 |
| # | time | 5 | 0 | 0 | 1 | 1 | # | 20 | Small | paralle | 99 |
| # | interaction | 3 | 0 | 1 | 1 | 0.75 | # | 20 | Small | paralle | 99 |
| # | disease | 7 | 4 | 0 | 0.636363636 | 1 | # | 20 | Small | ancom | 99 |
| # | sex | 4 | 0 | 0 | 1 | 1 | # | 20 | Small | ancom | 99 |
| # | time | 5 | 3 | 0 | 0.625 | 1 | # | 20 | Small | ancom | 99 |
| # | interaction | 0 | 0 | 4 | 0 | 0 | # | 20 | Small | ancom | 99 |
| # | disease | 7 | 6 | 0 | 0.538461538 | 1 | # | 20 | Small | sided2 | 99 |
| # | sex | 4 | 1 | 0 | 0.8 | 1 | # | 20 | Small | sided2 | 99 |
| # | time | 5 | 5 | 0 | 0.5 | 1 | # | 20 | Small | sided2 | 99 |
| # | interaction | 0 | 0 | 4 | 0 | 0 | # | 20 | Small | sided2 | 99 |
| # | disease | 7 | 6 | 0 | 0.538461538 | 1 | # | 20 | Baseline | paralle | 99 |
| # | sex | 4 | 6 | 0 | 0.4 | 1 | # | 20 | Baseline | paralle | 99 |
| # | time | 5 | 11 | 0 | 0.3125 | 1 | # | 20 | Baseline | paralle | 99 |
| # | interaction | 4 | 7 | 0 | 0.363636364 | 1 | # | 20 | Baseline | paralle | 99 |
| # | disease | 7 | 4 | 0 | 0.636363636 | 1 | # | 20 | Baseline | ancom | 99 |
| # | sex | 4 | 0 | 0 | 1 | 1 | # | 20 | Baseline | ancom | 99 |
| # | time | 5 | 4 | 0 | 0.555555556 | 1 | # | 20 | Baseline | ancom | 99 |
| # | interaction | 0 | 0 | 4 | 0 | 0 | # | 20 | Baseline | ancom | 99 |
| # | disease | 7 | 46 | 0 | 0.132075472 | 1 | # | 20 | Baseline | sided2 | 99 |
| # | sex | 4 | 13 | 0 | 0.235294118 | 1 | # | 20 | Baseline | sided2 | 99 |
| # | time | 5 | 33 | 0 | 0.131578947 | 1 | # | 20 | Baseline | sided2 | 99 |
| # | interaction | 0 | 0 | 4 | 0 | 0 | # | 20 | Baseline | sided2 | 99 |
| # | disease | 7 | 0 | 0 | 1 | 1 | # | 20 | Small | paralle | 99 |
| # | sex | 4 | 0 | 0 | 1 | 1 | # | 20 | Small | paralle | 99 |
| # | time | 5 | 0 | 0 | 1 | 1 | # | 20 | Small | paralle | 99 |
| # | interaction | 3 | 0 | 1 | 1 | 0.75 | # | 20 | Small | paralle | 99 |
| # | disease | 7 | 4 | 0 | 0.636363636 | 1 | # | 20 | Small | ancom | 99 |
| # | sex | 4 | 0 | 0 | 1 | 1 | # | 20 | Small | ancom | 99 |
| # | time | 5 | 4 | 0 | 0.555555556 | 1 | # | 20 | Small | ancom | 99 |
| # | interaction | 0 | 0 | 4 | 0 | 0 | # | 20 | Small | ancom | 99 |
| # | disease | 7 | 9 | 0 | 0.4375 | 1 | # | 20 | Small | sided2 | 99 |
| # | sex | 4 | 2 | 0 | 0.666666667 | 1 | # | 20 | Small | sided2 | 99 |
| # | time | 5 | 4 | 0 | 0.555555556 | 1 | # | 20 | Small | sided2 | 99 |
| # | interaction | 0 | 0 | 4 | 0 | 0 | # | 20 | Small | sided2 | 99 |
| # | disease | 7 | 9 | 0 | 0.4375 | 1 | # | 20 | Baseline | paralle | 99 |
| # | sex | 4 | 8 | 0 | 0.333333333 | 1 | # | 20 | Baseline | paralle | 99 |
| # | time | 5 | 6 | 0 | 0.454545455 | 1 | # | 20 | Baseline | paralle | 99 |
| # | interaction | 4 | 6 | 0 | 0.4 | 1 | # | 20 | Baseline | paralle | 99 |
| # | disease | 7 | 4 | 0 | 0.636363636 | 1 | # | 20 | Baseline | ancom | 99 |
| # | sex | 4 | 0 | 0 | 1 | 1 | # | 20 | Baseline | ancom | 99 |
| # | time | 5 | 4 | 0 | 0.555555556 | 1 | # | 20 | Baseline | ancom | 99 |
| # | interaction | 0 | 0 | 4 | 0 | 0 | # | 20 | Baseline | ancom | 99 |
| # | disease | 7 | 54 | 0 | 0.114754098 | 1 | # | 20 | Baseline | sided2 | 99 |
| # | sex | 4 | 16 | 0 | 0.2 | 1 | # | 20 | Baseline | sided2 | 99 |
| # | time | 5 | 39 | 0 | 0.113636364 | 1 | # | 20 | Baseline | sided2 | 99 |
| # | interaction | 0 | 0 | 4 | 0 | 0 | # | 20 | Baseline | sided2 | 99 |
| # | disease | 7 | 0 | 0 | 1 | 1 | # | 20 | Small | paralle | 99 |
| # | sex | 4 | 0 | 0 | 1 | 1 | # | 20 | Small | paralle | 99 |
| # | time | 5 | 0 | 0 | 1 | 1 | # | 20 | Small | paralle | 99 |
| # | interaction | 4 | 0 | 0 | 1 | 1 | # | 20 | Small | paralle | 99 |
| # | disease | 7 | 4 | 0 | 0.636363636 | 1 | # | 20 | Small | ancom | 99 |
| # | sex | 4 | 0 | 0 | 1 | 1 | # | 20 | Small | ancom | 99 |
| # | time | 5 | 4 | 0 | 0.555555556 | 1 | # | 20 | Small | ancom | 99 |
| # | interaction | 0 | 0 | 4 | 0 | 0 | # | 20 | Small | ancom | 99 |
| # | disease | 7 | 13 | 0 | 0.35 | 1 | # | 20 | Small | sided2 | 99 |
| # | sex | 4 | 3 | 0 | 0.571428571 | 1 | # | 20 | Small | sided2 | 99 |
| # | time | 5 | 3 | 0 | 0.625 | 1 | # | 20 | Small | sided2 | 99 |
| # | interaction | 0 | 0 | 4 | 0 | 0 | # | 20 | Small | sided2 | 99 |
| # | disease | 4 | 9 | 0 | 0.307692308 | 1 | # | 10 | Baseline | paralle | 100 |
| # | sex | 2 | 9 | 0 | 0.181818182 | 1 | # | 10 | Baseline | paralle | 100 |
| # | time | 2 | 4 | 0 | 0.333333333 | 1 | # | 10 | Baseline | paralle | 100 |
| # | interaction | 1 | 10 | 1 | 0.090909091 | 0.5 | # | 10 | Baseline | paralle | 100 |
| # | disease | 4 | 1 | 0 | 0.8 | 1 | # | 10 | Baseline | ancom | 100 |
| # | sex | 2 | 0 | 0 | 1 | 1 | # | 10 | Baseline | ancom | 100 |
| # | time | 2 | 0 | 0 | 1 | 1 | # | 10 | Baseline | ancom | 100 |
| # | interaction | 0 | 0 | 2 | 0 | 0 | # | 10 | Baseline | ancom | 100 |
| # | disease | 4 | 8 | 0 | 0.333333333 | 1 | # | 10 | Baseline | sided2 | 100 |
| # | sex | 2 | 3 | 0 | 0.4 | 1 | # | 10 | Baseline | sided2 | 100 |
| # | time | 2 | 8 | 0 | 0.2 | 1 | # | 10 | Baseline | sided2 | 100 |
| # | interaction | 0 | 0 | 2 | 0 | 0 | # | 10 | Baseline | sided2 | 100 |
| # | disease | 4 | 0 | 0 | 1 | 1 | # | 10 | Small | paralle | 100 |
| # | sex | 2 | 1 | 0 | 0.666666667 | 1 | # | 10 | Small | paralle | 100 |
| # | time | 2 | 0 | 0 | 1 | 1 | # | 10 | Small | paralle | 100 |
| # | interaction | 1 | 0 | 1 | 1 | 0.5 | # | 10 | Small | paralle | 100 |
| # | disease | 4 | 0 | 0 | 1 | 1 | # | 10 | Small | ancom | 100 |
| # | sex | 2 | 0 | 0 | 1 | 1 | # | 10 | Small | ancom | 100 |
| # | time | 2 | 0 | 0 | 1 | 1 | # | 10 | Small | ancom | 100 |
| # | interaction | 0 | 0 | 2 | 0 | 0 | # | 10 | Small | ancom | 100 |
| # | disease | 4 | 1 | 0 | 0.8 | 1 | # | 10 | Small | sided2 | 100 |
| # | sex | 2 | 0 | 0 | 1 | 1 | # | 10 | Small | sided2 | 100 |
| # | time | 2 | 1 | 0 | 0.666666667 | 1 | # | 10 | Small | sided2 | 100 |
| # | interaction | 0 | 0 | 2 | 0 | 0 | # | 10 | Small | sided2 | 100 |
| # | disease | 4 | 7 | 0 | 0.363636364 | 1 | # | 10 | Baseline | paralle | 100 |
| # | sex | 2 | 8 | 0 | 0.2 | 1 | # | 10 | Baseline | paralle | 100 |
| # | time | 2 | 8 | 0 | 0.2 | 1 | # | 10 | Baseline | paralle | 100 |
| # | interaction | 2 | 9 | 0 | 0.181818182 | 1 | # | 10 | Baseline | paralle | 100 |
| # | disease | 4 | 1 | 0 | 0.8 | 1 | # | 10 | Baseline | ancom | 100 |
| # | sex | 2 | 0 | 0 | 1 | 1 | # | 10 | Baseline | ancom | 100 |
| # | time | 2 | 1 | 0 | 0.666666667 | 1 | # | 10 | Baseline | ancom | 100 |
| # | interaction | 0 | 0 | 2 | 0 | 0 | # | 10 | Baseline | ancom | 100 |
| # | disease | 4 | 14 | 0 | 0.222222222 | 1 | # | 10 | Baseline | sided2 | 100 |
| # | sex | 2 | 12 | 0 | 0.142857143 | 1 | # | 10 | Baseline | sided2 | 100 |
| # | time | 2 | 16 | 0 | 0.111111111 | 1 | # | 10 | Baseline | sided2 | 100 |
| # | interaction | 0 | 0 | 2 | 0 | 0 | # | 10 | Baseline | sided2 | 100 |
| # | disease | 4 | 1 | 0 | 0.8 | 1 | # | 10 | Small | paralle | 100 |
| # | sex | 2 | 1 | 0 | 0.666666667 | 1 | # | 10 | Small | paralle | 100 |
| # | time | 2 | 0 | 0 | 1 | 1 | # | 10 | Small | paralle | 100 |
| # | interaction | 1 | 2 | 1 | 0.333333333 | 0.5 | # | 10 | Small | paralle | 100 |
| # | disease | 4 | 1 | 0 | 0.8 | 1 | # | 10 | Small | ancom | 100 |
| # | sex | 2 | 0 | 0 | 1 | 1 | # | 10 | Small | ancom | 100 |
| # | time | 2 | 1 | 0 | 0.666666667 | 1 | # | 10 | Small | ancom | 100 |
| # | interaction | 0 | 0 | 2 | 0 | 0 | # | 10 | Small | ancom | 100 |
| # | disease | 4 | 2 | 0 | 0.666666667 | 1 | # | 10 | Small | sided2 | 100 |
| # | sex | 2 | 0 | 0 | 1 | 1 | # | 10 | Small | sided2 | 100 |
| # | time | 2 | 2 | 0 | 0.5 | 1 | # | 10 | Small | sided2 | 100 |
| # | interaction | 0 | 0 | 2 | 0 | 0 | # | 10 | Small | sided2 | 100 |
| # | disease | 4 | 4 | 0 | 0.5 | 1 | # | 10 | Baseline | paralle | 100 |
| # | sex | 2 | 5 | 0 | 0.285714286 | 1 | # | 10 | Baseline | paralle | 100 |
| # | time | 2 | 5 | 0 | 0.285714286 | 1 | # | 10 | Baseline | paralle | 100 |
| # | interaction | 2 | 10 | 0 | 0.166666667 | 1 | # | 10 | Baseline | paralle | 100 |
| # | disease | 4 | 1 | 0 | 0.8 | 1 | # | 10 | Baseline | ancom | 100 |
| # | sex | 2 | 0 | 0 | 1 | 1 | # | 10 | Baseline | ancom | 100 |
| # | time | 2 | 2 | 0 | 0.5 | 1 | # | 10 | Baseline | ancom | 100 |
| # | interaction | 0 | 0 | 2 | 0 | 0 | # | 10 | Baseline | ancom | 100 |
| # | disease | 4 | 19 | 0 | 0.173913043 | 1 | # | 10 | Baseline | sided2 | 100 |
| # | sex | 2 | 5 | 0 | 0.285714286 | 1 | # | 10 | Baseline | sided2 | 100 |
| # | time | 2 | 13 | 0 | 0.133333333 | 1 | # | 10 | Baseline | sided2 | 100 |
| # | interaction | 0 | 0 | 2 | 0 | 0 | # | 10 | Baseline | sided2 | 100 |
| # | disease | 4 | 0 | 0 | 1 | 1 | # | 10 | Small | paralle | 100 |
| # | sex | 2 | 0 | 0 | 1 | 1 | # | 10 | Small | paralle | 100 |
| # | time | 2 | 0 | 0 | 1 | 1 | # | 10 | Small | paralle | 100 |
| # | interaction | 2 | 1 | 0 | 0.666666667 | 1 | # | 10 | Small | paralle | 100 |
| # | disease | 4 | 1 | 0 | 0.8 | 1 | # | 10 | Small | ancom | 100 |
| # | sex | 2 | 0 | 0 | 1 | 1 | # | 10 | Small | ancom | 100 |
| # | time | 2 | 2 | 0 | 0.5 | 1 | # | 10 | Small | ancom | 100 |
| # | interaction | 0 | 0 | 2 | 0 | 0 | # | 10 | Small | ancom | 100 |
| # | disease | 4 | 2 | 0 | 0.666666667 | 1 | # | 10 | Small | sided2 | 100 |
| # | sex | 2 | 0 | 0 | 1 | 1 | # | 10 | Small | sided2 | 100 |
| # | time | 2 | 3 | 0 | 0.4 | 1 | # | 10 | Small | sided2 | 100 |
| # | interaction | 0 | 0 | 2 | 0 | 0 | # | 10 | Small | sided2 | 100 |
| # | disease | 4 | 3 | 0 | 0.571428571 | 1 | # | 10 | Baseline | paralle | 100 |
| # | sex | 2 | 6 | 0 | 0.25 | 1 | # | 10 | Baseline | paralle | 100 |
| # | time | 2 | 4 | 0 | 0.333333333 | 1 | # | 10 | Baseline | paralle | 100 |
| # | interaction | 2 | 6 | 0 | 0.25 | 1 | # | 10 | Baseline | paralle | 100 |
| # | disease | 4 | 2 | 0 | 0.666666667 | 1 | # | 10 | Baseline | ancom | 100 |
| # | sex | 2 | 0 | 0 | 1 | 1 | # | 10 | Baseline | ancom | 100 |
| # | time | 2 | 2 | 0 | 0.5 | 1 | # | 10 | Baseline | ancom | 100 |
| # | interaction | 0 | 0 | 2 | 0 | 0 | # | 10 | Baseline | ancom | 100 |
| # | disease | 4 | 28 | 0 | 0.125 | 1 | # | 10 | Baseline | sided2 | 100 |
| # | sex | 2 | 7 | 0 | 0.222222222 | 1 | # | 10 | Baseline | sided2 | 100 |
| # | time | 2 | 17 | 0 | 0.105263158 | 1 | # | 10 | Baseline | sided2 | 100 |
| # | interaction | 0 | 0 | 2 | 0 | 0 | # | 10 | Baseline | sided2 | 100 |
| # | disease | 4 | 0 | 0 | 1 | 1 | # | 10 | Small | paralle | 100 |
| # | sex | 2 | 0 | 0 | 1 | 1 | # | 10 | Small | paralle | 100 |
| # | time | 2 | 0 | 0 | 1 | 1 | # | 10 | Small | paralle | 100 |
| # | interaction | 2 | 1 | 0 | 0.666666667 | 1 | # | 10 | Small | paralle | 100 |
| # | disease | 4 | 2 | 0 | 0.666666667 | 1 | # | 10 | Small | ancom | 100 |
| # | sex | 2 | 0 | 0 | 1 | 1 | # | 10 | Small | ancom | 100 |
| # | time | 2 | 2 | 0 | 0.5 | 1 | # | 10 | Small | ancom | 100 |
| # | interaction | 0 | 0 | 2 | 0 | 0 | # | 10 | Small | ancom | 100 |
| # | disease | 4 | 2 | 0 | 0.666666667 | 1 | # | 10 | Small | sided2 | 100 |
| # | sex | 2 | 0 | 0 | 1 | 1 | # | 10 | Small | sided2 | 100 |
| # | time | 2 | 3 | 0 | 0.4 | 1 | # | 10 | Small | sided2 | 100 |
| # | interaction | 0 | 0 | 2 | 0 | 0 | # | 10 | Small | sided2 | 100 |
| # | disease | 7 | 12 | 0 | 0.368421053 | 1 | # | 20 | Baseline | paralle | 100 |
| # | sex | 4 | 6 | 0 | 0.4 | 1 | # | 20 | Baseline | paralle | 100 |
| # | time | 5 | 7 | 0 | 0.416666667 | 1 | # | 20 | Baseline | paralle | 100 |
| # | interaction | 4 | 15 | 0 | 0.210526316 | 1 | # | 20 | Baseline | paralle | 100 |
| # | disease | 7 | 1 | 0 | 0.875 | 1 | # | 20 | Baseline | ancom | 100 |
| # | sex | 4 | 0 | 0 | 1 | 1 | # | 20 | Baseline | ancom | 100 |
| # | time | 5 | 2 | 0 | 0.714285714 | 1 | # | 20 | Baseline | ancom | 100 |
| # | interaction | 0 | 0 | 4 | 0 | 0 | # | 20 | Baseline | ancom | 100 |
| # | disease | 7 | 15 | 0 | 0.318181818 | 1 | # | 20 | Baseline | sided2 | 100 |
| # | sex | 4 | 9 | 0 | 0.307692308 | 1 | # | 20 | Baseline | sided2 | 100 |
| # | time | 5 | 13 | 0 | 0.277777778 | 1 | # | 20 | Baseline | sided2 | 100 |
| # | interaction | 0 | 0 | 4 | 0 | 0 | # | 20 | Baseline | sided2 | 100 |
| # | disease | 7 | 0 | 0 | 1 | 1 | # | 20 | Small | paralle | 100 |
| # | sex | 4 | 0 | 0 | 1 | 1 | # | 20 | Small | paralle | 100 |
| # | time | 5 | 0 | 0 | 1 | 1 | # | 20 | Small | paralle | 100 |
| # | interaction | 3 | 1 | 1 | 0.75 | 0.75 | # | 20 | Small | paralle | 100 |
| # | disease | 7 | 1 | 0 | 0.875 | 1 | # | 20 | Small | ancom | 100 |
| # | sex | 4 | 0 | 0 | 1 | 1 | # | 20 | Small | ancom | 100 |
| # | time | 5 | 1 | 0 | 0.833333333 | 1 | # | 20 | Small | ancom | 100 |
| # | interaction | 0 | 0 | 4 | 0 | 0 | # | 20 | Small | ancom | 100 |
| # | disease | 7 | 1 | 0 | 0.875 | 1 | # | 20 | Small | sided2 | 100 |
| # | sex | 4 | 0 | 0 | 1 | 1 | # | 20 | Small | sided2 | 100 |
| # | time | 5 | 0 | 0 | 1 | 1 | # | 20 | Small | sided2 | 100 |
| # | interaction | 0 | 0 | 4 | 0 | 0 | # | 20 | Small | sided2 | 100 |
| # | disease | 7 | 11 | 0 | 0.388888889 | 1 | # | 20 | Baseline | paralle | 100 |
| # | sex | 4 | 5 | 0 | 0.444444444 | 1 | # | 20 | Baseline | paralle | 100 |
| # | time | 5 | 7 | 0 | 0.416666667 | 1 | # | 20 | Baseline | paralle | 100 |
| # | interaction | 4 | 5 | 0 | 0.444444444 | 1 | # | 20 | Baseline | paralle | 100 |
| # | disease | 7 | 3 | 0 | 0.7 | 1 | # | 20 | Baseline | ancom | 100 |
| # | sex | 4 | 0 | 0 | 1 | 1 | # | 20 | Baseline | ancom | 100 |
| # | time | 5 | 3 | 0 | 0.625 | 1 | # | 20 | Baseline | ancom | 100 |
| # | interaction | 0 | 0 | 4 | 0 | 0 | # | 20 | Baseline | ancom | 100 |
| # | disease | 7 | 62 | 0 | 0.142857143 | 1 | # | 20 | Baseline | sided2 | 100 |
| # | sex | 4 | 11 | 0 | 0.266666667 | 1 | # | 20 | Baseline | sided2 | 100 |
| # | time | 5 | 27 | 0 | 0.15625 | 1 | # | 20 | Baseline | sided2 | 100 |
| # | interaction | 0 | 0 | 4 | 0 | 0 | # | 20 | Baseline | sided2 | 100 |
| # | disease | 7 | 0 | 0 | 1 | 1 | # | 20 | Small | paralle | 100 |
| # | sex | 4 | 0 | 0 | 1 | 1 | # | 20 | Small | paralle | 100 |
| # | time | 5 | 0 | 0 | 1 | 1 | # | 20 | Small | paralle | 100 |
| # | interaction | 4 | 1 | 0 | 0.8 | 1 | # | 20 | Small | paralle | 100 |
| # | disease | 7 | 3 | 0 | 0.7 | 1 | # | 20 | Small | ancom | 100 |
| # | sex | 4 | 0 | 0 | 1 | 1 | # | 20 | Small | ancom | 100 |
| # | time | 5 | 3 | 0 | 0.625 | 1 | # | 20 | Small | ancom | 100 |
| # | interaction | 0 | 0 | 4 | 0 | 0 | # | 20 | Small | ancom | 100 |
| # | disease | 7 | 4 | 0 | 0.636363636 | 1 | # | 20 | Small | sided2 | 100 |
| # | sex | 4 | 0 | 0 | 1 | 1 | # | 20 | Small | sided2 | 100 |
| # | time | 5 | 4 | 0 | 0.555555556 | 1 | # | 20 | Small | sided2 | 100 |
| # | interaction | 0 | 0 | 4 | 0 | 0 | # | 20 | Small | sided2 | 100 |
| # | disease | 7 | 12 | 0 | 0.368421053 | 1 | # | 20 | Baseline | paralle | 100 |
| # | sex | 4 | 5 | 0 | 0.444444444 | 1 | # | 20 | Baseline | paralle | 100 |
| # | time | 5 | 7 | 0 | 0.416666667 | 1 | # | 20 | Baseline | paralle | 100 |
| # | interaction | 4 | 5 | 0 | 0.444444444 | 1 | # | 20 | Baseline | paralle | 100 |
| # | disease | 7 | 4 | 0 | 0.636363636 | 1 | # | 20 | Baseline | ancom | 100 |
| # | sex | 4 | 0 | 0 | 1 | 1 | # | 20 | Baseline | ancom | 100 |
| # | time | 5 | 4 | 0 | 0.555555556 | 1 | # | 20 | Baseline | ancom | 100 |
| # | interaction | 0 | 0 | 4 | 0 | 0 | # | 20 | Baseline | ancom | 100 |
| # | disease | 7 | 59 | 0 | 0.106060606 | 1 | # | 20 | Baseline | sided2 | 100 |
| # | sex | 4 | 15 | 0 | 0.210526316 | 1 | # | 20 | Baseline | sided2 | 100 |
| # | time | 5 | 35 | 0 | 0.125 | 1 | # | 20 | Baseline | sided2 | 100 |
| # | interaction | 0 | 0 | 4 | 0 | 0 | # | 20 | Baseline | sided2 | 100 |
| # | disease | 7 | 0 | 0 | 1 | 1 | # | 20 | Small | paralle | 100 |
| # | sex | 4 | 0 | 0 | 1 | 1 | # | 20 | Small | paralle | 100 |
| # | time | 5 | 0 | 0 | 1 | 1 | # | 20 | Small | paralle | 100 |
| # | interaction | 4 | 0 | 0 | 1 | 1 | # | 20 | Small | paralle | 100 |
| # | disease | 7 | 4 | 0 | 0.636363636 | 1 | # | 20 | Small | ancom | 100 |
| # | sex | 4 | 0 | 0 | 1 | 1 | # | 20 | Small | ancom | 100 |
| # | time | 5 | 4 | 0 | 0.555555556 | 1 | # | 20 | Small | ancom | 100 |
| # | interaction | 0 | 0 | 4 | 0 | 0 | # | 20 | Small | ancom | 100 |
| # | disease | 7 | 3 | 0 | 0.7 | 1 | # | 20 | Small | sided2 | 100 |
| # | sex | 4 | 1 | 0 | 0.8 | 1 | # | 20 | Small | sided2 | 100 |
| # | time | 5 | 5 | 0 | 0.5 | 1 | # | 20 | Small | sided2 | 100 |
| # | interaction | 0 | 0 | 4 | 0 | 0 | # | 20 | Small | sided2 | 100 |
| # | disease | 7 | 16 | 0 | 0.304347826 | 1 | # | 20 | Baseline | paralle | 100 |
| # | sex | 4 | 9 | 0 | 0.307692308 | 1 | # | 20 | Baseline | paralle | 100 |
| # | time | 5 | 7 | 0 | 0.416666667 | 1 | # | 20 | Baseline | paralle | 100 |
| # | interaction | 4 | 7 | 0 | 0.363636364 | 1 | # | 20 | Baseline | paralle | 100 |
| # | disease | 7 | 4 | 0 | 0.636363636 | 1 | # | 20 | Baseline | ancom | 100 |
| # | sex | 4 | 0 | 0 | 1 | 1 | # | 20 | Baseline | ancom | 100 |
| # | time | 5 | 4 | 0 | 0.555555556 | 1 | # | 20 | Baseline | ancom | 100 |
| # | interaction | 0 | 0 | 4 | 0 | 0 | # | 20 | Baseline | ancom | 100 |
| # | disease | 7 | 64 | 0 | 0.098591549 | 1 | # | 20 | Baseline | sided2 | 100 |
| # | sex | 4 | 18 | 0 | 0.181818182 | 1 | # | 20 | Baseline | sided2 | 100 |
| # | time | 5 | 65 | 0 | 0.1 | 1 | # | 20 | Baseline | sided2 | 100 |
| # | interaction | 0 | 0 | 4 | 0 | 0 | # | 20 | Baseline | sided2 | 100 |
| # | disease | 7 | 0 | 0 | 1 | 1 | # | 20 | Small | paralle | 100 |
| # | sex | 4 | 0 | 0 | 1 | 1 | # | 20 | Small | paralle | 100 |
| # | time | 5 | 0 | 0 | 1 | 1 | # | 20 | Small | paralle | 100 |
| # | interaction | 4 | 0 | 0 | 1 | 1 | # | 20 | Small | paralle | 100 |
| # | disease | 7 | 4 | 0 | 0.636363636 | 1 | # | 20 | Small | ancom | 100 |
| # | sex | 4 | 0 | 0 | 1 | 1 | # | 20 | Small | ancom | 100 |
| # | time | 5 | 4 | 0 | 0.555555556 | 1 | # | 20 | Small | ancom | 100 |
| # | interaction | 0 | 0 | 4 | 0 | 0 | # | 20 | Small | ancom | 100 |
| # | disease | 7 | 10 | 0 | 0.411764706 | 1 | # | 20 | Small | sided2 | 100 |
| # | sex | 4 | 3 | 0 | 0.571428571 | 1 | # | 20 | Small | sided2 | 100 |
| # | time | 5 | 9 | 0 | 0.357142857 | 1 | # | 20 | Small | sided2 | 100 |
| # | interaction | 0 | 0 | 4 | 0 | 0 | # | 20 | Small | sided2 | 100 |
